## Supplementary material for "Chemokine Receptor 1 and its associated immune pathway are downregulated in SF3B1^MT^ blood and non-blood cancers": Table_S11

**Table S11. GSEA analysis in $SF3B1^{MT}$ and $SF3B1^{WT}$ AML patients.** See Table S16 for patient information.

**GSEA Positive** Hematopoietic/Immune pathways are highlighted in red.

| Gene sets | SIZE | ES | NES | NOM p-val | FDR q-val |
|---|---|---|---|---|---|
| GOBP_HOMOPHILIC_CELL_ADHESION_VIA_PLASMA_MEMBRANE_ADHESION_MOLECULE | 103 | 0.569891 | 1.972815 | 0 | 0.048365 |
| GOBP_CALCIUM_DEPENDENT_CELL_CELL_ADHESION_VIA_PLASMA_MEMBRANE_CELL_ | 23 | 0.728076 | 1.933065 | 0.001901 | 0.051056 |
| GOBP_NOTOCHORD_DEVELOPMENT | 15 | 0.767038 | 1.769344 | 0 | 0.50009 |
| GOBP_CAMERA_TYPE_EYE_PHOTORECEPTOR_CELL_DIFFERENTIATION | 16 | 0.712429 | 1.749367 | 0.01626 | 0.49215 |
| GOBP_COCHLEA_MORPHOGENESIS | 15 | 0.725908 | 1.745972 | 0.005769 | 0.409495 |
| GOBP_CELL_PROLIFERATION_IN_FOREBRAIN | 15 | 0.717513 | 1.74123 | 0.002028 | 0.363584 |
| GOBP_COCHLEA_DEVELOPMENT | 31 | 0.616195 | 1.729923 | 0.003891 | 0.35996 |
| GOBP_GASTRULATION_WITH_MOUTH_FORMING_SECOND | 18 | 0.689699 | 1.716622 | 0.007605 | 0.368825 |
| GOBP_NATURAL_KILLER_CELL_DIFFERENTIATION | 17 | 0.686719 | 1.687783 | 0.007843 | 0.451157 |
| GOBP_RENAL_FILTRATION | 21 | 0.6524 | 1.682955 | 0.007491 | 0.428412 |
| GOBP_POSITIVE_REGULATION_OF_NATURAL_KILLER_CELL_MEDIATED_IMMUNITY | 27 | 0.62481 | 1.673276 | 0.011858 | 0.432578 |
| GOBP_SYNAPTONEMAL_COMPLEX_ORGANIZATION | 19 | 0.655295 | 1.660279 | 0.012658 | 0.459704 |
| GOBP_CELLULAR_RESPONSE_TO_VITAMIN_D | 17 | 0.6691 | 1.637167 | 0.004065 | 0.541587 |
| GOBP_INTEGRIN_ACTIVATION | 22 | 0.644442 | 1.636794 | 0.003891 | 0.504826 |
| GOBP_ANION_HOMEOSTASIS | 23 | 0.620292 | 1.635986 | 0.013359 | 0.475051 |
| GOBP_REGULATION_OF_NATURAL_KILLER_CELL_MEDIATED_IMMUNITY | 42 | 0.553406 | 1.632868 | 0.005814 | 0.460731 |
| GOBP_NEGATIVE_REGULATION_OF_CARTILAGE_DEVELOPMENT | 16 | 0.665556 | 1.623485 | 0.017682 | 0.475796 |
| GOBP_POSITIVE_REGULATION_OF_MULTI_ORGANISM_PROCESS | 17 | 0.658602 | 1.60816 | 0.013566 | 0.522994 |
| GOBP_AXON_ENSHEATHMENT_IN_CENTRAL_NERVOUS_SYSTEM | 16 | 0.645589 | 1.607309 | 0.017953 | 0.500032 |
| GOBP_SKELETAL_MUSCLE_CONTRACTION | 30 | 0.580498 | 1.607072 | 0.020446 | 0.476258 |
| GOBP_REGULATION_OF_CARDIAC_MUSCLE_CONTRACTION_BY_REGULATION_OF_THE_ | 17 | 0.649071 | 1.605244 | 0.013699 | 0.460993 |
| GOBP_SENSORY_PERCEPTION_OF_BITTER_TASTE | 23 | 0.608696 | 1.60499 | 0.013133 | 0.441128 |
| GOBP_REGULATION_OF_NUCLEOBASE_CONTAINING_COMPOUND_TRANSPORT | 15 | 0.670642 | 1.604631 | 0.029575 | 0.423653 |
| GOBP_MAMMARY_GLAND_MORPHOGENESIS | 31 | 0.559322 | 1.593992 | 0.031308 | 0.448233 |
| GOBP_CEREBELLAR_CORTEX_MORPHOGENESIS | 25 | 0.590544 | 1.593763 | 0.009191 | 0.431145 |
| GOBP_POSITIVE_REGULATION_OF_NATURAL_KILLER_CELL_MEDIATED_CYTOTOXICITY | 23 | 0.604282 | 1.585539 | 0.027985 | 0.447155 |
| GOBP_THYMOCYTE_APOPTOTIC_PROCESS | 15 | 0.664792 | 1.583323 | 0.028195 | 0.439697 |
| GOBP_HINDBRAIN_MORPHOGENESIS | 34 | 0.552648 | 1.580422 | 0.017176 | 0.435678 |
| GOBP_MAMMARY_GLAND_DUCT_MORPHOGENESIS | 22 | 0.598422 | 1.579263 | 0.023762 | 0.42513 |
| GOBP_ACYLGLYCEROL_HOMEOSTASIS | 22 | 0.601483 | 1.578639 | 0.025896 | 0.413391 |
| GOBP_NEGATIVE_REGULATION_OF_SMOOTHENED_SIGNALING_PATHWAY | 28 | 0.566029 | 1.570879 | 0.030075 | 0.428914 |
| GOBP_REGULATION_OF_CARTILAGE_DEVELOPMENT | 50 | 0.509877 | 1.569828 | 0.013158 | 0.419168 |
| GOBP_DETECTION_OF_STIMULUS_INVOLVED_IN_SENSORY_PERCEPTION | 91 | 0.455099 | 1.565195 | 0.009579 | 0.424206 |
| GOBP_CELL_JUNCTION_ASSEMBLY | 309 | 0.395213 | 1.564019 | 0.001783 | 0.416021 |
| GOBP_DETECTION_OF_CHEMICAL_STIMULUS | 73 | 0.479994 | 1.556115 | 0.009488 | 0.433609 |
| GOBP_MITOCHONDRIAL_DEPOLARIZATION | 19 | 0.618753 | 1.554276 | 0.036398 | 0.428257 |
| GOBP_FOREBRAIN_NEURON_DEVELOPMENT | 17 | 0.632189 | 1.554164 | 0.033465 | 0.417027 |
| GOBP_BONE_GROWTH | 19 | 0.6179 | 1.544549 | 0.037109 | 0.441104 |
| GOBP_CELL_CELL_ADHESION_VIA_PLASMA_MEMBRANE_ADHESION_MOLECULES | 173 | 0.413439 | 1.538299 | 0.001866 | 0.452845 |
| GOBP_POLARIZED_EPITHELIAL_CELL_DIFFERENTIATION | 20 | 0.610189 | 1.533244 | 0.029354 | 0.461208 |
| GOBP_DETECTION_OF_CHEMICAL_STIMULUS_INVOLVED_IN_SENSORY_PERCEPTION_O | 22 | 0.581048 | 1.530362 | 0.030797 | 0.460632 |
| GOBP_FOREBRAIN_CELL_MIGRATION | 40 | 0.51965 | 1.53 | 0.021195 | 0.451051 |
| GOBP_ANTIGEN_PROCESSING_AND_PRESENTATION_OF_ENDOGENOUS_ANTIGEN | 24 | 0.578525 | 1.528285 | 0.020367 | 0.446949 |
| GOBP_REGULATION_OF_LEUKOCYTE_MEDIATED_CYTOTOXICITY | 73 | 0.463998 | 1.520841 | 0.011278 | 0.46522 |
| GOBP_SUBPALLIUM_DEVELOPMENT | 17 | 0.614836 | 1.516731 | 0.044146 | 0.470557 |
| GOBP_COTRANSLATIONAL_PROTEIN_TARGETING_TO_MEMBRANE | 102 | 0.435841 | 1.513533 | 0.001923 | 0.473411 |
| GOBP_POSITIVE_REGULATION_OF_CELL_KILLING | 58 | 0.479495 | 1.511902 | 0.019268 | 0.469612 |
| GOBP_MULTICELLULAR_ORGANISMAL_MOVEMENT | 37 | 0.521581 | 1.511743 | 0.039698 | 0.460541 |
| GOBP_CEREBRAL_CORTEX_DEVELOPMENT | 87 | 0.444976 | 1.502247 | 0.01145 | 0.487438 |
| GOBP_SENSORY_PERCEPTION_OF_SMELL | 39 | 0.51285 | 1.501813 | 0.035363 | 0.479366 |
| GOBP_REGULATION_OF_STRIATED_MUSCLE_CONTRACTION | 60 | 0.478396 | 1.501555 | 0.017731 | 0.47103 |
| GOBP_ACIDIC_AMINO_ACID_TRANSPORT | 54 | 0.473879 | 1.498638 | 0.026515 | 0.472458 |
| GOBP_TELENCEPHALON_DEVELOPMENT | 174 | 0.403946 | 1.494162 | 0.003509 | 0.480771 |
| GOBP_ADHERENS_JUNCTION_ORGANIZATION | 54 | 0.48015 | 1.490375 | 0.023339 | 0.486755 |
| GOBP_POSITIVE_REGULATION_OF_AXON_EXTENSION | 33 | 0.523403 | 1.485516 | 0.040619 | 0.496866 |
| GOBP_TELENCEPHALON_GLIAL_CELL_MIGRATION | 19 | 0.57626 | 1.483716 | 0.049808 | 0.494738 |
| GOBP_NEGATIVE_REGULATION_OF_CHONDROCYTE_DIFFERENTIATION | 15 | 0.637766 | 1.481937 | 0.047244 | 0.492928 |
| GOBP_CAMERA_TYPE_EYE_MORPHOGENESIS | 76 | 0.448871 | 1.481262 | 0.026266 | 0.487055 |
| GOBP_POSITIVE_REGULATION_OF_MITOCHONDRIAL_FISSION | 17 | 0.60564 | 1.478809 | 0.056452 | 0.487778 |
| GOBP_POTASSIUM_ION_HOMEOSTASIS | 24 | 0.557233 | 1.475236 | 0.038182 | 0.493509 |
| GOBP_NEGATIVE_REGULATION_OF_LEUKOCYTE_MIGRATION | 40 | 0.50002 | 1.468137 | 0.036697 | 0.512505 |
| GOBP_LIMBIC_SYSTEM_DEVELOPMENT | 66 | 0.447259 | 1.46546 | 0.028302 | 0.514113 |
| GOBP_ESTABLISHMENT_OR_MAINTENANCE_OF_MONOPOLAR_CELL_POLARITY | 20 | 0.561716 | 1.463636 | 0.069098 | 0.512877 |
| GOBP_DETECTION_OF_STIMULUS | 221 | 0.384901 | 1.461332 | 0.003552 | 0.513636 |
| GOBP_CENTRAL_NERVOUS_SYSTEM_NEURON_DEVELOPMENT | 63 | 0.458922 | 1.457047 | 0.028829 | 0.522481 |
| GOBP_REGULATION_OF_BICELLULAR_TIGHT_JUNCTION_ASSEMBLY | 20 | 0.578438 | 1.45399 | 0.051429 | 0.526387 |
| GOBP_VASCULOGENESIS | 64 | 0.443323 | 1.449656 | 0.038889 | 0.53515 |
| GOBP_REGULATION_OF_MEMBRANE_DEPOLARIZATION | 29 | 0.522137 | 1.449405 | 0.052529 | 0.528323 |
| GOBP_SODIUM_ION_HOMEOSTASIS | 35 | 0.501012 | 1.446761 | 0.036166 | 0.530944 |
| GOBP_NEGATIVE_REGULATION_OF_MYOBLAST_DIFFERENTIATION | 18 | 0.566161 | 1.443471 | 0.068273 | 0.536395 |
| GOBP_NEGATIVE_REGULATION_OF_MRNA_CATABOLIC_PROCESS | 48 | 0.462488 | 1.442478 | 0.044534 | 0.532853 |
| GOBP_NEURAL_RETINA_DEVELOPMENT | 46 | 0.475126 | 1.44145 | 0.042279 | 0.529264 |
| GOBP_SENSORY_PERCEPTION_OF_CHEMICAL_STIMULUS | 81 | 0.42865 | 1.440086 | 0.026616 | 0.527281 |
| GOBP_REGULATION_OF_CARDIAC_MUSCLE_CONTRACTION_BY_CALCIUM_ION_SIGNALI | 21 | 0.543111 | 1.438685 | 0.078095 | 0.525454 |
| GOBP_ARACHIDONIC_ACID_SECRETION | 15 | 0.587292 | 1.437919 | 0.081349 | 0.521462 |
| GOBP_NEGATIVE_REGULATION_OF_MUSCLE_CONTRACTION | 18 | 0.581561 | 1.434777 | 0.079523 | 0.526454 |
| GOBP_SECONDARY_METABOLITE_BIOSYNTHETIC_PROCESS | 17 | 0.583466 | 1.434399 | 0.064 | 0.521042 |
| GOBP_POSITIVE_REGULATION_OF_SYNAPSE_ASSEMBLY | 38 | 0.492315 | 1.43332 | 0.052224 | 0.518494 |
| GOBP_CELL_SURFACE_RECEPTOR_SIGNALING_PATHWAY_INVOLVED_IN_HEART_DEVEL | 20 | 0.560085 | 1.432235 | 0.068834 | 0.515948 |
| GOBP_GLAND_MORPHOGENESIS | 86 | 0.419594 | 1.431173 | 0.026975 | 0.513364 |
| GOBP_REGULATION_OF_CELL_JUNCTION_ASSEMBLY | 148 | 0.387582 | 1.427365 | 0.007394 | 0.521204 |
| GOBP_INNER_EAR_RECEPTOR_CELL_DEVELOPMENT | 31 | 0.509725 | 1.425671 | 0.052158 | 0.521202 |
| GOBP_EXTRACELLULAR_MATRIX_ASSEMBLY | 37 | 0.497454 | 1.424583 | 0.041502 | 0.519064 |
| GOBP_NEURAL_CREST_CELL_DIFFERENTIATION | 63 | 0.4478 | 1.422113 | 0.032197 | 0.522421 |
| GOBP_CARBOHYDRATE_DERIVATIVE_TRANSPORT | 74 | 0.434647 | 1.421755 | 0.03125 | 0.517564 |
| GOBP_LONG_TERM_SYNAPTIC_DEPRESSION | 22 | 0.532232 | 1.420467 | 0.081136 | 0.516639 |
| GOBP_VENTRICULAR_SEPTUM_MORPHOGENESIS | 29 | 0.513855 | 1.420347 | 0.082031 | 0.51109 |
| GOBP_POSITIVE_REGULATION_OF_STEM_CELL_PROLIFERATION | 17 | 0.579668 | 1.419096 | 0.090395 | 0.509498 |
| GOBP_POSITIVE_REGULATION_OF_AXONOGENESIS | 65 | 0.440302 | 1.418218 | 0.047619 | 0.506866 |
| GOBP_BINDING_OF_SPERM_TO_ZONA_PELLUCIDA | 27 | 0.520786 | 1.417848 | 0.057915 | 0.50269 |
| GOBP_CELL_COMMUNICATION_BY_ELECTRICAL_COUPLING_INVOLVED_IN_CARDIAC_C | 19 | 0.558365 | 1.417016 | 0.09369 | 0.500084 |
| GOBP_NEGATIVE_REGULATION_OF_BLOOD_CIRCULATION | 16 | 0.580754 | 1.416816 | 0.088632 | 0.495447 |
| GOBP_MUSCLE_CELL_CELLULAR_HOMEOSTASIS | 17 | 0.573165 | 1.412892 | 0.086042 | 0.503751 |
| GOBP_POSITIVE_REGULATION_OF_AMINE_TRANSPORT | 27 | 0.515458 | 1.412682 | 0.077821 | 0.499153 |
| GOBP_ASPARTATE_FAMILY_AMINO_ACID_CATABOLIC_PROCESS | 22 | 0.535909 | 1.411873 | 0.092131 | 0.496717 |
| GOBP_CELL_SUBSTRATE_JUNCTION_ORGANIZATION | 93 | 0.411295 | 1.411754 | 0.024074 | 0.491901 |
| GOBP_PROSTATE_GLAND_DEVELOPMENT | 32 | 0.502561 | 1.408822 | 0.068223 | 0.497542 |
| GOBP_CEREBRAL_CORTEX_RADIALLY_ORIENTED_CELL_MIGRATION | 24 | 0.528045 | 1.407135 | 0.074766 | 0.498203 |
| GOBP_GLUTAMATE_SECRETION | 33 | 0.500389 | 1.404437 | 0.065476 | 0.50249 |
| GOBP_NEGATIVE_REGULATION_OF_PEPTIDE_HORMONE_SECRETION | 28 | 0.514199 | 1.40309 | 0.067039 | 0.502071 |
| GOBP_NEGATIVE_REGULATION_OF_RNA_CATABOLIC_PROCESS | 59 | 0.437004 | 1.398166 | 0.033457 | 0.51431 |
| GOBP_POSITIVE_REGULATION_OF_CELL_JUNCTION_ASSEMBLY | 65 | 0.438121 | 1.397269 | 0.038391 | 0.512469 |
| GOBP_REGULATION_OF_NATURAL_KILLER_CELL_ACTIVATION | 30 | 0.506992 | 1.39639 | 0.068898 | 0.510587 |
| GOBP_SENSORY_PERCEPTION_OF_TASTE | 38 | 0.484606 | 1.392883 | 0.062157 | 0.517738 |
| GOBP_SYNAPSE_ASSEMBLY | 112 | 0.395204 | 1.392094 | 0.032028 | 0.515407 |
| GOBP_PROTEOGLYCAN_BIOSYNTHETIC_PROCESS | 55 | 0.442405 | 1.390741 | 0.054264 | 0.515317 |
| GOBP_PROTEIN_HETEROOLIGOMERIZATION | 21 | 0.543204 | 1.389909 | 0.089054 | 0.513431 |
| GOBP_REGULATION_OF_CHONDROCYTE_DIFFERENTIATION | 38 | 0.479505 | 1.389586 | 0.06968 | 0.50986 |
| GOBP_POSITIVE_REGULATION_OF_PHOSPHATIDYLINOSITOL_3_KINASE_SIGNALING | 79 | 0.417704 | 1.388482 | 0.052023 | 0.508909 |
| GOBP_THYMIC_T_CELL_SELECTION | 21 | 0.549688 | 1.382807 | 0.066922 | 0.523755 |
| GOBP_AXON_EXTENSION | 100 | 0.398944 | 1.381078 | 0.028419 | 0.524944 |
| GOBP_SKIN_EPIDERMIS_DEVELOPMENT | 67 | 0.416611 | 1.378923 | 0.056391 | 0.527711 |
| GOBP_NEUROBLAST_PROLIFERATION | 38 | 0.473414 | 1.377622 | 0.053903 | 0.527674 |
| GOBP_POSITIVE_REGULATION_OF_NATURAL_KILLER_CELL_ACTIVATION | 19 | 0.550096 | 1.377293 | 0.104449 | 0.524258 |
| GOBP_FOREBRAIN_GENERATION_OF_NEURONS | 33 | 0.474313 | 1.376044 | 0.07874 | 0.523972 |
| GOBP_REGULATION_OF_EXTENT_OF_CELL_GROWTH | 86 | 0.406326 | 1.374144 | 0.041509 | 0.525751 |
| GOBP_POSITIVE_REGULATION_OF_NERVOUS_SYSTEM_DEVELOPMENT | 209 | 0.356532 | 1.369505 | 0.015625 | 0.537501 |
| GOBP_KERATINIZATION | 51 | 0.438701 | 1.36715 | 0.062264 | 0.541317 |
| GOBP_ACTIN_MEDIATED_CELL_CONTRACTION | 81 | 0.40044 | 1.365651 | 0.037244 | 0.542125 |
| GOBP_EYE_MORPHOGENESIS | 97 | 0.398453 | 1.364654 | 0.041971 | 0.541119 |
| GOBP_NATURAL_KILLER_CELL_MEDIATED_IMMUNITY | 59 | 0.432997 | 1.360369 | 0.056711 | 0.551612 |
| GOBP_CELL_CELL_JUNCTION_ORGANIZATION | 163 | 0.37106 | 1.358355 | 0.023381 | 0.554154 |
| GOBP_NEGATIVE_REGULATION_OF_HORMONE_SECRETION | 43 | 0.454009 | 1.358192 | 0.072222 | 0.550271 |
| GOBP_REGULATION_OF_OSTEOBLAST_PROLIFERATION | 20 | 0.539901 | 1.35752 | 0.105159 | 0.548142 |
| GOBP_REGULATION_OF_PLATELET_DERIVED_GROWTH_FACTOR_RECEPTOR_SIGNALIN | 20 | 0.520676 | 1.356395 | 0.104046 | 0.547717 |
| GOBP_REGULATION_OF_SYNAPTIC_PLASTICITY | 128 | 0.375872 | 1.356195 | 0.027473 | 0.544039 |
| GOBP_RESPONSE_TO_AUDITORY_STIMULUS | 18 | 0.551252 | 1.355607 | 0.138943 | 0.541717 |
| GOBP_T_CELL_SELECTION | 46 | 0.456671 | 1.354051 | 0.073529 | 0.542964 |
| GOBP_REGULATION_OF_CELL_KILLING | 90 | 0.396696 | 1.353537 | 0.063869 | 0.540495 |
| GOBP_REGULATION_OF_RESPONSE_TO_EXTRACELLULAR_STIMULUS | 17 | 0.533982 | 1.353213 | 0.116505 | 0.537479 |
| GOBP_NEURAL_PRECURSOR_CELL_PROLIFERATION | 99 | 0.387975 | 1.352442 | 0.045541 | 0.536089 |
| GOBP_SYNAPSE_ORGANIZATION | 309 | 0.342013 | 1.352356 | 0.005208 | 0.532313 |
| GOBP_CORNIFICATION | 41 | 0.449199 | 1.350778 | 0.09434 | 0.533674 |
| GOBP_MYOBLAST_PROLIFERATION | 15 | 0.57381 | 1.350015 | 0.12782 | 0.532529 |
| GOBP_CELL_CELL_JUNCTION_ASSEMBLY | 105 | 0.386193 | 1.349757 | 0.034926 | 0.52942 |
| GOBP_ANTIGEN_PROCESSING_AND_PRESENTATION_OF_PEPTIDE_OR_POLYSACCHARID | 98 | 0.391546 | 1.348924 | 0.042279 | 0.528411 |
| GOBP_NEGATIVE_REGULATION_OF_EPITHELIAL_CELL_MIGRATION | 62 | 0.418793 | 1.348814 | 0.063197 | 0.524912 |
| GOBP_OLIGODENDROCYTE_DEVELOPMENT | 35 | 0.463876 | 1.348649 | 0.08046 | 0.521711 |
| GOBP_NEURON_PROJECTION_EXTENSION_INVOLVED_IN_NEURON_PROJECTION_GUIDA | 31 | 0.48045 | 1.343108 | 0.095685 | 0.536693 |
| GOBP_MUSCLE_FILAMENT_SLIDING | 24 | 0.502272 | 1.342173 | 0.120858 | 0.536056 |
| GOBP_RETINA_MORPHOGENESIS_IN_CAMERA_TYPE_EYE | 40 | 0.456825 | 1.341459 | 0.095057 | 0.534628 |
| GOBP_ADULT_BEHAVIOR | 79 | 0.401116 | 1.340094 | 0.064961 | 0.535492 |
| GOBP_PROSTATE_GLAND_MORPHOGENESIS | 19 | 0.536084 | 1.336381 | 0.126654 | 0.54439 |
| GOBP_CEREBRAL_CORTEX_CELL_MIGRATION | 31 | 0.481324 | 1.335784 | 0.089127 | 0.542581 |
| GOBP_CALCIUM_ION_REGULATED_EXOCYTOSIS | 49 | 0.432831 | 1.335001 | 0.07326 | 0.541447 |
| GOBP_HOMOLOGOUS_CHROMOSOME_PAIRING_AT_MEIOSIS | 37 | 0.460655 | 1.33325 | 0.079245 | 0.543671 |
| GOBP_RETINA_LAYER_FORMATION | 18 | 0.535151 | 1.328988 | 0.147874 | 0.5549 |
| GOBP_HEMATOPOIETIC_STEM_CELL_PROLIFERATION | 23 | 0.50633 | 1.327925 | 0.126838 | 0.554702 |
| GOBP_CENTRAL_NERVOUS_SYSTEM_PROJECTION_NEURON_AXONOGENESIS | 24 | 0.501549 | 1.327588 | 0.119608 | 0.552168 |
| GOBP_REGULATION_OF_CARDIAC_MUSCLE_CONTRACTION | 51 | 0.425239 | 1.325532 | 0.081481 | 0.555408 |
| GOBP_RENAL_SYSTEM_PROCESS_INVOLVED_IN_REGULATION_OF_SYSTEMIC_ARTERIA | 15 | 0.552525 | 1.32093 | 0.150301 | 0.568042 |
| GOBP_PROTEIN_ACTIVATION_CASCADE | 18 | 0.526654 | 1.320483 | 0.161597 | 0.565924 |
| GOBP_NEGATIVE_REGULATION_OF_ANOIKIS | 17 | 0.530696 | 1.319158 | 0.145914 | 0.566796 |

**GSEA Negative** Hematopoietic/Immune pathways are highlighted in green.

| Gene sets | SIZE | ES | NES | NOM p-val | FDR q-val |
|---|---|---|---|---|---|
| GOBP_POSITIVE_REGULATION_OF_NITRIC_OXIDE_METABOLIC_PROCESS | 35 | -0.766579 | -2.313989 | 0 | 0 |
| GOBP_POSITIVE_REGULATION_OF_REACTIVE_OXYGEN_SPECIES_BIOSYNTHETIC_PROCESS | 48 | -0.723554 | -2.263968 | 0 | 0 |
| GOBP_ANTIMICROBIAL_HUMORAL_IMMUNE_RESPONSE_MEDIATED_BY_ANTIMICROBIAL_PE | 41 | -0.745615 | -2.228916 | 0 | 0 |
| GOBP_POSITIVE_REGULATION_OF_REACTIVE_OXYGEN_SPECIES_METABOLIC_PROCESS | 86 | -0.639453 | -2.209405 | 0 | 0 |
| GOBP_NEUTROPHIL_MIGRATION | 97 | -0.602404 | -2.120387 | 0 | 6.31E-04 |
| GOBP_NEUTROPHIL_CHEMOTAXIS | 79 | -0.614944 | -2.109463 | 0 | 7.86E-04 |
| GOBP_ANTIMICROBIAL_HUMORAL_RESPONSE | 75 | -0.614675 | -2.08316 | 0 | 0.003849 |
| GOBP_RESPONSE_TO_HYDROGEN_PEROXIDE | 120 | -0.570016 | -2.074214 | 0 | 0.00416 |
| GOBP_HUMORAL_IMMUNE_RESPONSE | 249 | -0.520479 | -2.062616 | 0 | 0.004409 |
| GOBP_DEFENSE_RESPONSE_TO_BACTERIUM | 212 | -0.524117 | -2.054695 | 0 | 0.00428 |
| GOBP_RESPONSE_TO_CHEMOKINE | 72 | -0.611805 | -2.046238 | 0 | 0.00503 |
| GOBP_MYELOID_LEUKOCYTE_MIGRATION | 177 | -0.531071 | -2.026159 | 0 | 0.006846 |
| GOBP_REACTIVE_NITROGEN_SPECIES_METABOLIC_PROCESS | 68 | -0.605656 | -2.02357 | 0 | 0.006441 |
| GOBP_REACTIVE_OXYGEN_SPECIES_BIOSYNTHETIC_PROCESS | 102 | -0.55779 | -2.022639 | 0 | 0.006094 |
| GOBP_DEFENSE_RESPONSE_TO_GRAM_NEGATIVE_BACTERIUM | 50 | -0.644258 | -2.02166 | 0 | 0.00579 |
| GOBP_RESPONSE_TO_DSRNA | 35 | -0.673868 | -2.011268 | 0 | 0.006912 |
| GOBP_REGULATION_OF_REACTIVE_OXYGEN_SPECIES_METABOLIC_PROCESS | 159 | -0.528293 | -1.993392 | 0 | 0.010243 |
| GOBP_GRANULOCYTE_MIGRATION | 119 | -0.55066 | -1.990967 | 0 | 0.010378 |
| GOBP_BROWN_FAT_CELL_DIFFERENTIATION | 37 | -0.662345 | -1.986776 | 0 | 0.011002 |
| GOBP_GAS_TRANSPORT | 17 | -0.786775 | -1.981134 | 0 | 0.011559 |
| GOBP_REGULATION_OF_REACTIVE_OXYGEN_SPECIES_BIOSYNTHETIC_PROCESS | 81 | -0.569851 | -1.972837 | 0 | 0.012443 |
| GOBP_TRANSPOSITION | 23 | -0.733237 | -1.971824 | 0 | 0.012237 |
| GOBP_TETRAPYRROLE_BIOSYNTHETIC_PROCESS | 28 | -0.6976 | -1.969311 | 0 | 0.011915 |
| GOBP_RESPONSE_TO_ACTIVITY | 54 | -0.620855 | -1.967292 | 0 | 0.01201 |
| GOBP_REGULATION_OF_NITRIC_OXIDE_METABOLIC_PROCESS | 52 | -0.622076 | -1.961783 | 0.002146 | 0.013117 |
| GOBP_NEGATIVE_REGULATION_OF_MAP_KINASE_ACTIVITY | 61 | -0.595629 | -1.960103 | 0 | 0.012856 |
| GOBP_MICROGLIAL_CELL_ACTIVATION | 41 | -0.634913 | -1.958834 | 0 | 0.012792 |
| GOBP_GRANULOCYTE_CHEMOTAXIS | 98 | -0.554095 | -1.95837 | 0 | 0.012447 |
| GOBP_ERYTHROCYTE_DEVELOPMENT | 31 | -0.693233 | -1.955155 | 0 | 0.012452 |
| GOBP_INNATE_IMMUNE_RESPONSE_IN_MUCOSA | 19 | -0.75075 | -1.947939 | 0 | 0.013623 |
| GOBP_REGULATION_OF_COMPLEMENT_ACTIVATION | 91 | -0.556389 | -1.943047 | 0 | 0.014409 |
| GOBP_BICARBONATE_TRANSPORT | 36 | -0.65404 | -1.941703 | 0 | 0.014553 |
| GOBP_NUCLEOTIDE_BINDING_OLIGOMERIZATION_DOMAIN_CONTAINING_2_SIGNALING_PAT | 16 | -0.775841 | -1.932865 | 0 | 0.015934 |
| GOBP_RESPONSE_TO_LEUKEMIA_INHIBITORY_FACTOR | 72 | -0.55903 | -1.92351 | 0 | 0.018171 |
| GOBP_THYROID_GLAND_DEVELOPMENT | 16 | -0.769434 | -1.914979 | 0 | 0.019824 |
| GOBP_MYELOID_CELL_HOMEOSTASIS | 141 | -0.515608 | -1.914648 | 0 | 0.019318 |
| GOBP_REACTIVE_OXYGEN_SPECIES_METABOLIC_PROCESS | 237 | -0.486617 | -1.911472 | 0 | 0.019527 |
| GOBP_REGULATION_OF_TRANSPOSITION | 21 | -0.710758 | -1.907648 | 0 | 0.020429 |
| GOBP_REGULATION_OF_HUMORAL_IMMUNE_RESPONSE | 105 | -0.538641 | -1.899131 | 0 | 0.022743 |
| GOBP_SNRNA_PROCESSING | 30 | -0.666877 | -1.898174 | 0.002049 | 0.022645 |
| GOBP_ERYTHROCYTE_HOMEOSTASIS | 119 | -0.528919 | -1.896695 | 0 | 0.022439 |
| GOBP_MEMBRANE_REPOLARIZATION_DURING_ACTION_POTENTIAL | 20 | -0.715305 | -1.879515 | 0 | 0.028966 |
| GOBP_HEMOGLOBIN_METABOLIC_PROCESS | 16 | -0.768424 | -1.87821 | 0 | 0.029067 |
| GOBP_COMPLEMENT_ACTIVATION | 132 | -0.511792 | -1.876716 | 0 | 0.029017 |
| GOBP_POSITIVE_REGULATION_OF_MITOTIC_NUCLEAR_DIVISION | 38 | -0.639155 | -1.871335 | 0 | 0.030691 |
| GOBP_RECEPTOR_MEDIATED_ENDOCYTOSIS | 273 | -0.464418 | -1.871293 | 0 | 0.030057 |
| GOBP_ANTIBACTERIAL_HUMORAL_RESPONSE | 37 | -0.625927 | -1.870632 | 0 | 0.029691 |
| GOBP_ORGAN_OR_TISSUE_SPECIFIC_IMMUNE_RESPONSE | 28 | -0.656367 | -1.859137 | 0.002083 | 0.034219 |
| GOBP_POLYSACCHARIDE_CATABOLIC_PROCESS | 24 | -0.689077 | -1.857652 | 0.002058 | 0.034035 |
| GOBP_NEGATIVE_REGULATION_OF_VIRAL_GENOME_REPLICATION | 44 | -0.602546 | -1.85498 | 0.00211 | 0.034806 |
| GOBP_CELLULAR_RESPONSE_TO_EXOGENOUS_DSRNA | 17 | -0.739138 | -1.853563 | 0 | 0.034867 |
| GOBP_POSITIVE_REGULATION_OF_ACUTE_INFLAMMATORY_RESPONSE | 22 | -0.683258 | -1.852346 | 0.002066 | 0.034931 |
| GOBP_REGULATION_OF_MAP_KINASE_ACTIVITY | 264 | -0.46021 | -1.848862 | 0 | 0.035947 |
| GOBP_RESPONSE_TO_IMMOBILIZATION_STRESS | 18 | -0.734961 | -1.848621 | 0 | 0.035369 |
| GOBP_REGULATION_OF_MITOTIC_NUCLEAR_DIVISION | 102 | -0.521533 | -1.843176 | 0 | 0.038063 |
| GOBP_RESPONSE_TO_COLD | 41 | -0.609478 | -1.840613 | 0 | 0.038826 |
| GOBP_MUSCLE_CELL_MIGRATION | 69 | -0.543314 | -1.840508 | 0 | 0.038229 |
| GOBP_POSITIVE_REGULATION_OF_MICROTUBULE_POLYMERIZATION_OR_DEPOLYMERIZAT | 29 | -0.655487 | -1.839469 | 0 | 0.038306 |
| GOBP_NEGATIVE_REGULATION_OF_EXTRINSIC_APOPTOTIC_SIGNALING_PATHWAY | 89 | -0.528273 | -1.829416 | 0 | 0.043462 |
| GOBP_POSITIVE_REGULATION_OF_SMOOTH_MUSCLE_CELL_MIGRATION | 33 | -0.62943 | -1.82887 | 0.002193 | 0.042816 |
| GOBP_DEFENSE_RESPONSE_TO_FUNGUS | 18 | -0.724569 | -1.827767 | 0.006061 | 0.042942 |
| GOBP_LEUKOCYTE_CHEMOTAXIS | 182 | -0.473454 | -1.825278 | 0 | 0.04345 |
| GOBP_NEGATIVE_REGULATION_OF_EXTRINSIC_APOPTOTIC_SIGNALING_PATHWAY_VIA_DE | 36 | -0.61345 | -1.824918 | 0 | 0.043014 |
| GOBP_NEGATIVE_REGULATION_OF_PROTEIN_MODIFICATION_BY_SMALL_PROTEIN_CONJU | 84 | -0.532171 | -1.824395 | 0 | 0.042612 |
| GOBP_CELLULAR_RESPONSE_TO_DSRNA | 21 | -0.692576 | -1.820337 | 0 | 0.044201 |
| GOBP_POSITIVE_REGULATION_OF_FATTY_ACID_METABOLIC_PROCESS | 29 | -0.655775 | -1.819784 | 0.002114 | 0.043769 |
| GOBP_SYNAPTIC_VESICLE_TRANSPORT | 42 | -0.59058 | -1.819426 | 0.002188 | 0.043377 |
| GOBP_HEAT_GENERATION | 15 | -0.752657 | -1.817792 | 0.001972 | 0.043761 |
| GOBP_POSITIVE_REGULATION_OF_CHEMOTAXIS | 115 | -0.507048 | -1.814717 | 0 | 0.045128 |
| GOBP_SPLEEN_DEVELOPMENT | 27 | -0.64706 | -1.810646 | 0 | 0.046767 |
| GOBP_DOPAMINE_METABOLIC_PROCESS | 21 | -0.684137 | -1.809866 | 0.004348 | 0.04671 |
| GOBP_SMOOTH_MUSCLE_CELL_MIGRATION | 57 | -0.56125 | -1.809787 | 0.004329 | 0.046105 |
| GOBP_REGULATION_OF_CHROMOSOME_SEPARATION | 69 | -0.53678 | -1.807718 | 0 | 0.046688 |
| GOBP_RESPONSE_TO_REACTIVE_OXYGEN_SPECIES | 193 | -0.468372 | -1.806308 | 0 | 0.047103 |
| GOBP_RESPONSE_TO_GLUCAGON | 31 | -0.629131 | -1.799572 | 0 | 0.050762 |
| GOBP_MITOTIC_CELL_CYCLE_ARREST | 15 | -0.740306 | -1.798632 | 0 | 0.050636 |
| GOBP_PEPTIDYL_THREONINE_DEPHOSPHORYLATION | 21 | -0.693121 | -1.798333 | 0.002049 | 0.050164 |
| GOBP_METAPHASE_ANAPHASE_TRANSITION_OF_CELL_CYCLE | 62 | -0.548498 | -1.797024 | 0 | 0.0505 |
| GOBP_RESPONSE_TO_MUSCLE_ACTIVITY | 18 | -0.710435 | -1.792331 | 0.002079 | 0.053142 |
| GOBP_PEPTIDYL_ARGININE_MODIFICATION | 21 | -0.683435 | -1.792263 | 0.004202 | 0.052517 |
| GOBP_REGULATION_OF_RESPONSE_TO_INTERFERON_GAMMA | 24 | -0.638044 | -1.785187 | 0.002169 | 0.056636 |
| GOBP_PEROXISOME_PROLIFERATOR_ACTIVATED_RECEPTOR_SIGNALING_PATHWAY | 17 | -0.709231 | -1.784231 | 0.003992 | 0.056663 |
| GOBP_NEUTRAL_AMINO_ACID_TRANSPORT | 36 | -0.595351 | -1.780497 | 0.004405 | 0.058553 |
| GOBP_RESPONSE_TO_ISOQUINOLINE_ALKALOID | 20 | -0.679306 | -1.780025 | 0.002092 | 0.058046 |
| GOBP_MODULATION_OF_PROCESS_OF_OTHER_ORGANISM | 107 | -0.4929 | -1.778966 | 0 | 0.058258 |
| GOBP_EOSINOPHIL_MIGRATION | 15 | -0.739632 | -1.776646 | 0.002114 | 0.059146 |
| GOBP_MYELOID_CELL_DEVELOPMENT | 62 | -0.548025 | -1.77434 | 0 | 0.060105 |
| GOBP_EMBRYONIC_SKELETAL_SYSTEM_MORPHOGENESIS | 65 | -0.526902 | -1.773759 | 0.004376 | 0.059889 |
| GOBP_NEGATIVE_REGULATION_OF_VIRAL_PROCESS | 72 | -0.530378 | -1.770951 | 0 | 0.061283 |
| GOBP_RESPONSE_TO_PAIN | 16 | -0.725868 | -1.770746 | 0.003953 | 0.060673 |
| GOBP_CELLULAR_RESPONSE_TO_GLUCAGON_STIMULUS | 20 | -0.691903 | -1.768698 | 0.004049 | 0.061048 |
| GOBP_SPERM_CAPACITATION | 15 | -0.71901 | -1.766816 | 0.003984 | 0.061605 |
| GOBP_NEGATIVE_REGULATION_OF_MYELOID_CELL_DIFFERENTIATION | 80 | -0.515634 | -1.764487 | 0 | 0.0626 |
| GOBP_HYDROGEN_PEROXIDE_CATABOLIC_PROCESS | 28 | -0.628897 | -1.763005 | 0.005917 | 0.062946 |
| GOBP_RESPONSE_TO_LITHIUM_ION | 17 | -0.718597 | -1.758509 | 0.004338 | 0.066142 |
| GOBP_REGULATION_OF_MICROGLIAL_CELL_ACTIVATION | 15 | -0.73112 | -1.7579 | 0.004167 | 0.065964 |
| GOBP_SNRNA_METABOLIC_PROCESS | 41 | -0.585533 | -1.757846 | 0.004386 | 0.065317 |
| GOBP_REGULATION_OF_VIRAL_GENOME_REPLICATION | 73 | -0.521206 | -1.756892 | 0 | 0.065411 |
| GOBP_CHAPERONE_MEDIATED_PROTEIN_FOLDING | 60 | -0.543001 | -1.756324 | 0 | 0.065375 |
| GOBP_REGULATION_OF_PROTEIN_MODIFICATION_BY_SMALL_PROTEIN_CONJUGATION_OR | 222 | -0.444426 | -1.754788 | 0 | 0.066052 |
| GOBP_SEQUESTERING_OF_TRIGLYCERIDE | 15 | -0.699297 | -1.749771 | 0.008677 | 0.069444 |
| GOBP_POSITIVE_REGULATION_OF_WOUND_HEALING | 45 | -0.568621 | -1.747756 | 0 | 0.070429 |
| GOBP_RESPONSE_TO_FUNGUS | 31 | -0.618185 | -1.747598 | 0.002217 | 0.069869 |
| GOBP_NEGATIVE_REGULATION_OF_RESPONSE_TO_CYTOKINE_STIMULUS | 54 | -0.541124 | -1.747447 | 0 | 0.069289 |
| GOBP_DNA_REPLICATION_INDEPENDENT_NUCLEOSOME_ORGANIZATION | 51 | -0.543457 | -1.742225 | 0.006 | 0.073477 |
| GOBP_REGULATION_OF_POSITIVE_CHEMOTAXIS | 21 | -0.665368 | -1.738659 | 0.006048 | 0.075865 |
| GOBP_METANEPHROS_MORPHOGENESIS | 20 | -0.676869 | -1.736396 | 0.00616 | 0.077377 |
| GOBP_REGULATION_OF_RESPONSE_TO_CYTOKINE_STIMULUS | 160 | -0.45764 | -1.729882 | 0 | 0.082707 |
| GOBP_RESPONSE_TO_MANGANESE_ION | 17 | -0.675073 | -1.729753 | 0.008475 | 0.08208 |
| GOBP_RESPONSE_TO_LEPTIN | 19 | -0.681624 | -1.729518 | 0.003906 | 0.081464 |
| GOBP_REGULATION_OF_OSTEOCLAST_DIFFERENTIATION | 53 | -0.551713 | -1.72901 | 0.002101 | 0.081044 |
| GOBP_RESPONSE_TO_CAMP | 78 | -0.507871 | -1.728711 | 0 | 0.080574 |
| GOBP_RESPONSE_TO_EXTRACELLULAR_STIMULUS | 377 | -0.415484 | -1.728283 | 0 | 0.080323 |
| GOBP_POSITIVE_REGULATION_OF_OSTEOCLAST_DIFFERENTIATION | 22 | -0.65073 | -1.727003 | 0.003992 | 0.08056 |
| GOBP_VESICLE_BUDDING_FROM_MEMBRANE | 106 | -0.484513 | -1.723558 | 0 | 0.08286 |
| GOBP_IMMUNOGLOBULIN_PRODUCTION | 168 | -0.456857 | -1.722474 | 0 | 0.083075 |
| GOBP_POSITIVE_REGULATION_OF_MAP_KINASE_ACTIVITY | 200 | -0.442681 | -1.722289 | 0 | 0.082527 |
| GOBP_CELLULAR_RESPONSE_TO_BIOTIC_STIMULUS | 196 | -0.446298 | -1.721497 | 0 | 0.0825 |
| GOBP_MONOCYTE_CHEMOTAXIS | 44 | -0.556497 | -1.720647 | 0.00211 | 0.082525 |
| GOBP_KIDNEY_MORPHOGENESIS | 60 | -0.527851 | -1.720008 | 0.006424 | 0.082352 |
| GOBP_VASCULAR_ASSOCIATED_SMOOTH_MUSCLE_CONTRACTION | 19 | -0.669974 | -1.719583 | 0.00404 | 0.081919 |
| GOBP_CELL_CYCLE_DNA_REPLICATION | 64 | -0.525857 | -1.71874 | 0.004338 | 0.081938 |
| GOBP_POSITIVE_REGULATION_OF_EXCITATORY_POSTSYNAPTIC_POTENTIAL | 24 | -0.648167 | -1.717922 | 0.002079 | 0.082185 |
| GOBP_KILLING_OF_CELLS_IN_OTHER_ORGANISM_INVOLVED_IN_SYMBIOTIC_INTERACTION | 15 | -0.701735 | -1.717271 | 0.01217 | 0.08211 |
| GOBP_NEGATIVE_REGULATION_OF_RECEPTOR_MEDIATED_ENDOCYTOSIS | 24 | -0.636237 | -1.716906 | 0.008475 | 0.081784 |
| GOBP_CELLULAR_RESPONSE_TO_MOLECULE_OF_BACTERIAL_ORIGIN | 173 | -0.453788 | -1.713352 | 0 | 0.084176 |
| GOBP_MESONEPHROS_DEVELOPMENT | 62 | -0.514305 | -1.713127 | 0.002141 | 0.083738 |
| GOBP_REGULATION_OF_RECEPTOR_LOCALIZATION_TO_SYNAPSE | 17 | -0.686428 | -1.711709 | 0.005988 | 0.08446 |
| GOBP_CELL_CHEMOTAXIS | 242 | -0.432719 | -1.711539 | 0 | 0.08394 |
| GOBP_THYMUS_DEVELOPMENT | 38 | -0.577189 | -1.711406 | 0.002096 | 0.083344 |
| GOBP_MYELOID_CELL_DIFFERENTIATION | 376 | -0.413613 | -1.710845 | 0 | 0.083266 |
| GOBP_HYDROGEN_PEROXIDE_METABOLIC_PROCESS | 48 | -0.557646 | -1.709955 | 0.004141 | 0.083401 |
| GOBP_NUCLEOTIDE_BINDING_DOMAIN_LEUCINE_RICH_REPEAT_CONTAINING_RECEPTOR_ | 41 | -0.571077 | -1.708633 | 0.002128 | 0.083918 |
| GOBP_LEUKOCYTE_MIGRATION | 426 | -0.403437 | -1.707669 | 0 | 0.084119 |
| GOBP_RESPONSE_TO_CORTICOSTEROID | 113 | -0.469123 | -1.707485 | 0 | 0.083671 |
| GOBP_POSITIVE_REGULATION_OF_MYELOID_LEUKOCYTE_MEDIATED_IMMUNITY | 17 | -0.677202 | -1.706554 | 0.012371 | 0.08387 |
| GOBP_MITOTIC_CELL_CYCLE_CHECKPOINT | 154 | -0.452617 | -1.706323 | 0 | 0.083453 |
| GOBP_CELLULAR_RESPONSE_TO_FATTY_ACID | 28 | -0.611457 | -1.705831 | 0.010753 | 0.083216 |
| GOBP_SUBSTANTIA_NIGRA_DEVELOPMENT | 40 | -0.569213 | -1.705804 | 0.004264 | 0.08264 |
| GOBP_PHAGOCYTOSIS_RECOGNITION | 71 | -0.506189 | -1.70465 | 0.002092 | 0.083046 |
| GOBP_HOMEOSTASIS_OF_NUMBER_OF_CELLS | 235 | -0.433196 | -1.704274 | 0 | 0.08267 |
| GOBP_TAXIS | 499 | -0.399336 | -1.702208 | 0 | 0.083908 |
| GOBP_AUTOPHAGY_OF_MITOCHONDRION | 73 | -0.502785 | -1.701568 | 0.004264 | 0.083854 |
| GOBP_POSITIVE_REGULATION_OF_INFLAMMATORY_RESPONSE | 101 | -0.478798 | -1.701018 | 0 | 0.083656 |
| GOBP_RESPONSE_TO_GONADOTROPIN | 19 | -0.655267 | -1.700518 | 0.008677 | 0.08356 |
| GOBP_CHROMATIN_REMODELING_AT_CENTROMERE | 44 | -0.550962 | -1.699901 | 0.006289 | 0.083636 |
| GOBP_RIBONUCLEOSIDE_CATABOLIC_PROCESS | 21 | -0.647945 | -1.698869 | 0.007921 | 0.083917 |
| GOBP_POSITIVE_REGULATION_OF_PROTEIN_MODIFICATION_BY_SMALL_PROTEIN_CONJUG | 128 | -0.46709 | -1.696915 | 0.002217 | 0.084936 |
| GOBP_RESPONSE_TO_MOLECULE_OF_BACTERIAL_ORIGIN | 284 | -0.420656 | -1.694601 | 0 | 0.086398 |
| GOBP_BILE_ACID_AND_BILE_SALT_TRANSPORT | 20 | -0.646606 | -1.694281 | 0.017391 | 0.086105 |
| GOBP_REGULATION_OF_PROTEIN_SERINE_THREONINE_KINASE_ACTIVITY | 425 | -0.402166 | -1.693727 | 0 | 0.085944 |
| GOBP_POSITIVE_REGULATION_OF_MAPK_CASCADE | 418 | -0.403584 | -1.692885 | 0 | 0.086078 |
| GOBP_POSITIVE_REGULATION_OF_RESPONSE_TO_WOUNDING | 55 | -0.532728 | -1.692842 | 0.002203 | 0.085609 |

| | | | | | |
|---|---|---|---|---|---|
| GOBP_REGULATION_OF_ACTIN_FILAMENT_BASED_MOVEMENT | 26 | 0.484381 | 1.318864 | 0.150735 | 0.564136 |
| GOBP_NEURAL_CREST_CELL_MIGRATION | 36 | 0.453196 | 1.317516 | 0.093023 | 0.565067 |
| GOBP_POSITIVE_REGULATION_OF_NEUROTRANSMITTER_SECRETION | 16 | 0.538598 | 1.314595 | 0.133072 | 0.571656 |
| GOBP_REGULATION_OF_ANOIKIS | 23 | 0.485449 | 1.312301 | 0.127701 | 0.576204 |
| GOBP_INNER_EAR_MORPHOGENESIS | 63 | 0.403784 | 1.311936 | 0.081749 | 0.573858 |
| GOBP_REGULATION_OF_STEM_CELL_PROLIFERATION | 32 | 0.469462 | 1.309648 | 0.118644 | 0.578424 |
| GOBP_RESPONSE_TO_VITAMIN_D | 24 | 0.490736 | 1.308697 | 0.128155 | 0.57793 |
| GOBP_MELANOCYTE_DIFFERENTIATION | 16 | 0.538912 | 1.308179 | 0.152294 | 0.576112 |
| GOBP_MULTICELLULAR_ORGANISMAL_SIGNALING | 143 | 0.355813 | 1.307617 | 0.052252 | 0.57451 |
| GOBP_NEUROTRANSMITTER_REUPTAKE | 22 | 0.497036 | 1.307529 | 0.121857 | 0.571279 |
| GOBP_POSITIVE_REGULATION_OF_STRESS_FIBER_ASSEMBLY | 44 | 0.432132 | 1.307518 | 0.103053 | 0.567816 |
| GOBP_REGULATION_OF_AXONOGENESIS | 128 | 0.36319 | 1.306158 | 0.059727 | 0.569255 |
| GOBP_PALLIUM_DEVELOPMENT | 119 | 0.365818 | 1.303417 | 0.051661 | 0.575371 |
| GOBP_POSITIVE_REGULATION_OF_CELL_SUBSTRATE_ADHESION | 96 | 0.373884 | 1.302971 | 0.067179 | 0.573554 |
| GOBP_REGULATION_OF_MESENCHYMAL_CELL_PROLIFERATION | 16 | 0.54356 | 1.30131 | 0.169118 | 0.575635 |
| GOBP_FOREBRAIN_DEVELOPMENT | 262 | 0.332983 | 1.300386 | 0.014159 | 0.575278 |
| GOBP_SENSORY_PERCEPTION | 368 | 0.322055 | 1.297087 | 0.017762 | 0.583444 |
| GOBP_CEREBELLAR_CORTEX_DEVELOPMENT | 34 | 0.456521 | 1.295971 | 0.103704 | 0.583864 |
| GOBP_NEGATIVE_REGULATION_OF_NATURAL_KILLER_CELL_MEDIATED_IMMUNITY | 18 | 0.513754 | 1.293013 | 0.153257 | 0.590848 |
| GOBP_REGULATION_OF_NEURAL_PRECURSOR_CELL_PROLIFERATION | 55 | 0.403659 | 1.292873 | 0.11413 | 0.587919 |
| GOBP_PEPTIDYL_GLUTAMIC_ACID_MODIFICATION | 27 | 0.474671 | 1.292728 | 0.141988 | 0.585052 |
| GOBP_SKIN_DEVELOPMENT | 214 | 0.337686 | 1.290552 | 0.026786 | 0.589226 |
| GOBP_SYNAPTIC_VESICLE_PRIMING | 16 | 0.53572 | 1.287852 | 0.167279 | 0.595536 |
| GOBP_REGULATION_OF_CELL_SIZE | 150 | 0.349654 | 1.287229 | 0.04947 | 0.594407 |
| GOBP_PROTEIN_CONTAINING_COMPLEX_REMODELING | 17 | 0.51828 | 1.285831 | 0.1643 | 0.59584 |
| GOBP_ENDODERMAL_CELL_DIFFERENTIATION | 36 | 0.43967 | 1.284572 | 0.122905 | 0.596909 |
| GOBP_MECHANORECEPTOR_DIFFERENTIATION | 48 | 0.427425 | 1.283984 | 0.11068 | 0.595621 |
| GOBP_REGULATION_OF_ACTIN_NUCLEATION | 27 | 0.461136 | 1.283086 | 0.156673 | 0.59542 |
| GOBP_AUTONOMIC_NERVOUS_SYSTEM_DEVELOPMENT | 29 | 0.462304 | 1.282728 | 0.132673 | 0.593468 |
| GOBP_PROTEOGLYCAN_METABOLIC_PROCESS | 72 | 0.395253 | 1.282573 | 0.105751 | 0.590749 |
| GOBP_AUDITORY_RECEPTOR_CELL_DEVELOPMENT | 16 | 0.527176 | 1.281392 | 0.171322 | 0.591482 |
| GOBP_ENDOTHELIAL_CELL_PROLIFERATION | 126 | 0.360328 | 1.280531 | 0.046552 | 0.591264 |
| GOBP_STEM_CELL_PROLIFERATION | 53 | 0.409928 | 1.279853 | 0.124521 | 0.590461 |
| GOBP_APPENDAGE_DEVELOPMENT | 122 | 0.363278 | 1.27939 | 0.079279 | 0.588952 |
| GOBP_POSITIVE_REGULATION_OF_NEUROTRANSMITTER_TRANSPORT | 20 | 0.499021 | 1.279135 | 0.16632 | 0.586672 |
| GOBP_DORSAL_VENTRAL_NEURAL_TUBE_PATTERNING | 18 | 0.509967 | 1.273444 | 0.163265 | 0.603491 |
| GOBP_VENTRICULAR_SEPTUM_DEVELOPMENT | 50 | 0.417171 | 1.273153 | 0.113173 | 0.601393 |
| GOBP_MOLTING_CYCLE | 79 | 0.380765 | 1.271562 | 0.097328 | 0.603759 |
| GOBP_RETINAL_GANGLION_CELL_AXON_GUIDANCE | 15 | 0.524878 | 1.271158 | 0.176136 | 0.602028 |
| GOBP_ENDODERM_FORMATION | 43 | 0.426326 | 1.270574 | 0.12758 | 0.600967 |
| GOBP_EMBRYONIC_EYE_MORPHOGENESIS | 20 | 0.498667 | 1.269237 | 0.185771 | 0.602346 |
| GOBP_EYE_PHOTORECEPTOR_CELL_DIFFERENTIATION | 28 | 0.451953 | 1.265807 | 0.150905 | 0.611663 |
| GOBP_NEGATIVE_REGULATION_OF_SMOOTH_MUSCLE_CELL_PROLIFERATION | 37 | 0.439085 | 1.265195 | 0.15879 | 0.610647 |
| GOBP_REGULATION_OF_NERVOUS_SYSTEM_DEVELOPMENT | 331 | 0.314082 | 1.263401 | 0.031034 | 0.613904 |
| GOBP_TRANSMISSION_OF_NERVE_IMPULSE | 43 | 0.425439 | 1.26286 | 0.136194 | 0.612761 |
| GOBP_REGULATION_OF_ENDOTHELIAL_CELL_MIGRATION | 147 | 0.345858 | 1.261339 | 0.080342 | 0.615035 |
| GOBP_DENDRITIC_SPINE_MORPHOGENESIS | 55 | 0.405754 | 1.259881 | 0.130435 | 0.616999 |
| GOBP_NEGATIVE_REGULATION_OF_LIPID_METABOLIC_PROCESS | 73 | 0.38168 | 1.258766 | 0.127306 | 0.61792 |
| GOBP_GLANDULAR_EPITHELIAL_CELL_DEVELOPMENT | 17 | 0.50652 | 1.257831 | 0.181989 | 0.618166 |
| GOBP_HOMOLOGOUS_CHROMOSOME_SEGREGATION | 46 | 0.416893 | 1.255253 | 0.152416 | 0.62426 |
| GOBP_POSITIVE_REGULATION_OF_VASOCONSTRICTION | 18 | 0.508999 | 1.25487 | 0.178503 | 0.622457 |
| GOBP_INOSITOL_LIPID_MEDIATED_SIGNALING | 166 | 0.337958 | 1.253117 | 0.072897 | 0.625801 |
| GOBP_NEGATIVE_REGULATION_OF_NEURAL_PRECURSOR_CELL_PROLIFERATION | 15 | 0.525986 | 1.253104 | 0.189409 | 0.622792 |
| GOBP_BRAIN_MORPHOGENESIS | 29 | 0.454347 | 1.251875 | 0.163904 | 0.624184 |
| GOBP_DOPAMINE_TRANSPORT | 28 | 0.448023 | 1.251275 | 0.167347 | 0.62324 |
| GOBP_CELL_FATE_SPECIFICATION | 51 | 0.395287 | 1.250827 | 0.114901 | 0.62184 |
| GOBP_REGULATION_OF_PHOSPHATIDYLINOSITOL_3_KINASE_SIGNALING | 109 | 0.353706 | 1.248168 | 0.082437 | 0.628287 |
| GOBP_CELL_VOLUME_HOMEOSTASIS | 25 | 0.456027 | 1.248043 | 0.182171 | 0.625711 |
| GOBP_ADULT_LOCOMOTORY_BEHAVIOR | 42 | 0.419993 | 1.247608 | 0.164751 | 0.624373 |
| GOBP_POSITIVE_REGULATION_OF_NEUROGENESIS | 179 | 0.333627 | 1.247064 | 0.070588 | 0.623388 |
| GOBP_INOSITOL_PHOSPHATE_METABOLIC_PROCESS | 64 | 0.382989 | 1.247048 | 0.112523 | 0.620539 |
| GOBP_SISTER_CHROMATID_COHESION | 59 | 0.390207 | 1.245389 | 0.135472 | 0.623535 |
| GOBP_DICARBOXYLIC_ACID_TRANSPORT | 74 | 0.37537 | 1.242528 | 0.115686 | 0.630901 |
| GOBP_NEURON_PROJECTION_REGENERATION | 48 | 0.41077 | 1.242479 | 0.143396 | 0.628174 |
| GOBP_SECONDARY_METABOLIC_PROCESS | 35 | 0.432335 | 1.242325 | 0.178637 | 0.625866 |
| GOBP_REGULATION_OF_ANATOMICAL_STRUCTURE_SIZE | 404 | 0.303579 | 1.24167 | 0.024712 | 0.625331 |
| GOBP_ENDODERM_DEVELOPMENT | 58 | 0.399651 | 1.241658 | 0.12233 | 0.622533 |
| GOBP_APICAL_JUNCTION_ASSEMBLY | 63 | 0.390876 | 1.241471 | 0.118095 | 0.620356 |
| GOBP_CELL_SUBSTRATE_ADHESION | 297 | 0.312832 | 1.239794 | 0.036851 | 0.623426 |
| GOBP_RELAXATION_OF_MUSCLE | 23 | 0.483876 | 1.239152 | 0.193424 | 0.622857 |
| GOBP_NEGATIVE_REGULATION_OF_PEPTIDE_SECRETION | 52 | 0.397883 | 1.237397 | 0.144465 | 0.626213 |
| GOBP_POSITIVE_REGULATION_OF_ERBB_SIGNALING_PATHWAY | 37 | 0.420179 | 1.236956 | 0.147002 | 0.624939 |
| GOBP_PIGMENTATION | 75 | 0.369778 | 1.236381 | 0.128492 | 0.624109 |
| GOBP_SECRETION_BY_TISSUE | 25 | 0.458679 | 1.236007 | 0.176817 | 0.622661 |
| GOBP_NEURON_PROJECTION_ORGANIZATION | 82 | 0.366984 | 1.234658 | 0.121569 | 0.624495 |
| GOBP_SENSORY_ORGAN_MORPHOGENESIS | 170 | 0.330964 | 1.233814 | 0.078153 | 0.624677 |
| GOBP_LEUKOCYTE_MEDIATED_CYTOTOXICITY | 102 | 0.354998 | 1.232298 | 0.105691 | 0.627139 |
| GOBP_SYNAPTIC_TRANSMISSION_GLUTAMATERGIC | 59 | 0.389846 | 1.23166 | 0.168279 | 0.626544 |
| GOBP_SENSORY_PERCEPTION_OF_LIGHT_STIMULUS | 129 | 0.339258 | 1.231077 | 0.086176 | 0.625816 |
| GOBP_NUCLEAR_TRANSCRIBED_MRNA_CATABOLIC_PROCESS_NONSENSE_MEDIATED_ | 120 | 0.340792 | 1.228741 | 0.089767 | 0.631375 |
| GOBP_PHOSPHATIDYLINOSITOL_3_KINASE_SIGNALING | 131 | 0.342237 | 1.227512 | 0.112727 | 0.633191 |
| GOBP_POSITIVE_REGULATION_OF_CARTILAGE_DEVELOPMENT | 26 | 0.447465 | 1.226328 | 0.191095 | 0.634926 |
| GOBP_DIGESTIVE_TRACT_MORPHOGENESIS | 27 | 0.43705 | 1.223375 | 0.189243 | 0.642227 |
| GOBP_ENDOTHELIAL_CELL_MIGRATION | 189 | 0.324387 | 1.22248 | 0.08813 | 0.642697 |
| GOBP_ACTIN_FILAMENT_BASED_MOVEMENT | 109 | 0.35155 | 1.222041 | 0.116906 | 0.641483 |
| GOBP_NEGATIVE_REGULATION_OF_MRNA_METABOLIC_PROCESS | 76 | 0.368915 | 1.221757 | 0.140511 | 0.639797 |
| GOBP_SPERM_EGG_RECOGNITION | 39 | 0.414442 | 1.220971 | 0.178641 | 0.639967 |
| GOBP_TRANSFORMING_GROWTH_FACTOR_BETA_PRODUCTION | 38 | 0.412128 | 1.220558 | 0.182879 | 0.638844 |
| GOBP_SOMITE_DEVELOPMENT | 62 | 0.377135 | 1.220416 | 0.144712 | 0.636713 |
| GOBP_RETROGRADE_AXONAL_TRANSPORT | 18 | 0.483579 | 1.219834 | 0.224371 | 0.636123 |
| GOBP_CELL_AGGREGATION | 15 | 0.501281 | 1.21963 | 0.222445 | 0.634133 |
| GOBP_ACTIVATION_OF_PROTEIN_KINASE_B_ACTIVITY | 27 | 0.447802 | 1.219591 | 0.188192 | 0.631676 |
| GOBP_EMBRYONIC_AXIS_SPECIFICATION | 27 | 0.451398 | 1.219297 | 0.184158 | 0.630151 |
| GOBP_REGULATION_OF_DENDRITIC_SPINE_MORPHOGENESIS | 40 | 0.407614 | 1.218632 | 0.17451 | 0.629853 |
| GOBP_CHROMOSOME_ORGANIZATION_INVOLVED_IN_MEIOTIC_CELL_CYCLE | 52 | 0.392087 | 1.217701 | 0.173432 | 0.63058 |
| GOBP_MESENCHYME_MORPHOGENESIS | 35 | 0.417609 | 1.217296 | 0.18677 | 0.629439 |
| GOBP_ESTABLISHMENT_OF_PROTEIN_LOCALIZATION_TO_ENDOPLASMIC_RETICULUM | 116 | 0.343832 | 1.215746 | 0.124283 | 0.632265 |
| GOBP_BODY_FLUID_SECRETION | 68 | 0.370378 | 1.21452 | 0.144712 | 0.633965 |
| GOBP_EPIDERMIS_DEVELOPMENT | 243 | 0.316093 | 1.213761 | 0.085284 | 0.634159 |
| GOBP_PEPTIDYL_PROLINE_HYDROXYLATION | 15 | 0.50529 | 1.213331 | 0.234 | 0.633217 |
| GOBP_MATERNAL_PLACENTA_DEVELOPMENT | 22 | 0.468421 | 1.211938 | 0.218045 | 0.63564 |
| GOBP_FORMATION_OF_PRIMARY_GERM_LAYER | 88 | 0.353668 | 1.21109 | 0.140713 | 0.636162 |
| GOBP_EAR_MORPHOGENESIS | 80 | 0.360564 | 1.209645 | 0.143636 | 0.638678 |
| GOBP_NEURAL_TUBE_DEVELOPMENT | 126 | 0.33686 | 1.209403 | 0.131119 | 0.637099 |
| GOBP_REGULATION_OF_NEURONAL_SYNAPTIC_PLASTICITY | 34 | 0.422423 | 1.209352 | 0.206827 | 0.634793 |
| GOBP_NEGATIVE_REGULATION_OF_LEUKOCYTE_MEDIATED_CYTOTOXICITY | 21 | 0.472045 | 1.209031 | 0.207767 | 0.633466 |
| GOBP_REGULATION_OF_LONG_TERM_SYNAPTIC_POTENTIATION | 35 | 0.417936 | 1.208269 | 0.198516 | 0.633606 |
| GOBP_LENS_DEVELOPMENT_IN_CAMERA_TYPE_EYE | 53 | 0.382585 | 1.207886 | 0.172662 | 0.632487 |
| GOBP_REGULATION_OF_DNA_DAMAGE_RESPONSE_SIGNAL_TRANSDUCTION_BY_P53_C | 33 | 0.427612 | 1.207661 | 0.19434 | 0.630858 |
| GOBP_MESENCHYMAL_CELL_PROLIFERATION | 23 | 0.463369 | 1.207611 | 0.234834 | 0.628635 |
| GOBP_AMINE_TRANSPORT | 61 | 0.377597 | 1.206742 | 0.170455 | 0.629122 |
| GOBP_BEHAVIOR | 383 | 0.29879 | 1.205352 | 0.058407 | 0.631622 |
| GOBP_CENTRAL_NERVOUS_SYSTEM_NEURON_AXONOGENESIS | 32 | 0.429392 | 1.204909 | 0.201536 | 0.630756 |
| GOBP_CELLULAR_COMPONENT_DISASSEMBLY_INVOLVED_IN_EXECUTION_PHASE_OF_A | 30 | 0.430586 | 1.204759 | 0.204409 | 0.628902 |
| GOBP_LONG_TERM_SYNAPTIC_POTENTIATION | 61 | 0.380554 | 1.203953 | 0.185393 | 0.629306 |
| GOBP_POSTSYNAPSE_ORGANIZATION | 135 | 0.329559 | 1.203325 | 0.123412 | 0.62914 |
| GOBP_FACE_DEVELOPMENT | 37 | 0.414343 | 1.200226 | 0.202578 | 0.637428 |
| GOBP_OSTEOBLAST_PROLIFERATION | 25 | 0.437922 | 1.200219 | 0.220532 | 0.635085 |
| GOBP_ATP_TRANSPORT | 15 | 0.506094 | 1.199239 | 0.229084 | 0.636237 |
| GOBP_REGULATION_OF_MUSCLE_CONTRACTION | 106 | 0.340847 | 1.199227 | 0.131059 | 0.633948 |
| GOBP_REGULATION_OF_SYNAPTIC_TRANSMISSION_GLUTAMATERGIC | 44 | 0.399824 | 1.19922 | 0.202729 | 0.631656 |
| GOBP_RESPONSE_TO_EPIDERMAL_GROWTH_FACTOR | 40 | 0.402704 | 1.199177 | 0.188867 | 0.629477 |
| GOBP_AXON_DEVELOPMENT | 405 | 0.297504 | 1.197726 | 0.061856 | 0.632148 |
| GOBP_REGULATION_OF_CELL_SHAPE | 135 | 0.332174 | 1.197388 | 0.134892 | 0.631024 |
| GOBP_AMEBOIDAL_TYPE_CELL_MIGRATION | 351 | 0.300231 | 1.196835 | 0.061962 | 0.630591 |
| GOBP_FEAR_RESPONSE | 24 | 0.454873 | 1.196533 | 0.238185 | 0.629348 |
| GOBP_POSITIVE_REGULATION_OF_SYNAPTIC_TRANSMISSION | 105 | 0.348645 | 1.195158 | 0.150362 | 0.631806 |
| GOBP_HEART_VALVE_DEVELOPMENT | 48 | 0.393324 | 1.194138 | 0.201149 | 0.633147 |
| GOBP_OLIGODENDROCYTE_DIFFERENTIATION | 74 | 0.358047 | 1.192865 | 0.173913 | 0.635056 |
| GOBP_SKELETAL_MUSCLE_ADAPTATION | 16 | 0.489411 | 1.192735 | 0.238281 | 0.633224 |
| GOBP_POSITIVE_REGULATION_OF_STEROL_TRANSPORT | 23 | 0.45453 | 1.192231 | 0.236893 | 0.632663 |
| GOBP_LEARNING | 96 | 0.342212 | 1.190283 | 0.16185 | 0.637032 |
| GOBP_POSITIVE_REGULATION_OF_PROTEIN_LOCALIZATION_TO_CELL_PERIPHERY | 57 | 0.376208 | 1.190244 | 0.173664 | 0.634942 |
| GOBP_NEGATIVE_REGULATION_OF_CELL_JUNCTION_ASSEMBLY | 26 | 0.440206 | 1.189187 | 0.217221 | 0.636358 |
| GOBP_REGULATION_OF_INTEGRIN_ACTIVATION | 15 | 0.491844 | 1.188706 | 0.246988 | 0.635777 |
| GOBP_POSITIVE_T_CELL_SELECTION | 34 | 0.414257 | 1.188608 | 0.218412 | 0.633908 |
| GOBP_POSITIVE_REGULATION_OF_GLYCOPROTEIN_METABOLIC_PROCESS | 22 | 0.452213 | 1.188067 | 0.234592 | 0.633458 |
| GOBP_SOMITOGENESIS | 49 | 0.386442 | 1.186889 | 0.193613 | 0.635237 |
| GOBP_STEM_CELL_DIFFERENTIATION | 202 | 0.312781 | 1.186241 | 0.106164 | 0.635122 |
| GOBP_APPENDAGE_MORPHOGENESIS | 101 | 0.345886 | 1.186099 | 0.157407 | 0.633442 |
| GOBP_REGULATION_OF_CARDIAC_MUSCLE_CELL_PROLIFERATION | 35 | 0.415628 | 1.185942 | 0.201117 | 0.631857 |
| GOBP_NEGATIVE_REGULATION_OF_VASCULATURE_DEVELOPMENT | 79 | 0.358709 | 1.185313 | 0.194192 | 0.631846 |
| GOBP_CELL_JUNCTION_DISASSEMBLY | 18 | 0.472104 | 1.18518 | 0.254579 | 0.630157 |
| GOBP_MAMMARY_GLAND_EPITHELIAL_CELL_PROLIFERATION | 26 | 0.43555 | 1.184498 | 0.243243 | 0.630151 |
| GOBP_SODIUM_ION_TRANSMEMBRANE_TRANSPORT | 119 | 0.333477 | 1.183096 | 0.146903 | 0.632609 |
| GOBP_DOPAMINE_SECRETION | 18 | 0.467403 | 1.182974 | 0.246654 | 0.630926 |
| GOBP_REGULATION_OF_KERATINOCYTE_PROLIFERATION | 27 | 0.435541 | 1.182821 | 0.221359 | 0.629352 |
| GOBP_REGULATION_OF_SYNAPSE_ASSEMBLY | 64 | 0.367135 | 1.182611 | 0.203884 | 0.627969 |
| GOBP_METANEPHRIC_NEPHRON_DEVELOPMENT | 23 | 0.447745 | 1.180402 | 0.2607 | 0.633064 |
| GOBP_FATTY_ACID_TRANSPORT | 114 | 0.334968 | 1.179406 | 0.16171 | 0.634303 |
| GOBP_REGULATION_OF_BLOOD_CIRCULATION | 197 | 0.31113 | 1.17864 | 0.116071 | 0.634873 |
| GOBP_XENOBIOTIC_TRANSPORT | 37 | 0.402178 | 1.178403 | 0.229091 | 0.633599 |
| GOBP_POSITIVE_REGULATION_OF_INSULIN_SECRETION | 51 | 0.378953 | 1.178203 | 0.193246 | 0.632202 |
| GOBP_EMBRYONIC_DIGIT_MORPHOGENESIS | 41 | 0.39504 | 1.177573 | 0.217391 | 0.632309 |
| GOBP_GLUTATHIONE_DERIVATIVE_METABOLIC_PROCESS | 17 | 0.477514 | 1.176767 | 0.257253 | 0.632882 |
| GOBP_T_CELL_MEDIATED_CYTOTOXICITY | 44 | 0.387665 | 1.175753 | 0.209765 | 0.63422 |
| GOBP_NEGATIVE_REGULATION_OF_LOCOMOTION | 264 | 0.296296 | 1.174549 | 0.111732 | 0.636172 |

| | | | | | |
|---|---|---|---|---|---|
| GOBP_REGIONALIZATION | 225 | -0.427112 | -1.692165 | 0 | 0.0856 |
| GOBP_POSITIVE_REGULATION_OF_TUMOR_NECROSIS_FACTOR_SUPERFAMILY_CYTOKINE_ | 77 | -0.497657 | -1.691334 | 0 | 0.085691 |
| GOBP_PORPHYRIN_CONTAINING_COMPOUND_METABOLIC_PROCESS | 36 | -0.573309 | -1.691288 | 0.002141 | 0.085173 |
| GOBP_POSITIVE_REGULATION_OF_COAGULATION | 20 | -0.653526 | -1.689474 | 0.008351 | 0.086278 |
| GOBP_KILLING_OF_CELLS_OF_OTHER_ORGANISM | 37 | -0.576111 | -1.685949 | 0.006652 | 0.088466 |
| GOBP_HUMORAL_IMMUNE_RESPONSE_MEDIATED_BY_CIRCULATING_IMMUNOGLOBULIN | 122 | -0.461764 | -1.684703 | 0 | 0.088966 |
| GOBP_REGULATION_OF_ERYTHROCYTE_DIFFERENTIATION | 47 | -0.540539 | -1.683561 | 0.004283 | 0.089541 |
| GOBP_POSITIVE_REGULATION_OF_CELL_ADHESION_MEDIATED_BY_INTEGRIN | 18 | -0.662442 | -1.683463 | 0.0125 | 0.089053 |
| GOBP_REGULATION_OF_NUCLEAR_DIVISION | 122 | -0.458605 | -1.683236 | 0 | 0.088739 |
| GOBP_RESPONSE_TO_STARVATION | 173 | -0.440353 | -1.682555 | 0 | 0.088718 |
| GOBP_POSITIVE_REGULATION_OF_B_CELL_ACTIVATION | 126 | -0.455959 | -1.682493 | 0.002114 | 0.088255 |
| GOBP_FAT_CELL_DIFFERENTIATION | 181 | -0.438557 | -1.682455 | 0 | 0.087759 |
| GOBP_CHAPERONE_COFACTOR_DEPENDENT_PROTEIN_REFOLDING | 34 | -0.574458 | -1.680424 | 0.009634 | 0.089228 |
| GOBP_POSITIVE_REGULATION_OF_LIPID_STORAGE | 20 | -0.64681 | -1.679859 | 0.014403 | 0.089206 |
| GOBP_P38MAPK_CASCADE | 39 | -0.567706 | -1.679754 | 0.00625 | 0.088789 |
| GOBP_POSITIVE_REGULATION_OF_TYPE_I_INTERFERON_PRODUCTION | 75 | -0.496965 | -1.673084 | 0.006637 | 0.094513 |
| GOBP_SIGNAL_TRANSDUCTION_INVOLVED_IN_CELL_CYCLE_CHECKPOINT | 71 | -0.493902 | -1.672866 | 0.006637 | 0.094181 |
| GOBP_CATECHOL_CONTAINING_COMPOUND_METABOLIC_PROCESS | 30 | -0.597661 | -1.668987 | 0.002198 | 0.097413 |
| GOBP_PRODUCTION_OF_MOLECULAR_MEDIATOR_OF_IMMUNE_RESPONSE | 241 | -0.421728 | -1.668482 | 0 | 0.097343 |
| GOBP_INCLUSION_BODY_ASSEMBLY | 20 | -0.647195 | -1.667038 | 0.014989 | 0.098124 |
| GOBP_REGULATION_OF_CELL_DIVISION | 138 | -0.448511 | -1.66677 | 0 | 0.097768 |
| GOBP_REGULATION_OF_CHROMOSOME_SEGREGATION | 84 | -0.486245 | -1.666559 | 0.002155 | 0.09739 |
| GOBP_POSITIVE_REGULATION_OF_ATPASE_ACTIVITY | 52 | -0.528398 | -1.664148 | 0.002146 | 0.099116 |
| GOBP_CELLULAR_RESPONSE_TO_EXTRACELLULAR_STIMULUS | 204 | -0.427629 | -1.662774 | 0 | 0.100017 |
| GOBP_RETROGRADE_VESICLE_MEDIATED_TRANSPORT_GOLGI_TO_ENDOPLASMIC_RETICU | 84 | -0.479344 | -1.659849 | 0.002288 | 0.102365 |
| GOBP_ACTIVATION_OF_MAPK_ACTIVITY | 135 | -0.450083 | -1.656187 | 0 | 0.105489 |
| GOBP_MITOTIC_NUCLEAR_DIVISION | 284 | -0.409028 | -1.654171 | 0 | 0.107035 |
| GOBP_TEMPERATURE_HOMEOSTASIS | 143 | -0.443941 | -1.653762 | 0 | 0.106716 |
| GOBP_POSITIVE_REGULATION_OF_UBIQUITIN_PROTEIN_TRANSFERASE_ACTIVITY | 30 | -0.576539 | -1.652287 | 0.010101 | 0.107678 |
| GOBP_CENTROMERE_COMPLEX_ASSEMBLY | 53 | -0.519169 | -1.652115 | 0.008368 | 0.107289 |
| GOBP_CD4_POSITIVE_ALPHA_BETA_T_CELL_CYTOKINE_PRODUCTION | 15 | -0.684461 | -1.651829 | 0.012072 | 0.106964 |
| GOBP_REGULATION_OF_VIRAL_LIFE_CYCLE | 127 | -0.445161 | -1.6512 | 0 | 0.107011 |
| GOBP_POSITIVE_REGULATION_OF_PROTEIN_SERINE_THREONINE_KINASE_ACTIVITY | 270 | -0.409195 | -1.650505 | 0 | 0.107203 |
| GOBP_EMBRYONIC_SKELETAL_SYSTEM_DEVELOPMENT | 82 | -0.476583 | -1.649906 | 0.008565 | 0.107215 |
| GOBP_NEGATIVE_REGULATION_OF_SMOOTH_MUSCLE_CELL_MIGRATION | 17 | -0.656416 | -1.649895 | 0.016667 | 0.106679 |
| GOBP_REGULATION_OF_SPINDLE_ASSEMBLY | 25 | -0.602969 | -1.647273 | 0.010352 | 0.108988 |
| GOBP_REGULATION_OF_BIOLOGICAL_PROCESS_INVOLVED_IN_SYMBIOTIC_INTERACTION | 173 | -0.434452 | -1.646648 | 0 | 0.108989 |
| GOBP_MULTI_MULTICELLULAR_ORGANISM_PROCESS | 151 | -0.439641 | -1.646171 | 0 | 0.10888 |
| GOBP_NEGATIVE_REGULATION_OF_PROTEIN_DEPHOSPHORYLATION | 60 | -0.504654 | -1.645216 | 0.006316 | 0.10918 |
| GOBP_REGULATION_OF_GLIAL_CELL_MIGRATION | 17 | -0.655591 | -1.645109 | 0.010593 | 0.108746 |
| GOBP_POSITIVE_REGULATION_OF_PRI_MIRNA_TRANSCRIPTION_BY_RNA_POLYMERASE_II | 39 | -0.548065 | -1.644996 | 0.002092 | 0.10834 |
| GOBP_NEGATIVE_REGULATION_OF_METAPHASE_ANAPHASE_TRANSITION_OF_CELL_CYCL | 40 | -0.551521 | -1.644018 | 0.013514 | 0.10876 |
| GOBP_REGULATION_OF_MYELOID_LEUKOCYTE_DIFFERENTIATION | 99 | -0.46215 | -1.643145 | 0.00431 | 0.109076 |
| GOBP_REGULATION_OF_VENTRICULAR_CARDIAC_MUSCLE_CELL_MEMBRANE_REPOLARIZ | 16 | -0.674378 | -1.642113 | 0.016807 | 0.109851 |
| GOBP_REGULATION_OF_CHEMOTAXIS | 182 | -0.430703 | -1.641613 | 0 | 0.10987 |
| GOBP_POSITIVE_REGULATION_OF_NUCLEOTIDE_BIOSYNTHETIC_PROCESS | 16 | -0.654569 | -1.641092 | 0.016461 | 0.109902 |
| GOBP_EXTRINSIC_APOPTOTIC_SIGNALING_PATHWAY | 192 | -0.425162 | -1.640442 | 0 | 0.110051 |
| GOBP_RESPONSE_TO_TUMOR_NECROSIS_FACTOR | 267 | -0.408353 | -1.639782 | 0 | 0.110087 |
| GOBP_ACUTE_INFLAMMATORY_RESPONSE | 81 | -0.473811 | -1.639203 | 0.004454 | 0.110185 |
| GOBP_ORGANIC_ANION_TRANSPORT | 278 | -0.407519 | -1.638326 | 0 | 0.110507 |
| GOBP_PATTERN_SPECIFICATION_PROCESS | 299 | -0.401213 | -1.638227 | 0 | 0.110036 |
| GOBP_REGULATION_OF_ORGANIC_ACID_TRANSPORT | 48 | -0.527712 | -1.637262 | 0.012739 | 0.110451 |
| GOBP_NUCLEOSIDE_CATABOLIC_PROCESS | 30 | -0.579722 | -1.634585 | 0.012474 | 0.112693 |
| GOBP_POSITIVE_REGULATION_OF_VASCULAR_ENDOTHELIAL_GROWTH_FACTOR_PRODUC | 21 | -0.643292 | -1.633886 | 0.008639 | 0.112869 |
| GOBP_IMPORT_ACROSS_PLASMA_MEMBRANE | 122 | -0.451745 | -1.633222 | 0 | 0.112921 |
| GOBP_RESPONSE_TO_COPPER_ION | 30 | -0.571771 | -1.633007 | 0.016985 | 0.112698 |
| GOBP_REGULATION_OF_ACTIVIN_RECEPTOR_SIGNALING_PATHWAY | 19 | -0.635451 | -1.632723 | 0.012397 | 0.11244 |
| GOBP_APOPTOTIC_PROCESS_INVOLVED_IN_MORPHOGENESIS | 17 | -0.658465 | -1.632226 | 0.020161 | 0.112334 |
| GOBP_REGULATION_OF_B_CELL_ACTIVATION | 168 | -0.42964 | -1.631972 | 0 | 0.112007 |
| GOBP_MITOTIC_G1_S_TRANSITION_CHECKPOINT | 60 | -0.508373 | -1.631545 | 0.002315 | 0.111882 |
| GOBP_DE_NOVO_PROTEIN_FOLDING | 43 | -0.533359 | -1.631398 | 0.008602 | 0.111523 |
| GOBP_GLUCOSE_IMPORT | 57 | -0.508019 | -1.631397 | 0.01217 | 0.111012 |
| GOBP_POSITIVE_REGULATION_OF_GLUCOSE_TRANSMEMBRANE_TRANSPORT | 35 | -0.549726 | -1.628761 | 0.012295 | 0.113221 |
| GOBP_NEGATIVE_REGULATION_OF_PROTEIN_SERINE_THREONINE_KINASE_ACTIVITY | 121 | -0.451728 | -1.628732 | 0.002101 | 0.112722 |
| GOBP_RESPONSE_TO_TOPOLOGICALLY_INCORRECT_PROTEIN | 190 | -0.419933 | -1.628514 | 0 | 0.11246 |
| GOBP_REGULATION_OF_MEMBRANE_REPOLARIZATION | 25 | -0.594667 | -1.628467 | 0.015453 | 0.112005 |
| GOBP_POSITIVE_REGULATION_OF_ANION_TRANSMEMBRANE_TRANSPORT | 49 | -0.524443 | -1.627536 | 0.006224 | 0.112417 |
| GOBP_NCRNA_3_END_PROCESSING | 44 | -0.529323 | -1.62695 | 0.008584 | 0.112502 |
| GOBP_POSITIVE_REGULATION_OF_ORGANIC_ACID_TRANSPORT | 30 | -0.587575 | -1.625617 | 0.014199 | 0.113377 |
| GOBP_POSITIVE_REGULATION_OF_INTERFERON_BETA_PRODUCTION | 32 | -0.573533 | -1.625554 | 0.010893 | 0.112912 |
| GOBP_CELLULAR_RESPONSE_TO_PEPTIDE | 329 | -0.400191 | -1.625352 | 0 | 0.112577 |
| GOBP_PYRIMIDINE_NUCLEOSIDE_CATABOLIC_PROCESS | 20 | -0.62696 | -1.624968 | 0.010246 | 0.112449 |
| GOBP_REGULATION_OF_LIPID_LOCALIZATION | 133 | -0.441772 | -1.624846 | 0.0025 | 0.112057 |
| GOBP_REGULATION_OF_PHAGOCYTOSIS | 83 | -0.467947 | -1.624828 | 0.008421 | 0.111584 |
| GOBP_RESPONSE_TO_CAFFEINE | 15 | -0.666487 | -1.623636 | 0.017822 | 0.112527 |
| GOBP_SYNAPTIC_VESICLE_LOCALIZATION | 51 | -0.517513 | -1.623075 | 0.002083 | 0.112605 |
| GOBP_ACTIVIN_RECEPTOR_SIGNALING_PATHWAY | 33 | -0.567451 | -1.622002 | 0.016913 | 0.113197 |
| GOBP_NEGATIVE_REGULATION_OF_PROTEIN_MODIFICATION_PROCESS | 484 | -0.382377 | -1.621982 | 0 | 0.112728 |
| GOBP_REGULATION_OF_UBIQUITIN_PROTEIN_TRANSFERASE_ACTIVITY | 49 | -0.51641 | -1.621969 | 0.006224 | 0.112249 |
| GOBP_CHROMOSOME_SEPARATION | 90 | -0.473262 | -1.621435 | 0 | 0.112358 |
| GOBP_POSITIVE_REGULATION_OF_GLUCOSE_IMPORT | 29 | -0.570214 | -1.620092 | 0.022449 | 0.113288 |
| GOBP_GLOMERULAR_EPITHELIAL_CELL_DIFFERENTIATION | 15 | -0.670105 | -1.61989 | 0.0125 | 0.113056 |
| GOBP_VENTRICULAR_CARDIAC_MUSCLE_CELL_MEMBRANE_REPOLARIZATION | 18 | -0.644454 | -1.618574 | 0.022088 | 0.113845 |
| GOBP_AMYLOID_BETA_CLEARANCE | 34 | -0.554251 | -1.618327 | 0.020964 | 0.113632 |
| GOBP_CELLULAR_RESPONSE_TO_COPPER_ION | 17 | -0.638811 | -1.61797 | 0.012422 | 0.113654 |
| GOBP_ENERGY_HOMEOSTASIS | 32 | -0.560562 | -1.617586 | 0.006148 | 0.113568 |
| GOBP_LIPOPOLYSACCHARIDE_MEDIATED_SIGNALING_PATHWAY | 53 | -0.505379 | -1.615658 | 0.004032 | 0.115202 |
| GOBP_CELL_MATURATION | 125 | -0.43628 | -1.615582 | 0 | 0.114776 |
| GOBP_VESICLE_TARGETING | 91 | -0.454351 | -1.613317 | 0.006211 | 0.116759 |
| GOBP_B_CELL_MEDIATED_IMMUNITY | 190 | -0.417717 | -1.61292 | 0 | 0.116656 |
| GOBP_RESPONSE_TO_LEAD_ION | 19 | -0.61559 | -1.612201 | 0.014614 | 0.117009 |
| GOBP_RESPONSE_TO_OXYGEN_LEVELS | 324 | -0.395043 | -1.611644 | 0 | 0.117037 |
| GOBP_POSITIVE_REGULATION_OF_RESPONSE_TO_EXTERNAL_STIMULUS | 419 | -0.384625 | -1.611499 | 0 | 0.11669 |
| GOBP_NEGATIVE_REGULATION_OF_PROTEIN_BINDING | 82 | -0.465266 | -1.611484 | 0.012448 | 0.116224 |
| GOBP_AUTOPHAGOSOME_MATURATION | 40 | -0.532137 | -1.610916 | 0.010081 | 0.116318 |
| GOBP_NEGATIVE_REGULATION_OF_PROTEIN_LOCALIZATION_TO_MEMBRANE | 24 | -0.593513 | -1.610889 | 0.014894 | 0.115883 |
| GOBP_POSITIVE_REGULATION_OF_CELL_DIVISION | 61 | -0.495016 | -1.610693 | 0.006329 | 0.115615 |
| GOBP_FC_RECEPTOR_MEDIATED_STIMULATORY_SIGNALING_PATHWAY | 134 | -0.435999 | -1.610635 | 0 | 0.115187 |
| GOBP_MEMBRANE_INVAGINATION | 120 | -0.440534 | -1.608811 | 0 | 0.116806 |
| GOBP_B_CELL_ACTIVATION | 279 | -0.394177 | -1.607986 | 0 | 0.117353 |
| GOBP_ACUTE_PHASE_RESPONSE | 30 | -0.555116 | -1.607513 | 0.019737 | 0.11741 |
| GOBP_NEGATIVE_REGULATION_OF_REGULATED_SECRETORY_PATHWAY | 19 | -0.623956 | -1.607304 | 0.022917 | 0.117212 |
| GOBP_REGULATION_OF_RECEPTOR_MEDIATED_ENDOCYTOSIS | 81 | -0.462867 | -1.607134 | 0.002179 | 0.11697 |
| GOBP_RESPONSE_TO_DEXAMETHASONE | 31 | -0.555428 | -1.606741 | 0.004255 | 0.11686 |
| GOBP_SKELETAL_SYSTEM_MORPHOGENESIS | 158 | -0.428663 | -1.60637 | 0 | 0.116763 |
| GOBP_REGULATION_OF_CELL_CYCLE_ARREST | 95 | -0.453516 | -1.60435 | 0.002058 | 0.11856 |
| GOBP_ORGANIC_ACID_TRANSMEMBRANE_TRANSPORT | 121 | -0.439939 | -1.603339 | 0 | 0.119276 |
| GOBP_NUCLEOBASE_CONTAINING_SMALL_MOLECULE_CATABOLIC_PROCESS | 45 | -0.516718 | -1.603088 | 0.010395 | 0.119135 |
| GOBP_RESPONSE_TO_INTERLEUKIN_1 | 175 | -0.418568 | -1.60159 | 0 | 0.120224 |
| GOBP_PROTEIN_ADP_RIBOSYLATION | 32 | -0.555283 | -1.60118 | 0.020704 | 0.120284 |
| GOBP_ENTRY_INTO_HOST | 134 | -0.433857 | -1.599405 | 0 | 0.121642 |
| GOBP_REGULATION_OF_GLUCOSE_TRANSMEMBRANE_TRANSPORT | 60 | -0.493443 | -1.59825 | 0.010965 | 0.122542 |
| GOBP_SULFATION | 17 | -0.642663 | -1.596944 | 0.01232 | 0.12352 |
| GOBP_POSITIVE_REGULATION_OF_NEUTROPHIL_MIGRATION | 27 | -0.581708 | -1.596649 | 0.022965 | 0.123396 |
| GOBP_NEGATIVE_REGULATION_OF_APOPTOTIC_SIGNALING_PATHWAY | 195 | -0.409761 | -1.595983 | 0 | 0.123662 |
| GOBP_NEGATIVE_REGULATION_OF_MYELOID_LEUKOCYTE_DIFFERENTIATION | 41 | -0.526162 | -1.595962 | 0.012195 | 0.123226 |
| GOBP_REGULATION_OF_RECEPTOR_SIGNALING_PATHWAY_VIA_STAT | 72 | -0.480768 | -1.595962 | 0.012793 | 0.12277 |
| GOBP_REGULATION_OF_MITOTIC_SISTER_CHROMATID_SEGREGATION | 45 | -0.515723 | -1.595789 | 0.008547 | 0.122492 |
| GOBP_RESPONSE_TO_OSMOTIC_STRESS | 70 | -0.477144 | -1.595199 | 0.004464 | 0.122874 |
| GOBP_POSITIVE_REGULATION_OF_DNA_BINDING | 46 | -0.51266 | -1.595177 | 0.010246 | 0.122447 |
| GOBP_REGULATION_OF_LIPID_STORAGE | 43 | -0.522195 | -1.595177 | 0.019724 | 0.122001 |
| GOBP_MODULATION_BY_HOST_OF_SYMBIONT_PROCESS | 55 | -0.492471 | -1.593713 | 0.004098 | 0.122979 |
| GOBP_CELLULAR_RESPONSE_TO_PEPTIDE_HORMONE_STIMULUS | 273 | -0.39887 | -1.593593 | 0 | 0.122655 |
| GOBP_DEFENSE_RESPONSE_TO_GRAM_POSITIVE_BACTERIUM | 62 | -0.490853 | -1.592628 | 0.008602 | 0.123333 |
| GOBP_POSITIVE_REGULATION_OF_LIPID_LOCALIZATION | 86 | -0.463597 | -1.592204 | 0.006593 | 0.123357 |
| GOBP_INACTIVATION_OF_MAPK_ACTIVITY | 22 | -0.614955 | -1.591736 | 0.012821 | 0.123449 |
| GOBP_CELL_CYCLE_ARREST | 209 | -0.407525 | -1.591074 | 0 | 0.123721 |
| GOBP_VESICLE_TARGETING_TO_FROM_OR_WITHIN_GOLGI | 71 | -0.47275 | -1.590822 | 0.00616 | 0.12354 |
| GOBP_CELL_FATE_COMMITMENT_INVOLVED_IN_FORMATION_OF_PRIMARY_GERM_LAYER | 19 | -0.610128 | -1.590109 | 0.027888 | 0.123889 |
| GOBP_PROXIMAL_DISTAL_PATTERN_FORMATION | 18 | -0.633101 | -1.589594 | 0.027083 | 0.12406 |
| GOBP_POSITIVE_REGULATION_OF_CELL_CYCLE_ARREST | 74 | -0.467076 | -1.589209 | 0.008386 | 0.124019 |
| GOBP_DIGESTION | 77 | -0.468011 | -1.588507 | 0.014085 | 0.124386 |
| GOBP_REGULATION_OF_EXTRINSIC_APOPTOTIC_SIGNALING_PATHWAY | 132 | -0.43363 | -1.588146 | 0 | 0.124328 |
| GOBP_POSITIVE_REGULATION_OF_PROTEASOMAL_UBIQUITIN_DEPENDENT_PROTEIN_CAT/ | 83 | -0.465112 | -1.586737 | 0.010331 | 0.12538 |
| GOBP_APOPTOTIC_MITOCHONDRIAL_CHANGES | 108 | -0.442859 | -1.586399 | 0.006211 | 0.125292 |
| GOBP_CARDIAC_MUSCLE_CELL_MEMBRANE_REPOLARIZATION | 26 | -0.5811 | -1.586329 | 0.008696 | 0.124919 |
| GOBP_REGULATION_OF_APOPTOTIC_SIGNALING_PATHWAY | 305 | -0.389683 | -1.585178 | 0 | 0.125612 |
| GOBP_REGULATION_OF_MITOTIC_SPINDLE_ASSEMBLY | 18 | -0.626114 | -1.58507 | 0.026694 | 0.12529 |
| GOBP_POSITIVE_REGULATION_OF_PHAGOCYTOSIS | 59 | -0.488493 | -1.584991 | 0.008949 | 0.124943 |
| GOBP_MYELOID_LEUKOCYTE_DIFFERENTIATION | 177 | -0.411921 | -1.584922 | 0 | 0.124554 |
| GOBP_POSITIVE_REGULATION_OF_G1_S_TRANSITION_OF_MITOTIC_CELL_CYCLE | 35 | -0.536697 | -1.584251 | 0.015021 | 0.124942 |
| GOBP_LEFT_RIGHT_PATTERN_FORMATION | 18 | -0.624116 | -1.583718 | 0.033333 | 0.125098 |
| GOBP_EXTRINSIC_APOPTOTIC_SIGNALING_PATHWAY_VIA_DEATH_DOMAIN_RECEPTORS | 75 | -0.471225 | -1.583191 | 0.009153 | 0.125354 |
| GOBP_CELLULAR_CARBOHYDRATE_CATABOLIC_PROCESS | 40 | -0.535292 | -1.582279 | 0.012821 | 0.12595 |
| GOBP_POSITIVE_REGULATION_OF_LEUKOCYTE_ADHESION_TO_VASCULAR_ENDOTHELIAL_ | 19 | -0.625144 | -1.582194 | 0.03125 | 0.125633 |
| GOBP_REGULATION_OF_CELLULAR_RESPONSE_TO_TRANSFORMING_GROWTH_FACTOR_E | 112 | -0.440583 | -1.579956 | 0.010917 | 0.127584 |
| GOBP_POSITIVE_REGULATION_OF_LIPID_METABOLIC_PROCESS | 116 | -0.43877 | -1.579619 | 0 | 0.12755 |
| GOBP_POSITIVE_REGULATION_OF_P38MAPK_CASCADE | 22 | -0.595392 | -1.578756 | 0.026971 | 0.128155 |
| GOBP_RESPONSE_TO_PURINE_CONTAINING_COMPOUND | 124 | -0.426692 | -1.578221 | 0 | 0.128387 |
| GOBP_RESPONSE_TO_ALKALOID | 76 | -0.461233 | -1.577602 | 0.002232 | 0.128658 |
| GOBP_NEGATIVE_REGULATION_OF_CELL_DIFFERENTIATION | 492 | -0.371065 | -1.577359 | 0 | 0.128511 |
| GOBP_MACROPHAGE_ACTIVATION | 87 | -0.453916 | -1.576976 | 0.006522 | 0.128552 |
| GOBP_CELLULAR_RESPONSE_TO_STARVATION | 142 | -0.420701 | -1.576965 | 0.002174 | 0.128152 |
| GOBP_POSITIVE_REGULATION_OF_FAT_CELL_DIFFERENTIATION | 48 | -0.503708 | -1.576661 | 0.006289 | 0.128092 |
| GOBP_OLIGOSACCHARIDE_METABOLIC_PROCESS | 44 | -0.505557 | -1.575591 | 0.006329 | 0.128798 |
| GOBP_MONONUCLEAR_CELL_MIGRATION | 156 | -0.419585 | -1.575055 | 0.002398 | 0.128923 |
| GOBP_CELL_FATE_COMMITMENT | 161 | -0.414115 | -1.575 | 0 | 0.128605 |

| | | | | | |
|---|---|---|---|---|---|
| GOBP_POSITIVE_REGULATION_OF_ALCOHOL_BIOSYNTHETIC_PROCESS | 16 | 0.479728 | 1.173762 | 0.271805 | 0.636826 |
| GOBP_KERATINOCYTE_DIFFERENTIATION | 116 | 0.329924 | 1.173639 | 0.163303 | 0.635209 |
| GOBP_AXIS_ELONGATION | 20 | 0.467866 | 1.172033 | 0.242884 | 0.638475 |
| GOBP_HEART_PROCESS | 200 | 0.309748 | 1.171856 | 0.118243 | 0.637057 |
| GOBP_REGULATION_OF_HEART_CONTRACTION | 167 | 0.315399 | 1.170654 | 0.131086 | 0.638881 |
| GOBP_REGULATION_OF_CELLULAR_COMPONENT_SIZE | 325 | 0.292522 | 1.170419 | 0.090586 | 0.637589 |
| GOBP_GLYCOPROTEIN_CATABOLIC_PROCESS | 21 | 0.454499 | 1.169775 | 0.247619 | 0.637724 |
| GOBP_REGULATION_OF_TRANS_SYNAPTIC_SIGNALING | 303 | 0.295204 | 1.16899 | 0.11453 | 0.638348 |
| GOBP_NEGATIVE_REGULATION_OF_NEURON_PROJECTION_DEVELOPMENT | 111 | 0.33065 | 1.166818 | 0.170819 | 0.643666 |
| GOBP_CARDIAC_CONDUCTION | 106 | 0.329305 | 1.166287 | 0.17603 | 0.643479 |
| GOBP_REGULATION_OF_EPITHELIAL_CELL_MIGRATION | 203 | 0.308433 | 1.165578 | 0.142599 | 0.643833 |
| GOBP_CELL_MORPHOGENESIS_INVOLVED_IN_NEURON_DIFFERENTIATION | 469 | 0.284858 | 1.164871 | 0.076923 | 0.644197 |
| GOBP_PLATELET_DERIVED_GROWTH_FACTOR_RECEPTOR_SIGNALING_PATHWAY | 51 | 0.378702 | 1.16473 | 0.206287 | 0.642706 |
| GOBP_NEGATIVE_REGULATION_OF_CANONICAL_WNT_SIGNALING_PATHWAY | 155 | 0.318145 | 1.164218 | 0.156604 | 0.642466 |
| GOBP_POSITIVE_REGULATION_OF_RHO_PROTEIN_SIGNAL_TRANSDUCTION | 30 | 0.41188 | 1.164166 | 0.239464 | 0.64065 |
| GOBP_HEMATOPOIETIC_STEM_CELL_HOMEOSTASIS | 15 | 0.479027 | 1.164041 | 0.264151 | 0.639102 |
| GOBP_NEGATIVE_REGULATION_OF_INTRINSIC_APOPTOTIC_SIGNALING_PATHWAY_IN_RE | 30 | 0.416675 | 1.16382 | 0.249493 | 0.637868 |
| GOBP_RESPONSE_TO_ATP | 28 | 0.425632 | 1.163672 | 0.226744 | 0.63643 |
| GOBP_MONOAMINE_TRANSPORT | 50 | 0.371212 | 1.163299 | 0.208487 | 0.63575 |
| GOBP_REGULATION_OF_ADENYLATE_CYCLASE_ACTIVITY | 20 | 0.454262 | 1.161982 | 0.26945 | 0.638155 |
| GOBP_PHENOL_CONTAINING_COMPOUND_BIOSYNTHETIC_PROCESS | 26 | 0.420558 | 1.161823 | 0.255769 | 0.636738 |
| GOBP_HISTONE_H3_K9_METHYLATION | 30 | 0.41109 | 1.161202 | 0.243408 | 0.636861 |
| GOBP_NEGATIVE_REGULATION_OF_AXONOGENESIS | 56 | 0.374374 | 1.160816 | 0.207885 | 0.636209 |
| GOBP_ANOIKIS | 33 | 0.413333 | 1.160666 | 0.22366 | 0.634809 |
| GOBP_NEGATIVE_REGULATION_OF_LIPID_BIOSYNTHETIC_PROCESS | 40 | 0.39311 | 1.160199 | 0.243762 | 0.634433 |
| GOBP_REGULATION_OF_CARDIAC_CONDUCTION | 44 | 0.38947 | 1.159633 | 0.221359 | 0.634299 |
| GOBP_ARACHIDONIC_ACID_METABOLIC_PROCESS | 41 | 0.391811 | 1.15928 | 0.25046 | 0.63355 |
| GOBP_REGULATION_OF_INTRINSIC_APOPTOTIC_SIGNALING_PATHWAY_IN_RESPONSE_T | 36 | 0.398264 | 1.159127 | 0.255769 | 0.632178 |
| GOBP_WNT_SIGNALING_PATHWAY_CALCIUM_MODULATING_PATHWAY | 35 | 0.398644 | 1.15664 | 0.259184 | 0.638311 |
| GOBP_REGULATION_OF_T_CELL_MEDIATED_CYTOTOXICITY | 36 | 0.392198 | 1.155757 | 0.225326 | 0.639332 |
| GOBP_POSTSYNAPSE_ASSEMBLY | 23 | 0.435892 | 1.153951 | 0.266038 | 0.643432 |
| GOBP_ISOPRENOID_BIOSYNTHETIC_PROCESS | 24 | 0.43288 | 1.153839 | 0.275142 | 0.641876 |
| GOBP_POSITIVE_REGULATION_OF_T_CELL_MEDIATED_CYTOTOXICITY | 27 | 0.423566 | 1.153778 | 0.262264 | 0.6402 |
| GOBP_EPIDERMAL_CELL_DIFFERENTIATION | 162 | 0.311729 | 1.152578 | 0.145133 | 0.64223 |
| GOBP_NEURON_PROJECTION_GUIDANCE | 211 | 0.299349 | 1.152536 | 0.138053 | 0.640507 |
| GOBP_REGULATION_OF_MORPHOGENESIS_OF_A_BRANCHING_STRUCTURE | 30 | 0.408018 | 1.152234 | 0.259191 | 0.639659 |
| GOBP_MESODERM_MORPHOGENESIS | 50 | 0.371582 | 1.151725 | 0.238462 | 0.639448 |
| GOBP_POSITIVE_REGULATION_OF_MORPHOGENESIS_OF_AN_EPITHELIUM | 23 | 0.426617 | 1.150928 | 0.252768 | 0.640168 |
| GOBP_NEGATIVE_REGULATION_OF_EPITHELIAL_CELL_PROLIFERATION | 106 | 0.33017 | 1.150551 | 0.187617 | 0.639522 |
| GOBP_POSITIVE_REGULATION_OF_PEPTIDYL_THREONINE_PHOSPHORYLATION | 26 | 0.420022 | 1.149316 | 0.291585 | 0.641634 |
| GOBP_VESICLE_FUSION_TO_PLASMA_MEMBRANE | 17 | 0.468242 | 1.148909 | 0.293774 | 0.641143 |
| GOBP_REGULATION_OF_CALCIUM_ION_DEPENDENT_EXOCYTOSIS | 30 | 0.409878 | 1.14814 | 0.264479 | 0.641809 |
| GOBP_REGULATION_OF_NEUROGENESIS | 278 | 0.293176 | 1.148125 | 0.124343 | 0.640022 |
| GOBP_GENITALIA_DEVELOPMENT | 31 | 0.413532 | 1.146843 | 0.275391 | 0.64235 |
| GOBP_POSITIVE_REGULATION_OF_CHOLESTEROL_EFFLUX | 17 | 0.467096 | 1.146838 | 0.307692 | 0.640556 |
| GOBP_CARDIAC_ATRIUM_MORPHOGENESIS | 19 | 0.455974 | 1.146664 | 0.261122 | 0.639307 |
| GOBP_NEGATIVE_REGULATION_OF_AXON_EXTENSION_INVOLVED_IN_AXON_GUIDANCE | 22 | 0.442257 | 1.146395 | 0.284387 | 0.638355 |
| GOBP_BRANCHED_CHAIN_AMINO_ACID_METABOLIC_PROCESS | 23 | 0.448208 | 1.145291 | 0.282243 | 0.640037 |
| GOBP_STRIATED_MUSCLE_CONTRACTION | 119 | 0.325079 | 1.145163 | 0.185965 | 0.638656 |
| GOBP_REGULATION_OF_RENAL_SYSTEM_PROCESS | 19 | 0.460221 | 1.145058 | 0.319066 | 0.63723 |
| GOBP_NEGATIVE_REGULATION_OF_DNA_DAMAGE_RESPONSE_SIGNAL_TRANSDUCTION | 15 | 0.470967 | 1.144757 | 0.315589 | 0.6364 |
| GOBP_NEGATIVE_CHEMOTAXIS | 34 | 0.405059 | 1.144567 | 0.28932 | 0.63529 |
| GOBP_NEGATIVE_REGULATION_OF_STEROID_METABOLIC_PROCESS | 17 | 0.471644 | 1.144359 | 0.303263 | 0.634153 |
| GOBP_REGULATION_OF_TYPE_I_INTERFERON_MEDIATED_SIGNALING_PATHWAY | 31 | 0.405746 | 1.143967 | 0.273585 | 0.633678 |
| GOBP_RHYTHMIC_BEHAVIOR | 31 | 0.398071 | 1.143182 | 0.270169 | 0.634377 |
| GOBP_ACTIN_NUCLEATION | 46 | 0.375308 | 1.142485 | 0.248092 | 0.634943 |
| GOBP_CHONDROCYTE_DIFFERENTIATION | 86 | 0.341336 | 1.142448 | 0.219153 | 0.633366 |
| GOBP_EMBRYONIC_PATTERN_SPECIFICATION | 48 | 0.367617 | 1.142422 | 0.242366 | 0.631717 |
| GOBP_REGULATION_OF_SPROUTING_ANGIOGENESIS | 35 | 0.393756 | 1.141364 | 0.249057 | 0.633349 |
| GOBP_PRIMARY_NEURAL_TUBE_FORMATION | 78 | 0.34375 | 1.141333 | 0.20381 | 0.631739 |
| GOBP_LOCOMOTORY_BEHAVIOR | 109 | 0.323531 | 1.140797 | 0.199288 | 0.631705 |
| GOBP_IMMUNOLOGICAL_MEMORY_PROCESS | 15 | 0.47334 | 1.140016 | 0.314672 | 0.632458 |
| GOBP_SENSORY_SYSTEM_DEVELOPMENT | 261 | 0.291824 | 1.139118 | 0.140221 | 0.633525 |
| GOBP_MEMBRANE_DEPOLARIZATION_DURING_ACTION_POTENTIAL | 26 | 0.427752 | 1.138859 | 0.279245 | 0.63271 |
| GOBP_ATRIAL_SEPTUM_DEVELOPMENT | 15 | 0.471243 | 1.138191 | 0.295019 | 0.633038 |
| GOBP_CARDIAC_ATRIUM_DEVELOPMENT | 23 | 0.426214 | 1.137386 | 0.293358 | 0.633797 |
| GOBP_SENSORY_ORGAN_DEVELOPMENT | 375 | 0.286263 | 1.137266 | 0.134354 | 0.632498 |
| GOBP_MUSCLE_CELL_DEVELOPMENT | 121 | 0.319187 | 1.137224 | 0.212644 | 0.630963 |
| GOBP_ADENYLATE_CYCLASE_ACTIVATING_G_PROTEIN_COUPLED_RECEPTOR_SIGNALI | 91 | 0.337516 | 1.136819 | 0.215074 | 0.630584 |
| GOBP_INTERMEDIATE_FILAMENT_ORGANIZATION | 15 | 0.466643 | 1.136755 | 0.29899 | 0.629135 |
| GOBP_REGULATION_OF_MUSCLE_SYSTEM_PROCESS | 165 | 0.303737 | 1.136674 | 0.177817 | 0.627744 |
| GOBP_NEGATIVE_REGULATION_OF_AXON_EXTENSION | 38 | 0.394185 | 1.136111 | 0.278986 | 0.627838 |
| GOBP_POTASSIUM_ION_IMPORT_ACROSS_PLASMA_MEMBRANE | 33 | 0.401146 | 1.133573 | 0.27853 | 0.634112 |
| GOBP_POSITIVE_REGULATION_OF_PEPTIDE_HORMONE_SECRETION | 65 | 0.351337 | 1.132967 | 0.212844 | 0.634419 |
| GOBP_NEGATIVE_REGULATION_OF_CARDIAC_MUSCLE_TISSUE_GROWTH | 21 | 0.439116 | 1.132412 | 0.296642 | 0.634517 |
| GOBP_MUSCLE_TISSUE_DEVELOPMENT | 274 | 0.287578 | 1.130692 | 0.162544 | 0.638237 |
| GOBP_BLOOD_VESSEL_ENDOTHELIAL_CELL_MIGRATION | 105 | 0.32372 | 1.130445 | 0.231061 | 0.637428 |
| GOBP_REGULATION_OF_LONG_TERM_NEURONAL_SYNAPTIC_PLASTICITY | 18 | 0.451858 | 1.129875 | 0.318868 | 0.637553 |
| GOBP_L_GLUTAMATE_TRANSMEMBRANE_TRANSPORT | 19 | 0.446335 | 1.129643 | 0.305019 | 0.636644 |
| GOBP_SYNAPTIC_SIGNALING | 485 | 0.272968 | 1.128512 | 0.109855 | 0.638514 |
| GOBP_PHOSPHATE_ION_TRANSPORT | 20 | 0.450839 | 1.128408 | 0.308489 | 0.637212 |
| GOBP_NEGATIVE_REGULATION_OF_SYNAPTIC_TRANSMISSION | 47 | 0.369916 | 1.127742 | 0.262357 | 0.637581 |
| GOBP_TISSUE_MIGRATION | 264 | 0.289568 | 1.127652 | 0.191126 | 0.636229 |
| GOBP_CATECHOLAMINE_SECRETION | 31 | 0.405461 | 1.125519 | 0.295455 | 0.6412 |
| GOBP_NEGATIVE_REGULATION_OF_REPRODUCTIVE_PROCESS | 39 | 0.381208 | 1.12461 | 0.281716 | 0.642481 |
| GOBP_NEUROTRANSMITTER_UPTAKE | 26 | 0.415131 | 1.124206 | 0.312727 | 0.642152 |
| GOBP_NEGATIVE_REGULATION_OF_RESPONSE_TO_WOUNDING | 66 | 0.343134 | 1.124018 | 0.242647 | 0.641131 |
| GOBP_REGULATION_OF_CELL_SUBSTRATE_JUNCTION_ORGANIZATION | 60 | 0.344932 | 1.12245 | 0.243636 | 0.644501 |
| GOBP_T_CELL_RECEPTOR_SIGNALING_PATHWAY | 194 | 0.298346 | 1.122402 | 0.209009 | 0.643031 |
| GOBP_MAMMARY_GLAND_EPITHELIUM_DEVELOPMENT | 53 | 0.361464 | 1.121682 | 0.282243 | 0.643683 |
| GOBP_APOPTOTIC_DNA_FRAGMENTATION | 20 | 0.436288 | 1.121618 | 0.306613 | 0.642287 |
| GOBP_CELL_COMMUNICATION_BY_ELECTRICAL_COUPLING | 24 | 0.424597 | 1.120348 | 0.296435 | 0.644526 |
| GOBP_REGULATION_OF_NEURON_PROJECTION_DEVELOPMENT | 351 | 0.277524 | 1.120181 | 0.164384 | 0.643458 |
| GOBP_REGULATION_OF_VASOCONSTRICTION | 40 | 0.381527 | 1.119838 | 0.294798 | 0.642898 |
| GOBP_REGULATION_OF_SYNAPSE_STRUCTURE_OR_ACTIVITY | 159 | 0.302537 | 1.119718 | 0.209009 | 0.641673 |
| GOBP_PEPTIDE_HORMONE_SECRETION | 181 | 0.300402 | 1.119324 | 0.20696 | 0.641315 |
| GOBP_ARTERY_DEVELOPMENT | 65 | 0.345253 | 1.119058 | 0.270588 | 0.64057 |
| GOBP_HORMONE_TRANSPORT | 229 | 0.2919 | 1.119 | 0.215827 | 0.639184 |
| GOBP_REGULATION_OF_HORMONE_LEVELS | 357 | 0.277597 | 1.118979 | 0.139803 | 0.637674 |
| GOBP_MORPHOGENESIS_OF_EMBRYONIC_EPITHELIUM | 112 | 0.3217 | 1.117966 | 0.240809 | 0.639241 |
| GOBP_RELEASE_OF_SEQUESTERED_CALCIUM_ION_INTO_CYTOSOL_BY_ENDOPLASMIC | 27 | 0.415339 | 1.117546 | 0.293651 | 0.639013 |
| GOBP_ENTEROENDOCRINE_CELL_DIFFERENTIATION | 17 | 0.457372 | 1.117292 | 0.304519 | 0.638321 |
| GOBP_CELL_MATRIX_ADHESION | 190 | 0.300541 | 1.116907 | 0.215017 | 0.637998 |
| GOBP_ERBB_SIGNALING_PATHWAY | 127 | 0.308954 | 1.116826 | 0.236234 | 0.636725 |
| GOBP_PRESYNAPSE_ORGANIZATION | 32 | 0.394197 | 1.11611 | 0.310941 | 0.637412 |
| GOBP_POSITIVE_REGULATION_OF_LIPASE_ACTIVITY | 53 | 0.361481 | 1.115327 | 0.285714 | 0.638323 |
| GOBP_CAMERA_TYPE_EYE_DEVELOPMENT | 217 | 0.290877 | 1.115085 | 0.224779 | 0.637596 |
| GOBP_CHONDROITIN_SULFATE_PROTEOGLYCAN_BIOSYNTHETIC_PROCESS | 27 | 0.412948 | 1.114089 | 0.297445 | 0.639177 |
| GOBP_CELLULAR_RESPONSE_TO_ATP | 16 | 0.463793 | 1.113883 | 0.339882 | 0.638333 |
| GOBP_MUSCLE_CONTRACTION | 236 | 0.288663 | 1.113845 | 0.207815 | 0.636921 |
| GOBP_GASTRULATION | 141 | 0.304991 | 1.113502 | 0.23475 | 0.636492 |
| GOBP_EMBRYONIC_HEART_TUBE_MORPHOGENESIS | 51 | 0.357249 | 1.112925 | 0.296089 | 0.636786 |
| GOBP_CARDIAC_MUSCLE_TISSUE_DEVELOPMENT | 147 | 0.302837 | 1.112882 | 0.227778 | 0.635404 |
| GOBP_LYMPH_VESSEL_DEVELOPMENT | 22 | 0.425874 | 1.111854 | 0.320565 | 0.637053 |
| GOBP_CELL_COMMUNICATION_INVOLVED_IN_CARDIAC_CONDUCTION | 41 | 0.373448 | 1.111276 | 0.29982 | 0.637357 |
| GOBP_MEMBRANE_DEPOLARIZATION | 66 | 0.34295 | 1.110915 | 0.246777 | 0.636989 |
| GOBP_CARDIAC_SEPTUM_MORPHOGENESIS | 47 | 0.366102 | 1.110807 | 0.281426 | 0.635825 |
| GOBP_REGULATION_OF_POSTSYNAPSE_ORGANIZATION | 70 | 0.345965 | 1.110622 | 0.275797 | 0.634895 |
| GOBP_REGULATION_OF_MEMBRANE_POTENTIAL | 298 | 0.281921 | 1.108683 | 0.189046 | 0.639205 |
| GOBP_CARDIAC_EPITHELIAL_TO_MESENCHYMAL_TRANSITION | 25 | 0.420471 | 1.108673 | 0.317121 | 0.637757 |
| GOBP_MITOCHONDRIAL_CALCIUM_ION_TRANSMEMBRANE_TRANSPORT | 25 | 0.415349 | 1.108579 | 0.277778 | 0.636518 |
| GOBP_VASODILATION | 25 | 0.414862 | 1.107912 | 0.334608 | 0.637012 |
| GOBP_RESPONSE_TO_AXON_INJURY | 64 | 0.345716 | 1.107121 | 0.286486 | 0.637988 |
| GOBP_BARBED_END_ACTIN_FILAMENT_CAPPING | 24 | 0.424147 | 1.106818 | 0.321361 | 0.637467 |
| GOBP_SARCOPLASMIC_RETICULUM_CALCIUM_ION_TRANSPORT | 29 | 0.391555 | 1.105778 | 0.30754 | 0.639127 |
| GOBP_PEPTIDE_CROSS_LINKING | 20 | 0.434618 | 1.101759 | 0.345865 | 0.649968 |
| GOBP_ENSHEATHMENT_OF_NEURONS | 109 | 0.313459 | 1.101571 | 0.27037 | 0.64908 |
| GOBP_POSITIVE_REGULATION_OF_ACTIN_FILAMENT_POLYMERIZATION | 84 | 0.327904 | 1.101447 | 0.282648 | 0.647987 |
| GOBP_CHONDROITIN_SULFATE_PROTEOGLYCAN_METABOLIC_PROCESS | 37 | 0.380065 | 1.100349 | 0.327778 | 0.649953 |
| GOBP_NEGATIVE_REGULATION_OF_WOUND_HEALING | 52 | 0.352148 | 1.100104 | 0.296763 | 0.649247 |
| GOBP_NEGATIVE_REGULATION_OF_CHEMOTAXIS | 50 | 0.357494 | 1.099935 | 0.319703 | 0.648292 |
| GOBP_FERTILIZATION | 122 | 0.311629 | 1.099888 | 0.255009 | 0.646977 |
| GOBP_STRIATED_MUSCLE_CELL_PROLIFERATION | 55 | 0.349828 | 1.099664 | 0.300188 | 0.646186 |
| GOBP_ACROSOME_REACTION | 25 | 0.41012 | 1.09824 | 0.335206 | 0.648986 |
| GOBP_LYMPH_NODE_DEVELOPMENT | 15 | 0.458718 | 1.097902 | 0.353814 | 0.648587 |
| GOBP_NEURON_MIGRATION | 96 | 0.315353 | 1.097722 | 0.277064 | 0.647659 |
| GOBP_ENDOCARDIAL_CUSHION_FORMATION | 18 | 0.443102 | 1.097387 | 0.330739 | 0.64717 |
| GOBP_LEUKOCYTE_APOPTOTIC_PROCESS | 91 | 0.318817 | 1.096548 | 0.274864 | 0.648229 |
| GOBP_POSITIVE_REGULATION_OF_ACTIN_FILAMENT_BUNDLE_ASSEMBLY | 51 | 0.358578 | 1.095907 | 0.296223 | 0.648762 |
| GOBP_INNER_EAR_RECEPTOR_CELL_STEREOCILIUM_ORGANIZATION | 25 | 0.40664 | 1.095686 | 0.339286 | 0.648002 |
| GOBP_ACTION_POTENTIAL | 96 | 0.315725 | 1.095191 | 0.275093 | 0.648129 |
| GOBP_POSITIVE_REGULATION_OF_PROTEIN_TYROSINE_KINASE_ACTIVITY | 51 | 0.35449 | 1.094286 | 0.311284 | 0.649471 |
| GOBP_REGULATION_OF_LIPASE_ACTIVITY | 72 | 0.331416 | 1.094233 | 0.274545 | 0.648214 |
| GOBP_NEGATIVE_REGULATION_OF_NEURON_APOPTOTIC_PROCESS | 102 | 0.312123 | 1.093952 | 0.286232 | 0.647573 |
| GOBP_EPIDERMAL_GROWTH_FACTOR_RECEPTOR_SIGNALING_PATHWAY | 109 | 0.309194 | 1.093346 | 0.263258 | 0.64797 |
| GOBP_SLEEP | 20 | 0.418019 | 1.093125 | 0.336714 | 0.64721 |
| GOBP_DENDRITIC_SPINE_DEVELOPMENT | 92 | 0.323092 | 1.092967 | 0.26945 | 0.646259 |
| GOBP_LYMPHOCYTE_APOPTOTIC_PROCESS | 65 | 0.336579 | 1.09274 | 0.287402 | 0.645578 |
| GOBP_DENDRITE_MORPHOGENESIS | 122 | 0.305863 | 1.092144 | 0.246212 | 0.645965 |
| GOBP_INTERFERON_GAMMA_PRODUCTION | 97 | 0.316082 | 1.091902 | 0.296846 | 0.645323 |
| GOBP_CELLULAR_SODIUM_ION_HOMEOSTASIS | 15 | 0.455295 | 1.091728 | 0.343689 | 0.644472 |
| GOBP_NEURAL_TUBE_FORMATION | 85 | 0.324353 | 1.091682 | 0.303474 | 0.643236 |
| GOBP_ASPARTATE_FAMILY_AMINO_ACID_METABOLIC_PROCESS | 45 | 0.363563 | 1.090828 | 0.317857 | 0.644422 |
| GOBP_REGULATION_OF_CELL_CELL_ADHESION_MEDIATED_BY_CADHERIN | 18 | 0.45097 | 1.088548 | 0.373016 | 0.649973 |
| GOBP_CARDIAC_CELL_DEVELOPMENT | 58 | 0.340777 | 1.088479 | 0.328413 | 0.64879 |
| GOBP_NEGATIVE_REGULATION_OF_SECRETION | 116 | 0.303468 | 1.087847 | 0.287594 | 0.649375 |
| GOBP_POSITIVE_REGULATION_OF_MEIOTIC_CELL_CYCLE | 17 | 0.443186 | 1.087763 | 0.345725 | 0.648219 |

| | | | | | |
|---|---|---|---|---|---|
| GOBP_NEGATIVE_REGULATION_OF_PHOSPHORUS_METABOLIC_PROCESS | 450 | -0.371201 | -1.574407 | 0 | 0.128813 |
| GOBP_NLS_BEARING_PROTEIN_IMPORT_INTO_NUCLEUS | 18 | -0.625593 | -1.573602 | 0.02544 | 0.129322 |
| GOBP_POSITIVE_REGULATION_OF_STRESS_ACTIVATED_PROTEIN_KINASE_SIGNALING_CAS | 105 | -0.439417 | -1.572081 | 0.002179 | 0.130666 |
| GOBP_REGULATION_OF_CARDIAC_MUSCLE_CELL_MEMBRANE_REPOLARIZATION | 19 | -0.622495 | -1.571438 | 0.019108 | 0.130906 |
| GOBP_VESICLE_LOCALIZATION | 210 | -0.398841 | -1.569675 | 0 | 0.132513 |
| GOBP_ASTROCYTE_DEVELOPMENT | 34 | -0.533564 | -1.56775 | 0.010101 | 0.134292 |
| GOBP_TETRAPYRROLE_METABOLIC_PROCESS | 53 | -0.49774 | -1.567518 | 0.010309 | 0.134144 |
| GOBP_ADAPTIVE_IMMUNE_RESPONSE_BASED_ON_SOMATIC_RECOMBINATION_OF_IMMUN | 317 | -0.382595 | -1.565661 | 0 | 0.135801 |
| GOBP_POSITIVE_REGULATION_OF_NUCLEAR_DIVISION | 48 | -0.500652 | -1.565446 | 0.004132 | 0.135668 |
| GOBP_MYD88_DEPENDENT_TOLL_LIKE_RECEPTOR_SIGNALING_PATHWAY | 34 | -0.529236 | -1.565051 | 0.026369 | 0.135739 |
| GOBP_POSITIVE_REGULATION_OF_COLD_INDUCED_THERMOGENESIS | 82 | -0.452413 | -1.564959 | 0 | 0.135415 |
| GOBP_RESPONSE_TO_INORGANIC_SUBSTANCE | 429 | -0.374776 | -1.564759 | 0 | 0.13525 |
| GOBP_LYMPHOCYTE_CHEMOTAXIS | 41 | -0.524904 | -1.564642 | 0.014644 | 0.134949 |
| GOBP_REGULATION_OF_FATTY_ACID_BETA_OXIDATION | 19 | -0.602796 | -1.564606 | 0.021008 | 0.134567 |
| GOBP_REGULATION_OF_BONE_MINERALIZATION | 57 | -0.495622 | -1.563427 | 0.010661 | 0.135408 |
| GOBP_POSITIVE_REGULATION_OF_KERATINOCYTE_DIFFERENTIATION | 15 | -0.627873 | -1.561679 | 0.035941 | 0.136866 |
| GOBP_METANEPHROS_DEVELOPMENT | 55 | -0.495453 | -1.559617 | 0.010684 | 0.138905 |
| GOBP_POSITIVE_REGULATION_OF_CELL_CYCLE_G1_S_PHASE_TRANSITION | 44 | -0.501159 | -1.559582 | 0.014768 | 0.138511 |
| GOBP_NEGATIVE_REGULATION_OF_CYSTEINE_TYPE_ENDOPEPTIDASE_ACTIVITY | 66 | -0.4712 | -1.559439 | 0.004246 | 0.138259 |
| GOBP_REGULATION_OF_MRNA_PROCESSING | 126 | -0.428642 | -1.559308 | 0.004255 | 0.137993 |
| GOBP_POSITIVE_REGULATION_OF_TRANSCRIPTION_FROM_RNA_POLYMERASE_II_PROMOT | 23 | -0.588297 | -1.559035 | 0.032653 | 0.137911 |
| GOBP_BIOLOGICAL_PROCESS_INVOLVED_IN_INTERACTION_WITH_HOST | 196 | -0.403672 | -1.558818 | 0 | 0.137739 |
| GOBP_RESPONSE_TO_PEPTIDE | 433 | -0.373985 | -1.558461 | 0 | 0.13776 |
| GOBP_VASCULAR_PROCESS_IN_CIRCULATORY_SYSTEM | 186 | -0.407556 | -1.55769 | 0 | 0.138342 |
| GOBP_COLLAGEN_BIOSYNTHETIC_PROCESS | 38 | -0.518331 | -1.557627 | 0.020747 | 0.137986 |
| GOBP_ASTROCYTE_ACTIVATION | 20 | -0.617578 | -1.556469 | 0.029787 | 0.138812 |
| GOBP_DIGESTIVE_SYSTEM_DEVELOPMENT | 95 | -0.444014 | -1.555686 | 0.002096 | 0.139328 |
| GOBP_POSITIVE_REGULATION_OF_LOCOMOTION | 459 | -0.369718 | -1.555191 | 0 | 0.139493 |
| GOBP_EXTRACELLULAR_TRANSPORT | 30 | -0.545291 | -1.554392 | 0.014989 | 0.139997 |
| GOBP_NEGATIVE_REGULATION_OF_HEMOPOIESIS | 92 | -0.441352 | -1.554328 | 0 | 0.139664 |
| GOBP_RESPONSE_TO_TOXIC_SUBSTANCE | 194 | -0.40639 | -1.553678 | 0 | 0.139947 |
| GOBP_INTRINSIC_APOPTOTIC_SIGNALING_PATHWAY_IN_RESPONSE_TO_ENDOPLASMIC_RI | 56 | -0.488167 | -1.553241 | 0.008602 | 0.139954 |
| GOBP_ORGANIC_ACID_TRANSPORT | 237 | -0.392846 | -1.552299 | 0 | 0.140585 |
| GOBP_RNA_3_END_PROCESSING | 143 | -0.419353 | -1.552291 | 0.002364 | 0.140181 |
| GOBP_HEME_METABOLIC_PROCESS | 30 | -0.548858 | -1.550983 | 0.023305 | 0.141273 |
| GOBP_MITOTIC_DNA_REPLICATION | 16 | -0.63991 | -1.550546 | 0.028455 | 0.141446 |
| GOBP_VIRAL_LIFE_CYCLE | 311 | -0.379653 | -1.550415 | 0 | 0.141181 |
| GOBP_SIGNAL_TRANSDUCTION_IN_ABSENCE_OF_LIGAND | 59 | -0.4805 | -1.549933 | 0.00813 | 0.141245 |
| GOBP_NEGATIVE_REGULATION_OF_PEPTIDYL_TYROSINE_PHOSPHORYLATION | 49 | -0.492151 | -1.54975 | 0.023758 | 0.141071 |
| GOBP_REGULATION_OF_ANIMAL_ORGAN_MORPHOGENESIS | 143 | -0.41801 | -1.549643 | 0.004608 | 0.140835 |
| GOBP_REGULATION_OF_MYELOID_CELL_DIFFERENTIATION | 225 | -0.39384 | -1.549185 | 0.004662 | 0.140902 |
| GOBP_CELLULAR_RESPONSE_TO_TOXIC_SUBSTANCE | 101 | -0.442065 | -1.548516 | 0.004484 | 0.14114 |
| GOBP_INTRACILIARY_TRANSPORT | 52 | -0.48807 | -1.545982 | 0.023707 | 0.143678 |
| GOBP_REGULATION_OF_ANION_TRANSMEMBRANE_TRANSPORT | 89 | -0.443715 | -1.544623 | 0.004405 | 0.145028 |
| GOBP_REGULATION_OF_MICROTUBULE_BASED_MOVEMENT | 38 | -0.521136 | -1.543833 | 0.014957 | 0.145582 |
| GOBP_MODULATION_OF_PROCESS_OF_OTHER_ORGANISM_INVOLVED_IN_SYMBIOTIC_INTE | 89 | -0.443298 | -1.542377 | 0.00216 | 0.147035 |
| GOBP_MITOTIC_METAPHASE_PLATE_CONGRESSION | 50 | -0.48565 | -1.542325 | 0.012632 | 0.146734 |
| GOBP_HISTONE_EXCHANGE | 51 | -0.493088 | -1.541187 | 0.014831 | 0.147733 |
| GOBP_POSITIVE_REGULATION_OF_SMOOTH_MUSCLE_CELL_PROLIFERATION | 72 | -0.45782 | -1.540606 | 0.006397 | 0.147997 |
| GOBP_POSITIVE_REGULATION_OF_LEUKOCYTE_MIGRATION | 112 | -0.432099 | -1.539864 | 0.002398 | 0.148479 |
| GOBP_POSITIVE_CHEMOTAXIS | 46 | -0.501477 | -1.539653 | 0.016427 | 0.148283 |
| GOBP_RECEPTOR_SIGNALING_PATHWAY_VIA_STAT | 124 | -0.425218 | -1.539124 | 0.004444 | 0.148509 |
| GOBP_MOVEMENT_IN_HOST_ENVIRONMENT | 155 | -0.409583 | -1.538974 | 0 | 0.148322 |
| GOBP_POSITIVE_REGULATION_OF_CELLULAR_RESPONSE_TO_TRANSFORMING_GROWTH_ | 26 | -0.558094 | -1.538127 | 0.023207 | 0.14886 |
| GOBP_POSITIVE_REGULATION_OF_EPIDERMAL_CELL_DIFFERENTIATION | 21 | -0.582421 | -1.53723 | 0.037578 | 0.149551 |
| GOBP_ACTIVATION_OF_PROTEIN_KINASE_ACTIVITY | 285 | -0.378513 | -1.537163 | 0 | 0.149237 |
| GOBP_REGULATION_OF_EXTRINSIC_APOPTOTIC_SIGNALING_PATHWAY_VIA_DEATH_DOMA | 51 | -0.487507 | -1.537082 | 0.023707 | 0.148943 |
| GOBP_CELLULAR_RESPONSE_TO_EXTERNAL_STIMULUS | 267 | -0.380424 | -1.535736 | 0 | 0.150185 |
| GOBP_ACTIVATION_OF_CYSTEINE_TYPE_ENDOPEPTIDASE_ACTIVITY_INVOLVED_IN_APOPT | 80 | -0.450238 | -1.534848 | 0.010989 | 0.150787 |
| GOBP_NEGATIVE_REGULATION_OF_TRANSFERASE_ACTIVITY | 261 | -0.379658 | -1.533858 | 0 | 0.151448 |
| GOBP_POLYSACCHARIDE_METABOLIC_PROCESS | 99 | -0.427276 | -1.53304 | 0.004065 | 0.152022 |
| GOBP_POSITIVE_REGULATION_OF_INTERLEUKIN_6_PRODUCTION | 77 | -0.448798 | -1.532355 | 0.020225 | 0.152411 |
| GOBP_DEFENSE_RESPONSE_TO_VIRUS | 213 | -0.387499 | -1.53095 | 0.002268 | 0.153724 |
| GOBP_DEVELOPMENT_OF_PRIMARY_SEXUAL_CHARACTERISTICS | 166 | -0.399918 | -1.530735 | 0 | 0.153593 |
| GOBP_CIRCULATORY_SYSTEM_PROCESS | 441 | -0.364399 | -1.530517 | 0 | 0.153395 |
| GOBP_REGULATION_OF_UBIQUITIN_PROTEIN_LIGASE_ACTIVITY | 22 | -0.576504 | -1.530224 | 0.035088 | 0.153307 |
| GOBP_NEGATIVE_REGULATION_OF_MITOTIC_CELL_CYCLE | 283 | -0.378303 | -1.529869 | 0 | 0.153338 |
| GOBP_GLAND_DEVELOPMENT | 319 | -0.376115 | -1.529425 | 0 | 0.153446 |
| GOBP_CELLULAR_RESPONSE_TO_OSMOTIC_STRESS | 35 | -0.520831 | -1.526953 | 0.032328 | 0.156051 |
| GOBP_ANATOMICAL_STRUCTURE_MATURATION | 167 | -0.397569 | -1.52567 | 0.00216 | 0.157237 |
| GOBP_POSITIVE_REGULATION_OF_SMOOTH_MUSCLE_CONTRACTION | 20 | -0.5913 | -1.525553 | 0.031746 | 0.156969 |
| GOBP_INTERLEUKIN_1_MEDIATED_SIGNALING_PATHWAY | 100 | -0.430312 | -1.525337 | 0.006522 | 0.156775 |
| GOBP_POSITIVE_REGULATION_OF_CYTOKINE_PRODUCTION | 374 | -0.37124 | -1.524258 | 0 | 0.157849 |
| GOBP_NEGATIVE_REGULATION_OF_CELL_SUBSTRATE_ADHESION | 53 | -0.477254 | -1.522208 | 0.016701 | 0.160082 |
| GOBP_NEGATIVE_REGULATION_OF_PHOSPHORYLATION | 349 | -0.366131 | -1.522052 | 0 | 0.159883 |
| GOBP_PYRIMIDINE_NUCLEOSIDE_METABOLIC_PROCESS | 33 | -0.519926 | -1.521846 | 0.040773 | 0.159748 |
| GOBP_T_CELL_CYTOKINE_PRODUCTION | 32 | -0.52689 | -1.521434 | 0.034343 | 0.159782 |
| GOBP_AMINO_ACID_TRANSMEMBRANE_TRANSPORT | 72 | -0.452849 | -1.520215 | 0.013333 | 0.160829 |
| GOBP_ORGANELLE_DISASSEMBLY | 104 | -0.427367 | -1.518936 | 0.012739 | 0.161898 |
| GOBP_PHAGOCYTOSIS | 332 | -0.369053 | -1.51748 | 0 | 0.163291 |
| GOBP_GLYCOSYL_COMPOUND_CATABOLIC_PROCESS | 40 | -0.5052 | -1.516838 | 0.025424 | 0.163764 |
| GOBP_CARBOHYDRATE_CATABOLIC_PROCESS | 173 | -0.398128 | -1.516588 | 0.004808 | 0.163658 |
| GOBP_HEART_TRABECULA_MORPHOGENESIS | 24 | -0.561575 | -1.516259 | 0.039046 | 0.163649 |
| GOBP_REGULATION_OF_MITOCHONDRIAL_MEMBRANE_PERMEABILITY | 68 | -0.460887 | -1.515807 | 0.00885 | 0.163868 |
| GOBP_MICROTUBULE_NUCLEATION | 29 | -0.535781 | -1.514038 | 0.024096 | 0.165721 |
| GOBP_NEGATIVE_REGULATION_OF_TRANSFORMING_GROWTH_FACTOR_BETA_RECEPTOR | 75 | -0.448736 | -1.513862 | 0.016985 | 0.165515 |
| GOBP_IMPORT_INTO_CELL | 174 | -0.394145 | -1.512967 | 0.002268 | 0.166296 |
| GOBP_REGULATION_OF_MITOCHONDRIAL_OUTER_MEMBRANE_PERMEABILIZATION_INVOL | 43 | -0.484812 | -1.5126 | 0.016129 | 0.166375 |
| GOBP_HEPATICOBILIARY_SYSTEM_DEVELOPMENT | 115 | -0.412692 | -1.51236 | 0.002028 | 0.166308 |
| GOBP_SEX_DIFFERENTIATION | 196 | -0.392578 | -1.511501 | 0 | 0.167009 |
| GOBP_CELL_CYCLE_CHECKPOINT | 200 | -0.388109 | -1.510746 | 0.009238 | 0.16754 |
| GOBP_CARBOHYDRATE_TRANSPORT | 122 | -0.418493 | -1.510172 | 0.004219 | 0.167793 |
| GOBP_CELLULAR_RESPONSE_TO_CORTICOSTEROID_STIMULUS | 45 | -0.487591 | -1.509383 | 0.017241 | 0.168521 |
| GOBP_RESPONSE_TO_OXIDATIVE_STRESS | 385 | -0.359924 | -1.508811 | 0 | 0.168908 |
| GOBP_RESPONSE_TO_DRUG | 282 | -0.37563 | -1.507729 | 0 | 0.170003 |
| GOBP_REGULATION_OF_EPITHELIAL_TO_MESENCHYMAL_TRANSITION | 81 | -0.43605 | -1.507645 | 0.015119 | 0.169732 |
| GOBP_RESPONSE_TO_ENDOPLASMIC_RETICULUM_STRESS | 277 | -0.372055 | -1.507309 | 0 | 0.169806 |
| GOBP_NEGATIVE_REGULATION_OF_PROTEIN_LOCALIZATION_TO_CELL_PERIPHERY | 20 | -0.576473 | -1.507236 | 0.03268 | 0.169506 |
| GOBP_REGULATION_OF_TRANSLATION_IN_RESPONSE_TO_STRESS | 21 | -0.562499 | -1.506136 | 0.036876 | 0.170489 |
| GOBP_REGULATION_OF_CILIUM_MOVEMENT | 24 | -0.554094 | -1.504643 | 0.035491 | 0.17204 |
| GOBP_REGULATION_OF_EXECUTION_PHASE_OF_APOPTOSIS | 15 | -0.604616 | -1.504526 | 0.066239 | 0.171794 |
| GOBP_COPII_COATED_VESICLE_BUDDING | 70 | -0.450934 | -1.503154 | 0.012552 | 0.173055 |
| GOBP_POSITIVE_REGULATION_OF_SMOOTHENED_SIGNALING_PATHWAY | 29 | -0.531646 | -1.503132 | 0.042735 | 0.172678 |
| GOBP_REGULATION_OF_ATPASE_ACTIVITY | 71 | -0.449847 | -1.502779 | 0.010707 | 0.172739 |
| GOBP_CELLULAR_OXIDANT_DETOXIFICATION | 85 | -0.436794 | -1.502474 | 0.006742 | 0.172754 |
| GOBP_REGULATION_OF_ESTABLISHMENT_OF_PROTEIN_LOCALIZATION_TO_MITOCHONDRI | 70 | -0.44581 | -1.501881 | 0.013514 | 0.173102 |
| GOBP_POSITIVE_REGULATION_OF_PROTEIN_KINASE_ACTIVITY | 453 | -0.356426 | -1.500852 | 0 | 0.174086 |
| GOBP_POSITIVE_REGULATION_OF_CHEMOKINE_PRODUCTION | 52 | -0.471841 | -1.500498 | 0.030738 | 0.174162 |
| GOBP_GLYCOLYTIC_PROCESS_THROUGH_FRUCTOSE_6_PHOSPHATE | 26 | -0.537652 | -1.498779 | 0.050633 | 0.176064 |
| GOBP_CELLULAR_RESPONSE_TO_OXYGEN_LEVELS | 194 | -0.387206 | -1.498435 | 0.002212 | 0.176128 |
| GOBP_REGULATION_OF_BIOMINERALIZATION | 71 | -0.448455 | -1.49772 | 0.006438 | 0.176702 |
| GOBP_CELL_PROLIFERATION_INVOLVED_IN_KIDNEY_DEVELOPMENT | 15 | -0.609153 | -1.497461 | 0.059305 | 0.176622 |
| GOBP_POSITIVE_REGULATION_OF_TRANSMEMBRANE_RECEPTOR_PROTEIN_SERINE_THRE | 87 | -0.432515 | -1.497439 | 0.013363 | 0.176231 |
| GOBP_NEGATIVE_REGULATION_OF_CELL_DEVELOPMENT | 130 | -0.405069 | -1.496572 | 0.010331 | 0.177014 |
| GOBP_POSITIVE_REGULATION_OF_CELL_CYCLE_PHASE_TRANSITION | 90 | -0.428973 | -1.496509 | 0.011737 | 0.176698 |
| GOBP_REGULATION_OF_SPINDLE_CHECKPOINT | 15 | -0.609582 | -1.496342 | 0.050403 | 0.176461 |
| GOBP_POSITIVE_REGULATION_OF_AMINO_ACID_TRANSPORT | 15 | -0.615983 | -1.495656 | 0.04918 | 0.176971 |
| GOBP_RESPONSE_TO_ACETYLCHOLINE | 22 | -0.563502 | -1.495492 | 0.044534 | 0.176724 |
| GOBP_LABYRINTHINE_LAYER_BLOOD_VESSEL_DEVELOPMENT | 16 | -0.612924 | -1.494855 | 0.05144 | 0.17726 |
| GOBP_POSITIVE_REGULATION_OF_CELL_ACTIVATION | 364 | -0.362889 | -1.494015 | 0 | 0.177917 |
| GOBP_ANION_TRANSMEMBRANE_TRANSPORT | 422 | -0.355981 | -1.493729 | 0 | 0.177841 |
| GOBP_MITOCHONDRIAL_OUTER_MEMBRANE_PERMEABILIZATION | 54 | -0.466028 | -1.49357 | 0.002273 | 0.177664 |
| GOBP_REGULATION_OF_MULTICELLULAR_ORGANISM_GROWTH | 48 | -0.486372 | -1.493338 | 0.027897 | 0.177597 |
| GOBP_UROGENITAL_SYSTEM_DEVELOPMENT | 244 | -0.375712 | -1.492988 | 0.002358 | 0.177619 |
| GOBP_POSITIVE_REGULATION_OF_RECEPTOR_SIGNALING_PATHWAY_VIA_STAT | 37 | -0.496236 | -1.492959 | 0.026432 | 0.177247 |
| GOBP_TRANSEPITHELIAL_TRANSPORT | 24 | -0.551612 | -1.492259 | 0.04329 | 0.177745 |
| GOBP_PULMONARY_VALVE_DEVELOPMENT | 16 | -0.60655 | -1.49201 | 0.048458 | 0.177675 |
| GOBP_POSITIVE_REGULATION_OF_MYELOID_CELL_DIFFERENTIATION | 91 | -0.434493 | -1.49191 | 0.012903 | 0.177432 |
| GOBP_CELL_CYCLE_G1_S_PHASE_TRANSITION | 255 | -0.375203 | -1.491808 | 0.002364 | 0.177151 |
| GOBP_MITOTIC_SISTER_CHROMATID_SEGREGATION | 161 | -0.39462 | -1.491404 | 0.004301 | 0.177387 |
| GOBP_CHAPERONE_MEDIATED_AUTOPHAGY | 16 | -0.615302 | -1.491286 | 0.059448 | 0.177089 |
| GOBP_REGULATION_OF_DNA_BINDING_TRANSCRIPTION_FACTOR_ACTIVITY | 369 | -0.359787 | -1.490986 | 0 | 0.177062 |
| GOBP_REGULATION_OF_FATTY_ACID_METABOLIC_PROCESS | 76 | -0.441421 | -1.4902 | 0.015217 | 0.177671 |
| GOBP_FEMALE_SEX_DIFFERENTIATION | 87 | -0.427421 | -1.489842 | 0.015982 | 0.177727 |
| GOBP_REGULATION_OF_PH | 89 | -0.434664 | -1.4896 | 0.011442 | 0.177627 |
| GOBP_MEMBRANE_REPOLARIZATION | 33 | -0.518877 | -1.489456 | 0.034908 | 0.177424 |
| GOBP_POSITIVE_REGULATION_OF_EPIDERMIS_DEVELOPMENT | 26 | -0.534846 | -1.4883 | 0.046843 | 0.178657 |
| GOBP_REGULATION_OF_LIPID_TRANSPORT | 107 | -0.418281 | -1.487801 | 0.010571 | 0.178912 |
| GOBP_CYTOPLASMIC_PATTERN_RECOGNITION_RECEPTOR_SIGNALING_PATHWAY | 69 | -0.444658 | -1.487209 | 0.019737 | 0.179324 |
| GOBP_PROTEIN_PROCESSING | 173 | -0.38763 | -1.487106 | 0 | 0.179042 |
| GOBP_REGULATION_OF_TRANSMEMBRANE_RECEPTOR_PROTEIN_SERINE_THREONINE_KI | 203 | -0.382077 | -1.486546 | 0 | 0.179375 |
| GOBP_DNA_DEPENDENT_DNA_REPLICATION | 155 | -0.395987 | -1.485556 | 0.002114 | 0.180303 |
| GOBP_BLOOD_VESSEL_REMODELING | 30 | -0.519459 | -1.485058 | 0.036585 | 0.18062 |
| GOBP_NEGATIVE_REGULATION_OF_PROTEIN_MATURATION | 22 | -0.564945 | -1.484927 | 0.038462 | 0.180418 |
| GOBP_RESPONSE_TO_NUTRIENT | 129 | -0.406384 | -1.484704 | 0.013514 | 0.180296 |
| GOBP_DNA_STRAND_ELONGATION_INVOLVED_IN_DNA_REPLICATION | 19 | -0.577699 | -1.48467 | 0.044025 | 0.179963 |
| GOBP_CALCIUM_MEDIATED_SIGNALING_USING_INTRACELLULAR_CALCIUM_SOURCE | 18 | -0.577814 | -1.48458 | 0.060797 | 0.179659 |
| GOBP_MICROTUBULE_ORGANIZING_CENTER_ORGANIZATION | 134 | -0.403497 | -1.483769 | 0.002278 | 0.180388 |
| GOBP_TISSUE_REGENERATION | 56 | -0.457825 | -1.483545 | 0.029228 | 0.180268 |
| GOBP_DIGESTIVE_SYSTEM_PROCESS | 65 | -0.449606 | -1.4835 | 0.027586 | 0.179931 |
| GOBP_TYPE_B_PANCREATIC_CELL_PROLIFERATION | 16 | -0.603294 | -1.483176 | 0.072549 | 0.179956 |
| GOBP_KERATINOCYTE_MIGRATION | 15 | -0.612557 | -1.482896 | 0.068966 | 0.179943 |
| GOBP_REGULATION_OF_MRNA_SPLICING_VIA_SPLICEOSOME | 87 | -0.430076 | -1.482503 | 0.014614 | 0.180047 |
| GOBP_REGULATION_OF_KERATINOCYTE_DIFFERENTIATION | 32 | -0.503691 | -1.482317 | 0.041408 | 0.179888 |
| GOBP_NEGATIVE_REGULATION_OF_DNA_DEPENDENT_DNA_REPLICATION | 17 | -0.598381 | -1.482253 | 0.044681 | 0.17958 |
| GOBP_VIRAL_GENOME_REPLICATION | 116 | -0.410464 | -1.481275 | 0.008949 | 0.180505 |
| GOBP_CLATHRIN_DEPENDENT_ENDOCYTOSIS | 45 | -0.471773 | -1.48098 | 0.03956 | 0.180571 |

| | | | | | |
|---|---|---|---|---|---|
| GOBP_CELL_CELL_SIGNALING_INVOLVED_IN_CARDIAC_CONDUCTION | 22 | 0.421031 | 1.087308 | 0.362403 | 0.648221 |
| GOBP_SCHWANN_CELL_DEVELOPMENT | 25 | 0.402686 | 1.087244 | 0.356863 | 0.64703 |
| GOBP_OLEFINIC_COMPOUND_METABOLIC_PROCESS | 84 | 0.319529 | 1.08705 | 0.308851 | 0.64622 |
| GOBP_POSITIVE_REGULATION_OF_CELL_DEVELOPMENT | 236 | 0.281145 | 1.086334 | 0.240296 | 0.647018 |
| GOBP_NEURON_PROJECTION_EXTENSION | 138 | 0.29941 | 1.085094 | 0.279923 | 0.649441 |
| GOBP_REGULATION_OF_MITOCHONDRIAL_FISSION | 27 | 0.404165 | 1.085081 | 0.338403 | 0.648102 |
| GOBP_REGULATION_OF_NEUROBLAST_PROLIFERATION | 20 | 0.422576 | 1.083851 | 0.344106 | 0.650447 |
| GOBP_INORGANIC_ANION_TRANSPORT | 120 | 0.306066 | 1.083711 | 0.292419 | 0.649467 |
| GOBP_LONG_CHAIN_FATTY_ACID_METABOLIC_PROCESS | 92 | 0.314197 | 1.083284 | 0.286252 | 0.649342 |
| GOBP_MYELIN_MAINTENANCE | 16 | 0.453787 | 1.08328 | 0.359528 | 0.64799 |
| GOBP_ACTIVATION_OF_PHOSPHOLIPASE_C_ACTIVITY | 23 | 0.410302 | 1.08112 | 0.351449 | 0.653146 |
| GOBP_NEGATIVE_REGULATION_OF_CELL_KILLING | 24 | 0.408984 | 1.08044 | 0.328063 | 0.653794 |
| GOBP_POSITIVE_REGULATION_OF_SODIUM_ION_TRANSMEMBRANE_TRANSPORTER_AC | 15 | 0.446567 | 1.080285 | 0.36 | 0.652896 |
| GOBP_REGULATION_OF_PEPTIDE_HORMONE_SECRETION | 152 | 0.296447 | 1.08014 | 0.303371 | 0.651941 |
| GOBP_POSITIVE_REGULATION_OF_NEUROBLAST_PROLIFERATION | 15 | 0.455673 | 1.079782 | 0.353053 | 0.651657 |
| GOBP_POSITIVE_REGULATION_OF_SODIUM_ION_TRANSPORT | 27 | 0.400968 | 1.07935 | 0.355641 | 0.651616 |
| GOBP_PROTEIN_LIPID_COMPLEX_SUBUNIT_ORGANIZATION | 33 | 0.381312 | 1.079317 | 0.336576 | 0.650355 |
| GOBP_CELL_CELL_RECOGNITION | 53 | 0.348198 | 1.078842 | 0.313609 | 0.650427 |
| GOBP_MONOCARBOXYLIC_ACID_BIOSYNTHETIC_PROCESS | 170 | 0.288507 | 1.077469 | 0.283122 | 0.653292 |
| GOBP_PROTEIN_HYDROXYLATION | 26 | 0.390788 | 1.076734 | 0.330961 | 0.654095 |
| GOBP_REGULATION_OF_MYOBLAST_DIFFERENTIATION | 36 | 0.368104 | 1.076488 | 0.327172 | 0.653453 |
| GOBP_CARDIAC_MUSCLE_CELL_ACTION_POTENTIAL | 55 | 0.34637 | 1.075023 | 0.325411 | 0.656578 |
| GOBP_POSITIVE_REGULATION_OF_MACROAUTOPHAGY | 57 | 0.337766 | 1.0731 | 0.330869 | 0.66104 |
| GOBP_NEGATIVE_REGULATION_OF_FAT_CELL_DIFFERENTIATION | 42 | 0.359219 | 1.072725 | 0.341509 | 0.660779 |
| GOBP_INTERLEUKIN_5_PRODUCTION | 16 | 0.442834 | 1.072419 | 0.35249 | 0.660358 |
| GOBP_REGULATION_OF_CELL_FATE_COMMITMENT | 19 | 0.424542 | 1.071978 | 0.382524 | 0.660314 |
| GOBP_NEGATIVE_REGULATION_OF_DEVELOPMENTAL_GROWTH | 86 | 0.31448 | 1.070445 | 0.31903 | 0.663514 |
| GOBP_ACTIVATION_OF_JUN_KINASE_ACTIVITY | 36 | 0.371653 | 1.06995 | 0.346863 | 0.663694 |
| GOBP_EXOCYTIC_PROCESS | 68 | 0.328496 | 1.069803 | 0.316667 | 0.662803 |
| GOBP_CELL_CELL_ADHESION_MEDIATED_BY_CADHERIN | 25 | 0.396917 | 1.069733 | 0.366412 | 0.661667 |
| GOBP_REGULATION_OF_ASTROCYTE_DIFFERENTIATION | 17 | 0.431223 | 1.069231 | 0.39243 | 0.661828 |
| GOBP_NEGATIVE_REGULATION_OF_LYASE_ACTIVITY | 16 | 0.435753 | 1.069161 | 0.356473 | 0.660686 |
| GOBP_VENTRICULAR_CARDIAC_MUSCLE_TISSUE_DEVELOPMENT | 34 | 0.380558 | 1.069125 | 0.368121 | 0.659474 |
| GOBP_CORPUS_CALLOSUM_DEVELOPMENT | 15 | 0.443551 | 1.068848 | 0.395753 | 0.658978 |
| GOBP_ETHER_METABOLIC_PROCESS | 22 | 0.411543 | 1.068359 | 0.373606 | 0.659114 |
| GOBP_ENDOCARDIAL_CUSHION_MORPHOGENESIS | 26 | 0.397104 | 1.064763 | 0.382524 | 0.668636 |
| GOBP_ICOSANOID_METABOLIC_PROCESS | 87 | 0.310761 | 1.064291 | 0.347664 | 0.668725 |
| GOBP_CYTOLYSIS | 19 | 0.422157 | 1.06418 | 0.369942 | 0.667726 |
| GOBP_CENTRAL_NERVOUS_SYSTEM_NEURON_DIFFERENTIATION | 121 | 0.299395 | 1.062445 | 0.336397 | 0.671545 |
| GOBP_COGNITION | 202 | 0.279407 | 1.061095 | 0.32963 | 0.674308 |
| GOBP_REGULATION_OF_LEUKOCYTE_APOPTOTIC_PROCESS | 69 | 0.327334 | 1.060475 | 0.365672 | 0.674796 |
| GOBP_NEUROMUSCULAR_SYNAPTIC_TRANSMISSION | 15 | 0.451592 | 1.060351 | 0.404382 | 0.67389 |
| GOBP_POSITIVE_REGULATION_OF_ENDOTHELIAL_CELL_APOPTOTIC_PROCESS | 16 | 0.436939 | 1.060249 | 0.379958 | 0.672898 |
| GOBP_NEUROMUSCULAR_PROCESS | 58 | 0.33847 | 1.059688 | 0.364504 | 0.673259 |
| GOBP_REGULATION_OF_PROTEIN_TYROSINE_KINASE_ACTIVITY | 83 | 0.315254 | 1.05863 | 0.346369 | 0.675127 |
| GOBP_NEURONAL_ACTION_POTENTIAL | 19 | 0.424418 | 1.05817 | 0.385633 | 0.675241 |
| GOBP_CELLULAR_RESPONSE_TO_VITAMIN | 23 | 0.395758 | 1.057467 | 0.389535 | 0.676026 |
| GOBP_NEURAL_TUBE_PATTERNING | 28 | 0.389641 | 1.056277 | 0.389121 | 0.678296 |
| GOBP_LEUKOCYTE_TETHERING_OR_ROLLING | 25 | 0.40055 | 1.054735 | 0.392661 | 0.681676 |
| GOBP_REGULATION_OF_TRANSLATIONAL_FIDELITY | 20 | 0.410866 | 1.053727 | 0.379242 | 0.683435 |
| GOBP_PIGMENT_CELL_DIFFERENTIATION | 26 | 0.384453 | 1.053482 | 0.359922 | 0.682821 |
| GOBP_ACTIN_FILAMENT_ORGANIZATION | 371 | 0.259411 | 1.052108 | 0.27212 | 0.685692 |
| GOBP_NEGATIVE_REGULATION_OF_INTERFERON_GAMMA_PRODUCTION | 33 | 0.374166 | 1.051352 | 0.378327 | 0.686751 |
| GOBP_T_CELL_APOPTOTIC_PROCESS | 43 | 0.353139 | 1.050993 | 0.384328 | 0.686506 |
| GOBP_CARDIAC_MUSCLE_CELL_CONTRACTION | 50 | 0.343068 | 1.050589 | 0.366541 | 0.686442 |
| GOBP_POST_TRANSLATIONAL_PROTEIN_MODIFICATION | 311 | 0.264261 | 1.045685 | 0.326014 | 0.699957 |
| GOBP_REGULATION_OF_CYCLASE_ACTIVITY | 28 | 0.38813 | 1.045402 | 0.405714 | 0.699467 |
| GOBP_REGULATION_OF_CELL_SUBSTRATE_ADHESION | 175 | 0.28208 | 1.044918 | 0.338889 | 0.699585 |
| GOBP_MEIOTIC_CHROMOSOME_SEGREGATION | 72 | 0.31882 | 1.044171 | 0.365079 | 0.70048 |
| GOBP_REGULATION_OF_CAMP_MEDIATED_SIGNALING | 25 | 0.389671 | 1.043674 | 0.421663 | 0.70069 |
| GOBP_EMBRYONIC_APPENDAGE_MORPHOGENESIS | 81 | 0.312228 | 1.042698 | 0.378676 | 0.702299 |
| GOBP_NEGATIVE_REGULATION_OF_CELLULAR_RESPONSE_TO_GROWTH_FACTOR_STIM | 77 | 0.316236 | 1.042647 | 0.3902 | 0.701138 |
| GOBP_UNSATURATED_FATTY_ACID_METABOLIC_PROCESS | 85 | 0.305813 | 1.0425 | 0.371634 | 0.700296 |
| GOBP_CARDIAC_SEPTUM_DEVELOPMENT | 75 | 0.314111 | 1.042283 | 0.374765 | 0.699657 |
| GOBP_TOXIN_TRANSPORT | 37 | 0.361319 | 1.041562 | 0.380309 | 0.700509 |
| GOBP_RESPONSE_TO_HYDROPEROXIDE | 15 | 0.429879 | 1.040994 | 0.397938 | 0.700967 |
| GOBP_REGULATION_OF_ENDOCRINE_PROCESS | 24 | 0.393227 | 1.039602 | 0.412548 | 0.703968 |
| GOBP_REGULATION_OF_T_CELL_MIGRATION | 37 | 0.352944 | 1.039483 | 0.378428 | 0.703026 |
| GOBP_N_GLYCAN_PROCESSING | 17 | 0.411381 | 1.039357 | 0.412698 | 0.702087 |
| GOBP_MYOBLAST_FUSION | 33 | 0.357373 | 1.0393 | 0.391969 | 0.700944 |
| GOBP_PROTEIN_LOCALIZATION_TO_ENDOPLASMIC_RETICULUM | 143 | 0.286247 | 1.0376 | 0.379648 | 0.704774 |
| GOBP_MACROPHAGE_CYTOKINE_PRODUCTION | 15 | 0.431125 | 1.036951 | 0.414498 | 0.705394 |
| GOBP_REGULATION_OF_ENDOPLASMIC_RETICULUM_UNFOLDED_PROTEIN_RESPONSE | 29 | 0.371824 | 1.036869 | 0.408663 | 0.704327 |
| GOBP_REGULATION_OF_LIPOPROTEIN_LIPASE_ACTIVITY | 15 | 0.439188 | 1.036617 | 0.414365 | 0.703767 |
| GOBP_POSITIVE_REGULATION_OF_CHONDROCYTE_DIFFERENTIATION | 16 | 0.425215 | 1.035233 | 0.411765 | 0.706712 |
| GOBP_SUBSTRATE_DEPENDENT_CELL_MIGRATION | 22 | 0.391932 | 1.035073 | 0.418605 | 0.705895 |
| GOBP_CELLULAR_RESPONSE_TO_CALCIUM_ION | 60 | 0.329281 | 1.034821 | 0.40458 | 0.705372 |
| GOBP_PROTEIN_TRIMERIZATION | 16 | 0.427372 | 1.034707 | 0.420233 | 0.704415 |
| GOBP_NEGATIVE_REGULATION_OF_CELL_PROJECTION_ORGANIZATION | 150 | 0.284055 | 1.033956 | 0.379747 | 0.705388 |
| GOBP_RENAL_SYSTEM_PROCESS | 86 | 0.30223 | 1.033548 | 0.38716 | 0.705314 |
| GOBP_AORTIC_VALVE_MORPHOGENESIS | 23 | 0.395046 | 1.032802 | 0.393657 | 0.706306 |
| GOBP_REGULATION_OF_PEPTIDE_SECRETION | 235 | 0.269272 | 1.03236 | 0.36136 | 0.706332 |
| GOBP_REGULATION_OF_LYASE_ACTIVITY | 30 | 0.36892 | 1.030443 | 0.412331 | 0.710851 |
| GOBP_RESPIRATORY_SYSTEM_PROCESS | 26 | 0.383036 | 1.030173 | 0.416667 | 0.710382 |
| GOBP_DECIDUALIZATION | 15 | 0.428762 | 1.029825 | 0.443478 | 0.710166 |
| GOBP_MEMBRANE_DEPOLARIZATION_DURING_CARDIAC_MUSCLE_CELL_ACTION_POTE | 16 | 0.425781 | 1.02923 | 0.428571 | 0.710673 |
| GOBP_MAINTENANCE_OF_BLOOD_BRAIN_BARRIER | 27 | 0.372615 | 1.029229 | 0.43222 | 0.709395 |
| GOBP_CHONDROITIN_SULFATE_BIOSYNTHETIC_PROCESS | 22 | 0.391929 | 1.028543 | 0.407767 | 0.71021 |
| GOBP_REGULATION_OF_HAIR_CYCLE | 23 | 0.387746 | 1.027873 | 0.430168 | 0.710918 |
| GOBP_NUCLEUS_LOCALIZATION | 21 | 0.400784 | 1.027327 | 0.433198 | 0.711271 |
| GOBP_NEGATIVE_REGULATION_OF_WNT_SIGNALING_PATHWAY | 184 | 0.272138 | 1.026398 | 0.391771 | 0.712842 |
| GOBP_EXTRACELLULAR_MATRIX_DISASSEMBLY | 59 | 0.318557 | 1.026131 | 0.407273 | 0.712392 |
| GOBP_SPROUTING_ANGIOGENESIS | 107 | 0.292932 | 1.025495 | 0.389286 | 0.713022 |
| GOBP_DETECTION_OF_BIOTIC_STIMULUS | 31 | 0.367374 | 1.024608 | 0.41109 | 0.71436 |
| GOBP_RHO_PROTEIN_SIGNAL_TRANSDUCTION | 118 | 0.289246 | 1.02342 | 0.404293 | 0.716568 |
| GOBP_EMBRYONIC_HEMOPOIESIS | 20 | 0.397306 | 1.021916 | 0.437372 | 0.719816 |
| GOBP_CYTOPLASMIC_TRANSLATIONAL_INITIATION | 34 | 0.358965 | 1.021559 | 0.40691 | 0.719595 |
| GOBP_HYALURONAN_METABOLIC_PROCESS | 32 | 0.361363 | 1.020916 | 0.418561 | 0.720331 |
| GOBP_VITAMIN_BIOSYNTHETIC_PROCESS | 17 | 0.419801 | 1.020504 | 0.433333 | 0.720255 |
| GOBP_POSITIVE_REGULATION_OF_BLOOD_PRESSURE | 22 | 0.39093 | 1.020068 | 0.435294 | 0.720355 |
| GOBP_REGULATION_OF_MITOCHONDRIAL_MEMBRANE_POTENTIAL | 65 | 0.309009 | 1.018755 | 0.413979 | 0.72302 |
| GOBP_CRANIAL_NERVE_DEVELOPMENT | 31 | 0.361607 | 1.018506 | 0.448617 | 0.722529 |
| GOBP_INTERLEUKIN_13_PRODUCTION | 18 | 0.403463 | 1.01846 | 0.452158 | 0.721415 |
| GOBP_DENDRITE_DEVELOPMENT | 199 | 0.270208 | 1.018266 | 0.411335 | 0.720778 |
| GOBP_HOMEOSTASIS_OF_NUMBER_OF_CELLS_WITHIN_A_TISSUE | 23 | 0.391583 | 1.01704 | 0.440613 | 0.723206 |
| GOBP_ORGANIC_ACID_BIOSYNTHETIC_PROCESS | 263 | 0.258019 | 1.015573 | 0.426396 | 0.726418 |
| GOBP_REGULATION_OF_CHOLESTEROL_METABOLIC_PROCESS | 49 | 0.327504 | 1.015085 | 0.42243 | 0.726611 |
| GOBP_PROTEIN_O_LINKED_GLYCOSYLATION | 88 | 0.300113 | 1.014491 | 0.420765 | 0.727157 |
| GOBP_NEGATIVE_REGULATION_OF_B_CELL_PROLIFERATION | 17 | 0.410964 | 1.014026 | 0.444846 | 0.727236 |
| GOBP_NEGATIVE_REGULATION_OF_EPITHELIAL_CELL_APOPTOTIC_PROCESS | 36 | 0.350035 | 1.012935 | 0.443787 | 0.729327 |
| GOBP_POSITIVE_REGULATION_OF_CARDIAC_MUSCLE_CELL_PROLIFERATION | 21 | 0.393082 | 1.012842 | 0.424419 | 0.728339 |
| GOBP_NEGATIVE_REGULATION_OF_ORGAN_GROWTH | 28 | 0.372825 | 1.011928 | 0.438078 | 0.729775 |
| GOBP_NATURAL_KILLER_CELL_ACTIVATION | 62 | 0.315996 | 1.011588 | 0.431942 | 0.729536 |
| GOBP_FATTY_ACID_BIOSYNTHETIC_PROCESS | 130 | 0.278094 | 1.009969 | 0.426199 | 0.733093 |
| GOBP_POSITIVE_REGULATION_OF_NEURAL_PRECURSOR_CELL_PROLIFERATION | 32 | 0.355758 | 1.009762 | 0.425688 | 0.732454 |
| GOBP_MUCOPOLYSACCHARIDE_METABOLIC_PROCESS | 96 | 0.294642 | 1.009689 | 0.418692 | 0.731411 |
| GOBP_POSTSYNAPTIC_MEMBRANE_ORGANIZATION | 29 | 0.362257 | 1.009415 | 0.460803 | 0.731005 |
| GOBP_FAT_SOLUBLE_VITAMIN_METABOLIC_PROCESS | 29 | 0.36561 | 1.008931 | 0.45 | 0.731245 |
| GOBP_SODIUM_ION_TRANSPORT | 164 | 0.2718 | 1.008891 | 0.435101 | 0.73012 |
| GOBP_REGULATION_OF_ACTIN_FILAMENT_BUNDLE_ASSEMBLY | 84 | 0.303227 | 1.008359 | 0.444444 | 0.73046 |
| GOBP_INNER_EAR_AUDITORY_RECEPTOR_CELL_DIFFERENTIATION | 28 | 0.35865 | 1.00815 | 0.44 | 0.729854 |
| GOBP_NEGATIVE_REGULATION_OF_BMP_SIGNALING_PATHWAY | 40 | 0.334028 | 1.007995 | 0.418651 | 0.729062 |
| GOBP_POSITIVE_REGULATION_OF_MITOCHONDRIAL_TRANSLATION | 16 | 0.413414 | 1.007544 | 0.453307 | 0.729146 |
| GOBP_DNA_CATABOLIC_PROCESS_ENDONUCLEOLYTIC | 27 | 0.380475 | 1.007 | 0.469062 | 0.729518 |
| GOBP_PROTEIN_LOCALIZATION_TO_MICROTUBULE | 17 | 0.397337 | 1.006855 | 0.434363 | 0.728712 |
| GOBP_OLFACTORY_LOBE_DEVELOPMENT | 18 | 0.403571 | 1.006605 | 0.424901 | 0.728257 |
| GOBP_SINGLE_FERTILIZATION | 96 | 0.292797 | 1.005621 | 0.438261 | 0.730022 |
| GOBP_ANTIGEN_PROCESSING_AND_PRESENTATION_OF_PEPTIDE_ANTIGEN | 184 | 0.269361 | 1.00469 | 0.432862 | 0.731552 |
| GOBP_REGULATION_OF_THE_FORCE_OF_HEART_CONTRACTION | 17 | 0.407544 | 1.00468 | 0.473373 | 0.73036 |
| GOBP_AORTA_MORPHOGENESIS | 25 | 0.373575 | 1.004184 | 0.434608 | 0.730535 |
| GOBP_BLASTOCYST_FORMATION | 28 | 0.367544 | 1.004052 | 0.452282 | 0.729705 |
| GOBP_STRESS_FIBER_ASSEMBLY | 89 | 0.291968 | 1.003978 | 0.431227 | 0.728697 |
| GOBP_CARDIAC_MUSCLE_CELL_PROLIFERATION | 43 | 0.332162 | 1.003436 | 0.476723 | 0.72908 |
| GOBP_REGULATION_OF_SMAD_PROTEIN_SIGNAL_TRANSDUCTION | 22 | 0.379915 | 1.001823 | 0.463415 | 0.732617 |
| GOBP_EMBRYONIC_CAMERA_TYPE_EYE_DEVELOPMENT | 23 | 0.380843 | 1.001194 | 0.473585 | 0.733249 |
| GOBP_PEPTIDYL_TYROSINE_AUTOPHOSPHORYLATION | 16 | 0.401943 | 1.001068 | 0.45776 | 0.732399 |
| GOBP_SCHWANN_CELL_DIFFERENTIATION | 33 | 0.352341 | 1.00093 | 0.482213 | 0.731598 |
| GOBP_REGULATION_OF_SODIUM_ION_TRANSMEMBRANE_TRANSPORTER_ACTIVITY | 40 | 0.334975 | 1.00028 | 0.451613 | 0.732299 |
| GOBP_REGULATION_OF_CARDIAC_MUSCLE_CELL_ACTION_POTENTIAL | 20 | 0.390794 | 0.999695 | 0.472119 | 0.732819 |
| GOBP_STEROL_HOMEOSTASIS | 60 | 0.316531 | 0.999348 | 0.454023 | 0.732632 |
| GOBP_EMBRYONIC_CRANIAL_SKELETON_MORPHOGENESIS | 33 | 0.351718 | 0.999243 | 0.441392 | 0.731709 |
| GOBP_RETINOIC_ACID_RECEPTOR_SIGNALING_PATHWAY | 23 | 0.371539 | 0.99896 | 0.465421 | 0.731335 |
| GOBP_ACTIN_FILAMENT_BUNDLE_ORGANIZATION | 132 | 0.276452 | 0.998763 | 0.447788 | 0.730674 |
| GOBP_EPITHELIAL_TO_MESENCHYMAL_TRANSITION | 124 | 0.28033 | 0.998378 | 0.453069 | 0.730568 |
| GOBP_ANTIGEN_PROCESSING_AND_PRESENTATION_OF_ENDOGENOUS_PEPTIDE_ANTI( | 17 | 0.399109 | 0.997803 | 0.489837 | 0.731057 |
| GOBP_VESICLE_DOCKING | 61 | 0.313201 | 0.996282 | 0.450485 | 0.734379 |
| GOBP_REGULATION_OF_RELEASE_OF_SEQUESTERED_CALCIUM_ION_INTO_CYTOSOL | 59 | 0.308079 | 0.995188 | 0.461233 | 0.73635 |
| GOBP_REGULATION_OF_HORMONE_SECRETION | 188 | 0.267856 | 0.994702 | 0.481482 | 0.736523 |
| GOBP_CELLULAR_SENESCENCE | 64 | 0.31043 | 0.994594 | 0.439164 | 0.735646 |
| GOBP_SPERM_AXONEME_ASSEMBLY | 18 | 0.401488 | 0.992663 | 0.468451 | 0.740185 |
| GOBP_VASOCONSTRICTION | 51 | 0.32184 | 0.992012 | 0.456814 | 0.740903 |
| GOBP_VIRAL_TRANSLATION | 15 | 0.406615 | 0.991876 | 0.47048 | 0.740117 |
| GOBP_POSITIVE_REGULATION_OF_T_CELL_MIGRATION | 25 | 0.370987 | 0.990446 | 0.481905 | 0.743121 |
| GOBP_CARDIAC_MUSCLE_TISSUE_MORPHOGENESIS | 37 | 0.338655 | 0.990284 | 0.486166 | 0.7424 |
| GOBP_POSITIVE_REGULATION_OF_INTERFERON_GAMMA_PRODUCTION | 58 | 0.311242 | 0.98956 | 0.452158 | 0.743366 |
| GOBP_DNA_CATABOLIC_PROCESS | 34 | 0.344085 | 0.988109 | 0.498024 | 0.746393 |
| GOBP_PINOCYTOSIS | 21 | 0.383019 | 0.987075 | 0.493902 | 0.748315 |

| | | | | | |
|---|---|---|---|---|---|
| GOBP_REGULATION_OF_HYDROGEN_PEROXIDE_METABOLIC_PROCESS | 18 | -0.59758 | -1.479902 | 0.03854 | 0.181575 |
| GOBP_CYTOKINETIC_PROCESS | 38 | -0.486598 | -1.479725 | 0.038462 | 0.181445 |
| GOBP_REGULATION_OF_STRESS_ACTIVATED_PROTEIN_KINASE_SIGNALING_CASCADE | 159 | -0.396371 | -1.479719 | 0.006711 | 0.181066 |
| GOBP_ENERGY_RESERVE_METABOLIC_PROCESS | 72 | -0.437126 | -1.478164 | 0.017058 | 0.182849 |
| GOBP_NEGATIVE_REGULATION_OF_OSTEOCLAST_DIFFERENTIATION | 23 | -0.549614 | -1.477459 | 0.046512 | 0.183391 |
| GOBP_HETEROTYPIC_CELL_CELL_ADHESION | 49 | -0.476494 | -1.477092 | 0.033186 | 0.183439 |
| GOBP_REGULATION_OF_FATTY_ACID_TRANSPORT | 39 | -0.490672 | -1.476706 | 0.033264 | 0.18352 |
| GOBP_NEPHRON_DEVELOPMENT | 98 | -0.412598 | -1.476616 | 0.010571 | 0.18326 |
| GOBP_NEGATIVE_REGULATION_OF_MUSCLE_CELL_DIFFERENTIATION | 49 | -0.475847 | -1.476242 | 0.033755 | 0.183413 |
| GOBP_REGULATION_OF_CLATHRIN_DEPENDENT_ENDOCYTOSIS | 18 | -0.585216 | -1.475683 | 0.070213 | 0.183772 |
| GOBP_REGULATION_OF_EPIDERMIS_DEVELOPMENT | 55 | -0.455941 | -1.475266 | 0.032397 | 0.18394 |
| GOBP_AMINO_ACID_IMPORT | 38 | -0.499481 | -1.475105 | 0.030801 | 0.18377 |
| GOBP_T_CELL_HOMEOSTASIS | 34 | -0.505794 | -1.474497 | 0.050847 | 0.184189 |
| GOBP_DEVELOPMENT_OF_PRIMARY_FEMALE_SEXUAL_CHARACTERISTICS | 75 | -0.432633 | -1.474276 | 0.012658 | 0.184162 |
| GOBP_FIBRINOLYSIS | 16 | -0.59636 | -1.474011 | 0.049702 | 0.184138 |
| GOBP_POSITIVE_REGULATION_OF_CELL_CYCLE_PROCESS | 258 | -0.370056 | -1.473524 | 0.002545 | 0.184373 |
| GOBP_MULTICELLULAR_ORGANISMAL_HOMEOSTASIS | 416 | -0.350684 | -1.473188 | 0 | 0.184434 |
| GOBP_POSITIVE_REGULATION_OF_LIPID_KINASE_ACTIVITY | 33 | -0.512517 | -1.473122 | 0.057613 | 0.184146 |
| GOBP_ALTERNATIVE_MRNA_SPLICING_VIA_SPLICEOSOME | 62 | -0.446712 | -1.472899 | 0.01919 | 0.184059 |
| GOBP_REGULATION_OF_SUBSTRATE_ADHESION_DEPENDENT_CELL_SPREADING | 49 | -0.472065 | -1.471914 | 0.023707 | 0.184961 |
| GOBP_ADAPTIVE_THERMOGENESIS | 128 | -0.397437 | -1.47147 | 0.004608 | 0.185177 |
| GOBP_FOAM_CELL_DIFFERENTIATION | 25 | -0.546412 | -1.470807 | 0.058577 | 0.185592 |
| GOBP_NEGATIVE_REGULATION_OF_SMOOTH_MUSCLE_CELL_DIFFERENTIATION | 16 | -0.613614 | -1.470343 | 0.066955 | 0.18579 |
| GOBP_NEGATIVE_REGULATION_OF_LIPID_STORAGE | 17 | -0.579593 | -1.470237 | 0.057018 | 0.185544 |
| GOBP_MITOTIC_CYTOKINESIS | 69 | -0.436712 | -1.469876 | 0.01919 | 0.185694 |
| GOBP_REGULATION_OF_AMYLOID_BETA_CLEARANCE | 15 | -0.610773 | -1.469831 | 0.083658 | 0.185366 |
| GOBP_POSITIVE_REGULATION_OF_POTASSIUM_ION_TRANSMEMBRANE_TRANSPORT | 28 | -0.518465 | -1.469829 | 0.046512 | 0.184994 |
| GOBP_REGULATION_OF_INFLAMMATORY_RESPONSE | 272 | -0.365571 | -1.469302 | 0 | 0.185276 |
| GOBP_ORGANIC_HYDROXY_COMPOUND_CATABOLIC_PROCESS | 56 | -0.461811 | -1.468862 | 0.022587 | 0.185503 |
| GOBP_NEGATIVE_REGULATION_OF_POTASSIUM_ION_TRANSMEMBRANE_TRANSPORTER_ACTIVITY | 43 | -0.488246 | -1.468386 | 0.032967 | 0.185793 |
| GOBP_DNA_STRAND_ELONGATION | 26 | -0.536415 | -1.467851 | 0.048936 | 0.186208 |
| GOBP_IONOTROPIC_GLUTAMATE_RECEPTOR_SIGNALING_PATHWAY | 16 | -0.597333 | -1.467843 | 0.070994 | 0.185857 |
| GOBP_CELLULAR_RESPONSE_TO_HYDROGEN_PEROXIDE | 76 | -0.439163 | -1.467601 | 0.011442 | 0.185828 |
| GOBP_POSITIVE_REGULATION_OF_INTERLEUKIN_10_PRODUCTION | 35 | -0.50261 | -1.466685 | 0.041068 | 0.186587 |
| GOBP_G_PROTEIN_COUPLED_ACETYLCHOLINE_RECEPTOR_SIGNALING_PATHWAY | 15 | -0.608341 | -1.46618 | 0.069869 | 0.186876 |
| GOBP_POSITIVE_REGULATION_OF_LIPID_TRANSPORT | 67 | -0.440377 | -1.465893 | 0.021834 | 0.186898 |
| GOBP_NEGATIVE_REGULATION_OF_T_CELL_MEDIATED_IMMUNITY | 21 | -0.561613 | -1.465585 | 0.052154 | 0.186911 |
| GOBP_REGULATION_OF_ACUTE_INFLAMMATORY_RESPONSE | 39 | -0.492923 | -1.464712 | 0.033898 | 0.187738 |
| GOBP_NEGATIVE_REGULATION_OF_CELL_CYCLE_PHASE_TRANSITION | 235 | -0.370517 | -1.464462 | 0.004454 | 0.187684 |
| GOBP_INTERLEUKIN_6_PRODUCTION | 125 | -0.40225 | -1.464255 | 0.006536 | 0.18758 |
| GOBP_REGULATION_OF_HORMONE_BIOSYNTHETIC_PROCESS | 16 | -0.601159 | -1.463434 | 0.057325 | 0.188387 |
| GOBP_SENSORY_PERCEPTION_OF_PAIN | 59 | -0.443943 | -1.463157 | 0.036559 | 0.188447 |
| GOBP_BASE_CONVERSION_OR_SUBSTITUTION_EDITING | 16 | -0.60235 | -1.463105 | 0.066955 | 0.188137 |
| GOBP_RESPONSE_TO_PEPTIDE_HORMONE | 365 | -0.356429 | -1.462445 | 0 | 0.188676 |
| GOBP_LEUKOCYTE_HOMEOSTASIS | 74 | -0.42859 | -1.462267 | 0.021186 | 0.188555 |
| GOBP_REGULATION_OF_FATTY_ACID_OXIDATION | 31 | -0.516917 | -1.462122 | 0.036797 | 0.188404 |
| GOBP_INTERLEUKIN_1_PRODUCTION | 94 | -0.417125 | -1.462032 | 0.010917 | 0.188146 |
| GOBP_RECEPTOR_LOCALIZATION_TO_SYNAPSE | 43 | -0.473425 | -1.460126 | 0.04251 | 0.190271 |
| GOBP_DEVELOPMENTAL_MATURATION | 212 | -0.374813 | -1.460116 | 0.004566 | 0.189914 |
| GOBP_LONG_CHAIN_FATTY_ACID_IMPORT_INTO_CELL | 16 | -0.597907 | -1.459607 | 0.076605 | 0.190268 |
| GOBP_POSITIVE_REGULATION_OF_PROTEIN_KINASE_B_SIGNALING | 134 | -0.39129 | -1.459275 | 0.013043 | 0.190332 |
| GOBP_REGULATION_OF_DNA_BINDING | 103 | -0.412779 | -1.459019 | 0.004545 | 0.190304 |
| GOBP_POSITIVE_REGULATION_OF_EPITHELIAL_CELL_DIFFERENTIATION | 40 | -0.484018 | -1.458917 | 0.037611 | 0.19009 |
| GOBP_CELLULAR_RESPONSE_TO_ORGANIC_CYCLIC_COMPOUND | 441 | -0.34555 | -1.457911 | 0 | 0.191176 |
| GOBP_POSITIVE_REGULATION_OF_ERYTHROCYTE_DIFFERENTIATION | 32 | -0.506884 | -1.457687 | 0.036797 | 0.191065 |
| GOBP_BIOLOGICAL_PROCESS_INVOLVED_IN_INTERACTION_WITH_SYMBIONT | 72 | -0.434794 | -1.457495 | 0.019694 | 0.190996 |
| GOBP_AUTOPHAGOSOME_ORGANIZATION | 96 | -0.411346 | -1.457066 | 0.019355 | 0.191264 |
| GOBP_CELLULAR_METABOLIC_COMPOUND_SALVAGE | 32 | -0.500878 | -1.455922 | 0.045259 | 0.192456 |
| GOBP_POSITIVE_REGULATION_OF_JNK_CASCADE | 79 | -0.425636 | -1.45589 | 0.012685 | 0.192142 |
| GOBP_REGULATION_OF_ATP_BIOSYNTHETIC_PROCESS | 20 | -0.557205 | -1.455399 | 0.061441 | 0.192415 |
| GOBP_CELLULAR_RESPONSE_TO_CAMP | 44 | -0.487392 | -1.455056 | 0.041408 | 0.192536 |
| GOBP_CELLULAR_RESPONSE_TO_CHEMICAL_STRESS | 299 | -0.356087 | -1.455047 | 0 | 0.192181 |
| GOBP_POSITIVE_REGULATION_OF_BONE_MINERALIZATION | 30 | -0.508911 | -1.454618 | 0.052521 | 0.192377 |
| GOBP_POSITIVE_REGULATION_OF_PEPTIDYL_LYSINE_ACETYLATION | 31 | -0.510362 | -1.454037 | 0.045643 | 0.192759 |
| GOBP_REGULATION_OF_DIGESTIVE_SYSTEM_PROCESS | 24 | -0.535329 | -1.453479 | 0.064718 | 0.193326 |
| GOBP_LIPID_DROPLET_ORGANIZATION | 24 | -0.55243 | -1.452767 | 0.072805 | 0.193958 |
| GOBP_REPRODUCTIVE_SYSTEM_DEVELOPMENT | 319 | -0.357296 | -1.452735 | 0 | 0.193647 |
| GOBP_NEGATIVE_REGULATION_OF_MUSCLE_CELL_APOPTOTIC_PROCESS | 27 | -0.519564 | -1.45264 | 0.053079 | 0.193435 |
| GOBP_POSITIVE_REGULATION_OF_MYELOID_LEUKOCYTE_DIFFERENTIATION | 48 | -0.473771 | -1.452511 | 0.03913 | 0.193195 |
| GOBP_REGULATION_OF_CATION_CHANNEL_ACTIVITY | 123 | -0.396046 | -1.452266 | 0.007968 | 0.193185 |
| GOBP_ACUTE_INFLAMMATORY_RESPONSE_TO_ANTIGENIC_STIMULUS | 19 | -0.582604 | -1.452114 | 0.058577 | 0.193059 |
| GOBP_PHAGOSOME_ACIDIFICATION | 25 | -0.536967 | -1.451592 | 0.064583 | 0.193499 |
| GOBP_ASYMMETRIC_CELL_DIVISION | 15 | -0.602441 | -1.450097 | 0.062112 | 0.195298 |
| GOBP_REGULATION_OF_NEURON_DIFFERENTIATION | 130 | -0.391469 | -1.45001 | 0.004695 | 0.195092 |
| GOBP_POSITIVE_REGULATION_OF_T_CELL_CYTOKINE_PRODUCTION | 18 | -0.573486 | -1.449988 | 0.068273 | 0.194764 |
| GOBP_PROTEIN_REFOLDING | 22 | -0.547082 | -1.449827 | 0.061983 | 0.194618 |
| GOBP_POSITIVE_REGULATION_OF_MUSCLE_CONTRACTION | 29 | -0.519084 | -1.44979 | 0.046709 | 0.19431 |
| GOBP_RESPONSE_TO_TRANSFORMING_GROWTH_FACTOR_BETA | 215 | -0.365336 | -1.449779 | 0 | 0.193974 |
| GOBP_INTRINSIC_APOPTOTIC_SIGNALING_PATHWAY | 261 | -0.361846 | -1.449527 | 0.002347 | 0.193964 |
| GOBP_CELLULAR_RESPONSE_TO_TOPOLOGICALLY_INCORRECT_PROTEIN | 158 | -0.37918 | -1.449403 | 0.002212 | 0.193763 |
| GOBP_NEGATIVE_REGULATION_OF_DNA_REPLICATION | 30 | -0.516525 | -1.449397 | 0.060484 | 0.193427 |
| GOBP_POSITIVE_REGULATION_OF_MRNA_METABOLIC_PROCESS | 75 | -0.429993 | -1.449117 | 0.022272 | 0.193432 |
| GOBP_NEGATIVE_REGULATION_OF_SIGNAL_TRANSDUCTION_IN_ABSENCE_OF_LIGAND | 27 | -0.517198 | -1.448941 | 0.048421 | 0.193325 |
| GOBP_REGULATION_OF_KIDNEY_DEVELOPMENT | 18 | -0.570661 | -1.448668 | 0.068323 | 0.193348 |
| GOBP_REGULATION_OF_CELL_CYCLE_PHASE_TRANSITION | 421 | -0.347539 | -1.448064 | 0 | 0.193808 |
| GOBP_REGULATION_OF_DNA_DEPENDENT_DNA_REPLICATION | 47 | -0.471466 | -1.447024 | 0.038117 | 0.1949 |
| GOBP_RESPONSE_TO_TYPE_I_INTERFERON | 79 | -0.419477 | -1.446327 | 0.023707 | 0.195461 |
| GOBP_SPLICEOSOMAL_TRI_SNRNP_COMPLEX_ASSEMBLY | 23 | -0.546785 | -1.446293 | 0.065306 | 0.195141 |
| GOBP_POSITIVE_REGULATION_OF_ERK1_AND_ERK2_CASCADE | 150 | -0.389397 | -1.445711 | 0.006593 | 0.195634 |
| GOBP_DNA_REPLICATION_INITIATION | 39 | -0.487415 | -1.445295 | 0.045553 | 0.195906 |
| GOBP_DETOXIFICATION | 110 | -0.403148 | -1.444807 | 0.007009 | 0.196237 |
| GOBP_ENDOPLASMIC_RETICULUM_MANNOSE_TRIMMING | 16 | -0.591026 | -1.443795 | 0.066098 | 0.197234 |
| GOBP_REGULATION_OF_DNA_TEMPLATED_TRANSCRIPTION_INITIATION | 39 | -0.483088 | -1.443716 | 0.039337 | 0.197026 |
| GOBP_POSITIVE_REGULATION_OF_MEMBRANE_PERMEABILITY | 62 | -0.444276 | -1.443694 | 0.026726 | 0.196708 |
| GOBP_MAINTENANCE_OF_CELL_POLARITY | 17 | -0.575412 | -1.443145 | 0.066798 | 0.197134 |
| GOBP_NEGATIVE_REGULATION_OF_EPITHELIAL_TO_MESENCHYMAL_TRANSITION | 30 | -0.508937 | -1.443029 | 0.052846 | 0.196952 |
| GOBP_PYRIDINE_CONTAINING_COMPOUND_BIOSYNTHETIC_PROCESS | 30 | -0.509467 | -1.442568 | 0.044543 | 0.197342 |
| GOBP_FC_EPSILON_RECEPTOR_SIGNALING_PATHWAY | 158 | -0.38234 | -1.441436 | 0.00655 | 0.198468 |
| GOBP_REGULATION_OF_TYPE_2_IMMUNE_RESPONSE | 27 | -0.521764 | -1.440829 | 0.066532 | 0.199066 |
| GOBP_RESPONSE_TO_FIBROBLAST_GROWTH_FACTOR | 110 | -0.402175 | -1.440725 | 0.025114 | 0.198869 |
| GOBP_POSITIVE_REGULATION_OF_DNA_BINDING_TRANSCRIPTION_FACTOR_ACTIVITY | 230 | -0.363984 | -1.440674 | 0.002299 | 0.198575 |
| GOBP_EMBRYONIC_ORGAN_MORPHOGENESIS | 201 | -0.365749 | -1.440131 | 0 | 0.199005 |
| GOBP_CELLULAR_GLUCAN_METABOLIC_PROCESS | 67 | -0.44041 | -1.439919 | 0.031983 | 0.198937 |
| GOBP_CELLULAR_RESPONSE_TO_HORMONE_STIMULUS | 498 | -0.340923 | -1.439533 | 0 | 0.199079 |
| GOBP_POSITIVE_REGULATION_OF_MITOTIC_CELL_CYCLE | 103 | -0.403804 | -1.439501 | 0.018648 | 0.198778 |
| GOBP_NEGATIVE_REGULATION_OF_STRIATED_MUSCLE_CELL_APOPTOTIC_PROCESS | 17 | -0.591688 | -1.438797 | 0.057082 | 0.199439 |
| GOBP_GRANULOCYTE_DIFFERENTIATION | 29 | -0.505755 | -1.437068 | 0.05726 | 0.201584 |
| GOBP_REGULATION_OF_RNA_SPLICING | 129 | -0.391665 | -1.436824 | 0.010799 | 0.201611 |
| GOBP_REGULATION_OF_OSSIFICATION | 83 | -0.417373 | -1.436106 | 0.030108 | 0.202322 |
| GOBP_POSTSYNAPTIC_SIGNAL_TRANSDUCTION | 26 | -0.519085 | -1.436003 | 0.059448 | 0.202114 |
| GOBP_REGULATION_OF_LYMPHOCYTE_ACTIVATION | 452 | -0.343378 | -1.435687 | 0 | 0.202196 |
| GOBP_EMBRYONIC_ORGAN_DEVELOPMENT | 315 | -0.355113 | -1.434842 | 0 | 0.203075 |
| GOBP_TUMOR_NECROSIS_FACTOR_SUPERFAMILY_CYTOKINE_PRODUCTION | 128 | -0.389524 | -1.434618 | 0.011848 | 0.203044 |
| GOBP_RESPONSE_TO_METAL_ION | 272 | -0.356671 | -1.434252 | 0 | 0.203194 |
| GOBP_SECOND_MESSENGER_MEDIATED_SIGNALING | 230 | -0.363127 | -1.434 | 0.002309 | 0.203189 |
| GOBP_REGULATION_OF_PEPTIDYL_TYROSINE_PHOSPHORYLATION | 210 | -0.363007 | -1.433204 | 0.002242 | 0.203955 |
| GOBP_NEGATIVE_REGULATION_OF_CELL_CYCLE_PROCESS | 314 | -0.348139 | -1.433141 | 0.00237 | 0.203658 |
| GOBP_POSITIVE_REGULATION_OF_LEUKOCYTE_PROLIFERATION | 131 | -0.389035 | -1.432295 | 0.011111 | 0.20457 |
| GOBP_REGULATION_OF_CAMP_DEPENDENT_PROTEIN_KINASE_ACTIVITY | 18 | -0.564736 | -1.432262 | 0.076923 | 0.204254 |
| GOBP_REGULATION_OF_OLIGODENDROCYTE_DIFFERENTIATION | 28 | -0.509046 | -1.432232 | 0.067538 | 0.203952 |
| GOBP_RESPONSE_TO_PROTOZOAN | 20 | -0.549544 | -1.432033 | 0.066667 | 0.203926 |
| GOBP_DNA_REPLICATION_CHECKPOINT | 17 | -0.578285 | -1.431813 | 0.066132 | 0.203891 |
| GOBP_CELLULAR_CARBOHYDRATE_METABOLIC_PROCESS | 250 | -0.357388 | -1.430399 | 0.002155 | 0.205535 |
| GOBP_MYELOID_CELL_APOPTOTIC_PROCESS | 23 | -0.527285 | -1.429853 | 0.070393 | 0.205956 |
| GOBP_POSITIVE_REGULATION_OF_ESTABLISHMENT_OF_PROTEIN_LOCALIZATION_TO_MITC | 56 | -0.447371 | -1.429266 | 0.038618 | 0.206475 |
| GOBP_INTERFERON_BETA_PRODUCTION | 51 | -0.451392 | -1.428737 | 0.04908 | 0.206871 |
| GOBP_EMBRYONIC_MORPHOGENESIS | 437 | -0.342463 | -1.426937 | 0.002427 | 0.209127 |
| GOBP_SINGLE_STRANDED_VIRAL_RNA_REPLICATION_VIA_DOUBLE_STRANDED_DNA_INTE | 20 | -0.531736 | -1.426702 | 0.079665 | 0.209073 |
| GOBP_INTESTINAL_EPITHELIAL_CELL_DIFFERENTIATION | 17 | -0.573909 | -1.426206 | 0.074074 | 0.209474 |
| GOBP_C4_DICARBOXYLATE_TRANSPORT | 20 | -0.546645 | -1.425933 | 0.071584 | 0.209471 |
| GOBP_SYNAPTIC_MEMBRANE_ADHESION | 15 | -0.578385 | -1.425442 | 0.076433 | 0.209759 |
| GOBP_REGULATION_OF_CELL_MORPHOGENESIS_INVOLVED_IN_DIFFERENTIATION | 83 | -0.414998 | -1.425263 | 0.026432 | 0.209689 |
| GOBP_TYROSINE_PHOSPHORYLATION_OF_STAT_PROTEIN | 67 | -0.423161 | -1.425191 | 0.044743 | 0.209464 |
| GOBP_VASCULAR_ASSOCIATED_SMOOTH_MUSCLE_CELL_MIGRATION | 18 | -0.555563 | -1.425188 | 0.072805 | 0.209118 |
| GOBP_REGULATION_OF_TRANSCRIPTION_ELONGATION_FROM_RNA_POLYMERASE_II_PRO | 29 | -0.505856 | -1.423903 | 0.080169 | 0.21052 |
| GOBP_MITOTIC_DNA_INTEGRITY_CHECKPOINT | 103 | -0.400394 | -1.423172 | 0.016632 | 0.211136 |
| GOBP_REGULATION_OF_TRANSLATIONAL_INITIATION_IN_RESPONSE_TO_STRESS | 15 | -0.57517 | -1.422702 | 0.084388 | 0.21146 |
| GOBP_MITOTIC_CYTOKINETIC_PROCESS | 23 | -0.53923 | -1.421304 | 0.067901 | 0.213181 |
| GOBP_POSITIVE_REGULATION_OF_LEUKOCYTE_CHEMOTAXIS | 76 | -0.417161 | -1.421228 | 0.035165 | 0.212957 |
| GOBP_VACUOLAR_ACIDIFICATION | 22 | -0.535137 | -1.421219 | 0.065315 | 0.212614 |
| GOBP_RESPONSE_TO_VIRUS | 297 | -0.352586 | -1.41867 | 0.002257 | 0.215799 |
| GOBP_METAL_ION_HOMEOSTASIS | 492 | -0.335299 | -1.418204 | 0 | 0.21618 |
| GOBP_CELLULAR_RESPONSE_TO_LIPID | 447 | -0.33716 | -1.417876 | 0.00241 | 0.216345 |
| GOBP_ATTACHMENT_OF_SPINDLE_MICROTUBULES_TO_KINETOCHORE | 35 | -0.47496 | -1.417737 | 0.06405 | 0.21616 |
| GOBP_MOTILE_CILIUM_ASSEMBLY | 19 | -0.55855 | -1.417683 | 0.083333 | 0.215905 |
| GOBP_EMBRYONIC_PLACENTA_MORPHOGENESIS | 19 | -0.540296 | -1.417188 | 0.068966 | 0.216284 |
| GOBP_DNA_REPLICATION | 272 | -0.350668 | -1.41564 | 0.002193 | 0.218211 |
| GOBP_NEGATIVE_REGULATION_OF_PLASMA_MEMBRANE_BOUNDED_CELL_PROJECTION_ | 27 | -0.519295 | -1.415557 | 0.072687 | 0.218009 |
| GOBP_REGULATION_OF_EXTRINSIC_APOPTOTIC_SIGNALING_PATHWAY_IN_ABSENCE_OF_ | 36 | -0.489901 | -1.415065 | 0.060797 | 0.218348 |
| GOBP_MICROTUBULE_POLYMERIZATION | 70 | -0.420397 | -1.41482 | 0.039911 | 0.218362 |
| GOBP_NEGATIVE_REGULATION_OF_PROTEIN_ACETYLATION | 19 | -0.560033 | -1.414037 | 0.079108 | 0.21927 |
| GOBP_RESPONSE_TO_ANGIOTENSIN | 19 | -0.545629 | -1.413909 | 0.077236 | 0.219071 |
| GOBP_RESPONSE_TO_INTERFERON_BETA | 26 | -0.503681 | -1.41335 | 0.071429 | 0.219614 |
| GOBP_B_CELL_PROLIFERATION | 77 | -0.410335 | -1.412439 | 0.028747 | 0.220655 |
| GOBP_CALCIUM_MEDIATED_SIGNALING | 161 | -0.369985 | -1.412348 | 0.019737 | 0.22042 |
| GOBP_PRI_MIRNA_TRANSCRIPTION_BY_RNA_POLYMERASE_II | 50 | -0.448276 | -1.41221 | 0.052846 | 0.220247 |
| GOBP_POSITIVE_REGULATION_OF_VASCULAR_ASSOCIATED_SMOOTH_MUSCLE_CELL_MIG | 15 | -0.581986 | -1.412195 | 0.098501 | 0.219906 |

| | | | | | |
|---|---|---|---|---|---|
| GOBP_REGULATION_OF_LYMPHOCYTE_DIFFERENTIATION | 161 | 0.267995 | 0.986781 | 0.493783 | 0.747993 |
| GOBP_ANTIGEN_PROCESSING_AND_PRESENTATION | 221 | 0.256279 | 0.986222 | 0.477509 | 0.748407 |
| GOBP_REGULATION_OF_LYMPHOCYTE_MEDIATED_IMMUNITY | 144 | 0.273227 | 0.985333 | 0.456364 | 0.749765 |
| GOBP_REGULATION_OF_SODIUM_ION_TRANSPORT | 64 | 0.298386 | 0.985198 | 0.497041 | 0.748971 |
| GOBP_HISTONE_H3_K9_MODIFICATION | 41 | 0.337563 | 0.982555 | 0.478846 | 0.75557 |
| GOBP_PLATELET_AGGREGATION | 50 | 0.315044 | 0.98124 | 0.46833 | 0.758258 |
| GOBP_THYROID_HORMONE_METABOLIC_PROCESS | 18 | 0.388972 | 0.979602 | 0.478927 | 0.761955 |
| GOBP_PROTEIN_KINASE_C_SIGNALING | 24 | 0.371887 | 0.979488 | 0.488414 | 0.761086 |
| GOBP_MATERNAL_PROCESS_INVOLVED_IN_FEMALE_PREGNANCY | 43 | 0.323068 | 0.97922 | 0.478505 | 0.760669 |
| GOBP_ENDOCRINE_PANCREAS_DEVELOPMENT | 28 | 0.354677 | 0.979024 | 0.484252 | 0.760059 |
| GOBP_COLLAGEN_CATABOLIC_PROCESS | 34 | 0.341034 | 0.978789 | 0.510516 | 0.759538 |
| GOBP_REGULATION_OF_SYNAPTIC_TRANSMISSION_GABAERGIC | 19 | 0.393271 | 0.978376 | 0.46507 | 0.759563 |
| GOBP_POSITIVE_REGULATION_OF_LEUKOCYTE_MEDIATED_IMMUNITY | 116 | 0.2798 | 0.977132 | 0.497248 | 0.761953 |
| GOBP_REGULATION_OF_FIBROBLAST_GROWTH_FACTOR_RECEPTOR_SIGNALING_PATHW | 22 | 0.371162 | 0.977085 | 0.487365 | 0.760892 |
| GOBP_POSITIVE_REGULATION_OF_PEPTIDE_SECRETION | 117 | 0.275666 | 0.975153 | 0.517668 | 0.765294 |
| GOBP_NEUROMUSCULAR_PROCESS_CONTROLLING_BALANCE | 29 | 0.356062 | 0.974592 | 0.492366 | 0.765731 |
| GOBP_FEMALE_MEIOTIC_NUCLEAR_DIVISION | 24 | 0.367548 | 0.974463 | 0.484849 | 0.764926 |
| GOBP_REGULATION_OF_T_CELL_DIFFERENTIATION | 134 | 0.267507 | 0.974238 | 0.513011 | 0.764338 |
| GOBP_CELL_JUNCTION_MAINTENANCE | 26 | 0.360579 | 0.973579 | 0.49053 | 0.765022 |
| GOBP_POSITIVE_REGULATION_OF_CELL_PROJECTION_ORGANIZATION | 287 | 0.246628 | 0.973191 | 0.535714 | 0.765028 |
| GOBP_REGULATION_OF_NMDA_RECEPTOR_ACTIVITY | 25 | 0.36006 | 0.972822 | 0.483209 | 0.764975 |
| GOBP_REGULATION_OF_T_CELL_APOPTOTIC_PROCESS | 28 | 0.355912 | 0.970664 | 0.483685 | 0.770062 |
| GOBP_PROTEIN_DNA_COMPLEX_DISASSEMBLY | 18 | 0.37843 | 0.969833 | 0.488706 | 0.771301 |
| GOBP_STRIATED_MUSCLE_ADAPTATION | 32 | 0.33621 | 0.969763 | 0.501035 | 0.770322 |
| GOBP_ACTIN_FILAMENT_SEVERING | 15 | 0.405765 | 0.968078 | 0.51434 | 0.774027 |
| GOBP_EXPORT_ACROSS_PLASMA_MEMBRANE | 49 | 0.314222 | 0.967554 | 0.520599 | 0.774341 |
| GOBP_NEGATIVE_REGULATION_OF_NEURON_DEATH | 156 | 0.261184 | 0.966532 | 0.539286 | 0.776121 |
| GOBP_REGULATION_OF_RHO_PROTEIN_SIGNAL_TRANSDUCTION | 75 | 0.285161 | 0.966055 | 0.529301 | 0.776318 |
| GOBP_REGULATION_OF_AMPA_RECEPTOR_ACTIVITY | 18 | 0.388223 | 0.96601 | 0.528942 | 0.775259 |
| GOBP_DRUG_METABOLIC_PROCESS | 27 | 0.348861 | 0.963985 | 0.513514 | 0.78 |
| GOBP_NEGATIVE_REGULATION_OF_SIGNAL_TRANSDUCTION_BY_P53_CLASS_MEDIATOF | 29 | 0.348348 | 0.963952 | 0.489022 | 0.778905 |
| GOBP_EXPLORATION_BEHAVIOR | 15 | 0.40681 | 0.963459 | 0.527205 | 0.779094 |
| GOBP_ACTOMYOSIN_STRUCTURE_ORGANIZATION | 152 | 0.264497 | 0.962942 | 0.552028 | 0.779396 |
| GOBP_NEGATIVE_REGULATION_OF_MONONUCLEAR_CELL_MIGRATION | 21 | 0.367153 | 0.962415 | 0.488142 | 0.779767 |
| GOBP_PHOSPHATIDYLINOSITOL_DEPHOSPHORYLATION | 30 | 0.347105 | 0.959865 | 0.532491 | 0.785958 |
| GOBP_HAIR_CELL_DIFFERENTIATION | 33 | 0.340417 | 0.95957 | 0.535922 | 0.785596 |
| GOBP_REGULATION_OF_CELLULAR_RESPONSE_TO_VASCULAR_ENDOTHELIAL_GROWTI | 17 | 0.385546 | 0.959459 | 0.490079 | 0.784722 |
| GOBP_APOPTOTIC_CELL_CLEARANCE | 46 | 0.30943 | 0.956541 | 0.519455 | 0.791837 |
| GOBP_TRANSLATIONAL_INITIATION | 186 | 0.255019 | 0.954889 | 0.563055 | 0.795392 |
| GOBP_MITOTIC_SISTER_CHROMATID_COHESION | 26 | 0.348969 | 0.954603 | 0.526606 | 0.795018 |
| GOBP_RENAL_SYSTEM_VASCULATURE_DEVELOPMENT | 19 | 0.376194 | 0.953798 | 0.518664 | 0.796132 |
| GOBP_NEGATIVE_REGULATION_OF_LEUKOCYTE_CHEMOTAXIS | 17 | 0.388402 | 0.952494 | 0.495088 | 0.798791 |
| GOBP_NEGATIVE_REGULATION_OF_LEUKOCYTE_APOPTOTIC_PROCESS | 40 | 0.321456 | 0.952377 | 0.518095 | 0.797945 |
| GOBP_POSITIVE_REGULATION_OF_NEURON_PROJECTION_DEVELOPMENT | 127 | 0.264358 | 0.952031 | 0.572183 | 0.797756 |
| GOBP_REGULATION_OF_LYMPHOCYTE_APOPTOTIC_PROCESS | 48 | 0.303636 | 0.951876 | 0.542039 | 0.79702 |
| GOBP_PURINE_NUCLEOSIDE_BIOSYNTHETIC_PROCESS | 20 | 0.377818 | 0.951704 | 0.537374 | 0.796361 |
| GOBP_SYNCYTIUM_FORMATION | 46 | 0.309989 | 0.951597 | 0.532567 | 0.795481 |
| GOBP_POSITIVE_REGULATION_OF_GLYCOGEN_METABOLIC_PROCESS | 15 | 0.398976 | 0.951413 | 0.527778 | 0.794798 |
| GOBP_POSITIVE_REGULATION_OF_RESPONSE_TO_ENDOPLASMIC_RETICULUM_STRESS | 35 | 0.335902 | 0.950016 | 0.515094 | 0.797596 |
| GOBP_METENCEPHALON_DEVELOPMENT | 74 | 0.285857 | 0.948454 | 0.577982 | 0.800904 |
| GOBP_FORMATION_OF_CYTOPLASMIC_TRANSLATION_INITIATION_COMPLEX | 16 | 0.397473 | 0.946816 | 0.521569 | 0.804384 |
| GOBP_NEGATIVE_REGULATION_OF_LYMPHOCYTE_MIGRATION | 15 | 0.396729 | 0.946641 | 0.513292 | 0.803746 |
| GOBP_LYMPH_VESSEL_MORPHOGENESIS | 17 | 0.379186 | 0.946557 | 0.535922 | 0.802807 |
| GOBP_REGULATION_OF_HAIR_FOLLICLE_DEVELOPMENT | 15 | 0.400216 | 0.946205 | 0.535294 | 0.80261 |
| GOBP_REGULATION_OF_MYELINATION | 34 | 0.337949 | 0.946188 | 0.569201 | 0.801478 |
| GOBP_REGULATION_OF_PROTEIN_IMPORT | 54 | 0.300726 | 0.945498 | 0.543021 | 0.802298 |
| GOBP_CALCINEURIN_MEDIATED_SIGNALING | 38 | 0.321974 | 0.945362 | 0.56917 | 0.801531 |
| GOBP_EMBRYONIC_HEART_TUBE_DEVELOPMENT | 56 | 0.296822 | 0.942766 | 0.551657 | 0.807563 |
| GOBP_REGULATION_OF_MEIOTIC_NUCLEAR_DIVISION | 24 | 0.352213 | 0.941128 | 0.537453 | 0.8111 |
| GOBP_PITUITARY_GLAND_DEVELOPMENT | 18 | 0.384057 | 0.940189 | 0.539305 | 0.812662 |
| GOBP_CALCIUM_ION_IMPORT | 61 | 0.297864 | 0.939224 | 0.586022 | 0.814242 |
| GOBP_MAINTENANCE_OF_PROTEIN_LOCATION_IN_CELL | 59 | 0.292609 | 0.938706 | 0.548507 | 0.814533 |
| GOBP_POSITIVE_REGULATION_OF_MYELOID_LEUKOCYTE_CYTOKINE_PRODUCTION_INV | 16 | 0.384968 | 0.938601 | 0.528265 | 0.813642 |
| GOBP_REGULATION_OF_SENSORY_PERCEPTION | 23 | 0.356359 | 0.938059 | 0.564639 | 0.814018 |
| GOBP_DEVELOPMENTAL_PIGMENTATION | 33 | 0.328915 | 0.937623 | 0.551136 | 0.814061 |
| GOBP_ENDOTHELIAL_CELL_APOPTOTIC_PROCESS | 48 | 0.30281 | 0.937057 | 0.537037 | 0.814524 |
| GOBP_CELLULAR_ALDEHYDE_METABOLIC_PROCESS | 50 | 0.29881 | 0.936122 | 0.585127 | 0.81606 |
| GOBP_NEGATIVE_REGULATION_OF_CELL_SUBSTRATE_JUNCTION_ORGANIZATION | 17 | 0.378055 | 0.935655 | 0.560706 | 0.81623 |
| GOBP_PURINE_NUCLEOTIDE_TRANSPORT | 23 | 0.355127 | 0.935107 | 0.564961 | 0.816601 |
| GOBP_RETINA_DEVELOPMENT_IN_CAMERA_TYPE_EYE | 103 | 0.267646 | 0.93493 | 0.590074 | 0.815925 |
| GOBP_REGULATION_OF_MULTI_ORGANISM_PROCESS | 38 | 0.316741 | 0.933766 | 0.55642 | 0.817988 |
| GOBP_POSITIVE_REGULATION_OF_CELLULAR_COMPONENT_BIOGENESIS | 447 | 0.227524 | 0.931861 | 0.73366 | 0.822116 |
| GOBP_REGULATION_OF_DENDRITE_DEVELOPMENT | 82 | 0.272619 | 0.929376 | 0.632353 | 0.827912 |
| GOBP_EXCITATORY_SYNAPSE_ASSEMBLY | 19 | 0.363425 | 0.929331 | 0.559055 | 0.826875 |
| GOBP_REGULATION_OF_ERBB_SIGNALING_PATHWAY | 85 | 0.271656 | 0.929268 | 0.614841 | 0.82587 |
| GOBP_POLYOL_METABOLIC_PROCESS | 113 | 0.263908 | 0.928221 | 0.613678 | 0.827632 |
| GOBP_COPULATION | 16 | 0.385921 | 0.928206 | 0.544747 | 0.826499 |
| GOBP_INSULIN_SECRETION | 155 | 0.254233 | 0.926551 | 0.623711 | 0.829961 |
| GOBP_NEGATIVE_REGULATION_OF_TUMOR_NECROSIS_FACTOR_SUPERFAMILY_CYTOK | 49 | 0.300267 | 0.921855 | 0.594041 | 0.841982 |
| GOBP_O_GLYCAN_PROCESSING | 45 | 0.305579 | 0.921711 | 0.579737 | 0.841187 |
| GOBP_HIPPOCAMPUS_DEVELOPMENT | 53 | 0.292323 | 0.921406 | 0.587703 | 0.840862 |
| GOBP_VENTRAL_SPINAL_CORD_DEVELOPMENT | 23 | 0.349919 | 0.920935 | 0.587669 | 0.841003 |
| GOBP_MITOCHONDRIAL_CALCIUM_ION_HOMEOSTASIS | 24 | 0.34234 | 0.920786 | 0.573585 | 0.84023 |
| GOBP_NEGATIVE_REGULATION_OF_CYTOKINE_PRODUCTION_INVOLVED_IN_INFLAMMAT | 24 | 0.347858 | 0.918549 | 0.573308 | 0.845171 |
| GOBP_SEMAPHORIN_PLEXIN_SIGNALING_PATHWAY | 35 | 0.322511 | 0.918497 | 0.577982 | 0.844133 |
| GOBP_RECEPTOR_RECYCLING | 30 | 0.332914 | 0.918381 | 0.593573 | 0.843259 |
| GOBP_POSITIVE_REGULATION_OF_BLOOD_VESSEL_ENDOTHELIAL_CELL_MIGRATION | 51 | 0.294327 | 0.916494 | 0.606403 | 0.847274 |
| GOBP_ACTIVATION_OF_GTPASE_ACTIVITY | 98 | 0.26388 | 0.91529 | 0.626118 | 0.849342 |
| GOBP_ACTIVATION_OF_MAPKK_ACTIVITY | 45 | 0.301717 | 0.915184 | 0.590994 | 0.848448 |
| GOBP_POSITIVE_REGULATION_OF_REGULATED_SECRETORY_PATHWAY | 43 | 0.305382 | 0.914738 | 0.6 | 0.848509 |
| GOBP_REGULATION_OF_MACROPHAGE_CHEMOTAXIS | 26 | 0.34543 | 0.914219 | 0.581006 | 0.848752 |
| GOBP_GLUTATHIONE_METABOLIC_PROCESS | 49 | 0.297216 | 0.914157 | 0.606285 | 0.847749 |
| GOBP_LAMELLIPODIUM_ASSEMBLY | 63 | 0.282199 | 0.913636 | 0.616216 | 0.847937 |
| GOBP_PLASMA_MEMBRANE_TUBULATION | 19 | 0.362867 | 0.913301 | 0.586275 | 0.847692 |
| GOBP_SYNAPTIC_VESICLE_RECYCLING | 63 | 0.284948 | 0.912581 | 0.59252 | 0.848494 |
| GOBP_REGULATION_OF_CHOLESTEROL_EFFLUX | 27 | 0.334905 | 0.912244 | 0.604418 | 0.848199 |
| GOBP_WALKING_BEHAVIOR | 18 | 0.368703 | 0.911131 | 0.597786 | 0.850076 |
| GOBP_NEGATIVE_REGULATION_OF_TRANSCRIPTION_REGULATORY_REGION_DNA_BIND | 16 | 0.384185 | 0.908768 | 0.604555 | 0.855302 |
| GOBP_LONG_CHAIN_FATTY_ACID_BIOSYNTHETIC_PROCESS | 28 | 0.329253 | 0.908099 | 0.58891 | 0.855954 |
| GOBP_AMMONIUM_ION_METABOLIC_PROCESS | 18 | 0.363842 | 0.90791 | 0.597744 | 0.855288 |
| GOBP_INDOLE_CONTAINING_COMPOUND_METABOLIC_PROCESS | 16 | 0.371737 | 0.9076 | 0.611111 | 0.854946 |
| GOBP_REGULATION_OF_ACTOMYOSIN_STRUCTURE_ORGANIZATION | 80 | 0.268269 | 0.906616 | 0.672828 | 0.856433 |
| GOBP_MYOBLAST_DIFFERENTIATION | 61 | 0.280094 | 0.904132 | 0.629771 | 0.86182 |
| GOBP_POSITIVE_REGULATION_OF_PROTEIN_BINDING | 78 | 0.271329 | 0.903942 | 0.651246 | 0.861143 |
| GOBP_FOCAL_ADHESION_ASSEMBLY | 75 | 0.275632 | 0.902799 | 0.664729 | 0.863019 |
| GOBP_POSITIVE_REGULATION_OF_NEURON_MIGRATION | 15 | 0.379935 | 0.902402 | 0.597077 | 0.862917 |
| GOBP_POSITIVE_REGULATION_OF_CELL_AGING | 15 | 0.375856 | 0.901477 | 0.580882 | 0.864198 |
| GOBP_GENETIC_IMPRINTING | 20 | 0.348334 | 0.900537 | 0.604463 | 0.865563 |
| GOBP_REGULATION_OF_CALCIUM_ION_TRANSPORT_INTO_CYTOSOL | 73 | 0.272041 | 0.900463 | 0.650198 | 0.864571 |
| GOBP_NEGATIVE_REGULATION_OF_CALCIUM_ION_TRANSMEMBRANE_TRANSPORT | 28 | 0.324105 | 0.900442 | 0.637427 | 0.86345 |
| GOBP_NEUROTROPHIN_TRK_RECEPTOR_SIGNALING_PATHWAY | 24 | 0.339263 | 0.898519 | 0.59596 | 0.867333 |
| GOBP_NEUROTROPHIN_SIGNALING_PATHWAY | 30 | 0.324044 | 0.897167 | 0.624754 | 0.869703 |
| GOBP_ESTABLISHMENT_OR_MAINTENANCE_OF_BIPOLAR_CELL_POLARITY | 42 | 0.296817 | 0.896343 | 0.622857 | 0.870673 |
| GOBP_REGULATION_OF_SODIUM_ION_TRANSMEMBRANE_TRANSPORT | 47 | 0.291685 | 0.89492 | 0.626953 | 0.873208 |
| GOBP_EPITHELIAL_CELL_MORPHOGENESIS | 22 | 0.349201 | 0.894523 | 0.593688 | 0.87306 |
| GOBP_PHOSPHOLIPID_DEPHOSPHORYLATION | 41 | 0.304132 | 0.894439 | 0.641326 | 0.872096 |
| GOBP_NUCLEOTIDE_TRANSPORT | 30 | 0.321313 | 0.892233 | 0.614931 | 0.876647 |
| GOBP_REGULATION_OF_NEURON_MIGRATION | 27 | 0.327227 | 0.890945 | 0.634146 | 0.878836 |
| GOBP_NEGATIVE_REGULATION_OF_CHEMOKINE_PRODUCTION | 23 | 0.335001 | 0.889942 | 0.617409 | 0.880249 |
| GOBP_GLYCOPROTEIN_BIOSYNTHETIC_PROCESS | 291 | 0.225078 | 0.889241 | 0.824057 | 0.880896 |
| GOBP_POSITIVE_REGULATION_OF_SODIUM_ION_TRANSMEMBRANE_TRANSPORT | 19 | 0.354641 | 0.888523 | 0.624266 | 0.881535 |
| GOBP_REGULATION_OF_PHOSPHOLIPASE_C_ACTIVITY | 36 | 0.315229 | 0.887491 | 0.648598 | 0.882971 |
| GOBP_PROTEIN_LOCALIZATION_TO_PLASMA_MEMBRANE | 243 | 0.229933 | 0.887383 | 0.799645 | 0.882066 |
| GOBP_THIOESTER_METABOLIC_PROCESS | 93 | 0.258314 | 0.886694 | 0.683186 | 0.882628 |
| GOBP_CALCIUM_ION_IMPORT_INTO_CYTOSOL | 16 | 0.366412 | 0.886211 | 0.651685 | 0.882686 |
| GOBP_MITOCHONDRIAL_FISSION | 41 | 0.290363 | 0.885974 | 0.67027 | 0.882123 |
| GOBP_GLYCOPROTEIN_METABOLIC_PROCESS | 347 | 0.218946 | 0.880829 | 0.860465 | 0.89401 |
| GOBP_NEGATIVE_REGULATION_OF_MUSCLE_HYPERTROPHY | 26 | 0.324519 | 0.879293 | 0.661448 | 0.896596 |
| GOBP_POSITIVE_REGULATION_OF_EXTRINSIC_APOPTOTIC_SIGNALING_PATHWAY | 41 | 0.296 | 0.87752 | 0.665984 | 0.899871 |
| GOBP_NEGATIVE_REGULATION_OF_REACTIVE_OXYGEN_SPECIES_BIOSYNTHETIC_PROC | 25 | 0.324506 | 0.875285 | 0.630394 | 0.904195 |
| GOBP_REGULATION_OF_RECEPTOR_RECYCLING | 21 | 0.339735 | 0.874602 | 0.642293 | 0.904639 |
| GOBP_NEGATIVE_REGULATION_OF_AMYLOID_PRECURSOR_PROTEIN_CATABOLIC_PROC | 22 | 0.334948 | 0.873107 | 0.626747 | 0.907042 |
| GOBP_SPINAL_CORD_MOTOR_NEURON_DIFFERENTIATION | 18 | 0.358373 | 0.871815 | 0.618026 | 0.908968 |
| GOBP_NEGATIVE_REGULATION_OF_NIK_NF_KAPPAB_SIGNALING | 24 | 0.324295 | 0.871751 | 0.655738 | 0.907916 |
| GOBP_POSITIVE_REGULATION_OF_LYMPHOCYTE_DIFFERENTIATION | 95 | 0.252747 | 0.871714 | 0.737736 | 0.90682 |
| GOBP_NEGATIVE_REGULATION_OF_RESPONSE_TO_DNA_DAMAGE_STIMULUS | 82 | 0.256802 | 0.871529 | 0.727103 | 0.906104 |
| GOBP_REGULATION_OF_ACTIN_FILAMENT_ORGANIZATION | 235 | 0.226711 | 0.870847 | 0.826825 | 0.906575 |
| GOBP_RESPONSE_TO_NERVE_GROWTH_FACTOR | 40 | 0.29202 | 0.869898 | 0.688931 | 0.907634 |
| GOBP_NEGATIVE_REGULATION_OF_PEPTIDYL_SERINE_PHOSPHORYLATION | 25 | 0.323842 | 0.869476 | 0.664207 | 0.907466 |
| GOBP_REGULATION_OF_MITOCHONDRIAL_GENE_EXPRESSION | 29 | 0.317212 | 0.868895 | 0.653772 | 0.9077 |
| GOBP_POSITIVE_REGULATION_OF_TOR_SIGNALING | 38 | 0.290154 | 0.86657 | 0.704501 | 0.91212 |
| GOBP_COLUMNAR_CUBOIDAL_EPITHELIAL_CELL_DEVELOPMENT | 28 | 0.315039 | 0.865629 | 0.678917 | 0.913216 |
| GOBP_MATING | 28 | 0.315918 | 0.865512 | 0.669866 | 0.912288 |
| GOBP_RETINOIC_ACID_METABOLIC_PROCESS | 15 | 0.364508 | 0.864914 | 0.65019 | 0.912557 |
| GOBP_DETECTION_OF_LIGHT_STIMULUS | 48 | 0.281261 | 0.864732 | 0.686425 | 0.911787 |
| GOBP_CELLULAR_RESPONSE_TO_STEROL | 19 | 0.339757 | 0.863823 | 0.678309 | 0.912767 |
| GOBP_SECONDARY_PALATE_DEVELOPMENT | 20 | 0.338214 | 0.863656 | 0.670061 | 0.911967 |
| GOBP_EPITHELIAL_CELL_CELL_ADHESION | 15 | 0.362922 | 0.859387 | 0.664729 | 0.920692 |
| GOBP_CYCLIC_NUCLEOTIDE_METABOLIC_PROCESS | 28 | 0.310833 | 0.855461 | 0.683908 | 0.92852 |
| GOBP_CYTOPLASMIC_TRANSLATION | 99 | 0.247471 | 0.85544 | 0.767225 | 0.927364 |
| GOBP_VITAMIN_D_METABOLIC_PROCESS | 17 | 0.35711 | 0.850905 | 0.705305 | 0.936302 |
| GOBP_ACTIN_POLYMERIZATION_OR_DEPOLYMERIZATION | 191 | 0.224798 | 0.850807 | 0.865724 | 0.935307 |
| GOBP_PHOTOTRANSDUCTION | 35 | 0.291111 | 0.84886 | 0.705104 | 0.938553 |
| GOBP_BONE_REMODELING | 68 | 0.26129 | 0.847172 | 0.785579 | 0.94108 |
| GOBP_POSITIVE_REGULATION_OF_SPROUTING_ANGIOGENESIS | 19 | 0.331822 | 0.846285 | 0.679775 | 0.941788 |
| GOBP_REGULATION_OF_MUSCLE_ORGAN_DEVELOPMENT | 29 | 0.300724 | 0.846123 | 0.68906 | 0.940939 |
| GOBP_ORGANIC_HYDROXY_COMPOUND_BIOSYNTHETIC_PROCESS | 198 | 0.223108 | 0.845615 | 0.883424 | 0.940841 |

| | | | | | |
|---|---|---|---|---|---|
| GOBP_POSITIVE_REGULATION_OF_CELL_GROWTH | 135 | -0.382212 | -1.411675 | 0.013605 | 0.220341 |
| GOBP_CARBOXYLIC_ACID_TRANSPORT | 202 | -0.365547 | -1.411005 | 0.004608 | 0.220929 |
| GOBP_RESPONSE_TO_AMINO_ACID_STARVATION | 48 | -0.4514 | -1.41089 | 0.055066 | 0.220758 |
| GOBP_NEGATIVE_REGULATION_OF_KINASE_ACTIVITY | 233 | -0.356453 | -1.409758 | 0.007177 | 0.222009 |
| GOBP_MITOTIC_SPINDLE_ASSEMBLY | 65 | -0.43324 | -1.409597 | 0.052154 | 0.221915 |
| GOBP_IRON_ION_HOMEOSTASIS | 73 | -0.421153 | -1.409232 | 0.048387 | 0.222079 |
| GOBP_STRIATED_MUSCLE_CELL_APOPTOTIC_PROCESS | 27 | -0.508659 | -1.4087 | 0.075594 | 0.222483 |
| GOBP_NEGATIVE_REGULATION_OF_HISTONE_ACETYLATION | 15 | -0.588326 | -1.40656 | 0.077079 | 0.225245 |
| GOBP_NEGATIVE_REGULATION_OF_TRANSMEMBRANE_TRANSPORT | 89 | -0.404882 | -1.406441 | 0.022075 | 0.225094 |
| GOBP_POSITIVE_REGULATION_OF_MITOCHONDRIAL_OUTER_MEMBRANE_PERMEABILIZATI | 35 | -0.472115 | -1.405805 | 0.071429 | 0.225735 |
| GOBP_REGULATION_OF_ALTERNATIVE_MRNA_SPLICING_VIA_SPLICEOSOME | 48 | -0.449518 | -1.405013 | 0.046122 | 0.226652 |
| GOBP_NEGATIVE_REGULATION_OF_BINDING | 135 | -0.379125 | -1.404665 | 0.021322 | 0.226861 |
| GOBP_POSITIVE_REGULATION_OF_OSTEOBLAST_DIFFERENTIATION | 48 | -0.447511 | -1.4042 | 0.059574 | 0.227166 |
| GOBP_REGULATION_OF_PROTEASOMAL_UBIQUITIN_DEPENDENT_PROTEIN_CATABOLIC_PI | 123 | -0.387536 | -1.404118 | 0.028953 | 0.226929 |
| GOBP_NEGATIVE_REGULATION_OF_CELL_CYCLE_ARREST | 19 | -0.559477 | -1.403322 | 0.082645 | 0.227714 |
| GOBP_ANTERIOR_POSTERIOR_PATTERN_SPECIFICATION | 140 | -0.375705 | -1.402145 | 0.01373 | 0.229109 |
| GOBP_NEGATIVE_REGULATION_OF_CATION_CHANNEL_ACTIVITY | 31 | -0.482711 | -1.402027 | 0.065173 | 0.228964 |
| GOBP_PROTEIN_TRANSPORT_ALONG_MICROTUBULE | 65 | -0.422674 | -1.401951 | 0.043478 | 0.22869 |
| GOBP_CILIUM_MOVEMENT | 114 | -0.386948 | -1.401854 | 0.02069 | 0.228482 |
| GOBP_RESPONSE_TO_DIETARY_EXCESS | 15 | -0.579404 | -1.401824 | 0.102669 | 0.228172 |
| GOBP_POSITIVE_REGULATION_OF_LEUKOCYTE_CELL_CELL_ADHESION | 215 | -0.354315 | -1.401652 | 0.006961 | 0.228085 |
| GOBP_NEGATIVE_REGULATION_OF_PEPTIDYL_LYSINE_ACETYLATION | 17 | -0.56125 | -1.401476 | 0.101266 | 0.227977 |
| GOBP_MRNA_3_END_PROCESSING | 97 | -0.393342 | -1.400119 | 0.028571 | 0.229741 |
| GOBP_SNARE_COMPLEX_ASSEMBLY | 18 | -0.557687 | -1.399899 | 0.092715 | 0.229718 |
| GOBP_PROTEIN_CONTAINING_COMPLEX_LOCALIZATION | 261 | -0.345778 | -1.399384 | 0.004866 | 0.230144 |
| GOBP_GLUCOSE_6_PHOSPHATE_METABOLIC_PROCESS | 22 | -0.530061 | -1.398324 | 0.075594 | 0.231393 |
| GOBP_ATRIOVENTRICULAR_VALVE_DEVELOPMENT | 20 | -0.533038 | -1.398312 | 0.093878 | 0.231054 |
| GOBP_RESPONSE_TO_STEROID_HORMONE | 262 | -0.349983 | -1.398235 | 0.004684 | 0.230783 |
| GOBP_INTERLEUKIN_1_BETA_PRODUCTION | 82 | -0.410329 | -1.397853 | 0.029289 | 0.23108 |
| GOBP_REGULATION_OF_CHROMOSOME_ORGANIZATION | 258 | -0.350134 | -1.397534 | 0.002278 | 0.231151 |
| GOBP_EPITHELIAL_CELL_DIFFERENTIATION | 463 | -0.329987 | -1.396777 | 0 | 0.231932 |
| GOBP_LEUKOCYTE_PROLIFERATION | 265 | -0.351553 | -1.396638 | 0.004706 | 0.231793 |
| GOBP_INTERLEUKIN_10_PRODUCTION | 56 | -0.435305 | -1.396486 | 0.047619 | 0.231672 |
| GOBP_REGULATION_OF_CYSTEINE_TYPE_ENDOPEPTIDASE_ACTIVITY | 199 | -0.359764 | -1.396259 | 0.002387 | 0.231647 |
| GOBP_INTRACILIARY_TRANSPORT_INVOLVED_IN_CILIUM_ASSEMBLY | 39 | -0.462792 | -1.39586 | 0.05794 | 0.231975 |
| GOBP_PYRIMIDINE_CONTAINING_COMPOUND_CATABOLIC_PROCESS | 36 | -0.47119 | -1.395272 | 0.076271 | 0.232572 |
| GOBP_REGULATION_OF_SYNCYTIUM_FORMATION_BY_PLASMA_MEMBRANE_FUSION | 23 | -0.524981 | -1.394968 | 0.087475 | 0.232697 |
| GOBP_CARBOHYDRATE_TRANSMEMBRANE_TRANSPORT | 92 | -0.403414 | -1.394748 | 0.021142 | 0.232642 |
| GOBP_REGULATION_OF_CYCLIN_DEPENDENT_PROTEIN_KINASE_ACTIVITY | 92 | -0.398227 | -1.394048 | 0.028634 | 0.233364 |
| GOBP_POSITIVE_REGULATION_OF_CELL_CELL_ADHESION | 244 | -0.352222 | -1.393907 | 0.00978 | 0.233249 |
| GOBP_NEGATIVE_REGULATION_OF_CATION_TRANSMEMBRANE_TRANSPORT | 63 | -0.423089 | -1.393838 | 0.04 | 0.232989 |
| GOBP_B_CELL_RECEPTOR_SIGNALING_PATHWAY | 109 | -0.390355 | -1.393621 | 0.025862 | 0.232968 |
| GOBP_REGULATION_OF_NEUTROPHIL_MIGRATION | 39 | -0.464731 | -1.393597 | 0.07947 | 0.232685 |
| GOBP_REGULATION_OF_HEART_RATE | 62 | -0.4287 | -1.393566 | 0.042222 | 0.2324 |
| GOBP_REGULATION_OF_COLLAGEN_METABOLIC_PROCESS | 33 | -0.470745 | -1.393516 | 0.044543 | 0.232127 |
| GOBP_KIDNEY_EPITHELIUM_DEVELOPMENT | 91 | -0.398823 | -1.393062 | 0.022472 | 0.232524 |
| GOBP_NEGATIVE_REGULATION_OF_RESPONSE_TO_ENDOPLASMIC_RETICULUM_STRESS | 42 | -0.463895 | -1.392519 | 0.052863 | 0.232974 |
| GOBP_LUNG_ALVEOLUS_DEVELOPMENT | 27 | -0.500036 | -1.392489 | 0.07234 | 0.232656 |
| GOBP_CELLULAR_RESPONSE_TO_DEXAMETHASONE_STIMULUS | 25 | -0.496451 | -1.39226 | 0.089397 | 0.232666 |
| GOBP_RESPONSE_TO_INSULIN | 238 | -0.348796 | -1.390988 | 0.004474 | 0.23427 |
| GOBP_POSITIVE_REGULATION_OF_SYNAPTIC_TRANSMISSION_GLUTAMATERGIC | 20 | -0.533598 | -1.390979 | 0.067982 | 0.233933 |
| GOBP_POSITIVE_REGULATION_OF_INTERLEUKIN_1_PRODUCTION | 54 | -0.437311 | -1.390609 | 0.05071 | 0.234203 |
| GOBP_ACTIN_CYTOSKELETON_REORGANIZATION | 90 | -0.400611 | -1.390567 | 0.032407 | 0.233907 |
| GOBP_REGULATION_OF_CELL_CYCLE_CHECKPOINT | 30 | -0.496871 | -1.390527 | 0.077083 | 0.233644 |
| GOBP_PROTEIN_LOCALIZATION_TO_CILIUM | 54 | -0.434819 | -1.39052 | 0.057377 | 0.233304 |
| GOBP_STRESS_ACTIVATED_PROTEIN_KINASE_SIGNALING_CASCADE | 241 | -0.347947 | -1.390492 | 0.004866 | 0.232994 |
| GOBP_NEGATIVE_REGULATION_OF_UBIQUITIN_PROTEIN_TRANSFERASE_ACTIVITY | 16 | -0.579114 | -1.390476 | 0.101833 | 0.232682 |
| GOBP_REGULATION_OF_LIPID_KINASE_ACTIVITY | 62 | -0.433204 | -1.39031 | 0.03956 | 0.232603 |
| GOBP_NEUROINFLAMMATORY_RESPONSE | 31 | -0.482488 | -1.389627 | 0.063694 | 0.233349 |
| GOBP_LIPID_LOCALIZATION | 384 | -0.335245 | -1.389304 | 0.002237 | 0.233449 |
| GOBP_APOPTOTIC_PROCESS_INVOLVED_IN_DEVELOPMENT | 25 | -0.501997 | -1.389289 | 0.072978 | 0.233127 |
| GOBP_POSITIVE_REGULATION_OF_MRNA_CATABOLIC_PROCESS | 50 | -0.442465 | -1.389107 | 0.061966 | 0.233052 |
| GOBP_REGULATION_OF_GENE_EXPRESSION_EPIGENETIC | 177 | -0.362684 | -1.389036 | 0.017582 | 0.232817 |
| GOBP_POSITIVE_REGULATION_OF_LIPID_BIOSYNTHETIC_PROCESS | 65 | -0.420949 | -1.388967 | 0.063736 | 0.232589 |
| GOBP_NEPHRON_EPITHELIUM_DEVELOPMENT | 73 | -0.413413 | -1.38776 | 0.028513 | 0.234106 |
| GOBP_REGULATION_OF_PHOSPHOPROTEIN_PHOSPHATASE_ACTIVITY | 110 | -0.391998 | -1.387756 | 0.033058 | 0.233767 |
| GOBP_LYMPHOCYTE_MEDIATED_IMMUNITY | 308 | -0.341905 | -1.387201 | 0.004598 | 0.234293 |
| GOBP_REGULATION_OF_LEUKOCYTE_PROLIFERATION | 212 | -0.35569 | -1.387038 | 0.007229 | 0.234211 |
| GOBP_NEGATIVE_REGULATION_OF_POTASSIUM_ION_TRANSMEMBRANE_TRANSPORT | 18 | -0.55351 | -1.386923 | 0.112871 | 0.234066 |
| GOBP_POSITIVE_REGULATION_OF_B_CELL_PROLIFERATION | 37 | -0.464967 | -1.386662 | 0.063063 | 0.23414 |
| GOBP_NEGATIVE_REGULATION_OF_CELL_CYCLE_G1_S_PHASE_TRANSITION | 97 | -0.387948 | -1.386353 | 0.016807 | 0.2343 |
| GOBP_POSITIVE_REGULATION_OF_NIK_NF_KAPPAB_SIGNALING | 59 | -0.432873 | -1.385702 | 0.044776 | 0.234944 |
| GOBP_REGULATION_OF_MACROPHAGE_DERIVED_FOAM_CELL_DIFFERENTIATION | 19 | -0.537259 | -1.385632 | 0.087576 | 0.234693 |
| GOBP_CELL_RECOGNITION | 164 | -0.367576 | -1.385293 | 0.016055 | 0.234888 |
| GOBP_NEGATIVE_REGULATION_OF_NUCLEAR_DIVISION | 50 | -0.452233 | -1.385249 | 0.053419 | 0.234592 |
| GOBP_ODONTOGENESIS | 93 | -0.392033 | -1.384969 | 0.028571 | 0.234713 |
| GOBP_AXONEMAL_DYNEIN_COMPLEX_ASSEMBLY | 28 | -0.489375 | -1.384791 | 0.077381 | 0.234656 |
| GOBP_RESPONSE_TO_RETINOIC_ACID | 77 | -0.405914 | -1.384752 | 0.041758 | 0.234367 |
| GOBP_RESPONSE_TO_ORGANOPHOSPHORUS | 112 | -0.379623 | -1.383791 | 0.029412 | 0.235496 |
| GOBP_DEFINITIVE_HEMOPOIESIS | 19 | -0.545362 | -1.383546 | 0.091295 | 0.235548 |
| GOBP_NEGATIVE_REGULATION_OF_CELL_MATRIX_ADHESION | 32 | -0.470907 | -1.383063 | 0.075388 | 0.235954 |
| GOBP_CYCLIC_NUCLEOTIDE_MEDIATED_SIGNALING | 58 | -0.427855 | -1.382878 | 0.045259 | 0.235893 |
| GOBP_I_KAPPAB_PHOSPHORYLATION | 18 | -0.556352 | -1.381407 | 0.113208 | 0.237918 |
| GOBP_POSITIVE_REGULATION_OF_APOPTOTIC_SIGNALING_PATHWAY | 111 | -0.390621 | -1.381145 | 0.033333 | 0.238023 |
| GOBP_MICROTUBULE_BASED_MOVEMENT | 310 | -0.339055 | -1.380805 | 0.004651 | 0.238213 |
| GOBP_KINETOCHORE_ORGANIZATION | 22 | -0.523247 | -1.380706 | 0.10124 | 0.238014 |
| GOBP_NEGATIVE_REGULATION_OF_INTRACELLULAR_STEROID_HORMONE_RECEPTOR_SIC | 30 | -0.483665 | -1.380623 | 0.087475 | 0.23778 |
| GOBP_ACTIVATED_T_CELL_PROLIFERATION | 41 | -0.446876 | -1.380535 | 0.065817 | 0.237589 |
| GOBP_NEGATIVE_REGULATION_OF_EXOCYTOSIS | 26 | -0.502187 | -1.380395 | 0.074946 | 0.237471 |
| GOBP_REGULATION_OF_VASCULAR_ENDOTHELIAL_GROWTH_FACTOR_RECEPTOR_SIGNAI | 25 | -0.516139 | -1.380316 | 0.079108 | 0.237236 |
| GOBP_ORGANIC_HYDROXY_COMPOUND_TRANSPORT | 180 | -0.356865 | -1.380151 | 0.009501 | 0.237114 |
| GOBP_NEUROTRANSMITTER_METABOLIC_PROCESS | 18 | -0.54328 | -1.379705 | 0.110169 | 0.237401 |
| GOBP_REGULATION_OF_PROTEIN_MATURATION | 51 | -0.432 | -1.379105 | 0.049462 | 0.237946 |
| GOBP_REGULATION_OF_NEUTROPHIL_CHEMOTAXIS | 29 | -0.496766 | -1.378545 | 0.083512 | 0.238448 |
| GOBP_PROTEIN_INSERTION_INTO_MITOCHONDRIAL_MEMBRANE_INVOLVED_IN_APOPTOTI | 30 | -0.491692 | -1.378403 | 0.0818 | 0.238345 |
| GOBP_POSITIVE_REGULATION_OF_DEFENSE_RESPONSE | 302 | -0.339251 | -1.37799 | 0.009412 | 0.238625 |
| GOBP_NEGATIVE_REGULATION_OF_SMALL_GTPASE_MEDIATED_SIGNAL_TRANSDUCTION | 51 | -0.437882 | -1.377984 | 0.060976 | 0.238299 |
| GOBP_INTERLEUKIN_17_PRODUCTION | 30 | -0.495485 | -1.377596 | 0.10664 | 0.238581 |
| GOBP_T_HELPER_2_CELL_DIFFERENTIATION | 15 | -0.570125 | -1.37749 | 0.101695 | 0.238442 |
| GOBP_REGULATION_OF_PROTEIN_TARGETING | 74 | -0.404287 | -1.377446 | 0.02766 | 0.238165 |
| GOBP_PROTEIN_FOLDING | 211 | -0.356987 | -1.377373 | 0.009029 | 0.237953 |
| GOBP_INTERLEUKIN_8_PRODUCTION | 77 | -0.400902 | -1.376879 | 0.040449 | 0.23832 |
| GOBP_MITOCHONDRIAL_MEMBRANE_ORGANIZATION | 137 | -0.370374 | -1.376872 | 0.01978 | 0.238003 |
| GOBP_POSITIVE_REGULATION_OF_NEURON_APOPTOTIC_PROCESS | 47 | -0.446772 | -1.375072 | 0.057026 | 0.240561 |
| GOBP_POSITIVE_REGULATION_OF_ACTIVATED_T_CELL_PROLIFERATION | 23 | -0.516454 | -1.37488 | 0.061538 | 0.240491 |
| GOBP_CHEMOKINE_PRODUCTION | 85 | -0.40184 | -1.37437 | 0.047423 | 0.240991 |
| GOBP_VASCULAR_ENDOTHELIAL_GROWTH_FACTOR_PRODUCTION | 30 | -0.485276 | -1.374276 | 0.07551 | 0.240758 |
| GOBP_ENDOPLASMIC_RETICULUM_TO_GOLGI_VESICLE_MEDIATED_TRANSPORT | 190 | -0.354103 | -1.374184 | 0.009434 | 0.240538 |
| GOBP_GLIAL_CELL_PROLIFERATION | 39 | -0.455637 | -1.374022 | 0.059574 | 0.240437 |
| GOBP_NEGATIVE_REGULATION_OF_TRANSPORT | 332 | -0.335074 | -1.373927 | 0.006818 | 0.240258 |
| GOBP_PH_REDUCTION | 51 | -0.43973 | -1.373846 | 0.053533 | 0.240052 |
| GOBP_REGULATION_OF_IMMUNOGLOBULIN_PRODUCTION | 63 | -0.424318 | -1.373328 | 0.044681 | 0.240532 |
| GOBP_REGULATION_OF_BINDING | 314 | -0.335605 | -1.37216 | 0 | 0.241983 |
| GOBP_EPITHELIAL_STRUCTURE_MAINTENANCE | 20 | -0.528343 | -1.37212 | 0.099617 | 0.241724 |
| GOBP_NEGATIVE_REGULATION_OF_ORGANELLE_ORGANIZATION | 312 | -0.337055 | -1.371977 | 0.009259 | 0.241631 |
| GOBP_REGULATION_OF_PEPTIDYL_LYSINE_ACETYLATION | 56 | -0.422056 | -1.371828 | 0.065217 | 0.241568 |
| GOBP_REGULATION_OF_TELOMERASE_ACTIVITY | 44 | -0.452327 | -1.370995 | 0.06405 | 0.242558 |
| GOBP_NEGATIVE_REGULATION_OF_PHAGOCYTOSIS | 20 | -0.527573 | -1.370851 | 0.091858 | 0.242399 |
| GOBP_REGULATION_OF_HORMONE_METABOLIC_PROCESS | 27 | -0.496056 | -1.370841 | 0.07582 | 0.242094 |
| GOBP_CELLULAR_RESPONSE_TO_ARSENIC_CONTAINING_SUBSTANCE | 18 | -0.54104 | -1.370619 | 0.09776 | 0.242111 |
| GOBP_TRABECULA_FORMATION | 17 | -0.541397 | -1.37003 | 0.112676 | 0.242659 |
| GOBP_CARDIAC_CHAMBER_MORPHOGENESIS | 78 | -0.40165 | -1.36977 | 0.044625 | 0.242741 |
| GOBP_RNA_DESTABILIZATION | 35 | -0.456447 | -1.369608 | 0.058696 | 0.242641 |
| GOBP_RESPONSE_TO_TEMPERATURE_STIMULUS | 198 | -0.347022 | -1.369296 | 0.014085 | 0.242797 |
| GOBP_TRANSITION_METAL_ION_HOMEOSTASIS | 111 | -0.379903 | -1.368874 | 0.027842 | 0.243144 |
| GOBP_POSITIVE_REGULATION_OF_NUCLEOTIDE_METABOLIC_PROCESS | 35 | -0.467526 | -1.36878 | 0.078431 | 0.242985 |
| GOBP_VASCULAR_TRANSPORT | 74 | -0.408481 | -1.368722 | 0.04185 | 0.242756 |
| GOBP_OSTEOCLAST_DIFFERENTIATION | 80 | -0.400171 | -1.368051 | 0.04955 | 0.243511 |
| GOBP_POSITIVE_REGULATION_OF_NEURON_DIFFERENTIATION | 59 | -0.423319 | -1.367806 | 0.060215 | 0.243577 |
| GOBP_REGULATION_OF_GLUCONEOGENESIS | 39 | -0.459992 | -1.367751 | 0.066667 | 0.243353 |
| GOBP_REGULATION_OF_LEUKOCYTE_MIGRATION | 176 | -0.35831 | -1.366768 | 0.018265 | 0.244512 |
| GOBP_REGULATION_OF_INTRINSIC_APOPTOTIC_SIGNALING_PATHWAY | 144 | -0.367532 | -1.366692 | 0.024943 | 0.244331 |
| GOBP_INFLAMMATORY_RESPONSE_TO_ANTIGENIC_STIMULUS | 44 | -0.443697 | -1.366133 | 0.083512 | 0.244807 |
| GOBP_POSITIVE_REGULATION_OF_INTRINSIC_APOPTOTIC_SIGNALING_PATHWAY | 53 | -0.428082 | -1.365966 | 0.04793 | 0.244733 |
| GOBP_NEGATIVE_REGULATION_OF_CHROMOSOME_ORGANIZATION | 87 | -0.392118 | -1.365565 | 0.034483 | 0.245039 |
| GOBP_T_HELPER_1_TYPE_IMMUNE_RESPONSE | 36 | -0.471264 | -1.365522 | 0.083521 | 0.244806 |
| GOBP_NEGATIVE_REGULATION_OF_DNA_BINDING_TRANSCRIPTION_FACTOR_ACTIVITY | 140 | -0.370026 | -1.365268 | 0.027897 | 0.244903 |
| GOBP_POSITIVE_REGULATION_OF_HEART_RATE | 16 | -0.554172 | -1.36458 | 0.093254 | 0.245678 |
| GOBP_REGULATION_OF_CELLULAR_RESPONSE_TO_GROWTH_FACTOR_STIMULUS | 228 | -0.347597 | -1.364198 | 0.009456 | 0.245951 |
| GOBP_ERK1_AND_ERK2_CASCADE | 242 | -0.344603 | -1.363902 | 0.012077 | 0.246111 |
| GOBP_LOW_DENSITY_LIPOPROTEIN_PARTICLE_CLEARANCE | 28 | -0.488325 | -1.363322 | 0.09465 | 0.246673 |
| GOBP_LABYRINTHINE_LAYER_MORPHOGENESIS | 15 | -0.563142 | -1.363102 | 0.10998 | 0.246731 |
| GOBP_SIALYLATION | 17 | -0.540964 | -1.362572 | 0.115768 | 0.247184 |
| GOBP_RESPONSE_TO_AMINE | 35 | -0.462002 | -1.362436 | 0.102564 | 0.247071 |
| GOBP_NEPHRON_MORPHOGENESIS | 46 | -0.438354 | -1.362307 | 0.076271 | 0.246976 |
| GOBP_REGULATION_OF_PROTEIN_ACETYLATION | 66 | -0.409116 | -1.362106 | 0.047826 | 0.246956 |
| GOBP_GLOMERULAR_EPITHELIUM_DEVELOPMENT | 18 | -0.542652 | -1.361897 | 0.106029 | 0.246948 |
| GOBP_POSITIVE_REGULATION_OF_IMMUNOGLOBULIN_PRODUCTION | 43 | -0.433462 | -1.361673 | 0.085653 | 0.246971 |
| GOBP_REGULATION_OF_IMMUNE_EFFECTOR_PROCESS | 404 | -0.326414 | -1.361319 | 0 | 0.247233 |
| GOBP_REGULATION_OF_PROTEIN_TARGETING_TO_MITOCHONDRION | 42 | -0.445257 | -1.361159 | 0.077586 | 0.247138 |
| GOBP_RESPONSE_TO_MECHANICAL_STIMULUS | 171 | -0.35724 | -1.360652 | 0.013423 | 0.247587 |
| GOBP_NADPH_REGENERATION | 18 | -0.531401 | -1.360369 | 0.116183 | 0.247712 |
| GOBP_NEGATIVE_REGULATION_OF_LIPID_LOCALIZATION | 37 | -0.450089 | -1.359708 | 0.100642 | 0.248441 |
| GOBP_REGULATION_OF_NOTCH_SIGNALING_PATHWAY | 93 | -0.381781 | -1.359607 | 0.038877 | 0.248292 |
| GOBP_REGULATION_OF_CELL_MIGRATION_INVOLVED_IN_SPROUTING_ANGIOGENESIS | 35 | -0.468563 | -1.359204 | 0.083665 | 0.248573 |
| GOBP_LYMPHOCYTE_MIGRATION | 94 | -0.388686 | -1.358537 | 0.034409 | 0.249326 |
| GOBP_NEGATIVE_REGULATION_OF_CELLULAR_PROTEIN_LOCALIZATION | 100 | -0.379745 | -1.357989 | 0.043668 | 0.249823 |

| | | | | | |
|---|---|---|---|---|---|
| GOBP_CELLULAR_DEFENSE_RESPONSE | 47 | 0.280226 | 0.844909 | 0.730129 | 0.941197 |
| GOBP_NUCLEOLAR_LARGE_RRNA_TRANSCRIPTION_BY_RNA_POLYMERASE_I | 17 | 0.344994 | 0.844151 | 0.682186 | 0.941629 |
| GOBP_RETINA_HOMEOSTASIS | 59 | 0.266594 | 0.84403 | 0.760618 | 0.940694 |
| GOBP_POSITIVE_REGULATION_OF_NITRIC_OXIDE_SYNTHASE_BIOSYNTHETIC_PROCESS | 15 | 0.35778 | 0.843238 | 0.664078 | 0.941176 |
| GOBP_PHOSPHOLIPID_METABOLIC_PROCESS | 367 | 0.208034 | 0.842733 | 0.952465 | 0.941103 |
| GOBP_TRANSCYTOSIS | 19 | 0.333477 | 0.842205 | 0.69505 | 0.941049 |
| GOBP_CHROMATIN_DISASSEMBLY | 18 | 0.344514 | 0.841885 | 0.685039 | 0.940548 |
| GOBP_SPINDLE_ORGANIZATION | 177 | 0.223521 | 0.841779 | 0.865961 | 0.939593 |
| GOBP_OOGENESIS | 66 | 0.261938 | 0.840566 | 0.766234 | 0.940922 |
| GOBP_POSITIVE_REGULATION_OF_FOCAL_ADHESION_ASSEMBLY | 19 | 0.334099 | 0.835902 | 0.702206 | 0.949461 |
| GOBP_POSITIVE_REGULATION_OF_CELL_SUBSTRATE_JUNCTION_ORGANIZATION | 24 | 0.315139 | 0.834179 | 0.703488 | 0.951804 |
| GOBP_NEURON_PROJECTION_ARBORIZATION | 25 | 0.309323 | 0.833228 | 0.704174 | 0.952505 |
| GOBP_POSITIVE_REGULATION_OF_PROTEIN_IMPORT | 37 | 0.287005 | 0.833009 | 0.745763 | 0.951738 |
| GOBP_REGULATION_OF_CHROMATIN_SILENCING | 15 | 0.347197 | 0.831791 | 0.699262 | 0.953011 |
| GOBP_PLATELET_ACTIVATION | 133 | 0.229195 | 0.82985 | 0.869489 | 0.955756 |
| GOBP_INTERFERON_ALPHA_PRODUCTION | 26 | 0.307185 | 0.829763 | 0.734962 | 0.954726 |
| GOBP_CHROMOSOME_CONDENSATION | 37 | 0.28552 | 0.829471 | 0.756052 | 0.954098 |
| GOBP_POSITIVE_REGULATION_OF_MYELINATION | 15 | 0.352399 | 0.828605 | 0.70099 | 0.954667 |
| GOBP_POSITIVE_REGULATION_OF_PROTEIN_CATABOLIC_PROCESS | 209 | 0.216129 | 0.828291 | 0.930153 | 0.954082 |
| GOBP_PHOSPHATIDYLETHANOLAMINE_METABOLIC_PROCESS | 27 | 0.307359 | 0.827042 | 0.714556 | 0.955347 |
| GOBP_ANTIGEN_RECEPTOR_MEDIATED_SIGNALING_PATHWAY | 288 | 0.208997 | 0.826791 | 0.932432 | 0.954642 |
| GOBP_PROTEIN_LOCALIZATION_TO_CELL_PERIPHERY | 287 | 0.207942 | 0.826726 | 0.924429 | 0.953581 |
| GOBP_POSITIVE_REGULATION_OF_PROTEIN_DEPOLYMERIZATION | 15 | 0.351108 | 0.826092 | 0.676413 | 0.953612 |
| GOBP_POSITIVE_REGULATION_OF_SIGNAL_TRANSDUCTION_BY_P53_CLASS_MEDIATOR | 21 | 0.322757 | 0.824802 | 0.706237 | 0.954971 |
| GOBP_GLYCEROPHOSPHOLIPID_METABOLIC_PROCESS | 310 | 0.204914 | 0.819142 | 0.962963 | 0.964747 |
| GOBP_PEPTIDYL_PROLINE_MODIFICATION | 51 | 0.264223 | 0.816426 | 0.780115 | 0.968619 |
| GOBP_INTRACELLULAR_LIPID_TRANSPORT | 39 | 0.277132 | 0.81588 | 0.774621 | 0.968458 |
| GOBP_POSITIVE_REGULATION_OF_RESPONSE_TO_BIOTIC_STIMULUS | 222 | 0.213414 | 0.811941 | 0.935829 | 0.974467 |
| GOBP_NEGATIVE_REGULATION_OF_CALCIUM_ION_TRANSMEMBRANE_TRANSPORTER_A | 24 | 0.306577 | 0.810542 | 0.754098 | 0.97582 |
| GOBP_SPHINGOLIPID_MEDIATED_SIGNALING_PATHWAY | 15 | 0.340798 | 0.810369 | 0.727447 | 0.974926 |
| GOBP_POSITIVE_REGULATION_OF_LEUKOCYTE_APOPTOTIC_PROCESS | 22 | 0.306992 | 0.808695 | 0.759152 | 0.976718 |
| GOBP_MAINTENANCE_OF_PROTEIN_LOCALIZATION_IN_ORGANELLE | 39 | 0.268291 | 0.803064 | 0.788044 | 0.985503 |
| GOBP_REGULATION_OF_RESPIRATORY_BURST | 15 | 0.334385 | 0.803026 | 0.736842 | 0.984357 |
| GOBP_RRNA_TRANSCRIPTION | 33 | 0.280888 | 0.802418 | 0.787194 | 0.984175 |
| GOBP_NEGATIVE_REGULATION_OF_TUMOR_NECROSIS_FACTOR_MEDIATED_SIGNALING | 16 | 0.330235 | 0.801683 | 0.731061 | 0.984243 |
| GOBP_CORTICAL_CYTOSKELETON_ORGANIZATION | 56 | 0.256927 | 0.799914 | 0.811651 | 0.986045 |
| GOBP_IMMUNE_RESPONSE_TO_TUMOR_CELL | 18 | 0.313777 | 0.79689 | 0.74031 | 0.989841 |
| GOBP_INOSITOL_PHOSPHATE_MEDIATED_SIGNALING | 43 | 0.261561 | 0.796513 | 0.812741 | 0.989245 |
| GOBP_REGULATION_OF_PROTEIN_LOCALIZATION_TO_PLASMA_MEMBRANE | 84 | 0.235329 | 0.796357 | 0.863636 | 0.988301 |
| GOBP_PHOSPHOLIPID_BIOSYNTHETIC_PROCESS | 243 | 0.206432 | 0.796116 | 0.954225 | 0.987482 |
| GOBP_PRIMARY_ALCOHOL_METABOLIC_PROCESS | 64 | 0.24774 | 0.795469 | 0.876384 | 0.987377 |
| GOBP_NEGATIVE_REGULATION_OF_NECROTIC_CELL_DEATH | 16 | 0.325225 | 0.792157 | 0.754352 | 0.991665 |
| GOBP_REGULATION_OF_WATER_LOSS_VIA_SKIN | 21 | 0.304784 | 0.792086 | 0.754286 | 0.990577 |
| GOBP_BLASTOCYST_DEVELOPMENT | 84 | 0.233041 | 0.791364 | 0.878307 | 0.990519 |
| GOBP_ACETYL_COA_METABOLIC_PROCESS | 35 | 0.27749 | 0.790448 | 0.823423 | 0.990791 |
| GOBP_PROTEIN_LOCALIZATION_TO_CELL_SURFACE | 59 | 0.251636 | 0.788738 | 0.850467 | 0.992223 |
| GOBP_LAMELLIPODIUM_ORGANIZATION | 78 | 0.23542 | 0.788697 | 0.863216 | 0.991082 |
| GOBP_REGULATION_OF_MEMBRANE_PROTEIN_ECTODOMAIN_PROTEOLYSIS | 23 | 0.294699 | 0.787257 | 0.774319 | 0.992137 |
| GOBP_HEPARAN_SULFATE_PROTEOGLYCAN_METABOLIC_PROCESS | 27 | 0.289535 | 0.785733 | 0.787659 | 0.993343 |
| GOBP_SULFUR_COMPOUND_METABOLIC_PROCESS | 319 | 0.196746 | 0.785373 | 0.994872 | 0.992721 |
| GOBP_POSITIVE_REGULATION_OF_JUN_KINASE_ACTIVITY | 62 | 0.244064 | 0.785003 | 0.859779 | 0.99207 |
| GOBP_HEPATOCYTE_GROWTH_FACTOR_RECEPTOR_SIGNALING_PATHWAY | 15 | 0.322434 | 0.784889 | 0.762452 | 0.991054 |
| GOBP_TRANSCRIPTION_PREINITIATION_COMPLEX_ASSEMBLY | 35 | 0.270136 | 0.783195 | 0.813187 | 0.99243 |
| GOBP_RIBOSOMAL_SMALL_SUBUNIT_ASSEMBLY | 21 | 0.30197 | 0.779909 | 0.76616 | 0.996084 |
| GOBP_HEMATOPOIETIC_PROGENITOR_CELL_DIFFERENTIATION | 150 | 0.21305 | 0.779405 | 0.931035 | 0.995636 |
| GOBP_NADP_METABOLIC_PROCESS | 33 | 0.271663 | 0.778397 | 0.791188 | 0.995922 |
| GOBP_POSITIVE_REGULATION_OF_CELL_MATRIX_ADHESION | 45 | 0.25744 | 0.778024 | 0.84375 | 0.995297 |
| GOBP_HEPARAN_SULFATE_PROTEOGLYCAN_BIOSYNTHETIC_PROCESS | 23 | 0.294562 | 0.772641 | 0.783109 | 1 |
| GOBP_POSITIVE_REGULATION_OF_CREB_TRANSCRIPTION_FACTOR_ACTIVITY | 15 | 0.325368 | 0.772095 | 0.770992 | 1 |
| GOBP_REGULATION_OF_ALCOHOL_BIOSYNTHETIC_PROCESS | 63 | 0.23911 | 0.769544 | 0.876611 | 1 |
| GOBP_POSITIVE_REGULATION_OF_NITRIC_OXIDE_SYNTHASE_ACTIVITY | 21 | 0.297239 | 0.769159 | 0.826742 | 1 |
| GOBP_NEGATIVE_REGULATION_OF_FIBROBLAST_PROLIFERATION | 26 | 0.282895 | 0.765726 | 0.806216 | 1 |
| GOBP_RNA_POLYMERASE_II_PREINITIATION_COMPLEX_ASSEMBLY | 23 | 0.288739 | 0.765537 | 0.797131 | 1 |
| GOBP_NEGATIVE_REGULATION_OF_T_CELL_RECEPTOR_SIGNALING_PATHWAY | 20 | 0.297894 | 0.765232 | 0.792181 | 1 |
| GOBP_ENDOCYTIC_RECYCLING | 48 | 0.247918 | 0.763846 | 0.86692 | 1 |
| GOBP_POSITIVE_REGULATION_OF_PROTEIN_DEPHOSPHORYLATION | 42 | 0.254939 | 0.76146 | 0.845324 | 1 |
| GOBP_TOLL_LIKE_RECEPTOR_4_SIGNALING_PATHWAY | 35 | 0.266348 | 0.757527 | 0.84252 | 1 |
| GOBP_PHOSPHATIDIC_ACID_METABOLIC_PROCESS | 40 | 0.253983 | 0.757317 | 0.857692 | 1 |
| GOBP_ALCOHOL_METABOLIC_PROCESS | 295 | 0.193263 | 0.757017 | 0.996403 | 1 |
| GOBP_HISTONE_H2A_ACETYLATION | 17 | 0.300428 | 0.75237 | 0.79925 | 1 |
| GOBP_POSITIVE_REGULATION_OF_ANTIGEN_RECEPTOR_MEDIATED_SIGNALING_PATHW | 21 | 0.294215 | 0.749481 | 0.821632 | 1 |
| GOBP_CD4_POSITIVE_OR_CD8_POSITIVE_ALPHA_BETA_T_CELL_LINEAGE_COMMITMENT | 21 | 0.285069 | 0.748067 | 0.785448 | 1 |
| GOBP_REGULATION_OF_CYSTEINE_TYPE_ENDOPEPTIDASE_ACTIVITY_INVOLVED_IN_AP | 16 | 0.302833 | 0.743885 | 0.808743 | 1 |
| GOBP_INTERLEUKIN_4_PRODUCTION | 31 | 0.262152 | 0.74296 | 0.872928 | 1 |
| GOBP_REGULATION_OF_POLYSACCHARIDE_BIOSYNTHETIC_PROCESS | 32 | 0.258062 | 0.742806 | 0.863905 | 1 |
| GOBP_CHOLESTEROL_EFFLUX | 38 | 0.253035 | 0.742262 | 0.876611 | 1 |
| GOBP_3_UTR_MEDIATED_MRNA_STABILIZATION | 18 | 0.299672 | 0.740338 | 0.826996 | 1 |
| GOBP_POSITIVE_REGULATION_OF_TRANSCRIPTION_INITIATION_FROM_RNA_POLYMERAS | 25 | 0.270856 | 0.73946 | 0.851272 | 1 |
| GOBP_VERY_LONG_CHAIN_FATTY_ACID_METABOLIC_PROCESS | 32 | 0.255147 | 0.737501 | 0.865878 | 1 |
| GOBP_SULFUR_COMPOUND_BIOSYNTHETIC_PROCESS | 164 | 0.198165 | 0.736811 | 0.986965 | 1 |
| GOBP_REGULATION_OF_ANTIGEN_RECEPTOR_MEDIATED_SIGNALING_PATHWAY | 61 | 0.230572 | 0.736795 | 0.904135 | 1 |
| GOBP_TELOMERASE_RNA_LOCALIZATION | 19 | 0.285783 | 0.731296 | 0.852391 | 1 |
| GOBP_T_HELPER_CELL_LINEAGE_COMMITMENT | 16 | 0.308551 | 0.731286 | 0.831094 | 1 |
| GOBP_REGULATION_OF_MRNA_CATABOLIC_PROCESS | 193 | 0.192333 | 0.729483 | 0.985586 | 1 |
| GOBP_VIRAL_GENE_EXPRESSION | 194 | 0.18955 | 0.721913 | 0.99297 | 1 |
| GOBP_COENZYME_A_METABOLIC_PROCESS | 15 | 0.293072 | 0.720865 | 0.846743 | 1 |
| GOBP_PEROXISOMAL_MEMBRANE_TRANSPORT | 19 | 0.285212 | 0.719892 | 0.837209 | 1 |
| GOBP_MITOCHONDRIAL_TRANSLATION | 133 | 0.198212 | 0.719273 | 0.98155 | 1 |
| GOBP_REGULATION_OF_T_CELL_RECEPTOR_SIGNALING_PATHWAY | 38 | 0.241634 | 0.717795 | 0.900192 | 1 |
| GOBP_HYALURONAN_CATABOLIC_PROCESS | 16 | 0.297151 | 0.71547 | 0.831683 | 1 |
| GOBP_RESPONSE_TO_IRON_ION | 24 | 0.270887 | 0.715057 | 0.868885 | 1 |
| GOBP_PSEUDOURIDINE_SYNTHESIS | 18 | 0.286059 | 0.710752 | 0.83463 | 1 |
| GOBP_BETA_CATENIN_TCF_COMPLEX_ASSEMBLY | 31 | 0.251759 | 0.708495 | 0.897143 | 1 |
| GOBP_POSITIVE_REGULATION_OF_INTERLEUKIN_4_PRODUCTION | 24 | 0.266912 | 0.706931 | 0.895911 | 1 |
| GOBP_PHOSPHORYLATED_CARBOHYDRATE_DEPHOSPHORYLATION | 16 | 0.296821 | 0.706901 | 0.846296 | 1 |
| GOBP_MYELIN_ASSEMBLY | 18 | 0.283271 | 0.706464 | 0.856867 | 1 |
| GOBP_PHOSPHATIDYLINOSITOL_METABOLIC_PROCESS | 166 | 0.189996 | 0.706011 | 0.996344 | 1 |
| GOBP_NUCLEOTIDE_EXCISION_REPAIR_PREINCISION_COMPLEX_STABILIZATION | 21 | 0.275529 | 0.704356 | 0.866534 | 1 |
| GOBP_REGULATION_OF_ARP2_3_COMPLEX_MEDIATED_ACTIN_NUCLEATION | 20 | 0.275587 | 0.703786 | 0.858491 | 1 |
| GOBP_RESPONSE_TO_TUMOR_CELL | 23 | 0.265785 | 0.699001 | 0.870301 | 1 |
| GOBP_LOCALIZATION_WITHIN_MEMBRANE | 61 | 0.218553 | 0.69895 | 0.946886 | 1 |
| GOBP_REGULATION_OF_CHOLESTEROL_BIOSYNTHETIC_PROCESS | 40 | 0.231281 | 0.69694 | 0.931298 | 1 |
| GOBP_PRODUCTION_OF_SMALL_RNA_INVOLVED_IN_GENE_SILENCING_BY_RNA | 50 | 0.227038 | 0.694206 | 0.938517 | 1 |
| GOBP_RECEPTOR_CLUSTERING | 43 | 0.23357 | 0.692325 | 0.94347 | 1 |
| GOBP_CELL_DEATH_IN_RESPONSE_TO_HYDROGEN_PEROXIDE | 22 | 0.263294 | 0.687486 | 0.898406 | 1 |
| GOBP_GLUTAMATE_METABOLIC_PROCESS | 27 | 0.255595 | 0.687107 | 0.905882 | 1 |
| GOBP_REGULATION_OF_GLUCAN_BIOSYNTHETIC_PROCESS | 25 | 0.255232 | 0.685114 | 0.897143 | 1 |
| GOBP_POSITIVE_REGULATION_OF_TELOMERASE_RNA_LOCALIZATION_TO_CAJAL_BODY | 15 | 0.285728 | 0.678643 | 0.870079 | 1 |
| GOBP_RIBOSOMAL_LARGE_SUBUNIT_ASSEMBLY | 28 | 0.248631 | 0.677474 | 0.917355 | 1 |
| GOBP_REGULATION_OF_VIRAL_INDUCED_CYTOPLASMIC_PATTERN_RECOGNITION_REC | 21 | 0.261636 | 0.676422 | 0.906667 | 1 |
| GOBP_HISTONE_H4_K5_ACETYLATION | 16 | 0.282727 | 0.672961 | 0.864542 | 1 |
| GOBP_NEGATIVE_REGULATION_OF_PROTEIN_KINASE_B_SIGNALING | 41 | 0.227469 | 0.672322 | 0.957916 | 1 |
| GOBP_REGULATION_OF_CELLULAR_AMINO_ACID_METABOLIC_PROCESS | 54 | 0.214804 | 0.671905 | 0.973129 | 1 |
| GOBP_GLOBAL_GENOME_NUCLEOTIDE_EXCISION_REPAIR | 26 | 0.239653 | 0.669904 | 0.936508 | 1 |
| GOBP_MRNA_CLEAVAGE | 21 | 0.261623 | 0.664064 | 0.898039 | 1 |
| GOBP_GLUTAMINE_FAMILY_AMINO_ACID_CATABOLIC_PROCESS | 22 | 0.25681 | 0.663752 | 0.895582 | 1 |
| GOBP_CORTICAL_ACTIN_CYTOSKELETON_ORGANIZATION | 37 | 0.228849 | 0.660947 | 0.958167 | 1 |
| GOBP_REGULATION_OF_STEROID_BIOSYNTHETIC_PROCESS | 70 | 0.201662 | 0.660447 | 0.988806 | 1 |
| GOBP_REGULATION_OF_CALCINEURIN_MEDIATED_SIGNALING | 26 | 0.245752 | 0.658629 | 0.947674 | 1 |
| GOBP_NUCLEAR_MIGRATION | 18 | 0.274317 | 0.658307 | 0.902622 | 1 |
| GOBP_ESTABLISHMENT_OF_PIGMENT_GRANULE_LOCALIZATION | 21 | 0.252688 | 0.656398 | 0.911488 | 1 |
| GOBP_FATTY_ACYL_COA_BIOSYNTHETIC_PROCESS | 29 | 0.238127 | 0.652409 | 0.936782 | 1 |
| GOBP_REGULATION_OF_TOLL_LIKE_RECEPTOR_4_SIGNALING_PATHWAY | 19 | 0.254523 | 0.647829 | 0.929524 | 1 |
| GOBP_PEROXISOMAL_TRANSPORT | 66 | 0.198329 | 0.64584 | 0.994129 | 1 |
| GOBP_DIACYLGLYCEROL_METABOLIC_PROCESS | 19 | 0.258302 | 0.645327 | 0.912351 | 1 |
| GOBP_HISTONE_H4_K16_ACETYLATION | 21 | 0.245411 | 0.641733 | 0.930368 | 1 |
| GOBP_RNA_INTERFERENCE | 16 | 0.265304 | 0.641096 | 0.937262 | 1 |
| GOBP_REGULATION_OF_INTRACELLULAR_ESTROGEN_RECEPTOR_SIGNALING_PATHWA | 36 | 0.219072 | 0.640988 | 0.946 | 1 |
| GOBP_MITOCHONDRIAL_CYTOCHROME_C_OXIDASE_ASSEMBLY | 21 | 0.243616 | 0.639656 | 0.922925 | 1 |
| GOBP_RESPONSE_TO_STEROL | 28 | 0.232094 | 0.638334 | 0.946215 | 1 |
| GOBP_INSULIN_SECRETION_INVOLVED_IN_CELLULAR_RESPONSE_TO_GLUCOSE_STIMU | 52 | 0.206411 | 0.636911 | 0.972527 | 1 |
| GOBP_PROTEIN_AUTOPROCESSING | 17 | 0.262118 | 0.633833 | 0.924953 | 1 |
| GOBP_NUCLEOSIDE_BISPHOSPHATE_METABOLIC_PROCESS | 122 | 0.178421 | 0.633087 | 0.998236 | 1 |
| GOBP_V_D_J_RECOMBINATION | 16 | 0.258844 | 0.631736 | 0.914611 | 1 |
| GOBP_REGULATION_OF_CELLULAR_AMINE_METABOLIC_PROCESS | 65 | 0.195272 | 0.628404 | 0.985455 | 1 |
| GOBP_PIGMENT_GRANULE_LOCALIZATION | 23 | 0.241144 | 0.624864 | 0.955684 | 1 |
| GOBP_PROTEIN_TARGETING_TO_MEMBRANE | 192 | 0.164079 | 0.623685 | 1 | 1 |
| GOBP_REGULATION_OF_AMYLOID_PRECURSOR_PROTEIN_CATABOLIC_PROCESS | 41 | 0.207771 | 0.620041 | 0.984496 | 1 |
| GOBP_TRANSCRIPTION_INITIATION_FROM_RNA_POLYMERASE_I_PROMOTER | 38 | 0.209383 | 0.619025 | 0.972277 | 1 |
| GOBP_NUCLEOTIDE_EXCISION_REPAIR_DNA_DUPLEX_UNWINDING | 22 | 0.236431 | 0.618244 | 0.950593 | 1 |
| GOBP_SHORT_CHAIN_FATTY_ACID_METABOLIC_PROCESS | 15 | 0.255532 | 0.616212 | 0.934 | 1 |
| GOBP_NEGATIVE_REGULATION_OF_DEFENSE_RESPONSE_TO_VIRUS | 24 | 0.231688 | 0.61097 | 0.96008 | 1 |
| GOBP_NUCLEOTIDE_EXCISION_REPAIR_DNA_INCISION | 39 | 0.209926 | 0.61019 | 0.973231 | 1 |
| GOBP_PROTEIN_LOCALIZATION_TO_GOLGI_APPARATUS | 27 | 0.22485 | 0.602588 | 0.964143 | 1 |
| GOBP_CELLULAR_RESPONSE_TO_STEROL_DEPLETION | 16 | 0.25367 | 0.601845 | 0.936759 | 1 |
| GOBP_MITOCHONDRIAL_TRANSCRIPTION | 17 | 0.242946 | 0.600669 | 0.924528 | 1 |
| GOBP_NUCLEOTIDE_EXCISION_REPAIR_DNA_DAMAGE_RECOGNITION | 23 | 0.230208 | 0.598363 | 0.958559 | 1 |
| GOBP_PEROXISOME_ORGANIZATION | 78 | 0.178 | 0.591238 | 0.996473 | 1 |
| GOBP_RECEPTOR_CATABOLIC_PROCESS | 33 | 0.206153 | 0.589497 | 0.974265 | 1 |
| GOBP_RIBOSOME_ASSEMBLY | 63 | 0.183589 | 0.588508 | 0.992453 | 1 |
| GOBP_MITOCHONDRIAL_GENE_EXPRESSION | 164 | 0.155592 | 0.580018 | 1 | 1 |
| GOBP_AMYLOID_FIBRIL_FORMATION | 22 | 0.223829 | 0.576569 | 0.963168 | 1 |
| GOBP_NUCLEOTIDE_EXCISION_REPAIR_PREINCISION_COMPLEX_ASSEMBLY | 29 | 0.209824 | 0.575243 | 0.969582 | 1 |
| GOBP_MITOCHONDRIAL_TRANSLATIONAL_TERMINATION | 88 | 0.168504 | 0.570848 | 1 | 1 |
| GOBP_HISTONE_METHYLATION | 118 | 0.1602 | 0.566164 | 1 | 1 |
| GOBP_STRESS_GRANULE_ASSEMBLY | 21 | 0.221728 | 0.566072 | 0.971319 | 1 |
| GOBP_PHOTOPERIODISM | 29 | 0.207453 | 0.563468 | 0.984704 | 1 |

| | | | | | |
|---|---|---|---|---|---|
| GOBP_LEUKOCYTE_CELL_CELL_ADHESION | 329 | -0.331939 | -1.357948 | 0.011962 | 0.249585 |
| GOBP_POSITIVE_REGULATION_OF_TRANSMEMBRANE_TRANSPORT | 161 | -0.358104 | -1.35736 | 0.025478 | 0.250201 |
| GOBP_KERATAN_SULFATE_METABOLIC_PROCESS | 26 | -0.495556 | -1.357044 | 0.112798 | 0.250378 |
| GOBP_POSITIVE_REGULATION_OF_EPITHELIAL_CELL_PROLIFERATION | 144 | -0.363192 | -1.356682 | 0.028986 | 0.250671 |
| GOBP_POSITIVE_REGULATION_OF_PHOSPHATASE_ACTIVITY | 30 | -0.478297 | -1.354038 | 0.094949 | 0.254644 |
| GOBP_REGULATION_OF_AMINO_ACID_TRANSPORT | 25 | -0.495157 | -1.353863 | 0.102823 | 0.254588 |
| GOBP_MULTICELLULAR_ORGANISMAL_RESPONSE_TO_STRESS | 44 | -0.450509 | -1.353593 | 0.063786 | 0.254675 |
| GOBP_TAIL_ANCHORED_MEMBRANE_PROTEIN_INSERTION_INTO_ER_MEMBRANE | 16 | -0.546291 | -1.353552 | 0.13278 | 0.254424 |
| GOBP_VIRAL_RNA_GENOME_REPLICATION | 33 | -0.459261 | -1.353452 | 0.081448 | 0.254261 |
| GOBP_NEGATIVE_REGULATION_OF_CARBOHYDRATE_METABOLIC_PROCESS | 42 | -0.441426 | -1.352922 | 0.076152 | 0.254773 |
| GOBP_LYMPHOCYTE_HOMEOSTASIS | 54 | -0.43405 | -1.352747 | 0.07064 | 0.254717 |
| GOBP_NUCLEOTIDE_PHOSPHORYLATION | 118 | -0.375398 | -1.352321 | 0.035635 | 0.255045 |
| GOBP_RESPONSE_TO_ZINC_ION | 36 | -0.45428 | -1.351932 | 0.063158 | 0.255431 |
| GOBP_POSITIVE_REGULATION_OF_BMP_SIGNALING_PATHWAY | 26 | -0.493913 | -1.351144 | 0.089431 | 0.256291 |
| GOBP_NEGATIVE_REGULATION_OF_INTERLEUKIN_8_PRODUCTION | 21 | -0.504865 | -1.350553 | 0.11245 | 0.257004 |
| GOBP_REGULATION_OF_PROTEASOMAL_PROTEIN_CATABOLIC_PROCESS | 169 | -0.353241 | -1.350302 | 0.018824 | 0.257079 |
| GOBP_REGULATION_OF_DNA_REPLICATION | 102 | -0.381977 | -1.350227 | 0.039216 | 0.256854 |
| GOBP_TRABECULA_MORPHOGENESIS | 34 | -0.462129 | -1.349995 | 0.089109 | 0.256893 |
| GOBP_ORGANELLE_FISSION | 445 | -0.320021 | -1.349865 | 0 | 0.256788 |
| GOBP_ESTABLISHMENT_OF_ORGANELLE_LOCALIZATION | 402 | -0.32262 | -1.349787 | 0.004878 | 0.256566 |
| GOBP_EPITHELIAL_CELL_PROLIFERATION | 307 | -0.329266 | -1.349688 | 0 | 0.256436 |
| GOBP_ESTABLISHMENT_OF_MITOTIC_SPINDLE_LOCALIZATION | 32 | -0.468183 | -1.348164 | 0.109054 | 0.258525 |
| GOBP_POSITIVE_REGULATION_OF_PROTEIN_ACETYLATION | 40 | -0.452568 | -1.347955 | 0.079002 | 0.258574 |
| GOBP_PEPTIDYL_SERINE_DEPHOSPHORYLATION | 17 | -0.531646 | -1.347389 | 0.10643 | 0.259142 |
| GOBP_AXONEME_ASSEMBLY | 58 | -0.409418 | -1.34732 | 0.056645 | 0.258924 |
| GOBP_REGULATION_OF_ORGANELLE_ASSEMBLY | 167 | -0.357353 | -1.347114 | 0.031818 | 0.258909 |
| GOBP_FOLIC_ACID_METABOLIC_PROCESS | 18 | -0.532866 | -1.346208 | 0.1 | 0.25999 |
| GOBP_OVARIAN_FOLLICLE_DEVELOPMENT | 42 | -0.44137 | -1.346191 | 0.099783 | 0.259699 |
| GOBP_NUCLEOSOME_ASSEMBLY | 114 | -0.374816 | -1.346084 | 0.043478 | 0.259531 |
| GOBP_NEGATIVE_REGULATION_OF_MITOCHONDRION_ORGANIZATION | 44 | -0.440233 | -1.345983 | 0.067762 | 0.259358 |
| GOBP_CELLULAR_RESPONSE_TO_KETONE | 74 | -0.402326 | -1.345952 | 0.04918 | 0.25909 |
| GOBP_CHROMATIN_ORGANIZATION_INVOLVED_IN_REGULATION_OF_TRANSCRIPTION | 130 | -0.364912 | -1.345812 | 0.023605 | 0.259007 |
| GOBP_MIDBODY_ABSCISSION | 17 | -0.545693 | -1.345706 | 0.143177 | 0.258875 |
| GOBP_RENAL_TUBULE_DEVELOPMENT | 61 | -0.406723 | -1.345693 | 0.061441 | 0.258579 |
| GOBP_REGULATION_OF_LEUKOCYTE_CHEMOTAXIS | 102 | -0.377581 | -1.345414 | 0.047414 | 0.258703 |
| GOBP_NEGATIVE_REGULATION_OF_GENE_EXPRESSION_EPIGENETIC | 102 | -0.375315 | -1.345149 | 0.037037 | 0.258829 |
| GOBP_REGULATION_OF_VASCULAR_PERMEABILITY | 35 | -0.46691 | -1.344906 | 0.093555 | 0.258919 |
| GOBP_OUTER_DYNEIN_ARM_ASSEMBLY | 15 | -0.548623 | -1.344815 | 0.106029 | 0.25872 |
| GOBP_REGULATION_OF_JNK_CASCADE | 116 | -0.371444 | -1.343358 | 0.034935 | 0.260809 |
| GOBP_REGULATION_OF_GLUCOSE_IMPORT | 45 | -0.435713 | -1.343324 | 0.099537 | 0.260551 |
| GOBP_MESODERMAL_CELL_DIFFERENTIATION | 20 | -0.518691 | -1.343229 | 0.114919 | 0.260392 |
| GOBP_GLYCOSYLCERAMIDE_METABOLIC_PROCESS | 17 | -0.536049 | -1.34316 | 0.136364 | 0.260212 |
| GOBP_SUPEROXIDE_METABOLIC_PROCESS | 56 | -0.41711 | -1.342654 | 0.078891 | 0.260835 |
| GOBP_RELEASE_OF_CYTOCHROME_C_FROM_MITOCHONDRIA | 52 | -0.422946 | -1.342533 | 0.065678 | 0.260704 |
| GOBP_RESPONSE_TO_CATECHOLAMINE | 68 | -0.400963 | -1.339955 | 0.070776 | 0.264553 |
| GOBP_MITOCHONDRIAL_TRANSPORT | 256 | -0.332909 | -1.339328 | 0.009569 | 0.265201 |
| GOBP_MAGNESIUM_ION_TRANSPORT | 15 | -0.554196 | -1.338992 | 0.125498 | 0.265396 |
| GOBP_GENERATION_OF_PRECURSOR_METABOLITES_AND_ENERGY | 465 | -0.31492 | -1.338739 | 0 | 0.265474 |
| GOBP_RESPONSE_TO_KETONE | 156 | -0.357922 | -1.338571 | 0.024331 | 0.265429 |
| GOBP_REGULATION_OF_METAL_ION_TRANSPORT | 189 | -0.344795 | -1.338103 | 0.015801 | 0.265812 |
| GOBP_PROTEASOMAL_PROTEIN_CATABOLIC_PROCESS | 456 | -0.316954 | -1.337262 | 0 | 0.266805 |
| GOBP_VITAMIN_TRANSPORT | 30 | -0.465014 | -1.337086 | 0.099567 | 0.266751 |
| GOBP_RECEPTOR_INTERNALIZATION | 86 | -0.379153 | -1.336184 | 0.058455 | 0.267928 |
| GOBP_L_AMINO_ACID_TRANSPORT | 52 | -0.423819 | -1.336172 | 0.078723 | 0.267626 |
| GOBP_GLIOGENESIS | 232 | -0.336867 | -1.336096 | 0.020737 | 0.267425 |
| GOBP_HYDROGEN_PEROXIDE_BIOSYNTHETIC_PROCESS | 15 | -0.555252 | -1.335235 | 0.138158 | 0.268464 |
| GOBP_CELLULAR_PROCESS_INVOLVED_IN_REPRODUCTION_IN_MULTICELLULAR_ORGANI | 249 | -0.337647 | -1.335072 | 0.011287 | 0.268422 |
| GOBP_REGULATION_OF_CELL_CYCLE_G1_S_PHASE_TRANSITION | 155 | -0.34884 | -1.334692 | 0.039301 | 0.268747 |
| GOBP_REGULATION_OF_CELL_CELL_ADHESION | 379 | -0.319521 | -1.334277 | 0.004717 | 0.269131 |
| GOBP_LABYRINTHINE_LAYER_DEVELOPMENT | 36 | -0.451404 | -1.333204 | 0.096907 | 0.270545 |
| GOBP_DNA_METHYLATION_OR_DEMETHYLATION | 80 | -0.385347 | -1.332884 | 0.046154 | 0.270795 |
| GOBP_CELLULAR_RESPONSE_TO_STEROID_HORMONE_STIMULUS | 162 | -0.347884 | -1.332288 | 0.028139 | 0.271476 |
| GOBP_POSITIVE_REGULATION_OF_T_CELL_PROLIFERATION | 89 | -0.379762 | -1.329971 | 0.042283 | 0.274946 |
| GOBP_UBIQUITIN_DEPENDENT_ERAD_PATHWAY | 75 | -0.388732 | -1.329872 | 0.055066 | 0.274804 |
| GOBP_REGULATION_OF_VASCULAR_ASSOCIATED_SMOOTH_MUSCLE_CELL_PROLIFERATI | 47 | -0.419492 | -1.329833 | 0.0783 | 0.274534 |
| GOBP_MICROTUBULE_BASED_TRANSPORT | 178 | -0.345822 | -1.329405 | 0.014799 | 0.274911 |
| GOBP_REPLACEMENT_OSSIFICATION | 26 | -0.483355 | -1.329332 | 0.112705 | 0.274705 |
| GOBP_NEGATIVE_REGULATION_OF_ION_TRANSPORT | 234 | -0.337919 | -1.328446 | 0.023419 | 0.275745 |
| GOBP_LONG_TERM_MEMORY | 21 | -0.512007 | -1.328039 | 0.12095 | 0.27606 |
| GOBP_POSITIVE_REGULATION_OF_DNA_METABOLIC_PROCESS | 185 | -0.345062 | -1.327416 | 0.028369 | 0.276801 |
| GOBP_REGULATION_OF_NUCLEOTIDE_BIOSYNTHETIC_PROCESS | 33 | -0.463159 | -1.327279 | 0.099594 | 0.276733 |
| GOBP_TRANSFORMING_GROWTH_FACTOR_BETA_RECEPTOR_SIGNALING_PATHWAY | 171 | -0.347848 | -1.327004 | 0.033333 | 0.276833 |
| GOBP_MRNA_EXPORT_FROM_NUCLEUS | 107 | -0.363337 | -1.326636 | 0.037363 | 0.27716 |
| GOBP_B_CELL_DIFFERENTIATION | 115 | -0.367298 | -1.326286 | 0.05298 | 0.277386 |
| GOBP_RESPONSE_TO_CADMIUM_ION | 47 | -0.427826 | -1.3261 | 0.078775 | 0.277417 |
| GOBP_NEGATIVE_REGULATION_OF_NF_KAPPAB_TRANSCRIPTION_FACTOR_ACTIVITY | 75 | -0.392095 | -1.325765 | 0.069663 | 0.277671 |
| GOBP_NEGATIVE_REGULATION_OF_COAGULATION | 38 | -0.43969 | -1.325613 | 0.082073 | 0.277609 |
| GOBP_RESPONSE_TO_INTERLEUKIN_6 | 36 | -0.447453 | -1.325394 | 0.114224 | 0.277638 |
| GOBP_CELLULAR_RESPONSE_TO_INTERFERON_BETA | 18 | -0.515825 | -1.325336 | 0.109731 | 0.277419 |
| GOBP_POSTSYNAPTIC_SPECIALIZATION_ORGANIZATION | 24 | -0.493745 | -1.325126 | 0.120332 | 0.277456 |
| GOBP_POSITIVE_REGULATION_OF_INTERLEUKIN_8_PRODUCTION | 52 | -0.419143 | -1.324984 | 0.082627 | 0.277391 |
| GOBP_CELLULAR_RESPONSE_TO_REACTIVE_OXYGEN_SPECIES | 133 | -0.355273 | -1.323967 | 0.036876 | 0.278747 |
| GOBP_POSITIVE_REGULATION_OF_INTERLEUKIN_17_PRODUCTION | 15 | -0.556135 | -1.323819 | 0.12831 | 0.278672 |
| GOBP_REGULATION_OF_VASCULATURE_DEVELOPMENT | 231 | -0.336851 | -1.323817 | 0.028571 | 0.278357 |
| GOBP_CILIUM_ORGANIZATION | 362 | -0.3214 | -1.323604 | 0.002353 | 0.278436 |
| GOBP_CELLULAR_RESPONSE_TO_LOW_DENSITY_LIPOPROTEIN_PARTICLE_STIMULUS | 15 | -0.551751 | -1.323138 | 0.134538 | 0.278818 |
| GOBP_POSITIVE_REGULATION_OF_BIOMINERALIZATION | 37 | -0.438121 | -1.322958 | 0.097826 | 0.278802 |
| GOBP_LENS_FIBER_CELL_DIFFERENTIATION | 24 | -0.493096 | -1.322551 | 0.110879 | 0.279183 |
| GOBP_REGULATION_OF_EPITHELIAL_CELL_DIFFERENTIATION | 110 | -0.367259 | -1.322473 | 0.065217 | 0.279009 |
| GOBP_REGULATION_OF_VESICLE_MEDIATED_TRANSPORT | 437 | -0.312056 | -1.32224 | 0.002336 | 0.279102 |
| GOBP_PHYSIOLOGICAL_CARDIAC_MUSCLE_HYPERTROPHY | 21 | -0.495144 | -1.320838 | 0.126531 | 0.281124 |
| GOBP_L_ALPHA_AMINO_ACID_TRANSMEMBRANE_TRANSPORT | 48 | -0.424013 | -1.320535 | 0.081633 | 0.281304 |
| GOBP_AXIS_SPECIFICATION | 58 | -0.407498 | -1.320482 | 0.067485 | 0.281101 |
| GOBP_POSITIVE_REGULATION_OF_RELEASE_OF_CYTOCHROME_C_FROM_MITOCHONDRIA | 26 | -0.475902 | -1.32033 | 0.133603 | 0.281024 |
| GOBP_FEEDING_BEHAVIOR | 52 | -0.418846 | -1.319881 | 0.077263 | 0.2815 |
| GOBP_CARDIAC_CHAMBER_DEVELOPMENT | 107 | -0.365376 | -1.319853 | 0.064665 | 0.281228 |
| GOBP_REGULATION_OF_SYSTEMIC_ARTERIAL_BLOOD_PRESSURE_MEDIATED_BY_A_CHEM | 30 | -0.467266 | -1.319838 | 0.120163 | 0.280926 |
| GOBP_POSITIVE_REGULATION_OF_CELL_CYCLE | 338 | -0.321262 | -1.319336 | 0.01 | 0.281383 |
| GOBP_POSITIVE_REGULATION_OF_NOTCH_SIGNALING_PATHWAY | 48 | -0.418779 | -1.319066 | 0.078723 | 0.281477 |
| GOBP_POSITIVE_REGULATION_OF_VIRAL_PROCESS | 90 | -0.376828 | -1.318879 | 0.058036 | 0.281483 |
| GOBP_REGULATION_OF_CYTOKINESIS | 76 | -0.390163 | -1.318591 | 0.06263 | 0.281637 |
| GOBP_REGULATION_OF_POSTSYNAPTIC_MEMBRANE_NEUROTRANSMITTER_RECEPTOR_L | 46 | -0.424332 | -1.318327 | 0.074689 | 0.281762 |
| GOBP_NUCLEAR_TRANSPORT | 319 | -0.319815 | -1.318258 | 0.011848 | 0.281538 |
| GOBP_REGULATION_OF_REPRODUCTIVE_PROCESS | 115 | -0.364952 | -1.318172 | 0.061785 | 0.281378 |
| GOBP_PROTEIN_LOCALIZATION_TO_MITOCHONDRION | 143 | -0.360092 | -1.317979 | 0.033482 | 0.281388 |
| GOBP_HISTONE_MONOUBIQUITINATION | 28 | -0.458524 | -1.317875 | 0.135011 | 0.281224 |
| GOBP_REGULATION_OF_TRANSMEMBRANE_TRANSPORT | 412 | -0.315714 | -1.317847 | 0.002347 | 0.280954 |
| GOBP_OSTEOBLAST_DEVELOPMENT | 17 | -0.524795 | -1.315951 | 0.125541 | 0.283867 |
| GOBP_CELLULAR_RESPONSE_TO_RETINOIC_ACID | 47 | -0.428199 | -1.315859 | 0.09434 | 0.283671 |
| GOBP_NEGATIVE_REGULATION_OF_T_HELPER_CELL_DIFFERENTIATION | 18 | -0.522268 | -1.315827 | 0.15 | 0.283413 |
| GOBP_LIPID_STORAGE | 69 | -0.39051 | -1.314903 | 0.074074 | 0.284707 |
| GOBP_PROTEIN_MODIFICATION_BY_SMALL_PROTEIN_REMOVAL | 268 | -0.327618 | -1.31486 | 0.01373 | 0.284442 |
| GOBP_SPERM_MOTILITY | 66 | -0.388345 | -1.314625 | 0.057971 | 0.284543 |
| GOBP_SMOOTH_MUSCLE_CELL_PROLIFERATION | 113 | -0.359878 | -1.3146 | 0.041126 | 0.284277 |
| GOBP_REGULATION_OF_SYSTEM_PROCESS | 410 | -0.317134 | -1.313917 | 0.016787 | 0.285141 |
| GOBP_REGULATION_OF_MACROPHAGE_DIFFERENTIATION | 18 | -0.522479 | -1.313891 | 0.138365 | 0.284864 |
| GOBP_NUCLEAR_EXPORT | 190 | -0.344298 | -1.313821 | 0.021978 | 0.28467 |
| GOBP_POSITIVE_REGULATION_OF_NERVOUS_SYSTEM_PROCESS | 27 | -0.463483 | -1.312876 | 0.128692 | 0.286004 |
| GOBP_MRNA_PROCESSING | 497 | -0.308872 | -1.312875 | 0.004739 | 0.285686 |
| GOBP_ERAD_PATHWAY | 97 | -0.371434 | -1.312316 | 0.070213 | 0.286329 |
| GOBP_MODULATION_BY_SYMBIONT_OF_ENTRY_INTO_HOST | 40 | -0.435692 | -1.31194 | 0.107505 | 0.286625 |
| GOBP_REGULATION_OF_ENDOPLASMIC_RETICULUM_STRESS_INDUCED_INTRINSIC_APOPT | 28 | -0.470963 | -1.311221 | 0.140167 | 0.287558 |
| GOBP_REGULATION_OF_DENDRITE_EXTENSION | 19 | -0.500754 | -1.310885 | 0.15678 | 0.287828 |
| GOBP_BONE_MORPHOGENESIS | 69 | -0.396598 | -1.310627 | 0.063008 | 0.287944 |
| GOBP_CELLULAR_RESPONSE_TO_VASCULAR_ENDOTHELIAL_GROWTH_FACTOR_STIMULU | 53 | -0.416186 | -1.310548 | 0.099567 | 0.287766 |
| GOBP_POSITIVE_REGULATION_OF_GROWTH | 200 | -0.341258 | -1.310256 | 0.021231 | 0.287882 |
| GOBP_CELLULAR_RESPONSE_TO_LIPOPROTEIN_PARTICLE_STIMULUS | 26 | -0.480761 | -1.309497 | 0.119588 | 0.288834 |
| GOBP_REGULATION_OF_CELL_PROJECTION_ASSEMBLY | 160 | -0.346975 | -1.309017 | 0.031111 | 0.289394 |
| GOBP_MONOSACCHARIDE_BIOSYNTHETIC_PROCESS | 83 | -0.379111 | -1.308234 | 0.060134 | 0.290422 |
| GOBP_TRANSFERRIN_TRANSPORT | 33 | -0.455015 | -1.30802 | 0.117043 | 0.290487 |
| GOBP_NECROPTOTIC_PROCESS | 36 | -0.445071 | -1.307667 | 0.124444 | 0.290816 |
| GOBP_PROTEIN_N_LINKED_GLYCOSYLATION | 71 | -0.392573 | -1.30706 | 0.084967 | 0.29153 |
| GOBP_REGULATION_OF_NERVOUS_SYSTEM_PROCESS | 104 | -0.360786 | -1.306806 | 0.046875 | 0.29164 |
| GOBP_T_CELL_PROLIFERATION | 176 | -0.338325 | -1.306774 | 0.030093 | 0.291366 |
| GOBP_RHYTHMIC_PROCESS | 238 | -0.328023 | -1.306726 | 0.021951 | 0.291131 |
| GOBP_POSITIVE_REGULATION_OF_ATP_METABOLIC_PROCESS | 33 | -0.443086 | -1.306292 | 0.12 | 0.291528 |
| GOBP_NEGATIVE_REGULATION_OF_TRANSMEMBRANE_RECEPTOR_PROTEIN_SERINE_THR | 109 | -0.360265 | -1.306175 | 0.039648 | 0.291435 |
| GOBP_MUSCLE_CELL_APOPTOTIC_PROCESS | 53 | -0.407563 | -1.305835 | 0.092975 | 0.291685 |
| GOBP_CELLULAR_RESPONSE_TO_INSULIN_STIMULUS | 196 | -0.335706 | -1.305772 | 0.026608 | 0.2915 |
| GOBP_DIVALENT_INORGANIC_CATION_HOMEOSTASIS | 373 | -0.317606 | -1.305557 | 0.007463 | 0.291559 |
| GOBP_EXTERNAL_ENCAPSULATING_STRUCTURE_ORGANIZATION | 293 | -0.319598 | -1.305403 | 0.014851 | 0.29152 |
| GOBP_REGULATION_OF_MRNA_3_END_PROCESSING | 28 | -0.468849 | -1.305374 | 0.120408 | 0.291238 |
| GOBP_ENDOPLASMIC_RETICULUM_UNFOLDED_PROTEIN_RESPONSE | 120 | -0.357157 | -1.304933 | 0.061538 | 0.291684 |
| GOBP_KERATAN_SULFATE_BIOSYNTHETIC_PROCESS | 21 | -0.492766 | -1.304593 | 0.146881 | 0.291996 |
| GOBP_NUCLEAR_ENVELOPE_REASSEMBLY | 18 | -0.527032 | -1.304472 | 0.159041 | 0.291888 |
| GOBP_RESPONSE_TO_HYPEROXIA | 20 | -0.50688 | -1.304398 | 0.131635 | 0.291727 |
| GOBP_DNA_REPLICATION_DEPENDENT_NUCLEOSOME_ORGANIZATION | 25 | -0.479445 | -1.304347 | 0.136646 | 0.291481 |
| GOBP_INFLAMMATORY_CELL_APOPTOTIC_PROCESS | 18 | -0.51868 | -1.304039 | 0.176221 | 0.291709 |
| GOBP_MULTIVESICULAR_BODY_ORGANIZATION | 31 | -0.458766 | -1.304035 | 0.119284 | 0.291396 |
| GOBP_ENDOPLASMIC_RETICULUM_ORGANIZATION | 82 | -0.377993 | -1.303915 | 0.054585 | 0.291299 |
| GOBP_MEMBRANE_DISASSEMBLY | 15 | -0.516068 | -1.303709 | 0.150215 | 0.291318 |
| GOBP_REGULATION_OF_CENTROSOME_CYCLE | 46 | -0.419901 | -1.303579 | 0.115632 | 0.291244 |
| GOBP_TYPE_2_IMMUNE_RESPONSE | 33 | -0.440269 | -1.302985 | 0.115304 | 0.291995 |
| GOBP_ANDROGEN_RECEPTOR_SIGNALING_PATHWAY | 32 | -0.463821 | -1.302976 | 0.118998 | 0.291701 |
| GOBP_POSITIVE_REGULATION_OF_INTERFERON_ALPHA_PRODUCTION | 20 | -0.50315 | -1.302535 | 0.139959 | 0.292173 |
| GOBP_LINOLEIC_ACID_METABOLIC_PROCESS | 17 | -0.521567 | -1.302478 | 0.163599 | 0.291945 |
| GOBP_NCRNA_CATABOLIC_PROCESS | 34 | -0.447615 | -1.302055 | 0.099379 | 0.292395 |

| | | | | | |
|---|---|---|---|---|---|
| GOBP_REGULATION_OF_ESTABLISHMENT_OR_MAINTENANCE_OF_CELL_POLARITY | 24 | 0.209077 | 0.562176 | 0.973734 | 1 |
| GOBP_ASPARTATE_FAMILY_AMINO_ACID_BIOSYNTHETIC_PROCESS | 17 | 0.223814 | 0.560111 | 0.963636 | 1 |
| GOBP_CYTOCHROME_COMPLEX_ASSEMBLY | 35 | 0.19055 | 0.556721 | 0.983178 | 1 |
| GOBP_RRNA_METHYLATION | 28 | 0.200124 | 0.550282 | 0.986717 | 1 |
| GOBP_NUCLEAR_TRANSCRIBED_MRNA_CATABOLIC_PROCESS | 206 | 0.143749 | 0.548645 | 1 | 1 |
| GOBP_RRNA_MODIFICATION | 40 | 0.186179 | 0.54847 | 0.992308 | 1 |
| GOBP_TRICARBOXYLIC_ACID_CYCLE | 33 | 0.193062 | 0.545025 | 0.988506 | 1 |
| GOBP_REGULATION_OF_PROTEIN_LOCALIZATION_TO_CELL_SURFACE | 33 | 0.189295 | 0.537333 | 0.994175 | 1 |
| GOBP_TOR_SIGNALING | 110 | 0.152844 | 0.536807 | 1 | 1 |
| GOBP_HISTONE_H4_ACETYLATION | 66 | 0.167796 | 0.53669 | 0.998024 | 1 |
| GOBP_NEGATIVE_REGULATION_OF_HISTONE_METHYLATION | 20 | 0.205603 | 0.534423 | 0.974903 | 1 |
| GOBP_TRNA_5_END_PROCESSING | 16 | 0.218534 | 0.534335 | 0.980952 | 1 |
| GOBP_PHOSPHATIDYLINOSITOL_PHOSPHORYLATION | 54 | 0.17139 | 0.532383 | 0.998192 | 1 |
| GOBP_MICROTUBULE_ORGANIZING_CENTER_LOCALIZATION | 29 | 0.188205 | 0.527334 | 0.985047 | 1 |
| GOBP_REGULATION_OF_B_CELL_RECEPTOR_SIGNALING_PATHWAY | 25 | 0.196483 | 0.526019 | 0.986381 | 1 |
| GOBP_N_TERMINAL_PROTEIN_AMINO_ACID_ACETYLATION | 16 | 0.21689 | 0.523106 | 0.982284 | 1 |
| GOBP_REGULATION_OF_TRANSCRIPTION_INITIATION_FROM_RNA_POLYMERASE_II_PRO | 31 | 0.188718 | 0.519431 | 0.994536 | 1 |
| GOBP_REGULATION_OF_TOR_SIGNALING | 88 | 0.149659 | 0.50936 | 1 | 1 |
| GOBP_REGULATION_OF_DEFENSE_RESPONSE_TO_VIRUS_BY_VIRUS | 29 | 0.183566 | 0.500781 | 1 | 1 |
| GOBP_GLYCOSIDE_METABOLIC_PROCESS | 16 | 0.205767 | 0.497802 | 0.982692 | 1 |
| GOBP_REGULATION_OF_OXIDATIVE_PHOSPHORYLATION | 20 | 0.194745 | 0.492962 | 0.987928 | 1 |
| GOBP_RESPIRATORY_CHAIN_COMPLEX_IV_ASSEMBLY | 25 | 0.186224 | 0.489897 | 0.988 | 1 |
| GOBP_FATTY_ACYL_COA_METABOLIC_PROCESS | 39 | 0.165487 | 0.483355 | 1 | 1 |
| GOBP_MATURATION_OF_LSU_RRNA | 28 | 0.174426 | 0.477462 | 0.992509 | 1 |
| GOBP_THIOESTER_BIOSYNTHETIC_PROCESS | 50 | 0.152952 | 0.474799 | 1 | 1 |
| GOBP_REGULATION_OF_INTRINSIC_APOPTOTIC_SIGNALING_PATHWAY_IN_RESPONSE_T | 17 | 0.194726 | 0.474775 | 0.990329 | 1 |
| GOBP_RIBOSOMAL_LARGE_SUBUNIT_BIOGENESIS | 72 | 0.143231 | 0.474486 | 1 | 1 |
| GOBP_IRON_SULFUR_CLUSTER_ASSEMBLY | 23 | 0.169195 | 0.448873 | 0.996183 | 1 |
| GOBP_HISTONE_H3_K27_METHYLATION | 16 | 0.181433 | 0.445634 | 0.996132 | 1 |
| GOBP_POSITIVE_REGULATION_OF_DOUBLE_STRAND_BREAK_REPAIR | 39 | 0.146577 | 0.425149 | 1 | 1 |
| GOBP_REGULATION_OF_GOLGI_ORGANIZATION | 17 | 0.171924 | 0.424013 | 0.994106 | 1 |
| GOBP_SCF_DEPENDENT_PROTEASOMAL_UBIQUITIN_DEPENDENT_PROTEIN_CATABOLIC | 87 | 0.121282 | 0.408531 | 1 | 1 |
| GOBP_RNA_METHYLATION | 84 | 0.118231 | 0.401938 | 1 | 1 |
| GOBP_N_TERMINAL_PROTEIN_AMINO_ACID_MODIFICATION | 26 | 0.144874 | 0.394424 | 0.998145 | 1 |
| GOBP_NEGATIVE_REGULATION_OF_PROTEIN_TYROSINE_KINASE_ACTIVITY | 28 | 0.134542 | 0.376841 | 1 | 1 |
| GOBP_TRNA_METABOLIC_PROCESS | 177 | 0.097851 | 0.360631 | 1 | 1 |
| GOBP_PHOSPHATIDYLGLYCEROL_METABOLIC_PROCESS | 28 | 0.109727 | 0.299368 | 1 | 0.999997 |

| | | | | | |
|---|---|---|---|---|---|
| GOBP_LUNG_CELL_DIFFERENTIATION | 18 | -0.505993 | -1.301614 | 0.142857 | 0.292887 |
| GOBP_ASTROCYTE_DIFFERENTIATION | 60 | -0.409384 | -1.301545 | 0.090708 | 0.292699 |
| GOBP_POSITIVE_REGULATION_OF_DNA_TEMPLATED_TRANSCRIPTION_ELONGATION | 25 | -0.478048 | -1.301268 | 0.158537 | 0.292867 |
| GOBP_REGULATION_OF_MICROTUBULE_BASED_PROCESS | 199 | -0.332484 | -1.299442 | 0.019956 | 0.295757 |
| GOBP_ENDOCRINE_SYSTEM_DEVELOPMENT | 73 | -0.387265 | -1.299131 | 0.098434 | 0.295909 |
| GOBP_REGULATION_OF_GLIOGENESIS | 70 | -0.392969 | -1.298916 | 0.08952 | 0.295974 |
| GOBP_GOLGI_VESICLE_TRANSPORT | 350 | -0.315741 | -1.298747 | 0.017632 | 0.295964 |
| GOBP_CILIUM_OR_FLAGELLUM_DEPENDENT_CELL_MOTILITY | 89 | -0.366848 | -1.298656 | 0.067416 | 0.295802 |
| GOBP_REGULATION_OF_LIPID_CATABOLIC_PROCESS | 47 | -0.415964 | -1.298656 | 0.090336 | 0.29549 |
| GOBP_SMALL_MOLECULE_CATABOLIC_PROCESS | 370 | -0.313141 | -1.298457 | 0.013158 | 0.295541 |
| GOBP_TRANSMEMBRANE_RECEPTOR_PROTEIN_SERINE_THREONINE_KINASE_SIGNALING_ | 289 | -0.322711 | -1.297878 | 0.007042 | 0.29621 |
| GOBP_REGULATION_OF_PROTEIN_CONTAINING_COMPLEX_DISASSEMBLY | 106 | -0.368656 | -1.297696 | 0.059081 | 0.296238 |
| GOBP_PROTEIN_LOCALIZATION_TO_NUCLEUS | 250 | -0.324071 | -1.297646 | 0.023923 | 0.296008 |
| GOBP_TRANSITION_METAL_ION_TRANSPORT | 101 | -0.362951 | -1.297486 | 0.056561 | 0.295957 |
| GOBP_REGULATION_OF_PRODUCTION_OF_MOLECULAR_MEDIATOR_OF_IMMUNE_RESPON | 129 | -0.352944 | -1.297336 | 0.066372 | 0.295956 |
| GOBP_PROTEIN_INSERTION_INTO_ER_MEMBRANE | 18 | -0.508658 | -1.296793 | 0.166316 | 0.296619 |
| GOBP_POSITIVE_REGULATION_OF_CELLULAR_AMIDE_METABOLIC_PROCESS | 146 | -0.347024 | -1.296566 | 0.028761 | 0.296725 |
| GOBP_INTRA_GOLGI_VESICLE_MEDIATED_TRANSPORT | 32 | -0.452274 | -1.295991 | 0.11022 | 0.29744 |
| GOBP_PROTEIN_LOCALIZATION_TO_POSTSYNAPSE | 34 | -0.445291 | -1.295962 | 0.124 | 0.297182 |
| GOBP_FC_RECEPTOR_SIGNALING_PATHWAY | 227 | -0.327594 | -1.29574 | 0.023697 | 0.297272 |
| GOBP_PROTEIN_MATURATION | 239 | -0.327941 | -1.295224 | 0.018476 | 0.297894 |
| GOBP_NEGATIVE_REGULATION_OF_HYDROLASE_ACTIVITY | 334 | -0.317289 | -1.294547 | 0.016949 | 0.298728 |
| GOBP_NEGATIVE_REGULATION_OF_ERBB_SIGNALING_PATHWAY | 48 | -0.414539 | -1.294324 | 0.128692 | 0.298811 |
| GOBP_PROTEIN_QUALITY_CONTROL_FOR_MISFOLDED_OR_INCOMPLETELY_SYNTHESIZED | 28 | -0.457293 | -1.294288 | 0.138277 | 0.298568 |
| GOBP_REGULATION_OF_VOLTAGE_GATED_CALCIUM_CHANNEL_ACTIVITY | 25 | -0.477695 | -1.293943 | 0.125275 | 0.298868 |
| GOBP_TERPENOID_METABOLIC_PROCESS | 71 | -0.382688 | -1.293763 | 0.069507 | 0.298877 |
| GOBP_MIDBRAIN_DEVELOPMENT | 69 | -0.386098 | -1.293721 | 0.086316 | 0.298625 |
| GOBP_DNA_DAMAGE_RESPONSE_SIGNAL_TRANSDUCTION_BY_P53_CLASS_MEDIATOR | 103 | -0.364134 | -1.293337 | 0.04888 | 0.299012 |
| GOBP_MORPHOGENESIS_OF_A_BRANCHING_STRUCTURE | 129 | -0.352345 | -1.293266 | 0.063918 | 0.298808 |
| GOBP_REGULATION_OF_HIPPO_SIGNALING | 17 | -0.505296 | -1.293159 | 0.153518 | 0.298669 |
| GOBP_NEUROPEPTIDE_SIGNALING_PATHWAY | 48 | -0.419366 | -1.292781 | 0.120614 | 0.29902 |
| GOBP_REGULATION_OF_BLOOD_PRESSURE | 118 | -0.355138 | -1.291525 | 0.048246 | 0.300951 |
| GOBP_PATTERN_RECOGNITION_RECEPTOR_SIGNALING_PATHWAY | 185 | -0.336572 | -1.291402 | 0.043478 | 0.300895 |
| GOBP_LIVER_MORPHOGENESIS | 16 | -0.51915 | -1.290925 | 0.150628 | 0.301394 |
| GOBP_NAD_BIOSYNTHETIC_PROCESS | 24 | -0.474418 | -1.290884 | 0.129103 | 0.301156 |
| GOBP_COLLAGEN_FIBRIL_ORGANIZATION | 40 | -0.426383 | -1.290819 | 0.117914 | 0.300957 |
| GOBP_DETERMINATION_OF_ADULT_LIFESPAN | 15 | -0.537878 | -1.290761 | 0.187755 | 0.300747 |
| GOBP_PHOSPHATIDYLCHOLINE_ACYL_CHAIN_REMODELING | 21 | -0.498169 | -1.290729 | 0.124736 | 0.300482 |
| GOBP_GLOMERULUS_DEVELOPMENT | 45 | -0.416764 | -1.290672 | 0.136161 | 0.300295 |
| GOBP_POSTREPLICATION_REPAIR | 52 | -0.404225 | -1.290344 | 0.106472 | 0.300563 |
| GOBP_NEGATIVE_REGULATION_OF_CELL_ACTIVATION | 172 | -0.337126 | -1.290039 | 0.040909 | 0.300781 |
| GOBP_SIGNAL_TRANSDUCTION_IN_RESPONSE_TO_DNA_DAMAGE | 126 | -0.353279 | -1.289679 | 0.049356 | 0.301046 |
| GOBP_RESPONSE_TO_INTERFERON_GAMMA | 164 | -0.337086 | -1.289225 | 0.04955 | 0.301562 |
| GOBP_DOSAGE_COMPENSATION | 17 | -0.510598 | -1.288927 | 0.150495 | 0.301763 |
| GOBP_DNA_SYNTHESIS_INVOLVED_IN_DNA_REPAIR | 50 | -0.413363 | -1.2888 | 0.100604 | 0.301643 |
| GOBP_PROTEIN_PHOSPHOPANTETHEINYLATION | 284 | -0.317047 | -1.288565 | 0.009324 | 0.30175 |
| GOBP_POSITIVE_REGULATION_OF_CHROMATIN_ORGANIZATION | 93 | -0.369626 | -1.288486 | 0.08656 | 0.301598 |
| GOBP_REGULATION_OF_CARBOHYDRATE_CATABOLIC_PROCESS | 79 | -0.376106 | -1.288185 | 0.087146 | 0.301802 |
| GOBP_DEVELOPMENTAL_GROWTH_INVOLVED_IN_MORPHOGENESIS | 184 | -0.33493 | -1.28784 | 0.044706 | 0.302056 |
| GOBP_POSITIVE_REGULATION_OF_VIRAL_GENOME_REPLICATION | 31 | -0.453966 | -1.287573 | 0.12743 | 0.302195 |
| GOBP_POSITIVE_REGULATION_OF_GRANULOCYTE_CHEMOTAXIS | 24 | -0.471589 | -1.287511 | 0.141372 | 0.301974 |
| GOBP_WOUND_HEALING | 431 | -0.307448 | -1.286882 | 0.012315 | 0.302858 |
| GOBP_POSITIVE_REGULATION_OF_ORGAN_GROWTH | 40 | -0.417963 | -1.286842 | 0.117137 | 0.302615 |
| GOBP_POSITIVE_REGULATION_OF_HORMONE_SECRETION | 88 | -0.368399 | -1.286664 | 0.089172 | 0.302596 |
| GOBP_MALE_GAMETE_GENERATION | 399 | -0.307724 | -1.286399 | 0.024691 | 0.302781 |
| GOBP_REGULATION_OF_DNA_METABOLIC_PROCESS | 323 | -0.314042 | -1.286383 | 0.013605 | 0.302508 |
| GOBP_POSITIVE_REGULATION_OF_UBIQUITIN_DEPENDENT_PROTEIN_CATABOLIC_PROCES | 97 | -0.36425 | -1.28632 | 0.09396 | 0.30233 |
| GOBP_ONE_CARBON_METABOLIC_PROCESS | 34 | -0.442946 | -1.286276 | 0.142251 | 0.302108 |
| GOBP_MESENCHYME_DEVELOPMENT | 214 | -0.328254 | -1.286109 | 0.02716 | 0.302045 |
| GOBP_RESPONSE_TO_ESTRADIOL | 99 | -0.36292 | -1.285783 | 0.052402 | 0.302333 |
| GOBP_LEUKOCYTE_DIFFERENTIATION | 460 | -0.303639 | -1.284917 | 0.006897 | 0.303457 |
| GOBP_NEGATIVE_REGULATION_OF_ANION_TRANSPORT | 158 | -0.341242 | -1.284889 | 0.040964 | 0.303206 |
| GOBP_REGULATION_OF_CYTOSOLIC_CALCIUM_ION_CONCENTRATION | 256 | -0.320096 | -1.284453 | 0.035047 | 0.303739 |
| GOBP_DOPAMINERGIC_NEURON_DIFFERENTIATION | 21 | -0.485518 | -1.284326 | 0.154176 | 0.303637 |
| GOBP_PROTEIN_EXIT_FROM_ENDOPLASMIC_RETICULUM | 45 | -0.417029 | -1.284032 | 0.096311 | 0.303903 |
| GOBP_NECROTIC_CELL_DEATH | 55 | -0.400351 | -1.283496 | 0.102908 | 0.304556 |
| GOBP_NEGATIVE_REGULATION_OF_NUCLEOTIDE_METABOLIC_PROCESS | 18 | -0.519372 | -1.283163 | 0.169772 | 0.304805 |
| GOBP_COAGULATION | 280 | -0.319191 | -1.282678 | 0.014388 | 0.305379 |
| GOBP_CHEMOKINE_C_X_C_MOTIF_LIGAND_2_PRODUCTION | 16 | -0.524191 | -1.282532 | 0.158004 | 0.305288 |
| GOBP_REGULATION_OF_FAT_CELL_DIFFERENTIATION | 104 | -0.360217 | -1.282301 | 0.0625 | 0.305416 |
| GOBP_NUCLEOSIDE_DIPHOSPHATE_METABOLIC_PROCESS | 140 | -0.345572 | -1.282064 | 0.048565 | 0.305509 |
| GOBP_EMBRYO_DEVELOPMENT_ENDING_IN_BIRTH_OR_EGG_HATCHING | 494 | -0.300223 | -1.281646 | 0.009479 | 0.305928 |
| GOBP_POSITIVE_REGULATION_OF_GLIOGENESIS | 52 | -0.40135 | -1.281143 | 0.107066 | 0.306537 |
| GOBP_MESONEPHRIC_TUBULE_MORPHOGENESIS | 38 | -0.428575 | -1.280824 | 0.119247 | 0.306796 |
| GOBP_DEVELOPMENTAL_GROWTH | 487 | -0.301598 | -1.280764 | 0.007353 | 0.306598 |
| GOBP_REGULATION_OF_PHOSPHATIDYLINOSITOL_3_KINASE_ACTIVITY | 50 | -0.407534 | -1.280245 | 0.108 | 0.307228 |
| GOBP_CELLULAR_BIOGENIC_AMINE_METABOLIC_PROCESS | 76 | -0.372656 | -1.280048 | 0.078829 | 0.307298 |
| GOBP_CELLULAR_RESPONSE_TO_ABIOTIC_STIMULUS | 280 | -0.314774 | -1.279765 | 0.030162 | 0.307517 |
| GOBP_DNA_INTEGRITY_CHECKPOINT | 151 | -0.341714 | -1.279484 | 0.053398 | 0.307702 |
| GOBP_CELLULAR_RESPONSE_TO_GROWTH_HORMONE_STIMULUS | 21 | -0.492878 | -1.279273 | 0.158798 | 0.307785 |
| GOBP_RDNA_HETEROCHROMATIN_ASSEMBLY | 30 | -0.452166 | -1.278671 | 0.135903 | 0.30849 |
| GOBP_PEPTIDYL_TYROSINE_MODIFICATION | 307 | -0.315379 | -1.278412 | 0.014184 | 0.308597 |
| GOBP_DETECTION_OF_MECHANICAL_STIMULUS | 37 | -0.430798 | -1.278362 | 0.141053 | 0.308393 |
| GOBP_REGULATION_OF_COAGULATION | 56 | -0.405742 | -1.277852 | 0.099366 | 0.308999 |
| GOBP_TISSUE_MORPHOGENESIS | 493 | -0.303722 | -1.277835 | 0.017677 | 0.308725 |
| GOBP_REGULATION_OF_SECRETION | 473 | -0.300015 | -1.277057 | 0.010309 | 0.3098 |
| GOBP_REGULATION_OF_GLUCOSE_METABOLIC_PROCESS | 103 | -0.356491 | -1.276873 | 0.064159 | 0.309871 |
| GOBP_FATTY_ACID_BETA_OXIDATION | 71 | -0.379225 | -1.27672 | 0.093617 | 0.309846 |
| GOBP_ESTABLISHMENT_OF_PROTEIN_LOCALIZATION_TO_MITOCHONDRIAL_MEMBRANE | 53 | -0.399882 | -1.275848 | 0.111597 | 0.311167 |
| GOBP_REGULATION_OF_HEMOPOIESIS | 368 | -0.306982 | -1.275775 | 0.014634 | 0.311014 |
| GOBP_RESPONSE_TO_MURAMYL_DIPEPTIDE | 20 | -0.490773 | -1.275553 | 0.173038 | 0.311132 |
| GOBP_REGULATION_OF_SUPEROXIDE_METABOLIC_PROCESS | 25 | -0.462222 | -1.275154 | 0.158333 | 0.31155 |
| GOBP_NEGATIVE_REGULATION_OF_DEPHOSPHORYLATION | 97 | -0.36099 | -1.274342 | 0.080357 | 0.312656 |
| GOBP_RESPIRATORY_BURST | 34 | -0.440669 | -1.274306 | 0.157667 | 0.31242 |
| GOBP_POSITIVE_REGULATION_OF_RECEPTOR_MEDIATED_ENDOCYTOSIS | 40 | -0.424573 | -1.274165 | 0.127083 | 0.312363 |
| GOBP_POSITIVE_REGULATION_OF_OLIGODENDROCYTE_DIFFERENTIATION | 18 | -0.50254 | -1.273666 | 0.151951 | 0.312919 |
| GOBP_3_UTR_MEDIATED_MRNA_DESTABILIZATION | 17 | -0.516328 | -1.273605 | 0.157113 | 0.312727 |
| GOBP_PEPTIDYL_ASPARAGINE_MODIFICATION | 29 | -0.451596 | -1.273314 | 0.143737 | 0.312906 |
| GOBP_POSITIVE_REGULATION_OF_SECRETION | 235 | -0.318747 | -1.273139 | 0.036866 | 0.312916 |
| GOBP_NEURAL_NUCLEUS_DEVELOPMENT | 48 | -0.402251 | -1.272968 | 0.118503 | 0.312953 |
| GOBP_MONOCYTE_DIFFERENTIATION | 32 | -0.446102 | -1.272646 | 0.137712 | 0.313206 |
| GOBP_BIOMINERALIZATION | 118 | -0.351099 | -1.272603 | 0.063966 | 0.312969 |
| GOBP_CD4_POSITIVE_ALPHA_BETA_T_CELL_DIFFERENTIATION | 74 | -0.376794 | -1.272532 | 0.090713 | 0.312804 |
| GOBP_PROTEIN_MANNOSYLATION | 23 | -0.472571 | -1.272265 | 0.153527 | 0.312976 |
| GOBP_FIBROBLAST_APOPTOTIC_PROCESS | 19 | -0.500834 | -1.271862 | 0.177866 | 0.313348 |
| GOBP_PROTEIN_HOMOTETRAMERIZATION | 45 | -0.41252 | -1.271129 | 0.112971 | 0.314352 |
| GOBP_REGULATION_OF_WOUND_HEALING | 98 | -0.360154 | -1.270995 | 0.08114 | 0.314262 |
| GOBP_TYPE_I_INTERFERON_PRODUCTION | 122 | -0.352736 | -1.27098 | 0.049661 | 0.313994 |
| GOBP_RESPONSE_TO_INCREASED_OXYGEN_LEVELS | 27 | -0.458938 | -1.27092 | 0.165577 | 0.313785 |
| GOBP_REGULATION_OF_PHOSPHATASE_ACTIVITY | 157 | -0.338885 | -1.270842 | 0.054176 | 0.31365 |
| GOBP_TELOMERE_MAINTENANCE_VIA_SEMI_CONSERVATIVE_REPLICATION | 27 | -0.461744 | -1.270238 | 0.141935 | 0.314439 |
| GOBP_COLUMNAR_CUBOIDAL_EPITHELIAL_CELL_DIFFERENTIATION | 53 | -0.406288 | -1.270028 | 0.138298 | 0.314586 |
| GOBP_REGULATION_OF_SUPEROXIDE_ANION_GENERATION | 17 | -0.514645 | -1.269858 | 0.164634 | 0.314582 |
| GOBP_BLOOD_VESSEL_MORPHOGENESIS | 486 | -0.299735 | -1.269582 | 0.007075 | 0.314779 |
| GOBP_ENDOCHONDRAL_BONE_MORPHOGENESIS | 44 | -0.413138 | -1.26939 | 0.141935 | 0.314838 |
| GOBP_RESPONSE_TO_GAMMA_RADIATION | 49 | -0.4024 | -1.269288 | 0.118503 | 0.314686 |
| GOBP_RESPONSE_TO_ANTIBIOTIC | 41 | -0.423625 | -1.269254 | 0.135593 | 0.314461 |
| GOBP_INTRACELLULAR_ESTROGEN_RECEPTOR_SIGNALING_PATHWAY | 52 | -0.399371 | -1.269121 | 0.11435 | 0.314394 |
| GOBP_C21_STEROID_HORMONE_METABOLIC_PROCESS | 26 | -0.465691 | -1.26895 | 0.143158 | 0.314389 |
| GOBP_CELLULAR_MODIFIED_AMINO_ACID_BIOSYNTHETIC_PROCESS | 40 | -0.42046 | -1.267963 | 0.148387 | 0.315807 |
| GOBP_REGULATION_OF_CATION_TRANSMEMBRANE_TRANSPORT | 252 | -0.317222 | -1.267766 | 0.035794 | 0.315827 |
| GOBP_REGULATION_OF_MAST_CELL_ACTIVATION | 38 | -0.422497 | -1.2677 | 0.141949 | 0.315648 |
| GOBP_LATE_ENDOSOME_TO_VACUOLE_TRANSPORT | 24 | -0.468271 | -1.267266 | 0.16632 | 0.316154 |
| GOBP_ENERGY_DERIVATION_BY_OXIDATION_OF_ORGANIC_COMPOUNDS | 252 | -0.319273 | -1.266926 | 0.047393 | 0.316439 |
| GOBP_REGULATION_OF_HEART_RATE_BY_CARDIAC_CONDUCTION | 28 | -0.450977 | -1.2666 | 0.140115 | 0.316786 |
| GOBP_TRANSLESION_SYNTHESIS | 42 | -0.414048 | -1.266579 | 0.133183 | 0.316515 |
| GOBP_PROTEIN_TETRAMERIZATION | 68 | -0.378324 | -1.266289 | 0.119748 | 0.316726 |
| GOBP_DETECTION_OF_MECHANICAL_STIMULUS_INVOLVED_IN_SENSORY_PERCEPTION | 23 | -0.476774 | -1.266284 | 0.169043 | 0.316435 |
| GOBP_MALE_SEX_DIFFERENTIATION | 123 | -0.340905 | -1.266028 | 0.058315 | 0.316575 |
| GOBP_T_HELPER_1_CELL_DIFFERENTIATION | 16 | -0.509538 | -1.265662 | 0.172414 | 0.316928 |
| GOBP_MAMMARY_GLAND_DEVELOPMENT | 109 | -0.350062 | -1.26543 | 0.070815 | 0.317102 |
| GOBP_POST_EMBRYONIC_DEVELOPMENT | 67 | -0.374636 | -1.264755 | 0.103093 | 0.318003 |
| GOBP_OVULATION_CYCLE_PROCESS | 33 | -0.44446 | -1.264754 | 0.150905 | 0.317708 |
| GOBP_REGULATION_OF_NUCLEAR_TRANSCRIBED_MRNA_CATABOLIC_PROCESS_DEADEN | 16 | -0.507167 | -1.264681 | 0.189621 | 0.31754 |
| GOBP_REGULATION_OF_DNA_TEMPLATED_TRANSCRIPTION_ELONGATION | 49 | -0.406714 | -1.264673 | 0.116484 | 0.317262 |
| GOBP_SKELETAL_SYSTEM_DEVELOPMENT | 373 | -0.306059 | -1.264324 | 0.00939 | 0.317613 |
| GOBP_TELOMERE_CAPPING | 50 | -0.402628 | -1.264033 | 0.129512 | 0.317804 |
| GOBP_NONRIBOSOMAL_PEPTIDE_BIOSYNTHETIC_PROCESS | 16 | -0.514807 | -1.263998 | 0.184049 | 0.317562 |
| GOBP_REGULATION_OF_POSTSYNAPTIC_MEMBRANE_POTENTIAL | 82 | -0.364717 | -1.263439 | 0.102128 | 0.318288 |
| GOBP_POSITIVE_REGULATION_OF_POTASSIUM_ION_TRANSMEMBRANE_TRANSPORTER_A( | 17 | -0.506346 | -1.262716 | 0.180467 | 0.319256 |
| GOBP_CARBOHYDRATE_HOMEOSTASIS | 171 | -0.330686 | -1.262462 | 0.047085 | 0.319393 |
| GOBP_ENDOCRINE_PROCESS | 60 | -0.387694 | -1.262297 | 0.113208 | 0.319379 |
| GOBP_ARGININE_METABOLIC_PROCESS | 17 | -0.506538 | -1.261737 | 0.155319 | 0.320133 |
| GOBP_REGULATION_OF_CELLULAR_CARBOHYDRATE_METABOLIC_PROCESS | 126 | -0.341379 | -1.260887 | 0.062084 | 0.321316 |
| GOBP_EMBRYONIC_PLACENTA_DEVELOPMENT | 67 | -0.387417 | -1.259868 | 0.11991 | 0.322848 |
| GOBP_NEGATIVE_REGULATION_OF_CELL_MIGRATION_INVOLVED_IN_SPROUTING_ANGIOGE | 15 | -0.50954 | -1.259862 | 0.186992 | 0.322559 |
| GOBP_POSITIVE_REGULATION_OF_SMALL_MOLECULE_METABOLIC_PROCESS | 109 | -0.346942 | -1.259637 | 0.063457 | 0.322688 |
| GOBP_REGULATION_OF_MYOBLAST_FUSION | 16 | -0.505866 | -1.259538 | 0.207819 | 0.322552 |
| GOBP_REGULATION_OF_NEUROTRANSMITTER_LEVELS | 163 | -0.330375 | -1.259529 | 0.05467 | 0.32228 |
| GOBP_EMBRYONIC_FORELIMB_MORPHOGENESIS | 21 | -0.491617 | -1.25938 | 0.160677 | 0.322243 |
| GOBP_REGULATION_OF_B_CELL_APOPTOTIC_PROCESS | 19 | -0.494889 | -1.259342 | 0.188034 | 0.322012 |
| GOBP_DETECTION_OF_STIMULUS_INVOLVED_IN_SENSORY_PERCEPTION_OF_PAIN | 17 | -0.503018 | -1.25881 | 0.190184 | 0.32268 |
| GOBP_POSITIVE_REGULATION_OF_EPITHELIAL_TO_MESENCHYMAL_TRANSITION | 44 | -0.413051 | -1.2586 | 0.137168 | 0.322774 |
| GOBP_INTRINSIC_APOPTOTIC_SIGNALING_PATHWAY_IN_RESPONSE_TO_DNA_DAMAGE_BY | 43 | -0.41157 | -1.258553 | 0.142549 | 0.322561 |
| GOBP_NEGATIVE_REGULATION_OF_POSTTRANSCRIPTIONAL_GENE_SILENCING | 20 | -0.492389 | -1.258488 | 0.185039 | 0.322382 |
| GOBP_MODULATION_OF_EXCITATORY_POSTSYNAPTIC_POTENTIAL | 36 | -0.42694 | -1.258028 | 0.15102 | 0.322958 |
| GOBP_MUSCLE_CELL_PROLIFERATION | 157 | -0.338275 | -1.257976 | 0.068465 | 0.322771 |

| | | | | | |
|---|---|---|---|---|---|
| GOBP_NEGATIVE_REGULATION_OF_CYTOKINE_PRODUCTION | 259 | -0.314403 | -1.257491 | 0.036408 | 0.32331 |
| GOBP_VACUOLE_ORGANIZATION | 163 | -0.333281 | -1.257359 | 0.056769 | 0.323248 |
| GOBP_METANEPHRIC_EPITHELIUM_DEVELOPMENT | 15 | -0.525368 | -1.257118 | 0.201299 | 0.32339 |
| GOBP_REGULATION_OF_CHROMATIN_ORGANIZATION | 169 | -0.329537 | -1.25691 | 0.056818 | 0.323455 |
| GOBP_REGULATION_OF_NUCLEOTIDE_METABOLIC_PROCESS | 102 | -0.358634 | -1.256771 | 0.07947 | 0.323415 |
| GOBP_VASCULAR_ASSOCIATED_SMOOTH_MUSCLE_CELL_DIFFERENTIATION | 25 | -0.462777 | -1.256583 | 0.168724 | 0.323502 |
| GOBP_INTERLEUKIN_2_PRODUCTION | 57 | -0.392387 | -1.256253 | 0.130977 | 0.323836 |
| GOBP_HEART_GROWTH | 73 | -0.374583 | -1.256127 | 0.098468 | 0.323739 |
| GOBP_VIRION_ASSEMBLY | 40 | -0.42314 | -1.255999 | 0.136771 | 0.323687 |
| GOBP_REGULATION_OF_PROTEIN_BINDING | 175 | -0.32505 | -1.255618 | 0.053215 | 0.324083 |
| GOBP_VIRAL_BUDDING | 26 | -0.458687 | -1.255555 | 0.164063 | 0.3239 |
| GOBP_REGULATION_OF_PROTEIN_TARGETING_TO_MEMBRANE | 31 | -0.44358 | -1.255392 | 0.161504 | 0.323916 |
| GOBP_CELL_ADHESION_MEDIATED_BY_INTEGRIN | 63 | -0.372899 | -1.255225 | 0.095541 | 0.323944 |
| GOBP_POSITIVE_REGULATION_OF_DEFENSE_RESPONSE_TO_VIRUS_BY_HOST | 29 | -0.442176 | -1.254914 | 0.171429 | 0.324206 |
| GOBP_SERINE_FAMILY_AMINO_ACID_BIOSYNTHETIC_PROCESS | 18 | -0.497194 | -1.254879 | 0.190669 | 0.323987 |
| GOBP_LIPID_EXPORT_FROM_CELL | 63 | -0.380943 | -1.254803 | 0.114224 | 0.32384 |
| GOBP_BONE_MATURATION | 18 | -0.494176 | -1.254794 | 0.189899 | 0.323569 |
| GOBP_REGULATION_OF_POTASSIUM_ION_TRANSPORT | 69 | -0.371071 | -1.254721 | 0.11828 | 0.323401 |
| GOBP_POSITIVE_REGULATION_OF_DNA_BIOSYNTHETIC_PROCESS | 62 | -0.38206 | -1.254316 | 0.093817 | 0.32381 |
| GOBP_AROMATIC_AMINO_ACID_FAMILY_METABOLIC_PROCESS | 24 | -0.46564 | -1.25276 | 0.182844 | 0.326273 |
| GOBP_RNA_SURVEILLANCE | 15 | -0.521996 | -1.252062 | 0.180723 | 0.32725 |
| GOBP_CENTROSOME_DUPLICATION | 68 | -0.375823 | -1.252044 | 0.110647 | 0.326985 |
| GOBP_REGULATION_OF_RESPONSE_TO_WOUNDING | 126 | -0.341336 | -1.251918 | 0.066098 | 0.326946 |
| GOBP_CERAMIDE_BIOSYNTHETIC_PROCESS | 64 | -0.382549 | -1.251884 | 0.122622 | 0.326713 |
| GOBP_POTASSIUM_ION_TRANSPORT | 147 | -0.331974 | -1.251828 | 0.05 | 0.326522 |
| GOBP_CD4_POSITIVE_ALPHA_BETA_T_CELL_ACTIVATION | 89 | -0.355232 | -1.250689 | 0.110329 | 0.328267 |
| GOBP_REGULATION_OF_T_CELL_MEDIATED_IMMUNITY | 71 | -0.372334 | -1.250561 | 0.109705 | 0.328208 |
| GOBP_CHROMATIN_SILENCING | 63 | -0.385333 | -1.250514 | 0.121593 | 0.327989 |
| GOBP_NEGATIVE_REGULATION_OF_MRNA_SPLICING_VIA_SPLICEOSOME | 19 | -0.485503 | -1.250499 | 0.168467 | 0.327728 |
| GOBP_NEGATIVE_REGULATION_OF_PROTEOLYSIS | 241 | -0.314046 | -1.25041 | 0.047009 | 0.327627 |
| GOBP_MONOSACCHARIDE_CATABOLIC_PROCESS | 53 | -0.388686 | -1.249686 | 0.135076 | 0.328721 |
| GOBP_NEGATIVE_REGULATION_OF_PEPTIDYL_THREONINE_PHOSPHORYLATION | 19 | -0.494789 | -1.249392 | 0.184265 | 0.328975 |
| GOBP_EXIT_FROM_MITOSIS | 26 | -0.450526 | -1.249321 | 0.171779 | 0.328807 |
| GOBP_CIRCADIAN_RHYTHM | 170 | -0.3257 | -1.24924 | 0.054705 | 0.328669 |
| GOBP_NEGATIVE_REGULATION_OF_LYMPHOCYTE_DIFFERENTIATION | 48 | -0.398829 | -1.24917 | 0.14916 | 0.328515 |
| GOBP_NEUROTRANSMITTER_SECRETION | 122 | -0.341643 | -1.248994 | 0.083333 | 0.328557 |
| GOBP_POSITIVE_REGULATION_OF_CATION_TRANSMEMBRANE_TRANSPORT | 108 | -0.348782 | -1.248891 | 0.076037 | 0.328487 |
| GOBP_DEVELOPMENTAL_INDUCTION | 18 | -0.507968 | -1.248856 | 0.19 | 0.328266 |
| GOBP_SUPEROXIDE_ANION_GENERATION | 28 | -0.450173 | -1.248581 | 0.173469 | 0.328455 |
| GOBP_NEGATIVE_REGULATION_OF_LIPID_CATABOLIC_PROCESS | 20 | -0.479716 | -1.248449 | 0.175197 | 0.328422 |
| GOBP_GLIAL_CELL_DIFFERENTIATION | 171 | -0.324094 | -1.248273 | 0.063319 | 0.328429 |
| GOBP_PROTEIN_LOCALIZATION_TO_KINETOCHORE | 19 | -0.489686 | -1.248015 | 0.194332 | 0.328598 |
| GOBP_REGULATION_OF_MEMBRANE_INVAGINATION | 16 | -0.506594 | -1.248001 | 0.194503 | 0.32834 |
| GOBP_DEOXYRIBONUCLEOSIDE_MONOPHOSPHATE_METABOLIC_PROCESS | 15 | -0.517537 | -1.247948 | 0.197938 | 0.328152 |
| GOBP_NEGATIVE_REGULATION_OF_SIGNALING_RECEPTOR_ACTIVITY | 41 | -0.415085 | -1.247803 | 0.146608 | 0.328111 |
| GOBP_LUNG_EPITHELIUM_DEVELOPMENT | 28 | -0.446254 | -1.247487 | 0.185501 | 0.328401 |
| GOBP_REGULATION_OF_T_CELL_ACTIVATION | 298 | -0.305366 | -1.247217 | 0.030691 | 0.328559 |
| GOBP_PROTEIN_LOCALIZATION_TO_CHROMOSOME_CENTROMERIC_REGION | 25 | -0.459127 | -1.24652 | 0.161885 | 0.329525 |
| GOBP_ADRENERGIC_RECEPTOR_SIGNALING_PATHWAY | 17 | -0.507424 | -1.246422 | 0.190476 | 0.329415 |
| GOBP_AROMATIC_AMINO_ACID_FAMILY_CATABOLIC_PROCESS | 16 | -0.508069 | -1.246374 | 0.181275 | 0.329234 |
| GOBP_NEGATIVE_REGULATION_OF_MAPK_CASCADE | 148 | -0.33166 | -1.246371 | 0.0625 | 0.328953 |
| GOBP_NIK_NF_KAPPAB_SIGNALING | 159 | -0.328106 | -1.24624 | 0.083146 | 0.328946 |
| GOBP_RNA_EXPORT_FROM_NUCLEUS | 132 | -0.338174 | -1.246157 | 0.061002 | 0.328822 |
| GOBP_ORGAN_GROWTH | 126 | -0.341789 | -1.246086 | 0.071594 | 0.328666 |
| GOBP_DENDRITIC_SPINE_MAINTENANCE | 17 | -0.499088 | -1.246072 | 0.198758 | 0.328409 |
| GOBP_REGULATION_OF_TRANSPORTER_ACTIVITY | 207 | -0.319783 | -1.245651 | 0.048837 | 0.328929 |
| GOBP_IRON_ION_TRANSPORT | 66 | -0.380133 | -1.245385 | 0.12035 | 0.329146 |
| GOBP_ESTABLISHMENT_OF_MITOTIC_SPINDLE_ORIENTATION | 27 | -0.449487 | -1.245275 | 0.175299 | 0.32907 |
| GOBP_METAL_ION_TRANSPORT | 380 | -0.297663 | -1.244843 | 0.021378 | 0.329533 |
| GOBP_PYRIMIDINE_NUCLEOSIDE_BIOSYNTHETIC_PROCESS | 16 | -0.497304 | -1.244736 | 0.187364 | 0.329418 |
| GOBP_REGULATION_OF_HISTONE_MODIFICATION | 129 | -0.339336 | -1.244608 | 0.073826 | 0.329379 |
| GOBP_REGULATION_OF_MITOCHONDRION_ORGANIZATION | 146 | -0.336377 | -1.244338 | 0.041475 | 0.329593 |
| GOBP_DNA_METHYLATION | 58 | -0.386365 | -1.243898 | 0.142562 | 0.330071 |
| GOBP_RESPIRATORY_SYSTEM_DEVELOPMENT | 148 | -0.329579 | -1.243359 | 0.062907 | 0.330725 |
| GOBP_MESENCHYMAL_CELL_DIFFERENTIATION | 176 | -0.323756 | -1.24328 | 0.065315 | 0.33057 |
| GOBP_NEGATIVE_REGULATION_OF_CD4_POSITIVE_ALPHA_BETA_T_CELL_DIFFERENTIATIO | 22 | -0.47814 | -1.243175 | 0.1875 | 0.330473 |
| GOBP_POSITIVE_REGULATION_OF_ANIMAL_ORGAN_MORPHOGENESIS | 18 | -0.489986 | -1.243128 | 0.188017 | 0.330281 |
| GOBP_CONNECTIVE_TISSUE_DEVELOPMENT | 192 | -0.318631 | -1.242805 | 0.056893 | 0.330609 |
| GOBP_FATTY_ACID_DERIVATIVE_METABOLIC_PROCESS | 68 | -0.368923 | -1.242586 | 0.135417 | 0.33073 |
| GOBP_NEGATIVE_REGULATION_OF_ENDOPLASMIC_RETICULUM_STRESS_INDUCED_INTRIN | 16 | -0.494546 | -1.242316 | 0.187373 | 0.331014 |
| GOBP_CYTOKINE_PRODUCTION_INVOLVED_IN_IMMUNE_RESPONSE | 84 | -0.363153 | -1.242278 | 0.116228 | 0.330794 |
| GOBP_PRODUCTION_OF_MOLECULAR_MEDIATOR_INVOLVED_IN_INFLAMMATORY_RESPON | 66 | -0.378015 | -1.241879 | 0.14375 | 0.331279 |
| GOBP_CERAMIDE_METABOLIC_PROCESS | 95 | -0.347927 | -1.241724 | 0.109375 | 0.331305 |
| GOBP_REGULATION_OF_BROWN_FAT_CELL_DIFFERENTIATION | 17 | -0.499127 | -1.241199 | 0.232409 | 0.331971 |
| GOBP_NEGATIVE_REGULATION_OF_CALCIUM_MEDIATED_SIGNALING | 16 | -0.500975 | -1.24103 | 0.191268 | 0.331999 |
| GOBP_MACROPHAGE_MIGRATION | 46 | -0.393484 | -1.240866 | 0.133612 | 0.33198 |
| GOBP_CHROMATIN_ASSEMBLY_OR_DISASSEMBLY | 187 | -0.318161 | -1.2406 | 0.070175 | 0.332189 |
| GOBP_CELL_CYCLE_G2_M_PHASE_TRANSITION | 263 | -0.30839 | -1.240379 | 0.05531 | 0.332283 |
| GOBP_POSITIVE_REGULATION_OF_T_HELPER_CELL_DIFFERENTIATION | 19 | -0.487066 | -1.23993 | 0.189621 | 0.332825 |
| GOBP_3_PHOSPHOADENOSINE_5_PHOSPHOSULFATE_METABOLIC_PROCESS | 19 | -0.494115 | -1.23967 | 0.17234 | 0.333023 |
| GOBP_SIGNALING_RECEPTOR_LIGAND_PRECURSOR_PROCESSING | 21 | -0.481087 | -1.239657 | 0.193617 | 0.332762 |
| GOBP_JNK_CASCADE | 170 | -0.324731 | -1.238983 | 0.061927 | 0.333677 |
| GOBP_REGENERATION | 159 | -0.332601 | -1.238596 | 0.076577 | 0.334072 |
| GOBP_PROTEIN_STABILIZATION | 175 | -0.327209 | -1.238483 | 0.048352 | 0.334029 |
| GOBP_NEGATIVE_REGULATION_OF_INTRACELLULAR_PROTEIN_TRANSPORT | 39 | -0.413038 | -1.238175 | 0.162791 | 0.334347 |
| GOBP_REGULATION_OF_SIGNALING_RECEPTOR_ACTIVITY | 136 | -0.340755 | -1.237378 | 0.062069 | 0.335472 |
| GOBP_ODONTOGENESIS_OF_DENTIN_CONTAINING_TOOTH | 62 | -0.373102 | -1.237332 | 0.135193 | 0.335247 |
| GOBP_REGULATION_OF_CELLULAR_KETONE_METABOLIC_PROCESS | 150 | -0.329302 | -1.237047 | 0.068584 | 0.335481 |
| GOBP_POSITIVE_REGULATION_OF_CALCIUM_ION_TRANSPORT | 89 | -0.349515 | -1.236385 | 0.109865 | 0.336366 |
| GOBP_ADENYLATE_CYCLASE_MODULATING_G_PROTEIN_COUPLED_RECEPTOR_SIGNALIN | 147 | -0.330445 | -1.235943 | 0.082969 | 0.336956 |
| GOBP_POSITIVE_REGULATION_OF_MUSCLE_TISSUE_DEVELOPMENT | 15 | -0.501325 | -1.235513 | 0.201613 | 0.337396 |
| GOBP_POSITIVE_REGULATION_OF_TRANSLATION | 119 | -0.343293 | -1.235195 | 0.100629 | 0.337702 |
| GOBP_REGULATION_OF_INTRACELLULAR_PROTEIN_TRANSPORT | 230 | -0.309757 | -1.234652 | 0.027523 | 0.338477 |
| GOBP_POSITIVE_REGULATION_OF_TYROSINE_PHOSPHORYLATION_OF_STAT_PROTEIN | 52 | -0.390597 | -1.233758 | 0.15121 | 0.339903 |
| GOBP_REGULATION_OF_MICROTUBULE_POLYMERIZATION | 45 | -0.402195 | -1.233365 | 0.168182 | 0.340326 |
| GOBP_REGULATION_OF_FEEDING_BEHAVIOR | 16 | -0.501276 | -1.233219 | 0.216049 | 0.340311 |
| GOBP_ERBB2_SIGNALING_PATHWAY | 27 | -0.444153 | -1.233109 | 0.1917 | 0.340222 |
| GOBP_POSITIVE_REGULATION_OF_PEPTIDYL_TYROSINE_PHOSPHORYLATION | 153 | -0.324304 | -1.233076 | 0.079727 | 0.339999 |
| GOBP_PHOSPHATIDYLSERINE_METABOLIC_PROCESS | 27 | -0.449587 | -1.232654 | 0.194323 | 0.340502 |
| GOBP_NEGATIVE_REGULATION_OF_ENDOCYTOSIS | 41 | -0.408099 | -1.232353 | 0.158333 | 0.340778 |
| GOBP_REGULATION_OF_MEMBRANE_PERMEABILITY | 77 | -0.365095 | -1.232169 | 0.144086 | 0.340853 |
| GOBP_AMINO_ACID_TRANSPORT | 114 | -0.342292 | -1.232082 | 0.097917 | 0.340723 |
| GOBP_MULTICELLULAR_ORGANISM_AGING | 24 | -0.462504 | -1.231625 | 0.204969 | 0.341267 |
| GOBP_POSITIVE_REGULATION_OF_VASCULAR_ASSOCIATED_SMOOTH_MUSCLE_CELL_PR | 32 | -0.429079 | -1.231569 | 0.166667 | 0.341077 |
| GOBP_BONE_MINERALIZATION | 87 | -0.351726 | -1.231481 | 0.120253 | 0.340944 |
| GOBP_MACROPHAGE_DIFFERENTIATION | 40 | -0.406653 | -1.231435 | 0.176972 | 0.340739 |
| GOBP_GLUCOSE_CATABOLIC_PROCESS | 30 | -0.429382 | -1.231283 | 0.180498 | 0.340708 |
| GOBP_REGULATION_OF_NIK_NF_KAPPAB_SIGNALING | 85 | -0.357277 | -1.231254 | 0.117137 | 0.340467 |
| GOBP_BILE_ACID_BIOSYNTHETIC_PROCESS | 25 | -0.450899 | -1.23087 | 0.194274 | 0.340943 |
| GOBP_ANIMAL_ORGAN_FORMATION | 38 | -0.413978 | -1.230698 | 0.172131 | 0.340987 |
| GOBP_REGULATION_OF_DNA_METHYLATION | 18 | -0.484795 | -1.23064 | 0.210526 | 0.340841 |
| GOBP_PURINE_CONTAINING_COMPOUND_SALVAGE | 15 | -0.518215 | -1.230634 | 0.210309 | 0.340574 |
| GOBP_POSITIVE_REGULATION_OF_MUSCLE_HYPERTROPHY | 23 | -0.462637 | -1.230573 | 0.187368 | 0.340404 |
| GOBP_PYRIDINE_CONTAINING_COMPOUND_METABOLIC_PROCESS | 36 | -0.423896 | -1.229578 | 0.148472 | 0.341955 |
| GOBP_POSITIVE_REGULATION_OF_RESPONSE_TO_CYTOKINE_STIMULUS | 54 | -0.385055 | -1.229272 | 0.145089 | 0.342314 |
| GOBP_INTRACELLULAR_RECEPTOR_SIGNALING_PATHWAY | 236 | -0.312339 | -1.228734 | 0.046154 | 0.343029 |
| GOBP_VESICLE_CYTOSKELETAL_TRAFFICKING | 67 | -0.373841 | -1.228484 | 0.164179 | 0.343189 |
| GOBP_REGULATION_OF_DNA_BIOSYNTHETIC_PROCESS | 99 | -0.343727 | -1.228423 | 0.101322 | 0.343027 |
| GOBP_NEGATIVE_REGULATION_OF_INFLAMMATORY_RESPONSE | 109 | -0.347777 | -1.228267 | 0.105023 | 0.343005 |
| GOBP_DNA_MODIFICATION | 101 | -0.350309 | -1.228169 | 0.104478 | 0.342898 |
| GOBP_POSITIVE_REGULATION_OF_PATTERN_RECOGNITION_RECEPTOR_SIGNALING_PATH | 39 | -0.417022 | -1.228151 | 0.173034 | 0.342658 |
| GOBP_NEGATIVE_REGULATION_OF_G_PROTEIN_COUPLED_RECEPTOR_SIGNALING_PATHW | 37 | -0.421299 | -1.227995 | 0.17119 | 0.342688 |
| GOBP_PEPTIDYL_TYROSINE_DEPHOSPHORYLATION | 88 | -0.357961 | -1.227701 | 0.123043 | 0.342943 |
| GOBP_REGULATION_OF_PROTEIN_LOCALIZATION_TO_SYNAPSE | 25 | -0.459025 | -1.227558 | 0.16309 | 0.342926 |
| GOBP_NEGATIVE_REGULATION_OF_NEURON_DIFFERENTIATION | 40 | -0.406395 | -1.226875 | 0.170259 | 0.343895 |
| GOBP_TRIGLYCERIDE_CATABOLIC_PROCESS | 20 | -0.473376 | -1.226781 | 0.175476 | 0.343786 |
| GOBP_REGULATION_OF_MICROTUBULE_DEPOLYMERIZATION | 24 | -0.450852 | -1.226777 | 0.179266 | 0.343512 |
| GOBP_NEGATIVE_REGULATION_OF_ANION_TRANSMEMBRANE_TRANSPORT | 27 | -0.441783 | -1.226755 | 0.18259 | 0.343268 |
| GOBP_REGULATION_OF_MACROPHAGE_ACTIVATION | 48 | -0.399212 | -1.226738 | 0.162637 | 0.343009 |
| GOBP_REGULATION_OF_GLIAL_CELL_PROLIFERATION | 25 | -0.460749 | -1.226571 | 0.177551 | 0.343037 |
| GOBP_REGULATION_OF_GLIAL_CELL_DIFFERENTIATION | 50 | -0.389267 | -1.226349 | 0.137421 | 0.343216 |
| GOBP_T_CELL_ACTIVATION_INVOLVED_IN_IMMUNE_RESPONSE | 89 | -0.347586 | -1.2263 | 0.119097 | 0.343016 |
| GOBP_VENTRICULAR_CARDIAC_MUSCLE_CELL_ACTION_POTENTIAL | 24 | -0.458061 | -1.225745 | 0.18107 | 0.343818 |
| GOBP_B_CELL_ACTIVATION_INVOLVED_IN_IMMUNE_RESPONSE | 78 | -0.361641 | -1.225402 | 0.153518 | 0.344098 |
| GOBP_REGULATION_OF_TRANSCRIPTION_REGULATORY_REGION_DNA_BINDING | 41 | -0.404071 | -1.225264 | 0.175214 | 0.344057 |
| GOBP_INTRINSIC_APOPTOTIC_SIGNALING_PATHWAY_BY_P53_CLASS_MEDIATOR | 70 | -0.367695 | -1.225064 | 0.122622 | 0.344151 |
| GOBP_NEGATIVE_REGULATION_OF_B_CELL_ACTIVATION | 32 | -0.423261 | -1.225013 | 0.2 | 0.34397 |
| GOBP_NEGATIVE_REGULATION_OF_AUTOPHAGY | 75 | -0.360616 | -1.22494 | 0.133603 | 0.34383 |
| GOBP_NEGATIVE_REGULATION_OF_LYMPHOCYTE_ACTIVATION | 137 | -0.331006 | -1.224178 | 0.086316 | 0.344972 |
| GOBP_EMBRYONIC_HINDLIMB_MORPHOGENESIS | 17 | -0.496749 | -1.224112 | 0.224066 | 0.344796 |
| GOBP_MYELOID_DENDRITIC_CELL_DIFFERENTIATION | 18 | -0.482547 | -1.223819 | 0.231884 | 0.345081 |
| GOBP_REGULATION_OF_DEVELOPMENTAL_GROWTH | 247 | -0.308135 | -1.22323 | 0.067757 | 0.346023 |
| GOBP_POSITIVE_REGULATION_OF_GLIAL_CELL_PROLIFERATION | 16 | -0.496772 | -1.22287 | 0.209544 | 0.34639 |
| GOBP_VIRAL_RELEASE_FROM_HOST_CELL | 31 | -0.421195 | -1.222406 | 0.187225 | 0.346988 |
| GOBP_ESTABLISHMENT_OF_SPINDLE_ORIENTATION | 31 | -0.430416 | -1.222208 | 0.190381 | 0.347078 |
| GOBP_CELL_REDOX_HOMEOSTASIS | 41 | -0.40786 | -1.222101 | 0.170732 | 0.347008 |
| GOBP_DNA_PACKAGING | 197 | -0.31614 | -1.222004 | 0.080831 | 0.346918 |
| GOBP_INTEGRIN_MEDIATED_SIGNALING_PATHWAY | 96 | -0.344553 | -1.221618 | 0.122768 | 0.347339 |
| GOBP_REGULATION_OF_ORGAN_GROWTH | 78 | -0.360926 | -1.221447 | 0.131183 | 0.347374 |
| GOBP_REGULATION_OF_AUTOPHAGY | 309 | -0.298039 | -1.221397 | 0.04401 | 0.347193 |
| GOBP_PROTEIN_INSERTION_INTO_MEMBRANE | 72 | -0.362858 | -1.221252 | 0.150628 | 0.347193 |
| GOBP_ORGANIC_CATION_TRANSPORT | 27 | -0.441137 | -1.221222 | 0.207392 | 0.346992 |
| GOBP_AMINO_SUGAR_METABOLIC_PROCESS | 37 | -0.413812 | -1.221189 | 0.177419 | 0.346779 |
| GOBP_ACID_SECRETION | 27 | -0.439104 | -1.221064 | 0.223629 | 0.346731 |
| GOBP_RNA_SPLICING | 440 | -0.289002 | -1.220791 | 0.028369 | 0.346965 |
| GOBP_VIRAL_BUDDING_VIA_HOST_ESCRT_COMPLEX | 21 | -0.458982 | -1.220338 | 0.190476 | 0.347518 |
| GOBP_POSITIVE_REGULATION_OF_PROTEOLYSIS | 329 | -0.295335 | -1.220286 | 0.034803 | 0.347332 |

| | | | | | |
|---|---|---|---|---|---|
| GOBP_NEGATIVE_REGULATION_OF_MRNA_PROCESSING | 28 | -0.445893 | -1.220163 | 0.193613 | 0.347301 |
| GOBP_POSITIVE_REGULATION_OF_TRANSCRIPTION_REGULATORY_REGION_DNA_BINDING | 20 | -0.475313 | -1.219843 | 0.200828 | 0.347603 |
| GOBP_WATER_SOLUBLE_VITAMIN_METABOLIC_PROCESS | 75 | -0.364452 | -1.219814 | 0.151899 | 0.347381 |
| GOBP_RESPONSE_TO_MISFOLDED_PROTEIN | 24 | -0.455998 | -1.219786 | 0.216102 | 0.34716 |
| GOBP_CELL_DIFFERENTIATION_INVOLVED_IN_EMBRYONIC_PLACENTA_DEVELOPMENT | 19 | -0.479992 | -1.219341 | 0.223214 | 0.347715 |
| GOBP_CELLULAR_RESPONSE_TO_UV | 80 | -0.358604 | -1.218933 | 0.127572 | 0.348162 |
| GOBP_RNA_SPLICING_VIA_TRANSESTERIFICATION_REACTIONS | 352 | -0.29574 | -1.218736 | 0.036446 | 0.348219 |
| GOBP_CENTRIOLE_ASSEMBLY | 41 | -0.40055 | -1.218385 | 0.207739 | 0.348632 |
| GOBP_NEGATIVE_REGULATION_OF_RNA_SPLICING | 24 | -0.462653 | -1.21828 | 0.212185 | 0.348542 |
| GOBP_PROTEIN_KINASE_C_ACTIVATING_G_PROTEIN_COUPLED_RECEPTOR_SIGNALING_P | 19 | -0.481191 | -1.218218 | 0.216842 | 0.348368 |
| GOBP_MORPHOGENESIS_OF_AN_EPITHELIUM | 424 | -0.29109 | -1.217986 | 0.039906 | 0.348514 |
| GOBP_ACTIVATION_OF_PROTEIN_KINASE_A_ACTIVITY | 15 | -0.505411 | -1.217908 | 0.226782 | 0.348369 |
| GOBP_NON_CANONICAL_WNT_SIGNALING_PATHWAY | 138 | -0.331497 | -1.217371 | 0.103371 | 0.349059 |
| GOBP_CELLULAR_RESPONSE_TO_IONIZING_RADIATION | 61 | -0.375092 | -1.21688 | 0.135965 | 0.349674 |
| GOBP_PROTEIN_LOCALIZATION_TO_MICROTUBULE_ORGANIZING_CENTER | 35 | -0.424713 | -1.216764 | 0.170259 | 0.349583 |
| GOBP_MAINTENANCE_OF_LOCATION | 266 | -0.304578 | -1.216466 | 0.048611 | 0.349876 |
| GOBP_POSITIVE_REGULATION_OF_CELLULAR_PROTEIN_CATABOLIC_PROCESS | 139 | -0.329698 | -1.215313 | 0.093817 | 0.351881 |
| GOBP_REGULATION_OF_SPINDLE_ORGANIZATION | 39 | -0.403684 | -1.215199 | 0.180467 | 0.351842 |
| GOBP_REGULATION_OF_PROTEIN_LOCALIZATION_TO_MEMBRANE | 167 | -0.319216 | -1.214992 | 0.079832 | 0.351955 |
| GOBP_POSITIVE_REGULATION_OF_OXIDOREDUCTASE_ACTIVITY | 47 | -0.388958 | -1.214193 | 0.169892 | 0.353166 |
| GOBP_HISTONE_PHOSPHORYLATION | 38 | -0.412006 | -1.214187 | 0.190265 | 0.352906 |
| GOBP_DEOXYRIBONUCLEOTIDE_METABOLIC_PROCESS | 38 | -0.411013 | -1.214146 | 0.164557 | 0.352704 |
| GOBP_NEGATIVE_REGULATION_OF_STRIATED_MUSCLE_CELL_DIFFERENTIATION | 29 | -0.425762 | -1.21387 | 0.206478 | 0.352893 |
| GOBP_POSITIVE_REGULATION_OF_MEMBRANE_PROTEIN_ECTODOMAIN_PROTEOLYSIS | 15 | -0.497456 | -1.213234 | 0.207317 | 0.353798 |
| GOBP_POSITIVE_REGULATION_OF_CHROMOSOME_SEPARATION | 17 | -0.483978 | -1.21303 | 0.210317 | 0.353944 |
| GOBP_NAD_METABOLIC_PROCESS | 45 | -0.391221 | -1.21302 | 0.168444 | 0.353685 |
| GOBP_SEQUESTERING_OF_CALCIUM_ION | 96 | -0.348578 | -1.212126 | 0.108108 | 0.355088 |
| GOBP_NEGATIVE_REGULATION_OF_SINGLE_STRANDED_VIRAL_RNA_REPLICATION_VIA_DO | 16 | -0.493986 | -1.212094 | 0.229958 | 0.354873 |
| GOBP_MUSCLE_STRUCTURE_DEVELOPMENT | 466 | -0.286361 | -1.211945 | 0.026005 | 0.354869 |
| GOBP_MODULATION_BY_HOST_OF_VIRAL_PROCESS | 26 | -0.444375 | -1.211846 | 0.213693 | 0.354779 |
| GOBP_T_CELL_ACTIVATION | 423 | -0.291133 | -1.211815 | 0.043478 | 0.35456 |
| GOBP_TOOTH_MINERALIZATION | 18 | -0.472637 | -1.21165 | 0.219409 | 0.354615 |
| GOBP_REGULATION_OF_SYSTEMIC_ARTERIAL_BLOOD_PRESSURE | 56 | -0.382457 | -1.211414 | 0.159596 | 0.354788 |
| GOBP_OSSIFICATION | 314 | -0.296379 | -1.211386 | 0.061364 | 0.354576 |
| GOBP_PYRIMIDINE_CONTAINING_COMPOUND_METABOLIC_PROCESS | 78 | -0.354063 | -1.211227 | 0.148936 | 0.354596 |
| GOBP_PHOSPHOLIPASE_C_ACTIVATING_G_PROTEIN_COUPLED_RECEPTOR_SIGNALING_PA | 62 | -0.367859 | -1.21108 | 0.158014 | 0.3546 |
| GOBP_PROTEIN_MONOUBIQUITINATION | 64 | -0.367517 | -1.210198 | 0.145336 | 0.356026 |
| GOBP_CARDIOCYTE_DIFFERENTIATION | 96 | -0.341027 | -1.21004 | 0.125 | 0.356051 |
| GOBP_REGULATION_OF_PROTEIN_STABILITY | 273 | -0.303094 | -1.209342 | 0.058005 | 0.3571 |
| GOBP_NEGATIVE_REGULATION_OF_COLD_INDUCED_THERMOGENESIS | 38 | -0.398086 | -1.20907 | 0.179283 | 0.357308 |
| GOBP_NUCLEAR_ENVELOPE_ORGANIZATION | 51 | -0.388763 | -1.208967 | 0.152361 | 0.357209 |
| GOBP_POSITIVE_REGULATION_OF_REPRODUCTIVE_PROCESS | 59 | -0.376406 | -1.208598 | 0.169231 | 0.357663 |
| GOBP_MEIOTIC_CELL_CYCLE | 211 | -0.311122 | -1.208407 | 0.067916 | 0.357705 |
| GOBP_ATTACHMENT_OF_MITOTIC_SPINDLE_MICROTUBULES_TO_KINETOCHORE | 15 | -0.507219 | -1.208394 | 0.253192 | 0.357447 |
| GOBP_PLACENTA_DEVELOPMENT | 108 | -0.339118 | -1.208278 | 0.105495 | 0.357391 |
| GOBP_TRIGLYCERIDE_METABOLIC_PROCESS | 72 | -0.362105 | -1.208237 | 0.159274 | 0.357174 |
| GOBP_POSITIVE_REGULATION_OF_VASCULATURE_DEVELOPMENT | 133 | -0.331246 | -1.208062 | 0.111111 | 0.35724 |
| GOBP_CELL_DIFFERENTIATION_IN_SPINAL_CORD | 27 | -0.435952 | -1.20796 | 0.211207 | 0.357175 |
| GOBP_MUSCLE_SYSTEM_PROCESS | 305 | -0.296555 | -1.207823 | 0.051402 | 0.357152 |
| GOBP_RESPONSE_TO_SALT_STRESS | 22 | -0.446778 | -1.207657 | 0.199153 | 0.357209 |
| GOBP_NEGATIVE_REGULATION_OF_INTRACELLULAR_SIGNAL_TRANSDUCTION | 435 | -0.286697 | -1.207621 | 0.034913 | 0.356995 |
| GOBP_INSULIN_RECEPTOR_SIGNALING_PATHWAY | 124 | -0.327449 | -1.207284 | 0.119403 | 0.357346 |
| GOBP_REGULATION_OF_MYELOID_LEUKOCYTE_MEDIATED_IMMUNITY | 48 | -0.383142 | -1.206625 | 0.166302 | 0.358345 |
| GOBP_REGULATION_OF_TRANSCRIPTION_INVOLVED_IN_G1_S_TRANSITION_OF_MITOTIC_C | 35 | -0.412045 | -1.20633 | 0.193833 | 0.358605 |
| GOBP_MYELOID_DENDRITIC_CELL_ACTIVATION | 27 | -0.438009 | -1.206053 | 0.198732 | 0.358861 |
| GOBP_POSITIVE_REGULATION_OF_GTPASE_ACTIVITY | 353 | -0.293851 | -1.205924 | 0.053269 | 0.358815 |
| GOBP_REGULATION_OF_OXIDATIVE_STRESS_INDUCED_INTRINSIC_APOPTOTIC_SIGNALING | 25 | -0.442148 | -1.20573 | 0.222467 | 0.35893 |
| GOBP_PHAGOSOME_MATURATION | 41 | -0.405519 | -1.205394 | 0.175153 | 0.359227 |
| GOBP_T_CELL_MEDIATED_IMMUNITY | 94 | -0.345175 | -1.205391 | 0.151079 | 0.35896 |
| GOBP_CELL_CELL_SIGNALING_BY_WNT | 447 | -0.284636 | -1.205238 | 0.033333 | 0.359012 |
| GOBP_REGULATION_OF_DNA_TEMPLATED_TRANSCRIPTION_IN_RESPONSE_TO_STRESS | 111 | -0.3316 | -1.204796 | 0.110879 | 0.359576 |
| GOBP_ORGANELLE_FUSION | 132 | -0.326723 | -1.204474 | 0.098253 | 0.359939 |
| GOBP_MULTI_ORGANISM_CELLULAR_PROCESS | 16 | -0.489459 | -1.20427 | 0.219462 | 0.360092 |
| GOBP_POSITIVE_REGULATION_OF_CELL_MIGRATION_INVOLVED_IN_SPROUTING_ANGIOGE | 19 | -0.474242 | -1.204087 | 0.221774 | 0.360164 |
| GOBP_CARDIAC_MUSCLE_CELL_ACTION_POTENTIAL_INVOLVED_IN_CONTRACTION | 37 | -0.398366 | -1.204072 | 0.176955 | 0.359933 |
| GOBP_POSITIVE_REGULATION_OF_PROTEOLYSIS_INVOLVED_IN_CELLULAR_PROTEIN_CAT | 118 | -0.328079 | -1.203843 | 0.125 | 0.360071 |
| GOBP_EXCRETION | 34 | -0.41076 | -1.203418 | 0.181619 | 0.360639 |
| GOBP_MACROAUTOPHAGY | 301 | -0.296222 | -1.203202 | 0.069378 | 0.360756 |
| GOBP_AMINE_CATABOLIC_PROCESS | 24 | -0.446756 | -1.203079 | 0.196121 | 0.360706 |
| GOBP_VESICLE_ORGANIZATION | 316 | -0.295396 | -1.202845 | 0.065375 | 0.360861 |
| GOBP_POSITIVE_REGULATION_OF_MONONUCLEAR_CELL_MIGRATION | 53 | -0.381361 | -1.202396 | 0.152361 | 0.36137 |
| GOBP_ISOPRENOID_METABOLIC_PROCESS | 89 | -0.352059 | -1.202358 | 0.159574 | 0.361163 |
| GOBP_AMINOGLYCAN_BIOSYNTHETIC_PROCESS | 97 | -0.338192 | -1.20188 | 0.114407 | 0.361784 |
| GOBP_REGULATION_OF_TISSUE_REMODELING | 54 | -0.371715 | -1.201761 | 0.161491 | 0.361725 |
| GOBP_REGULATION_OF_ENDOCYTOSIS | 175 | -0.313486 | -1.200824 | 0.105012 | 0.363276 |
| GOBP_KINETOCHORE_ASSEMBLY | 17 | -0.494079 | -1.200567 | 0.223849 | 0.363524 |
| GOBP_REGULATION_OF_GLYCOGEN_METABOLIC_PROCESS | 32 | -0.416937 | -1.20038 | 0.197938 | 0.363593 |
| GOBP_GLUTAMATE_RECEPTOR_SIGNALING_PATHWAY | 31 | -0.421191 | -1.200321 | 0.208511 | 0.363416 |
| GOBP_REGULATION_OF_LYSOSOMAL_LUMEN_PH | 15 | -0.500921 | -1.200307 | 0.269978 | 0.363182 |
| GOBP_REGULATION_OF_BODY_FLUID_LEVELS | 400 | -0.284789 | -1.20026 | 0.025575 | 0.362997 |
| GOBP_MRNA_TRANSPORT | 142 | -0.32183 | -1.199941 | 0.120087 | 0.363321 |
| GOBP_PROTEIN_POLYMERIZATION | 260 | -0.297556 | -1.199832 | 0.060538 | 0.363267 |
| GOBP_METAPHASE_PLATE_CONGRESSION | 63 | -0.368863 | -1.199318 | 0.183299 | 0.36397 |
| GOBP_POSITIVE_REGULATION_OF_CHROMOSOME_ORGANIZATION | 153 | -0.31909 | -1.19888 | 0.121348 | 0.364514 |
| GOBP_RESPONSE_TO_ALCOHOL | 182 | -0.313249 | -1.198434 | 0.094923 | 0.365068 |
| GOBP_REGULATION_OF_PROTEIN_DEPHOSPHORYLATION | 132 | -0.330886 | -1.198052 | 0.105856 | 0.365487 |
| GOBP_CELL_FATE_DETERMINATION | 23 | -0.446138 | -1.197879 | 0.243187 | 0.365577 |
| GOBP_RESPONSE_TO_VITAMIN | 66 | -0.359518 | -1.197024 | 0.165179 | 0.36694 |
| GOBP_MEMBRANE_LIPID_METABOLIC_PROCESS | 194 | -0.308934 | -1.19693 | 0.098947 | 0.366847 |
| GOBP_CELLULAR_MONOVALENT_INORGANIC_CATION_HOMEOSTASIS | 100 | -0.336768 | -1.196924 | 0.123644 | 0.366581 |
| GOBP_REGULATION_OF_INTRACELLULAR_TRANSPORT | 319 | -0.292228 | -1.196899 | 0.057554 | 0.366359 |
| GOBP_REGULATION_OF_PEPTIDASE_ACTIVITY | 338 | -0.291343 | -1.196476 | 0.053571 | 0.366951 |
| GOBP_SENSORY_PERCEPTION_OF_MECHANICAL_STIMULUS | 118 | -0.326572 | -1.196472 | 0.135983 | 0.366681 |
| GOBP_NEGATIVE_REGULATION_OF_GROWTH | 198 | -0.308574 | -1.196178 | 0.086168 | 0.367007 |
| GOBP_BRANCHING_INVOLVED_IN_BLOOD_VESSEL_MORPHOGENESIS | 23 | -0.444848 | -1.196121 | 0.220302 | 0.366839 |
| GOBP_HOMOTYPIC_CELL_CELL_ADHESION | 70 | -0.356341 | -1.19599 | 0.165158 | 0.366846 |
| GOBP_MUSCLE_CELL_DIFFERENTIATION | 273 | -0.296273 | -1.195791 | 0.081281 | 0.366968 |
| GOBP_VESICLE_CARGO_LOADING | 22 | -0.459934 | -1.195689 | 0.215321 | 0.366895 |
| GOBP_TISSUE_HOMEOSTASIS | 207 | -0.30853 | -1.195628 | 0.082251 | 0.36675 |
| GOBP_POSITIVE_REGULATION_OF_PEPTIDASE_ACTIVITY | 175 | -0.311778 | -1.195624 | 0.104019 | 0.366484 |
| GOBP_CARDIAC_VENTRICLE_MORPHOGENESIS | 45 | -0.381232 | -1.195567 | 0.182018 | 0.366307 |
| GOBP_NUCLEOSOME_ORGANIZATION | 146 | -0.322369 | -1.195367 | 0.111392 | 0.366431 |
| GOBP_REGULATION_OF_PRESYNAPSE_ORGANIZATION | 17 | -0.476664 | -1.194846 | 0.259794 | 0.367108 |
| GOBP_CELLULAR_EXTRAVASATION | 60 | -0.363783 | -1.194368 | 0.153226 | 0.367695 |
| GOBP_NOREPINEPHRINE_TRANSPORT | 16 | -0.480941 | -1.193746 | 0.242798 | 0.368653 |
| GOBP_SISTER_CHROMATID_SEGREGATION | 193 | -0.310094 | -1.193718 | 0.090465 | 0.368422 |
| GOBP_MONONUCLEAR_CELL_DIFFERENTIATION | 364 | -0.289675 | -1.193685 | 0.060325 | 0.368211 |
| GOBP_POSITIVE_REGULATION_OF_OSSIFICATION | 37 | -0.405399 | -1.193532 | 0.188797 | 0.368214 |
| GOBP_NITRIC_OXIDE_MEDIATED_SIGNAL_TRANSDUCTION | 15 | -0.486028 | -1.19301 | 0.251004 | 0.368883 |
| GOBP_MODIFIED_AMINO_ACID_TRANSPORT | 26 | -0.426986 | -1.192792 | 0.195699 | 0.369032 |
| GOBP_NEGATIVE_REGULATION_OF_INTRINSIC_APOPTOTIC_SIGNALING_PATHWAY | 86 | -0.343129 | -1.19269 | 0.157143 | 0.368957 |
| GOBP_REGULATION_OF_TUMOR_NECROSIS_FACTOR_MEDIATED_SIGNALING_PATHWAY | 55 | -0.372399 | -1.191796 | 0.172043 | 0.370389 |
| GOBP_POSITIVE_REGULATION_OF_CELL_ADHESION | 366 | -0.286935 | -1.191605 | 0.047368 | 0.370518 |
| GOBP_NEGATIVE_REGULATION_OF_PEPTIDASE_ACTIVITY | 163 | -0.31244 | -1.190961 | 0.096491 | 0.371509 |
| GOBP_RESPONSE_TO_X_RAY | 31 | -0.415372 | -1.190896 | 0.193277 | 0.37135 |
| GOBP_REGULATION_OF_PROTEIN_SUMOYLATION | 22 | -0.441239 | -1.190204 | 0.22664 | 0.3724 |
| GOBP_REGULATION_OF_CELL_DEVELOPMENT | 387 | -0.28638 | -1.189883 | 0.054374 | 0.372735 |
| GOBP_DEMETHYLATION | 63 | -0.360259 | -1.189041 | 0.178495 | 0.374069 |
| GOBP_REGULATION_OF_TUBE_SIZE | 86 | -0.344951 | -1.188994 | 0.138004 | 0.373865 |
| GOBP_REGULATION_OF_INSULIN_RECEPTOR_SIGNALING_PATHWAY | 59 | -0.362149 | -1.188991 | 0.180851 | 0.373596 |
| GOBP_SMOOTH_MUSCLE_CONTRACTION | 66 | -0.359634 | -1.188847 | 0.173121 | 0.37361 |
| GOBP_REGULATION_OF_GENERATION_OF_PRECURSOR_METABOLITES_AND_ENERGY | 143 | -0.320535 | -1.18854 | 0.126362 | 0.373936 |
| GOBP_REGULATION_OF_NEUROTRANSMITTER_RECEPTOR_ACTIVITY | 58 | -0.362184 | -1.188488 | 0.197339 | 0.373757 |
| GOBP_PHOSPHOLIPID_TRANSPORT | 75 | -0.350768 | -1.188283 | 0.157563 | 0.373854 |
| GOBP_REGULATION_OF_T_HELPER_1_TYPE_IMMUNE_RESPONSE | 22 | -0.445717 | -1.187613 | 0.240557 | 0.37497 |
| GOBP_TUMOR_NECROSIS_FACTOR_MEDIATED_SIGNALING_PATHWAY | 159 | -0.313239 | -1.187457 | 0.102908 | 0.374992 |
| GOBP_AMP_METABOLIC_PROCESS | 16 | -0.477876 | -1.18738 | 0.260246 | 0.374868 |
| GOBP_NEGATIVE_REGULATION_OF_PRODUCTION_OF_MOLECULAR_MEDIATOR_OF_IMMUN | 34 | -0.405673 | -1.187157 | 0.21978 | 0.375008 |
| GOBP_RIBONUCLEOSIDE_METABOLIC_PROCESS | 65 | -0.355719 | -1.186997 | 0.163482 | 0.37508 |
| GOBP_NEGATIVE_REGULATION_OF_ADAPTIVE_IMMUNE_RESPONSE | 51 | -0.375536 | -1.185665 | 0.181628 | 0.377426 |
| GOBP_MULTI_ORGANISM_LOCALIZATION | 65 | -0.359539 | -1.185464 | 0.174603 | 0.377601 |
| GOBP_POSITIVE_REGULATION_OF_NF_KAPPAB_TRANSCRIPTION_FACTOR_ACTIVITY | 143 | -0.317641 | -1.185353 | 0.135484 | 0.377562 |
| GOBP_POSITIVE_REGULATION_OF_PEPTIDYL_SERINE_PHOSPHORYLATION | 77 | -0.344169 | -1.185337 | 0.173362 | 0.37732 |
| GOBP_NEGATIVE_REGULATION_OF_ERK1_AND_ERK2_CASCADE | 64 | -0.359792 | -1.18493 | 0.17833 | 0.377884 |
| GOBP_MONOCARBOXYLIC_ACID_TRANSPORT | 126 | -0.321933 | -1.184902 | 0.135965 | 0.377671 |
| GOBP_POSITIVE_REGULATION_BY_HOST_OF_VIRAL_TRANSCRIPTION | 15 | -0.485395 | -1.184498 | 0.252193 | 0.378202 |
| GOBP_AGING | 248 | -0.298143 | -1.184237 | 0.083333 | 0.378466 |
| GOBP_REGULATION_OF_STEM_CELL_DIFFERENTIATION | 100 | -0.33692 | -1.184206 | 0.149184 | 0.378265 |
| GOBP_LIPOPROTEIN_METABOLIC_PROCESS | 115 | -0.326264 | -1.18398 | 0.145652 | 0.378456 |
| GOBP_REGULATION_OF_MONONUCLEAR_CELL_MIGRATION | 98 | -0.335862 | -1.183821 | 0.146288 | 0.378498 |
| GOBP_CARTILAGE_DEVELOPMENT | 142 | -0.318425 | -1.183613 | 0.131222 | 0.378651 |
| GOBP_NEGATIVE_REGULATION_OF_BLOOD_PRESSURE | 23 | -0.435845 | -1.183489 | 0.23374 | 0.378612 |
| GOBP_REGULATION_OF_MICROTUBULE_POLYMERIZATION_OR_DEPOLYMERIZATION | 71 | -0.351755 | -1.182912 | 0.170306 | 0.37949 |
| GOBP_ANIMAL_ORGAN_MATURATION | 21 | -0.454807 | -1.182688 | 0.227083 | 0.379664 |
| GOBP_REGULATION_OF_RELEASE_OF_CYTOCHROME_C_FROM_MITOCHONDRIA | 43 | -0.39207 | -1.182247 | 0.20339 | 0.380213 |
| GOBP_POSITIVE_REGULATION_OF_BLOOD_CIRCULATION | 27 | -0.432718 | -1.1821 | 0.224033 | 0.380252 |
| GOBP_POSITIVE_REGULATION_OF_G_PROTEIN_COUPLED_RECEPTOR_SIGNALING_PATHWA | 17 | -0.472318 | -1.182048 | 0.249497 | 0.380085 |
| GOBP_POSITIVE_REGULATION_OF_DNA_TEMPLATED_TRANSCRIPTION_INITIATION | 32 | -0.409828 | -1.182027 | 0.212185 | 0.379859 |
| GOBP_CELLULAR_RESPONSE_TO_THYROID_HORMONE_STIMULUS | 15 | -0.485369 | -1.181874 | 0.246862 | 0.379883 |
| GOBP_CELL_GROWTH | 391 | -0.284734 | -1.181573 | 0.06264 | 0.380163 |
| GOBP_BRANCHING_MORPHOGENESIS_OF_AN_EPITHELIAL_TUBE | 100 | -0.333721 | -1.180926 | 0.160944 | 0.38117 |
| GOBP_CELLULAR_RESPONSE_TO_RADIATION | 162 | -0.312118 | -1.180567 | 0.111364 | 0.381631 |
| GOBP_CAMP_MEDIATED_SIGNALING | 45 | -0.383136 | -1.180156 | 0.203426 | 0.382198 |
| GOBP_CYTOKINESIS | 153 | -0.30983 | -1.180141 | 0.125551 | 0.381948 |
| GOBP_REGULATION_OF_STRIATED_MUSCLE_CELL_DIFFERENTIATION | 63 | -0.355838 | -1.180046 | 0.178571 | 0.381848 |
| GOBP_CELL_MIGRATION_INVOLVED_IN_SPROUTING_ANGIOGENESIS | 49 | -0.375109 | -1.179852 | 0.194969 | 0.381946 |
| GOBP_REGULATION_OF_B_CELL_PROLIFERATION | 58 | -0.363813 | -1.179836 | 0.182796 | 0.381693 |

| | | | | | |
|---|---|---|---|---|---|
| GOBP_REGULATION_OF_TRIGLYCERIDE_BIOSYNTHETIC_PROCESS | 17 | -0.478111 | -1.179816 | 0.216895 | 0.381469 |
| GOBP_DENDRITE_EXTENSION | 27 | -0.430108 | -1.179638 | 0.225873 | 0.381531 |
| GOBP_CARDIAC_MUSCLE_CELL_DIFFERENTIATION | 78 | -0.343426 | -1.179596 | 0.163636 | 0.381357 |
| GOBP_NEGATIVE_REGULATION_OF_TRANSPORTER_ACTIVITY | 56 | -0.364077 | -1.17951 | 0.198178 | 0.381232 |
| GOBP_PROTEIN_KINASE_B_SIGNALING | 206 | -0.303674 | -1.179359 | 0.10625 | 0.381255 |
| GOBP_PROTEIN_DEPHOSPHORYLATION | 294 | -0.291477 | -1.178299 | 0.086162 | 0.38294 |
| GOBP_STEROID_HORMONE_MEDIATED_SIGNALING_PATHWAY | 111 | -0.326473 | -1.178017 | 0.144522 | 0.383207 |
| GOBP_POSITIVE_REGULATION_OF_PROTEIN_TARGETING_TO_MITOCHONDRION | 30 | -0.420092 | -1.177546 | 0.243243 | 0.383902 |
| GOBP_T_CELL_DIFFERENTIATION_INVOLVED_IN_IMMUNE_RESPONSE | 67 | -0.356305 | -1.177481 | 0.193206 | 0.383766 |
| GOBP_FIBROBLAST_GROWTH_FACTOR_RECEPTOR_SIGNALING_PATHWAY | 85 | -0.342038 | -1.177419 | 0.170168 | 0.383622 |
| GOBP_BIOLOGICAL_PROCESS_INVOLVED_IN_INTRASPECIES_INTERACTION_BETWEEN_OR | 38 | -0.395473 | -1.177379 | 0.207364 | 0.383421 |
| GOBP_REGULATION_OF_LIPID_METABOLIC_PROCESS | 325 | -0.288017 | -1.176534 | 0.069544 | 0.384755 |
| GOBP_LIPID_OXIDATION | 101 | -0.335019 | -1.176449 | 0.153153 | 0.384667 |
| GOBP_CELLULAR_RESPONSE_TO_DRUG | 52 | -0.371046 | -1.17611 | 0.192641 | 0.385141 |
| GOBP_ALDITOL_METABOLIC_PROCESS | 17 | -0.47966 | -1.175913 | 0.251984 | 0.385259 |
| GOBP_GLUCOSAMINE_CONTAINING_COMPOUND_METABOLIC_PROCESS | 24 | -0.439968 | -1.175815 | 0.238589 | 0.385188 |
| GOBP_PROTEIN_HOMOOLIGOMERIZATION | 136 | -0.318633 | -1.175731 | 0.133489 | 0.385085 |
| GOBP_CANONICAL_WNT_SIGNALING_PATHWAY | 286 | -0.292704 | -1.175341 | 0.104019 | 0.385606 |
| GOBP_DENDRITIC_CELL_DIFFERENTIATION | 43 | -0.389505 | -1.175028 | 0.230932 | 0.385962 |
| GOBP_NEURON_APOPTOTIC_PROCESS | 185 | -0.303122 | -1.174865 | 0.092593 | 0.386029 |
| GOBP_EXOCRINE_SYSTEM_DEVELOPMENT | 33 | -0.400651 | -1.174738 | 0.227172 | 0.385986 |
| GOBP_METAL_ION_EXPORT | 38 | -0.399375 | -1.174465 | 0.239726 | 0.386262 |
| GOBP_REGULATION_OF_MYELOID_CELL_APOPTOTIC_PROCESS | 18 | -0.466234 | -1.174276 | 0.271186 | 0.386371 |
| GOBP_REGULATION_OF_MUSCLE_CELL_DIFFERENTIATION | 114 | -0.323522 | -1.174252 | 0.141553 | 0.38615 |
| GOBP_MUSCLE_HYPERTROPHY_IN_RESPONSE_TO_STRESS | 17 | -0.458237 | -1.174236 | 0.245763 | 0.385909 |
| GOBP_MICROTUBULE_CYTOSKELETON_ORGANIZATION_INVOLVED_IN_MITOSIS | 139 | -0.313851 | -1.17302 | 0.11954 | 0.388096 |
| GOBP_DENDRITIC_CELL_MIGRATION | 26 | -0.424231 | -1.172803 | 0.259875 | 0.38825 |
| GOBP_REGULATION_OF_SMOOTH_MUSCLE_CONTRACTION | 40 | -0.389104 | -1.17273 | 0.208 | 0.388112 |
| GOBP_MOTOR_NEURON_AXON_GUIDANCE | 24 | -0.433759 | -1.172604 | 0.255365 | 0.388104 |
| GOBP_ESTABLISHMENT_OR_MAINTENANCE_OF_CELL_POLARITY | 198 | -0.300203 | -1.172539 | 0.110874 | 0.387967 |
| GOBP_CALCIUM_ION_TRANSPORT_INTO_CYTOSOL | 117 | -0.326597 | -1.171852 | 0.148741 | 0.388983 |
| GOBP_SIGNAL_RELEASE | 361 | -0.282529 | -1.17099 | 0.068293 | 0.390525 |
| GOBP_MALE_MEIOSIS_I | 17 | -0.483577 | -1.16956 | 0.277433 | 0.39302 |
| GOBP_REGULATION_OF_EXOCYTOSIS | 162 | -0.307656 | -1.169072 | 0.154709 | 0.393706 |
| GOBP_MITOCHONDRION_MORPHOGENESIS | 18 | -0.460004 | -1.168819 | 0.278373 | 0.393923 |
| GOBP_MESODERM_DEVELOPMENT | 92 | -0.334546 | -1.168383 | 0.189474 | 0.394543 |
| GOBP_POSITIVE_REGULATION_OF_BINDING | 152 | -0.308702 | -1.168256 | 0.14059 | 0.394498 |
| GOBP_EPITHELIAL_CELL_DEVELOPMENT | 167 | -0.309837 | -1.168232 | 0.136364 | 0.394271 |
| GOBP_GLYCEROLIPID_CATABOLIC_PROCESS | 48 | -0.368284 | -1.168021 | 0.207965 | 0.394424 |
| GOBP_POSITIVE_REGULATION_OF_DENDRITIC_SPINE_DEVELOPMENT | 40 | -0.384717 | -1.167911 | 0.233624 | 0.394382 |
| GOBP_NEGATIVE_REGULATION_OF_ENDOTHELIAL_CELL_PROLIFERATION | 37 | -0.391924 | -1.167745 | 0.219959 | 0.394412 |
| GOBP_NEGATIVE_REGULATION_OF_GLIOGENESIS | 20 | -0.456206 | -1.167661 | 0.244635 | 0.394317 |
| GOBP_NEGATIVE_REGULATION_OF_LYMPHOCYTE_APOPTOTIC_PROCESS | 29 | -0.420083 | -1.167569 | 0.228033 | 0.394237 |
| GOBP_CARTILAGE_DEVELOPMENT_INVOLVED_IN_ENDOCHONDRAL_BONE_MORPHOGENE | 20 | -0.451533 | -1.167105 | 0.265107 | 0.394939 |
| GOBP_OOCYTE_MATURATION | 22 | -0.442725 | -1.166724 | 0.237113 | 0.395447 |
| GOBP_REGULATION_OF_PEPTIDYL_SERINE_PHOSPHORYLATION | 106 | -0.3247 | -1.166583 | 0.163866 | 0.395477 |
| GOBP_OSTEOBLAST_DIFFERENTIATION | 177 | -0.306427 | -1.166381 | 0.131519 | 0.395603 |
| GOBP_STRIATED_MUSCLE_CELL_DIFFERENTIATION | 201 | -0.303089 | -1.166098 | 0.128319 | 0.395864 |
| GOBP_GLYCOSPHINGOLIPID_BIOSYNTHETIC_PROCESS | 25 | -0.431695 | -1.166077 | 0.235417 | 0.395632 |
| GOBP_RESPONSE_TO_UV | 131 | -0.313486 | -1.165991 | 0.137168 | 0.39555 |
| GOBP_REGULATION_OF_CIRCADIAN_RHYTHM | 94 | -0.335656 | -1.16541 | 0.187364 | 0.396509 |
| GOBP_REGULATION_OF_REGULATED_SECRETORY_PATHWAY | 112 | -0.321791 | -1.165305 | 0.171558 | 0.396435 |
| GOBP_PYRIMIDINE_NUCLEOSIDE_MONOPHOSPHATE_METABOLIC_PROCESS | 18 | -0.463706 | -1.164932 | 0.271008 | 0.396932 |
| GOBP_PHOTORECEPTOR_CELL_DEVELOPMENT | 30 | -0.4098 | -1.16488 | 0.246939 | 0.396771 |
| GOBP_NEGATIVE_REGULATION_OF_GENE_SILENCING | 31 | -0.405914 | -1.164853 | 0.22833 | 0.396555 |
| GOBP_LIVER_REGENERATION | 27 | -0.424393 | -1.164834 | 0.21281 | 0.396331 |
| GOBP_CHEMICAL_SYNAPTIC_TRANSMISSION_POSTSYNAPTIC | 73 | -0.340825 | -1.164399 | 0.18595 | 0.396892 |
| GOBP_ADENYLATE_CYCLASE_INHIBITING_G_PROTEIN_COUPLED_RECEPTOR_SIGNALING_ | 50 | -0.362411 | -1.164334 | 0.218818 | 0.396717 |
| GOBP_BONE_DEVELOPMENT | 159 | -0.306048 | -1.164201 | 0.144231 | 0.396731 |
| GOBP_SMAD_PROTEIN_SIGNAL_TRANSDUCTION | 62 | -0.356634 | -1.1639 | 0.209713 | 0.397109 |
| GOBP_STEROID_CATABOLIC_PROCESS | 16 | -0.466773 | -1.163748 | 0.243187 | 0.39715 |
| GOBP_FATTY_ACID_DERIVATIVE_BIOSYNTHETIC_PROCESS | 52 | -0.37677 | -1.16374 | 0.229299 | 0.396893 |
| GOBP_EAR_DEVELOPMENT | 145 | -0.309765 | -1.163549 | 0.139293 | 0.397008 |
| GOBP_CELLULAR_CARBOHYDRATE_BIOSYNTHETIC_PROCESS | 70 | -0.344444 | -1.163118 | 0.18427 | 0.397563 |
| GOBP_MONOSACCHARIDE_METABOLIC_PROCESS | 238 | -0.291579 | -1.162939 | 0.094563 | 0.397665 |
| GOBP_RESPONSE_TO_FOOD | 21 | -0.433497 | -1.162863 | 0.244399 | 0.397541 |
| GOBP_POSITIVE_REGULATION_OF_AMYLOID_BETA_FORMATION | 17 | -0.462471 | -1.162844 | 0.290254 | 0.397319 |
| GOBP_COLLAGEN_METABOLIC_PROCESS | 82 | -0.33658 | -1.16201 | 0.18319 | 0.398643 |
| GOBP_MYOTUBE_DIFFERENTIATION | 85 | -0.338732 | -1.161922 | 0.185682 | 0.398539 |
| GOBP_ALCOHOL_CATABOLIC_PROCESS | 41 | -0.38188 | -1.161921 | 0.246725 | 0.39827 |
| GOBP_RESPONSE_TO_FATTY_ACID | 47 | -0.376136 | -1.161246 | 0.225263 | 0.399392 |
| GOBP_EMBRYO_IMPLANTATION | 36 | -0.391778 | -1.160619 | 0.223819 | 0.400406 |
| GOBP_CYTOSKELETON_DEPENDENT_INTRACELLULAR_TRANSPORT | 187 | -0.29979 | -1.160599 | 0.146572 | 0.400188 |
| GOBP_T_CELL_CHEMOTAXIS | 20 | -0.454562 | -1.160503 | 0.282158 | 0.40011 |
| GOBP_NEGATIVE_REGULATION_OF_INTRACELLULAR_TRANSPORT | 47 | -0.37446 | -1.160396 | 0.208421 | 0.400055 |
| GOBP_VASCULAR_ENDOTHELIAL_CELL_PROLIFERATION | 22 | -0.44512 | -1.160308 | 0.248366 | 0.399952 |
| GOBP_POSITIVE_REGULATION_OF_FATTY_ACID_BIOSYNTHETIC_PROCESS | 17 | -0.465189 | -1.160268 | 0.272541 | 0.399769 |
| GOBP_MEMBRANE_FUSION | 145 | -0.308502 | -1.160042 | 0.131818 | 0.399959 |
| GOBP_NEGATIVE_REGULATION_OF_PROTEIN_CONTAINING_COMPLEX_DISASSEMBLY | 70 | -0.345205 | -1.16001 | 0.18319 | 0.399754 |
| GOBP_MODULATION_BY_SYMBIONT_OF_HOST_PROCESS | 35 | -0.393654 | -1.159821 | 0.213836 | 0.399866 |
| GOBP_REGULATION_OF_EXOSOMAL_SECRETION | 16 | -0.469705 | -1.159008 | 0.299389 | 0.401224 |
| GOBP_G_PROTEIN_COUPLED_RECEPTOR_SIGNALING_PATHWAY_COUPLED_TO_CYCLIC_N | 26 | -0.426623 | -1.158421 | 0.261339 | 0.402153 |
| GOBP_PURINERGIC_NUCLEOTIDE_RECEPTOR_SIGNALING_PATHWAY | 28 | -0.414535 | -1.158215 | 0.255717 | 0.402293 |
| GOBP_NEGATIVE_REGULATION_OF_INTERLEUKIN_6_PRODUCTION | 40 | -0.398987 | -1.158168 | 0.247401 | 0.402102 |
| GOBP_NERVE_DEVELOPMENT | 46 | -0.373517 | -1.157911 | 0.213415 | 0.402356 |
| GOBP_REGULATION_OF_HETEROTYPIC_CELL_CELL_ADHESION | 20 | -0.453582 | -1.156537 | 0.280412 | 0.405002 |
| GOBP_ATP_BIOSYNTHETIC_PROCESS | 55 | -0.360766 | -1.156459 | 0.216471 | 0.404869 |
| GOBP_NEURON_DEATH_IN_RESPONSE_TO_OXIDATIVE_STRESS | 25 | -0.41988 | -1.156409 | 0.262055 | 0.404698 |
| GOBP_SPHINGOLIPID_METABOLIC_PROCESS | 154 | -0.308111 | -1.156369 | 0.145055 | 0.404513 |
| GOBP_POSITIVE_REGULATION_OF_MITOCHONDRION_ORGANIZATION | 82 | -0.339923 | -1.15626 | 0.220408 | 0.404453 |
| GOBP_POSITIVE_REGULATION_OF_TELOMERASE_ACTIVITY | 31 | -0.411009 | -1.156167 | 0.275269 | 0.404363 |
| GOBP_CYTOSOLIC_CALCIUM_ION_TRANSPORT | 138 | -0.302763 | -1.155791 | 0.128378 | 0.404843 |
| GOBP_NEGATIVE_REGULATION_OF_CYTOKINE_PRODUCTION_INVOLVED_IN_IMMUNE_RES | 24 | -0.417806 | -1.155387 | 0.269231 | 0.405369 |
| GOBP_SPERMATID_DIFFERENTIATION | 109 | -0.319746 | -1.155085 | 0.166667 | 0.405682 |
| GOBP_NEGATIVE_REGULATION_OF_LYMPHOCYTE_MEDIATED_IMMUNITY | 43 | -0.376986 | -1.155045 | 0.226374 | 0.405494 |
| GOBP_SKELETAL_MUSCLE_ORGAN_DEVELOPMENT | 117 | -0.316099 | -1.154751 | 0.179545 | 0.405776 |
| GOBP_DNA_CONFORMATION_CHANGE | 307 | -0.28393 | -1.154614 | 0.096852 | 0.40579 |
| GOBP_REGULATION_OF_OXIDOREDUCTASE_ACTIVITY | 86 | -0.331958 | -1.15445 | 0.177824 | 0.405845 |
| GOBP_GTP_METABOLIC_PROCESS | 22 | -0.434832 | -1.154231 | 0.261224 | 0.406041 |
| GOBP_FIBROBLAST_PROLIFERATION | 69 | -0.345228 | -1.15423 | 0.181425 | 0.405774 |
| GOBP_CELLULAR_RESPONSE_TO_AMYLOID_BETA | 34 | -0.394338 | -1.154186 | 0.253192 | 0.405584 |
| GOBP_NEGATIVE_REGULATION_OF_CD4_POSITIVE_ALPHA_BETA_T_CELL_ACTIVATION | 29 | -0.408801 | -1.153617 | 0.266129 | 0.40645 |
| GOBP_MACROPHAGE_CHEMOTAXIS | 35 | -0.388153 | -1.153454 | 0.2527 | 0.406533 |
| GOBP_NEGATIVE_REGULATION_OF_I_KAPPAB_KINASE_NF_KAPPAB_SIGNALING | 42 | -0.378708 | -1.153404 | 0.24714 | 0.406385 |
| GOBP_NEGATIVE_REGULATION_OF_SMALL_MOLECULE_METABOLIC_PROCESS | 73 | -0.3413 | -1.153191 | 0.201754 | 0.406526 |
| GOBP_MATURE_B_CELL_DIFFERENTIATION_INVOLVED_IN_IMMUNE_RESPONSE | 25 | -0.428781 | -1.152558 | 0.271552 | 0.407644 |
| GOBP_PROGRAMMED_NECROTIC_CELL_DEATH | 44 | -0.367318 | -1.152075 | 0.236786 | 0.408363 |
| GOBP_LUNG_MORPHOGENESIS | 36 | -0.385056 | -1.151411 | 0.249489 | 0.409404 |
| GOBP_N_ACETYLGLUCOSAMINE_METABOLIC_PROCESS | 18 | -0.450974 | -1.150436 | 0.281513 | 0.411095 |
| GOBP_EMBRYONIC_DIGESTIVE_TRACT_DEVELOPMENT | 20 | -0.445012 | -1.150332 | 0.294118 | 0.411024 |
| GOBP_MONOCARBOXYLIC_ACID_CATABOLIC_PROCESS | 114 | -0.318533 | -1.150173 | 0.209302 | 0.411062 |
| GOBP_INTERLEUKIN_7_MEDIATED_SIGNALING_PATHWAY | 24 | -0.423249 | -1.150161 | 0.266529 | 0.410818 |
| GOBP_ICOSANOID_BIOSYNTHETIC_PROCESS | 41 | -0.385476 | -1.149997 | 0.237136 | 0.41087 |
| GOBP_NEGATIVE_REGULATION_OF_ACTIN_FILAMENT_POLYMERIZATION | 56 | -0.356967 | -1.14976 | 0.209052 | 0.411074 |
| GOBP_POSITIVE_REGULATION_OF_CYCLIN_DEPENDENT_PROTEIN_KINASE_ACTIVITY | 28 | -0.418351 | -1.149352 | 0.253579 | 0.411641 |
| GOBP_REGULATION_OF_PROTEIN_AUTOPHOSPHORYLATION | 41 | -0.370728 | -1.149236 | 0.240449 | 0.411609 |
| GOBP_PROTEIN_TRANSMEMBRANE_TRANSPORT | 57 | -0.353926 | -1.1489 | 0.205821 | 0.412021 |
| GOBP_LIPOPROTEIN_BIOSYNTHETIC_PROCESS | 86 | -0.333219 | -1.148537 | 0.209402 | 0.41249 |
| GOBP_POSITIVE_REGULATION_OF_INTRACELLULAR_PROTEIN_TRANSPORT | 166 | -0.299906 | -1.147561 | 0.158129 | 0.414246 |
| GOBP_NEURON_MATURATION | 35 | -0.392147 | -1.14743 | 0.246888 | 0.414257 |
| GOBP_REGULATION_OF_ACTION_POTENTIAL | 37 | -0.385019 | -1.147405 | 0.256522 | 0.414024 |
| GOBP_SELECTIVE_AUTOPHAGY | 60 | -0.348861 | -1.147304 | 0.216274 | 0.413968 |
| GOBP_REGULATION_OF_LEUKOCYTE_DIFFERENTIATION | 248 | -0.288169 | -1.147181 | 0.134259 | 0.413969 |
| GOBP_NEGATIVE_REGULATION_OF_ATP_METABOLIC_PROCESS | 25 | -0.431088 | -1.147178 | 0.292683 | 0.413703 |
| GOBP_REGULATION_OF_DENDRITIC_SPINE_DEVELOPMENT | 56 | -0.352899 | -1.146089 | 0.231441 | 0.415636 |
| GOBP_POSITIVE_REGULATION_OF_MAST_CELL_ACTIVATION | 18 | -0.457507 | -1.145313 | 0.298925 | 0.417006 |
| GOBP_CELLULAR_RESPONSE_TO_GLUCOSE_STARVATION | 46 | -0.363416 | -1.145102 | 0.232653 | 0.417169 |
| GOBP_PROTEIN_TARGETING_TO_MITOCHONDRION | 98 | -0.323745 | -1.144987 | 0.199536 | 0.417128 |
| GOBP_ESTABLISHMENT_OF_TISSUE_POLARITY | 113 | -0.317627 | -1.144816 | 0.203704 | 0.417177 |
| GOBP_REGULATION_OF_NITRIC_OXIDE_SYNTHASE_ACTIVITY | 41 | -0.37796 | -1.144673 | 0.248408 | 0.417209 |
| GOBP_OVULATION_CYCLE | 46 | -0.369266 | -1.14467 | 0.257143 | 0.416942 |
| GOBP_POSITIVE_REGULATION_OF_EPITHELIAL_CELL_APOPTOTIC_PROCESS | 31 | -0.393208 | -1.144647 | 0.275862 | 0.416711 |
| GOBP_RESPIRATORY_ELECTRON_TRANSPORT_CHAIN | 110 | -0.320626 | -1.144471 | 0.181818 | 0.416795 |
| GOBP_POSITIVE_REGULATION_OF_ANION_TRANSPORT | 405 | -0.276 | -1.144424 | 0.103614 | 0.416627 |
| GOBP_DENDRITIC_CELL_CHEMOTAXIS | 21 | -0.438625 | -1.144065 | 0.289916 | 0.417166 |
| GOBP_POSITIVE_REGULATION_OF_PRODUCTION_OF_MOLECULAR_MEDIATOR_OF_IMMUNE | 89 | -0.326987 | -1.143569 | 0.205128 | 0.417867 |
| GOBP_NEGATIVE_REGULATION_OF_LIPID_TRANSPORT | 23 | -0.425389 | -1.14355 | 0.276596 | 0.417641 |
| GOBP_PROTEIN_O_LINKED_MANNOSYLATION | 18 | -0.451514 | -1.14328 | 0.27027 | 0.417952 |
| GOBP_REGULATION_OF_EMBRYONIC_DEVELOPMENT | 48 | -0.361402 | -1.14311 | 0.231263 | 0.418061 |
| GOBP_RESPONSE_TO_AMYLOID_BETA | 35 | -0.389924 | -1.14281 | 0.249509 | 0.418426 |
| GOBP_POSITIVE_REGULATION_OF_MUSCLE_CELL_APOPTOTIC_PROCESS | 21 | -0.429174 | -1.142736 | 0.280335 | 0.418289 |
| GOBP_RESPONSE_TO_ACID_CHEMICAL | 101 | -0.325103 | -1.142572 | 0.200864 | 0.418324 |
| GOBP_NEGATIVE_REGULATION_OF_CELL_CELL_ADHESION | 165 | -0.298587 | -1.142518 | 0.165605 | 0.418165 |
| GOBP_NEGATIVE_REGULATION_OF_STRESS_ACTIVATED_PROTEIN_KINASE_SIGNALING_CA | 41 | -0.377711 | -1.142333 | 0.236253 | 0.418296 |
| GOBP_POSITIVE_REGULATION_OF_ORGANELLE_ASSEMBLY | 60 | -0.351868 | -1.142168 | 0.219565 | 0.418386 |
| GOBP_RESPONSE_TO_RADIATION | 380 | -0.274928 | -1.142077 | 0.105398 | 0.418295 |
| GOBP_ATP_METABOLIC_PROCESS | 283 | -0.282142 | -1.141707 | 0.127315 | 0.418791 |
| GOBP_C21_STEROID_HORMONE_BIOSYNTHETIC_PROCESS | 19 | -0.441657 | -1.140869 | 0.274227 | 0.420203 |
| GOBP_AMINOGLYCAN_METABOLIC_PROCESS | 142 | -0.310261 | -1.14038 | 0.172949 | 0.420963 |
| GOBP_SPLICEOSOMAL_SNRNP_ASSEMBLY | 50 | -0.362306 | -1.139719 | 0.23871 | 0.422087 |
| GOBP_HISTONE_UBIQUITINATION | 45 | -0.36463 | -1.13954 | 0.243421 | 0.422201 |
| GOBP_COLLATERAL_SPROUTING | 17 | -0.461525 | -1.13902 | 0.278455 | 0.422915 |
| GOBP_SEGMENTATION | 69 | -0.337607 | -1.138775 | 0.219828 | 0.423186 |
| GOBP_CARBOHYDRATE_DERIVATIVE_CATABOLIC_PROCESS | 165 | -0.301857 | -1.138627 | 0.179825 | 0.423206 |
| GOBP_POSITIVE_REGULATION_OF_HEMOPOIESIS | 139 | -0.304335 | -1.138605 | 0.165116 | 0.422981 |
| GOBP_NUCLEOSIDE_TRIPHOSPHATE_METABOLIC_PROCESS | 104 | -0.315171 | -1.138201 | 0.21978 | 0.423545 |
| GOBP_TOLL_LIKE_RECEPTOR_SIGNALING_PATHWAY | 137 | -0.307507 | -1.137955 | 0.197248 | 0.423777 |

| | | | | | |
|---|---|---|---|---|---|
| GOBP_REGULATION_OF_NEUROTRANSMITTER_TRANSPORT | 79 | -0.335867 | -1.137677 | 0.236897 | 0.424102 |
| GOBP_REGULATION_OF_BMP_SIGNALING_PATHWAY | 72 | -0.335468 | -1.137346 | 0.243644 | 0.42453 |
| GOBP_POSITIVE_REGULATION_OF_HISTONE_DEACETYLATION | 15 | -0.466385 | -1.1371 | 0.306968 | 0.424782 |
| GOBP_PYRIMIDINE_RIBONUCLEOTIDE_METABOLIC_PROCESS | 27 | -0.406542 | -1.137072 | 0.282828 | 0.424555 |
| GOBP_REGULATION_OF_NON_CANONICAL_WNT_SIGNALING_PATHWAY | 19 | -0.445129 | -1.136956 | 0.287846 | 0.4245 |
| GOBP_MATING_BEHAVIOR | 15 | -0.472898 | -1.136046 | 0.306483 | 0.426155 |
| GOBP_MICROTUBULE_POLYMERIZATION_OR_DEPOLYMERIZATION | 101 | -0.318967 | -1.135704 | 0.205011 | 0.426623 |
| GOBP_REGULATION_OF_ATP_METABOLIC_PROCESS | 105 | -0.320371 | -1.135415 | 0.19181 | 0.426988 |
| GOBP_VACUOLAR_TRANSPORT | 146 | -0.30452 | -1.135363 | 0.209932 | 0.426812 |
| GOBP_VENTRICULAR_SYSTEM_DEVELOPMENT | 21 | -0.437399 | -1.135102 | 0.283702 | 0.427082 |
| GOBP_POSITIVE_REGULATION_OF_HEART_GROWTH | 32 | -0.399079 | -1.134903 | 0.286017 | 0.427222 |
| GOBP_LEUKOTRIENE_BIOSYNTHETIC_PROCESS | 16 | -0.463877 | -1.134857 | 0.311871 | 0.427056 |
| GOBP_CELLULAR_RESPONSE_TO_INORGANIC_SUBSTANCE | 165 | -0.299631 | -1.134651 | 0.187215 | 0.427215 |
| GOBP_REGULATION_OF_SYSTEMIC_ARTERIAL_BLOOD_PRESSURE_BY_HORMONE | 24 | -0.42288 | -1.134536 | 0.28125 | 0.427191 |
| GOBP_NEGATIVE_REGULATION_OF_PROTEASOMAL_PROTEIN_CATABOLIC_PROCESS | 44 | -0.374019 | -1.134294 | 0.257928 | 0.427434 |
| GOBP_BILE_ACID_METABOLIC_PROCESS | 32 | -0.391344 | -1.134072 | 0.280514 | 0.427645 |
| GOBP_POSITIVE_REGULATION_OF_GLUCOSE_METABOLIC_PROCESS | 34 | -0.387312 | -1.134051 | 0.272177 | 0.427415 |
| GOBP_POSITIVE_REGULATION_OF_INTRACELLULAR_TRANSPORT | 203 | -0.293594 | -1.134008 | 0.153153 | 0.427248 |
| GOBP_REGULATION_OF_CD8_POSITIVE_ALPHA_BETA_T_CELL_ACTIVATION | 18 | -0.465325 | -1.133643 | 0.302128 | 0.427741 |
| GOBP_RIBONUCLEOSIDE_TRIPHOSPHATE_METABOLIC_PROCESS | 84 | -0.328858 | -1.133446 | 0.223881 | 0.427875 |
| GOBP_FATTY_ACID_CATABOLIC_PROCESS | 99 | -0.317487 | -1.133259 | 0.211329 | 0.42804 |
| GOBP_CARBOHYDRATE_BIOSYNTHETIC_PROCESS | 180 | -0.292342 | -1.133215 | 0.171674 | 0.427854 |
| GOBP_REGULATION_OF_UBIQUITIN_DEPENDENT_PROTEIN_CATABOLIC_PROCESS | 149 | -0.30327 | -1.133019 | 0.191781 | 0.428022 |
| GOBP_HEART_DEVELOPMENT | 419 | -0.274361 | -1.132885 | 0.090465 | 0.428018 |
| GOBP_ACTIVATION_OF_NF_KAPPAB_INDUCING_KINASE_ACTIVITY | 18 | -0.444438 | -1.132835 | 0.289796 | 0.427852 |
| GOBP_NEUROTRANSMITTER_TRANSPORT | 156 | -0.302078 | -1.132738 | 0.180587 | 0.427793 |
| GOBP_POSITIVE_REGULATION_OF_CYTOKINE_PRODUCTION_INVOLVED_IN_INFLAMMATORY | 16 | -0.470569 | -1.132663 | 0.321792 | 0.427674 |
| GOBP_MEIOTIC_CELL_CYCLE_PROCESS | 159 | -0.297617 | -1.132228 | 0.165179 | 0.428353 |
| GOBP_POSITIVE_REGULATION_OF_PROTEIN_AUTOPHOSPHORYLATION | 24 | -0.414984 | -1.132197 | 0.273731 | 0.428147 |
| GOBP_INTRACELLULAR_STEROID_HORMONE_RECEPTOR_SIGNALING_PATHWAY | 97 | -0.321238 | -1.131765 | 0.207188 | 0.428754 |
| GOBP_REGULATION_OF_DEFENSE_RESPONSE_TO_VIRUS | 64 | -0.346296 | -1.131015 | 0.251111 | 0.430042 |
| GOBP_REGULATION_OF_GENE_SILENCING | 124 | -0.307826 | -1.130755 | 0.196305 | 0.43033 |
| GOBP_POSITIVE_REGULATION_OF_DEVELOPMENTAL_GROWTH | 128 | -0.309617 | -1.13065 | 0.197425 | 0.430255 |
| GOBP_POSITIVE_REGULATION_OF_TRANSFORMING_GROWTH_FACTOR_BETA_PRODUCTIOI | 17 | -0.45514 | -1.12993 | 0.310924 | 0.431478 |
| GOBP_PROTEIN_DEMANNOSYLATION | 20 | -0.438326 | -1.129908 | 0.28903 | 0.431242 |
| GOBP_TISSUE_REMODELING | 122 | -0.314287 | -1.129862 | 0.202299 | 0.43106 |
| GOBP_CELLULAR_RESPONSE_TO_CADMIUM_ION | 26 | -0.403532 | -1.129415 | 0.285097 | 0.431737 |
| GOBP_REGULATION_OF_FATTY_ACID_BIOSYNTHETIC_PROCESS | 37 | -0.376294 | -1.129414 | 0.259714 | 0.431468 |
| GOBP_CIRCADIAN_REGULATION_OF_GENE_EXPRESSION | 58 | -0.349251 | -1.129187 | 0.26226 | 0.431682 |
| GOBP_PHOTORECEPTOR_CELL_MAINTENANCE | 33 | -0.400495 | -1.1288 | 0.260965 | 0.432278 |
| GOBP_CELLULAR_TRANSITION_METAL_ION_HOMEOSTASIS | 93 | -0.318268 | -1.128258 | 0.219955 | 0.433205 |
| GOBP_POSITIVE_REGULATION_OF_RNA_SPLICING | 31 | -0.397751 | -1.127765 | 0.291304 | 0.433999 |
| GOBP_REGULATION_OF_TRANSLATIONAL_ELONGATION | 17 | -0.454921 | -1.127553 | 0.313559 | 0.434211 |
| GOBP_ENDOMEMBRANE_SYSTEM_ORGANIZATION | 441 | -0.267715 | -1.12733 | 0.098237 | 0.434399 |
| GOBP_CALCIUM_ION_TRANSPORT | 316 | -0.278372 | -1.12727 | 0.126386 | 0.434267 |
| GOBP_NEGATIVE_REGULATION_OF_RESPONSE_TO_EXTERNAL_STIMULUS | 292 | -0.273811 | -1.124954 | 0.135338 | 0.43889 |
| GOBP_REGULATION_OF_CELLULAR_AMIDE_METABOLIC_PROCESS | 399 | -0.270872 | -1.12487 | 0.1067 | 0.438802 |
| GOBP_PROTON_TRANSMEMBRANE_TRANSPORT | 133 | -0.309182 | -1.124347 | 0.209567 | 0.439611 |
| GOBP_HEART_MORPHOGENESIS | 170 | -0.29428 | -1.123821 | 0.203991 | 0.440459 |
| GOBP_NEGATIVE_REGULATION_OF_AMINE_TRANSPORT | 16 | -0.454948 | -1.123646 | 0.300211 | 0.440588 |
| GOBP_TRANSCRIPTION_ELONGATION_FROM_RNA_POLYMERASE_II_PROMOTER | 82 | -0.32502 | -1.123618 | 0.239224 | 0.440385 |
| GOBP_VASCULAR_ENDOTHELIAL_GROWTH_FACTOR_SIGNALING_PATHWAY | 35 | -0.382171 | -1.123588 | 0.270541 | 0.440192 |
| GOBP_NEGATIVE_REGULATION_OF_CELL_ADHESION | 249 | -0.282591 | -1.123476 | 0.139423 | 0.440165 |
| GOBP_RAB_PROTEIN_SIGNAL_TRANSDUCTION | 21 | -0.427323 | -1.122656 | 0.314103 | 0.441741 |
| GOBP_POSITIVE_REGULATION_OF_REGULATORY_T_CELL_DIFFERENTIATION | 16 | -0.457288 | -1.122648 | 0.314103 | 0.441485 |
| GOBP_REGULATION_OF_SMOOTH_MUSCLE_CELL_DIFFERENTIATION | 28 | -0.401166 | -1.122309 | 0.294363 | 0.441964 |
| GOBP_SERINE_FAMILY_AMINO_ACID_METABOLIC_PROCESS | 36 | -0.379975 | -1.12228 | 0.281125 | 0.44174 |
| GOBP_AMINOGLYCAN_CATABOLIC_PROCESS | 53 | -0.354787 | -1.121877 | 0.244353 | 0.442327 |
| GOBP_DEVELOPMENTAL_CELL_GROWTH | 179 | -0.292408 | -1.12149 | 0.21659 | 0.442884 |
| GOBP_TELOMERE_ORGANIZATION | 160 | -0.296633 | -1.121291 | 0.194131 | 0.44302 |
| GOBP_MICROTUBULE_BUNDLE_FORMATION | 88 | -0.322291 | -1.121224 | 0.21822 | 0.442887 |
| GOBP_MONOVALENT_INORGANIC_CATION_HOMEOSTASIS | 129 | -0.302213 | -1.121219 | 0.229437 | 0.442626 |
| GOBP_LIPID_IMPORT_INTO_CELL | 30 | -0.404366 | -1.121103 | 0.309524 | 0.442589 |
| GOBP_PROTEIN_DEMETHYLATION | 32 | -0.388625 | -1.120597 | 0.287912 | 0.443436 |
| GOBP_RESPONSE_TO_AMPHETAMINE | 24 | -0.413225 | -1.120457 | 0.305051 | 0.443492 |
| GOBP_DNA_DEMETHYLATION | 25 | -0.410403 | -1.120383 | 0.3125 | 0.443395 |
| GOBP_TOLL_LIKE_RECEPTOR_2_SIGNALING_PATHWAY | 16 | -0.460594 | -1.120236 | 0.317829 | 0.443452 |
| GOBP_MITOCHONDRIAL_FUSION | 28 | -0.39797 | -1.120127 | 0.317172 | 0.443441 |
| GOBP_LIPID_MODIFICATION | 241 | -0.27987 | -1.119919 | 0.182448 | 0.443583 |
| GOBP_REGULATION_OF_INSULIN_LIKE_GROWTH_FACTOR_RECEPTOR_SIGNALING_PATHW | 16 | -0.450464 | -1.119418 | 0.309478 | 0.444437 |
| GOBP_PYRIMIDINE_NUCLEOTIDE_METABOLIC_PROCESS | 47 | -0.355138 | -1.119376 | 0.270042 | 0.444259 |
| GOBP_INORGANIC_ION_IMPORT_ACROSS_PLASMA_MEMBRANE | 69 | -0.337483 | -1.119185 | 0.265795 | 0.444418 |
| GOBP_POSITIVE_REGULATION_OF_CYTOSOLIC_CALCIUM_ION_CONCENTRATION_INVOLVEI | 24 | -0.407427 | -1.118727 | 0.321932 | 0.445091 |
| GOBP_SNRNA_TRANSCRIPTION | 74 | -0.328855 | -1.118457 | 0.24086 | 0.445399 |
| GOBP_OUTFLOW_TRACT_MORPHOGENESIS | 50 | -0.357326 | -1.118358 | 0.281465 | 0.445357 |
| GOBP_RESPONSE_TO_IONIZING_RADIATION | 134 | -0.301211 | -1.118027 | 0.196507 | 0.445806 |
| GOBP_NEGATIVE_REGULATION_OF_AUTOPHAGOSOME_ASSEMBLY | 15 | -0.453389 | -1.117366 | 0.334702 | 0.44698 |
| GOBP_ANDROGEN_METABOLIC_PROCESS | 22 | -0.416203 | -1.116957 | 0.317992 | 0.447604 |
| GOBP_SECRETORY_GRANULE_ORGANIZATION | 27 | -0.41218 | -1.116887 | 0.297727 | 0.447484 |
| GOBP_HORMONE_MEDIATED_SIGNALING_PATHWAY | 145 | -0.297536 | -1.116879 | 0.196468 | 0.447238 |
| GOBP_REGULATION_OF_OSTEOBLAST_DIFFERENTIATION | 97 | -0.318543 | -1.116793 | 0.227176 | 0.447148 |
| GOBP_POSITIVE_REGULATION_OF_NEURON_DEATH | 83 | -0.324134 | -1.1167 | 0.266075 | 0.447075 |
| GOBP_REGULATION_OF_GRANULOCYTE_CHEMOTAXIS | 44 | -0.364224 | -1.116445 | 0.275424 | 0.44736 |
| GOBP_PROTEIN_K11_LINKED_UBIQUITINATION | 29 | -0.397743 | -1.116422 | 0.287785 | 0.447135 |
| GOBP_GENE_SILENCING | 293 | -0.273773 | -1.116344 | 0.159524 | 0.447061 |
| GOBP_NEGATIVE_REGULATION_OF_INTERLEUKIN_1_PRODUCTION | 29 | -0.394584 | -1.116206 | 0.308489 | 0.447055 |
| GOBP_PROTEIN_LOCALIZATION_TO_SYNAPSE | 65 | -0.333931 | -1.115915 | 0.251656 | 0.447429 |
| GOBP_NEGATIVE_REGULATION_OF_LEUKOCYTE_CELL_CELL_ADHESION | 119 | -0.306034 | -1.115808 | 0.248879 | 0.447374 |
| GOBP_GLUTAMINE_FAMILY_AMINO_ACID_METABOLIC_PROCESS | 62 | -0.342425 | -1.115732 | 0.276639 | 0.447241 |
| GOBP_CELLULAR_RESPONSE_TO_FLUID_SHEAR_STRESS | 16 | -0.436325 | -1.115154 | 0.334831 | 0.448228 |
| GOBP_REGULATION_OF_DEPHOSPHORYLATION | 190 | -0.285758 | -1.115021 | 0.189765 | 0.448251 |
| GOBP_NEGATIVE_REGULATION_OF_LEUKOCYTE_MEDIATED_IMMUNITY | 48 | -0.361827 | -1.114412 | 0.284254 | 0.449281 |
| GOBP_ORGANELLE_MEMBRANE_FUSION | 100 | -0.313571 | -1.113767 | 0.253933 | 0.45043 |
| GOBP_MYOTUBE_CELL_DEVELOPMENT | 23 | -0.411936 | -1.113501 | 0.316917 | 0.450751 |
| GOBP_POSITIVE_REGULATION_OF_MACROPHAGE_MIGRATION | 22 | -0.422447 | -1.113381 | 0.321138 | 0.450742 |
| GOBP_REGULATION_OF_LEUKOCYTE_ADHESION_TO_VASCULAR_ENDOTHELIAL_CELL | 27 | -0.403023 | -1.113334 | 0.279476 | 0.450573 |
| GOBP_INSULIN_LIKE_GROWTH_FACTOR_RECEPTOR_SIGNALING_PATHWAY | 27 | -0.398104 | -1.113243 | 0.292339 | 0.450487 |
| GOBP_BASEMENT_MEMBRANE_ORGANIZATION | 26 | -0.397623 | -1.112847 | 0.29979 | 0.451066 |
| GOBP_REGULATION_OF_TOLL_LIKE_RECEPTOR_SIGNALING_PATHWAY | 66 | -0.331197 | -1.112601 | 0.299776 | 0.45133 |
| GOBP_EPIDERMIS_MORPHOGENESIS | 24 | -0.407968 | -1.112443 | 0.318471 | 0.451414 |
| GOBP_POSITIVE_REGULATION_OF_CD4_POSITIVE_ALPHA_BETA_T_CELL_DIFFERENTIATION | 29 | -0.399531 | -1.112378 | 0.301455 | 0.451276 |
| GOBP_CELLULAR_IRON_ION_HOMEOSTASIS | 60 | -0.343296 | -1.112119 | 0.264108 | 0.451563 |
| GOBP_GASTRIC_ACID_SECRETION | 15 | -0.454698 | -1.111919 | 0.331155 | 0.451698 |
| GOBP_REGULATION_OF_B_CELL_DIFFERENTIATION | 29 | -0.393114 | -1.111541 | 0.303393 | 0.452237 |
| GOBP_NEURON_RECOGNITION | 32 | -0.393135 | -1.110855 | 0.297917 | 0.453467 |
| GOBP_REGULATION_OF_MORPHOGENESIS_OF_AN_EPITHELIUM | 40 | -0.373575 | -1.109571 | 0.292517 | 0.456005 |
| GOBP_CELLULAR_MODIFIED_AMINO_ACID_METABOLIC_PROCESS | 154 | -0.292142 | -1.109531 | 0.208421 | 0.455799 |
| GOBP_REGULATION_OF_MEIOTIC_CELL_CYCLE | 38 | -0.373579 | -1.109476 | 0.307536 | 0.455645 |
| GOBP_GLUCOSE_METABOLIC_PROCESS | 182 | -0.290689 | -1.109323 | 0.212528 | 0.455697 |
| GOBP_ESTROGEN_METABOLIC_PROCESS | 22 | -0.420833 | -1.108452 | 0.305447 | 0.457386 |
| GOBP_IMPORT_INTO_NUCLEUS | 146 | -0.297545 | -1.108329 | 0.236957 | 0.457381 |
| GOBP_CERAMIDE_CATABOLIC_PROCESS | 20 | -0.430283 | -1.108066 | 0.334008 | 0.457698 |
| GOBP_REGULATION_OF_GTPASE_ACTIVITY | 421 | -0.263594 | -1.107964 | 0.131313 | 0.457649 |
| GOBP_REGULATION_OF_CARBOHYDRATE_METABOLIC_PROCESS | 182 | -0.291537 | -1.107485 | 0.228637 | 0.45845 |
| GOBP_CELL_DIFFERENTIATION_INVOLVED_IN_KIDNEY_DEVELOPMENT | 40 | -0.367563 | -1.107189 | 0.318367 | 0.458838 |
| GOBP_POSITIVE_REGULATION_OF_LYMPHOCYTE_MIGRATION | 31 | -0.384699 | -1.106894 | 0.295688 | 0.459174 |
| GOBP_REPLICATIVE_SENESCENCE | 15 | -0.456663 | -1.106817 | 0.342612 | 0.459095 |
| GOBP_CARDIAC_MUSCLE_CONTRACTION | 93 | -0.315156 | -1.106155 | 0.250564 | 0.460299 |
| GOBP_DNA_TEMPLATED_TRANSCRIPTION_INITIATION | 223 | -0.280362 | -1.105784 | 0.207944 | 0.460842 |
| GOBP_POSITIVE_REGULATION_OF_INTERLEUKIN_2_PRODUCTION | 29 | -0.392389 | -1.105245 | 0.305328 | 0.4618 |
| GOBP_REGULATION_OF_CELL_MATRIX_ADHESION | 103 | -0.309124 | -1.105164 | 0.253363 | 0.461723 |
| GOBP_BODY_MORPHOGENESIS | 39 | -0.367696 | -1.104945 | 0.306346 | 0.461943 |
| GOBP_NITRIC_OXIDE_SYNTHASE_BIOSYNTHETIC_PROCESS | 19 | -0.427062 | -1.104882 | 0.33617 | 0.461799 |
| GOBP_POSITIVE_REGULATION_OF_PROTEIN_CONTAINING_COMPLEX_DISASSEMBLY | 31 | -0.387329 | -1.104871 | 0.323974 | 0.461547 |
| GOBP_PROTEIN_DNA_COMPLEX_SUBUNIT_ORGANIZATION | 241 | -0.279134 | -1.10409 | 0.179724 | 0.462954 |
| GOBP_ROOF_OF_MOUTH_DEVELOPMENT | 68 | -0.333413 | -1.104089 | 0.246085 | 0.462683 |
| GOBP_RNA_CAPPING | 34 | -0.383211 | -1.103557 | 0.322034 | 0.463582 |
| GOBP_REGULATION_OF_CELL_MATURATION | 18 | -0.433955 | -1.10279 | 0.330645 | 0.464986 |
| GOBP_REGULATION_OF_RAS_PROTEIN_SIGNAL_TRANSDUCTION | 174 | -0.291994 | -1.102553 | 0.226328 | 0.465231 |
| GOBP_ACTIVATION_OF_IMMUNE_RESPONSE | 492 | -0.260375 | -1.102476 | 0.134409 | 0.465144 |
| GOBP_CHROMOSOME_SEGREGATION | 314 | -0.269659 | -1.102367 | 0.186893 | 0.465128 |
| GOBP_MALE_MEIOTIC_NUCLEAR_DIVISION | 34 | -0.376113 | -1.102364 | 0.287912 | 0.464863 |
| GOBP_RRNA_CATABOLIC_PROCESS | 19 | -0.436361 | -1.10231 | 0.344898 | 0.464708 |
| GOBP_LYMPHOID_PROGENITOR_CELL_DIFFERENTIATION | 18 | -0.438102 | -1.102072 | 0.325431 | 0.464969 |
| GOBP_ICOSANOID_SECRETION | 25 | -0.398639 | -1.10207 | 0.317269 | 0.464698 |
| GOBP_REGULATION_OF_PROTEIN_EXIT_FROM_ENDOPLASMIC_RETICULUM | 26 | -0.392944 | -1.100938 | 0.317125 | 0.466925 |
| GOBP_SUBSTRATE_ADHESION_DEPENDENT_CELL_SPREADING | 94 | -0.3154 | -1.100774 | 0.242553 | 0.466993 |
| GOBP_HORMONE_METABOLIC_PROCESS | 143 | -0.296699 | -1.100696 | 0.233032 | 0.466881 |
| GOBP_ZYMOGEN_ACTIVATION | 42 | -0.361709 | -1.1006 | 0.320879 | 0.466817 |
| GOBP_CYTOPLASMIC_SEQUESTERING_OF_TRANSCRIPTION_FACTOR | 15 | -0.44747 | -1.100356 | 0.338947 | 0.467076 |
| GOBP_PLASMA_MEMBRANE_ORGANIZATION | 97 | -0.314748 | -1.100026 | 0.268966 | 0.467529 |
| GOBP_RESPIRATORY_GASEOUS_EXCHANGE_BY_RESPIRATORY_SYSTEM | 51 | -0.346581 | -1.099832 | 0.2949 | 0.467668 |
| GOBP_POSITIVE_REGULATION_OF_CYSTEINE_TYPE_ENDOPEPTIDASE_ACTIVITY | 135 | -0.296465 | -1.099692 | 0.253247 | 0.467707 |
| GOBP_MICROVILLUS_ORGANIZATION | 20 | -0.422607 | -1.099536 | 0.329876 | 0.467756 |
| GOBP_REGULATION_OF_CYTOPLASMIC_TRANSLATION | 19 | -0.426311 | -1.099466 | 0.346809 | 0.467637 |
| GOBP_PHOSPHATIDYLETHANOLAMINE_ACYL_CHAIN_REMODELING | 17 | -0.451275 | -1.099417 | 0.338776 | 0.467468 |
| GOBP_POSITIVE_REGULATION_OF_VIRAL_TRANSCRIPTION | 26 | -0.4113 | -1.099365 | 0.325956 | 0.467308 |
| GOBP_STEROID_HORMONE_BIOSYNTHETIC_PROCESS | 36 | -0.364374 | -1.09935 | 0.301927 | 0.467064 |
| GOBP_PEPTIDE_SECRETION | 310 | -0.269408 | -1.099277 | 0.172986 | 0.466927 |
| GOBP_REGULATION_OF_OXIDATIVE_STRESS_INDUCED_NEURON_DEATH | 21 | -0.406833 | -1.099121 | 0.331337 | 0.466979 |
| GOBP_PYRIMIDINE_CONTAINING_COMPOUND_BIOSYNTHETIC_PROCESS | 37 | -0.373142 | -1.098857 | 0.297521 | 0.467314 |
| GOBP_SIGNAL_TRANSDUCTION_BY_P53_CLASS_MEDIATOR | 252 | -0.275731 | -1.098852 | 0.196388 | 0.467054 |
| GOBP_REGULATION_OF_CELL_CYCLE_G2_M_PHASE_TRANSITION | 208 | -0.279872 | -1.098724 | 0.208716 | 0.467073 |
| GOBP_PYRUVATE_METABOLIC_PROCESS | 131 | -0.297156 | -1.09843 | 0.227766 | 0.467462 |
| GOBP_POSITIVE_REGULATION_OF_TRIGLYCERIDE_METABOLIC_PROCESS | 17 | -0.438118 | -1.098371 | 0.325203 | 0.467315 |
| GOBP_REGULATION_OF_CHROMATIN_ASSEMBLY_OR_DISASSEMBLY | 24 | -0.419141 | -1.097586 | 0.317328 | 0.468794 |
| GOBP_EPITHELIAL_TUBE_MORPHOGENESIS | 236 | -0.275215 | -1.097459 | 0.210989 | 0.468783 |

| | | | | | |
|---|---|---|---|---|---|
| GOBP_REGULATION_OF_MACROPHAGE_MIGRATION | 36 | -0.372682 | -1.097269 | 0.288934 | 0.468936 |
| GOBP_MUSCLE_HYPERTROPHY | 67 | -0.331165 | -1.097193 | 0.27451 | 0.468831 |
| GOBP_BLOOD_VESSEL_ENDOTHELIAL_CELL_PROLIFERATION_INVOLVED_IN_SPROUTING_ | 21 | -0.418297 | -1.096917 | 0.343874 | 0.469114 |
| GOBP_DEOXYRIBOSE_PHOSPHATE_CATABOLIC_PROCESS | 25 | -0.398056 | -1.096848 | 0.329918 | 0.46902 |
| GOBP_CYTOKINE_PRODUCTION_INVOLVED_IN_INFLAMMATORY_RESPONSE | 48 | -0.347403 | -1.096764 | 0.322654 | 0.46894 |
| GOBP_FOLIC_ACID_CONTAINING_COMPOUND_METABOLIC_PROCESS | 25 | -0.397034 | -1.096234 | 0.329764 | 0.469864 |
| GOBP_NEGATIVE_REGULATION_OF_INTERLEUKIN_2_PRODUCTION | 26 | -0.400589 | -1.09616 | 0.344468 | 0.469771 |
| GOBP_CALCIUM_ION_TRANSMEMBRANE_TRANSPORT | 236 | -0.275345 | -1.095985 | 0.200445 | 0.469866 |
| GOBP_PYRIMIDINE_DEOXYRIBONUCLEOTIDE_METABOLIC_PROCESS | 22 | -0.415215 | -1.09586 | 0.322449 | 0.469876 |
| GOBP_ANATOMICAL_STRUCTURE_HOMEOSTASIS | 391 | -0.262018 | -1.095825 | 0.165865 | 0.469687 |
| GOBP_ATP_SYNTHESIS_COUPLED_ELECTRON_TRANSPORT | 95 | -0.308067 | -1.095493 | 0.288421 | 0.470118 |
| GOBP_NUCLEOBASE_CONTAINING_SMALL_MOLECULE_INTERCONVERSION | 27 | -0.397001 | -1.094764 | 0.309524 | 0.471475 |
| GOBP_POLYSACCHARIDE_BIOSYNTHETIC_PROCESS | 66 | -0.33055 | -1.094583 | 0.300668 | 0.471601 |
| GOBP_REGULATION_OF_RESPONSE_TO_BIOTIC_STIMULUS | 367 | -0.263037 | -1.094318 | 0.182243 | 0.471918 |
| GOBP_VESICLE_TRANSPORT_ALONG_MICROTUBULE | 44 | -0.352026 | -1.094151 | 0.295503 | 0.472024 |
| GOBP_NEGATIVE_REGULATION_OF_DEFENSE_RESPONSE | 176 | -0.28361 | -1.094054 | 0.23356 | 0.471975 |
| GOBP_NEGATIVE_REGULATION_OF_STEM_CELL_DIFFERENTIATION | 18 | -0.440243 | -1.093781 | 0.32981 | 0.472283 |
| GOBP_REGULATION_OF_CARDIOCYTE_DIFFERENTIATION | 19 | -0.434031 | -1.09378 | 0.33125 | 0.472013 |
| GOBP_CELL_KILLING | 136 | -0.292181 | -1.093633 | 0.244444 | 0.472054 |
| GOBP_REGULATION_OF_ADAPTIVE_IMMUNE_RESPONSE | 158 | -0.292729 | -1.093424 | 0.253763 | 0.472225 |
| GOBP_GLYCOSPHINGOLIPID_METABOLIC_PROCESS | 62 | -0.329176 | -1.093164 | 0.302632 | 0.472503 |
| GOBP_REGULATION_OF_MICROTUBULE_CYTOSKELETON_ORGANIZATION | 123 | -0.298107 | -1.092421 | 0.243363 | 0.473822 |
| GOBP_PLACENTA_BLOOD_VESSEL_DEVELOPMENT | 26 | -0.394181 | -1.092362 | 0.313726 | 0.473673 |
| GOBP_PROTEIN_NEDDYLATION | 17 | -0.442254 | -1.092084 | 0.341004 | 0.473998 |
| GOBP_ATP_DEPENDENT_CHROMATIN_REMODELING | 81 | -0.315513 | -1.091828 | 0.298137 | 0.474266 |
| GOBP_STEROL_TRANSPORT | 84 | -0.322542 | -1.091783 | 0.277186 | 0.474103 |
| GOBP_PROTEIN_PALMITOYLATION | 30 | -0.383966 | -1.091532 | 0.312621 | 0.474406 |
| GOBP_CARBOHYDRATE_PHOSPHORYLATION | 22 | -0.409676 | -1.091272 | 0.337054 | 0.47472 |
| GOBP_NEGATIVE_REGULATION_OF_OXIDATIVE_STRESS_INDUCED_INTRINSIC_APOPTOTIC_ | 18 | -0.430493 | -1.090596 | 0.335317 | 0.475953 |
| GOBP_POSITIVE_REGULATION_OF_MRNA_PROCESSING | 27 | -0.392814 | -1.09042 | 0.352475 | 0.476079 |
| GOBP_ESTABLISHMENT_OF_RNA_LOCALIZATION | 186 | -0.286122 | -1.09012 | 0.260181 | 0.476494 |
| GOBP_REGULATION_OF_AUTOPHAGOSOME_ASSEMBLY | 39 | -0.372689 | -1.090026 | 0.318083 | 0.476438 |
| GOBP_PANCREAS_DEVELOPMENT | 47 | -0.353737 | -1.089421 | 0.298643 | 0.477524 |
| GOBP_ALPHA_BETA_T_CELL_DIFFERENTIATION | 98 | -0.306437 | -1.089387 | 0.283447 | 0.477326 |
| GOBP_NEUTRAL_LIPID_BIOSYNTHETIC_PROCESS | 38 | -0.363786 | -1.088937 | 0.319915 | 0.4781 |
| GOBP_MATURE_B_CELL_DIFFERENTIATION | 30 | -0.386953 | -1.088886 | 0.33121 | 0.477941 |
| GOBP_CYTOSOLIC_TRANSPORT | 159 | -0.28574 | -1.088806 | 0.259508 | 0.47784 |
| GOBP_STEROID_METABOLIC_PROCESS | 235 | -0.277836 | -1.088622 | 0.217284 | 0.477975 |
| GOBP_PIGMENT_METABOLIC_PROCESS | 58 | -0.326631 | -1.088572 | 0.321429 | 0.477834 |
| GOBP_LIPOSACCHARIDE_METABOLIC_PROCESS | 100 | -0.309475 | -1.088249 | 0.267416 | 0.478278 |
| GOBP_PYRIMIDINE_NUCLEOTIDE_CATABOLIC_PROCESS | 19 | -0.429943 | -1.088106 | 0.348178 | 0.478351 |
| GOBP_T_HELPER_17_TYPE_IMMUNE_RESPONSE | 30 | -0.380291 | -1.0875 | 0.331224 | 0.479441 |
| GOBP_RESPONSE_TO_BMP | 130 | -0.298169 | -1.086885 | 0.252796 | 0.480538 |
| GOBP_POSITIVE_REGULATION_OF_EXTRACELLULAR_MATRIX_ORGANIZATION | 19 | -0.418922 | -1.086312 | 0.341463 | 0.481576 |
| GOBP_CELLULAR_RESPONSE_TO_HEAT | 111 | -0.304651 | -1.086264 | 0.270325 | 0.481411 |
| GOBP_REGULATION_OF_PROTEIN_DEPOLYMERIZATION | 72 | -0.319536 | -1.085638 | 0.296943 | 0.482537 |
| GOBP_LATE_ENDOSOME_TO_VACUOLE_TRANSPORT_VIA_MULTIVESICULAR_BODY_SORTII | 17 | -0.432405 | -1.085339 | 0.346405 | 0.482942 |
| GOBP_REGULATION_OF_CELLULAR_RESPIRATION | 20 | -0.41795 | -1.083723 | 0.342159 | 0.486346 |
| GOBP_SPECIFICATION_OF_SYMMETRY | 93 | -0.310836 | -1.083584 | 0.315098 | 0.486377 |
| GOBP_REGULATION_OF_PLATELET_ACTIVATION | 29 | -0.38431 | -1.083228 | 0.353659 | 0.486908 |
| GOBP_CORONARY_VASCULATURE_DEVELOPMENT | 37 | -0.365763 | -1.082514 | 0.341463 | 0.488214 |
| GOBP_REGULATION_OF_INNATE_IMMUNE_RESPONSE | 279 | -0.268321 | -1.082044 | 0.213152 | 0.489015 |
| GOBP_REGULATION_OF_CALCIUM_ION_TRANSMEMBRANE_TRANSPORTER_ACTIVITY | 61 | -0.328341 | -1.081664 | 0.316562 | 0.489602 |
| GOBP_TRANSPORT_ALONG_MICROTUBULE | 151 | -0.290766 | -1.081635 | 0.233184 | 0.489384 |
| GOBP_GERM_CELL_DEVELOPMENT | 182 | -0.280448 | -1.08143 | 0.273138 | 0.489541 |
| GOBP_RESPONSE_TO_CALCIUM_ION | 108 | -0.301903 | -1.081361 | 0.27533 | 0.489423 |
| GOBP_REGULATION_OF_SYNAPTIC_VESICLE_EXOCYTOSIS | 46 | -0.349551 | -1.081213 | 0.336756 | 0.489481 |
| GOBP_PROTEIN_CONTAINING_COMPLEX_DISASSEMBLY | 303 | -0.26895 | -1.081068 | 0.219048 | 0.489531 |
| GOBP_LONG_CHAIN_FATTY_ACID_TRANSPORT | 53 | -0.339996 | -1.079643 | 0.328571 | 0.492457 |
| GOBP_NEGATIVE_REGULATION_OF_SUPRAMOLECULAR_FIBER_ORGANIZATION | 135 | -0.295394 | -1.079552 | 0.277647 | 0.492385 |
| GOBP_POSITIVE_REGULATION_OF_POTASSIUM_ION_TRANSPORT | 34 | -0.363718 | -1.079263 | 0.325843 | 0.492779 |
| GOBP_ENDOCRINE_HORMONE_SECRETION | 36 | -0.370727 | -1.079169 | 0.336864 | 0.492724 |
| GOBP_NEGATIVE_REGULATION_OF_NERVOUS_SYSTEM_DEVELOPMENT | 102 | -0.305156 | -1.079072 | 0.301518 | 0.492671 |
| GOBP_CHLORIDE_TRANSPORT | 69 | -0.318663 | -1.078857 | 0.283517 | 0.492872 |
| GOBP_CARDIAC_VENTRICLE_DEVELOPMENT | 78 | -0.315784 | -1.078662 | 0.290541 | 0.493054 |
| GOBP_ENDOTHELIUM_DEVELOPMENT | 104 | -0.298547 | -1.078624 | 0.292906 | 0.49287 |
| GOBP_REGULATION_OF_CELLULAR_RESPONSE_TO_INSULIN_STIMULUS | 57 | -0.335367 | -1.078473 | 0.318872 | 0.492918 |
| GOBP_REGULATION_OF_ENDOTHELIAL_CELL_DIFFERENTIATION | 29 | -0.393392 | -1.078469 | 0.346154 | 0.492655 |
| GOBP_RESPONSE_TO_NICOTINE | 31 | -0.373754 | -1.078163 | 0.334061 | 0.493069 |
| GOBP_VASCULAR_ENDOTHELIAL_GROWTH_FACTOR_RECEPTOR_SIGNALING_PATHWAY | 84 | -0.306888 | -1.077705 | 0.31441 | 0.493867 |
| GOBP_CHROMOSOME_LOCALIZATION | 75 | -0.319571 | -1.077669 | 0.333333 | 0.493664 |
| GOBP_SALIVARY_GLAND_DEVELOPMENT | 26 | -0.393781 | -1.07746 | 0.363071 | 0.493838 |
| GOBP_REGULATION_OF_GENE_SILENCING_BY_RNA | 108 | -0.305633 | -1.076807 | 0.283224 | 0.495103 |
| GOBP_REGULATION_OF_MONOOXYGENASE_ACTIVITY | 51 | -0.340788 | -1.076752 | 0.307856 | 0.494936 |
| GOBP_POSITIVE_REGULATION_OF_PHOSPHOPROTEIN_PHOSPHATASE_ACTIVITY | 19 | -0.415493 | -1.076683 | 0.378378 | 0.494812 |
| GOBP_CILIARY_BASAL_BODY_PLASMA_MEMBRANE_DOCKING | 94 | -0.307341 | -1.076478 | 0.312073 | 0.495027 |
| GOBP_TUBE_FORMATION | 115 | -0.295469 | -1.07634 | 0.304825 | 0.495034 |
| GOBP_PROTEIN_DEGLYCOSYLATION | 28 | -0.37854 | -1.075422 | 0.339169 | 0.496898 |
| GOBP_POSITIVE_REGULATION_OF_WNT_SIGNALING_PATHWAY | 159 | -0.280155 | -1.074967 | 0.276923 | 0.497629 |
| GOBP_POSITIVE_REGULATION_OF_TRANSLATIONAL_INITIATION | 24 | -0.393761 | -1.07493 | 0.350951 | 0.497446 |
| GOBP_REGULATION_OF_CELL_MORPHOGENESIS | 270 | -0.267203 | -1.073892 | 0.251163 | 0.499574 |
| GOBP_AORTA_DEVELOPMENT | 44 | -0.347616 | -1.073805 | 0.356574 | 0.499482 |
| GOBP_NEGATIVE_REGULATION_OF_OXIDATIVE_STRESS_INDUCED_CELL_DEATH | 35 | -0.365476 | -1.073709 | 0.343035 | 0.499441 |
| GOBP_ELECTRON_TRANSPORT_CHAIN | 159 | -0.287749 | -1.072865 | 0.286638 | 0.501097 |
| GOBP_REGULATION_OF_TRIGLYCERIDE_METABOLIC_PROCESS | 28 | -0.384881 | -1.072667 | 0.336083 | 0.501281 |
| GOBP_POSITIVE_REGULATION_OF_CANONICAL_WNT_SIGNALING_PATHWAY | 131 | -0.291099 | -1.072549 | 0.276087 | 0.501289 |
| GOBP_B_CELL_APOPTOTIC_PROCESS | 25 | -0.391083 | -1.072395 | 0.369388 | 0.501365 |
| GOBP_REGULATION_OF_LYMPHOCYTE_CHEMOTAXIS | 20 | -0.415852 | -1.072315 | 0.389002 | 0.501261 |
| GOBP_NEGATIVE_REGULATION_OF_RHO_PROTEIN_SIGNAL_TRANSDUCTION | 19 | -0.413879 | -1.072245 | 0.348684 | 0.501138 |
| GOBP_UNSATURATED_FATTY_ACID_BIOSYNTHETIC_PROCESS | 42 | -0.351735 | -1.071583 | 0.336694 | 0.502325 |
| GOBP_REGULATION_OF_DEFENSE_RESPONSE_TO_VIRUS_BY_HOST | 37 | -0.356446 | -1.071496 | 0.345572 | 0.502262 |
| GOBP_RECOMBINATIONAL_REPAIR | 135 | -0.2922 | -1.071466 | 0.294248 | 0.502053 |
| GOBP_REGULATION_OF_PROTEIN_POLYMERIZATION | 197 | -0.275789 | -1.071369 | 0.243119 | 0.502024 |
| GOBP_POSITIVE_REGULATION_OF_EPITHELIAL_CELL_MIGRATION | 134 | -0.285423 | -1.071133 | 0.294118 | 0.502275 |
| GOBP_PTERIDINE_CONTAINING_COMPOUND_METABOLIC_PROCESS | 31 | -0.374227 | -1.070983 | 0.359408 | 0.502336 |
| GOBP_CELLULAR_RESPONSE_TO_LIGHT_STIMULUS | 108 | -0.301993 | -1.070601 | 0.310195 | 0.502927 |
| GOBP_NEGATIVE_REGULATION_OF_IMMUNE_RESPONSE | 124 | -0.291274 | -1.07055 | 0.284404 | 0.502776 |
| GOBP_RESPONSE_TO_MUSCLE_STRETCH | 18 | -0.418521 | -1.070479 | 0.383673 | 0.502647 |
| GOBP_HISTONE_H3_K4_METHYLATION | 49 | -0.346847 | -1.070405 | 0.332613 | 0.502547 |
| GOBP_HINDBRAIN_DEVELOPMENT | 102 | -0.303795 | -1.070322 | 0.306488 | 0.502475 |
| GOBP_CRANIAL_SKELETAL_SYSTEM_DEVELOPMENT | 46 | -0.346512 | -1.070043 | 0.32996 | 0.502808 |
| GOBP_CEREBELLAR_PURKINJE_CELL_LAYER_DEVELOPMENT | 18 | -0.424249 | -1.069894 | 0.357143 | 0.50286 |
| GOBP_REGULATION_OF_CELLULAR_PROTEIN_CATABOLIC_PROCESS | 231 | -0.271133 | -1.069667 | 0.266979 | 0.50313 |
| GOBP_INORGANIC_ANION_TRANSMEMBRANE_TRANSPORT | 77 | -0.31656 | -1.069627 | 0.326316 | 0.502937 |
| GOBP_POSITIVE_REGULATION_OF_STEROID_METABOLIC_PROCESS | 16 | -0.433374 | -1.069474 | 0.365112 | 0.503008 |
| GOBP_POSITIVE_REGULATION_OF_DENDRITIC_SPINE_MORPHOGENESIS | 18 | -0.420973 | -1.069101 | 0.353659 | 0.503568 |
| GOBP_ENDOPLASMIC_RETICULUM_TO_CYTOSOL_TRANSPORT | 28 | -0.388269 | -1.068796 | 0.366316 | 0.503981 |
| GOBP_CELLULAR_LIPID_CATABOLIC_PROCESS | 188 | -0.276675 | -1.068763 | 0.269841 | 0.503785 |
| GOBP_POSITIVE_REGULATION_OF_PROTEIN_DEACETYLATION | 20 | -0.414165 | -1.068676 | 0.396146 | 0.503719 |
| GOBP_NUCLEOTIDE_EXCISION_REPAIR_DNA_GAP_FILLING | 23 | -0.401263 | -1.067941 | 0.395299 | 0.505079 |
| GOBP_ERROR_FREE_TRANSLESION_SYNTHESIS | 22 | -0.403502 | -1.067526 | 0.373444 | 0.505724 |
| GOBP_EPITHELIAL_CELL_DIFFERENTIATION_INVOLVED_IN_KIDNEY_DEVELOPMENT | 33 | -0.377578 | -1.067464 | 0.371179 | 0.505596 |
| GOBP_NEGATIVE_REGULATION_OF_ESTABLISHMENT_OF_PROTEIN_LOCALIZATION | 102 | -0.304018 | -1.066988 | 0.297872 | 0.506436 |
| GOBP_REGULATION_OF_SMALL_MOLECULE_METABOLIC_PROCESS | 371 | -0.259157 | -1.066452 | 0.241784 | 0.507406 |
| GOBP_NEGATIVE_REGULATION_OF_POTASSIUM_ION_TRANSPORT | 22 | -0.40751 | -1.066222 | 0.367615 | 0.507645 |
| GOBP_HEAD_MORPHOGENESIS | 27 | -0.388169 | -1.066002 | 0.376299 | 0.507936 |
| GOBP_GLUTAMINE_FAMILY_AMINO_ACID_BIOSYNTHETIC_PROCESS | 16 | -0.437722 | -1.065992 | 0.376448 | 0.507686 |
| GOBP_NEGATIVE_REGULATION_OF_CYCLIN_DEPENDENT_PROTEIN_KINASE_ACTIVITY | 29 | -0.380179 | -1.065944 | 0.366316 | 0.507526 |
| GOBP_MEIOTIC_CHROMOSOME_SEPARATION | 21 | -0.408989 | -1.065669 | 0.37037 | 0.507888 |
| GOBP_NEGATIVE_REGULATION_OF_PROTEIN_LOCALIZATION_TO_NUCLEUS | 27 | -0.387007 | -1.065661 | 0.340909 | 0.507626 |
| GOBP_CYTOSKELETON_DEPENDENT_CYTOKINESIS | 93 | -0.305372 | -1.065066 | 0.312088 | 0.508703 |
| GOBP_IN_UTERO_EMBRYONIC_DEVELOPMENT | 296 | -0.263466 | -1.064282 | 0.270396 | 0.510244 |
| GOBP_ARTERY_MORPHOGENESIS | 43 | -0.350048 | -1.064035 | 0.341102 | 0.510565 |
| GOBP_RAS_PROTEIN_SIGNAL_TRANSDUCTION | 312 | -0.261766 | -1.063755 | 0.261845 | 0.510942 |
| GOBP_NUCLEOSIDE_PHOSPHATE_BIOSYNTHETIC_PROCESS | 240 | -0.26743 | -1.063745 | 0.271264 | 0.510691 |
| GOBP_RESPONSE_TO_GROWTH_HORMONE | 33 | -0.359537 | -1.063614 | 0.331887 | 0.510701 |
| GOBP_RNA_LOCALIZATION | 219 | -0.268272 | -1.06324 | 0.288372 | 0.511254 |
| GOBP_MEMBRANE_PROTEIN_INTRACELLULAR_DOMAIN_PROTEOLYSIS | 20 | -0.417483 | -1.062906 | 0.364629 | 0.511719 |
| GOBP_DNA_TEMPLATED_TRANSCRIPTION_ELONGATION | 111 | -0.294392 | -1.062616 | 0.325103 | 0.512101 |
| GOBP_SKELETAL_MUSCLE_CELL_DIFFERENTIATION | 47 | -0.3339 | -1.062339 | 0.364979 | 0.512497 |
| GOBP_NEGATIVE_REGULATION_OF_BIOMINERALIZATION | 22 | -0.396127 | -1.062263 | 0.365503 | 0.512408 |
| GOBP_POSITIVE_REGULATION_OF_AMYLOID_PRECURSOR_PROTEIN_CATABOLIC_PROCES | 22 | -0.39689 | -1.061815 | 0.361111 | 0.513127 |
| GOBP_PHOSPHATIDYLCHOLINE_BIOSYNTHETIC_PROCESS | 37 | -0.358381 | -1.061666 | 0.362069 | 0.513191 |
| GOBP_NUCLEOSIDE_METABOLIC_PROCESS | 97 | -0.309187 | -1.061299 | 0.331096 | 0.513741 |
| GOBP_REGULATION_OF_RESPONSE_TO_ENDOPLASMIC_RETICULUM_STRESS | 79 | -0.311786 | -1.061167 | 0.325726 | 0.513787 |
| GOBP_FLUID_TRANSPORT | 22 | -0.398196 | -1.060835 | 0.383505 | 0.514233 |
| GOBP_REGULATION_OF_PEPTIDE_TRANSPORT | 484 | -0.248449 | -1.060377 | 0.229219 | 0.515019 |
| GOBP_FORELIMB_MORPHOGENESIS | 27 | -0.390753 | -1.060354 | 0.372671 | 0.514797 |
| GOBP_STRIATED_MUSCLE_CELL_DEVELOPMENT | 73 | -0.310889 | -1.060287 | 0.338057 | 0.514687 |
| GOBP_POSITIVE_REGULATION_OF_FIBROBLAST_PROLIFERATION | 40 | -0.350913 | -1.060262 | 0.371369 | 0.51447 |
| GOBP_DEPHOSPHORYLATION | 435 | -0.24947 | -1.060099 | 0.253886 | 0.514593 |
| GOBP_NEGATIVE_REGULATION_OF_VIRAL_ENTRY_INTO_HOST_CELL | 16 | -0.425859 | -1.059703 | 0.379098 | 0.515234 |
| GOBP_RESPONSE_TO_ESTROGEN | 60 | -0.323436 | -1.059611 | 0.337662 | 0.515169 |
| GOBP_ENDOSOME_TRANSPORT_VIA_MULTIVESICULAR_BODY_SORTING_PATHWAY | 31 | -0.370678 | -1.059395 | 0.354167 | 0.515388 |
| GOBP_ICOSANOID_TRANSPORT | 33 | -0.362566 | -1.059279 | 0.340611 | 0.515382 |
| GOBP_REGULATION_OF_CYTOSKELETON_ORGANIZATION | 444 | -0.252532 | -1.059246 | 0.246851 | 0.515165 |
| GOBP_CELLULAR_KETONE_METABOLIC_PROCESS | 206 | -0.269777 | -1.059231 | 0.28246 | 0.514925 |
| GOBP_CD40_SIGNALING_PATHWAY | 15 | -0.43479 | -1.059151 | 0.383117 | 0.514848 |
| GOBP_ESTABLISHMENT_OF_ENDOTHELIAL_BARRIER | 44 | -0.351083 | -1.058938 | 0.344828 | 0.515063 |
| GOBP_CHAPERONE_MEDIATED_PROTEIN_COMPLEX_ASSEMBLY | 20 | -0.415302 | -1.058402 | 0.409292 | 0.516048 |
| GOBP_ENDOCARDIAL_CUSHION_DEVELOPMENT | 31 | -0.370768 | -1.05832 | 0.385439 | 0.515962 |
| GOBP_RESPONSE_TO_INTERLEUKIN_7 | 34 | -0.364861 | -1.058234 | 0.384615 | 0.515861 |
| GOBP_VITAMIN_METABOLIC_PROCESS | 105 | -0.293948 | -1.057734 | 0.339667 | 0.516735 |
| GOBP_POSITIVE_REGULATION_OF_ESTABLISHMENT_OF_PROTEIN_LOCALIZATION | 288 | -0.259761 | -1.057688 | 0.27 | 0.516571 |
| GOBP_SMOOTH_MUSCLE_CELL_DIFFERENTIATION | 50 | -0.336139 | -1.057331 | 0.351779 | 0.517151 |
| GOBP_EPITHELIAL_CELL_APOPTOTIC_PROCESS | 92 | -0.305785 | -1.057316 | 0.343374 | 0.51691 |
| GOBP_REGULATION_OF_CELLULAR_PROTEIN_LOCALIZATION | 492 | -0.248747 | -1.057122 | 0.250597 | 0.517088 |

| | | | | | |
|---|---|---|---|---|---|
| GOBP_REGULATION_OF_ER_TO_GOLGI_VESICLE_MEDIATED_TRANSPORT | 15 | -0.443161 | -1.056964 | 0.393004 | 0.517186 |
| GOBP_NEUTRAL_LIPID_METABOLIC_PROCESS | 94 | -0.302747 | -1.056877 | 0.335556 | 0.517137 |
| GOBP_CELLULAR_RESPONSE_TO_ALKALOID | 22 | -0.398015 | -1.056385 | 0.39413 | 0.518016 |
| GOBP_PROSTANOID_BIOSYNTHETIC_PROCESS | 24 | -0.397306 | -1.05626 | 0.390192 | 0.51803 |
| GOBP_POSITIVE_REGULATION_OF_ENDOTHELIAL_CELL_PROLIFERATION | 76 | -0.310507 | -1.055881 | 0.331897 | 0.518602 |
| GOBP_POSITIVE_REGULATION_OF_TRANSCRIPTION_FROM_RNA_POLYMERASE_II_PROMOT | 20 | -0.405651 | -1.055726 | 0.384615 | 0.518675 |
| GOBP_ESTABLISHMENT_OF_MITOCHONDRION_LOCALIZATION | 23 | -0.397605 | -1.055608 | 0.403397 | 0.518682 |
| GOBP_MUSCLE_ORGAN_DEVELOPMENT | 239 | -0.266892 | -1.055518 | 0.304556 | 0.518606 |
| GOBP_MAST_CELL_ACTIVATION | 54 | -0.333994 | -1.055205 | 0.365904 | 0.519052 |
| GOBP_NUCLEOSIDE_TRIPHOSPHATE_BIOSYNTHETIC_PROCESS | 81 | -0.309951 | -1.055085 | 0.332618 | 0.519064 |
| GOBP_RESPONSE_TO_MINERALOCORTICOID | 24 | -0.396132 | -1.054955 | 0.37422 | 0.519109 |
| GOBP_NEGATIVE_REGULATION_OF_HISTONE_MODIFICATION | 41 | -0.346331 | -1.054407 | 0.359743 | 0.520079 |
| GOBP_REGULATION_OF_HEART_GROWTH | 56 | -0.32621 | -1.053649 | 0.370763 | 0.521567 |
| GOBP_EPITHELIAL_TUBE_FORMATION | 102 | -0.294758 | -1.053608 | 0.305618 | 0.521383 |
| GOBP_SMOOTH_MUSCLE_CELL_APOPTOTIC_PROCESS | 21 | -0.406962 | -1.053475 | 0.384306 | 0.521434 |
| GOBP_ORGANOPHOSPHATE_ESTER_TRANSPORT | 111 | -0.29148 | -1.052898 | 0.357466 | 0.522473 |
| GOBP_PYRIMIDINE_NUCLEOTIDE_BIOSYNTHETIC_PROCESS | 27 | -0.375507 | -1.052752 | 0.371622 | 0.522591 |
| GOBP_SPHINGOLIPID_BIOSYNTHETIC_PROCESS | 102 | -0.296928 | -1.052692 | 0.343478 | 0.522459 |
| GOBP_REGULATION_OF_PROTON_TRANSPORT | 22 | -0.398027 | -1.052645 | 0.404612 | 0.522287 |
| GOBP_DIENCEPHALON_DEVELOPMENT | 36 | -0.355677 | -1.051759 | 0.387295 | 0.524119 |
| GOBP_FEMALE_GAMETE_GENERATION | 102 | -0.297508 | -1.051667 | 0.326797 | 0.524053 |
| GOBP_MEMBRANE_LIPID_BIOSYNTHETIC_PROCESS | 136 | -0.27971 | -1.051309 | 0.353982 | 0.524574 |
| GOBP_NOTCH_SIGNALING_PATHWAY | 155 | -0.278566 | -1.051006 | 0.309051 | 0.525027 |
| GOBP_POSITIVE_REGULATION_OF_VIRAL_LIFE_CYCLE | 31 | -0.372698 | -1.05098 | 0.373984 | 0.524813 |
| GOBP_PEPTIDYL_CYSTEINE_MODIFICATION | 43 | -0.340052 | -1.050629 | 0.365344 | 0.525388 |
| GOBP_REGULATION_OF_VASCULAR_ASSOCIATED_SMOOTH_MUSCLE_CELL_DIFFERENTIAT | 15 | -0.439016 | -1.050289 | 0.365751 | 0.525933 |
| GOBP_FATTY_ACID_METABOLIC_PROCESS | 313 | -0.25664 | -1.050129 | 0.273399 | 0.526019 |
| GOBP_ENDOTHELIAL_CELL_DEVELOPMENT | 59 | -0.326399 | -1.050113 | 0.366516 | 0.52578 |
| GOBP_I_KAPPAB_KINASE_NF_KAPPAB_SIGNALING | 261 | -0.261057 | -1.049759 | 0.319202 | 0.526335 |
| GOBP_CELLULAR_RESPONSE_TO_NUTRIENT | 33 | -0.367439 | -1.049601 | 0.362705 | 0.526434 |
| GOBP_NEURON_CELLULAR_HOMEOSTASIS | 23 | -0.385166 | -1.049572 | 0.397149 | 0.526229 |
| GOBP_ALPHA_BETA_T_CELL_ACTIVATION | 134 | -0.285783 | -1.049485 | 0.313063 | 0.526175 |
| GOBP_POSITIVE_REGULATION_OF_PROTEIN_EXPORT_FROM_NUCLEUS | 20 | -0.411778 | -1.048995 | 0.40501 | 0.527015 |
| GOBP_REGULATION_OF_EPITHELIAL_CELL_APOPTOTIC_PROCESS | 71 | -0.313672 | -1.048799 | 0.343413 | 0.527201 |
| GOBP_MAINTENANCE_OF_CELL_NUMBER | 125 | -0.285401 | -1.04787 | 0.349558 | 0.529096 |
| GOBP_RESPONSE_TO_ARSENIC_CONTAINING_SUBSTANCE | 29 | -0.371441 | -1.047631 | 0.392405 | 0.529353 |
| GOBP_GLANDULAR_EPITHELIAL_CELL_DIFFERENTIATION | 31 | -0.363116 | -1.047605 | 0.369658 | 0.529137 |
| GOBP_NEGATIVE_REGULATION_OF_CHROMATIN_ORGANIZATION | 56 | -0.322403 | -1.04735 | 0.359081 | 0.529492 |
| GOBP_ANIMAL_ORGAN_REGENERATION | 65 | -0.31247 | -1.047259 | 0.37694 | 0.529429 |
| GOBP_SMALL_GTPASE_MEDIATED_SIGNAL_TRANSDUCTION | 466 | -0.246593 | -1.047184 | 0.282443 | 0.529343 |
| GOBP_CELL_AGING | 97 | -0.296646 | -1.047166 | 0.352818 | 0.529118 |
| GOBP_REGULATION_OF_PATTERN_RECOGNITION_RECEPTOR_SIGNALING_PATHWAY | 93 | -0.304003 | -1.045326 | 0.3878 | 0.53314 |
| GOBP_RESPONSE_TO_LIGHT_STIMULUS | 262 | -0.259241 | -1.045305 | 0.315909 | 0.53291 |
| GOBP_REGULATION_OF_CELL_ADHESION_MEDIATED_BY_INTEGRIN | 43 | -0.339291 | -1.045289 | 0.365546 | 0.532684 |
| GOBP_NEGATIVE_REGULATION_OF_DNA_BIOSYNTHETIC_PROCESS | 34 | -0.355486 | -1.044791 | 0.388186 | 0.533602 |
| GOBP_REGULATION_OF_POLYSACCHARIDE_METABOLIC_PROCESS | 39 | -0.349828 | -1.044642 | 0.382479 | 0.533663 |
| GOBP_CELLULAR_HORMONE_METABOLIC_PROCESS | 85 | -0.308341 | -1.043815 | 0.37395 | 0.535336 |
| GOBP_REGULATION_OF_WNT_SIGNALING_PATHWAY | 314 | -0.256661 | -1.043695 | 0.31738 | 0.535349 |
| GOBP_SENSORY_PERCEPTION_OF_TEMPERATURE_STIMULUS | 15 | -0.424538 | -1.043409 | 0.40678 | 0.535716 |
| GOBP_PROTEIN_LOCALIZATION_TO_CELL_JUNCTION | 92 | -0.300465 | -1.043276 | 0.375796 | 0.535723 |
| GOBP_REGULATION_OF_HISTONE_DEACETYLATION | 24 | -0.380182 | -1.043075 | 0.375 | 0.535917 |
| GOBP_REGULATION_OF_PROTEIN_CONTAINING_COMPLEX_ASSEMBLY | 393 | -0.251062 | -1.04296 | 0.301932 | 0.535926 |
| GOBP_POSITIVE_REGULATION_OF_DENDRITE_MORPHOGENESIS | 32 | -0.364354 | -1.042863 | 0.409871 | 0.535864 |
| GOBP_INTRINSIC_APOPTOTIC_SIGNALING_PATHWAY_IN_RESPONSE_TO_OXIDATIVE_STRES | 40 | -0.346616 | -1.042543 | 0.386037 | 0.536346 |
| GOBP_CHROMATIN_REMODELING | 190 | -0.267811 | -1.042514 | 0.350446 | 0.536143 |
| GOBP_REGULATION_OF_MEMBRANE_LIPID_DISTRIBUTION | 46 | -0.3361 | -1.042314 | 0.385093 | 0.536316 |
| GOBP_RESPONSE_TO_ELECTRICAL_STIMULUS | 34 | -0.357549 | -1.042211 | 0.366803 | 0.536298 |
| GOBP_NEURON_DEATH | 288 | -0.257715 | -1.041784 | 0.32093 | 0.537071 |
| GOBP_REGULATION_OF_PROTEIN_LOCALIZATION_TO_NUCLEUS | 120 | -0.290237 | -1.041214 | 0.363409 | 0.538118 |
| GOBP_RNA_PHOSPHODIESTER_BOND_HYDROLYSIS_EXONUCLEOLYTIC | 42 | -0.339585 | -1.041039 | 0.384106 | 0.538246 |
| GOBP_CYTOPLASMIC_SEQUESTERING_OF_PROTEIN | 21 | -0.393964 | -1.039878 | 0.400794 | 0.540696 |
| GOBP_REGULATION_OF_HOMOTYPIC_CELL_CELL_ADHESION | 24 | -0.392675 | -1.039567 | 0.419355 | 0.541153 |
| GOBP_RESPONSE_TO_DOPAMINE | 51 | -0.328919 | -1.039116 | 0.349174 | 0.541978 |
| GOBP_NEGATIVE_REGULATION_OF_INTERLEUKIN_10_PRODUCTION | 18 | -0.413401 | -1.037646 | 0.395349 | 0.545176 |
| GOBP_REGULATION_OF_LIPID_BIOSYNTHETIC_PROCESS | 156 | -0.27515 | -1.037607 | 0.347032 | 0.54499 |
| GOBP_POSITIVE_REGULATION_OF_GLIAL_CELL_DIFFERENTIATION | 34 | -0.354075 | -1.037602 | 0.403433 | 0.54472 |
| GOBP_REGULATION_OF_NEURON_PROJECTION_REGENERATION | 26 | -0.376139 | -1.037104 | 0.372745 | 0.545591 |
| GOBP_REGULATION_OF_CD4_POSITIVE_ALPHA_BETA_T_CELL_DIFFERENTIATION | 48 | -0.337556 | -1.037049 | 0.367304 | 0.545447 |
| GOBP_IRON_ION_TRANSMEMBRANE_TRANSPORT | 18 | -0.412746 | -1.036832 | 0.429769 | 0.54572 |
| GOBP_REGULATION_OF_SKELETAL_MUSCLE_TISSUE_DEVELOPMENT | 17 | -0.409641 | -1.035672 | 0.420619 | 0.548157 |
| GOBP_SPINDLE_LOCALIZATION | 48 | -0.33272 | -1.035644 | 0.404661 | 0.547945 |
| GOBP_OOCYTE_DIFFERENTIATION | 39 | -0.345169 | -1.035546 | 0.400853 | 0.547904 |
| GOBP_REGULATION_OF_PRODUCTION_OF_SMALL_RNA_INVOLVED_IN_GENE_SILENCING_I | 20 | -0.405001 | -1.035181 | 0.421359 | 0.548504 |
| GOBP_POSITIVE_REGULATION_OF_PROTEIN_CONTAINING_COMPLEX_ASSEMBLY | 225 | -0.259689 | -1.035138 | 0.331126 | 0.548331 |
| GOBP_LIPID_TRANSLOCATION | 41 | -0.343199 | -1.03483 | 0.388548 | 0.548798 |
| GOBP_REGULATION_OF_MRNA_METABOLIC_PROCESS | 303 | -0.256339 | -1.034517 | 0.323672 | 0.54925 |
| GOBP_REGULATION_OF_CILIUM_ASSEMBLY | 50 | -0.328139 | -1.034093 | 0.405462 | 0.549928 |
| GOBP_PROTEIN_IMPORT | 176 | -0.26933 | -1.033113 | 0.366359 | 0.551957 |
| GOBP_REGULATION_OF_ANDROGEN_RECEPTOR_SIGNALING_PATHWAY | 22 | -0.407851 | -1.032881 | 0.424242 | 0.552228 |
| GOBP_REGULATION_OF_SPHINGOLIPID_BIOSYNTHETIC_PROCESS | 17 | -0.410863 | -1.032808 | 0.434322 | 0.55212 |
| GOBP_ENDOSOMAL_TRANSPORT | 233 | -0.261982 | -1.032758 | 0.37296 | 0.551957 |
| GOBP_PHOSPHATIDYLCHOLINE_METABOLIC_PROCESS | 73 | -0.304501 | -1.032216 | 0.397895 | 0.55299 |
| GOBP_NEGATIVE_REGULATION_OF_BLOOD_VESSEL_ENDOTHELIAL_CELL_MIGRATION | 31 | -0.353903 | -1.032103 | 0.415385 | 0.552982 |
| GOBP_MUSCLE_ORGAN_MORPHOGENESIS | 47 | -0.324162 | -1.03144 | 0.406504 | 0.554358 |
| GOBP_NEGATIVE_REGULATION_OF_CALCIUM_ION_TRANSPORT | 41 | -0.346677 | -1.031359 | 0.395745 | 0.554275 |
| GOBP_BONE_RESORPTION | 50 | -0.321717 | -1.031205 | 0.4 | 0.554371 |
| GOBP_NEGATIVE_REGULATION_OF_DOUBLE_STRAND_BREAK_REPAIR_VIA_HOMOLOGOUS | 19 | -0.400796 | -1.030812 | 0.410101 | 0.55501 |
| GOBP_REGULATION_OF_CARBOHYDRATE_BIOSYNTHETIC_PROCESS | 83 | -0.298295 | -1.030703 | 0.395299 | 0.554985 |
| GOBP_POSITIVE_REGULATION_OF_DNA_REPLICATION | 36 | -0.339825 | -1.029687 | 0.421286 | 0.557112 |
| GOBP_REGULATION_OF_CALCIUM_ION_IMPORT | 25 | -0.383864 | -1.028119 | 0.421842 | 0.560635 |
| GOBP_NUCLEAR_TRANSCRIBED_MRNA_CATABOLIC_PROCESS_DEADENYLATION_DEPEND | 75 | -0.305018 | -1.027892 | 0.396104 | 0.560922 |
| GOBP_POSITIVE_REGULATION_OF_CD4_POSITIVE_ALPHA_BETA_T_CELL_ACTIVATION | 35 | -0.359816 | -1.027735 | 0.405405 | 0.561003 |
| GOBP_POSITIVE_REGULATION_OF_SIGNALING_RECEPTOR_ACTIVITY | 39 | -0.346347 | -1.026944 | 0.391481 | 0.56269 |
| GOBP_EPITHELIAL_TUBE_BRANCHING_INVOLVED_IN_LUNG_MORPHOGENESIS | 20 | -0.392886 | -1.026934 | 0.424 | 0.562436 |
| GOBP_ORGANIC_ACID_CATABOLIC_PROCESS | 224 | -0.264025 | -1.026481 | 0.366667 | 0.563241 |
| GOBP_ENDOPLASMIC_RETICULUM_TUBULAR_NETWORK_ORGANIZATION | 18 | -0.410514 | -1.02539 | 0.430672 | 0.565602 |
| GOBP_NEGATIVE_REGULATION_OF_CYTOSKELETON_ORGANIZATION | 135 | -0.273929 | -1.025046 | 0.375281 | 0.566164 |
| GOBP_POSITIVE_REGULATION_OF_PATHWAY_RESTRICTED_SMAD_PROTEIN_PHOSPHORYL | 37 | -0.346495 | -1.024646 | 0.399586 | 0.56684 |
| GOBP_T_CELL_DIFFERENTIATION | 226 | -0.263445 | -1.024282 | 0.373832 | 0.567437 |
| GOBP_PLATELET_MORPHOGENESIS | 21 | -0.387367 | -1.024048 | 0.428571 | 0.567702 |
| GOBP_MEMBRANE_PROTEIN_PROTEOLYSIS | 60 | -0.318175 | -1.023799 | 0.417051 | 0.56802 |
| GOBP_RESPONSE_TO_ETHANOL | 97 | -0.289408 | -1.023591 | 0.388764 | 0.568222 |
| GOBP_GLYCINE_METABOLIC_PROCESS | 16 | -0.424131 | -1.023552 | 0.446154 | 0.568038 |
| GOBP_NEGATIVE_REGULATION_OF_FIBROBLAST_GROWTH_FACTOR_RECEPTOR_SIGNALIN | 16 | -0.413966 | -1.023331 | 0.417166 | 0.568239 |
| GOBP_NUCLEOBASE_METABOLIC_PROCESS | 31 | -0.359215 | -1.02321 | 0.422505 | 0.568249 |
| GOBP_FATTY_ACID_BETA_OXIDATION_USING_ACYL_COA_OXIDASE | 15 | -0.425186 | -1.02216 | 0.418067 | 0.570495 |
| GOBP_PEPTIDYL_SERINE_MODIFICATION | 279 | -0.251071 | -1.021966 | 0.362998 | 0.570652 |
| GOBP_POSITIVE_REGULATION_OF_ENDOTHELIAL_CELL_MIGRATION | 96 | -0.290583 | -1.021869 | 0.401361 | 0.570616 |
| GOBP_REGULATION_OF_CD4_POSITIVE_ALPHA_BETA_T_CELL_ACTIVATION | 60 | -0.311074 | -1.02104 | 0.390456 | 0.572348 |
| GOBP_PROSTANOID_METABOLIC_PROCESS | 36 | -0.346361 | -1.021037 | 0.424176 | 0.572065 |
| GOBP_POSITIVE_REGULATION_OF_METAPHASE_ANAPHASE_TRANSITION_OF_CELL_CYCLE | 15 | -0.429546 | -1.020937 | 0.414784 | 0.572035 |
| GOBP_NEGATIVE_REGULATION_OF_VIRAL_LIFE_CYCLE | 19 | -0.398922 | -1.020452 | 0.435789 | 0.572895 |
| GOBP_PEPTIDYL_LYSINE_ACETYLATION | 152 | -0.273948 | -1.019839 | 0.407323 | 0.574137 |
| GOBP_IMMUNE_RESPONSE_REGULATING_SIGNALING_PATHWAY | 461 | -0.24213 | -1.019723 | 0.375335 | 0.574146 |
| GOBP_REGULATION_OF_PROTEIN_DEACETYLATION | 38 | -0.346348 | -1.019501 | 0.446764 | 0.574365 |
| GOBP_NEGATIVE_REGULATION_OF_ENDOTHELIAL_CELL_APOPTOTIC_PROCESS | 25 | -0.369782 | -1.019251 | 0.425662 | 0.574722 |
| GOBP_AMYLOID_PRECURSOR_PROTEIN_METABOLIC_PROCESS | 70 | -0.307109 | -1.018799 | 0.403017 | 0.575551 |
| GOBP_NEGATIVE_REGULATION_OF_CELLULAR_AMIDE_METABOLIC_PROCESS | 159 | -0.26968 | -1.018449 | 0.390558 | 0.576122 |
| GOBP_PHENOL_CONTAINING_COMPOUND_METABOLIC_PROCESS | 70 | -0.301819 | -1.018438 | 0.426407 | 0.575862 |
| GOBP_RESPONSE_TO_INTERLEUKIN_12 | 47 | -0.325343 | -1.018328 | 0.412281 | 0.575858 |
| GOBP_POSITIVE_REGULATION_OF_EXOCYTOSIS | 71 | -0.308556 | -1.01831 | 0.421842 | 0.575612 |
| GOBP_REGULATION_OF_LEUKOCYTE_DEGRANULATION | 39 | -0.337595 | -1.018248 | 0.425764 | 0.575465 |
| GOBP_RESPONSE_TO_MITOCHONDRIAL_DEPOLARISATION | 20 | -0.396739 | -1.018062 | 0.432866 | 0.575637 |
| GOBP_ANTERIOR_POSTERIOR_AXIS_SPECIFICATION | 31 | -0.360281 | -1.017522 | 0.442886 | 0.576671 |
| GOBP_REGULATION_OF_LIPOPOLYSACCHARIDE_MEDIATED_SIGNALING_PATHWAY | 20 | -0.394984 | -1.017422 | 0.433684 | 0.576639 |
| GOBP_REGULATION_OF_FILOPODIUM_ASSEMBLY | 39 | -0.343489 | -1.017412 | 0.431965 | 0.576376 |
| GOBP_NEURON_FATE_COMMITMENT | 27 | -0.360665 | -1.017312 | 0.435789 | 0.576334 |
| GOBP_NEGATIVE_REGULATION_OF_ALPHA_BETA_T_CELL_DIFFERENTIATION | 25 | -0.38388 | -1.017139 | 0.428274 | 0.57647 |
| GOBP_NEGATIVE_REGULATION_OF_CATABOLIC_PROCESS | 270 | -0.25363 | -1.016444 | 0.394619 | 0.577867 |
| GOBP_ADIPOSE_TISSUE_DEVELOPMENT | 33 | -0.352883 | -1.016402 | 0.440404 | 0.577693 |
| GOBP_EXECUTION_PHASE_OF_APOPTOSIS | 71 | -0.299271 | -1.01626 | 0.432099 | 0.577768 |
| GOBP_T_CELL_MIGRATION | 57 | -0.313517 | -1.016078 | 0.410596 | 0.577959 |
| GOBP_REGULATION_OF_AUTOPHAGY_OF_MITOCHONDRION | 31 | -0.353919 | -1.0159 | 0.419913 | 0.5781 |
| GOBP_LACTATION | 39 | -0.341009 | -1.015487 | 0.418112 | 0.578852 |
| GOBP_POSITIVE_REGULATION_OF_COLLAGEN_METABOLIC_PROCESS | 21 | -0.392186 | -1.015302 | 0.449791 | 0.579026 |
| GOBP_REGULATION_OF_BEHAVIOR | 40 | -0.334347 | -1.014975 | 0.433486 | 0.579544 |
| GOBP_REGULATION_OF_SUPRAMOLECULAR_FIBER_ORGANIZATION | 317 | -0.249266 | -1.014888 | 0.404018 | 0.579487 |
| GOBP_PROTEIN_ACYLATION | 217 | -0.258671 | -1.014582 | 0.405963 | 0.579939 |
| GOBP_REGULATION_OF_SMALL_GTPASE_MEDIATED_SIGNAL_TRANSDUCTION | 294 | -0.249317 | -1.014432 | 0.392345 | 0.58002 |
| GOBP_SPECIFICATION_OF_ANIMAL_ORGAN_IDENTITY | 20 | -0.394448 | -1.014282 | 0.442231 | 0.580087 |
| GOBP_NEGATIVE_REGULATION_OF_IMMUNE_SYSTEM_PROCESS | 340 | -0.247547 | -1.014083 | 0.374101 | 0.580273 |
| GOBP_RESPONSE_TO_ACIDIC_PH | 17 | -0.414273 | -1.013898 | 0.455852 | 0.580418 |
| GOBP_REPRODUCTIVE_BEHAVIOR | 19 | -0.397048 | -1.013163 | 0.426559 | 0.581916 |
| GOBP_NUCLEOTIDE_SUGAR_METABOLIC_PROCESS | 36 | -0.342947 | -1.01283 | 0.405229 | 0.582424 |
| GOBP_RESPONSE_TO_PROSTAGLANDIN | 23 | -0.381341 | -1.012587 | 0.437363 | 0.582727 |
| GOBP_NEGATIVE_REGULATION_OF_CELLULAR_RESPONSE_TO_INSULIN_STIMULUS | 35 | -0.347845 | -1.012498 | 0.428875 | 0.582664 |
| GOBP_MONOCYTE_CHEMOTACTIC_PROTEIN_1_PRODUCTION | 15 | -0.416781 | -1.012098 | 0.453763 | 0.583405 |
| GOBP_LEUKOTRIENE_METABOLIC_PROCESS | 28 | -0.365912 | -1.011955 | 0.431818 | 0.583462 |
| GOBP_CELLULAR_MODIFIED_AMINO_ACID_CATABOLIC_PROCESS | 23 | -0.3804 | -1.011903 | 0.440426 | 0.583301 |
| GOBP_ANGIOGENESIS_INVOLVED_IN_WOUND_HEALING | 21 | -0.389265 | -1.011893 | 0.452991 | 0.58304 |
| GOBP_POSITIVE_REGULATION_OF_CALCIUM_ION_TRANSPORT_INTO_CYTOSOL | 40 | -0.333585 | -1.010843 | 0.433962 | 0.585297 |
| GOBP_REGULATION_OF_RECEPTOR_INTERNALIZATION | 39 | -0.337174 | -1.010544 | 0.429448 | 0.58579 |
| GOBP_REGULATION_OF_MRNA_POLYADENYLATION | 17 | -0.393806 | -1.010433 | 0.455301 | 0.585763 |
| GOBP_NEGATIVE_REGULATION_OF_ORGANELLE_ASSEMBLY | 40 | -0.330999 | -1.010416 | 0.412903 | 0.585519 |
| GOBP_REGULATION_OF_PHOSPHOLIPASE_ACTIVITY | 54 | -0.313919 | -1.00848 | 0.4375 | 0.589943 |
| GOBP_POSITIVE_REGULATION_OF_CELLULAR_CARBOHYDRATE_METABOLIC_PROCESS | 46 | -0.328312 | -1.00828 | 0.431579 | 0.590149 |
| GOBP_REGULATION_OF_CELL_GROWTH_INVOLVED_IN_CARDIAC_MUSCLE_CELL_DEVELO | 16 | -0.41338 | -1.007655 | 0.455487 | 0.591411 |

| | | | | | |
|---|---|---|---|---|---|
| GOBP_GLYCOSYL_COMPOUND_METABOLIC_PROCESS | 117 | -0.280532 | -1.007122 | 0.429806 | 0.592482 |
| GOBP_CELLULAR_RESPONSE_TO_MECHANICAL_STIMULUS | 69 | -0.301264 | -1.00707 | 0.447826 | 0.592332 |
| GOBP_SYNAPTIC_TRANSMISSION_GABAERGIC | 25 | -0.372438 | -1.00622 | 0.449485 | 0.59414 |
| GOBP_PLASMA_MEMBRANE_PHOSPHOLIPID_SCRAMBLING | 15 | -0.420624 | -1.006046 | 0.464078 | 0.594312 |
| GOBP_MODULATION_BY_SYMBIONT_OF_HOST_CELLULAR_PROCESS | 23 | -0.372207 | -1.005953 | 0.42827 | 0.59422 |
| GOBP_REGULATION_OF_JUN_KINASE_ACTIVITY | 74 | -0.298629 | -1.005771 | 0.429185 | 0.594366 |
| GOBP_POSITIVE_REGULATION_OF_TRANSPORTER_ACTIVITY | 88 | -0.29296 | -1.005375 | 0.436559 | 0.595088 |
| GOBP_INTEGRATED_STRESS_RESPONSE_SIGNALING | 26 | -0.366978 | -1.005299 | 0.447084 | 0.594972 |
| GOBP_POSITIVE_REGULATION_OF_ENDOCYTOSIS | 85 | -0.293095 | -1.005219 | 0.431493 | 0.594874 |
| GOBP_SYNAPTIC_TRANSMISSION_CHOLINERGIC | 19 | -0.395394 | -1.005053 | 0.453125 | 0.594999 |
| GOBP_HETEROCHROMATIN_ORGANIZATION | 70 | -0.2958 | -1.004909 | 0.431555 | 0.595066 |
| GOBP_VESICLE_MEDIATED_TRANSPORT_IN_SYNAPSE | 156 | -0.267567 | -1.004538 | 0.430233 | 0.595708 |
| GOBP_MODULATION_BY_HOST_OF_VIRAL_GENOME_REPLICATION | 17 | -0.402579 | -1.004353 | 0.470339 | 0.595895 |
| GOBP_NEGATIVE_REGULATION_OF_MACROAUTOPHAGY | 35 | -0.340675 | -1.004136 | 0.450111 | 0.596138 |
| GOBP_POSITIVE_REGULATION_OF_MONOOXYGENASE_ACTIVITY | 28 | -0.355026 | -1.004037 | 0.438114 | 0.596071 |
| GOBP_CRANIAL_NERVE_MORPHOGENESIS | 18 | -0.407104 | -1.003555 | 0.476395 | 0.596978 |
| GOBP_PROTEIN_COMPLEX_OLIGOMERIZATION | 184 | -0.263557 | -1.002932 | 0.440449 | 0.598213 |
| GOBP_NEGATIVE_REGULATION_OF_OSSIFICATION | 25 | -0.369075 | -1.002604 | 0.433134 | 0.59876 |
| GOBP_NEUROTRANSMITTER_RECEPTOR_TRANSPORT | 36 | -0.337844 | -1.002447 | 0.443564 | 0.598847 |
| GOBP_RESPONSE_TO_ISCHEMIA | 41 | -0.328155 | -1.002177 | 0.451064 | 0.599202 |
| GOBP_DETECTION_OF_ABIOTIC_STIMULUS | 95 | -0.283684 | -1.00179 | 0.429844 | 0.599902 |
| GOBP_PLATELET_DEGRANULATION | 105 | -0.282034 | -1.00162 | 0.429487 | 0.600027 |
| GOBP_GLYCOLIPID_BIOSYNTHETIC_PROCESS | 59 | -0.307884 | -1.001473 | 0.433839 | 0.600093 |
| GOBP_MITOCHONDRIAL_ELECTRON_TRANSPORT_NADH_TO_UBIQUINONE | 52 | -0.310172 | -1.001284 | 0.444444 | 0.600285 |
| GOBP_POSITIVE_REGULATION_OF_GENE_EXPRESSION_EPIGENETIC | 55 | -0.312648 | -1.000185 | 0.453799 | 0.602691 |
| GOBP_DNA_DEALKYLATION | 31 | -0.351295 | -0.999946 | 0.46748 | 0.602986 |
| GOBP_TRIGLYCERIDE_BIOSYNTHETIC_PROCESS | 33 | -0.345813 | -0.999926 | 0.434879 | 0.602734 |
| GOBP_NUCLEAR_CHROMOSOME_SEGREGATION | 253 | -0.250021 | -0.999687 | 0.453917 | 0.602979 |
| GOBP_NUCLEAR_MEMBRANE_ORGANIZATION | 17 | -0.40849 | -0.999612 | 0.463415 | 0.602885 |
| GOBP_NUCLEOSIDE_MONOPHOSPHATE_METABOLIC_PROCESS | 72 | -0.295411 | -0.999189 | 0.439024 | 0.603666 |
| GOBP_KERATINOCYTE_PROLIFERATION | 34 | -0.346145 | -0.998756 | 0.455487 | 0.604461 |
| GOBP_ENDOSOME_ORGANIZATION | 80 | -0.288652 | -0.997709 | 0.462203 | 0.606772 |
| GOBP_PEPTIDYL_L_CYSTEINE_S_PALMITOYLATION | 22 | -0.370716 | -0.997436 | 0.472505 | 0.607132 |
| GOBP_MITOCHONDRION_LOCALIZATION | 44 | -0.322 | -0.996988 | 0.461378 | 0.607905 |
| GOBP_POSITIVE_REGULATION_OF_STRIATED_MUSCLE_CELL_DIFFERENTIATION | 28 | -0.355066 | -0.996436 | 0.482618 | 0.608992 |
| GOBP_REGULATION_OF_T_HELPER_CELL_DIFFERENTIATION | 36 | -0.332638 | -0.996314 | 0.418219 | 0.608995 |
| GOBP_RESPONSE_TO_PROSTAGLANDIN_E | 19 | -0.39295 | -0.995679 | 0.463115 | 0.610327 |
| GOBP_NEGATIVE_REGULATION_OF_VASCULAR_ASSOCIATED_SMOOTH_MUSCLE_CELL_PR | 16 | -0.392477 | -0.99547 | 0.454728 | 0.610579 |
| GOBP_POSITIVE_REGULATION_OF_CELL_CYCLE_G2_M_PHASE_TRANSITION | 27 | -0.363159 | -0.99478 | 0.462527 | 0.612023 |
| GOBP_CELLULAR_RESPONSE_TO_VIRUS | 57 | -0.313065 | -0.994435 | 0.443975 | 0.612545 |
| GOBP_SOMATIC_STEM_CELL_POPULATION_MAINTENANCE | 56 | -0.31184 | -0.993491 | 0.4814 | 0.61463 |
| GOBP_RESPONSE_TO_THYROID_HORMONE | 21 | -0.382729 | -0.992647 | 0.460084 | 0.616537 |
| GOBP_IRE1_MEDIATED_UNFOLDED_PROTEIN_RESPONSE | 63 | -0.307467 | -0.992439 | 0.461702 | 0.616763 |
| GOBP_NEGATIVE_REGULATION_OF_LEUKOCYTE_PROLIFERATION | 76 | -0.295205 | -0.992225 | 0.457906 | 0.617016 |
| GOBP_CELLULAR_COMPONENT_MAINTENANCE | 52 | -0.311781 | -0.99198 | 0.467811 | 0.617304 |
| GOBP_SEMI_LUNAR_VALVE_DEVELOPMENT | 31 | -0.346012 | -0.991871 | 0.456103 | 0.617268 |
| GOBP_PROTEIN_TARGETING_TO_LYSOSOME | 23 | -0.380066 | -0.991799 | 0.45182 | 0.617154 |
| GOBP_CELLULAR_RESPONSE_TO_OXYGEN_RADICAL | 19 | -0.388156 | -0.99179 | 0.457661 | 0.616885 |
| GOBP_MAINTENANCE_OF_LOCATION_IN_CELL | 175 | -0.258668 | -0.991775 | 0.467562 | 0.616614 |
| GOBP_PATHWAY_RESTRICTED_SMAD_PROTEIN_PHOSPHORYLATION | 52 | -0.312736 | -0.991638 | 0.465596 | 0.616641 |
| GOBP_POSITIVE_REGULATION_OF_RELEASE_OF_SEQUESTERED_CALCIUM_ION_INTO_CYT | 31 | -0.344394 | -0.991156 | 0.472689 | 0.617516 |
| GOBP_MORPHOGENESIS_OF_A_POLARIZED_EPITHELIUM | 132 | -0.270126 | -0.991139 | 0.473333 | 0.617258 |
| GOBP_POLYOL_CATABOLIC_PROCESS | 24 | -0.366992 | -0.991126 | 0.470954 | 0.616998 |
| GOBP_REGULATION_OF_BONE_REMODELING | 37 | -0.335742 | -0.991073 | 0.471739 | 0.616826 |
| GOBP_BRANCHING_INVOLVED_IN_SALIVARY_GLAND_MORPHOGENESIS | 15 | -0.403052 | -0.991032 | 0.478351 | 0.616626 |
| GOBP_B_CELL_HOMEOSTASIS | 28 | -0.358969 | -0.990963 | 0.460581 | 0.616496 |
| GOBP_CELLULAR_RESPONSE_TO_PH | 16 | -0.403814 | -0.990925 | 0.47379 | 0.616304 |
| GOBP_AMINE_METABOLIC_PROCESS | 128 | -0.272367 | -0.990551 | 0.476636 | 0.616936 |
| GOBP_REGULATION_OF_CALCIUM_ION_TRANSMEMBRANE_TRANSPORT | 107 | -0.274046 | -0.990447 | 0.45815 | 0.616923 |
| GOBP_FATTY_ACID_TRANSMEMBRANE_TRANSPORT | 42 | -0.325508 | -0.989551 | 0.471338 | 0.618924 |
| GOBP_INTRINSIC_APOPTOTIC_SIGNALING_PATHWAY_IN_RESPONSE_TO_DNA_DAMAGE | 98 | -0.275008 | -0.989133 | 0.470339 | 0.619636 |
| GOBP_REGULATION_OF_OXIDATIVE_STRESS_INDUCED_CELL_DEATH | 50 | -0.31747 | -0.988969 | 0.490566 | 0.619739 |
| GOBP_ALPHA_AMINO_ACID_METABOLIC_PROCESS | 159 | -0.264381 | -0.988802 | 0.490323 | 0.619909 |
| GOBP_LYMPHOCYTE_ACTIVATION_INVOLVED_IN_IMMUNE_RESPONSE | 165 | -0.258936 | -0.9879 | 0.479482 | 0.621908 |
| GOBP_REGULATION_OF_CELLULAR_RESPONSE_TO_HEAT | 75 | -0.292097 | -0.987484 | 0.475375 | 0.622629 |
| GOBP_MEGAKARYOCYTE_DIFFERENTIATION | 85 | -0.284092 | -0.986836 | 0.478458 | 0.623915 |
| GOBP_INNERVATION | 17 | -0.385946 | -0.986553 | 0.467091 | 0.624313 |
| GOBP_REGULATION_OF_URINE_VOLUME | 16 | -0.393413 | -0.986278 | 0.476386 | 0.624667 |
| GOBP_OUTFLOW_TRACT_SEPTUM_MORPHOGENESIS | 18 | -0.386243 | -0.985217 | 0.472917 | 0.627045 |
| GOBP_PROTEIN_DEPOLYMERIZATION | 96 | -0.27815 | -0.984638 | 0.478448 | 0.62822 |
| GOBP_POSITIVE_REGULATION_OF_CYTOKINE_PRODUCTION_INVOLVED_IN_IMMUNE_RESP | 48 | -0.31747 | -0.984253 | 0.498952 | 0.628877 |
| GOBP_MITOTIC_RECOMBINATION | 23 | -0.367603 | -0.983518 | 0.475207 | 0.630416 |
| GOBP_LEUKOCYTE_ADHESION_TO_VASCULAR_ENDOTHELIAL_CELL | 42 | -0.32075 | -0.983314 | 0.493363 | 0.630643 |
| GOBP_REGULATION_OF_MONOCYTE_CHEMOTAXIS | 20 | -0.378715 | -0.983271 | 0.502119 | 0.630446 |
| GOBP_CHONDROCYTE_DEVELOPMENT | 24 | -0.364583 | -0.983221 | 0.482759 | 0.630269 |
| GOBP_MACROPHAGE_ACTIVATION_INVOLVED_IN_IMMUNE_RESPONSE | 17 | -0.390526 | -0.982805 | 0.481799 | 0.63108 |
| GOBP_PROTEIN_K48_LINKED_DEUBIQUITINATION | 33 | -0.341568 | -0.982598 | 0.498911 | 0.631325 |
| GOBP_NEGATIVE_REGULATION_OF_RECEPTOR_SIGNALING_PATHWAY_VIA_STAT | 22 | -0.366706 | -0.98187 | 0.470348 | 0.632833 |
| GOBP_NEGATIVE_REGULATION_OF_JNK_CASCADE | 31 | -0.349379 | -0.981864 | 0.469512 | 0.632553 |
| GOBP_REGULATION_OF_ALPHA_BETA_T_CELL_ACTIVATION | 92 | -0.278037 | -0.981109 | 0.469957 | 0.634101 |
| GOBP_REGULATION_OF_MAST_CELL_ACTIVATION_INVOLVED_IN_IMMUNE_RESPONSE | 27 | -0.356717 | -0.98102 | 0.496957 | 0.634036 |
| GOBP_NEGATIVE_REGULATION_OF_INTERLEUKIN_12_PRODUCTION | 17 | -0.397357 | -0.979816 | 0.464066 | 0.636773 |
| GOBP_CELLULAR_COMPONENT_ASSEMBLY_INVOLVED_IN_MORPHOGENESIS | 72 | -0.291145 | -0.979377 | 0.48218 | 0.63758 |
| GOBP_POSITIVE_REGULATION_OF_PLASMA_MEMBRANE_BOUNDED_CELL_PROJECTION_AS | 90 | -0.279783 | -0.977309 | 0.511312 | 0.642548 |
| GOBP_GLIAL_CELL_DEVELOPMENT | 95 | -0.277884 | -0.977295 | 0.497778 | 0.642283 |
| GOBP_POSITIVE_REGULATION_OF_TRANSCRIPTION_BY_RNA_POLYMERASE_I | 22 | -0.367753 | -0.976897 | 0.485106 | 0.642984 |
| GOBP_MUSCLE_ADAPTATION | 83 | -0.28254 | -0.976411 | 0.50655 | 0.643863 |
| GOBP_MEMBRANE_PROTEIN_ECTODOMAIN_PROTEOLYSIS | 42 | -0.325105 | -0.975196 | 0.488706 | 0.646678 |
| GOBP_DORSAL_VENTRAL_PATTERN_FORMATION | 48 | -0.316269 | -0.975053 | 0.490644 | 0.646745 |
| GOBP_NEGATIVE_REGULATION_OF_RECEPTOR_SIGNALING_PATHWAY_VIA_JAK_STAT | 22 | -0.366706 | -0.97438 | 0.492724 | 0.648141 |
| GOBP_REGULATION_OF_DNA_REPAIR | 125 | -0.266075 | -0.974325 | 0.501109 | 0.647966 |
| GOBP_EPIBOLY | 30 | -0.346884 | -0.97397 | 0.489837 | 0.648517 |
| GOBP_PEPTIDYL_THREONINE_MODIFICATION | 114 | -0.269741 | -0.973623 | 0.507338 | 0.649103 |
| GOBP_HORMONE_BIOSYNTHETIC_PROCESS | 51 | -0.313301 | -0.973307 | 0.502101 | 0.649587 |
| GOBP_BONE_CELL_DEVELOPMENT | 29 | -0.349876 | -0.97322 | 0.492537 | 0.649501 |
| GOBP_RIBONUCLEOSIDE_TRIPHOSPHATE_BIOSYNTHETIC_PROCESS | 70 | -0.29129 | -0.97258 | 0.496732 | 0.650793 |
| GOBP_REGULATION_OF_TRANSLATIONAL_INITIATION | 74 | -0.290314 | -0.972435 | 0.507761 | 0.65086 |
| GOBP_SYNAPTIC_VESICLE_EXOCYTOSIS | 80 | -0.283371 | -0.972234 | 0.50905 | 0.651046 |
| GOBP_REGULATION_OF_NECROTIC_CELL_DEATH | 31 | -0.341051 | -0.971971 | 0.511579 | 0.651447 |
| GOBP_SULFUR_AMINO_ACID_METABOLIC_PROCESS | 28 | -0.347189 | -0.971444 | 0.495088 | 0.652489 |
| GOBP_NEGATIVE_REGULATION_OF_ALPHA_BETA_T_CELL_ACTIVATION | 37 | -0.32336 | -0.970439 | 0.514644 | 0.654664 |
| GOBP_NCRNA_EXPORT_FROM_NUCLEUS | 37 | -0.323888 | -0.969465 | 0.5 | 0.656835 |
| GOBP_RIBONUCLEOSIDE_MONOPHOSPHATE_BIOSYNTHETIC_PROCESS | 34 | -0.327355 | -0.969462 | 0.511435 | 0.656536 |
| GOBP_POSITIVE_REGULATION_OF_SMALL_GTPASE_MEDIATED_SIGNAL_TRANSDUCTION | 69 | -0.28878 | -0.968715 | 0.518931 | 0.658212 |
| GOBP_NEGATIVE_REGULATION_OF_CELL_GROWTH | 150 | -0.258027 | -0.968379 | 0.524946 | 0.658779 |
| GOBP_RAC_PROTEIN_SIGNAL_TRANSDUCTION | 37 | -0.330252 | -0.968156 | 0.491837 | 0.659081 |
| GOBP_NEUTRAL_LIPID_CATABOLIC_PROCESS | 28 | -0.345917 | -0.968004 | 0.492754 | 0.659142 |
| GOBP_GPI_ANCHOR_METABOLIC_PROCESS | 30 | -0.345605 | -0.967831 | 0.502137 | 0.659276 |
| GOBP_RESPONSE_TO_PROGESTERONE | 38 | -0.324873 | -0.966653 | 0.497934 | 0.661959 |
| GOBP_REGULATION_OF_CALCIUM_MEDIATED_SIGNALING | 52 | -0.302531 | -0.966306 | 0.531183 | 0.662574 |
| GOBP_ARF_PROTEIN_SIGNAL_TRANSDUCTION | 19 | -0.372323 | -0.965837 | 0.489712 | 0.663429 |
| GOBP_CELLULAR_AMINO_ACID_BIOSYNTHETIC_PROCESS | 65 | -0.292822 | -0.965825 | 0.549234 | 0.66315 |
| GOBP_NUCLEUS_ORGANIZATION | 113 | -0.267793 | -0.965487 | 0.519912 | 0.663703 |
| GOBP_REGULATION_OF_LEUKOCYTE_MEDIATED_IMMUNITY | 193 | -0.249772 | -0.96478 | 0.555315 | 0.665225 |
| GOBP_CAMP_METABOLIC_PROCESS | 18 | -0.381871 | -0.964545 | 0.521484 | 0.665544 |
| GOBP_REGULATION_OF_RECEPTOR_BINDING | 19 | -0.383754 | -0.964535 | 0.489178 | 0.665253 |
| GOBP_POSITIVE_REGULATION_OF_MUSCLE_CELL_DIFFERENTIATION | 55 | -0.30177 | -0.963889 | 0.502359 | 0.666629 |
| GOBP_HISTONE_H2A_UBIQUITINATION | 25 | -0.349777 | -0.963833 | 0.523622 | 0.666482 |
| GOBP_SMOOTHENED_SIGNALING_PATHWAY | 112 | -0.269414 | -0.963571 | 0.551648 | 0.666823 |
| GOBP_REGULATION_OF_SMOOTHENED_SIGNALING_PATHWAY | 66 | -0.288081 | -0.962785 | 0.504237 | 0.668531 |
| GOBP_WOUND_HEALING_SPREADING_OF_EPIDERMAL_CELLS | 16 | -0.384667 | -0.96263 | 0.517316 | 0.668623 |
| GOBP_GLUCAN_BIOSYNTHETIC_PROCESS | 39 | -0.319915 | -0.960744 | 0.521561 | 0.67313 |
| GOBP_REGULATION_OF_NUCLEASE_ACTIVITY | 21 | -0.371537 | -0.960242 | 0.537815 | 0.674097 |
| GOBP_POSITIVE_REGULATION_OF_CYTOSKELETON_ORGANIZATION | 188 | -0.246815 | -0.958469 | 0.548673 | 0.678229 |
| GOBP_POSITIVE_REGULATION_OF_CALCIUM_ION_TRANSMEMBRANE_TRANSPORT | 53 | -0.301875 | -0.958394 | 0.515991 | 0.678119 |
| GOBP_GLIAL_CELL_MIGRATION | 44 | -0.311093 | -0.95823 | 0.53144 | 0.678228 |
| GOBP_POSITIVE_REGULATION_OF_IMMUNE_EFFECTOR_PROCESS | 192 | -0.250906 | -0.957453 | 0.546988 | 0.67985 |
| GOBP_NEGATIVE_REGULATION_OF_PROTEOLYSIS_INVOLVED_IN_CELLULAR_PROTEIN_CA | 58 | -0.296148 | -0.956387 | 0.541667 | 0.682233 |
| GOBP_PURINE_CONTAINING_COMPOUND_BIOSYNTHETIC_PROCESS | 187 | -0.245152 | -0.956024 | 0.59081 | 0.682819 |
| GOBP_STEROID_BIOSYNTHETIC_PROCESS | 141 | -0.25722 | -0.956011 | 0.534935 | 0.682544 |
| GOBP_ALPHA_AMINO_ACID_BIOSYNTHETIC_PROCESS | 58 | -0.29319 | -0.955587 | 0.534831 | 0.683315 |
| GOBP_SOMATIC_DIVERSIFICATION_OF_IMMUNE_RECEPTORS_VIA_SOMATIC_MUTATION | 15 | -0.400512 | -0.954502 | 0.506198 | 0.685871 |
| GOBP_HINDLIMB_MORPHOGENESIS | 22 | -0.36634 | -0.954461 | 0.53112 | 0.685664 |
| GOBP_MEMBRANE_LIPID_CATABOLIC_PROCESS | 34 | -0.331032 | -0.954109 | 0.530786 | 0.686203 |
| GOBP_ER_NUCLEUS_SIGNALING_PATHWAY | 52 | -0.298675 | -0.954043 | 0.526206 | 0.686056 |
| GOBP_GOLGI_TO_ENDOSOME_TRANSPORT | 17 | -0.378213 | -0.9537 | 0.495727 | 0.686654 |
| GOBP_MUSCLE_FIBER_DEVELOPMENT | 38 | -0.317577 | -0.953611 | 0.521008 | 0.686551 |
| GOBP_HIPPO_SIGNALING | 42 | -0.309859 | -0.953286 | 0.5 | 0.687078 |
| GOBP_MICROTUBULE_DEPOLYMERIZATION | 36 | -0.320477 | -0.953272 | 0.530271 | 0.686806 |
| GOBP_REGULATION_OF_T_HELPER_17_TYPE_IMMUNE_RESPONSE | 19 | -0.374972 | -0.953271 | 0.540984 | 0.686497 |
| GOBP_ESTABLISHMENT_OF_CELL_POLARITY | 126 | -0.257318 | -0.95286 | 0.584821 | 0.687241 |
| GOBP_DNA_RECOMBINATION | 273 | -0.240609 | -0.952026 | 0.62037 | 0.689108 |
| GOBP_REGULATION_OF_STEM_CELL_POPULATION_MAINTENANCE | 24 | -0.350735 | -0.951741 | 0.519068 | 0.68948 |
| GOBP_MEMORY | 71 | -0.287085 | -0.95167 | 0.535398 | 0.689343 |
| GOBP_INTERMEDIATE_FILAMENT_BASED_PROCESS | 39 | -0.320181 | -0.951224 | 0.539256 | 0.690199 |
| GOBP_PROTEIN_TARGETING_TO_VACUOLE | 40 | -0.315793 | -0.950865 | 0.560484 | 0.690786 |
| GOBP_POSITIVE_REGULATION_OF_LONG_TERM_SYNAPTIC_POTENTIATION | 20 | -0.369329 | -0.950844 | 0.509474 | 0.690524 |
| GOBP_REGULATION_OF_RESPONSE_TO_DNA_DAMAGE_STIMULUS | 213 | -0.242445 | -0.950811 | 0.631696 | 0.690302 |
| GOBP_LIPID_CATABOLIC_PROCESS | 263 | -0.23806 | -0.950463 | 0.634259 | 0.690877 |
| GOBP_POSITIVE_REGULATION_OF_CELL_MORPHOGENESIS_INVOLVED_IN_DIFFERENTIATIO | 69 | -0.28651 | -0.950352 | 0.549884 | 0.690845 |
| GOBP_PROTEIN_K48_LINKED_UBIQUITINATION | 61 | -0.294749 | -0.95001 | 0.534188 | 0.691397 |
| GOBP_PEPTIDE_CATABOLIC_PROCESS | 25 | -0.349 | -0.949166 | 0.533623 | 0.693239 |
| GOBP_NEGATIVE_REGULATION_OF_OXIDOREDUCTASE_ACTIVITY | 21 | -0.370495 | -0.947495 | 0.569002 | 0.69731 |
| GOBP_LYSOSOMAL_TRANSPORT | 104 | -0.262146 | -0.946624 | 0.588636 | 0.699304 |
| GOBP_DOPAMINE_RECEPTOR_SIGNALING_PATHWAY | 30 | -0.337683 | -0.946512 | 0.535865 | 0.699303 |
| GOBP_MULTIVESICULAR_BODY_SORTING_PATHWAY | 37 | -0.322082 | -0.946496 | 0.563492 | 0.699017 |
| GOBP_REGULATION_OF_MUSCLE_ADAPTATION | 68 | -0.283919 | -0.945918 | 0.530172 | 0.700167 |

| | | | | | |
|---|---|---|---|---|---|
| GOBP_REGULATION_OF_EXTRACELLULAR_MATRIX_ORGANIZATION | 37 | -0.316957 | -0.944333 | 0.554192 | 0.703927 |
| GOBP_POSITIVE_REGULATION_OF_PROTEIN_POLYMERIZATION | 114 | -0.26295 | -0.943888 | 0.594595 | 0.704753 |
| GOBP_HEART_FORMATION | 17 | -0.377086 | -0.943496 | 0.526971 | 0.705453 |
| GOBP_REGULATION_OF_LYMPHOCYTE_MIGRATION | 56 | -0.293297 | -0.943316 | 0.569388 | 0.705565 |
| GOBP_POSITIVE_REGULATION_OF_LIPID_CATABOLIC_PROCESS | 18 | -0.378567 | -0.942678 | 0.558282 | 0.706878 |
| GOBP_REGULATION_OF_PLASMA_LIPOPROTEIN_PARTICLE_LEVELS | 64 | -0.284512 | -0.942376 | 0.578366 | 0.707298 |
| GOBP_REGULATION_OF_PHOSPHOLIPID_METABOLIC_PROCESS | 27 | -0.345201 | -0.941787 | 0.544444 | 0.708488 |
| GOBP_RIBOSE_PHOSPHATE_BIOSYNTHETIC_PROCESS | 182 | -0.243892 | -0.941518 | 0.605791 | 0.708894 |
| GOBP_ANTIGEN_PROCESSING_AND_PRESENTATION_OF_EXOGENOUS_PEPTIDE_ANTIGEN_ | 77 | -0.277308 | -0.94123 | 0.569937 | 0.709318 |
| GOBP_MITOCHONDRIAL_GENOME_MAINTENANCE | 21 | -0.361413 | -0.941035 | 0.539554 | 0.709537 |
| GOBP_RESPONSE_TO_TESTOSTERONE | 37 | -0.321727 | -0.940674 | 0.529412 | 0.710124 |
| GOBP_MEIOSIS_I_CELL_CYCLE_PROCESS | 105 | -0.268424 | -0.940642 | 0.568282 | 0.709897 |
| GOBP_POSITIVE_REGULATION_OF_I_KAPPAB_KINASE_NF_KAPPAB_SIGNALING | 169 | -0.248832 | -0.940332 | 0.605856 | 0.710361 |
| GOBP_DIOL_BIOSYNTHETIC_PROCESS | 22 | -0.364207 | -0.939383 | 0.545825 | 0.712408 |
| GOBP_NEGATIVE_REGULATION_OF_ACTIN_FILAMENT_DEPOLYMERIZATION | 39 | -0.313096 | -0.939289 | 0.532468 | 0.712308 |
| GOBP_REGULATION_OF_ACTIN_FILAMENT_BASED_PROCESS | 326 | -0.230477 | -0.939276 | 0.678082 | 0.712019 |
| GOBP_REGULATION_OF_MONOCYTE_DIFFERENTIATION | 19 | -0.367722 | -0.939034 | 0.543086 | 0.712331 |
| GOBP_RIBONUCLEOSIDE_MONOPHOSPHATE_METABOLIC_PROCESS | 55 | -0.292825 | -0.938825 | 0.572034 | 0.712562 |
| GOBP_SNO_S_RNA_METABOLIC_PROCESS | 15 | -0.385944 | -0.937234 | 0.532129 | 0.716276 |
| GOBP_ORGANIC_HYDROXY_COMPOUND_METABOLIC_PROCESS | 425 | -0.224485 | -0.936522 | 0.71599 | 0.717816 |
| GOBP_REGULATION_OF_PROTEIN_LOCALIZATION_TO_CELL_PERIPHERY | 103 | -0.263648 | -0.936423 | 0.587234 | 0.717764 |
| GOBP_POSITIVE_REGULATION_OF_LEUKOCYTE_DEGRANULATION | 22 | -0.351562 | -0.935682 | 0.565306 | 0.719328 |
| GOBP_RECEPTOR_METABOLIC_PROCESS | 141 | -0.251 | -0.935531 | 0.63109 | 0.719394 |
| GOBP_PROTEIN_AUTOPHOSPHORYLATION | 203 | -0.240188 | -0.935288 | 0.634259 | 0.71972 |
| GOBP_RESPONSE_TO_XENOBIOTIC_STIMULUS | 85 | -0.265743 | -0.935029 | 0.621505 | 0.720081 |
| GOBP_DNA_DOUBLE_STRAND_BREAK_PROCESSING | 21 | -0.359987 | -0.934811 | 0.55414 | 0.720292 |
| GOBP_POSITIVE_REGULATION_OF_CATABOLIC_PROCESS | 410 | -0.223809 | -0.934111 | 0.745501 | 0.721708 |
| GOBP_TOLL_LIKE_RECEPTOR_3_SIGNALING_PATHWAY | 19 | -0.366996 | -0.933988 | 0.547009 | 0.721698 |
| GOBP_RESPONSE_TO_INTERFERON_ALPHA | 19 | -0.364407 | -0.932878 | 0.558333 | 0.724233 |
| GOBP_POSITIVE_REGULATION_OF_ADAPTIVE_IMMUNE_RESPONSE | 94 | -0.268112 | -0.932698 | 0.609808 | 0.72436 |
| GOBP_POSITIVE_REGULATION_OF_TOLL_LIKE_RECEPTOR_SIGNALING_PATHWAY | 25 | -0.347581 | -0.932582 | 0.561983 | 0.72436 |
| GOBP_SULFUR_COMPOUND_CATABOLIC_PROCESS | 39 | -0.30979 | -0.932268 | 0.548117 | 0.724848 |
| GOBP_ORGANELLE_TRANSPORT_ALONG_MICROTUBULE | 78 | -0.27214 | -0.930887 | 0.613248 | 0.728032 |
| GOBP_NEGATIVE_REGULATION_OF_DNA_BINDING | 47 | -0.304416 | -0.930449 | 0.582796 | 0.72886 |
| GOBP_INTERLEUKIN_6_MEDIATED_SIGNALING_PATHWAY | 23 | -0.354716 | -0.928972 | 0.56427 | 0.732221 |
| GOBP_TRNA_TRANSPORT | 36 | -0.314936 | -0.928513 | 0.550102 | 0.733056 |
| GOBP_WATER_HOMEOSTASIS | 59 | -0.288818 | -0.928268 | 0.5875 | 0.733384 |
| GOBP_GLYCEROLIPID_METABOLIC_PROCESS | 362 | -0.22446 | -0.928102 | 0.694175 | 0.73347 |
| GOBP_CHOLESTEROL_STORAGE | 16 | -0.373625 | -0.926814 | 0.548134 | 0.736417 |
| GOBP_PHOTORECEPTOR_CELL_DIFFERENTIATION | 41 | -0.305906 | -0.926684 | 0.576017 | 0.736425 |
| GOBP_POSITIVE_REGULATION_OF_CELLULAR_PROTEIN_LOCALIZATION | 267 | -0.232544 | -0.926508 | 0.669663 | 0.736563 |
| GOBP_NEGATIVE_REGULATION_OF_OXIDATIVE_STRESS_INDUCED_NEURON_DEATH | 15 | -0.380336 | -0.926313 | 0.527233 | 0.736695 |
| GOBP_ACTIN_FILAMENT_BASED_TRANSPORT | 20 | -0.351904 | -0.92609 | 0.556044 | 0.736923 |
| GOBP_TELOMERE_MAINTENANCE_VIA_RECOMBINATION | 15 | -0.374109 | -0.925935 | 0.517895 | 0.73699 |
| GOBP_REGULATION_OF_VESICLE_FUSION | 21 | -0.354067 | -0.925557 | 0.586639 | 0.737647 |
| GOBP_BLASTOCYST_GROWTH | 19 | -0.370833 | -0.925468 | 0.583153 | 0.737548 |
| GOBP_DETECTION_OF_OTHER_ORGANISM | 16 | -0.369794 | -0.925057 | 0.556485 | 0.738306 |
| GOBP_REGULATION_OF_HEMATOPOIETIC_STEM_CELL_DIFFERENTIATION | 72 | -0.275141 | -0.924642 | 0.616034 | 0.739059 |
| GOBP_GOLGI_ORGANIZATION | 138 | -0.249163 | -0.924501 | 0.610022 | 0.739088 |
| GOBP_CELL_DEATH_IN_RESPONSE_TO_OXIDATIVE_STRESS | 70 | -0.27685 | -0.924471 | 0.597826 | 0.738836 |
| GOBP_UBIQUITIN_DEPENDENT_PROTEIN_CATABOLIC_PROCESS_VIA_THE_MULTIVESICULA | 23 | -0.345816 | -0.92425 | 0.589912 | 0.738995 |
| GOBP_RESPONSE_TO_INTERLEUKIN_4 | 32 | -0.319448 | -0.924105 | 0.604888 | 0.73904 |
| GOBP_CELLULAR_RESPONSE_TO_ESTRADIOL_STIMULUS | 27 | -0.334811 | -0.923781 | 0.579477 | 0.739529 |
| GOBP_NEGATIVE_REGULATION_OF_IMMUNE_EFFECTOR_PROCESS | 115 | -0.256698 | -0.923103 | 0.611494 | 0.740876 |
| GOBP_SPINDLE_ASSEMBLY | 113 | -0.253584 | -0.922491 | 0.654945 | 0.742121 |
| GOBP_POSITIVE_REGULATION_OF_TRANSCRIPTION_OF_NOTCH_RECEPTOR_TARGET | 17 | -0.37409 | -0.922455 | 0.562762 | 0.741882 |
| GOBP_PYRIMIDINE_NUCLEOBASE_METABOLIC_PROCESS | 16 | -0.375037 | -0.922065 | 0.563126 | 0.742561 |
| GOBP_RESPONSE_TO_OXYGEN_RADICAL | 23 | -0.343324 | -0.921681 | 0.5625 | 0.743131 |
| GOBP_NEGATIVE_REGULATION_OF_PROTEIN_POLYMERIZATION | 69 | -0.27667 | -0.921321 | 0.621145 | 0.743674 |
| GOBP_REGULATION_OF_LAMELLIPODIUM_ASSEMBLY | 35 | -0.316589 | -0.920975 | 0.586279 | 0.744178 |
| GOBP_MYELOID_LEUKOCYTE_CYTOKINE_PRODUCTION | 25 | -0.341752 | -0.920894 | 0.556468 | 0.74406 |
| GOBP_POSITIVE_REGULATION_OF_PROTEIN_LOCALIZATION_TO_MEMBRANE | 112 | -0.254693 | -0.920796 | 0.637149 | 0.743974 |
| GOBP_INTERLEUKIN_12_PRODUCTION | 55 | -0.29166 | -0.920649 | 0.611814 | 0.743993 |
| GOBP_PROTEIN_SUMOYLATION | 75 | -0.269275 | -0.920355 | 0.61242 | 0.744348 |
| GOBP_POSITIVE_REGULATION_OF_CELLULAR_RESPONSE_TO_INSULIN_STIMULUS | 20 | -0.35084 | -0.91981 | 0.552809 | 0.74534 |
| GOBP_POSITIVE_REGULATION_OF_LAMELLIPODIUM_ASSEMBLY | 26 | -0.3462 | -0.918996 | 0.576923 | 0.746997 |
| GOBP_INTERFERON_GAMMA_MEDIATED_SIGNALING_PATHWAY | 83 | -0.265782 | -0.918446 | 0.625 | 0.748046 |
| GOBP_REGULATION_OF_DNA_RECOMBINATION | 99 | -0.260458 | -0.918262 | 0.633124 | 0.748195 |
| GOBP_MEMBRANE_RAFT_ORGANIZATION | 23 | -0.347324 | -0.917888 | 0.570842 | 0.748781 |
| GOBP_NEGATIVE_REGULATION_OF_NUCLEOCYTOPLASMIC_TRANSPORT | 20 | -0.349034 | -0.917869 | 0.580851 | 0.748498 |
| GOBP_CYCLIC_NUCLEOTIDE_BIOSYNTHETIC_PROCESS | 15 | -0.38826 | -0.917615 | 0.57764 | 0.748769 |
| GOBP_NEUROMUSCULAR_JUNCTION_DEVELOPMENT | 38 | -0.31034 | -0.917378 | 0.56875 | 0.749033 |
| GOBP_FIBROBLAST_MIGRATION | 40 | -0.304612 | -0.917139 | 0.585837 | 0.749284 |
| GOBP_INTERSTRAND_CROSS_LINK_REPAIR | 54 | -0.292787 | -0.916894 | 0.588865 | 0.749524 |
| GOBP_REGULATION_OF_CELLULAR_EXTRAVASATION | 28 | -0.321219 | -0.916499 | 0.57384 | 0.750138 |
| GOBP_AMINE_BIOSYNTHETIC_PROCESS | 27 | -0.33484 | -0.915905 | 0.558887 | 0.751275 |
| GOBP_STEM_CELL_DIVISION | 22 | -0.349247 | -0.915613 | 0.58517 | 0.751674 |
| GOBP_GANGLIOSIDE_METABOLIC_PROCESS | 21 | -0.35063 | -0.915442 | 0.570499 | 0.75176 |
| GOBP_TRANSCRIPTION_COUPLED_NUCLEOTIDE_EXCISION_REPAIR | 73 | -0.276301 | -0.914778 | 0.618026 | 0.75306 |
| GOBP_POSITIVE_REGULATION_OF_CATION_CHANNEL_ACTIVITY | 52 | -0.287952 | -0.914435 | 0.601732 | 0.753586 |
| GOBP_POSITIVE_REGULATION_OF_RESPONSE_TO_DNA_DAMAGE_STIMULUS | 101 | -0.260615 | -0.914115 | 0.632054 | 0.754013 |
| GOBP_NEGATIVE_REGULATION_OF_RESPONSE_TO_BIOTIC_STIMULUS | 86 | -0.263927 | -0.913941 | 0.655172 | 0.754146 |
| GOBP_POSITIVE_REGULATION_OF_DOUBLE_STRAND_BREAK_REPAIR_VIA_NONHOMOLOG( | 16 | -0.369148 | -0.913887 | 0.561674 | 0.753957 |
| GOBP_REGULATION_OF_NUCLEOCYTOPLASMIC_TRANSPORT | 97 | -0.254791 | -0.913263 | 0.64375 | 0.755239 |
| GOBP_REGULATION_OF_ANTIGEN_PROCESSING_AND_PRESENTATION | 18 | -0.365336 | -0.912649 | 0.58351 | 0.756463 |
| GOBP_POSITIVE_REGULATION_OF_DNA_REPAIR | 70 | -0.272965 | -0.912543 | 0.628755 | 0.756404 |
| GOBP_POSITIVE_REGULATION_OF_SUBSTRATE_ADHESION_DEPENDENT_CELL_SPREADIN( | 35 | -0.31674 | -0.9116 | 0.583333 | 0.758386 |
| GOBP_GLYCEROLIPID_BIOSYNTHETIC_PROCESS | 231 | -0.227939 | -0.911152 | 0.734266 | 0.759154 |
| GOBP_PIGMENT_BIOSYNTHETIC_PROCESS | 50 | -0.28861 | -0.910141 | 0.628505 | 0.761235 |
| GOBP_POSITIVE_REGULATION_OF_MULTICELLULAR_ORGANISM_GROWTH | 19 | -0.354398 | -0.909983 | 0.59481 | 0.761288 |
| GOBP_GAMMA_DELTA_T_CELL_ACTIVATION | 19 | -0.357105 | -0.909495 | 0.591398 | 0.762148 |
| GOBP_REGULATION_OF_HEMATOPOIETIC_PROGENITOR_CELL_DIFFERENTIATION | 85 | -0.263406 | -0.908863 | 0.641026 | 0.763344 |
| GOBP_POSITIVE_REGULATION_OF_MACROPHAGE_CHEMOTAXIS | 16 | -0.37388 | -0.908513 | 0.596838 | 0.763857 |
| GOBP_DNA_BIOSYNTHETIC_PROCESS | 177 | -0.239311 | -0.907449 | 0.723256 | 0.766111 |
| GOBP_HETEROPHILIC_CELL_CELL_ADHESION_VIA_PLASMA_MEMBRANE_CELL_ADHESION | 32 | -0.312491 | -0.906265 | 0.609504 | 0.768629 |
| GOBP_KETONE_BIOSYNTHETIC_PROCESS | 35 | -0.313469 | -0.906187 | 0.618182 | 0.768491 |
| GOBP_CELLULAR_RESPONSE_TO_CARBOHYDRATE_STIMULUS | 107 | -0.25221 | -0.906015 | 0.662946 | 0.768574 |
| GOBP_DETECTION_OF_EXTERNAL_BIOTIC_STIMULUS | 21 | -0.346424 | -0.905394 | 0.598778 | 0.769779 |
| GOBP_POSITIVE_REGULATION_OF_B_CELL_MEDIATED_IMMUNITY | 37 | -0.306308 | -0.904864 | 0.609959 | 0.770742 |
| GOBP_REGULATION_OF_PROTEIN_EXPORT_FROM_NUCLEUS | 31 | -0.313486 | -0.904452 | 0.609071 | 0.771382 |
| GOBP_VISUAL_BEHAVIOR | 36 | -0.308861 | -0.904257 | 0.599156 | 0.771513 |
| GOBP_NEGATIVE_REGULATION_OF_NOTCH_SIGNALING_PATHWAY | 32 | -0.318624 | -0.904129 | 0.59751 | 0.771494 |
| GOBP_DNA_UNWINDING_INVOLVED_IN_DNA_REPLICATION | 16 | -0.358905 | -0.904082 | 0.606589 | 0.771273 |
| GOBP_NEGATIVE_REGULATION_OF_OSTEOBLAST_DIFFERENTIATION | 38 | -0.301381 | -0.903254 | 0.639913 | 0.772896 |
| GOBP_REGULATION_OF_RAC_PROTEIN_SIGNAL_TRANSDUCTION | 17 | -0.365306 | -0.902976 | 0.611465 | 0.773208 |
| GOBP_RESPONSE_TO_CARBOHYDRATE | 171 | -0.237476 | -0.902881 | 0.754967 | 0.773093 |
| GOBP_PROTEIN_LIPID_COMPLEX_ASSEMBLY | 23 | -0.337476 | -0.902384 | 0.590814 | 0.773955 |
| GOBP_RESPONSE_TO_FLUID_SHEAR_STRESS | 28 | -0.325029 | -0.902135 | 0.616205 | 0.77422 |
| GOBP_ALPHA_AMINO_ACID_CATABOLIC_PROCESS | 73 | -0.268641 | -0.901132 | 0.675732 | 0.776338 |
| GOBP_LIPID_HOMEOSTASIS | 113 | -0.248664 | -0.89864 | 0.718341 | 0.781891 |
| GOBP_NEGATIVE_REGULATION_OF_ACTIN_FILAMENT_BUNDLE_ASSEMBLY | 26 | -0.326949 | -0.897773 | 0.60771 | 0.783662 |
| GOBP_MODULATION_BY_VIRUS_OF_HOST_PROCESS | 28 | -0.3231 | -0.897329 | 0.644491 | 0.784384 |
| GOBP_PROTEIN_LOCALIZATION_TO_ENDOSOME | 25 | -0.331754 | -0.896716 | 0.592292 | 0.785476 |
| GOBP_DICARBOXYLIC_ACID_METABOLIC_PROCESS | 82 | -0.262306 | -0.896636 | 0.656904 | 0.785332 |
| GOBP_AMYLOID_PRECURSOR_PROTEIN_CATABOLIC_PROCESS | 53 | -0.279998 | -0.896446 | 0.62419 | 0.785444 |
| GOBP_POSITIVE_REGULATION_OF_T_CELL_MEDIATED_IMMUNITY | 47 | -0.288964 | -0.896307 | 0.634409 | 0.785447 |
| GOBP_RESPONSE_TO_HEAT | 145 | -0.240452 | -0.895841 | 0.736487 | 0.786166 |
| GOBP_POLYOL_BIOSYNTHETIC_PROCESS | 57 | -0.274989 | -0.894626 | 0.634956 | 0.788749 |
| GOBP_NEGATIVE_REGULATION_OF_DNA_TEMPLATED_TRANSCRIPTION_ELONGATION | 17 | -0.356648 | -0.894539 | 0.589958 | 0.788641 |
| GOBP_COVALENT_CHROMATIN_MODIFICATION | 426 | -0.214553 | -0.893988 | 0.875 | 0.789593 |
| GOBP_PROTEIN_ACETYLATION | 178 | -0.233607 | -0.893977 | 0.767442 | 0.789277 |
| GOBP_PURINE_CONTAINING_COMPOUND_METABOLIC_PROCESS | 400 | -0.213894 | -0.893335 | 0.859189 | 0.790436 |
| GOBP_POSITIVE_REGULATION_OF_PROTEIN_LOCALIZATION_TO_NUCLEUS | 80 | -0.264608 | -0.892781 | 0.695932 | 0.791392 |
| GOBP_RESPONSE_TO_MONOSACCHARIDE | 152 | -0.2375 | -0.892733 | 0.707944 | 0.791157 |
| GOBP_POSITIVE_REGULATION_OF_CARBOHYDRATE_METABOLIC_PROCESS | 65 | -0.268961 | -0.891793 | 0.664583 | 0.793067 |
| GOBP_MICROTUBULE_ANCHORING | 26 | -0.322846 | -0.890666 | 0.647436 | 0.795392 |
| GOBP_REGULATION_OF_MYOTUBE_DIFFERENTIATION | 27 | -0.327106 | -0.890539 | 0.62931 | 0.795345 |
| GOBP_AMYLOID_PRECURSOR_PROTEIN_BIOSYNTHETIC_PROCESS | 15 | -0.373627 | -0.889384 | 0.617647 | 0.797698 |
| GOBP_POSITIVE_REGULATION_OF_PROTEIN_TARGETING_TO_MEMBRANE | 26 | -0.326691 | -0.889193 | 0.660044 | 0.797802 |
| GOBP_CELLULAR_AMINO_ACID_CATABOLIC_PROCESS | 90 | -0.253901 | -0.888258 | 0.680851 | 0.799656 |
| GOBP_ENGULFMENT_OF_APOPTOTIC_CELL | 18 | -0.353938 | -0.888086 | 0.616803 | 0.799697 |
| GOBP_CELLULAR_RESPONSE_TO_ALCOHOL | 70 | -0.267547 | -0.88794 | 0.674897 | 0.799675 |
| GOBP_REGULATION_OF_MEGAKARYOCYTE_DIFFERENTIATION | 69 | -0.267814 | -0.887915 | 0.665871 | 0.799399 |
| GOBP_RETINOL_METABOLIC_PROCESS | 27 | -0.324331 | -0.887907 | 0.632568 | 0.799081 |
| GOBP_MYOFIBRIL_ASSEMBLY | 40 | -0.295398 | -0.887627 | 0.644958 | 0.799377 |
| GOBP_DEOXYRIBONUCLEOSIDE_TRIPHOSPHATE_METABOLIC_PROCESS | 15 | -0.37314 | -0.886968 | 0.608421 | 0.800546 |
| GOBP_MITOTIC_G2_M_TRANSITION_CHECKPOINT | 36 | -0.297262 | -0.886017 | 0.644958 | 0.802386 |
| GOBP_POSITIVE_REGULATION_OF_ALPHA_BETA_T_CELL_ACTIVATION | 60 | -0.272309 | -0.885816 | 0.660131 | 0.802499 |
| GOBP_REGULATION_OF_INTRACELLULAR_STEROID_HORMONE_RECEPTOR_SIGNALING_P/ | 64 | -0.272061 | -0.885699 | 0.681435 | 0.802408 |
| GOBP_SPINAL_CORD_DEVELOPMENT | 62 | -0.266588 | -0.885478 | 0.682377 | 0.802575 |
| GOBP_NEGATIVE_REGULATION_OF_DNA_METABOLIC_PROCESS | 109 | -0.249023 | -0.885315 | 0.725225 | 0.802617 |
| GOBP_NEGATIVE_REGULATION_OF_EPITHELIAL_CELL_DIFFERENTIATION | 29 | -0.32302 | -0.885218 | 0.620915 | 0.802507 |
| GOBP_PYRIMIDINE_NUCLEOSIDE_TRIPHOSPHATE_METABOLIC_PROCESS | 22 | -0.336779 | -0.88507 | 0.635762 | 0.802507 |
| GOBP_CELLULAR_RESPIRATION | 176 | -0.230117 | -0.884171 | 0.790094 | 0.804247 |
| GOBP_NEGATIVE_REGULATION_OF_GTPASE_ACTIVITY | 41 | -0.288976 | -0.884105 | 0.667364 | 0.804052 |
| GOBP_GLYCEROL_ETHER_METABOLIC_PROCESS | 17 | -0.34667 | -0.883203 | 0.613108 | 0.805749 |
| GOBP_NEGATIVE_REGULATION_OF_CELLULAR_PROTEIN_CATABOLIC_PROCESS | 68 | -0.266423 | -0.883067 | 0.680851 | 0.805738 |
| GOBP_SYNAPSE_MATURATION | 21 | -0.337203 | -0.882716 | 0.62253 | 0.80619 |
| GOBP_REGULATION_OF_G_PROTEIN_COUPLED_RECEPTOR_SIGNALING_PATHWAY | 105 | -0.248478 | -0.882663 | 0.734694 | 0.805954 |
| GOBP_REGULATION_OF_MUSCLE_HYPERTROPHY | 49 | -0.277255 | -0.882489 | 0.668076 | 0.805992 |
| GOBP_NEGATIVE_REGULATION_OF_INNATE_IMMUNE_RESPONSE | 55 | -0.277018 | -0.881821 | 0.658947 | 0.807154 |
| GOBP_AMYLOID_BETA_METABOLIC_PROCESS | 53 | -0.277131 | -0.880037 | 0.684989 | 0.810905 |
| GOBP_REGULATION_OF_MACROAUTOPHAGY | 155 | -0.232649 | -0.879371 | 0.77305 | 0.812056 |
| GOBP_POSITIVE_REGULATION_OF_INTERLEUKIN_12_PRODUCTION | 36 | -0.293626 | -0.87904 | 0.633262 | 0.812466 |
| GOBP_METANEPHRIC_NEPHRON_MORPHOGENESIS | 15 | -0.362719 | -0.878987 | 0.62987 | 0.812231 |
| GOBP_RNA_PHOSPHODIESTER_BOND_HYDROLYSIS | 142 | -0.235894 | -0.878005 | 0.788546 | 0.814142 |
| GOBP_NEGATIVE_REGULATION_OF_CELLULAR_SENESCENCE | 18 | -0.34343 | -0.877966 | 0.644491 | 0.813886 |

| | | | | | |
|---|---|---|---|---|---|
| GOBP_NEUROTRANSMITTER_RECEPTOR_INTERNALIZATION | 16 | -0.355452 | -0.877825 | 0.631027 | 0.813832 |
| GOBP_REGULATION_OF_VACUOLE_ORGANIZATION | 46 | -0.283627 | -0.877421 | 0.66318 | 0.814399 |
| GOBP_MIRNA_METABOLIC_PROCESS | 27 | -0.319518 | -0.877012 | 0.637527 | 0.81499 |
| GOBP_RESPONSE_TO_PLATELET_DERIVED_GROWTH_FACTOR | 16 | -0.354912 | -0.876388 | 0.634731 | 0.816064 |
| GOBP_NEGATIVE_REGULATION_OF_FATTY_ACID_METABOLIC_PROCESS | 24 | -0.323515 | -0.875549 | 0.6625 | 0.81757 |
| GOBP_RESPONSE_TO_PH | 29 | -0.308407 | -0.875495 | 0.657676 | 0.81734 |
| GOBP_ESTABLISHMENT_OF_PROTEIN_LOCALIZATION_TO_VACUOLE | 48 | -0.279884 | -0.875398 | 0.676471 | 0.817225 |
| GOBP_RENAL_WATER_HOMEOSTASIS | 28 | -0.314457 | -0.874911 | 0.645514 | 0.818001 |
| GOBP_ACTIVATION_OF_ADENYLATE_CYCLASE_ACTIVITY | 23 | -0.324221 | -0.874537 | 0.658103 | 0.818486 |
| GOBP_REGULATION_OF_DOUBLE_STRAND_BREAK_REPAIR_VIA_HOMOLOGOUS_RECOMBI | 47 | -0.284859 | -0.874513 | 0.689278 | 0.818189 |
| GOBP_REGULATION_OF_CELL_AGING | 48 | -0.284441 | -0.873465 | 0.677083 | 0.820136 |
| GOBP_NEGATIVE_REGULATION_OF_REACTIVE_OXYGEN_SPECIES_METABOLIC_PROCESS | 47 | -0.279869 | -0.873003 | 0.68357 | 0.820785 |
| GOBP_REGULATION_OF_EPIDERMAL_GROWTH_FACTOR_ACTIVATED_RECEPTOR_ACTIVITY | 29 | -0.31628 | -0.872784 | 0.655844 | 0.820937 |
| GOBP_INOSITOL_PHOSPHATE_CATABOLIC_PROCESS | 18 | -0.345889 | -0.871928 | 0.626033 | 0.822491 |
| GOBP_MITOTIC_CHROMOSOME_CONDENSATION | 15 | -0.359633 | -0.871784 | 0.654945 | 0.822482 |
| GOBP_PROTEIN_LOCALIZATION_TO_CYTOSKELETON | 56 | -0.274447 | -0.871699 | 0.708609 | 0.822324 |
| GOBP_NEGATIVE_REGULATION_OF_TORC1_SIGNALING | 16 | -0.35638 | -0.871189 | 0.627803 | 0.823066 |
| GOBP_SOMATIC_DIVERSIFICATION_OF_IMMUNOGLOBULINS | 63 | -0.263785 | -0.870509 | 0.734649 | 0.824225 |
| GOBP_REGULATION_OF_ALPHA_BETA_T_CELL_DIFFERENTIATION | 63 | -0.262742 | -0.870237 | 0.715177 | 0.824447 |
| GOBP_PLASMA_LIPOPROTEIN_PARTICLE_CLEARANCE | 42 | -0.291009 | -0.869771 | 0.680672 | 0.825099 |
| GOBP_POSITIVE_REGULATION_OF_DEPHOSPHORYLATION | 54 | -0.272685 | -0.86882 | 0.729508 | 0.826822 |
| GOBP_NON_RECOMBINATIONAL_REPAIR | 100 | -0.249395 | -0.867941 | 0.770492 | 0.828336 |
| GOBP_REGULATION_OF_HISTONE_H3_K4_METHYLATION | 26 | -0.309836 | -0.867016 | 0.665298 | 0.830021 |
| GOBP_PROTEIN_AUTOUBIQUITINATION | 68 | -0.256454 | -0.866978 | 0.748344 | 0.829757 |
| GOBP_POSITIVE_REGULATION_OF_RECEPTOR_INTERNALIZATION | 18 | -0.34847 | -0.866659 | 0.66736 | 0.830124 |
| GOBP_PURINE_NUCLEOBASE_METABOLIC_PROCESS | 16 | -0.354369 | -0.865798 | 0.65121 | 0.831679 |
| GOBP_REGULATION_OF_RYANODINE_SENSITIVE_CALCIUM_RELEASE_CHANNEL_ACTIVITY | 19 | -0.339149 | -0.86565 | 0.663918 | 0.831653 |
| GOBP_AMELOGENESIS | 17 | -0.347443 | -0.865102 | 0.638889 | 0.832424 |
| GOBP_HYPEROSMOTIC_RESPONSE | 23 | -0.320714 | -0.864316 | 0.654321 | 0.833766 |
| GOBP_MEGAKARYOCYTE_DEVELOPMENT | 15 | -0.35689 | -0.864 | 0.628931 | 0.834108 |
| GOBP_PROTEIN_K63_LINKED_UBIQUITINATION | 54 | -0.274408 | -0.863986 | 0.698482 | 0.833799 |
| GOBP_OLEFINIC_COMPOUND_BIOSYNTHETIC_PROCESS | 16 | -0.356377 | -0.862861 | 0.669388 | 0.83586 |
| GOBP_NEGATIVE_REGULATION_OF_CELLULAR_CATABOLIC_PROCESS | 223 | -0.216903 | -0.862532 | 0.878161 | 0.83625 |
| GOBP_MULTICELLULAR_ORGANISM_GROWTH | 110 | -0.239567 | -0.862034 | 0.774775 | 0.836957 |
| GOBP_PROTEIN_KINASE_A_SIGNALING | 28 | -0.314477 | -0.861401 | 0.667364 | 0.83792 |
| GOBP_PURINE_NUCLEOSIDE_MONOPHOSPHATE_BIOSYNTHETIC_PROCESS | 23 | -0.322261 | -0.861002 | 0.675159 | 0.83844 |
| GOBP_ACTIN_FILAMENT_DEPOLYMERIZATION | 52 | -0.268436 | -0.860892 | 0.714571 | 0.838307 |
| GOBP_DNA_METHYLATION_DEPENDENT_HETEROCHROMATIN_ASSEMBLY | 19 | -0.334745 | -0.860628 | 0.648544 | 0.838516 |
| GOBP_REGULATION_OF_PEPTIDYL_THREONINE_PHOSPHORYLATION | 42 | -0.281083 | -0.859748 | 0.700214 | 0.84 |
| GOBP_HISTONE_H2A_MONOUBIQUITINATION | 16 | -0.341524 | -0.859527 | 0.642857 | 0.84011 |
| GOBP_ACTIVATION_OF_INNATE_IMMUNE_RESPONSE | 132 | -0.234487 | -0.858926 | 0.803167 | 0.841033 |
| GOBP_PYRIMIDINE_RIBONUCLEOTIDE_BIOSYNTHETIC_PROCESS | 20 | -0.331523 | -0.858532 | 0.66805 | 0.841483 |
| GOBP_SKELETAL_MUSCLE_TISSUE_REGENERATION | 31 | -0.295246 | -0.858439 | 0.716418 | 0.84132 |
| GOBP_ESTABLISHMENT_OF_EPITHELIAL_CELL_POLARITY | 26 | -0.3111 | -0.857904 | 0.666667 | 0.842055 |
| GOBP_SARCOMERE_ORGANIZATION | 25 | -0.321142 | -0.856917 | 0.686475 | 0.843686 |
| GOBP_RUFFLE_ORGANIZATION | 50 | -0.276592 | -0.856823 | 0.743852 | 0.843545 |
| GOBP_REGULATION_OF_CENTRIOLE_REPLICATION | 20 | -0.326778 | -0.856368 | 0.668803 | 0.844087 |
| GOBP_REGULATION_OF_INTRINSIC_APOPTOTIC_SIGNALING_PATHWAY_BY_P53_CLASS_ME | 26 | -0.310714 | -0.856232 | 0.663774 | 0.844032 |
| GOBP_CELLULAR_GLUCOSE_HOMEOSTASIS | 111 | -0.236916 | -0.856084 | 0.792735 | 0.843993 |
| GOBP_BETA_CATENIN_DESTRUCTION_COMPLEX_DISASSEMBLY | 18 | -0.342407 | -0.85537 | 0.678161 | 0.845147 |
| GOBP_MORPHOGENESIS_OF_AN_EPITHELIAL_SHEET | 47 | -0.274398 | -0.855327 | 0.724576 | 0.844883 |
| GOBP_NEGATIVE_REGULATION_OF_DNA_RECOMBINATION | 40 | -0.282676 | -0.85513 | 0.714894 | 0.844928 |
| GOBP_TRANSCRIPTION_INITIATION_FROM_RNA_POLYMERASE_II_PROMOTER | 165 | -0.224387 | -0.855064 | 0.830626 | 0.844715 |
| GOBP_ORGANOPHOSPHATE_CATABOLIC_PROCESS | 129 | -0.233248 | -0.854791 | 0.817972 | 0.844871 |
| GOBP_EYE_PHOTORECEPTOR_CELL_DEVELOPMENT | 20 | -0.331099 | -0.852919 | 0.688525 | 0.848313 |
| GOBP_REGULATION_OF_LAMELLIPODIUM_ORGANIZATION | 45 | -0.275314 | -0.851812 | 0.728195 | 0.850209 |
| GOBP_REGULATION_OF_PROGRAMMED_NECROTIC_CELL_DEATH | 21 | -0.320601 | -0.851143 | 0.67126 | 0.851226 |
| GOBP_REGULATION_OF_EARLY_ENDOSOME_TO_LATE_ENDOSOME_TRANSPORT | 18 | -0.331918 | -0.85105 | 0.670707 | 0.851066 |
| GOBP_MEMBRANE_BIOGENESIS | 35 | -0.2881 | -0.850306 | 0.712215 | 0.852238 |
| GOBP_EATING_BEHAVIOR | 18 | -0.338328 | -0.848327 | 0.690678 | 0.855883 |
| GOBP_EXTRACELLULAR_VESICLE_BIOGENESIS | 22 | -0.321431 | -0.848244 | 0.68595 | 0.855699 |
| GOBP_REGULATION_OF_MAMMARY_GLAND_EPITHELIAL_CELL_PROLIFERATION | 17 | -0.341229 | -0.848191 | 0.674134 | 0.855453 |
| GOBP_PROTEIN_TRANSMEMBRANE_IMPORT_INTO_INTRACELLULAR_ORGANELLE | 35 | -0.286536 | -0.847085 | 0.716981 | 0.85728 |
| GOBP_REGULATION_OF_FIBROBLAST_MIGRATION | 32 | -0.292993 | -0.846744 | 0.712274 | 0.857589 |
| GOBP_REGULATION_OF_B_CELL_MEDIATED_IMMUNITY | 53 | -0.267418 | -0.846587 | 0.731651 | 0.857569 |
| GOBP_REGULATION_OF_SIGNAL_TRANSDUCTION_BY_P53_CLASS_MEDIATOR | 170 | -0.220726 | -0.846536 | 0.87965 | 0.857301 |
| GOBP_NEGATIVE_REGULATION_OF_DNA_REPAIR | 36 | -0.287717 | -0.845559 | 0.728051 | 0.858808 |
| GOBP_COBALAMIN_METABOLIC_PROCESS | 17 | -0.344551 | -0.845361 | 0.654167 | 0.858853 |
| GOBP_MEMBRANE_DOCKING | 175 | -0.221533 | -0.844931 | 0.876923 | 0.859356 |
| GOBP_REGULATION_OF_RUFFLE_ASSEMBLY | 26 | -0.308913 | -0.844695 | 0.726337 | 0.859443 |
| GOBP_POSITIVE_REGULATION_OF_HISTONE_H3_K4_METHYLATION | 17 | -0.338505 | -0.844486 | 0.685417 | 0.859483 |
| GOBP_PRESYNAPTIC_ENDOCYTOSIS | 53 | -0.270119 | -0.843644 | 0.728 | 0.860701 |
| GOBP_POSITIVE_REGULATION_OF_HISTONE_METHYLATION | 33 | -0.286578 | -0.842849 | 0.725108 | 0.86189 |
| GOBP_POSITIVE_REGULATION_OF_ISOTYPE_SWITCHING | 26 | -0.306242 | -0.842703 | 0.714579 | 0.861816 |
| GOBP_INOSITOL_PHOSPHATE_BIOSYNTHETIC_PROCESS | 24 | -0.31409 | -0.841758 | 0.699153 | 0.863318 |
| GOBP_NEGATIVE_REGULATION_OF_PLATELET_ACTIVATION | 15 | -0.34871 | -0.841564 | 0.707983 | 0.863356 |
| GOBP_MITOTIC_SPINDLE_ORGANIZATION | 115 | -0.233409 | -0.841231 | 0.864679 | 0.863647 |
| GOBP_INTRA_S_DNA_DAMAGE_CHECKPOINT | 17 | -0.339878 | -0.84113 | 0.663866 | 0.863478 |
| GOBP_REGULATION_OF_DNA_DAMAGE_CHECKPOINT | 15 | -0.349231 | -0.840722 | 0.683652 | 0.863888 |
| GOBP_NADH_METABOLIC_PROCESS | 38 | -0.284006 | -0.840677 | 0.725773 | 0.863606 |
| GOBP_HISTONE_H3_K4_TRIMETHYLATION | 16 | -0.34648 | -0.840566 | 0.693387 | 0.863449 |
| GOBP_HOMOLOGOUS_RECOMBINATION | 50 | -0.267652 | -0.839901 | 0.756208 | 0.864359 |
| GOBP_POSITIVE_REGULATION_OF_LAMELLIPODIUM_ORGANIZATION | 33 | -0.28936 | -0.83966 | 0.757282 | 0.864455 |
| GOBP_RNA_POLYADENYLATION | 45 | -0.270582 | -0.839541 | 0.75052 | 0.864347 |
| GOBP_GLYCOSYLATION | 230 | -0.213173 | -0.839402 | 0.904872 | 0.864256 |
| GOBP_REGULATION_OF_RESPONSE_TO_OXIDATIVE_STRESS | 72 | -0.253826 | -0.839077 | 0.771619 | 0.864506 |
| GOBP_NEGATIVE_REGULATION_OF_PROTEASOMAL_UBIQUITIN_DEPENDENT_PROTEIN_CA | 32 | -0.289896 | -0.838404 | 0.763486 | 0.865453 |
| GOBP_ZINC_ION_HOMEOSTASIS | 27 | -0.300055 | -0.837193 | 0.689516 | 0.867357 |
| GOBP_ENDOSOME_TO_LYSOSOME_TRANSPORT | 51 | -0.265434 | -0.835815 | 0.798742 | 0.86966 |
| GOBP_REGULATION_OF_ENDOTHELIAL_CELL_DEVELOPMENT | 16 | -0.340934 | -0.834618 | 0.677228 | 0.871493 |
| GOBP_T_CELL_DIFFERENTIATION_IN_THYMUS | 68 | -0.251455 | -0.834425 | 0.797802 | 0.871532 |
| GOBP_MODIFICATION_OF_SYNAPTIC_STRUCTURE | 17 | -0.343813 | -0.834328 | 0.704684 | 0.871379 |
| GOBP_DOUBLE_STRAND_BREAK_REPAIR | 253 | -0.210773 | -0.834197 | 0.943529 | 0.871274 |
| GOBP_INTESTINAL_ABSORPTION | 24 | -0.305392 | -0.833132 | 0.734818 | 0.87293 |
| GOBP_REGULATION_OF_TRANSLATION_NCRNA_MEDIATED | 24 | -0.308998 | -0.832728 | 0.701754 | 0.873314 |
| GOBP_REGULATION_OF_ACTIN_FILAMENT_LENGTH | 164 | -0.218471 | -0.831043 | 0.888158 | 0.876045 |
| GOBP_ASSOCIATIVE_LEARNING | 49 | -0.263163 | -0.830311 | 0.760684 | 0.877034 |
| GOBP_DNA_TEMPLATED_TRANSCRIPTION_TERMINATION | 75 | -0.244254 | -0.829507 | 0.817204 | 0.878109 |
| GOBP_NUCLEOTIDE_SUGAR_BIOSYNTHETIC_PROCESS | 23 | -0.315285 | -0.828238 | 0.692632 | 0.880103 |
| GOBP_MITOCHONDRIAL_ELECTRON_TRANSPORT_CYTOCHROME_C_TO_OXYGEN | 19 | -0.321197 | -0.828067 | 0.714876 | 0.880043 |
| GOBP_REGULATION_OF_INFLAMMATORY_RESPONSE_TO_ANTIGENIC_STIMULUS | 23 | -0.307927 | -0.827722 | 0.695122 | 0.880319 |
| GOBP_PROTEIN_DESTABILIZATION | 45 | -0.265149 | -0.826176 | 0.776181 | 0.882724 |
| GOBP_DNA_DEPENDENT_DNA_REPLICATION_MAINTENANCE_OF_FIDELITY | 48 | -0.265368 | -0.825475 | 0.770386 | 0.883612 |
| GOBP_OXIDATIVE_PHOSPHORYLATION | 136 | -0.226208 | -0.824794 | 0.892202 | 0.884491 |
| GOBP_GLYCEROPHOSPHOLIPID_CATABOLIC_PROCESS | 24 | -0.299159 | -0.824756 | 0.718367 | 0.884191 |
| GOBP_POSITIVE_REGULATION_OF_MACROPHAGE_ACTIVATION | 23 | -0.316272 | -0.823132 | 0.720322 | 0.886742 |
| GOBP_CELLULAR_AMINO_ACID_METABOLIC_PROCESS | 291 | -0.203624 | -0.822944 | 0.972286 | 0.886683 |
| GOBP_LYSOSOME_LOCALIZATION | 68 | -0.248562 | -0.822888 | 0.810753 | 0.886424 |
| GOBP_PROTEIN_LOCALIZATION_TO_CHROMOSOME | 90 | -0.236603 | -0.822745 | 0.820628 | 0.886294 |
| GOBP_MHC_CLASS_II_BIOSYNTHETIC_PROCESS | 16 | -0.339501 | -0.822583 | 0.678208 | 0.886233 |
| GOBP_TRANSCRIPTION_BY_RNA_POLYMERASE_III | 45 | -0.265479 | -0.821347 | 0.769072 | 0.888026 |
| GOBP_POSITIVE_REGULATION_OF_ALPHA_BETA_T_CELL_DIFFERENTIATION | 45 | -0.268435 | -0.820431 | 0.777538 | 0.889267 |
| GOBP_ANTEROGRADE_AXONAL_TRANSPORT | 44 | -0.267787 | -0.819832 | 0.777538 | 0.8899 |
| GOBP_CD8_POSITIVE_ALPHA_BETA_T_CELL_ACTIVATION | 24 | -0.307675 | -0.81951 | 0.740286 | 0.890109 |
| GOBP_CLATHRIN_COAT_ASSEMBLY | 17 | -0.32553 | -0.817759 | 0.719409 | 0.892774 |
| GOBP_CELLULAR_RESPONSE_TO_GAMMA_RADIATION | 25 | -0.30184 | -0.817616 | 0.723014 | 0.89265 |
| GOBP_ANTIGEN_PROCESSING_AND_PRESENTATION_OF_PEPTIDE_ANTIGEN_VIA_MHC_CLA | 94 | -0.234827 | -0.817432 | 0.863341 | 0.8926 |
| GOBP_NEGATIVE_REGULATION_OF_TOLL_LIKE_RECEPTOR_SIGNALING_PATHWAY | 36 | -0.277976 | -0.81714 | 0.785263 | 0.892739 |
| GOBP_PEPTIDYL_LYSINE_MODIFICATION | 356 | -0.19766 | -0.816355 | 0.982456 | 0.893742 |
| GOBP_REGULATION_OF_CHROMATIN_BINDING | 21 | -0.309935 | -0.815506 | 0.710372 | 0.894756 |
| GOBP_NUCLEOSIDE_PHOSPHATE_CATABOLIC_PROCESS | 72 | -0.241794 | -0.81531 | 0.847917 | 0.894743 |
| GOBP_NEGATIVE_REGULATION_OF_CELL_AGING | 24 | -0.300981 | -0.815193 | 0.748466 | 0.894564 |
| GOBP_REGULATION_OF_GLYCOPROTEIN_METABOLIC_PROCESS | 43 | -0.265196 | -0.815052 | 0.79089 | 0.894434 |
| GOBP_NEGATIVE_REGULATION_OF_PROTEIN_CONTAINING_COMPLEX_ASSEMBLY | 120 | -0.22479 | -0.813415 | 0.860674 | 0.896814 |
| GOBP_RNA_DECAPPING | 16 | -0.334231 | -0.813113 | 0.698545 | 0.896963 |
| GOBP_POSITIVE_REGULATION_OF_ACTIN_CYTOSKELETON_REORGANIZATION | 17 | -0.328845 | -0.812729 | 0.72885 | 0.897211 |
| GOBP_CELLULAR_RESPONSE_TO_ACID_CHEMICAL | 62 | -0.24493 | -0.812451 | 0.825328 | 0.89732 |
| GOBP_MITOCHONDRIAL_TRANSMEMBRANE_TRANSPORT | 101 | -0.232348 | -0.811943 | 0.867238 | 0.897789 |
| GOBP_NEGATIVE_REGULATION_OF_T_CELL_PROLIFERATION | 58 | -0.254594 | -0.811783 | 0.798165 | 0.897703 |
| GOBP_MATURATION_OF_5_8S_RRNA_FROM_TRICISTRONIC_RRNA_TRANSCRIPT_SSU_RRN | 24 | -0.294669 | -0.811113 | 0.743215 | 0.898445 |
| GOBP_TERMINATION_OF_RNA_POLYMERASE_II_TRANSCRIPTION | 35 | -0.276198 | -0.810211 | 0.775109 | 0.899569 |
| GOBP_POSITIVE_REGULATION_OF_VASCULAR_ENDOTHELIAL_CELL_PROLIFERATION | 16 | -0.336181 | -0.810189 | 0.708678 | 0.899246 |
| GOBP_MITOPHAGY | 24 | -0.301211 | -0.809871 | 0.75891 | 0.899419 |
| GOBP_SULFUR_COMPOUND_TRANSPORT | 42 | -0.267013 | -0.809384 | 0.803419 | 0.899826 |
| GOBP_POSITIVE_REGULATION_OF_AUTOPHAGY | 114 | -0.223524 | -0.808996 | 0.887387 | 0.900075 |
| GOBP_POSITIVE_REGULATION_OF_NUCLEOCYTOPLASMIC_TRANSPORT | 58 | -0.251706 | -0.808778 | 0.821739 | 0.900077 |
| GOBP_ESTABLISHMENT_OF_PROTEIN_LOCALIZATION_TO_PLASMA_MEMBRANE | 56 | -0.25196 | -0.80874 | 0.824034 | 0.89978 |
| GOBP_POSITIVE_REGULATION_OF_TYPE_2_IMMUNE_RESPONSE | 15 | -0.337451 | -0.808683 | 0.737844 | 0.899514 |
| GOBP_POSITIVE_REGULATION_OF_TELOMERE_CAPPING | 17 | -0.31834 | -0.806699 | 0.722222 | 0.90241 |
| GOBP_GOLGI_TO_PLASMA_MEMBRANE_PROTEIN_TRANSPORT | 38 | -0.26645 | -0.806399 | 0.784431 | 0.902534 |
| GOBP_NUCLEOBASE_CONTAINING_COMPOUND_TRANSPORT | 233 | -0.203945 | -0.805462 | 0.963049 | 0.903698 |
| GOBP_MAINTENANCE_OF_PROTEIN_LOCATION_IN_NUCLEUS | 21 | -0.311195 | -0.803987 | 0.73374 | 0.905709 |
| GOBP_G2_DNA_DAMAGE_CHECKPOINT | 34 | -0.277386 | -0.80391 | 0.795259 | 0.905461 |
| GOBP_STEROL_METABOLIC_PROCESS | 125 | -0.219338 | -0.802643 | 0.91048 | 0.907101 |
| GOBP_MODULATION_BY_VIRUS_OF_HOST_CELLULAR_PROCESS | 17 | -0.328346 | -0.800707 | 0.762004 | 0.909711 |
| GOBP_VESICLE_MEDIATED_TRANSPORT_BETWEEN_ENDOSOMAL_COMPARTMENTS | 43 | -0.264199 | -0.800478 | 0.806452 | 0.909699 |
| GOBP_PROTEIN_CATABOLIC_PROCESS_IN_THE_VACUOLE | 16 | -0.328591 | -0.799995 | 0.758985 | 0.910086 |
| GOBP_REGULATION_OF_PROTEIN_KINASE_A_SIGNALING | 17 | -0.32829 | -0.799856 | 0.738589 | 0.909956 |
| GOBP_DNA_GEOMETRIC_CHANGE | 112 | -0.222026 | -0.799193 | 0.907489 | 0.910611 |
| GOBP_REGULATION_OF_CYTOPLASMIC_TRANSPORT | 27 | -0.285503 | -0.798824 | 0.766284 | 0.910803 |
| GOBP_FILOPODIUM_ASSEMBLY | 53 | -0.248826 | -0.798741 | 0.793878 | 0.910566 |
| GOBP_CELLULAR_PROTEIN_COMPLEX_DISASSEMBLY | 202 | -0.204864 | -0.797971 | 0.988453 | 0.911396 |
| GOBP_CYTOPLASMIC_MICROTUBULE_ORGANIZATION | 59 | -0.245501 | -0.797617 | 0.836559 | 0.911548 |
| GOBP_MANNOSYLATION | 36 | -0.272923 | -0.795124 | 0.824242 | 0.914982 |
| GOBP_SPHINGOID_METABOLIC_PROCESS | 21 | -0.30131 | -0.79484 | 0.769384 | 0.915072 |
| GOBP_HISTONE_MRNA_METABOLIC_PROCESS | 24 | -0.293305 | -0.794622 | 0.778781 | 0.915051 |
| GOBP_NEGATIVE_REGULATION_OF_CELL_CYCLE_G2_M_PHASE_TRANSITION | 105 | -0.222529 | -0.794537 | 0.873874 | 0.914806 |

| | | | | | |
|---|---|---|---|---|---|
| GOBP_POSITIVE_REGULATION_OF_CYTOKINESIS | 28 | -0.282109 | -0.794118 | 0.76087 | 0.915082 |
| GOBP_REGULATION_OF_DENDRITE_MORPHOGENESIS | 56 | -0.242878 | -0.790024 | 0.827511 | 0.920807 |
| GOBP_PERIPHERAL_NERVOUS_SYSTEM_DEVELOPMENT | 57 | -0.244384 | -0.789261 | 0.858351 | 0.921572 |
| GOBP_RUFFLE_ASSEMBLY | 39 | -0.26128 | -0.788215 | 0.824153 | 0.922688 |
| GOBP_NEGATIVE_REGULATION_OF_INTRINSIC_APOPTOTIC_SIGNALING_PATHWAY_BY_P53 | 19 | -0.315224 | -0.787925 | 0.751592 | 0.922738 |
| GOBP_NEGATIVE_REGULATION_OF_MICROTUBULE_POLYMERIZATION_OR_DEPOLYMERIZA | 33 | -0.277744 | -0.787875 | 0.789916 | 0.922441 |
| GOBP_LYTIC_VACUOLE_ORGANIZATION | 63 | -0.240353 | -0.786572 | 0.871636 | 0.923918 |
| GOBP_REGULATION_OF_STEROL_TRANSPORT | 39 | -0.262156 | -0.784587 | 0.833333 | 0.926345 |
| GOBP_RAP_PROTEIN_SIGNAL_TRANSDUCTION | 15 | -0.320095 | -0.784345 | 0.76569 | 0.926307 |
| GOBP_TRANSCRIPTION_ELONGATION_FROM_RNA_POLYMERASE_I_PROMOTER | 30 | -0.276558 | -0.783998 | 0.828157 | 0.926406 |
| GOBP_INNATE_IMMUNE_RESPONSE_ACTIVATING_SIGNAL_TRANSDUCTION | 104 | -0.220949 | -0.783959 | 0.912017 | 0.92609 |
| GOBP_SOMATIC_RECOMBINATION_OF_IMMUNOGLOBULIN_GENE_SEGMENTS | 54 | -0.247828 | -0.783925 | 0.851111 | 0.925763 |
| GOBP_PROTEASOMAL_UBIQUITIN_INDEPENDENT_PROTEIN_CATABOLIC_PROCESS | 21 | -0.298256 | -0.782845 | 0.774892 | 0.926933 |
| GOBP_REGULATORY_T_CELL_DIFFERENTIATION | 34 | -0.27309 | -0.782602 | 0.779661 | 0.9269 |
| GOBP_RETROGRADE_TRANSPORT_ENDOSOME_TO_GOLGI | 88 | -0.225049 | -0.782413 | 0.902386 | 0.926788 |
| GOBP_POSITIVE_REGULATION_OF_MRNA_SPLICING_VIA_SPLICEOSOME | 17 | -0.311795 | -0.781165 | 0.752918 | 0.9281 |
| GOBP_IMMUNOGLOBULIN_PRODUCTION_INVOLVED_IN_IMMUNOGLOBULIN_MEDIATED_IMM | 54 | -0.247572 | -0.780379 | 0.838174 | 0.928815 |
| GOBP_PYRIMIDINE_NUCLEOSIDE_TRIPHOSPHATE_BIOSYNTHETIC_PROCESS | 17 | -0.322611 | -0.780353 | 0.784736 | 0.928479 |
| GOBP_METHYLATION | 321 | -0.190894 | -0.780106 | 0.997602 | 0.928454 |
| GOBP_LONG_CHAIN_FATTY_ACYL_COA_BIOSYNTHETIC_PROCESS | 17 | -0.315002 | -0.779062 | 0.776 | 0.929533 |
| GOBP_VESICLE_DOCKING_INVOLVED_IN_EXOCYTOSIS | 41 | -0.258712 | -0.778897 | 0.83871 | 0.929376 |
| GOBP_2_OXOGLUTARATE_METABOLIC_PROCESS | 15 | -0.32442 | -0.776769 | 0.761905 | 0.93189 |
| GOBP_ENDOTHELIAL_CELL_CHEMOTAXIS | 22 | -0.291623 | -0.776453 | 0.775051 | 0.931924 |
| GOBP_INTERMEMBRANE_LIPID_TRANSFER | 39 | -0.261577 | -0.775469 | 0.83004 | 0.932942 |
| GOBP_REGULATION_OF_PROTEIN_POLYUBIQUITINATION | 23 | -0.294051 | -0.775266 | 0.776181 | 0.932851 |
| GOBP_REGULATION_OF_TRANSCRIPTION_BY_RNA_POLYMERASE_III | 21 | -0.298551 | -0.774554 | 0.8 | 0.93346 |
| GOBP_PROTEIN_POLYUBIQUITINATION | 317 | -0.19067 | -0.773934 | 0.997561 | 0.933908 |
| GOBP_PROTEIN_TARGETING | 411 | -0.184411 | -0.773038 | 1 | 0.934667 |
| GOBP_MRNA_METHYLATION | 16 | -0.312661 | -0.772864 | 0.773006 | 0.934509 |
| GOBP_CELLULAR_PROTEIN_CONTAINING_COMPLEX_LOCALIZATION | 20 | -0.297436 | -0.772754 | 0.789168 | 0.934273 |
| GOBP_REGULATION_OF_PROTEIN_CATABOLIC_PROCESS | 352 | -0.187119 | -0.772701 | 1 | 0.933961 |
| GOBP_REGULATION_OF_TORC1_SIGNALING | 38 | -0.255792 | -0.772287 | 0.847134 | 0.934141 |
| GOBP_RIG_I_SIGNALING_PATHWAY | 24 | -0.289741 | -0.771621 | 0.802966 | 0.934646 |
| GOBP_MACROMOLECULE_METHYLATION | 275 | -0.191847 | -0.770167 | 0.986395 | 0.936177 |
| GOBP_PHOSPHOLIPID_CATABOLIC_PROCESS | 36 | -0.257124 | -0.765977 | 0.852248 | 0.941113 |
| GOBP_AXO_DENDRITIC_TRANSPORT | 66 | -0.227323 | -0.765098 | 0.884774 | 0.941845 |
| GOBP_NEGATIVE_REGULATION_OF_RESPONSE_TO_OXIDATIVE_STRESS | 18 | -0.302738 | -0.764932 | 0.795309 | 0.941669 |
| GOBP_POSITIVE_REGULATION_OF_POSTTRANSCRIPTIONAL_GENE_SILENCING | 26 | -0.279246 | -0.764537 | 0.813025 | 0.941821 |
| GOBP_UBIQUINONE_METABOLIC_PROCESS | 18 | -0.302259 | -0.764328 | 0.760163 | 0.941701 |
| GOBP_NEGATIVE_REGULATION_OF_ANTIGEN_RECEPTOR_MEDIATED_SIGNALING_PATHWA | 27 | -0.271286 | -0.762183 | 0.84413 | 0.944037 |
| GOBP_DIOL_METABOLIC_PROCESS | 28 | -0.272812 | -0.761387 | 0.849174 | 0.944622 |
| GOBP_MAINTENANCE_OF_PROTEIN_LOCATION | 85 | -0.220452 | -0.760809 | 0.927313 | 0.944987 |
| GOBP_PROTEIN_K63_LINKED_DEUBIQUITINATION | 34 | -0.26292 | -0.759627 | 0.864407 | 0.946014 |
| GOBP_ERROR_PRONE_TRANSLESION_SYNTHESIS | 21 | -0.298458 | -0.759046 | 0.819444 | 0.946345 |
| GOBP_REGULATION_OF_STEROID_METABOLIC_PROCESS | 91 | -0.219045 | -0.758425 | 0.939262 | 0.946707 |
| GOBP_REGULATION_OF_TRANSCRIPTION_FROM_RNA_POLYMERASE_II_PROMOTER_IN_RE | 76 | -0.224413 | -0.757788 | 0.906179 | 0.947074 |
| GOBP_MRNA_MODIFICATION | 22 | -0.28447 | -0.756791 | 0.809524 | 0.947875 |
| GOBP_REGULATION_OF_SKELETAL_MUSCLE_CELL_DIFFERENTIATION | 15 | -0.320763 | -0.756576 | 0.770624 | 0.947757 |
| GOBP_POSITIVE_REGULATION_OF_DNA_RECOMBINATION | 44 | -0.24751 | -0.753991 | 0.855072 | 0.950379 |
| GOBP_REGULATION_OF_NUCLEAR_TRANSCRIBED_MRNA_POLY_A_TAIL_SHORTENING | 15 | -0.313352 | -0.752944 | 0.808247 | 0.951199 |
| GOBP_RNA_MODIFICATION | 163 | -0.198791 | -0.75256 | 0.991011 | 0.951289 |
| GOBP_ORGANISM_EMERGENCE_FROM_PROTECTIVE_STRUCTURE | 21 | -0.286387 | -0.752505 | 0.812109 | 0.950973 |
| GOBP_NEGATIVE_REGULATION_OF_RELEASE_OF_CYTOCHROME_C_FROM_MITOCHONDRIA | 17 | -0.302324 | -0.750518 | 0.810753 | 0.952836 |
| GOBP_NEGATIVE_REGULATION_OF_GLYCOPROTEIN_METABOLIC_PROCESS | 16 | -0.311361 | -0.749703 | 0.792657 | 0.953405 |
| GOBP_MYD88_INDEPENDENT_TOLL_LIKE_RECEPTOR_SIGNALING_PATHWAY | 34 | -0.259196 | -0.749611 | 0.87472 | 0.95313 |
| GOBP_ACETYL_COA_BIOSYNTHETIC_PROCESS | 21 | -0.290316 | -0.749125 | 0.823762 | 0.953309 |
| GOBP_POSITIVE_REGULATION_OF_ALPHA_BETA_T_CELL_PROLIFERATION | 19 | -0.293928 | -0.748037 | 0.787942 | 0.954137 |
| GOBP_HISTONE_H3_DEACETYLATION | 19 | -0.291471 | -0.74798 | 0.805328 | 0.953823 |
| GOBP_REGULATION_OF_CHROMATIN_ASSEMBLY | 19 | -0.283834 | -0.7478 | 0.848018 | 0.953642 |
| GOBP_PROTEIN_IMPORT_INTO_MITOCHONDRIAL_MATRIX | 20 | -0.286329 | -0.747474 | 0.811741 | 0.953639 |
| GOBP_REGULATION_OF_RESPONSE_TO_REACTIVE_OXYGEN_SPECIES | 27 | -0.273536 | -0.746859 | 0.826804 | 0.953948 |
| GOBP_RIBONUCLEOTIDE_CATABOLIC_PROCESS | 34 | -0.256491 | -0.746482 | 0.851111 | 0.953993 |
| GOBP_REGULATION_OF_DOUBLE_STRAND_BREAK_REPAIR | 83 | -0.215937 | -0.745475 | 0.952965 | 0.954699 |
| GOBP_PYROPTOSIS | 16 | -0.306074 | -0.7432 | 0.793926 | 0.95674 |
| GOBP_LYMPHOCYTE_COSTIMULATION | 56 | -0.230047 | -0.742624 | 0.928889 | 0.957003 |
| GOBP_NUCLEOTIDE_EXCISION_REPAIR | 106 | -0.206996 | -0.742333 | 0.967811 | 0.956957 |
| GOBP_POLY_A_PLUS_MRNA_EXPORT_FROM_NUCLEUS | 18 | -0.293935 | -0.741411 | 0.830472 | 0.957585 |
| GOBP_VESICLE_MEDIATED_TRANSPORT_TO_THE_PLASMA_MEMBRANE | 86 | -0.215117 | -0.740686 | 0.948198 | 0.957976 |
| GOBP_NEGATIVE_REGULATION_OF_UBIQUITIN_DEPENDENT_PROTEIN_CATABOLIC_PROCE | 44 | -0.239559 | -0.74013 | 0.906054 | 0.958167 |
| GOBP_RESPONSE_TO_NITRIC_OXIDE | 15 | -0.304258 | -0.739275 | 0.820116 | 0.958645 |
| GOBP_POSITIVE_REGULATION_OF_TORC1_SIGNALING | 17 | -0.294979 | -0.738998 | 0.815789 | 0.958551 |
| GOBP_CEREBELLAR_CORTEX_FORMATION | 16 | -0.305247 | -0.736745 | 0.819277 | 0.960461 |
| GOBP_REGULATION_OF_TRANSCRIPTION_BY_RNA_POLYMERASE_I | 32 | -0.257265 | -0.735418 | 0.896774 | 0.961444 |
| GOBP_ESTABLISHMENT_OF_PROTEIN_LOCALIZATION_TO_MEMBRANE | 330 | -0.179539 | -0.735284 | 1 | 0.961223 |
| GOBP_MAST_CELL_ACTIVATION_INVOLVED_IN_IMMUNE_RESPONSE | 42 | -0.242899 | -0.734511 | 0.902692 | 0.961663 |
| GOBP_POST_GOLGI_VESICLE_MEDIATED_TRANSPORT | 102 | -0.205264 | -0.733977 | 0.971739 | 0.961813 |
| GOBP_POLYAMINE_METABOLIC_PROCESS | 17 | -0.293246 | -0.733276 | 0.838115 | 0.962113 |
| GOBP_INTRACELLULAR_STEROL_TRANSPORT | 21 | -0.271739 | -0.732952 | 0.827161 | 0.96207 |
| GOBP_STORE_OPERATED_CALCIUM_ENTRY | 17 | -0.292867 | -0.732816 | 0.829218 | 0.961851 |
| GOBP_POSITIVE_REGULATION_OF_BONE_RESORPTION | 16 | -0.302312 | -0.732335 | 0.82449 | 0.961939 |
| GOBP_POSITIVE_REGULATION_OF_FILOPODIUM_ASSEMBLY | 24 | -0.271457 | -0.731938 | 0.857831 | 0.961931 |
| GOBP_POSITIVE_REGULATION_OF_CALCIUM_ION_TRANSMEMBRANE_TRANSPORTER_ACTI | 30 | -0.255179 | -0.731746 | 0.861345 | 0.961756 |
| GOBP_NCRNA_PROCESSING | 386 | -0.175728 | -0.731458 | 1 | 0.961675 |
| GOBP_ENDONUCLEOLYTIC_CLEAVAGE_INVOLVED_IN_RRNA_PROCESSING | 15 | -0.287902 | -0.731234 | 0.83 | 0.96154 |
| GOBP_SPLICEOSOMAL_COMPLEX_ASSEMBLY | 65 | -0.224812 | -0.728799 | 0.943694 | 0.96347 |
| GOBP_PIGMENT_GRANULE_ORGANIZATION | 25 | -0.266447 | -0.727913 | 0.829004 | 0.963951 |
| GOBP_RESPONSE_TO_COCAINE | 31 | -0.252702 | -0.727647 | 0.892276 | 0.963851 |
| GOBP_POSITIVE_REGULATION_OF_INSULIN_SECRETION_INVOLVED_IN_CELLULAR_RESPO | 26 | -0.263016 | -0.727301 | 0.855446 | 0.963786 |
| GOBP_G0_TO_G1_TRANSITION | 43 | -0.239664 | -0.725251 | 0.89441 | 0.965327 |
| GOBP_CELLULAR_RESPONSE_TO_PROSTAGLANDIN_STIMULUS | 15 | -0.300686 | -0.724862 | 0.840909 | 0.965321 |
| GOBP_TETRAHYDROFOLATE_METABOLIC_PROCESS | 18 | -0.2887 | -0.723828 | 0.839662 | 0.965918 |
| GOBP_CELLULAR_PIGMENTATION | 47 | -0.232036 | -0.721204 | 0.916493 | 0.967939 |
| GOBP_REGULATION_OF_KILLING_OF_CELLS_OF_OTHER_ORGANISM | 15 | -0.297765 | -0.720556 | 0.857732 | 0.968135 |
| GOBP_SOMATIC_DIVERSIFICATION_OF_IMMUNE_RECEPTORS | 74 | -0.212439 | -0.720145 | 0.969365 | 0.968127 |
| GOBP_NUCLEAR_TRANSCRIBED_MRNA_POLY_A_TAIL_SHORTENING | 34 | -0.247905 | -0.720106 | 0.892276 | 0.967791 |
| GOBP_BASE_EXCISION_REPAIR | 39 | -0.239939 | -0.717163 | 0.914405 | 0.969975 |
| GOBP_REGULATION_OF_ISOTYPE_SWITCHING | 35 | -0.239474 | -0.713551 | 0.928717 | 0.972734 |
| GOBP_NEGATIVE_REGULATION_OF_TYPE_I_INTERFERON_PRODUCTION | 43 | -0.234682 | -0.711076 | 0.911063 | 0.974412 |
| GOBP_POSITIVE_REGULATION_OF_CALCIUM_MEDIATED_SIGNALING | 23 | -0.264409 | -0.710226 | 0.881857 | 0.974744 |
| GOBP_ALPHA_BETA_T_CELL_PROLIFERATION | 33 | -0.247812 | -0.709301 | 0.90389 | 0.975122 |
| GOBP_REGULATION_OF_VIRAL_TRANSCRIPTION | 41 | -0.232831 | -0.709081 | 0.935841 | 0.974957 |
| GOBP_TOLERANCE_INDUCTION | 25 | -0.262861 | -0.709078 | 0.876596 | 0.974587 |
| GOBP_CELL_CELL_ADHESION_MEDIATED_BY_INTEGRIN | 15 | -0.292394 | -0.707755 | 0.845691 | 0.9753 |
| GOBP_PURINE_NUCLEOSIDE_METABOLIC_PROCESS | 56 | -0.213937 | -0.707737 | 0.965885 | 0.97494 |
| GOBP_NEGATIVE_REGULATION_OF_T_CELL_APOPTOTIC_PROCESS | 18 | -0.281254 | -0.70739 | 0.868535 | 0.974846 |
| GOBP_REGULATION_OF_TELOMERE_MAINTENANCE | 75 | -0.210394 | -0.70736 | 0.954733 | 0.974492 |
| GOBP_PROTEIN_LOCALIZATION_TO_CELL_CELL_JUNCTION | 21 | -0.270211 | -0.707215 | 0.858351 | 0.974238 |
| GOBP_MATURATION_OF_SSU_RRNA_FROM_TRICISTRONIC_RRNA_TRANSCRIPT_SSU_RRN | 37 | -0.236184 | -0.707174 | 0.904555 | 0.973903 |
| GOBP_LONG_CHAIN_FATTY_ACYL_COA_METABOLIC_PROCESS | 23 | -0.256439 | -0.704504 | 0.864865 | 0.975669 |
| GOBP_QUINONE_METABOLIC_PROCESS | 29 | -0.251399 | -0.702735 | 0.899177 | 0.976656 |
| GOBP_NCRNA_METABOLIC_PROCESS | 472 | -0.16639 | -0.701462 | 1 | 0.977267 |
| GOBP_RRNA_CONTAINING_RIBONUCLEOPROTEIN_COMPLEX_EXPORT_FROM_NUCLEUS | 15 | -0.282636 | -0.699881 | 0.855131 | 0.978143 |
| GOBP_POSITIVE_REGULATION_OF_TELOMERE_MAINTENANCE | 48 | -0.223773 | -0.6996 | 0.933619 | 0.977989 |
| GOBP_GOLGI_TO_PLASMA_MEMBRANE_TRANSPORT | 56 | -0.218218 | -0.699329 | 0.943845 | 0.977835 |
| GOBP_NEGATIVE_REGULATION_OF_INTERLEUKIN_1_BETA_PRODUCTION | 24 | -0.257662 | -0.697887 | 0.875776 | 0.97857 |
| GOBP_REGULATION_OF_HISTONE_METHYLATION | 56 | -0.213292 | -0.69756 | 0.950495 | 0.978448 |
| GOBP_GLYCOSYL_COMPOUND_BIOSYNTHETIC_PROCESS | 40 | -0.233273 | -0.697318 | 0.944681 | 0.978232 |
| GOBP_MRNA_5_SPLICE_SITE_RECOGNITION | 16 | -0.284202 | -0.697187 | 0.856582 | 0.977952 |
| GOBP_ESTABLISHMENT_OF_PROTEIN_LOCALIZATION_TO_CHROMOSOME | 27 | -0.254186 | -0.696494 | 0.90249 | 0.978091 |
| GOBP_RESOLUTION_OF_MEIOTIC_RECOMBINATION_INTERMEDIATES | 15 | -0.289468 | -0.695797 | 0.879276 | 0.978202 |
| GOBP_NUCLEOBASE_CONTAINING_SMALL_MOLECULE_BIOSYNTHETIC_PROCESS | 102 | -0.194774 | -0.695099 | 0.991525 | 0.978358 |
| GOBP_RNA_PHOSPHODIESTER_BOND_HYDROLYSIS_ENDONUCLEOLYTIC | 74 | -0.205812 | -0.692763 | 0.980263 | 0.979611 |
| GOBP_ZINC_ION_TRANSPORT | 21 | -0.268947 | -0.692375 | 0.88172 | 0.979534 |
| GOBP_ALCOHOL_BIOSYNTHETIC_PROCESS | 130 | -0.189788 | -0.692359 | 0.990676 | 0.979171 |
| GOBP_NUCLEOSIDE_SALVAGE | 16 | -0.284257 | -0.689621 | 0.879447 | 0.980677 |
| GOBP_NUCLEOSIDE_MONOPHOSPHATE_BIOSYNTHETIC_PROCESS | 42 | -0.230921 | -0.688242 | 0.929032 | 0.981242 |
| GOBP_MATURATION_OF_5_8S_RRNA | 35 | -0.232179 | -0.686036 | 0.926778 | 0.982368 |
| GOBP_HISTONE_DEUBIQUITINATION | 23 | -0.255415 | -0.685096 | 0.886831 | 0.982618 |
| GOBP_T_CELL_LINEAGE_COMMITMENT | 25 | -0.251253 | -0.684257 | 0.919662 | 0.982815 |
| GOBP_POSITIVE_REGULATION_OF_GLYCOLYTIC_PROCESS | 19 | -0.26626 | -0.683931 | 0.906639 | 0.982657 |
| GOBP_REGULATION_OF_T_CELL_DIFFERENTIATION_IN_THYMUS | 19 | -0.267772 | -0.68391 | 0.897233 | 0.982298 |
| GOBP_HEMATOPOIETIC_STEM_CELL_DIFFERENTIATION | 84 | -0.200823 | -0.683725 | 0.995745 | 0.982049 |
| GOBP_REGULATION_OF_PLATELET_AGGREGATION | 16 | -0.278495 | -0.683579 | 0.859575 | 0.981778 |
| GOBP_EPHRIN_RECEPTOR_SIGNALING_PATHWAY | 76 | -0.199371 | -0.683194 | 0.980089 | 0.981653 |
| GOBP_REGULATION_OF_ERAD_PATHWAY | 19 | -0.260176 | -0.678747 | 0.902808 | 0.984182 |
| GOBP_PROTEIN_PEPTIDYL_PROLYL_ISOMERIZATION | 35 | -0.232213 | -0.676221 | 0.93988 | 0.985396 |
| GOBP_PROTEIN_LOCALIZATION_TO_VACUOLE | 65 | -0.202715 | -0.674346 | 0.973684 | 0.986152 |
| GOBP_NADH_DEHYDROGENASE_COMPLEX_ASSEMBLY | 64 | -0.204465 | -0.674067 | 0.985685 | 0.985944 |
| GOBP_TOLL_LIKE_RECEPTOR_9_SIGNALING_PATHWAY | 24 | -0.253576 | -0.673269 | 0.910828 | 0.986057 |
| GOBP_NEGATIVE_REGULATION_OF_PROTEIN_CATABOLIC_PROCESS | 106 | -0.188795 | -0.67245 | 1 | 0.986207 |
| GOBP_MITOCHONDRIAL_RNA_METABOLIC_PROCESS | 47 | -0.218184 | -0.672166 | 0.960334 | 0.986 |
| GOBP_MATURATION_OF_LSU_RRNA_FROM_TRICISTRONIC_RRNA_TRANSCRIPT_SSU_RRN | 15 | -0.271137 | -0.672149 | 0.89613 | 0.98564 |
| GOBP_HISTONE_H3_ACETYLATION | 58 | -0.207906 | -0.669308 | 0.968085 | 0.986945 |
| GOBP_PREASSEMBLY_OF_GPI_ANCHOR_IN_ER_MEMBRANE | 15 | -0.282173 | -0.66921 | 0.861167 | 0.986626 |
| GOBP_NUCLEIC_ACID_PHOSPHODIESTER_BOND_HYDROLYSIS | 277 | -0.165848 | -0.668692 | 1 | 0.986548 |
| GOBP_NCRNA_TRANSCRIPTION | 107 | -0.186593 | -0.668203 | 0.993492 | 0.986448 |
| GOBP_MATURATION_OF_SSU_RRNA | 50 | -0.213086 | -0.668055 | 0.970954 | 0.986153 |
| GOBP_POSITIVE_REGULATION_OF_PROTEIN_MATURATION | 21 | -0.258766 | -0.666853 | 0.917927 | 0.986457 |
| GOBP_MRNA_SPLICE_SITE_SELECTION | 39 | -0.22067 | -0.665266 | 0.966518 | 0.986982 |
| GOBP_DNA_LIGATION | 15 | -0.278515 | -0.66265 | 0.892925 | 0.988036 |
| GOBP_DNA_DAMAGE_RESPONSE_SIGNAL_TRANSDUCTION_RESULTING_IN_TRANSCRIPTIO | 18 | -0.26325 | -0.662123 | 0.899598 | 0.987957 |
| GOBP_RETROGRADE_TRANSPORT_ENDOSOME_TO_PLASMA_MEMBRANE | 28 | -0.240508 | -0.661692 | 0.927536 | 0.98782 |
| GOBP_NUCLEOBASE_BIOSYNTHETIC_PROCESS | 18 | -0.256286 | -0.661098 | 0.907173 | 0.987774 |
| GOBP_PYRIMIDINE_RIBONUCLEOSIDE_TRIPHOSPHATE_METABOLIC_PROCESS | 16 | -0.267446 | -0.659888 | 0.895492 | 0.988041 |
| GOBP_REGULATION_OF_TOLERANCE_INDUCTION | 17 | -0.265532 | -0.658289 | 0.928726 | 0.988518 |
| GOBP_RHODOPSIN_MEDIATED_SIGNALING_PATHWAY | 20 | -0.244037 | -0.650241 | 0.908511 | 0.992239 |
| GOBP_POSITIVE_REGULATION_OF_B_CELL_DIFFERENTIATION | 15 | -0.271735 | -0.648976 | 0.916667 | 0.992473 |

| | | | | | |
|---|---|---|---|---|---|
| GOBP_REGULATION_OF_SYNAPTIC_VESICLE_RECYCLING | 21 | -0.248832 | -0.647304 | 0.914712 | 0.992914 |
| GOBP_PEPTIDYL_LYSINE_TRIMETHYLATION | 42 | -0.213304 | -0.647156 | 0.965293 | 0.992603 |
| GOBP_AMINO_ACID_ACTIVATION | 49 | -0.205768 | -0.645608 | 0.973523 | 0.992912 |
| GOBP_REGULATION_OF_TELOMERE_CAPPING | 26 | -0.235502 | -0.644882 | 0.944086 | 0.992878 |
| GOBP_MITOTIC_G2_DNA_DAMAGE_CHECKPOINT | 23 | -0.236407 | -0.636295 | 0.948718 | 0.996284 |
| GOBP_UDP_N_ACETYLGLUCOSAMINE_METABOLIC_PROCESS | 15 | -0.261835 | -0.632619 | 0.925253 | 0.997438 |
| GOBP_PTERIDINE_CONTAINING_COMPOUND_BIOSYNTHETIC_PROCESS | 15 | -0.259639 | -0.631502 | 0.932806 | 0.997522 |
| GOBP_PROTEIN_LOCALIZATION_TO_CHROMOSOME_TELOMERIC_REGION | 30 | -0.223033 | -0.631178 | 0.962451 | 0.99727 |
| GOBP_LIPID_PHOSPHORYLATION | 67 | -0.189574 | -0.629041 | 0.991525 | 0.997723 |
| GOBP_COPPER_ION_HOMEOSTASIS | 15 | -0.259541 | -0.627682 | 0.904459 | 0.99785 |
| GOBP_TERMINATION_OF_RNA_POLYMERASE_I_TRANSCRIPTION | 31 | -0.221994 | -0.624234 | 0.955128 | 0.998764 |
| GOBP_ANAPHASE_PROMOTING_COMPLEX_DEPENDENT_CATABOLIC_PROCESS | 80 | -0.181542 | -0.621895 | 0.995465 | 0.999245 |
| GOBP_GLUTAMINE_METABOLIC_PROCESS | 19 | -0.239716 | -0.621767 | 0.945148 | 0.998915 |
| GOBP_TRANSCRIPTION_BY_RNA_POLYMERASE_I | 67 | -0.185015 | -0.615675 | 0.993507 | 1 |
| GOBP_HISTONE_DEACETYLATION | 72 | -0.183563 | -0.614038 | 0.995565 | 1 |
| GOBP_REGULATION_OF_ACTIN_CYTOSKELETON_REORGANIZATION | 34 | -0.204636 | -0.608579 | 0.972746 | 1 |
| GOBP_ESTABLISHMENT_OF_PROTEIN_LOCALIZATION_TO_TELOMERE | 18 | -0.238898 | -0.607782 | 0.950749 | 1 |
| GOBP_MACROMOLECULE_DEACYLATION | 102 | -0.173005 | -0.60735 | 0.997868 | 1 |
| GOBP_NUCLEAR_TRANSCRIBED_MRNA_CATABOLIC_PROCESS_EXONUCLEOLYTIC | 35 | -0.204019 | -0.606604 | 0.978541 | 1 |
| GOBP_PURINE_CONTAINING_COMPOUND_CATABOLIC_PROCESS | 42 | -0.198564 | -0.601534 | 0.983264 | 1 |
| GOBP_NEGATIVE_REGULATION_OF_TELOMERE_MAINTENANCE_VIA_TELOMERASE | 19 | -0.237436 | -0.601167 | 0.940816 | 1 |
| GOBP_MITOCHONDRIAL_RNA_PROCESSING | 17 | -0.243922 | -0.600544 | 0.944773 | 1 |
| GOBP_RNA_SPLICING_VIA_ENDONUCLEOLYTIC_CLEAVAGE_AND_LIGATION | 16 | -0.244237 | -0.600406 | 0.943845 | 1 |
| GOBP_PURINE_NUCLEOSIDE_MONOPHOSPHATE_METABOLIC_PROCESS | 41 | -0.19688 | -0.599016 | 0.984979 | 1 |
| GOBP_PHOTOTRANSDUCTION_VISIBLE_LIGHT | 22 | -0.224696 | -0.598079 | 0.954545 | 1 |
| GOBP_CLEAVAGE_INVOLVED_IN_RRNA_PROCESSING | 27 | -0.212849 | -0.593557 | 0.968128 | 1 |
| GOBP_TRNA_MODIFICATION | 88 | -0.170958 | -0.592863 | 1 | 1 |
| GOBP_REGULATION_OF_RIG_I_SIGNALING_PATHWAY | 18 | -0.233063 | -0.590265 | 0.953157 | 1 |
| GOBP_CYTOPLASMIC_PATTERN_RECOGNITION_RECEPTOR_SIGNALING_PATHWAY_IN_RES | 29 | -0.207901 | -0.589634 | 0.976744 | 1 |
| GOBP_TELOMERE_MAINTENANCE_VIA_TELOMERE_LENGTHENING | 75 | -0.176453 | -0.589391 | 0.99789 | 1 |
| GOBP_GUANOSINE_CONTAINING_COMPOUND_METABOLIC_PROCESS | 37 | -0.202535 | -0.58823 | 0.980973 | 1 |
| GOBP_REPLICATION_FORK_PROCESSING | 39 | -0.197731 | -0.58581 | 0.987552 | 1 |
| GOBP_BENZENE_CONTAINING_COMPOUND_METABOLIC_PROCESS | 20 | -0.220909 | -0.580143 | 0.974082 | 1 |
| GOBP_MRNA_CIS_SPLICING_VIA_SPLICEOSOME | 30 | -0.203852 | -0.577593 | 0.986456 | 1 |
| GOBP_PEPTIDYL_LYSINE_DIMETHYLATION | 20 | -0.221041 | -0.577369 | 0.974104 | 1 |
| GOBP_RESPONSE_TO_STEROL_DEPLETION | 18 | -0.229977 | -0.573378 | 0.971074 | 1 |
| GOBP_DICARBOXYLIC_ACID_CATABOLIC_PROCESS | 15 | -0.231624 | -0.568761 | 0.96822 | 1 |
| GOBP_RNA_CATABOLIC_PROCESS | 392 | -0.134865 | -0.565964 | 1 | 1 |
| GOBP_INNER_MITOCHONDRIAL_MEMBRANE_ORGANIZATION | 56 | -0.174631 | -0.564625 | 0.998004 | 1 |
| GOBP_RIBOSOMAL_SMALL_SUBUNIT_BIOGENESIS | 75 | -0.167603 | -0.564359 | 1 | 1 |
| GOBP_PROTEIN_METHYLATION | 154 | -0.150293 | -0.563751 | 1 | 1 |
| GOBP_GOLGI_TO_VACUOLE_TRANSPORT | 19 | -0.211851 | -0.558959 | 0.980263 | 1 |
| GOBP_PROTEIN_LOCALIZATION_TO_CHROMATIN | 29 | -0.201104 | -0.55566 | 0.970402 | 1 |
| GOBP_PEPTIDYL_LYSINE_METHYLATION | 112 | -0.153192 | -0.545666 | 1 | 1 |
| GOBP_REGULATION_OF_HISTONE_H3_K9_METHYLATION | 18 | -0.213651 | -0.544203 | 0.979253 | 1 |
| GOBP_TORC1_SIGNALING | 46 | -0.172565 | -0.542163 | 0.995671 | 1 |
| GOBP_POSITIVE_REGULATION_OF_TELOMERE_MAINTENANCE_VIA_TELOMERE_LENGTHEN | 36 | -0.181898 | -0.54186 | 0.989247 | 1 |
| GOBP_NEGATIVE_REGULATION_OF_TELOMERE_MAINTENANCE | 32 | -0.188772 | -0.537033 | 0.995842 | 1 |
| GOBP_STEROL_BIOSYNTHETIC_PROCESS | 63 | -0.165139 | -0.536524 | 1 | 1 |
| GOBP_RRNA_METABOLIC_PROCESS | 234 | -0.135417 | -0.536076 | 1 | 1 |
| GOBP_TRNA_METHYLATION | 40 | -0.174996 | -0.535399 | 0.995851 | 1 |
| GOBP_RNA_DEPENDENT_DNA_BIOSYNTHETIC_PROCESS | 68 | -0.159893 | -0.533128 | 0.997947 | 1 |
| GOBP_ARP2_3_COMPLEX_MEDIATED_ACTIN_NUCLEATION | 37 | -0.179679 | -0.532366 | 0.995763 | 1 |
| GOBP_MISMATCH_REPAIR | 36 | -0.177606 | -0.527946 | 0.99384 | 1 |
| GOBP_TRANSLATIONAL_ELONGATION | 129 | -0.144095 | -0.526221 | 1 | 1 |
| GOBP_SYNAPTIC_VESICLE_CYTOSKELETAL_TRANSPORT | 19 | -0.205035 | -0.523616 | 0.978175 | 1 |
| GOBP_ENDOPLASMIC_RETICULUM_CALCIUM_ION_HOMEOSTASIS | 23 | -0.188104 | -0.513027 | 0.983505 | 1 |
| GOBP_REGULATION_OF_TELOMERE_MAINTENANCE_VIA_TELOMERE_LENGTHENING | 57 | -0.159808 | -0.512571 | 1 | 1 |
| GOBP_PROTEIN_LOCALIZATION_TO_LYSOSOME | 41 | -0.171097 | -0.510724 | 1 | 1 |
| GOBP_ATP_SYNTHESIS_COUPLED_PROTON_TRANSPORT | 27 | -0.182493 | -0.508904 | 0.993852 | 1 |
| GOBP_NEGATIVE_REGULATION_OF_TELOMERE_MAINTENANCE_VIA_TELOMERE_LENGTHEI | 24 | -0.188699 | -0.507982 | 0.988987 | 1 |
| GOBP_MITOCHONDRIAL_ATP_SYNTHESIS_COUPLED_PROTON_TRANSPORT | 22 | -0.190037 | -0.501228 | 0.987755 | 1 |
| GOBP_PHOSPHATIDYLINOSITOL_BIOSYNTHETIC_PROCESS | 107 | -0.138531 | -0.493179 | 1 | 1 |
| GOBP_OLIGOSACCHARIDE_LIPID_INTERMEDIATE_BIOSYNTHETIC_PROCESS | 21 | -0.187947 | -0.49265 | 0.993814 | 1 |
| GOBP_REGULATION_OF_DOUBLE_STRAND_BREAK_REPAIR_VIA_NONHOMOLOGOUS_END_ | 28 | -0.175331 | -0.488662 | 0.997807 | 1 |
| GOBP_NEGATIVE_REGULATION_OF_TRANSLATIONAL_INITIATION | 19 | -0.193582 | -0.488337 | 0.989177 | 1 |
| GOBP_DNA_DAMAGE_RESPONSE_DETECTION_OF_DNA_DAMAGE | 38 | -0.162463 | -0.478979 | 0.993435 | 1 |
| GOBP_PROTEIN_MATURATION_BY_IRON_SULFUR_CLUSTER_TRANSFER | 15 | -0.198115 | -0.473295 | 0.990253 | 1 |
| GOBP_RIBONUCLEOPROTEIN_COMPLEX_SUBUNIT_ORGANIZATION | 214 | -0.116712 | -0.458736 | 1 | 1 |
| GOBP_POSITIVE_REGULATION_OF_CILIUM_ASSEMBLY | 20 | -0.175259 | -0.457975 | 0.997886 | 1 |
| GOBP_TRNA_WOBBLE_BASE_MODIFICATION | 20 | -0.176775 | -0.455705 | 0.995662 | 1 |
| GOBP_TRANSLATIONAL_TERMINATION | 103 | -0.127705 | -0.45518 | 1 | 1 |
| GOBP_RIBONUCLEOPROTEIN_COMPLEX_BIOGENESIS | 452 | -0.107947 | -0.454471 | 1 | 1 |
| GOBP_MITOCHONDRIAL_RESPIRATORY_CHAIN_COMPLEX_ASSEMBLY | 99 | -0.128252 | -0.448088 | 1 | 1 |
| GOBP_NON_MOTILE_CILIUM_ASSEMBLY | 51 | -0.139429 | -0.447105 | 1 | 1 |
| GOBP_CRISTAE_FORMATION | 35 | -0.151695 | -0.444814 | 1 | 1 |
| GOBP_CARDIOLIPIN_METABOLIC_PROCESS | 16 | -0.181524 | -0.443486 | 0.985597 | 1 |
| GOBP_NEGATIVE_REGULATION_OF_TOR_SIGNALING | 39 | -0.147916 | -0.43817 | 1 | 1 |
| GOBP_RIBOSOME_BIOGENESIS | 303 | -0.106299 | -0.433465 | 1 | 1 |
| GOBP_ENTRAINMENT_OF_CIRCADIAN_CLOCK | 29 | -0.15537 | -0.431949 | 1 | 1 |
| GOBP_AEROBIC_RESPIRATION | 82 | -0.122157 | -0.425227 | 1 | 1 |
| GOBP_PORE_COMPLEX_ASSEMBLY | 19 | -0.164292 | -0.424802 | 0.993802 | 1 |
| GOBP_NUCLEOSIDE_BISPHOSPHATE_BIOSYNTHETIC_PROCESS | 64 | -0.126924 | -0.419915 | 1 | 1 |
| GOBP_TRNA_PROCESSING | 128 | -0.11235 | -0.413737 | 1 | 1 |
| GOBP_P_BODY_ASSEMBLY | 21 | -0.154483 | -0.402244 | 1 | 1 |
| GOBP_REGULATION_OF_SISTER_CHROMATID_COHESION | 22 | -0.152119 | -0.392317 | 1 | 1 |
| GOBP_RNA_5_END_PROCESSING | 23 | -0.138379 | -0.37273 | 1 | 0.999827 |

**Table S11. GSEA analysis in $SF3B1^{MT}$ and $SF3B1^{WT}$ CLL patients.** See Table S16 for patient information.

**GSEA Positive** Hematopoietic/Immune pathways are highlighted in red.

| Gene sets | SIZE | ES | NES | NOM p-val | FDR q-val |
|---|---|---|---|---|---|
| GOBP_POSITIVE_REGULATION_OF_NEURON_DIFFERENTIATION | 48 | 0.685306 | 2.002231 | 0 | 0.414537 |
| GOBP_EMBRYONIC_CAMERA_TYPE_EYE_MORPHOGENESIS | 15 | 0.84743 | 1.998656 | 0.002618 | 0.219241 |
| GOBP_REGULATION_OF_CELL_CELL_ADHESION_MEDIATED_BY_CADHERIN | 15 | 0.835616 | 1.967665 | 0 | 0.218959 |
| GOBP_POLARIZED_EPITHELIAL_CELL_DIFFERENTIATION | 16 | 0.797035 | 1.940391 | 0.005236 | 0.232676 |
| GOBP_EMBRYONIC_EYE_MORPHOGENESIS | 18 | 0.81238 | 1.922514 | 0.002646 | 0.228837 |
| GOBP_EMBRYONIC_CAMERA_TYPE_EYE_DEVELOPMENT | 21 | 0.793838 | 1.916275 | 0 | 0.204646 |
| GOBP_RESPONSE_TO_TYPE_I_INTERFERON | 84 | 0.585378 | 1.860135 | 0 | 0.305264 |
| GOBP_CELL_CELL_ADHESION_MEDIATED_BY_CADHERIN | 20 | 0.734494 | 1.859773 | 0.002882 | 0.267996 |
| GOBP_PTERIDINE_CONTAINING_COMPOUND_METABOLIC_PROCESS | 31 | 0.682842 | 1.85412 | 0.002907 | 0.253642 |
| GOBP_CARDIAC_MUSCLE_TISSUE_MORPHOGENESIS | 31 | 0.70039 | 1.847827 | 0 | 0.243898 |
| GOBP_CAMERA_TYPE_EYE_MORPHOGENESIS | 65 | 0.599491 | 1.833472 | 0 | 0.253465 |
| GOBP_EYE_MORPHOGENESIS | 81 | 0.571948 | 1.812767 | 0 | 0.281331 |
| GOBP_CARDIAC_ATRIUM_MORPHOGENESIS | 17 | 0.746075 | 1.796733 | 0.002519 | 0.299403 |
| GOBP_REGULATION_OF_TYPE_I_INTERFERON_MEDIATED_SIGNALING_PATHWAY | 32 | 0.665325 | 1.793537 | 0.009804 | 0.285473 |
| GOBP_VENTRICULAR_SEPTUM_DEVELOPMENT | 40 | 0.649353 | 1.791327 | 0.003205 | 0.27153 |
| GOBP_SKELETAL_MUSCLE_CELL_DIFFERENTIATION | 40 | 0.632861 | 1.788546 | 0.003236 | 0.261647 |
| GOBP_MUSCLE_ORGAN_MORPHOGENESIS | 41 | 0.632834 | 1.78787 | 0.009036 | 0.247754 |
| GOBP_PTERIDINE_CONTAINING_COMPOUND_BIOSYNTHETIC_PROCESS | 15 | 0.780447 | 1.787848 | 0.007634 | 0.23399 |
| GOBP_PHOSPHATIDYLSERINE_METABOLIC_PROCESS | 24 | 0.680278 | 1.776249 | 0.011765 | 0.24458 |
| GOBP_REGULATION_OF_MORPHOGENESIS_OF_A_BRANCHING_STRUCTURE | 22 | 0.708671 | 1.770684 | 0.00554 | 0.243534 |
| GOBP_VENTRICULAR_SEPTUM_MORPHOGENESIS | 24 | 0.684735 | 1.765244 | 0.005602 | 0.242112 |
| GOBP_RETINA_MORPHOGENESIS_IN_CAMERA_TYPE_EYE | 32 | 0.639151 | 1.748021 | 0 | 0.267143 |
| GOBP_GENITALIA_DEVELOPMENT | 20 | 0.703183 | 1.738386 | 0.005291 | 0.276612 |
| GOBP_NEGATIVE_REGULATION_OF_NEURAL_PRECURSOR_CELL_PROLIFERATION | 15 | 0.769058 | 1.726784 | 0.005333 | 0.291996 |
| GOBP_REGULATION_OF_MEIOTIC_NUCLEAR_DIVISION | 21 | 0.683329 | 1.721654 | 0.011364 | 0.291285 |
| GOBP_LENS_DEVELOPMENT_IN_CAMERA_TYPE_EYE | 45 | 0.594929 | 1.716691 | 0.006515 | 0.291667 |
| GOBP_VENTRICULAR_CARDIAC_MUSCLE_TISSUE_DEVELOPMENT | 27 | 0.646152 | 1.715393 | 0.005319 | 0.283896 |
| GOBP_CARDIAC_VENTRICLE_DEVELOPMENT | 67 | 0.566637 | 1.713314 | 0.003745 | 0.278449 |
| GOBP_CARDIAC_VENTRICLE_MORPHOGENESIS | 38 | 0.610687 | 1.704145 | 0.016611 | 0.289051 |
| GOBP_FOLIC_ACID_CONTAINING_COMPOUND_METABOLIC_PROCESS | 25 | 0.666768 | 1.681338 | 0.008499 | 0.335255 |
| GOBP_BRAIN_MORPHOGENESIS | 24 | 0.664536 | 1.676704 | 0.0125 | 0.335841 |
| GOBP_CARDIAC_ATRIUM_DEVELOPMENT | 21 | 0.672721 | 1.668944 | 0.025714 | 0.344027 |
| GOBP_SULFUR_COMPOUND_CATABOLIC_PROCESS | 33 | 0.619208 | 1.661052 | 0.018576 | 0.354471 |
| GOBP_STEM_CELL_PROLIFERATION | 39 | 0.581681 | 1.655665 | 0.01 | 0.360015 |
| GOBP_SECONDARY_PALATE_DEVELOPMENT | 18 | 0.696606 | 1.650559 | 0.010959 | 0.363521 |
| GOBP_GLANDULAR_EPITHELIAL_CELL_DIFFERENTIATION | 25 | 0.636637 | 1.643331 | 0.022727 | 0.373617 |
| GOBP_REGULATION_OF_PRODUCTION_OF_SMALL_RNA_INVOLVED_IN_GENE_SILENC | 20 | 0.687618 | 1.642144 | 0.019553 | 0.366993 |
| GOBP_EMBRYONIC_HEMOPOIESIS | 18 | 0.678995 | 1.627553 | 0.033241 | 0.398045 |
| GOBP_ACTIVATION_OF_PROTEIN_KINASE_B_ACTIVITY | 24 | 0.65272 | 1.626569 | 0.019337 | 0.390787 |
| GOBP_ANGIOGENESIS_INVOLVED_IN_WOUND_HEALING | 20 | 0.663385 | 1.626452 | 0.027919 | 0.381486 |
| GOBP_TETRAHYDROFOLATE_METABOLIC_PROCESS | 18 | 0.681686 | 1.625549 | 0.030471 | 0.374684 |
| GOBP_NEGATIVE_REGULATION_OF_BLOOD_PRESSURE | 23 | 0.633545 | 1.624766 | 0.028818 | 0.367787 |
| GOBP_ESTABLISHMENT_OR_MAINTENANCE_OF_MONOPOLAR_CELL_POLARITY | 17 | 0.683291 | 1.617233 | 0.026385 | 0.380178 |
| GOBP_REGULATION_OF_HAIR_CYCLE | 17 | 0.662887 | 1.612158 | 0.024931 | 0.386091 |
| GOBP_REGULATION_OF_MEIOTIC_CELL_CYCLE | 33 | 0.59674 | 1.610229 | 0.025806 | 0.383097 |
| GOBP_KERATINIZATION | 26 | 0.620551 | 1.601093 | 0.0317 | 0.400033 |
| GOBP_SARCOMERE_ORGANIZATION | 20 | 0.646259 | 1.589004 | 0.036939 | 0.426721 |
| GOBP_EPITHELIAL_TUBE_FORMATION | 88 | 0.495805 | 1.583088 | 0.009174 | 0.435929 |
| GOBP_PROTEIN_LOCALIZATION_TO_GOLGI_APPARATUS | 26 | 0.595134 | 1.570087 | 0.027174 | 0.467807 |
| GOBP_CORNIFICATION | 16 | 0.677397 | 1.565963 | 0.040506 | 0.471799 |
| GOBP_COLUMNAR_CUBOIDAL_EPITHELIAL_CELL_DIFFERENTIATION | 47 | 0.549102 | 1.562435 | 0.024476 | 0.473939 |
| GOBP_NEGATIVE_REGULATION_OF_VIRAL_GENOME_REPLICATION | 46 | 0.538583 | 1.560074 | 0.029412 | 0.472433 |
| GOBP_POSITIVE_REGULATION_OF_OSSIFICATION | 30 | 0.579661 | 1.55953 | 0.04451 | 0.465378 |
| GOBP_CARDIAC_MUSCLE_TISSUE_DEVELOPMENT | 109 | 0.47066 | 1.556613 | 0.014151 | 0.465637 |
| GOBP_SENSORY_ORGAN_MORPHOGENESIS | 134 | 0.456336 | 1.555715 | 0.015873 | 0.460099 |
| GOBP_TUBE_FORMATION | 99 | 0.469854 | 1.553783 | 0.017391 | 0.457656 |
| GOBP_MEMBRANE_DISASSEMBLY | 15 | 0.677846 | 1.547428 | 0.068421 | 0.469422 |
| GOBP_DRUG_METABOLIC_PROCESS | 24 | 0.615905 | 1.546571 | 0.053824 | 0.463939 |
| GOBP_SPINAL_CORD_DEVELOPMENT | 50 | 0.531076 | 1.546139 | 0.013841 | 0.457234 |
| GOBP_POSITIVE_REGULATION_OF_LYMPHOCYTE_MIGRATION | 27 | 0.593918 | 1.545461 | 0.046921 | 0.451615 |
| GOBP_POSITIVE_REGULATION_OF_STRIATED_MUSCLE_CELL_DIFFERENTIATION | 26 | 0.617677 | 1.544242 | 0.032258 | 0.448105 |
| GOBP_REGULATION_OF_CHROMATIN_ASSEMBLY_OR_DISASSEMBLY | 23 | 0.609382 | 1.542691 | 0.049708 | 0.445158 |
| GOBP_GLANDULAR_EPITHELIAL_CELL_DEVELOPMENT | 16 | 0.670073 | 1.535781 | 0.041667 | 0.458756 |
| GOBP_HOMOPHILIC_CELL_ADHESION_VIA_PLASMA_MEMBRANE_ADHESION_MOLECU | 66 | 0.50634 | 1.53359 | 0.036 | 0.457652 |
| GOBP_REGULATION_OF_STEM_CELL_POPULATION_MAINTENANCE | 22 | 0.609381 | 1.531029 | 0.060172 | 0.458139 |
| GOBP_REGULATION_OF_WATER_LOSS_VIA_SKIN | 18 | 0.644692 | 1.529854 | 0.053846 | 0.454838 |
| GOBP_COLUMNAR_CUBOIDAL_EPITHELIAL_CELL_DEVELOPMENT | 24 | 0.60169 | 1.524575 | 0.047486 | 0.463971 |
| GOBP_EYE_PHOTORECEPTOR_CELL_DIFFERENTIATION | 26 | 0.590689 | 1.522671 | 0.053977 | 0.46235 |
| GOBP_MESENCHYMAL_CELL_PROLIFERATION | 15 | 0.666548 | 1.522614 | 0.040506 | 0.455868 |
| GOBP_PYRIDINE_CONTAINING_COMPOUND_BIOSYNTHETIC_PROCESS | 28 | 0.593811 | 1.519247 | 0.032353 | 0.45909 |
| GOBP_LYMPH_VESSEL_DEVELOPMENT | 17 | 0.645136 | 1.518517 | 0.063613 | 0.454694 |
| GOBP_CARDIAC_CHAMBER_DEVELOPMENT | 89 | 0.473279 | 1.517154 | 0.023148 | 0.452021 |
| GOBP_POSITIVE_REGULATION_OF_REPRODUCTIVE_PROCESS | 42 | 0.539651 | 1.515374 | 0.038462 | 0.451084 |
| GOBP_POSITIVE_REGULATION_OF_OSTEOBLAST_DIFFERENTIATION | 37 | 0.543365 | 1.508373 | 0.031746 | 0.465124 |
| GOBP_FACE_DEVELOPMENT | 33 | 0.558021 | 1.480201 | 0.029316 | 0.544556 |
| GOBP_PROTEIN_HYDROXYLATION | 25 | 0.562991 | 1.478804 | 0.085635 | 0.542011 |
| GOBP_CARDIAC_CELL_DEVELOPMENT | 42 | 0.507458 | 1.475586 | 0.039088 | 0.545302 |
| GOBP_MORPHOGENESIS_OF_EMBRYONIC_EPITHELIUM | 96 | 0.452105 | 1.474452 | 0.017937 | 0.541989 |
| GOBP_POSITIVE_REGULATION_OF_INTERLEUKIN_1_PRODUCTION | 48 | 0.512364 | 1.47307 | 0.026846 | 0.539819 |
| GOBP_NAD_BIOSYNTHETIC_PROCESS | 22 | 0.579314 | 1.470785 | 0.058824 | 0.540661 |
| GOBP_REGULATION_OF_STEM_CELL_PROLIFERATION | 22 | 0.585788 | 1.467597 | 0.058355 | 0.544017 |
| GOBP_FOLIC_ACID_METABOLIC_PROCESS | 18 | 0.616882 | 1.464479 | 0.063452 | 0.547445 |
| GOBP_MOLTING_CYCLE | 55 | 0.488943 | 1.453536 | 0.026403 | 0.576212 |
| GOBP_POSITIVE_REGULATION_OF_CYCLIN_DEPENDENT_PROTEIN_KINASE_ACTIVITY | 30 | 0.545863 | 1.450023 | 0.059829 | 0.581395 |
| GOBP_POSITIVE_REGULATION_OF_MUSCLE_CELL_DIFFERENTIATION | 48 | 0.511514 | 1.448411 | 0.054608 | 0.580056 |
| GOBP_CARDIAC_MUSCLE_CELL_DIFFERENTIATION | 60 | 0.479227 | 1.446269 | 0.061224 | 0.580087 |
| GOBP_DIGESTIVE_TRACT_MORPHOGENESIS | 19 | 0.589309 | 1.445655 | 0.084656 | 0.57517 |
| GOBP_PYRIDINE_CONTAINING_COMPOUND_METABOLIC_PROCESS | 33 | 0.533044 | 1.445586 | 0.053371 | 0.568856 |
| GOBP_ESTABLISHMENT_OF_PROTEIN_LOCALIZATION_TO_TELOMERE | 18 | 0.588393 | 1.444064 | 0.073604 | 0.567557 |
| GOBP_CARDIAC_SEPTUM_DEVELOPMENT | 63 | 0.475857 | 1.443671 | 0.050781 | 0.562644 |
| GOBP_NEGATIVE_REGULATION_OF_REPRODUCTIVE_PROCESS | 34 | 0.532101 | 1.43936 | 0.066465 | 0.570336 |
| GOBP_RNA_INTERFERENCE | 16 | 0.620889 | 1.436246 | 0.093909 | 0.573919 |
| GOBP_PINOCYTOSIS | 21 | 0.569777 | 1.435282 | 0.097686 | 0.570646 |
| GOBP_CRISTAE_FORMATION | 35 | 0.524251 | 1.42429 | 0.079179 | 0.600747 |
| GOBP_NEUROTRANSMITTER_RECEPTOR_INTERNALIZATION | 16 | 0.601403 | 1.424228 | 0.102828 | 0.5946 |
| GOBP_SOMATIC_DIVERSIFICATION_OF_IMMUNE_RECEPTORS_VIA_SOMATIC_MUTATIO | 16 | 0.615027 | 1.423353 | 0.104878 | 0.591304 |
| GOBP_WATER_SOLUBLE_VITAMIN_METABOLIC_PROCESS | 73 | 0.46535 | 1.421251 | 0.036437 | 0.592095 |
| GOBP_CELLULAR_MODIFIED_AMINO_ACID_CATABOLIC_PROCESS | 16 | 0.605946 | 1.421071 | 0.110818 | 0.586552 |
| GOBP_APPENDAGE_MORPHOGENESIS | 71 | 0.455068 | 1.414131 | 0.033457 | 0.603794 |
| GOBP_CARDIAC_CHAMBER_MORPHOGENESIS | 60 | 0.480411 | 1.412496 | 0.064639 | 0.602485 |
| GOBP_VERY_LONG_CHAIN_FATTY_ACID_METABOLIC_PROCESS | 25 | 0.55851 | 1.410966 | 0.074074 | 0.601466 |
| GOBP_MITOCHONDRIAL_ATP_SYNTHESIS_COUPLED_PROTON_TRANSPORT | 22 | 0.56003 | 1.410483 | 0.096866 | 0.597267 |
| GOBP_RESPONSE_TO_INTERFERON_BETA | 26 | 0.548206 | 1.395773 | 0.09375 | 0.640882 |
| GOBP_POSITIVE_REGULATION_OF_HEART_GROWTH | 21 | 0.56314 | 1.393968 | 0.104839 | 0.640861 |
| GOBP_ESTABLISHMENT_OR_MAINTENANCE_OF_BIPOLAR_CELL_POLARITY | 34 | 0.509812 | 1.385594 | 0.0721 | 0.66319 |
| GOBP_MAINTENANCE_OF_CELL_NUMBER | 111 | 0.415126 | 1.385229 | 0.03125 | 0.658128 |
| GOBP_SENSORY_PERCEPTION_OF_SMELL | 23 | 0.555692 | 1.385055 | 0.119318 | 0.652598 |
| GOBP_MYOFIBRIL_ASSEMBLY | 36 | 0.503181 | 1.37896 | 0.101796 | 0.667216 |
| GOBP_CARDIAC_SEPTUM_MORPHOGENESIS | 37 | 0.491458 | 1.376081 | 0.092357 | 0.67146 |
| GOBP_REGULATION_OF_REPRODUCTIVE_PROCESS | 91 | 0.429061 | 1.37324 | 0.044898 | 0.675406 |
| GOBP_SKIN_EPIDERMIS_DEVELOPMENT | 44 | 0.484204 | 1.372952 | 0.095941 | 0.670389 |
| GOBP_REGULATION_OF_POSTSYNAPTIC_MEMBRANE_POTENTIAL | 71 | 0.435205 | 1.372494 | 0.061776 | 0.666001 |
| GOBP_MORPHOGENESIS_OF_AN_EPITHELIAL_SHEET | 45 | 0.48243 | 1.36952 | 0.085409 | 0.670862 |
| GOBP_MATING | 23 | 0.546834 | 1.368031 | 0.101648 | 0.670153 |
| GOBP_EYE_PHOTORECEPTOR_CELL_DEVELOPMENT | 19 | 0.573876 | 1.364787 | 0.11831 | 0.675664 |
| GOBP_EAR_MORPHOGENESIS | 60 | 0.452956 | 1.359394 | 0.060714 | 0.688811 |
| GOBP_SIGNAL_TRANSDUCTION_INVOLVED_IN_CELL_CYCLE_CHECKPOINT | 70 | 0.437457 | 1.357508 | 0.037736 | 0.689621 |
| GOBP_GASTRULATION_WITH_MOUTH_FORMING_SECOND | 18 | 0.569874 | 1.357339 | 0.126344 | 0.68428 |
| GOBP_INNER_EAR_MORPHOGENESIS | 49 | 0.469489 | 1.352722 | 0.081356 | 0.695044 |
| GOBP_REGULATION_OF_MORPHOGENESIS_OF_AN_EPITHELIUM | 29 | 0.510705 | 1.343001 | 0.107042 | 0.724267 |
| GOBP_SOMITE_DEVELOPMENT | 53 | 0.450811 | 1.342978 | 0.107383 | 0.718391 |
| GOBP_ATP_SYNTHESIS_COUPLED_PROTON_TRANSPORT | 26 | 0.532669 | 1.341797 | 0.146552 | 0.716674 |
| GOBP_PROTEIN_KINASE_C_SIGNALING | 21 | 0.538751 | 1.339225 | 0.132791 | 0.720276 |
| GOBP_REGULATION_OF_HEART_GROWTH | 41 | 0.468989 | 1.337654 | 0.093373 | 0.719965 |
| GOBP_POSITIVE_REGULATION_OF_T_CELL_MIGRATION | 22 | 0.530391 | 1.335493 | 0.138418 | 0.721987 |
| GOBP_ACTIVATION_OF_JUN_KINASE_ACTIVITY | 35 | 0.476176 | 1.335188 | 0.11875 | 0.717412 |
| GOBP_RIBOSOMAL_LARGE_SUBUNIT_BIOGENESIS | 71 | 0.436865 | 1.334432 | 0.045082 | 0.714394 |
| GOBP_PROTEIN_LOCALIZATION_TO_CHROMOSOME_TELOMERIC_REGION | 30 | 0.49809 | 1.332221 | 0.102804 | 0.716852 |
| GOBP_COTRANSLATIONAL_PROTEIN_TARGETING_TO_MEMBRANE | 102 | 0.40892 | 1.330644 | 0.083333 | 0.716931 |
| GOBP_PHOTORECEPTOR_CELL_DIFFERENTIATION | 37 | 0.482314 | 1.330309 | 0.105096 | 0.712653 |
| GOBP_NEGATIVE_REGULATION_OF_STEROID_METABOLIC_PROCESS | 15 | 0.56686 | 1.329657 | 0.150259 | 0.709611 |
| GOBP_INNER_MITOCHONDRIAL_MEMBRANE_ORGANIZATION | 56 | 0.446915 | 1.328058 | 0.077519 | 0.709757 |
| GOBP_PRODUCTION_OF_SMALL_RNA_INVOLVED_IN_GENE_SILENCING_BY_RNA | 47 | 0.448979 | 1.327181 | 0.07074 | 0.707826 |
| GOBP_POSITIVE_REGULATION_OF_ERYTHROCYTE_DIFFERENTIATION | 29 | 0.507614 | 1.32467 | 0.153846 | 0.711593 |
| GOBP_NEURAL_RETINA_DEVELOPMENT | 40 | 0.463206 | 1.321879 | 0.101818 | 0.716194 |
| GOBP_POSITIVE_REGULATION_OF_TOLL_LIKE_RECEPTOR_SIGNALING_PATHWAY | 21 | 0.53727 | 1.320907 | 0.13079 | 0.714507 |
| GOBP_ANIMAL_ORGAN_FORMATION | 24 | 0.510805 | 1.319928 | 0.157895 | 0.712697 |
| GOBP_EMBRYONIC_APPENDAGE_MORPHOGENESIS | 59 | 0.437266 | 1.319708 | 0.095941 | 0.708245 |
| GOBP_POSITIVE_REGULATION_OF_RESPONSE_TO_ENDOPLASMIC_RETICULUM_STRE | 33 | 0.497918 | 1.317395 | 0.129338 | 0.711424 |
| GOBP_HEMATOPOIETIC_STEM_CELL_PROLIFERATION | 20 | 0.526063 | 1.316912 | 0.151899 | 0.708132 |
| GOBP_COCHLEA_DEVELOPMENT | 24 | 0.520251 | 1.315535 | 0.115819 | 0.70784 |
| GOBP_EMBRYONIC_DIGIT_MORPHOGENESIS | 26 | 0.515361 | 1.315214 | 0.131737 | 0.703869 |
| GOBP_REGULATION_OF_NEURON_DIFFERENTIATION | 108 | 0.398459 | 1.311616 | 0.072115 | 0.711493 |

**GSEA Negative** Hematopoietic/Immune pathways are highlighted in green.

| Gene sets | SIZE | ES | NES | NOM p-val | FDR q-val |
|---|---|---|---|---|---|
| GOBP_POSITIVE_T_CELL_SELECTION | 30 | -0.886485 | -2.043992 | 0 | 0.0073 |
| GOBP_T_CELL_DIFFERENTIATION | 204 | -0.706959 | -2.040995 | 0 | 0.00365 |
| GOBP_T_CELL_SELECTION | 40 | -0.843596 | -2.026661 | 0 | 0.003467 |
| GOBP_T_CELL_ACTIVATION | 383 | -0.661276 | -2.011879 | 0 | 0.003381 |
| GOBP_T_CELL_DIFFERENTIATION_IN_THYMUS | 62 | -0.796837 | -2.011497 | 0 | 0.002705 |
| GOBP_T_CELL_RECEPTOR_SIGNALING_PATHWAY | 186 | -0.696443 | -2.001647 | 0 | 0.002428 |
| GOBP_POSITIVE_REGULATION_OF_INTERLEUKIN_2_PRODUCTION | 30 | -0.869094 | -1.982524 | 0 | 0.003726 |
| GOBP_CYTOLYSIS | 16 | -0.94006 | -1.971139 | 0 | 0.005083 |
| GOBP_MONONUCLEAR_CELL_DIFFERENTIATION | 329 | -0.660593 | -1.967392 | 0 | 0.005332 |
| GOBP_ALPHA_BETA_T_CELL_ACTIVATION | 120 | -0.698146 | -1.933444 | 0 | 0.013771 |
| GOBP_LEUKOCYTE_DIFFERENTIATION | 417 | -0.631443 | -1.928443 | 0 | 0.013561 |
| GOBP_GRANULOCYTE_MIGRATION | 95 | -0.715653 | -1.919634 | 0 | 0.016261 |
| GOBP_POSITIVE_REGULATION_OF_ERK1_AND_ERK2_CASCADE | 118 | -0.695368 | -1.91318 | 0 | 0.018455 |
| GOBP_INTERLEUKIN_2_PRODUCTION | 56 | -0.762183 | -1.911992 | 0.001377 | 0.017436 |
| GOBP_GRANULOCYTE_CHEMOTAXIS | 77 | -0.7178 | -1.895872 | 0 | 0.025108 |
| GOBP_RESPONSE_TO_CHEMOKINE | 51 | -0.762075 | -1.887222 | 0 | 0.030979 |
| GOBP_POSITIVE_REGULATION_OF_HEMOPOIESIS | 125 | -0.676844 | -1.886377 | 0 | 0.03002 |
| GOBP_MYELOID_LEUKOCYTE_MIGRATION | 140 | -0.681915 | -1.882169 | 0 | 0.03161 |
| GOBP_REGULATION_OF_MACROPHAGE_CHEMOTAXIS | 21 | -0.868799 | -1.877691 | 0 | 0.033631 |
| GOBP_AMYLOID_BETA_CLEARANCE | 29 | -0.809181 | -1.876571 | 0 | 0.032681 |
| GOBP_NEGATIVE_REGULATION_OF_INNATE_IMMUNE_RESPONSE | 51 | -0.759212 | -1.873417 | 0.001361 | 0.033412 |
| GOBP_ERK1_AND_ERK2_CASCADE | 195 | -0.650078 | -1.867199 | 0 | 0.037112 |
| GOBP_REGULATION_OF_PHAGOCYTOSIS | 74 | -0.703542 | -1.866893 | 0 | 0.035543 |
| GOBP_REGULATION_OF_T_CELL_ACTIVATION | 265 | -0.628792 | -1.862288 | 0 | 0.038715 |
| GOBP_CELLULAR_DEFENSE_RESPONSE | 36 | -0.794919 | -1.860032 | 0.001462 | 0.039041 |
| GOBP_NEGATIVE_REGULATION_OF_IMMUNE_EFFECTOR_PROCESS | 100 | -0.68571 | -1.858622 | 0 | 0.038626 |
| GOBP_MONONUCLEAR_CELL_MIGRATION | 129 | -0.667517 | -1.855846 | 0 | 0.039594 |
| GOBP_REGULATION_OF_MACROPHAGE_MIGRATION | 29 | -0.822279 | -1.854186 | 0.001553 | 0.03986 |
| GOBP_MACROPHAGE_MIGRATION | 36 | -0.78491 | -1.851247 | 0.001427 | 0.040938 |
| GOBP_POSITIVE_REGULATION_OF_CELL_MATRIX_ADHESION | 39 | -0.763174 | -1.85077 | 0 | 0.039851 |
| GOBP_POSITIVE_REGULATION_OF_LEUKOCYTE_CELL_CELL_ADHESION | 194 | -0.636983 | -1.84993 | 0 | 0.039376 |
| GOBP_POSITIVE_REGULATION_OF_T_CELL_PROLIFERATION | 79 | -0.694515 | -1.843569 | 0 | 0.044087 |
| GOBP_THYMIC_T_CELL_SELECTION | 17 | -0.879913 | -1.841067 | 0 | 0.044712 |
| GOBP_ALPHA_BETA_T_CELL_DIFFERENTIATION | 84 | -0.702416 | -1.839133 | 0 | 0.045123 |
| GOBP_NEGATIVE_REGULATION_OF_IMMUNE_RESPONSE | 112 | -0.675599 | -1.835883 | 0.001271 | 0.046732 |
| GOBP_LEUKOCYTE_CELL_CELL_ADHESION | 293 | -0.619339 | -1.828649 | 0 | 0.052082 |
| GOBP_APOPTOTIC_CELL_CLEARANCE | 41 | -0.760672 | -1.828544 | 0.00146 | 0.050899 |
| GOBP_REGULATION_OF_LEUKOCYTE_DIFFERENTIATION | 222 | -0.622061 | -1.827081 | 0 | 0.050878 |
| GOBP_TAXIS | 403 | -0.604986 | -1.825302 | 0 | 0.05134 |
| GOBP_NEGATIVE_REGULATION_OF_DEFENSE_RESPONSE | 146 | -0.639141 | -1.821389 | 0 | 0.053976 |
| GOBP_CELL_CHEMOTAXIS | 192 | -0.627056 | -1.818719 | 0 | 0.055715 |
| GOBP_POSITIVE_REGULATION_OF_LEUKOCYTE_PROLIFERATION | 116 | -0.657607 | -1.818309 | 0.001224 | 0.054761 |
| GOBP_INFLAMMATORY_RESPONSE | 492 | -0.597094 | -1.812227 | 0 | 0.059606 |
| GOBP_REGULATION_OF_ANTIGEN_RECEPTOR_MEDIATED_SIGNALING_PATHWA | 62 | -0.712735 | -1.812171 | 0.001333 | 0.058322 |
| GOBP_GLUCOSAMINE_CONTAINING_COMPOUND_METABOLIC_PROCESS | 21 | -0.821928 | -1.810121 | 0.006547 | 0.05965 |
| GOBP_MACROPHAGE_CHEMOTAXIS | 27 | -0.797525 | -1.809417 | 0.003175 | 0.05917 |
| GOBP_REGULATION_OF_MONOCYTE_CHEMOTAXIS | 16 | -0.846858 | -1.806253 | 0.0016 | 0.061378 |
| GOBP_POSITIVE_REGULATION_OF_NITRIC_OXIDE_METABOLIC_PROCESS | 31 | -0.780675 | -1.803104 | 0.001513 | 0.063731 |
| GOBP_NEUTROPHIL_MIGRATION | 77 | -0.692837 | -1.801472 | 0 | 0.064203 |
| GOBP_MONOCYTE_CHEMOTAXIS | 32 | -0.773753 | -1.793616 | 0.001497 | 0.071439 |
| GOBP_POSTSYNAPTIC_MEMBRANE_ORGANIZATION | 25 | -0.805897 | -1.792664 | 0.004644 | 0.070898 |
| GOBP_POSITIVE_REGULATION_OF_MACROPHAGE_MIGRATION | 19 | -0.84675 | -1.792093 | 0.00491 | 0.070096 |
| GOBP_POSITIVE_REGULATION_OF_MYELOID_LEUKOCYTE_DIFFERENTIATION | 43 | -0.729266 | -1.79175 | 0.002882 | 0.069049 |
| GOBP_REGULATION_OF_NUCLEASE_ACTIVITY | 22 | -0.817137 | -1.790355 | 0.00625 | 0.069456 |
| GOBP_THYMUS_DEVELOPMENT | 33 | -0.765514 | -1.788961 | 0.004587 | 0.070073 |
| GOBP_LEUKOCYTE_CHEMOTAXIS | 146 | -0.638879 | -1.784172 | 0 | 0.074344 |
| GOBP_NEGATIVE_REGULATION_OF_RESPONSE_TO_BIOTIC_STIMULUS | 79 | -0.676466 | -1.78331 | 0 | 0.074066 |
| GOBP_T_CELL_PROLIFERATION | 159 | -0.629112 | -1.779037 | 0 | 0.07767 |
| GOBP_REGULATION_OF_LYMPHOCYTE_MEDIATED_IMMUNITY | 128 | -0.645432 | -1.777074 | 0 | 0.078688 |
| GOBP_POSITIVE_REGULATION_OF_PHAGOCYTOSIS | 55 | -0.71401 | -1.775754 | 0.001381 | 0.079134 |
| GOBP_NEGATIVE_REGULATION_OF_ANTIGEN_RECEPTOR_MEDIATED_SIGNALIN | 28 | -0.787345 | -1.773679 | 0.004464 | 0.080406 |
| GOBP_REGULATION_OF_DEFENSE_RESPONSE | 497 | -0.576511 | -1.772559 | 0 | 0.080628 |
| GOBP_PYROPTOSIS | 17 | -0.850561 | -1.772019 | 0.001522 | 0.080012 |
| GOBP_V_D_J_RECOMBINATION | 15 | -0.88276 | -1.769949 | 0.005155 | 0.081229 |
| GOBP_MATURE_B_CELL_DIFFERENTIATION | 24 | -0.798256 | -1.769389 | 0.001629 | 0.080702 |
| GOBP_ENTRY_INTO_HOST | 125 | -0.642971 | -1.767148 | 0 | 0.081978 |
| GOBP_DOSAGE_COMPENSATION | 17 | -0.838647 | -1.766953 | 0.004754 | 0.081019 |
| GOBP_REGULATION_OF_CELL_MATRIX_ADHESION | 89 | -0.654318 | -1.764522 | 0 | 0.082469 |
| GOBP_REGULATION_OF_LEUKOCYTE_MIGRATION | 149 | -0.621743 | -1.75948 | 0 | 0.087197 |
| GOBP_MONOCYTE_DIFFERENTIATION | 28 | -0.78017 | -1.759247 | 0.003072 | 0.08616 |
| GOBP_REGULATION_OF_T_HELPER_1_TYPE_IMMUNE_RESPONSE | 20 | -0.803316 | -1.745442 | 0.0067 | 0.103163 |
| GOBP_REGULATION_OF_ACTIN_FILAMENT_ORGANIZATION | 215 | -0.594562 | -1.744889 | 0 | 0.102528 |
| GOBP_NEGATIVE_REGULATION_OF_IMMUNE_SYSTEM_PROCESS | 299 | -0.583808 | -1.743528 | 0 | 0.102998 |
| GOBP_VASCULAR_TRANSPORT | 59 | -0.693999 | -1.7408 | 0.001342 | 0.105599 |
| GOBP_CHOLESTEROL_STORAGE | 16 | -0.835419 | -1.739503 | 0.006547 | 0.1061 |
| GOBP_POSITIVE_REGULATION_OF_DNA_BINDING | 42 | -0.73232 | -1.738927 | 0.005806 | 0.105514 |
| GOBP_POSITIVE_REGULATION_OF_LEUKOCYTE_MEDIATED_IMMUNITY | 105 | -0.646515 | -1.738025 | 0.001284 | 0.105489 |
| GOBP_NEUTROPHIL_CHEMOTAXIS | 62 | -0.693902 | -1.737633 | 0 | 0.104577 |
| GOBP_ACTIVATION_OF_IMMUNE_RESPONSE | 410 | -0.571514 | -1.736606 | 0 | 0.104602 |
| GOBP_ANTIGEN_RECEPTOR_MEDIATED_SIGNALING_PATHWAY | 256 | -0.590432 | -1.736418 | 0 | 0.103619 |
| GOBP_VASCULAR_ENDOTHELIAL_GROWTH_FACTOR_PRODUCTION | 29 | -0.76855 | -1.736217 | 0.002886 | 0.102598 |
| GOBP_REGULATION_OF_GRANULOCYTE_CHEMOTAXIS | 39 | -0.724014 | -1.734187 | 0.004354 | 0.104202 |
| GOBP_PHOSPHOLIPASE_C_ACTIVATING_G_PROTEIN_COUPLED_RECEPTOR_SI | 42 | -0.708839 | -1.733463 | 0.004386 | 0.103956 |
| GOBP_POSITIVE_REGULATION_OF_FILOPODIUM_ASSEMBLY | 25 | -0.786901 | -1.732164 | 0.00939 | 0.10492 |
| GOBP_REGULATION_OF_INFLAMMATORY_RESPONSE | 228 | -0.594674 | -1.727632 | 0 | 0.110003 |
| GOBP_CELL_KILLING | 110 | -0.626693 | -1.726222 | 0 | 0.110714 |
| GOBP_REGULATION_OF_MONOCYTE_DIFFERENTIATION | 17 | -0.83814 | -1.724674 | 0.004878 | 0.111617 |
| GOBP_REGULATION_OF_LYMPHOCYTE_DIFFERENTIATION | 142 | -0.616623 | -1.724286 | 0 | 0.110871 |
| GOBP_REACTIVE_NITROGEN_SPECIES_METABOLIC_PROCESS | 61 | -0.680604 | -1.72301 | 0.001393 | 0.111045 |
| GOBP_NEGATIVE_REGULATION_OF_LEUKOCYTE_MIGRATION | 26 | -0.769952 | -1.722671 | 0.006279 | 0.110194 |
| GOBP_NATURAL_KILLER_CELL_MEDIATED_IMMUNITY | 51 | -0.6888 | -1.722217 | 0.002869 | 0.109649 |
| GOBP_ASTROCYTE_DEVELOPMENT | 30 | -0.748747 | -1.721757 | 0.005908 | 0.109115 |
| GOBP_REGULATION_OF_DEFENSE_RESPONSE_TO_VIRUS_BY_VIRUS | 28 | -0.765499 | -1.718173 | 0.00905 | 0.112975 |
| GOBP_REGULATION_OF_T_CELL_RECEPTOR_SIGNALING_PATHWAY | 39 | -0.714389 | -1.717479 | 0.011461 | 0.112875 |
| GOBP_MOVEMENT_IN_HOST_ENVIRONMENT | 145 | -0.607244 | -1.717254 | 0 | 0.11196 |
| GOBP_REGULATION_OF_HEMOPOIESIS | 327 | -0.570996 | -1.716291 | 0 | 0.112174 |
| GOBP_POSITIVE_REGULATION_OF_VASCULAR_ENDOTHELIAL_GROWTH_FACTO | 21 | -0.797606 | -1.715569 | 0.00625 | 0.111857 |
| GOBP_REGULATION_OF_VASCULATURE_DEVELOPMENT | 189 | -0.597578 | -1.713512 | 0 | 0.113615 |
| GOBP_CD8_POSITIVE_ALPHA_BETA_T_CELL_ACTIVATION | 24 | -0.770449 | -1.712495 | 0.00936 | 0.113913 |
| GOBP_POSITIVE_REGULATION_OF_LEUKOCYTE_MIGRATION | 100 | -0.641673 | -1.710917 | 0.003793 | 0.114788 |
| GOBP_MULTI_ORGANISM_CELLULAR_PROCESS | 15 | -0.815067 | -1.70828 | 0.006452 | 0.117491 |
| GOBP_REGULATION_OF_B_CELL_MEDIATED_IMMUNITY | 50 | -0.69285 | -1.707918 | 0.004172 | 0.116819 |
| GOBP_POSITIVE_REGULATION_OF_CELL_ADHESION | 324 | -0.564278 | -1.707541 | 0 | 0.116232 |
| GOBP_REGULATION_OF_NITRIC_OXIDE_METABOLIC_PROCESS | 48 | -0.699386 | -1.704824 | 0.002833 | 0.118503 |
| GOBP_POSITIVE_REGULATION_OF_CELL_CELL_ADHESION | 220 | -0.58104 | -1.704674 | 0 | 0.117524 |
| GOBP_POSITIVE_REGULATION_OF_REACTIVE_OXYGEN_SPECIES_BIOSYNTHETI | 42 | -0.712572 | -1.704345 | 0.011429 | 0.116769 |
| GOBP_POSITIVE_REGULATION_OF_B_CELL_MEDIATED_IMMUNITY | 36 | -0.714563 | -1.7042 | 0.010145 | 0.11595 |
| GOBP_BILE_ACID_AND_BILE_SALT_TRANSPORT | 18 | -0.802954 | -1.703971 | 0.004854 | 0.115156 |
| GOBP_NEGATIVE_REGULATION_OF_RESPONSE_TO_EXTERNAL_STIMULUS | 242 | -0.586082 | -1.703614 | 0 | 0.114569 |
| GOBP_REGULATION_OF_MACROPHAGE_DERIVED_FOAM_CELL_DIFFERENTIATIO | 20 | -0.783656 | -1.702267 | 0.004823 | 0.115217 |
| GOBP_REGULATION_OF_CYSTEINE_TYPE_ENDOPEPTIDASE_ACTIVITY_INVOLVE | 16 | -0.821975 | -1.702097 | 0.016717 | 0.114471 |
| GOBP_LEUKOCYTE_PROLIFERATION | 242 | -0.579761 | -1.700117 | 0 | 0.116281 |
| GOBP_POSITIVE_REGULATION_OF_RESPONSE_TO_EXTERNAL_STIMULUS | 368 | -0.568672 | -1.699921 | 0 | 0.115529 |
| GOBP_REGULATION_OF_ADAPTIVE_IMMUNE_RESPONSE | 143 | -0.603703 | -1.699395 | 0 | 0.115184 |
| GOBP_POSITIVE_REGULATION_OF_CELL_KILLING | 49 | -0.691765 | -1.697707 | 0.004286 | 0.116199 |
| GOBP_REGULATION_OF_NATURAL_KILLER_CELL_MEDIATED_IMMUNITY | 36 | -0.724253 | -1.697608 | 0.005839 | 0.115288 |
| GOBP_NERVE_DEVELOPMENT | 35 | -0.730932 | -1.697463 | 0.006163 | 0.114526 |
| GOBP_REGULATION_OF_B_CELL_RECEPTOR_SIGNALING_PATHWAY | 25 | -0.745471 | -1.69736 | 0.012678 | 0.11367 |
| GOBP_REGULATION_OF_LEUKOCYTE_PROLIFERATION | 189 | -0.593663 | -1.697121 | 0 | 0.113136 |
| GOBP_IMMUNE_RESPONSE_REGULATING_SIGNALING_PATHWAY | 385 | -0.559982 | -1.696994 | 0 | 0.112332 |
| GOBP_POSITIVE_REGULATION_OF_CELL_SUBSTRATE_ADHESION | 81 | -0.631115 | -1.696209 | 0.002621 | 0.112344 |
| GOBP_REGULATION_OF_INNATE_IMMUNE_RESPONSE | 261 | -0.570952 | -1.691995 | 0 | 0.116764 |
| GOBP_REGULATION_OF_ALPHA_BETA_T_CELL_ACTIVATION | 83 | -0.640934 | -1.691156 | 0.007833 | 0.116935 |
| GOBP_REGULATION_OF_CELL_KILLING | 73 | -0.658733 | -1.691115 | 0.002759 | 0.116034 |
| GOBP_REGULATION_OF_FILOPODIUM_ASSEMBLY | 37 | -0.711242 | -1.69083 | 0.008523 | 0.115623 |
| GOBP_RESPONSE_TO_INTERLEUKIN_1 | 158 | -0.595455 | -1.690619 | 0 | 0.114996 |
| GOBP_REGULATION_OF_DNA_BINDING | 89 | -0.633722 | -1.68975 | 0.00129 | 0.115193 |
| GOBP_REGULATION_OF_T_CELL_DIFFERENTIATION | 118 | -0.615266 | -1.689259 | 0.002525 | 0.11488 |
| GOBP_POSITIVE_REGULATION_OF_LYMPHOCYTE_DIFFERENTIATION | 85 | -0.632248 | -1.689066 | 0.001325 | 0.114273 |
| GOBP_ADAPTIVE_IMMUNE_RESPONSE | 404 | -0.559362 | -1.688229 | 0 | 0.114639 |
| GOBP_POSITIVE_REGULATION_OF_DEFENSE_RESPONSE | 269 | -0.571707 | -1.68732 | 0 | 0.114943 |
| GOBP_REGULATION_OF_SYNCYTIUM_FORMATION_BY_PLASMA_MEMBRANE_FU | 18 | -0.796083 | -1.686864 | 0.009756 | 0.114515 |
| GOBP_EXTRACELLULAR_MATRIX_DISASSEMBLY | 55 | -0.669174 | -1.685812 | 0.002762 | 0.11487 |
| GOBP_REGULATION_OF_LEUKOCYTE_MEDIATED_CYTOTOXICITY | 61 | -0.660103 | -1.683824 | 0.006925 | 0.116515 |
| GOBP_POSITIVE_REGULATION_OF_VASCULATURE_DEVELOPMENT | 105 | -0.621211 | -1.682854 | 0.002478 | 0.116726 |
| GOBP_POSITIVE_REGULATION_OF_CYTOSOLIC_CALCIUM_ION_CONCENTRATIO | 19 | -0.781457 | -1.681366 | 0.008197 | 0.117841 |
| GOBP_COLLAGEN_CATABOLIC_PROCESS | 29 | -0.741345 | -1.681099 | 0.004622 | 0.117354 |
| GOBP_DENDRITIC_CELL_MIGRATION | 24 | -0.751377 | -1.680593 | 0.012539 | 0.116988 |
| GOBP_PLACENTA_BLOOD_VESSEL_DEVELOPMENT | 19 | -0.787604 | -1.679526 | 0.011236 | 0.117437 |
| GOBP_NEGATIVE_REGULATION_OF_INTERLEUKIN_12_PRODUCTION | 16 | -0.807821 | -1.678216 | 0.015437 | 0.118315 |
| GOBP_POSITIVE_REGULATION_OF_REACTIVE_OXYGEN_SPECIES_METABOLIC_F | 74 | -0.644661 | -1.677629 | 0.002695 | 0.11821 |
| GOBP_BLOOD_VESSEL_MORPHOGENESIS | 397 | -0.554463 | -1.677062 | 0 | 0.118024 |
| GOBP_MACROPHAGE_DIFFERENTIATION | 33 | -0.718574 | -1.676987 | 0.007634 | 0.117286 |

| | | | | | |
|---|---|---|---|---|---|
| GOBP_MATURATION_OF_LSU_RRNA_FROM_TRICISTRONIC_RRNA_TRANSCRIPT_SSU_ | 15 | 0.562548 | 1.311282 | 0.173028 | 0.707745 |
| GOBP_REGULATION_OF_NEURAL_PRECURSOR_CELL_PROLIFERATION | 48 | 0.451405 | 1.309809 | 0.109155 | 0.7081 |
| GOBP_LIMBIC_SYSTEM_DEVELOPMENT | 52 | 0.440852 | 1.306572 | 0.105263 | 0.714231 |
| GOBP_NEURAL_TUBE_FORMATION | 78 | 0.411532 | 1.303974 | 0.084871 | 0.718344 |
| GOBP_RETROGRADE_AXONAL_TRANSPORT | 18 | 0.552706 | 1.300703 | 0.161202 | 0.724534 |
| GOBP_FATTY_ACYL_COA_BIOSYNTHETIC_PROCESS | 27 | 0.489649 | 1.297769 | 0.130178 | 0.730065 |
| GOBP_POSITIVE_REGULATION_OF_PEPTIDYL_SERINE_PHOSPHORYLATION | 72 | 0.423083 | 1.293925 | 0.09375 | 0.7386 |
| GOBP_CYTOPLASMIC_TRANSLATIONAL_INITIATION | 34 | 0.476956 | 1.292116 | 0.131343 | 0.739567 |
| GOBP_NEGATIVE_REGULATION_OF_ENDOTHELIAL_CELL_APOPTOTIC_PROCESS | 17 | 0.531403 | 1.291344 | 0.173697 | 0.737322 |
| GOBP_CELLULAR_RESPONSE_TO_CALCIUM_ION | 56 | 0.442768 | 1.290644 | 0.08 | 0.735042 |
| GOBP_AMINO_ACID_ACTIVATION | 49 | 0.443059 | 1.289722 | 0.111111 | 0.733387 |
| GOBP_ENDOTHELIUM_DEVELOPMENT | 88 | 0.399479 | 1.289089 | 0.065844 | 0.730773 |
| GOBP_RHODOPSIN_MEDIATED_SIGNALING_PATHWAY | 17 | 0.536121 | 1.28902 | 0.197861 | 0.726373 |
| GOBP_REGULATION_OF_HIPPO_SIGNALING | 16 | 0.558331 | 1.285125 | 0.18984 | 0.735545 |
| GOBP_HEART_GROWTH | 54 | 0.427979 | 1.28434 | 0.087121 | 0.733504 |
| GOBP_ANTERIOR_POSTERIOR_AXIS_SPECIFICATION | 24 | 0.50375 | 1.283734 | 0.145897 | 0.731123 |
| GOBP_PEPTIDYL_PROLINE_MODIFICATION | 48 | 0.444884 | 1.283636 | 0.132258 | 0.72684 |
| GOBP_TRANSLATIONAL_TERMINATION | 104 | 0.391165 | 1.283104 | 0.068273 | 0.724155 |
| GOBP_DOPAMINERGIC_NEURON_DIFFERENTIATION | 16 | 0.567275 | 1.279578 | 0.2 | 0.731623 |
| GOBP_HIPPOCAMPUS_DEVELOPMENT | 43 | 0.453848 | 1.278365 | 0.12462 | 0.731264 |
| GOBP_IONOTROPIC_GLUTAMATE_RECEPTOR_SIGNALING_PATHWAY | 17 | 0.544664 | 1.274646 | 0.162291 | 0.739625 |
| GOBP_LONG_CHAIN_FATTY_ACID_BIOSYNTHETIC_PROCESS | 25 | 0.494629 | 1.273329 | 0.15493 | 0.739759 |
| GOBP_THIOESTER_METABOLIC_PROCESS | 89 | 0.395813 | 1.27072 | 0.095023 | 0.744588 |
| GOBP_AROMATIC_AMINO_ACID_FAMILY_METABOLIC_PROCESS | 23 | 0.504845 | 1.270322 | 0.168865 | 0.741494 |
| GOBP_LUNG_MORPHOGENESIS | 27 | 0.482322 | 1.260243 | 0.190736 | 0.772003 |
| GOBP_REGULATION_OF_CELL_CYCLE_CHECKPOINT | 30 | 0.466854 | 1.25947 | 0.152589 | 0.770216 |
| GOBP_DNA_METHYLATION_DEPENDENT_HETEROCHROMATIN_ASSEMBLY | 19 | 0.52133 | 1.259086 | 0.204082 | 0.767157 |
| GOBP_COENZYME_A_METABOLIC_PROCESS | 15 | 0.538679 | 1.258522 | 0.19 | 0.764487 |
| GOBP_FC_EPSILON_RECEPTOR_SIGNALING_PATHWAY | 127 | 0.374333 | 1.258096 | 0.087209 | 0.761467 |
| GOBP_REGULATION_OF_STEROID_BIOSYNTHETIC_PROCESS | 65 | 0.414329 | 1.253182 | 0.143939 | 0.774955 |
| GOBP_PITUITARY_GLAND_DEVELOPMENT | 15 | 0.529543 | 1.251966 | 0.19898 | 0.775102 |
| GOBP_ESTABLISHMENT_OF_PROTEIN_LOCALIZATION_TO_CHROMOSOME | 26 | 0.493345 | 1.251144 | 0.161111 | 0.773503 |
| GOBP_REGULATION_OF_PEPTIDYL_SERINE_PHOSPHORYLATION | 101 | 0.379477 | 1.250364 | 0.110553 | 0.771977 |
| GOBP_FC_RECEPTOR_MEDIATED_STIMULATORY_SIGNALING_PATHWAY | 99 | 0.381826 | 1.250097 | 0.126761 | 0.768524 |
| GOBP_ROOF_OF_MOUTH_DEVELOPMENT | 50 | 0.427077 | 1.249838 | 0.143345 | 0.765162 |
| GOBP_NEGATIVE_REGULATION_OF_UBIQUITIN_PROTEIN_TRANSFERASE_ACTIVITY | 16 | 0.528457 | 1.249315 | 0.21123 | 0.762739 |
| GOBP_T_CELL_CHEMOTAXIS | 17 | 0.507929 | 1.2493 | 0.197861 | 0.758541 |
| GOBP_ESTABLISHMENT_OF_PROTEIN_LOCALIZATION_TO_ENDOPLASMIC_RETICULUM | 115 | 0.377678 | 1.249268 | 0.090909 | 0.754448 |
| GOBP_PYRIMIDINE_NUCLEOSIDE_TRIPHOSPHATE_BIOSYNTHETIC_PROCESS | 17 | 0.524524 | 1.247499 | 0.202899 | 0.756252 |
| GOBP_IMMUNOGLOBULIN_PRODUCTION | 110 | 0.372999 | 1.246799 | 0.089286 | 0.754638 |
| GOBP_REGULATION_OF_HEMATOPOIETIC_STEM_CELL_DIFFERENTIATION | 70 | 0.398035 | 1.242678 | 0.100806 | 0.764721 |
| GOBP_MUSCLE_TISSUE_DEVELOPMENT | 219 | 0.344629 | 1.242027 | 0.026549 | 0.762546 |
| GOBP_SYNAPTIC_VESICLE_PRIMING | 15 | 0.531025 | 1.241798 | 0.233032 | 0.759279 |
| GOBP_POSITIVE_REGULATION_OF_WNT_SIGNALING_PATHWAY | 137 | 0.363308 | 1.239346 | 0.08589 | 0.764051 |
| GOBP_HIPPO_SIGNALING | 35 | 0.447126 | 1.239303 | 0.177632 | 0.760137 |
| GOBP_PRODUCTION_OF_MOLECULAR_MEDIATOR_OF_IMMUNE_RESPONSE | 177 | 0.353261 | 1.238144 | 0.080745 | 0.759863 |
| GOBP_SOMATIC_STEM_CELL_POPULATION_MAINTENANCE | 46 | 0.430976 | 1.237361 | 0.16 | 0.758642 |
| GOBP_POSITIVE_REGULATION_OF_ORGAN_GROWTH | 27 | 0.485353 | 1.236485 | 0.174033 | 0.757717 |
| GOBP_EMBRYONIC_AXIS_SPECIFICATION | 19 | 0.516855 | 1.236442 | 0.201058 | 0.753908 |
| GOBP_RESPONSE_TO_INTERFERON_ALPHA | 21 | 0.491326 | 1.235607 | 0.195187 | 0.752952 |
| GOBP_MITOCHONDRIAL_ELECTRON_TRANSPORT_CYTOCHROME_C_TO_OXYGEN | 19 | 0.515836 | 1.233439 | 0.220109 | 0.756683 |
| GOBP_APOPTOTIC_DNA_FRAGMENTATION | 19 | 0.511283 | 1.233306 | 0.220386 | 0.753204 |
| GOBP_CEREBELLAR_CORTEX_MORPHOGENESIS | 20 | 0.495558 | 1.233147 | 0.220708 | 0.749852 |
| GOBP_NEURAL_TUBE_DEVELOPMENT | 108 | 0.368966 | 1.232033 | 0.119048 | 0.749919 |
| GOBP_DNA_STRAND_ELONGATION | 26 | 0.475744 | 1.231473 | 0.214502 | 0.748059 |
| GOBP_NEGATIVE_REGULATION_OF_LYASE_ACTIVITY | 15 | 0.525956 | 1.22863 | 0.232911 | 0.753729 |
| GOBP_PROTEIN_LOCALIZATION_TO_SYNAPSE | 55 | 0.417375 | 1.225772 | 0.127273 | 0.759593 |
| GOBP_ALPHA_AMINO_ACID_BIOSYNTHETIC_PROCESS | 55 | 0.418932 | 1.225287 | 0.157143 | 0.757381 |
| GOBP_MODULATION_OF_EXCITATORY_POSTSYNAPTIC_POTENTIAL | 26 | 0.480822 | 1.224711 | 0.196237 | 0.755635 |
| GOBP_ESTABLISHMENT_OF_EPITHELIAL_CELL_POLARITY | 21 | 0.505106 | 1.224394 | 0.176471 | 0.752988 |
| GOBP_MAMMARY_GLAND_MORPHOGENESIS | 24 | 0.490171 | 1.220874 | 0.206304 | 0.761237 |
| GOBP_REGULATION_OF_T_CELL_MIGRATION | 32 | 0.443898 | 1.220344 | 0.20068 | 0.759272 |
| GOBP_REGULATION_OF_STRIATED_MUSCLE_CELL_DIFFERENTIATION | 54 | 0.418462 | 1.219386 | 0.179211 | 0.758968 |
| GOBP_LEUKOTRIENE_METABOLIC_PROCESS | 18 | 0.516175 | 1.218201 | 0.233516 | 0.759549 |
| GOBP_ADENYLATE_CYCLASE_INHIBITING_G_PROTEIN_COUPLED_RECEPTOR_SIGNAL | 40 | 0.440519 | 1.216401 | 0.194079 | 0.762144 |
| GOBP_REGULATION_OF_ANIMAL_ORGAN_MORPHOGENESIS | 121 | 0.363748 | 1.216132 | 0.115385 | 0.759573 |
| GOBP_STRIATED_MUSCLE_ADAPTATION | 27 | 0.457905 | 1.2139 | 0.1875 | 0.763726 |
| GOBP_ANTIBACTERIAL_HUMORAL_RESPONSE | 22 | 0.487219 | 1.211988 | 0.21447 | 0.766794 |
| GOBP_FATTY_ACYL_COA_METABOLIC_PROCESS | 37 | 0.451064 | 1.209556 | 0.200647 | 0.771539 |
| GOBP_RETINA_DEVELOPMENT_IN_CAMERA_TYPE_EYE | 89 | 0.371146 | 1.206296 | 0.14978 | 0.778961 |
| GOBP_DEOXYRIBONUCLEOSIDE_TRIPHOSPHATE_METABOLIC_PROCESS | 15 | 0.523647 | 1.20528 | 0.254593 | 0.778967 |
| GOBP_NEURON_MATURATION | 30 | 0.456423 | 1.205104 | 0.217391 | 0.776047 |
| GOBP_POSITIVE_REGULATION_OF_TRANSCRIPTION_OF_NOTCH_RECEPTOR_TARGET | 16 | 0.527098 | 1.204326 | 0.237598 | 0.775085 |
| GOBP_PROTEIN_LOCALIZATION_TO_CELL_JUNCTION | 80 | 0.373679 | 1.203071 | 0.148148 | 0.775888 |
| GOBP_REGULATION_OF_NON_CANONICAL_WNT_SIGNALING_PATHWAY | 18 | 0.505409 | 1.202789 | 0.216495 | 0.773253 |
| GOBP_VIRAL_BUDDING | 25 | 0.468103 | 1.201766 | 0.22093 | 0.773183 |
| GOBP_NEGATIVE_REGULATION_OF_NEURON_DIFFERENTIATION | 31 | 0.446418 | 1.200927 | 0.245014 | 0.772623 |
| GOBP_APOPTOTIC_PROCESS_INVOLVED_IN_DEVELOPMENT | 21 | 0.480213 | 1.199755 | 0.238227 | 0.773042 |
| GOBP_REGULATION_OF_VASCULAR_PERMEABILITY | 24 | 0.468124 | 1.198904 | 0.22293 | 0.7724 |
| GOBP_NUCLEAR_TRANSCRIBED_MRNA_CATABOLIC_PROCESS_NONSENSE_MEDIATE | 120 | 0.351679 | 1.197349 | 0.156863 | 0.774112 |
| GOBP_COPII_COATED_VESICLE_CARGO_LOADING | 15 | 0.517758 | 1.197003 | 0.257143 | 0.771796 |
| GOBP_ALPHA_AMINO_ACID_METABOLIC_PROCESS | 150 | 0.342071 | 1.195047 | 0.082902 | 0.774877 |
| GOBP_POSITIVE_REGULATION_OF_ENDOTHELIAL_CELL_MIGRATION | 80 | 0.367166 | 1.194887 | 0.170732 | 0.771978 |
| GOBP_NUCLEOTIDE_BINDING_OLIGOMERIZATION_DOMAIN_CONTAINING_2_SIGNALING | 16 | 0.51853 | 1.19407 | 0.262211 | 0.771362 |
| GOBP_NLS_BEARING_PROTEIN_IMPORT_INTO_NUCLEUS | 16 | 0.525665 | 1.193668 | 0.230769 | 0.769391 |
| GOBP_REGULATION_OF_ORGAN_GROWTH | 60 | 0.390085 | 1.192636 | 0.149091 | 0.769482 |
| GOBP_PYRIMIDINE_NUCLEOSIDE_TRIPHOSPHATE_METABOLIC_PROCESS | 22 | 0.473605 | 1.192527 | 0.243243 | 0.766489 |
| GOBP_APPENDAGE_DEVELOPMENT | 93 | 0.373296 | 1.192476 | 0.147679 | 0.763327 |
| GOBP_TRICARBOXYLIC_ACID_CYCLE | 31 | 0.441828 | 1.191878 | 0.209726 | 0.761963 |
| GOBP_CENTRAL_NERVOUS_SYSTEM_NEURON_DEVELOPMENT | 45 | 0.414335 | 1.190799 | 0.198653 | 0.762342 |
| GOBP_SOMITOGENESIS | 39 | 0.419531 | 1.188549 | 0.222222 | 0.766637 |
| GOBP_FEMALE_SEX_DIFFERENTIATION | 68 | 0.385438 | 1.186785 | 0.182156 | 0.769434 |
| GOBP_EMBRYONIC_SKELETAL_SYSTEM_DEVELOPMENT | 60 | 0.390835 | 1.186395 | 0.200692 | 0.767446 |
| GOBP_NUCLEOSIDE_BISPHOSPHATE_METABOLIC_PROCESS | 112 | 0.358645 | 1.183579 | 0.14218 | 0.773678 |
| GOBP_EXCITATORY_SYNAPSE_ASSEMBLY | 15 | 0.530357 | 1.182905 | 0.255102 | 0.772806 |
| GOBP_ADHERENS_JUNCTION_ORGANIZATION | 39 | 0.419343 | 1.181411 | 0.229358 | 0.774647 |
| GOBP_MAMMARY_GLAND_DUCT_MORPHOGENESIS | 18 | 0.499762 | 1.180002 | 0.253333 | 0.776001 |
| GOBP_MONOCARBOXYLIC_ACID_CATABOLIC_PROCESS | 103 | 0.357921 | 1.178361 | 0.138756 | 0.778166 |
| GOBP_FATTY_ACID_DERIVATIVE_BIOSYNTHETIC_PROCESS | 42 | 0.416849 | 1.17648 | 0.169329 | 0.781289 |
| GOBP_SENSORY_PERCEPTION_OF_MECHANICAL_STIMULUS | 105 | 0.351814 | 1.175631 | 0.116279 | 0.780927 |
| GOBP_GLAND_MORPHOGENESIS | 67 | 0.370061 | 1.174622 | 0.206349 | 0.781093 |
| GOBP_PHOTORECEPTOR_CELL_DEVELOPMENT | 29 | 0.445658 | 1.172533 | 0.25072 | 0.785009 |
| GOBP_FORMATION_OF_CYTOPLASMIC_TRANSLATION_INITIATION_COMPLEX | 16 | 0.496495 | 1.172149 | 0.274074 | 0.783188 |
| GOBP_MITOCHONDRIAL_TRANSLATION | 133 | 0.348825 | 1.171763 | 0.154639 | 0.781202 |
| GOBP_POSITIVE_REGULATION_OF_TRANSCRIPTION_FROM_RNA_POLYMERASE_II_PR | 20 | 0.472105 | 1.170557 | 0.297521 | 0.782011 |
| GOBP_PEROXISOME_ORGANIZATION | 76 | 0.372401 | 1.170434 | 0.20438 | 0.779285 |
| GOBP_MITOCHONDRIAL_TRANSLATIONAL_TERMINATION | 88 | 0.367981 | 1.168202 | 0.140351 | 0.783591 |
| GOBP_TETRAPYRROLE_BIOSYNTHETIC_PROCESS | 27 | 0.453532 | 1.163959 | 0.248447 | 0.794542 |
| GOBP_LUNG_EPITHELIUM_DEVELOPMENT | 15 | 0.50975 | 1.163233 | 0.283465 | 0.793802 |
| GOBP_POSITIVE_REGULATION_OF_NEURON_DEATH | 73 | 0.369456 | 1.162383 | 0.181452 | 0.793654 |
| GOBP_CONNECTIVE_TISSUE_DEVELOPMENT | 140 | 0.339758 | 1.159744 | 0.146067 | 0.799455 |
| GOBP_PROTEIN_LOCALIZATION_TO_ENDOPLASMIC_RETICULUM | 139 | 0.340618 | 1.157116 | 0.183246 | 0.805081 |
| GOBP_MORPHOGENESIS_OF_A_BRANCHING_STRUCTURE | 101 | 0.356761 | 1.156108 | 0.220339 | 0.805402 |
| GOBP_NEGATIVE_REGULATION_OF_VIRAL_PROCESS | 73 | 0.369579 | 1.155511 | 0.215328 | 0.804208 |
| GOBP_DIGESTIVE_SYSTEM_DEVELOPMENT | 65 | 0.373536 | 1.154634 | 0.214008 | 0.803961 |
| GOBP_REGULATION_OF_TRANSLATIONAL_FIDELITY | 20 | 0.476064 | 1.152822 | 0.298734 | 0.806892 |
| GOBP_REGULATION_OF_BMP_SIGNALING_PATHWAY | 48 | 0.400608 | 1.152118 | 0.204152 | 0.806202 |
| GOBP_POSITIVE_REGULATION_OF_TELOMERASE_RNA_LOCALIZATION_TO_CAJAL_BOI | 15 | 0.494769 | 1.149595 | 0.32 | 0.811556 |
| GOBP_ACTIVATION_OF_MAPK_ACTIVITY | 116 | 0.345461 | 1.14915 | 0.191176 | 0.809858 |
| GOBP_SERINE_FAMILY_AMINO_ACID_BIOSYNTHETIC_PROCESS | 17 | 0.478194 | 1.147289 | 0.280928 | 0.813322 |
| GOBP_WNT_SIGNALING_PATHWAY_CALCIUM_MODULATING_PATHWAY | 32 | 0.429451 | 1.145868 | 0.263804 | 0.815125 |
| GOBP_NAD_METABOLIC_PROCESS | 47 | 0.394437 | 1.144634 | 0.227106 | 0.81616 |
| GOBP_MESENCHYME_MORPHOGENESIS | 23 | 0.460842 | 1.14397 | 0.300268 | 0.815314 |
| GOBP_PROSTATE_GLAND_DEVELOPMENT | 21 | 0.463521 | 1.1427 | 0.279006 | 0.816291 |
| GOBP_RESPONSE_TO_DEXAMETHASONE | 27 | 0.438177 | 1.141842 | 0.293578 | 0.81613 |
| GOBP_CEREBELLAR_CORTEX_DEVELOPMENT | 30 | 0.432771 | 1.137976 | 0.302857 | 0.826454 |
| GOBP_REGULATION_OF_CARTILAGE_DEVELOPMENT | 32 | 0.430482 | 1.136615 | 0.295598 | 0.828015 |
| GOBP_VIRAL_RELEASE_FROM_HOST_CELL | 29 | 0.422949 | 1.135284 | 0.268882 | 0.829546 |
| GOBP_REGULATION_OF_CHROMATIN_ASSEMBLY | 19 | 0.471051 | 1.133201 | 0.292479 | 0.833754 |
| GOBP_FEEDING_BEHAVIOR | 36 | 0.420873 | 1.131712 | 0.291793 | 0.83599 |
| GOBP_STRIATED_MUSCLE_CELL_DEVELOPMENT | 63 | 0.381043 | 1.127394 | 0.24498 | 0.847494 |
| GOBP_WATER_HOMEOSTASIS | 48 | 0.381299 | 1.127028 | 0.297203 | 0.845703 |
| GOBP_NEGATIVE_REGULATION_OF_EPITHELIAL_CELL_APOPTOTIC_PROCESS | 26 | 0.438818 | 1.125028 | 0.292479 | 0.849813 |
| GOBP_REGULATION_OF_NEURON_MIGRATION | 25 | 0.431934 | 1.124547 | 0.306233 | 0.848388 |
| GOBP_LIVER_MORPHOGENESIS | 15 | 0.505582 | 1.124151 | 0.316667 | 0.846691 |
| GOBP_POSITIVE_REGULATION_OF_PATTERN_RECOGNITION_RECEPTOR_SIGNALING_I | 34 | 0.422041 | 1.123978 | 0.277955 | 0.844232 |
| GOBP_PYRIMIDINE_DEOXYRIBONUCLEOTIDE_METABOLIC_PROCESS | 21 | 0.44681 | 1.123851 | 0.307692 | 0.841654 |
| GOBP_RETINA_LAYER_FORMATION | 15 | 0.485057 | 1.123261 | 0.323155 | 0.840547 |
| GOBP_REGULATION_OF_OSSIFICATION | 70 | 0.361222 | 1.121292 | 0.242291 | 0.844478 |
| GOBP_RESPONSE_TO_CALCIUM_ION | 96 | 0.342655 | 1.120912 | 0.245763 | 0.842895 |
| GOBP_MYELOID_LEUKOCYTE_CYTOKINE_PRODUCTION | 24 | 0.44121 | 1.119483 | 0.302632 | 0.844916 |
| GOBP_DEFENSE_RESPONSE_TO_VIRUS | 208 | 0.315864 | 1.1179 | 0.169014 | 0.847327 |
| GOBP_TRNA_5_END_PROCESSING | 16 | 0.480086 | 1.117764 | 0.331695 | 0.8448 |
| GOBP_NEURON_RECOGNITION | 22 | 0.446252 | 1.115211 | 0.291545 | 0.850685 |
| GOBP_REGULATION_OF_MUSCLE_CELL_DIFFERENTIATION | 96 | 0.33405 | 1.114473 | 0.237885 | 0.850338 |
| GOBP_REGULATION_OF_VIRAL_GENOME_REPLICATION | 74 | 0.344038 | 1.113693 | 0.24498 | 0.850059 |
| GOBP_DETECTION_OF_CHEMICAL_STIMULUS_INVOLVED_IN_SENSORY_PERCEPTION_ | 17 | 0.471547 | 1.112265 | 0.326683 | 0.852125 |

| | | | | | |
|---|---|---|---|---|---|
| GOBP_REGULATION_OF_MEMBRANE_INVAGINATION | 15 | -0.803282 | -1.674762 | 0.012422 | 0.119099 |
| GOBP_REGULATION_OF_LEUKOCYTE_MEDIATED_IMMUNITY | 173 | -0.586525 | -1.674333 | 0 | 0.118766 |
| GOBP_MYELOID_CELL_APOPTOTIC_PROCESS | 21 | -0.781811 | -1.672799 | 0.011094 | 0.119798 |
| GOBP_POSITIVE_REGULATION_OF_ALPHA_BETA_T_CELL_ACTIVATION | 55 | -0.664169 | -1.671662 | 0.004127 | 0.120156 |
| GOBP_REGULATION_OF_CALCIUM_MEDIATED_SIGNALING | 51 | -0.679699 | -1.670759 | 0.004144 | 0.120601 |
| GOBP_INFLAMMATORY_CELL_APOPTOTIC_PROCESS | 16 | -0.821551 | -1.670555 | 0.013311 | 0.120009 |
| GOBP_CALCIUM_MEDIATED_SIGNALING | 144 | -0.606138 | -1.667841 | 0.00123 | 0.122369 |
| GOBP_REGULATION_OF_ACTIN_CYTOSKELETON_REORGANIZATION | 30 | -0.717923 | -1.667674 | 0.015291 | 0.12178 |
| GOBP_POSITIVE_REGULATION_OF_CATION_CHANNEL_ACTIVITY | 43 | -0.690693 | -1.666714 | 0.005806 | 0.122218 |
| GOBP_POSITIVE_REGULATION_OF_LIPID_STORAGE | 20 | -0.783146 | -1.663789 | 0.016556 | 0.125209 |
| GOBP_POSITIVE_REGULATION_OF_SMALL_GTPASE_MEDIATED_SIGNAL_TRANSI | 55 | -0.669546 | -1.663767 | 0.006729 | 0.12443 |
| GOBP_REGULATION_OF_CELL_SHAPE | 115 | -0.605028 | -1.662743 | 0.002442 | 0.124995 |
| GOBP_POSITIVE_REGULATION_OF_IMMUNE_EFFECTOR_PROCESS | 175 | -0.581708 | -1.662264 | 0 | 0.124776 |
| GOBP_T_CELL_DIFFERENTIATION_INVOLVED_IN_IMMUNE_RESPONSE | 57 | -0.656233 | -1.660331 | 0.01084 | 0.126302 |
| GOBP_REGULATION_OF_CHEMOTAXIS | 151 | -0.591826 | -1.659739 | 0 | 0.126461 |
| GOBP_POSITIVE_REGULATION_OF_FATTY_ACID_METABOLIC_PROCESS | 23 | -0.775376 | -1.657108 | 0.009449 | 0.128929 |
| GOBP_CELLULAR_RESPONSE_TO_CADMIUM_ION | 25 | -0.734735 | -1.656591 | 0.007813 | 0.128775 |
| GOBP_STEROL_TRANSPORT | 79 | -0.63658 | -1.656387 | 0.003989 | 0.128261 |
| GOBP_SOMATIC_DIVERSIFICATION_OF_IMMUNE_RECEPTORS | 72 | -0.635396 | -1.655594 | 0.006793 | 0.128623 |
| GOBP_NEGATIVE_REGULATION_OF_INFLAMMATORY_RESPONSE | 86 | -0.622358 | -1.654014 | 0.007989 | 0.129979 |
| GOBP_NEUROINFLAMMATORY_RESPONSE | 23 | -0.750358 | -1.653327 | 0.013889 | 0.130162 |
| GOBP_INTEGRATED_STRESS_RESPONSE_SIGNALING | 23 | -0.755911 | -1.651205 | 0.012289 | 0.132341 |
| GOBP_EPITHELIAL_CELL_MORPHOGENESIS | 19 | -0.784704 | -1.650706 | 0.00639 | 0.132166 |
| GOBP_NEGATIVE_REGULATION_OF_T_CELL_RECEPTOR_SIGNALING_PATHWAY | 21 | -0.767967 | -1.649514 | 0.017296 | 0.133071 |
| GOBP_LEUKOCYTE_MEDIATED_CYTOTOXICITY | 85 | -0.613416 | -1.649441 | 0.002567 | 0.132334 |
| GOBP_POSITIVE_REGULATION_OF_GLIAL_CELL_PROLIFERATION | 15 | -0.796538 | -1.64944 | 0.016611 | 0.131551 |
| GOBP_POSITIVE_REGULATION_OF_NATURAL_KILLER_CELL_MEDIATED_IMMUNI | 23 | -0.751122 | -1.649186 | 0.018072 | 0.131041 |
| GOBP_ASTROCYTE_ACTIVATION | 17 | -0.803231 | -1.648377 | 0.013093 | 0.131349 |
| GOBP_GLYCEROPHOSPHOLIPID_CATABOLIC_PROCESS | 22 | -0.74768 | -1.647175 | 0.017685 | 0.132281 |
| GOBP_MONOCARBOXYLIC_ACID_TRANSPORT | 106 | -0.606427 | -1.645985 | 0.002525 | 0.133119 |
| GOBP_POSITIVE_REGULATION_OF_SIGNALING_RECEPTOR_ACTIVITY | 28 | -0.726852 | -1.645982 | 0.012403 | 0.132365 |
| GOBP_POSITIVE_REGULATION_OF_DNA_TEMPLATED_TRANSCRIPTION_INITIATIC | 31 | -0.71389 | -1.644375 | 0.016541 | 0.133902 |
| GOBP_TISSUE_REMODELING | 114 | -0.604275 | -1.644332 | 0.002558 | 0.133236 |
| GOBP_POSITIVE_REGULATION_OF_ERBB_SIGNALING_PATHWAY | 27 | -0.736851 | -1.644278 | 0.012739 | 0.13256 |
| GOBP_SECOND_MESSENGER_MEDIATED_SIGNALING | 197 | -0.570456 | -1.643732 | 0.001153 | 0.132495 |
| GOBP_REGULATION_OF_RHO_PROTEIN_SIGNAL_TRANSDUCTION | 64 | -0.640666 | -1.643701 | 0.00534 | 0.131778 |
| GOBP_REGULATION_OF_FIBROBLAST_MIGRATION | 30 | -0.716582 | -1.64326 | 0.007519 | 0.131678 |
| GOBP_FOAM_CELL_DIFFERENTIATION | 25 | -0.751703 | -1.642891 | 0.012403 | 0.131458 |
| GOBP_ACTIN_POLYMERIZATION_OR_DEPOLYMERIZATION | 174 | -0.572688 | -1.642438 | 0 | 0.131383 |
| GOBP_MODIFICATION_OF_SYNAPTIC_STRUCTURE | 16 | -0.784205 | -1.642022 | 0.020505 | 0.131247 |
| GOBP_POSITIVE_REGULATION_OF_OSTEOCLAST_DIFFERENTIATION | 19 | -0.77266 | -1.641673 | 0.01248 | 0.130954 |
| GOBP_TOLL_LIKE_RECEPTOR_SIGNALING_PATHWAY | 130 | -0.595474 | -1.640629 | 0.001188 | 0.131493 |
| GOBP_NEGATIVE_REGULATION_OF_DNA_METABOLIC_PROCESS | 99 | -0.603657 | -1.639972 | 0.010485 | 0.131578 |
| GOBP_REGULATION_OF_CELL_SUBSTRATE_ADHESION | 147 | -0.577422 | -1.639794 | 0.003593 | 0.131081 |
| GOBP_DENDRITIC_CELL_CHEMOTAXIS | 19 | -0.762887 | -1.639472 | 0.023411 | 0.130851 |
| GOBP_REGULATION_OF_REACTIVE_OXYGEN_SPECIES_BIOSYNTHETIC_PROCES | 74 | -0.62681 | -1.638213 | 0.014765 | 0.131976 |
| GOBP_LABYRINTHINE_LAYER_DEVELOPMENT | 29 | -0.711495 | -1.637666 | 0.016369 | 0.131936 |
| GOBP_ATP_TRANSPORT | 15 | -0.793239 | -1.636544 | 0.023689 | 0.132613 |
| GOBP_REGULATION_OF_SUPEROXIDE_ANION_GENERATION | 15 | -0.796442 | -1.636226 | 0.025806 | 0.132309 |
| GOBP_ACUTE_INFLAMMATORY_RESPONSE | 65 | -0.632016 | -1.636031 | 0.006916 | 0.131862 |
| GOBP_NEGATIVE_REGULATION_OF_G_PROTEIN_COUPLED_RECEPTOR_SIGNAL | 34 | -0.703999 | -1.635691 | 0.022825 | 0.131606 |
| GOBP_PLASMA_LIPOPROTEIN_PARTICLE_CLEARANCE | 39 | -0.688985 | -1.635254 | 0.010448 | 0.131467 |
| GOBP_POSITIVE_REGULATION_OF_CYTOKINE_PRODUCTION | 340 | -0.544433 | -1.634464 | 0 | 0.131809 |
| GOBP_POSITIVE_REGULATION_OF_RHO_PROTEIN_SIGNAL_TRANSDUCTION | 22 | -0.748188 | -1.634143 | 0.019637 | 0.131644 |
| GOBP_POSITIVE_REGULATION_OF_LOCOMOTION | 396 | -0.541634 | -1.632613 | 0 | 0.132868 |
| GOBP_REGULATION_OF_CELL_SUBSTRATE_JUNCTION_ORGANIZATION | 55 | -0.657285 | -1.630202 | 0.014225 | 0.135367 |
| GOBP_MYD88_DEPENDENT_TOLL_LIKE_RECEPTOR_SIGNALING_PATHWAY | 33 | -0.712372 | -1.630004 | 0.010432 | 0.134946 |
| GOBP_PATTERN_RECOGNITION_RECEPTOR_SIGNALING_PATHWAY | 177 | -0.563106 | -1.629814 | 0 | 0.134535 |
| GOBP_BIOLOGICAL_PROCESS_INVOLVED_IN_INTERACTION_WITH_HOST | 184 | -0.568496 | -1.627954 | 0.002326 | 0.136092 |
| GOBP_POSITIVE_REGULATION_OF_RECEPTOR_SIGNALING_PATHWAY_VIA_STAT | 29 | -0.715649 | -1.627638 | 0.017161 | 0.135859 |
| GOBP_INTERLEUKIN_12_PRODUCTION | 50 | -0.66837 | -1.625992 | 0.009915 | 0.137395 |
| GOBP_REGULATION_OF_LYMPHOCYTE_ACTIVATION | 388 | -0.536305 | -1.625954 | 0 | 0.136765 |
| GOBP_POST_EMBRYONIC_DEVELOPMENT | 58 | -0.639228 | -1.625895 | 0.011019 | 0.136218 |
| GOBP_REGULATION_OF_DNA_TEMPLATED_TRANSCRIPTION_INITIATION | 36 | -0.683992 | -1.624911 | 0.016591 | 0.136858 |
| GOBP_NEGATIVE_REGULATION_OF_OXIDATIVE_STRESS_INDUCED_CELL_DEAT | 36 | -0.683209 | -1.623366 | 0.010249 | 0.138222 |
| GOBP_ODONTOGENESIS_OF_DENTIN_CONTAINING_TOOTH | 43 | -0.666599 | -1.623252 | 0.012987 | 0.13774 |
| GOBP_ENDODERMAL_CELL_DIFFERENTIATION | 31 | -0.706458 | -1.622349 | 0.027273 | 0.138328 |
| GOBP_FIBROBLAST_MIGRATION | 38 | -0.676805 | -1.620575 | 0.023426 | 0.139766 |
| GOBP_POSITIVE_REGULATION_OF_CHEMOTAXIS | 97 | -0.595326 | -1.618947 | 0.005025 | 0.141412 |
| GOBP_NEGATIVE_REGULATION_OF_LEUKOCYTE_CELL_CELL_ADHESION | 104 | -0.607703 | -1.618405 | 0.006418 | 0.141539 |
| GOBP_POSITIVE_REGULATION_OF_NEUTROPHIL_MIGRATION | 26 | -0.723981 | -1.618191 | 0.018462 | 0.141137 |
| GOBP_NEGATIVE_REGULATION_OF_LYMPHOCYTE_MEDIATED_IMMUNITY | 37 | -0.688 | -1.617635 | 0.012251 | 0.14123 |
| GOBP_NEGATIVE_REGULATION_OF_OXIDOREDUCTASE_ACTIVITY | 15 | -0.784937 | -1.617559 | 0.019078 | 0.140716 |
| GOBP_RESPONSE_TO_HYDROGEN_PEROXIDE | 108 | -0.597409 | -1.616015 | 0.003822 | 0.142056 |
| GOBP_REGULATION_OF_ACTIN_FILAMENT_LENGTH | 150 | -0.571228 | -1.615474 | 0.003654 | 0.142108 |
| GOBP_POSITIVE_REGULATION_OF_PROTEIN_POLYMERIZATION | 111 | -0.5882 | -1.615085 | 0.006281 | 0.141955 |
| GOBP_RESPIRATORY_BURST | 32 | -0.688692 | -1.614654 | 0.019146 | 0.141866 |
| GOBP_REGULATION_OF_MYELOID_LEUKOCYTE_DIFFERENTIATION | 88 | -0.600321 | -1.613018 | 0.004021 | 0.143275 |
| GOBP_POSITIVE_REGULATION_OF_ACTIN_FILAMENT_POLYMERIZATION | 80 | -0.620831 | -1.612642 | 0.009211 | 0.143119 |
| GOBP_ENERGY_HOMEOSTASIS | 23 | -0.738903 | -1.611046 | 0.020701 | 0.144645 |
| GOBP_RESPONSE_TO_AMYLOID_BETA | 31 | -0.693871 | -1.611023 | 0.014793 | 0.144033 |
| GOBP_PLACENTA_DEVELOPMENT | 85 | -0.604345 | -1.610431 | 0.011436 | 0.144172 |
| GOBP_REGULATION_OF_MONONUCLEAR_CELL_MIGRATION | 83 | -0.604719 | -1.610364 | 0.008986 | 0.143604 |
| GOBP_MATURE_B_CELL_DIFFERENTIATION_INVOLVED_IN_IMMUNE_RESPONSE | 19 | -0.767121 | -1.610261 | 0.020537 | 0.143156 |
| GOBP_REGULATION_OF_OXIDOREDUCTASE_ACTIVITY | 76 | -0.61761 | -1.609542 | 0.009383 | 0.143426 |
| GOBP_POSITIVE_REGULATION_OF_CALCIUM_MEDIATED_SIGNALING | 21 | -0.752114 | -1.60872 | 0.021242 | 0.143889 |
| GOBP_POSITIVE_REGULATION_OF_CELL_ACTIVATION | 305 | -0.541128 | -1.608555 | 0 | 0.143468 |
| GOBP_REGULATION_OF_LIPID_STORAGE | 43 | -0.668374 | -1.608022 | 0.028169 | 0.143574 |
| GOBP_SUPEROXIDE_ANION_GENERATION | 25 | -0.709388 | -1.607777 | 0.02795 | 0.143266 |
| GOBP_POSITIVE_REGULATION_OF_PHOSPHOPROTEIN_PHOSPHATASE_ACTIVITY | 19 | -0.745749 | -1.605825 | 0.032051 | 0.145245 |
| GOBP_BLOOD_VESSEL_REMODELING | 28 | -0.709288 | -1.605342 | 0.016975 | 0.145276 |
| GOBP_NEGATIVE_REGULATION_OF_LEUKOCYTE_MEDIATED_IMMUNITY | 43 | -0.665589 | -1.60384 | 0.02446 | 0.146641 |
| GOBP_KERATINOCYTE_PROLIFERATION | 27 | -0.71556 | -1.603632 | 0.023184 | 0.146267 |
| GOBP_NEGATIVE_REGULATION_OF_OXIDATIVE_STRESS_INDUCED_NEURON_DI | 16 | -0.780295 | -1.603489 | 0.03 | 0.145853 |
| GOBP_REGULATION_OF_PROTEIN_LOCALIZATION_TO_PLASMA_MEMBRANE | 76 | -0.620967 | -1.603286 | 0.005188 | 0.145573 |
| GOBP_NEGATIVE_REGULATION_OF_CELL_ADHESION | 208 | -0.548967 | -1.602475 | 0.001139 | 0.145983 |
| GOBP_REGULATION_OF_ANTIGEN_PROCESSING_AND_PRESENTATION | 18 | -0.756816 | -1.600729 | 0.030945 | 0.147718 |
| GOBP_REGULATION_OF_RESPONSE_TO_WOUNDING | 106 | -0.5887 | -1.600611 | 0.002532 | 0.147236 |
| GOBP_POSITIVE_REGULATION_OF_CYSTEINE_TYPE_ENDOPEPTIDASE_ACTIVITY | 120 | -0.580906 | -1.600039 | 0.004944 | 0.147448 |
| GOBP_CELL_MATRIX_ADHESION | 162 | -0.566489 | -1.599961 | 0.001188 | 0.146961 |
| GOBP_NEGATIVE_REGULATION_OF_BLOOD_VESSEL_ENDOTHELIAL_CELL_MIGF | 27 | -0.718949 | -1.599288 | 0.015748 | 0.147244 |
| GOBP_NEGATIVE_REGULATION_OF_ADAPTIVE_IMMUNE_RESPONSE | 41 | -0.663953 | -1.598126 | 0.022727 | 0.148255 |
| GOBP_REGULATION_OF_RECEPTOR_BINDING | 18 | -0.769861 | -1.597123 | 0.02454 | 0.148921 |
| GOBP_VENTRICULAR_SYSTEM_DEVELOPMENT | 18 | -0.757398 | -1.596277 | 0.038647 | 0.149476 |
| GOBP_PHAGOCYTOSIS | 261 | -0.542057 | -1.596248 | 0 | 0.148911 |
| GOBP_SEMAPHORIN_PLEXIN_SIGNALING_PATHWAY | 31 | -0.692336 | -1.595869 | 0.020833 | 0.148724 |
| GOBP_NEURON_PROJECTION_GUIDANCE | 170 | -0.557946 | -1.595498 | 0.001176 | 0.148597 |
| GOBP_ICOSANOID_TRANSPORT | 27 | -0.710651 | -1.594847 | 0.023328 | 0.148862 |
| GOBP_POSITIVE_REGULATION_OF_PROTEIN_LOCALIZATION_TO_CELL_PERIPHE | 50 | -0.632254 | -1.594139 | 0.016997 | 0.149286 |
| GOBP_T_HELPER_1_TYPE_IMMUNE_RESPONSE | 31 | -0.686061 | -1.593962 | 0.025605 | 0.148916 |
| GOBP_EPHRIN_RECEPTOR_SIGNALING_PATHWAY | 68 | -0.617175 | -1.593655 | 0.011065 | 0.148753 |
| GOBP_REGULATION_OF_B_CELL_DIFFERENTIATION | 26 | -0.705617 | -1.593183 | 0.027149 | 0.148808 |
| GOBP_ORGANIC_HYDROXY_COMPOUND_TRANSPORT | 154 | -0.566163 | -1.591976 | 0.00609 | 0.149937 |
| GOBP_REGULATION_OF_NEUTROPHIL_MIGRATION | 36 | -0.670186 | -1.591894 | 0.022599 | 0.149479 |
| GOBP_VASCULATURE_DEVELOPMENT | 471 | -0.520753 | -1.591578 | 0 | 0.149441 |
| GOBP_NEURON_CELLULAR_HOMEOSTASIS | 23 | -0.717782 | -1.590096 | 0.020833 | 0.15086 |
| GOBP_LEUKOCYTE_MIGRATION | 328 | -0.53503 | -1.589392 | 0.001088 | 0.151241 |
| GOBP_NEGATIVE_REGULATION_OF_LEUKOCYTE_MEDIATED_CYTOTOXICITY | 18 | -0.752699 | -1.587435 | 0.019672 | 0.153534 |
| GOBP_HORMONE_METABOLIC_PROCESS | 114 | -0.579797 | -1.583975 | 0.008861 | 0.15775 |
| GOBP_REGULATION_OF_CELL_CELL_ADHESION | 333 | -0.535338 | -1.583955 | 0 | 0.157182 |
| GOBP_NEGATIVE_REGULATION_OF_CELL_KILLING | 19 | -0.749673 | -1.583907 | 0.030111 | 0.156658 |
| GOBP_CELL_FATE_COMMITMENT | 125 | -0.575245 | -1.583623 | 0.006258 | 0.156461 |
| GOBP_REGULATION_OF_LEUKOCYTE_CHEMOTAXIS | 84 | -0.600623 | -1.58316 | 0.006562 | 0.156541 |
| GOBP_REGULATION_OF_RESPONSE_TO_BIOTIC_STIMULUS | 340 | -0.529137 | -1.582865 | 0 | 0.156381 |
| GOBP_OSTEOCLAST_DIFFERENTIATION | 72 | -0.605247 | -1.581714 | 0.019074 | 0.157451 |
| GOBP_NEURON_FATE_COMMITMENT | 17 | -0.744087 | -1.581517 | 0.034884 | 0.15718 |
| GOBP_SMOOTH_MUSCLE_CELL_PROLIFERATION | 95 | -0.588504 | -1.579918 | 0.005284 | 0.159039 |
| GOBP_REGULATION_OF_ACTIN_FILAMENT_BUNDLE_ASSEMBLY | 78 | -0.606576 | -1.579355 | 0.012097 | 0.159246 |
| GOBP_ASTROCYTE_DIFFERENTIATION | 50 | -0.642114 | -1.579071 | 0.025175 | 0.159052 |
| GOBP_POSITIVE_REGULATION_OF_TRANSPORTER_ACTIVITY | 78 | -0.603913 | -1.576617 | 0.012162 | 0.161769 |
| GOBP_NEGATIVE_REGULATION_OF_INTERLEUKIN_2_PRODUCTION | 24 | -0.698965 | -1.574464 | 0.031866 | 0.164245 |
| GOBP_CRANIAL_NERVE_DEVELOPMENT | 22 | -0.726279 | -1.574396 | 0.024653 | 0.16377 |
| GOBP_NEGATIVE_REGULATION_OF_CATION_CHANNEL_ACTIVITY | 27 | -0.685473 | -1.573303 | 0.032012 | 0.164777 |
| GOBP_NEGATIVE_REGULATION_OF_LOCOMOTION | 222 | -0.54147 | -1.573266 | 0.004515 | 0.164246 |
| GOBP_POSITIVE_REGULATION_OF_NATURAL_KILLER_CELL_MEDIATED_CYTOTC | 20 | -0.727127 | -1.572915 | 0.038647 | 0.16417 |
| GOBP_MYELOID_CELL_DIFFERENTIATION | 337 | -0.522161 | -1.572747 | 0 | 0.163866 |
| GOBP_NEGATIVE_REGULATION_OF_CYTOKINE_PRODUCTION | 222 | -0.539142 | -1.572436 | 0 | 0.16374 |
| GOBP_CELLULAR_RESPONSE_TO_FLUID_SHEAR_STRESS | 15 | -0.772272 | -1.571722 | 0.038585 | 0.164139 |
| GOBP_MYELOID_LEUKOCYTE_DIFFERENTIATION | 155 | -0.56013 | -1.571566 | 0.002448 | 0.163749 |
| GOBP_ACTIN_FILAMENT_BUNDLE_ORGANIZATION | 121 | -0.567435 | -1.571198 | 0.004854 | 0.163728 |
| GOBP_POSITIVE_REGULATION_OF_PEPTIDYL_TYROSINE_PHOSPHORYLATION | 121 | -0.57382 | -1.569662 | 0.004884 | 0.165413 |
| GOBP_ANDROGEN_METABOLIC_PROCESS | 18 | -0.741618 | -1.568897 | 0.026646 | 0.165999 |
| GOBP_NEGATIVE_REGULATION_OF_NEURON_DEATH | 135 | -0.56656 | -1.568512 | 0.006061 | 0.165974 |
| GOBP_ORGANIC_ACID_TRANSPORT | 206 | -0.538058 | -1.568 | 0 | 0.166101 |
| GOBP_POSITIVE_REGULATION_OF_PEPTIDASE_ACTIVITY | 151 | -0.553349 | -1.564687 | 0.003632 | 0.170412 |
| GOBP_REGULATION_OF_ACTIN_FILAMENT_BASED_PROCESS | 292 | -0.527131 | -1.564051 | 0 | 0.170834 |
| GOBP_REGULATION_OF_PLATELET_DERIVED_GROWTH_FACTOR_RECEPTOR_S | 18 | -0.73937 | -1.562612 | 0.024917 | 0.172478 |

| | | | | |
|---|---|---|---|---|
| GOBP_GOLGI_TO_PLASMA_MEMBRANE_TRANSPORT | 55 | 0.372795 | 1.110461 | 0.240283 | 0.855637 |
| GOBP_LONG_CHAIN_FATTY_ACYL_COA_BIOSYNTHETIC_PROCESS | 15 | 0.481611 | 1.107627 | 0.353659 | 0.862733 |
| GOBP_NEGATIVE_REGULATION_OF_FAT_CELL_DIFFERENTIATION | 35 | 0.396451 | 1.106941 | 0.290221 | 0.862259 |
| GOBP_CELL_CYCLE_CHECKPOINT | 197 | 0.307122 | 1.106618 | 0.176471 | 0.860362 |
| GOBP_HEART_TRABECULA_MORPHOGENESIS | 18 | 0.466713 | 1.106034 | 0.34715 | 0.859625 |
| GOBP_PYRIMIDINE_NUCLEOTIDE_BIOSYNTHETIC_PROCESS | 25 | 0.433325 | 1.104547 | 0.305476 | 0.86189 |
| GOBP_BRANCHING_MORPHOGENESIS_OF_AN_EPITHELIAL_TUBE | 76 | 0.352916 | 1.103189 | 0.264463 | 0.86378 |
| GOBP_CELL_DIFFERENTIATION_IN_SPINAL_CORD | 21 | 0.448552 | 1.101345 | 0.345345 | 0.867287 |
| GOBP_DIENCEPHALON_DEVELOPMENT | 27 | 0.43496 | 1.101173 | 0.323699 | 0.865008 |
| GOBP_POSITIVE_REGULATION_OF_MYELOID_LEUKOCYTE_CYTOKINE_PRODUCTION_II | 15 | 0.488845 | 1.097315 | 0.339572 | 0.875679 |
| GOBP_RUFFLE_ASSEMBLY | 35 | 0.401426 | 1.096645 | 0.283388 | 0.875139 |
| GOBP_EMBRYONIC_SKELETAL_SYSTEM_MORPHOGENESIS | 45 | 0.381882 | 1.09561 | 0.313589 | 0.875916 |
| GOBP_THIOESTER_BIOSYNTHETIC_PROCESS | 48 | 0.377533 | 1.095433 | 0.288889 | 0.873663 |
| GOBP_NUCLEOTIDE_SUGAR_BIOSYNTHETIC_PROCESS | 22 | 0.433491 | 1.094789 | 0.319892 | 0.87308 |
| GOBP_REGULATION_OF_PLATELET_ACTIVATION | 25 | 0.429806 | 1.092749 | 0.333333 | 0.877576 |
| GOBP_POSITIVE_REGULATION_OF_PEPTIDYL_THREONINE_PHOSPHORYLATION | 25 | 0.4333 | 1.092692 | 0.349432 | 0.874928 |
| GOBP_POSITIVE_REGULATION_OF_BMP_SIGNALING_PATHWAY | 18 | 0.448054 | 1.091387 | 0.349594 | 0.876861 |
| GOBP_MAMMARY_GLAND_EPITHELIAL_CELL_PROLIFERATION | 23 | 0.432482 | 1.091292 | 0.325843 | 0.874318 |
| GOBP_MUSCLE_ORGAN_DEVELOPMENT | 204 | 0.29957 | 1.091146 | 0.159292 | 0.871987 |
| GOBP_POSITIVE_REGULATION_OF_JUN_KINASE_ACTIVITY | 56 | 0.362385 | 1.090561 | 0.301418 | 0.871205 |
| GOBP_REGULATION_OF_PROTEIN_LOCALIZATION_TO_SYNAPSE | 22 | 0.438929 | 1.090244 | 0.349432 | 0.869489 |
| GOBP_FATTY_ACID_DERIVATIVE_METABOLIC_PROCESS | 55 | 0.374076 | 1.089614 | 0.327138 | 0.868794 |
| GOBP_OUTFLOW_TRACT_MORPHOGENESIS | 37 | 0.391421 | 1.089574 | 0.32107 | 0.8662 |
| GOBP_CARDIOCYTE_DIFFERENTIATION | 73 | 0.354451 | 1.088313 | 0.278431 | 0.867909 |
| GOBP_RIBOSOMAL_LARGE_SUBUNIT_ASSEMBLY | 27 | 0.416123 | 1.088019 | 0.331325 | 0.866074 |
| GOBP_PROTEIN_DNA_COMPLEX_DISASSEMBLY | 18 | 0.454303 | 1.084824 | 0.358247 | 0.874652 |
| GOBP_OLIGODENDROCYTE_DEVELOPMENT | 28 | 0.421465 | 1.084617 | 0.338753 | 0.872623 |
| GOBP_REGULATION_OF_LYMPHOCYTE_CHEMOTAXIS | 15 | 0.464837 | 1.084539 | 0.363171 | 0.870165 |
| GOBP_REGULATION_OF_MYOBLAST_DIFFERENTIATION | 27 | 0.426032 | 1.083385 | 0.351429 | 0.871536 |
| GOBP_SULFUR_AMINO_ACID_METABOLIC_PROCESS | 28 | 0.409705 | 1.083266 | 0.352601 | 0.869228 |
| GOBP_CYTOPLASMIC_TRANSLATION | 98 | 0.331492 | 1.082332 | 0.313305 | 0.869696 |
| GOBP_ACTIVATION_OF_GTPASE_ACTIVITY | 93 | 0.333028 | 1.082209 | 0.293103 | 0.867417 |
| GOBP_CORONARY_VASCULATURE_DEVELOPMENT | 31 | 0.40457 | 1.079094 | 0.325444 | 0.875506 |
| GOBP_POSITIVE_REGULATION_OF_TRANSMEMBRANE_RECEPTOR_PROTEIN_SERINE_ | 72 | 0.34949 | 1.078047 | 0.294118 | 0.876473 |
| GOBP_POSITIVE_REGULATION_OF_EPITHELIAL_CELL_DIFFERENTIATION | 32 | 0.400197 | 1.075638 | 0.345345 | 0.882101 |
| GOBP_SENSORY_PERCEPTION_OF_LIGHT_STIMULUS | 115 | 0.32225 | 1.074173 | 0.240437 | 0.884786 |
| GOBP_POSITIVE_REGULATION_OF_FIBROBLAST_PROLIFERATION | 32 | 0.398051 | 1.07311 | 0.347701 | 0.885888 |
| GOBP_REGULATION_OF_PEPTIDYL_THREONINE_PHOSPHORYLATION | 40 | 0.389102 | 1.071662 | 0.328076 | 0.888225 |
| GOBP_REGULATION_OF_LYMPHOCYTE_MIGRATION | 47 | 0.363204 | 1.071524 | 0.344156 | 0.886058 |
| GOBP_REGULATION_OF_TRANSCRIPTION_REGULATORY_REGION_DNA_BINDING | 32 | 0.401781 | 1.071364 | 0.346749 | 0.883882 |
| GOBP_REGULATION_OF_T_CELL_APOPTOTIC_PROCESS | 28 | 0.404705 | 1.070991 | 0.353276 | 0.882485 |
| GOBP_REGULATION_OF_GOLGI_ORGANIZATION | 16 | 0.456023 | 1.070903 | 0.377953 | 0.880116 |
| GOBP_RESPONSE_TO_MURAMYL_DIPEPTIDE | 20 | 0.444098 | 1.070111 | 0.359673 | 0.880055 |
| GOBP_MEIOTIC_CELL_CYCLE | 189 | 0.302093 | 1.068811 | 0.277027 | 0.881962 |
| GOBP_POSITIVE_REGULATION_OF_PATHWAY_RESTRICTED_SMAD_PROTEIN_PHOSPH | 30 | 0.395157 | 1.068597 | 0.330303 | 0.880071 |
| GOBP_MALE_SEX_DIFFERENTIATION | 90 | 0.334703 | 1.068341 | 0.331967 | 0.878295 |
| GOBP_TRANSCYTOSIS | 16 | 0.45377 | 1.067068 | 0.369509 | 0.879997 |
| GOBP_REGULATION_OF_CYCLIN_DEPENDENT_PROTEIN_KINASE_ACTIVITY | 91 | 0.340695 | 1.066255 | 0.314159 | 0.880185 |
| GOBP_TELOMERASE_RNA_LOCALIZATION | 19 | 0.435036 | 1.064493 | 0.384831 | 0.88369 |
| GOBP_GLOMERULAR_EPITHELIUM_DEVELOPMENT | 17 | 0.451243 | 1.063814 | 0.387654 | 0.883568 |
| GOBP_MATURATION_OF_SSU_RRNA | 50 | 0.358734 | 1.063535 | 0.327138 | 0.881965 |
| GOBP_MATURATION_OF_LSU_RRNA | 28 | 0.406391 | 1.063455 | 0.370149 | 0.8797 |
| GOBP_REGULATION_OF_ARP2_3_COMPLEX_MEDIATED_ACTIN_NUCLEATION | 18 | 0.441663 | 1.063303 | 0.402089 | 0.877651 |
| GOBP_MEIOTIC_CELL_CYCLE_PROCESS | 146 | 0.311834 | 1.062819 | 0.287293 | 0.876778 |
| GOBP_AEROBIC_RESPIRATION | 81 | 0.334629 | 1.061018 | 0.291139 | 0.880418 |
| GOBP_CELLULAR_AMINO_ACID_BIOSYNTHETIC_PROCESS | 61 | 0.350875 | 1.058729 | 0.329412 | 0.88571 |
| GOBP_FIBROBLAST_GROWTH_FACTOR_RECEPTOR_SIGNALING_PATHWAY | 70 | 0.34532 | 1.057112 | 0.310219 | 0.888736 |
| GOBP_NEGATIVE_REGULATION_OF_SMALL_MOLECULE_METABOLIC_PROCESS | 66 | 0.346962 | 1.05684 | 0.326923 | 0.887075 |
| GOBP_POSITIVE_REGULATION_OF_GLUCOSE_IMPORT | 27 | 0.411109 | 1.056829 | 0.369697 | 0.884568 |
| GOBP_POSITIVE_REGULATION_OF_MITOCHONDRIAL_TRANSLATION | 16 | 0.449849 | 1.054166 | 0.376316 | 0.8912 |
| GOBP_MITOCHONDRIAL_RESPIRATORY_CHAIN_COMPLEX_ASSEMBLY | 101 | 0.32224 | 1.054161 | 0.3 | 0.888688 |
| GOBP_IRON_ION_HOMEOSTASIS | 72 | 0.34408 | 1.052144 | 0.364341 | 0.893195 |
| GOBP_MISMATCH_REPAIR | 35 | 0.38724 | 1.051459 | 0.362776 | 0.893118 |
| GOBP_T_CELL_APOPTOTIC_PROCESS | 40 | 0.375501 | 1.051367 | 0.356913 | 0.890935 |
| GOBP_PSEUDOURIDINE_SYNTHESIS | 18 | 0.440293 | 1.050635 | 0.40404 | 0.890925 |
| GOBP_CLEAVAGE_INVOLVED_IN_RRNA_PROCESSING | 27 | 0.40409 | 1.050526 | 0.364672 | 0.888794 |
| GOBP_NEGATIVE_CHEMOTAXIS | 29 | 0.397464 | 1.04907 | 0.41543 | 0.891304 |
| GOBP_VITAMIN_METABOLIC_PROCESS | 92 | 0.329448 | 1.048985 | 0.36087 | 0.889064 |
| GOBP_BONE_MORPHOGENESIS | 48 | 0.360648 | 1.04836 | 0.35461 | 0.888762 |
| GOBP_BILE_ACID_METABOLIC_PROCESS | 29 | 0.399473 | 1.048263 | 0.385965 | 0.886611 |
| GOBP_APOPTOTIC_PROCESS_INVOLVED_IN_MORPHOGENESIS | 16 | 0.453725 | 1.047817 | 0.424157 | 0.885737 |
| GOBP_REGULATION_OF_SYNAPTIC_TRANSMISSION_GABAERGIC | 18 | 0.439207 | 1.047448 | 0.380208 | 0.884541 |
| GOBP_ONE_CARBON_METABOLIC_PROCESS | 31 | 0.390597 | 1.04713 | 0.381232 | 0.883189 |
| GOBP_ESTABLISHMENT_OF_PIGMENT_GRANULE_LOCALIZATION | 20 | 0.426805 | 1.04702 | 0.397368 | 0.881101 |
| GOBP_PHOSPHATIDYLETHANOLAMINE_METABOLIC_PROCESS | 24 | 0.410552 | 1.046956 | 0.394521 | 0.878936 |
| GOBP_LACTATION | 35 | 0.382519 | 1.046682 | 0.380665 | 0.877481 |
| GOBP_GLYCOSYL_COMPOUND_CATABOLIC_PROCESS | 36 | 0.385717 | 1.04659 | 0.376147 | 0.875382 |
| GOBP_POSITIVE_REGULATION_OF_EXCITATORY_POSTSYNAPTIC_POTENTIAL | 15 | 0.455677 | 1.044814 | 0.388325 | 0.879141 |
| GOBP_PEROXISOMAL_TRANSPORT | 66 | 0.343178 | 1.044755 | 0.380952 | 0.876987 |
| GOBP_CELLULAR_RESPONSE_TO_GAMMA_RADIATION | 23 | 0.412004 | 1.044514 | 0.41779 | 0.875398 |
| GOBP_POSITIVE_REGULATION_OF_HORMONE_SECRETION | 77 | 0.333868 | 1.043742 | 0.368421 | 0.875666 |
| GOBP_AROMATIC_AMINO_ACID_FAMILY_CATABOLIC_PROCESS | 15 | 0.454704 | 1.043627 | 0.411765 | 0.873664 |
| GOBP_TRNA_METABOLIC_PROCESS | 176 | 0.293237 | 1.041979 | 0.292994 | 0.876905 |
| GOBP_REGULATION_OF_ENDOCRINE_PROCESS | 16 | 0.448434 | 1.040088 | 0.411268 | 0.880988 |
| GOBP_NEGATIVE_REGULATION_OF_MICROTUBULE_POLYMERIZATION_OR_DEPOLYME | 29 | 0.382661 | 1.040077 | 0.373887 | 0.878665 |
| GOBP_REGULATION_OF_MICROTUBULE_DEPOLYMERIZATION | 23 | 0.411652 | 1.039876 | 0.397933 | 0.877001 |
| GOBP_ORGANIC_HYDROXY_COMPOUND_BIOSYNTHETIC_PROCESS | 178 | 0.299774 | 1.039756 | 0.348993 | 0.875014 |
| GOBP_RRNA_CONTAINING_RIBONUCLEOPROTEIN_COMPLEX_EXPORT_FROM_NUCLE | 15 | 0.465657 | 1.039264 | 0.416268 | 0.874325 |
| GOBP_ENDOPLASMIC_RETICULUM_MANNOSE_TRIMMING | 16 | 0.438544 | 1.03899 | 0.41623 | 0.872952 |
| GOBP_RETROGRADE_VESICLE_MEDIATED_TRANSPORT_GOLGI_TO_ENDOPLASMIC_R | 82 | 0.325693 | 1.038884 | 0.32549 | 0.871054 |
| GOBP_REGULATION_OF_FATTY_ACID_BETA_OXIDATION | 16 | 0.447303 | 1.038844 | 0.395161 | 0.868871 |
| GOBP_PIGMENT_GRANULE_LOCALIZATION | 22 | 0.42201 | 1.034394 | 0.408108 | 0.881598 |
| GOBP_NEGATIVE_REGULATION_OF_PROTEIN_ACETYLATION | 18 | 0.432598 | 1.032677 | 0.432643 | 0.884989 |
| GOBP_PIGMENT_BIOSYNTHETIC_PROCESS | 50 | 0.346028 | 1.032364 | 0.368243 | 0.883799 |
| GOBP_BILE_ACID_BIOSYNTHETIC_PROCESS | 22 | 0.405077 | 1.032166 | 0.422652 | 0.882144 |
| GOBP_POSITIVE_REGULATION_OF_CHEMOKINE_PRODUCTION | 41 | 0.370787 | 1.031604 | 0.395349 | 0.881803 |
| GOBP_CELLULAR_RESPONSE_TO_INORGANIC_SUBSTANCE | 151 | 0.29529 | 1.030307 | 0.343374 | 0.88383 |
| GOBP_OXIDATIVE_DEMETHYLATION | 16 | 0.436624 | 1.030069 | 0.416459 | 0.882408 |
| GOBP_FATTY_ACID_CATABOLIC_PROCESS | 95 | 0.314301 | 1.029872 | 0.391705 | 0.880776 |
| GOBP_VESICLE_MEDIATED_TRANSPORT_TO_THE_PLASMA_MEMBRANE | 83 | 0.323421 | 1.02899 | 0.429167 | 0.881512 |
| GOBP_AUTONOMIC_NERVOUS_SYSTEM_DEVELOPMENT | 22 | 0.407751 | 1.028946 | 0.408219 | 0.879378 |
| GOBP_MICROTUBULE_DEPOLYMERIZATION | 32 | 0.379839 | 1.028447 | 0.387692 | 0.878836 |
| GOBP_REGULATION_OF_ATPASE_ACTIVITY | 71 | 0.331334 | 1.027414 | 0.392593 | 0.879975 |
| GOBP_SKELETAL_SYSTEM_MORPHOGENESIS | 115 | 0.304279 | 1.026513 | 0.382353 | 0.880755 |
| GOBP_LONG_CHAIN_FATTY_ACYL_COA_METABOLIC_PROCESS | 21 | 0.414906 | 1.025696 | 0.41092 | 0.881335 |
| GOBP_SMAD_PROTEIN_SIGNAL_TRANSDUCTION | 47 | 0.362285 | 1.02253 | 0.398693 | 0.889489 |
| GOBP_MATURATION_OF_5_8S_RRNA_FROM_TRICISTRONIC_RRNA_TRANSCRIPT_SSU | 24 | 0.402587 | 1.022164 | 0.40056 | 0.88841 |
| GOBP_PATHWAY_RESTRICTED_SMAD_PROTEIN_PHOSPHORYLATION | 42 | 0.365889 | 1.02105 | 0.428135 | 0.889883 |
| GOBP_ACUTE_INFLAMMATORY_RESPONSE_TO_ANTIGENIC_STIMULUS | 17 | 0.431405 | 1.020212 | 0.416459 | 0.890377 |
| GOBP_DNA_CATABOLIC_PROCESS_ENDONUCLEOLYTIC | 26 | 0.399723 | 1.019369 | 0.425 | 0.890974 |
| GOBP_REGULATION_OF_ERAD_PATHWAY | 19 | 0.428148 | 1.019277 | 0.425316 | 0.889054 |
| GOBP_UBIQUITIN_DEPENDENT_ERAD_PATHWAY | 75 | 0.323591 | 1.01859 | 0.395161 | 0.889114 |
| GOBP_EPITHELIAL_STRUCTURE_MAINTENANCE | 15 | 0.433211 | 1.016997 | 0.42228 | 0.892152 |
| GOBP_REPLACEMENT_OSSIFICATION | 19 | 0.422876 | 1.015791 | 0.433702 | 0.893975 |
| GOBP_REGULATION_OF_BEHAVIOR | 32 | 0.379406 | 1.015365 | 0.431611 | 0.893307 |
| GOBP_RIBOSOME_BIOGENESIS | 302 | 0.268365 | 1.015195 | 0.414141 | 0.891652 |
| GOBP_GLUTAMINE_FAMILY_AMINO_ACID_BIOSYNTHETIC_PROCESS | 16 | 0.430762 | 1.014347 | 0.44389 | 0.892363 |
| GOBP_NEGATIVE_REGULATION_OF_PEPTIDYL_LYSINE_ACETYLATION | 16 | 0.44076 | 1.009362 | 0.46114 | 0.907242 |
| GOBP_TRABECULA_MORPHOGENESIS | 27 | 0.388662 | 1.008715 | 0.421538 | 0.907245 |
| GOBP_REGULATION_OF_STEROID_METABOLIC_PROCESS | 82 | 0.312188 | 1.006028 | 0.404082 | 0.914088 |
| GOBP_PYRIMIDINE_CONTAINING_COMPOUND_BIOSYNTHETIC_PROCESS | 33 | 0.37371 | 1.005601 | 0.442424 | 0.913369 |
| GOBP_PEPTIDE_CATABOLIC_PROCESS | 22 | 0.397021 | 1.004608 | 0.448276 | 0.914744 |
| GOBP_ENDONUCLEOLYTIC_CLEAVAGE_INVOLVED_IN_RRNA_PROCESSING | 15 | 0.447904 | 1.002467 | 0.464646 | 0.919764 |
| GOBP_NEGATIVE_REGULATION_OF_CHROMATIN_ORGANIZATION | 52 | 0.332721 | 1.000919 | 0.460317 | 0.92279 |
| GOBP_REGULATION_OF_FIBROBLAST_GROWTH_FACTOR_RECEPTOR_SIGNALING_PAT | 17 | 0.42684 | 0.999498 | 0.458445 | 0.925307 |
| GOBP_POSITIVE_REGULATION_OF_JNK_CASCADE | 69 | 0.32071 | 0.996073 | 0.453061 | 0.934711 |
| GOBP_REGULATION_OF_HEMATOPOIETIC_PROGENITOR_CELL_DIFFERENTIATION | 83 | 0.316788 | 0.995687 | 0.442149 | 0.933801 |
| GOBP_CALCIUM_DEPENDENT_CELL_CELL_ADHESION_VIA_PLASMA_MEMBRANE_CEL | 17 | 0.429171 | 0.995189 | 0.453826 | 0.933276 |
| GOBP_VESICLE_CARGO_LOADING | 20 | 0.398897 | 0.993765 | 0.4375 | 0.935955 |
| GOBP_SECONDARY_METABOLITE_BIOSYNTHETIC_PROCESS | 17 | 0.426683 | 0.992647 | 0.463476 | 0.93761 |
| GOBP_NUCLEOSIDE_BISPHOSPHATE_BIOSYNTHETIC_PROCESS | 62 | 0.321488 | 0.992288 | 0.492126 | 0.936638 |
| GOBP_RESPONSE_TO_RETINOIC_ACID | 62 | 0.320581 | 0.991101 | 0.450909 | 0.938516 |
| GOBP_REGULATION_OF_JNK_CASCADE | 105 | 0.306573 | 0.99012 | 0.445415 | 0.939735 |
| GOBP_PEPTIDYL_SERINE_MODIFICATION | 260 | 0.271727 | 0.989356 | 0.479592 | 0.940124 |
| GOBP_ORGANIC_ACID_CATABOLIC_PROCESS | 205 | 0.271554 | 0.986736 | 0.491936 | 0.946893 |
| GOBP_SYNAPTIC_TRANSMISSION_GABAERGIC | 22 | 0.391665 | 0.986146 | 0.483696 | 0.946554 |
| GOBP_CYTOKINETIC_PROCESS | 37 | 0.359552 | 0.986135 | 0.474006 | 0.944366 |
| GOBP_PURINE_CONTAINING_COMPOUND_CATABOLIC_PROCESS | 42 | 0.349728 | 0.985743 | 0.462069 | 0.943512 |
| GOBP_REGULATION_OF_CYCLASE_ACTIVITY | 25 | 0.398902 | 0.984618 | 0.468208 | 0.945265 |
| GOBP_NEURON_PROJECTION_ARBORIZATION | 21 | 0.390905 | 0.98349 | 0.477778 | 0.946789 |
| GOBP_CYTOCHROME_COMPLEX_ASSEMBLY | 36 | 0.355001 | 0.983489 | 0.447284 | 0.944599 |
| GOBP_VESICLE_FUSION_TO_PLASMA_MEMBRANE | 16 | 0.421553 | 0.982895 | 0.457746 | 0.944459 |
| GOBP_AORTA_DEVELOPMENT | 36 | 0.358275 | 0.982866 | 0.478125 | 0.942369 |
| GOBP_RNA_5_END_PROCESSING | 23 | 0.382912 | 0.980559 | 0.480447 | 0.947944 |
| GOBP_REGULATION_OF_PLATELET_AGGREGATION | 15 | 0.431822 | 0.980193 | 0.495935 | 0.947034 |
| GOBP_ORGAN_GROWTH | 96 | 0.298759 | 0.974112 | 0.547511 | 0.965672 |
| GOBP_STRIATED_MUSCLE_CELL_DIFFERENTIATION | 158 | 0.284746 | 0.974033 | 0.517647 | 0.963752 |

| | | | | | |
|---|---|---|---|---|---|
| GOBP_POSITIVE_REGULATION_OF_TRANSCRIPTION_REGULATORY_REGION_DN | 16 | -0.748066 | -1.562391 | 0.037767 | 0.172227 |
| GOBP_G_PROTEIN_COUPLED_RECEPTOR_SIGNALING_PATHWAY_COUPLED_TC | 16 | -0.752128 | -1.561202 | 0.031667 | 0.173402 |
| GOBP_REGULATION_OF_PEPTIDASE_ACTIVITY | 292 | -0.524752 | -1.559217 | 0.00108 | 0.175872 |
| GOBP_AMINO_SUGAR_METABOLIC_PROCESS | 33 | -0.675872 | -1.558601 | 0.031915 | 0.176151 |
| GOBP_FATTY_ACID_TRANSMEMBRANE_TRANSPORT | 39 | -0.663954 | -1.557538 | 0.024566 | 0.177248 |
| GOBP_REGULATION_OF_CELL_ACTIVATION | 475 | -0.515093 | -1.555246 | 0 | 0.180108 |
| GOBP_CELLULAR_MONOVALENT_INORGANIC_CATION_HOMEOSTASIS | 86 | -0.587784 | -1.554937 | 0.007682 | 0.179976 |
| GOBP_MEMBRANE_RAFT_ORGANIZATION | 22 | -0.711441 | -1.554202 | 0.033281 | 0.180522 |
| GOBP_REGULATION_OF_KERATINOCYTE_PROLIFERATION | 21 | -0.711111 | -1.554092 | 0.030817 | 0.180118 |
| GOBP_POSITIVE_REGULATION_OF_INTERLEUKIN_4_PRODUCTION | 22 | -0.704613 | -1.552371 | 0.027464 | 0.182197 |
| GOBP_ANTIMICROBIAL_HUMORAL_IMMUNE_RESPONSE_MEDIATED_BY_ANTIMIC | 24 | -0.700603 | -1.552081 | 0.031797 | 0.182081 |
| GOBP_POSITIVE_REGULATION_OF_PHOSPHATASE_ACTIVITY | 28 | -0.683846 | -1.551192 | 0.027778 | 0.182851 |
| GOBP_PROSTANOID_BIOSYNTHETIC_PROCESS | 17 | -0.747126 | -1.550706 | 0.036741 | 0.182983 |
| GOBP_ENDODERM_FORMATION | 38 | -0.649686 | -1.55013 | 0.020927 | 0.183298 |
| GOBP_RESPONSE_TO_WOUNDING | 441 | -0.510515 | -1.549966 | 0 | 0.182954 |
| GOBP_RESPONSE_TO_TUMOR_NECROSIS_FACTOR | 245 | -0.524096 | -1.549005 | 0 | 0.183922 |
| GOBP_NEGATIVE_REGULATION_OF_VASCULATURE_DEVELOPMENT | 69 | -0.606162 | -1.548536 | 0.01368 | 0.184079 |
| GOBP_POSITIVE_REGULATION_OF_MITOTIC_NUCLEAR_DIVISION | 31 | -0.669335 | -1.547539 | 0.022157 | 0.18515 |
| GOBP_POSITIVE_REGULATION_OF_ANTIGEN_RECEPTOR_MEDIATED_SIGNALING | 21 | -0.719083 | -1.547391 | 0.037441 | 0.1848 |
| GOBP_FOCAL_ADHESION_ASSEMBLY | 69 | -0.609148 | -1.547 | 0.012032 | 0.184793 |
| GOBP_POSITIVE_REGULATION_OF_LEUKOCYTE_CHEMOTAXIS | 66 | -0.598601 | -1.544577 | 0.024259 | 0.188034 |
| GOBP_LOW_DENSITY_LIPOPROTEIN_PARTICLE_CLEARANCE | 27 | -0.692145 | -1.544538 | 0.029895 | 0.187488 |
| GOBP_ACTIN_FILAMENT_ORGANIZATION | 336 | -0.514283 | -1.544472 | 0.001072 | 0.187009 |
| GOBP_TOLERANCE_INDUCTION | 23 | -0.699477 | -1.544073 | 0.042386 | 0.187033 |
| GOBP_REGULATION_OF_TRIGLYCERIDE_BIOSYNTHETIC_PROCESS | 16 | -0.748946 | -1.543296 | 0.029549 | 0.187583 |
| GOBP_POSITIVE_REGULATION_OF_MAPK_CASCADE | 347 | -0.51599 | -1.542177 | 0 | 0.188897 |
| GOBP_NEGATIVE_REGULATION_OF_DNA_BIOSYNTHETIC_PROCESS | 29 | -0.681165 | -1.539964 | 0.023952 | 0.191941 |
| GOBP_NEGATIVE_REGULATION_OF_PRODUCTION_OF_MOLECULAR_MEDIATOR_ | 32 | -0.677818 | -1.539949 | 0.027818 | 0.191371 |
| GOBP_NEGATIVE_REGULATION_OF_NEURON_APOPTOTIC_PROCESS | 87 | -0.580698 | -1.5397 | 0.012016 | 0.191177 |
| GOBP_NEGATIVE_REGULATION_OF_NF_KAPPAB_TRANSCRIPTION_FACTOR_ACT | 73 | -0.596794 | -1.539038 | 0.019048 | 0.191536 |
| GOBP_N_ACETYLGLUCOSAMINE_METABOLIC_PROCESS | 15 | -0.751135 | -1.538776 | 0.034976 | 0.191307 |
| GOBP_SMOOTH_MUSCLE_CELL_APOPTOTIC_PROCESS | 21 | -0.713786 | -1.538486 | 0.039002 | 0.191164 |
| GOBP_LYMPHOCYTE_ACTIVATION_INVOLVED_IN_IMMUNE_RESPONSE | 149 | -0.551838 | -1.538402 | 0.003546 | 0.190734 |
| GOBP_REGULATION_OF_CELL_ADHESION_MEDIATED_BY_INTEGRIN | 36 | -0.66472 | -1.537722 | 0.019006 | 0.191215 |
| GOBP_INNATE_IMMUNE_RESPONSE_ACTIVATING_SIGNAL_TRANSDUCTION | 96 | -0.576155 | -1.537676 | 0.015113 | 0.190707 |
| GOBP_RECEPTOR_SIGNALING_PATHWAY_VIA_STAT | 105 | -0.565452 | -1.537156 | 0.012594 | 0.19094 |
| GOBP_REGULATION_OF_SUPEROXIDE_METABOLIC_PROCESS | 22 | -0.713808 | -1.536775 | 0.036309 | 0.190953 |
| GOBP_RESPONSE_TO_BACTERIUM | 413 | -0.506768 | -1.536664 | 0 | 0.190556 |
| GOBP_LIPID_STORAGE | 67 | -0.593822 | -1.536384 | 0.025435 | 0.190501 |
| GOBP_ANTIMICROBIAL_HUMORAL_RESPONSE | 46 | -0.633393 | -1.535406 | 0.027536 | 0.191394 |
| GOBP_REGULATION_OF_IMMUNOGLOBULIN_PRODUCTION | 58 | -0.620034 | -1.535339 | 0.01931 | 0.190917 |
| GOBP_POSITIVE_REGULATION_OF_ISOTYPE_SWITCHING | 24 | -0.695341 | -1.53511 | 0.033228 | 0.190678 |
| GOBP_PEPTIDYL_TYROSINE_MODIFICATION | 258 | -0.519757 | -1.534478 | 0.001131 | 0.191069 |
| GOBP_RESPONSE_TO_INTERLEUKIN_6 | 28 | -0.668674 | -1.53367 | 0.03882 | 0.191669 |
| GOBP_REGULATION_OF_LIPID_LOCALIZATION | 111 | -0.564719 | -1.533616 | 0.017566 | 0.191203 |
| GOBP_MYELOID_LEUKOCYTE_MEDIATED_IMMUNITY | 465 | -0.503405 | -1.53129 | 0 | 0.194293 |
| GOBP_REGULATION_OF_MAPK_CASCADE | 480 | -0.502802 | -1.530721 | 0 | 0.194633 |
| GOBP_DETECTION_OF_BIOTIC_STIMULUS | 26 | -0.687054 | -1.529749 | 0.037936 | 0.195628 |
| GOBP_INTERLEUKIN_4_PRODUCTION | 29 | -0.667269 | -1.529104 | 0.039394 | 0.19607 |
| GOBP_ORGANIC_ANION_TRANSPORT | 235 | -0.522278 | -1.529001 | 0.001109 | 0.195626 |
| GOBP_POSITIVE_REGULATION_OF_SMOOTH_MUSCLE_CELL_PROLIFERATION | 62 | -0.592197 | -1.527718 | 0.024725 | 0.197052 |
| GOBP_CELL_SUBSTRATE_JUNCTION_ORGANIZATION | 84 | -0.58423 | -1.5264 | 0.022107 | 0.198669 |
| GOBP_REGULATION_OF_LIPID_METABOLIC_PROCESS | 294 | -0.513458 | -1.525532 | 0.00222 | 0.199508 |
| GOBP_PROTEIN_AUTOPROCESSING | 15 | -0.742997 | -1.524786 | 0.047541 | 0.200082 |
| GOBP_REGULATION_OF_MYELOID_CELL_APOPTOTIC_PROCESS | 17 | -0.717992 | -1.524381 | 0.050736 | 0.200122 |
| GOBP_REGULATION_OF_CD4_POSITIVE_ALPHA_BETA_T_CELL_DIFFERENTIATIO | 42 | -0.619804 | -1.524364 | 0.029806 | 0.199568 |
| GOBP_REGULATION_OF_CD8_POSITIVE_ALPHA_BETA_T_CELL_ACTIVATION | 19 | -0.720094 | -1.523727 | 0.037855 | 0.199932 |
| GOBP_LONG_CHAIN_FATTY_ACID_TRANSPORT | 49 | -0.61927 | -1.52249 | 0.031792 | 0.201393 |
| GOBP_REGULATION_OF_SUPRAMOLECULAR_FIBER_ORGANIZATION | 296 | -0.515093 | -1.522148 | 0.001105 | 0.201377 |
| GOBP_CELLULAR_RESPONSE_TO_LOW_DENSITY_LIPOPROTEIN_PARTICLE_STI | 16 | -0.738916 | -1.520804 | 0.030894 | 0.202868 |
| GOBP_POSITIVE_REGULATION_OF_LIPID_LOCALIZATION | 73 | -0.579276 | -1.520073 | 0.02046 | 0.20345 |
| GOBP_MEMBRANE_FUSION | 136 | -0.541048 | -1.520071 | 0.011933 | 0.202881 |
| GOBP_LIPID_IMPORT_INTO_CELL | 28 | -0.673549 | -1.519028 | 0.052795 | 0.203983 |
| GOBP_RESPIRATORY_GASEOUS_EXCHANGE_BY_RESPIRATORY_SYSTEM | 42 | -0.638452 | -1.518875 | 0.035971 | 0.20369 |
| GOBP_REGULATION_OF_TELOMERASE_ACTIVITY | 43 | -0.627567 | -1.518791 | 0.046154 | 0.203284 |
| GOBP_REGULATION_OF_PROTEIN_LOCALIZATION_TO_CELL_PERIPHERY | 91 | -0.571639 | -1.518786 | 0.014139 | 0.202719 |
| GOBP_REGULATION_OF_TISSUE_REMODELING | 51 | -0.597238 | -1.518726 | 0.039492 | 0.202248 |
| GOBP_RESPONSE_TO_MOLECULE_OF_BACTERIAL_ORIGIN | 238 | -0.521241 | -1.518483 | 0.003348 | 0.202053 |
| GOBP_REGULATION_OF_ALPHA_BETA_T_CELL_DIFFERENTIATION | 53 | -0.607436 | -1.518254 | 0.026352 | 0.201884 |
| GOBP_PHOSPHOLIPID_CATABOLIC_PROCESS | 33 | -0.656231 | -1.517807 | 0.045386 | 0.202051 |
| GOBP_EXTERNAL_ENCAPSULATING_STRUCTURE_ORGANIZATION | 239 | -0.518362 | -1.517046 | 0.003375 | 0.202649 |
| GOBP_REGULATION_OF_TRANSPORTER_ACTIVITY | 184 | -0.529241 | -1.516597 | 0.00813 | 0.202893 |
| GOBP_L_GLUTAMATE_TRANSMEMBRANE_TRANSPORT | 16 | -0.740794 | -1.516109 | 0.047541 | 0.203057 |
| GOBP_POSITIVE_REGULATION_OF_ACTIN_FILAMENT_BUNDLE_ASSEMBLY | 50 | -0.60973 | -1.515367 | 0.032922 | 0.203881 |
| GOBP_T_CELL_ACTIVATION_INVOLVED_IN_IMMUNE_RESPONSE | 79 | -0.574958 | -1.515318 | 0.021711 | 0.20342 |
| GOBP_T_HELPER_1_CELL_DIFFERENTIATION | 15 | -0.739051 | -1.514524 | 0.05141 | 0.204025 |
| GOBP_CD4_POSITIVE_ALPHA_BETA_T_CELL_DIFFERENTIATION | 64 | -0.597318 | -1.514443 | 0.022663 | 0.203596 |
| GOBP_POSITIVE_REGULATION_OF_INFLAMMATORY_RESPONSE | 84 | -0.568477 | -1.51438 | 0.016086 | 0.203137 |
| GOBP_POSITIVE_REGULATION_OF_GRANULOCYTE_CHEMOTAXIS | 22 | -0.693392 | -1.513979 | 0.034483 | 0.203281 |
| GOBP_OVULATION_CYCLE_PROCESS | 25 | -0.663263 | -1.511617 | 0.048544 | 0.206613 |
| GOBP_NEGATIVE_REGULATION_OF_INTERLEUKIN_8_PRODUCTION | 18 | -0.711796 | -1.51135 | 0.058917 | 0.206445 |
| GOBP_REGULATION_OF_CYSTEINE_TYPE_ENDOPEPTIDASE_ACTIVITY | 182 | -0.524664 | -1.511224 | 0.011655 | 0.206112 |
| GOBP_POSITIVE_REGULATION_OF_TYROSINE_PHOSPHORYLATION_OF_STAT_PI | 40 | -0.629009 | -1.511219 | 0.051205 | 0.205567 |
| GOBP_POSITIVE_REGULATION_OF_CYTOSKELETON_ORGANIZATION | 179 | -0.528136 | -1.510503 | 0.013064 | 0.206253 |
| GOBP_POSITIVE_REGULATION_OF_ALPHA_BETA_T_CELL_PROLIFERATION | 19 | -0.713869 | -1.509937 | 0.039246 | 0.206566 |
| GOBP_POSITIVE_REGULATION_OF_COAGULATION | 15 | -0.728845 | -1.509912 | 0.048232 | 0.206055 |
| GOBP_REGULATION_OF_DNA_BINDING_TRANSCRIPTION_FACTOR_ACTIVITY | 344 | -0.505699 | -1.509903 | 0.001067 | 0.205528 |
| GOBP_ZYMOGEN_ACTIVATION | 38 | -0.632667 | -1.509738 | 0.045655 | 0.205277 |
| GOBP_REGULATION_OF_AMYLOID_PRECURSOR_PROTEIN_CATABOLIC_PROCES | 34 | -0.635061 | -1.509153 | 0.040404 | 0.205738 |
| GOBP_POSITIVE_REGULATION_OF_VIRAL_GENOME_REPLICATION | 30 | -0.650571 | -1.508785 | 0.033846 | 0.20582 |
| GOBP_REGULATION_OF_LONG_TERM_SYNAPTIC_POTENTIATION | 28 | -0.660211 | -1.508464 | 0.032407 | 0.205809 |
| GOBP_REGULATION_OF_MYELOID_CELL_DIFFERENTIATION | 199 | -0.522959 | -1.508025 | 0.002273 | 0.205935 |
| GOBP_REGULATION_OF_PROTEIN_POLYMERIZATION | 184 | -0.523231 | -1.507585 | 0.005862 | 0.20607 |
| GOBP_ESTROGEN_METABOLIC_PROCESS | 21 | -0.701624 | -1.50742 | 0.037559 | 0.205814 |
| GOBP_REACTIVE_OXYGEN_SPECIES_BIOSYNTHETIC_PROCESS | 91 | -0.566187 | -1.506642 | 0.019206 | 0.206537 |
| GOBP_HUMORAL_IMMUNE_RESPONSE | 161 | -0.538033 | -1.506165 | 0.007308 | 0.206766 |
| GOBP_REGULATION_OF_ISOTYPE_SWITCHING | 33 | -0.646672 | -1.504706 | 0.049924 | 0.208669 |
| GOBP_CARBOXYLIC_ACID_TRANSPORT | 171 | -0.526267 | -1.504544 | 0.008304 | 0.208409 |
| GOBP_NEGATIVE_REGULATION_OF_ENDOTHELIAL_CELL_PROLIFERATION | 29 | -0.653778 | -1.50409 | 0.045872 | 0.208615 |
| GOBP_ENTRAINMENT_OF_CIRCADIAN_CLOCK | 29 | -0.65675 | -1.503976 | 0.046269 | 0.208305 |
| GOBP_MICROGLIAL_CELL_ACTIVATION | 32 | -0.653444 | -1.503568 | 0.045845 | 0.20844 |
| GOBP_REGULATION_OF_CYTOSKELETON_ORGANIZATION | 408 | -0.497286 | -1.503347 | 0 | 0.208273 |
| GOBP_NEGATIVE_REGULATION_OF_OSTEOBLAST_DIFFERENTIATION | 27 | -0.662757 | -1.502659 | 0.037975 | 0.208774 |
| GOBP_NEGATIVE_REGULATION_OF_NATURAL_KILLER_CELL_MEDIATED_IMMUN | 15 | -0.729313 | -1.502639 | 0.052805 | 0.208297 |
| GOBP_LIPOPOLYSACCHARIDE_MEDIATED_SIGNALING_PATHWAY | 50 | -0.611146 | -1.502514 | 0.04507 | 0.207968 |
| GOBP_NEGATIVE_REGULATION_OF_DNA_BINDING_TRANSCRIPTION_FACTOR_A | 132 | -0.5394 | -1.500467 | 0.013366 | 0.210792 |
| GOBP_REGULATION_OF_TRIGLYCERIDE_METABOLIC_PROCESS | 26 | -0.666756 | -1.500436 | 0.036866 | 0.2103 |
| GOBP_OVULATION_CYCLE | 37 | -0.634937 | -1.500395 | 0.044053 | 0.209846 |
| GOBP_ACUTE_PHASE_RESPONSE | 25 | -0.680693 | -1.500082 | 0.03438 | 0.209846 |
| GOBP_POSITIVE_REGULATION_OF_CD4_POSITIVE_ALPHA_BETA_T_CELL_ACTIV | 33 | -0.651717 | -1.49926 | 0.038292 | 0.210631 |
| GOBP_MAINTENANCE_OF_LOCATION | 236 | -0.516319 | -1.49925 | 0.004614 | 0.210124 |
| GOBP_REGULATION_OF_NEURON_PROJECTION_REGENERATION | 22 | -0.688712 | -1.499183 | 0.045161 | 0.209685 |
| GOBP_INTERLEUKIN_8_PRODUCTION | 67 | -0.58241 | -1.499182 | 0.029333 | 0.209172 |
| GOBP_VIRAL_LIFE_CYCLE | 305 | -0.497809 | -1.498468 | 0.003282 | 0.20997 |
| GOBP_DEFENSE_RESPONSE_TO_GRAM_POSITIVE_BACTERIUM | 51 | -0.596873 | -1.49735 | 0.038082 | 0.211414 |
| GOBP_POSITIVE_REGULATION_OF_PLASMA_MEMBRANE_BOUNDED_CELL_PRO | 85 | -0.558267 | -1.496896 | 0.016624 | 0.211653 |
| GOBP_REGULATION_OF_TOLERANCE_INDUCTION | 15 | -0.730466 | -1.496497 | 0.039931 | 0.211808 |
| GOBP_RHO_PROTEIN_SIGNAL_TRANSDUCTION | 103 | -0.553222 | -1.496426 | 0.010309 | 0.211404 |
| GOBP_EMBRYONIC_PLACENTA_DEVELOPMENT | 55 | -0.602543 | -1.496059 | 0.037657 | 0.211529 |
| GOBP_REGULATION_OF_FATTY_ACID_BIOSYNTHETIC_PROCESS | 33 | -0.649719 | -1.494293 | 0.045994 | 0.21388 |
| GOBP_ANTIGEN_PROCESSING_AND_PRESENTATION | 210 | -0.511288 | -1.493668 | 0.005727 | 0.214379 |
| GOBP_REGULATION_OF_PEPTIDYL_TYROSINE_PHOSPHORYLATION | 175 | -0.520062 | -1.493172 | 0.015403 | 0.214651 |
| GOBP_PROTEIN_LOCALIZATION_TO_CELL_SURFACE | 55 | -0.599986 | -1.492291 | 0.032033 | 0.215623 |
| GOBP_CHOLESTEROL_EFFLUX | 35 | -0.637146 | -1.492026 | 0.044881 | 0.215597 |
| GOBP_REGULATION_OF_ATP_BIOSYNTHETIC_PROCESS | 16 | -0.722833 | -1.490878 | 0.055988 | 0.21693 |
| GOBP_INTERFERON_GAMMA_PRODUCTION | 89 | -0.566586 | -1.490793 | 0.019557 | 0.216541 |
| GOBP_REGULATION_OF_REACTIVE_OXYGEN_SPECIES_METABOLIC_PROCESS | 144 | -0.526009 | -1.48959 | 0.013333 | 0.218046 |
| GOBP_STRESS_FIBER_ASSEMBLY | 83 | -0.561432 | -1.48904 | 0.019011 | 0.218479 |
| GOBP_MEMBRANE_PROTEIN_ECTODOMAIN_PROTEOLYSIS | 36 | -0.634493 | -1.489004 | 0.044248 | 0.218017 |
| GOBP_ENDODERM_DEVELOPMENT | 54 | -0.59603 | -1.488615 | 0.04742 | 0.21817 |
| GOBP_SYNCYTIUM_FORMATION | 38 | -0.622768 | -1.488291 | 0.044993 | 0.218181 |
| GOBP_POSITIVE_REGULATION_OF_FAT_CELL_DIFFERENTIATION | 42 | -0.616284 | -1.487142 | 0.037736 | 0.219658 |
| GOBP_MUSCLE_CELL_PROLIFERATION | 125 | -0.534254 | -1.486816 | 0.013301 | 0.21967 |
| GOBP_REGULATION_OF_EPIDERMIS_DEVELOPMENT | 46 | -0.606971 | -1.486498 | 0.045911 | 0.219741 |
| GOBP_NEURAL_TUBE_PATTERNING | 22 | -0.698088 | -1.486286 | 0.06793 | 0.219588 |
| GOBP_CELL_JUNCTION_ORGANIZATION | 449 | -0.486845 | -1.485902 | 0 | 0.219808 |
| GOBP_NEGATIVE_REGULATION_OF_HYDROLASE_ACTIVITY | 289 | -0.499685 | -1.485856 | 0.005507 | 0.219391 |
| GOBP_REGULATION_OF_CELL_JUNCTION_ASSEMBLY | 117 | -0.539243 | -1.485671 | 0.011139 | 0.219171 |
| GOBP_CD4_POSITIVE_ALPHA_BETA_T_CELL_ACTIVATION | 80 | -0.561639 | -1.485484 | 0.027888 | 0.218928 |
| GOBP_POSITIVE_REGULATION_OF_CATION_TRANSMEMBRANE_TRANSPORT | 87 | -0.554259 | -1.485091 | 0.033333 | 0.219139 |
| GOBP_DEFENSE_RESPONSE_TO_BACTERIUM | 158 | -0.522858 | -1.485058 | 0.00479 | 0.218687 |
| GOBP_POSITIVE_REGULATION_OF_ADAPTIVE_IMMUNE_RESPONSE | 90 | -0.564154 | -1.485015 | 0.018963 | 0.21825 |
| GOBP_POSITIVE_REGULATION_OF_INTERFERON_GAMMA_PRODUCTION | 55 | -0.595714 | -1.485014 | 0.024725 | 0.217749 |
| GOBP_CELL_MIGRATION_INVOLVED_IN_SPROUTING_ANGIOGENESIS | 37 | -0.619554 | -1.483469 | 0.048632 | 0.219926 |
| GOBP_NEGATIVE_REGULATION_OF_RESPONSE_TO_CYTOKINE_STIMULUS | 48 | -0.599827 | -1.483118 | 0.044097 | 0.219903 |
| GOBP_POSITIVE_REGULATION_OF_RESPONSE_TO_WOUNDING | 47 | -0.609017 | -1.482322 | 0.04661 | 0.220749 |
| GOBP_RAS_PROTEIN_SIGNAL_TRANSDUCTION | 289 | -0.498708 | -1.482163 | 0 | 0.220545 |

| | | | | | |
|---|---|---|---|---|---|
| GOBP_MATURATION_OF_5_8S_RRNA | 35 | 0.367636 | 0.973568 | 0.5 | 0.963174 |
| GOBP_NEGATIVE_REGULATION_OF_RECEPTOR_SIGNALING_PATHWAY_VIA_JAK_STAT | 18 | 0.404144 | 0.97282 | 0.477833 | 0.963489 |
| GOBP_RESPONSE_TO_IRON_ION | 20 | 0.398991 | 0.972748 | 0.491003 | 0.961545 |
| GOBP_NEGATIVE_REGULATION_OF_HISTONE_METHYLATION | 21 | 0.397003 | 0.972634 | 0.494475 | 0.959759 |
| GOBP_PURINE_CONTAINING_COMPOUND_BIOSYNTHETIC_PROCESS | 174 | 0.2829 | 0.972389 | 0.479452 | 0.958418 |
| GOBP_GANGLIOSIDE_METABOLIC_PROCESS | 18 | 0.41538 | 0.972204 | 0.505181 | 0.956842 |
| GOBP_ASPARTATE_FAMILY_AMINO_ACID_CATABOLIC_PROCESS | 17 | 0.420043 | 0.971905 | 0.506427 | 0.95572 |
| GOBP_POSITIVE_REGULATION_OF_NEURAL_PRECURSOR_CELL_PROLIFERATION | 27 | 0.369447 | 0.971259 | 0.507246 | 0.955656 |
| GOBP_MITOCHONDRIAL_GENE_EXPRESSION | 164 | 0.280639 | 0.970097 | 0.503106 | 0.957574 |
| GOBP_CELLULAR_AMINO_ACID_METABOLIC_PROCESS | 279 | 0.260985 | 0.969279 | 0.541177 | 0.95814 |
| GOBP_REGULATION_OF_JUN_KINASE_ACTIVITY | 68 | 0.317691 | 0.968515 | 0.544061 | 0.958483 |
| GOBP_RESPONSE_TO_VIRUS | 283 | 0.258955 | 0.96676 | 0.59375 | 0.962361 |
| GOBP_CELLULAR_RESPONSE_TO_DEXAMETHASONE_STIMULUS | 23 | 0.385516 | 0.965842 | 0.493036 | 0.963237 |
| GOBP_CHROMATIN_DISASSEMBLY | 17 | 0.403257 | 0.965483 | 0.506204 | 0.962326 |
| GOBP_POSITIVE_REGULATION_OF_INTERFERON_ALPHA_PRODUCTION | 20 | 0.391036 | 0.96427 | 0.488432 | 0.964263 |
| GOBP_INTRINSIC_APOPTOTIC_SIGNALING_PATHWAY_IN_RESPONSE_TO_DNA_DAMAG | 41 | 0.337556 | 0.963875 | 0.531353 | 0.963465 |
| GOBP_PEROXISOMAL_MEMBRANE_TRANSPORT | 20 | 0.394331 | 0.962857 | 0.496241 | 0.964782 |
| GOBP_REGULATION_OF_INTRACELLULAR_ESTROGEN_RECEPTOR_SIGNALING_PATHW | 30 | 0.357895 | 0.962802 | 0.521472 | 0.962831 |
| GOBP_NEGATIVE_REGULATION_OF_RECEPTOR_SIGNALING_PATHWAY_VIA_STAT | 18 | 0.404144 | 0.959895 | 0.506427 | 0.970473 |
| GOBP_PROTEIN_TETRAMERIZATION | 62 | 0.319339 | 0.957771 | 0.487544 | 0.975349 |
| GOBP_NEGATIVE_REGULATION_OF_ORGAN_GROWTH | 25 | 0.383813 | 0.956839 | 0.525641 | 0.976289 |
| GOBP_RIBONUCLEOTIDE_CATABOLIC_PROCESS | 32 | 0.348606 | 0.956666 | 0.527508 | 0.974831 |
| GOBP_POSITIVE_REGULATION_OF_BONE_MINERALIZATION | 22 | 0.379703 | 0.956562 | 0.491892 | 0.973032 |
| GOBP_POSITIVE_REGULATION_OF_MORPHOGENESIS_OF_AN_EPITHELIUM | 19 | 0.40164 | 0.956175 | 0.519663 | 0.972203 |
| GOBP_HEMATOPOIETIC_STEM_CELL_DIFFERENTIATION | 81 | 0.296109 | 0.955499 | 0.538462 | 0.972245 |
| GOBP_ENDOCHONDRAL_BONE_MORPHOGENESIS | 31 | 0.355933 | 0.951939 | 0.523121 | 0.982122 |
| GOBP_INTRINSIC_APOPTOTIC_SIGNALING_PATHWAY_IN_RESPONSE_TO_ENDOPLASM | 55 | 0.329995 | 0.951259 | 0.521898 | 0.982132 |
| GOBP_POSITIVE_REGULATION_OF_CELL_CYCLE_G2_M_PHASE_TRANSITION | 28 | 0.368644 | 0.951082 | 0.516129 | 0.980606 |
| GOBP_FIBROBLAST_APOPTOTIC_PROCESS | 19 | 0.38901 | 0.948683 | 0.535065 | 0.986389 |
| GOBP_REGULATION_OF_ADENYLATE_CYCLASE_ACTIVITY | 18 | 0.398031 | 0.948651 | 0.526455 | 0.984397 |
| GOBP_INTERMEDIATE_FILAMENT_BASED_PROCESS | 23 | 0.371325 | 0.948495 | 0.567568 | 0.982802 |
| GOBP_OLIGOSACCHARIDE_LIPID_INTERMEDIATE_BIOSYNTHETIC_PROCESS | 21 | 0.384701 | 0.947611 | 0.528302 | 0.983533 |
| GOBP_CELLULAR_RESPONSE_TO_DRUG | 50 | 0.318885 | 0.946693 | 0.55814 | 0.984475 |
| GOBP_BIOLOGICAL_PROCESS_INVOLVED_IN_INTERACTION_WITH_SYMBIONT | 62 | 0.313853 | 0.946399 | 0.567669 | 0.983366 |
| GOBP_MYOBLAST_DIFFERENTIATION | 49 | 0.324971 | 0.941264 | 0.539855 | 0.998172 |
| GOBP_METANEPHRIC_NEPHRON_DEVELOPMENT | 17 | 0.400679 | 0.941079 | 0.531017 | 0.996702 |
| GOBP_CELLULAR_RESPONSE_TO_INTERFERON_BETA | 17 | 0.394035 | 0.940658 | 0.515152 | 0.995889 |
| GOBP_CHEMICAL_SYNAPTIC_TRANSMISSION_POSTSYNAPTIC | 64 | 0.312869 | 0.9401 | 0.53125 | 0.995634 |
| GOBP_POSITIVE_REGULATION_OF_POSTTRANSCRIPTIONAL_GENE_SILENCING | 24 | 0.374583 | 0.939045 | 0.534759 | 0.996948 |
| GOBP_RESPONSE_TO_PLATELET_DERIVED_GROWTH_FACTOR | 16 | 0.408732 | 0.938931 | 0.55291 | 0.99524 |
| GOBP_DIOL_METABOLIC_PROCESS | 26 | 0.370336 | 0.937513 | 0.557471 | 0.997749 |
| GOBP_SULFUR_COMPOUND_METABOLIC_PROCESS | 287 | 0.248374 | 0.937082 | 0.722892 | 0.997031 |
| GOBP_PIGMENT_METABOLIC_PROCESS | 57 | 0.313071 | 0.936987 | 0.577558 | 0.995231 |
| GOBP_REGULATION_OF_LYASE_ACTIVITY | 28 | 0.35651 | 0.936507 | 0.527933 | 0.994596 |
| GOBP_NEGATIVE_REGULATION_OF_CELLULAR_SENESCENCE | 15 | 0.406144 | 0.935087 | 0.549223 | 0.997089 |
| GOBP_PURINE_NUCLEOSIDE_MONOPHOSPHATE_METABOLIC_PROCESS | 40 | 0.322651 | 0.933174 | 0.550162 | 1 |
| GOBP_PROTEASOMAL_UBIQUITIN_INDEPENDENT_PROTEIN_CATABOLIC_PROCESS | 21 | 0.378225 | 0.931578 | 0.534946 | 1 |
| GOBP_GLUCOSE_CATABOLIC_PROCESS | 33 | 0.339929 | 0.931532 | 0.560472 | 1 |
| GOBP_NUCLEOTIDE_SUGAR_METABOLIC_PROCESS | 35 | 0.336826 | 0.931348 | 0.567308 | 1 |
| GOBP_NEGATIVE_REGULATION_OF_PEPTIDYL_SERINE_PHOSPHORYLATION | 24 | 0.372855 | 0.931258 | 0.557103 | 0.998861 |
| GOBP_RIBOSOMAL_SMALL_SUBUNIT_BIOGENESIS | 75 | 0.295791 | 0.930393 | 0.572549 | 0.999448 |
| GOBP_NUCLEOSIDE_PHOSPHATE_BIOSYNTHETIC_PROCESS | 222 | 0.256409 | 0.929856 | 0.637931 | 0.999111 |
| GOBP_VASODILATION | 21 | 0.374425 | 0.92878 | 0.547315 | 1 |
| GOBP_NEGATIVE_REGULATION_OF_PLASMA_MEMBRANE_BOUNDED_CELL_PROJECTI | 24 | 0.358639 | 0.928049 | 0.560976 | 1 |
| GOBP_PEPTIDYL_SERINE_DEPHOSPHORYLATION | 16 | 0.402858 | 0.927252 | 0.574413 | 1 |
| GOBP_DETECTION_OF_STIMULUS_INVOLVED_IN_SENSORY_PERCEPTION | 66 | 0.313525 | 0.92565 | 0.581481 | 1 |
| GOBP_ACETYL_COA_METABOLIC_PROCESS | 33 | 0.342779 | 0.921399 | 0.5625 | 1 |
| GOBP_MITOTIC_CYTOKINESIS | 66 | 0.299331 | 0.920558 | 0.585271 | 1 |
| GOBP_REGULATION_OF_PROTEIN_MATURATION | 50 | 0.315742 | 0.918783 | 0.633562 | 1 |
| GOBP_ERAD_PATHWAY | 97 | 0.275759 | 0.918693 | 0.600939 | 1 |
| GOBP_NEGATIVE_REGULATION_OF_CELL_CYCLE_ARREST | 15 | 0.402045 | 0.91848 | 0.566327 | 1 |
| GOBP_NEGATIVE_REGULATION_OF_GTPASE_ACTIVITY | 38 | 0.333534 | 0.918233 | 0.573379 | 1 |
| GOBP_LIPID_DROPLET_ORGANIZATION | 21 | 0.372117 | 0.916155 | 0.5427 | 1 |
| GOBP_RIBOSE_PHOSPHATE_BIOSYNTHETIC_PROCESS | 168 | 0.260852 | 0.916149 | 0.680272 | 1 |
| GOBP_REPRODUCTIVE_BEHAVIOR | 18 | 0.387507 | 0.915168 | 0.568 | 1 |
| GOBP_ENDOPLASMIC_RETICULUM_TO_CYTOSOL_TRANSPORT | 28 | 0.347177 | 0.91506 | 0.572222 | 1 |
| GOBP_NEGATIVE_REGULATION_OF_PROTEIN_TYROSINE_KINASE_ACTIVITY | 27 | 0.351577 | 0.914053 | 0.575 | 1 |
| GOBP_I_KAPPAB_PHOSPHORYLATION | 17 | 0.382752 | 0.913151 | 0.572482 | 1 |
| GOBP_PRIMARY_NEURAL_TUBE_FORMATION | 70 | 0.295084 | 0.912612 | 0.639344 | 1 |
| GOBP_MATURATION_OF_SSU_RRNA_FROM_TRICISTRONIC_RRNA_TRANSCRIPT_SSU_ | 37 | 0.33087 | 0.912572 | 0.600649 | 1 |
| GOBP_REGULATION_OF_CARDIAC_MUSCLE_CELL_PROLIFERATION | 22 | 0.360383 | 0.912557 | 0.585492 | 1 |
| GOBP_NCRNA_3_END_PROCESSING | 38 | 0.3317 | 0.911978 | 0.59596 | 1 |
| GOBP_PROTEIN_MATURATION_BY_IRON_SULFUR_CLUSTER_TRANSFER | 16 | 0.383898 | 0.911353 | 0.542234 | 1 |
| GOBP_SHORT_CHAIN_FATTY_ACID_METABOLIC_PROCESS | 15 | 0.396711 | 0.90889 | 0.564403 | 1 |
| GOBP_CYTOSKELETON_DEPENDENT_CYTOKINESIS | 88 | 0.277393 | 0.906625 | 0.664 | 1 |
| GOBP_ESTABLISHMENT_OF_TISSUE_POLARITY | 105 | 0.270823 | 0.906142 | 0.691542 | 1 |
| GOBP_CELLULAR_PIGMENTATION | 46 | 0.314423 | 0.903434 | 0.6 | 1 |
| GOBP_CELLULAR_GLUCOSE_HOMEOSTASIS | 106 | 0.272852 | 0.901386 | 0.678392 | 1 |
| GOBP_POSITIVE_REGULATION_OF_TRANSCRIPTION_FROM_RNA_POLYMERASE_II_PR( | 18 | 0.381137 | 0.900842 | 0.581967 | 1 |
| GOBP_DNA_CATABOLIC_PROCESS | 34 | 0.327491 | 0.899851 | 0.579105 | 1 |
| GOBP_REGULATION_OF_ENDOTHELIAL_CELL_DIFFERENTIATION | 25 | 0.354778 | 0.897924 | 0.60221 | 1 |
| GOBP_MIDBRAIN_DEVELOPMENT | 59 | 0.301709 | 0.896276 | 0.675472 | 1 |
| GOBP_PEPTIDYL_THREONINE_MODIFICATION | 112 | 0.266697 | 0.896165 | 0.703518 | 1 |
| GOBP_PHOSPHATIDYLCHOLINE_ACYL_CHAIN_REMODELING | 19 | 0.363926 | 0.895807 | 0.595855 | 1 |
| GOBP_NUCLEOBASE_BIOSYNTHETIC_PROCESS | 17 | 0.371777 | 0.894766 | 0.581683 | 1 |
| GOBP_CHONDROITIN_SULFATE_BIOSYNTHETIC_PROCESS | 17 | 0.376531 | 0.894513 | 0.563636 | 1 |
| GOBP_FEAR_RESPONSE | 19 | 0.372962 | 0.894253 | 0.555233 | 1 |
| GOBP_CHROMATIN_ASSEMBLY_OR_DISASSEMBLY | 166 | 0.257402 | 0.893794 | 0.766917 | 1 |
| GOBP_POSITIVE_REGULATION_OF_LEUKOCYTE_ADHESION_TO_VASCULAR_ENDOTHE | 18 | 0.369788 | 0.893322 | 0.582633 | 1 |
| GOBP_MITOTIC_G1_S_TRANSITION_CHECKPOINT | 58 | 0.29444 | 0.88813 | 0.683849 | 1 |
| GOBP_REGULATION_OF_GLUCOSE_IMPORT | 40 | 0.320111 | 0.886077 | 0.628931 | 1 |
| GOBP_POSITIVE_REGULATION_OF_NITRIC_OXIDE_SYNTHASE_ACTIVITY | 20 | 0.368367 | 0.885501 | 0.600544 | 1 |
| GOBP_ACTIVATION_OF_ADENYLATE_CYCLASE_ACTIVITY | 19 | 0.369305 | 0.885018 | 0.654255 | 1 |
| GOBP_ASPARTATE_FAMILY_AMINO_ACID_BIOSYNTHETIC_PROCESS | 17 | 0.367725 | 0.883905 | 0.601604 | 1 |
| GOBP_POSITIVE_REGULATION_OF_PROTEIN_CONTAINING_COMPLEX_DISASSEMBLY | 27 | 0.346247 | 0.883385 | 0.614493 | 1 |
| GOBP_RESPONSE_TO_ATP | 23 | 0.354382 | 0.88331 | 0.615819 | 1 |
| GOBP_REGULATION_OF_ENDOPLASMIC_RETICULUM_UNFOLDED_PROTEIN_RESPONS | 27 | 0.338984 | 0.878559 | 0.642442 | 1 |
| GOBP_DEOXYRIBONUCLEOTIDE_METABOLIC_PROCESS | 35 | 0.317321 | 0.877909 | 0.625 | 1 |
| GOBP_IRON_ION_TRANSPORT | 63 | 0.291495 | 0.876204 | 0.664063 | 1 |
| GOBP_NADH_DEHYDROGENASE_COMPLEX_ASSEMBLY | 65 | 0.284739 | 0.875457 | 0.681452 | 1 |
| GOBP_PROTEIN_EXIT_FROM_ENDOPLASMIC_RETICULUM | 40 | 0.313092 | 0.875324 | 0.631579 | 1 |
| GOBP_POSITIVE_REGULATION_OF_CELL_CYCLE_ARREST | 70 | 0.285528 | 0.872642 | 0.682657 | 1 |
| GOBP_NEGATIVE_REGULATION_OF_MUSCLE_HYPERTROPHY | 23 | 0.346617 | 0.871426 | 0.651351 | 1 |
| GOBP_AXO_DENDRITIC_TRANSPORT | 62 | 0.287468 | 0.870967 | 0.688172 | 1 |
| GOBP_NUCLEAR_ENVELOPE_ORGANIZATION | 49 | 0.303902 | 0.870802 | 0.708475 | 1 |
| GOBP_ESTABLISHMENT_OF_ENDOTHELIAL_BARRIER | 38 | 0.308271 | 0.869057 | 0.649068 | 1 |
| GOBP_VIRAL_BUDDING_VIA_HOST_ESCRT_COMPLEX | 20 | 0.358897 | 0.868293 | 0.615776 | 1 |
| GOBP_TRNA_METHYLATION | 39 | 0.305838 | 0.86756 | 0.695364 | 1 |
| GOBP_MITOTIC_CYTOKINETIC_PROCESS | 23 | 0.342987 | 0.866792 | 0.639205 | 1 |
| GOBP_DNA_REPLICATION_INDEPENDENT_NUCLEOSOME_ORGANIZATION | 47 | 0.293114 | 0.866279 | 0.702614 | 1 |
| GOBP_REGULATION_OF_CHOLESTEROL_BIOSYNTHETIC_PROCESS | 40 | 0.305626 | 0.864931 | 0.69 | 1 |
| GOBP_MIDBODY_ABSCISSION | 17 | 0.37368 | 0.863901 | 0.624 | 1 |
| GOBP_SPHINGOLIPID_METABOLIC_PROCESS | 133 | 0.253437 | 0.863713 | 0.797619 | 1 |
| GOBP_OSTEOBLAST_PROLIFERATION | 18 | 0.367393 | 0.862351 | 0.663889 | 1 |
| GOBP_CHROMOSOME_ORGANIZATION_INVOLVED_IN_MEIOTIC_CELL_CYCLE | 50 | 0.297363 | 0.859011 | 0.680272 | 1 |
| GOBP_REGULATION_OF_TRANSLATIONAL_ELONGATION | 18 | 0.359123 | 0.858288 | 0.637641 | 1 |
| GOBP_POSTSYNAPTIC_NEUROTRANSMITTER_RECEPTOR_INTERNALIZATION | 15 | 0.373906 | 0.856559 | 0.628916 | 1 |
| GOBP_RIBONUCLEOPROTEIN_COMPLEX_BIOGENESIS | 436 | 0.22241 | 0.855576 | 0.961538 | 1 |
| GOBP_GENETIC_IMPRINTING | 20 | 0.338627 | 0.853574 | 0.682099 | 1 |
| GOBP_LAMELLIPODIUM_ORGANIZATION | 76 | 0.273392 | 0.853413 | 0.743363 | 1 |
| GOBP_DENDRITIC_SPINE_MAINTENANCE | 18 | 0.355775 | 0.853184 | 0.664 | 1 |
| GOBP_TERMINATION_OF_RNA_POLYMERASE_I_TRANSCRIPTION | 30 | 0.327803 | 0.85259 | 0.653061 | 1 |
| GOBP_NUCLEOBASE_METABOLIC_PROCESS | 28 | 0.331481 | 0.851731 | 0.67147 | 1 |
| GOBP_NEGATIVE_REGULATION_OF_REACTIVE_OXYGEN_SPECIES_METABOLIC_PROC | 43 | 0.293731 | 0.851708 | 0.712375 | 1 |
| GOBP_CELLULAR_CARBOHYDRATE_CATABOLIC_PROCESS | 39 | 0.303448 | 0.851682 | 0.673203 | 1 |
| GOBP_NEGATIVE_REGULATION_OF_NECROTIC_CELL_DEATH | 15 | 0.363846 | 0.851571 | 0.668342 | 1 |
| GOBP_SPINDLE_ASSEMBLY | 112 | 0.257403 | 0.851109 | 0.832461 | 1 |
| GOBP_REGULATION_OF_CELL_CYCLE_ARREST | 87 | 0.266824 | 0.85092 | 0.79638 | 1 |
| GOBP_PURINE_CONTAINING_COMPOUND_METABOLIC_PROCESS | 370 | 0.225618 | 0.849537 | 0.941177 | 1 |
| GOBP_NUCLEOTIDE_BINDING_DOMAIN_LEUCINE_RICH_REPEAT_CONTAINING_RECEP | 41 | 0.297748 | 0.849021 | 0.702875 | 1 |
| GOBP_NUCLEUS_ORGANIZATION | 113 | 0.251893 | 0.847167 | 0.830097 | 1 |
| GOBP_NEGATIVE_REGULATION_OF_NUCLEAR_DIVISION | 50 | 0.289425 | 0.847133 | 0.727273 | 1 |
| GOBP_NEGATIVE_REGULATION_OF_CARDIAC_MUSCLE_TISSUE_GROWTH | 18 | 0.356298 | 0.846792 | 0.66838 | 1 |
| GOBP_MULTICELLULAR_ORGANISM_AGING | 21 | 0.356961 | 0.846131 | 0.698061 | 1 |
| GOBP_RIBONUCLEOSIDE_MONOPHOSPHATE_METABOLIC_PROCESS | 51 | 0.291842 | 0.846011 | 0.744615 | 1 |
| GOBP_POSITIVE_REGULATION_OF_RNA_SPLICING | 34 | 0.309421 | 0.844332 | 0.667665 | 1 |
| GOBP_REGULATION_OF_RESPONSE_TO_ENDOPLASMIC_RETICULUM_STRESS | 75 | 0.270402 | 0.844225 | 0.793522 | 1 |
| GOBP_PYRIMIDINE_CONTAINING_COMPOUND_METABOLIC_PROCESS | 73 | 0.261435 | 0.844181 | 0.79476 | 1 |
| GOBP_PROTEIN_LOCALIZATION_TO_CHROMOSOME | 89 | 0.261938 | 0.844076 | 0.776316 | 1 |
| GOBP_MORPHOGENESIS_OF_A_POLARIZED_EPITHELIUM | 119 | 0.256396 | 0.843271 | 0.814815 | 1 |
| GOBP_MAINTENANCE_OF_BLOOD_BRAIN_BARRIER | 22 | 0.338446 | 0.843184 | 0.665635 | 1 |
| GOBP_REGULATION_OF_RAC_PROTEIN_SIGNAL_TRANSDUCTION | 18 | 0.347126 | 0.842802 | 0.663185 | 1 |
| GOBP_REGULATION_OF_PEPTIDYL_LYSINE_ACETYLATION | 54 | 0.284205 | 0.837138 | 0.74359 | 1 |
| GOBP_REGULATION_OF_SENSORY_PERCEPTION | 18 | 0.344882 | 0.83554 | 0.654891 | 1 |
| GOBP_GOLGI_ORGANIZATION | 131 | 0.240347 | 0.835317 | 0.876405 | 1 |
| GOBP_HEAD_MORPHOGENESIS | 24 | 0.333762 | 0.833696 | 0.684066 | 1 |
| GOBP_REGULATION_OF_MITOCHONDRIAL_GENE_EXPRESSION | 29 | 0.31235 | 0.832784 | 0.714286 | 1 |

| | | | | | |
|---|---|---|---|---|---|
| GOBP_LYMPHOCYTE_COSTIMULATION | 51 | -0.595145 | -1.482076 | 0.033287 | 0.220194 |
| GOBP_SEMI_LUNAR_VALVE_DEVELOPMENT | 20 | -0.697747 | -1.481615 | 0.047096 | 0.22049 |
| GOBP_REGULATION_OF_GLIAL_CELL_DIFFERENTIATION | 37 | -0.625695 | -1.480829 | 0.054676 | 0.221254 |
| GOBP_T_CELL_MEDIATED_CYTOTOXICITY | 38 | -0.619237 | -1.48008 | 0.051593 | 0.222007 |
| GOBP_POSITIVE_REGULATION_OF_WOUND_HEALING | 38 | -0.631921 | -1.479857 | 0.045845 | 0.2219 |
| GOBP_POSITIVE_REGULATION_OF_MYELOID_CELL_DIFFERENTIATION | 80 | -0.559599 | -1.479416 | 0.032852 | 0.222126 |
| GOBP_FATTY_ACID_TRANSPORT | 98 | -0.544247 | -1.479171 | 0.029151 | 0.222074 |
| GOBP_COLLAGEN_METABOLIC_PROCESS | 69 | -0.570833 | -1.479099 | 0.023904 | 0.22169 |
| GOBP_CELLULAR_RESPONSE_TO_HYDROGEN_PEROXIDE | 74 | -0.575819 | -1.478836 | 0.036635 | 0.221643 |
| GOBP_PROTEIN_KINASE_B_SIGNALING | 175 | -0.518181 | -1.478392 | 0.008216 | 0.221878 |
| GOBP_MEMBRANE_BIOGENESIS | 30 | -0.640491 | -1.478369 | 0.056489 | 0.221427 |
| GOBP_T_CELL_MEDIATED_IMMUNITY | 84 | -0.553112 | -1.477936 | 0.029062 | 0.221639 |
| GOBP_NEGATIVE_REGULATION_OF_PEPTIDASE_ACTIVITY | 138 | -0.532584 | -1.477897 | 0.013317 | 0.221212 |
| GOBP_PEPTIDYL_CYSTEINE_MODIFICATION | 41 | -0.609973 | -1.477773 | 0.07278 | 0.220913 |
| GOBP_CHEMOKINE_PRODUCTION | 67 | -0.571285 | -1.476193 | 0.036986 | 0.222869 |
| GOBP_CELLULAR_RESPONSE_TO_ALCOHOL | 61 | -0.589691 | -1.475989 | 0.037448 | 0.222743 |
| GOBP_NEGATIVE_REGULATION_OF_B_CELL_ACTIVATION | 29 | -0.640271 | -1.475575 | 0.052308 | 0.222922 |
| GOBP_POSITIVE_REGULATION_OF_T_HELPER_CELL_DIFFERENTIATION | 18 | -0.693313 | -1.474652 | 0.058537 | 0.223861 |
| GOBP_CELLULAR_RESPONSE_TO_LIPOPROTEIN_PARTICLE_STIMULUS | 27 | -0.658829 | -1.474605 | 0.054688 | 0.223457 |
| GOBP_SIGNALING_RECEPTOR_LIGAND_PRECURSOR_PROCESSING | 18 | -0.686344 | -1.473458 | 0.072115 | 0.224789 |
| GOBP_PROSTANOID_METABOLIC_PROCESS | 26 | -0.665285 | -1.473204 | 0.062305 | 0.224784 |
| GOBP_O_GLYCAN_PROCESSING | 40 | -0.60365 | -1.472676 | 0.066766 | 0.225128 |
| GOBP_CELLULAR_RESPONSE_TO_STEROL | 16 | -0.700069 | -1.472619 | 0.067545 | 0.224744 |
| GOBP_REGULATION_OF_CELL_MORPHOGENESIS | 237 | -0.505311 | -1.472284 | 0.005656 | 0.224818 |
| GOBP_REGULATION_OF_PROTEIN_POLYUBIQUITINATION | 22 | -0.674971 | -1.471921 | 0.068071 | 0.224948 |
| GOBP_REGULATION_OF_PLASMA_LIPOPROTEIN_PARTICLE_LEVELS | 63 | -0.584407 | -1.470172 | 0.038199 | 0.227505 |
| GOBP_PLASMA_MEMBRANE_ORGANIZATION | 85 | -0.554975 | -1.47017 | 0.030383 | 0.227014 |
| GOBP_NEGATIVE_REGULATION_OF_T_CELL_MEDIATED_IMMUNITY | 19 | -0.693817 | -1.470059 | 0.072013 | 0.226733 |
| GOBP_REGULATION_OF_WOUND_HEALING | 82 | -0.559391 | -1.469954 | 0.035762 | 0.226426 |
| GOBP_TERPENOID_METABOLIC_PROCESS | 62 | -0.575314 | -1.468174 | 0.038147 | 0.229016 |
| GOBP_POSITIVE_REGULATION_OF_IMMUNOGLOBULIN_PRODUCTION | 37 | -0.61743 | -1.467955 | 0.054834 | 0.228915 |
| GOBP_NEGATIVE_REGULATION_OF_INTRINSIC_APOPTOTIC_SIGNALING_PATHW | 80 | -0.558958 | -1.467398 | 0.030223 | 0.229408 |
| GOBP_CELL_SUBSTRATE_ADHESION | 251 | -0.499505 | -1.467289 | 0.006834 | 0.229111 |
| GOBP_AXON_DEVELOPMENT | 334 | -0.49133 | -1.466889 | 0 | 0.229308 |
| GOBP_LIPID_LOCALIZATION | 336 | -0.496104 | -1.466217 | 0.004315 | 0.230022 |
| GOBP_NEGATIVE_REGULATION_OF_CELL_POPULATION_PROLIFERATION | 489 | -0.480977 | -1.465321 | 0.001038 | 0.231081 |
| GOBP_REGULATION_OF_PROTEIN_LOCALIZATION_TO_MEMBRANE | 154 | -0.521325 | -1.46499 | 0.01993 | 0.231156 |
| GOBP_DIACYLGLYCEROL_METABOLIC_PROCESS | 20 | -0.672109 | -1.464571 | 0.053712 | 0.231428 |
| GOBP_REGULATION_OF_ENDOCYTOSIS | 162 | -0.513868 | -1.463313 | 0.008383 | 0.2331 |
| GOBP_ENDOTHELIAL_CELL_CHEMOTAXIS | 19 | -0.69644 | -1.462459 | 0.059908 | 0.234109 |
| GOBP_RESPONSE_TO_LEUKEMIA_INHIBITORY_FACTOR | 65 | -0.564811 | -1.462353 | 0.039894 | 0.233811 |
| GOBP_POSITIVE_REGULATION_OF_RELEASE_OF_CYTOCHROME_C_FROM_MITC | 25 | -0.656083 | -1.462093 | 0.060372 | 0.233774 |
| GOBP_POSITIVE_REGULATION_OF_LIPID_METABOLIC_PROCESS | 100 | -0.541417 | -1.461757 | 0.030573 | 0.233862 |
| GOBP_REGULATION_OF_VESICLE_MEDIATED_TRANSPORT | 394 | -0.490326 | -1.461752 | 0.003202 | 0.233383 |
| GOBP_REGULATION_OF_VASCULAR_ASSOCIATED_SMOOTH_MUSCLE_CELL_PF | 39 | -0.613012 | -1.459992 | 0.044379 | 0.235684 |
| GOBP_REGULATION_OF_ERBB_SIGNALING_PATHWAY | 74 | -0.553013 | -1.459021 | 0.044619 | 0.23691 |
| GOBP_REGULATION_OF_BICELLULAR_TIGHT_JUNCTION_ASSEMBLY | 15 | -0.716322 | -1.458957 | 0.064516 | 0.236536 |
| GOBP_REGULATION_OF_PHOSPHOLIPID_METABOLIC_PROCESS | 28 | -0.640278 | -1.458933 | 0.06585 | 0.236102 |
| GOBP_LYMPHOCYTE_MEDIATED_IMMUNITY | 237 | -0.498489 | -1.457993 | 0.006764 | 0.237207 |
| GOBP_GLYCEROLIPID_CATABOLIC_PROCESS | 41 | -0.609466 | -1.457848 | 0.05132 | 0.236971 |
| GOBP_MODULATION_BY_VIRUS_OF_HOST_PROCESS | 28 | -0.65268 | -1.457772 | 0.058552 | 0.236642 |
| GOBP_ACTIVATION_OF_PHOSPHOLIPASE_C_ACTIVITY | 20 | -0.685167 | -1.457644 | 0.067524 | 0.236361 |
| GOBP_POSITIVE_REGULATION_OF_CD4_POSITIVE_ALPHA_BETA_T_CELL_DIFFE | 27 | -0.646839 | -1.457077 | 0.054217 | 0.236958 |
| GOBP_RESPONSE_TO_STEROL | 24 | -0.659133 | -1.454885 | 0.066236 | 0.240314 |
| GOBP_SUPEROXIDE_METABOLIC_PROCESS | 51 | -0.586493 | -1.453347 | 0.04507 | 0.242307 |
| GOBP_PROTEIN_O_LINKED_GLYCOSYLATION | 79 | -0.549176 | -1.452845 | 0.040107 | 0.242693 |
| GOBP_CELLULAR_RESPONSE_TO_TOXIC_SUBSTANCE | 85 | -0.545691 | -1.451962 | 0.039948 | 0.243613 |
| GOBP_NEGATIVE_REGULATION_OF_LIPID_LOCALIZATION | 30 | -0.638882 | -1.451765 | 0.057229 | 0.243488 |
| GOBP_B_CELL_ACTIVATION_INVOLVED_IN_IMMUNE_RESPONSE | 70 | -0.559762 | -1.450674 | 0.037889 | 0.244953 |
| GOBP_POSITIVE_REGULATION_OF_MYELOID_LEUKOCYTE_MEDIATED_IMMUNITY | 16 | -0.691374 | -1.450611 | 0.086379 | 0.244587 |
| GOBP_REGULATION_OF_KERATINOCYTE_DIFFERENTIATION | 29 | -0.629383 | -1.450072 | 0.066256 | 0.244945 |
| GOBP_CELLULAR_RESPONSE_TO_BIOTIC_STIMULUS | 175 | -0.503961 | -1.450066 | 0.01649 | 0.244467 |
| GOBP_REGULATORY_T_CELL_DIFFERENTIATION | 31 | -0.632429 | -1.449599 | 0.069016 | 0.244783 |
| GOBP_BLOOD_VESSEL_ENDOTHELIAL_CELL_MIGRATION | 88 | -0.552378 | -1.448884 | 0.030544 | 0.245636 |
| GOBP_CELL_PART_MORPHOGENESIS | 461 | -0.474636 | -1.448758 | 0 | 0.245365 |
| GOBP_REGULATION_OF_CATION_CHANNEL_ACTIVITY | 109 | -0.528099 | -1.448549 | 0.043534 | 0.245241 |
| GOBP_DENDRITIC_CELL_DIFFERENTIATION | 37 | -0.598279 | -1.447094 | 0.067976 | 0.247385 |
| GOBP_REGULATION_OF_OSTEOCLAST_DIFFERENTIATION | 47 | -0.588614 | -1.446397 | 0.056577 | 0.248092 |
| GOBP_POSITIVE_REGULATION_OF_ALPHA_BETA_T_CELL_DIFFERENTIATION | 40 | -0.601481 | -1.445973 | 0.061404 | 0.248363 |
| GOBP_CELLULAR_RESPONSE_TO_MOLECULE_OF_BACTERIAL_ORIGIN | 152 | -0.513416 | -1.445464 | 0.019048 | 0.248779 |
| GOBP_CLATHRIN_DEPENDENT_ENDOCYTOSIS | 41 | -0.59981 | -1.445463 | 0.068917 | 0.248294 |
| GOBP_POSITIVE_REGULATION_OF_SMALL_MOLECULE_METABOLIC_PROCESS | 92 | -0.547127 | -1.444686 | 0.03268 | 0.249232 |
| GOBP_REGULATION_OF_IMMUNE_EFFECTOR_PROCESS | 331 | -0.482672 | -1.444493 | 0.002148 | 0.249114 |
| GOBP_ALPHA_BETA_T_CELL_PROLIFERATION | 34 | -0.615105 | -1.444172 | 0.05772 | 0.24919 |
| GOBP_CELLULAR_RESPONSE_TO_CHEMICAL_STRESS | 287 | -0.486372 | -1.443762 | 0.006445 | 0.249464 |
| GOBP_NEGATIVE_REGULATION_OF_SMOOTH_MUSCLE_CELL_PROLIFERATION | 31 | -0.638218 | -1.443328 | 0.062691 | 0.249776 |
| GOBP_PROTEIN_POLYMERIZATION | 243 | -0.492811 | -1.443181 | 0.008959 | 0.249576 |
| GOBP_AMYLOID_FIBRIL_FORMATION | 22 | -0.669102 | -1.442918 | 0.061224 | 0.249558 |
| GOBP_SYNAPSE_ORGANIZATION | 255 | -0.488115 | -1.442787 | 0.008879 | 0.249315 |
| GOBP_REGULATION_OF_RAS_PROTEIN_SIGNAL_TRANSDUCTION | 158 | -0.511161 | -1.442647 | 0.019139 | 0.249057 |
| GOBP_WOUND_HEALING | 359 | -0.48474 | -1.442566 | 0.005336 | 0.248721 |
| GOBP_REGULATION_OF_OLIGODENDROCYTE_DIFFERENTIATION | 19 | -0.671109 | -1.441653 | 0.077176 | 0.249884 |
| GOBP_NLRP3_INFLAMMASOME_COMPLEX_ASSEMBLY | 15 | -0.718301 | -1.441326 | 0.063652 | 0.250059 |
| GOBP_NEGATIVE_REGULATION_OF_NUCLEOCYTOPLASMIC_TRANSPORT | 17 | -0.705067 | -1.439975 | 0.076175 | 0.251993 |
| GOBP_NEGATIVE_REGULATION_OF_T_CELL_PROLIFERATION | 54 | -0.580123 | -1.439729 | 0.055021 | 0.251986 |
| GOBP_REGULATION_OF_EMBRYONIC_DEVELOPMENT | 43 | -0.608712 | -1.4391 | 0.060294 | 0.252653 |
| GOBP_LYMPHOCYTE_MIGRATION | 76 | -0.556351 | -1.438493 | 0.058511 | 0.253344 |
| GOBP_ACTIVATION_OF_CYSTEINE_TYPE_ENDOPEPTIDASE_ACTIVITY_INVOLVED | 72 | -0.551251 | -1.438264 | 0.058824 | 0.25327 |
| GOBP_POSITIVE_REGULATION_OF_CELL_SUBSTRATE_JUNCTION_ORGANIZATIO | 23 | -0.646571 | -1.437587 | 0.066256 | 0.25397 |
| GOBP_REGULATION_OF_TRANSMEMBRANE_TRANSPORT | 357 | -0.480771 | -1.437484 | 0.001064 | 0.253681 |
| GOBP_REGULATION_OF_PH | 82 | -0.542321 | -1.437157 | 0.047493 | 0.253819 |
| GOBP_NEGATIVE_REGULATION_OF_LYMPHOCYTE_ACTIVATION | 123 | -0.517021 | -1.437003 | 0.018182 | 0.253582 |
| GOBP_REGULATION_OF_EPIDERMAL_GROWTH_FACTOR_ACTIVATED_RECEPTO | 21 | -0.660428 | -1.436863 | 0.085761 | 0.253323 |
| GOBP_NEGATIVE_REGULATION_OF_PROTEOLYSIS | 208 | -0.492502 | -1.436721 | 0.019563 | 0.253109 |
| GOBP_RESPONSE_TO_FUNGUS | 26 | -0.6379 | -1.436096 | 0.091324 | 0.253856 |
| GOBP_POSITIVE_REGULATION_OF_STRESS_FIBER_ASSEMBLY | 43 | -0.58586 | -1.435424 | 0.064516 | 0.254624 |
| GOBP_RESPONSE_TO_REACTIVE_OXYGEN_SPECIES | 178 | -0.504821 | -1.435149 | 0.019699 | 0.254636 |
| GOBP_AMINE_CATABOLIC_PROCESS | 19 | -0.664116 | -1.435106 | 0.087859 | 0.254239 |
| GOBP_NEGATIVE_REGULATION_OF_CHEMOTAXIS | 40 | -0.595938 | -1.434471 | 0.073913 | 0.254941 |
| GOBP_REGULATION_OF_RESPONSE_TO_REACTIVE_OXYGEN_SPECIES | 26 | -0.630432 | -1.432243 | 0.082043 | 0.258628 |
| GOBP_MODULATION_BY_VIRUS_OF_HOST_CELLULAR_PROCESS | 17 | -0.675153 | -1.431797 | 0.089577 | 0.258927 |
| GOBP_MODULATION_BY_SYMBIONT_OF_ENTRY_INTO_HOST | 37 | -0.603823 | -1.431584 | 0.071856 | 0.258897 |
| GOBP_LYMPH_NODE_DEVELOPMENT | 15 | -0.704339 | -1.431404 | 0.087163 | 0.258759 |
| GOBP_POSITIVE_REGULATION_OF_GLIAL_CELL_DIFFERENTIATION | 27 | -0.626681 | -1.430206 | 0.074516 | 0.260456 |
| GOBP_REGULATION_OF_SIGNALING_RECEPTOR_ACTIVITY | 105 | -0.526032 | -1.429855 | 0.035851 | 0.260662 |
| GOBP_TYROSINE_PHOSPHORYLATION_OF_STAT_PROTEIN | 53 | -0.568135 | -1.429685 | 0.066856 | 0.260455 |
| GOBP_CELL_MORPHOGENESIS_INVOLVED_IN_NEURON_DIFFERENTIATION | 390 | -0.476639 | -1.429492 | 0.003205 | 0.26034 |
| GOBP_REGULATION_OF_RESPONSE_TO_INTERFERON_GAMMA | 22 | -0.650422 | -1.428969 | 0.075235 | 0.260766 |
| GOBP_GLIOGENESIS | 198 | -0.497165 | -1.428779 | 0.013746 | 0.260602 |
| GOBP_CALCINEURIN_MEDIATED_SIGNALING | 35 | -0.614527 | -1.42868 | 0.078261 | 0.260326 |
| GOBP_CELL_DEATH_IN_RESPONSE_TO_OXIDATIVE_STRESS | 71 | -0.546631 | -1.428675 | 0.05548 | 0.259855 |
| GOBP_NEGATIVE_REGULATION_OF_EPITHELIAL_CELL_MIGRATION | 50 | -0.579345 | -1.42705 | 0.06768 | 0.262343 |
| GOBP_ORGANIC_ACID_TRANSMEMBRANE_TRANSPORT | 107 | -0.529393 | -1.426917 | 0.032746 | 0.262098 |
| GOBP_RESPONSE_TO_FLUID_SHEAR_STRESS | 25 | -0.639029 | -1.426857 | 0.071212 | 0.261747 |
| GOBP_POSITIVE_REGULATION_OF_GLIOGENESIS | 44 | -0.591747 | -1.426422 | 0.077033 | 0.262095 |
| GOBP_POSITIVE_REGULATION_OF_DENDRITIC_SPINE_DEVELOPMENT | 31 | -0.619213 | -1.425862 | 0.092846 | 0.26267 |
| GOBP_NEGATIVE_REGULATION_OF_CELL_SUBSTRATE_JUNCTION_ORGANIZATI( | 15 | -0.694931 | -1.424649 | 0.079867 | 0.264404 |
| GOBP_EXCRETION | 29 | -0.624109 | -1.424449 | 0.070571 | 0.264307 |
| GOBP_GLIAL_CELL_PROLIFERATION | 37 | -0.593905 | -1.423441 | 0.064194 | 0.265692 |
| GOBP_EPITHELIAL_CELL_PROLIFERATION | 252 | -0.487357 | -1.422807 | 0.010204 | 0.266346 |
| GOBP_POSITIVE_REGULATION_OF_TRANSLATIONAL_INITIATION | 24 | -0.650076 | -1.42246 | 0.076923 | 0.266445 |
| GOBP_REGULATION_OF_PHOSPHOLIPASE_ACTIVITY | 42 | -0.583028 | -1.422449 | 0.073864 | 0.265981 |
| GOBP_NEGATIVE_REGULATION_OF_INTERLEUKIN_1_BETA_PRODUCTION | 20 | -0.665632 | -1.422425 | 0.079032 | 0.265538 |
| GOBP_NEGATIVE_REGULATION_OF_HEMOPOIESIS | 77 | -0.541657 | -1.422216 | 0.045514 | 0.265474 |
| GOBP_NEGATIVE_REGULATION_OF_INTERLEUKIN_1_PRODUCTION | 24 | -0.646764 | -1.42172 | 0.067176 | 0.265913 |
| GOBP_L_AMINO_ACID_TRANSPORT | 44 | -0.583302 | -1.420139 | 0.063274 | 0.268377 |
| GOBP_GLIAL_CELL_DEVELOPMENT | 80 | -0.536801 | -1.41991 | 0.054902 | 0.268354 |
| GOBP_DIVALENT_INORGANIC_CATION_HOMEOSTASIS | 307 | -0.47534 | -1.418492 | 0.003297 | 0.270531 |
| GOBP_CELLULAR_HORMONE_METABOLIC_PROCESS | 71 | -0.553636 | -1.416976 | 0.055851 | 0.272924 |
| GOBP_REGULATION_OF_LIPASE_ACTIVITY | 58 | -0.56091 | -1.41687 | 0.06392 | 0.272635 |
| GOBP_RETINOL_METABOLIC_PROCESS | 27 | -0.639008 | -1.416517 | 0.09403 | 0.27285 |
| GOBP_REGULATION_OF_CELLULAR_COMPONENT_SIZE | 294 | -0.476691 | -1.415835 | 0.009934 | 0.273607 |
| GOBP_POSITIVE_REGULATION_OF_PROTEIN_LOCALIZATION_TO_MEMBRANE | 107 | -0.517886 | -1.415303 | 0.037736 | 0.274173 |
| GOBP_ACTION_POTENTIAL | 68 | -0.549133 | -1.415256 | 0.062834 | 0.273771 |
| GOBP_ANIMAL_ORGAN_REGENERATION | 57 | -0.557426 | -1.415197 | 0.069012 | 0.273411 |
| GOBP_ACYLGLYCEROL_HOMEOSTASIS | 17 | -0.677315 | -1.414834 | 0.101613 | 0.273573 |
| GOBP_POSITIVE_REGULATION_OF_VIRAL_PROCESS | 86 | -0.538028 | -1.414126 | 0.049544 | 0.274495 |
| GOBP_REGULATION_OF_TRANSCRIPTION_BY_RNA_POLYMERASE_III | 22 | -0.641743 | -1.413597 | 0.083591 | 0.275053 |
| GOBP_RESPONSE_TO_NUTRIENT | 111 | -0.517461 | -1.413383 | 0.041353 | 0.274958 |
| GOBP_POSITIVE_REGULATION_OF_ENDOCYTOSIS | 77 | -0.536409 | -1.412992 | 0.049519 | 0.275258 |
| GOBP_CELLULAR_ION_HOMEOSTASIS | 439 | -0.468757 | -1.412781 | 0.001054 | 0.27518 |
| GOBP_ORGANIC_HYDROXY_COMPOUND_CATABOLIC_PROCESS | 43 | -0.573656 | -1.4125 | 0.080357 | 0.275244 |
| GOBP_FAT_SOLUBLE_VITAMIN_METABOLIC_PROCESS | 18 | -0.678494 | -1.412289 | 0.112676 | 0.275182 |
| GOBP_REGULATION_OF_OXIDATIVE_STRESS_INDUCED_CELL_DEATH | 52 | -0.568398 | -1.411816 | 0.073826 | 0.275582 |
| GOBP_REGULATION_OF_RELEASE_OF_CYTOCHROME_C_FROM_MITOCHONDRI/ | 39 | -0.58837 | -1.411475 | 0.075444 | 0.27577 |
| GOBP_CENTRAL_NERVOUS_SYSTEM_NEURON_AXONOGENESIS | 21 | -0.660312 | -1.410663 | 0.083587 | 0.276771 |
| GOBP_MODULATION_BY_SYMBIONT_OF_HOST_PROCESS | 34 | -0.605866 | -1.409342 | 0.072593 | 0.278748 |

| | | | | | |
|---|---|---|---|---|---|
| GOBP_MITOCHONDRIAL_MEMBRANE_ORGANIZATION | 135 | 0.241731 | 0.831908 | 0.881988 | 1 |
| GOBP_TETRAPYRROLE_METABOLIC_PROCESS | 51 | 0.278381 | 0.831155 | 0.790441 | 1 |
| GOBP_PEPTIDYL_GLUTAMIC_ACID_MODIFICATION | 23 | 0.331817 | 0.830528 | 0.666667 | 1 |
| GOBP_ERYTHROCYTE_DEVELOPMENT | 27 | 0.328374 | 0.829747 | 0.684507 | 1 |
| GOBP_REGULATION_OF_NEUROBLAST_PROLIFERATION | 17 | 0.355177 | 0.829585 | 0.682692 | 1 |
| GOBP_INTRINSIC_APOPTOTIC_SIGNALING_PATHWAY_BY_P53_CLASS_MEDIATOR | 65 | 0.27168 | 0.828835 | 0.777778 | 1 |
| GOBP_ATTACHMENT_OF_SPINDLE_MICROTUBULES_TO_KINETOCHORE | 35 | 0.309945 | 0.827382 | 0.744548 | 1 |
| GOBP_BETA_CATENIN_TCF_COMPLEX_ASSEMBLY | 29 | 0.312057 | 0.82709 | 0.696875 | 1 |
| GOBP_CENTROMERE_COMPLEX_ASSEMBLY | 48 | 0.283219 | 0.827012 | 0.77193 | 1 |
| GOBP_REGULATION_OF_CAMP_DEPENDENT_PROTEIN_KINASE_ACTIVITY | 17 | 0.349056 | 0.826193 | 0.667546 | 1 |
| GOBP_REGULATION_OF_POSTSYNAPTIC_MEMBRANE_NEUROTRANSMITTER_RECEPT( | 42 | 0.285967 | 0.826084 | 0.747634 | 1 |
| GOBP_INCLUSION_BODY_ASSEMBLY | 20 | 0.333554 | 0.825117 | 0.691057 | 1 |
| GOBP_INTERMEMBRANE_LIPID_TRANSFER | 38 | 0.289313 | 0.824926 | 0.748485 | 1 |
| GOBP_TOXIN_TRANSPORT | 35 | 0.304961 | 0.824526 | 0.707042 | 1 |
| GOBP_CHROMATIN_REMODELING | 179 | 0.232915 | 0.823848 | 0.895105 | 1 |
| GOBP_PROTEIN_KINASE_A_SIGNALING | 23 | 0.326816 | 0.823451 | 0.714286 | 1 |
| GOBP_REGULATION_OF_CHROMATIN_ORGANIZATION | 162 | 0.234223 | 0.822869 | 0.85625 | 1 |
| GOBP_BRANCHED_CHAIN_AMINO_ACID_METABOLIC_PROCESS | 22 | 0.336775 | 0.82262 | 0.709402 | 1 |
| GOBP_POSITIVE_REGULATION_OF_BIOMINERALIZATION | 28 | 0.322561 | 0.820092 | 0.722388 | 1 |
| GOBP_COBALAMIN_METABOLIC_PROCESS | 17 | 0.342122 | 0.819298 | 0.672589 | 1 |
| GOBP_RNA_CAPPING | 34 | 0.301083 | 0.817878 | 0.737013 | 1 |
| GOBP_NUCLEOSOME_ASSEMBLY | 96 | 0.248042 | 0.816566 | 0.833333 | 1 |
| GOBP_POSITIVE_REGULATION_OF_DEFENSE_RESPONSE_TO_VIRUS_BY_HOST | 28 | 0.309603 | 0.814968 | 0.719764 | 1 |
| GOBP_REGULATION_OF_POLYSACCHARIDE_METABOLIC_PROCESS | 32 | 0.306068 | 0.814835 | 0.772472 | 1 |
| GOBP_REGULATION_OF_ALCOHOL_BIOSYNTHETIC_PROCESS | 56 | 0.274939 | 0.811286 | 0.797753 | 1 |
| GOBP_POSITIVE_REGULATION_OF_CELL_CYCLE_PHASE_TRANSITION | 84 | 0.255436 | 0.807268 | 0.883117 | 1 |
| GOBP_FATTY_ACID_BETA_OXIDATION | 67 | 0.264218 | 0.807256 | 0.836431 | 1 |
| GOBP_UBIQUITIN_DEPENDENT_PROTEIN_CATABOLIC_PROCESS_VIA_THE_MULTIVESI( | 22 | 0.323834 | 0.806343 | 0.707246 | 1 |
| GOBP_POST_GOLGI_VESICLE_MEDIATED_TRANSPORT | 98 | 0.245292 | 0.805521 | 0.868182 | 1 |
| GOBP_RIBOSOME_ASSEMBLY | 62 | 0.265004 | 0.803398 | 0.81203 | 1 |
| GOBP_REGULATION_OF_PROGRAMMED_NECROTIC_CELL_DEATH | 20 | 0.331665 | 0.802909 | 0.733146 | 1 |
| GOBP_IRON_SULFUR_CLUSTER_ASSEMBLY | 24 | 0.309637 | 0.800568 | 0.719136 | 1 |
| GOBP_HOMOLOGOUS_CHROMOSOME_PAIRING_AT_MEIOSIS | 35 | 0.292056 | 0.800319 | 0.769886 | 1 |
| GOBP_SYNAPTIC_VESICLE_CYTOSKELETAL_TRANSPORT | 18 | 0.327899 | 0.799442 | 0.726582 | 1 |
| GOBP_POSITIVE_REGULATION_OF_ATPASE_ACTIVITY | 52 | 0.273965 | 0.799328 | 0.836576 | 1 |
| GOBP_NEUROMUSCULAR_PROCESS_CONTROLLING_BALANCE | 24 | 0.314147 | 0.799309 | 0.751462 | 1 |
| GOBP_TRANSLATIONAL_ELONGATION | 129 | 0.24045 | 0.797493 | 0.909091 | 1 |
| GOBP_NUCLEOSOME_ORGANIZATION | 129 | 0.234232 | 0.795916 | 0.923858 | 1 |
| GOBP_VIRION_ASSEMBLY | 39 | 0.292605 | 0.793907 | 0.74375 | 1 |
| GOBP_SISTER_CHROMATID_COHESION | 58 | 0.265324 | 0.793056 | 0.820513 | 1 |
| GOBP_POSITIVE_REGULATION_OF_PROTEIN_MODIFICATION_BY_SMALL_PROTEIN_CO | 125 | 0.234984 | 0.792718 | 0.921053 | 1 |
| GOBP_ATTACHMENT_OF_MITOTIC_SPINDLE_MICROTUBULES_TO_KINETOCHORE | 15 | 0.343855 | 0.792458 | 0.74092 | 1 |
| GOBP_TRANSLATIONAL_INITIATION | 186 | 0.221908 | 0.791352 | 0.963504 | 1 |
| GOBP_TRNA_PROCESSING | 127 | 0.235144 | 0.790657 | 0.933333 | 1 |
| GOBP_CELLULAR_COMPONENT_DISASSEMBLY_INVOLVED_IN_EXECUTION_PHASE_OI | 28 | 0.305714 | 0.789841 | 0.771084 | 1 |
| GOBP_PHOTOTRANSDUCTION_VISIBLE_LIGHT | 20 | 0.318947 | 0.788233 | 0.744 | 1 |
| GOBP_RHYTHMIC_BEHAVIOR | 26 | 0.309145 | 0.787366 | 0.746439 | 1 |
| GOBP_RRNA_METABOLIC_PROCESS | 234 | 0.216723 | 0.787351 | 0.992126 | 1 |
| GOBP_CHROMATIN_REMODELING_AT_CENTROMERE | 40 | 0.279311 | 0.787308 | 0.780645 | 1 |
| GOBP_SISTER_CHROMATID_SEGREGATION | 188 | 0.221521 | 0.786454 | 0.962687 | 1 |
| GOBP_MRNA_METHYLATION | 16 | 0.333646 | 0.784007 | 0.713555 | 1 |
| GOBP_MITOCHONDRIAL_FUSION | 27 | 0.302767 | 0.782602 | 0.764012 | 1 |
| GOBP_TRNA_MODIFICATION | 87 | 0.240316 | 0.779742 | 0.905172 | 1 |
| GOBP_GLUTAMINE_METABOLIC_PROCESS | 20 | 0.314228 | 0.779408 | 0.772472 | 1 |
| GOBP_REGULATION_OF_CELLULAR_EXTRAVASATION | 25 | 0.304767 | 0.776458 | 0.773743 | 1 |
| GOBP_POSITIVE_REGULATION_OF_CELL_CYCLE_G1_S_PHASE_TRANSITION | 38 | 0.282215 | 0.775549 | 0.831776 | 1 |
| GOBP_SCHWANN_CELL_DIFFERENTIATION | 31 | 0.292065 | 0.775045 | 0.804154 | 1 |
| GOBP_LATE_ENDOSOME_TO_VACUOLE_TRANSPORT_VIA_MULTIVESICULAR_BODY_S( | 17 | 0.327021 | 0.77451 | 0.746231 | 1 |
| GOBP_REGULATION_OF_RUFFLE_ASSEMBLY | 22 | 0.315558 | 0.774459 | 0.775726 | 1 |
| GOBP_NUCLEAR_CHROMOSOME_SEGREGATION | 246 | 0.211204 | 0.77393 | 1 | 1 |
| GOBP_RESPONSE_TO_GLUCAGON | 24 | 0.30192 | 0.767973 | 0.777778 | 1 |
| GOBP_REGULATION_OF_MRNA_POLYADENYLATION | 17 | 0.324869 | 0.766848 | 0.75 | 1 |
| GOBP_INTRACELLULAR_ESTROGEN_RECEPTOR_SIGNALING_PATHWAY | 42 | 0.274269 | 0.766586 | 0.848684 | 1 |
| GOBP_POSITIVE_REGULATION_OF_MITOCHONDRIAL_OUTER_MEMBRANE_PERMEABIL | 35 | 0.272826 | 0.764886 | 0.824074 | 1 |
| GOBP_LATE_ENDOSOME_TO_VACUOLE_TRANSPORT | 24 | 0.295512 | 0.762296 | 0.786127 | 1 |
| GOBP_PROTEIN_TARGETING_TO_LYSOSOME | 22 | 0.301752 | 0.761671 | 0.811111 | 1 |
| GOBP_ASPARTATE_FAMILY_AMINO_ACID_METABOLIC_PROCESS | 40 | 0.26765 | 0.760254 | 0.832808 | 1 |
| GOBP_MITOTIC_SISTER_CHROMATID_SEGREGATION | 157 | 0.221029 | 0.759416 | 0.973154 | 1 |
| GOBP_N_TERMINAL_PROTEIN_AMINO_ACID_MODIFICATION | 26 | 0.2934 | 0.75916 | 0.795918 | 1 |
| GOBP_REGULATION_OF_CHROMOSOME_SEGREGATION | 81 | 0.244677 | 0.758378 | 0.911647 | 1 |
| GOBP_NUCLEOSIDE_MONOPHOSPHATE_BIOSYNTHETIC_PROCESS | 38 | 0.275388 | 0.756091 | 0.837662 | 1 |
| GOBP_CHROMOSOME_SEGREGATION | 305 | 0.207578 | 0.75172 | 1 | 1 |
| GOBP_REGULATION_OF_PROTEIN_ACETYLATION | 64 | 0.248222 | 0.751102 | 0.904215 | 1 |
| GOBP_MULTIVESICULAR_BODY_ORGANIZATION | 30 | 0.283166 | 0.748053 | 0.831884 | 1 |
| GOBP_RNA_DEPENDENT_DNA_BIOSYNTHETIC_PROCESS | 68 | 0.243754 | 0.747892 | 0.887597 | 1 |
| GOBP_REGULATION_OF_MITOTIC_SPINDLE_ASSEMBLY | 18 | 0.316192 | 0.747862 | 0.795213 | 1 |
| GOBP_REGULATION_OF_CYTOKINESIS | 71 | 0.239536 | 0.744406 | 0.910569 | 1 |
| GOBP_CHROMOSOME_LOCALIZATION | 74 | 0.238769 | 0.742323 | 0.933333 | 1 |
| GOBP_VIRAL_GENE_EXPRESSION | 194 | 0.209894 | 0.74176 | 0.986577 | 1 |
| GOBP_N_TERMINAL_PROTEIN_AMINO_ACID_ACETYLATION | 16 | 0.319931 | 0.740383 | 0.817232 | 1 |
| GOBP_HOMOLOGOUS_CHROMOSOME_SEGREGATION | 43 | 0.265 | 0.738291 | 0.865979 | 1 |
| GOBP_HEME_METABOLIC_PROCESS | 30 | 0.281921 | 0.736919 | 0.848315 | 1 |
| GOBP_REGULATION_OF_MEMBRANE_PERMEABILITY | 73 | 0.236621 | 0.736345 | 0.939759 | 1 |
| GOBP_MEIOTIC_CHROMOSOME_SEGREGATION | 69 | 0.239036 | 0.736261 | 0.931559 | 1 |
| GOBP_REGULATION_OF_ACTIVIN_RECEPTOR_SIGNALING_PATHWAY | 16 | 0.321264 | 0.734524 | 0.775068 | 1 |
| GOBP_STRIATED_MUSCLE_CELL_PROLIFERATION | 37 | 0.264534 | 0.733717 | 0.888563 | 1 |
| GOBP_PROTEIN_N_LINKED_GLYCOSYLATION | 66 | 0.242269 | 0.733373 | 0.92674 | 1 |
| GOBP_PYRIMIDINE_NUCLEOTIDE_METABOLIC_PROCESS | 44 | 0.256842 | 0.73211 | 0.914191 | 1 |
| GOBP_POTASSIUM_ION_IMPORT_ACROSS_PLASMA_MEMBRANE | 24 | 0.288105 | 0.731831 | 0.828125 | 1 |
| GOBP_NCRNA_PROCESSING | 378 | 0.191371 | 0.731428 | 1 | 1 |
| GOBP_TRANSFERRIN_TRANSPORT | 31 | 0.272265 | 0.730565 | 0.854786 | 1 |
| GOBP_HISTONE_EXCHANGE | 47 | 0.258076 | 0.729747 | 0.895683 | 1 |
| GOBP_REGULATION_OF_SISTER_CHROMATID_COHESION | 22 | 0.288354 | 0.728466 | 0.826087 | 1 |
| GOBP_ALPHA_AMINO_ACID_CATABOLIC_PROCESS | 64 | 0.233624 | 0.728161 | 0.919847 | 1 |
| GOBP_DNA_DOUBLE_STRAND_BREAK_PROCESSING | 21 | 0.297559 | 0.7278 | 0.8125 | 1 |
| GOBP_REGULATION_OF_POLYSACCHARIDE_BIOSYNTHETIC_PROCESS | 27 | 0.282648 | 0.727134 | 0.863777 | 1 |
| GOBP_MRNA_MODIFICATION | 24 | 0.2891 | 0.724791 | 0.850877 | 1 |
| GOBP_REGULATION_OF_SPINDLE_ASSEMBLY | 25 | 0.284455 | 0.724613 | 0.854651 | 1 |
| GOBP_LAMELLIPODIUM_ASSEMBLY | 60 | 0.242574 | 0.723458 | 0.909774 | 1 |
| GOBP_MITOTIC_METAPHASE_PLATE_CONGRESSION | 50 | 0.249625 | 0.722531 | 0.924915 | 1 |
| GOBP_CLATHRIN_COAT_ASSEMBLY | 15 | 0.312047 | 0.720811 | 0.802057 | 1 |
| GOBP_REGULATION_OF_MITOCHONDRIAL_OUTER_MEMBRANE_PERMEABILIZATION_IN | 42 | 0.251298 | 0.720363 | 0.893238 | 1 |
| GOBP_PHOSPHATIDYLINOSITOL_PHOSPHORYLATION | 53 | 0.247214 | 0.718016 | 0.906574 | 1 |
| GOBP_CYTOKINESIS | 143 | 0.207853 | 0.717704 | 1 | 1 |
| GOBP_CELLULAR_RESPONSE_TO_ALKALOID | 19 | 0.298408 | 0.717487 | 0.797872 | 1 |
| GOBP_HISTONE_H4_K5_ACETYLATION | 16 | 0.309839 | 0.717307 | 0.802703 | 1 |
| GOBP_DNA_LIGATION | 15 | 0.311104 | 0.711099 | 0.819121 | 1 |
| GOBP_PORPHYRIN_CONTAINING_COMPOUND_METABOLIC_PROCESS | 34 | 0.266841 | 0.710885 | 0.876161 | 1 |
| GOBP_PHOSPHOLIPID_DEPHOSPHORYLATION | 38 | 0.253018 | 0.710303 | 0.902736 | 1 |
| GOBP_REGULATION_OF_CHROMOSOME_SEPARATION | 67 | 0.226009 | 0.707637 | 0.959108 | 1 |
| GOBP_MITOTIC_SPINDLE_ASSEMBLY | 63 | 0.230084 | 0.707469 | 0.957854 | 1 |
| GOBP_RESPONSE_TO_FOOD | 19 | 0.292727 | 0.705659 | 0.837912 | 1 |
| GOBP_INTRA_GOLGI_VESICLE_MEDIATED_TRANSPORT | 31 | 0.264656 | 0.705635 | 0.894587 | 1 |
| GOBP_POSITIVE_REGULATION_OF_PROTEIN_LOCALIZATION_TO_NUCLEUS | 72 | 0.223618 | 0.702226 | 0.96371 | 1 |
| GOBP_METAPHASE_PLATE_CONGRESSION | 63 | 0.229143 | 0.702213 | 0.944223 | 1 |
| GOBP_RNA_METHYLATION | 82 | 0.220231 | 0.70204 | 0.973684 | 1 |
| GOBP_PROTEIN_TRANSMEMBRANE_IMPORT_INTO_INTRACELLULAR_ORGANELLE | 36 | 0.257672 | 0.701776 | 0.913044 | 1 |
| GOBP_MITOCHONDRIAL_TRANSMEMBRANE_TRANSPORT | 101 | 0.215462 | 0.701675 | 0.985507 | 1 |
| GOBP_NEGATIVE_REGULATION_OF_FATTY_ACID_METABOLIC_PROCESS | 21 | 0.28333 | 0.699183 | 0.866469 | 1 |
| GOBP_IRE1_MEDIATED_UNFOLDED_PROTEIN_RESPONSE | 63 | 0.2245 | 0.697308 | 0.908425 | 1 |
| GOBP_RRNA_TRANSCRIPTION | 32 | 0.249095 | 0.69704 | 0.907652 | 1 |
| GOBP_ATP_SYNTHESIS_COUPLED_ELECTRON_TRANSPORT | 96 | 0.217179 | 0.696751 | 0.970443 | 1 |
| GOBP_SCHWANN_CELL_DEVELOPMENT | 23 | 0.282248 | 0.696132 | 0.878187 | 1 |
| GOBP_RESPIRATORY_ELECTRON_TRANSPORT_CHAIN | 112 | 0.209444 | 0.69571 | 0.981818 | 1 |
| GOBP_TELOMERE_MAINTENANCE_VIA_TELOMERE_LENGTHENING | 75 | 0.220344 | 0.695376 | 0.97318 | 1 |
| GOBP_TRNA_TRANSPORT | 37 | 0.250926 | 0.694949 | 0.913313 | 1 |
| GOBP_HOMOLOGOUS_RECOMBINATION | 51 | 0.235663 | 0.693472 | 0.950166 | 1 |
| GOBP_POSITIVE_REGULATION_OF_INTERFERON_BETA_PRODUCTION | 32 | 0.256901 | 0.69293 | 0.896552 | 1 |
| GOBP_TELOMERE_MAINTENANCE_VIA_RECOMBINATION | 15 | 0.305879 | 0.689206 | 0.831633 | 1 |
| GOBP_REGULATION_OF_MACROAUTOPHAGY | 149 | 0.19394 | 0.688884 | 0.99375 | 1 |
| GOBP_POSITIVE_REGULATION_OF_COLLAGEN_METABOLIC_PROCESS | 19 | 0.28147 | 0.688592 | 0.853786 | 1 |
| GOBP_NUCLEOTIDE_EXCISION_REPAIR_PREINCISION_COMPLEX_STABILIZATION | 21 | 0.284365 | 0.687043 | 0.88172 | 1 |
| GOBP_REGULATION_OF_RNA_SPLICING | 128 | 0.201947 | 0.685964 | 0.994737 | 1 |
| GOBP_G0_TO_G1_TRANSITION | 42 | 0.239992 | 0.682317 | 0.921502 | 1 |
| GOBP_PROTEIN_LOCALIZATION_TO_CYTOSKELETON | 57 | 0.224468 | 0.681987 | 0.944649 | 1 |
| GOBP_REGULATION_OF_MITOCHONDRIAL_MEMBRANE_POTENTIAL | 64 | 0.221387 | 0.680793 | 0.95302 | 1 |
| GOBP_PROTEIN_COMPLEX_OLIGOMERIZATION | 172 | 0.194499 | 0.680443 | 1 | 1 |
| GOBP_PROTEIN_DEMANNOSYLATION | 20 | 0.281271 | 0.679211 | 0.858696 | 1 |
| GOBP_ESTABLISHMENT_OF_PROTEIN_LOCALIZATION_TO_PLASMA_MEMBRANE | 53 | 0.226651 | 0.679181 | 0.962963 | 1 |
| GOBP_DNA_PACKAGING | 177 | 0.192047 | 0.676529 | 1 | 1 |
| GOBP_REGULATION_OF_ENDOPLASMIC_RETICULUM_STRESS_INDUCED_INTRINSIC_A | 27 | 0.25869 | 0.673822 | 0.919643 | 1 |
| GOBP_REGULATION_OF_DNA_TEMPLATED_TRANSCRIPTION_IN_RESPONSE_TO_STRE | 105 | 0.208571 | 0.673442 | 0.990338 | 1 |
| GOBP_ENDOSOME_TRANSPORT_VIA_MULTIVESICULAR_BODY_SORTING_PATHWAY | 30 | 0.257 | 0.673337 | 0.936937 | 1 |
| GOBP_CERAMIDE_CATABOLIC_PROCESS | 17 | 0.279982 | 0.671909 | 0.857881 | 1 |
| GOBP_PROTEIN_TARGETING_TO_VACUOLE | 38 | 0.232096 | 0.670946 | 0.950725 | 1 |
| GOBP_POTASSIUM_ION_HOMEOSTASIS | 15 | 0.295763 | 0.669012 | 0.871134 | 1 |
| GOBP_RESPIRATORY_CHAIN_COMPLEX_IV_ASSEMBLY | 26 | 0.256786 | 0.668014 | 0.911765 | 1 |

| | | | | | |
|---|---|---|---|---|---|
| GOBP_NEGATIVE_REGULATION_OF_CELL_ACTIVATION | 152 | -0.498088 | -1.408327 | 0.028986 | 0.280309 |
| GOBP_REGULATION_OF_INTRINSIC_APOPTOTIC_SIGNALING_PATHWAY | 139 | -0.502338 | -1.407308 | 0.033835 | 0.281817 |
| GOBP_NEGATIVE_REGULATION_OF_CELL_CELL_ADHESION | 141 | -0.504742 | -1.406634 | 0.035971 | 0.2826 |
| GOBP_MACROPHAGE_ACTIVATION | 68 | -0.54842 | -1.406509 | 0.048452 | 0.282344 |
| GOBP_PURINE_NUCLEOTIDE_TRANSPORT | 22 | -0.6423 | -1.406397 | 0.099844 | 0.282109 |
| GOBP_REGULATION_OF_POSITIVE_CHEMOTAXIS | 16 | -0.679683 | -1.405748 | 0.10467 | 0.282903 |
| GOBP_REGULATION_OF_TRANSLATION_NCRNA_MEDIATED | 21 | -0.633303 | -1.405512 | 0.107425 | 0.282858 |
| GOBP_REGULATION_OF_PRODUCTION_OF_MOLECULAR_MEDIATOR_OF_IMMUN | 118 | -0.507928 | -1.405474 | 0.04305 | 0.282436 |
| GOBP_AMINOGLYCAN_METABOLIC_PROCESS | 118 | -0.513838 | -1.405426 | 0.035309 | 0.282036 |
| GOBP_NEGATIVE_REGULATION_OF_AMYLOID_PRECURSOR_PROTEIN_CATABOL | 17 | -0.672324 | -1.405362 | 0.10989 | 0.281681 |
| GOBP_HEAD_DEVELOPMENT | 484 | -0.459313 | -1.404745 | 0.004175 | 0.282424 |
| GOBP_REGULATION_OF_CATION_TRANSMEMBRANE_TRANSPORT | 218 | -0.481849 | -1.404188 | 0.02265 | 0.282991 |
| GOBP_REGULATION_OF_SMOOTHENED_SIGNALING_PATHWAY | 57 | -0.558654 | -1.403892 | 0.081492 | 0.283078 |
| GOBP_POSITIVE_REGULATION_OF_CELLULAR_RESPONSE_TO_TRANSFORMING | 24 | -0.641566 | -1.403532 | 0.085758 | 0.28335 |
| GOBP_REGULATION_OF_MAMMARY_GLAND_EPITHELIAL_CELL_PROLIFERATION | 15 | -0.685088 | -1.402604 | 0.117647 | 0.284667 |
| GOBP_POSITIVE_REGULATION_OF_LIPID_BIOSYNTHETIC_PROCESS | 58 | -0.55159 | -1.402329 | 0.07367 | 0.284749 |
| GOBP_LOCALIZATION_WITHIN_MEMBRANE | 57 | -0.560805 | -1.402135 | 0.072289 | 0.28463 |
| GOBP_REACTIVE_OXYGEN_SPECIES_METABOLIC_PROCESS | 203 | -0.486021 | -1.402059 | 0.024249 | 0.284279 |
| GOBP_NEGATIVE_REGULATION_OF_SMOOTHENED_SIGNALING_PATHWAY | 25 | -0.635081 | -1.401905 | 0.1 | 0.284074 |
| GOBP_REGULATION_OF_CD4_POSITIVE_ALPHA_BETA_T_CELL_ACTIVATION | 55 | -0.564549 | -1.401756 | 0.065753 | 0.283903 |
| GOBP_POSITIVE_REGULATION_OF_CELLULAR_COMPONENT_BIOGENESIS | 413 | -0.463583 | -1.399967 | 0.007345 | 0.286894 |
| GOBP_ANDROGEN_RECEPTOR_SIGNALING_PATHWAY | 32 | -0.614294 | -1.399459 | 0.084615 | 0.287386 |
| GOBP_POSITIVE_REGULATION_OF_PEPTIDE_SECRETION | 104 | -0.511114 | -1.399441 | 0.037037 | 0.286938 |
| GOBP_EMBRYO_IMPLANTATION | 31 | -0.598334 | -1.398359 | 0.098802 | 0.288584 |
| GOBP_POSITIVE_REGULATION_OF_AXON_EXTENSION | 32 | -0.598465 | -1.398004 | 0.068452 | 0.288764 |
| GOBP_POSITIVE_REGULATION_OF_BINDING | 138 | -0.497026 | -1.397232 | 0.040097 | 0.289748 |
| GOBP_CELLULAR_OXIDANT_DETOXIFICATION | 71 | -0.539776 | -1.397124 | 0.054201 | 0.289483 |
| GOBP_MYELOID_DENDRITIC_CELL_ACTIVATION | 22 | -0.634754 | -1.39678 | 0.100787 | 0.289701 |
| GOBP_CELLULAR_RESPONSE_TO_AMYLOID_BETA | 28 | -0.622543 | -1.395043 | 0.097859 | 0.292485 |
| GOBP_REGULATION_OF_CLATHRIN_DEPENDENT_ENDOCYTOSIS | 16 | -0.679661 | -1.395016 | 0.104037 | 0.292053 |
| GOBP_SPROUTING_ANGIOGENESIS | 79 | -0.530955 | -1.393688 | 0.065246 | 0.294206 |
| GOBP_ODONTOGENESIS | 62 | -0.541093 | -1.393635 | 0.06291 | 0.293831 |
| GOBP_MUSCLE_CELL_APOPTOTIC_PROCESS | 51 | -0.562345 | -1.393405 | 0.09484 | 0.293806 |
| GOBP_MODULATION_OF_PROCESS_OF_OTHER_ORGANISM | 94 | -0.524336 | -1.392198 | 0.053985 | 0.295717 |
| GOBP_INTESTINAL_ABSORPTION | 23 | -0.644044 | -1.390036 | 0.112164 | 0.299466 |
| GOBP_PRODUCTION_OF_MOLECULAR_MEDIATOR_INVOLVED_IN_INFLAMMATOF | 60 | -0.54747 | -1.389086 | 0.070201 | 0.300824 |
| GOBP_POSITIVE_REGULATION_OF_PROTEOLYSIS | 298 | -0.468477 | -1.388917 | 0.005482 | 0.300702 |
| GOBP_ER_NUCLEUS_SIGNALING_PATHWAY | 49 | -0.560612 | -1.388739 | 0.082386 | 0.300569 |
| GOBP_SMALL_GTPASE_MEDIATED_SIGNAL_TRANSDUCTION | 430 | -0.454554 | -1.388369 | 0.006303 | 0.300803 |
| GOBP_POSITIVE_REGULATION_OF_CELL_JUNCTION_ASSEMBLY | 47 | -0.564157 | -1.388141 | 0.078635 | 0.300732 |
| GOBP_PH_REDUCTION | 47 | -0.566294 | -1.388075 | 0.079208 | 0.30039 |
| GOBP_ACTIVATION_OF_INNATE_IMMUNE_RESPONSE | 124 | -0.504051 | -1.387681 | 0.041565 | 0.300675 |
| GOBP_POSITIVE_REGULATION_OF_PROTEIN_KINASE_B_SIGNALING | 111 | -0.507791 | -1.387203 | 0.041131 | 0.30113 |
| GOBP_ADAPTIVE_IMMUNE_RESPONSE_BASED_ON_SOMATIC_RECOMBINATION_ | 247 | -0.472726 | -1.38705 | 0.011236 | 0.30095 |
| GOBP_NEURON_DEATH_IN_RESPONSE_TO_OXIDATIVE_STRESS | 25 | -0.625224 | -1.386754 | 0.108896 | 0.301031 |
| GOBP_HYPEROSMOTIC_RESPONSE | 18 | -0.653487 | -1.386494 | 0.095935 | 0.30108 |
| GOBP_REGULATION_OF_T_HELPER_CELL_DIFFERENTIATION | 31 | -0.593866 | -1.385628 | 0.105505 | 0.302333 |
| GOBP_ADENYLATE_CYCLASE_ACTIVATING_G_PROTEIN_COUPLED_RECEPTOR_ | 61 | -0.535831 | -1.383924 | 0.084388 | 0.30515 |
| GOBP_CIRCULATORY_SYSTEM_PROCESS | 349 | -0.461456 | -1.383345 | 0.013072 | 0.305763 |
| GOBP_BASEMENT_MEMBRANE_ORGANIZATION | 20 | -0.65297 | -1.38322 | 0.105263 | 0.305524 |
| GOBP_HETEROPHILIC_CELL_CELL_ADHESION_VIA_PLASMA_MEMBRANE_CELL_ | 22 | -0.635498 | -1.382548 | 0.103339 | 0.306445 |
| GOBP_BIOLOGICAL_PROCESS_INVOLVED_IN_INTRASPECIES_INTERACTION_BE | 31 | -0.601799 | -1.381501 | 0.103659 | 0.30806 |
| GOBP_POSITIVE_REGULATION_OF_NF_KAPPAB_TRANSCRIPTION_FACTOR_ACTI | 139 | -0.497469 | -1.381491 | 0.042118 | 0.307586 |
| GOBP_ANTIGEN_PROCESSING_AND_PRESENTATION_OF_PEPTIDE_ANTIGEN | 178 | -0.478545 | -1.381301 | 0.030769 | 0.307487 |
| GOBP_REGULATION_OF_MEMBRANE_PROTEIN_ECTODOMAIN_PROTEOLYSIS | 18 | -0.651077 | -1.381217 | 0.116203 | 0.307199 |
| GOBP_NEGATIVE_REGULATION_OF_PROTEIN_LOCALIZATION_TO_MEMBRANE | 21 | -0.640958 | -1.381034 | 0.118565 | 0.307149 |
| GOBP_MONOVALENT_INORGANIC_CATION_HOMEOSTASIS | 109 | -0.508628 | -1.381002 | 0.0475 | 0.306739 |
| GOBP_REGULATION_OF_OXIDATIVE_STRESS_INDUCED_NEURON_DEATH | 22 | -0.644054 | -1.380008 | 0.109501 | 0.308148 |
| GOBP_NEGATIVE_REGULATION_OF_FIBROBLAST_PROLIFERATION | 25 | -0.611173 | -1.379154 | 0.111801 | 0.309467 |
| GOBP_IN_UTERO_EMBRYONIC_DEVELOPMENT | 260 | -0.469037 | -1.378427 | 0.016873 | 0.310517 |
| GOBP_NEGATIVE_REGULATION_OF_CELL_CYCLE_G1_S_PHASE_TRANSITION | 95 | -0.512872 | -1.377931 | 0.047865 | 0.311077 |
| GOBP_ISOPRENOID_METABOLIC_PROCESS | 79 | -0.517387 | -1.377468 | 0.071979 | 0.311449 |
| GOBP_INTEGRIN_MEDIATED_SIGNALING_PATHWAY | 85 | -0.519803 | -1.377132 | 0.063492 | 0.311618 |
| GOBP_ACTIN_FILAMENT_DEPOLYMERIZATION | 46 | -0.560073 | -1.376603 | 0.082305 | 0.312207 |
| GOBP_NEGATIVE_REGULATION_OF_MUSCLE_CELL_APOPTOTIC_PROCESS | 26 | -0.623188 | -1.376476 | 0.115265 | 0.31202 |
| GOBP_REGULATION_OF_STEROL_TRANSPORT | 37 | -0.583905 | -1.37622 | 0.088445 | 0.312029 |
| GOBP_TRIGLYCERIDE_BIOSYNTHETIC_PROCESS | 31 | -0.596622 | -1.37494 | 0.107143 | 0.314174 |
| GOBP_REGULATION_OF_RECEPTOR_MEDIATED_ENDOCYTOSIS | 74 | -0.525654 | -1.374816 | 0.058902 | 0.313934 |
| GOBP_REGULATION_OF_T_CELL_DIFFERENTIATION_IN_THYMUS | 16 | -0.670337 | -1.374553 | 0.131988 | 0.313958 |
| GOBP_REGULATION_OF_RECEPTOR_RECYCLING | 16 | -0.670507 | -1.374463 | 0.1232 | 0.313647 |
| GOBP_POSITIVE_REGULATION_OF_VASCULAR_ASSOCIATED_SMOOTH_MUSCLE | 25 | -0.616001 | -1.373913 | 0.115152 | 0.314284 |
| GOBP_NEGATIVE_REGULATION_OF_APOPTOTIC_SIGNALING_PATHWAY | 177 | -0.482283 | -1.373552 | 0.035294 | 0.314601 |
| GOBP_POSITIVE_REGULATION_OF_PROTEIN_TARGETING_TO_MEMBRANE | 29 | -0.609534 | -1.373206 | 0.115097 | 0.314905 |
| GOBP_NEGATIVE_REGULATION_OF_ACTIN_FILAMENT_DEPOLYMERIZATION | 34 | -0.578658 | -1.37264 | 0.118571 | 0.315581 |
| GOBP_SALIVARY_GLAND_DEVELOPMENT | 18 | -0.645603 | -1.372423 | 0.131707 | 0.315534 |
| GOBP_NONRIBOSOMAL_PEPTIDE_BIOSYNTHETIC_PROCESS | 16 | -0.661209 | -1.371774 | 0.122977 | 0.316363 |
| GOBP_MAINTENANCE_OF_LOCATION_IN_CELL | 152 | -0.48084 | -1.371487 | 0.042755 | 0.316454 |
| GOBP_POSITIVE_REGULATION_OF_AMYLOID_BETA_FORMATION | 15 | -0.669358 | -1.371213 | 0.101942 | 0.316509 |
| GOBP_DETECTION_OF_EXTERNAL_BIOTIC_STIMULUS | 19 | -0.642611 | -1.369795 | 0.116585 | 0.31887 |
| GOBP_CELL_ADHESION_MEDIATED_BY_INTEGRIN | 53 | -0.553465 | -1.369033 | 0.08642 | 0.320023 |
| GOBP_POSITIVE_REGULATION_OF_NIK_NF_KAPPAB_SIGNALING | 55 | -0.538332 | -1.369007 | 0.086301 | 0.319603 |
| GOBP_REGULATION_OF_BINDING | 284 | -0.46091 | -1.368365 | 0.014493 | 0.320414 |
| GOBP_RESPONSE_TO_ALCOHOL | 152 | -0.481752 | -1.368025 | 0.040984 | 0.320633 |
| GOBP_ARACHIDONIC_ACID_METABOLIC_PROCESS | 32 | -0.601735 | -1.367726 | 0.113293 | 0.320777 |
| GOBP_POSITIVE_REGULATION_OF_CELL_DEATH | 495 | -0.448217 | -1.36743 | 0.004149 | 0.320922 |
| GOBP_POSITIVE_REGULATION_OF_FOCAL_ADHESION_ASSEMBLY | 18 | -0.656684 | -1.366431 | 0.1008 | 0.322432 |
| GOBP_REGULATION_OF_NEUTROPHIL_CHEMOTAXIS | 27 | -0.59923 | -1.366062 | 0.113354 | 0.322691 |
| GOBP_EPIDERMAL_CELL_DIFFERENTIATION | 121 | -0.493184 | -1.365723 | 0.046482 | 0.322875 |
| GOBP_RESPONSE_TO_HYDROPEROXIDE | 16 | -0.653023 | -1.364922 | 0.124409 | 0.323917 |
| GOBP_REGULATION_OF_RESPONSE_TO_OXIDATIVE_STRESS | 72 | -0.530698 | -1.364642 | 0.075168 | 0.324002 |
| GOBP_PROTEIN_CONTAINING_COMPLEX_REMODELING | 17 | -0.658658 | -1.364416 | 0.130856 | 0.323994 |
| GOBP_FAT_CELL_DIFFERENTIATION | 152 | -0.482642 | -1.364371 | 0.033981 | 0.323607 |
| GOBP_REGULATION_OF_SPINDLE_CHECKPOINT | 15 | -0.660451 | -1.362279 | 0.134426 | 0.327482 |
| GOBP_TUMOR_NECROSIS_FACTOR_SUPERFAMILY_CYTOKINE_PRODUCTION | 115 | -0.494555 | -1.361339 | 0.042735 | 0.328946 |
| GOBP_T_CELL_LINEAGE_COMMITMENT | 21 | -0.635389 | -1.36123 | 0.124803 | 0.328678 |
| GOBP_GLYCOPROTEIN_CATABOLIC_PROCESS | 18 | -0.642636 | -1.361027 | 0.112164 | 0.32868 |
| GOBP_NEGATIVE_REGULATION_OF_LYMPHOCYTE_DIFFERENTIATION | 37 | -0.573668 | -1.360792 | 0.101449 | 0.32866 |
| GOBP_POSITIVE_REGULATION_OF_NUCLEOTIDE_METABOLIC_PROCESS | 27 | -0.607071 | -1.360435 | 0.104478 | 0.328956 |
| GOBP_POSITIVE_REGULATION_OF_PROTEIN_CONTAINING_COMPLEX_ASSEMBL | 218 | -0.468031 | -1.360352 | 0.028571 | 0.328616 |
| GOBP_AMEBOIDAL_TYPE_CELL_MIGRATION | 295 | -0.455803 | -1.359025 | 0.020675 | 0.33091 |
| GOBP_CELLULAR_RESPONSE_TO_REACTIVE_OXYGEN_SPECIES | 128 | -0.49233 | -1.358794 | 0.045906 | 0.3309 |
| GOBP_REGULATION_OF_CYTOSOLIC_CALCIUM_ION_CONCENTRATION | 201 | -0.474709 | -1.358772 | 0.032295 | 0.330472 |
| GOBP_RELEASE_OF_CYTOCHROME_C_FROM_MITOCHONDRIA | 48 | -0.562714 | -1.358363 | 0.097087 | 0.330869 |
| GOBP_VACUOLAR_ACIDIFICATION | 21 | -0.618187 | -1.358279 | 0.124595 | 0.330594 |
| GOBP_NEGATIVE_REGULATION_OF_ACTIN_FILAMENT_BUNDLE_ASSEMBLY | 21 | -0.627957 | -1.358132 | 0.129032 | 0.330461 |
| GOBP_REGULATION_OF_ANATOMICAL_STRUCTURE_SIZE | 356 | -0.451342 | -1.357966 | 0.019587 | 0.330331 |
| GOBP_LONG_CHAIN_FATTY_ACID_IMPORT_INTO_CELL | 17 | -0.643261 | -1.357448 | 0.111293 | 0.33102 |
| GOBP_CELLULAR_RESPONSE_TO_ORGANIC_CYCLIC_COMPOUND | 379 | -0.452582 | -1.357212 | 0.012766 | 0.331046 |
| GOBP_APOPTOTIC_SIGNALING_PATHWAY | 478 | -0.445391 | -1.35682 | 0.009404 | 0.331317 |
| GOBP_NEGATIVE_REGULATION_OF_PHAGOCYTOSIS | 17 | -0.648387 | -1.356652 | 0.137821 | 0.331213 |
| GOBP_COAGULATION | 236 | -0.461454 | -1.356098 | 0.025316 | 0.331926 |
| GOBP_MEMBRANE_PROTEIN_PROTEOLYSIS | 52 | -0.548254 | -1.355833 | 0.074286 | 0.331985 |
| GOBP_REGULATION_OF_T_CELL_MEDIATED_CYTOTOXICITY | 31 | -0.584962 | -1.355185 | 0.094453 | 0.332812 |
| GOBP_NEGATIVE_REGULATION_OF_LYMPHOCYTE_APOPTOTIC_PROCESS | 27 | -0.594425 | -1.354975 | 0.121069 | 0.332786 |
| GOBP_INTRACELLULAR_STEROL_TRANSPORT | 21 | -0.636017 | -1.354768 | 0.121352 | 0.332767 |
| GOBP_AMYLOID_BETA_METABOLIC_PROCESS | 43 | -0.54762 | -1.354675 | 0.09607 | 0.332478 |
| GOBP_DETOXIFICATION | 92 | -0.510188 | -1.354524 | 0.069767 | 0.33233 |
| GOBP_LEUKOCYTE_APOPTOTIC_PROCESS | 84 | -0.516065 | -1.354355 | 0.084856 | 0.332226 |
| GOBP_NEGATIVE_REGULATION_OF_DNA_BINDING | 38 | -0.575005 | -1.353835 | 0.097525 | 0.332894 |
| GOBP_RESPONSE_TO_ACETYLCHOLINE | 17 | -0.650903 | -1.353363 | 0.135823 | 0.333367 |
| GOBP_MEMBRANE_INVAGINATION | 89 | -0.510453 | -1.352827 | 0.078406 | 0.334045 |
| GOBP_NEGATIVE_REGULATION_OF_RESPONSE_TO_WOUNDING | 51 | -0.538904 | -1.352485 | 0.110632 | 0.334286 |
| GOBP_AMELOGENESIS | 15 | -0.650413 | -1.352065 | 0.151104 | 0.334705 |
| GOBP_POSITIVE_REGULATION_OF_RESPONSE_TO_BIOTIC_STIMULUS | 207 | -0.468709 | -1.351998 | 0.032768 | 0.33439 |
| GOBP_NEGATIVE_REGULATION_OF_CHEMOKINE_PRODUCTION | 19 | -0.640069 | -1.351981 | 0.142157 | 0.333952 |
| GOBP_NEGATIVE_REGULATION_OF_LEUKOCYTE_PROLIFERATION | 70 | -0.524698 | -1.351443 | 0.087483 | 0.334625 |
| GOBP_NEGATIVE_REGULATION_OF_CYTOKINE_PRODUCTION_INVOLVED_IN_IMI | 22 | -0.621337 | -1.351347 | 0.132376 | 0.334353 |
| GOBP_POSITIVE_REGULATION_OF_TRANSFORMING_GROWTH_FACTOR_BETA_F | 16 | -0.65336 | -1.351011 | 0.1408 | 0.334577 |
| GOBP_NEUROTRANSMITTER_REUPTAKE | 17 | -0.635493 | -1.350873 | 0.123377 | 0.334419 |
| GOBP_CD4_POSITIVE_OR_CD8_POSITIVE_ALPHA_BETA_T_CELL_LINEAGE_COM | 18 | -0.652366 | -1.350584 | 0.152846 | 0.33455 |
| GOBP_POSITIVE_REGULATION_OF_ION_TRANSPORT | 480 | -0.446762 | -1.350569 | 0.012346 | 0.334114 |
| GOBP_CD40_SIGNALING_PATHWAY | 15 | -0.646399 | -1.350259 | 0.127731 | 0.334298 |
| GOBP_REGULATION_OF_GLIAL_CELL_MIGRATION | 15 | -0.666952 | -1.350155 | 0.126645 | 0.334039 |
| GOBP_REGULATION_OF_COAGULATION | 45 | -0.560417 | -1.350116 | 0.097525 | 0.33364 |
| GOBP_REGULATION_OF_EARLY_ENDOSOME_TO_LATE_ENDOSOME_TRANSPOR | 18 | -0.645354 | -1.349513 | 0.140625 | 0.33448 |
| GOBP_FILOPODIUM_ASSEMBLY | 52 | -0.547634 | -1.349491 | 0.093151 | 0.334052 |
| GOBP_NEGATIVE_REGULATION_OF_VASCULAR_ASSOCIATED_SMOOTH_MUSCL | 15 | -0.659323 | -1.3485 | 0.139693 | 0.335676 |
| GOBP_REGENERATION | 136 | -0.47672 | -1.347829 | 0.067925 | 0.336613 |
| GOBP_CATECHOL_CONTAINING_COMPOUND_METABOLIC_PROCESS | 25 | -0.60681 | -1.346567 | 0.126697 | 0.338834 |
| GOBP_POSITIVE_REGULATION_OF_ATP_METABOLIC_PROCESS | 27 | -0.593779 | -1.346388 | 0.137134 | 0.33876 |
| GOBP_ENDOCYTOSIS | 473 | -0.439011 | -1.345846 | 0.010482 | 0.339437 |
| GOBP_POSITIVE_REGULATION_OF_EPITHELIAL_CELL_APOPTOTIC_PROCESS | 22 | -0.623852 | -1.344232 | 0.124417 | 0.342403 |
| GOBP_POSITIVE_REGULATION_OF_CHOLESTEROL_EFFLUX | 18 | -0.626665 | -1.344221 | 0.12013 | 0.341945 |
| GOBP_REGULATION_OF_TRANSPOSITION | 19 | -0.627403 | -1.343166 | 0.124006 | 0.343685 |
| GOBP_NEGATIVE_REGULATION_OF_TRANSMEMBRANE_TRANSPORT | 78 | -0.512308 | -1.343066 | 0.090433 | 0.343444 |
| GOBP_HETEROTYPIC_CELL_CELL_ADHESION | 38 | -0.568331 | -1.34301 | 0.106725 | 0.343108 |
| GOBP_CYCLIC_NUCLEOTIDE_MEDIATED_SIGNALING | 41 | -0.562288 | -1.342511 | 0.116809 | 0.343738 |

| | | | | | |
|---|---|---|---|---|---|
| GOBP_PEPTIDYL_ASPARAGINE_MODIFICATION | 27 | 0.25371 | 0.667086 | 0.916201 | 1 |
| GOBP_AXONEMAL_DYNEIN_COMPLEX_ASSEMBLY | 21 | 0.263425 | 0.66572 | 0.894587 | 1 |
| GOBP_REGULATION_OF_NECROTIC_CELL_DEATH | 30 | 0.252652 | 0.664638 | 0.919753 | 1 |
| GOBP_NCRNA_METABOLIC_PROCESS | 458 | 0.1726 | 0.664249 | 1 | 1 |
| GOBP_CELL_VOLUME_HOMEOSTASIS | 20 | 0.272269 | 0.662463 | 0.897959 | 1 |
| GOBP_RESPONSE_TO_AMINO_ACID_STARVATION | 45 | 0.23504 | 0.660285 | 0.936242 | 1 |
| GOBP_NEGATIVE_REGULATION_OF_TUMOR_NECROSIS_FACTOR_MEDIATED_SIGNALIN | 16 | 0.287189 | 0.659529 | 0.858586 | 1 |
| GOBP_NEGATIVE_REGULATION_OF_HISTONE_MODIFICATION | 40 | 0.234804 | 0.659457 | 0.931677 | 1 |
| GOBP_POSITIVE_REGULATION_OF_TORC1_SIGNALING | 17 | 0.276229 | 0.658093 | 0.893048 | 1 |
| GOBP_REGULATION_OF_TRANSLATION_IN_RESPONSE_TO_STRESS | 20 | 0.265015 | 0.657914 | 0.887052 | 1 |
| GOBP_STEROL_BIOSYNTHETIC_PROCESS | 63 | 0.215225 | 0.657703 | 0.976378 | 1 |
| GOBP_PIGMENT_GRANULE_ORGANIZATION | 25 | 0.26214 | 0.656295 | 0.887283 | 1 |
| GOBP_SPINDLE_ORGANIZATION | 171 | 0.187691 | 0.655979 | 1 | 1 |
| GOBP_MONOSACCHARIDE_BIOSYNTHETIC_PROCESS | 78 | 0.207077 | 0.655543 | 0.977012 | 1 |
| GOBP_POSITIVE_REGULATION_OF_CANONICAL_WNT_SIGNALING_PATHWAY | 111 | 0.201399 | 0.655158 | 0.985646 | 1 |
| GOBP_REGULATION_OF_PROTEIN_EXIT_FROM_ENDOPLASMIC_RETICULUM | 21 | 0.267325 | 0.654287 | 0.885246 | 1 |
| GOBP_DNA_DAMAGE_RESPONSE_SIGNAL_TRANSDUCTION_BY_P53_CLASS_MEDIATO | 92 | 0.202819 | 0.652853 | 0.991936 | 1 |
| GOBP_PHOSPHATIDYLGLYCEROL_METABOLIC_PROCESS | 28 | 0.245894 | 0.651556 | 0.926471 | 1 |
| GOBP_PROTEIN_LOCALIZATION_TO_CELL_CELL_JUNCTION | 18 | 0.274059 | 0.651494 | 0.887701 | 1 |
| GOBP_ENDOPLASMIC_RETICULUM_TUBULAR_NETWORK_ORGANIZATION | 18 | 0.270566 | 0.651457 | 0.912935 | 1 |
| GOBP_PURINE_NUCLEOSIDE_BIOSYNTHETIC_PROCESS | 20 | 0.261889 | 0.650971 | 0.894879 | 1 |
| GOBP_PROTEIN_INSERTION_INTO_MEMBRANE | 71 | 0.210338 | 0.645677 | 0.984556 | 1 |
| GOBP_PURINE_NUCLEOSIDE_MONOPHOSPHATE_BIOSYNTHETIC_PROCESS | 22 | 0.255122 | 0.64507 | 0.909333 | 1 |
| GOBP_RRNA_MODIFICATION | 39 | 0.230585 | 0.64389 | 0.963934 | 1 |
| GOBP_FEMALE_MEIOTIC_NUCLEAR_DIVISION | 25 | 0.246287 | 0.64366 | 0.912807 | 1 |
| GOBP_PYRIMIDINE_RIBONUCLEOTIDE_METABOLIC_PROCESS | 24 | 0.254733 | 0.643605 | 0.911765 | 1 |
| GOBP_SPLICEOSOMAL_SNRNP_ASSEMBLY | 40 | 0.231547 | 0.642089 | 0.965625 | 1 |
| GOBP_RNA_POLYADENYLATION | 45 | 0.225022 | 0.641249 | 0.95189 | 1 |
| GOBP_REGULATION_OF_PROTEIN_STABILITY | 259 | 0.176697 | 0.639319 | 1 | 1 |
| GOBP_RIBOSOMAL_SMALL_SUBUNIT_ASSEMBLY | 21 | 0.26616 | 0.637481 | 0.889197 | 1 |
| GOBP_RESPONSE_TO_NERVE_GROWTH_FACTOR | 38 | 0.230971 | 0.636772 | 0.954248 | 1 |
| GOBP_ATP_DEPENDENT_CHROMATIN_REMODELING | 77 | 0.200055 | 0.634308 | 0.992032 | 1 |
| GOBP_NCRNA_EXPORT_FROM_NUCLEUS | 38 | 0.222488 | 0.630466 | 0.949206 | 1 |
| GOBP_MITOTIC_CELL_CYCLE_CHECKPOINT | 149 | 0.184292 | 0.629595 | 0.994792 | 1 |
| GOBP_SNRNA_METABOLIC_PROCESS | 36 | 0.223435 | 0.62622 | 0.958457 | 1 |
| GOBP_RESPONSE_TO_GAMMA_RADIATION | 46 | 0.216155 | 0.623305 | 0.953488 | 1 |
| GOBP_REGULATION_OF_MITOCHONDRIAL_MEMBRANE_PERMEABILITY | 66 | 0.201577 | 0.619352 | 0.988048 | 1 |
| GOBP_OXIDATIVE_PHOSPHORYLATION | 137 | 0.182818 | 0.618625 | 1 | 1 |
| GOBP_TRANSCRIPTION_ELONGATION_FROM_RNA_POLYMERASE_I_PROMOTER | 29 | 0.229188 | 0.612164 | 0.968912 | 1 |
| GOBP_VIRAL_TRANSLATION | 16 | 0.259966 | 0.611554 | 0.917313 | 1 |
| GOBP_REGULATION_OF_TRANSCRIPTION_FROM_RNA_POLYMERASE_II_PROMOTER_I | 73 | 0.194222 | 0.606947 | 0.991525 | 1 |
| GOBP_MULTIVESICULAR_BODY_SORTING_PATHWAY | 35 | 0.223013 | 0.603305 | 0.962848 | 1 |
| GOBP_MITOCHONDRIAL_DEPOLARIZATION | 17 | 0.249639 | 0.602195 | 0.928382 | 1 |
| GOBP_PROTEIN_QUALITY_CONTROL_FOR_MISFOLDED_OR_INCOMPLETELY_SYNTHES | 28 | 0.22914 | 0.600136 | 0.973988 | 1 |
| GOBP_RNA_SURVEILLANCE | 15 | 0.255937 | 0.598371 | 0.924871 | 1 |
| GOBP_SIGNAL_TRANSDUCTION_BY_P53_CLASS_MEDIATOR | 235 | 0.165496 | 0.59602 | 1 | 1 |
| GOBP_NEGATIVE_REGULATION_OF_TOR_SIGNALING | 40 | 0.210882 | 0.595582 | 0.972561 | 1 |
| GOBP_REGULATION_OF_HISTONE_MODIFICATION | 126 | 0.175876 | 0.595353 | 1 | 1 |
| GOBP_PROTEIN_HETEROOLIGOMERIZATION | 19 | 0.24655 | 0.595262 | 0.950685 | 1 |
| GOBP_REGULATION_OF_DEFENSE_RESPONSE_TO_VIRUS_BY_HOST | 36 | 0.217271 | 0.59074 | 0.97205 | 1 |
| GOBP_SNRNA_PROCESSING | 25 | 0.22908 | 0.589943 | 0.936464 | 1 |
| GOBP_ESTABLISHMENT_OF_MITOTIC_SPINDLE_ORIENTATION | 22 | 0.238746 | 0.589511 | 0.953804 | 1 |
| GOBP_PROTEIN_LOCALIZATION_TO_CHROMOSOME_CENTROMERIC_REGION | 25 | 0.229994 | 0.587985 | 0.938416 | 1 |
| GOBP_PROTEIN_LOCALIZATION_TO_LYSOSOME | 38 | 0.214567 | 0.587859 | 0.984663 | 1 |
| GOBP_NUCLEOTIDE_EXCISION_REPAIR_PREINCISION_COMPLEX_ASSEMBLY | 29 | 0.222752 | 0.586737 | 0.974719 | 1 |
| GOBP_SCF_DEPENDENT_PROTEASOMAL_UBIQUITIN_DEPENDENT_PROTEIN_CATABOL | 88 | 0.183786 | 0.583381 | 1 | 1 |
| GOBP_RNA_MODIFICATION | 161 | 0.167757 | 0.582216 | 1 | 1 |
| GOBP_ANAPHASE_PROMOTING_COMPLEX_DEPENDENT_CATABOLIC_PROCESS | 79 | 0.18497 | 0.581533 | 0.995671 | 1 |
| GOBP_PROTEIN_INSERTION_INTO_ER_MEMBRANE | 18 | 0.244937 | 0.580192 | 0.937143 | 1 |
| GOBP_ESTABLISHMENT_OF_PROTEIN_LOCALIZATION_TO_VACUOLE | 44 | 0.203638 | 0.579782 | 0.990506 | 1 |
| GOBP_REGULATION_OF_HISTONE_METHYLATION | 56 | 0.196375 | 0.577122 | 0.983278 | 1 |
| GOBP_PROTON_TRANSMEMBRANE_TRANSPORT | 127 | 0.171366 | 0.574789 | 0.995122 | 1 |
| GOBP_ESTABLISHMENT_OF_MITOTIC_SPINDLE_LOCALIZATION | 27 | 0.213988 | 0.572103 | 0.953488 | 1 |
| GOBP_REGULATION_OF_PROTEIN_KINASE_A_SIGNALING | 15 | 0.256347 | 0.571807 | 0.941953 | 1 |
| GOBP_REGULATION_OF_CENTRIOLE_REPLICATION | 19 | 0.232142 | 0.570839 | 0.948787 | 1 |
| GOBP_GLOBAL_GENOME_NUCLEOTIDE_EXCISION_REPAIR | 26 | 0.214602 | 0.567555 | 0.986376 | 1 |
| GOBP_PYRIMIDINE_RIBONUCLEOSIDE_TRIPHOSPHATE_METABOLIC_PROCESS | 16 | 0.243701 | 0.565824 | 0.956633 | 1 |
| GOBP_PROTEIN_DEGLYCOSYLATION | 28 | 0.210809 | 0.557506 | 0.985955 | 1 |
| GOBP_RNA_PHOSPHODIESTER_BOND_HYDROLYSIS_ENDONUCLEOLYTIC | 71 | 0.180236 | 0.552723 | 1 | 1 |
| GOBP_REGULATION_OF_CELLULAR_RESPONSE_TO_HEAT | 76 | 0.17571 | 0.548561 | 0.995652 | 1 |
| GOBP_TRANSCRIPTION_INITIATION_FROM_RNA_POLYMERASE_I_PROMOTER | 37 | 0.196074 | 0.536498 | 0.99262 | 1 |
| GOBP_ARP2_3_COMPLEX_MEDIATED_ACTIN_NUCLEATION | 35 | 0.196628 | 0.535185 | 0.993976 | 1 |
| GOBP_NEGATIVE_REGULATION_OF_CELL_CYCLE_G2_M_PHASE_TRANSITION | 103 | 0.16644 | 0.534861 | 0.995327 | 1 |
| GOBP_HISTONE_DEUBIQUITINATION | 23 | 0.211025 | 0.534787 | 0.981928 | 1 |
| GOBP_MITOCHONDRIAL_FISSION | 39 | 0.191825 | 0.534713 | 0.993939 | 1 |
| GOBP_GPI_ANCHOR_METABOLIC_PROCESS | 30 | 0.200239 | 0.530663 | 0.982507 | 1 |
| GOBP_REGULATION_OF_AUTOPHAGY_OF_MITOCHONDRION | 31 | 0.197799 | 0.530634 | 0.991202 | 1 |
| GOBP_NUCLEAR_MIGRATION | 18 | 0.222089 | 0.528194 | 0.970803 | 1 |
| GOBP_PROTEIN_IMPORT_INTO_MITOCHONDRIAL_MATRIX | 20 | 0.215601 | 0.525062 | 0.97541 | 1 |
| GOBP_PREASSEMBLY_OF_GPI_ANCHOR_IN_ER_MEMBRANE | 16 | 0.226972 | 0.523218 | 0.95641 | 1 |
| GOBP_PEPTIDYL_ARGININE_MODIFICATION | 15 | 0.230783 | 0.521803 | 0.967828 | 1 |
| GOBP_MITOCHONDRIAL_CYTOCHROME_C_OXIDASE_ASSEMBLY | 22 | 0.203338 | 0.511523 | 0.985836 | 1 |
| GOBP_POLYOL_CATABOLIC_PROCESS | 23 | 0.203878 | 0.510922 | 0.982558 | 1 |
| GOBP_MRNA_CIS_SPLICING_VIA_SPLICEOSOME | 23 | 0.209911 | 0.509977 | 0.982659 | 1 |
| GOBP_RNA_SPLICING_VIA_ENDONUCLEOLYTIC_CLEAVAGE_AND_LIGATION | 16 | 0.214494 | 0.509047 | 0.981912 | 1 |
| GOBP_ENDOSOME_ORGANIZATION | 75 | 0.160707 | 0.507135 | 1 | 1 |
| GOBP_PROTEIN_ADP_RIBOSYLATION | 29 | 0.195527 | 0.505249 | 0.993548 | 1 |
| GOBP_RRNA_METHYLATION | 27 | 0.195531 | 0.504705 | 0.985714 | 1 |
| GOBP_NEGATIVE_REGULATION_OF_DOUBLE_STRAND_BREAK_REPAIR_VIA_HOMOLOG | 18 | 0.210474 | 0.496891 | 0.981675 | 1 |
| GOBP_NUCLEAR_TRANSCRIBED_MRNA_CATABOLIC_PROCESS | 204 | 0.137047 | 0.489749 | 1 | 1 |
| GOBP_RESPONSE_TO_ARSENIC_CONTAINING_SUBSTANCE | 26 | 0.190319 | 0.486261 | 0.991304 | 1 |
| GOBP_RIBONUCLEOPROTEIN_COMPLEX_SUBUNIT_ORGANIZATION | 198 | 0.133076 | 0.482452 | 1 | 1 |
| GOBP_KINETOCHORE_ASSEMBLY | 16 | 0.202876 | 0.473893 | 0.98533 | 1 |
| GOBP_HISTONE_MONOUBIQUITINATION | 29 | 0.17954 | 0.466951 | 0.997135 | 1 |
| GOBP_PROTEIN_HOMOTETRAMERIZATION | 41 | 0.166038 | 0.464839 | 1 | 1 |
| GOBP_MITOCHONDRIAL_RNA_PROCESSING | 17 | 0.199316 | 0.463431 | 0.994638 | 1 |
| GOBP_NUCLEOTIDE_EXCISION_REPAIR_DNA_DUPLEX_UNWINDING | 22 | 0.191707 | 0.46272 | 0.98961 | 1 |
| GOBP_RNA_EXPORT_FROM_NUCLEUS | 135 | 0.134278 | 0.461664 | 1 | 1 |
| GOBP_NCRNA_TRANSCRIPTION | 106 | 0.138411 | 0.459163 | 1 | 1 |
| GOBP_TAIL_ANCHORED_MEMBRANE_PROTEIN_INSERTION_INTO_ER_MEMBRANE | 16 | 0.190571 | 0.454515 | 0.994898 | 1 |
| GOBP_NEGATIVE_REGULATION_OF_DNA_REPAIR | 33 | 0.166952 | 0.451351 | 0.993921 | 1 |
| GOBP_MAINTENANCE_OF_PROTEIN_LOCALIZATION_IN_ORGANELLE | 39 | 0.155749 | 0.447205 | 1 | 1 |
| GOBP_RNA_3_END_PROCESSING | 137 | 0.12953 | 0.446325 | 1 | 1 |
| GOBP_POSITIVE_REGULATION_OF_MACROAUTOPHAGY | 54 | 0.151761 | 0.441314 | 1 | 1 |
| GOBP_SNRNA_TRANSCRIPTION | 74 | 0.13815 | 0.429197 | 1 | 1 |
| GOBP_DNA_TEMPLATED_TRANSCRIPTION_TERMINATION | 74 | 0.135232 | 0.428649 | 1 | 1 |
| GOBP_POSTREPLICATION_REPAIR | 51 | 0.148246 | 0.426568 | 1 | 1 |
| GOBP_TRANSLESION_SYNTHESIS | 41 | 0.148159 | 0.425264 | 1 | 1 |
| GOBP_NEGATIVE_REGULATION_OF_MRNA_PROCESSING | 29 | 0.158265 | 0.417593 | 1 | 1 |
| GOBP_REGULATION_OF_UBIQUITIN_PROTEIN_LIGASE_ACTIVITY | 22 | 0.166168 | 0.416319 | 1 | 1 |
| GOBP_NEGATIVE_REGULATION_OF_RNA_SPLICING | 25 | 0.158228 | 0.415909 | 1 | 1 |
| GOBP_PROTEIN_TRANSMEMBRANE_TRANSPORT | 58 | 0.137314 | 0.413573 | 1 | 1 |
| GOBP_SPLICEOSOMAL_COMPLEX_ASSEMBLY | 58 | 0.135552 | 0.408431 | 1 | 1 |
| GOBP_NON_RECOMBINATIONAL_REPAIR | 94 | 0.125904 | 0.406897 | 1 | 1 |
| GOBP_UBIQUINONE_METABOLIC_PROCESS | 18 | 0.170408 | 0.406266 | 0.994624 | 1 |
| GOBP_CARDIOLIPIN_METABOLIC_PROCESS | 16 | 0.166471 | 0.386499 | 0.99729 | 1 |
| GOBP_RNA_SPLICING_VIA_TRANSESTERIFICATION_REACTIONS | 336 | 0.101677 | 0.384726 | 1 | 1 |
| GOBP_NUCLEOTIDE_EXCISION_REPAIR_DNA_INCISION | 39 | 0.132222 | 0.381083 | 1 | 1 |
| GOBP_HISTONE_H4_ACETYLATION | 65 | 0.123644 | 0.376924 | 1 | 1 |
| GOBP_RESPONSE_TO_MISFOLDED_PROTEIN | 23 | 0.148241 | 0.37044 | 1 | 1 |
| GOBP_HISTONE_UBIQUITINATION | 46 | 0.128289 | 0.367972 | 1 | 1 |
| GOBP_RNA_PHOSPHODIESTER_BOND_HYDROLYSIS | 138 | 0.110444 | 0.367028 | 1 | 1 |
| GOBP_MRNA_PROCESSING | 479 | 0.092196 | 0.354807 | 1 | 1 |
| GOBP_ESTABLISHMENT_OF_RNA_LOCALIZATION | 189 | 0.09566 | 0.341376 | 1 | 1 |
| GOBP_TRANSCRIPTION_COUPLED_NUCLEOTIDE_EXCISION_REPAIR | 73 | 0.106372 | 0.332819 | 1 | 1 |
| GOBP_REGULATION_OF_ALTERNATIVE_MRNA_SPLICING_VIA_SPLICEOSOME | 46 | 0.116033 | 0.327957 | 1 | 1 |
| GOBP_TELOMERE_MAINTENANCE_VIA_SEMI_CONSERVATIVE_REPLICATION | 27 | 0.127068 | 0.325558 | 1 | 0.999935 |

| | | | | | |
|---|---|---|---|---|---|
| GOBP_DENDRITE_DEVELOPMENT | 170 | -0.470973 | -1.341918 | 0.042353 | 0.34456 |
| GOBP_AMINOGLYCAN_BIOSYNTHETIC_PROCESS | 76 | -0.511551 | -1.339301 | 0.088312 | 0.349568 |
| GOBP_REGULATION_OF_EXTRACELLULAR_MATRIX_ORGANIZATION | 35 | -0.571897 | -1.338233 | 0.109012 | 0.351414 |
| GOBP_ZINC_ION_HOMEOSTASIS | 25 | -0.603315 | -1.338194 | 0.135938 | 0.351012 |
| GOBP_CELLULAR_RESPONSE_TO_KETONE | 68 | -0.5145 | -1.338003 | 0.08971 | 0.350984 |
| GOBP_REGULATION_OF_NIK_NF_KAPPAB_SIGNALING | 75 | -0.507126 | -1.337848 | 0.072072 | 0.350833 |
| GOBP_MODIFIED_AMINO_ACID_TRANSPORT | 22 | -0.614902 | -1.337773 | 0.103286 | 0.350516 |
| GOBP_BONE_REMODELING | 61 | -0.527357 | -1.337265 | 0.086436 | 0.351157 |
| GOBP_POSITIVE_REGULATION_OF_LONG_TERM_SYNAPTIC_POTENTIATION | 15 | -0.655854 | -1.337254 | 0.151515 | 0.350705 |
| GOBP_RESPONSE_TO_OXIDATIVE_STRESS | 362 | -0.447935 | -1.337175 | 0.022556 | 0.350388 |
| GOBP_LUNG_ALVEOLUS_DEVELOPMENT | 20 | -0.613835 | -1.336565 | 0.145833 | 0.351167 |
| GOBP_KERATINOCYTE_DIFFERENTIATION | 85 | -0.498428 | -1.336154 | 0.099237 | 0.351604 |
| GOBP_CELLULAR_RESPONSE_TO_LIPID | 393 | -0.440962 | -1.33612 | 0.022246 | 0.351228 |
| GOBP_REGULATION_OF_CHOLESTEROL_EFFLUX | 26 | -0.601517 | -1.335491 | 0.142202 | 0.352061 |
| GOBP_ENSHEATHMENT_OF_NEURONS | 97 | -0.496179 | -1.334371 | 0.07875 | 0.353936 |
| GOBP_REGULATION_OF_LIPID_TRANSPORT | 87 | -0.50358 | -1.334183 | 0.083223 | 0.353847 |
| GOBP_NEGATIVE_REGULATION_OF_CALCIUM_ION_TRANSMEMBRANE_TRANSP( | 22 | -0.606388 | -1.333981 | 0.146457 | 0.35385 |
| GOBP_PROTEIN_O_LINKED_MANNOSYLATION | 17 | -0.65359 | -1.333454 | 0.147708 | 0.35455 |
| GOBP_POSITIVE_REGULATION_OF_T_CELL_CYTOKINE_PRODUCTION | 18 | -0.642984 | -1.333273 | 0.154742 | 0.354451 |
| GOBP_SULFUR_COMPOUND_TRANSPORT | 40 | -0.552276 | -1.332963 | 0.124077 | 0.354642 |
| GOBP_POSITIVE_REGULATION_OF_DNA_BINDING_TRANSCRIPTION_FACTOR_AC | 215 | -0.458052 | -1.331394 | 0.050847 | 0.357557 |
| GOBP_NEURAL_CREST_CELL_MIGRATION | 30 | -0.586656 | -1.331019 | 0.130699 | 0.357961 |
| GOBP_GLIAL_CELL_DIFFERENTIATION | 141 | -0.468068 | -1.330972 | 0.076271 | 0.357573 |
| GOBP_REGULATION_OF_RECEPTOR_SIGNALING_PATHWAY_VIA_STAT | 58 | -0.532628 | -1.330925 | 0.100551 | 0.357201 |
| GOBP_POSITIVE_REGULATION_OF_RECEPTOR_MEDIATED_ENDOCYTOSIS | 36 | -0.556663 | -1.330714 | 0.124088 | 0.357224 |
| GOBP_POSITIVE_REGULATION_OF_EXOCYTOSIS | 62 | -0.526955 | -1.329553 | 0.096234 | 0.359272 |
| GOBP_SMOOTHENED_SIGNALING_PATHWAY | 96 | -0.494597 | -1.329396 | 0.097222 | 0.359132 |
| GOBP_ICOSANOID_SECRETION | 21 | -0.61894 | -1.329128 | 0.129485 | 0.359248 |
| GOBP_OLEFINIC_COMPOUND_METABOLIC_PROCESS | 67 | -0.514579 | -1.328672 | 0.115747 | 0.359738 |
| GOBP_REGULATION_OF_EPITHELIAL_CELL_DIFFERENTIATION | 90 | -0.48834 | -1.328107 | 0.09264 | 0.360467 |
| GOBP_NIK_NF_KAPPAB_SIGNALING | 145 | -0.471964 | -1.32741 | 0.068263 | 0.36144 |
| GOBP_NEUTRAL_LIPID_BIOSYNTHETIC_PROCESS | 36 | -0.559943 | -1.326749 | 0.148256 | 0.362425 |
| GOBP_LYMPHOCYTE_CHEMOTAXIS | 28 | -0.583724 | -1.326507 | 0.125 | 0.362443 |
| GOBP_NUCLEOTIDE_TRANSPORT | 28 | -0.573517 | -1.326108 | 0.126888 | 0.362795 |
| GOBP_POSITIVE_REGULATION_OF_MITOCHONDRION_ORGANIZATION | 78 | -0.503704 | -1.325761 | 0.097368 | 0.363133 |
| GOBP_REGULATION_OF_CELL_CYCLE_G1_S_PHASE_TRANSITION | 146 | -0.469083 | -1.32508 | 0.058612 | 0.364076 |
| GOBP_RESPONSE_TO_INTERFERON_GAMMA | 154 | -0.469578 | -1.324449 | 0.069628 | 0.364993 |
| GOBP_IMMUNOGLOBULIN_PRODUCTION_INVOLVED_IN_IMMUNOGLOBULIN_MEI | 52 | -0.53783 | -1.324265 | 0.099281 | 0.364938 |
| GOBP_MALE_MEIOSIS_I | 16 | -0.644308 | -1.324142 | 0.138978 | 0.364744 |
| GOBP_TELENCEPHALON_DEVELOPMENT | 139 | -0.472981 | -1.323907 | 0.07034 | 0.364817 |
| GOBP_AMINE_METABOLIC_PROCESS | 115 | -0.471638 | -1.323193 | 0.080446 | 0.3659 |
| GOBP_REGULATION_OF_EPITHELIAL_CELL_MIGRATION | 169 | -0.466531 | -1.322355 | 0.071096 | 0.367223 |
| GOBP_EPITHELIAL_CELL_DIFFERENTIATION | 378 | -0.439307 | -1.322346 | 0.023454 | 0.366777 |
| GOBP_REGULATION_OF_SPROUTING_ANGIOGENESIS | 27 | -0.589699 | -1.322213 | 0.140673 | 0.366631 |
| GOBP_TRANSFORMING_GROWTH_FACTOR_BETA_RECEPTOR_SIGNALING_PATH | 146 | -0.472125 | -1.321616 | 0.07049 | 0.36743 |
| GOBP_REGULATION_OF_VOLTAGE_GATED_CALCIUM_CHANNEL_ACTIVITY | 24 | -0.598179 | -1.320853 | 0.140458 | 0.368668 |
| GOBP_POSITIVE_REGULATION_OF_DNA_RECOMBINATION | 41 | -0.552027 | -1.320514 | 0.144681 | 0.368904 |
| GOBP_RECEPTOR_RECYCLING | 23 | -0.609226 | -1.320337 | 0.147975 | 0.368762 |
| GOBP_CARBOHYDRATE_DERIVATIVE_TRANSPORT | 70 | -0.505395 | -1.319552 | 0.090541 | 0.370007 |
| GOBP_CELL_JUNCTION_ASSEMBLY | 250 | -0.447046 | -1.319352 | 0.037862 | 0.369962 |
| GOBP_REGULATION_OF_DNA_METABOLIC_PROCESS | 297 | -0.444062 | -1.318902 | 0.029476 | 0.370428 |
| GOBP_NEGATIVE_REGULATION_OF_PROTEIN_LOCALIZATION_TO_NUCLEUS | 22 | -0.60553 | -1.31837 | 0.151274 | 0.371098 |
| GOBP_POSITIVE_CHEMOTAXIS | 34 | -0.56496 | -1.317869 | 0.11782 | 0.371647 |
| GOBP_MULTI_ORGANISM_LOCALIZATION | 63 | -0.520784 | -1.317506 | 0.104435 | 0.371892 |
| GOBP_AMINO_ACID_TRANSMEMBRANE_TRANSPORT | 58 | -0.52371 | -1.31708 | 0.121379 | 0.372318 |
| GOBP_CELL_CELL_JUNCTION_ASSEMBLY | 88 | -0.494876 | -1.316622 | 0.101316 | 0.372958 |
| GOBP_SMOOTH_MUSCLE_CELL_MIGRATION | 48 | -0.535443 | -1.316573 | 0.140473 | 0.372591 |
| GOBP_REGULATION_OF_FAT_CELL_DIFFERENTIATION | 90 | -0.492541 | -1.316125 | 0.098296 | 0.373089 |
| GOBP_INTERLEUKIN_6_PRODUCTION | 109 | -0.483793 | -1.315957 | 0.086634 | 0.372981 |
| GOBP_BIOMINERALIZATION | 92 | -0.485843 | -1.315579 | 0.102302 | 0.373321 |
| GOBP_TISSUE_MIGRATION | 219 | -0.454588 | -1.315206 | 0.049438 | 0.373691 |
| GOBP_FOREBRAIN_DEVELOPMENT | 211 | -0.456013 | -1.315138 | 0.058553 | 0.373357 |
| GOBP_DNA_RECOMBINATION | 263 | -0.447361 | -1.315135 | 0.042391 | 0.372892 |
| GOBP_BICARBONATE_TRANSPORT | 26 | -0.593753 | -1.314804 | 0.140379 | 0.373109 |
| GOBP_METAL_ION_HOMEOSTASIS | 415 | -0.435954 | -1.31478 | 0.029443 | 0.372688 |
| GOBP_POSITIVE_REGULATION_OF_CELL_GROWTH | 122 | -0.478871 | -1.31463 | 0.085859 | 0.372558 |
| GOBP_ENDOCARDIAL_CUSHION_DEVELOPMENT | 17 | -0.636716 | -1.3146 | 0.177966 | 0.372164 |
| GOBP_PROTEIN_LOCALIZATION_TO_PLASMA_MEMBRANE | 223 | -0.451103 | -1.314516 | 0.053118 | 0.37187 |
| GOBP_NEGATIVE_REGULATION_OF_STRESS_ACTIVATED_PROTEIN_KINASE_SIG | 40 | -0.539369 | -1.314416 | 0.12574 | 0.371616 |
| GOBP_REGULATION_OF_PROTEIN_TARGETING_TO_MEMBRANE | 32 | -0.57227 | -1.314341 | 0.140719 | 0.371303 |
| GOBP_REGULATION_OF_EXTRACELLULAR_MATRIX_DISASSEMBLY | 15 | -0.655551 | -1.314145 | 0.164782 | 0.371255 |
| GOBP_ADULT_LOCOMOTORY_BEHAVIOR | 35 | -0.553134 | -1.313792 | 0.132939 | 0.371541 |
| GOBP_CELL_MORPHOGENESIS_INVOLVED_IN_DIFFERENTIATION | 499 | -0.431414 | -1.313555 | 0.012526 | 0.371603 |
| GOBP_REGULATION_OF_CALCIUM_ION_DEPENDENT_EXOCYTOSIS | 24 | -0.603536 | -1.313449 | 0.156951 | 0.37137 |
| GOBP_RESPONSE_TO_MUSCLE_STRETCH | 15 | -0.654895 | -1.312877 | 0.160331 | 0.372193 |
| GOBP_POSITIVE_REGULATION_OF_ANION_TRANSPORT | 366 | -0.431851 | -1.312627 | 0.025751 | 0.37227 |
| GOBP_APICAL_JUNCTION_ASSEMBLY | 55 | -0.5373 | -1.312494 | 0.120956 | 0.372101 |
| GOBP_REGULATION_OF_PROTEIN_TARGETING | 74 | -0.50066 | -1.31214 | 0.108871 | 0.372361 |
| GOBP_REGULATION_OF_EXOCYTOSIS | 139 | -0.468645 | -1.311995 | 0.068044 | 0.37223 |
| GOBP_CELL_RECOGNITION | 116 | -0.478606 | -1.311512 | 0.086404 | 0.372798 |
| GOBP_EMBRYONIC_PLACENTA_MORPHOGENESIS | 15 | -0.64005 | -1.311421 | 0.191781 | 0.372523 |
| GOBP_MUSCLE_CELL_MIGRATION | 59 | -0.519097 | -1.31111 | 0.122995 | 0.372729 |
| GOBP_AMINO_ACID_IMPORT | 31 | -0.568463 | -1.310683 | 0.147692 | 0.373207 |
| GOBP_FLUID_TRANSPORT | 16 | -0.632292 | -1.310489 | 0.162479 | 0.373182 |
| GOBP_REGULATION_OF_NATURAL_KILLER_CELL_ACTIVATION | 27 | -0.574923 | -1.309575 | 0.159306 | 0.374671 |
| GOBP_REGULATION_OF_APOPTOTIC_SIGNALING_PATHWAY | 277 | -0.443866 | -1.309553 | 0.042841 | 0.374254 |
| GOBP_PROTEIN_LOCALIZATION_TO_ENDOSOME | 23 | -0.587493 | -1.309228 | 0.162461 | 0.374504 |
| GOBP_VASCULAR_ENDOTHELIAL_CELL_PROLIFERATION | 17 | -0.637802 | -1.308811 | 0.166139 | 0.374968 |
| GOBP_NEGATIVE_REGULATION_OF_AUTOPHAGY | 69 | -0.505533 | -1.308325 | 0.103586 | 0.375607 |
| GOBP_REGULATION_OF_DNA_RECOMBINATION | 90 | -0.487448 | -1.307196 | 0.094072 | 0.377614 |
| GOBP_MEMBRANE_LIPID_CATABOLIC_PROCESS | 31 | -0.562721 | -1.307148 | 0.135135 | 0.377248 |
| GOBP_GLAND_DEVELOPMENT | 276 | -0.440339 | -1.306983 | 0.031111 | 0.377146 |
| GOBP_NEURON_APOPTOTIC_PROCESS | 165 | -0.45678 | -1.306821 | 0.065321 | 0.37702 |
| GOBP_REGULATION_OF_SMALL_GTPASE_MEDIATED_SIGNAL_TRANSDUCTION | 271 | -0.440631 | -1.30647 | 0.057206 | 0.377299 |
| GOBP_NEGATIVE_REGULATION_OF_ION_TRANSPORT | 199 | -0.454342 | -1.305927 | 0.068685 | 0.378069 |
| GOBP_PATTERN_SPECIFICATION_PROCESS | 214 | -0.445934 | -1.305745 | 0.063854 | 0.378028 |
| GOBP_EPIDERMIS_DEVELOPMENT | 178 | -0.450462 | -1.305469 | 0.080189 | 0.378197 |
| GOBP_ALCOHOL_CATABOLIC_PROCESS | 36 | -0.55414 | -1.305216 | 0.158126 | 0.378319 |
| GOBP_PEPTIDE_SECRETION | 279 | -0.435867 | -1.30512 | 0.04 | 0.378069 |
| GOBP_NEGATIVE_REGULATION_OF_CELL_DIFFERENTIATION | 398 | -0.432185 | -1.30501 | 0.027719 | 0.37786 |
| GOBP_REGULATION_OF_CELL_MIGRATION_INVOLVED_IN_SPROUTING_ANGIOGI | 27 | -0.577632 | -1.304146 | 0.149701 | 0.37932 |
| GOBP_B_CELL_DIFFERENTIATION | 105 | -0.47105 | -1.303497 | 0.103185 | 0.380319 |
| GOBP_ACID_SECRETION | 19 | -0.611979 | -1.303489 | 0.157724 | 0.379881 |
| GOBP_REGULATION_OF_AXONOGENESIS | 107 | -0.474763 | -1.303156 | 0.101652 | 0.38017 |
| GOBP_REGULATION_OF_T_CELL_MEDIATED_IMMUNITY | 64 | -0.509257 | -1.302785 | 0.121786 | 0.380541 |
| GOBP_REGULATION_OF_BONE_MINERALIZATION | 45 | -0.540781 | -1.302592 | 0.140845 | 0.380503 |
| GOBP_RESPONSE_TO_X_RAY | 28 | -0.569479 | -1.302472 | 0.136432 | 0.380273 |
| GOBP_CENTRAL_NERVOUS_SYSTEM_NEURON_DIFFERENTIATION | 92 | -0.484343 | -1.301799 | 0.104987 | 0.381321 |
| GOBP_REGULATION_OF_DENDRITIC_SPINE_DEVELOPMENT | 45 | -0.531044 | -1.301657 | 0.147564 | 0.381201 |
| GOBP_NEGATIVE_REGULATION_OF_ALPHA_BETA_T_CELL_DIFFERENTIATION | 20 | -0.606612 | -1.301637 | 0.16748 | 0.380798 |
| GOBP_RESPONSE_TO_TOXIC_SUBSTANCE | 166 | -0.455813 | -1.301486 | 0.063232 | 0.380655 |
| GOBP_CELLULAR_EXTRAVASATION | 54 | -0.519672 | -1.301323 | 0.135854 | 0.380562 |
| GOBP_INORGANIC_ANION_TRANSPORT | 103 | -0.478933 | -1.300832 | 0.114943 | 0.381223 |
| GOBP_POSITIVE_REGULATION_OF_ORGANELLE_ORGANIZATION | 497 | -0.426711 | -1.300787 | 0.02079 | 0.380868 |
| GOBP_NUCLEAR_TRANSCRIBED_MRNA_POLY_A_TAIL_SHORTENING | 34 | -0.562635 | -1.299884 | 0.164425 | 0.382371 |
| GOBP_RESPONSE_TO_AXON_INJURY | 52 | -0.523822 | -1.299483 | 0.129125 | 0.382781 |
| GOBP_REGULATION_OF_PROTEIN_CONTAINING_COMPLEX_ASSEMBLY | 373 | -0.431374 | -1.299186 | 0.029979 | 0.382978 |
| GOBP_SODIUM_ION_TRANSPORT | 138 | -0.461919 | -1.298497 | 0.077019 | 0.384077 |
| GOBP_CELL_DEATH_IN_RESPONSE_TO_HYDROGEN_PEROXIDE | 22 | -0.584057 | -1.298281 | 0.1776 | 0.384105 |
| GOBP_B_CELL_APOPTOTIC_PROCESS | 24 | -0.591454 | -1.298045 | 0.15361 | 0.384115 |
| GOBP_NEGATIVE_REGULATION_OF_CALCIUM_ION_TRANSMEMBRANE_TRANSP( | 25 | -0.581419 | -1.298018 | 0.163297 | 0.383728 |
| GOBP_TOLL_LIKE_RECEPTOR_3_SIGNALING_PATHWAY | 18 | -0.60379 | -1.297935 | 0.164179 | 0.383442 |
| GOBP_POSITIVE_REGULATION_OF_VIRAL_LIFE_CYCLE | 28 | -0.579537 | -1.297482 | 0.141159 | 0.383938 |
| GOBP_REGULATION_OF_PROTEIN_LOCALIZATION_TO_CELL_SURFACE | 34 | -0.561327 | -1.296923 | 0.151604 | 0.3847 |
| GOBP_VASCULAR_PROCESS_IN_CIRCULATORY_SYSTEM | 144 | -0.463942 | -1.296772 | 0.11236 | 0.384585 |
| GOBP_REGULATION_OF_GLYCOPROTEIN_METABOLIC_PROCESS | 36 | -0.545036 | -1.296649 | 0.145138 | 0.384424 |
| GOBP_NEGATIVE_REGULATION_OF_T_CELL_APOPTOTIC_PROCESS | 17 | -0.6143 | -1.296446 | 0.176759 | 0.38444 |
| GOBP_PEPTIDYL_THREONINE_DEPHOSPHORYLATION | 21 | -0.606453 | -1.29631 | 0.154321 | 0.384278 |
| GOBP_NEGATIVE_REGULATION_OF_STEM_CELL_DIFFERENTIATION | 16 | -0.618393 | -1.295939 | 0.192243 | 0.384641 |
| GOBP_LIPID_HOMEOSTASIS | 99 | -0.485827 | -1.294826 | 0.099111 | 0.3867 |
| GOBP_CELLULAR_BIOGENIC_AMINE_METABOLIC_PROCESS | 65 | -0.518367 | -1.29479 | 0.120332 | 0.386329 |
| GOBP_REGULATION_OF_NERVOUS_SYSTEM_DEVELOPMENT | 274 | -0.430812 | -1.293739 | 0.06044 | 0.388244 |
| GOBP_POSITIVE_REGULATION_OF_AMYLOID_PRECURSOR_PROTEIN_CATABOLI | 20 | -0.616428 | -1.29256 | 0.149837 | 0.390381 |
| GOBP_RESPONSE_TO_CORTICOSTEROID | 92 | -0.48203 | -1.2925 | 0.115885 | 0.390056 |
| GOBP_NEGATIVE_REGULATION_OF_CATION_TRANSMEMBRANE_TRANSPORT | 53 | -0.514182 | -1.292201 | 0.127536 | 0.390254 |
| GOBP_AMYLOID_PRECURSOR_PROTEIN_METABOLIC_PROCESS | 59 | -0.510428 | -1.291515 | 0.124147 | 0.391363 |
| GOBP_CELLULAR_RESPONSE_TO_VASCULAR_ENDOTHELIAL_GROWTH_FACTO | 41 | -0.530555 | -1.291273 | 0.153285 | 0.391447 |
| GOBP_NEGATIVE_REGULATION_OF_CELL_GROWTH | 135 | -0.466792 | -1.291024 | 0.080586 | 0.391533 |
| GOBP_RHYTHMIC_PROCESS | 212 | -0.443165 | -1.290548 | 0.072832 | 0.392145 |
| GOBP_REGULATION_OF_BIOMINERALIZATION | 56 | -0.514341 | -1.290456 | 0.113314 | 0.391908 |
| GOBP_NEGATIVE_REGULATION_OF_MYELOID_LEUKOCYTE_DIFFERENTIATION | 35 | -0.550955 | -1.289745 | 0.164425 | 0.393056 |
| GOBP_AMINE_BIOSYNTHETIC_PROCESS | 23 | -0.594293 | -1.289254 | 0.176829 | 0.393715 |
| GOBP_KILLING_OF_CELLS_OF_OTHER_ORGANISM | 29 | -0.555984 | -1.289235 | 0.143731 | 0.393303 |
| GOBP_POSITIVE_REGULATION_OF_TRANSMEMBRANE_TRANSPORT | 137 | -0.456875 | -1.288631 | 0.083231 | 0.394177 |
| GOBP_NEGATIVE_REGULATION_OF_B_CELL_PROLIFERATION | 17 | -0.620236 | -1.28854 | 0.175896 | 0.393914 |
| GOBP_FORMATION_OF_PRIMARY_GERM_LAYER | 73 | -0.49736 | -1.287748 | 0.12269 | 0.395262 |
| GOBP_NEGATIVE_REGULATION_OF_TRANSPORTER_ACTIVITY | 48 | -0.526673 | -1.286627 | 0.130737 | 0.39722 |

| | | | | | |
|---|---|---|---|---|---|
| GOBP_INOSITOL_LIPID_MEDIATED_SIGNALING | 141 | -0.46108 | -1.286182 | 0.085502 | 0.397742 |
| GOBP_EXTRACELLULAR_MATRIX_ASSEMBLY | 32 | -0.549518 | -1.285393 | 0.15512 | 0.399065 |
| GOBP_MALE_MEIOTIC_NUCLEAR_DIVISION | 28 | -0.563192 | -1.284848 | 0.154993 | 0.399845 |
| GOBP_REGULATION_OF_G_PROTEIN_COUPLED_RECEPTOR_SIGNALING_PATHW | 97 | -0.481678 | -1.284724 | 0.109434 | 0.399648 |
| GOBP_NEGATIVE_REGULATION_OF_CD4_POSITIVE_ALPHA_BETA_T_CELL_DIFFE | 19 | -0.612595 | -1.284109 | 0.188272 | 0.400628 |
| GOBP_NEGATIVE_REGULATION_OF_T_HELPER_CELL_DIFFERENTIATION | 15 | -0.635205 | -1.284105 | 0.188312 | 0.400184 |
| GOBP_POSITIVE_REGULATION_OF_STEROL_TRANSPORT | 24 | -0.592011 | -1.283496 | 0.162687 | 0.401074 |
| GOBP_XENOBIOTIC_TRANSPORT | 30 | -0.547117 | -1.28314 | 0.15994 | 0.401408 |
| GOBP_POSITIVE_REGULATION_OF_EPITHELIAL_TO_MESENCHYMAL_TRANSITIO | 37 | -0.539883 | -1.28297 | 0.132159 | 0.401294 |
| GOBP_NEGATIVE_REGULATION_OF_INTRACELLULAR_SIGNAL_TRANSDUCTION | 407 | -0.425039 | -1.28252 | 0.034298 | 0.401892 |
| GOBP_REGULATION_OF_PROTEIN_IMPORT | 45 | -0.527871 | -1.282514 | 0.157817 | 0.40146 |
| GOBP_AMINE_TRANSPORT | 41 | -0.535019 | -1.282379 | 0.152758 | 0.401287 |
| GOBP_POSITIVE_REGULATION_OF_INTRACELLULAR_TRANSPORT | 195 | -0.441521 | -1.281996 | 0.081911 | 0.401737 |
| GOBP_DIGESTIVE_SYSTEM_PROCESS | 52 | -0.51119 | -1.281899 | 0.15491 | 0.401513 |
| GOBP_KIDNEY_MORPHOGENESIS | 50 | -0.515009 | -1.281434 | 0.129477 | 0.402082 |
| GOBP_CARDIAC_MUSCLE_CELL_CONTRACTION | 34 | -0.551331 | -1.281136 | 0.154545 | 0.402389 |
| GOBP_MODULATION_BY_SYMBIONT_OF_HOST_CELLULAR_PROCESS | 23 | -0.583649 | -1.280681 | 0.186728 | 0.402895 |
| GOBP_NEGATIVE_REGULATION_OF_TRANSPORT | 289 | -0.431001 | -1.279837 | 0.063736 | 0.404357 |
| GOBP_MEMBRANE_PROTEIN_INTRACELLULAR_DOMAIN_PROTEOLYSIS | 19 | -0.601513 | -1.279741 | 0.1968 | 0.404133 |
| GOBP_RESPONSE_TO_TUMOR_CELL | 24 | -0.57747 | -1.279736 | 0.169059 | 0.403688 |
| GOBP_GLIAL_CELL_MIGRATION | 39 | -0.534209 | -1.279149 | 0.144105 | 0.404569 |
| GOBP_REGULATION_OF_COLLAGEN_METABOLIC_PROCESS | 30 | -0.560292 | -1.278939 | 0.168129 | 0.404563 |
| GOBP_MYELIN_ASSEMBLY | 16 | -0.614337 | -1.278896 | 0.19063 | 0.404219 |
| GOBP_ACTIVATION_OF_MAPKK_ACTIVITY | 41 | -0.533059 | -1.278403 | 0.156385 | 0.404872 |
| GOBP_EMBRYONIC_HEART_TUBE_MORPHOGENESIS | 42 | -0.521829 | -1.278065 | 0.161932 | 0.405147 |
| GOBP_POSITIVE_REGULATION_OF_MRNA_CATABOLIC_PROCESS | 46 | -0.531756 | -1.277372 | 0.152682 | 0.40631 |
| GOBP_AMINO_ACID_TRANSPORT | 91 | -0.479467 | -1.277307 | 0.129887 | 0.405996 |
| GOBP_NEGATIVE_REGULATION_OF_EPITHELIAL_CELL_PROLIFERATION | 88 | -0.476999 | -1.277113 | 0.114474 | 0.405985 |
| GOBP_CALCIUM_ION_IMPORT | 50 | -0.511496 | -1.276699 | 0.155433 | 0.406486 |
| GOBP_CELL_CELL_JUNCTION_ORGANIZATION | 132 | -0.458189 | -1.27669 | 0.107317 | 0.406058 |
| GOBP_NEURON_PROJECTION_REGENERATION | 40 | -0.523459 | -1.276425 | 0.15495 | 0.406213 |
| GOBP_ACTIN_CYTOSKELETON_REORGANIZATION | 78 | -0.483905 | -1.276266 | 0.104636 | 0.406114 |
| GOBP_RESPONSE_TO_GROWTH_FACTOR | 473 | -0.419421 | -1.275202 | 0.028213 | 0.407986 |
| GOBP_REGIONALIZATION | 159 | -0.450471 | -1.275178 | 0.085106 | 0.407605 |
| GOBP_REGULATION_OF_NEURON_PROJECTION_DEVELOPMENT | 313 | -0.426446 | -1.274904 | 0.055127 | 0.407803 |
| GOBP_REGULATION_OF_HORMONE_LEVELS | 308 | -0.426782 | -1.273718 | 0.06423 | 0.410031 |
| GOBP_CARDIAC_MUSCLE_CELL_MEMBRANE_REPOLARIZATION | 19 | -0.603481 | -1.273576 | 0.170654 | 0.409908 |
| GOBP_REGULATION_OF_TYPE_2_IMMUNE_RESPONSE | 23 | -0.587023 | -1.273547 | 0.186186 | 0.409514 |
| GOBP_NEGATIVE_REGULATION_OF_WOUND_HEALING | 40 | -0.533037 | -1.273381 | 0.175953 | 0.409469 |
| GOBP_REGULATION_OF_LEUKOCYTE_APOPTOTIC_PROCESS | 66 | -0.495537 | -1.273286 | 0.14941 | 0.409225 |
| GOBP_NEGATIVE_REGULATION_OF_LIPID_TRANSPORT | 17 | -0.607608 | -1.272395 | 0.194489 | 0.410794 |
| GOBP_PHAGOCYTOSIS_RECOGNITION | 44 | -0.526529 | -1.271869 | 0.154494 | 0.411479 |
| GOBP_REGULATION_OF_MAST_CELL_ACTIVATION | 34 | -0.547176 | -1.271828 | 0.178363 | 0.411128 |
| GOBP_PHENOL_CONTAINING_COMPOUND_METABOLIC_PROCESS | 60 | -0.508681 | -1.270658 | 0.137363 | 0.413332 |
| GOBP_METAL_ION_TRANSPORT | 319 | -0.425148 | -1.270561 | 0.062366 | 0.41312 |
| GOBP_REGULATION_OF_TOLL_LIKE_RECEPTOR_SIGNALING_PATHWAY | 62 | -0.503744 | -1.269259 | 0.13807 | 0.41555 |
| GOBP_BONE_MINERALIZATION | 68 | -0.485447 | -1.268731 | 0.13272 | 0.416302 |
| GOBP_NEUTRAL_LIPID_METABOLIC_PROCESS | 85 | -0.475265 | -1.268162 | 0.141333 | 0.417084 |
| GOBP_NEGATIVE_REGULATION_OF_ALPHA_BETA_T_CELL_ACTIVATION | 33 | -0.549119 | -1.267994 | 0.183359 | 0.417016 |
| GOBP_RESPONSE_TO_INTERLEUKIN_4 | 33 | -0.55733 | -1.266686 | 0.185401 | 0.419579 |
| GOBP_TOLL_LIKE_RECEPTOR_9_SIGNALING_PATHWAY | 25 | -0.567742 | -1.266146 | 0.194099 | 0.420361 |
| GOBP_PHOTOPERIODISM | 28 | -0.561128 | -1.265981 | 0.193009 | 0.420302 |
| GOBP_AMYLOID_PRECURSOR_PROTEIN_CATABOLIC_PROCESS | 46 | -0.519931 | -1.265858 | 0.149573 | 0.420149 |
| GOBP_REGULATION_OF_B_CELL_ACTIVATION | 136 | -0.451821 | -1.265841 | 0.113139 | 0.419744 |
| GOBP_NEGATIVE_REGULATION_OF_ESTABLISHMENT_OF_PROTEIN_LOCALIZATI | 92 | -0.472886 | -1.265774 | 0.139257 | 0.419423 |
| GOBP_POSITIVE_REGULATION_OF_SECRETION | 194 | -0.436882 | -1.265604 | 0.09546 | 0.41941 |
| GOBP_POSITIVE_REGULATION_OF_INTRACELLULAR_PROTEIN_TRANSPORT | 161 | -0.445607 | -1.265477 | 0.099526 | 0.419243 |
| GOBP_REPRODUCTIVE_SYSTEM_DEVELOPMENT | 249 | -0.434691 | -1.26539 | 0.08324 | 0.418988 |
| GOBP_NEGATIVE_REGULATION_OF_PROTEIN_LOCALIZATION_TO_CELL_PERIPH | 18 | -0.597593 | -1.265339 | 0.184039 | 0.418653 |
| GOBP_REGULATION_OF_ACTOMYOSIN_STRUCTURE_ORGANIZATION | 76 | -0.478261 | -1.265162 | 0.14094 | 0.418601 |
| GOBP_POSITIVE_REGULATION_OF_ESTABLISHMENT_OF_PROTEIN_LOCALIZATIO | 271 | -0.429118 | -1.264808 | 0.060936 | 0.418939 |
| GOBP_CYTOPLASMIC_SEQUESTERING_OF_PROTEIN | 20 | -0.605721 | -1.264628 | 0.201278 | 0.418885 |
| GOBP_REGULATION_OF_OSTEOBLAST_DIFFERENTIATION | 72 | -0.480565 | -1.263818 | 0.146309 | 0.420251 |
| GOBP_RETINOIC_ACID_RECEPTOR_SIGNALING_PATHWAY | 23 | -0.57019 | -1.263136 | 0.200972 | 0.421385 |
| GOBP_NEGATIVE_REGULATION_OF_MAPK_CASCADE | 137 | -0.451739 | -1.262759 | 0.119077 | 0.421804 |
| GOBP_CIRCADIAN_REGULATION_OF_GENE_EXPRESSION | 55 | -0.505892 | -1.262028 | 0.160935 | 0.423125 |
| GOBP_PHOSPHATIDYLINOSITOL_3_KINASE_SIGNALING | 108 | -0.455477 | -1.261372 | 0.115629 | 0.424214 |
| GOBP_RESPONSE_TO_PURINE_CONTAINING_COMPOUND | 100 | -0.467196 | -1.26099 | 0.129032 | 0.424657 |
| GOBP_POSITIVE_REGULATION_OF_SYNAPSE_ASSEMBLY | 24 | -0.579403 | -1.26042 | 0.175719 | 0.425455 |
| GOBP_TRANSFORMING_GROWTH_FACTOR_BETA_PRODUCTION | 29 | -0.55494 | -1.259585 | 0.198198 | 0.426911 |
| GOBP_RESPONSE_TO_CADMIUM_ION | 46 | -0.510311 | -1.259276 | 0.157319 | 0.427151 |
| GOBP_REGULATION_OF_SYNAPSE_STRUCTURE_OR_ACTIVITY | 139 | -0.450218 | -1.259197 | 0.113721 | 0.42688 |
| GOBP_REGULATION_OF_CAMP_MEDIATED_SIGNALING | 21 | -0.577662 | -1.259192 | 0.191291 | 0.426442 |
| GOBP_SKIN_DEVELOPMENT | 147 | -0.443323 | -1.258862 | 0.101327 | 0.426743 |
| GOBP_RESPONSE_TO_CARBOHYDRATE | 162 | -0.439729 | -1.25885 | 0.108434 | 0.426324 |
| GOBP_POSITIVE_REGULATION_OF_PRODUCTION_OF_MOLECULAR_MEDIATOR_ | 81 | -0.481874 | -1.258692 | 0.148438 | 0.426249 |
| GOBP_CALCIUM_ION_REGULATED_EXOCYTOSIS | 36 | -0.534065 | -1.258528 | 0.178102 | 0.426196 |
| GOBP_REGULATION_OF_LIPID_KINASE_ACTIVITY | 56 | -0.500408 | -1.258259 | 0.160689 | 0.426396 |
| GOBP_RESPONSE_TO_VITAMIN | 53 | -0.51231 | -1.258003 | 0.165498 | 0.426575 |
| GOBP_NEGATIVE_REGULATION_OF_CELL_PROJECTION_ORGANIZATION | 131 | -0.444957 | -1.256952 | 0.117216 | 0.42855 |
| GOBP_REGULATION_OF_GLIOGENESIS | 59 | -0.49189 | -1.256702 | 0.155028 | 0.428689 |
| GOBP_ENDOTHELIAL_CELL_PROLIFERATION | 101 | -0.465714 | -1.256629 | 0.130208 | 0.428435 |
| GOBP_NEGATIVE_REGULATION_OF_LIPID_STORAGE | 17 | -0.610623 | -1.256375 | 0.207071 | 0.428577 |
| GOBP_REGULATION_OF_GTPASE_ACTIVITY | 389 | -0.415342 | -1.256276 | 0.062633 | 0.428364 |
| GOBP_POSTSYNAPTIC_SIGNAL_TRANSDUCTION | 21 | -0.584373 | -1.255411 | 0.209936 | 0.429853 |
| GOBP_RECEPTOR_MEDIATED_ENDOCYTOSIS | 213 | -0.434841 | -1.25516 | 0.092677 | 0.429976 |
| GOBP_POSITIVE_REGULATION_OF_TUMOR_NECROSIS_FACTOR_SUPERFAMILY_ | 70 | -0.477974 | -1.255039 | 0.158879 | 0.429798 |
| GOBP_DENDRITE_MORPHOGENESIS | 99 | -0.473949 | -1.254692 | 0.141009 | 0.430112 |
| GOBP_REGULATION_OF_PATTERN_RECOGNITION_RECEPTOR_SIGNALING_PATH | 87 | -0.468566 | -1.254493 | 0.131854 | 0.430143 |
| GOBP_REGULATION_OF_PEPTIDE_SECRETION | 214 | -0.43179 | -1.254381 | 0.095512 | 0.429975 |
| GOBP_REGULATION_OF_METAL_ION_TRANSPORT | 161 | -0.436139 | -1.254258 | 0.108076 | 0.429815 |
| GOBP_POSITIVE_REGULATION_OF_MUSCLE_CELL_APOPTOTIC_PROCESS | 21 | -0.567099 | -1.253516 | 0.203498 | 0.431109 |
| GOBP_CELLULAR_RESPONSE_TO_GROWTH_HORMONE_STIMULUS | 18 | -0.614356 | -1.253449 | 0.22508 | 0.430803 |
| GOBP_ICOSANOID_BIOSYNTHETIC_PROCESS | 29 | -0.56022 | -1.253307 | 0.19084 | 0.43069 |
| GOBP_POSITIVE_REGULATION_OF_POTASSIUM_ION_TRANSPORT | 26 | -0.568108 | -1.253207 | 0.192366 | 0.430468 |
| GOBP_B_CELL_RECEPTOR_SIGNALING_PATHWAY | 84 | -0.465726 | -1.253157 | 0.161333 | 0.430134 |
| GOBP_REGULATION_OF_BONE_REMODELING | 34 | -0.541811 | -1.253107 | 0.192082 | 0.429815 |
| GOBP_REGULATION_OF_CALCIUM_ION_TRANSMEMBRANE_TRANSPORTER_ACT | 55 | -0.497296 | -1.25214 | 0.152893 | 0.431581 |
| GOBP_NEGATIVE_REGULATION_OF_NEURON_PROJECTION_DEVELOPMENT | 95 | -0.475412 | -1.251506 | 0.144686 | 0.432611 |
| GOBP_ENDOSOME_TO_LYSOSOME_TRANSPORT | 50 | -0.516257 | -1.249751 | 0.181818 | 0.436139 |
| GOBP_POSITIVE_REGULATION_OF_DEPHOSPHORYLATION | 50 | -0.515086 | -1.249636 | 0.161017 | 0.435958 |
| GOBP_CELLULAR_RESPONSE_TO_VITAMIN | 17 | -0.592296 | -1.249479 | 0.222395 | 0.435849 |
| GOBP_REGULATION_OF_INTRACELLULAR_PROTEIN_TRANSPORT | 220 | -0.42717 | -1.247441 | 0.08059 | 0.439959 |
| GOBP_MULTICELLULAR_ORGANISMAL_SIGNALING | 102 | -0.462992 | -1.246499 | 0.155759 | 0.44168 |
| GOBP_POSITIVE_REGULATION_OF_TRIGLYCERIDE_METABOLIC_PROCESS | 16 | -0.6063 | -1.246437 | 0.217666 | 0.441391 |
| GOBP_REGULATION_OF_CELL_PROJECTION_ORGANIZATION | 475 | -0.407787 | -1.246278 | 0.047518 | 0.441338 |
| GOBP_REGULATION_OF_PHOSPHOPROTEIN_PHOSPHATASE_ACTIVITY | 98 | -0.462687 | -1.24574 | 0.138232 | 0.442215 |
| GOBP_REGULATION_OF_NEUROGENESIS | 234 | -0.42795 | -1.245185 | 0.098398 | 0.443086 |
| GOBP_NEGATIVE_REGULATION_OF_BINDING | 124 | -0.455244 | -1.245086 | 0.139057 | 0.442865 |
| GOBP_RESPONSE_TO_INORGANIC_SUBSTANCE | 385 | -0.41493 | -1.244942 | 0.069892 | 0.442766 |
| GOBP_POSITIVE_REGULATION_OF_GTPASE_ACTIVITY | 331 | -0.417162 | -1.244066 | 0.073197 | 0.444332 |
| GOBP_POSITIVE_REGULATION_OF_SYNAPTIC_TRANSMISSION | 87 | -0.465382 | -1.243998 | 0.158163 | 0.444043 |
| GOBP_CELL_FATE_SPECIFICATION | 31 | -0.538026 | -1.243494 | 0.195231 | 0.444841 |
| GOBP_PLATELET_DEGRANULATION | 97 | -0.463839 | -1.243165 | 0.147477 | 0.445166 |
| GOBP_DNA_REPLICATION_CHECKPOINT | 16 | -0.596831 | -1.242836 | 0.225649 | 0.445505 |
| GOBP_CELLULAR_COMPONENT_DISASSEMBLY | 481 | -0.409336 | -1.242407 | 0.058031 | 0.445991 |
| GOBP_PROTEIN_LIPID_COMPLEX_SUBUNIT_ORGANIZATION | 32 | -0.538306 | -1.242357 | 0.212166 | 0.445663 |
| GOBP_MODULATION_OF_PROCESS_OF_OTHER_ORGANISM_INVOLVED_IN_SYM | 83 | -0.467856 | -1.242025 | 0.157623 | 0.445969 |
| GOBP_POSITIVE_REGULATION_OF_LIPASE_ACTIVITY | 43 | -0.522346 | -1.241023 | 0.177215 | 0.447804 |
| GOBP_PIGMENT_CELL_DIFFERENTIATION | 24 | -0.569838 | -1.24087 | 0.215596 | 0.447708 |
| GOBP_REGULATION_OF_FATTY_ACID_METABOLIC_PROCESS | 66 | -0.482326 | -1.240868 | 0.150203 | 0.447261 |
| GOBP_REGULATION_OF_BODY_FLUID_LEVELS | 331 | -0.414115 | -1.240598 | 0.06681 | 0.44745 |
| GOBP_GASTRULATION | 121 | -0.445911 | -1.240362 | 0.145939 | 0.447532 |
| GOBP_STEROL_HOMEOSTASIS | 52 | -0.497091 | -1.240122 | 0.194406 | 0.447615 |
| GOBP_PHAGOSOME_ACIDIFICATION | 24 | -0.561234 | -1.240117 | 0.210123 | 0.447177 |
| GOBP_CELL_CELL_RECOGNITION | 45 | -0.51443 | -1.239743 | 0.176136 | 0.447571 |
| GOBP_EPIDERMAL_GROWTH_FACTOR_RECEPTOR_SIGNALING_PATHWAY | 94 | -0.465049 | -1.238732 | 0.156838 | 0.44942 |
| GOBP_PRI_MIRNA_TRANSCRIPTION_BY_RNA_POLYMERASE_II | 38 | -0.528264 | -1.238394 | 0.176912 | 0.449812 |
| GOBP_POSTSYNAPSE_ORGANIZATION | 118 | -0.451385 | -1.238384 | 0.151625 | 0.449383 |
| GOBP_ERBB_SIGNALING_PATHWAY | 112 | -0.46262 | -1.237412 | 0.14801 | 0.451132 |
| GOBP_CELL_JUNCTION_MAINTENANCE | 19 | -0.585338 | -1.237385 | 0.234513 | 0.450748 |
| GOBP_REGULATION_OF_LIPID_CATABOLIC_PROCESS | 38 | -0.52544 | -1.237055 | 0.165932 | 0.451054 |
| GOBP_POSITIVE_REGULATION_OF_PROTEIN_LOCALIZATION_TO_CELL_SURFAC | 15 | -0.608746 | -1.236576 | 0.252855 | 0.451704 |
| GOBP_REGULATION_OF_NITRIC_OXIDE_SYNTHASE_ACTIVITY | 39 | -0.517365 | -1.236363 | 0.183119 | 0.451762 |
| GOBP_NEGATIVE_REGULATION_OF_TOLL_LIKE_RECEPTOR_SIGNALING_PATHW | 34 | -0.529124 | -1.236354 | 0.193353 | 0.451335 |
| GOBP_B_CELL_ACTIVATION | 241 | -0.421837 | -1.236087 | 0.10962 | 0.451492 |
| GOBP_REGULATION_OF_PEPTIDE_TRANSPORT | 452 | -0.40773 | -1.235829 | 0.060797 | 0.451627 |
| GOBP_ANION_TRANSMEMBRANE_TRANSPORT | 379 | -0.408274 | -1.23566 | 0.068303 | 0.451581 |
| GOBP_L_ALPHA_AMINO_ACID_TRANSMEMBRANE_TRANSPORT | 40 | -0.513254 | -1.23503 | 0.189802 | 0.452672 |
| GOBP_T_CELL_CYTOKINE_PRODUCTION | 30 | -0.535048 | -1.234976 | 0.204651 | 0.452351 |
| GOBP_NEGATIVE_REGULATION_OF_EXOCYTOSIS | 23 | -0.554487 | -1.23457 | 0.212074 | 0.452832 |
| GOBP_NEGATIVE_REGULATION_OF_INTRACELLULAR_TRANSPORT | 43 | -0.508866 | -1.233585 | 0.19152 | 0.45457 |
| GOBP_NEGATIVE_REGULATION_OF_INTRACELLULAR_PROTEIN_TRANSPORT | 34 | -0.523923 | -1.233378 | 0.204173 | 0.454608 |
| GOBP_REGULATION_OF_DENDRITE_MORPHOGENESIS | 49 | -0.497236 | -1.233357 | 0.200546 | 0.45421 |
| GOBP_PLATELET_ACTIVATION | 115 | -0.447641 | -1.232294 | 0.136536 | 0.45629 |
| GOBP_POSITIVE_REGULATION_OF_NERVOUS_SYSTEM_DEVELOPMENT | 174 | -0.429081 | -1.23173 | 0.128957 | 0.457169 |
| GOBP_POSITIVE_REGULATION_OF_INTERLEUKIN_8_PRODUCTION | 45 | -0.513003 | -1.231096 | 0.174825 | 0.458175 |

| | | | | | |
|---|---|---|---|---|---|
| GOBP_POSITIVE_REGULATION_OF_AXONOGENESIS | 59 | -0.490221 | -1.229903 | 0.200561 | 0.460504 |
| GOBP_DENDRITIC_SPINE_DEVELOPMENT | 75 | -0.473609 | -1.229859 | 0.183288 | 0.460145 |
| GOBP_GAMMA_DELTA_T_CELL_ACTIVATION | 15 | -0.6092 | -1.229171 | 0.240401 | 0.461228 |
| GOBP_NEGATIVE_REGULATION_OF_PEPTIDE_SECRETION | 49 | -0.498191 | -1.228957 | 0.202878 | 0.461273 |
| GOBP_POSITIVE_REGULATION_OF_PHOSPHATIDYLINOSITOL_3_KINASE_SIGNAL | 59 | -0.488585 | -1.228175 | 0.186992 | 0.462658 |
| GOBP_LEARNING | 79 | -0.465478 | -1.228095 | 0.183099 | 0.462392 |
| GOBP_NEGATIVE_REGULATION_OF_NERVOUS_SYSTEM_DEVELOPMENT | 86 | -0.457998 | -1.227811 | 0.174905 | 0.462619 |
| GOBP_CALCIUM_ION_TRANSPORT | 265 | -0.418319 | -1.227805 | 0.091212 | 0.462178 |
| GOBP_MYELIN_MAINTENANCE | 16 | -0.598868 | -1.227712 | 0.250814 | 0.461972 |
| GOBP_PROTEIN_LOCALIZATION_TO_CELL_PERIPHERY | 260 | -0.41611 | -1.226955 | 0.098889 | 0.463374 |
| GOBP_POSITIVE_REGULATION_OF_NEUROGENESIS | 156 | -0.435149 | -1.226685 | 0.141119 | 0.463543 |
| GOBP_CELL_GROWTH | 352 | -0.406359 | -1.22643 | 0.080786 | 0.463719 |
| GOBP_PROTEIN_NEDDYLATION | 17 | -0.595495 | -1.226337 | 0.25585 | 0.463505 |
| GOBP_GLYCOPROTEIN_METABOLIC_PROCESS | 302 | -0.412908 | -1.225698 | 0.087241 | 0.464639 |
| GOBP_LONG_TERM_MEMORY | 21 | -0.566762 | -1.225071 | 0.218299 | 0.465673 |
| GOBP_CARDIAC_CONDUCTION | 77 | -0.476419 | -1.225058 | 0.167794 | 0.465249 |
| GOBP_NEGATIVE_REGULATION_OF_MYELOID_CELL_DIFFERENTIATION | 65 | -0.481113 | -1.224985 | 0.19699 | 0.464968 |
| GOBP_ANTIGEN_PROCESSING_AND_PRESENTATION_OF_PEPTIDE_ANTIGEN_VI | 93 | -0.459259 | -1.224962 | 0.160875 | 0.464567 |
| GOBP_MITOCHONDRIAL_CALCIUM_ION_HOMEOSTASIS | 24 | -0.557999 | -1.224714 | 0.238897 | 0.464697 |
| GOBP_NEGATIVE_REGULATION_OF_RESPONSE_TO_OXIDATIVE_STRESS | 17 | -0.585975 | -1.224641 | 0.229642 | 0.464406 |
| GOBP_IMPORT_ACROSS_PLASMA_MEMBRANE | 101 | -0.448731 | -1.224189 | 0.150845 | 0.464998 |
| GOBP_MYD88_INDEPENDENT_TOLL_LIKE_RECEPTOR_SIGNALING_PATHWAY | 31 | -0.532489 | -1.223721 | 0.215022 | 0.465662 |
| GOBP_REGULATION_OF_CHROMATIN_BINDING | 20 | -0.569326 | -1.223218 | 0.245283 | 0.466394 |
| GOBP_PHOSPHATIDYLCHOLINE_METABOLIC_PROCESS | 65 | -0.473346 | -1.223203 | 0.191197 | 0.465985 |
| GOBP_CELL_DIFFERENTIATION_INVOLVED_IN_KIDNEY_DEVELOPMENT | 35 | -0.518011 | -1.222955 | 0.229412 | 0.466114 |
| GOBP_NEGATIVE_REGULATION_OF_ANION_TRANSPORT | 135 | -0.434369 | -1.222773 | 0.160434 | 0.466081 |
| GOBP_ENDOTHELIAL_CELL_MIGRATION | 157 | -0.433376 | -1.222692 | 0.15858 | 0.465813 |
| GOBP_GTP_METABOLIC_PROCESS | 22 | -0.554772 | -1.22204 | 0.2288 | 0.466903 |
| GOBP_CEREBELLAR_PURKINJE_CELL_LAYER_DEVELOPMENT | 18 | -0.58717 | -1.222003 | 0.234146 | 0.466544 |
| GOBP_RESPONSE_TO_FATTY_ACID | 43 | -0.503152 | -1.221977 | 0.224894 | 0.466146 |
| GOBP_NEURON_DEATH | 255 | -0.41802 | -1.221827 | 0.114699 | 0.466065 |
| GOBP_CHAPERONE_MEDIATED_PROTEIN_COMPLEX_ASSEMBLY | 20 | -0.572714 | -1.221663 | 0.249589 | 0.46601 |
| GOBP_NEGATIVE_REGULATION_OF_CALCIUM_ION_TRANSPORT | 36 | -0.514256 | -1.22126 | 0.210526 | 0.466486 |
| GOBP_HOMEOSTASIS_OF_NUMBER_OF_CELLS | 215 | -0.424264 | -1.220663 | 0.129505 | 0.46743 |
| GOBP_POSITIVE_REGULATION_OF_PRI_MIRNA_TRANSCRIPTION_BY_RNA_POLY | 29 | -0.53333 | -1.220604 | 0.235669 | 0.467107 |
| GOBP_REGULATION_OF_MACROPHAGE_ACTIVATION | 39 | -0.504765 | -1.220522 | 0.189802 | 0.466876 |
| GOBP_LONG_TERM_SYNAPTIC_POTENTIATION | 51 | -0.49404 | -1.22052 | 0.196581 | 0.466449 |
| GOBP_NEGATIVE_REGULATION_OF_GROWTH | 176 | -0.424877 | -1.220479 | 0.155952 | 0.466111 |
| GOBP_ANTIGEN_PROCESSING_AND_PRESENTATION_OF_EXOGENOUS_PEPTIDE | 75 | -0.46617 | -1.219613 | 0.181092 | 0.467664 |
| GOBP_POSITIVE_REGULATION_OF_EPITHELIAL_CELL_MIGRATION | 113 | -0.446238 | -1.219176 | 0.161893 | 0.468271 |
| GOBP_NEGATIVE_REGULATION_OF_CELL_SUBSTRATE_ADHESION | 41 | -0.504097 | -1.219159 | 0.22063 | 0.467878 |
| GOBP_NEGATIVE_REGULATION_OF_TUMOR_NECROSIS_FACTOR_SUPERFAMILY | 42 | -0.500733 | -1.218158 | 0.194525 | 0.469738 |
| GOBP_POSITIVE_REGULATION_OF_POTASSIUM_ION_TRANSMEMBRANE_TRANS | 22 | -0.568484 | -1.218036 | 0.240527 | 0.469554 |
| GOBP_CELLULAR_RESPONSE_TO_ACID_CHEMICAL | 54 | -0.484856 | -1.217255 | 0.176224 | 0.470987 |
| GOBP_HISTONE_H3_DEACETYLATION | 19 | -0.567746 | -1.217161 | 0.253125 | 0.470769 |
| GOBP_REGULATION_OF_VIRAL_LIFE_CYCLE | 124 | -0.438772 | -1.216386 | 0.173198 | 0.472171 |
| GOBP_REGULATION_OF_RESPONSE_TO_EXTRACELLULAR_STIMULUS | 16 | -0.592873 | -1.21625 | 0.22504 | 0.472039 |
| GOBP_REGULATION_OF_INTRACELLULAR_TRANSPORT | 304 | -0.409879 | -1.216113 | 0.118093 | 0.471966 |
| GOBP_POSITIVE_REGULATION_OF_SMOOTH_MUSCLE_CELL_MIGRATION | 27 | -0.541278 | -1.216094 | 0.216867 | 0.471576 |
| GOBP_NEGATIVE_REGULATION_OF_AXONOGENESIS | 46 | -0.494617 | -1.215617 | 0.216885 | 0.472262 |
| GOBP_COMPLEMENT_ACTIVATION | 74 | -0.468502 | -1.214206 | 0.169514 | 0.475126 |
| GOBP_INTERLEUKIN_1_PRODUCTION | 85 | -0.462744 | -1.213324 | 0.179045 | 0.476793 |
| GOBP_POSITIVE_REGULATION_OF_TRANSCRIPTION_INITIATION_FROM_RNA_PO | 26 | -0.53012 | -1.212941 | 0.237288 | 0.477237 |
| GOBP_SODIUM_ION_HOMEOSTASIS | 21 | -0.562906 | -1.21286 | 0.268489 | 0.476989 |
| GOBP_REGULATION_OF_GLIAL_CELL_PROLIFERATION | 26 | -0.534373 | -1.212003 | 0.217791 | 0.478614 |
| GOBP_PHAGOSOME_MATURATION | 39 | -0.510112 | -1.211223 | 0.223704 | 0.480002 |
| GOBP_AMINOGLYCAN_CATABOLIC_PROCESS | 43 | -0.502585 | -1.210098 | 0.196676 | 0.482135 |
| GOBP_RECEPTOR_METABOLIC_PROCESS | 128 | -0.43793 | -1.209853 | 0.158596 | 0.482274 |
| GOBP_EMBRYONIC_HEART_TUBE_DEVELOPMENT | 47 | -0.49319 | -1.209701 | 0.220315 | 0.48217 |
| GOBP_RESPONSE_TO_MECHANICAL_STIMULUS | 139 | -0.436558 | -1.209584 | 0.152334 | 0.481981 |
| GOBP_ANTIGEN_PROCESSING_AND_PRESENTATION_OF_PEPTIDE_OR_POLYSA | 94 | -0.452216 | -1.20953 | 0.170186 | 0.481676 |
| GOBP_RESPONSE_TO_PEPTIDE | 385 | -0.400629 | -1.208013 | 0.091102 | 0.484797 |
| GOBP_POSITIVE_REGULATION_OF_CELLULAR_PROTEIN_LOCALIZATION | 254 | -0.412533 | -1.207735 | 0.126257 | 0.485013 |
| GOBP_REGULATION_OF_PHOSPHATIDYLINOSITOL_3_KINASE_ACTIVITY | 45 | -0.504017 | -1.206404 | 0.219274 | 0.487774 |
| GOBP_NEGATIVE_REGULATION_OF_INTERLEUKIN_10_PRODUCTION | 15 | -0.585289 | -1.206323 | 0.272583 | 0.487517 |
| GOBP_MESENCHYMAL_CELL_DIFFERENTIATION | 151 | -0.427861 | -1.206201 | 0.150602 | 0.487381 |
| GOBP_ACTOMYOSIN_STRUCTURE_ORGANIZATION | 142 | -0.428184 | -1.20528 | 0.179487 | 0.489162 |
| GOBP_NEGATIVE_REGULATION_OF_EXTRINSIC_APOPTOTIC_SIGNALING_PATHW | 75 | -0.463112 | -1.204761 | 0.189987 | 0.489984 |
| GOBP_HISTONE_H3_K9_METHYLATION | 29 | -0.523688 | -1.204575 | 0.234493 | 0.489997 |
| GOBP_REGULATION_OF_ANDROGEN_RECEPTOR_SIGNALING_PATHWAY | 21 | -0.549511 | -1.204174 | 0.229167 | 0.490488 |
| GOBP_ORGANOPHOSPHATE_CATABOLIC_PROCESS | 121 | -0.435902 | -1.204153 | 0.180025 | 0.490084 |
| GOBP_REGULATION_OF_PHOSPHATIDYLINOSITOL_3_KINASE_SIGNALING | 89 | -0.448802 | -1.203054 | 0.185987 | 0.492217 |
| GOBP_GLYCEROL_ETHER_METABOLIC_PROCESS | 19 | -0.554288 | -1.202687 | 0.257009 | 0.492686 |
| GOBP_SINGLE_STRANDED_VIRAL_RNA_REPLICATION_VIA_DOUBLE_STRANDED | 18 | -0.570484 | -1.202208 | 0.248387 | 0.493326 |
| GOBP_REGULATION_OF_EPITHELIAL_TO_MESENCHYMAL_TRANSITION | 69 | -0.471783 | -1.201996 | 0.196053 | 0.493414 |
| GOBP_ANION_HOMEOSTASIS | 18 | -0.56497 | -1.201846 | 0.245696 | 0.493351 |
| GOBP_NEGATIVE_REGULATION_OF_INTERFERON_GAMMA_PRODUCTION | 29 | -0.525952 | -1.201792 | 0.239394 | 0.493027 |
| GOBP_POSITIVE_REGULATION_OF_CALCIUM_ION_TRANSPORT | 72 | -0.46462 | -1.201699 | 0.200271 | 0.492825 |
| GOBP_RESPONSE_TO_TRANSFORMING_GROWTH_FACTOR_BETA | 181 | -0.417653 | -1.200257 | 0.159437 | 0.495901 |
| GOBP_NEGATIVE_REGULATION_OF_CYSTEINE_TYPE_ENDOPEPTIDASE_ACTIVIT | 62 | -0.470219 | -1.200004 | 0.217332 | 0.496043 |
| GOBP_RESPONSE_TO_OSMOTIC_STRESS | 63 | -0.46981 | -1.19972 | 0.21751 | 0.496303 |
| GOBP_MESONEPHROS_DEVELOPMENT | 44 | -0.49688 | -1.199596 | 0.210294 | 0.496135 |
| GOBP_NEGATIVE_REGULATION_OF_TRANSFORMING_GROWTH_FACTOR_BETA_ | 62 | -0.470108 | -1.199573 | 0.204082 | 0.495746 |
| GOBP_REGULATION_OF_CALCINEURIN_MEDIATED_SIGNALING | 24 | -0.539034 | -1.198578 | 0.257716 | 0.497646 |
| GOBP_NEGATIVE_REGULATION_OF_INTERLEUKIN_6_PRODUCTION | 33 | -0.501643 | -1.198504 | 0.236256 | 0.497374 |
| GOBP_NEGATIVE_REGULATION_OF_PEPTIDE_HORMONE_SECRETION | 24 | -0.548041 | -1.197747 | 0.261398 | 0.498739 |
| GOBP_REGULATION_OF_CALCIUM_ION_TRANSMEMBRANE_TRANSPORT | 91 | -0.448755 | -1.197349 | 0.201602 | 0.499285 |
| GOBP_POSITIVE_REGULATION_OF_PROTEIN_KINASE_ACTIVITY | 401 | -0.400039 | -1.197331 | 0.108974 | 0.498884 |
| GOBP_SODIUM_ION_TRANSMEMBRANE_TRANSPORT | 99 | -0.447686 | -1.196944 | 0.211854 | 0.499367 |
| GOBP_CELLULAR_RESPONSE_TO_NITROGEN_COMPOUND | 486 | -0.394652 | -1.196704 | 0.099052 | 0.499514 |
| GOBP_GLYCOPROTEIN_BIOSYNTHETIC_PROCESS | 252 | -0.407682 | -1.196468 | 0.143973 | 0.499617 |
| GOBP_BONE_RESORPTION | 46 | -0.496185 | -1.196435 | 0.228883 | 0.499256 |
| GOBP_REGULATION_OF_RECEPTOR_LOCALIZATION_TO_SYNAPSE | 15 | -0.591022 | -1.196203 | 0.264559 | 0.499339 |
| GOBP_RESPONSE_TO_ISCHEMIA | 37 | -0.501651 | -1.196117 | 0.239163 | 0.499109 |
| GOBP_RELEASE_OF_SEQUESTERED_CALCIUM_ION_INTO_CYTOSOL_BY_ENDOI | 22 | -0.548714 | -1.196095 | 0.26183 | 0.498723 |
| GOBP_HINDBRAIN_DEVELOPMENT | 85 | -0.458483 | -1.195849 | 0.192513 | 0.498895 |
| GOBP_RESPONSE_TO_DRUG | 246 | -0.410333 | -1.194996 | 0.129425 | 0.500449 |
| GOBP_CELL_CELL_ADHESION_VIA_PLASMA_MEMBRANE_ADHESION_MOLECUL | 125 | -0.429025 | -1.194821 | 0.177778 | 0.500405 |
| GOBP_SYNAPTIC_VESICLE_RECYCLING | 56 | -0.482011 | -1.194626 | 0.226573 | 0.500428 |
| GOBP_RESPONSE_TO_FIBROBLAST_GROWTH_FACTOR | 91 | -0.441052 | -1.193903 | 0.193122 | 0.50175 |
| GOBP_NEGATIVE_REGULATION_OF_SUPRAMOLECULAR_FIBER_ORGANIZATION | 118 | -0.433115 | -1.193593 | 0.192786 | 0.502039 |
| GOBP_POSITIVE_REGULATION_OF_INTERLEUKIN_12_PRODUCTION | 32 | -0.525025 | -1.193537 | 0.243939 | 0.501721 |
| GOBP_ORGANOPHOSPHATE_ESTER_TRANSPORT | 101 | -0.438541 | -1.193371 | 0.18285 | 0.501673 |
| GOBP_SOMATIC_RECOMBINATION_OF_IMMUNOGLOBULIN_GENE_SEGMENTS | 52 | -0.474654 | -1.193133 | 0.211172 | 0.501785 |
| GOBP_POSITIVE_REGULATION_OF_CALCIUM_ION_TRANSPORT_INTO_CYTOSOL | 26 | -0.525944 | -1.192602 | 0.25 | 0.502603 |
| GOBP_CELL_CYCLE_G1_S_PHASE_TRANSITION | 241 | -0.403145 | -1.191374 | 0.130533 | 0.504968 |
| GOBP_TRIGLYCERIDE_METABOLIC_PROCESS | 65 | -0.459635 | -1.191225 | 0.218407 | 0.504912 |
| GOBP_INOSITOL_PHOSPHATE_MEDIATED_SIGNALING | 40 | -0.502899 | -1.191021 | 0.234133 | 0.504993 |
| GOBP_REGULATION_OF_CYTOPLASMIC_TRANSPORT | 25 | -0.534062 | -1.19049 | 0.261346 | 0.505779 |
| GOBP_REGULATION_OF_NMDA_RECEPTOR_ACTIVITY | 23 | -0.546138 | -1.190435 | 0.258165 | 0.505455 |
| GOBP_NEGATIVE_REGULATION_OF_CELLULAR_AMIDE_METABOLIC_PROCESS | 147 | -0.420599 | -1.190271 | 0.179302 | 0.505367 |
| GOBP_NITRIC_OXIDE_SYNTHASE_BIOSYNTHETIC_PROCESS | 16 | -0.584204 | -1.190256 | 0.291181 | 0.50496 |
| GOBP_NATURAL_KILLER_CELL_ACTIVATION | 57 | -0.474842 | -1.189467 | 0.234568 | 0.506459 |
| GOBP_MESENCHYME_DEVELOPMENT | 175 | -0.417602 | -1.18915 | 0.177246 | 0.50678 |
| GOBP_NEGATIVE_REGULATION_OF_LEUKOCYTE_APOPTOTIC_PROCESS | 37 | -0.495575 | -1.189115 | 0.237814 | 0.506414 |
| GOBP_TYPE_2_IMMUNE_RESPONSE | 27 | -0.527615 | -1.188878 | 0.250386 | 0.506527 |
| GOBP_CARDIAC_MUSCLE_CELL_ACTION_POTENTIAL | 37 | -0.506475 | -1.187941 | 0.224432 | 0.50834 |
| GOBP_REGULATION_OF_B_CELL_APOPTOTIC_PROCESS | 18 | -0.571617 | -1.187785 | 0.272583 | 0.508249 |
| GOBP_B_CELL_PROLIFERATION | 73 | -0.452337 | -1.187368 | 0.214929 | 0.508797 |
| GOBP_INTERLEUKIN_1_MEDIATED_SIGNALING_PATHWAY | 94 | -0.444069 | -1.187329 | 0.211613 | 0.508462 |
| GOBP_REGULATION_OF_CELL_DEVELOPMENT | 328 | -0.398009 | -1.187114 | 0.133909 | 0.508556 |
| GOBP_POSITIVE_REGULATION_OF_PROTEIN_DEPHOSPHORYLATION | 40 | -0.501325 | -1.186464 | 0.229607 | 0.50969 |
| GOBP_TRANSEPITHELIAL_TRANSPORT | 18 | -0.566113 | -1.186345 | 0.263554 | 0.509503 |
| GOBP_EMBRYO_DEVELOPMENT_ENDING_IN_BIRTH_OR_EGG_HATCHING | 418 | -0.392425 | -1.186299 | 0.116081 | 0.50917 |
| GOBP_REGULATION_OF_CELL_PROJECTION_ASSEMBLY | 149 | -0.42212 | -1.186291 | 0.176116 | 0.508746 |
| GOBP_RESPONSE_TO_ANGIOTENSIN | 17 | -0.565963 | -1.186048 | 0.266116 | 0.508858 |
| GOBP_VENTRICULAR_CARDIAC_MUSCLE_CELL_ACTION_POTENTIAL | 15 | -0.582797 | -1.185976 | 0.294309 | 0.508586 |
| GOBP_PROTEIN_PROCESSING | 153 | -0.417217 | -1.185953 | 0.189964 | 0.508211 |
| GOBP_KIDNEY_EPITHELIUM_DEVELOPMENT | 72 | -0.457422 | -1.185368 | 0.222964 | 0.509136 |
| GOBP_NEGATIVE_REGULATION_OF_RECEPTOR_MEDIATED_ENDOCYTOSIS | 22 | -0.544158 | -1.185254 | 0.27673 | 0.508988 |
| GOBP_POSITIVE_REGULATION_OF_TELOMERASE_ACTIVITY | 31 | -0.51786 | -1.184548 | 0.235205 | 0.510262 |
| GOBP_BRANCHING_INVOLVED_IN_BLOOD_VESSEL_MORPHOGENESIS | 20 | -0.548461 | -1.183881 | 0.289833 | 0.511413 |
| GOBP_REGULATION_OF_MONOOXYGENASE_ACTIVITY | 46 | -0.484696 | -1.183583 | 0.24212 | 0.511689 |
| GOBP_POSITIVE_REGULATION_OF_NOTCH_SIGNALING_PATHWAY | 42 | -0.488395 | -1.182932 | 0.237482 | 0.51277 |
| GOBP_DETECTION_OF_ABIOTIC_STIMULUS | 74 | -0.457227 | -1.182817 | 0.195448 | 0.512611 |
| GOBP_TEMPERATURE_HOMEOSTASIS | 122 | -0.430312 | -1.182405 | 0.203202 | 0.513129 |
| GOBP_OSSIFICATION | 254 | -0.402715 | -1.182166 | 0.148984 | 0.513309 |
| GOBP_POSITIVE_REGULATION_OF_EPIDERMAL_CELL_DIFFERENTIATION | 18 | -0.558965 | -1.181832 | 0.274959 | 0.513669 |
| GOBP_RENAL_SYSTEM_PROCESS | 66 | -0.466588 | -1.181461 | 0.218421 | 0.514137 |
| GOBP_RESPONSE_TO_KETONE | 129 | -0.423018 | -1.18111 | 0.202166 | 0.514562 |
| GOBP_RESPONSE_TO_CAMP | 60 | -0.469927 | -1.18033 | 0.231176 | 0.515959 |
| GOBP_ACTIN_MEDIATED_CELL_CONTRACTION | 62 | -0.463503 | -1.180187 | 0.238095 | 0.515849 |
| GOBP_HYDROGEN_PEROXIDE_CATABOLIC_PROCESS | 16 | -0.576597 | -1.179709 | 0.289557 | 0.516527 |
| GOBP_MULTI_MULTICELLULAR_ORGANISM_PROCESS | 124 | -0.426697 | -1.179443 | 0.2 | 0.516734 |
| GOBP_SENSORY_ORGAN_DEVELOPMENT | 305 | -0.398112 | -1.178423 | 0.152747 | 0.518784 |
| GOBP_REGULATION_OF_HETEROTYPIC_CELL_CELL_ADHESION | 17 | -0.569347 | -1.176852 | 0.276083 | 0.522225 |
| GOBP_NEGATIVE_REGULATION_OF_HORMONE_SECRETION | 36 | -0.502764 | -1.176652 | 0.259146 | 0.522277 |

| | | | | | |
|---|---|---|---|---|---|
| GOBP_REGULATION_OF_DIGESTIVE_SYSTEM_PROCESS | 18 | -0.561018 | -1.17637 | 0.279435 | 0.522532 |
| GOBP_PHOSPHATIDIC_ACID_METABOLIC_PROCESS | 37 | -0.504604 | -1.176233 | 0.270802 | 0.522405 |
| GOBP_RESPONSE_TO_STEROID_HORMONE | 223 | -0.403119 | -1.175913 | 0.165541 | 0.522736 |
| GOBP_EMBRYONIC_MORPHOGENESIS | 349 | -0.390812 | -1.175785 | 0.148268 | 0.522604 |
| GOBP_REGULATION_OF_NEUROTRANSMITTER_RECEPTOR_ACTIVITY | 45 | -0.478127 | -1.1757 | 0.252137 | 0.522392 |
| GOBP_CELLULAR_RESPONSE_TO_PEPTIDE | 293 | -0.396187 | -1.175133 | 0.154185 | 0.523338 |
| GOBP_NEGATIVE_REGULATION_OF_SIGNALING_RECEPTOR_ACTIVITY | 32 | -0.516705 | -1.174927 | 0.25268 | 0.523393 |
| GOBP_COLLAGEN_FIBRIL_ORGANIZATION | 32 | -0.51447 | -1.174789 | 0.254844 | 0.523272 |
| GOBP_EMBRYONIC_ORGAN_DEVELOPMENT | 245 | -0.401094 | -1.174644 | 0.167592 | 0.523191 |
| GOBP_NEGATIVE_REGULATION_OF_COAGULATION | 29 | -0.51935 | -1.174585 | 0.289355 | 0.522901 |
| GOBP_C21_STEROID_HORMONE_METABOLIC_PROCESS | 21 | -0.543379 | -1.17453 | 0.283931 | 0.52258 |
| GOBP_POSITIVE_REGULATION_OF_PROTEIN_TYROSINE_KINASE_ACTIVITY | 38 | -0.493871 | -1.174244 | 0.246617 | 0.522779 |
| GOBP_REGULATION_OF_MEGAKARYOCYTE_DIFFERENTIATION | 63 | -0.45936 | -1.174157 | 0.222535 | 0.522565 |
| GOBP_POSITIVE_REGULATION_OF_GROWTH | 172 | -0.414113 | -1.173019 | 0.190083 | 0.524824 |
| GOBP_FOREBRAIN_GENERATION_OF_NEURONS | 23 | -0.531452 | -1.172985 | 0.287781 | 0.524462 |
| GOBP_REGULATION_OF_CELLULAR_RESPIRATION | 18 | -0.559867 | -1.172927 | 0.29717 | 0.524183 |
| GOBP_DNA_METHYLATION_OR_DEMETHYLATION | 73 | -0.447603 | -1.17248 | 0.212283 | 0.524853 |
| GOBP_TRANSCRIPTION_BY_RNA_POLYMERASE_III | 46 | -0.477951 | -1.172013 | 0.249643 | 0.525573 |
| GOBP_REGULATION_OF_GROWTH | 457 | -0.385279 | -1.171819 | 0.112288 | 0.525589 |
| GOBP_CALCIUM_ION_TRANSMEMBRANE_TRANSPORT | 197 | -0.406368 | -1.171399 | 0.175644 | 0.526118 |
| GOBP_STEROL_METABOLIC_PROCESS | 120 | -0.425746 | -1.171132 | 0.214461 | 0.526289 |
| GOBP_ETHER_METABOLIC_PROCESS | 23 | -0.538629 | -1.170897 | 0.28439 | 0.526406 |
| GOBP_SENSORY_SYSTEM_DEVELOPMENT | 214 | -0.402284 | -1.170344 | 0.168743 | 0.527386 |
| GOBP_POSITIVE_REGULATION_OF_INTERLEUKIN_6_PRODUCTION | 68 | -0.4598 | -1.170008 | 0.229282 | 0.527738 |
| GOBP_POSITIVE_REGULATION_OF_CELL_DEVELOPMENT | 210 | -0.403414 | -1.169918 | 0.204833 | 0.527525 |
| GOBP_HEART_PROCESS | 158 | -0.409779 | -1.169592 | 0.207776 | 0.52785 |
| GOBP_DORSAL_VENTRAL_PATTERN_FORMATION | 33 | -0.493989 | -1.168705 | 0.27786 | 0.529584 |
| GOBP_POSITIVE_REGULATION_OF_NUCLEAR_DIVISION | 39 | -0.490254 | -1.168412 | 0.241774 | 0.529843 |
| GOBP_REGULATION_OF_RESPONSE_TO_CYTOKINE_STIMULUS | 148 | -0.417595 | -1.167694 | 0.193587 | 0.531191 |
| GOBP_PERIPHERAL_NERVOUS_SYSTEM_DEVELOPMENT | 50 | -0.470732 | -1.167325 | 0.257022 | 0.531599 |
| GOBP_REGULATION_OF_MITOTIC_NUCLEAR_DIVISION | 90 | -0.440929 | -1.167162 | 0.260483 | 0.5315 |
| GOBP_RECEPTOR_CLUSTERING | 40 | -0.492712 | -1.166905 | 0.268477 | 0.531699 |
| GOBP_NEGATIVE_REGULATION_OF_EPITHELIAL_CELL_DIFFERENTIATION | 22 | -0.542419 | -1.166625 | 0.283048 | 0.531924 |
| GOBP_REGULATION_OF_SODIUM_ION_TRANSPORT | 58 | -0.464185 | -1.165537 | 0.242466 | 0.534194 |
| GOBP_SEQUESTERING_OF_CALCIUM_ION | 77 | -0.436538 | -1.164806 | 0.250328 | 0.535485 |
| GOBP_REGULATION_OF_BLOOD_CIRCULATION | 152 | -0.414276 | -1.1643 | 0.215733 | 0.53625 |
| GOBP_ACTIVATED_T_CELL_PROLIFERATION | 33 | -0.508363 | -1.163952 | 0.277126 | 0.536655 |
| GOBP_POSITIVE_REGULATION_OF_AMINE_TRANSPORT | 19 | -0.548933 | -1.163356 | 0.284091 | 0.537653 |
| GOBP_PLATELET_DERIVED_GROWTH_FACTOR_RECEPTOR_SIGNALING_PATHW | 47 | -0.473501 | -1.163257 | 0.25788 | 0.537475 |
| GOBP_SARCOPLASMIC_RETICULUM_CALCIUM_ION_TRANSPORT | 25 | -0.526235 | -1.163033 | 0.273148 | 0.537575 |
| GOBP_UNSATURATED_FATTY_ACID_METABOLIC_PROCESS | 68 | -0.45112 | -1.162802 | 0.214481 | 0.537682 |
| GOBP_RENAL_TUBULE_DEVELOPMENT | 48 | -0.481255 | -1.162307 | 0.264664 | 0.538444 |
| GOBP_HAIR_CELL_DIFFERENTIATION | 26 | -0.515929 | -1.162069 | 0.302795 | 0.53857 |
| GOBP_TOLL_LIKE_RECEPTOR_4_SIGNALING_PATHWAY | 31 | -0.501977 | -1.161885 | 0.26209 | 0.538554 |
| GOBP_REGULATION_OF_VESICLE_FUSION | 17 | -0.556819 | -1.161706 | 0.282927 | 0.538515 |
| GOBP_INTERLEUKIN_1_BETA_PRODUCTION | 73 | -0.443773 | -1.161259 | 0.230345 | 0.5392 |
| GOBP_ADENYLATE_CYCLASE_MODULATING_G_PROTEIN_COUPLED_RECEPTOF | 105 | -0.42494 | -1.160399 | 0.21883 | 0.540854 |
| GOBP_INTRINSIC_APOPTOTIC_SIGNALING_PATHWAY | 249 | -0.392685 | -1.160271 | 0.189189 | 0.540712 |
| GOBP_BARBED_END_ACTIN_FILAMENT_CAPPING | 20 | -0.539108 | -1.160148 | 0.29595 | 0.540561 |
| GOBP_REGULATION_OF_CELL_AGING | 42 | -0.483673 | -1.159967 | 0.271552 | 0.540572 |
| GOBP_LYMPHOID_PROGENITOR_CELL_DIFFERENTIATION | 17 | -0.556245 | -1.159913 | 0.283892 | 0.540264 |
| GOBP_EXOCRINE_SYSTEM_DEVELOPMENT | 23 | -0.531448 | -1.158943 | 0.288316 | 0.542128 |
| GOBP_REGULATION_OF_SECRETION | 406 | -0.378891 | -1.158603 | 0.150106 | 0.54247 |
| GOBP_DENDRITIC_SPINE_MORPHOGENESIS | 44 | -0.474485 | -1.157674 | 0.272859 | 0.544238 |
| GOBP_REGULATION_OF_EXTRINSIC_APOPTOTIC_SIGNALING_PATHWAY | 113 | -0.424713 | -1.15742 | 0.229193 | 0.544427 |
| GOBP_HOMOTYPIC_CELL_CELL_ADHESION | 63 | -0.457448 | -1.157314 | 0.244709 | 0.544238 |
| GOBP_INTERLEUKIN_10_PRODUCTION | 46 | -0.470658 | -1.157159 | 0.26204 | 0.544188 |
| GOBP_REGULATION_OF_NUCLEOTIDE_BIOSYNTHETIC_PROCESS | 30 | -0.506988 | -1.156856 | 0.269865 | 0.544497 |
| GOBP_PRESYNAPTIC_ENDOCYTOSIS | 47 | -0.4837 | -1.156552 | 0.280802 | 0.544797 |
| GOBP_REGULATION_OF_MYELINATION | 30 | -0.504509 | -1.155639 | 0.267356 | 0.546661 |
| GOBP_POSITIVE_REGULATION_OF_CELLULAR_PROTEIN_CATABOLIC_PROCESS | 133 | -0.417789 | -1.155087 | 0.218335 | 0.547541 |
| GOBP_CARDIAC_MUSCLE_CELL_ACTION_POTENTIAL_INVOLVED_IN_CONTRACT | 23 | -0.530192 | -1.155033 | 0.302508 | 0.547222 |
| GOBP_ELECTRON_TRANSPORT_CHAIN | 159 | -0.409093 | -1.154047 | 0.214022 | 0.549274 |
| GOBP_CELLULAR_RESPONSE_TO_UV | 78 | -0.43942 | -1.15323 | 0.249351 | 0.550871 |
| GOBP_POSITIVE_REGULATION_OF_COLD_INDUCED_THERMOGENESIS | 67 | -0.448632 | -1.153178 | 0.245232 | 0.550549 |
| GOBP_MUSCLE_CELL_CELLULAR_HOMEOSTASIS | 17 | -0.556529 | -1.152748 | 0.288889 | 0.551146 |
| GOBP_REGULATION_OF_CELLULAR_PROTEIN_LOCALIZATION | 463 | -0.37854 | -1.152655 | 0.157173 | 0.550922 |
| GOBP_DNA_DEMETHYLATION | 25 | -0.515109 | -1.152478 | 0.292188 | 0.55088 |
| GOBP_RESPONSE_TO_PROTOZOAN | 19 | -0.543119 | -1.151687 | 0.33441 | 0.552372 |
| GOBP_VASCULAR_ENDOTHELIAL_GROWTH_FACTOR_RECEPTOR_SIGNALING_P | 75 | -0.437625 | -1.151027 | 0.236807 | 0.55355 |
| GOBP_POSITIVE_REGULATION_OF_LAMELLIPODIUM_ORGANIZATION | 33 | -0.489287 | -1.150939 | 0.286525 | 0.553316 |
| GOBP_REGULATION_OF_ORGANIC_ACID_TRANSPORT | 36 | -0.484467 | -1.150927 | 0.290043 | 0.552904 |
| GOBP_GLYCOSYLATION | 209 | -0.398467 | -1.150066 | 0.198157 | 0.5546 |
| GOBP_NEGATIVE_REGULATION_OF_CELLULAR_PROTEIN_LOCALIZATION | 91 | -0.434406 | -1.150017 | 0.265092 | 0.554293 |
| GOBP_RESPONSE_TO_ETHANOL | 81 | -0.436771 | -1.149572 | 0.269133 | 0.55498 |
| GOBP_ARTERY_MORPHOGENESIS | 38 | -0.492583 | -1.149369 | 0.292945 | 0.555032 |
| GOBP_PRIMARY_ALCOHOL_METABOLIC_PROCESS | 55 | -0.458806 | -1.148793 | 0.264336 | 0.55601 |
| GOBP_CELLULAR_RESPONSE_TO_EXOGENOUS_DSRNA | 15 | -0.558024 | -1.14747 | 0.330632 | 0.558779 |
| GOBP_POSITIVE_REGULATION_OF_PROTEIN_TARGETING_TO_MITOCHONDRION | 30 | -0.499284 | -1.147339 | 0.275811 | 0.558639 |
| GOBP_CIRCADIAN_RHYTHM | 155 | -0.402927 | -1.146672 | 0.218225 | 0.559817 |
| GOBP_REGULATION_OF_ANOIKIS | 20 | -0.530005 | -1.145966 | 0.319805 | 0.561089 |
| GOBP_SMOOTH_MUSCLE_CELL_DIFFERENTIATION | 38 | -0.484395 | -1.145885 | 0.282051 | 0.560829 |
| GOBP_REGULATION_OF_NOTCH_SIGNALING_PATHWAY | 78 | -0.442195 | -1.14588 | 0.2711 | 0.560395 |
| GOBP_NEPHRON_DEVELOPMENT | 81 | -0.429909 | -1.145172 | 0.249664 | 0.561669 |
| GOBP_NEGATIVE_REGULATION_OF_VIRAL_ENTRY_INTO_HOST_CELL | 15 | -0.549485 | -1.144585 | 0.323383 | 0.562711 |
| GOBP_CAMERA_TYPE_EYE_DEVELOPMENT | 181 | -0.394718 | -1.144084 | 0.224199 | 0.563473 |
| GOBP_NEGATIVE_REGULATION_OF_MAP_KINASE_ACTIVITY | 59 | -0.453433 | -1.143966 | 0.274064 | 0.563311 |
| GOBP_POSITIVE_REGULATION_OF_SUBSTRATE_ADHESION_DEPENDENT_CELL_ | 34 | -0.489171 | -1.143306 | 0.282014 | 0.56447 |
| GOBP_POSITIVE_REGULATION_OF_REGULATED_SECRETORY_PATHWAY | 37 | -0.491048 | -1.143262 | 0.293255 | 0.56413 |
| GOBP_RESPONSE_TO_EXTRACELLULAR_STIMULUS | 342 | -0.37931 | -1.143242 | 0.194384 | 0.563723 |
| GOBP_REGULATION_OF_ENDOTHELIAL_CELL_MIGRATION | 121 | -0.414831 | -1.143145 | 0.240338 | 0.563508 |
| GOBP_DOPAMINE_METABOLIC_PROCESS | 18 | -0.542161 | -1.142954 | 0.308307 | 0.563524 |
| GOBP_CHLORIDE_TRANSPORT | 58 | -0.453416 | -1.142745 | 0.28273 | 0.563568 |
| GOBP_REGULATION_OF_HOMOTYPIC_CELL_CELL_ADHESION | 23 | -0.524539 | -1.142702 | 0.322884 | 0.56323 |
| GOBP_REGULATION_OF_REGULATED_SECRETORY_PATHWAY | 97 | -0.422303 | -1.142555 | 0.248684 | 0.563168 |
| GOBP_REGULATION_OF_PHOSPHOLIPASE_C_ACTIVITY | 30 | -0.505803 | -1.142504 | 0.28852 | 0.562862 |
| GOBP_INNER_EAR_AUDITORY_RECEPTOR_CELL_DIFFERENTIATION | 23 | -0.525999 | -1.14245 | 0.318885 | 0.562553 |
| GOBP_B_CELL_MEDIATED_IMMUNITY | 136 | -0.410715 | -1.142338 | 0.244772 | 0.56241 |
| GOBP_SYNAPTIC_SIGNALING | 410 | -0.378138 | -1.141889 | 0.178155 | 0.563055 |
| GOBP_N_GLYCAN_PROCESSING | 17 | -0.562859 | -1.141694 | 0.317152 | 0.563123 |
| GOBP_RESPONSE_TO_PROSTAGLANDIN | 22 | -0.535268 | -1.141517 | 0.314729 | 0.563138 |
| GOBP_REGULATION_OF_PHOSPHATASE_ACTIVITY | 139 | -0.409011 | -1.141027 | 0.252774 | 0.563909 |
| GOBP_LIVER_REGENERATION | 23 | -0.519463 | -1.14085 | 0.300813 | 0.563888 |
| GOBP_RESPONSE_TO_VITAMIN_D | 17 | -0.53691 | -1.140629 | 0.315133 | 0.56399 |
| GOBP_MYOBLAST_FUSION | 24 | -0.517413 | -1.140513 | 0.312989 | 0.563818 |
| GOBP_NEGATIVE_REGULATION_OF_CELL_MATRIX_ADHESION | 27 | -0.508939 | -1.140475 | 0.298742 | 0.563464 |
| GOBP_DNA_REPLICATION_INITIATION | 39 | -0.475133 | -1.138625 | 0.302391 | 0.567658 |
| GOBP_REGULATION_OF_TRANSCRIPTION_INITIATION_FROM_RNA_POLYMERASI | 31 | -0.483981 | -1.138471 | 0.311728 | 0.567571 |
| GOBP_CELLULAR_PROTEIN_CONTAINING_COMPLEX_LOCALIZATION | 19 | -0.532052 | -1.138437 | 0.318108 | 0.567201 |
| GOBP_REGULATION_OF_B_CELL_PROLIFERATION | 54 | -0.451987 | -1.137845 | 0.277393 | 0.568215 |
| GOBP_DEFENSE_RESPONSE_TO_GRAM_NEGATIVE_BACTERIUM | 37 | -0.471145 | -1.137673 | 0.28655 | 0.568223 |
| GOBP_COGNITION | 170 | -0.391971 | -1.137583 | 0.237126 | 0.567993 |
| GOBP_FIBROBLAST_PROLIFERATION | 60 | -0.448917 | -1.137116 | 0.26614 | 0.568743 |
| GOBP_METAL_ION_EXPORT | 26 | -0.51601 | -1.136656 | 0.314417 | 0.569435 |
| GOBP_REGULATION_OF_CELL_MORPHOGENESIS_INVOLVED_IN_DIFFERENTIATI | 75 | -0.428825 | -1.136398 | 0.264276 | 0.569628 |
| GOBP_REGULATION_OF_CELLULAR_AMINE_METABOLIC_PROCESS | 66 | -0.440608 | -1.136352 | 0.288435 | 0.569294 |
| GOBP_REGULATION_OF_TRANS_SYNAPTIC_SIGNALING | 258 | -0.385106 | -1.13618 | 0.229834 | 0.569254 |
| GOBP_RESPONSE_TO_TEMPERATURE_STIMULUS | 180 | -0.397022 | -1.13609 | 0.224797 | 0.569031 |
| GOBP_DNA_TEMPLATED_TRANSCRIPTION_INITIATION | 208 | -0.393346 | -1.135892 | 0.229714 | 0.569088 |
| GOBP_ACTIN_FILAMENT_BASED_MOVEMENT | 88 | -0.431277 | -1.135809 | 0.275689 | 0.568841 |
| GOBP_GLYCOSPHINGOLIPID_BIOSYNTHETIC_PROCESS | 20 | -0.535295 | -1.135783 | 0.328526 | 0.568472 |
| GOBP_IMPORT_INTO_CELL | 148 | -0.401321 | -1.135641 | 0.241587 | 0.568378 |
| GOBP_POSITIVE_REGULATION_OF_GLYCOLYTIC_PROCESS | 15 | -0.559464 | -1.135476 | 0.3408 | 0.568344 |
| GOBP_REGULATION_OF_LYMPHOCYTE_APOPTOTIC_PROCESS | 46 | -0.46272 | -1.134852 | 0.296671 | 0.569435 |
| GOBP_VESICLE_MEDIATED_TRANSPORT_IN_SYNAPSE | 135 | -0.403437 | -1.134105 | 0.264339 | 0.570852 |
| GOBP_LIPID_CATABOLIC_PROCESS | 233 | -0.391194 | -1.134066 | 0.230944 | 0.570515 |
| GOBP_REGULATION_OF_CALCIUM_ION_TRANSPORT_INTO_CYTOSOL | 55 | -0.449485 | -1.133973 | 0.277174 | 0.570317 |
| GOBP_REGULATION_OF_ACUTE_INFLAMMATORY_RESPONSE | 29 | -0.506298 | -1.133839 | 0.313869 | 0.57022 |
| GOBP_REGULATION_OF_SUBSTRATE_ADHESION_DEPENDENT_CELL_SPREADII | 45 | -0.459043 | -1.133827 | 0.299291 | 0.569805 |
| GOBP_SYNAPTIC_VESICLE_LOCALIZATION | 43 | -0.468015 | -1.133613 | 0.296512 | 0.569911 |
| GOBP_PEPTIDE_HORMONE_SECRETION | 166 | -0.391499 | -1.133199 | 0.242823 | 0.570451 |
| GOBP_DEVELOPMENTAL_PIGMENTATION | 31 | -0.491774 | -1.133156 | 0.286567 | 0.570115 |
| GOBP_NEGATIVE_REGULATION_OF_ERK1_AND_ERK2_CASCADE | 56 | -0.453868 | -1.132937 | 0.285124 | 0.570211 |
| GOBP_POSITIVE_REGULATION_OF_RELEASE_OF_SEQUESTERED_CALCIUM_ION | 20 | -0.536001 | -1.13249 | 0.336923 | 0.570859 |
| GOBP_RESPONSE_TO_GROWTH_HORMONE | 27 | -0.502472 | -1.1324 | 0.333333 | 0.570648 |
| GOBP_POSITIVE_REGULATION_OF_PROTEIN_AUTOPHOSPHORYLATION | 23 | -0.512405 | -1.13201 | 0.327586 | 0.571196 |
| GOBP_ICOSANOID_METABOLIC_PROCESS | 67 | -0.439778 | -1.131953 | 0.305365 | 0.57091 |
| GOBP_HEPATICOBILIARY_SYSTEM_DEVELOPMENT | 105 | -0.411378 | -1.131592 | 0.281602 | 0.571312 |
| GOBP_MYELOID_CELL_HOMEOSTASIS | 127 | -0.409302 | -1.131583 | 0.247273 | 0.570898 |
| GOBP_REGULATION_OF_CELLULAR_RESPONSE_TO_TRANSFORMING_GROWTH | 96 | -0.417093 | -1.131545 | 0.266922 | 0.570553 |
| GOBP_REGULATION_OF_ATP_METABOLIC_PROCESS | 98 | -0.419055 | -1.130805 | 0.271808 | 0.571971 |
| GOBP_MUSCLE_CONTRACTION | 201 | -0.390766 | -1.130729 | 0.23322 | 0.571731 |
| GOBP_WALKING_BEHAVIOR | 15 | -0.54778 | -1.130657 | 0.341256 | 0.57146 |
| GOBP_HEART_MORPHOGENESIS | 130 | -0.4047 | -1.130448 | 0.27439 | 0.571538 |
| GOBP_POLYOL_BIOSYNTHETIC_PROCESS | 49 | -0.456068 | -1.130236 | 0.296703 | 0.571613 |
| GOBP_NEGATIVE_REGULATION_OF_TRANSFERASE_ACTIVITY | 250 | -0.388665 | -1.13022 | 0.230769 | 0.571212 |
| GOBP_RESPONSE_TO_PROSTAGLANDIN_E | 19 | -0.537648 | -1.130197 | 0.333868 | 0.570824 |

| | | | | | |
|---|---|---|---|---|---|
| GOBP_NEGATIVE_REGULATION_OF_ANION_TRANSMEMBRANE_TRANSPORT | 26 | -0.496792 | -1.130162 | 0.322485 | 0.570462 |
| GOBP_POSITIVE_REGULATION_OF_APOPTOTIC_SIGNALING_PATHWAY | 103 | -0.412993 | -1.130103 | 0.256082 | 0.570164 |
| GOBP_DOPAMINE_TRANSPORT | 20 | -0.517759 | -1.12927 | 0.340557 | 0.571816 |
| GOBP_NEGATIVE_REGULATION_OF_ERBB_SIGNALING_PATHWAY | 41 | -0.48034 | -1.129238 | 0.288762 | 0.571458 |
| GOBP_CARDIAC_MUSCLE_CONTRACTION | 73 | -0.433675 | -1.12893 | 0.292225 | 0.571815 |
| GOBP_REGULATION_OF_MEMBRANE_POTENTIAL | 248 | -0.383581 | -1.128851 | 0.241611 | 0.571566 |
| GOBP_STEROID_METABOLIC_PROCESS | 212 | -0.388003 | -1.12813 | 0.223516 | 0.572893 |
| GOBP_NEURAL_CREST_CELL_DIFFERENTIATION | 54 | -0.453832 | -1.127488 | 0.296454 | 0.574032 |
| GOBP_INTRINSIC_APOPTOTIC_SIGNALING_PATHWAY_IN_RESPONSE_TO_OXIDA | 41 | -0.465482 | -1.127 | 0.30219 | 0.574799 |
| GOBP_NEGATIVE_REGULATION_OF_CELL_DEVELOPMENT | 105 | -0.416909 | -1.12696 | 0.277411 | 0.574461 |
| GOBP_POSITIVE_REGULATION_OF_EXTRINSIC_APOPTOTIC_SIGNALING_PATHWA | 36 | -0.47863 | -1.126873 | 0.314949 | 0.574245 |
| GOBP_DETECTION_OF_MECHANICAL_STIMULUS | 30 | -0.490098 | -1.126087 | 0.291727 | 0.575709 |
| GOBP_RESPONSE_TO_COPPER_ION | 24 | -0.507646 | -1.126032 | 0.322632 | 0.57542 |
| GOBP_REGULATION_OF_HEART_RATE | 44 | -0.46208 | -1.126017 | 0.294618 | 0.575032 |
| GOBP_MYOTUBE_DIFFERENTIATION | 68 | -0.432166 | -1.125129 | 0.280749 | 0.576777 |
| GOBP_MUSCLE_SYSTEM_PROCESS | 261 | -0.383717 | -1.124848 | 0.244693 | 0.577018 |
| GOBP_EXTRINSIC_APOPTOTIC_SIGNALING_PATHWAY | 168 | -0.399177 | -1.124826 | 0.252392 | 0.576634 |
| GOBP_NEGATIVE_REGULATION_OF_ENDOCYTOSIS | 40 | -0.478905 | -1.124105 | 0.308229 | 0.577976 |
| GOBP_POSITIVE_REGULATION_OF_NATURAL_KILLER_CELL_ACTIVATION | 18 | -0.532963 | -1.12409 | 0.347334 | 0.577579 |
| GOBP_NEGATIVE_REGULATION_OF_BIOMINERALIZATION | 16 | -0.541044 | -1.123952 | 0.362385 | 0.57747 |
| GOBP_NEPHRON_EPITHELIUM_DEVELOPMENT | 61 | -0.446976 | -1.123813 | 0.296496 | 0.577394 |
| GOBP_REGULATION_OF_BLOOD_PRESSURE | 99 | -0.416617 | -1.123698 | 0.282995 | 0.577231 |
| GOBP_GRANULOCYTE_DIFFERENTIATION | 27 | -0.498181 | -1.122599 | 0.339312 | 0.579564 |
| GOBP_OSTEOBLAST_DIFFERENTIATION | 136 | -0.398036 | -1.121736 | 0.262515 | 0.581298 |
| GOBP_INORGANIC_ANION_TRANSMEMBRANE_TRANSPORT | 68 | -0.436364 | -1.121156 | 0.300279 | 0.582312 |
| GOBP_CELLULAR_RESPONSE_TO_NUTRIENT | 27 | -0.491327 | -1.120565 | 0.345736 | 0.583325 |
| GOBP_CELLULAR_RESPONSE_TO_CORTICOSTEROID_STIMULUS | 41 | -0.470165 | -1.120121 | 0.322674 | 0.583964 |
| GOBP_NEGATIVE_REGULATION_OF_OXIDATIVE_STRESS_INDUCED_INTRINSIC_A | 18 | -0.524231 | -1.119962 | 0.343333 | 0.583905 |
| GOBP_REGULATION_OF_CELL_SIZE | 135 | -0.398383 | -1.11933 | 0.268585 | 0.585078 |
| GOBP_NEGATIVE_REGULATION_OF_AXON_EXTENSION_INVOLVED_IN_AXON_GU | 20 | -0.517869 | -1.11754 | 0.334375 | 0.589148 |
| GOBP_ANATOMICAL_STRUCTURE_HOMEOSTASIS | 358 | -0.373411 | -1.117087 | 0.226254 | 0.589754 |
| GOBP_REGULATION_OF_ACTION_POTENTIAL | 28 | -0.495542 | -1.116791 | 0.335843 | 0.590025 |
| GOBP_POSITIVE_REGULATION_OF_LIPID_TRANSPORT | 55 | -0.451183 | -1.116749 | 0.294925 | 0.589707 |
| GOBP_NEUROMUSCULAR_JUNCTION_DEVELOPMENT | 28 | -0.491441 | -1.116589 | 0.342383 | 0.589599 |
| GOBP_HISTONE_H3_K9_MODIFICATION | 39 | -0.469133 | -1.115374 | 0.315108 | 0.59217 |
| GOBP_SECRETORY_GRANULE_ORGANIZATION | 27 | -0.495688 | -1.11496 | 0.33595 | 0.59272 |
| GOBP_REGULATION_OF_RYANODINE_SENSITIVE_CALCIUM_RELEASE_CHANNE | 17 | -0.536141 | -1.114834 | 0.345219 | 0.59259 |
| GOBP_NEGATIVE_REGULATION_OF_LIPID_CATABOLIC_PROCESS | 15 | -0.543246 | -1.11464 | 0.362916 | 0.592638 |
| GOBP_POSITIVE_REGULATION_OF_BLOOD_CIRCULATION | 21 | -0.527604 | -1.112744 | 0.344 | 0.59692 |
| GOBP_PROTEIN_AUTOPHOSPHORYLATION | 193 | -0.384901 | -1.112547 | 0.26122 | 0.596952 |
| GOBP_NEGATIVE_REGULATION_OF_MITOTIC_CELL_CYCLE | 276 | -0.375979 | -1.112477 | 0.252212 | 0.596681 |
| GOBP_LYMPHOCYTE_APOPTOTIC_PROCESS | 60 | -0.441494 | -1.111508 | 0.317073 | 0.598655 |
| GOBP_NEURON_PROJECTION_ORGANIZATION | 73 | -0.429921 | -1.1111 | 0.295172 | 0.59915 |
| GOBP_LOCOMOTORY_BEHAVIOR | 91 | -0.417231 | -1.110965 | 0.309429 | 0.599081 |
| GOBP_EPITHELIAL_TO_MESENCHYMAL_TRANSITION | 105 | -0.411453 | -1.110965 | 0.287742 | 0.598642 |
| GOBP_NEGATIVE_REGULATION_OF_TRANSMEMBRANE_RECEPTOR_PROTEIN_S | 85 | -0.418832 | -1.110819 | 0.290323 | 0.598554 |
| GOBP_REGULATION_OF_PROTEIN_TARGETING_TO_MITOCHONDRION | 41 | -0.465209 | -1.110788 | 0.312413 | 0.5982 |
| GOBP_VIRAL_GENOME_REPLICATION | 117 | -0.401987 | -1.110379 | 0.293399 | 0.598731 |
| GOBP_REGULATION_OF_GLUCOSE_METABOLIC_PROCESS | 94 | -0.412329 | -1.11019 | 0.312102 | 0.59877 |
| GOBP_REGULATION_OF_DNA_BIOSYNTHETIC_PROCESS | 91 | -0.411822 | -1.109021 | 0.307494 | 0.601183 |
| GOBP_POSITIVE_REGULATION_OF_CELL_MORPHOGENESIS_INVOLVED_IN_DIFF | 64 | -0.434348 | -1.108942 | 0.311671 | 0.600932 |
| GOBP_NEGATIVE_REGULATION_OF_PROTEIN_CONTAINING_COMPLEX_ASSEMB | 108 | -0.407243 | -1.108938 | 0.31114 | 0.600503 |
| GOBP_IMMUNE_RESPONSE_TO_TUMOR_CELL | 20 | -0.514217 | -1.108126 | 0.347032 | 0.602082 |
| GOBP_REGULATION_OF_SPHINGOLIPID_BIOSYNTHETIC_PROCESS | 16 | -0.544376 | -1.10808 | 0.357254 | 0.601777 |
| GOBP_AGING | 216 | -0.380856 | -1.107053 | 0.253136 | 0.603873 |
| GOBP_MEMORY | 64 | -0.433014 | -1.107038 | 0.306958 | 0.603474 |
| GOBP_NEPHRON_MORPHOGENESIS | 36 | -0.461815 | -1.10593 | 0.329819 | 0.605746 |
| GOBP_POSITIVE_REGULATION_OF_INTERLEUKIN_10_PRODUCTION | 30 | -0.476921 | -1.105134 | 0.335901 | 0.607274 |
| GOBP_PLATELET_AGGREGATION | 48 | -0.442242 | -1.104367 | 0.311813 | 0.608725 |
| GOBP_POLYSACCHARIDE_CATABOLIC_PROCESS | 23 | -0.512937 | -1.104156 | 0.348367 | 0.60878 |
| GOBP_CAMP_MEDIATED_SIGNALING | 30 | -0.487243 | -1.10343 | 0.344361 | 0.610153 |
| GOBP_REGULATION_OF_MAST_CELL_ACTIVATION_INVOLVED_IN_IMMUNE_RESI | 25 | -0.491589 | -1.102973 | 0.357143 | 0.610909 |
| GOBP_RESPONSE_TO_OXYGEN_LEVELS | 300 | -0.364104 | -1.102879 | 0.258772 | 0.610687 |
| GOBP_POSITIVE_REGULATION_OF_INTRINSIC_APOPTOTIC_SIGNALING_PATHWA | 54 | -0.450922 | -1.102556 | 0.325858 | 0.611063 |
| GOBP_RESPONSE_TO_ANTIBIOTIC | 37 | -0.473629 | -1.101946 | 0.33142 | 0.612154 |
| GOBP_ADAPTIVE_THERMOGENESIS | 110 | -0.402558 | -1.101816 | 0.312736 | 0.612009 |
| GOBP_DEVELOPMENTAL_GROWTH | 414 | -0.36498 | -1.101635 | 0.25 | 0.612 |
| GOBP_NEGATIVE_REGULATION_OF_CELL_CYCLE_PHASE_TRANSITION | 231 | -0.373348 | -1.100428 | 0.276451 | 0.614564 |
| GOBP_ENDOPLASMIC_RETICULUM_CALCIUM_ION_HOMEOSTASIS | 25 | -0.494961 | -1.100258 | 0.357035 | 0.614559 |
| GOBP_CELLULAR_TRANSITION_METAL_ION_HOMEOSTASIS | 89 | -0.412341 | -1.100087 | 0.306995 | 0.614551 |
| GOBP_CELLULAR_RESPONSE_TO_GLUCOSE_STARVATION | 44 | -0.457883 | -1.099938 | 0.337535 | 0.61447 |
| GOBP_NEGATIVE_REGULATION_OF_PHOSPHORUS_METABOLIC_PROCESS | 408 | -0.364719 | -1.099756 | 0.258167 | 0.614478 |
| GOBP_NEGATIVE_REGULATION_OF_VIRAL_LIFE_CYCLE | 17 | -0.539139 | -1.099373 | 0.388034 | 0.61499 |
| GOBP_REGULATION_OF_LIPID_BIOSYNTHETIC_PROCESS | 143 | -0.389634 | -1.099181 | 0.288533 | 0.615028 |
| GOBP_REGULATION_OF_MULTICELLULAR_ORGANISM_GROWTH | 42 | -0.457319 | -1.09898 | 0.334307 | 0.615091 |
| GOBP_NEGATIVE_REGULATION_OF_PROTEIN_CATABOLIC_PROCESS | 99 | -0.405935 | -1.098156 | 0.318008 | 0.616693 |
| GOBP_REGULATION_OF_PROTEIN_DEPHOSPHORYLATION | 118 | -0.396905 | -1.098011 | 0.314465 | 0.616598 |
| GOBP_NUCLEOSIDE_TRIPHOSPHATE_METABOLIC_PROCESS | 96 | -0.405633 | -1.097972 | 0.294041 | 0.616253 |
| GOBP_MITOCHONDRION_LOCALIZATION | 42 | -0.462264 | -1.097678 | 0.349782 | 0.616558 |
| GOBP_POST_TRANSLATIONAL_PROTEIN_MODIFICATION | 293 | -0.371654 | -1.097486 | 0.284104 | 0.616572 |
| GOBP_POSITIVE_REGULATION_OF_STEROID_METABOLIC_PROCESS | 15 | -0.539785 | -1.097397 | 0.393791 | 0.61637 |
| GOBP_REGULATION_OF_ESTABLISHMENT_OR_MAINTENANCE_OF_CELL_POLAR | 23 | -0.5027 | -1.096651 | 0.35061 | 0.6178 |
| GOBP_REGULATION_OF_CELLULAR_PROTEIN_CATABOLIC_PROCESS | 221 | -0.37673 | -1.096098 | 0.284223 | 0.618785 |
| GOBP_REGULATION_OF_FATTY_ACID_TRANSPORT | 29 | -0.486678 | -1.095997 | 0.369436 | 0.618573 |
| GOBP_SKELETAL_SYSTEM_DEVELOPMENT | 285 | -0.37111 | -1.095589 | 0.277778 | 0.619144 |
| GOBP_POSITIVE_REGULATION_OF_MRNA_METABOLIC_PROCESS | 74 | -0.426243 | -1.094796 | 0.336375 | 0.620708 |
| GOBP_GLYCEROLIPID_BIOSYNTHETIC_PROCESS | 220 | -0.37993 | -1.094664 | 0.311628 | 0.620597 |
| GOBP_REGULATION_OF_BIOLOGICAL_PROCESS_INVOLVED_IN_SYMBIOTIC_INTE | 168 | -0.384451 | -1.094636 | 0.300242 | 0.62021 |
| GOBP_EXPORT_ACROSS_PLASMA_MEMBRANE | 35 | -0.470955 | -1.094232 | 0.342065 | 0.62084 |
| GOBP_NEGATIVE_REGULATION_OF_KINASE_ACTIVITY | 223 | -0.374714 | -1.094106 | 0.295633 | 0.620751 |
| GOBP_CELL_REDOX_HOMEOSTASIS | 39 | -0.455373 | -1.093274 | 0.352339 | 0.622335 |
| GOBP_MESONEPHRIC_TUBULE_MORPHOGENESIS | 29 | -0.47545 | -1.093216 | 0.369969 | 0.622026 |
| GOBP_DETECTION_OF_LIGHT_STIMULUS | 39 | -0.457487 | -1.093211 | 0.351695 | 0.621604 |
| GOBP_TOR_SIGNALING | 112 | -0.395252 | -1.092261 | 0.328801 | 0.623513 |
| GOBP_BEHAVIOR | 309 | -0.368516 | -1.092217 | 0.284464 | 0.623178 |
| GOBP_GENE_SILENCING | 235 | -0.369608 | -1.091506 | 0.296629 | 0.624471 |
| GOBP_PROTEIN_KINASE_C_ACTIVATING_G_PROTEIN_COUPLED_RECEPTOR_SI | 16 | -0.516377 | -1.091065 | 0.367347 | 0.625135 |
| GOBP_TRANSMEMBRANE_RECEPTOR_PROTEIN_SERINE_THREONINE_KINASE_ | 233 | -0.370686 | -1.090976 | 0.287794 | 0.624911 |
| GOBP_MOTOR_NEURON_AXON_GUIDANCE | 19 | -0.508367 | -1.089325 | 0.380727 | 0.628617 |
| GOBP_REGULATION_OF_TUMOR_NECROSIS_FACTOR_MEDIATED_SIGNALING_P | 54 | -0.434884 | -1.089195 | 0.358726 | 0.628503 |
| GOBP_EMBRYONIC_ORGAN_MORPHOGENESIS | 152 | -0.39375 | -1.088345 | 0.315976 | 0.63015 |
| GOBP_REGULATION_OF_RELEASE_OF_SEQUESTERED_CALCIUM_ION_INTO_CY | 44 | -0.442849 | -1.088145 | 0.353111 | 0.630186 |
| GOBP_CELLULAR_RESPONSE_TO_HEAT | 106 | -0.394664 | -1.088021 | 0.320102 | 0.630053 |
| GOBP_REGULATION_OF_SYNAPTIC_PLASTICITY | 116 | -0.397712 | -1.087892 | 0.329643 | 0.629933 |
| GOBP_BROWN_FAT_CELL_DIFFERENTIATION | 31 | -0.469542 | -1.086718 | 0.368889 | 0.632412 |
| GOBP_DEVELOPMENTAL_GROWTH_INVOLVED_IN_MORPHOGENESIS | 161 | -0.380802 | -1.086026 | 0.315851 | 0.633714 |
| GOBP_CHONDROCYTE_DIFFERENTIATION | 57 | -0.430234 | -1.085983 | 0.346774 | 0.633358 |
| GOBP_P38MAPK_CASCADE | 36 | -0.457407 | -1.085981 | 0.332842 | 0.632917 |
| GOBP_POTASSIUM_ION_TRANSPORT | 122 | -0.391986 | -1.085885 | 0.324291 | 0.632698 |
| GOBP_LENS_FIBER_CELL_DIFFERENTIATION | 21 | -0.493719 | -1.085802 | 0.363777 | 0.632468 |
| GOBP_POSITIVE_REGULATION_OF_EXTRACELLULAR_MATRIX_ORGANIZATION | 19 | -0.509071 | -1.085697 | 0.377419 | 0.632278 |
| GOBP_NEGATIVE_REGULATION_OF_TORC1_SIGNALING | 15 | -0.525875 | -1.085608 | 0.377451 | 0.632041 |
| GOBP_MORPHOGENESIS_OF_AN_EPITHELIUM | 354 | -0.364237 | -1.085271 | 0.283243 | 0.632454 |
| GOBP_TRANSCRIPTION_INITIATION_FROM_RNA_POLYMERASE_II_PROMOTER | 158 | -0.379103 | -1.085217 | 0.327014 | 0.632142 |
| GOBP_RESPONSE_TO_ORGANOPHOSPHORUS | 88 | -0.404001 | -1.083909 | 0.330739 | 0.634954 |
| GOBP_RESPONSE_TO_INTERLEUKIN_7 | 29 | -0.466601 | -1.08373 | 0.362963 | 0.634949 |
| GOBP_NEGATIVE_REGULATION_OF_PHOSPHORYLATION | 319 | -0.361429 | -1.082646 | 0.303591 | 0.637166 |
| GOBP_NEGATIVE_REGULATION_OF_AXON_EXTENSION | 34 | -0.459212 | -1.081388 | 0.354449 | 0.63988 |
| GOBP_CELLULAR_RESPONSE_TO_STEROID_HORMONE_STIMULUS | 148 | -0.384466 | -1.081248 | 0.326165 | 0.639781 |
| GOBP_REGULATION_OF_HEART_CONTRACTION | 128 | -0.392781 | -1.081044 | 0.331725 | 0.639842 |
| GOBP_MAST_CELL_ACTIVATION | 47 | -0.436168 | -1.080912 | 0.356932 | 0.639702 |
| GOBP_EPITHELIAL_TUBE_MORPHOGENESIS | 194 | -0.378609 | -1.080484 | 0.338863 | 0.640297 |
| GOBP_POSITIVE_REGULATION_OF_B_CELL_ACTIVATION | 97 | -0.401232 | -1.080481 | 0.331633 | 0.639862 |
| GOBP_POSITIVE_REGULATION_OF_SODIUM_ION_TRANSPORT | 26 | -0.473405 | -1.080357 | 0.390977 | 0.639734 |
| GOBP_INTRACELLULAR_LIPID_TRANSPORT | 39 | -0.446209 | -1.080296 | 0.36119 | 0.639431 |
| GOBP_POSITIVE_REGULATION_OF_CYTOKINE_PRODUCTION_INVOLVED_IN_IMM | 46 | -0.439873 | -1.080209 | 0.363248 | 0.639205 |
| GOBP_REGULATION_OF_CELLULAR_CARBOHYDRATE_METABOLIC_PROCESS | 109 | -0.397103 | -1.078361 | 0.339109 | 0.643302 |
| GOBP_MEMBRANE_REPOLARIZATION | 23 | -0.484546 | -1.078312 | 0.392345 | 0.642976 |
| GOBP_ANOIKIS | 30 | -0.459175 | -1.077784 | 0.375951 | 0.643851 |
| GOBP_NEGATIVE_REGULATION_OF_SIGNAL_TRANSDUCTION_IN_ABSENCE_OF_ | 20 | -0.492339 | -1.077547 | 0.392384 | 0.643967 |
| GOBP_HEART_DEVELOPMENT | 337 | -0.360258 | -1.07753 | 0.308361 | 0.643564 |
| GOBP_MONOCARBOXYLIC_ACID_METABOLIC_PROCESS | 467 | -0.353483 | -1.077295 | 0.293933 | 0.643722 |
| GOBP_KERATAN_SULFATE_BIOSYNTHETIC_PROCESS | 18 | -0.515318 | -1.07727 | 0.401945 | 0.643342 |
| GOBP_SUBSTRATE_ADHESION_DEPENDENT_CELL_SPREADING | 84 | -0.409286 | -1.077112 | 0.344737 | 0.643328 |
| GOBP_TISSUE_MORPHOGENESIS | 410 | -0.355332 | -1.076815 | 0.298209 | 0.643615 |
| GOBP_REGULATION_OF_CELLULAR_AMIDE_METABOLIC_PROCESS | 376 | -0.355038 | -1.075937 | 0.308271 | 0.645319 |
| GOBP_RESPONSE_TO_ALKALOID | 66 | -0.413146 | -1.075852 | 0.351678 | 0.645092 |
| GOBP_RESPONSE_TO_METAL_ION | 242 | -0.369535 | -1.075441 | 0.320628 | 0.645647 |
| GOBP_DNA_MODIFICATION | 93 | -0.397196 | -1.075163 | 0.351634 | 0.645885 |
| GOBP_NEGATIVE_REGULATION_OF_UBIQUITIN_DEPENDENT_PROTEIN_CATABO | 45 | -0.441433 | -1.074864 | 0.366762 | 0.646182 |
| GOBP_DNA_DEALKYLATION | 31 | -0.468365 | -1.073988 | 0.390649 | 0.647845 |
| GOBP_INTERLEUKIN_7_MEDIATED_SIGNALING_PATHWAY | 19 | -0.502898 | -1.073557 | 0.383628 | 0.648433 |
| GOBP_RESPONSE_TO_NICOTINE | 25 | -0.483239 | -1.073514 | 0.382263 | 0.648093 |
| GOBP_REGULATION_OF_TRANSCRIPTION_INVOLVED_IN_G1_S_TRANSITION_OF_ | 32 | -0.461899 | -1.073464 | 0.361482 | 0.647772 |
| GOBP_REGULATION_OF_SYSTEMIC_ARTERIAL_BLOOD_PRESSURE | 45 | -0.439416 | -1.072935 | 0.392241 | 0.648672 |
| GOBP_CALCIUM_ION_TRANSPORT_INTO_CYTOSOL | 91 | -0.407629 | -1.072216 | 0.355263 | 0.650037 |
| GOBP_NEGATIVE_REGULATION_OF_CATABOLIC_PROCESS | 247 | -0.368612 | -1.071897 | 0.330717 | 0.650357 |

| | | | | | |
|---|---|---|---|---|---|
| GOBP_POSITIVE_REGULATION_OF_GLYCOPROTEIN_METABOLIC_PROCESS | 18 | -0.507283 | -1.071796 | 0.394904 | 0.650186 |
| GOBP_COLLAGEN_BIOSYNTHETIC_PROCESS | 30 | -0.464502 | -1.071402 | 0.377444 | 0.650725 |
| GOBP_NEGATIVE_REGULATION_OF_CYTOKINE_PRODUCTION_INVOLVED_IN_INF | 20 | -0.501129 | -1.071125 | 0.388889 | 0.650965 |
| GOBP_RESPIRATORY_SYSTEM_DEVELOPMENT | 120 | -0.393691 | -1.070763 | 0.336709 | 0.651421 |
| GOBP_T_HELPER_17_TYPE_IMMUNE_RESPONSE | 25 | -0.476111 | -1.070687 | 0.4096 | 0.651159 |
| GOBP_ISOPRENOID_BIOSYNTHETIC_PROCESS | 21 | -0.491059 | -1.070563 | 0.39685 | 0.651016 |
| GOBP_TUMOR_NECROSIS_FACTOR_MEDIATED_SIGNALING_PATHWAY | 152 | -0.380074 | -1.070012 | 0.333734 | 0.651919 |
| GOBP_POSITIVE_REGULATION_OF_INSULIN_SECRETION_INVOLVED_IN_CELLUL | 26 | -0.476987 | -1.069848 | 0.387755 | 0.651872 |
| GOBP_RESPONSE_TO_ESTRADIOL | 76 | -0.401704 | -1.069578 | 0.345745 | 0.652074 |
| GOBP_NEGATIVE_REGULATION_OF_CD4_POSITIVE_ALPHA_BETA_T_CELL_ACTI | 27 | -0.479634 | -1.068763 | 0.385093 | 0.653701 |
| GOBP_ERBB2_SIGNALING_PATHWAY | 25 | -0.476953 | -1.06868 | 0.380282 | 0.653452 |
| GOBP_VASOCONSTRICTION | 35 | -0.454228 | -1.068525 | 0.39302 | 0.653363 |
| GOBP_CYTOSOLIC_CALCIUM_ION_TRANSPORT | 110 | -0.394906 | -1.068018 | 0.352287 | 0.654105 |
| GOBP_REGULATION_OF_EXTRINSIC_APOPTOTIC_SIGNALING_PATHWAY_VIA_DE | 44 | -0.445755 | -1.068008 | 0.373737 | 0.653688 |
| GOBP_REGULATION_OF_VASOCONSTRICTION | 29 | -0.472368 | -1.067338 | 0.405926 | 0.654932 |
| GOBP_VIRAL_RNA_GENOME_REPLICATION | 32 | -0.467454 | -1.067326 | 0.390909 | 0.654521 |
| GOBP_CELLULAR_COMPONENT_MAINTENANCE | 47 | -0.433693 | -1.0671 | 0.39233 | 0.654627 |
| GOBP_POSITIVE_REGULATION_OF_CELLULAR_RESPONSE_TO_INSULIN_STIMUL | 18 | -0.507078 | -1.067004 | 0.402477 | 0.654434 |
| GOBP_POSITIVE_REGULATION_OF_MICROTUBULE_POLYMERIZATION_OR_DEPO | 29 | -0.475227 | -1.06686 | 0.394303 | 0.654361 |
| GOBP_CELL_COMMUNICATION_INVOLVED_IN_CARDIAC_CONDUCTION | 27 | -0.474864 | -1.065932 | 0.392366 | 0.656216 |
| GOBP_NEGATIVE_REGULATION_OF_CELLULAR_RESPONSE_TO_GROWTH_FACT | 57 | -0.418809 | -1.065153 | 0.376502 | 0.657743 |
| GOBP_ANTIGEN_PROCESSING_AND_PRESENTATION_OF_ENDOGENOUS_ANTIG | 21 | -0.49478 | -1.065039 | 0.386513 | 0.657562 |
| GOBP_POSITIVE_REGULATION_OF_EPIDERMIS_DEVELOPMENT | 22 | -0.499528 | -1.064672 | 0.420063 | 0.658031 |
| GOBP_REGULATION_OF_DENDRITE_EXTENSION | 17 | -0.507325 | -1.064667 | 0.384117 | 0.657601 |
| GOBP_NEURON_PROJECTION_EXTENSION_INVOLVED_IN_NEURON_PROJECTIO | 26 | -0.47844 | -1.063607 | 0.4003 | 0.659795 |
| GOBP_CELLULAR_SENESCENCE | 57 | -0.43497 | -1.063394 | 0.379408 | 0.659846 |
| GOBP_NEGATIVE_REGULATION_OF_OSSIFICATION | 19 | -0.499124 | -1.063146 | 0.382445 | 0.660013 |
| GOBP_REGULATION_OF_MAP_KINASE_ACTIVITY | 236 | -0.365485 | -1.062714 | 0.36991 | 0.660606 |
| GOBP_CELLULAR_RESPONSE_TO_DSRNA | 19 | -0.502167 | -1.062412 | 0.425775 | 0.660915 |
| GOBP_VASCULAR_ENDOTHELIAL_GROWTH_FACTOR_SIGNALING_PATHWAY | 27 | -0.467018 | -1.06227 | 0.409295 | 0.660813 |
| GOBP_GLUTAMATE_METABOLIC_PROCESS | 25 | -0.477749 | -1.060905 | 0.400612 | 0.663707 |
| GOBP_HEART_VALVE_DEVELOPMENT | 35 | -0.454504 | -1.060825 | 0.390244 | 0.663472 |
| GOBP_POSITIVE_REGULATION_OF_CELL_PROJECTION_ORGANIZATION | 263 | -0.363568 | -1.0608 | 0.344062 | 0.663094 |
| GOBP_CYTOKINE_PRODUCTION_INVOLVED_IN_INFLAMMATORY_RESPONSE | 44 | -0.433149 | -1.060731 | 0.398204 | 0.662802 |
| GOBP_REGULATION_OF_NEUROTRANSMITTER_LEVELS | 132 | -0.381178 | -1.059901 | 0.377614 | 0.664415 |
| GOBP_RESPONSE_TO_HEAT | 137 | -0.381249 | -1.059656 | 0.361244 | 0.664581 |
| GOBP_ORGANIC_HYDROXY_COMPOUND_METABOLIC_PROCESS | 383 | -0.351718 | -1.059499 | 0.350706 | 0.664519 |
| GOBP_NEGATIVE_REGULATION_OF_LIPID_METABOLIC_PROCESS | 63 | -0.412926 | -1.058711 | 0.390042 | 0.666023 |
| GOBP_PROTEIN_MATURATION | 218 | -0.365274 | -1.058461 | 0.364475 | 0.666167 |
| GOBP_ADULT_BEHAVIOR | 65 | -0.41555 | -1.058179 | 0.383217 | 0.666431 |
| GOBP_REGULATION_OF_DEPHOSPHORYLATION | 170 | -0.370902 | -1.057919 | 0.357907 | 0.666614 |
| GOBP_NEURON_PROJECTION_EXTENSION | 126 | -0.382316 | -1.057331 | 0.369458 | 0.667593 |
| GOBP_POSITIVE_REGULATION_OF_DENDRITIC_SPINE_MORPHOGENESIS | 15 | -0.515548 | -1.057314 | 0.413735 | 0.667195 |
| GOBP_ACIDIC_AMINO_ACID_TRANSPORT | 42 | -0.447295 | -1.056956 | 0.397727 | 0.667596 |
| GOBP_SPERM_AXONEME_ASSEMBLY | 16 | -0.516364 | -1.056883 | 0.413399 | 0.667341 |
| GOBP_ACTIVATION_OF_PROTEIN_KINASE_ACTIVITY | 247 | -0.364033 | -1.056624 | 0.364641 | 0.667532 |
| GOBP_CELLULAR_KETONE_METABOLIC_PROCESS | 188 | -0.369506 | -1.056522 | 0.368605 | 0.66734 |
| GOBP_PEPTIDYL_TYROSINE_DEPHOSPHORYLATION | 82 | -0.399978 | -1.056067 | 0.387399 | 0.668035 |
| GOBP_POSITIVE_REGULATION_OF_PROTEIN_BINDING | 70 | -0.403552 | -1.055854 | 0.383065 | 0.668164 |
| GOBP_REGULATION_OF_PROTEIN_TYROSINE_KINASE_ACTIVITY | 69 | -0.405091 | -1.055619 | 0.388186 | 0.66829 |
| GOBP_RESPIRATORY_SYSTEM_PROCESS | 20 | -0.488311 | -1.054881 | 0.392188 | 0.669678 |
| GOBP_NEGATIVE_REGULATION_OF_PROTEIN_SERINE_THREONINE_KINASE_AC | 116 | -0.383286 | -1.054234 | 0.373134 | 0.670828 |
| GOBP_PURINERGIC_NUCLEOTIDE_RECEPTOR_SIGNALING_PATHWAY | 23 | -0.473538 | -1.054012 | 0.394081 | 0.670901 |
| GOBP_POSITIVE_REGULATION_OF_DEVELOPMENTAL_GROWTH | 109 | -0.383934 | -1.05387 | 0.353884 | 0.670835 |
| GOBP_MEGAKARYOCYTE_DIFFERENTIATION | 79 | -0.395581 | -1.053815 | 0.381201 | 0.670542 |
| GOBP_CARBOHYDRATE_DERIVATIVE_CATABOLIC_PROCESS | 147 | -0.373415 | -1.053627 | 0.384154 | 0.670577 |
| GOBP_LIPID_OXIDATION | 93 | -0.388056 | -1.053076 | 0.360202 | 0.67146 |
| GOBP_STRIATED_MUSCLE_CELL_APOPTOTIC_PROCESS | 25 | -0.463949 | -1.052406 | 0.415902 | 0.672677 |
| GOBP_REGULATION_OF_PROTEIN_BINDING | 161 | -0.370682 | -1.052172 | 0.384346 | 0.672821 |
| GOBP_NEGATIVE_REGULATION_OF_PROTEIN_DEPHOSPHORYLATION | 51 | -0.428034 | -1.052079 | 0.379913 | 0.672607 |
| GOBP_RESPONSE_TO_HYPEROXIA | 18 | -0.479886 | -1.051455 | 0.418719 | 0.673656 |
| GOBP_CYTOKINE_PRODUCTION_INVOLVED_IN_IMMUNE_RESPONSE | 77 | -0.403634 | -1.05071 | 0.385445 | 0.675052 |
| GOBP_RIBONUCLEOSIDE_TRIPHOSPHATE_METABOLIC_PROCESS | 78 | -0.400395 | -1.050566 | 0.386973 | 0.674963 |
| GOBP_MUSCLE_STRUCTURE_DEVELOPMENT | 388 | -0.347828 | -1.050378 | 0.360129 | 0.674939 |
| GOBP_MULTICELLULAR_ORGANISM_GROWTH | 97 | -0.388627 | -1.05025 | 0.378205 | 0.674812 |
| GOBP_REGULATION_OF_FATTY_ACID_OXIDATION | 26 | -0.466486 | -1.050014 | 0.402439 | 0.674956 |
| GOBP_PROTEIN_K63_LINKED_UBIQUITINATION | 53 | -0.417724 | -1.049999 | 0.402685 | 0.674552 |
| GOBP_REGULATION_OF_SYNAPTIC_VESICLE_RECYCLING | 20 | -0.487588 | -1.049991 | 0.42236 | 0.674139 |
| GOBP_AXON_EXTENSION | 91 | -0.387585 | -1.049686 | 0.387306 | 0.674476 |
| GOBP_NEGATIVE_REGULATION_OF_SECRETION | 98 | -0.382984 | -1.049394 | 0.375949 | 0.674753 |
| GOBP_RESPONSE_TO_CATECHOLAMINE | 54 | -0.416452 | -1.049361 | 0.381487 | 0.6744 |
| GOBP_LEUKOCYTE_HOMEOSTASIS | 70 | -0.405219 | -1.049136 | 0.391478 | 0.674507 |
| GOBP_SIGNAL_RELEASE | 306 | -0.350387 | -1.048861 | 0.374172 | 0.674712 |
| GOBP_REGULATION_OF_CARDIAC_CONDUCTION | 35 | -0.447015 | -1.048187 | 0.416918 | 0.675991 |
| GOBP_MESODERM_DEVELOPMENT | 72 | -0.40291 | -1.047614 | 0.414115 | 0.676935 |
| GOBP_MITOTIC_RECOMBINATION | 23 | -0.47641 | -1.046507 | 0.417702 | 0.679181 |
| GOBP_TRANSMISSION_OF_NERVE_IMPULSE | 29 | -0.460101 | -1.04622 | 0.427457 | 0.679449 |
| GOBP_REGULATION_OF_DENDRITIC_SPINE_MORPHOGENESIS | 36 | -0.444318 | -1.045401 | 0.402635 | 0.681016 |
| GOBP_TRANSPOSITION | 21 | -0.492155 | -1.044989 | 0.428571 | 0.681624 |
| GOBP_ALCOHOL_METABOLIC_PROCESS | 273 | -0.352515 | -1.044531 | 0.399556 | 0.682303 |
| GOBP_INTEGRIN_ACTIVATION | 15 | -0.504316 | -1.044093 | 0.435484 | 0.682926 |
| GOBP_RESPONSE_TO_COLD | 36 | -0.450207 | -1.044042 | 0.408571 | 0.682601 |
| GOBP_NEGATIVE_REGULATION_OF_TYPE_I_INTERFERON_PRODUCTION | 44 | -0.424826 | -1.043795 | 0.406828 | 0.682776 |
| GOBP_ORGANISM_EMERGENCE_FROM_PROTECTIVE_STRUCTURE | 16 | -0.500216 | -1.043702 | 0.420886 | 0.682574 |
| GOBP_REGULATION_OF_CELLULAR_RESPONSE_TO_GROWTH_FACTOR_STIMUL | 176 | -0.361882 | -1.04347 | 0.395376 | 0.682681 |
| GOBP_REGULATION_OF_SYSTEM_PROCESS | 322 | -0.349504 | -1.043374 | 0.388949 | 0.682471 |
| GOBP_POSITIVE_REGULATION_OF_ACUTE_INFLAMMATORY_RESPONSE | 18 | -0.494495 | -1.043019 | 0.471729 | 0.682908 |
| GOBP_EXTRACELLULAR_TRANSPORT | 25 | -0.469045 | -1.04265 | 0.429675 | 0.683339 |
| GOBP_ANATOMICAL_STRUCTURE_MATURATION | 140 | -0.371325 | -1.042539 | 0.393162 | 0.683153 |
| GOBP_NEURON_MIGRATION | 84 | -0.391119 | -1.042092 | 0.392388 | 0.683833 |
| GOBP_RESPONSE_TO_ELECTRICAL_STIMULUS | 30 | -0.448581 | -1.041904 | 0.427914 | 0.683851 |
| GOBP_NEGATIVE_REGULATION_OF_CYTOSKELETON_ORGANIZATION | 118 | -0.385153 | -1.041904 | 0.397243 | 0.683416 |
| GOBP_ORGANELLE_MEMBRANE_FUSION | 94 | -0.390265 | -1.041679 | 0.400501 | 0.683514 |
| GOBP_LEUKOCYTE_ADHESION_TO_VASCULAR_ENDOTHELIAL_CELL | 36 | -0.443717 | -1.041519 | 0.434018 | 0.683464 |
| GOBP_ESTABLISHMENT_OR_MAINTENANCE_OF_CELL_POLARITY | 171 | -0.36351 | -1.041409 | 0.395294 | 0.683302 |
| GOBP_DETECTION_OF_STIMULUS | 176 | -0.364226 | -1.041223 | 0.381295 | 0.683349 |
| GOBP_UROGENITAL_SYSTEM_DEVELOPMENT | 200 | -0.360197 | -1.040836 | 0.387022 | 0.683853 |
| GOBP_CELLULAR_LIPID_CATABOLIC_PROCESS | 173 | -0.365052 | -1.040769 | 0.387707 | 0.683572 |
| GOBP_RESPONSE_TO_MANGANESE_ION | 16 | -0.505756 | -1.040742 | 0.448947 | 0.683202 |
| GOBP_MEMBRANE_DEPOLARIZATION_DURING_ACTION_POTENTIAL | 20 | -0.500186 | -1.040425 | 0.466777 | 0.683533 |
| GOBP_REGULATION_OF_SODIUM_ION_TRANSMEMBRANE_TRANSPORT | 45 | -0.425175 | -1.038843 | 0.412342 | 0.686931 |
| GOBP_GLYCEROLIPID_METABOLIC_PROCESS | 340 | -0.348339 | -1.0386 | 0.388528 | 0.687092 |
| GOBP_PROTEIN_DEPOLYMERIZATION | 85 | -0.390734 | -1.03845 | 0.409987 | 0.68698 |
| GOBP_PROTEIN_REFOLDING | 23 | -0.475171 | -1.037524 | 0.433642 | 0.688869 |
| GOBP_EPITHELIAL_CELL_DIFFERENTIATION_INVOLVED_IN_KIDNEY_DEVELOPMI | 29 | -0.451048 | -1.036837 | 0.433483 | 0.690094 |
| GOBP_POSITIVE_REGULATION_OF_PEPTIDYL_LYSINE_ACETYLATION | 29 | -0.445495 | -1.036516 | 0.436834 | 0.690426 |
| GOBP_RESPONSE_TO_ZINC_ION | 32 | -0.447081 | -1.036026 | 0.42522 | 0.691183 |
| GOBP_PROTEIN_LIPID_COMPLEX_ASSEMBLY | 22 | -0.475541 | -1.035489 | 0.429907 | 0.692092 |
| GOBP_ENDOCRINE_SYSTEM_DEVELOPMENT | 59 | -0.410666 | -1.035259 | 0.431293 | 0.692225 |
| GOBP_RESPONSE_TO_AMINE | 32 | -0.453972 | -1.034983 | 0.422323 | 0.692469 |
| GOBP_REGULATION_OF_EXTENT_OF_CELL_GROWTH | 79 | -0.391857 | -1.034911 | 0.415803 | 0.692204 |
| GOBP_RESPONSE_TO_XENOBIOTIC_STIMULUS | 74 | -0.400508 | -1.034484 | 0.406085 | 0.692785 |
| GOBP_LIPID_TRANSLOCATION | 34 | -0.446748 | -1.03435 | 0.41716 | 0.692675 |
| GOBP_FEMALE_GAMETE_GENERATION | 88 | -0.387444 | -1.034253 | 0.406045 | 0.692474 |
| GOBP_ANTERIOR_POSTERIOR_PATTERN_SPECIFICATION | 101 | -0.381433 | -1.033909 | 0.411613 | 0.692867 |
| GOBP_METENCEPHALON_DEVELOPMENT | 66 | -0.403932 | -1.033772 | 0.401628 | 0.692757 |
| GOBP_RESPONSE_TO_LIGHT_STIMULUS | 244 | -0.351824 | -1.033216 | 0.40678 | 0.69363 |
| GOBP_ARTERY_DEVELOPMENT | 55 | -0.410225 | -1.032732 | 0.413605 | 0.69433 |
| GOBP_POSITIVE_REGULATION_OF_ORGANIC_ACID_TRANSPORT | 21 | -0.48548 | -1.032565 | 0.44898 | 0.694301 |
| GOBP_POSITIVE_REGULATION_OF_CALCIUM_ION_TRANSMEMBRANE_TRANSPO | 26 | -0.468396 | -1.031987 | 0.446032 | 0.695256 |
| GOBP_PANCREAS_DEVELOPMENT | 34 | -0.439146 | -1.031506 | 0.434911 | 0.695958 |
| GOBP_REGULATION_OF_OXIDATIVE_PHOSPHORYLATION | 20 | -0.472532 | -1.031294 | 0.445498 | 0.69601 |
| GOBP_MULTICELLULAR_ORGANISMAL_HOMEOSTASIS | 349 | -0.345758 | -1.030714 | 0.395945 | 0.696998 |
| GOBP_PHOSPHATE_ION_TRANSPORT | 20 | -0.48185 | -1.030562 | 0.421138 | 0.696925 |
| GOBP_REGULATION_OF_CIRCADIAN_RHYTHM | 87 | -0.384766 | -1.030362 | 0.409333 | 0.696965 |
| GOBP_NEGATIVE_REGULATION_OF_PROTEIN_MODIFICATION_PROCESS | 451 | -0.340229 | -1.030351 | 0.393494 | 0.696555 |
| GOBP_REGULATION_OF_MYELOID_LEUKOCYTE_MEDIATED_IMMUNITY | 45 | -0.4215 | -1.029905 | 0.41831 | 0.697202 |
| GOBP_ANIMAL_ORGAN_MATURATION | 15 | -0.501965 | -1.029745 | 0.449251 | 0.697143 |
| GOBP_NEGATIVE_REGULATION_OF_CALCIUM_MEDIATED_SIGNALING | 17 | -0.480017 | -1.029474 | 0.436667 | 0.697367 |
| GOBP_ENDOCRINE_PANCREAS_DEVELOPMENT | 22 | -0.468522 | -1.028361 | 0.445498 | 0.699645 |
| GOBP_SPERM_MOTILITY | 54 | -0.402166 | -1.027989 | 0.426966 | 0.700108 |
| GOBP_REGULATION_OF_SYNAPSE_ASSEMBLY | 45 | -0.417243 | -1.02798 | 0.420613 | 0.699686 |
| GOBP_INNER_EAR_RECEPTOR_CELL_DEVELOPMENT | 29 | -0.455817 | -1.027733 | 0.460857 | 0.699856 |
| GOBP_REGULATION_OF_GENE_EXPRESSION_EPIGENETIC | 155 | -0.362094 | -1.026881 | 0.409412 | 0.701455 |
| GOBP_MUSCLE_FIBER_DEVELOPMENT | 34 | -0.439631 | -1.026543 | 0.458841 | 0.701901 |
| GOBP_ESTABLISHMENT_OF_MITOCHONDRION_LOCALIZATION | 23 | -0.472271 | -1.026294 | 0.451314 | 0.702026 |
| GOBP_SENSORY_PERCEPTION_OF_PAIN | 47 | -0.420747 | -1.026177 | 0.432081 | 0.701869 |
| GOBP_CELLULAR_RESPONSE_TO_CAMP | 35 | -0.429725 | -1.025693 | 0.445415 | 0.702602 |
| GOBP_REGULATION_OF_CALCIUM_ION_IMPORT | 22 | -0.465434 | -1.025579 | 0.451314 | 0.702446 |
| GOBP_KERATAN_SULFATE_METABOLIC_PROCESS | 23 | -0.463015 | -1.024773 | 0.443574 | 0.70388 |
| GOBP_APOPTOTIC_MITOCHONDRIAL_CHANGES | 102 | -0.372831 | -1.024003 | 0.401985 | 0.705271 |
| GOBP_DEVELOPMENTAL_CELL_GROWTH | 160 | -0.35905 | -1.023643 | 0.420238 | 0.705681 |
| GOBP_DNA_METHYLATION | 51 | -0.412457 | -1.023194 | 0.419944 | 0.706344 |
| GOBP_POSITIVE_REGULATION_OF_TRANSLATION | 114 | -0.37056 | -1.023153 | 0.439451 | 0.706002 |
| GOBP_NEURONAL_ACTION_POTENTIAL | 15 | -0.501389 | -1.022625 | 0.479933 | 0.706802 |
| GOBP_PROTEIN_DEPHOSPHORYLATION | 271 | -0.347847 | -1.022521 | 0.427163 | 0.706589 |
| GOBP_REGULATION_OF_MITOCHONDRION_ORGANIZATION | 138 | -0.365333 | -1.021927 | 0.450185 | 0.707613 |

| | | | | | |
|---|---|---|---|---|---|
| GOBP_POSITIVE_REGULATION_OF_LAMELLIPODIUM_ASSEMBLY | 26 | -0.454754 | -1.021398 | 0.421947 | 0.708455 |
| GOBP_REGULATION_OF_MEMBRANE_LIPID_DISTRIBUTION | 40 | -0.425634 | -1.021337 | 0.450216 | 0.70815 |
| GOBP_NEGATIVE_REGULATION_OF_DEVELOPMENTAL_GROWTH | 78 | -0.386179 | -1.021291 | 0.431968 | 0.70783 |
| GOBP_NEGATIVE_REGULATION_OF_DNA_TEMPLATED_TRANSCRIPTION_ELONG | 15 | -0.495863 | -1.021234 | 0.466561 | 0.707524 |
| GOBP_DEPHOSPHORYLATION | 403 | -0.340196 | -1.021038 | 0.436709 | 0.70752 |
| GOBP_MAMMARY_GLAND_EPITHELIUM_DEVELOPMENT | 46 | -0.414515 | -1.020485 | 0.426554 | 0.708409 |
| GOBP_NEGATIVE_REGULATION_OF_NOTCH_SIGNALING_PATHWAY | 25 | -0.462631 | -1.020118 | 0.456061 | 0.708867 |
| GOBP_PROTEIN_PHOSPHOPANTETHEINYLATION | 271 | -0.341983 | -1.019839 | 0.433815 | 0.70915 |
| GOBP_MUSCLE_CELL_DIFFERENTIATION | 221 | -0.350156 | -1.019771 | 0.441964 | 0.708867 |
| GOBP_CELLULAR_RESPONSE_TO_OXYGEN_LEVELS | 185 | -0.354703 | -1.01953 | 0.42623 | 0.709029 |
| GOBP_NEGATIVE_REGULATION_OF_EPITHELIAL_TO_MESENCHYMAL_TRANSITIO | 25 | -0.463338 | -1.019254 | 0.437788 | 0.70924 |
| GOBP_DEVELOPMENT_OF_PRIMARY_FEMALE_SEXUAL_CHARACTERISTICS | 60 | -0.404007 | -1.018472 | 0.410014 | 0.710651 |
| GOBP_POSITIVE_REGULATION_OF_BLOOD_PRESSURE | 15 | -0.497324 | -1.01843 | 0.448331 | 0.710335 |
| GOBP_REGULATION_OF_PROTEIN_AUTOPHOSPHORYLATION | 39 | -0.425167 | -1.018187 | 0.443038 | 0.710503 |
| GOBP_REGULATION_OF_TRANSMEMBRANE_RECEPTOR_PROTEIN_SERINE_THR | 159 | -0.357875 | -1.017926 | 0.443654 | 0.710682 |
| GOBP_CELL_PROJECTION_ASSEMBLY | 493 | -0.331545 | -1.017668 | 0.42029 | 0.710876 |
| GOBP_BLASTOCYST_DEVELOPMENT | 77 | -0.389482 | -1.017309 | 0.430429 | 0.711299 |
| GOBP_PROTEIN_K48_LINKED_DEUBIQUITINATION | 31 | -0.436121 | -1.017004 | 0.451761 | 0.711602 |
| GOBP_FC_RECEPTOR_SIGNALING_PATHWAY | 188 | -0.355343 | -1.016953 | 0.443169 | 0.71129 |
| GOBP_MESODERM_MORPHOGENESIS | 39 | -0.426952 | -1.016736 | 0.43299 | 0.711349 |
| GOBP_MATERNAL_PROCESS_INVOLVED_IN_FEMALE_PREGNANCY | 36 | -0.428513 | -1.016563 | 0.454023 | 0.711331 |
| GOBP_REGULATION_OF_TOR_SIGNALING | 90 | -0.378716 | -1.016245 | 0.433071 | 0.71166 |
| GOBP_SIGNAL_TRANSDUCTION_IN_ABSENCE_OF_LIGAND | 47 | -0.417398 | -1.015984 | 0.442441 | 0.711824 |
| GOBP_NUCLEAR_TRANSCRIBED_MRNA_CATABOLIC_PROCESS_DEADENYLATIO | 73 | -0.387705 | -1.015796 | 0.448953 | 0.711851 |
| GOBP_POSITIVE_REGULATION_OF_NEURON_APOPTOTIC_PROCESS | 42 | -0.425389 | -1.015183 | 0.437775 | 0.712846 |
| GOBP_GLUTATHIONE_METABOLIC_PROCESS | 44 | -0.410331 | -1.014674 | 0.463768 | 0.713606 |
| GOBP_NEGATIVE_REGULATION_OF_CELLULAR_CATABOLIC_PROCESS | 204 | -0.351443 | -1.01353 | 0.451429 | 0.715887 |
| GOBP_NEGATIVE_REGULATION_OF_PROTEASOMAL_UBIQUITIN_DEPENDENT_PF | 32 | -0.444011 | -1.013089 | 0.441558 | 0.716527 |
| GOBP_REGULATION_OF_BROWN_FAT_CELL_DIFFERENTIATION | 15 | -0.490271 | -1.012794 | 0.460033 | 0.716814 |
| GOBP_REGULATION_OF_DEVELOPMENTAL_GROWTH | 213 | -0.349025 | -1.012608 | 0.448909 | 0.716826 |
| GOBP_MUSCLE_CELL_DEVELOPMENT | 98 | -0.375055 | -1.012215 | 0.40678 | 0.717362 |
| GOBP_CHAPERONE_COFACTOR_DEPENDENT_PROTEIN_REFOLDING | 34 | -0.432778 | -1.011458 | 0.446043 | 0.718665 |
| GOBP_REGULATION_OF_ANION_TRANSMEMBRANE_TRANSPORT | 82 | -0.384245 | -1.011172 | 0.449141 | 0.718886 |
| GOBP_RESPONSE_TO_ACID_CHEMICAL | 86 | -0.38469 | -1.011164 | 0.459279 | 0.718469 |
| GOBP_GLYCOLIPID_BIOSYNTHETIC_PROCESS | 54 | -0.397271 | -1.010922 | 0.439227 | 0.718606 |
| GOBP_RESPONSE_TO_RADIATION | 350 | -0.338082 | -1.010551 | 0.459315 | 0.719081 |
| GOBP_INTRACELLULAR_RECEPTOR_SIGNALING_PATHWAY | 211 | -0.3476 | -1.0104 | 0.449827 | 0.719034 |
| GOBP_POSITIVE_REGULATION_OF_SODIUM_ION_TRANSMEMBRANE_TRANSPOR | 18 | -0.478175 | -1.010265 | 0.45114 | 0.718931 |
| GOBP_STEM_CELL_DIFFERENTIATION | 176 | -0.354083 | -1.009843 | 0.452962 | 0.719516 |
| GOBP_CELLULAR_CARBOHYDRATE_METABOLIC_PROCESS | 223 | -0.347045 | -1.009221 | 0.461364 | 0.720522 |
| GOBP_POSITIVE_REGULATION_OF_TOR_SIGNALING | 39 | -0.424122 | -1.009192 | 0.454545 | 0.720159 |
| GOBP_POSITIVE_REGULATION_OF_CALCIUM_ION_TRANSMEMBRANE_TRANSPO | 40 | -0.419242 | -1.009181 | 0.438316 | 0.719752 |
| GOBP_EPIBOLY | 27 | -0.449208 | -1.00913 | 0.459618 | 0.719422 |
| GOBP_I_KAPPAB_KINASE_NF_KAPPAB_SIGNALING | 244 | -0.34354 | -1.009084 | 0.462754 | 0.7191 |
| GOBP_GAMETE_GENERATION | 437 | -0.332847 | -1.008877 | 0.455882 | 0.719134 |
| GOBP_TISSUE_REGENERATION | 48 | -0.419023 | -1.008488 | 0.453731 | 0.719589 |
| GOBP_EPITHELIAL_CELL_DEVELOPMENT | 141 | -0.359065 | -1.008454 | 0.453892 | 0.719233 |
| GOBP_REGULATION_OF_EPITHELIAL_CELL_APOPTOTIC_PROCESS | 52 | -0.402569 | -1.008431 | 0.454294 | 0.718862 |
| GOBP_ATP_BIOSYNTHETIC_PROCESS | 49 | -0.406322 | -1.008131 | 0.446602 | 0.719167 |
| GOBP_POSITIVE_REGULATION_OF_T_CELL_MEDIATED_IMMUNITY | 43 | -0.419846 | -1.007757 | 0.478261 | 0.719632 |
| GOBP_VESICLE_LOCALIZATION | 191 | -0.349726 | -1.007566 | 0.445358 | 0.719672 |
| GOBP_REGULATION_OF_NUCLEOTIDE_METABOLIC_PROCESS | 94 | -0.375571 | -1.006869 | 0.483161 | 0.720828 |
| GOBP_HORMONE_TRANSPORT | 203 | -0.346034 | -1.006844 | 0.468822 | 0.720461 |
| GOBP_HUMORAL_IMMUNE_RESPONSE_MEDIATED_BY_CIRCULATING_IMMUNOG | 70 | -0.388686 | -1.006545 | 0.455497 | 0.720738 |
| GOBP_REGULATION_OF_SMALL_MOLECULE_METABOLIC_PROCESS | 341 | -0.336874 | -1.006214 | 0.445629 | 0.721102 |
| GOBP_METANEPHROS_MORPHOGENESIS | 16 | -0.48342 | -1.006099 | 0.466667 | 0.720942 |
| GOBP_NEGATIVE_REGULATION_OF_DEPHOSPHORYLATION | 84 | -0.379712 | -1.005888 | 0.431498 | 0.721051 |
| GOBP_ESTABLISHMENT_OF_CELL_POLARITY | 108 | -0.366946 | -1.005798 | 0.448669 | 0.720816 |
| GOBP_NOTCH_SIGNALING_PATHWAY | 124 | -0.364284 | -1.005515 | 0.449631 | 0.721048 |
| GOBP_POSITIVE_REGULATION_OF_MAP_KINASE_ACTIVITY | 174 | -0.348493 | -1.005347 | 0.46083 | 0.721029 |
| GOBP_POSITIVE_REGULATION_OF_ENDOTHELIAL_CELL_PROLIFERATION | 63 | -0.391994 | -1.004971 | 0.456693 | 0.721488 |
| GOBP_CELLULAR_RESPONSE_TO_LIGHT_STIMULUS | 101 | -0.36842 | -1.004693 | 0.44171 | 0.721702 |
| GOBP_NATURAL_KILLER_CELL_DIFFERENTIATION | 15 | -0.495027 | -1.004342 | 0.483389 | 0.722147 |
| GOBP_PROTEIN_TRANSPORT_ALONG_MICROTUBULE | 65 | -0.393996 | -1.004283 | 0.455041 | 0.721851 |
| GOBP_RESPONSE_TO_INCREASED_OXYGEN_LEVELS | 25 | -0.449778 | -1.004183 | 0.475969 | 0.72165 |
| GOBP_SUBSTRATE_DEPENDENT_CELL_MIGRATION | 17 | -0.470205 | -1.003319 | 0.483444 | 0.723265 |
| GOBP_NEGATIVE_REGULATION_OF_PROTEOLYSIS_INVOLVED_IN_CELLULAR_PI | 56 | -0.404596 | -1.001447 | 0.444288 | 0.727238 |
| GOBP_CELLULAR_MODIFIED_AMINO_ACID_BIOSYNTHETIC_PROCESS | 41 | -0.414036 | -1.00138 | 0.463483 | 0.726965 |
| GOBP_TRANSITION_METAL_ION_HOMEOSTASIS | 108 | -0.368073 | -1.001337 | 0.456821 | 0.726629 |
| GOBP_NEUROTRANSMITTER_TRANSPORT | 126 | -0.360487 | -1.00085 | 0.479012 | 0.727354 |
| GOBP_CELLULAR_RESPONSE_TO_HORMONE_STIMULUS | 436 | -0.328157 | -1.00083 | 0.484785 | 0.726975 |
| GOBP_DNA_GEOMETRIC_CHANGE | 111 | -0.366936 | -0.999445 | 0.468165 | 0.729768 |
| GOBP_RESPONSE_TO_PEPTIDE_HORMONE | 321 | -0.336324 | -0.998712 | 0.4791 | 0.731077 |
| GOBP_VESICLE_DOCKING | 57 | -0.395416 | -0.99858 | 0.458564 | 0.730939 |
| GOBP_OLIGODENDROCYTE_DIFFERENTIATION | 55 | -0.397738 | -0.998157 | 0.476839 | 0.731476 |
| GOBP_REGULATION_OF_CELLULAR_AMINO_ACID_METABOLIC_PROCESS | 56 | -0.394244 | -0.99749 | 0.481119 | 0.732586 |
| GOBP_CELL_MATURATION | 105 | -0.366658 | -0.997319 | 0.480154 | 0.732557 |
| GOBP_REGULATION_OF_NUCLEOCYTOPLASMIC_TRANSPORT | 90 | -0.377051 | -0.997227 | 0.472513 | 0.732336 |
| GOBP_NEGATIVE_REGULATION_OF_CYCLIN_DEPENDENT_PROTEIN_KINASE_AC | 28 | -0.4449 | -0.997184 | 0.490284 | 0.732001 |
| GOBP_POSITIVE_REGULATION_OF_B_CELL_PROLIFERATION | 33 | -0.427264 | -0.997054 | 0.462222 | 0.731895 |
| GOBP_PLASMA_MEMBRANE_TUBULATION | 17 | -0.476859 | -0.99698 | 0.494272 | 0.731643 |
| GOBP_REGULATION_OF_PEPTIDE_HORMONE_SECRETION | 139 | -0.357318 | -0.996063 | 0.465455 | 0.733417 |
| GOBP_COPII_COATED_VESICLE_BUDDING | 67 | -0.382978 | -0.996008 | 0.479361 | 0.733131 |
| GOBP_MODULATION_BY_HOST_OF_VIRAL_PROCESS | 25 | -0.450513 | -0.995139 | 0.475113 | 0.734796 |
| GOBP_POSITIVE_REGULATION_BY_HOST_OF_VIRAL_TRANSCRIPTION | 15 | -0.477661 | -0.995092 | 0.459099 | 0.734479 |
| GOBP_CARBOHYDRATE_METABOLIC_PROCESS | 493 | -0.327863 | -0.99504 | 0.49793 | 0.734174 |
| GOBP_CYTOSOLIC_TRANSPORT | 153 | -0.351019 | -0.994474 | 0.481743 | 0.735062 |
| GOBP_NEGATIVE_REGULATION_OF_NIK_NF_KAPPAB_SIGNALING | 18 | -0.472058 | -0.994417 | 0.494436 | 0.734766 |
| GOBP_CANONICAL_WNT_SIGNALING_PATHWAY | 245 | -0.339294 | -0.994349 | 0.501116 | 0.734489 |
| GOBP_REGULATION_OF_DENDRITE_DEVELOPMENT | 73 | -0.376026 | -0.993979 | 0.475676 | 0.73494 |
| GOBP_INFLAMMATORY_RESPONSE_TO_ANTIGENIC_STIMULUS | 41 | -0.41263 | -0.992828 | 0.469846 | 0.737158 |
| GOBP_GLUTAMATE_RECEPTOR_SIGNALING_PATHWAY | 30 | -0.431872 | -0.992496 | 0.483013 | 0.737471 |
| GOBP_POSITIVE_REGULATION_OF_CELLULAR_CARBOHYDRATE_METABOLIC_PI | 37 | -0.426877 | -0.992298 | 0.482808 | 0.737505 |
| GOBP_UNSATURATED_FATTY_ACID_BIOSYNTHETIC_PROCESS | 31 | -0.430764 | -0.991673 | 0.479821 | 0.738484 |
| GOBP_PHOSPHOLIPID_METABOLIC_PROCESS | 340 | -0.329471 | -0.991591 | 0.492358 | 0.738243 |
| GOBP_POSITIVE_REGULATION_OF_PROTEIN_ACETYLATION | 38 | -0.418932 | -0.991556 | 0.483099 | 0.737897 |
| GOBP_SYNAPSE_MATURATION | 16 | -0.483794 | -0.991351 | 0.4768 | 0.737941 |
| GOBP_REGULATION_OF_MUSCLE_ORGAN_DEVELOPMENT | 26 | -0.44129 | -0.991345 | 0.454128 | 0.737525 |
| GOBP_GLOMERULUS_DEVELOPMENT | 39 | -0.417664 | -0.990364 | 0.479109 | 0.739378 |
| GOBP_RESPONSE_TO_ESTROGEN | 50 | -0.403885 | -0.990284 | 0.453061 | 0.739124 |
| GOBP_HYDROGEN_PEROXIDE_METABOLIC_PROCESS | 33 | -0.425028 | -0.990037 | 0.474474 | 0.739262 |
| GOBP_LYSOSOMAL_TRANSPORT | 100 | -0.37359 | -0.990022 | 0.490272 | 0.738866 |
| GOBP_NEGATIVE_REGULATION_OF_PROTEIN_CONTAINING_COMPLEX_DISASSE | 64 | -0.387269 | -0.989798 | 0.477178 | 0.738943 |
| GOBP_REGULATION_OF_DNA_DAMAGE_RESPONSE_SIGNAL_TRANSDUCTION_B | 27 | -0.43592 | -0.98911 | 0.473101 | 0.740143 |
| GOBP_CILIARY_BASAL_BODY_PLASMA_MEMBRANE_DOCKING | 95 | -0.359405 | -0.98774 | 0.49017 | 0.74292 |
| GOBP_T_CELL_MIGRATION | 50 | -0.40441 | -0.987493 | 0.479433 | 0.743087 |
| GOBP_FATTY_ACID_METABOLIC_PROCESS | 283 | -0.332258 | -0.987208 | 0.510544 | 0.743313 |
| GOBP_PROTEIN_FOLDING | 205 | -0.341065 | -0.986877 | 0.491972 | 0.74364 |
| GOBP_ORGANELLE_FUSION | 126 | -0.35903 | -0.985882 | 0.492629 | 0.745508 |
| GOBP_NEGATIVE_REGULATION_OF_CELL_JUNCTION_ASSEMBLY | 24 | -0.448781 | -0.985299 | 0.479199 | 0.746427 |
| GOBP_REGULATION_OF_ORGANELLE_ASSEMBLY | 165 | -0.34365 | -0.984893 | 0.48284 | 0.746965 |
| GOBP_STRESS_ACTIVATED_PROTEIN_KINASE_SIGNALING_CASCADE | 226 | -0.33823 | -0.983659 | 0.511973 | 0.749519 |
| GOBP_POSITIVE_REGULATION_OF_MONONUCLEAR_CELL_MIGRATION | 47 | -0.392665 | -0.982869 | 0.485673 | 0.750939 |
| GOBP_RECEPTOR_CATABOLIC_PROCESS | 34 | -0.42602 | -0.982639 | 0.494118 | 0.751029 |
| GOBP_REGULATION_OF_SYSTEMIC_ARTERIAL_BLOOD_PRESSURE_BY_HORMO | 19 | -0.472376 | -0.982542 | 0.514754 | 0.750839 |
| GOBP_INSULIN_SECRETION | 142 | -0.349099 | -0.9817 | 0.507426 | 0.752301 |
| GOBP_VESICLE_DOCKING_INVOLVED_IN_EXOCYTOSIS | 40 | -0.414901 | -0.981284 | 0.506456 | 0.752808 |
| GOBP_ENDOCRINE_PROCESS | 44 | -0.399764 | -0.981085 | 0.507396 | 0.752852 |
| GOBP_SECONDARY_METABOLIC_PROCESS | 36 | -0.418932 | -0.980607 | 0.502203 | 0.753476 |
| GOBP_POSITIVE_REGULATION_OF_PEPTIDE_HORMONE_SECRETION | 61 | -0.389419 | -0.979794 | 0.480315 | 0.754932 |
| GOBP_POSITIVE_REGULATION_OF_LIPID_KINASE_ACTIVITY | 27 | -0.435556 | -0.979463 | 0.495549 | 0.755286 |
| GOBP_POSITIVE_REGULATION_OF_PROTEIN_EXPORT_FROM_NUCLEUS | 20 | -0.456448 | -0.978339 | 0.514241 | 0.757512 |
| GOBP_PHOSPHOLIPID_TRANSPORT | 69 | -0.386854 | -0.978325 | 0.489851 | 0.757117 |
| GOBP_CELLULAR_COMPONENT_ASSEMBLY_INVOLVED_IN_MORPHOGENESIS | 67 | -0.381984 | -0.978037 | 0.498013 | 0.757343 |
| GOBP_STEROID_HORMONE_BIOSYNTHETIC_PROCESS | 29 | -0.425382 | -0.977363 | 0.495441 | 0.758414 |
| GOBP_LIPOSACCHARIDE_METABOLIC_PROCESS | 92 | -0.364719 | -0.977305 | 0.494751 | 0.758124 |
| GOBP_NEGATIVE_REGULATION_OF_REGULATED_SECRETORY_PATHWAY | 18 | -0.462531 | -0.977179 | 0.518293 | 0.757981 |
| GOBP_CELL_FATE_DETERMINATION | 20 | -0.456168 | -0.975165 | 0.50234 | 0.762248 |
| GOBP_MALE_GAMETE_GENERATION | 349 | -0.327224 | -0.974928 | 0.538713 | 0.762354 |
| GOBP_PROTEIN_MANNOSYLATION | 22 | -0.443236 | -0.974572 | 0.496885 | 0.762714 |
| GOBP_BENZENE_CONTAINING_COMPOUND_METABOLIC_PROCESS | 17 | -0.462745 | -0.974548 | 0.517028 | 0.762334 |
| GOBP_VITAMIN_BIOSYNTHETIC_PROCESS | 15 | -0.489651 | -0.974179 | 0.520767 | 0.762713 |
| GOBP_MEMBRANE_DOCKING | 170 | -0.341095 | -0.973879 | 0.493552 | 0.763008 |
| GOBP_REGULATION_OF_INFLAMMATORY_RESPONSE_TO_ANTIGENIC_STIMULUS | 24 | -0.441445 | -0.973769 | 0.513932 | 0.762827 |
| GOBP_HINDBRAIN_MORPHOGENESIS | 27 | -0.433814 | -0.973383 | 0.515198 | 0.763231 |
| GOBP_POSITIVE_REGULATION_OF_I_KAPPAB_KINASE_NF_KAPPAB_SIGNALING | 158 | -0.343494 | -0.973252 | 0.522209 | 0.763105 |
| GOBP_POSITIVE_REGULATION_OF_SODIUM_ION_TRANSMEMBRANE_TRANSPOR | 15 | -0.480914 | -0.973112 | 0.519608 | 0.763024 |
| GOBP_TISSUE_HOMEOSTASIS | 177 | -0.338868 | -0.972934 | 0.526995 | 0.762989 |
| GOBP_MECHANORECEPTOR_DIFFERENTIATION | 37 | -0.408548 | -0.971952 | 0.51073 | 0.764875 |
| GOBP_ERYTHROCYTE_HOMEOSTASIS | 106 | -0.356837 | -0.971797 | 0.503145 | 0.764784 |
| GOBP_RNA_DESTABILIZATION | 32 | -0.420035 | -0.971698 | 0.504373 | 0.764574 |
| GOBP_REGULATION_OF_PROTEIN_CATABOLIC_PROCESS | 328 | -0.327205 | -0.971683 | 0.549403 | 0.764173 |
| GOBP_POSITIVE_REGULATION_OF_GLUCOSE_METABOLIC_PROCESS | 31 | -0.417583 | -0.970071 | 0.509517 | 0.76747 |
| GOBP_LYTIC_VACUOLE_ORGANIZATION | 62 | -0.387846 | -0.96988 | 0.525815 | 0.767478 |
| GOBP_GLYCOSPHINGOLIPID_METABOLIC_PROCESS | 53 | -0.38538 | -0.969676 | 0.507824 | 0.76751 |
| GOBP_INTERLEUKIN_17_PRODUCTION | 27 | -0.437001 | -0.969069 | 0.53973 | 0.768449 |
| GOBP_REGULATION_OF_COMPLEMENT_ACTIVATION | 51 | -0.388097 | -0.968981 | 0.502755 | 0.768213 |

| | | | | | |
|---|---|---|---|---|---|
| GOBP_RECEPTOR_INTERNALIZATION | 77 | -0.372751 | -0.968955 | 0.502618 | 0.767836 |
| GOBP_REGULATION_OF_CELLULAR_KETONE_METABOLIC_PROCESS | 142 | -0.346585 | -0.968894 | 0.517923 | 0.76755 |
| GOBP_REGULATION_OF_EXECUTION_PHASE_OF_APOPTOSIS | 15 | -0.468349 | -0.968739 | 0.513245 | 0.767476 |
| GOBP_SPECIFICATION_OF_SYMMETRY | 76 | -0.365367 | -0.968055 | 0.503303 | 0.768659 |
| GOBP_REGULATION_OF_ESTABLISHMENT_OF_PROTEIN_LOCALIZATION_TO_MIT | 69 | -0.37445 | -0.967786 | 0.516043 | 0.768788 |
| GOBP_OLIGOSACCHARIDE_METABOLIC_PROCESS | 38 | -0.407937 | -0.967568 | 0.499274 | 0.768846 |
| GOBP_MONOAMINE_TRANSPORT | 37 | -0.396519 | -0.966132 | 0.502236 | 0.771667 |
| GOBP_LIPID_MODIFICATION | 223 | -0.329905 | -0.966093 | 0.54504 | 0.771324 |
| GOBP_MYOTUBE_CELL_DEVELOPMENT | 23 | -0.430358 | -0.96587 | 0.508346 | 0.771434 |
| GOBP_POSITIVE_REGULATION_OF_DNA_METABOLIC_PROCESS | 172 | -0.338038 | -0.965817 | 0.531435 | 0.771122 |
| GOBP_NEGATIVE_REGULATION_OF_CELL_AGING | 20 | -0.444301 | -0.965581 | 0.535658 | 0.771201 |
| GOBP_POSITIVE_REGULATION_OF_CELLULAR_AMIDE_METABOLIC_PROCESS | 138 | -0.347485 | -0.965061 | 0.51986 | 0.771921 |
| GOBP_NADP_METABOLIC_PROCESS | 34 | -0.415348 | -0.964082 | 0.519481 | 0.773723 |
| GOBP_POSITIVE_REGULATION_OF_EPITHELIAL_CELL_PROLIFERATION | 117 | -0.349811 | -0.963853 | 0.534678 | 0.773812 |
| GOBP_REGULATION_OF_HUMORAL_IMMUNE_RESPONSE | 63 | -0.371612 | -0.963552 | 0.521246 | 0.774059 |
| GOBP_POSITIVE_REGULATION_OF_MUSCLE_HYPERTROPHY | 19 | -0.453735 | -0.963365 | 0.543831 | 0.774047 |
| GOBP_CELL_COMMUNICATION_BY_ELECTRICAL_COUPLING | 18 | -0.459407 | -0.961739 | 0.515599 | 0.777369 |
| GOBP_INOSITOL_PHOSPHATE_BIOSYNTHETIC_PROCESS | 17 | -0.457485 | -0.961708 | 0.508885 | 0.777008 |
| GOBP_BINDING_OF_SPERM_TO_ZONA_PELLUCIDA | 24 | -0.423492 | -0.960413 | 0.531646 | 0.779515 |
| GOBP_CELL_AGING | 88 | -0.359755 | -0.959733 | 0.526115 | 0.780609 |
| GOBP_VESICLE_MEDIATED_TRANSPORT_BETWEEN_ENDOSOMAL_COMPARTME | 41 | -0.400722 | -0.959135 | 0.512744 | 0.781518 |
| GOBP_NEGATIVE_REGULATION_OF_STRIATED_MUSCLE_CELL_APOPTOTIC_PRO | 15 | -0.467506 | -0.959089 | 0.513732 | 0.781193 |
| GOBP_ANTIGEN_PROCESSING_AND_PRESENTATION_OF_ENDOGENOUS_PEPTI | 18 | -0.459772 | -0.959003 | 0.52496 | 0.78095 |
| GOBP_EXTRACELLULAR_VESICLE_BIOGENESIS | 20 | -0.459236 | -0.958947 | 0.54372 | 0.780626 |
| GOBP_REGULATION_OF_T_HELPER_17_TYPE_IMMUNE_RESPONSE | 16 | -0.46372 | -0.958587 | 0.507911 | 0.781014 |
| GOBP_CELLULAR_ALDEHYDE_METABOLIC_PROCESS | 45 | -0.3892 | -0.957671 | 0.53662 | 0.782621 |
| GOBP_POSITIVE_REGULATION_OF_CELL_DIVISION | 50 | -0.382817 | -0.957608 | 0.536903 | 0.78233 |
| GOBP_PHOSPHOLIPID_BIOSYNTHETIC_PROCESS | 228 | -0.328991 | -0.957417 | 0.573059 | 0.782329 |
| GOBP_NUCLEOBASE_CONTAINING_SMALL_MOLECULE_INTERCONVERSION | 25 | -0.429981 | -0.957255 | 0.528951 | 0.782249 |
| GOBP_RESPONSE_TO_SALT_STRESS | 17 | -0.459673 | -0.957018 | 0.532154 | 0.782345 |
| GOBP_REGULATION_OF_EXTRINSIC_APOPTOTIC_SIGNALING_PATHWAY_IN_ABS | 26 | -0.42595 | -0.956748 | 0.532362 | 0.782569 |
| GOBP_DIGESTION | 62 | -0.375628 | -0.956462 | 0.513699 | 0.782766 |
| GOBP_PHENOL_CONTAINING_COMPOUND_BIOSYNTHETIC_PROCESS | 24 | -0.433482 | -0.955916 | 0.525502 | 0.783593 |
| GOBP_DEVELOPMENTAL_MATURATION | 179 | -0.337121 | -0.955761 | 0.558167 | 0.783512 |
| GOBP_SOMATIC_DIVERSIFICATION_OF_IMMUNOGLOBULINS | 62 | -0.372473 | -0.955408 | 0.537234 | 0.783885 |
| GOBP_REGULATION_OF_MYOTUBE_DIFFERENTIATION | 25 | -0.429069 | -0.954221 | 0.540146 | 0.786141 |
| GOBP_REGULATION_OF_SMAD_PROTEIN_SIGNAL_TRANSDUCTION | 20 | -0.44545 | -0.953917 | 0.565916 | 0.786368 |
| GOBP_NEGATIVE_REGULATION_OF_TELOMERE_MAINTENANCE_VIA_TELOMERA | 19 | -0.450639 | -0.953725 | 0.548536 | 0.786352 |
| GOBP_SKELETAL_MUSCLE_TISSUE_REGENERATION | 29 | -0.420887 | -0.953103 | 0.553223 | 0.787334 |
| GOBP_REGULATION_OF_RESPONSE_TO_TUMOR_CELL | 16 | -0.462924 | -0.952547 | 0.539867 | 0.788126 |
| GOBP_MAST_CELL_ACTIVATION_INVOLVED_IN_IMMUNE_RESPONSE | 37 | -0.405397 | -0.950913 | 0.533724 | 0.791397 |
| GOBP_MAINTENANCE_OF_PROTEIN_LOCATION_IN_CELL | 57 | -0.376862 | -0.950282 | 0.557312 | 0.792362 |
| GOBP_REGULATION_OF_CHONDROCYTE_DIFFERENTIATION | 23 | -0.429589 | -0.950215 | 0.530864 | 0.792077 |
| GOBP_GLUTAMINE_FAMILY_AMINO_ACID_METABOLIC_PROCESS | 60 | -0.378555 | -0.950111 | 0.545328 | 0.791891 |
| GOBP_SEGMENTATION | 54 | -0.381738 | -0.949011 | 0.554017 | 0.79398 |
| GOBP_MITOTIC_G2_DNA_DAMAGE_CHECKPOINT | 21 | -0.433742 | -0.948902 | 0.545455 | 0.793797 |
| GOBP_CEREBRAL_CORTEX_DEVELOPMENT | 74 | -0.366213 | -0.948512 | 0.54244 | 0.794229 |
| GOBP_POSITIVE_REGULATION_OF_DNA_REPLICATION | 32 | -0.408228 | -0.947941 | 0.566358 | 0.795057 |
| GOBP_SLEEP | 15 | -0.460543 | -0.947209 | 0.524126 | 0.796214 |
| GOBP_REGULATION_OF_PROTEIN_DEPOLYMERIZATION | 64 | -0.373572 | -0.947147 | 0.541667 | 0.795911 |
| GOBP_SENSORY_PERCEPTION_OF_TASTE | 29 | -0.411199 | -0.946291 | 0.541056 | 0.79744 |
| GOBP_REGULATION_OF_SODIUM_ION_TRANSMEMBRANE_TRANSPORTER_ACTI | 38 | -0.393936 | -0.945943 | 0.529155 | 0.79778 |
| GOBP_POSITIVE_REGULATION_OF_ORGANELLE_ASSEMBLY | 59 | -0.378058 | -0.945396 | 0.562753 | 0.798574 |
| GOBP_INACTIVATION_OF_MAPK_ACTIVITY | 23 | -0.429919 | -0.945154 | 0.550861 | 0.798673 |
| GOBP_POLYOL_METABOLIC_PROCESS | 104 | -0.348404 | -0.944166 | 0.569604 | 0.800469 |
| GOBP_REGULATION_OF_NEURONAL_SYNAPTIC_PLASTICITY | 35 | -0.403278 | -0.944088 | 0.528963 | 0.800209 |
| GOBP_ORGAN_OR_TISSUE_SPECIFIC_IMMUNE_RESPONSE | 17 | -0.444729 | -0.94374 | 0.534733 | 0.800517 |
| GOBP_REGULATION_OF_LIPOPOLYSACCHARIDE_MEDIATED_SIGNALING_PATHW | 19 | -0.434264 | -0.942705 | 0.56536 | 0.802423 |
| GOBP_NEGATIVE_REGULATION_OF_CELL_CYCLE_PROCESS | 305 | -0.316099 | -0.941511 | 0.610629 | 0.804567 |
| GOBP_ADRENERGIC_RECEPTOR_SIGNALING_PATHWAY | 15 | -0.473596 | -0.940727 | 0.552013 | 0.805879 |
| GOBP_NEGATIVE_REGULATION_OF_PROTEIN_BINDING | 80 | -0.363301 | -0.940635 | 0.546296 | 0.805644 |
| GOBP_REGULATION_OF_PROTEIN_SERINE_THREONINE_KINASE_ACTIVITY | 390 | -0.312166 | -0.940299 | 0.618997 | 0.805959 |
| GOBP_REGULATION_OF_CELL_CYCLE_PHASE_TRANSITION | 411 | -0.309876 | -0.939947 | 0.636173 | 0.806302 |
| GOBP_STEROID_HORMONE_MEDIATED_SIGNALING_PATHWAY | 101 | -0.344965 | -0.93859 | 0.571063 | 0.808873 |
| GOBP_REGULATION_OF_STRESS_ACTIVATED_PROTEIN_KINASE_SIGNALING_CA | 146 | -0.334505 | -0.938517 | 0.584816 | 0.808593 |
| GOBP_REGULATION_OF_ERYTHROCYTE_DIFFERENTIATION | 42 | -0.389567 | -0.937403 | 0.545455 | 0.810584 |
| GOBP_RESPONSE_TO_BMP | 95 | -0.349157 | -0.937019 | 0.561006 | 0.810975 |
| GOBP_OVARIAN_FOLLICLE_DEVELOPMENT | 39 | -0.392113 | -0.935913 | 0.553009 | 0.812974 |
| GOBP_REGULATION_OF_LAMELLIPODIUM_ORGANIZATION | 44 | -0.383707 | -0.935648 | 0.556164 | 0.813136 |
| GOBP_INTRACELLULAR_STEROID_HORMONE_RECEPTOR_SIGNALING_PATHWA | 87 | -0.355175 | -0.935565 | 0.563218 | 0.81287 |
| GOBP_SELECTIVE_AUTOPHAGY | 59 | -0.367752 | -0.935363 | 0.554348 | 0.812875 |
| GOBP_REGULATION_OF_POTASSIUM_ION_TRANSMEMBRANE_TRANSPORTER_A | 36 | -0.391984 | -0.93519 | 0.552174 | 0.81278 |
| GOBP_GLYCEROPHOSPHOLIPID_METABOLIC_PROCESS | 292 | -0.316294 | -0.934924 | 0.629464 | 0.81294 |
| GOBP_MODULATION_BY_HOST_OF_SYMBIONT_PROCESS | 50 | -0.375648 | -0.934702 | 0.559322 | 0.812981 |
| GOBP_POSITIVE_REGULATION_OF_PROTEIN_SERINE_THREONINE_KINASE_ACT | 243 | -0.320311 | -0.934689 | 0.617813 | 0.812569 |
| GOBP_STRIATED_MUSCLE_CONTRACTION | 96 | -0.345196 | -0.934675 | 0.58125 | 0.81215 |
| GOBP_REGULATION_OF_PROTEASOMAL_PROTEIN_CATABOLIC_PROCESS | 161 | -0.331205 | -0.934088 | 0.613744 | 0.813021 |
| GOBP_NEGATIVE_REGULATION_OF_INTRACELLULAR_STEROID_HORMONE_REC | 27 | -0.418076 | -0.933981 | 0.553846 | 0.812811 |
| GOBP_INTRINSIC_APOPTOTIC_SIGNALING_PATHWAY_IN_RESPONSE_TO_DNA_D | 90 | -0.352211 | -0.933407 | 0.569211 | 0.813651 |
| GOBP_REGULATION_OF_MUSCLE_SYSTEM_PROCESS | 144 | -0.333587 | -0.93307 | 0.590629 | 0.814017 |
| GOBP_REGULATION_OF_NUCLEAR_DIVISION | 107 | -0.344919 | -0.93302 | 0.563131 | 0.813674 |
| GOBP_RESPONSE_TO_EPIDERMAL_GROWTH_FACTOR | 37 | -0.392368 | -0.932609 | 0.564328 | 0.814183 |
| GOBP_CORTICAL_ACTIN_CYTOSKELETON_ORGANIZATION | 33 | -0.402902 | -0.932581 | 0.554545 | 0.813806 |
| GOBP_HORMONE_MEDIATED_SIGNALING_PATHWAY | 131 | -0.335302 | -0.932087 | 0.602706 | 0.814489 |
| GOBP_POSITIVE_REGULATION_OF_PROTEIN_CATABOLIC_PROCESS | 195 | -0.324067 | -0.93057 | 0.609195 | 0.817399 |
| GOBP_REGULATION_OF_CARBOHYDRATE_METABOLIC_PROCESS | 162 | -0.328328 | -0.930426 | 0.619277 | 0.817262 |
| GOBP_PROTEOGLYCAN_METABOLIC_PROCESS | 54 | -0.376747 | -0.930295 | 0.578501 | 0.817085 |
| GOBP_EAR_DEVELOPMENT | 111 | -0.34386 | -0.930289 | 0.576923 | 0.816655 |
| GOBP_REGULATION_OF_PROTON_TRANSPORT | 20 | -0.432785 | -0.930281 | 0.557947 | 0.81623 |
| GOBP_CELLULAR_RESPONSE_TO_PEPTIDE_HORMONE_STIMULUS | 243 | -0.319785 | -0.930251 | 0.617117 | 0.81586 |
| GOBP_SPLEEN_DEVELOPMENT | 22 | -0.426076 | -0.930176 | 0.580997 | 0.815573 |
| GOBP_NEGATIVE_REGULATION_OF_INTRINSIC_APOPTOTIC_SIGNALING_PATHW | 25 | -0.41844 | -0.929632 | 0.578009 | 0.816325 |
| GOBP_REGULATION_OF_MUSCLE_CONTRACTION | 93 | -0.347178 | -0.929164 | 0.582908 | 0.816895 |
| GOBP_IRON_ION_TRANSMEMBRANE_TRANSPORT | 17 | -0.441741 | -0.928701 | 0.57189 | 0.817479 |
| GOBP_PHOSPHATIDYLINOSITOL_BIOSYNTHETIC_PROCESS | 105 | -0.340645 | -0.927599 | 0.595985 | 0.819429 |
| GOBP_NEGATIVE_REGULATION_OF_TRANSLATIONAL_INITIATION | 19 | -0.432225 | -0.927289 | 0.568038 | 0.819683 |
| GOBP_NEGATIVE_REGULATION_OF_DNA_DEPENDENT_DNA_REPLICATION | 17 | -0.450218 | -0.926908 | 0.558029 | 0.820082 |
| GOBP_PRESYNAPSE_ORGANIZATION | 22 | -0.431671 | -0.926627 | 0.589984 | 0.820213 |
| GOBP_POSITIVE_REGULATION_OF_SMOOTH_MUSCLE_CONTRACTION | 17 | -0.439053 | -0.92593 | 0.569767 | 0.82128 |
| GOBP_CELLULAR_RESPONSE_TO_CARBOHYDRATE_STIMULUS | 102 | -0.348509 | -0.925909 | 0.581218 | 0.82088 |
| GOBP_PYRIMIDINE_NUCLEOSIDE_CATABOLIC_PROCESS | 17 | -0.437848 | -0.925877 | 0.557845 | 0.820503 |
| GOBP_AXIS_SPECIFICATION | 42 | -0.38097 | -0.925866 | 0.571633 | 0.820092 |
| GOBP_METANEPHROS_DEVELOPMENT | 38 | -0.388388 | -0.925688 | 0.578279 | 0.820036 |
| GOBP_MIRNA_METABOLIC_PROCESS | 26 | -0.418233 | -0.925637 | 0.565354 | 0.819707 |
| GOBP_PIGMENTATION | 69 | -0.356799 | -0.925615 | 0.571238 | 0.819322 |
| GOBP_DICARBOXYLIC_ACID_TRANSPORT | 60 | -0.360968 | -0.924944 | 0.569149 | 0.820321 |
| GOBP_STEM_CELL_DIVISION | 18 | -0.436986 | -0.924849 | 0.563981 | 0.820082 |
| GOBP_CELLULAR_RESPONSE_TO_OXYGEN_RADICAL | 18 | -0.447254 | -0.924786 | 0.590985 | 0.819776 |
| GOBP_POSITIVE_REGULATION_OF_CATABOLIC_PROCESS | 382 | -0.30941 | -0.92463 | 0.683544 | 0.819685 |
| GOBP_LONG_CHAIN_FATTY_ACID_METABOLIC_PROCESS | 80 | -0.347665 | -0.924347 | 0.588235 | 0.819887 |
| GOBP_VESICLE_TARGETING_TO_FROM_OR_WITHIN_GOLGI | 68 | -0.366776 | -0.924258 | 0.572414 | 0.819641 |
| GOBP_EMBRYONIC_PATTERN_SPECIFICATION | 30 | -0.401549 | -0.923855 | 0.552511 | 0.820092 |
| GOBP_REGULATION_OF_GLUCOSE_TRANSMEMBRANE_TRANSPORT | 53 | -0.374489 | -0.923248 | 0.578947 | 0.820959 |
| GOBP_MEMBRANE_DEPOLARIZATION | 55 | -0.364741 | -0.921954 | 0.574409 | 0.823419 |
| GOBP_CHROMATIN_SILENCING | 51 | -0.368365 | -0.920949 | 0.572022 | 0.825102 |
| GOBP_MONOCARBOXYLIC_ACID_BIOSYNTHETIC_PROCESS | 149 | -0.327343 | -0.920154 | 0.624402 | 0.826403 |
| GOBP_RESPONSE_TO_INTERLEUKIN_12 | 45 | -0.376112 | -0.919593 | 0.568871 | 0.827189 |
| GOBP_RESPONSE_TO_OXYGEN_RADICAL | 22 | -0.416008 | -0.918889 | 0.567485 | 0.828232 |
| GOBP_PROTEOGLYCAN_BIOSYNTHETIC_PROCESS | 41 | -0.385963 | -0.918592 | 0.587209 | 0.828427 |
| GOBP_CARTILAGE_DEVELOPMENT | 98 | -0.339298 | -0.917537 | 0.621967 | 0.83027 |
| GOBP_CILIUM_MOVEMENT | 96 | -0.345288 | -0.917036 | 0.603581 | 0.830886 |
| GOBP_DE_NOVO_PROTEIN_FOLDING | 43 | -0.375134 | -0.916991 | 0.578561 | 0.830536 |
| GOBP_C21_STEROID_HORMONE_BIOSYNTHETIC_PROCESS | 16 | -0.429776 | -0.916891 | 0.538591 | 0.83032 |
| GOBP_PYRUVATE_METABOLIC_PROCESS | 123 | -0.330812 | -0.916608 | 0.606396 | 0.830481 |
| GOBP_NEGATIVE_REGULATION_OF_PROTEIN_POLYMERIZATION | 59 | -0.35919 | -0.916159 | 0.586835 | 0.831063 |
| GOBP_CARBOHYDRATE_TRANSPORT | 105 | -0.339681 | -0.91563 | 0.605634 | 0.831757 |
| GOBP_NEGATIVE_REGULATION_OF_ACTIN_FILAMENT_POLYMERIZATION | 49 | -0.367501 | -0.915465 | 0.579832 | 0.83168 |
| GOBP_DNA_DEPENDENT_DNA_REPLICATION_MAINTENANCE_OF_FIDELITY | 46 | -0.375071 | -0.915393 | 0.592806 | 0.831392 |
| GOBP_CORTICAL_CYTOSKELETON_ORGANIZATION | 48 | -0.369071 | -0.913126 | 0.569832 | 0.835768 |
| GOBP_OOCYTE_MATURATION | 20 | -0.423895 | -0.91287 | 0.578462 | 0.835867 |
| GOBP_TYPE_I_INTERFERON_PRODUCTION | 122 | -0.333296 | -0.912475 | 0.618582 | 0.836278 |
| GOBP_INNER_EAR_RECEPTOR_CELL_STEREOCILIUM_ORGANIZATION | 22 | -0.420068 | -0.911189 | 0.598784 | 0.83855 |
| GOBP_SPHINGOLIPID_BIOSYNTHETIC_PROCESS | 86 | -0.340532 | -0.910425 | 0.615588 | 0.839802 |
| GOBP_CELLULAR_MODIFIED_AMINO_ACID_METABOLIC_PROCESS | 140 | -0.32341 | -0.910209 | 0.647971 | 0.839816 |
| GOBP_NEGATIVE_REGULATION_OF_PROTEIN_KINASE_B_SIGNALING | 40 | -0.383493 | -0.909997 | 0.596264 | 0.839815 |
| GOBP_POSITIVE_REGULATION_OF_DENDRITE_MORPHOGENESIS | 28 | -0.400215 | -0.909637 | 0.57622 | 0.840159 |
| GOBP_STORE_OPERATED_CALCIUM_ENTRY | 16 | -0.437958 | -0.909573 | 0.586922 | 0.839852 |
| GOBP_NECROPTOTIC_PROCESS | 33 | -0.400079 | -0.909313 | 0.583691 | 0.83997 |
| GOBP_POSITIVE_REGULATION_OF_PROTEOLYSIS_INVOLVED_IN_CELLULAR_PR | 113 | -0.329893 | -0.909085 | 0.606329 | 0.840024 |
| GOBP_NEUROTRANSMITTER_UPTAKE | 22 | -0.414932 | -0.908047 | 0.582258 | 0.841776 |
| GOBP_AORTA_MORPHOGENESIS | 20 | -0.424297 | -0.907885 | 0.593798 | 0.841651 |
| GOBP_ADIPOSE_TISSUE_DEVELOPMENT | 30 | -0.403168 | -0.907584 | 0.598811 | 0.841855 |
| GOBP_POSTSYNAPSE_ASSEMBLY | 16 | -0.432482 | -0.905523 | 0.579618 | 0.845849 |
| GOBP_HORMONE_BIOSYNTHETIC_PROCESS | 42 | -0.378013 | -0.904078 | 0.598303 | 0.848518 |
| GOBP_NEGATIVE_REGULATION_OF_DNA_REPLICATION | 29 | -0.401538 | -0.904003 | 0.619403 | 0.848249 |
| GOBP_PEPTIDYL_TYROSINE_AUTOPHOSPHORYLATION | 15 | -0.441449 | -0.903321 | 0.580431 | 0.849235 |
| GOBP_NEGATIVE_REGULATION_OF_CARBOHYDRATE_METABOLIC_PROCESS | 39 | -0.38279 | -0.903199 | 0.584416 | 0.849033 |

| | | | | | |
|---|---|---|---|---|---|
| GOBP_DEVELOPMENT_OF_PRIMARY_SEXUAL_CHARACTERISTICS | 132 | -0.320524 | -0.903154 | 0.637157 | 0.848695 |
| GOBP_VACUOLE_ORGANIZATION | 160 | -0.315257 | -0.902817 | 0.643192 | 0.848956 |
| GOBP_NEGATIVE_REGULATION_OF_SIGNAL_TRANSDUCTION_BY_P53_CLASS_M | 25 | -0.407121 | -0.902671 | 0.591049 | 0.848831 |
| GOBP_ENDOMEMBRANE_SYSTEM_ORGANIZATION | 411 | -0.298445 | -0.902226 | 0.716702 | 0.849336 |
| GOBP_REGULATION_OF_ACTIN_NUCLEATION | 24 | -0.407384 | -0.902206 | 0.590343 | 0.848929 |
| GOBP_MEMBRANE_LIPID_METABOLIC_PROCESS | 174 | -0.313148 | -0.901949 | 0.672454 | 0.849033 |
| GOBP_NEGATIVE_REGULATION_OF_DEFENSE_RESPONSE_TO_VIRUS | 21 | -0.413735 | -0.90191 | 0.601824 | 0.848669 |
| GOBP_INTERFERON_ALPHA_PRODUCTION | 26 | -0.39306 | -0.901459 | 0.598793 | 0.849146 |
| GOBP_REGULATION_OF_OXIDATIVE_STRESS_INDUCED_INTRINSIC_APOPTOTIC_ | 26 | -0.400035 | -0.901258 | 0.587057 | 0.849151 |
| GOBP_EXTRINSIC_APOPTOTIC_SIGNALING_PATHWAY_VIA_DEATH_DOMAIN_REC | 67 | -0.351089 | -0.900692 | 0.596317 | 0.849905 |
| GOBP_MAMMARY_GLAND_DEVELOPMENT | 93 | -0.336952 | -0.900625 | 0.626984 | 0.849609 |
| GOBP_POSITIVE_REGULATION_OF_P38MAPK_CASCADE | 19 | -0.417623 | -0.899898 | 0.594855 | 0.850729 |
| GOBP_ALCOHOL_BIOSYNTHETIC_PROCESS | 120 | -0.325877 | -0.89908 | 0.627139 | 0.851984 |
| GOBP_NEUROTROPHIN_TRK_RECEPTOR_SIGNALING_PATHWAY | 19 | -0.417852 | -0.897203 | 0.607874 | 0.855479 |
| GOBP_NEGATIVE_REGULATION_OF_WNT_SIGNALING_PATHWAY | 153 | -0.314992 | -0.897186 | 0.660839 | 0.855073 |
| GOBP_GLUTATHIONE_DERIVATIVE_METABOLIC_PROCESS | 16 | -0.439335 | -0.896611 | 0.588525 | 0.855823 |
| GOBP_INTRACILIARY_TRANSPORT | 52 | -0.360861 | -0.89594 | 0.619777 | 0.856791 |
| GOBP_VISUAL_BEHAVIOR | 32 | -0.388188 | -0.894674 | 0.637681 | 0.858949 |
| GOBP_INOSITOL_PHOSPHATE_METABOLIC_PROCESS | 56 | -0.35617 | -0.894377 | 0.618207 | 0.859136 |
| GOBP_NEURAL_PRECURSOR_CELL_PROLIFERATION | 86 | -0.340297 | -0.894286 | 0.620155 | 0.858878 |
| GOBP_RESPONSE_TO_LEAD_ION | 18 | -0.423848 | -0.893989 | 0.597738 | 0.85904 |
| GOBP_MAINTENANCE_OF_CELL_POLARITY | 17 | -0.426708 | -0.893674 | 0.599691 | 0.859271 |
| GOBP_POSITIVE_REGULATION_OF_MACROPHAGE_ACTIVATION | 18 | -0.423225 | -0.891529 | 0.605388 | 0.863362 |
| GOBP_MULTICELLULAR_ORGANISMAL_RESPONSE_TO_STRESS | 30 | -0.393063 | -0.891392 | 0.601199 | 0.863204 |
| GOBP_SECRETION_BY_TISSUE | 17 | -0.430413 | -0.891335 | 0.596405 | 0.862887 |
| GOBP_BONE_CELL_DEVELOPMENT | 28 | -0.384541 | -0.890591 | 0.597668 | 0.86397 |
| GOBP_NEGATIVE_REGULATION_OF_POSTTRANSCRIPTIONAL_GENE_SILENCING | 18 | -0.424382 | -0.890396 | 0.595674 | 0.863945 |
| GOBP_RETINA_HOMEOSTASIS | 50 | -0.355953 | -0.889891 | 0.637016 | 0.864575 |
| GOBP_REGULATION_OF_TUBE_SIZE | 66 | -0.348521 | -0.889372 | 0.639456 | 0.865233 |
| GOBP_CELL_SURFACE_RECEPTOR_SIGNALING_PATHWAY_INVOLVED_IN_CELL | 449 | -0.292937 | -0.888809 | 0.753977 | 0.865967 |
| GOBP_BLASTOCYST_FORMATION | 29 | -0.397297 | -0.887732 | 0.605926 | 0.867735 |
| GOBP_VESICLE_ORGANIZATION | 296 | -0.298099 | -0.88723 | 0.727273 | 0.868271 |
| GOBP_GENERATION_OF_PRECURSOR_METABOLITES_AND_ENERGY | 443 | -0.292461 | -0.887048 | 0.760167 | 0.868172 |
| GOBP_EXOCYTIC_PROCESS | 65 | -0.345898 | -0.886499 | 0.651535 | 0.868871 |
| GOBP_SKELETAL_MUSCLE_ORGAN_DEVELOPMENT | 101 | -0.331484 | -0.885705 | 0.686684 | 0.870072 |
| GOBP_POSTSYNAPTIC_SPECIALIZATION_ORGANIZATION | 19 | -0.412694 | -0.885406 | 0.61215 | 0.870249 |
| GOBP_VASCULAR_ASSOCIATED_SMOOTH_MUSCLE_CELL_MIGRATION | 16 | -0.422944 | -0.885043 | 0.61664 | 0.870536 |
| GOBP_POSITIVE_REGULATION_OF_MITOTIC_CELL_CYCLE | 94 | -0.326075 | -0.884967 | 0.650259 | 0.870248 |
| GOBP_RESPONSE_TO_PH | 24 | -0.396731 | -0.884567 | 0.625978 | 0.870595 |
| GOBP_MEMBRANE_LIPID_BIOSYNTHETIC_PROCESS | 120 | -0.319883 | -0.884518 | 0.653652 | 0.870249 |
| GOBP_RECEPTOR_LOCALIZATION_TO_SYNAPSE | 35 | -0.37598 | -0.884475 | 0.648567 | 0.869895 |
| GOBP_REGULATION_OF_MEMBRANE_REPOLARIZATION | 18 | -0.419635 | -0.883517 | 0.607201 | 0.871435 |
| GOBP_NEUROPEPTIDE_SIGNALING_PATHWAY | 33 | -0.381225 | -0.883196 | 0.608443 | 0.871671 |
| GOBP_POLYSACCHARIDE_METABOLIC_PROCESS | 83 | -0.331953 | -0.88284 | 0.631169 | 0.871933 |
| GOBP_MAINTENANCE_OF_PROTEIN_LOCATION | 78 | -0.340319 | -0.882236 | 0.654762 | 0.872743 |
| GOBP_DNA_DEPENDENT_DNA_REPLICATION | 151 | -0.310419 | -0.881979 | 0.682635 | 0.872815 |
| GOBP_POSITIVE_REGULATION_OF_UBIQUITIN_PROTEIN_TRANSFERASE_ACTIVIT | 30 | -0.376751 | -0.88181 | 0.609792 | 0.872715 |
| GOBP_NUCLEOTIDE_PHOSPHORYLATION | 108 | -0.324215 | -0.881515 | 0.647959 | 0.872881 |
| GOBP_POSITIVE_REGULATION_OF_INSULIN_SECRETION | 50 | -0.357533 | -0.881444 | 0.623395 | 0.872582 |
| GOBP_PROGRAMMED_NECROTIC_CELL_DEATH | 40 | -0.369583 | -0.880925 | 0.637168 | 0.873199 |
| GOBP_POSITIVE_REGULATION_OF_NEURON_PROJECTION_DEVELOPMENT | 116 | -0.318177 | -0.880789 | 0.660377 | 0.873015 |
| GOBP_NEGATIVE_REGULATION_OF_GENE_EXPRESSION_EPIGENETIC | 83 | -0.334421 | -0.880785 | 0.657143 | 0.872577 |
| GOBP_REGULATION_OF_LEUKOCYTE_ADHESION_TO_VASCULAR_ENDOTHELIAL | 22 | -0.406348 | -0.879643 | 0.611021 | 0.874539 |
| GOBP_REGULATION_OF_NERVOUS_SYSTEM_PROCESS | 76 | -0.337775 | -0.879307 | 0.645248 | 0.874779 |
| GOBP_NEGATIVE_REGULATION_OF_MUSCLE_CELL_DIFFERENTIATION | 39 | -0.368383 | -0.878971 | 0.624633 | 0.87502 |
| GOBP_CELLULAR_RESPONSE_TO_EXTERNAL_STIMULUS | 240 | -0.2998 | -0.878355 | 0.723714 | 0.875856 |
| GOBP_LYSOSOME_LOCALIZATION | 62 | -0.346576 | -0.87818 | 0.662125 | 0.875786 |
| GOBP_RESPONSE_TO_DOPAMINE | 42 | -0.359613 | -0.877896 | 0.631728 | 0.87593 |
| GOBP_POSITIVE_REGULATION_OF_ESTABLISHMENT_OF_PROTEIN_LOCALIZATIO | 56 | -0.355086 | -0.877179 | 0.667579 | 0.876949 |
| GOBP_REGULATION_OF_SMOOTH_MUSCLE_CELL_DIFFERENTIATION | 24 | -0.394928 | -0.876354 | 0.623053 | 0.878204 |
| GOBP_MUCOPOLYSACCHARIDE_METABOLIC_PROCESS | 80 | -0.332428 | -0.876034 | 0.659269 | 0.878407 |
| GOBP_SYNAPTIC_TRANSMISSION_CHOLINERGIC | 20 | -0.406258 | -0.875704 | 0.635401 | 0.878666 |
| GOBP_NEGATIVE_REGULATION_OF_LIPID_BIOSYNTHETIC_PROCESS | 35 | -0.379594 | -0.875356 | 0.636765 | 0.878912 |
| GOBP_ATP_METABOLIC_PROCESS | 269 | -0.296763 | -0.875246 | 0.737259 | 0.878677 |
| GOBP_CARBOHYDRATE_TRANSMEMBRANE_TRANSPORT | 78 | -0.331362 | -0.874868 | 0.670103 | 0.878976 |
| GOBP_RESPONSE_TO_MONOSACCHARIDE | 143 | -0.31125 | -0.874457 | 0.695035 | 0.879382 |
| GOBP_VACUOLAR_TRANSPORT | 140 | -0.3142 | -0.874409 | 0.69335 | 0.879027 |
| GOBP_GLUCOSE_METABOLIC_PROCESS | 168 | -0.308076 | -0.874239 | 0.712919 | 0.878949 |
| GOBP_POSITIVE_REGULATION_OF_PROTEIN_IMPORT | 31 | -0.378627 | -0.8741 | 0.648688 | 0.878791 |
| GOBP_REGULATION_OF_LEUKOCYTE_DEGRANULATION | 36 | -0.369073 | -0.873969 | 0.634393 | 0.87861 |
| GOBP_REGULATION_OF_NUCLEAR_TRANSCRIBED_MRNA_POLY_A_TAIL_SHORT | 15 | -0.426584 | -0.873507 | 0.630573 | 0.8791 |
| GOBP_JNK_CASCADE | 158 | -0.308439 | -0.873288 | 0.709563 | 0.879134 |
| GOBP_CARDIAC_EPITHELIAL_TO_MESENCHYMAL_TRANSITION | 16 | -0.421482 | -0.872745 | 0.623188 | 0.879827 |
| GOBP_CELL_CELL_SIGNALING_BY_WNT | 391 | -0.290723 | -0.872152 | 0.774262 | 0.880608 |
| GOBP_REGULATION_OF_INTRINSIC_APOPTOTIC_SIGNALING_PATHWAY_IN_RESI | 32 | -0.37422 | -0.871973 | 0.614934 | 0.880534 |
| GOBP_REPLICATION_FORK_PROCESSING | 37 | -0.370718 | -0.87161 | 0.642346 | 0.880825 |
| GOBP_REGULATION_OF_PROTEIN_EXPORT_FROM_NUCLEUS | 32 | -0.3701 | -0.870994 | 0.633333 | 0.881596 |
| GOBP_REGULATION_OF_TRANSLATIONAL_INITIATION | 74 | -0.333515 | -0.870763 | 0.675462 | 0.881618 |
| GOBP_REGULATION_OF_LONG_TERM_NEURONAL_SYNAPTIC_PLASTICITY | 17 | -0.420095 | -0.870754 | 0.619512 | 0.881189 |
| GOBP_RESPONSE_TO_THYROID_HORMONE | 17 | -0.414528 | -0.870672 | 0.624 | 0.880919 |
| GOBP_REGULATION_OF_PROTEASOMAL_UBIQUITIN_DEPENDENT_PROTEIN_CA1 | 119 | -0.316631 | -0.870352 | 0.673292 | 0.88112 |
| GOBP_REGULATION_OF_UBIQUITIN_DEPENDENT_PROTEIN_CATABOLIC_PROCE | 146 | -0.308541 | -0.87029 | 0.716687 | 0.880798 |
| GOBP_RESPONSE_TO_UV | 130 | -0.313457 | -0.870067 | 0.702539 | 0.880803 |
| GOBP_RESPONSE_TO_TOPOLOGICALLY_INCORRECT_PROTEIN | 184 | -0.30398 | -0.86902 | 0.714286 | 0.882476 |
| GOBP_VESICLE_CYTOSKELETAL_TRAFFICKING | 61 | -0.342056 | -0.868416 | 0.65847 | 0.883257 |
| GOBP_SYNAPTIC_VESICLE_TRANSPORT | 38 | -0.367976 | -0.867956 | 0.645604 | 0.883717 |
| GOBP_GUANOSINE_CONTAINING_COMPOUND_METABOLIC_PROCESS | 37 | -0.37571 | -0.86786 | 0.659664 | 0.883457 |
| GOBP_VASCULAR_ASSOCIATED_SMOOTH_MUSCLE_CONTRACTION | 15 | -0.427532 | -0.867569 | 0.64404 | 0.883585 |
| GOBP_SYNAPSE_ASSEMBLY | 80 | -0.330876 | -0.867289 | 0.673712 | 0.883673 |
| GOBP_RDNA_HETEROCHROMATIN_ASSEMBLY | 23 | -0.396231 | -0.867105 | 0.650078 | 0.883607 |
| GOBP_POSITIVE_REGULATION_OF_BLOOD_VESSEL_ENDOTHELIAL_CELL_MIGRA | 44 | -0.358939 | -0.866089 | 0.656566 | 0.885163 |
| GOBP_POSITIVE_REGULATION_OF_CARBOHYDRATE_METABOLIC_PROCESS | 53 | -0.352018 | -0.864874 | 0.663934 | 0.887167 |
| GOBP_CELLULAR_RESPONSE_TO_RADIATION | 149 | -0.310096 | -0.864683 | 0.721015 | 0.88711 |
| GOBP_REGULATION_OF_POSTSYNAPSE_ORGANIZATION | 65 | -0.341467 | -0.864421 | 0.671001 | 0.887158 |
| GOBP_ENDOTHELIAL_CELL_APOPTOTIC_PROCESS | 37 | -0.368563 | -0.864052 | 0.647145 | 0.887462 |
| GOBP_REGULATION_OF_POTASSIUM_ION_TRANSPORT | 53 | -0.347111 | -0.8639 | 0.675487 | 0.887338 |
| GOBP_NEUROBLAST_PROLIFERATION | 32 | -0.378273 | -0.863874 | 0.648773 | 0.886946 |
| GOBP_CHAPERONE_MEDIATED_PROTEIN_FOLDING | 59 | -0.33808 | -0.863294 | 0.660714 | 0.887664 |
| GOBP_CARBOHYDRATE_PHOSPHORYLATION | 23 | -0.39237 | -0.862934 | 0.63354 | 0.887931 |
| GOBP_NEGATIVE_REGULATION_OF_BMP_SIGNALING_PATHWAY | 28 | -0.384732 | -0.862737 | 0.653251 | 0.887886 |
| GOBP_CYCLIC_NUCLEOTIDE_METABOLIC_PROCESS | 23 | -0.383129 | -0.862166 | 0.635938 | 0.888566 |
| GOBP_NEGATIVE_REGULATION_OF_OSTEOCLAST_DIFFERENTIATION | 20 | -0.402214 | -0.861958 | 0.641745 | 0.888535 |
| GOBP_C4_DICARBOXYLATE_TRANSPORT | 18 | -0.411857 | -0.861361 | 0.62212 | 0.889252 |
| GOBP_FOREBRAIN_CELL_MIGRATION | 32 | -0.371836 | -0.860232 | 0.659969 | 0.891076 |
| GOBP_RESPONSE_TO_STARVATION | 161 | -0.301375 | -0.860083 | 0.738407 | 0.89093 |
| GOBP_CELLULAR_RESPONSE_TO_ABIOTIC_STIMULUS | 255 | -0.294177 | -0.860013 | 0.784825 | 0.890622 |
| GOBP_NECROTIC_CELL_DEATH | 51 | -0.341056 | -0.859759 | 0.657382 | 0.890673 |
| GOBP_POSITIVE_REGULATION_OF_RECEPTOR_INTERNALIZATION | 16 | -0.420471 | -0.85901 | 0.650519 | 0.891755 |
| GOBP_PROTEIN_LOCALIZATION_TO_POSTSYNAPSE | 28 | -0.381506 | -0.858791 | 0.654762 | 0.891738 |
| GOBP_REGULATION_OF_NEUROTRANSMITTER_TRANSPORT | 64 | -0.335673 | -0.858668 | 0.671662 | 0.891526 |
| GOBP_CELLULAR_RESPONSE_TO_EXTRACELLULAR_STIMULUS | 184 | -0.301056 | -0.858519 | 0.729093 | 0.891371 |
| GOBP_GLUCOSE_6_PHOSPHATE_METABOLIC_PROCESS | 21 | -0.399036 | -0.857941 | 0.639871 | 0.892089 |
| GOBP_NUCLEOSIDE_METABOLIC_PROCESS | 94 | -0.321377 | -0.857552 | 0.694301 | 0.892388 |
| GOBP_RESPONSE_TO_ACIDIC_PH | 15 | -0.424761 | -0.857424 | 0.664372 | 0.892192 |
| GOBP_MUSCLE_FILAMENT_SLIDING | 22 | -0.391775 | -0.857372 | 0.658574 | 0.891852 |
| GOBP_REGULATION_OF_GENERATION_OF_PRECURSOR_METABOLITES_AND_E | 133 | -0.313614 | -0.857159 | 0.706983 | 0.891835 |
| GOBP_MITOCHONDRIAL_GENOME_MAINTENANCE | 20 | -0.399309 | -0.857064 | 0.683007 | 0.891575 |
| GOBP_CEREBRAL_CORTEX_CELL_MIGRATION | 25 | -0.38051 | -0.856778 | 0.635802 | 0.891673 |
| GOBP_INTERFERON_GAMMA_MEDIATED_SIGNALING_PATHWAY | 81 | -0.322135 | -0.855589 | 0.700922 | 0.893534 |
| GOBP_RIBONUCLEOSIDE_CATABOLIC_PROCESS | 19 | -0.395578 | -0.855062 | 0.6544 | 0.89412 |
| GOBP_CELLULAR_GLUCAN_METABOLIC_PROCESS | 59 | -0.338529 | -0.85433 | 0.669916 | 0.895118 |
| GOBP_POSITIVE_REGULATION_OF_RESPONSE_TO_CYTOKINE_STIMULUS | 53 | -0.335537 | -0.854291 | 0.673851 | 0.894764 |
| GOBP_COLLATERAL_SPROUTING | 15 | -0.420009 | -0.854191 | 0.673333 | 0.894514 |
| GOBP_REGULATION_OF_HORMONE_SECRETION | 168 | -0.299061 | -0.853694 | 0.768065 | 0.895002 |
| GOBP_POSITIVE_REGULATION_OF_STRESS_ACTIVATED_PROTEIN_KINASE_SIGN | 94 | -0.317896 | -0.853232 | 0.700394 | 0.895439 |
| GOBP_NEGATIVE_REGULATION_OF_GLIOGENESIS | 18 | -0.404133 | -0.853045 | 0.649416 | 0.895393 |
| GOBP_POSITIVE_REGULATION_OF_LEUKOCYTE_APOPTOTIC_PROCESS | 21 | -0.390907 | -0.852762 | 0.653673 | 0.895498 |
| GOBP_NEGATIVE_REGULATION_OF_PROTEASOMAL_PROTEIN_CATABOLIC_PRO | 42 | -0.352333 | -0.852506 | 0.676946 | 0.895555 |
| GOBP_REGULATION_OF_CILIUM_ASSEMBLY | 50 | -0.349041 | -0.851502 | 0.687065 | 0.897055 |
| GOBP_AUDITORY_RECEPTOR_CELL_DEVELOPMENT | 15 | -0.413111 | -0.850666 | 0.636066 | 0.898207 |
| GOBP_HYALURONAN_METABOLIC_PROCESS | 29 | -0.364037 | -0.850403 | 0.654079 | 0.898276 |
| GOBP_EXIT_FROM_MITOSIS | 25 | -0.382323 | -0.850361 | 0.690994 | 0.89793 |
| GOBP_NEGATIVE_REGULATION_OF_JNK_CASCADE | 30 | -0.367302 | -0.849965 | 0.713405 | 0.898214 |
| GOBP_GERM_CELL_DEVELOPMENT | 163 | -0.300863 | -0.849913 | 0.749702 | 0.897873 |
| GOBP_PYRIMIDINE_NUCLEOSIDE_METABOLIC_PROCESS | 30 | -0.367528 | -0.84982 | 0.658915 | 0.897612 |
| GOBP_NEGATIVE_REGULATION_OF_COLD_INDUCED_THERMOGENESIS | 37 | -0.360686 | -0.849447 | 0.672414 | 0.897888 |
| GOBP_POSITIVE_REGULATION_OF_ANION_TRANSMEMBRANE_TRANSPORT | 46 | -0.349735 | -0.849049 | 0.66993 | 0.898249 |
| GOBP_REGULATION_OF_STRIATED_MUSCLE_CONTRACTION | 52 | -0.338274 | -0.848691 | 0.664316 | 0.898484 |
| GOBP_LONG_TERM_SYNAPTIC_DEPRESSION | 18 | -0.398989 | -0.848658 | 0.675283 | 0.8981 |
| GOBP_REGULATION_OF_SYSTEMIC_ARTERIAL_BLOOD_PRESSURE_MEDIATED_ | 25 | -0.378721 | -0.848591 | 0.659159 | 0.897774 |
| GOBP_BONE_DEVELOPMENT | 127 | -0.306977 | -0.847904 | 0.736585 | 0.898648 |
| GOBP_PYRIMIDINE_CONTAINING_COMPOUND_CATABOLIC_PROCESS | 33 | -0.364934 | -0.84783 | 0.667158 | 0.898338 |
| GOBP_NEGATIVE_REGULATION_OF_SMALL_GTPASE_MEDIATED_SIGNAL_TRANS | 49 | -0.344662 | -0.847209 | 0.69297 | 0.899104 |
| GOBP_CILIUM_ORGANIZATION | 341 | -0.284518 | -0.847024 | 0.800871 | 0.898995 |
| GOBP_VESICLE_BUDDING_FROM_MEMBRANE | 101 | -0.312363 | -0.846996 | 0.724005 | 0.898604 |
| GOBP_CHONDROITIN_SULFATE_PROTEOGLYCAN_BIOSYNTHETIC_PROCESS | 21 | -0.397792 | -0.846972 | 0.672414 | 0.898216 |
| GOBP_NEGATIVE_REGULATION_OF_CELLULAR_RESPONSE_TO_INSULIN_STIMU | 31 | -0.36672 | -0.846608 | 0.681048 | 0.898463 |

| | | | | | |
|---|---|---|---|---|---|
| GOBP_POSITIVE_REGULATION_OF_NEUROTRANSMITTER_TRANSPORT | 18 | -0.407706 | -0.845409 | 0.656151 | 0.900319 |
| GOBP_HISTONE_DEACETYLATION | 70 | -0.328026 | -0.845219 | 0.705718 | 0.900251 |
| GOBP_MONOSACCHARIDE_METABOLIC_PROCESS | 220 | -0.290946 | -0.844365 | 0.777652 | 0.901429 |
| GOBP_MUSCLE_ADAPTATION | 75 | -0.325475 | -0.844209 | 0.718912 | 0.901269 |
| GOBP_REGULATION_OF_SMOOTH_MUSCLE_CONTRACTION | 33 | -0.365689 | -0.84331 | 0.678779 | 0.902534 |
| GOBP_MICROTUBULE_NUCLEATION | 30 | -0.361687 | -0.843042 | 0.689605 | 0.902602 |
| GOBP_CELLULAR_PROCESS_INVOLVED_IN_REPRODUCTION_IN_MULTICELLUL | 211 | -0.295234 | -0.842597 | 0.779206 | 0.902976 |
| GOBP_SENSORY_PERCEPTION | 306 | -0.281888 | -0.840627 | 0.816017 | 0.906236 |
| GOBP_SMOOTH_MUSCLE_CONTRACTION | 53 | -0.338304 | -0.840457 | 0.711724 | 0.906121 |
| GOBP_POSITIVE_REGULATION_OF_CELL_CYCLE | 309 | -0.277408 | -0.838832 | 0.832794 | 0.908717 |
| GOBP_REGULATION_OF_AUTOPHAGOSOME_ASSEMBLY | 38 | -0.354614 | -0.838736 | 0.708861 | 0.908458 |
| GOBP_PHOTOTRANSDUCTION | 29 | -0.366543 | -0.838342 | 0.681402 | 0.908751 |
| GOBP_NEGATIVE_REGULATION_OF_CANONICAL_WNT_SIGNALING_PATHWAY | 131 | -0.299178 | -0.83832 | 0.737783 | 0.908357 |
| GOBP_PLATELET_MORPHOGENESIS | 20 | -0.394494 | -0.838101 | 0.691919 | 0.908328 |
| GOBP_NEGATIVE_REGULATION_OF_CELLULAR_PROTEIN_CATABOLIC_PROCES | 65 | -0.332918 | -0.837647 | 0.727763 | 0.908723 |
| GOBP_NEGATIVE_REGULATION_OF_GENE_SILENCING | 26 | -0.374603 | -0.837277 | 0.679274 | 0.90897 |
| GOBP_REGULATION_OF_GLUCAN_BIOSYNTHETIC_PROCESS | 21 | -0.38063 | -0.836922 | 0.6848 | 0.909195 |
| GOBP_CHONDROITIN_SULFATE_PROTEOGLYCAN_METABOLIC_PROCESS | 29 | -0.364958 | -0.836755 | 0.697289 | 0.909057 |
| GOBP_NEGATIVE_REGULATION_OF_MACROAUTOPHAGY | 33 | -0.359597 | -0.836546 | 0.696241 | 0.90899 |
| GOBP_REGULATION_OF_CARBOHYDRATE_BIOSYNTHETIC_PROCESS | 70 | -0.319689 | -0.836484 | 0.731973 | 0.90866 |
| GOBP_NUCLEOSIDE_TRIPHOSPHATE_BIOSYNTHETIC_PROCESS | 74 | -0.316566 | -0.836408 | 0.718121 | 0.908365 |
| GOBP_CARBOHYDRATE_HOMEOSTASIS | 158 | -0.292734 | -0.835797 | 0.780142 | 0.909056 |
| GOBP_REGULATION_OF_PROTEIN_CONTAINING_COMPLEX_DISASSEMBLY | 99 | -0.307802 | -0.835411 | 0.732051 | 0.909358 |
| GOBP_CYTOSKELETON_DEPENDENT_INTRACELLULAR_TRANSPORT | 178 | -0.293631 | -0.835364 | 0.778947 | 0.909005 |
| GOBP_ALDITOL_METABOLIC_PROCESS | 16 | -0.403381 | -0.834728 | 0.7031 | 0.909755 |
| GOBP_CELL_CYCLE_ARREST | 194 | -0.285407 | -0.834105 | 0.801615 | 0.91048 |
| GOBP_RUFFLE_ORGANIZATION | 46 | -0.341833 | -0.832179 | 0.715481 | 0.913558 |
| GOBP_MITOTIC_DNA_REPLICATION | 16 | -0.404476 | -0.831718 | 0.675241 | 0.913955 |
| GOBP_NEGATIVE_REGULATION_OF_PROTEIN_MODIFICATION_BY_SMALL_PROTE | 84 | -0.312475 | -0.831451 | 0.723545 | 0.913989 |
| GOBP_INSULIN_LIKE_GROWTH_FACTOR_RECEPTOR_SIGNALING_PATHWAY | 19 | -0.399189 | -0.831285 | 0.667208 | 0.913839 |
| GOBP_POSITIVE_REGULATION_OF_PROTEIN_MATURATION | 20 | -0.389655 | -0.831115 | 0.67737 | 0.913726 |
| GOBP_ACTIN_NUCLEATION | 44 | -0.337555 | -0.830995 | 0.720863 | 0.913509 |
| GOBP_EPITHELIAL_CELL_APOPTOTIC_PROCESS | 72 | -0.316079 | -0.830958 | 0.726081 | 0.913145 |
| GOBP_RIBONUCLEOSIDE_METABOLIC_PROCESS | 64 | -0.329696 | -0.830105 | 0.758808 | 0.91435 |
| GOBP_POSITIVE_REGULATION_OF_CELL_CYCLE_PROCESS | 239 | -0.281812 | -0.829912 | 0.817568 | 0.914271 |
| GOBP_LAMELLIPODIUM_MORPHOGENESIS | 15 | -0.415809 | -0.829548 | 0.697674 | 0.914498 |
| GOBP_CELLULAR_RESPONSE_TO_ESTRADIOL_STIMULUS | 25 | -0.377452 | -0.828128 | 0.687596 | 0.916653 |
| GOBP_SEX_DIFFERENTIATION | 153 | -0.293881 | -0.828047 | 0.77497 | 0.916349 |
| GOBP_DICARBOXYLIC_ACID_METABOLIC_PROCESS | 78 | -0.313397 | -0.828005 | 0.73613 | 0.915979 |
| GOBP_NEUROTRANSMITTER_RECEPTOR_TRANSPORT | 32 | -0.348834 | -0.827978 | 0.701183 | 0.915593 |
| GOBP_POSITIVE_REGULATION_OF_DNA_TEMPLATED_TRANSCRIPTION_ELONGA | 26 | -0.370559 | -0.826816 | 0.705401 | 0.917277 |
| GOBP_ASSOCIATIVE_LEARNING | 43 | -0.343142 | -0.826436 | 0.713267 | 0.91752 |
| GOBP_REGULATION_OF_SYNAPTIC_TRANSMISSION_GLUTAMATERGIC | 30 | -0.363105 | -0.826055 | 0.691843 | 0.91775 |
| GOBP_PROCESS_UTILIZING_AUTOPHAGIC_MECHANISM | 485 | -0.272714 | -0.824468 | 0.879328 | 0.920162 |
| GOBP_CELLULAR_CARBOHYDRATE_BIOSYNTHETIC_PROCESS | 60 | -0.331115 | -0.824257 | 0.727532 | 0.920104 |
| GOBP_ESTABLISHMENT_OF_ORGANELLE_LOCALIZATION | 373 | -0.274968 | -0.824253 | 0.861736 | 0.919674 |
| GOBP_CELLULAR_RESPONSE_TO_MECHANICAL_STIMULUS | 61 | -0.322312 | -0.824031 | 0.728395 | 0.919658 |
| GOBP_REGULATION_OF_SKELETAL_MUSCLE_TISSUE_DEVELOPMENT | 15 | -0.410628 | -0.823985 | 0.693038 | 0.9193 |
| GOBP_INTERLEUKIN_6_MEDIATED_SIGNALING_PATHWAY | 18 | -0.394536 | -0.822811 | 0.716828 | 0.92097 |
| GOBP_RESPONSE_TO_DSRNA | 36 | -0.348606 | -0.822785 | 0.733133 | 0.920577 |
| GOBP_VASCULAR_ASSOCIATED_SMOOTH_MUSCLE_CELL_DIFFERENTIATION | 21 | -0.384041 | -0.822309 | 0.704867 | 0.921037 |
| GOBP_HETEROCHROMATIN_ORGANIZATION | 59 | -0.322407 | -0.822028 | 0.711749 | 0.921118 |
| GOBP_CEREBRAL_CORTEX_RADIALLY_ORIENTED_CELL_MIGRATION | 21 | -0.378839 | -0.82192 | 0.698541 | 0.920882 |
| GOBP_MICROTUBULE_BASED_TRANSPORT | 168 | -0.289765 | -0.82109 | 0.777648 | 0.921926 |
| GOBP_ARGININE_METABOLIC_PROCESS | 16 | -0.39299 | -0.82104 | 0.677831 | 0.921576 |
| GOBP_DENDRITE_EXTENSION | 25 | -0.370654 | -0.820393 | 0.714286 | 0.922302 |
| GOBP_TELENCEPHALON_GLIAL_CELL_MIGRATION | 16 | -0.404415 | -0.820271 | 0.689655 | 0.92209 |
| GOBP_DEOXYRIBOSE_PHOSPHATE_CATABOLIC_PROCESS | 24 | -0.36824 | -0.819842 | 0.69102 | 0.922413 |
| GOBP_CELLULAR_RESPONSE_TO_RETINOIC_ACID | 37 | -0.346229 | -0.819223 | 0.728614 | 0.923086 |
| GOBP_REGULATION_OF_MICROTUBULE_POLYMERIZATION | 43 | -0.335513 | -0.81871 | 0.737288 | 0.923555 |
| GOBP_POSITIVE_REGULATION_OF_LEUKOCYTE_DEGRANULATION | 18 | -0.389351 | -0.818226 | 0.701954 | 0.923981 |
| GOBP_AUTOPHAGOSOME_ORGANIZATION | 93 | -0.306153 | -0.817183 | 0.76625 | 0.925405 |
| GOBP_CELLULAR_RESPONSE_TO_TOPOLOGICALLY_INCORRECT_PROTEIN | 151 | -0.292647 | -0.81611 | 0.793689 | 0.926898 |
| GOBP_SPHINGOID_METABOLIC_PROCESS | 19 | -0.381493 | -0.815297 | 0.710311 | 0.927865 |
| GOBP_NUCLEOSIDE_CATABOLIC_PROCESS | 27 | -0.359173 | -0.814633 | 0.72291 | 0.928581 |
| GOBP_MITOCHONDRIAL_TRANSCRIPTION | 17 | -0.384977 | -0.814586 | 0.711185 | 0.928234 |
| GOBP_MYELOID_CELL_DEVELOPMENT | 56 | -0.327232 | -0.814445 | 0.727528 | 0.928036 |
| GOBP_REGULATION_OF_INTRACELLULAR_STEROID_HORMONE_RECEPTOR_SI | 58 | -0.319159 | -0.813905 | 0.749296 | 0.92859 |
| GOBP_POSITIVE_REGULATION_OF_GLUCOSE_TRANSMEMBRANE_TRANSPORT | 32 | -0.356256 | -0.813886 | 0.731895 | 0.928188 |
| GOBP_DEMETHYLATION | 64 | -0.322086 | -0.813542 | 0.721024 | 0.928378 |
| GOBP_VESICLE_TARGETING | 86 | -0.308728 | -0.812678 | 0.792526 | 0.929456 |
| GOBP_ORGANIC_ACID_BIOSYNTHETIC_PROCESS | 231 | -0.277433 | -0.812084 | 0.84289 | 0.930068 |
| GOBP_MANNOSYLATION | 35 | -0.343842 | -0.811247 | 0.722628 | 0.931133 |
| GOBP_REGULATION_OF_SYNAPTIC_VESICLE_EXOCYTOSIS | 38 | -0.338351 | -0.810445 | 0.733432 | 0.932084 |
| GOBP_3_UTR_MEDIATED_MRNA_STABILIZATION | 17 | -0.392207 | -0.810077 | 0.703349 | 0.932282 |
| GOBP_RETROGRADE_TRANSPORT_ENDOSOME_TO_GOLGI | 86 | -0.303134 | -0.809446 | 0.788462 | 0.932928 |
| GOBP_ACROSOME_REACTION | 21 | -0.378618 | -0.808543 | 0.698587 | 0.934025 |
| GOBP_POSITIVE_REGULATION_OF_TYPE_I_INTERFERON_PRODUCTION | 74 | -0.310959 | -0.808534 | 0.759894 | 0.933602 |
| GOBP_REGULATION_OF_VACUOLE_ORGANIZATION | 45 | -0.330417 | -0.808335 | 0.757267 | 0.933485 |
| GOBP_SENSORY_PERCEPTION_OF_BITTER_TASTE | 18 | -0.386288 | -0.808141 | 0.717355 | 0.933383 |
| GOBP_PYRIMIDINE_NUCLEOTIDE_CATABOLIC_PROCESS | 18 | -0.381091 | -0.807646 | 0.705128 | 0.933777 |
| GOBP_REGULATION_OF_MICROTUBULE_BASED_PROCESS | 183 | -0.281615 | -0.80756 | 0.84881 | 0.933497 |
| GOBP_NEGATIVE_REGULATION_OF_RHO_PROTEIN_SIGNAL_TRANSDUCTION | 17 | -0.389281 | -0.806941 | 0.70915 | 0.934154 |
| GOBP_MICROTUBULE_ORGANIZING_CENTER_LOCALIZATION | 24 | -0.36625 | -0.806928 | 0.703928 | 0.933742 |
| GOBP_REGULATION_OF_CELLULAR_RESPONSE_TO_INSULIN_STIMULUS | 51 | -0.324816 | -0.806828 | 0.754458 | 0.933494 |
| GOBP_QUINONE_METABOLIC_PROCESS | 25 | -0.360823 | -0.80619 | 0.740122 | 0.934148 |
| GOBP_REGULATION_OF_CARDIAC_MUSCLE_CONTRACTION | 42 | -0.334141 | -0.802789 | 0.750355 | 0.939587 |
| GOBP_ARF_PROTEIN_SIGNAL_TRANSDUCTION | 17 | -0.385694 | -0.80272 | 0.719934 | 0.939259 |
| GOBP_POSITIVE_REGULATION_OF_VIRAL_TRANSCRIPTION | 26 | -0.350692 | -0.80111 | 0.726718 | 0.941601 |
| GOBP_PROTEIN_K11_LINKED_UBIQUITINATION | 29 | -0.354635 | -0.79993 | 0.722874 | 0.943189 |
| GOBP_COPPER_ION_HOMEOSTASIS | 15 | -0.385892 | -0.798359 | 0.72935 | 0.945426 |
| GOBP_NEGATIVE_REGULATION_OF_TELOMERE_MAINTENANCE_VIA_TELOMERE | 24 | -0.35698 | -0.798351 | 0.722552 | 0.944999 |
| GOBP_RAC_PROTEIN_SIGNAL_TRANSDUCTION | 38 | -0.334016 | -0.797751 | 0.723404 | 0.94559 |
| GOBP_REGULATION_OF_DNA_METHYLATION | 18 | -0.376871 | -0.797454 | 0.720379 | 0.945659 |
| GOBP_REGULATION_OF_RECEPTOR_INTERNALIZATION | 37 | -0.331591 | -0.796583 | 0.725664 | 0.946642 |
| GOBP_P_BODY_ASSEMBLY | 21 | -0.365927 | -0.795782 | 0.714946 | 0.947543 |
| GOBP_POLYAMINE_METABOLIC_PROCESS | 17 | -0.378781 | -0.795431 | 0.729299 | 0.947697 |
| GOBP_KETONE_BIOSYNTHETIC_PROCESS | 32 | -0.343697 | -0.795197 | 0.761103 | 0.947635 |
| GOBP_REGULATION_OF_TOLL_LIKE_RECEPTOR_4_SIGNALING_PATHWAY | 17 | -0.372975 | -0.794208 | 0.712195 | 0.948837 |
| GOBP_PHOTORECEPTOR_CELL_MAINTENANCE | 30 | -0.343227 | -0.792918 | 0.732116 | 0.950511 |
| GOBP_NEGATIVE_REGULATION_OF_MRNA_CATABOLIC_PROCESS | 44 | -0.323571 | -0.792775 | 0.767341 | 0.950297 |
| GOBP_NUCLEAR_MEMBRANE_ORGANIZATION | 16 | -0.382813 | -0.791014 | 0.736409 | 0.952778 |
| GOBP_PURINE_NUCLEOSIDE_METABOLIC_PROCESS | 57 | -0.316772 | -0.790901 | 0.793201 | 0.952508 |
| GOBP_NEUROTROPHIN_SIGNALING_PATHWAY | 25 | -0.354249 | -0.790629 | 0.758359 | 0.952508 |
| GOBP_REGULATION_OF_MUSCLE_ADAPTATION | 61 | -0.306738 | -0.790214 | 0.785814 | 0.952735 |
| GOBP_PROTEIN_PEPTIDYL_PROLYL_ISOMERIZATION | 33 | -0.337144 | -0.787949 | 0.748858 | 0.956062 |
| GOBP_ENDOCRINE_HORMONE_SECRETION | 24 | -0.356643 | -0.787593 | 0.755302 | 0.956215 |
| GOBP_CELLULAR_METABOLIC_COMPOUND_SALVAGE | 29 | -0.342913 | -0.787413 | 0.736446 | 0.95608 |
| GOBP_REGULATION_OF_TRANSCRIPTION_BY_RNA_POLYMERASE_I | 31 | -0.339049 | -0.787271 | 0.754947 | 0.95588 |
| GOBP_REGULATION_OF_HISTONE_H3_K4_METHYLATION | 25 | -0.361732 | -0.787096 | 0.742138 | 0.955734 |
| GOBP_NEUROTRANSMITTER_SECRETION | 99 | -0.291064 | -0.786619 | 0.818065 | 0.956063 |
| GOBP_ENDOPLASMIC_RETICULUM_UNFOLDED_PROTEIN_RESPONSE | 113 | -0.287545 | -0.786515 | 0.838072 | 0.955797 |
| GOBP_REGULATION_OF_CARDIAC_MUSCLE_CONTRACTION_BY_CALCIUM_ION_ | 18 | -0.369218 | -0.786348 | 0.746434 | 0.955614 |
| GOBP_LEUKOCYTE_TETHERING_OR_ROLLING | 22 | -0.360287 | -0.786244 | 0.773643 | 0.955358 |
| GOBP_INORGANIC_ION_IMPORT_ACROSS_PLASMA_MEMBRANE | 56 | -0.310275 | -0.78529 | 0.79617 | 0.95647 |
| GOBP_REGULATION_OF_LAMELLIPODIUM_ASSEMBLY | 33 | -0.341601 | -0.785185 | 0.735426 | 0.956198 |
| GOBP_POSITIVE_REGULATION_OF_MONOOXYGENASE_ACTIVITY | 26 | -0.346482 | -0.785145 | 0.748792 | 0.955818 |
| GOBP_CERAMIDE_BIOSYNTHETIC_PROCESS | 54 | -0.316408 | -0.784889 | 0.78022 | 0.955767 |
| GOBP_POSITIVE_REGULATION_OF_ACTIVATED_T_CELL_PROLIFERATION | 17 | -0.370119 | -0.784495 | 0.714516 | 0.955958 |
| GOBP_NEGATIVE_REGULATION_OF_ORGANELLE_ORGANIZATION | 286 | -0.264156 | -0.784326 | 0.919069 | 0.955785 |
| GOBP_ENERGY_RESERVE_METABOLIC_PROCESS | 61 | -0.307183 | -0.782797 | 0.799447 | 0.957761 |
| GOBP_POSITIVE_REGULATION_OF_G1_S_TRANSITION_OF_MITOTIC_CELL_CYCL | 29 | -0.335619 | -0.782713 | 0.752336 | 0.957457 |
| GOBP_REGULATION_OF_MRNA_CATABOLIC_PROCESS | 186 | -0.270012 | -0.781978 | 0.878613 | 0.958199 |
| GOBP_PROTEIN_AUTOUBIQUITINATION | 65 | -0.303066 | -0.781319 | 0.809652 | 0.958794 |
| GOBP_POSITIVE_REGULATION_OF_SMOOTHENED_SIGNALING_PATHWAY | 25 | -0.349154 | -0.781027 | 0.746836 | 0.958824 |
| GOBP_RIBONUCLEOSIDE_TRIPHOSPHATE_BIOSYNTHETIC_PROCESS | 64 | -0.303984 | -0.77983 | 0.832168 | 0.960251 |
| GOBP_NUCLEOSIDE_PHOSPHATE_CATABOLIC_PROCESS | 67 | -0.300073 | -0.777373 | 0.816167 | 0.963713 |
| GOBP_DETECTION_OF_CHEMICAL_STIMULUS | 52 | -0.312256 | -0.776597 | 0.779804 | 0.964461 |
| GOBP_CELLULAR_RESPONSE_TO_INSULIN_STIMULUS | 180 | -0.267012 | -0.776464 | 0.880562 | 0.964207 |
| GOBP_VASCULOGENESIS | 45 | -0.318848 | -0.7763 | 0.782793 | 0.964034 |
| GOBP_INTRA_S_DNA_DAMAGE_CHECKPOINT | 17 | -0.371554 | -0.776125 | 0.75 | 0.963853 |
| GOBP_NEGATIVE_REGULATION_OF_REACTIVE_OXYGEN_SPECIES_BIOSYNTHET | 24 | -0.349211 | -0.775843 | 0.763975 | 0.963849 |
| GOBP_REGULATION_OF_TORC1_SIGNALING | 37 | -0.331041 | -0.775548 | 0.775964 | 0.963875 |
| GOBP_REGULATION_OF_CELL_DIVISION | 115 | -0.277693 | -0.775219 | 0.843788 | 0.963941 |
| GOBP_POSITIVE_REGULATION_OF_MUSCLE_CONTRACTION | 25 | -0.350989 | -0.774999 | 0.75975 | 0.963832 |
| GOBP_NON_MOTILE_CILIUM_ASSEMBLY | 50 | -0.312289 | -0.774875 | 0.780453 | 0.96359 |
| GOBP_SULFUR_COMPOUND_BIOSYNTHETIC_PROCESS | 147 | -0.273817 | -0.7745 | 0.873939 | 0.963742 |
| GOBP_CARBOHYDRATE_BIOSYNTHETIC_PROCESS | 156 | -0.273127 | -0.7736 | 0.868605 | 0.964675 |
| GOBP_REGULATION_OF_VASCULAR_ENDOTHELIAL_GROWTH_FACTOR_RECEPT | 22 | -0.353829 | -0.772628 | 0.7743 | 0.965766 |
| GOBP_CHROMATIN_ORGANIZATION_INVOLVED_IN_REGULATION_OF_TRANSCRI | 108 | -0.286919 | -0.772107 | 0.849873 | 0.966144 |
| GOBP_DETECTION_OF_OTHER_ORGANISM | 15 | -0.383878 | -0.771944 | 0.755 | 0.965952 |
| GOBP_MITOTIC_CHROMOSOME_CONDENSATION | 15 | -0.382115 | -0.771594 | 0.738854 | 0.966055 |
| GOBP_DNA_BIOSYNTHETIC_PROCESS | 166 | -0.275023 | -0.771451 | 0.875147 | 0.965819 |
| GOBP_RECOMBINATIONAL_REPAIR | 133 | -0.274118 | -0.771426 | 0.865455 | 0.965417 |
| GOBP_STEROID_BIOSYNTHETIC_PROCESS | 128 | -0.275445 | -0.771076 | 0.853333 | 0.965521 |
| GOBP_REGULATION_OF_INSULIN_RECEPTOR_SIGNALING_PATHWAY | 51 | -0.311874 | -0.770569 | 0.795231 | 0.965895 |
| GOBP_SPERMATID_DIFFERENTIATION | 101 | -0.281961 | -0.770525 | 0.848101 | 0.965521 |

| | | | | | |
|---|---|---|---|---|---|
| GOBP_RESPONSE_TO_COCAINE | 28 | -0.345093 | -0.769797 | 0.783061 | 0.966195 |
| GOBP_CENTROSOME_DUPLICATION | 66 | -0.297773 | -0.769369 | 0.811828 | 0.966409 |
| GOBP_POSITIVE_REGULATION_OF_DNA_BIOSYNTHETIC_PROCESS | 59 | -0.305124 | -0.769054 | 0.807114 | 0.966424 |
| GOBP_DNA_CONFORMATION_CHANGE | 286 | -0.259027 | -0.768079 | 0.925556 | 0.967458 |
| GOBP_ORGANIC_CATION_TRANSPORT | 23 | -0.346031 | -0.766891 | 0.768868 | 0.968824 |
| GOBP_REGULATION_OF_MICROTUBULE_CYTOSKELETON_ORGANIZATION | 118 | -0.278802 | -0.766577 | 0.862794 | 0.968859 |
| GOBP_ORGANOPHOSPHATE_BIOSYNTHETIC_PROCESS | 487 | -0.252067 | -0.765563 | 0.961578 | 0.969946 |
| GOBP_REGULATION_OF_GLYCOGEN_METABOLIC_PROCESS | 26 | -0.345602 | -0.76504 | 0.781874 | 0.970314 |
| GOBP_CELLULAR_RESPONSE_TO_STARVATION | 132 | -0.27616 | -0.764593 | 0.883663 | 0.970541 |
| GOBP_CELL_COMMUNICATION_BY_ELECTRICAL_COUPLING_INVOLVED_IN_CAF | 15 | -0.381682 | -0.764576 | 0.766881 | 0.970122 |
| GOBP_CATECHOLAMINE_SECRETION | 20 | -0.356093 | -0.764369 | 0.769585 | 0.96998 |
| GOBP_REGULATION_OF_AMINO_ACID_TRANSPORT | 16 | -0.363674 | -0.764162 | 0.756714 | 0.969855 |
| GOBP_SMALL_MOLECULE_CATABOLIC_PROCESS | 340 | -0.254606 | -0.762654 | 0.936078 | 0.971612 |
| GOBP_MICROTUBULE_CYTOSKELETON_ORGANIZATION | 492 | -0.250737 | -0.762473 | 0.960375 | 0.971452 |
| GOBP_BASE_CONVERSION_OR_SUBSTITUTION_EDITING | 15 | -0.373094 | -0.762089 | 0.757724 | 0.971596 |
| GOBP_HISTONE_H3_K4_TRIMETHYLATION | 16 | -0.367831 | -0.761478 | 0.734177 | 0.972071 |
| GOBP_GLYCOSYL_COMPOUND_METABOLIC_PROCESS | 112 | -0.276449 | -0.760962 | 0.849817 | 0.972376 |
| GOBP_MITOTIC_CELL_CYCLE_ARREST | 16 | -0.36682 | -0.760651 | 0.773246 | 0.97238 |
| GOBP_SYNAPTIC_TRANSMISSION_GLUTAMATERGIC | 43 | -0.310299 | -0.759694 | 0.804035 | 0.973396 |
| GOBP_POSITIVE_REGULATION_OF_CYTOKINE_PRODUCTION_INVOLVED_IN_INFL | 16 | -0.376095 | -0.759337 | 0.73127 | 0.973506 |
| GOBP_ACTIN_FILAMENT_BASED_TRANSPORT | 19 | -0.352655 | -0.759274 | 0.758842 | 0.973156 |
| GOBP_PEPTIDYL_L_CYSTEINE_S_PALMITOYLATION | 21 | -0.352025 | -0.759217 | 0.75969 | 0.972794 |
| GOBP_POSITIVE_REGULATION_OF_PROTEASOMAL_UBIQUITIN_DEPENDENT_PR | 79 | -0.284144 | -0.758672 | 0.861111 | 0.973169 |
| GOBP_RELAXATION_OF_MUSCLE | 18 | -0.368026 | -0.758454 | 0.775081 | 0.973038 |
| GOBP_RESPONSE_TO_ENDOPLASMIC_RETICULUM_STRESS | 268 | -0.256861 | -0.758043 | 0.935094 | 0.97319 |
| GOBP_PROTEIN_LOCALIZATION_TO_CILIUM | 54 | -0.303984 | -0.757914 | 0.833568 | 0.972954 |
| GOBP_PHYSIOLOGICAL_CARDIAC_MUSCLE_HYPERTROPHY | 17 | -0.360226 | -0.757806 | 0.766169 | 0.972684 |
| GOBP_POSITIVE_REGULATION_OF_NERVOUS_SYSTEM_PROCESS | 20 | -0.355952 | -0.757292 | 0.7792 | 0.973032 |
| GOBP_METHYLATION | 308 | -0.25192 | -0.757243 | 0.943072 | 0.972666 |
| GOBP_SENSORY_PERCEPTION_OF_CHEMICAL_STIMULUS | 55 | -0.29925 | -0.757084 | 0.822888 | 0.972459 |
| GOBP_POSITIVE_REGULATION_OF_T_CELL_MEDIATED_CYTOTOXICITY | 23 | -0.341987 | -0.756451 | 0.779635 | 0.972974 |
| GOBP_RESPONSE_TO_LEPTIN | 16 | -0.368574 | -0.756065 | 0.771654 | 0.973094 |
| GOBP_HEMATOPOIETIC_PROGENITOR_CELL_DIFFERENTIATION | 137 | -0.269825 | -0.755919 | 0.888889 | 0.972885 |
| GOBP_REGULATION_OF_CILIUM_MOVEMENT | 16 | -0.360601 | -0.755625 | 0.757877 | 0.972888 |
| GOBP_SERINE_FAMILY_AMINO_ACID_METABOLIC_PROCESS | 35 | -0.324188 | -0.755545 | 0.807407 | 0.972566 |
| GOBP_PROTEIN_CONTAINING_COMPLEX_LOCALIZATION | 255 | -0.257251 | -0.754132 | 0.948124 | 0.974203 |
| GOBP_MICROTUBULE_BASED_MOVEMENT | 281 | -0.252314 | -0.75392 | 0.942077 | 0.974059 |
| GOBP_RESPONSE_TO_INSULIN | 222 | -0.255355 | -0.753633 | 0.935227 | 0.974018 |
| GOBP_NEGATIVE_REGULATION_OF_I_KAPPAB_KINASE_NF_KAPPAB_SIGNALING | 39 | -0.320029 | -0.752484 | 0.807092 | 0.975218 |
| GOBP_POSITIVE_REGULATION_OF_UBIQUITIN_DEPENDENT_PROTEIN_CATABOL | 93 | -0.276995 | -0.751879 | 0.872562 | 0.975635 |
| GOBP_POSITIVE_REGULATION_OF_TRANSCRIPTION_BY_RNA_POLYMERASE_I | 22 | -0.348453 | -0.751858 | 0.793323 | 0.975231 |
| GOBP_LIPID_PHOSPHORYLATION | 65 | -0.295926 | -0.750776 | 0.836592 | 0.976326 |
| GOBP_ZINC_ION_TRANSPORT | 21 | -0.339958 | -0.750506 | 0.768489 | 0.97627 |
| GOBP_DIOL_BIOSYNTHETIC_PROCESS | 20 | -0.345458 | -0.749531 | 0.787338 | 0.977242 |
| GOBP_NEGATIVE_REGULATION_OF_ORGANELLE_ASSEMBLY | 38 | -0.314688 | -0.749355 | 0.819103 | 0.977061 |
| GOBP_VITAMIN_TRANSPORT | 24 | -0.3416 | -0.749317 | 0.792126 | 0.976679 |
| GOBP_LIPOPROTEIN_METABOLIC_PROCESS | 113 | -0.276655 | -0.748898 | 0.875784 | 0.976815 |
| GOBP_REGULATION_OF_WNT_SIGNALING_PATHWAY | 269 | -0.251123 | -0.748884 | 0.952954 | 0.976398 |
| GOBP_RIG_I_SIGNALING_PATHWAY | 23 | -0.343676 | -0.747069 | 0.79289 | 0.978517 |
| GOBP_NUCLEAR_ENVELOPE_REASSEMBLY | 18 | -0.353698 | -0.745098 | 0.788274 | 0.980808 |
| GOBP_HEPARAN_SULFATE_PROTEOGLYCAN_METABOLIC_PROCESS | 24 | -0.3405 | -0.744823 | 0.786482 | 0.980753 |
| GOBP_REGULATION_OF_MULTI_ORGANISM_PROCESS | 32 | -0.321915 | -0.743679 | 0.822095 | 0.981855 |
| GOBP_REGULATION_OF_MEMBRANE_DEPOLARIZATION | 26 | -0.338765 | -0.742591 | 0.811728 | 0.982885 |
| GOBP_G2_DNA_DAMAGE_CHECKPOINT | 32 | -0.31712 | -0.742455 | 0.813253 | 0.982635 |
| GOBP_SYNAPTIC_VESICLE_EXOCYTOSIS | 68 | -0.291526 | -0.741775 | 0.871486 | 0.983113 |
| GOBP_POLYSACCHARIDE_BIOSYNTHETIC_PROCESS | 55 | -0.299119 | -0.741482 | 0.845833 | 0.983057 |
| GOBP_NUCLEOBASE_CONTAINING_SMALL_MOLECULE_CATABOLIC_PROCESS | 41 | -0.306202 | -0.741362 | 0.843517 | 0.982796 |
| GOBP_ANTEROGRADE_AXONAL_TRANSPORT | 43 | -0.304911 | -0.741233 | 0.82684 | 0.982543 |
| GOBP_INSULIN_SECRETION_INVOLVED_IN_CELLULAR_RESPONSE_TO_GLUCOS | 52 | -0.302187 | -0.740705 | 0.844101 | 0.982797 |
| GOBP_NEGATIVE_REGULATION_OF_SYNAPTIC_TRANSMISSION | 38 | -0.312218 | -0.740652 | 0.822653 | 0.982438 |
| GOBP_BODY_MORPHOGENESIS | 33 | -0.31833 | -0.739381 | 0.819259 | 0.983677 |
| GOBP_CHROMOSOME_CONDENSATION | 33 | -0.316891 | -0.73877 | 0.82009 | 0.98405 |
| GOBP_GLUCOSE_IMPORT | 50 | -0.298364 | -0.7387 | 0.844508 | 0.983712 |
| GOBP_NUCLEOSIDE_DIPHOSPHATE_METABOLIC_PROCESS | 129 | -0.265841 | -0.737266 | 0.903465 | 0.985204 |
| GOBP_NEGATIVE_REGULATION_OF_METAPHASE_ANAPHASE_TRANSITION_OF_( | 40 | -0.298086 | -0.736106 | 0.831412 | 0.986315 |
| GOBP_REGULATION_OF_PROTEIN_MODIFICATION_BY_SMALL_PROTEIN_CONJU | 219 | -0.253254 | -0.735662 | 0.945143 | 0.986474 |
| GOBP_PALLIUM_DEVELOPMENT | 100 | -0.272133 | -0.735225 | 0.897727 | 0.986599 |
| GOBP_CILIUM_OR_FLAGELLUM_DEPENDENT_CELL_MOTILITY | 74 | -0.284868 | -0.735054 | 0.875321 | 0.986395 |
| GOBP_NEGATIVE_REGULATION_OF_RNA_CATABOLIC_PROCESS | 55 | -0.291768 | -0.734055 | 0.847107 | 0.987247 |
| GOBP_DNA_REPLICATION | 260 | -0.2501 | -0.733821 | 0.955801 | 0.987102 |
| GOBP_MITOCHONDRIAL_CALCIUM_ION_TRANSMEMBRANE_TRANSPORT | 23 | -0.334473 | -0.732534 | 0.806107 | 0.988308 |
| GOBP_PHOSPHATIDYLINOSITOL_DEPHOSPHORYLATION | 29 | -0.325872 | -0.731717 | 0.807576 | 0.9889 |
| GOBP_PHOSPHATIDYLINOSITOL_METABOLIC_PROCESS | 160 | -0.257648 | -0.731713 | 0.939904 | 0.988469 |
| GOBP_CELLULAR_IRON_ION_HOMEOSTASIS | 58 | -0.289377 | -0.731128 | 0.851296 | 0.988814 |
| GOBP_GLYCOLYTIC_PROCESS_THROUGH_FRUCTOSE_6_PHOSPHATE | 25 | -0.332459 | -0.73042 | 0.829085 | 0.989293 |
| GOBP_INTRACILIARY_TRANSPORT_INVOLVED_IN_CILIUM_ASSEMBLY | 39 | -0.313271 | -0.73035 | 0.845152 | 0.988952 |
| GOBP_CARDIAC_MUSCLE_CELL_PROLIFERATION | 28 | -0.318468 | -0.727737 | 0.831071 | 0.991805 |
| GOBP_CELLULAR_AMINO_ACID_CATABOLIC_PROCESS | 80 | -0.274512 | -0.72635 | 0.88946 | 0.993049 |
| GOBP_LIPID_EXPORT_FROM_CELL | 43 | -0.299671 | -0.725771 | 0.84 | 0.993345 |
| GOBP_BONE_GROWTH | 15 | -0.361408 | -0.725502 | 0.795608 | 0.993228 |
| GOBP_HOMEOSTASIS_OF_NUMBER_OF_CELLS_WITHIN_A_TISSUE | 21 | -0.335928 | -0.72417 | 0.816746 | 0.994392 |
| GOBP_GLUTAMATE_SECRETION | 21 | -0.332486 | -0.723356 | 0.820919 | 0.994956 |
| GOBP_REGULATION_OF_GENE_SILENCING | 111 | -0.265542 | -0.722132 | 0.911729 | 0.995966 |
| GOBP_ENDOTHELIAL_CELL_DEVELOPMENT | 50 | -0.290637 | -0.721054 | 0.856552 | 0.996841 |
| GOBP_CERAMIDE_METABOLIC_PROCESS | 79 | -0.273571 | -0.720704 | 0.891051 | 0.996828 |
| GOBP_MACROMOLECULE_DEACYLATION | 96 | -0.270821 | -0.720314 | 0.895484 | 0.99685 |
| GOBP_NEGATIVE_REGULATION_OF_PROTEIN_MATURATION | 21 | -0.334753 | -0.720136 | 0.821483 | 0.996635 |
| GOBP_NADH_METABOLIC_PROCESS | 39 | -0.301338 | -0.720118 | 0.841246 | 0.996213 |
| GOBP_NEUROMUSCULAR_PROCESS | 49 | -0.292423 | -0.719296 | 0.874652 | 0.996765 |
| GOBP_MICROTUBULE_POLYMERIZATION | 66 | -0.279167 | -0.718438 | 0.889774 | 0.99732 |
| GOBP_POSITIVE_REGULATION_OF_CHROMOSOME_SEPARATION | 16 | -0.347898 | -0.717978 | 0.798046 | 0.997396 |
| GOBP_NEGATIVE_REGULATION_OF_RESPONSE_TO_DNA_DAMAGE_STIMULUS | 74 | -0.271312 | -0.717896 | 0.903567 | 0.997058 |
| GOBP_REGULATION_OF_MUSCLE_HYPERTROPHY | 42 | -0.293552 | -0.716989 | 0.869955 | 0.997696 |
| GOBP_PROTEIN_PALMITOYLATION | 29 | -0.311742 | -0.716711 | 0.830384 | 0.997603 |
| GOBP_TRANSPORT_ALONG_MICROTUBULE | 146 | -0.258177 | -0.714838 | 0.931076 | 0.999349 |
| GOBP_RESPONSE_TO_PROGESTERONE | 27 | -0.315567 | -0.714626 | 0.827327 | 0.999169 |
| GOBP_REGULATION_OF_PROTEIN_LOCALIZATION_TO_NUCLEUS | 107 | -0.261419 | -0.713604 | 0.915545 | 0.999922 |
| GOBP_GLYCINE_METABOLIC_PROCESS | 15 | -0.343575 | -0.712532 | 0.823817 | 1 |
| GOBP_REGULATION_OF_HORMONE_METABOLIC_PROCESS | 23 | -0.323024 | -0.710487 | 0.847962 | 1 |
| GOBP_REGULATION_OF_CYTOPLASMIC_TRANSLATION | 18 | -0.340625 | -0.709879 | 0.848 | 1 |
| GOBP_REGULATION_OF_MITOTIC_SISTER_CHROMATID_SEGREGATION | 44 | -0.293302 | -0.709076 | 0.883853 | 1 |
| GOBP_POSITIVE_REGULATION_OF_CHROMATIN_ORGANIZATION | 91 | -0.266612 | -0.708676 | 0.926489 | 1 |
| GOBP_REGULATION_OF_MICROTUBULE_POLYMERIZATION_OR_DEPOLYMERIZA | 69 | -0.27646 | -0.708265 | 0.905479 | 1 |
| GOBP_RESPONSE_TO_ACTIVITY | 49 | -0.288998 | -0.708162 | 0.871469 | 1 |
| GOBP_DOPAMINE_RECEPTOR_SIGNALING_PATHWAY | 27 | -0.319364 | -0.708136 | 0.832817 | 1 |
| GOBP_MITOTIC_G2_M_TRANSITION_CHECKPOINT | 34 | -0.308504 | -0.707946 | 0.856495 | 1 |
| GOBP_FATTY_ACID_BIOSYNTHETIC_PROCESS | 115 | -0.25815 | -0.706834 | 0.925971 | 1 |
| GOBP_NEGATIVE_REGULATION_OF_INTRINSIC_APOPTOTIC_SIGNALING_PATHW/ | 18 | -0.329383 | -0.706567 | 0.817035 | 1 |
| GOBP_MITOTIC_NUCLEAR_DIVISION | 267 | -0.240749 | -0.706361 | 0.973094 | 1 |
| GOBP_PROTEIN_POLYUBIQUITINATION | 311 | -0.237086 | -0.705055 | 0.992366 | 1 |
| GOBP_REGULATION_OF_TELOMERE_MAINTENANCE_VIA_TELOMERE_LENGTHEI | 57 | -0.283372 | -0.704645 | 0.886872 | 1 |
| GOBP_CYTOPLASMIC_MICROTUBULE_ORGANIZATION | 53 | -0.28616 | -0.704285 | 0.886846 | 1 |
| GOBP_REGULATION_OF_MICROTUBULE_BASED_MOVEMENT | 29 | -0.311029 | -0.704136 | 0.859031 | 1 |
| GOBP_RENAL_WATER_HOMEOSTASIS | 22 | -0.323089 | -0.703742 | 0.833595 | 1 |
| GOBP_HEPARAN_SULFATE_PROTEOGLYCAN_BIOSYNTHETIC_PROCESS | 20 | -0.321196 | -0.703622 | 0.830372 | 1 |
| GOBP_TRANSITION_METAL_ION_TRANSPORT | 96 | -0.258343 | -0.70331 | 0.926708 | 1 |
| GOBP_RAB_PROTEIN_SIGNAL_TRANSDUCTION | 21 | -0.321677 | -0.702763 | 0.818923 | 1 |
| GOBP_AXONEME_ASSEMBLY | 47 | -0.28317 | -0.700429 | 0.873596 | 1 |
| GOBP_T_CELL_HOMEOSTASIS | 33 | -0.29804 | -0.700361 | 0.85 | 1 |
| GOBP_NUCLEOSIDE_MONOPHOSPHATE_METABOLIC_PROCESS | 66 | -0.27245 | -0.700301 | 0.894452 | 1 |
| GOBP_PROTEIN_INSERTION_INTO_MITOCHONDRIAL_MEMBRANE_INVOLVED_IN_ | 30 | -0.308339 | -0.700256 | 0.865237 | 1 |
| GOBP_REGULATION_OF_GLUCONEOGENESIS | 38 | -0.294272 | -0.69955 | 0.868264 | 1 |
| GOBP_BODY_FLUID_SECRETION | 55 | -0.27729 | -0.69859 | 0.893646 | 1 |
| GOBP_MATERNAL_PLACENTA_DEVELOPMENT | 17 | -0.336245 | -0.698175 | 0.842623 | 1 |
| GOBP_CELLULAR_RESPONSE_TO_FATTY_ACID | 24 | -0.315466 | -0.698098 | 0.852308 | 1 |
| GOBP_MITOCHONDRION_MORPHOGENESIS | 17 | -0.329632 | -0.695284 | 0.847003 | 1 |
| GOBP_POSITIVE_REGULATION_OF_MULTICELLULAR_ORGANISM_GROWTH | 17 | -0.329648 | -0.69471 | 0.815873 | 1 |
| GOBP_VENTRAL_SPINAL_CORD_DEVELOPMENT | 17 | -0.330063 | -0.694026 | 0.814815 | 1 |
| GOBP_PROTEIN_LOCALIZATION_TO_MITOCHONDRION | 142 | -0.248943 | -0.69386 | 0.967153 | 1 |
| GOBP_REGULATION_OF_TRANSCRIPTION_ELONGATION_FROM_RNA_POLYMER/ | 29 | -0.306133 | -0.693033 | 0.854135 | 1 |
| GOBP_TELOMERE_CAPPING | 46 | -0.281378 | -0.692882 | 0.906815 | 1 |
| GOBP_GLUCAN_BIOSYNTHETIC_PROCESS | 34 | -0.293687 | -0.691742 | 0.87426 | 1 |
| GOBP_POSITIVE_REGULATION_OF_RESPONSE_TO_DNA_DAMAGE_STIMULUS | 92 | -0.260844 | -0.691571 | 0.930114 | 1 |
| GOBP_REGULATION_OF_GENE_SILENCING_BY_RNA | 98 | -0.254729 | -0.691422 | 0.94665 | 1 |
| GOBP_GLUTAMINE_FAMILY_AMINO_ACID_CATABOLIC_PROCESS | 20 | -0.323581 | -0.690358 | 0.833333 | 1 |
| GOBP_ORGANELLE_TRANSPORT_ALONG_MICROTUBULE | 75 | -0.268522 | -0.690172 | 0.928479 | 1 |
| GOBP_MUSCLE_HYPERTROPHY | 56 | -0.269283 | -0.689483 | 0.906849 | 1 |
| GOBP_MICROTUBULE_POLYMERIZATION_OR_DEPOLYMERIZATION | 95 | -0.258212 | -0.688891 | 0.926829 | 1 |
| GOBP_POSITIVE_REGULATION_OF_OXIDOREDUCTASE_ACTIVITY | 44 | -0.28704 | -0.688419 | 0.885057 | 1 |
| GOBP_EMBRYONIC_CRANIAL_SKELETON_MORPHOGENESIS | 28 | -0.300095 | -0.687596 | 0.860465 | 1 |
| GOBP_REGULATION_OF_STEM_CELL_DIFFERENTIATION | 91 | -0.255787 | -0.687448 | 0.947301 | 1 |
| GOBP_NEGATIVE_REGULATION_OF_STRIATED_MUSCLE_CELL_DIFFERENTIATIO | 23 | -0.313161 | -0.685646 | 0.858713 | 1 |
| GOBP_INTERSTRAND_CROSS_LINK_REPAIR | 54 | -0.272735 | -0.683141 | 0.906555 | 1 |
| GOBP_REGULATION_OF_RESPONSE_TO_DNA_DAMAGE_STIMULUS | 196 | -0.237369 | -0.682878 | 0.980747 | 1 |
| GOBP_REGULATION_OF_ACTIN_FILAMENT_BASED_MOVEMENT | 20 | -0.314408 | -0.682691 | 0.86465 | 1 |
| GOBP_REGULATION_OF_HEART_RATE_BY_CARDIAC_CONDUCTION | 17 | -0.321313 | -0.68152 | 0.849498 | 1 |
| GOBP_POSITIVE_REGULATION_OF_MRNA_SPLICING_VIA_SPLICEOSOME | 20 | -0.324955 | -0.680972 | 0.861371 | 1 |

| | | | | | |
|---|---|---|---|---|---|
| GOBP_REGULATION_OF_RIG_I_SIGNALING_PATHWAY | 17 | -0.322554 | -0.680784 | 0.839546 | 1 |
| GOBP_PROTEIN_TARGETING_TO_MITOCHONDRION | 97 | -0.251412 | -0.67904 | 0.948298 | 1 |
| GOBP_PROTEIN_K63_LINKED_DEUBIQUITINATION | 33 | -0.292784 | -0.67841 | 0.876106 | 1 |
| GOBP_BASE_EXCISION_REPAIR | 37 | -0.289345 | -0.67816 | 0.877143 | 1 |
| GOBP_POSITIVE_REGULATION_OF_TELOMERE_CAPPING | 17 | -0.329891 | -0.677388 | 0.843393 | 1 |
| GOBP_PHOSPHATIDYLCHOLINE_BIOSYNTHETIC_PROCESS | 31 | -0.295286 | -0.675984 | 0.883096 | 1 |
| GOBP_BETA_CATENIN_DESTRUCTION_COMPLEX_DISASSEMBLY | 19 | -0.313944 | -0.675685 | 0.846405 | 1 |
| GOBP_INSULIN_RECEPTOR_SIGNALING_PATHWAY | 112 | -0.242669 | -0.675672 | 0.961006 | 1 |
| GOBP_PROTEIN_LOCALIZATION_TO_KINETOCHORE | 19 | -0.319423 | -0.673435 | 0.853543 | 1 |
| GOBP_NEUTRAL_LIPID_CATABOLIC_PROCESS | 21 | -0.312227 | -0.673177 | 0.855721 | 1 |
| GOBP_INTERFERON_BETA_PRODUCTION | 50 | -0.26943 | -0.672206 | 0.921429 | 1 |
| GOBP_NEGATIVE_REGULATION_OF_PEPTIDYL_TYROSINE_PHOSPHORYLATION | 46 | -0.277024 | -0.671577 | 0.912979 | 1 |
| GOBP_NEGATIVE_REGULATION_OF_MITOCHONDRION_ORGANIZATION | 39 | -0.277035 | -0.669335 | 0.905045 | 1 |
| GOBP_MICROTUBULE_BUNDLE_FORMATION | 73 | -0.258273 | -0.668945 | 0.932969 | 1 |
| GOBP_CYTOPLASMIC_PATTERN_RECOGNITION_RECEPTOR_SIGNALING_PATHW | 28 | -0.295537 | -0.668783 | 0.895366 | 1 |
| GOBP_RNA_DECAPPING | 16 | -0.322191 | -0.668775 | 0.864297 | 1 |
| GOBP_POSITIVE_REGULATION_OF_CILIUM_ASSEMBLY | 20 | -0.307783 | -0.668334 | 0.868132 | 1 |
| GOBP_RESPONSE_TO_TESTOSTERONE | 32 | -0.28906 | -0.667118 | 0.888554 | 1 |
| GOBP_POSITIVE_REGULATION_OF_CHROMOSOME_ORGANIZATION | 152 | -0.23657 | -0.664697 | 0.976134 | 1 |
| GOBP_POSITIVE_REGULATION_OF_SIGNAL_TRANSDUCTION_BY_P53_CLASS_MI | 19 | -0.318167 | -0.66464 | 0.858748 | 1 |
| GOBP_POSITIVE_REGULATION_OF_HISTONE_H3_K4_METHYLATION | 16 | -0.323187 | -0.664183 | 0.8438 | 1 |
| GOBP_PYRIMIDINE_NUCLEOSIDE_MONOPHOSPHATE_METABOLIC_PROCESS | 16 | -0.319827 | -0.663654 | 0.851373 | 1 |
| GOBP_CENTRIOLE_ASSEMBLY | 40 | -0.277793 | -0.661055 | 0.90393 | 1 |
| GOBP_HISTONE_PHOSPHORYLATION | 35 | -0.275934 | -0.660026 | 0.910979 | 1 |
| GOBP_PROTEIN_DESTABILIZATION | 43 | -0.27361 | -0.658554 | 0.920548 | 1 |
| GOBP_REGULATION_OF_DNA_REPLICATION | 95 | -0.246862 | -0.657597 | 0.949239 | 1 |
| GOBP_OOGENESIS | 56 | -0.255336 | -0.657162 | 0.945302 | 1 |
| GOBP_POSITIVE_REGULATION_OF_NUCLEOCYTOPLASMIC_TRANSPORT | 53 | -0.263324 | -0.656051 | 0.924716 | 1 |
| GOBP_PROTEIN_HOMOOLIGOMERIZATION | 129 | -0.233781 | -0.654467 | 0.973171 | 1 |
| GOBP_MULTICELLULAR_ORGANISMAL_MOVEMENT | 35 | -0.274909 | -0.65409 | 0.903226 | 1 |
| GOBP_CARBOHYDRATE_CATABOLIC_PROCESS | 159 | -0.23339 | -0.654086 | 0.984831 | 1 |
| GOBP_RESPONSE_TO_IONIZING_RADIATION | 126 | -0.234996 | -0.65385 | 0.972488 | 1 |
| GOBP_MITOPHAGY | 24 | -0.297105 | -0.6532 | 0.889236 | 1 |
| GOBP_DETECTION_OF_MECHANICAL_STIMULUS_INVOLVED_IN_SENSORY_PER( | 19 | -0.306138 | -0.653124 | 0.872473 | 1 |
| GOBP_MITOTIC_SISTER_CHROMATID_COHESION | 26 | -0.28694 | -0.651552 | 0.896396 | 1 |
| GOBP_DNA_UNWINDING_INVOLVED_IN_DNA_REPLICATION | 16 | -0.317421 | -0.650369 | 0.859016 | 1 |
| GOBP_SPERM_EGG_RECOGNITION | 30 | -0.28117 | -0.649831 | 0.897281 | 1 |
| GOBP_MONOSACCHARIDE_CATABOLIC_PROCESS | 53 | -0.26388 | -0.649686 | 0.907231 | 1 |
| GOBP_VESICLE_TRANSPORT_ALONG_MICROTUBULE | 41 | -0.271011 | -0.649268 | 0.912333 | 1 |
| GOBP_ACTIVIN_RECEPTOR_SIGNALING_PATHWAY | 24 | -0.303374 | -0.648921 | 0.905573 | 1 |
| GOBP_EXECUTION_PHASE_OF_APOPTOSIS | 66 | -0.254961 | -0.648085 | 0.952767 | 1 |
| GOBP_TORC1_SIGNALING | 45 | -0.262934 | -0.647838 | 0.932584 | 1 |
| GOBP_HISTONE_H3_K4_METHYLATION | 48 | -0.264177 | -0.646925 | 0.92028 | 1 |
| GOBP_KINETOCHORE_ORGANIZATION | 21 | -0.292494 | -0.645657 | 0.912363 | 1 |
| GOBP_ENDOSOMAL_TRANSPORT | 218 | -0.224093 | -0.645639 | 0.990971 | 1 |
| GOBP_NEGATIVE_REGULATION_OF_EXTRINSIC_APOPTOTIC_SIGNALING_PATHW | 31 | -0.278551 | -0.644434 | 0.905547 | 1 |
| GOBP_B_CELL_HOMEOSTASIS | 28 | -0.281924 | -0.64228 | 0.906009 | 1 |
| GOBP_CHONDROCYTE_DEVELOPMENT | 17 | -0.307133 | -0.642145 | 0.902928 | 1 |
| GOBP_TOLL_LIKE_RECEPTOR_2_SIGNALING_PATHWAY | 16 | -0.310127 | -0.642142 | 0.884176 | 1 |
| GOBP_HISTONE_H2A_UBIQUITINATION | 26 | -0.290748 | -0.641581 | 0.898305 | 1 |
| GOBP_NUCLEOTIDE_EXCISION_REPAIR_DNA_GAP_FILLING | 23 | -0.290271 | -0.640709 | 0.895072 | 1 |
| GOBP_DOUBLE_STRAND_BREAK_REPAIR | 243 | -0.218731 | -0.640142 | 0.993355 | 1 |
| GOBP_HISTONE_METHYLATION | 116 | -0.233191 | -0.639703 | 0.98094 | 1 |
| GOBP_POSITIVE_REGULATION_OF_CYTOKINESIS | 27 | -0.279248 | -0.639642 | 0.895455 | 1 |
| GOBP_REGULATION_OF_DNA_TEMPLATED_TRANSCRIPTION_ELONGATION | 48 | -0.252423 | -0.637243 | 0.953957 | 1 |
| GOBP_MEIOSIS_I_CELL_CYCLE_PROCESS | 99 | -0.231557 | -0.635676 | 0.985 | 1 |
| GOBP_AUTOPHAGY_OF_MITOCHONDRION | 70 | -0.248985 | -0.635536 | 0.958612 | 1 |
| GOBP_CRANIAL_SKELETAL_SYSTEM_DEVELOPMENT | 40 | -0.265671 | -0.634667 | 0.918367 | 1 |
| GOBP_PEPTIDYL_LYSINE_METHYLATION | 112 | -0.232932 | -0.633993 | 0.983791 | 1 |
| GOBP_REGULATION_OF_AUTOPHAGY | 297 | -0.210989 | -0.630652 | 0.998901 | 1 |
| GOBP_DNA_DAMAGE_RESPONSE_SIGNAL_TRANSDUCTION_RESULTING_IN_TRA | 17 | -0.304842 | -0.630058 | 0.86645 | 1 |
| GOBP_POSITIVE_REGULATION_OF_DOUBLE_STRAND_BREAK_REPAIR | 38 | -0.261123 | -0.629327 | 0.946667 | 1 |
| GOBP_RESPONSE_TO_AMPHETAMINE | 22 | -0.292269 | -0.628991 | 0.904321 | 1 |
| GOBP_PEPTIDYL_LYSINE_TRIMETHYLATION | 43 | -0.258992 | -0.628643 | 0.965517 | 1 |
| GOBP_REGULATION_OF_VIRAL_INDUCED_CYTOPLASMIC_PATTERN_RECOGNITI | 19 | -0.296851 | -0.628055 | 0.88141 | 1 |
| GOBP_POSITIVE_REGULATION_OF_TELOMERE_MAINTENANCE_VIA_TELOMERE_ | 36 | -0.269688 | -0.628033 | 0.929088 | 1 |
| GOBP_NEGATIVE_REGULATION_OF_TELOMERE_MAINTENANCE | 32 | -0.271729 | -0.627672 | 0.93921 | 1 |
| GOBP_GLYCOSYL_COMPOUND_BIOSYNTHETIC_PROCESS | 38 | -0.265973 | -0.62765 | 0.940146 | 1 |
| GOBP_REGULATION_OF_DNA_DEPENDENT_DNA_REPLICATION | 45 | -0.256846 | -0.626181 | 0.948895 | 1 |
| GOBP_HISTONE_H2A_MONOUBIQUITINATION | 17 | -0.29762 | -0.62611 | 0.8867 | 1 |
| GOBP_RESPONSE_TO_ISOQUINOLINE_ALKALOID | 15 | -0.305464 | -0.624721 | 0.888147 | 1 |
| GOBP_DNA_INTEGRITY_CHECKPOINT | 146 | -0.221136 | -0.62366 | 0.996437 | 1 |
| GOBP_IMPORT_INTO_NUCLEUS | 138 | -0.221748 | -0.623228 | 0.989064 | 1 |
| GOBP_ORGANELLE_FISSION | 413 | -0.205151 | -0.622092 | 1 | 1 |
| GOBP_HISTONE_H3_K27_METHYLATION | 17 | -0.30332 | -0.621536 | 0.920398 | 1 |
| GOBP_PROTEIN_ACYLATION | 213 | -0.215016 | -0.620797 | 0.998851 | 1 |
| GOBP_PROTEIN_IMPORT | 167 | -0.218152 | -0.620022 | 0.992832 | 1 |
| GOBP_NON_CANONICAL_WNT_SIGNALING_PATHWAY | 127 | -0.224108 | -0.619227 | 0.98882 | 1 |
| GOBP_PROTEIN_MONOUBIQUITINATION | 66 | -0.241083 | -0.61894 | 0.963989 | 1 |
| GOBP_AMP_METABOLIC_PROCESS | 15 | -0.303605 | -0.614114 | 0.901454 | 1 |
| GOBP_ESTABLISHMENT_OF_PROTEIN_LOCALIZATION_TO_MITOCHONDRIAL_MEI | 53 | -0.245665 | -0.613842 | 0.963121 | 1 |
| GOBP_PYRIMIDINE_RIBONUCLEOTIDE_BIOSYNTHETIC_PROCESS | 18 | -0.289419 | -0.613521 | 0.908486 | 1 |
| GOBP_ENDOPLASMIC_RETICULUM_ORGANIZATION | 75 | -0.237402 | -0.612485 | 0.979452 | 1 |
| GOBP_LYMPHOCYTE_HOMEOSTASIS | 53 | -0.247159 | -0.611188 | 0.962025 | 1 |
| GOBP_HISTONE_H2A_ACETYLATION | 16 | -0.299109 | -0.608009 | 0.894828 | 1 |
| GOBP_CELL_CYCLE_DNA_REPLICATION | 62 | -0.235417 | -0.607661 | 0.969816 | 1 |
| GOBP_PROTEASOMAL_PROTEIN_CATABOLIC_PROCESS | 445 | -0.201315 | -0.607534 | 1 | 1 |
| GOBP_ORGANELLE_DISASSEMBLY | 100 | -0.223472 | -0.60717 | 0.992509 | 1 |
| GOBP_OOCYTE_DIFFERENTIATION | 34 | -0.256192 | -0.606663 | 0.955817 | 1 |
| GOBP_ENERGY_DERIVATION_BY_OXIDATION_OF_ORGANIC_COMPOUNDS | 240 | -0.208686 | -0.606618 | 1 | 1 |
| GOBP_POSITIVE_REGULATION_OF_DNA_REPAIR | 65 | -0.236097 | -0.606555 | 0.981767 | 1 |
| GOBP_POSITIVE_REGULATION_OF_MITOCHONDRIAL_FISSION | 15 | -0.291541 | -0.60639 | 0.914614 | 1 |
| GOBP_REGULATION_OF_MITOCHONDRIAL_FISSION | 24 | -0.27309 | -0.605907 | 0.931087 | 1 |
| GOBP_SINGLE_FERTILIZATION | 80 | -0.226322 | -0.605179 | 0.98248 | 1 |
| GOBP_NEGATIVE_REGULATION_OF_RESPONSE_TO_ENDOPLASMIC_RETICULUM | 40 | -0.24968 | -0.605026 | 0.954674 | 1 |
| GOBP_NEGATIVE_REGULATION_OF_MRNA_METABOLIC_PROCESS | 73 | -0.232039 | -0.604917 | 0.978836 | 1 |
| GOBP_NUCLEOBASE_CONTAINING_COMPOUND_TRANSPORT | 232 | -0.206629 | -0.604903 | 0.996603 | 1 |
| GOBP_PURINE_NUCLEOBASE_METABOLIC_PROCESS | 16 | -0.29436 | -0.604502 | 0.908646 | 1 |
| GOBP_NEURAL_NUCLEUS_DEVELOPMENT | 43 | -0.24613 | -0.603824 | 0.959941 | 1 |
| GOBP_NUCLEOBASE_CONTAINING_SMALL_MOLECULE_BIOSYNTHETIC_PROCES | 96 | -0.225101 | -0.603548 | 0.988491 | 1 |
| GOBP_TERMINATION_OF_RNA_POLYMERASE_II_TRANSCRIPTION | 35 | -0.258703 | -0.603379 | 0.941088 | 1 |
| GOBP_MODULATION_BY_HOST_OF_VIRAL_GENOME_REPLICATION | 17 | -0.284866 | -0.602942 | 0.896226 | 1 |
| GOBP_SNARE_COMPLEX_ASSEMBLY | 16 | -0.295451 | -0.602527 | 0.898305 | 1 |
| GOBP_FERTILIZATION | 101 | -0.22107 | -0.597929 | 0.99493 | 1 |
| GOBP_PROTEIN_STABILIZATION | 165 | -0.208097 | -0.597684 | 0.99521 | 1 |
| GOBP_POSITIVE_REGULATION_OF_AUTOPHAGY | 109 | -0.222246 | -0.595833 | 0.995037 | 1 |
| GOBP_PEPTIDYL_LYSINE_MODIFICATION | 354 | -0.199229 | -0.595269 | 1 | 1 |
| GOBP_PROTEIN_LOCALIZATION_TO_CHROMATIN | 28 | -0.261382 | -0.593865 | 0.937126 | 1 |
| GOBP_BLASTOCYST_GROWTH | 17 | -0.286758 | -0.593718 | 0.927445 | 1 |
| GOBP_REGULATION_OF_MRNA_METABOLIC_PROCESS | 297 | -0.199265 | -0.590162 | 1 | 1 |
| GOBP_REGULATION_OF_TELOMERE_CAPPING | 26 | -0.269308 | -0.588251 | 0.937097 | 1 |
| GOBP_GOLGI_TO_ENDOSOME_TRANSPORT | 17 | -0.281936 | -0.587682 | 0.926635 | 1 |
| GOBP_NEGATIVE_REGULATION_OF_PEPTIDYL_THREONINE_PHOSPHORYLATION | 18 | -0.270305 | -0.585828 | 0.920611 | 1 |
| GOBP_METAPHASE_ANAPHASE_TRANSITION_OF_CELL_CYCLE | 61 | -0.22976 | -0.585777 | 0.980085 | 1 |
| GOBP_ENDOPLASMIC_RETICULUM_TO_GOLGI_VESICLE_MEDIATED_TRANSPOR | 178 | -0.20722 | -0.58486 | 1 | 1 |
| GOBP_MACROMOLECULE_METHYLATION | 265 | -0.198936 | -0.584448 | 1 | 1 |
| GOBP_PROTEIN_ACETYLATION | 175 | -0.204877 | -0.583599 | 1 | 1 |
| GOBP_CELL_CYCLE_G2_M_PHASE_TRANSITION | 259 | -0.200073 | -0.582797 | 1 | 1 |
| GOBP_RNA_PHOSPHODIESTER_BOND_HYDROLYSIS_EXONUCLEOLYTIC | 43 | -0.238755 | -0.582244 | 0.967504 | 1 |
| GOBP_MAINTENANCE_OF_PROTEIN_LOCATION_IN_NUCLEUS | 21 | -0.269135 | -0.580837 | 0.93722 | 1 |
| GOBP_CYTOPLASMIC_PATTERN_RECOGNITION_RECEPTOR_SIGNALING_PATHW | 68 | -0.226636 | -0.58079 | 0.977573 | 1 |
| GOBP_MICROTUBULE_ANCHORING | 25 | -0.263684 | -0.580125 | 0.940989 | 1 |
| GOBP_PROTEIN_DNA_COMPLEX_SUBUNIT_ORGANIZATION | 222 | -0.198635 | -0.579826 | 1 | 1 |
| GOBP_REGULATION_OF_TELOMERE_MAINTENANCE | 76 | -0.223198 | -0.578918 | 0.989362 | 1 |
| GOBP_NEGATIVE_REGULATION_OF_ENDOPLASMIC_RETICULUM_STRESS_INDU | 15 | -0.278344 | -0.578836 | 0.913713 | 1 |
| GOBP_MITOCHONDRIAL_OUTER_MEMBRANE_PERMEABILIZATION | 52 | -0.230623 | -0.578225 | 0.980769 | 1 |
| GOBP_PROTEIN_TARGETING_TO_MEMBRANE | 189 | -0.199642 | -0.57771 | 0.998833 | 1 |
| GOBP_NEUTRAL_AMINO_ACID_TRANSPORT | 29 | -0.252792 | -0.576876 | 0.953593 | 1 |
| GOBP_REGULATION_OF_CARBOHYDRATE_CATABOLIC_PROCESS | 75 | -0.219569 | -0.575903 | 0.991946 | 1 |
| GOBP_MICROVILLUS_ORGANIZATION | 20 | -0.268785 | -0.574085 | 0.9344 | 1 |
| GOBP_CELLULAR_RESPONSE_TO_ATP | 15 | -0.278984 | -0.57405 | 0.927586 | 1 |
| GOBP_RIBONUCLEOSIDE_MONOPHOSPHATE_BIOSYNTHETIC_PROCESS | 31 | -0.239547 | -0.570462 | 0.957055 | 1 |
| GOBP_DNA_STRAND_ELONGATION_INVOLVED_IN_DNA_REPLICATION | 19 | -0.262558 | -0.570213 | 0.94822 | 1 |
| GOBP_GOLGI_TO_PLASMA_MEMBRANE_PROTEIN_TRANSPORT | 37 | -0.237099 | -0.566355 | 0.972384 | 1 |
| GOBP_PROTEIN_K48_LINKED_UBIQUITINATION | 60 | -0.221044 | -0.565839 | 0.982597 | 1 |
| GOBP_ESTABLISHMENT_OF_PROTEIN_LOCALIZATION_TO_MEMBRANE | 324 | -0.189211 | -0.565598 | 1 | 1 |
| GOBP_NUCLEIC_ACID_PHOSPHODIESTER_BOND_HYDROLYSIS | 269 | -0.190438 | -0.564891 | 1 | 1 |
| GOBP_POSITIVE_REGULATION_OF_HISTONE_DEACETYLATION | 16 | -0.272004 | -0.562737 | 0.948718 | 1 |
| GOBP_REGULATION_OF_INTRINSIC_APOPTOTIC_SIGNALING_PATHWAY_IN_RESI | 16 | -0.267722 | -0.562246 | 0.949097 | 1 |
| GOBP_HISTONE_H4_K16_ACETYLATION | 20 | -0.26619 | -0.562236 | 0.948113 | 1 |
| GOBP_DNA_SYNTHESIS_INVOLVED_IN_DNA_REPAIR | 49 | -0.225207 | -0.560102 | 0.984419 | 1 |
| GOBP_CELLULAR_RESPONSE_TO_OSMOTIC_STRESS | 31 | -0.239283 | -0.558468 | 0.974589 | 1 |
| GOBP_LIPOPROTEIN_BIOSYNTHETIC_PROCESS | 85 | -0.210253 | -0.558405 | 0.997375 | 1 |
| GOBP_REGULATION_OF_PROTEIN_DEACETYLATION | 37 | -0.235763 | -0.557952 | 0.982734 | 1 |
| GOBP_CELLULAR_RESPIRATION | 175 | -0.193817 | -0.557201 | 1 | 1 |
| GOBP_REGULATION_OF_DEFENSE_RESPONSE_TO_VIRUS | 60 | -0.219973 | -0.55646 | 0.991585 | 1 |
| GOBP_RETROGRADE_TRANSPORT_ENDOSOME_TO_PLASMA_MEMBRANE | 27 | -0.244357 | -0.555771 | 0.95814 | 1 |

| | | | | | |
|---|---|---|---|---|---|
| GOBP_SYNAPTONEMAL_COMPLEX_ORGANIZATION | 18 | -0.266287 | -0.554784 | 0.934084 | 1 |
| GOBP_REGULATION_OF_CHOLESTEROL_METABOLIC_PROCESS | 49 | -0.225364 | -0.552865 | 0.979138 | 1 |
| GOBP_CELLULAR_PROTEIN_COMPLEX_DISASSEMBLY | 193 | -0.191747 | -0.550948 | 1 | 1 |
| GOBP_REGULATION_OF_INTRINSIC_APOPTOTIC_SIGNALING_PATHWAY_BY_P53 | 24 | -0.245735 | -0.549921 | 0.954331 | 1 |
| GOBP_POSITIVE_REGULATION_OF_PROTEIN_DEACETYLATION | 19 | -0.258741 | -0.549476 | 0.952229 | 1 |
| GOBP_PROTEIN_CONTAINING_COMPLEX_DISASSEMBLY | 293 | -0.184569 | -0.548933 | 1 | 1 |
| GOBP_REGULATION_OF_CHROMOSOME_ORGANIZATION | 255 | -0.186383 | -0.548135 | 1 | 1 |
| GOBP_RNA_POLYMERASE_II_PREINITIATION_COMPLEX_ASSEMBLY | 22 | -0.247177 | -0.547233 | 0.959821 | 1 |
| GOBP_NUCLEUS_LOCALIZATION | 21 | -0.253731 | -0.54638 | 0.96672 | 1 |
| GOBP_DNA_REPLICATION_DEPENDENT_NUCLEOSOME_ORGANIZATION | 18 | -0.258607 | -0.544778 | 0.945736 | 1 |
| GOBP_CHROMOSOME_SEPARATION | 89 | -0.201737 | -0.543658 | 0.998686 | 1 |
| GOBP_SPINDLE_LOCALIZATION | 42 | -0.225804 | -0.542908 | 0.979109 | 1 |
| GOBP_PROTEIN_LOCALIZATION_TO_NUCLEUS | 231 | -0.184216 | -0.537754 | 1 | 1 |
| GOBP_SIGNAL_TRANSDUCTION_IN_RESPONSE_TO_DNA_DAMAGE | 115 | -0.194387 | -0.536985 | 1 | 1 |
| GOBP_REGULATION_OF_HISTONE_H3_K9_METHYLATION | 18 | -0.256616 | -0.534325 | 0.962791 | 1 |
| GOBP_ENDOCYTIC_RECYCLING | 43 | -0.22223 | -0.532269 | 0.989691 | 1 |
| GOBP_PROTEIN_METHYLATION | 153 | -0.184942 | -0.528239 | 1 | 1 |
| GOBP_POSITIVE_REGULATION_OF_TELOMERE_MAINTENANCE | 49 | -0.215163 | -0.527135 | 0.992837 | 1 |
| GOBP_REGULATION_OF_HISTONE_DEACETYLATION | 25 | -0.23664 | -0.525727 | 0.980153 | 1 |
| GOBP_MITOCHONDRIAL_TRANSPORT | 252 | -0.17743 | -0.523544 | 1 | 1 |
| GOBP_STRESS_GRANULE_ASSEMBLY | 22 | -0.240616 | -0.523424 | 0.97545 | 1 |
| GOBP_REGULATION_OF_CENTROSOME_CYCLE | 44 | -0.21755 | -0.523135 | 0.981455 | 1 |
| GOBP_POSITIVE_REGULATION_OF_MEMBRANE_PERMEABILITY | 60 | -0.206526 | -0.522746 | 0.993084 | 1 |
| GOBP_PROTEIN_LOCALIZATION_TO_MICROTUBULE_ORGANIZING_CENTER | 35 | -0.221042 | -0.522509 | 0.994118 | 1 |
| GOBP_MACROAUTOPHAGY | 290 | -0.175155 | -0.522338 | 1 | 1 |
| GOBP_POSITIVE_REGULATION_OF_HISTONE_METHYLATION | 32 | -0.223236 | -0.521257 | 0.980213 | 1 |
| GOBP_REGULATION_OF_DOUBLE_STRAND_BREAK_REPAIR_VIA_NONHOMOLOG | 27 | -0.232461 | -0.52073 | 0.971935 | 1 |
| GOBP_REGULATION_OF_UBIQUITIN_PROTEIN_TRANSFERASE_ACTIVITY | 49 | -0.213908 | -0.520525 | 0.988717 | 1 |
| GOBP_HISTONE_MRNA_METABOLIC_PROCESS | 24 | -0.237064 | -0.520283 | 0.978916 | 1 |
| GOBP_PROTEIN_LOCALIZATION_TO_VACUOLE | 61 | -0.204616 | -0.520184 | 0.997199 | 1 |
| GOBP_NEGATIVE_REGULATION_OF_CHROMOSOME_ORGANIZATION | 86 | -0.196801 | -0.512791 | 0.996109 | 1 |
| GOBP_SUBSTANTIA_NIGRA_DEVELOPMENT | 37 | -0.213687 | -0.509409 | 0.99133 | 1 |
| GOBP_MICROTUBULE_ORGANIZING_CENTER_ORGANIZATION | 130 | -0.180201 | -0.505004 | 1 | 1 |
| GOBP_REGULATION_OF_MRNA_SPLICING_VIA_SPLICEOSOME | 86 | -0.189919 | -0.504006 | 0.998728 | 1 |
| GOBP_COVALENT_CHROMATIN_MODIFICATION | 412 | -0.165476 | -0.499283 | 1 | 1 |
| GOBP_GOLGI_TO_VACUOLE_TRANSPORT | 19 | -0.235262 | -0.49754 | 0.966825 | 1 |
| GOBP_NUCLEAR_TRANSPORT | 310 | -0.166841 | -0.49744 | 1 | 1 |
| GOBP_AUTOPHAGOSOME_MATURATION | 38 | -0.20896 | -0.496579 | 0.994109 | 1 |
| GOBP_MITOTIC_DNA_INTEGRITY_CHECKPOINT | 98 | -0.184119 | -0.493928 | 0.997403 | 1 |
| GOBP_ESTABLISHMENT_OF_SPINDLE_ORIENTATION | 26 | -0.215054 | -0.490287 | 0.98941 | 1 |
| GOBP_NADPH_REGENERATION | 17 | -0.235593 | -0.487202 | 0.966981 | 1 |
| GOBP_REGULATION_OF_SPINDLE_ORGANIZATION | 37 | -0.205178 | -0.486192 | 0.989378 | 1 |
| GOBP_PEPTIDYL_LYSINE_ACETYLATION | 150 | -0.173 | -0.484186 | 1 | 1 |
| GOBP_GOLGI_VESICLE_TRANSPORT | 332 | -0.161639 | -0.481349 | 1 | 1 |
| GOBP_PROTEIN_TARGETING | 403 | -0.158999 | -0.478466 | 1 | 1 |
| GOBP_REGULATION_OF_CELL_CYCLE_G2_M_PHASE_TRANSITION | 208 | -0.165888 | -0.478005 | 1 | 1 |
| GOBP_POSITIVE_REGULATION_OF_DOUBLE_STRAND_BREAK_REPAIR_VIA_NON | 16 | -0.23177 | -0.474554 | 0.985099 | 1 |
| GOBP_RESOLUTION_OF_MEIOTIC_RECOMBINATION_INTERMEDIATES | 16 | -0.227645 | -0.472216 | 0.990385 | 1 |
| GOBP_RESPONSE_TO_MITOCHONDRIAL_DEPOLARISATION | 20 | -0.224003 | -0.46893 | 0.988818 | 1 |
| GOBP_MRNA_CLEAVAGE | 20 | -0.216899 | -0.468751 | 0.980539 | 1 |
| GOBP_CELLULAR_RESPONSE_TO_VIRUS | 54 | -0.183955 | -0.466154 | 0.998601 | 1 |
| GOBP_MEIOTIC_CHROMOSOME_SEPARATION | 22 | -0.211629 | -0.465276 | 0.990755 | 1 |
| GOBP_NEGATIVE_REGULATION_OF_DNA_RECOMBINATION | 36 | -0.200658 | -0.46514 | 0.994245 | 1 |
| GOBP_NEGATIVE_REGULATION_OF_MRNA_SPLICING_VIA_SPLICEOSOME | 20 | -0.220976 | -0.461377 | 0.984252 | 1 |
| GOBP_HISTONE_H3_ACETYLATION | 58 | -0.1809 | -0.4591 | 1 | 1 |
| GOBP_NCRNA_CATABOLIC_PROCESS | 34 | -0.1945 | -0.457903 | 0.992493 | 1 |
| GOBP_MRNA_SPLICE_SITE_SELECTION | 32 | -0.196479 | -0.45751 | 0.992636 | 1 |
| GOBP_PROTEIN_DEMETHYLATION | 32 | -0.196939 | -0.45634 | 0.99537 | 1 |
| GOBP_PROTEIN_LOCALIZATION_TO_MICROTUBULE | 18 | -0.214083 | -0.455123 | 0.976228 | 1 |
| GOBP_NEGATIVE_REGULATION_OF_ATP_METABOLIC_PROCESS | 20 | -0.211368 | -0.454179 | 0.983103 | 1 |
| GOBP_ERROR_FREE_TRANSLESION_SYNTHESIS | 22 | -0.207884 | -0.453944 | 0.982759 | 1 |
| GOBP_PHOSPHORYLATED_CARBOHYDRATE_DEPHOSPHORYLATION | 16 | -0.217342 | -0.448431 | 0.987382 | 1 |
| GOBP_ERROR_PRONE_TRANSLESION_SYNTHESIS | 21 | -0.207871 | -0.445716 | 0.987635 | 1 |
| GOBP_2_OXOGLUTARATE_METABOLIC_PROCESS | 15 | -0.222101 | -0.444762 | 0.988487 | 1 |
| GOBP_MITOTIC_SPINDLE_ORGANIZATION | 108 | -0.164699 | -0.44378 | 1 | 1 |
| GOBP_PROTEIN_SUMOYLATION | 76 | -0.169069 | -0.443047 | 1 | 1 |
| GOBP_REGULATION_OF_MRNA_PROCESSING | 125 | -0.161242 | -0.441785 | 1 | 1 |
| GOBP_TRANSCRIPTION_BY_RNA_POLYMERASE_I | 65 | -0.174842 | -0.441758 | 0.998634 | 1 |
| GOBP_CELLULAR_RESPONSE_TO_IONIZING_RADIATION | 56 | -0.176636 | -0.441152 | 0.998621 | 1 |
| GOBP_INOSITOL_PHOSPHATE_CATABOLIC_PROCESS | 17 | -0.214744 | -0.440946 | 0.988449 | 1 |
| GOBP_NUCLEOTIDE_EXCISION_REPAIR | 106 | -0.161017 | -0.439031 | 1 | 1 |
| GOBP_DNA_TEMPLATED_TRANSCRIPTION_ELONGATION | 111 | -0.157862 | -0.435439 | 1 | 1 |
| GOBP_RNA_CATABOLIC_PROCESS | 384 | -0.143682 | -0.435287 | 1 | 1 |
| GOBP_RRNA_CATABOLIC_PROCESS | 19 | -0.207358 | -0.434315 | 0.996689 | 1 |
| GOBP_POSITIVE_REGULATION_OF_MRNA_PROCESSING | 30 | -0.185981 | -0.431463 | 0.99403 | 1 |
| GOBP_MICROTUBULE_CYTOSKELETON_ORGANIZATION_INVOLVED_IN_MITOSIS | 128 | -0.154558 | -0.431327 | 1 | 1 |
| GOBP_REGULATION_OF_PROTEIN_SUMOYLATION | 22 | -0.200345 | -0.431224 | 0.987097 | 1 |
| GOBP_MITOCHONDRIAL_ELECTRON_TRANSPORT_NADH_TO_UBIQUINONE | 53 | -0.167773 | -0.430057 | 0.998611 | 1 |
| GOBP_NUCLEAR_TRANSCRIBED_MRNA_CATABOLIC_PROCESS_EXONUCLEOLY | 35 | -0.182057 | -0.427777 | 0.99854 | 1 |
| GOBP_TELOMERE_ORGANIZATION | 153 | -0.151413 | -0.42704 | 1 | 1 |
| GOBP_TRANSCRIPTION_ELONGATION_FROM_RNA_POLYMERASE_II_PROMOTER | 83 | -0.159121 | -0.420323 | 1 | 1 |
| GOBP_SKELETAL_MUSCLE_CONTRACTION | 27 | -0.185927 | -0.419707 | 0.996885 | 1 |
| GOBP_PORE_COMPLEX_ASSEMBLY | 20 | -0.196008 | -0.418625 | 0.995283 | 1 |
| GOBP_MRNA_TRANSPORT | 144 | -0.150954 | -0.415701 | 1 | 1 |
| GOBP_CELLULAR_RESPONSE_TO_ARSENIC_CONTAINING_SUBSTANCE | 17 | -0.201372 | -0.414927 | 0.988216 | 1 |
| GOBP_REGULATION_OF_SIGNAL_TRANSDUCTION_BY_P53_CLASS_MEDIATOR | 161 | -0.145935 | -0.41204 | 1 | 1 |
| GOBP_MRNA_EXPORT_FROM_NUCLEUS | 109 | -0.149475 | -0.409372 | 1 | 1 |
| GOBP_POSITIVE_REGULATION_OF_GENE_EXPRESSION_EPIGENETIC | 51 | -0.158308 | -0.396048 | 1 | 1 |
| GOBP_REGULATION_OF_DNA_REPAIR | 116 | -0.142211 | -0.39225 | 1 | 1 |
| GOBP_TRANSCRIPTION_PREINITIATION_COMPLEX_ASSEMBLY | 34 | -0.16852 | -0.391615 | 0.997097 | 1 |
| GOBP_PEPTIDYL_LYSINE_DIMETHYLATION | 20 | -0.181035 | -0.39123 | 0.996764 | 1 |
| GOBP_TRNA_WOBBLE_BASE_MODIFICATION | 19 | -0.184149 | -0.388419 | 0.996825 | 1 |
| GOBP_CELLULAR_RESPONSE_TO_STEROL_DEPLETION | 16 | -0.179606 | -0.381659 | 0.995276 | 1 |
| GOBP_RESPONSE_TO_STEROL_DEPLETION | 18 | -0.179627 | -0.381147 | 0.998382 | 1 |
| GOBP_REGULATION_OF_DOUBLE_STRAND_BREAK_REPAIR | 78 | -0.140906 | -0.373321 | 1 | 1 |
| GOBP_PROTEIN_MODIFICATION_BY_SMALL_PROTEIN_REMOVAL | 261 | -0.125845 | -0.371673 | 1 | 1 |
| GOBP_REGULATION_OF_VIRAL_TRANSCRIPTION | 41 | -0.153566 | -0.367532 | 0.998563 | 1 |
| GOBP_RNA_LOCALIZATION | 222 | -0.124514 | -0.364691 | 1 | 1 |
| GOBP_NUCLEOLAR_LARGE_RRNA_TRANSCRIPTION_BY_RNA_POLYMERASE_I | 17 | -0.171583 | -0.359501 | 0.998454 | 1 |
| GOBP_DNA_DAMAGE_RESPONSE_DETECTION_OF_DNA_DAMAGE | 37 | -0.149471 | -0.353307 | 1 | 1 |
| GOBP_ALTERNATIVE_MRNA_SPLICING_VIA_SPLICEOSOME | 60 | -0.13658 | -0.349311 | 1 | 1 |
| GOBP_REGULATION_OF_DOUBLE_STRAND_BREAK_REPAIR_VIA_HOMOLOGOUS | 43 | -0.142736 | -0.346575 | 1 | 1 |
| GOBP_REGULATION_OF_LYSOSOMAL_LUMEN_PH | 15 | -0.163506 | -0.337564 | 0.998374 | 1 |
| GOBP_NUCLEAR_EXPORT | 192 | -0.117943 | -0.335621 | 1 | 1 |
| GOBP_ACETYL_COA_BIOSYNTHETIC_PROCESS | 21 | -0.15029 | -0.326539 | 1 | 1 |
| GOBP_RNA_SPLICING | 421 | -0.103344 | -0.313999 | 1 | 1 |
| GOBP_POLY_A_PLUS_MRNA_EXPORT_FROM_NUCLEUS | 19 | -0.146194 | -0.313878 | 1 | 1 |
| GOBP_NUCLEOTIDE_EXCISION_REPAIR_DNA_DAMAGE_RECOGNITION | 23 | -0.143134 | -0.31031 | 1 | 1 |
| GOBP_REGULATION_OF_MRNA_3_END_PROCESSING | 28 | -0.133758 | -0.302439 | 1 | 1 |
| GOBP_MRNA_3_END_PROCESSING | 97 | -0.109136 | -0.291296 | 1 | 1 |
| GOBP_MITOCHONDRIAL_RNA_METABOLIC_PROCESS | 47 | -0.109638 | -0.26947 | 1 | 0.999994 |
