## Supplementary material for "Chemokine Receptor 1 and its associated immune pathway are downregulated in SF3B1^MT^ blood and non-blood cancers": Table_S12

**Table S12. GSEA analyses of RNA-seq data in $SF3B1^{MT}$ versus $SF3B1^{WT}$ BRCA patients.** See table S16 for patient information.

**GSEA Positive** Hematopoietic/Immune pathways are highlighted in red.

| Gene sets | SIZE | ES | NES | NOM p-val | FDR q-val |
|---|---|---|---|---|---|
| GOBP_RESPONSE_TO_FOOD | 32 | 0.627459 | 2.061886 | 0 | 0.359644 |
| GOBP_CELLULAR_RESPONSE_TO_COPPER_ION | 24 | 0.65176 | 1.999237 | 0 | 0.437998 |
| GOBP_CELLULAR_RESPONSE_TO_CADMIUM_ION | 34 | 0.599923 | 1.996225 | 0.004545 | 0.302714 |
| GOBP_ATRIOVENTRICULAR_VALVE_DEVELOPMENT | 25 | 0.655753 | 1.993761 | 0 | 0.233072 |
| GOBP_POSITIVE_REGULATION_OF_BMP_SIGNALING_PATHWAY | 31 | 0.613936 | 1.96783 | 0 | 0.258686 |
| GOBP_NEURON_CELL_CELL_ADHESION | 16 | 0.716162 | 1.94203 | 0 | 0.284973 |
| GOBP_STRESS_RESPONSE_TO_METAL_ION | 15 | 0.711264 | 1.926665 | 0.003521 | 0.285312 |
| GOBP_CEREBELLAR_PURKINJE_CELL_LAYER_DEVELOPMENT | 22 | 0.65883 | 1.922618 | 0 | 0.258229 |
| GOBP_CELLULAR_RESPONSE_TO_ZINC_ION | 20 | 0.659457 | 1.874444 | 0 | 0.372419 |
| GOBP_REGULATION_OF_RECEPTOR_BINDING | 25 | 0.62914 | 1.87104 | 0 | 0.344339 |
| GOBP_RESPONSE_TO_TYPE_I_INTERFERON | 85 | 0.489193 | 1.871025 | 0 | 0.313036 |
| GOBP_RESPONSE_TO_CADMIUM_ION | 56 | 0.503781 | 1.862757 | 0.005495 | 0.310157 |
| GOBP_RESPONSE_TO_COPPER_ION | 38 | 0.543597 | 1.852912 | 0 | 0.314777 |
| GOBP_CEREBELLAR_CORTEX_DEVELOPMENT | 43 | 0.520871 | 1.825189 | 0 | 0.37852 |
| GOBP_ADRENAL_GLAND_DEVELOPMENT | 21 | 0.621265 | 1.813379 | 0.003984 | 0.394781 |
| GOBP_ANTIGEN_PROCESSING_AND_PRESENTATION_OF_ENDOGENOUS | 19 | 0.635768 | 1.80989 | 0.006993 | 0.38156 |
| GOBP_REGULATION_OF_RESPONSE_TO_FOOD | 15 | 0.666635 | 1.804794 | 0.018315 | 0.375354 |
| GOBP_TYPE_B_PANCREATIC_CELL_DEVELOPMENT | 20 | 0.596444 | 1.802365 | 0 | 0.361532 |
| GOBP_POSITIVE_REGULATION_OF_LEUKOCYTE_DEGRANULATION | 23 | 0.603637 | 1.79155 | 0.011538 | 0.376778 |
| GOBP_CELL_DIFFERENTIATION_IN_HINDBRAIN | 18 | 0.635894 | 1.782532 | 0.007491 | 0.385345 |
| GOBP_RESPONSE_TO_LITHIUM_ION | 19 | 0.626503 | 1.771808 | 0.003876 | 0.405223 |
| GOBP_REGULATION_OF_BMP_SIGNALING_PATHWAY | 85 | 0.447122 | 1.758473 | 0 | 0.4273 |
| GOBP_DECIDUALIZATION | 19 | 0.615845 | 1.742614 | 0.021053 | 0.463816 |
| GOBP_URETER_DEVELOPMENT | 18 | 0.610455 | 1.740576 | 0.014545 | 0.451189 |
| GOBP_ACTIVATION_OF_PROTEIN_KINASE_A_ACTIVITY | 17 | 0.609496 | 1.738621 | 0.017544 | 0.439919 |
| GOBP_COLLAGEN_CATABOLIC_PROCESS | 46 | 0.494698 | 1.73454 | 0 | 0.437002 |
| GOBP_EXTRACELLULAR_MATRIX_DISASSEMBLY | 76 | 0.451373 | 1.730636 | 0 | 0.4342 |
| GOBP_DOPAMINE_SECRETION | 32 | 0.535515 | 1.719005 | 0.004184 | 0.457097 |
| GOBP_POSITIVE_REGULATION_OF_LIPID_STORAGE | 25 | 0.575992 | 1.712469 | 0.0181 | 0.463838 |
| GOBP_LUNG_CELL_DIFFERENTIATION | 20 | 0.592038 | 1.711065 | 0.010753 | 0.452577 |
| GOBP_REGULATION_OF_NEUROTRANSMITTER_TRANSPORT | 95 | 0.425596 | 1.701554 | 0 | 0.473471 |
| GOBP_REGULATION_OF_RESPONSE_TO_EXTRACELLULAR_STIMULUS | 22 | 0.568735 | 1.696127 | 0.008439 | 0.479066 |
| GOBP_RESPONSE_TO_OXYGEN_RADICAL | 26 | 0.555847 | 1.689051 | 0.019685 | 0.490736 |
| GOBP_REGULATION_OF_BROWN_FAT_CELL_DIFFERENTIATION | 21 | 0.568619 | 1.674391 | 0.010638 | 0.529733 |
| GOBP_ENTEROENDOCRINE_CELL_DIFFERENTIATION | 25 | 0.545145 | 1.666492 | 0.004739 | 0.546006 |
| GOBP_RESPONSE_TO_BMP | 154 | 0.394889 | 1.663435 | 0 | 0.544426 |
| GOBP_RESPONSE_TO_AMYLOID_BETA | 46 | 0.458775 | 1.654864 | 0.017341 | 0.563957 |
| GOBP_MODULATION_BY_SYMBIONT_OF_ENTRY_INTO_HOST | 45 | 0.473242 | 1.649952 | 0 | 0.569558 |
| GOBP_REGULATION_OF_KIDNEY_DEVELOPMENT | 30 | 0.51479 | 1.645348 | 0.022222 | 0.572294 |
| GOBP_EXTRACELLULAR_MATRIX_ASSEMBLY | 44 | 0.476057 | 1.644063 | 0 | 0.563062 |
| GOBP_ANDROGEN_RECEPTOR_SIGNALING_PATHWAY | 38 | 0.488325 | 1.639498 | 0.005236 | 0.567168 |
| GOBP_ANTIGEN_PROCESSING_AND_PRESENTATION_OF_ENDOGENOUS | 26 | 0.522256 | 1.638114 | 0.018349 | 0.559934 |
| GOBP_RESPONSE_TO_ZINC_ION | 52 | 0.46665 | 1.638103 | 0 | 0.547122 |
| GOBP_CAMP_METABOLIC_PROCESS | 21 | 0.546512 | 1.629112 | 0.016064 | 0.568874 |
| GOBP_ORGANISM_EMERGENCE_FROM_PROTECTIVE_STRUCTURE | 22 | 0.55804 | 1.629027 | 0.024291 | 0.556868 |
| GOBP_CATECHOLAMINE_SECRETION | 50 | 0.466519 | 1.628595 | 0 | 0.546326 |
| GOBP_RESPONSE_TO_VITAMIN | 82 | 0.420113 | 1.624607 | 0.007874 | 0.547828 |
| GOBP_AMINE_TRANSPORT | 90 | 0.405031 | 1.59875 | 0 | 0.640359 |
| GOBP_CARDIAC_MUSCLE_CELL_CONTRACTION | 69 | 0.430914 | 1.59838 | 0 | 0.629173 |
| GOBP_REGULATION_OF_SUPEROXIDE_METABOLIC_PROCESS | 30 | 0.492363 | 1.592475 | 0.018605 | 0.64017 |
| GOBP_POSITIVE_REGULATION_OF_MACROPHAGE_DERIVED_FOAM_CEL | 18 | 0.577539 | 1.587615 | 0.039427 | 0.647745 |
| GOBP_POSITIVE_REGULATION_OF_TRANSMEMBRANE_RECEPTOR_PRO | 99 | 0.401619 | 1.58696 | 0.009091 | 0.637558 |
| GOBP_REGULATION_OF_VESICLE_FUSION | 23 | 0.533678 | 1.586265 | 0.030189 | 0.628507 |
| GOBP_CARDIOBLAST_DIFFERENTIATION | 17 | 0.565058 | 1.582562 | 0.037879 | 0.632824 |
| GOBP_SECRETORY_GRANULE_ORGANIZATION | 31 | 0.492422 | 1.581362 | 0.009217 | 0.625705 |
| GOBP_CEREBELLAR_CORTEX_FORMATION | 22 | 0.52647 | 1.580847 | 0.018519 | 0.61675 |
| GOBP_POSTSYNAPTIC_MEMBRANE_ORGANIZATION | 36 | 0.502637 | 1.57777 | 0.038462 | 0.618696 |
| GOBP_RENAL_WATER_HOMEOSTASIS | 35 | 0.472113 | 1.572663 | 0.023923 | 0.626912 |
| GOBP_CARDIAC_RIGHT_VENTRICLE_MORPHOGENESIS | 19 | 0.548667 | 1.570008 | 0.03125 | 0.626833 |
| GOBP_PULMONARY_VALVE_MORPHOGENESIS | 17 | 0.567147 | 1.569029 | 0.050179 | 0.620089 |
| GOBP_ENDODERM_DEVELOPMENT | 74 | 0.415606 | 1.568263 | 0.014286 | 0.613487 |
| GOBP_POSITIVE_REGULATION_OF_BROWN_FAT_CELL_DIFFERENTIATIO | 15 | 0.58883 | 1.565038 | 0.043165 | 0.615796 |
| GOBP_POSITIVE_REGULATION_OF_NEUROTRANSMITTER_TRANSPORT | 22 | 0.530096 | 1.561635 | 0.022222 | 0.619269 |
| GOBP_POSITIVE_REGULATION_OF_REGULATED_SECRETORY_PATHWAY | 51 | 0.429674 | 1.561413 | 0.010526 | 0.610556 |
| GOBP_ATRIAL_SEPTUM_DEVELOPMENT | 22 | 0.524293 | 1.552092 | 0.052045 | 0.636175 |
| GOBP_ECTODERM_DEVELOPMENT | 18 | 0.536787 | 1.549165 | 0.030043 | 0.637547 |
| GOBP_NEGATIVE_REGULATION_OF_OXIDATIVE_STRESS_INDUCED_CEL | 40 | 0.460636 | 1.548571 | 0.029268 | 0.630318 |
| GOBP_REGULATION_OF_MACROPHAGE_DERIVED_FOAM_CELL_DIFFERE | 31 | 0.48388 | 1.544225 | 0.021459 | 0.638024 |
| GOBP_COLUMNAR_CUBOIDAL_EPITHELIAL_CELL_DIFFERENTIATION | 75 | 0.400752 | 1.543996 | 0 | 0.629759 |
| GOBP_ACTIN_MEDIATED_CELL_CONTRACTION | 116 | 0.371976 | 1.54283 | 0 | 0.625468 |
| GOBP_PERICARDIUM_DEVELOPMENT | 19 | 0.536645 | 1.542068 | 0.048276 | 0.619649 |
| GOBP_REGULATION_OF_LIPID_STORAGE | 52 | 0.43764 | 1.539637 | 0.011429 | 0.62054 |
| GOBP_MEGAKARYOCYTE_DEVELOPMENT | 16 | 0.566368 | 1.538924 | 0.048507 | 0.614551 |
| GOBP_TRANSMISSION_OF_NERVE_IMPULSE | 55 | 0.433554 | 1.53837 | 0.006369 | 0.608078 |
| GOBP_ATRIAL_SEPTUM_MORPHOGENESIS | 15 | 0.588513 | 1.534272 | 0.060284 | 0.614756 |
| GOBP_ZINC_ION_HOMEOSTASIS | 37 | 0.460897 | 1.534165 | 0.035556 | 0.60707 |
| GOBP_POLYSACCHARIDE_CATABOLIC_PROCESS | 24 | 0.52484 | 1.529102 | 0.05 | 0.618789 |
| GOBP_PULMONARY_VALVE_DEVELOPMENT | 21 | 0.528301 | 1.527463 | 0.036 | 0.616487 |
| GOBP_CARDIAC_ATRIUM_DEVELOPMENT | 34 | 0.461872 | 1.52547 | 0.022422 | 0.617116 |
| GOBP_METANEPHROS_MORPHOGENESIS | 31 | 0.474965 | 1.522385 | 0.02381 | 0.62077 |
| GOBP_PROTEIN_PALMITOYLATION | 33 | 0.468584 | 1.522048 | 0.037736 | 0.61448 |
| GOBP_NEGATIVE_REGULATION_OF_AMINE_TRANSPORT | 24 | 0.510879 | 1.521541 | 0.048 | 0.608434 |
| GOBP_SENSORY_PERCEPTION_OF_SMELL | 55 | 0.411764 | 1.517657 | 0.005464 | 0.615888 |
| GOBP_MONOAMINE_TRANSPORT | 75 | 0.389876 | 1.515785 | 0 | 0.615683 |
| GOBP_PRESYNAPTIC_ENDOCYTOSIS | 58 | 0.408972 | 1.514754 | 0.020408 | 0.611911 |
| GOBP_CELL_FATE_DETERMINATION | 37 | 0.463923 | 1.514362 | 0.033654 | 0.60593 |
| GOBP_SYNAPTIC_VESICLE_RECYCLING | 71 | 0.400179 | 1.512016 | 0.007092 | 0.607737 |
| GOBP_CARDIAC_ATRIUM_MORPHOGENESIS | 28 | 0.478913 | 1.508886 | 0.033195 | 0.611397 |
| GOBP_TISSUE_REGENERATION | 69 | 0.403741 | 1.508592 | 0.007463 | 0.60566 |
| GOBP_CELLULAR_RESPONSE_TO_OXYGEN_RADICAL | 22 | 0.507346 | 1.49996 | 0.051095 | 0.630321 |
| GOBP_DOPAMINE_TRANSPORT | 46 | 0.434714 | 1.496396 | 0.030928 | 0.637014 |
| GOBP_BRANCH_ELONGATION_OF_AN_EPITHELIUM | 17 | 0.54458 | 1.494993 | 0.094545 | 0.635099 |
| GOBP_POSTSYNAPSE_ASSEMBLY | 28 | 0.483864 | 1.491619 | 0.035714 | 0.641781 |
| GOBP_PRESYNAPSE_ORGANIZATION | 46 | 0.41 | 1.490367 | 0.018293 | 0.639696 |
| GOBP_AMYLOID_FIBRIL_FORMATION | 23 | 0.488948 | 1.485752 | 0.052419 | 0.651037 |
| GOBP_KERATINIZATION | 108 | 0.369421 | 1.485285 | 0 | 0.64607 |
| GOBP_CARDIAC_CELL_DEVELOPMENT | 71 | 0.37961 | 1.485175 | 0.007937 | 0.639835 |
| GOBP_METANEPHRIC_EPITHELIUM_DEVELOPMENT | 24 | 0.493058 | 1.484068 | 0.055118 | 0.63705 |
| GOBP_POSITIVE_REGULATION_OF_CELLULAR_RESPONSE_TO_INSULIN_ | 23 | 0.494916 | 1.481605 | 0.06015 | 0.639797 |
| GOBP_SYNAPTIC_MEMBRANE_ADHESION | 24 | 0.489247 | 1.481353 | 0.049808 | 0.63414 |
| GOBP_NEGATIVE_REGULATION_OF_DNA_DAMAGE_RESPONSE_SIGNAL_ | 15 | 0.556874 | 1.477484 | 0.05618 | 0.641287 |
| GOBP_REGULATION_OF_ACTIN_FILAMENT_BASED_MOVEMENT | 37 | 0.435292 | 1.477195 | 0.032407 | 0.63618 |
| GOBP_HEMATOPOIETIC_STEM_CELL_PROLIFERATION | 24 | 0.470536 | 1.474657 | 0.060714 | 0.639218 |
| GOBP_METENCEPHALON_DEVELOPMENT | 97 | 0.371904 | 1.473484 | 0.007692 | 0.637676 |
| GOBP_REGULATION_OF_BRANCHING_INVOLVED_IN_URETERIC_BUD_M | 20 | 0.498276 | 1.467738 | 0.05668 | 0.651987 |
| GOBP_PROTEIN_LOCALIZATION_TO_CELL_SURFACE | 65 | 0.398793 | 1.467365 | 0.018987 | 0.647486 |
| GOBP_HETEROPHILIC_CELL_CELL_ADHESION_VIA_PLASMA_MEMBRAN | 45 | 0.428315 | 1.466666 | 0.041667 | 0.643879 |
| GOBP_NEGATIVE_REGULATION_OF_INTRINSIC_APOPTOTIC_SIGNALING_ | 30 | 0.454785 | 1.466392 | 0.046154 | 0.638883 |
| GOBP_NEUROEPITHELIAL_CELL_DIFFERENTIATION | 19 | 0.5083 | 1.464924 | 0.067925 | 0.63826 |
| GOBP_CHAPERONE_MEDIATED_AUTOPHAGY | 16 | 0.538347 | 1.463747 | 0.075188 | 0.636337 |
| GOBP_GLANDULAR_EPITHELIAL_CELL_DIFFERENTIATION | 45 | 0.418564 | 1.461242 | 0.025641 | 0.640112 |
| GOBP_COLLAGEN_METABOLIC_PROCESS | 101 | 0.367411 | 1.460298 | 0 | 0.638541 |
| GOBP_REGULATION_OF_RENAL_SYSTEM_PROCESS | 29 | 0.462372 | 1.458922 | 0.044177 | 0.638466 |
| GOBP_NEGATIVE_REGULATION_OF_CARDIAC_MUSCLE_CELL_PROLIFER | 15 | 0.548719 | 1.458878 | 0.067376 | 0.632963 |
| GOBP_INTERMEDIATE_FILAMENT_BASED_PROCESS | 49 | 0.409534 | 1.458128 | 0.025641 | 0.629982 |
| GOBP_METANEPHROS_DEVELOPMENT | 81 | 0.37992 | 1.457652 | 0.008475 | 0.626431 |
| GOBP_CELLULAR_SODIUM_ION_HOMEOSTASIS | 18 | 0.514335 | 1.455573 | 0.082143 | 0.628278 |
| GOBP_LUNG_EPITHELIUM_DEVELOPMENT | 33 | 0.452118 | 1.453613 | 0.049505 | 0.630126 |
| GOBP_REGULATION_OF_INTRINSIC_APOPTOTIC_SIGNALING_PATHWAY_ | 37 | 0.437996 | 1.452935 | 0.053812 | 0.627322 |
| GOBP_NEGATIVE_REGULATION_OF_REGULATED_SECRETORY_PATHWA | 22 | 0.500687 | 1.452168 | 0.070588 | 0.625155 |
| GOBP_WALKING_BEHAVIOR | 24 | 0.483636 | 1.447507 | 0.070755 | 0.638736 |
| GOBP_WOUND_HEALING_SPREADING_OF_EPIDERMAL_CELLS | 18 | 0.505636 | 1.446578 | 0.081181 | 0.637656 |
| GOBP_ENDOCARDIAL_CUSHION_DEVELOPMENT | 42 | 0.425642 | 1.443917 | 0.020942 | 0.642045 |
| GOBP_NEGATIVE_REGULATION_OF_RECEPTOR_BINDING | 15 | 0.526329 | 1.443752 | 0.082031 | 0.63747 |
| GOBP_REGULATION_OF_EXTRINSIC_APOPTOTIC_SIGNALING_PATHWAY_ | 40 | 0.43274 | 1.441169 | 0.042781 | 0.641281 |
| GOBP_REGULATION_OF_PRESYNAPSE_ORGANIZATION | 29 | 0.460634 | 1.439456 | 0.046729 | 0.642523 |
| GOBP_REGULATION_OF_SYNAPTIC_TRANSMISSION_GABAERGIC | 30 | 0.452073 | 1.436323 | 0.083333 | 0.648572 |
| GOBP_NEGATIVE_REGULATION_OF_AUTOPHAGY | 81 | 0.369865 | 1.43293 | 0.015152 | 0.655713 |
| GOBP_NEGATIVE_REGULATION_OF_APOPTOTIC_SIGNALING_PATHWAY | 214 | 0.333424 | 1.432023 | 0 | 0.65412 |
| GOBP_REGULATION_OF_TRANSMEMBRANE_RECEPTOR_PROTEIN_SERI | 231 | 0.323679 | 1.430581 | 0 | 0.654011 |
| GOBP_RENAL_SODIUM_EXCRETION | 18 | 0.502736 | 1.428478 | 0.091633 | 0.656973 |
| GOBP_VESICLE_FUSION_TO_PLASMA_MEMBRANE | 20 | 0.483821 | 1.428307 | 0.081481 | 0.652608 |
| GOBP_REGULATION_OF_EXOCYTOSIS | 196 | 0.328746 | 1.423935 | 0 | 0.663417 |
| GOBP_STEROID_CATABOLIC_PROCESS | 24 | 0.464849 | 1.421885 | 0.099585 | 0.666325 |

**GSEA Negative** Hematopoietic/Immune pathways are highlighted in green.

| Gene sets | SIZE | ES | NES | NOM p-val | FDR q-val |
|---|---|---|---|---|---|
| GOBP_IMMUNOGLOBULIN_PRODUCTION | 184 | -0.612389 | -1.998158 | 0 | 0.454609 |
| GOBP_HUMORAL_IMMUNE_RESPONSE_MEDIATED_BY_CIRCULATING_IMMUN | 136 | -0.610603 | -1.952248 | 0 | 0.422982 |
| GOBP_COMPLEMENT_ACTIVATION | 151 | -0.584523 | -1.889209 | 0 | 0.494282 |
| GOBP_REGULATION_OF_HUMORAL_IMMUNE_RESPONSE | 121 | -0.587812 | -1.865636 | 0 | 0.4347 |
| GOBP_HUMORAL_IMMUNE_RESPONSE | 305 | -0.533638 | -1.799556 | 0 | 0.461809 |
| GOBP_PRODUCTION_OF_MOLECULAR_MEDIATOR_OF_IMMUNE_RESPONSE | 268 | -0.530955 | -1.770304 | 0 | 0.4995 |
| GOBP_B_CELL_RECEPTOR_SIGNALING_PATHWAY | 117 | -0.558401 | -1.764235 | 0.001076 | 0.457934 |
| GOBP_REGULATION_OF_COMPLEMENT_ACTIVATION | 101 | -0.563924 | -1.743289 | 0 | 0.482469 |
| GOBP_B_CELL_MEDIATED_IMMUNITY | 205 | -0.507858 | -1.670891 | 0 | 0.667833 |
| GOBP_ANTIBACTERIAL_HUMORAL_RESPONSE | 50 | -0.590313 | -1.662323 | 0.001222 | 0.676648 |
| GOBP_POSITIVE_REGULATION_OF_B_CELL_ACTIVATION | 135 | -0.522902 | -1.657498 | 0 | 0.663761 |
| GOBP_ANTIMICROBIAL_HUMORAL_IMMUNE_RESPONSE_MEDIATED_BY_ANTIM | 54 | -0.580444 | -1.654872 | 0.00361 | 0.637475 |
| GOBP_REGULATION_OF_B_CELL_ACTIVATION | 178 | -0.492559 | -1.600826 | 0 | 0.68381 |
| GOBP_ANTIMICROBIAL_HUMORAL_RESPONSE | 110 | -0.500851 | -1.589319 | 0.00221 | 0.666845 |
| GOBP_INNATE_IMMUNE_RESPONSE_IN_MUCOSA | 19 | -0.634749 | -1.535146 | 0.019553 | 0.785177 |
| GOBP_REGULATION_OF_B_CELL_PROLIFERATION | 61 | -0.526007 | -1.533498 | 0.010551 | 0.78086 |
| GOBP_LYMPHOCYTE_MEDIATED_IMMUNITY | 334 | -0.456174 | -1.53163 | 0 | 0.777206 |
| GOBP_B_CELL_ACTIVATION | 292 | -0.447549 | -1.511996 | 0 | 0.830211 |
| GOBP_B_CELL_PROLIFERATION | 80 | -0.498968 | -1.495192 | 0.014673 | 0.901692 |
| GOBP_CHRONIC_INFLAMMATORY_RESPONSE | 19 | -0.619299 | -1.478995 | 0.04533 | 0.977944 |
| GOBP_ADAPTIVE_IMMUNE_RESPONSE_BASED_ON_SOMATIC_RECOMBINATIO | 340 | -0.435794 | -1.477493 | 0.00101 | 0.974998 |
| GOBP_POSITIVE_REGULATION_OF_MONOCYTE_CHEMOTAXIS | 19 | -0.608546 | -1.454185 | 0.041779 | 1 |
| GOBP_REGULATION_OF_LYMPHOCYTE_ACTIVATION | 482 | -0.371932 | -1.266874 | 0.011033 | 1 |
| GOBP_REGULATION_OF_IMMUNE_EFFECTOR_PROCESS | 438 | -0.362042 | -1.235125 | 0.035176 | 1 |
| GOBP_LEUKOCYTE_MIGRATION | 480 | -0.359304 | -1.232723 | 0.021127 | 1 |
| GOBP_PHAGOCYTOSIS_RECOGNITION | 83 | -0.590163 | -1.807779 | 0 | 0.522227 |
| GOBP_NEGATIVE_REGULATION_OF_EXECUTION_PHASE_OF_APOPTOSIS | 15 | -0.740732 | -1.699132 | 0 | 0.657485 |
| GOBP_DNA_METHYLATION_INVOLVED_IN_GAMETE_GENERATION | 17 | -0.707133 | -1.682879 | 0.002786 | 0.698789 |
| GOBP_DNA_REPLICATION_DEPENDENT_NUCLEOSOME_ORGANIZATION | 25 | -0.654511 | -1.679785 | 0.006588 | 0.660386 |
| GOBP_DOPAMINERGIC_NEURON_DIFFERENTIATION | 34 | -0.617055 | -1.63871 | 0.004957 | 0.710456 |
| GOBP_DEFENSE_RESPONSE_TO_BACTERIUM | 261 | -0.488501 | -1.637951 | 0 | 0.676385 |
| GOBP_RESPONSE_TO_FUNGUS | 35 | -0.603759 | -1.637819 | 0.010204 | 0.641897 |
| GOBP_OLIGOSACCHARIDE_BIOSYNTHETIC_PROCESS | 17 | -0.70794 | -1.635113 | 0.011158 | 0.625918 |
| GOBP_DEFENSE_RESPONSE_TO_FUNGUS | 22 | -0.651531 | -1.623794 | 0.013333 | 0.670316 |
| GOBP_NEUROMUSCULAR_PROCESS_CONTROLLING_POSTURE | 15 | -0.707515 | -1.620173 | 0.01122 | 0.663354 |
| GOBP_NEGATIVE_REGULATION_OF_ADENYLATE_CYCLASE_ACTIVITY | 18 | -0.684231 | -1.612505 | 0.015172 | 0.687073 |
| GOBP_REGULATION_OF_EXECUTION_PHASE_OF_APOPTOSIS | 23 | -0.643665 | -1.601944 | 0.008253 | 0.73226 |
| GOBP_FC_EPSILON_RECEPTOR_SIGNALING_PATHWAY | 164 | -0.493057 | -1.60163 | 0.003165 | 0.705138 |
| GOBP_RDNA_HETEROCHROMATIN_ASSEMBLY | 30 | -0.608884 | -1.599354 | 0.014286 | 0.667913 |
| GOBP_CHROMATIN_ORGANIZATION_INVOLVED_IN_REGULATION_OF_TRANSC | 132 | -0.502632 | -1.596575 | 0.005382 | 0.662612 |
| GOBP_FC_RECEPTOR_MEDIATED_STIMULATORY_SIGNALING_PATHWAY | 141 | -0.498417 | -1.594619 | 0.004296 | 0.652846 |
| GOBP_SERINE_FAMILY_AMINO_ACID_BIOSYNTHETIC_PROCESS | 21 | -0.648687 | -1.584841 | 0.012162 | 0.676067 |
| GOBP_POSITIVE_REGULATION_OF_HUMORAL_IMMUNE_RESPONSE | 17 | -0.67715 | -1.575089 | 0.013736 | 0.725382 |
| GOBP_CIRCADIAN_SLEEP_WAKE_CYCLE_SLEEP | 17 | -0.66822 | -1.573235 | 0.013351 | 0.716451 |
| GOBP_FAT_SOLUBLE_VITAMIN_METABOLIC_PROCESS | 41 | -0.572045 | -1.571715 | 0.003778 | 0.705911 |
| GOBP_SLEEP | 27 | -0.607508 | -1.569271 | 0.016927 | 0.703032 |
| GOBP_PIRNA_METABOLIC_PROCESS | 18 | -0.651988 | -1.563067 | 0.019691 | 0.728123 |
| GOBP_SPERM_CAPACITATION | 22 | -0.629967 | -1.557716 | 0.015089 | 0.746705 |
| GOBP_FATTY_ACID_DERIVATIVE_METABOLIC_PROCESS | 80 | -0.509912 | -1.553898 | 0.003452 | 0.755106 |
| GOBP_IMMUNE_RESPONSE_REGULATING_SIGNALING_PATHWAY | 490 | -0.450975 | -1.548723 | 0 | 0.774845 |
| GOBP_VASODILATION | 34 | -0.569759 | -1.54789 | 0.009162 | 0.761776 |
| GOBP_POSITIVE_REGULATION_OF_ANTIGEN_RECEPTOR_MEDIATED_SIGNALI | 21 | -0.635875 | -1.545938 | 0.030263 | 0.757661 |
| GOBP_CIRCADIAN_SLEEP_WAKE_CYCLE | 22 | -0.619891 | -1.538036 | 0.028571 | 0.800501 |
| GOBP_ANTIGEN_RECEPTOR_MEDIATED_SIGNALING_PATHWAY | 302 | -0.457309 | -1.53574 | 0 | 0.798514 |
| GOBP_EPOXYGENASE_P450_PATHWAY | 18 | -0.639972 | -1.527593 | 0.023192 | 0.791155 |
| GOBP_NEGATIVE_REGULATION_OF_DNA_TEMPLATED_TRANSCRIPTION_ELON | 17 | -0.663175 | -1.524893 | 0.022973 | 0.794852 |
| GOBP_PERIPHERAL_NERVOUS_SYSTEM_DEVELOPMENT | 72 | -0.510077 | -1.513965 | 0.014994 | 0.865506 |
| GOBP_FATTY_ACID_DERIVATIVE_BIOSYNTHETIC_PROCESS | 60 | -0.523883 | -1.513921 | 0.011723 | 0.848618 |
| GOBP_DRUG_CATABOLIC_PROCESS | 23 | -0.612147 | -1.513423 | 0.031165 | 0.836007 |
| GOBP_NEGATIVE_REGULATION_OF_GENE_EXPRESSION_EPIGENETIC | 104 | -0.484477 | -1.510304 | 0.009978 | 0.828056 |
| GOBP_REGULATION_OF_CIRCADIAN_SLEEP_WAKE_CYCLE | 17 | -0.644032 | -1.500253 | 0.032877 | 0.891715 |
| GOBP_DRUG_METABOLIC_PROCESS | 42 | -0.540927 | -1.495645 | 0.027027 | 0.914186 |
| GOBP_NEGATIVE_REGULATION_OF_LYASE_ACTIVITY | 24 | -0.589274 | -1.485834 | 0.033898 | 0.966447 |
| GOBP_MEMBRANE_INVAGINATION | 129 | -0.465772 | -1.484917 | 0.006543 | 0.958336 |
| GOBP_DEOXYRIBONUCLEOSIDE_MONOPHOSPHATE_METABOLIC_PROCESS | 16 | -0.626078 | -1.484684 | 0.04749 | 0.944223 |
| GOBP_NEGATIVE_REGULATION_OF_LIPASE_ACTIVITY | 15 | -0.641292 | -1.474894 | 0.052113 | 0.982136 |
| GOBP_LYMPH_NODE_DEVELOPMENT | 16 | -0.632246 | -1.474157 | 0.054167 | 0.972832 |
| GOBP_OLFACTORY_LOBE_DEVELOPMENT | 26 | -0.564656 | -1.4646 | 0.039894 | 1 |
| GOBP_PHOSPHATIDYLETHANOLAMINE_BIOSYNTHETIC_PROCESS | 15 | -0.628508 | -1.461605 | 0.045723 | 1 |
| GOBP_FC_RECEPTOR_SIGNALING_PATHWAY | 234 | -0.441303 | -1.460147 | 0.001024 | 1 |
| GOBP_ORGAN_OR_TISSUE_SPECIFIC_IMMUNE_RESPONSE | 32 | -0.551769 | -1.458865 | 0.044271 | 1 |
| GOBP_DIENCEPHALON_DEVELOPMENT | 63 | -0.492596 | -1.457777 | 0.024362 | 1 |
| GOBP_RESPONSE_TO_XENOBIOTIC_STIMULUS | 118 | -0.457751 | -1.45705 | 0.018319 | 1 |
| GOBP_URONIC_ACID_METABOLIC_PROCESS | 15 | -0.638396 | -1.456776 | 0.049435 | 1 |
| GOBP_RHYTHMIC_BEHAVIOR | 40 | -0.527392 | -1.454319 | 0.034956 | 1 |
| GOBP_C21_STEROID_HORMONE_METABOLIC_PROCESS | 37 | -0.531819 | -1.450941 | 0.039851 | 1 |
| GOBP_UNSATURATED_FATTY_ACID_METABOLIC_PROCESS | 108 | -0.464235 | -1.450839 | 0.007795 | 1 |
| GOBP_NEGATIVE_REGULATION_OF_B_CELL_PROLIFERATION | 17 | -0.616062 | -1.448921 | 0.064343 | 1 |
| GOBP_POSITIVE_REGULATION_OF_TRIGLYCERIDE_METABOLIC_PROCESS | 20 | -0.602729 | -1.447515 | 0.053405 | 1 |
| GOBP_CALCIUM_DEPENDENT_CELL_CELL_ADHESION_VIA_PLASMA_MEMBRA | 41 | -0.526012 | -1.444136 | 0.034956 | 1 |
| GOBP_SPERMATID_NUCLEUS_DIFFERENTIATION | 16 | -0.623558 | -1.441362 | 0.075034 | 1 |
| GOBP_T_CELL_CHEMOTAXIS | 24 | -0.565604 | -1.440714 | 0.062918 | 1 |
| GOBP_SERINE_FAMILY_AMINO_ACID_METABOLIC_PROCESS | 43 | -0.513366 | -1.437632 | 0.041924 | 1 |
| GOBP_POSITIVE_REGULATION_OF_TRANSCRIPTION_REGULATORY_REGION_D | 23 | -0.577496 | -1.436347 | 0.04712 | 1 |
| GOBP_ARACHIDONIC_ACID_METABOLIC_PROCESS | 57 | -0.494743 | -1.43632 | 0.035545 | 1 |
| GOBP_POSITIVE_REGULATION_OF_MEIOTIC_CELL_CYCLE | 21 | -0.586408 | -1.43624 | 0.055256 | 1 |
| GOBP_REGULATION_OF_ANTIGEN_RECEPTOR_MEDIATED_SIGNALING_PATHW | 63 | -0.480779 | -1.435945 | 0.027123 | 1 |
| GOBP_HEAT_GENERATION | 16 | -0.616851 | -1.433474 | 0.064607 | 1 |
| GOBP_HISTONE_EXCHANGE | 51 | -0.503576 | -1.431823 | 0.042184 | 1 |
| GOBP_HIGH_DENSITY_LIPOPROTEIN_PARTICLE_REMODELING | 15 | -0.622487 | -1.430349 | 0.057061 | 1 |
| GOBP_NEURON_FATE_COMMITMENT | 46 | -0.514083 | -1.42881 | 0.03805 | 1 |
| GOBP_NEGATIVE_REGULATION_OF_GLIAL_CELL_DIFFERENTIATION | 17 | -0.603866 | -1.427158 | 0.053147 | 1 |
| GOBP_INTERLEUKIN_7_MEDIATED_SIGNALING_PATHWAY | 25 | -0.568043 | -1.420887 | 0.063144 | 1 |
| GOBP_POSITIVE_REGULATION_OF_B_CELL_PROLIFERATION | 40 | -0.517679 | -1.420638 | 0.051661 | 1 |
| GOBP_CELLULAR_RESPONSE_TO_STEROL | 21 | -0.575228 | -1.417011 | 0.06546 | 1 |
| GOBP_OLIGOSACCHARIDE_METABOLIC_PROCESS | 49 | -0.490312 | -1.416949 | 0.042857 | 1 |
| GOBP_MALE_MEIOTIC_NUCLEAR_DIVISION | 42 | -0.512429 | -1.416794 | 0.049261 | 1 |
| GOBP_PHAGOCYTOSIS | 358 | -0.418328 | -1.416731 | 0.001018 | 1 |
| GOBP_SULFUR_AMINO_ACID_BIOSYNTHETIC_PROCESS | 17 | -0.599201 | -1.4153 | 0.067227 | 1 |
| GOBP_OLEFINIC_COMPOUND_METABOLIC_PROCESS | 114 | -0.456211 | -1.414589 | 0.016411 | 1 |
| GOBP_TRANSCRIPTION_ELONGATION_FROM_RNA_POLYMERASE_I_PROMOTE | 30 | -0.53148 | -1.412644 | 0.053665 | 1 |
| GOBP_PURINERGIC_NUCLEOTIDE_RECEPTOR_SIGNALING_PATHWAY | 30 | -0.535709 | -1.412257 | 0.060802 | 1 |
| GOBP_RESPONSE_TO_INTERLEUKIN_7 | 35 | -0.516975 | -1.411323 | 0.056653 | 1 |
| GOBP_CELLULAR_RESPONSE_TO_CHOLESTEROL | 16 | -0.608311 | -1.404269 | 0.074022 | 1 |
| GOBP_POSITIVE_REGULATION_OF_NITRIC_OXIDE_SYNTHASE_BIOSYNTHETIC_ | 15 | -0.612679 | -1.402134 | 0.075391 | 1 |
| GOBP_ICOSANOID_METABOLIC_PROCESS | 114 | -0.448521 | -1.402018 | 0.023333 | 1 |
| GOBP_EXECUTION_PHASE_OF_APOPTOSIS | 86 | -0.458909 | -1.401826 | 0.036613 | 1 |
| GOBP_ATP_DEPENDENT_CHROMATIN_REMODELING | 81 | -0.463348 | -1.398249 | 0.027682 | 1 |
| GOBP_CHROMATIN_SILENCING | 65 | -0.474689 | -1.39731 | 0.038551 | 1 |
| GOBP_SULFUR_AMINO_ACID_METABOLIC_PROCESS | 37 | -0.513236 | -1.394881 | 0.06125 | 1 |
| GOBP_HETEROCHROMATIN_ORGANIZATION | 70 | -0.471417 | -1.393757 | 0.034762 | 1 |
| GOBP_REGULATION_OF_ADENYLATE_CYCLASE_ACTIVITY | 29 | -0.528085 | -1.391043 | 0.072751 | 1 |
| GOBP_OOCYTE_MATURATION | 23 | -0.54861 | -1.386986 | 0.087516 | 1 |
| GOBP_APOPTOTIC_DNA_FRAGMENTATION | 23 | -0.548802 | -1.384009 | 0.075198 | 1 |
| GOBP_GENE_SILENCING | 312 | -0.410804 | -1.382883 | 0.006129 | 1 |
| GOBP_NUCLEOSOME_ASSEMBLY | 119 | -0.436559 | -1.379781 | 0.034103 | 1 |
| GOBP_LEUKOCYTE_MIGRATION_INVOLVED_IN_INFLAMMATORY_RESPONSE | 18 | -0.586275 | -1.378873 | 0.099723 | 1 |
| GOBP_DNA_PACKAGING | 206 | -0.417667 | -1.375153 | 0.020747 | 1 |
| GOBP_LIPOPOLYSACCHARIDE_MEDIATED_SIGNALING_PATHWAY | 57 | -0.472806 | -1.372013 | 0.055147 | 1 |
| GOBP_EMBRYONIC_EYE_MORPHOGENESIS | 32 | -0.520417 | -1.371855 | 0.08312 | 1 |
| GOBP_STORE_OPERATED_CALCIUM_ENTRY | 18 | -0.576221 | -1.369517 | 0.105048 | 1 |
| GOBP_PYROPTOSIS | 18 | -0.581255 | -1.367099 | 0.096552 | 1 |
| GOBP_REGULATION_OF_FLAGELLATED_SPERM_MOTILITY | 15 | -0.597527 | -1.365762 | 0.096685 | 1 |
| GOBP_SPLICEOSOMAL_TRI_SNRNP_COMPLEX_ASSEMBLY | 19 | -0.569194 | -1.364116 | 0.097826 | 1 |
| GOBP_THYMIC_T_CELL_SELECTION | 22 | -0.551028 | -1.363729 | 0.094444 | 1 |
| GOBP_EMBRYONIC_CAMERA_TYPE_EYE_MORPHOGENESIS | 24 | -0.549809 | -1.362488 | 0.098202 | 1 |
| GOBP_MOTOR_NEURON_AXON_GUIDANCE | 29 | -0.52252 | -1.360567 | 0.098945 | 1 |
| GOBP_FATTY_ACID_DERIVATIVE_CATABOLIC_PROCESS | 16 | -0.588248 | -1.358857 | 0.108484 | 1 |
| GOBP_VITAMIN_BIOSYNTHETIC_PROCESS | 19 | -0.561356 | -1.357985 | 0.111413 | 1 |
| GOBP_DETECTION_OF_CHEMICAL_STIMULUS_INVOLVED_IN_SENSORY_PERC | 25 | -0.528602 | -1.353128 | 0.115486 | 1 |
| GOBP_REGULATION_OF_TRANSCRIPTION_BY_RNA_POLYMERASE_I | 33 | -0.507936 | -1.35288 | 0.105195 | 1 |
| GOBP_3_PHOSPHOADENOSINE_5_PHOSPHOSULFATE_METABOLIC_PROCESS | 22 | -0.546533 | -1.352692 | 0.105121 | 1 |
| GOBP_CELL_RECOGNITION | 210 | -0.405563 | -1.351635 | 0.024008 | 1 |
| GOBP_METAPHASE_ANAPHASE_TRANSITION_OF_CELL_CYCLE | 62 | -0.461998 | -1.350937 | 0.062428 | 1 |

| | | | | | |
|---|---|---|---|---|---|
| GOBP_METANEPHRIC_NEPHRON_MORPHOGENESIS | 25 | 0.465894 | 1.420804 | 0.048387 | 0.66526 |
| GOBP_SENSORY_PERCEPTION_OF_PAIN | 91 | 0.362942 | 1.420585 | 0.023622 | 0.661011 |
| GOBP_CELL_COMMUNICATION_BY_ELECTRICAL_COUPLING | 31 | 0.461872 | 1.420458 | 0.068182 | 0.656711 |
| GOBP_POSITIVE_REGULATION_OF_CARDIOCYTE_DIFFERENTIATION | 17 | 0.518887 | 1.419629 | 0.103321 | 0.654782 |
| GOBP_REGULATION_OF_SYNAPTIC_VESICLE_ENDOCYTOSIS | 15 | 0.530003 | 1.418835 | 0.069536 | 0.652728 |
| GOBP_NEGATIVE_REGULATION_OF_REPRODUCTIVE_PROCESS | 50 | 0.405262 | 1.418385 | 0.03012 | 0.649857 |
| GOBP_REGULATION_OF_EXTRACELLULAR_MATRIX_ASSEMBLY | 15 | 0.532227 | 1.416777 | 0.094276 | 0.651196 |
| GOBP_POSITIVE_REGULATION_OF_MORPHOGENESIS_OF_AN_EPITHELIU | 33 | 0.443431 | 1.416516 | 0.04 | 0.647468 |
| GOBP_EPIDERMIS_DEVELOPMENT | 337 | 0.303458 | 1.411898 | 0 | 0.659243 |
| GOBP_POSITIVE_REGULATION_OF_FILOPODIUM_ASSEMBLY | 30 | 0.444673 | 1.410751 | 0.04878 | 0.6586 |
| GOBP_ENDODERM_FORMATION | 53 | 0.392372 | 1.409328 | 0.035714 | 0.65951 |
| GOBP_CELLULAR_RESPONSE_TO_LEPTIN_STIMULUS | 18 | 0.516786 | 1.409319 | 0.085106 | 0.655083 |
| GOBP_HEART_GROWTH | 86 | 0.359059 | 1.40563 | 0.02439 | 0.663368 |
| GOBP_RESPONSE_TO_VITAMIN_A | 17 | 0.510438 | 1.400001 | 0.101399 | 0.680435 |
| GOBP_REGULATION_OF_EXCRETION | 19 | 0.481979 | 1.399649 | 0.076642 | 0.677214 |
| GOBP_REGULATION_OF_MYOBLAST_FUSION | 17 | 0.491637 | 1.394227 | 0.094891 | 0.692875 |
| GOBP_NEGATIVE_REGULATION_OF_PEPTIDASE_ACTIVITY | 222 | 0.309959 | 1.394039 | 0 | 0.688868 |
| GOBP_CORNIFICATION | 92 | 0.350882 | 1.390674 | 0.025 | 0.695968 |
| GOBP_AMINOGLYCAN_BIOSYNTHETIC_PROCESS | 113 | 0.346067 | 1.389785 | 0.037037 | 0.694701 |
| GOBP_CELL_CELL_RECOGNITION | 73 | 0.364474 | 1.389125 | 0.03937 | 0.692831 |
| GOBP_NEGATIVE_REGULATION_OF_SIGNAL_TRANSDUCTION_IN_ABSEN | 31 | 0.430096 | 1.387888 | 0.073059 | 0.692902 |
| GOBP_COPPER_ION_HOMEOSTASIS | 16 | 0.496171 | 1.387849 | 0.108209 | 0.6886 |
| GOBP_POSITIVE_REGULATION_OF_NEURON_APOPTOTIC_PROCESS | 52 | 0.393896 | 1.387265 | 0.028736 | 0.686284 |
| GOBP_PROSTATE_GLAND_MORPHOGENESIS | 27 | 0.462468 | 1.386562 | 0.084444 | 0.684594 |
| GOBP_EATING_BEHAVIOR | 24 | 0.456567 | 1.3863 | 0.075314 | 0.681354 |
| GOBP_CLATHRIN_COAT_ASSEMBLY | 18 | 0.481488 | 1.385097 | 0.096154 | 0.681366 |
| GOBP_NEGATIVE_REGULATION_OF_VIRAL_LIFE_CYCLE | 21 | 0.481437 | 1.38403 | 0.084677 | 0.681453 |
| GOBP_CELLULAR_RESPONSE_TO_GLUCAGON_STIMULUS | 23 | 0.474517 | 1.383116 | 0.093023 | 0.68055 |
| GOBP_NITRIC_OXIDE_MEDIATED_SIGNAL_TRANSDUCTION | 24 | 0.456676 | 1.381503 | 0.085714 | 0.682948 |
| GOBP_CELLULAR_RESPONSE_TO_AMYLOID_BETA | 37 | 0.409767 | 1.381448 | 0.072917 | 0.678941 |
| GOBP_RESPONSE_TO_METAL_ION | 333 | 0.300913 | 1.380796 | 0 | 0.677662 |
| GOBP_FOAM_CELL_DIFFERENTIATION | 37 | 0.432113 | 1.380552 | 0.075758 | 0.674381 |
| GOBP_CORTICOSTEROID_HORMONE_SECRETION | 15 | 0.506383 | 1.380042 | 0.103226 | 0.672118 |
| GOBP_CARDIOCYTE_DIFFERENTIATION | 124 | 0.338744 | 1.379904 | 0.041667 | 0.668508 |
| GOBP_CELL_CELL_ADHESION_VIA_PLASMA_MEMBRANE_ADHESION_MC | 266 | 0.295531 | 1.378614 | 0 | 0.66942 |
| GOBP_RENAL_SYSTEM_PROCESS | 112 | 0.336767 | 1.377939 | 0.020408 | 0.66787 |
| GOBP_SUPEROXIDE_METABOLIC_PROCESS | 64 | 0.365992 | 1.377837 | 0.044025 | 0.664258 |
| GOBP_REGULATION_OF_CARDIAC_MUSCLE_CELL_MEMBRANE_REPOLA | 24 | 0.455053 | 1.376596 | 0.091667 | 0.66456 |
| GOBP_NEGATIVE_REGULATION_OF_EXTRINSIC_APOPTOTIC_SIGNALING_ | 100 | 0.349488 | 1.372701 | 0.009259 | 0.674714 |
| GOBP_SYNAPTIC_TRANSMISSION_GABAERGIC | 43 | 0.39447 | 1.371692 | 0.055556 | 0.674243 |
| GOBP_POSITIVE_REGULATION_OF_NEUROTRANSMITTER_SECRETION | 17 | 0.495985 | 1.367402 | 0.089069 | 0.685968 |
| GOBP_METANEPHRIC_RENAL_VESICLE_MORPHOGENESIS | 15 | 0.486089 | 1.365396 | 0.100694 | 0.68939 |
| GOBP_DORSAL_VENTRAL_PATTERN_FORMATION | 69 | 0.372745 | 1.364754 | 0.037313 | 0.687686 |
| GOBP_CELLULAR_RESPONSE_TO_VITAMIN | 27 | 0.425745 | 1.361251 | 0.097561 | 0.696823 |
| GOBP_SPINAL_CORD_PATTERNING | 16 | 0.500002 | 1.360866 | 0.136667 | 0.694387 |
| GOBP_AORTIC_VALVE_MORPHOGENESIS | 31 | 0.422266 | 1.359598 | 0.087719 | 0.695265 |
| GOBP_FEMALE_GENITALIA_DEVELOPMENT | 16 | 0.492985 | 1.35943 | 0.129771 | 0.691911 |
| GOBP_RESPONSE_TO_GLUCAGON | 34 | 0.419621 | 1.358414 | 0.070352 | 0.691976 |
| GOBP_SEMI_LUNAR_VALVE_DEVELOPMENT | 41 | 0.395571 | 1.358069 | 0.066986 | 0.689331 |
| GOBP_CEREBELLAR_CORTEX_MORPHOGENESIS | 31 | 0.424939 | 1.356871 | 0.082677 | 0.689831 |
| GOBP_POSITIVE_REGULATION_OF_POTASSIUM_ION_TRANSMEMBRANE_ | 24 | 0.460005 | 1.352786 | 0.11157 | 0.700627 |
| GOBP_CARDIAC_MUSCLE_CELL_DIFFERENTIATION | 99 | 0.342301 | 1.351874 | 0.018519 | 0.700107 |
| GOBP_RESPONSE_TO_LEPTIN | 23 | 0.459901 | 1.350341 | 0.111588 | 0.701653 |
| GOBP_COLUMNAR_CUBOIDAL_EPITHELIAL_CELL_DEVELOPMENT | 38 | 0.401233 | 1.348419 | 0.061674 | 0.704878 |
| GOBP_CHONDROCYTE_DEVELOPMENT | 28 | 0.442498 | 1.346463 | 0.119617 | 0.708411 |
| GOBP_COBALAMIN_METABOLIC_PROCESS | 19 | 0.462505 | 1.3462 | 0.096491 | 0.705695 |
| GOBP_MATERNAL_PROCESS_INVOLVED_IN_FEMALE_PREGNANCY | 54 | 0.371772 | 1.344105 | 0.057471 | 0.709738 |
| GOBP_KERATINOCYTE_DIFFERENTIATION | 182 | 0.306105 | 1.343996 | 0 | 0.706537 |
| GOBP_PIGMENT_GRANULE_LOCALIZATION | 26 | 0.430364 | 1.343217 | 0.101887 | 0.705873 |
| GOBP_DERMATAN_SULFATE_PROTEOGLYCAN_METABOLIC_PROCESS | 16 | 0.488116 | 1.342633 | 0.132143 | 0.704455 |
| GOBP_NEUROPEPTIDE_SIGNALING_PATHWAY | 83 | 0.345212 | 1.340099 | 0.056 | 0.709766 |
| GOBP_ACTIN_FILAMENT_BASED_MOVEMENT | 146 | 0.31274 | 1.335641 | 0 | 0.722373 |
| GOBP_SKELETAL_MUSCLE_TISSUE_REGENERATION | 35 | 0.404875 | 1.333019 | 0.089623 | 0.728463 |
| GOBP_MULTICELLULAR_ORGANISMAL_SIGNALING | 186 | 0.315453 | 1.329647 | 0 | 0.737746 |
| GOBP_SPECIFICATION_OF_SYMMETRY | 116 | 0.327035 | 1.329379 | 0 | 0.735028 |
| GOBP_METANEPHRIC_NEPHRON_DEVELOPMENT | 38 | 0.392333 | 1.32909 | 0.095023 | 0.73248 |
| GOBP_NEGATIVE_REGULATION_OF_VIRAL_ENTRY_INTO_HOST_CELL | 18 | 0.47278 | 1.326737 | 0.15 | 0.737764 |
| GOBP_SENSORY_PERCEPTION_OF_CHEMICAL_STIMULUS | 104 | 0.33172 | 1.32666 | 0.023622 | 0.734371 |
| GOBP_POSITIVE_REGULATION_OF_T_CELL_MEDIATED_CYTOTOXICITY | 29 | 0.424859 | 1.322907 | 0.111111 | 0.744436 |
| GOBP_INTESTINAL_EPITHELIAL_CELL_DIFFERENTIATION | 19 | 0.465328 | 1.322344 | 0.148289 | 0.742836 |
| GOBP_GLANDULAR_EPITHELIAL_CELL_DEVELOPMENT | 25 | 0.446626 | 1.321024 | 0.127193 | 0.744087 |
| GOBP_PEPTIDYL_PROLINE_HYDROXYLATION | 15 | 0.498954 | 1.317671 | 0.132867 | 0.753056 |
| GOBP_HEART_PROCESS | 267 | 0.284433 | 1.311487 | 0 | 0.77334 |
| GOBP_POSITIVE_REGULATION_OF_EXOCYTOSIS | 85 | 0.338674 | 1.310672 | 0.008772 | 0.772696 |
| GOBP_NEGATIVE_REGULATION_OF_ANOIKIS | 17 | 0.462992 | 1.308544 | 0.140741 | 0.777847 |
| GOBP_NEGATIVE_REGULATION_OF_SECRETION | 140 | 0.313055 | 1.308339 | 0.014493 | 0.774954 |
| GOBP_REGULATION_OF_OXIDATIVE_STRESS_INDUCED_CELL_DEATH | 57 | 0.368381 | 1.308237 | 0.095541 | 0.771754 |
| GOBP_CELL_DEATH_IN_RESPONSE_TO_OXIDATIVE_STRESS | 78 | 0.341694 | 1.307441 | 0.060345 | 0.771087 |
| GOBP_REGULATION_OF_DNA_METHYLATION | 20 | 0.468545 | 1.303315 | 0.115538 | 0.784745 |
| GOBP_COCHLEA_DEVELOPMENT | 42 | 0.377027 | 1.302238 | 0.097297 | 0.785384 |
| GOBP_PATHWAY_RESTRICTED_SMAD_PROTEIN_PHOSPHORYLATION | 60 | 0.349686 | 1.300369 | 0.067164 | 0.789117 |
| GOBP_NEGATIVE_REGULATION_OF_BMP_SIGNALING_PATHWAY | 47 | 0.373788 | 1.299784 | 0.121053 | 0.787874 |
| GOBP_CEREBELLAR_PURKINJE_CELL_LAYER_MORPHOGENESIS | 15 | 0.486492 | 1.299055 | 0.132492 | 0.787101 |
| GOBP_CELLULAR_RESPONSE_TO_INORGANIC_SUBSTANCE | 204 | 0.299121 | 1.298088 | 0 | 0.787177 |
| GOBP_HINDBRAIN_MORPHOGENESIS | 40 | 0.38742 | 1.295233 | 0.102564 | 0.79501 |
| GOBP_EPIDERMAL_CELL_DIFFERENTIATION | 236 | 0.288627 | 1.294327 | 0 | 0.795358 |
| GOBP_PIGMENT_GRANULE_ORGANIZATION | 28 | 0.415699 | 1.292684 | 0.11811 | 0.797922 |
| GOBP_SODIUM_ION_HOMEOSTASIS | 48 | 0.369314 | 1.29102 | 0.071429 | 0.801053 |
| GOBP_CELLULAR_PIGMENTATION | 52 | 0.357048 | 1.29101 | 0.075269 | 0.7975 |
| GOBP_HEPARAN_SULFATE_PROTEOGLYCAN_BIOSYNTHETIC_PROCESS | 25 | 0.418205 | 1.289789 | 0.145522 | 0.798758 |
| GOBP_EMBRYONIC_AXIS_SPECIFICATION | 32 | 0.386703 | 1.286429 | 0.158879 | 0.808253 |
| GOBP_CELL_MIGRATION_INVOLVED_IN_HEART_DEVELOPMENT | 17 | 0.462949 | 1.28529 | 0.152727 | 0.809337 |
| GOBP_ADULT_BEHAVIOR | 110 | 0.321992 | 1.284113 | 0.028846 | 0.810389 |
| GOBP_REGULATION_OF_MEMBRANE_REPOLARIZATION | 31 | 0.401104 | 1.28152 | 0.121339 | 0.817331 |
| GOBP_REGULATION_OF_ACTIVIN_RECEPTOR_SIGNALING_PATHWAY | 23 | 0.425592 | 1.280652 | 0.130435 | 0.817332 |
| GOBP_SPERM_EGG_RECOGNITION | 54 | 0.355866 | 1.278392 | 0.116564 | 0.823004 |
| GOBP_CYCLIC_NUCLEOTIDE_BIOSYNTHETIC_PROCESS | 20 | 0.443734 | 1.277168 | 0.151163 | 0.82444 |
| GOBP_HOMOPHILIC_CELL_ADHESION_VIA_PLASMA_MEMBRANE_ADHES | 164 | 0.299081 | 1.276661 | 0.018868 | 0.822874 |
| GOBP_ESTABLISHMENT_OF_PIGMENT_GRANULE_LOCALIZATION | 24 | 0.428024 | 1.276169 | 0.142857 | 0.821126 |
| GOBP_REGULATION_OF_CELLULAR_RESPONSE_TO_GROWTH_FACTOR_ | 264 | 0.286081 | 1.275726 | 0 | 0.819535 |
| GOBP_REGULATION_OF_CARDIAC_MUSCLE_CELL_PROLIFERATION | 39 | 0.380524 | 1.275446 | 0.134021 | 0.817245 |
| GOBP_NEGATIVE_REGULATION_OF_PROTEOLYSIS | 306 | 0.280887 | 1.274733 | 0 | 0.816817 |
| GOBP_MYOBLAST_FUSION | 39 | 0.376218 | 1.274664 | 0.121053 | 0.813654 |
| GOBP_SIGNAL_TRANSDUCTION_INVOLVED_IN_REGULATION_OF_GENE_ | 15 | 0.458832 | 1.274564 | 0.151408 | 0.810702 |
| GOBP_ACTIVATION_OF_MAPKK_ACTIVITY | 50 | 0.365497 | 1.273889 | 0.079096 | 0.809874 |
| GOBP_NEGATIVE_REGULATION_OF_EXOCYTOSIS | 30 | 0.396669 | 1.273143 | 0.141026 | 0.809653 |
| GOBP_HYDROGEN_PEROXIDE_CATABOLIC_PROCESS | 26 | 0.417041 | 1.271945 | 0.129464 | 0.811142 |
| GOBP_EXTERNAL_ENCAPSULATING_STRUCTURE_ORGANIZATION | 383 | 0.285873 | 1.271735 | 0 | 0.808675 |
| GOBP_REGULATION_OF_VENTRICULAR_CARDIAC_MUSCLE_CELL_MEMI | 20 | 0.433754 | 1.27092 | 0.149194 | 0.808599 |
| GOBP_CATECHOL_CONTAINING_COMPOUND_BIOSYNTHETIC_PROCESS | 19 | 0.447473 | 1.267393 | 0.159533 | 0.819426 |
| GOBP_REGULATION_OF_SYNCYTIUM_FORMATION_BY_PLASMA_MEMBR | 25 | 0.427247 | 1.266333 | 0.163265 | 0.820578 |
| GOBP_SIGNAL_TRANSDUCTION_IN_ABSENCE_OF_LIGAND | 65 | 0.337413 | 1.26592 | 0.091603 | 0.818936 |
| GOBP_REGULATION_OF_SUPEROXIDE_ANION_GENERATION | 20 | 0.447625 | 1.26454 | 0.165414 | 0.821081 |
| GOBP_AUDITORY_RECEPTOR_CELL_DEVELOPMENT | 17 | 0.452661 | 1.262584 | 0.159259 | 0.825801 |
| GOBP_HEART_VALVE_DEVELOPMENT | 61 | 0.349139 | 1.261141 | 0.096591 | 0.828259 |
| GOBP_SKIN_DEVELOPMENT | 294 | 0.275967 | 1.261025 | 0 | 0.825433 |
| GOBP_SPINAL_CORD_DEVELOPMENT | 85 | 0.317359 | 1.260399 | 0.080645 | 0.824768 |
| GOBP_POSITIVE_REGULATION_OF_RECEPTOR_SIGNALING_PATHWAY_V | 44 | 0.36106 | 1.260001 | 0.118557 | 0.823253 |
| GOBP_CALCINEURIN_MEDIATED_SIGNALING | 44 | 0.363309 | 1.257929 | 0.103896 | 0.827775 |
| GOBP_POSITIVE_REGULATION_OF_TYPE_2_IMMUNE_RESPONSE | 16 | 0.471827 | 1.257388 | 0.180328 | 0.826749 |
| GOBP_POSITIVE_REGULATION_OF_PATHWAY_RESTRICTED_SMAD_PROT | 44 | 0.361164 | 1.256254 | 0.105 | 0.828198 |
| GOBP_NEURON_MIGRATION | 134 | 0.300782 | 1.25542 | 0.023529 | 0.828405 |
| GOBP_ESTABLISHMENT_OF_MITOTIC_SPINDLE_ORIENTATION | 28 | 0.41317 | 1.255243 | 0.17 | 0.825781 |
| GOBP_DIGESTIVE_SYSTEM_DEVELOPMENT | 121 | 0.305923 | 1.253907 | 0.043478 | 0.828406 |
| GOBP_LIPID_STORAGE | 81 | 0.322273 | 1.253833 | 0.056075 | 0.825451 |
| GOBP_RESPONSE_TO_AXON_INJURY | 72 | 0.329694 | 1.251991 | 0.067669 | 0.829678 |
| GOBP_POSITIVE_REGULATION_OF_OSSIFICATION | 47 | 0.36153 | 1.251499 | 0.093407 | 0.828548 |
| GOBP_CELL_DIFFERENTIATION_IN_SPINAL_CORD | 39 | 0.382896 | 1.251423 | 0.117647 | 0.825751 |
| GOBP_REGULATION_OF_APOPTOTIC_SIGNALING_PATHWAY | 332 | 0.279132 | 1.25139 | 0 | 0.822786 |
| GOBP_MATERNAL_PLACENTA_DEVELOPMENT | 27 | 0.395334 | 1.250869 | 0.156118 | 0.821736 |
| GOBP_STRIATED_MUSCLE_CELL_PROLIFERATION | 62 | 0.341059 | 1.249879 | 0.106145 | 0.822699 |
| GOBP_REGULATION_OF_CARDIOCYTE_DIFFERENTIATION | 30 | 0.405192 | 1.24977 | 0.149573 | 0.819984 |
| GOBP_NEGATIVE_REGULATION_OF_CARTILAGE_DEVELOPMENT | 25 | 0.419848 | 1.247844 | 0.164 | 0.824513 |
| GOBP_KILLING_OF_CELLS_IN_OTHER_ORGANISM_INVOLVED_IN_SYMBIO | 17 | 0.456612 | 1.247104 | 0.19084 | 0.824157 |
| GOBP_MUSCLE_CELL_DEVELOPMENT | 152 | 0.296646 | 1.246382 | 0.04878 | 0.823995 |
| GOBP_REGULATION_OF_HEART_CONTRACTION | 225 | 0.283415 | 1.24598 | 0.027778 | 0.822643 |
| GOBP_RESPONSE_TO_INTERFERON_ALPHA | 21 | 0.434778 | 1.245978 | 0.190661 | 0.819607 |
| GOBP_NEGATIVE_REGULATION_OF_STRIATED_MUSCLE_CELL_APOPTOT | 23 | 0.419703 | 1.24503 | 0.161765 | 0.820223 |

| | | | | | |
|---|---|---|---|---|---|
| GOBP_REGULATION_OF_BEHAVIOR | 56 | -0.462796 | -1.348419 | 0.065614 | 1 |
| GOBP_LONG_CHAIN_FATTY_ACID_METABOLIC_PROCESS | 118 | -0.427047 | -1.34816 | 0.032438 | 1 |
| GOBP_REGULATION_OF_GENE_SILENCING | 128 | -0.425458 | -1.347303 | 0.045752 | 1 |
| GOBP_NEGATIVE_REGULATION_OF_MRNA_SPLICING_VIA_SPLICEOSOME | 20 | -0.550435 | -1.344097 | 0.109018 | 1 |
| GOBP_GLYCEROLIPID_CATABOLIC_PROCESS | 65 | -0.455264 | -1.343634 | 0.07034 | 1 |
| GOBP_ANDROGEN_METABOLIC_PROCESS | 30 | -0.514265 | -1.342456 | 0.104302 | 1 |
| GOBP_OOCYTE_DIFFERENTIATION | 46 | -0.482578 | -1.341457 | 0.08384 | 1 |
| GOBP_MITOCHONDRIAL_GENOME_MAINTENANCE | 21 | -0.54796 | -1.340879 | 0.106901 | 1 |
| GOBP_NEUTRAL_AMINO_ACID_TRANSPORT | 40 | -0.485934 | -1.339784 | 0.088639 | 1 |
| GOBP_RESPONSE_TO_STEROL | 31 | -0.502825 | -1.339475 | 0.090426 | 1 |
| GOBP_REGULATION_OF_LYASE_ACTIVITY | 44 | -0.487089 | -1.338969 | 0.084951 | 1 |
| GOBP_POSITIVE_REGULATION_OF_INTERFERON_GAMMA_PRODUCTION | 65 | -0.453347 | -1.335177 | 0.082734 | 1 |
| GOBP_CELLULAR_RESPONSE_TO_ATP | 16 | -0.571055 | -1.334907 | 0.113416 | 1 |
| GOBP_WATER_TRANSPORT | 20 | -0.549379 | -1.334755 | 0.121986 | 1 |
| GOBP_DNA_CATABOLIC_PROCESS_ENDONUCLEOLYTIC | 30 | -0.505592 | -1.33335 | 0.099338 | 1 |
| GOBP_HYPOTHALAMUS_DEVELOPMENT | 16 | -0.57712 | -1.333104 | 0.124157 | 1 |
| GOBP_ACYLGLYCEROL_HOMEOSTASIS | 29 | -0.509008 | -1.332614 | 0.104222 | 1 |
| GOBP_RESPONSE_TO_CORTICOSTERONE | 18 | -0.552757 | -1.331046 | 0.135593 | 1 |
| GOBP_MONOCYTE_CHEMOTAXIS | 61 | -0.452075 | -1.33054 | 0.077283 | 1 |
| GOBP_POSITIVE_REGULATION_OF_INTERLEUKIN_17_PRODUCTION | 17 | -0.572177 | -1.330148 | 0.104312 | 1 |
| GOBP_REGULATION_OF_MULTI_ORGANISM_PROCESS | 47 | -0.464818 | -1.328161 | 0.085439 | 1 |
| GOBP_POSITIVE_REGULATION_OF_INTERLEUKIN_4_PRODUCTION | 24 | -0.522254 | -1.328114 | 0.113456 | 1 |
| GOBP_LEUKOTRIENE_METABOLIC_PROCESS | 33 | -0.498685 | -1.328084 | 0.096734 | 1 |
| GOBP_PROSTANOID_METABOLIC_PROCESS | 44 | -0.472229 | -1.324 | 0.101591 | 1 |
| GOBP_TRANSPOSITION | 29 | -0.509988 | -1.323987 | 0.099075 | 1 |
| GOBP_CHROMATIN_REMODELING_AT_CENTROMERE | 43 | -0.476546 | -1.323139 | 0.104218 | 1 |
| GOBP_POSITIVE_REGULATION_OF_LIPID_CATABOLIC_PROCESS | 23 | -0.531038 | -1.320719 | 0.123506 | 1 |
| GOBP_NATURAL_KILLER_CELL_MEDIATED_IMMUNITY | 66 | -0.445589 | -1.318389 | 0.079861 | 1 |
| GOBP_NITRIC_OXIDE_SYNTHASE_BIOSYNTHETIC_PROCESS | 19 | -0.542184 | -1.318213 | 0.118046 | 1 |
| GOBP_MEIOTIC_CELL_CYCLE_PROCESS | 179 | -0.405004 | -1.31576 | 0.041314 | 1 |
| GOBP_NEURON_CELLULAR_HOMEOSTASIS | 33 | -0.493811 | -1.314588 | 0.110256 | 1 |
| GOBP_REGULATION_OF_GENE_EXPRESSION_EPIGENETIC | 181 | -0.40056 | -1.313542 | 0.04375 | 1 |
| GOBP_PITUITARY_GLAND_DEVELOPMENT | 37 | -0.485357 | -1.313542 | 0.123171 | 1 |
| GOBP_LINOLEIC_ACID_METABOLIC_PROCESS | 21 | -0.531793 | -1.312794 | 0.139276 | 1 |
| GOBP_REGULATION_OF_PRESYNAPTIC_CYTOSOLIC_CALCIUM_ION_CONCENT | 16 | -0.566659 | -1.312104 | 0.15508 | 1 |
| GOBP_MATURE_B_CELL_DIFFERENTIATION_INVOLVED_IN_IMMUNE_RESPONS | 24 | -0.522385 | -1.31177 | 0.134846 | 1 |
| GOBP_SULFATION | 21 | -0.535931 | -1.311707 | 0.127572 | 1 |
| GOBP_SEROTONIN_RECEPTOR_SIGNALING_PATHWAY | 23 | -0.528733 | -1.310204 | 0.124665 | 1 |
| GOBP_PROTEIN_CONTAINING_COMPLEX_REMODELING | 29 | -0.495701 | -1.309782 | 0.119326 | 1 |
| GOBP_MEIOTIC_CELL_CYCLE | 233 | -0.391226 | -1.309652 | 0.030644 | 1 |
| GOBP_LIMBIC_SYSTEM_DEVELOPMENT | 88 | -0.429824 | -1.308007 | 0.077268 | 1 |
| GOBP_REGULATION_OF_B_CELL_RECEPTOR_SIGNALING_PATHWAY | 25 | -0.510919 | -1.307502 | 0.138852 | 1 |
| GOBP_RESOLUTION_OF_MEIOTIC_RECOMBINATION_INTERMEDIATES | 17 | -0.552897 | -1.304787 | 0.158263 | 1 |
| GOBP_ALPHA_BETA_T_CELL_ACTIVATION | 143 | -0.403747 | -1.30458 | 0.059524 | 1 |
| GOBP_CENTROMERE_COMPLEX_ASSEMBLY | 52 | -0.458312 | -1.304501 | 0.093976 | 1 |
| GOBP_NEGATIVE_REGULATION_OF_MACROPHAGE_ACTIVATION | 18 | -0.555158 | -1.303433 | 0.144966 | 1 |
| GOBP_THYROID_HORMONE_METABOLIC_PROCESS | 25 | -0.514159 | -1.300867 | 0.128514 | 1 |
| GOBP_COPULATION | 17 | -0.551626 | -1.300799 | 0.148611 | 1 |
| GOBP_FOREBRAIN_NEURON_DEVELOPMENT | 24 | -0.51367 | -1.30021 | 0.135246 | 1 |
| GOBP_LYMPHOCYTE_COSTIMULATION | 59 | -0.445467 | -1.298603 | 0.105516 | 1 |
| GOBP_POSITIVE_REGULATION_OF_OSTEOCLAST_DIFFERENTIATION | 25 | -0.509632 | -1.297773 | 0.132353 | 1 |
| GOBP_CELL_MATURATION | 155 | -0.395971 | -1.296099 | 0.054025 | 1 |
| GOBP_CELL_FATE_COMMITMENT_INVOLVED_IN_FORMATION_OF_PRIMARY_G | 28 | -0.501935 | -1.292637 | 0.13245 | 1 |
| GOBP_TOLERANCE_INDUCTION | 28 | -0.507798 | -1.292403 | 0.132147 | 1 |
| GOBP_INTERLEUKIN_17_PRODUCTION | 34 | -0.48007 | -1.29215 | 0.139594 | 1 |
| GOBP_FEEDING_BEHAVIOR | 79 | -0.426061 | -1.28966 | 0.097032 | 1 |
| GOBP_NEGATIVE_REGULATION_OF_CELL_DIVISION | 15 | -0.558513 | -1.28796 | 0.157083 | 1 |
| GOBP_POSITIVE_REGULATION_OF_MULTI_ORGANISM_PROCESS | 18 | -0.548036 | -1.285297 | 0.153642 | 1 |
| GOBP_REGULATION_OF_T_CELL_CHEMOTAXIS | 15 | -0.560256 | -1.285005 | 0.154728 | 1 |
| GOBP_POSITIVE_REGULATION_OF_STEROID_METABOLIC_PROCESS | 24 | -0.507422 | -1.283641 | 0.15102 | 1 |
| GOBP_POSITIVE_REGULATION_OF_PHOSPHATASE_ACTIVITY | 33 | -0.485722 | -1.283037 | 0.130818 | 1 |
| GOBP_ISOPRENOID_BIOSYNTHETIC_PROCESS | 27 | -0.502823 | -1.282878 | 0.159642 | 1 |
| GOBP_PURINE_NUCLEOBASE_METABOLIC_PROCESS | 19 | -0.533313 | -1.282805 | 0.1622 | 1 |
| GOBP_ACETYL_COA_METABOLIC_PROCESS | 37 | -0.46537 | -1.282794 | 0.131026 | 1 |
| GOBP_REGULATION_OF_B_CELL_DIFFERENTIATION | 29 | -0.487637 | -1.281865 | 0.14653 | 1 |
| GOBP_NEGATIVE_REGULATION_OF_LIPID_CATABOLIC_PROCESS | 22 | -0.508323 | -1.281849 | 0.158035 | 1 |
| GOBP_POSITIVE_REGULATION_OF_MUSCLE_CONTRACTION | 36 | -0.475115 | -1.281669 | 0.127024 | 1 |
| GOBP_REGULATION_OF_GLIAL_CELL_MIGRATION | 18 | -0.531218 | -1.280776 | 0.153333 | 1 |
| GOBP_TRIGLYCERIDE_CATABOLIC_PROCESS | 31 | -0.490155 | -1.280369 | 0.147135 | 1 |
| GOBP_REGULATION_OF_LIPOPROTEIN_LIPASE_ACTIVITY | 20 | -0.534812 | -1.278468 | 0.169607 | 1 |
| GOBP_REGULATION_OF_EXTRACELLULAR_MATRIX_DISASSEMBLY | 16 | -0.553468 | -1.277909 | 0.173913 | 1 |
| GOBP_REGULATION_OF_GENE_SILENCING_BY_RNA | 111 | -0.406703 | -1.2763 | 0.089227 | 1 |
| GOBP_CELLULAR_COMPONENT_DISASSEMBLY_INVOLVED_IN_EXECUTION_PI | 34 | -0.471 | -1.273306 | 0.134021 | 1 |
| GOBP_CHROMATIN_ASSEMBLY_OR_DISASSEMBLY | 192 | -0.386023 | -1.27328 | 0.051095 | 1 |
| GOBP_T_CELL_SELECTION | 48 | -0.448371 | -1.272491 | 0.133333 | 1 |
| GOBP_POSITIVE_T_CELL_SELECTION | 36 | -0.470805 | -1.272021 | 0.148194 | 1 |
| GOBP_ALPHA_BETA_T_CELL_DIFFERENTIATION | 104 | -0.405249 | -1.2719 | 0.093468 | 1 |
| GOBP_PRIMARY_ALCOHOL_METABOLIC_PROCESS | 89 | -0.421218 | -1.270363 | 0.096956 | 1 |
| GOBP_POSITIVE_REGULATION_OF_ORGANIC_ACID_TRANSPORT | 33 | -0.470575 | -1.27025 | 0.168798 | 1 |
| GOBP_T_CELL_RECEPTOR_SIGNALING_PATHWAY | 200 | -0.380544 | -1.268988 | 0.062305 | 1 |
| GOBP_REGULATION_OF_B_CELL_APOPTOTIC_PROCESS | 19 | -0.527636 | -1.268861 | 0.154391 | 1 |
| GOBP_T_CELL_MIGRATION | 62 | -0.432263 | -1.268082 | 0.119952 | 1 |
| GOBP_CHONDROCYTE_PROLIFERATION | 19 | -0.535225 | -1.265944 | 0.167553 | 1 |
| GOBP_FRUCTOSE_METABOLIC_PROCESS | 15 | -0.554669 | -1.262729 | 0.194444 | 1 |
| GOBP_SULFUR_COMPOUND_CATABOLIC_PROCESS | 54 | -0.439237 | -1.260858 | 0.147268 | 1 |
| GOBP_NEGATIVE_REGULATION_OF_SIGNALING_RECEPTOR_ACTIVITY | 56 | -0.43945 | -1.259208 | 0.129608 | 1 |
| GOBP_LYMPHOCYTE_CHEMOTAXIS | 55 | -0.431958 | -1.256996 | 0.146028 | 1 |
| GOBP_DNA_REPLICATION_INDEPENDENT_NUCLEOSOME_ORGANIZATION | 50 | -0.436167 | -1.25647 | 0.146732 | 1 |
| GOBP_DNA_REPLICATION_CHECKPOINT | 17 | -0.536246 | -1.253944 | 0.18707 | 1 |
| GOBP_POSITIVE_REGULATION_OF_PHOSPHOPROTEIN_PHOSPHATASE_ACTIVI | 21 | -0.503167 | -1.253937 | 0.179487 | 1 |
| GOBP_NEUTRAL_LIPID_CATABOLIC_PROCESS | 39 | -0.46906 | -1.252896 | 0.151592 | 1 |
| GOBP_RESPONSE_TO_AMPHETAMINE | 29 | -0.4837 | -1.252892 | 0.181092 | 1 |
| GOBP_INTERLEUKIN_4_PRODUCTION | 31 | -0.469907 | -1.251408 | 0.183136 | 1 |
| GOBP_HORMONE_METABOLIC_PROCESS | 207 | -0.378791 | -1.250872 | 0.066597 | 1 |
| GOBP_MEMBRANE_RAFT_ORGANIZATION | 24 | -0.491196 | -1.24986 | 0.17655 | 1 |
| GOBP_HYPEROSMOTIC_RESPONSE | 24 | -0.502007 | -1.249361 | 0.177015 | 1 |
| GOBP_POSITIVE_REGULATION_OF_T_CELL_PROLIFERATION | 96 | -0.405 | -1.249043 | 0.120805 | 1 |
| GOBP_REGULATION_OF_CHROMOSOME_SEPARATION | 68 | -0.422481 | -1.248823 | 0.136891 | 1 |
| GOBP_NEUTRAL_LIPID_BIOSYNTHETIC_PROCESS | 45 | -0.4481 | -1.247905 | 0.163975 | 1 |
| GOBP_NEGATIVE_REGULATION_OF_TUMOR_NECROSIS_FACTOR_SUPERFAMI | 56 | -0.429349 | -1.247699 | 0.138756 | 1 |
| GOBP_NEGATIVE_REGULATION_OF_ANDROGEN_RECEPTOR_SIGNALING_PATI | 15 | -0.542338 | -1.24664 | 0.196051 | 1 |
| GOBP_MATURE_B_CELL_DIFFERENTIATION | 29 | -0.481106 | -1.24521 | 0.159517 | 1 |
| GOBP_CELLULAR_HORMONE_METABOLIC_PROCESS | 128 | -0.395853 | -1.245069 | 0.092511 | 1 |
| GOBP_NUCLEOSOME_ORGANIZATION | 154 | -0.386306 | -1.244558 | 0.089827 | 1 |
| GOBP_GLYCEROPHOSPHOLIPID_CATABOLIC_PROCESS | 31 | -0.481561 | -1.244272 | 0.160875 | 1 |
| GOBP_THYROID_HORMONE_GENERATION | 21 | -0.511277 | -1.244009 | 0.193931 | 1 |
| GOBP_DIGESTIVE_SYSTEM_PROCESS | 86 | -0.4053 | -1.243848 | 0.128176 | 1 |
| GOBP_RESPONSE_TO_CAFFEINE | 17 | -0.527536 | -1.243645 | 0.204167 | 1 |
| GOBP_O_GLYCAN_PROCESSING | 58 | -0.425207 | -1.24268 | 0.152147 | 1 |
| GOBP_PHOSPHATIDYLETHANOLAMINE_METABOLIC_PROCESS | 32 | -0.475646 | -1.242075 | 0.179487 | 1 |
| GOBP_DNA_METHYLATION | 67 | -0.428128 | -1.241442 | 0.154028 | 1 |
| GOBP_BIOLOGICAL_PROCESS_INVOLVED_IN_INTRASPECIES_INTERACTION_B | 49 | -0.436121 | -1.239653 | 0.169903 | 1 |
| GOBP_MEIOSIS_I_CELL_CYCLE_PROCESS | 119 | -0.38697 | -1.238856 | 0.10341 | 1 |
| GOBP_MITOTIC_CHROMOSOME_CONDENSATION | 15 | -0.537872 | -1.238623 | 0.205797 | 1 |
| GOBP_G_PROTEIN_COUPLED_RECEPTOR_SIGNALING_PATHWAY_COUPLED_T | 47 | -0.44044 | -1.237615 | 0.174334 | 1 |
| GOBP_THIOESTER_BIOSYNTHETIC_PROCESS | 52 | -0.429764 | -1.236708 | 0.139506 | 1 |
| GOBP_NOREPINEPHRINE_TRANSPORT | 22 | -0.499853 | -1.235714 | 0.212081 | 1 |
| GOBP_DNA_CONFORMATION_CHANGE | 316 | -0.367933 | -1.235002 | 0.046701 | 1 |
| GOBP_RETINA_HOMEOSTASIS | 78 | -0.414026 | -1.234279 | 0.130081 | 1 |
| GOBP_NEGATIVE_REGULATION_OF_MULTI_ORGANISM_PROCESS | 19 | -0.520992 | -1.233797 | 0.219178 | 1 |
| GOBP_STEROL_BIOSYNTHETIC_PROCESS | 71 | -0.418546 | -1.232573 | 0.168009 | 1 |
| GOBP_MEIOTIC_CHROMOSOME_SEPARATION | 23 | -0.493384 | -1.231933 | 0.197333 | 1 |
| GOBP_FOREBRAIN_DEVELOPMENT | 333 | -0.361535 | -1.230648 | 0.046653 | 1 |
| GOBP_PEPTIDYL_GLUTAMIC_ACID_MODIFICATION | 29 | -0.468347 | -1.230498 | 0.198693 | 1 |
| GOBP_NEUROTRANSMITTER_RECEPTOR_LOCALIZATION_TO_POSTSYNAPTIC_ | 16 | -0.518016 | -1.228288 | 0.218707 | 1 |
| GOBP_POSITIVE_REGULATION_OF_LEUKOCYTE_APOPTOTIC_PROCESS | 24 | -0.487654 | -1.224976 | 0.211688 | 1 |
| GOBP_CELL_FATE_SPECIFICATION | 77 | -0.402839 | -1.224518 | 0.162573 | 1 |
| GOBP_REGULATION_OF_MONOCYTE_CHEMOTAXIS | 26 | -0.47757 | -1.224501 | 0.214379 | 1 |
| GOBP_NUCLEOSIDE_BISPHOSPHATE_METABOLIC_PROCESS | 136 | -0.381926 | -1.223724 | 0.112432 | 1 |
| GOBP_FEAR_RESPONSE | 34 | -0.453706 | -1.223323 | 0.181818 | 1 |
| GOBP_DETERMINATION_OF_ADULT_LIFESPAN | 17 | -0.519809 | -1.223296 | 0.226776 | 1 |
| GOBP_NLS_BEARING_PROTEIN_IMPORT_INTO_NUCLEUS | 18 | -0.503942 | -1.222979 | 0.203552 | 1 |
| GOBP_CELL_PROLIFERATION_IN_FOREBRAIN | 24 | -0.48836 | -1.221975 | 0.198953 | 1 |
| GOBP_CHROMOSOME_SEPARATION | 91 | -0.397395 | -1.220658 | 0.134312 | 1 |
| GOBP_ANATOMICAL_STRUCTURE_MATURATION | 205 | -0.369001 | -1.220116 | 0.102833 | 1 |
| GOBP_REGULATION_OF_MEIOTIC_CELL_CYCLE | 43 | -0.437113 | -1.219264 | 0.189937 | 1 |
| GOBP_NEUROMUSCULAR_PROCESS_CONTROLLING_BALANCE | 37 | -0.451851 | -1.219252 | 0.209677 | 1 |
| GOBP_FLUID_TRANSPORT | 30 | -0.46164 | -1.21917 | 0.195767 | 1 |

| | | | | | |
|---|---|---|---|---|---|
| GOBP_CARDIAC_SEPTUM_MORPHOGENESIS | 62 | 0.330435 | 1.244988 | 0.084507 | 0.817434 |
| GOBP_POSITIVE_REGULATION_OF_MRNA_SPLICING_VIA_SPLICEOSOME | 22 | 0.414205 | 1.244762 | 0.159259 | 0.815248 |
| GOBP_PEPTIDYL_L_CYSTEINE_S_PALMITOYLATION | 24 | 0.410818 | 1.243461 | 0.171429 | 0.817284 |
| GOBP_NEGATIVE_REGULATION_OF_COAGULATION | 47 | 0.359787 | 1.24283 | 0.115183 | 0.816876 |
| GOBP_REGULATION_OF_REGULATED_SECRETORY_PATHWAY | 136 | 0.302553 | 1.242202 | 0.079545 | 0.816315 |
| GOBP_GAS_TRANSPORT | 16 | 0.4556 | 1.241693 | 0.218391 | 0.815368 |
| GOBP_REGULATION_OF_MORPHOGENESIS_OF_AN_EPITHELIUM | 60 | 0.33902 | 1.239787 | 0.093567 | 0.819541 |
| GOBP_BASEMENT_MEMBRANE_ORGANIZATION | 30 | 0.393334 | 1.239005 | 0.152542 | 0.819748 |
| GOBP_OVULATION_CYCLE_PROCESS | 40 | 0.367899 | 1.23897 | 0.131868 | 0.816931 |
| GOBP_VESICLE_MEDIATED_TRANSPORT_IN_SYNAPSE | 188 | 0.285147 | 1.236841 | 0.019608 | 0.822638 |
| GOBP_GRANULOCYTE_MACROPHAGE_COLONY_STIMULATING_FACTOR | 15 | 0.454072 | 1.23647 | 0.202899 | 0.821333 |
| GOBP_NEGATIVE_REGULATION_OF_SIGNAL_TRANSDUCTION_BY_P53_C | 30 | 0.393448 | 1.236009 | 0.170732 | 0.820267 |
| GOBP_NOSE_DEVELOPMENT | 15 | 0.466186 | 1.235864 | 0.203922 | 0.817871 |
| GOBP_NEGATIVE_REGULATION_OF_CELLULAR_RESPONSE_TO_GROWT | 95 | 0.312151 | 1.23493 | 0.116505 | 0.818334 |
| GOBP_VENTRICULAR_SEPTUM_DEVELOPMENT | 63 | 0.337825 | 1.234633 | 0.090909 | 0.816666 |
| GOBP_EXTRACELLULAR_TRANSPORT | 33 | 0.388321 | 1.233447 | 0.146667 | 0.818525 |
| GOBP_EPIDERMIS_MORPHOGENESIS | 27 | 0.390016 | 1.233182 | 0.188976 | 0.81675 |
| GOBP_BASIC_AMINO_ACID_TRANSPORT | 18 | 0.432214 | 1.232893 | 0.175373 | 0.815014 |
| GOBP_CHAPERONE_MEDIATED_PROTEIN_FOLDING | 62 | 0.328801 | 1.23236 | 0.104575 | 0.814491 |
| GOBP_NEURON_PROJECTION_ARBORIZATION | 28 | 0.399304 | 1.231633 | 0.161826 | 0.814773 |
| GOBP_REGULATION_OF_ANOIKIS | 24 | 0.398884 | 1.231541 | 0.190909 | 0.812381 |
| GOBP_NEGATIVE_REGULATION_OF_STEROID_METABOLIC_PROCESS | 23 | 0.42038 | 1.230514 | 0.178182 | 0.813339 |
| GOBP_REGULATION_OF_LEUKOCYTE_DEGRANULATION | 43 | 0.356974 | 1.22986 | 0.121387 | 0.813277 |
| GOBP_REGULATION_OF_HEART_RATE_BY_CARDIAC_CONDUCTION | 39 | 0.367599 | 1.228797 | 0.13242 | 0.814469 |
| GOBP_NEGATIVE_REGULATION_OF_VIRAL_TRANSCRIPTION | 15 | 0.452874 | 1.22773 | 0.2 | 0.815858 |
| GOBP_MYELOID_LEUKOCYTE_CYTOKINE_PRODUCTION | 26 | 0.399538 | 1.227127 | 0.168033 | 0.815495 |
| GOBP_VENTRAL_SPINAL_CORD_DEVELOPMENT | 36 | 0.372031 | 1.226549 | 0.159624 | 0.814925 |
| GOBP_MESONEPHRIC_TUBULE_MORPHOGENESIS | 59 | 0.33766 | 1.22456 | 0.134146 | 0.819899 |
| GOBP_UTERUS_DEVELOPMENT | 18 | 0.440882 | 1.224077 | 0.206522 | 0.819097 |
| GOBP_ESTABLISHMENT_OF_SPINDLE_ORIENTATION | 33 | 0.3696 | 1.223648 | 0.157068 | 0.818124 |
| GOBP_PHYSIOLOGICAL_CARDIAC_MUSCLE_HYPERTROPHY | 26 | 0.384502 | 1.22338 | 0.168103 | 0.816485 |
| GOBP_REGULATION_OF_EXTRINSIC_APOPTOTIC_SIGNALING_PATHWAY | 149 | 0.296812 | 1.223322 | 0.065574 | 0.814081 |
| GOBP_INTERMEDIATE_FILAMENT_ORGANIZATION | 22 | 0.415429 | 1.222532 | 0.177122 | 0.814838 |
| GOBP_POSITIVE_REGULATION_OF_STEM_CELL_DIFFERENTIATION | 18 | 0.437813 | 1.221157 | 0.204301 | 0.817283 |
| GOBP_CARDIAC_CONDUCTION | 139 | 0.290908 | 1.219024 | 0.095238 | 0.822872 |
| GOBP_EMBRYO_IMPLANTATION | 45 | 0.346944 | 1.218666 | 0.129032 | 0.821721 |
| GOBP_MITOPHAGY | 24 | 0.41151 | 1.218399 | 0.215859 | 0.820247 |
| GOBP_NEURONAL_ACTION_POTENTIAL | 30 | 0.383448 | 1.215741 | 0.184906 | 0.8283 |
| GOBP_RESPONSE_TO_NITRIC_OXIDE | 19 | 0.434102 | 1.212173 | 0.205036 | 0.839435 |
| GOBP_LONG_TERM_MEMORY | 32 | 0.371818 | 1.212163 | 0.178423 | 0.836779 |
| GOBP_CARDIAC_MUSCLE_CELL_PROLIFERATION | 48 | 0.339948 | 1.209795 | 0.171598 | 0.843856 |
| GOBP_ACTIVATION_OF_PROTEIN_KINASE_ACTIVITY | 321 | 0.266825 | 1.209056 | 0 | 0.844246 |
| GOBP_TRANSMEMBRANE_RECEPTOR_PROTEIN_SERINE_THREONINE_K | 328 | 0.267983 | 1.208869 | 0 | 0.842332 |
| GOBP_POSTSYNAPTIC_SPECIALIZATION_ASSEMBLY | 20 | 0.424764 | 1.207152 | 0.231884 | 0.846374 |
| GOBP_REGULATION_OF_MACROPHAGE_DIFFERENTIATION | 20 | 0.42739 | 1.206299 | 0.246154 | 0.847093 |
| GOBP_REGULATION_OF_RESPONSE_TO_OXIDATIVE_STRESS | 80 | 0.312543 | 1.205467 | 0.111111 | 0.847649 |
| GOBP_REGULATION_OF_VASCULAR_ENDOTHELIAL_GROWTH_FACTOR_ | 29 | 0.396137 | 1.203881 | 0.209302 | 0.851298 |
| GOBP_POSITIVE_REGULATION_OF_OLIGODENDROCYTE_DIFFERENTIATI | 20 | 0.419634 | 1.203666 | 0.231061 | 0.849285 |
| GOBP_REGULATION_OF_MORPHOGENESIS_OF_A_BRANCHING_STRUCT | 48 | 0.340756 | 1.202861 | 0.144654 | 0.850045 |
| GOBP_CELL_KILLING | 15 | 0.436927 | 1.202763 | 0.220339 | 0.84779 |
| GOBP_MACROPHAGE_CYTOKINE_PRODUCTION | 39 | 0.354012 | 1.202208 | 0.169082 | 0.847366 |
| GOBP_PHENOL_CONTAINING_COMPOUND_BIOSYNTHETIC_PROCESS | 22 | 0.408765 | 1.201917 | 0.187773 | 0.845959 |
| GOBP_NEGATIVE_REGULATION_OF_CELL_CYCLE_ARREST | 54 | 0.338382 | 1.201406 | 0.153846 | 0.845432 |
| GOBP_CELL_COMMUNICATION_INVOLVED_IN_CARDIAC_CONDUCTION | 28 | 0.387133 | 1.201224 | 0.186957 | 0.843536 |
| GOBP_REGULATION_OF_LEUKOCYTE_ADHESION_TO_VASCULAR_ENDO | 152 | 0.278424 | 1.20095 | 0.046154 | 0.841977 |
| GOBP_POSITIVE_REGULATION_OF_DEVELOPMENTAL_GROWTH | 37 | 0.360177 | 1.199343 | 0.190476 | 0.845599 |
| GOBP_LUNG_ALVEOLUS_DEVELOPMENT | 41 | 0.359104 | 1.198952 | 0.179487 | 0.844544 |
| GOBP_REGULATION_OF_CALCIUM_ION_DEPENDENT_EXOCYTOSIS | 127 | 0.290766 | 1.198242 | 0.121951 | 0.844652 |
| GOBP_CARDIAC_MUSCLE_CONTRACTION | 26 | 0.394275 | 1.197205 | 0.203463 | 0.846369 |
| GOBP_POSITIVE_REGULATION_OF_P38MAPK_CASCADE | 43 | 0.338252 | 1.196996 | 0.168367 | 0.844635 |
| GOBP_CELLULAR_CARBOHYDRATE_CATABOLIC_PROCESS | 124 | 0.293274 | 1.196906 | 0.1 | 0.842433 |
| GOBP_DETOXIFICATION | 64 | 0.318541 | 1.196112 | 0.116959 | 0.842946 |
| GOBP_EMBRYONIC_HEART_TUBE_MORPHOGENESIS | 126 | 0.288328 | 1.194111 | 0.074627 | 0.848477 |
| GOBP_ACTION_POTENTIAL | 107 | 0.295817 | 1.193466 | 0.122222 | 0.848404 |
| GOBP_INTRACELLULAR_STEROID_HORMONE_RECEPTOR_SIGNALING_F | 33 | 0.364504 | 1.193407 | 0.188406 | 0.846185 |
| GOBP_ESTABLISHMENT_OF_MITOTIC_SPINDLE_LOCALIZATION | 57 | 0.328368 | 1.192282 | 0.152439 | 0.848179 |
| GOBP_MORPHOGENESIS_OF_AN_EPITHELIAL_SHEET | 90 | 0.303663 | 1.192056 | 0.125 | 0.846512 |
| GOBP_MESONEPHROS_DEVELOPMENT | 32 | 0.373118 | 1.191964 | 0.193694 | 0.844406 |
| GOBP_POSITIVE_REGULATION_OF_AMINE_TRANSPORT | 19 | 0.411781 | 1.189465 | 0.243137 | 0.852003 |
| GOBP_FUSION_OF_SPERM_TO_EGG_PLASMA_MEMBRANE_INVOLVED_IN | 89 | 0.29884 | 1.189289 | 0.110294 | 0.850291 |
| GOBP_NEGATIVE_REGULATION_OF_INTRINSIC_APOPTOTIC_SIGNALING_ | 61 | 0.323764 | 1.188863 | 0.166667 | 0.849681 |
| GOBP_PROTEOGLYCAN_BIOSYNTHETIC_PROCESS | 42 | 0.362033 | 1.188402 | 0.191919 | 0.849048 |
| GOBP_NEGATIVE_REGULATION_OF_RESPONSE_TO_ENDOPLASMIC_RET | 44 | 0.337454 | 1.186757 | 0.177083 | 0.853139 |
| GOBP_BROWN_FAT_CELL_DIFFERENTIATION | 132 | 0.28332 | 1.186757 | 0.030769 | 0.850673 |
| GOBP_HINDBRAIN_DEVELOPMENT | 30 | 0.366842 | 1.185812 | 0.215517 | 0.851945 |
| GOBP_POSTSYNAPTIC_SPECIALIZATION_ORGANIZATION | 22 | 0.395705 | 1.184272 | 0.196787 | 0.855425 |
| GOBP_PLASMA_MEMBRANE_FUSION | 21 | 0.40929 | 1.183736 | 0.275194 | 0.854943 |
| GOBP_NUCLEOTIDE_EXCISION_REPAIR_PREINCISION_COMPLEX_STABI | 19 | 0.411813 | 1.183694 | 0.220472 | 0.85263 |
| GOBP_KIDNEY_MESENCHYME_DEVELOPMENT | 152 | 0.276331 | 1.183293 | 0.071429 | 0.851993 |
| GOBP_RESPONSE_TO_NUTRIENT | 44 | 0.346503 | 1.183001 | 0.162562 | 0.850864 |
| GOBP_VESICLE_DOCKING_INVOLVED_IN_EXOCYTOSIS | 15 | 0.430786 | 1.18299 | 0.278388 | 0.848509 |
| GOBP_NEGATIVE_REGULATION_OF_BONE_RESORPTION | 108 | 0.293727 | 1.181252 | 0.078652 | 0.853037 |
| GOBP_FORMATION_OF_PRIMARY_GERM_LAYER | 43 | 0.339778 | 1.181137 | 0.198068 | 0.851084 |
| GOBP_ENDOCRINE_HORMONE_SECRETION | 148 | 0.28383 | 1.180805 | 0.086957 | 0.850014 |
| GOBP_NEUROTRANSMITTER_SECRETION | 65 | 0.320088 | 1.180535 | 0.180645 | 0.848655 |
| GOBP_REGULATION_OF_HEART_GROWTH | 138 | 0.283559 | 1.180376 | 0.1 | 0.846833 |
| GOBP_BRANCHING_MORPHOGENESIS_OF_AN_EPITHELIAL_TUBE | 17 | 0.421642 | 1.179151 | 0.271062 | 0.849358 |
| GOBP_CELLULAR_RESPONSE_TO_REACTIVE_NITROGEN_SPECIES | 212 | 0.275017 | 1.179049 | 0.055556 | 0.847364 |
| GOBP_EXTRINSIC_APOPTOTIC_SIGNALING_PATHWAY | 24 | 0.391837 | 1.178285 | 0.237154 | 0.848066 |
| GOBP_CELL_COMMUNICATION_BY_ELECTRICAL_COUPLING_INVOLVED_ | 20 | 0.401413 | 1.177532 | 0.254613 | 0.848832 |
| GOBP_MALE_GENITALIA_DEVELOPMENT | 122 | 0.275574 | 1.177166 | 0.083333 | 0.848088 |
| GOBP_PLATELET_DEGRANULATION | 231 | 0.267017 | 1.176757 | 0.1 | 0.847434 |
| GOBP_DENDRITE_DEVELOPMENT | 23 | 0.382607 | 1.176739 | 0.253846 | 0.845171 |
| GOBP_NUCLEOTIDE_SUGAR_BIOSYNTHETIC_PROCESS | 90 | 0.303795 | 1.175373 | 0.127273 | 0.848346 |
| GOBP_RENAL_TUBULE_DEVELOPMENT | 32 | 0.35995 | 1.175243 | 0.196721 | 0.846733 |
| GOBP_REGULATION_OF_NATURAL_KILLER_CELL_ACTIVATION | 83 | 0.294245 | 1.174536 | 0.135714 | 0.847115 |
| GOBP_ENERGY_RESERVE_METABOLIC_PROCESS | 15 | 0.417136 | 1.174506 | 0.263736 | 0.844914 |
| GOBP_EPITHELIAL_CELL_MATURATION | 409 | 0.249254 | 1.174165 | 0 | 0.843889 |
| GOBP_NEGATIVE_REGULATION_OF_HYDROLASE_ACTIVITY | 37 | 0.345939 | 1.173819 | 0.1875 | 0.843148 |
| GOBP_CYCLIC_NUCLEOTIDE_METABOLIC_PROCESS | 119 | 0.288176 | 1.172327 | 0.119565 | 0.846623 |
| GOBP_CELLULAR_RESPONSE_TO_HEAT | 111 | 0.289815 | 1.171943 | 0.128205 | 0.845916 |
| GOBP_CARDIAC_VENTRICLE_DEVELOPMENT | 18 | 0.411267 | 1.170019 | 0.24031 | 0.851211 |
| GOBP_REGULATION_OF_APPETITE | 65 | 0.313973 | 1.169234 | 0.203008 | 0.851915 |
| GOBP_RESPONSE_TO_ACTIVITY | 32 | 0.353691 | 1.169141 | 0.246305 | 0.849974 |
| GOBP_ORGANIC_CATION_TRANSPORT | 43 | 0.344454 | 1.169082 | 0.206349 | 0.847937 |
| GOBP_GLUCAN_BIOSYNTHETIC_PROCESS | 201 | 0.260858 | 1.168941 | 0.034483 | 0.84627 |
| GOBP_REGULATION_OF_NEUROTRANSMITTER_LEVELS | 29 | 0.373287 | 1.168374 | 0.230089 | 0.846192 |
| GOBP_NUCLEOTIDE_EXCISION_REPAIR_PREINCISION_COMPLEX_ASSEN | 71 | 0.306991 | 1.168323 | 0.20155 | 0.844148 |
| GOBP_NEPHRON_MORPHOGENESIS | 62 | 0.317057 | 1.168091 | 0.186207 | 0.842857 |
| GOBP_NEGATIVE_REGULATION_OF_PEPTIDE_SECRETION | 46 | 0.338389 | 1.167565 | 0.191011 | 0.842804 |
| GOBP_REGULATION_OF_FILOPODIUM_ASSEMBLY | 27 | 0.383869 | 1.167068 | 0.214286 | 0.842591 |
| GOBP_CELL_SURFACE_RECEPTOR_SIGNALING_PATHWAY_INVOLVED_IN | 280 | 0.250139 | 1.166528 | 0.037037 | 0.842514 |
| GOBP_POSITIVE_REGULATION_OF_SECRETION | 19 | 0.404622 | 1.166155 | 0.233962 | 0.841932 |
| GOBP_RENAL_VESICLE_DEVELOPMENT | 88 | 0.290223 | 1.164025 | 0.148148 | 0.848304 |
| GOBP_KIDNEY_MORPHOGENESIS | 26 | 0.371483 | 1.162069 | 0.223214 | 0.853861 |
| GOBP_NEGATIVE_REGULATION_OF_CARDIAC_MUSCLE_TISSUE_GROWT | 79 | 0.296198 | 1.161161 | 0.160714 | 0.855223 |
| GOBP_LONG_TERM_SYNAPTIC_POTENTIATION | 19 | 0.4104 | 1.160997 | 0.25188 | 0.853603 |
| GOBP_NEGATIVE_REGULATION_OF_TISSUE_REMODELING | 109 | 0.292425 | 1.160237 | 0.089744 | 0.854253 |
| GOBP_CELLULAR_TRANSITION_METAL_ION_HOMEOSTASIS | 24 | 0.389552 | 1.159272 | 0.275194 | 0.855921 |
| GOBP_REGULATION_OF_DENDRITE_EXTENSION | 23 | 0.386772 | 1.159197 | 0.24898 | 0.853989 |
| GOBP_PROTEIN_ACTIVATION_CASCADE | 84 | 0.292514 | 1.159135 | 0.123894 | 0.852055 |
| GOBP_INNER_EAR_MORPHOGENESIS | 19 | 0.40856 | 1.158929 | 0.240876 | 0.8507 |
| GOBP_MESENCHYMAL_TO_EPITHELIAL_TRANSITION | 148 | 0.27227 | 1.158809 | 0.089286 | 0.848968 |
| GOBP_ORGAN_GROWTH | 55 | 0.322972 | 1.158775 | 0.165714 | 0.846977 |
| GOBP_EXCRETION | 63 | 0.319051 | 1.158457 | 0.179641 | 0.846074 |
| GOBP_NEGATIVE_REGULATION_OF_SYNAPTIC_TRANSMISSION | 19 | 0.39447 | 1.158428 | 0.261194 | 0.844045 |
| GOBP_ENGULFMENT_OF_APOPTOTIC_CELL | 65 | 0.311255 | 1.158282 | 0.166667 | 0.842492 |
| GOBP_AMINOGLYCAN_CATABOLIC_PROCESS | 55 | 0.331235 | 1.157916 | 0.160584 | 0.842037 |
| GOBP_NEURON_PROJECTION_REGENERATION | 15 | 0.440679 | 1.157792 | 0.275618 | 0.840383 |
| GOBP_BASEMENT_MEMBRANE_ASSEMBLY | 27 | 0.374166 | 1.15779 | 0.271255 | 0.838315 |
| GOBP_AXIS_ELONGATION | 58 | 0.319754 | 1.156894 | 0.203704 | 0.83993 |
| GOBP_OVULATION_CYCLE | 245 | 0.259646 | 1.156102 | 0.076923 | 0.840941 |
| GOBP_STRIATED_MUSCLE_CELL_DIFFERENTIATION | 38 | 0.348766 | 1.155718 | 0.218905 | 0.840573 |
| GOBP_ACTIVIN_RECEPTOR_SIGNALING_PATHWAY | 22 | 0.375666 | 1.155482 | 0.243542 | 0.839493 |
| GOBP_CELL_DIFFERENTIATION_INVOLVED_IN_EMBRYONIC_PLACENTA_ | 23 | 0.396296 | 1.155362 | 0.25641 | 0.837909 |
| GOBP_REGULATION_OF_SYNAPTIC_VESICLE_RECYCLING | 60 | 0.322785 | 1.155177 | 0.204969 | 0.836484 |
| GOBP_POSITIVE_REGULATION_OF_FAT_CELL_DIFFERENTIATION | 24 | 0.377194 | 1.154904 | 0.256604 | 0.835678 |
| GOBP_NEGATIVE_REGULATION_OF_BIOMINERALIZATION | 39 | 0.341784 | 1.154356 | 0.221675 | 0.835673 |
| GOBP_NEGATIVE_REGULATION_OF_CATION_CHANNEL_ACTIVITY | | | | | |

| | | | | | |
|---|---|---|---|---|---|
| GOBP_DNA_METHYLATION_OR_DEMETHYLATION | 90 | -0.396846 | -1.219049 | 0.140924 | 1 |
| GOBP_REGULATION_OF_CYCLASE_ACTIVITY | 41 | -0.436699 | -1.218891 | 0.189633 | 1 |
| GOBP_MITOCHONDRIAL_TRANSCRIPTION | 17 | -0.517568 | -1.218537 | 0.218208 | 1 |
| GOBP_DEFENSE_RESPONSE_TO_GRAM_POSITIVE_BACTERIUM | 71 | -0.405929 | -1.217734 | 0.171395 | 1 |
| GOBP_NEGATIVE_REGULATION_OF_RNA_SPLICING | 25 | -0.471398 | -1.21754 | 0.216867 | 1 |
| GOBP_REGULATION_OF_CHROMATIN_SILENCING | 16 | -0.527696 | -1.21749 | 0.241144 | 1 |
| GOBP_REGULATION_OF_DNA_REPLICATION | 105 | -0.3864 | -1.217015 | 0.127109 | 1 |
| GOBP_POSITIVE_REGULATION_OF_COLD_INDUCED_THERMOGENESIS | 94 | -0.396298 | -1.216737 | 0.148106 | 1 |
| GOBP_REGULATION_OF_CILIUM_DEPENDENT_CELL_MOTILITY | 22 | -0.486496 | -1.216607 | 0.208499 | 1 |
| GOBP_REGULATION_OF_T_CELL_DIFFERENTIATION_IN_THYMUS | 22 | -0.492492 | -1.21513 | 0.215659 | 1 |
| GOBP_REGULATION_OF_TRANSCRIPTION_INVOLVED_IN_G1_S_TRANSITION_C | 35 | -0.444739 | -1.214173 | 0.193758 | 1 |
| GOBP_REGULATION_OF_LIPID_CATABOLIC_PROCESS | 54 | -0.423993 | -1.21388 | 0.177832 | 1 |
| GOBP_POSITIVE_REGULATION_OF_MEMBRANE_PROTEIN_ECTODOMAIN_PROT | 15 | -0.532954 | -1.213347 | 0.219346 | 1 |
| GOBP_NUCLEOSIDE_BISPHOSPHATE_BIOSYNTHETIC_PROCESS | 66 | -0.406711 | -1.21159 | 0.15808 | 1 |
| GOBP_DNA_CATABOLIC_PROCESS | 38 | -0.440895 | -1.211542 | 0.197201 | 1 |
| GOBP_B_CELL_ACTIVATION_INVOLVED_IN_IMMUNE_RESPONSE | 77 | -0.402787 | -1.211376 | 0.170237 | 1 |
| GOBP_CD4_POSITIVE_ALPHA_BETA_T_CELL_ACTIVATION | 96 | -0.392514 | -1.211167 | 0.160314 | 1 |
| GOBP_POSITIVE_REGULATION_OF_DNA_BINDING | 53 | -0.420052 | -1.210634 | 0.170552 | 1 |
| GOBP_POSITIVE_REGULATION_OF_AMINO_ACID_TRANSPORT | 15 | -0.521836 | -1.210229 | 0.242254 | 1 |
| GOBP_HOMOLOGOUS_CHROMOSOME_PAIRING_AT_MEIOSIS | 44 | -0.434565 | -1.210092 | 0.196822 | 1 |
| GOBP_HYDROGEN_PEROXIDE_BIOSYNTHETIC_PROCESS | 16 | -0.527659 | -1.209233 | 0.238621 | 1 |
| GOBP_ACTIVATION_OF_ADENYLATE_CYCLASE_ACTIVITY | 35 | -0.445189 | -1.206652 | 0.209158 | 1 |
| GOBP_RESPONSE_TO_ATP | 29 | -0.451926 | -1.206324 | 0.216389 | 1 |
| GOBP_POSITIVE_REGULATION_OF_NUCLEAR_DIVISION | 54 | -0.419473 | -1.206224 | 0.195889 | 1 |
| GOBP_POSITIVE_REGULATION_OF_INTERLEUKIN_1_PRODUCTION | 60 | -0.416281 | -1.205989 | 0.2 | 1 |
| GOBP_CELLULAR_DEFENSE_RESPONSE | 51 | -0.423569 | -1.205536 | 0.19521 | 1 |
| GOBP_REGULATION_OF_NATURAL_KILLER_CELL_MEDIATED_IMMUNITY | 48 | -0.430519 | -1.205288 | 0.200993 | 1 |
| GOBP_CELLULAR_PROCESS_INVOLVED_IN_REPRODUCTION_IN_MULTICELLU | 311 | -0.35992 | -1.204132 | 0.075988 | 1 |
| GOBP_SENSORY_PERCEPTION_OF_TASTE | 45 | -0.432218 | -1.203231 | 0.208333 | 1 |
| GOBP_AMINO_ACID_BETAINE_METABOLIC_PROCESS | 17 | -0.517918 | -1.202652 | 0.243056 | 1 |
| GOBP_POSITIVE_REGULATION_OF_MACROPHAGE_MIGRATION | 24 | -0.477626 | -1.201916 | 0.231076 | 1 |
| GOBP_REGULATION_OF_DNA_DEPENDENT_DNA_REPLICATION | 47 | -0.422266 | -1.201575 | 0.203349 | 1 |
| GOBP_NEUTRAL_LIPID_METABOLIC_PROCESS | 118 | -0.380736 | -1.200818 | 0.146205 | 1 |
| GOBP_RIBOSOMAL_LARGE_SUBUNIT_ASSEMBLY | 28 | -0.46384 | -1.200578 | 0.224227 | 1 |
| GOBP_GAMMA_AMINOBUTYRIC_ACID_SIGNALING_PATHWAY | 24 | -0.462845 | -1.199382 | 0.225764 | 1 |
| GOBP_N_TERMINAL_PROTEIN_AMINO_ACID_ACETYLATION | 17 | -0.513054 | -1.19838 | 0.252778 | 1 |
| GOBP_RETINOIC_ACID_RECEPTOR_SIGNALING_PATHWAY | 31 | -0.454451 | -1.198251 | 0.222791 | 1 |
| GOBP_FOREBRAIN_GENERATION_OF_NEURONS | 51 | -0.420515 | -1.198102 | 0.208086 | 1 |
| GOBP_CD4_POSITIVE_ALPHA_BETA_T_CELL_DIFFERENTIATION | 79 | -0.391879 | -1.197891 | 0.177329 | 1 |
| GOBP_REGULATION_OF_T_CELL_RECEPTOR_SIGNALING_PATHWAY | 40 | -0.4437 | -1.196889 | 0.217054 | 1 |
| GOBP_REGULATION_OF_HYDROGEN_PEROXIDE_METABOLIC_PROCESS | 19 | -0.498067 | -1.196343 | 0.247552 | 1 |
| GOBP_DNA_RECOMBINATION | 285 | -0.355265 | -1.194646 | 0.101626 | 1 |
| GOBP_CYTOLYSIS | 22 | -0.484178 | -1.193184 | 0.256614 | 1 |
| GOBP_LYMPHOCYTE_MIGRATION | 109 | -0.379284 | -1.192061 | 0.167778 | 1 |
| GOBP_REGULATION_OF_MACROPHAGE_MIGRATION | 38 | -0.434928 | -1.191702 | 0.224719 | 1 |
| GOBP_TRANSCYTOSIS | 20 | -0.488654 | -1.191477 | 0.243641 | 1 |
| GOBP_POSITIVE_REGULATION_OF_DNA_REPLICATION | 37 | -0.431903 | -1.1907 | 0.219849 | 1 |
| GOBP_VITAMIN_METABOLIC_PROCESS | 123 | -0.371754 | -1.190538 | 0.150163 | 1 |
| GOBP_REGULATION_OF_HISTONE_H3_K9_METHYLATION | 20 | -0.499049 | -1.189512 | 0.257556 | 1 |
| GOBP_CELL_KILLING | 150 | -0.368024 | -1.187864 | 0.155954 | 1 |
| GOBP_NEGATIVE_REGULATION_OF_PROTEIN_MATURATION | 26 | -0.463606 | -1.187809 | 0.247718 | 1 |
| GOBP_ANATOMICAL_STRUCTURE_ARRANGEMENT | 15 | -0.509864 | -1.187421 | 0.273894 | 1 |
| GOBP_REGULATION_OF_MITOTIC_NUCLEAR_DIVISION | 107 | -0.375675 | -1.186951 | 0.170595 | 1 |
| GOBP_RESPONSE_TO_PH | 36 | -0.439587 | -1.186896 | 0.227735 | 1 |
| GOBP_REGULATORY_T_CELL_DIFFERENTIATION | 34 | -0.440177 | -1.185582 | 0.248072 | 1 |
| GOBP_SOMATIC_STEM_CELL_POPULATION_MAINTENANCE | 65 | -0.401341 | -1.184898 | 0.19422 | 1 |
| GOBP_B_CELL_DIFFERENTIATION | 118 | -0.37301 | -1.184843 | 0.170652 | 1 |
| GOBP_NEGATIVE_REGULATION_OF_INTERLEUKIN_12_PRODUCTION | 17 | -0.502139 | -1.184157 | 0.258953 | 1 |
| GOBP_LIPOXYGENASE_PATHWAY | 16 | -0.510303 | -1.183602 | 0.269601 | 1 |
| GOBP_POSITIVE_REGULATION_OF_KERATINOCYTE_DIFFERENTIATION | 18 | -0.493338 | -1.182678 | 0.269774 | 1 |
| GOBP_SPERMATID_DIFFERENTIATION | 133 | -0.371057 | -1.182534 | 0.173913 | 1 |
| GOBP_NEUROTRANSMITTER_REUPTAKE | 30 | -0.445513 | -1.182208 | 0.230373 | 1 |
| GOBP_NEGATIVE_REGULATION_OF_INTERLEUKIN_6_PRODUCTION | 43 | -0.420671 | -1.182063 | 0.23494 | 1 |
| GOBP_T_CELL_DIFFERENTIATION_INVOLVED_IN_IMMUNE_RESPONSE | 72 | -0.394646 | -1.181988 | 0.19092 | 1 |
| GOBP_T_CELL_DIFFERENTIATION | 238 | -0.357294 | -1.1811 | 0.1294 | 1 |
| GOBP_RESPONSE_TO_INTERLEUKIN_4 | 34 | -0.441262 | -1.180219 | 0.23354 | 1 |
| GOBP_GLUCOSAMINE_CONTAINING_COMPOUND_METABOLIC_PROCESS | 25 | -0.464878 | -1.17941 | 0.261968 | 1 |
| GOBP_CHEMICAL_SYNAPTIC_TRANSMISSION_POSTSYNAPTIC | 97 | -0.383153 | -1.179389 | 0.175947 | 1 |
| GOBP_CELLULAR_RESPONSE_TO_PH | 20 | -0.48707 | -1.178707 | 0.265522 | 1 |
| GOBP_RESPONSE_TO_ACIDIC_PH | 22 | -0.47708 | -1.178267 | 0.250333 | 1 |
| GOBP_POSITIVE_REGULATION_OF_CELL_ACTIVATION | 388 | -0.34626 | -1.178161 | 0.075911 | 1 |
| GOBP_MULTICELLULAR_ORGANISMAL_RESPONSE_TO_STRESS | 63 | -0.401082 | -1.177758 | 0.207701 | 1 |
| GOBP_REGULATION_OF_ICOSANOID_SECRETION | 17 | -0.511709 | -1.177552 | 0.278779 | 1 |
| GOBP_POSITIVE_REGULATION_OF_SMOOTH_MUSCLE_CONTRACTION | 25 | -0.472666 | -1.176924 | 0.256242 | 1 |
| GOBP_ICOSANOID_BIOSYNTHETIC_PROCESS | 49 | -0.417675 | -1.175993 | 0.237139 | 1 |
| GOBP_POSITIVE_REGULATION_OF_CYTOSOLIC_CALCIUM_ION_CONCENTRATI | 32 | -0.450185 | -1.175734 | 0.241107 | 1 |
| GOBP_SMOOTH_MUSCLE_CONTRACTION | 91 | -0.383329 | -1.175377 | 0.205215 | 1 |
| GOBP_RETINOL_METABOLIC_PROCESS | 46 | -0.417875 | -1.174799 | 0.23733 | 1 |
| GOBP_POSITIVE_REGULATION_OF_FATTY_ACID_METABOLIC_PROCESS | 32 | -0.44012 | -1.174551 | 0.24934 | 1 |
| GOBP_OOGENESIS | 79 | -0.386946 | -1.174517 | 0.20023 | 1 |
| GOBP_GERM_CELL_DEVELOPMENT | 223 | -0.35705 | -1.174123 | 0.137681 | 1 |
| GOBP_DEFENSE_RESPONSE_TO_GRAM_NEGATIVE_BACTERIUM | 57 | -0.403352 | -1.174101 | 0.210526 | 1 |
| GOBP_RESPONSE_TO_INTERLEUKIN_6 | 41 | -0.429897 | -1.173821 | 0.219512 | 1 |
| GOBP_LEUKOCYTE_CHEMOTAXIS | 211 | -0.353113 | -1.173454 | 0.149688 | 1 |
| GOBP_FEMALE_MEIOTIC_NUCLEAR_DIVISION | 26 | -0.45811 | -1.173323 | 0.264052 | 1 |
| GOBP_NEGATIVE_REGULATION_OF_SODIUM_ION_TRANSPORT | 17 | -0.510762 | -1.172775 | 0.281081 | 1 |
| GOBP_POSITIVE_REGULATION_OF_REPRODUCTIVE_PROCESS | 72 | -0.387818 | -1.17271 | 0.206186 | 1 |
| GOBP_NEGATIVE_REGULATION_OF_NEURAL_PRECURSOR_CELL_PROLIFERA | 18 | -0.481988 | -1.172618 | 0.261224 | 1 |
| GOBP_FATTY_ACYL_COA_METABOLIC_PROCESS | 42 | -0.422853 | -1.171852 | 0.249691 | 1 |
| GOBP_ISOPRENOID_METABOLIC_PROCESS | 128 | -0.368605 | -1.170957 | 0.194175 | 1 |
| GOBP_RESPONSE_TO_MOLECULE_OF_BACTERIAL_ORIGIN | 315 | -0.34865 | -1.170604 | 0.110204 | 1 |
| GOBP_POSITIVE_REGULATION_OF_NATURAL_KILLER_CELL_MEDIATED_IMMU | 31 | -0.441551 | -1.17035 | 0.260026 | 1 |
| GOBP_KINETOCHORE_ORGANIZATION | 22 | -0.473018 | -1.170301 | 0.260516 | 1 |
| GOBP_GASTRIC_ACID_SECRETION | 17 | -0.491007 | -1.17003 | 0.275623 | 1 |
| GOBP_SERTOLI_CELL_DIFFERENTIATION | 15 | -0.514503 | -1.169851 | 0.280702 | 1 |
| GOBP_N_ACETYLGLUCOSAMINE_METABOLIC_PROCESS | 19 | -0.485019 | -1.169647 | 0.265278 | 1 |
| GOBP_BILE_ACID_AND_BILE_SALT_TRANSPORT | 29 | -0.454624 | -1.169383 | 0.254642 | 1 |
| GOBP_PYRIMIDINE_NUCLEOSIDE_MONOPHOSPHATE_METABOLIC_PROCESS | 18 | -0.494035 | -1.168615 | 0.26511 | 1 |
| GOBP_SKELETAL_MUSCLE_CELL_DIFFERENTIATION | 58 | -0.4075 | -1.168392 | 0.208038 | 1 |
| GOBP_INTERLEUKIN_1_PRODUCTION | 104 | -0.374763 | -1.167777 | 0.210699 | 1 |
| GOBP_TRANSCRIPTION_BY_RNA_POLYMERASE_I | 68 | -0.39738 | -1.16716 | 0.217241 | 1 |
| GOBP_REGULATION_OF_CHROMATIN_ASSEMBLY_OR_DISASSEMBLY | 24 | -0.469155 | -1.167067 | 0.255291 | 1 |
| GOBP_CELL_FATE_COMMITMENT | 222 | -0.352595 | -1.16627 | 0.163712 | 1 |
| GOBP_POLYAMINE_METABOLIC_PROCESS | 17 | -0.491506 | -1.166239 | 0.269542 | 1 |
| GOBP_REGULATION_OF_TOLL_LIKE_RECEPTOR_4_SIGNALING_PATHWAY | 22 | -0.468679 | -1.166235 | 0.283984 | 1 |
| GOBP_NEGATIVE_REGULATION_OF_DIGESTIVE_SYSTEM_PROCESS | 15 | -0.507697 | -1.165905 | 0.284138 | 1 |
| GOBP_POSITIVE_REGULATION_OF_METAPHASE_ANAPHASE_TRANSITION_OF_ | 15 | -0.501103 | -1.165427 | 0.272476 | 1 |
| GOBP_NEGATIVE_REGULATION_OF_GLYCOPROTEIN_METABOLIC_PROCESS | 15 | -0.510976 | -1.164052 | 0.288184 | 1 |
| GOBP_PHOSPHOLIPID_CATABOLIC_PROCESS | 47 | -0.406489 | -1.163725 | 0.258621 | 1 |
| GOBP_FATTY_ACYL_COA_BIOSYNTHETIC_PROCESS | 31 | -0.447807 | -1.162935 | 0.26536 | 1 |
| GOBP_THIOESTER_METABOLIC_PROCESS | 104 | -0.370125 | -1.162394 | 0.204036 | 1 |
| GOBP_MONONUCLEAR_CELL_MIGRATION | 178 | -0.354416 | -1.16209 | 0.165792 | 1 |
| GOBP_MEMBRANE_DISASSEMBLY | 15 | -0.511138 | -1.161915 | 0.282337 | 1 |
| GOBP_C4_DICARBOXYLATE_TRANSPORT | 24 | -0.462206 | -1.160723 | 0.254545 | 1 |
| GOBP_INTERLEUKIN_2_PRODUCTION | 58 | -0.403124 | -1.160547 | 0.233092 | 1 |
| GOBP_MEIOTIC_CHROMOSOME_SEGREGATION | 82 | -0.382895 | -1.160402 | 0.220588 | 1 |
| GOBP_POSITIVE_REGULATION_OF_INTERLEUKIN_2_PRODUCTION | 30 | -0.445941 | -1.160397 | 0.277562 | 1 |
| GOBP_TERPENOID_METABOLIC_PROCESS | 110 | -0.366423 | -1.159301 | 0.20242 | 1 |
| GOBP_REGULATION_OF_T_CELL_ACTIVATION | 316 | -0.343149 | -1.159153 | 0.143002 | 1 |
| GOBP_REGULATION_OF_TRIGLYCERIDE_METABOLIC_PROCESS | 33 | -0.429475 | -1.158285 | 0.285156 | 1 |
| GOBP_CELLULAR_RESPONSE_TO_FATTY_ACID | 35 | -0.431676 | -1.157895 | 0.250656 | 1 |
| GOBP_HIPPOCAMPUS_DEVELOPMENT | 67 | -0.389651 | -1.157684 | 0.227433 | 1 |
| GOBP_TERPENOID_BIOSYNTHETIC_PROCESS | 15 | -0.507019 | -1.156646 | 0.292072 | 1 |
| GOBP_GAMMA_DELTA_T_CELL_ACTIVATION | 19 | -0.482446 | -1.156596 | 0.263675 | 1 |
| GOBP_REGULATION_OF_RETINOIC_ACID_RECEPTOR_SIGNALING_PATHWAY | 16 | -0.500098 | -1.156503 | 0.279452 | 1 |
| GOBP_CELLULAR_MODIFIED_AMINO_ACID_CATABOLIC_PROCESS | 29 | -0.448303 | -1.156199 | 0.260131 | 1 |
| GOBP_CHROMOSOME_ORGANIZATION_INVOLVED_IN_MEIOTIC_CELL_CYCLE | 61 | -0.397998 | -1.155991 | 0.248492 | 1 |
| GOBP_VENTRICULAR_CARDIAC_MUSCLE_TISSUE_DEVELOPMENT | 48 | -0.404249 | -1.155697 | 0.246944 | 1 |
| GOBP_POSITIVE_REGULATION_OF_ALPHA_BETA_T_CELL_PROLIFERATION | 21 | -0.458495 | -1.155544 | 0.269737 | 1 |
| GOBP_REGULATION_OF_CILIUM_MOVEMENT | 32 | -0.43931 | -1.153658 | 0.24937 | 1 |
| GOBP_POSITIVE_REGULATION_OF_BLOOD_CIRCULATION | 39 | -0.425151 | -1.153449 | 0.277301 | 1 |
| GOBP_NEGATIVE_REGULATION_OF_NEURON_DIFFERENTIATION | 57 | -0.394999 | -1.151571 | 0.257757 | 1 |
| GOBP_GRANULOCYTE_CHEMOTAXIS | 117 | -0.366041 | -1.151143 | 0.208609 | 1 |
| GOBP_RESPONSE_TO_CHEMOKINE | 91 | -0.37876 | -1.150929 | 0.234483 | 1 |
| GOBP_RECEPTOR_MEDIATED_ENDOCYTOSIS | 314 | -0.340734 | -1.150444 | 0.153611 | 1 |
| GOBP_POSITIVE_REGULATION_OF_B_CELL_MEDIATED_IMMUNITY | 38 | -0.425494 | -1.150003 | 0.253086 | 1 |
| GOBP_POSITIVE_REGULATION_OF_INFLAMMATORY_RESPONSE | 122 | -0.360917 | -1.149674 | 0.226395 | 1 |
| GOBP_DETECTION_OF_CHEMICAL_STIMULUS | 93 | -0.37442 | -1.149364 | 0.215176 | 1 |

| | | | | | |
|---|---|---|---|---|---|
| GOBP_EMBRYONIC_ORGAN_MORPHOGENESIS | 264 | 0.243544 | 1.154301 | 0 | 0.833783 |
| GOBP_POSITIVE_REGULATION_OF_NATURAL_KILLER_CELL_ACTIVATION | 20 | 0.39691 | 1.153983 | 0.251012 | 0.833029 |
| GOBP_POSITIVE_REGULATION_OF_SYNAPTIC_TRANSMISSION | 135 | 0.285119 | 1.153701 | 0.135135 | 0.83214 |
| GOBP_NEGATIVE_REGULATION_OF_ANION_TRANSPORT | 185 | 0.261915 | 1.153331 | 0.113636 | 0.831587 |
| GOBP_PROSTATE_GLAND_DEVELOPMENT | 40 | 0.350984 | 1.15306 | 0.223404 | 0.830475 |
| GOBP_EXCITATORY_SYNAPSE_ASSEMBLY | 25 | 0.383042 | 1.152498 | 0.268293 | 0.830565 |
| GOBP_SECRETION_BY_TISSUE | 30 | 0.365583 | 1.151902 | 0.22381 | 0.830746 |
| GOBP_AMINOGLYCAN_METABOLIC_PROCESS | 167 | 0.271766 | 1.151413 | 0.122807 | 0.830479 |
| GOBP_STRIATED_MUSCLE_CONTRACTION | 164 | 0.270004 | 1.1514 | 0.12963 | 0.828539 |
| GOBP_MEMBRANE_FUSION | 167 | 0.277929 | 1.149046 | 0.126984 | 0.835423 |
| GOBP_MODULATION_BY_VIRUS_OF_HOST_PROCESS | 29 | 0.356604 | 1.148634 | 0.255 | 0.834966 |
| GOBP_MYOBLAST_DIFFERENTIATION | 75 | 0.305193 | 1.148138 | 0.20155 | 0.834931 |
| GOBP_EMBRYONIC_DIGIT_MORPHOGENESIS | 50 | 0.321615 | 1.1475 | 0.236842 | 0.835345 |
| GOBP_LIPID_DIGESTION | 19 | 0.39586 | 1.145138 | 0.25 | 0.842734 |
| GOBP_REGULATION_OF_NOTCH_SIGNALING_PATHWAY | 103 | 0.28147 | 1.144267 | 0.158537 | 0.844094 |
| GOBP_INTERFERON_GAMMA_MEDIATED_SIGNALING_PATHWAY | 88 | 0.297785 | 1.144136 | 0.181818 | 0.842566 |
| GOBP_TYPE_2_IMMUNE_RESPONSE | 35 | 0.349544 | 1.14355 | 0.238532 | 0.842817 |
| GOBP_SINGLE_FERTILIZATION | 132 | 0.275928 | 1.143413 | 0.155844 | 0.841434 |
| GOBP_CELL_MATRIX_ADHESION | 221 | 0.259048 | 1.143256 | 0.054054 | 0.840052 |
| GOBP_MACROPHAGE_DIFFERENTIATION | 44 | 0.333063 | 1.143142 | 0.220588 | 0.838497 |
| GOBP_DIGESTIVE_TRACT_MORPHOGENESIS | 41 | 0.332943 | 1.142028 | 0.233696 | 0.840823 |
| GOBP_HEMATOPOIETIC_STEM_CELL_HOMEOSTASIS | 15 | 0.420842 | 1.141867 | 0.286275 | 0.839417 |
| GOBP_NEGATIVE_REGULATION_OF_POTASSIUM_ION_TRANSPORT | 29 | 0.357523 | 1.141733 | 0.257511 | 0.838021 |
| GOBP_REGULATION_OF_EXTRINSIC_APOPTOTIC_SIGNALING_PATHWAY_ | 56 | 0.315419 | 1.141641 | 0.226744 | 0.836464 |
| GOBP_REPRODUCTIVE_SYSTEM_DEVELOPMENT | 378 | 0.243205 | 1.141578 | 0 | 0.834771 |
| GOBP_MONOCYTE_DIFFERENTIATION | 34 | 0.346202 | 1.141327 | 0.227273 | 0.833841 |
| GOBP_NEGATIVE_REGULATION_OF_MUSCLE_CELL_APOPTOTIC_PROCE | 35 | 0.355253 | 1.139635 | 0.270742 | 0.838766 |
| GOBP_POSITIVE_REGULATION_OF_NERVOUS_SYSTEM_PROCESS | 30 | 0.361372 | 1.139234 | 0.277778 | 0.838302 |
| GOBP_MICROVILLUS_ASSEMBLY | 16 | 0.432856 | 1.137581 | 0.273381 | 0.842619 |
| GOBP_AROMATIC_AMINO_ACID_FAMILY_CATABOLIC_PROCESS | 20 | 0.389377 | 1.137483 | 0.253788 | 0.84114 |
| GOBP_ACTIVATION_OF_JANUS_KINASE_ACTIVITY | 15 | 0.431088 | 1.136371 | 0.267797 | 0.843718 |
| GOBP_GLOMERULAR_MESANGIUM_DEVELOPMENT | 15 | 0.424376 | 1.136239 | 0.283784 | 0.842298 |
| GOBP_AUTONOMIC_NERVOUS_SYSTEM_DEVELOPMENT | 42 | 0.344776 | 1.135087 | 0.241206 | 0.844706 |
| GOBP_POSITIVE_REGULATION_OF_CELL_MATRIX_ADHESION | 53 | 0.313278 | 1.134914 | 0.236025 | 0.843525 |
| GOBP_ZINC_ION_TRANSPORT | 27 | 0.363366 | 1.134797 | 0.228916 | 0.842062 |
| GOBP_REGULATION_OF_STRIATED_MUSCLE_CONTRACTION | 84 | 0.293269 | 1.134072 | 0.185185 | 0.84305 |
| GOBP_REGULATION_OF_TRANS_SYNAPTIC_SIGNALING | 388 | 0.258452 | 1.133331 | 0.090909 | 0.844153 |
| GOBP_ENDODERMAL_CELL_DIFFERENTIATION | 44 | 0.329131 | 1.132885 | 0.245283 | 0.844102 |
| GOBP_PLACENTA_DEVELOPMENT | 128 | 0.265274 | 1.132623 | 0.177778 | 0.843306 |
| GOBP_REGULATION_OF_NERVOUS_SYSTEM_PROCESS | 125 | 0.274139 | 1.132144 | 0.181818 | 0.843255 |
| GOBP_EXPORT_ACROSS_PLASMA_MEMBRANE | 56 | 0.310879 | 1.13108 | 0.242938 | 0.845717 |
| GOBP_REGULATION_OF_CELLULAR_EXTRAVASATION | 33 | 0.353441 | 1.13038 | 0.273543 | 0.846685 |
| GOBP_ARTERY_DEVELOPMENT | 86 | 0.298217 | 1.129888 | 0.208333 | 0.846464 |
| GOBP_REGULATION_OF_SENSORY_PERCEPTION | 33 | 0.346152 | 1.129132 | 0.262009 | 0.847571 |
| GOBP_REGULATION_OF_BLOOD_CIRCULATION | 265 | 0.260348 | 1.128396 | 0.125 | 0.848458 |
| GOBP_POLYSACCHARIDE_BIOSYNTHETIC_PROCESS | 74 | 0.296029 | 1.128173 | 0.221477 | 0.84746 |
| GOBP_POSITIVE_REGULATION_OF_DEFENSE_RESPONSE_TO_VIRUS_BY | 31 | 0.345054 | 1.128125 | 0.254098 | 0.845744 |
| GOBP_MEMBRANE_BIOGENESIS | 39 | 0.333004 | 1.128052 | 0.25 | 0.844168 |
| GOBP_MEMORY | 105 | 0.27858 | 1.12799 | 0.180952 | 0.842553 |
| GOBP_MHC_CLASS_II_BIOSYNTHETIC_PROCESS | 16 | 0.413667 | 1.127467 | 0.304965 | 0.842785 |
| GOBP_GLUTAMINE_METABOLIC_PROCESS | 22 | 0.3833 | 1.127291 | 0.3125 | 0.841652 |
| GOBP_THYMUS_DEVELOPMENT | 44 | 0.329369 | 1.127161 | 0.26178 | 0.840367 |
| GOBP_MODULATION_BY_HOST_OF_VIRAL_GENOME_REPLICATION | 18 | 0.420843 | 1.126814 | 0.277567 | 0.83987 |
| GOBP_RESPONSE_TO_MONOSACCHARIDE | 180 | 0.256344 | 1.126568 | 0.181818 | 0.838972 |
| GOBP_RESPONSE_TO_UV_B | 16 | 0.412782 | 1.126521 | 0.301471 | 0.83737 |
| GOBP_POSITIVE_REGULATION_OF_CALCIUM_ION_DEPENDENT_EXOCYT | 19 | 0.400071 | 1.126444 | 0.29588 | 0.835893 |
| GOBP_SIGNAL_RELEASE | 449 | 0.238955 | 1.1264 | 0 | 0.83431 |
| GOBP_POSITIVE_REGULATION_OF_CYTOKINE_PRODUCTION_INVOLVED_ | 53 | 0.31231 | 1.126236 | 0.237805 | 0.833164 |
| GOBP_CALCIUM_ION_REGULATED_EXOCYTOSIS | 62 | 0.300265 | 1.126033 | 0.2375 | 0.832154 |
| GOBP_METANEPHRIC_TUBULE_DEVELOPMENT | 21 | 0.384731 | 1.125831 | 0.26971 | 0.831219 |
| GOBP_NEGATIVE_REGULATION_OF_EXTRINSIC_APOPTOTIC_SIGNALING_ | 39 | 0.342386 | 1.125557 | 0.262136 | 0.830522 |
| GOBP_HYDROGEN_PEROXIDE_METABOLIC_PROCESS | 49 | 0.328872 | 1.125414 | 0.258427 | 0.829266 |
| GOBP_NEGATIVE_REGULATION_OF_PEPTIDE_HORMONE_SECRETION | 37 | 0.334003 | 1.125258 | 0.214286 | 0.828126 |
| GOBP_REGULATION_OF_ANTIGEN_PROCESSING_AND_PRESENTATION | 20 | 0.393845 | 1.123787 | 0.276364 | 0.831918 |
| GOBP_ADENYLATE_CYCLASE_INHIBITING_G_PROTEIN_COUPLED_RECE | 73 | 0.298527 | 1.1221 | 0.201342 | 0.836497 |
| GOBP_RESPONSE_TO_INTERFERON_GAMMA | 190 | 0.250829 | 1.118786 | 0.162162 | 0.847439 |
| GOBP_EMBRYONIC_HEART_TUBE_DEVELOPMENT | 72 | 0.286265 | 1.118033 | 0.19403 | 0.848835 |
| GOBP_POSITIVE_REGULATION_OF_HEART_GROWTH | 35 | 0.341136 | 1.1168 | 0.271186 | 0.851918 |
| GOBP_CARDIAC_VENTRICLE_MORPHOGENESIS | 67 | 0.296217 | 1.116659 | 0.261438 | 0.850694 |
| GOBP_SENSORY_PERCEPTION_OF_MECHANICAL_STIMULUS | 157 | 0.263514 | 1.116362 | 0.169231 | 0.850037 |
| GOBP_MYOTUBE_DIFFERENTIATION | 102 | 0.277292 | 1.115604 | 0.2 | 0.851092 |
| GOBP_NEGATIVE_REGULATION_OF_ION_TRANSPORT | 277 | 0.244348 | 1.115554 | 0.125 | 0.84952 |
| GOBP_CARDIAC_SEPTUM_DEVELOPMENT | 94 | 0.285202 | 1.115502 | 0.186275 | 0.847975 |
| GOBP_ARTERY_MORPHOGENESIS | 61 | 0.298143 | 1.115098 | 0.23871 | 0.847741 |
| GOBP_RESPONSE_TO_MECHANICAL_STIMULUS | 197 | 0.24786 | 1.114805 | 0.212121 | 0.84716 |
| GOBP_CARDIAC_MUSCLE_TISSUE_DEVELOPMENT | 188 | 0.25406 | 1.113578 | 0.142857 | 0.85027 |
| GOBP_MORPHOGENESIS_OF_A_BRANCHING_STRUCTURE | 180 | 0.257023 | 1.113081 | 0.066667 | 0.850444 |
| GOBP_DEVELOPMENTAL_GROWTH_INVOLVED_IN_MORPHOGENESIS | 217 | 0.253431 | 1.113033 | 0.228571 | 0.848878 |
| GOBP_POSITIVE_REGULATION_OF_T_CELL_MEDIATED_IMMUNITY | 51 | 0.312451 | 1.112329 | 0.289617 | 0.850048 |
| GOBP_CHAPERONE_COFACTOR_DEPENDENT_PROTEIN_REFOLDING | 34 | 0.336042 | 1.112154 | 0.258373 | 0.848968 |
| GOBP_TRANSITION_METAL_ION_HOMEOSTASIS | 130 | 0.272711 | 1.11047 | 0.177215 | 0.853826 |
| GOBP_REGULATION_OF_DEFENSE_RESPONSE_TO_VIRUS_BY_HOST | 40 | 0.331342 | 1.109447 | 0.29 | 0.85585 |
| GOBP_POSITIVE_REGULATION_OF_NEURAL_PRECURSOR_CELL_PROLIF | 48 | 0.32065 | 1.109399 | 0.24 | 0.854312 |
| GOBP_POSTSYNAPSE_ORGANIZATION | 156 | 0.26898 | 1.108986 | 0.169811 | 0.854119 |
| GOBP_REGULATION_OF_COAGULATION | 66 | 0.300186 | 1.108053 | 0.243421 | 0.855968 |
| GOBP_NEUROTRANSMITTER_METABOLIC_PROCESS | 26 | 0.357903 | 1.108018 | 0.318548 | 0.854335 |
| GOBP_REGULATION_OF_POTASSIUM_ION_TRANSMEMBRANE_TRANSPO | 56 | 0.309865 | 1.107496 | 0.275862 | 0.85451 |
| GOBP_RENAL_SYSTEM_VASCULATURE_DEVELOPMENT | 25 | 0.353836 | 1.107337 | 0.261411 | 0.853392 |
| GOBP_HEPATOCYTE_DIFFERENTIATION | 16 | 0.397677 | 1.107299 | 0.316364 | 0.851825 |
| GOBP_POSITIVE_REGULATION_OF_INSULIN_SECRETION | 64 | 0.293593 | 1.107183 | 0.253521 | 0.850522 |
| GOBP_PANCREAS_DEVELOPMENT | 64 | 0.29678 | 1.106841 | 0.258993 | 0.850208 |
| GOBP_FAT_CELL_DIFFERENTIATION | 210 | 0.255951 | 1.106831 | 0.088889 | 0.848563 |
| GOBP_REGIONALIZATION | 301 | 0.242091 | 1.106071 | 0.181818 | 0.849662 |
| GOBP_STRIATED_MUSCLE_CELL_APOPTOTIC_PROCESS | 35 | 0.338676 | 1.105496 | 0.281553 | 0.850192 |
| GOBP_RESPIRATORY_SYSTEM_DEVELOPMENT | 183 | 0.260925 | 1.105333 | 0.196429 | 0.849097 |
| GOBP_REGULATION_OF_INTEGRIN_MEDIATED_SIGNALING_PATHWAY | 15 | 0.408095 | 1.104087 | 0.33101 | 0.852071 |
| GOBP_NEGATIVE_REGULATION_OF_INTRINSIC_APOPTOTIC_SIGNALING_ | 20 | 0.368269 | 1.103921 | 0.294545 | 0.851 |
| GOBP_REGULATION_OF_CALCINEURIN_MEDIATED_SIGNALING | 32 | 0.342444 | 1.103114 | 0.273973 | 0.852516 |
| GOBP_ANTIGEN_PROCESSING_AND_PRESENTATION_OF_PEPTIDE_ANTI | 98 | 0.279607 | 1.10302 | 0.216981 | 0.851243 |
| GOBP_ANTERIOR_POSTERIOR_AXIS_SPECIFICATION | 40 | 0.326088 | 1.102556 | 0.252294 | 0.851318 |
| GOBP_RESPONSE_TO_VITAMIN_D | 29 | 0.347876 | 1.101997 | 0.319048 | 0.851805 |
| GOBP_EPITHELIAL_CELL_DEVELOPMENT | 200 | 0.258864 | 1.101511 | 0.263158 | 0.851867 |
| GOBP_RETINA_LAYER_FORMATION | 24 | 0.373694 | 1.101292 | 0.300781 | 0.851078 |
| GOBP_NEGATIVE_REGULATION_OF_VASCULATURE_DEVELOPMENT | 98 | 0.276971 | 1.100952 | 0.221311 | 0.850821 |
| GOBP_RRNA_METHYLATION | 28 | 0.35397 | 1.100751 | 0.297872 | 0.849876 |
| GOBP_POSITIVE_REGULATION_OF_CREB_TRANSCRIPTION_FACTOR_AC | 19 | 0.395262 | 1.099571 | 0.323843 | 0.852765 |
| GOBP_REGULATION_OF_T_CELL_MEDIATED_CYTOTOXICITY | 38 | 0.325227 | 1.098939 | 0.244019 | 0.853696 |
| GOBP_NEGATIVE_REGULATION_OF_BONE_MINERALIZATION | 15 | 0.417743 | 1.098625 | 0.337884 | 0.85326 |
| GOBP_ORGANELLE_MEMBRANE_FUSION | 109 | 0.2696 | 1.097947 | 0.236559 | 0.854323 |
| GOBP_CELLULAR_RESPONSE_TO_CALCIUM_ION | 80 | 0.285799 | 1.097026 | 0.201681 | 0.856202 |
| GOBP_DENDRITE_EXTENSION | 33 | 0.34119 | 1.096453 | 0.286364 | 0.856781 |
| GOBP_AROMATIC_AMINO_ACID_FAMILY_METABOLIC_PROCESS | 30 | 0.349258 | 1.095872 | 0.32287 | 0.85738 |
| GOBP_REGENERATION | 182 | 0.25817 | 1.095375 | 0.217391 | 0.857673 |
| GOBP_SYMPATHETIC_NERVOUS_SYSTEM_DEVELOPMENT | 19 | 0.389381 | 1.093747 | 0.313253 | 0.862392 |
| GOBP_NERVE_DEVELOPMENT | 65 | 0.296743 | 1.093728 | 0.270968 | 0.860815 |
| GOBP_MUSCLE_FILAMENT_SLIDING | 38 | 0.324266 | 1.092679 | 0.299492 | 0.863211 |
| GOBP_DETECTION_OF_MECHANICAL_STIMULUS | 47 | 0.310658 | 1.091225 | 0.265957 | 0.867177 |
| GOBP_ENTRY_OF_BACTERIUM_INTO_HOST_CELL | 15 | 0.405657 | 1.091112 | 0.3367 | 0.865932 |
| GOBP_HORMONE_MEDIATED_SIGNALING_PATHWAY | 172 | 0.254457 | 1.090761 | 0.244444 | 0.865595 |
| GOBP_MOTILE_CILIUM_ASSEMBLY | 20 | 0.377212 | 1.089442 | 0.33574 | 0.868985 |
| GOBP_NEGATIVE_REGULATION_OF_POTASSIUM_ION_TRANSMEMBRANE | 23 | 0.362588 | 1.089367 | 0.349138 | 0.867587 |
| GOBP_NEGATIVE_REGULATION_OF_VIRAL_PROCESS | 83 | 0.277709 | 1.089342 | 0.241611 | 0.866018 |
| GOBP_METAL_ION_EXPORT | 44 | 0.319988 | 1.088784 | 0.327684 | 0.866507 |
| GOBP_DEVELOPMENTAL_CELL_GROWTH | 208 | 0.247924 | 1.088699 | 0.153846 | 0.865219 |
| GOBP_RESPONSE_TO_FLUID_SHEAR_STRESS | 30 | 0.339135 | 1.088563 | 0.318386 | 0.86405 |
| GOBP_POSITIVE_REGULATION_OF_MRNA_PROCESSING | 32 | 0.34402 | 1.087133 | 0.316279 | 0.867725 |
| GOBP_POSITIVE_REGULATION_OF_ORGAN_GROWTH | 44 | 0.315067 | 1.08669 | 0.294416 | 0.867798 |
| GOBP_POSITIVE_REGULATION_OF_RNA_SPLICING | 36 | 0.320427 | 1.086084 | 0.34434 | 0.868703 |
| GOBP_VASCULAR_ENDOTHELIAL_GROWTH_FACTOR_RECEPTOR_SIGNA | 92 | 0.271513 | 1.085848 | 0.268519 | 0.86799 |
| GOBP_REGULATION_OF_TYPE_2_IMMUNE_RESPONSE | 29 | 0.353575 | 1.084331 | 0.318182 | 0.872143 |
| GOBP_DORSAL_VENTRAL_NEURAL_TUBE_PATTERNING | 20 | 0.37137 | 1.084209 | 0.328467 | 0.870945 |
| GOBP_RESPONSE_TO_GONADOTROPIN | 22 | 0.377702 | 1.082521 | 0.347985 | 0.875665 |
| GOBP_MUSCLE_CONTRACTION | 328 | 0.237307 | 1.082075 | 0.181818 | 0.875922 |
| GOBP_MUSCLE_CELL_DIFFERENTIATION | 326 | 0.237172 | 1.082023 | 0.058824 | 0.874557 |
| GOBP_PROTEIN_LOCALIZATION_TO_CELL_CELL_JUNCTION | 21 | 0.376747 | 1.081964 | 0.355372 | 0.87316 |
| GOBP_POSITIVE_REGULATION_OF_PROTEIN_BINDING | 82 | 0.289499 | 1.081933 | 0.299145 | 0.871671 |
| GOBP_PEPTIDYL_PROLINE_MODIFICATION | 52 | 0.304732 | 1.081623 | 0.329412 | 0.871329 |
| GOBP_REGULATION_OF_ENDOCYTOSIS | 199 | 0.250695 | 1.081421 | 0.171429 | 0.870444 |

| | | | | | |
|---|---|---|---|---|---|
| GOBP_REGULATION_OF_NUCLEAR_DIVISION | 130 | -0.360583 | -1.149163 | 0.2 | 1 |
| GOBP_AMELOGENESIS | 22 | -0.460984 | -1.149151 | 0.278296 | 1 |
| GOBP_REGULATION_OF_SYSTEMIC_ARTERIAL_BLOOD_PRESSURE | 83 | -0.376389 | -1.148548 | 0.241918 | 1 |
| GOBP_TELENCEPHALON_GLIAL_CELL_MIGRATION | 22 | -0.472554 | -1.146635 | 0.299578 | 1 |
| GOBP_LEUKOTRIENE_BIOSYNTHETIC_PROCESS | 18 | -0.486226 | -1.146256 | 0.295198 | 1 |
| GOBP_REGULATION_OF_MEIOTIC_NUCLEAR_DIVISION | 28 | -0.439522 | -1.146068 | 0.306069 | 1 |
| GOBP_ENAMEL_MINERALIZATION | 15 | -0.491329 | -1.145917 | 0.301273 | 1 |
| GOBP_CHROMATIN_REMODELING | 194 | -0.347463 | -1.145914 | 0.189781 | 1 |
| GOBP_SIGNALING_RECEPTOR_LIGAND_PRECURSOR_PROCESSING | 30 | -0.438055 | -1.145455 | 0.274936 | 1 |
| GOBP_REGULATION_OF_LIPOPROTEIN_PARTICLE_CLEARANCE | 18 | -0.480437 | -1.145158 | 0.269179 | 1 |
| GOBP_REGULATION_OF_LEUKOCYTE_PROLIFERATION | 234 | -0.342725 | -1.143754 | 0.189495 | 1 |
| GOBP_NEGATIVE_REGULATION_OF_METAPHASE_ANAPHASE_TRANSITION_OF | 40 | -0.413564 | -1.142896 | 0.287879 | 1 |
| GOBP_NEGATIVE_REGULATION_OF_LEUKOCYTE_PROLIFERATION | 86 | -0.370234 | -1.142439 | 0.242873 | 1 |
| GOBP_DENDRITIC_CELL_MIGRATION | 29 | -0.435989 | -1.142129 | 0.27086 | 1 |
| GOBP_POSITIVE_REGULATION_OF_MYELINATION | 16 | -0.487221 | -1.141655 | 0.307246 | 1 |
| GOBP_NUCLEUS_LOCALIZATION | 24 | -0.44733 | -1.140492 | 0.285714 | 1 |
| GOBP_ESTROGEN_METABOLIC_PROCESS | 35 | -0.425876 | -1.140271 | 0.289373 | 1 |
| GOBP_HOMEOSTASIS_OF_NUMBER_OF_CELLS_WITHIN_A_TISSUE | 24 | -0.458659 | -1.140014 | 0.287979 | 1 |
| GOBP_REGULATION_OF_SMOOTH_MUSCLE_CONTRACTION | 53 | -0.398553 | -1.138885 | 0.277778 | 1 |
| GOBP_DNA_DEPENDENT_DNA_REPLICATION_MAINTENANCE_OF_FIDELITY | 48 | -0.404932 | -1.13884 | 0.292743 | 1 |
| GOBP_DNA_DEPENDENT_DNA_REPLICATION | 156 | -0.350724 | -1.138703 | 0.221041 | 1 |
| GOBP_REGULATION_OF_LIPOPOLYSACCHARIDE_MEDIATED_SIGNALING_PATH | 24 | -0.452613 | -1.138402 | 0.299338 | 1 |
| GOBP_NEGATIVE_REGULATION_OF_LYMPHOCYTE_APOPTOTIC_PROCESS | 31 | -0.425764 | -1.138172 | 0.295775 | 1 |
| GOBP_LONG_CHAIN_FATTY_ACID_BIOSYNTHETIC_PROCESS | 32 | -0.43282 | -1.138146 | 0.286853 | 1 |
| GOBP_REGULATION_OF_ODONTOGENESIS | 15 | -0.502444 | -1.137741 | 0.306407 | 1 |
| GOBP_T_CELL_ACTIVATION | 445 | -0.33297 | -1.137411 | 0.124373 | 1 |
| GOBP_POSITIVE_REGULATION_OF_EPIDERMIS_DEVELOPMENT | 30 | -0.435439 | -1.137071 | 0.30039 | 1 |
| GOBP_POSITIVE_REGULATION_OF_PROTEIN_TYROSINE_KINASE_ACTIVITY | 59 | -0.389245 | -1.136913 | 0.259172 | 1 |
| GOBP_REGULATION_OF_CELL_MIGRATION_INVOLVED_IN_SPROUTING_ANGIO | 45 | -0.405074 | -1.136649 | 0.277428 | 1 |
| GOBP_CELLULAR_RESPONSE_TO_OSMOTIC_STRESS | 39 | -0.417808 | -1.136388 | 0.277987 | 1 |
| GOBP_CD40_SIGNALING_PATHWAY | 15 | -0.488405 | -1.136387 | 0.308824 | 1 |
| GOBP_NEUROMUSCULAR_PROCESS | 82 | -0.375549 | -1.135717 | 0.263039 | 1 |
| GOBP_HORMONE_BIOSYNTHETIC_PROCESS | 66 | -0.38145 | -1.13519 | 0.267131 | 1 |
| GOBP_NEGATIVE_REGULATION_OF_GLIOGENESIS | 29 | -0.43534 | -1.134775 | 0.271795 | 1 |
| GOBP_REGULATION_OF_ORGANIC_ACID_TRANSPORT | 57 | -0.389344 | -1.134195 | 0.291415 | 1 |
| GOBP_REGULATION_OF_CYTOSOLIC_CALCIUM_ION_CONCENTRATION | 325 | -0.333355 | -1.133734 | 0.166667 | 1 |
| GOBP_METHIONINE_METABOLIC_PROCESS | 16 | -0.496218 | -1.133651 | 0.336066 | 1 |
| GOBP_REGULATION_OF_B_CELL_MEDIATED_IMMUNITY | 54 | -0.393167 | -1.131586 | 0.275656 | 1 |
| GOBP_CELLULAR_AMINO_ACID_BIOSYNTHETIC_PROCESS | 76 | -0.372834 | -1.131435 | 0.276023 | 1 |
| GOBP_CELL_CHEMOTAXIS | 279 | -0.338856 | -1.130974 | 0.201015 | 1 |
| GOBP_HOMOLOGOUS_RECOMBINATION | 57 | -0.387573 | -1.130508 | 0.305353 | 1 |
| GOBP_NEGATIVE_REGULATION_OF_DNA_DEPENDENT_DNA_REPLICATION | 17 | -0.485111 | -1.130119 | 0.321088 | 1 |
| GOBP_DENDRITIC_CELL_CHEMOTAXIS | 24 | -0.438368 | -1.129159 | 0.302949 | 1 |
| GOBP_NEUTROPHIL_CHEMOTAXIS | 96 | -0.36478 | -1.128038 | 0.264342 | 1 |
| GOBP_PARAXIAL_MESODERM_DEVELOPMENT | 17 | -0.479239 | -1.125822 | 0.330176 | 1 |
| GOBP_DNA_GEOMETRIC_CHANGE | 112 | -0.354259 | -1.124599 | 0.265351 | 1 |
| GOBP_GLUCOSE_6_PHOSPHATE_METABOLIC_PROCESS | 23 | -0.452982 | -1.123749 | 0.31912 | 1 |
| GOBP_PHASIC_SMOOTH_MUSCLE_CONTRACTION | 18 | -0.474837 | -1.123197 | 0.321526 | 1 |
| GOBP_ADHERENS_JUNCTION_ORGANIZATION | 65 | -0.379932 | -1.122142 | 0.269596 | 1 |
| GOBP_MITOTIC_SISTER_CHROMATID_SEGREGATION | 161 | -0.343882 | -1.121782 | 0.249732 | 1 |
| GOBP_REGULATION_OF_T_CELL_DIFFERENTIATION | 141 | -0.347103 | -1.120276 | 0.260259 | 1 |
| GOBP_NEGATIVE_REGULATION_OF_GENE_SILENCING | 31 | -0.427533 | -1.120053 | 0.303413 | 1 |
| GOBP_POSITIVE_REGULATION_OF_TRANSCRIPTION_INITIATION_FROM_RNA_P | 28 | -0.430508 | -1.119563 | 0.314433 | 1 |
| GOBP_PHOTORECEPTOR_CELL_DIFFERENTIATION | 59 | -0.383279 | -1.119336 | 0.290805 | 1 |
| GOBP_REGULATION_OF_CHROMATIN_ASSEMBLY | 19 | -0.468999 | -1.119086 | 0.325452 | 1 |
| GOBP_MODIFICATION_OF_SYNAPTIC_STRUCTURE | 19 | -0.479814 | -1.118509 | 0.354278 | 1 |
| GOBP_CEREBRAL_CORTEX_NEURON_DIFFERENTIATION | 19 | -0.463397 | -1.118448 | 0.338667 | 1 |
| GOBP_KILLING_OF_CELLS_OF_OTHER_ORGANISM | 43 | -0.404859 | -1.118432 | 0.313253 | 1 |
| GOBP_REGULATION_OF_LYMPHOCYTE_APOPTOTIC_PROCESS | 52 | -0.390898 | -1.118255 | 0.296206 | 1 |
| GOBP_TRIGLYCERIDE_METABOLIC_PROCESS | 91 | -0.362711 | -1.117332 | 0.272418 | 1 |
| GOBP_REGULATION_OF_POSTSYNAPTIC_MEMBRANE_POTENTIAL | 126 | -0.353121 | -1.11707 | 0.257525 | 1 |
| GOBP_POSITIVE_REGULATION_OF_SODIUM_ION_TRANSMEMBRANE_TRANSPO | 16 | -0.477677 | -1.116032 | 0.329773 | 1 |
| GOBP_INTERFERON_GAMMA_PRODUCTION | 107 | -0.355724 | -1.115857 | 0.262092 | 1 |
| GOBP_KERATINOCYTE_PROLIFERATION | 42 | -0.406429 | -1.115771 | 0.308733 | 1 |
| GOBP_SENSORY_PERCEPTION_OF_LIGHT_STIMULUS | 190 | -0.343538 | -1.115698 | 0.256545 | 1 |
| GOBP_MONONUCLEAR_CELL_DIFFERENTIATION | 384 | -0.326833 | -1.114921 | 0.186492 | 1 |
| GOBP_NADP_METABOLIC_PROCESS | 34 | -0.41272 | -1.114685 | 0.321883 | 1 |
| GOBP_POSITIVE_REGULATION_OF_DEFENSE_RESPONSE | 340 | -0.329177 | -1.114613 | 0.210366 | 1 |
| GOBP_CELLULAR_RESPONSE_TO_MOLECULE_OF_BACTERIAL_ORIGIN | 184 | -0.339144 | -1.114197 | 0.249218 | 1 |
| GOBP_RESPONSE_TO_MINERALOCORTICOID | 30 | -0.428717 | -1.11415 | 0.326772 | 1 |
| GOBP_PHOSPHATIDYLCHOLINE_METABOLIC_PROCESS | 86 | -0.364469 | -1.112968 | 0.288268 | 1 |
| GOBP_MRNA_TRANSCRIPTION | 15 | -0.489746 | -1.112143 | 0.348966 | 1 |
| GOBP_SPERM_MOTILITY | 86 | -0.360948 | -1.111893 | 0.276836 | 1 |
| GOBP_HEART_FIELD_SPECIFICATION | 18 | -0.472415 | -1.111794 | 0.341789 | 1 |
| GOBP_PROTEIN_TETRAMERIZATION | 83 | -0.363994 | -1.111732 | 0.282976 | 1 |
| GOBP_POSITIVE_REGULATION_OF_MONONUCLEAR_CELL_MIGRATION | 61 | -0.380424 | -1.110575 | 0.313167 | 1 |
| GOBP_BARBED_END_ACTIN_FILAMENT_CAPPING | 26 | -0.429623 | -1.11004 | 0.313078 | 1 |
| GOBP_SPECIFICATION_OF_ANIMAL_ORGAN_IDENTITY | 34 | -0.424717 | -1.110016 | 0.323641 | 1 |
| GOBP_POSITIVE_REGULATION_OF_MEMBRANE_POTENTIAL | 15 | -0.485128 | -1.109958 | 0.349501 | 1 |
| GOBP_NUCLEAR_MIGRATION | 19 | -0.46204 | -1.108662 | 0.341096 | 1 |
| GOBP_LIVER_REGENERATION | 30 | -0.421943 | -1.108503 | 0.335032 | 1 |
| GOBP_SIALYLATION | 19 | -0.458262 | -1.107303 | 0.33427 | 1 |
| GOBP_PROTEIN_AUTOPROCESSING | 23 | -0.447402 | -1.106381 | 0.357616 | 1 |
| GOBP_NEGATIVE_REGULATION_OF_B_CELL_ACTIVATION | 33 | -0.41084 | -1.106075 | 0.316794 | 1 |
| GOBP_REGULATION_OF_LYMPHOCYTE_DIFFERENTIATION | 169 | -0.337599 | -1.105232 | 0.272825 | 1 |
| GOBP_NEGATIVE_REGULATION_OF_NUCLEAR_DIVISION | 53 | -0.380166 | -1.105121 | 0.323208 | 1 |
| GOBP_NEGATIVE_REGULATION_OF_DNA_REPLICATION | 32 | -0.419037 | -1.102759 | 0.335878 | 1 |
| GOBP_INTEGRIN_ACTIVATION | 25 | -0.429334 | -1.101781 | 0.341207 | 1 |
| GOBP_NEGATIVE_REGULATION_OF_CELL_ACTIVATION | 190 | -0.335524 | -1.10138 | 0.273486 | 1 |
| GOBP_NEGATIVE_REGULATION_OF_LEUKOCYTE_MEDIATED_CYTOTOXICITY | 22 | -0.446518 | -1.101005 | 0.334651 | 1 |
| GOBP_GLIAL_CELL_MIGRATION | 49 | -0.383145 | -1.100263 | 0.324455 | 1 |
| GOBP_POSITIVE_REGULATION_OF_CYCLIN_DEPENDENT_PROTEIN_KINASE_A | 33 | -0.419061 | -1.100194 | 0.338562 | 1 |
| GOBP_POSITIVE_REGULATION_OF_ALCOHOL_BIOSYNTHETIC_PROCESS | 19 | -0.455335 | -1.100008 | 0.355705 | 1 |
| GOBP_FATTY_ACID_BIOSYNTHETIC_PROCESS | 152 | -0.341959 | -1.099868 | 0.293869 | 1 |
| GOBP_REGULATION_OF_PLASMA_LIPOPROTEIN_PARTICLE_LEVELS | 81 | -0.362259 | -1.099426 | 0.310345 | 1 |
| GOBP_EMBRYONIC_CAMERA_TYPE_EYE_DEVELOPMENT | 33 | -0.410754 | -1.098956 | 0.330251 | 1 |
| GOBP_RESPONSE_TO_DOPAMINE | 71 | -0.37141 | -1.098295 | 0.328655 | 1 |
| GOBP_NUCLEOBASE_METABOLIC_PROCESS | 34 | -0.409802 | -1.098096 | 0.344916 | 1 |
| GOBP_ICOSANOID_SECRETION | 37 | -0.40403 | -1.097881 | 0.337612 | 1 |
| GOBP_REGULATION_OF_TOLL_LIKE_RECEPTOR_SIGNALING_PATHWAY | 71 | -0.371416 | -1.097503 | 0.306433 | 1 |
| GOBP_ATTACHMENT_OF_SPINDLE_MICROTUBULES_TO_KINETOCHORE | 35 | -0.415175 | -1.097313 | 0.330357 | 1 |
| GOBP_REGULATION_OF_TRANSCRIPTION_REGULATORY_REGION_DNA_BINDI | 48 | -0.388046 | -1.096469 | 0.325123 | 1 |
| GOBP_PROTEIN_LIPID_COMPLEX_SUBUNIT_ORGANIZATION | 47 | -0.386747 | -1.096094 | 0.353086 | 1 |
| GOBP_CALCIUM_ION_TRANSPORT_INTO_CYTOSOL | 147 | -0.339045 | -1.095994 | 0.296017 | 1 |
| GOBP_POSITIVE_REGULATION_OF_ACTIN_CYTOSKELETON_REORGANIZATION | 18 | -0.462834 | -1.095259 | 0.362637 | 1 |
| GOBP_REGULATION_OF_CARDIAC_CONDUCTION | 56 | -0.372977 | -1.095238 | 0.326805 | 1 |
| GOBP_POSITIVE_REGULATION_OF_LYMPHOCYTE_CHEMOTAXIS | 19 | -0.46187 | -1.094073 | 0.38069 | 1 |
| GOBP_CALCIUM_INDEPENDENT_CELL_CELL_ADHESION_VIA_PLASMA_MEMBI | 21 | -0.445078 | -1.093981 | 0.363762 | 1 |
| GOBP_CEREBRAL_CORTEX_CELL_MIGRATION | 38 | -0.399586 | -1.093958 | 0.347447 | 1 |
| GOBP_CELLULAR_RESPONSE_TO_BIOTIC_STIMULUS | 208 | -0.328235 | -1.093396 | 0.271784 | 1 |
| GOBP_T_CELL_PROLIFERATION | 190 | -0.331399 | -1.092912 | 0.282766 | 1 |
| GOBP_PHOTOTRANSDUCTION_VISIBLE_LIGHT | 31 | -0.416685 | -1.091455 | 0.341495 | 1 |
| GOBP_POSITIVE_REGULATION_OF_NATURAL_KILLER_CELL_MEDIATED_CYTO | 26 | -0.429252 | -1.090753 | 0.364597 | 1 |
| GOBP_PHOTOTRANSDUCTION | 48 | -0.380926 | -1.089656 | 0.35169 | 1 |
| GOBP_REGULATION_OF_TRANSPOSITION | 25 | -0.426103 | -1.089276 | 0.362205 | 1 |
| GOBP_POSITIVE_REGULATION_OF_RESPONSE_TO_BIOTIC_STIMULUS | 243 | -0.327938 | -1.088884 | 0.290589 | 1 |
| GOBP_MAGNESIUM_ION_TRANSPORT | 17 | -0.460956 | -1.087006 | 0.366985 | 1 |
| GOBP_REGULATION_OF_SIGNALING_RECEPTOR_ACTIVITY | 169 | -0.336002 | -1.086462 | 0.295671 | 1 |
| GOBP_REGULATION_OF_LIPASE_ACTIVITY | 89 | -0.349156 | -1.085218 | 0.322652 | 1 |
| GOBP_ACUTE_PHASE_RESPONSE | 37 | -0.401735 | -1.085197 | 0.381313 | 1 |
| GOBP_REGULATION_OF_TRIGLYCERIDE_BIOSYNTHETIC_PROCESS | 18 | -0.458624 | -1.085127 | 0.357045 | 1 |
| GOBP_HOMOLOGOUS_CHROMOSOME_SEGREGATION | 54 | -0.374707 | -1.085082 | 0.355746 | 1 |
| GOBP_PROTEIN_LIPID_COMPLEX_ASSEMBLY | 29 | -0.418822 | -1.085046 | 0.369509 | 1 |
| GOBP_MULTICELLULAR_ORGANISM_AGING | 30 | -0.407608 | -1.084382 | 0.356129 | 1 |
| GOBP_DIGESTION | 111 | -0.346754 | -1.084334 | 0.319429 | 1 |
| GOBP_FATTY_ACID_METABOLIC_PROCESS | 370 | -0.321769 | -1.083846 | 0.252546 | 1 |
| GOBP_REGULATION_OF_CHROMOSOME_SEGREGATION | 83 | -0.356016 | -1.082439 | 0.340254 | 1 |
| GOBP_BETA_CATENIN_DESTRUCTION_COMPLEX_DISASSEMBLY | 20 | -0.440438 | -1.082159 | 0.379177 | 1 |
| GOBP_LEUKOCYTE_PROLIFERATION | 289 | -0.324755 | -1.08211 | 0.295918 | 1 |
| GOBP_TRANSEPITHELIAL_TRANSPORT | 30 | -0.417908 | -1.082102 | 0.364917 | 1 |
| GOBP_REPLICATION_FORK_PROCESSING | 39 | -0.396129 | -1.081503 | 0.359356 | 1 |
| GOBP_POSITIVE_REGULATION_OF_CELL_CELL_ADHESION | 269 | -0.321155 | -1.081231 | 0.295015 | 1 |
| GOBP_CYTOSOLIC_CALCIUM_ION_TRANSPORT | 168 | -0.332056 | -1.080871 | 0.294055 | 1 |
| GOBP_FORMATION_OF_EXTRACHROMOSOMAL_CIRCULAR_DNA | 15 | -0.464013 | -1.080843 | 0.373297 | 1 |
| GOBP_RESPIRATORY_BURST | 34 | -0.403375 | -1.079923 | 0.357326 | 1 |
| GOBP_CARTILAGE_DEVELOPMENT_INVOLVED_IN_ENDOCHONDRAL_BONE_M | 25 | -0.421224 | -1.079505 | 0.381016 | 1 |
| GOBP_SEQUESTERING_OF_CALCIUM_ION | 115 | -0.338602 | -1.078671 | 0.329658 | 1 |
| GOBP_KINETOCHORE_ASSEMBLY | 17 | -0.458434 | -1.078262 | 0.402817 | 1 |

| | | | | | |
|---|---|---|---|---|---|
| GOBP_REGULATION_OF_HETEROTYPIC_CELL_CELL_ADHESION | 26 | 0.357604 | 1.080028 | 0.317269 | 0.874358 |
| GOBP_CARDIAC_MUSCLE_CELL_ACTION_POTENTIAL_INVOLVED_IN_CO | 51 | 0.313008 | 1.080022 | 0.304813 | 0.872776 |
| GOBP_COLLECTING_DUCT_DEVELOPMENT | 16 | 0.385561 | 1.079974 | 0.363971 | 0.871414 |
| GOBP_NEGATIVE_REGULATION_OF_VIRAL_GENOME_REPLICATION | 52 | 0.294589 | 1.078962 | 0.273333 | 0.873886 |
| GOBP_NEGATIVE_REGULATION_OF_ORGANIC_ACID_TRANSPORT | 20 | 0.371245 | 1.078646 | 0.357143 | 0.87355 |
| GOBP_LIPOPROTEIN_BIOSYNTHETIC_PROCESS | 93 | 0.277214 | 1.078227 | 0.265957 | 0.873527 |
| GOBP_POSITIVE_REGULATION_OF_ENDOCYTOSIS | 97 | 0.271412 | 1.078211 | 0.276191 | 0.872024 |
| GOBP_EXTRINSIC_APOPTOTIC_SIGNALING_PATHWAY_VIA_DEATH_DOM | 81 | 0.281535 | 1.078205 | 0.260163 | 0.870471 |
| GOBP_ORGANIC_HYDROXY_COMPOUND_TRANSPORT | 246 | 0.245984 | 1.077415 | 0.107143 | 0.871841 |
| GOBP_NEGATIVE_REGULATION_OF_OSTEOCLAST_DIFFERENTIATION | 27 | 0.346605 | 1.076496 | 0.321429 | 0.873559 |
| GOBP_RESPONSE_TO_MUSCLE_ACTIVITY | 24 | 0.350665 | 1.075849 | 0.350195 | 0.874401 |
| GOBP_REGULATION_OF_FAT_CELL_DIFFERENTIATION | 125 | 0.260657 | 1.075285 | 0.227848 | 0.875051 |
| GOBP_REGULATION_OF_LIPOPROTEIN_METABOLIC_PROCESS | 15 | 0.399927 | 1.073766 | 0.380471 | 0.879187 |
| GOBP_REGULATION_OF_NEURAL_PRECURSOR_CELL_PROLIFERATION | 78 | 0.274366 | 1.072739 | 0.275591 | 0.881407 |
| GOBP_REGULATION_OF_EPITHELIAL_CELL_DIFFERENTIATION | 135 | 0.255014 | 1.072684 | 0.25 | 0.880052 |
| GOBP_REGULATION_OF_NEUROTRANSMITTER_UPTAKE | 15 | 0.395781 | 1.071317 | 0.383803 | 0.883875 |
| GOBP_NEGATIVE_REGULATION_OF_GROWTH | 235 | 0.237198 | 1.07081 | 0.242424 | 0.884303 |
| GOBP_RESPONSE_TO_CARBOHYDRATE | 203 | 0.239033 | 1.070476 | 0.212766 | 0.884045 |
| GOBP_DETECTION_OF_STIMULUS_INVOLVED_IN_SENSORY_PERCEPTIO | 24 | 0.359474 | 1.069205 | 0.34188 | 0.887481 |
| GOBP_CELLULAR_RESPONSE_TO_UV | 86 | 0.278434 | 1.068726 | 0.304348 | 0.887635 |
| GOBP_DETECTION_OF_LIGHT_STIMULUS_INVOLVED_IN_SENSORY_PER | 18 | 0.386992 | 1.068631 | 0.367041 | 0.88643 |
| GOBP_EAR_MORPHOGENESIS | 104 | 0.263893 | 1.066951 | 0.298851 | 0.891377 |
| GOBP_AORTA_DEVELOPMENT | 51 | 0.307006 | 1.06664 | 0.342391 | 0.891005 |
| GOBP_REGULATION_OF_COLLATERAL_SPROUTING | 17 | 0.382705 | 1.066284 | 0.399254 | 0.890892 |
| GOBP_POSITIVE_REGULATION_OF_VASCULAR_ENDOTHELIAL_GROWTH_ | 27 | 0.350665 | 1.065943 | 0.346154 | 0.890702 |
| GOBP_RESPONSE_TO_MAGNESIUM_ION | 17 | 0.379117 | 1.065811 | 0.337884 | 0.889648 |
| GOBP_MITOCHONDRIAL_DEPOLARIZATION | 22 | 0.365203 | 1.065256 | 0.365462 | 0.890307 |
| GOBP_CGMP_MEDIATED_SIGNALING | 25 | 0.34592 | 1.065028 | 0.354167 | 0.889636 |
| GOBP_REGULATION_OF_BODY_FLUID_LEVELS | 462 | 0.234479 | 1.065019 | 0 | 0.888136 |
| GOBP_BINDING_OF_SPERM_TO_ZONA_PELLUCIDA | 37 | 0.313718 | 1.064856 | 0.334783 | 0.887183 |
| GOBP_CARDIAC_MYOFIBRIL_ASSEMBLY | 16 | 0.40019 | 1.064323 | 0.373239 | 0.887641 |
| GOBP_BIOLOGICAL_PROCESS_INVOLVED_IN_INTERACTION_WITH_SYMB | 79 | 0.27884 | 1.064293 | 0.318841 | 0.886261 |
| GOBP_LIPID_EXPORT_FROM_CELL | 79 | 0.280018 | 1.06409 | 0.284615 | 0.885532 |
| GOBP_ACROSOME_REACTION | 33 | 0.325154 | 1.064028 | 0.335025 | 0.884218 |
| GOBP_RESPONSE_TO_STEROID_HORMONE | 311 | 0.231706 | 1.063828 | 0.142857 | 0.883508 |
| GOBP_NEGATIVE_REGULATION_OF_EPITHELIAL_CELL_DIFFERENTIATION | 39 | 0.307273 | 1.062713 | 0.358852 | 0.886091 |
| GOBP_REGULATION_OF_CELL_MATRIX_ADHESION | 117 | 0.25974 | 1.061773 | 0.247312 | 0.88827 |
| GOBP_REGULATION_OF_PEPTIDASE_ACTIVITY | 414 | 0.231089 | 1.06172 | 0.166667 | 0.886972 |
| GOBP_REGULATION_OF_CARDIAC_MUSCLE_CONTRACTION | 70 | 0.280276 | 1.060866 | 0.323308 | 0.889001 |
| GOBP_REGULATION_OF_HEART_RATE | 91 | 0.262651 | 1.060077 | 0.377551 | 0.890566 |
| GOBP_WOUND_HEALING | 485 | 0.216756 | 1.059706 | 0 | 0.890476 |
| GOBP_NEGATIVE_REGULATION_OF_BLOOD_CIRCULATION | 21 | 0.35613 | 1.059421 | 0.335821 | 0.890101 |
| GOBP_RENAL_ABSORPTION | 18 | 0.375296 | 1.059341 | 0.360294 | 0.888922 |
| GOBP_EPITHELIAL_STRUCTURE_MAINTENANCE | 27 | 0.341164 | 1.059 | 0.345833 | 0.888753 |
| GOBP_REGULATION_OF_NEURON_MIGRATION | 37 | 0.320111 | 1.05867 | 0.357143 | 0.888511 |
| GOBP_REGULATION_OF_RESPONSE_TO_REACTIVE_OXYGEN_SPECIES | 30 | 0.333805 | 1.057339 | 0.393013 | 0.892395 |
| GOBP_NEUROBLAST_PROLIFERATION | 52 | 0.297429 | 1.05715 | 0.324742 | 0.891634 |
| GOBP_REGULATION_OF_DEVELOPMENTAL_GROWTH | 296 | 0.234854 | 1.055463 | 0.333333 | 0.896384 |
| GOBP_RESPONSE_TO_NERVE_GROWTH_FACTOR | 50 | 0.300286 | 1.055284 | 0.361582 | 0.895515 |
| GOBP_REGULATION_OF_CAMP_DEPENDENT_PROTEIN_KINASE_ACTIVIT | 20 | 0.375916 | 1.054585 | 0.358108 | 0.896642 |
| GOBP_MALE_SEX_DIFFERENTIATION | 143 | 0.253758 | 1.054499 | 0.271429 | 0.895435 |
| GOBP_MODULATION_OF_EXCITATORY_POSTSYNAPTIC_POTENTIAL | 43 | 0.306604 | 1.054207 | 0.356383 | 0.894999 |
| GOBP_REGULATION_OF_ENDOCRINE_PROCESS | 32 | 0.337781 | 1.053353 | 0.358079 | 0.896622 |
| GOBP_NEGATIVE_REGULATION_OF_CELL_MATRIX_ADHESION | 36 | 0.323987 | 1.053244 | 0.377359 | 0.895568 |
| GOBP_EPITHELIAL_TUBE_MORPHOGENESIS | 298 | 0.23146 | 1.053017 | 0.2 | 0.894936 |
| GOBP_GLOBAL_GENOME_NUCLEOTIDE_EXCISION_REPAIR | 26 | 0.340616 | 1.052955 | 0.379747 | 0.893718 |
| GOBP_EAR_DEVELOPMENT | 189 | 0.245022 | 1.051875 | 0.289474 | 0.896266 |
| GOBP_PLASMINOGEN_ACTIVATION | 20 | 0.363701 | 1.050899 | 0.348148 | 0.898438 |
| GOBP_INOSITOL_PHOSPHATE_MEDIATED_SIGNALING | 52 | 0.296134 | 1.050822 | 0.347561 | 0.897242 |
| GOBP_BICARBONATE_TRANSPORT | 43 | 0.311424 | 1.050097 | 0.34359 | 0.898684 |
| GOBP_POSITIVE_REGULATION_OF_EXTRINSIC_APOPTOTIC_SIGNALING_ | 46 | 0.308741 | 1.050048 | 0.36413 | 0.89738 |
| GOBP_MYOBLAST_PROLIFERATION | 17 | 0.387743 | 1.049591 | 0.40604 | 0.897729 |
| GOBP_POSITIVE_REGULATION_OF_PEPTIDE_HORMONE_SECRETION | 87 | 0.266705 | 1.049038 | 0.284553 | 0.898241 |
| GOBP_REGULATION_OF_PROTEIN_LOCALIZATION_TO_SYNAPSE | 26 | 0.344062 | 1.048285 | 0.390909 | 0.899571 |
| GOBP_VASOCONSTRICTION | 73 | 0.271295 | 1.047971 | 0.353742 | 0.899411 |
| GOBP_STEROID_HORMONE_MEDIATED_SIGNALING_PATHWAY | 128 | 0.249897 | 1.047459 | 0.369048 | 0.899936 |
| GOBP_NEGATIVE_REGULATION_OF_TRANSPORT | 388 | 0.224697 | 1.04744 | 0.272727 | 0.898562 |
| GOBP_NEGATIVE_REGULATION_OF_HORMONE_SECRETION | 54 | 0.289674 | 1.047058 | 0.344633 | 0.898603 |
| GOBP_TONGUE_DEVELOPMENT | 19 | 0.367941 | 1.046098 | 0.381323 | 0.900839 |
| GOBP_NEGATIVE_REGULATION_OF_MYOBLAST_DIFFERENTIATION | 23 | 0.350533 | 1.045967 | 0.40249 | 0.899934 |
| GOBP_BONE_MINERALIZATION | 106 | 0.259853 | 1.045587 | 0.289474 | 0.900048 |
| GOBP_ATP_TRANSPORT | 16 | 0.39074 | 1.045411 | 0.378906 | 0.899303 |
| GOBP_RESPONSE_TO_ELECTRICAL_STIMULUS | 41 | 0.307527 | 1.044194 | 0.368687 | 0.902703 |
| GOBP_ENDOTHELIUM_DEVELOPMENT | 125 | 0.249755 | 1.042591 | 0.333333 | 0.907616 |
| GOBP_NEURAL_NUCLEUS_DEVELOPMENT | 56 | 0.281184 | 1.042143 | 0.341317 | 0.907914 |
| GOBP_VASCULOGENESIS | 76 | 0.266386 | 1.041771 | 0.375 | 0.907879 |
| GOBP_POSITIVE_REGULATION_OF_RUFFLE_ASSEMBLY | 15 | 0.386959 | 1.040853 | 0.390845 | 0.909911 |
| GOBP_POSITIVE_REGULATION_OF_LONG_TERM_SYNAPTIC_POTENTIATI | 22 | 0.362645 | 1.039937 | 0.351563 | 0.912119 |
| GOBP_POSITIVE_REGULATION_OF_DENDRITIC_SPINE_DEVELOPMENT | 42 | 0.303018 | 1.037475 | 0.37 | 0.920073 |
| GOBP_REGULATION_OF_DNA_DAMAGE_RESPONSE_SIGNAL_TRANSDUC | 34 | 0.328627 | 1.037164 | 0.410526 | 0.919849 |
| GOBP_PHARYNGEAL_SYSTEM_DEVELOPMENT | 23 | 0.34418 | 1.036835 | 0.40678 | 0.919626 |
| GOBP_RESPONSE_TO_EXTRACELLULAR_STIMULUS | 442 | 0.240654 | 1.03646 | 0.333333 | 0.919728 |
| GOBP_EMBRYONIC_ORGAN_DEVELOPMENT | 395 | 0.225819 | 1.03553 | 0.142857 | 0.921763 |
| GOBP_T_CELL_MEDIATED_CYTOTOXICITY | 46 | 0.294221 | 1.035167 | 0.404145 | 0.921663 |
| GOBP_POSITIVE_REGULATION_OF_EXCITATORY_POSTSYNAPTIC_POTEN | 27 | 0.331715 | 1.034952 | 0.407407 | 0.92101 |
| GOBP_ANTIGEN_PROCESSING_AND_PRESENTATION_OF_EXOGENOUS_ | 79 | 0.268315 | 1.034867 | 0.397059 | 0.919988 |
| GOBP_WATER_HOMEOSTASIS | 75 | 0.270079 | 1.034304 | 0.377953 | 0.920846 |
| GOBP_SYNCYTIUM_FORMATION | 54 | 0.277029 | 1.033172 | 0.396104 | 0.923639 |
| GOBP_COAGULATION | 313 | 0.218209 | 1.033041 | 0.230769 | 0.922678 |
| GOBP_MYOTUBE_CELL_DEVELOPMENT | 31 | 0.323662 | 1.032708 | 0.375546 | 0.922502 |
| GOBP_NEGATIVE_REGULATION_OF_TRANSMEMBRANE_RECEPTOR_PRO | 122 | 0.252492 | 1.032507 | 0.352941 | 0.92178 |
| GOBP_POSITIVE_REGULATION_OF_NEURON_DEATH | 90 | 0.263591 | 1.032502 | 0.354545 | 0.920357 |
| GOBP_CARBOHYDRATE_DERIVATIVE_CATABOLIC_PROCESS | 188 | 0.237233 | 1.032161 | 0.346939 | 0.920177 |
| GOBP_CARDIAC_MUSCLE_CELL_ACTION_POTENTIAL | 71 | 0.279234 | 1.031945 | 0.345912 | 0.919564 |
| GOBP_POSITIVE_REGULATION_OF_SYNAPSE_ASSEMBLY | 56 | 0.291045 | 1.031752 | 0.388535 | 0.918883 |
| GOBP_GENITALIA_DEVELOPMENT | 40 | 0.297673 | 1.031674 | 0.395939 | 0.917759 |
| GOBP_COLLATERAL_SPROUTING | 23 | 0.334894 | 1.031042 | 0.390625 | 0.918769 |
| GOBP_PATTERN_SPECIFICATION_PROCESS | 394 | 0.227664 | 1.030549 | 0.125 | 0.919326 |
| GOBP_POSITIVE_REGULATION_OF_MUSCLE_HYPERTROPHY | 26 | 0.342809 | 1.030012 | 0.373391 | 0.91997 |
| GOBP_CELL_PROLIFERATION_INVOLVED_IN_KIDNEY_DEVELOPMENT | 19 | 0.372585 | 1.03001 | 0.359259 | 0.918557 |
| GOBP_OUTFLOW_TRACT_MORPHOGENESIS | 71 | 0.279655 | 1.029717 | 0.333333 | 0.918307 |
| GOBP_EMBRYONIC_PLACENTA_MORPHOGENESIS | 24 | 0.340993 | 1.029494 | 0.395556 | 0.91777 |
| GOBP_POSITIVE_REGULATION_OF_BONE_MINERALIZATION | 38 | 0.312196 | 1.02932 | 0.391534 | 0.916986 |
| GOBP_CARDIAC_CHAMBER_DEVELOPMENT | 148 | 0.244768 | 1.028711 | 0.337838 | 0.91793 |
| GOBP_BRANCHING_INVOLVED_IN_BLOOD_VESSEL_MORPHOGENESIS | 31 | 0.317025 | 1.028467 | 0.451923 | 0.917516 |
| GOBP_ENDOCRINE_PANCREAS_DEVELOPMENT | 38 | 0.313504 | 1.028425 | 0.402985 | 0.916227 |
| GOBP_FEMALE_SEX_DIFFERENTIATION | 105 | 0.264548 | 1.028259 | 0.404255 | 0.915444 |
| GOBP_NEGATIVE_REGULATION_OF_CELL_SUBSTRATE_JUNCTION_ORG | 18 | 0.361723 | 1.027973 | 0.410909 | 0.91518 |
| GOBP_MUSCLE_CELL_CELLULAR_HOMEOSTASIS | 20 | 0.354146 | 1.027684 | 0.40856 | 0.91484 |
| GOBP_UBIQUITIN_DEPENDENT_PROTEIN_CATABOLIC_PROCESS_VIA_TH | 23 | 0.347839 | 1.026682 | 0.416 | 0.91745 |
| GOBP_RESPONSE_TO_TESTOSTERONE | 40 | 0.309528 | 1.026436 | 0.376289 | 0.917027 |
| GOBP_MEMBRANE_PROTEIN_PROTEOLYSIS | 61 | 0.275858 | 1.025386 | 0.409722 | 0.919755 |
| GOBP_INSULIN_SECRETION | 193 | 0.238193 | 1.025046 | 0.297297 | 0.91961 |
| GOBP_RESPONSE_TO_TOXIC_SUBSTANCE | 224 | 0.236402 | 1.024726 | 0.384615 | 0.919399 |
| GOBP_REGULATION_OF_PEPTIDYL_SERINE_PHOSPHORYLATION | 124 | 0.247031 | 1.024527 | 0.348315 | 0.9188 |
| GOBP_RRNA_MODIFICATION | 40 | 0.30519 | 1.023149 | 0.407609 | 0.922657 |
| GOBP_MESENCHYME_DEVELOPMENT | 269 | 0.226049 | 1.023049 | 0.333333 | 0.921687 |
| GOBP_ACIDIC_AMINO_ACID_TRANSPORT | 65 | 0.275001 | 1.022323 | 0.413793 | 0.922998 |
| GOBP_REGULATION_OF_INTRINSIC_APOPTOTIC_SIGNALING_PATHWAY_ | 17 | 0.372442 | 1.022288 | 0.429091 | 0.92173 |
| GOBP_CELLULAR_RESPONSE_TO_VITAMIN_D | 19 | 0.361991 | 1.022245 | 0.414449 | 0.92053 |
| GOBP_CD4_POSITIVE_ALPHA_BETA_T_CELL_CYTOKINE_PRODUCTION | 17 | 0.355273 | 1.021381 | 0.410448 | 0.92238 |
| GOBP_CARDIAC_MUSCLE_CELL_MEMBRANE_REPOLARIZATION | 34 | 0.305742 | 1.020106 | 0.371041 | 0.925766 |
| GOBP_PODOSOME_ASSEMBLY | 16 | 0.376351 | 1.019634 | 0.418327 | 0.926244 |
| GOBP_T_CELL_CYTOKINE_PRODUCTION | 35 | 0.307614 | 1.019232 | 0.430493 | 0.926328 |
| GOBP_DICARBOXYLIC_ACID_TRANSPORT | 90 | 0.25833 | 1.016681 | 0.411765 | 0.934463 |
| GOBP_POSITIVE_REGULATION_OF_CELL_JUNCTION_ASSEMBLY | 92 | 0.25124 | 1.016039 | 0.406015 | 0.935498 |
| GOBP_NEGATIVE_REGULATION_OF_LIPID_BIOSYNTHETIC_PROCESS | 50 | 0.287929 | 1.015328 | 0.411043 | 0.936756 |
| GOBP_OLIGODENDROCYTE_DEVELOPMENT | 40 | 0.293263 | 1.01514 | 0.419811 | 0.936093 |
| GOBP_AMINE_BIOSYNTHETIC_PROCESS | 36 | 0.311082 | 1.015121 | 0.42723 | 0.934767 |
| GOBP_STEROID_HORMONE_SECRETION | 20 | 0.347088 | 1.014861 | 0.409449 | 0.934301 |
| GOBP_REGULATION_OF_CELLULAR_RESPONSE_TO_HEAT | 79 | 0.265731 | 1.014207 | 0.442478 | 0.935466 |
| GOBP_INTRINSIC_APOPTOTIC_SIGNALING_PATHWAY_IN_RESPONSE_TO | 44 | 0.296828 | 1.013174 | 0.4 | 0.938198 |
| GOBP_REGULATION_OF_INSULIN_RECEPTOR_SIGNALING_PATHWAY | 63 | 0.273162 | 1.013126 | 0.43125 | 0.937011 |
| GOBP_NEGATIVE_REGULATION_OF_TUMOR_NECROSIS_FACTOR_MEDIA | 20 | 0.35279 | 1.012836 | 0.425197 | 0.936826 |
| GOBP_HEPARAN_SULFATE_PROTEOGLYCAN_METABOLIC_PROCESS | 31 | 0.322061 | 1.012645 | 0.435185 | 0.936129 |
| GOBP_POSITIVE_REGULATION_OF_PHOSPHATIDYLINOSITOL_3_KINASE_ | 87 | 0.258306 | 1.012213 | 0.411765 | 0.936322 |
| GOBP_CARDIAC_CHAMBER_MORPHOGENESIS | 112 | 0.250796 | 1.011511 | 0.468085 | 0.937615 |
| GOBP_PEROXISOME_PROLIFERATOR_ACTIVATED_RECEPTOR_SIGNALIN | 20 | 0.356791 | 1.011417 | 0.392226 | 0.936557 |

| | | | | | |
|---|---|---|---|---|---|
| GOBP_CRANIAL_SKELETAL_SYSTEM_DEVELOPMENT | 62 | -0.367905 | -1.077795 | 0.362471 | 1 |
| GOBP_DEVELOPMENTAL_MATURATION | 255 | -0.319222 | -1.07743 | 0.305556 | 1 |
| GOBP_T_CELL_DIFFERENTIATION_IN_THYMUS | 71 | -0.358198 | -1.077213 | 0.341601 | 1 |
| GOBP_CELLULAR_PROTEIN_CONTAINING_COMPLEX_LOCALIZATION | 20 | -0.436277 | -1.076918 | 0.38388 | 1 |
| GOBP_MATING | 34 | -0.405432 | -1.076507 | 0.376263 | 1 |
| GOBP_TISSUE_HOMEOSTASIS | 252 | -0.321338 | -1.075676 | 0.297631 | 1 |
| GOBP_PROTEIN_DNA_COMPLEX_SUBUNIT_ORGANIZATION | 250 | -0.320852 | -1.075663 | 0.305641 | 1 |
| GOBP_TELOMERE_CAPPING | 51 | -0.378814 | -1.075231 | 0.360437 | 1 |
| GOBP_PHOSPHATIDYLCHOLINE_ACYL_CHAIN_REMODELING | 29 | -0.424922 | -1.075031 | 0.379265 | 1 |
| GOBP_NEGATIVE_REGULATION_OF_MUSCLE_CONTRACTION | 20 | -0.440839 | -1.074692 | 0.384513 | 1 |
| GOBP_TELENCEPHALON_DEVELOPMENT | 218 | -0.325564 | -1.074484 | 0.309917 | 1 |
| GOBP_REGULATION_OF_LYMPHOCYTE_MEDIATED_IMMUNITY | 155 | -0.330101 | -1.074251 | 0.326923 | 1 |
| GOBP_NEGATIVE_REGULATION_OF_REACTIVE_OXYGEN_SPECIES_BIOSYNTHE | 27 | -0.417556 | -1.07424 | 0.363165 | 1 |
| GOBP_PHOSPHOLIPASE_C_ACTIVATING_G_PROTEIN_COUPLED_RECEPTOR_S | 93 | -0.351963 | -1.073555 | 0.349658 | 1 |
| GOBP_VERY_LONG_CHAIN_FATTY_ACID_METABOLIC_PROCESS | 33 | -0.401392 | -1.073433 | 0.376774 | 1 |
| GOBP_AMINO_SUGAR_METABOLIC_PROCESS | 40 | -0.386296 | -1.073408 | 0.358846 | 1 |
| GOBP_CELL_MIGRATION_INVOLVED_IN_SPROUTING_ANGIOGENESIS | 61 | -0.369333 | -1.072321 | 0.358513 | 1 |
| GOBP_DNA_UNWINDING_INVOLVED_IN_DNA_REPLICATION | 16 | -0.460417 | -1.072231 | 0.391489 | 1 |
| GOBP_POSITIVE_REGULATION_OF_STEROID_BIOSYNTHETIC_PROCESS | 16 | -0.458814 | -1.072176 | 0.39834 | 1 |
| GOBP_OVULATION | 15 | -0.468639 | -1.072032 | 0.400279 | 1 |
| GOBP_GRANULOCYTE_MIGRATION | 140 | -0.332626 | -1.071843 | 0.338727 | 1 |
| GOBP_EYE_PHOTORECEPTOR_CELL_DIFFERENTIATION | 42 | -0.382714 | -1.071701 | 0.361858 | 1 |
| GOBP_CALCIUM_MEDIATED_SIGNALING | 192 | -0.327811 | -1.071469 | 0.32906 | 1 |
| GOBP_PHOSPHATIDYLSERINE_ACYL_CHAIN_REMODELING | 21 | -0.437585 | -1.070649 | 0.389474 | 1 |
| GOBP_MESODERM_DEVELOPMENT | 109 | -0.339838 | -1.070264 | 0.338445 | 1 |
| GOBP_CGMP_METABOLIC_PROCESS | 15 | -0.460144 | -1.069549 | 0.388579 | 1 |
| GOBP_POSITIVE_REGULATION_OF_DEPHOSPHORYLATION | 58 | -0.365839 | -1.069468 | 0.359375 | 1 |
| GOBP_IRON_ION_TRANSPORT | 75 | -0.353555 | -1.06917 | 0.356725 | 1 |
| GOBP_RESPONSE_TO_DIETARY_EXCESS | 22 | -0.430478 | -1.068929 | 0.401639 | 1 |
| GOBP_REGULATION_OF_LYMPHOCYTE_CHEMOTAXIS | 25 | -0.419711 | -1.067534 | 0.386842 | 1 |
| GOBP_POSITIVE_REGULATION_OF_CHROMOSOME_SEPARATION | 17 | -0.44559 | -1.067142 | 0.388664 | 1 |
| GOBP_TOLL_LIKE_RECEPTOR_2_SIGNALING_PATHWAY | 17 | -0.450337 | -1.067037 | 0.382979 | 1 |
| GOBP_SCHWANN_CELL_DIFFERENTIATION | 36 | -0.395969 | -1.066987 | 0.367949 | 1 |
| GOBP_TETRAHYDROFOLATE_METABOLIC_PROCESS | 19 | -0.438525 | -1.066858 | 0.394231 | 1 |
| GOBP_MYELOID_LEUKOCYTE_MIGRATION | 206 | -0.32217 | -1.066283 | 0.336809 | 1 |
| GOBP_REGULATION_OF_FATTY_ACID_TRANSPORT | 45 | -0.375682 | -1.065302 | 0.372149 | 1 |
| GOBP_REGULATION_OF_FATTY_ACID_METABOLIC_PROCESS | 81 | -0.353304 | -1.065065 | 0.377727 | 1 |
| GOBP_PROTEIN_O_LINKED_GLYCOSYLATION | 102 | -0.343782 | -1.064759 | 0.350838 | 1 |
| GOBP_RESPONSE_TO_PURINE_CONTAINING_COMPOUND | 141 | -0.331173 | -1.064742 | 0.350858 | 1 |
| GOBP_RESPONSE_TO_ESTRADIOL | 119 | -0.339745 | -1.064598 | 0.349241 | 1 |
| GOBP_NEGATIVE_REGULATION_OF_CHROMOSOME_ORGANIZATION | 89 | -0.344086 | -1.064126 | 0.349261 | 1 |
| GOBP_DETECTION_OF_STIMULUS_INVOLVED_IN_SENSORY_PERCEPTION | 121 | -0.337512 | -1.064109 | 0.349119 | 1 |
| GOBP_ENSHEATHMENT_OF_NEURONS | 122 | -0.336173 | -1.062644 | 0.37041 | 1 |
| GOBP_INTERLEUKIN_1_BETA_PRODUCTION | 90 | -0.343451 | -1.062025 | 0.348365 | 1 |
| GOBP_STEM_CELL_DIVISION | 24 | -0.428321 | -1.062017 | 0.391534 | 1 |
| GOBP_CALCIUM_MEDIATED_SIGNALING_USING_INTRACELLULAR_CALCIUM_S | 22 | -0.423967 | -1.061451 | 0.38992 | 1 |
| GOBP_REGULATION_OF_MULTICELLULAR_ORGANISM_GROWTH | 58 | -0.366093 | -1.060515 | 0.381455 | 1 |
| GOBP_DNA_REPLICATION_INITIATION | 40 | -0.385317 | -1.06049 | 0.40399 | 1 |
| GOBP_REGULATION_OF_NEUROTRANSMITTER_RECEPTOR_ACTIVITY | 72 | -0.357175 | -1.060481 | 0.381776 | 1 |
| GOBP_NEPHRON_TUBULE_EPITHELIAL_CELL_DIFFERENTIATION | 15 | -0.472945 | -1.060455 | 0.421277 | 1 |
| GOBP_RETINAL_METABOLIC_PROCESS | 15 | -0.455703 | -1.059615 | 0.403226 | 1 |
| GOBP_PYRIMIDINE_NUCLEOSIDE_METABOLIC_PROCESS | 34 | -0.392034 | -1.059254 | 0.394771 | 1 |
| GOBP_EMBRYONIC_DIGESTIVE_TRACT_MORPHOGENESIS | 16 | -0.45993 | -1.059232 | 0.391967 | 1 |
| GOBP_DIACYLGLYCEROL_METABOLIC_PROCESS | 24 | -0.418033 | -1.058872 | 0.407602 | 1 |
| GOBP_REGULATION_OF_CIRCADIAN_RHYTHM | 109 | -0.33638 | -1.058405 | 0.378855 | 1 |
| GOBP_REGULATION_OF_AMINO_ACID_TRANSPORT | 30 | -0.401352 | -1.058382 | 0.38896 | 1 |
| GOBP_REGULATION_OF_ALPHA_BETA_T_CELL_DIFFERENTIATION | 66 | -0.357454 | -1.058366 | 0.383546 | 1 |
| GOBP_ORGANELLE_FISSION | 471 | -0.310806 | -1.05794 | 0.306225 | 1 |
| GOBP_IMPORT_INTO_CELL | 221 | -0.319703 | -1.057907 | 0.35447 | 1 |
| GOBP_MRNA_SPLICE_SITE_SELECTION | 39 | -0.389534 | -1.05756 | 0.388005 | 1 |
| GOBP_DNA_MODIFICATION | 110 | -0.33755 | -1.057154 | 0.375691 | 1 |
| GOBP_REGULATION_OF_TUBE_SIZE | 119 | -0.333564 | -1.055999 | 0.370933 | 1 |
| GOBP_POSITIVE_REGULATION_OF_ISOTYPE_SWITCHING | 26 | -0.413872 | -1.05562 | 0.39922 | 1 |
| GOBP_POSITIVE_REGULATION_OF_ACTIN_FILAMENT_POLYMERIZATION | 97 | -0.341257 | -1.055598 | 0.386667 | 1 |
| GOBP_PYRIMIDINE_NUCLEOSIDE_BIOSYNTHETIC_PROCESS | 16 | -0.454742 | -1.054156 | 0.413199 | 1 |
| GOBP_REGULATION_OF_SMAD_PROTEIN_SIGNAL_TRANSDUCTION | 25 | -0.421661 | -1.054022 | 0.409449 | 1 |
| GOBP_STEROID_HORMONE_BIOSYNTHETIC_PROCESS | 46 | -0.37256 | -1.054003 | 0.385276 | 1 |
| GOBP_DETECTION_OF_TEMPERATURE_STIMULUS_INVOLVED_IN_SENSORY_F | 17 | -0.451181 | -1.053934 | 0.40819 | 1 |
| GOBP_TEMPERATURE_HOMEOSTASIS | 166 | -0.323305 | -1.051429 | 0.376997 | 1 |
| GOBP_FOREBRAIN_CELL_MIGRATION | 52 | -0.363755 | -1.051238 | 0.376513 | 1 |
| GOBP_NEURON_FATE_SPECIFICATION | 19 | -0.440939 | -1.050496 | 0.417687 | 1 |
| GOBP_RESPONSE_TO_CATECHOLAMINE | 89 | -0.343864 | -1.050264 | 0.400911 | 1 |
| GOBP_CELLULAR_RESPONSE_TO_PROSTAGLANDIN_E_STIMULUS | 15 | -0.457246 | -1.050201 | 0.424501 | 1 |
| GOBP_PALLIUM_DEVELOPMENT | 145 | -0.326797 | -1.049923 | 0.365282 | 1 |
| GOBP_POSITIVE_REGULATION_OF_SYNAPTIC_TRANSMISSION_GLUTAMATERG | 25 | -0.414865 | -1.048069 | 0.423338 | 1 |
| GOBP_REGULATION_OF_PH | 100 | -0.334112 | -1.047686 | 0.391354 | 1 |
| GOBP_CENTRAL_NERVOUS_SYSTEM_NEURON_DIFFERENTIATION | 164 | -0.3238 | -1.047511 | 0.393283 | 1 |
| GOBP_RIBOSOME_ASSEMBLY | 64 | -0.356044 | -1.046883 | 0.413018 | 1 |
| GOBP_POSITIVE_REGULATION_OF_RESPONSE_TO_EXTERNAL_STIMULUS | 475 | -0.307081 | -1.046436 | 0.349699 | 1 |
| GOBP_REGULATION_OF_INNATE_IMMUNE_RESPONSE | 301 | -0.311044 | -1.046192 | 0.373225 | 1 |
| GOBP_ORGANIC_ACID_BIOSYNTHETIC_PROCESS | 313 | -0.311714 | -1.045703 | 0.388379 | 1 |
| GOBP_POSITIVE_REGULATION_OF_LEUKOCYTE_PROLIFERATION | 143 | -0.325587 | -1.045165 | 0.403433 | 1 |
| GOBP_NEUTROPHIL_MIGRATION | 116 | -0.331483 | -1.045126 | 0.391494 | 1 |
| GOBP_DNA_REPLICATION | 276 | -0.311558 | -1.044863 | 0.383044 | 1 |
| GOBP_LEUKOCYTE_DIFFERENTIATION | 493 | -0.307171 | -1.044491 | 0.361361 | 1 |
| GOBP_POSITIVE_REGULATION_OF_TOLL_LIKE_RECEPTOR_SIGNALING_PATHW | 27 | -0.410369 | -1.044281 | 0.434371 | 1 |
| GOBP_REGULATION_OF_LEUKOCYTE_MEDIATED_CYTOTOXICITY | 79 | -0.349198 | -1.044124 | 0.396119 | 1 |
| GOBP_NEUROTRANSMITTER_UPTAKE | 36 | -0.387375 | -1.043363 | 0.396947 | 1 |
| GOBP_NEGATIVE_REGULATION_OF_BLOOD_PRESSURE | 35 | -0.393475 | -1.043221 | 0.427848 | 1 |
| GOBP_LEUKOCYTE_HOMEOSTASIS | 78 | -0.344157 | -1.043089 | 0.396119 | 1 |
| GOBP_CAMERA_TYPE_EYE_PHOTORECEPTOR_CELL_DIFFERENTIATION | 25 | -0.413387 | -1.041677 | 0.411917 | 1 |
| GOBP_PROTEIN_LOCALIZATION_TO_KINETOCHORE | 19 | -0.434443 | -1.040286 | 0.409524 | 1 |
| GOBP_POSITIVE_REGULATION_OF_LEUKOCYTE_CHEMOTAXIS | 88 | -0.336609 | -1.040021 | 0.403429 | 1 |
| GOBP_CEREBRAL_CORTEX_RADIALLY_ORIENTED_CELL_MIGRATION | 29 | -0.404148 | -1.039799 | 0.439206 | 1 |
| GOBP_STEROID_ESTERIFICATION | 16 | -0.446636 | -1.0387 | 0.416785 | 1 |
| GOBP_CELLULAR_RESPONSE_TO_ALCOHOL | 82 | -0.343771 | -1.038154 | 0.41055 | 1 |
| GOBP_PEPTIDE_CROSS_LINKING | 33 | -0.390617 | -1.037863 | 0.426938 | 1 |
| GOBP_RESPONSE_TO_OSMOTIC_STRESS | 79 | -0.3422 | -1.037507 | 0.431235 | 1 |
| GOBP_CARBOXYLIC_ACID_TRANSPORT | 260 | -0.308478 | -1.037341 | 0.398349 | 1 |
| GOBP_POSITIVE_REGULATION_OF_MITOTIC_NUCLEAR_DIVISION | 41 | -0.379987 | -1.03729 | 0.427307 | 1 |
| GOBP_RESPONSE_TO_PROGESTERONE | 44 | -0.368526 | -1.03718 | 0.415347 | 1 |
| GOBP_MAMMARY_GLAND_EPITHELIAL_CELL_PROLIFERATION | 26 | -0.406167 | -1.036886 | 0.439216 | 1 |
| GOBP_SENSORY_PERCEPTION_OF_BITTER_TASTE | 24 | -0.411058 | -1.0368 | 0.405622 | 1 |
| GOBP_POSITIVE_REGULATION_OF_LIPID_TRANSPORT | 80 | -0.345433 | -1.036693 | 0.39627 | 1 |
| GOBP_REGULATION_OF_NEURON_DIFFERENTIATION | 174 | -0.316128 | -1.0366 | 0.388771 | 1 |
| GOBP_LEUKOCYTE_MEDIATED_CYTOTOXICITY | 110 | -0.32843 | -1.036259 | 0.419248 | 1 |
| GOBP_RETINAL_GANGLION_CELL_AXON_GUIDANCE | 18 | -0.437279 | -1.036244 | 0.456284 | 1 |
| GOBP_POSITIVE_REGULATION_OF_FATTY_ACID_BIOSYNTHETIC_PROCESS | 19 | -0.432351 | -1.035207 | 0.44337 | 1 |
| GOBP_BRAIN_MORPHOGENESIS | 34 | -0.385105 | -1.03354 | 0.415929 | 1 |
| GOBP_LEUKOCYTE_CELL_CELL_ADHESION | 347 | -0.303943 | -1.033378 | 0.39899 | 1 |
| GOBP_MESODERMAL_CELL_FATE_COMMITMENT | 16 | -0.45103 | -1.033197 | 0.465565 | 1 |
| GOBP_RESPONSE_TO_MURAMYL_DIPEPTIDE | 20 | -0.424255 | -1.032752 | 0.415323 | 1 |
| GOBP_MITOCHONDRIAL_RNA_METABOLIC_PROCESS | 47 | -0.363866 | -1.032408 | 0.43617 | 1 |
| GOBP_B_CELL_HOMEOSTASIS | 28 | -0.400872 | -1.031203 | 0.440053 | 1 |
| GOBP_POSITIVE_REGULATION_OF_RHO_PROTEIN_SIGNAL_TRANSDUCTION | 33 | -0.386032 | -1.031028 | 0.418367 | 1 |
| GOBP_SYNAPTIC_TRANSMISSION_DOPAMINERGIC | 21 | -0.423235 | -1.03094 | 0.431507 | 1 |
| GOBP_SISTER_CHROMATID_SEGREGATION | 194 | -0.315158 | -1.030806 | 0.422756 | 1 |
| GOBP_ICOSANOID_TRANSPORT | 45 | -0.369652 | -1.030605 | 0.424354 | 1 |
| GOBP_IMPORT_ACROSS_PLASMA_MEMBRANE | 157 | -0.317654 | -1.02999 | 0.430556 | 1 |
| GOBP_REGULATION_OF_PROTEIN_TYROSINE_KINASE_ACTIVITY | 93 | -0.333073 | -1.029696 | 0.411364 | 1 |
| GOBP_FOREBRAIN_REGIONALIZATION | 18 | -0.444969 | -1.029625 | 0.451303 | 1 |
| GOBP_ACETYL_COA_BIOSYNTHETIC_PROCESS | 21 | -0.422711 | -1.029554 | 0.433989 | 1 |
| GOBP_POSITIVE_REGULATION_OF_CALCIUM_ION_TRANSPORT_INTO_CYTOSO | 51 | -0.361776 | -1.029129 | 0.42665 | 1 |
| GOBP_POSITIVE_REGULATION_OF_DNA_RECOMBINATION | 44 | -0.369052 | -1.028991 | 0.431204 | 1 |
| GOBP_NEGATIVE_REGULATION_OF_MRNA_PROCESSING | 29 | -0.397155 | -1.028572 | 0.440415 | 1 |
| GOBP_MITOTIC_SISTER_CHROMATID_COHESION | 26 | -0.393509 | -1.027953 | 0.424516 | 1 |
| GOBP_REGULATION_OF_PROTEIN_KINASE_C_SIGNALING | 16 | -0.441278 | -1.027945 | 0.455056 | 1 |
| GOBP_NEGATIVE_REGULATION_OF_MRNA_METABOLIC_PROCESS | 82 | -0.337265 | -1.027677 | 0.426714 | 1 |
| GOBP_SECOND_MESSENGER_MEDIATED_SIGNALING | 289 | -0.304511 | -1.026917 | 0.410623 | 1 |
| GOBP_CARDIAC_MUSCLE_TISSUE_MORPHOGENESIS | 54 | -0.353909 | -1.026901 | 0.435042 | 1 |
| GOBP_NUCLEAR_CHROMOSOME_SEGREGATION | 263 | -0.310362 | -1.02638 | 0.436029 | 1 |
| GOBP_CHROMOSOME_CONDENSATION | 40 | -0.371307 | -1.025857 | 0.445693 | 1 |
| GOBP_ENDOPLASMIC_RETICULUM_ORGANIZATION | 86 | -0.337327 | -1.02573 | 0.423516 | 1 |
| GOBP_NEGATIVE_REGULATION_OF_IMMUNE_SYSTEM_PROCESS | 377 | -0.301909 | -1.02536 | 0.426263 | 1 |
| GOBP_ENDOTHELIAL_CELL_CHEMOTAXIS | 28 | -0.394301 | -1.025173 | 0.431552 | 1 |
| GOBP_NEGATIVE_REGULATION_OF_LEUKOCYTE_MEDIATED_IMMUNITY | 52 | -0.357614 | -1.025092 | 0.419198 | 1 |
| GOBP_PROXIMAL_DISTAL_PATTERN_FORMATION | 31 | -0.390141 | -1.025052 | 0.458839 | 1 |
| GOBP_NEGATIVE_REGULATION_OF_NEURON_APOPTOTIC_PROCESS | 129 | -0.322283 | -1.025024 | 0.433983 | 1 |

| | | | | |
|---|---|---|---|---|
| GOBP_REGULATION_OF_CELL_JUNCTION_ASSEMBLY | 186 | 0.227017 | 1.011175 | 0.446429 | 0.936058 |
| GOBP_PLATELET_MORPHOGENESIS | 21 | 0.335991 | 1.010966 | 0.424 | 0.935549 |
| GOBP_PROTEOGLYCAN_METABOLIC_PROCESS | 86 | 0.26174 | 1.010315 | 0.369748 | 0.936763 |
| GOBP_RESPONSE_TO_CALCIUM_ION | 137 | 0.243468 | 1.01025 | 0.439394 | 0.935661 |
| GOBP_POSITIVE_REGULATION_OF_VASCULAR_ASSOCIATED_SMOOTH_M | 36 | 0.307508 | 1.009573 | 0.4375 | 0.936835 |
| GOBP_POSITIVE_REGULATION_OF_HORMONE_SECRETION | 116 | 0.249074 | 1.00956 | 0.435185 | 0.93553 |
| GOBP_RELAXATION_OF_CARDIAC_MUSCLE | 17 | 0.368209 | 1.009444 | 0.44086 | 0.934618 |
| GOBP_CELL_ADHESION_MEDIATED_BY_INTEGRIN | 71 | 0.265896 | 1.008975 | 0.441667 | 0.935103 |
| GOBP_POSITIVE_REGULATION_OF_MYELOID_LEUKOCYTE_MEDIATED_IM | 17 | 0.367004 | 1.008466 | 0.441176 | 0.935758 |
| GOBP_MORPHOGENESIS_OF_AN_EPITHELIAL_BUD | 15 | 0.373005 | 1.008107 | 0.459364 | 0.935776 |
| GOBP_MEMBRANE_REPOLARIZATION | 42 | 0.298786 | 1.008022 | 0.428571 | 0.934759 |
| GOBP_REGULATION_OF_BIOLOGICAL_PROCESS_INVOLVED_IN_SYMBIOT | 189 | 0.242032 | 1.007551 | 0.393939 | 0.935314 |
| GOBP_PHOSPHATIDIC_ACID_METABOLIC_PROCESS | 48 | 0.283588 | 1.007022 | 0.455556 | 0.935943 |
| GOBP_NEUROTRANSMITTER_RECEPTOR_INTERNALIZATION | 20 | 0.348813 | 1.006926 | 0.415385 | 0.934976 |
| GOBP_NATURAL_KILLER_CELL_ACTIVATION_INVOLVED_IN_IMMUNE_RE | 15 | 0.378157 | 1.00555 | 0.456747 | 0.93881 |
| GOBP_MORPHOGENESIS_OF_AN_ENDOTHELIUM | 18 | 0.354343 | 1.005468 | 0.464567 | 0.937804 |
| GOBP_REGULATION_OF_MUSCLE_CONTRACTION | 148 | 0.240226 | 1.004891 | 0.492754 | 0.938577 |
| GOBP_CELL_CELL_SIGNALING_INVOLVED_IN_CARDIAC_CONDUCTION | 30 | 0.317403 | 1.003621 | 0.408163 | 0.94205 |
| GOBP_RESPONSE_TO_INTERFERON_BETA | 28 | 0.318144 | 1.002548 | 0.456897 | 0.944864 |
| GOBP_NOTOCHORD_DEVELOPMENT | 16 | 0.363554 | 1.001876 | 0.435374 | 0.94609 |
| GOBP_MODULATION_BY_SYMBIONT_OF_HOST_PROCESS | 35 | 0.309269 | 1.001284 | 0.428571 | 0.946966 |
| GOBP_MOLTING_CYCLE | 94 | 0.255986 | 1.000977 | 0.451613 | 0.946673 |
| GOBP_BENZENE_CONTAINING_COMPOUND_METABOLIC_PROCESS | 24 | 0.337749 | 1.00082 | 0.436735 | 0.945988 |
| GOBP_REGULATION_OF_VESICLE_MEDIATED_TRANSPORT | 499 | 0.220988 | 1.000703 | 0.666667 | 0.945088 |
| GOBP_REGULATION_OF_SYNAPSE_ASSEMBLY | 90 | 0.253934 | 1.000464 | 0.46 | 0.944763 |
| GOBP_REGULATION_OF_POSITIVE_CHEMOTAXIS | 25 | 0.329159 | 0.999018 | 0.445693 | 0.948946 |
| GOBP_GASTRULATION | 168 | 0.237006 | 0.998933 | 0.426471 | 0.947902 |
| GOBP_REGULATION_OF_PEPTIDE_SECRETION | 285 | 0.222633 | 0.998576 | 0.533333 | 0.948043 |
| GOBP_POSITIVE_REGULATION_OF_MACROAUTOPHAGY | 60 | 0.267779 | 0.998188 | 0.448276 | 0.948223 |
| GOBP_METANEPHRIC_MESENCHYME_DEVELOPMENT | 15 | 0.369453 | 0.997979 | 0.479866 | 0.947737 |
| GOBP_REGULATION_OF_FIBROBLAST_GROWTH_FACTOR_RECEPTOR_S | 28 | 0.326016 | 0.997253 | 0.477551 | 0.949193 |
| GOBP_DENDRITE_MORPHOGENESIS | 137 | 0.240664 | 0.997055 | 0.397059 | 0.94859 |
| GOBP_POSITIVE_REGULATION_OF_MAPK_CASCADE | 495 | 0.214017 | 0.994493 | 0.5 | 0.957242 |
| GOBP_NEGATIVE_REGULATION_OF_PROTEASOMAL_UBIQUITIN_DEPEND | 34 | 0.296106 | 0.994441 | 0.478261 | 0.956118 |
| GOBP_MUSCLE_HYPERTROPHY | 81 | 0.261436 | 0.993704 | 0.496183 | 0.957498 |
| GOBP_MESODERM_MORPHOGENESIS | 63 | 0.266723 | 0.99337 | 0.425806 | 0.957453 |
| GOBP_NEUROTRANSMITTER_BIOSYNTHETIC_PROCESS | 15 | 0.368048 | 0.993214 | 0.439114 | 0.956704 |
| GOBP_NEGATIVE_REGULATION_OF_RELEASE_OF_CYTOCHROME_C_FR | 16 | 0.357052 | 0.992884 | 0.458955 | 0.956618 |
| GOBP_NEGATIVE_REGULATION_OF_EPITHELIAL_CELL_APOPTOTIC_PRO | 43 | 0.289839 | 0.992512 | 0.490826 | 0.95665 |
| GOBP_ENDOCARDIAL_CUSHION_FORMATION | 24 | 0.32617 | 0.992511 | 0.506438 | 0.955332 |
| GOBP_REACTIVE_OXYGEN_SPECIES_METABOLIC_PROCESS | 259 | 0.217073 | 0.992494 | 0.461538 | 0.954098 |
| GOBP_REGULATION_OF_SKELETAL_MUSCLE_TISSUE_DEVELOPMENT | 23 | 0.334914 | 0.991652 | 0.473684 | 0.956057 |
| GOBP_ORGANELLE_FUSION | 144 | 0.234565 | 0.991177 | 0.45 | 0.956536 |
| GOBP_REGULATION_OF_CARTILAGE_DEVELOPMENT | 61 | 0.270912 | 0.991081 | 0.467456 | 0.955566 |
| GOBP_CELLULAR_GLUCAN_METABOLIC_PROCESS | 71 | 0.265465 | 0.990924 | 0.446154 | 0.954755 |
| GOBP_MEMBRANE_REPOLARIZATION_DURING_ACTION_POTENTIAL | 25 | 0.319071 | 0.989837 | 0.448133 | 0.95742 |
| GOBP_PROTEIN_HOMOOLIGOMERIZATION | 170 | 0.23133 | 0.986769 | 0.509091 | 0.967733 |
| GOBP_NEUROMUSCULAR_JUNCTION_DEVELOPMENT | 46 | 0.283875 | 0.986284 | 0.460227 | 0.968124 |
| GOBP_POSITIVE_REGULATION_OF_GLIAL_CELL_DIFFERENTIATION | 38 | 0.298987 | 0.985728 | 0.494792 | 0.9689 |
| GOBP_ANOIKIS | 34 | 0.295953 | 0.985713 | 0.473684 | 0.967622 |
| GOBP_NEURAL_CREST_CELL_MIGRATION | 53 | 0.278616 | 0.98515 | 0.468208 | 0.968395 |
| GOBP_ESTABLISHMENT_OF_CELL_POLARITY | 136 | 0.245765 | 0.985124 | 0.566667 | 0.967198 |
| GOBP_LEFT_RIGHT_PATTERN_FORMATION | 19 | 0.33066 | 0.985123 | 0.475836 | 0.965895 |
| GOBP_POSITIVE_REGULATION_OF_OSTEOBLAST_DIFFERENTIATION | 61 | 0.263406 | 0.984894 | 0.455556 | 0.965483 |
| GOBP_REGULATION_OF_RESPONSE_TO_ENDOPLASMIC_RETICULUM_ST | 79 | 0.249577 | 0.984421 | 0.464 | 0.965844 |
| GOBP_COPPER_ION_TRANSPORT | 16 | 0.352316 | 0.98421 | 0.498141 | 0.965271 |
| GOBP_POSITIVE_REGULATION_OF_MYELOID_LEUKOCYTE_CYTOKINE_PI | 16 | 0.370525 | 0.983403 | 0.458647 | 0.966961 |
| GOBP_DNA_DAMAGE_RESPONSE_SIGNAL_TRANSDUCTION_BY_P53_CL | 105 | 0.240809 | 0.983299 | 0.512821 | 0.966041 |
| GOBP_ANTERIOR_POSTERIOR_PATTERN_SPECIFICATION | 185 | 0.22785 | 0.982482 | 0.423077 | 0.967666 |
| GOBP_ARGININE_METABOLIC_PROCESS | 19 | 0.341704 | 0.98193 | 0.453532 | 0.968515 |
| GOBP_REGULATION_OF_INTRINSIC_APOPTOTIC_SIGNALING_PATHWAY_ | 27 | 0.322581 | 0.979775 | 0.439834 | 0.975384 |
| GOBP_PINOCYTOSIS | 21 | 0.338309 | 0.979675 | 0.47479 | 0.974495 |
| GOBP_SKIN_EPIDERMIS_DEVELOPMENT | 79 | 0.253387 | 0.979243 | 0.486957 | 0.974853 |
| GOBP_NEGATIVE_REGULATION_OF_DEVELOPMENTAL_GROWTH | 103 | 0.242929 | 0.979014 | 0.510204 | 0.974404 |
| GOBP_AMYLOID_BETA_CLEARANCE | 36 | 0.303826 | 0.977522 | 0.448113 | 0.97842 |
| GOBP_VASCULAR_ENDOTHELIAL_GROWTH_FACTOR_PRODUCTION | 36 | 0.301426 | 0.977497 | 0.479167 | 0.977208 |
| GOBP_REGULATION_OF_TUMOR_NECROSIS_FACTOR_MEDIATED_SIGNA | 59 | 0.266066 | 0.977209 | 0.512195 | 0.977 |
| GOBP_DIOL_METABOLIC_PROCESS | 29 | 0.313477 | 0.976954 | 0.476415 | 0.976654 |
| GOBP_CORPUS_CALLOSUM_DEVELOPMENT | 15 | 0.366157 | 0.975636 | 0.463415 | 0.980233 |
| GOBP_REGULATION_OF_NEUTROPHIL_CHEMOTAXIS | 31 | 0.305841 | 0.974371 | 0.471366 | 0.983734 |
| GOBP_MEMBRANE_DEPOLARIZATION_DURING_CARDIAC_MUSCLE_CELI | 21 | 0.327123 | 0.973924 | 0.5 | 0.984039 |
| GOBP_POSITIVE_REGULATION_OF_NERVOUS_SYSTEM_DEVELOPMENT | 251 | 0.212014 | 0.972514 | 0.52381 | 0.98819 |
| GOBP_DEVELOPMENTAL_PIGMENTATION | 41 | 0.285043 | 0.9719 | 0.502463 | 0.989319 |
| GOBP_SMOOTH_MUSCLE_CELL_DIFFERENTIATION | 57 | 0.267128 | 0.971025 | 0.5 | 0.991294 |
| GOBP_REGULATION_OF_CATION_CHANNEL_ACTIVITY | 164 | 0.227997 | 0.97057 | 0.54717 | 0.991525 |
| GOBP_RESPONSE_TO_MITOCHONDRIAL_DEPOLARISATION | 20 | 0.339204 | 0.96963 | 0.527778 | 0.9937 |
| GOBP_DIOL_BIOSYNTHETIC_PROCESS | 22 | 0.337412 | 0.969563 | 0.447257 | 0.992629 |
| GOBP_REGULATION_OF_INTRINSIC_APOPTOTIC_SIGNALING_PATHWAY | 152 | 0.22866 | 0.969279 | 0.626866 | 0.992408 |
| GOBP_POSITIVE_REGULATION_OF_EPITHELIAL_TO_MESENCHYMAL_TRA | 48 | 0.274985 | 0.969258 | 0.511628 | 0.991197 |
| GOBP_AXIS_SPECIFICATION | 73 | 0.257451 | 0.969103 | 0.492857 | 0.990534 |
| GOBP_POSITIVE_REGULATION_OF_POTASSIUM_ION_TRANSMEMBRANE_ | 37 | 0.295212 | 0.969041 | 0.480687 | 0.98947 |
| GOBP_COLLAGEN_FIBRIL_ORGANIZATION | 52 | 0.272767 | 0.968606 | 0.525773 | 0.989782 |
| GOBP_POSTSYNAPTIC_NEUROTRANSMITTER_RECEPTOR_INTERNALIZAT | 17 | 0.35635 | 0.968021 | 0.488806 | 0.99067 |
| GOBP_POSITIVE_REGULATION_OF_SMOOTH_MUSCLE_CELL_MIGRATION | 38 | 0.287145 | 0.967743 | 0.513761 | 0.990427 |
| GOBP_SNRNA_TRANSCRIPTION | 74 | 0.251602 | 0.966767 | 0.5 | 0.992569 |
| GOBP_EPITHELIAL_TUBE_BRANCHING_INVOLVED_IN_LUNG_MORPHOGE | 26 | 0.324262 | 0.966621 | 0.512931 | 0.991817 |
| GOBP_NEGATIVE_REGULATION_OF_GLUCONEOGENESIS | 17 | 0.34858 | 0.966618 | 0.501742 | 0.990562 |
| GOBP_POSITIVE_REGULATION_OF_LIPID_LOCALIZATION | 103 | 0.238274 | 0.966495 | 0.558824 | 0.989723 |
| GOBP_MUSCLE_SYSTEM_PROCESS | 412 | 0.212536 | 0.966155 | 0.6 | 0.989622 |
| GOBP_REGULATION_OF_ACTION_POTENTIAL | 49 | 0.27975 | 0.965233 | 0.491228 | 0.992104 |
| GOBP_NEGATIVE_REGULATION_OF_CALCIUM_ION_TRANSMEMBRANE_T | 30 | 0.308394 | 0.96522 | 0.486842 | 0.990896 |
| GOBP_NUCLEAR_MEMBRANE_ORGANIZATION | 17 | 0.351586 | 0.964861 | 0.498246 | 0.990885 |
| GOBP_PEPTIDE_SECRETION | 373 | 0.213096 | 0.964114 | 0.666667 | 0.992255 |
| GOBP_POLYSACCHARIDE_METABOLIC_PROCESS | 106 | 0.240709 | 0.963095 | 0.568421 | 0.994887 |
| GOBP_CELLULAR_RESPONSE_TO_STEROID_HORMONE_STIMULUS | 190 | 0.221918 | 0.962916 | 0.617021 | 0.994297 |
| GOBP_NEGATIVE_REGULATION_OF_RESPONSE_TO_DNA_DAMAGE_STIM | 82 | 0.249638 | 0.96286 | 0.589286 | 0.993223 |
| GOBP_ANION_HOMEOSTASIS | 31 | 0.295917 | 0.962721 | 0.502392 | 0.992454 |
| GOBP_ANTIGEN_PROCESSING_AND_PRESENTATION_OF_PEPTIDE_ANTI | 190 | 0.218837 | 0.960549 | 0.666667 | 0.999386 |
| GOBP_CORONARY_VASCULATURE_DEVELOPMENT | 42 | 0.275971 | 0.960368 | 0.547739 | 0.998772 |
| GOBP_CATECHOL_CONTAINING_COMPOUND_METABOLIC_PROCESS | 47 | 0.265507 | 0.959907 | 0.528409 | 0.999311 |
| GOBP_EXOCYTIC_PROCESS | 77 | 0.255412 | 0.959301 | 0.53125 | 1 |
| GOBP_CELL_DIFFERENTIATION_INVOLVED_IN_KIDNEY_DEVELOPMENT | 57 | 0.269568 | 0.959255 | 0.538462 | 0.999288 |
| GOBP_RELEASE_OF_CYTOCHROME_C_FROM_MITOCHONDRIA | 54 | 0.267744 | 0.958785 | 0.525641 | 0.999699 |
| GOBP_REGULATION_OF_RELEASE_OF_CYTOCHROME_C_FROM_MITOCH | 43 | 0.276624 | 0.958556 | 0.540404 | 0.999241 |
| GOBP_REGULATION_OF_SPROUTING_ANGIOGENESIS | 38 | 0.29309 | 0.957922 | 0.557789 | 1 |
| GOBP_REGULATION_OF_DEFENSE_RESPONSE_TO_VIRUS | 66 | 0.250158 | 0.957282 | 0.523077 | 1 |
| GOBP_REGULATION_OF_SYNAPSE_STRUCTURE_OR_ACTIVITY | 206 | 0.225526 | 0.955755 | 0.666667 | 1 |
| GOBP_INNER_EAR_AUDITORY_RECEPTOR_CELL_DIFFERENTIATION | 31 | 0.303453 | 0.955672 | 0.514286 | 1 |
| GOBP_NEGATIVE_REGULATION_OF_BLOOD_VESSEL_ENDOTHELIAL_CEI | 37 | 0.284992 | 0.954669 | 0.530233 | 1 |
| GOBP_NUCLEAR_ENVELOPE_REASSEMBLY | 18 | 0.336322 | 0.954214 | 0.535055 | 1 |
| GOBP_POSITIVE_REGULATION_OF_REACTIVE_OXYGEN_SPECIES_META | 96 | 0.243586 | 0.953351 | 0.651376 | 1 |
| GOBP_GLUTAMATE_SECRETION | 39 | 0.290752 | 0.953233 | 0.530055 | 1 |
| GOBP_ANTIGEN_PROCESSING_AND_PRESENTATION | 229 | 0.215204 | 0.952249 | 0.615385 | 1 |
| GOBP_NEGATIVE_REGULATION_OF_CHONDROCYTE_DIFFERENTIATION | 21 | 0.326659 | 0.951637 | 0.501901 | 1 |
| GOBP_HEART_MORPHOGENESIS | 226 | 0.219326 | 0.951505 | 0.55 | 1 |
| GOBP_REGULATION_OF_AMYLOID_BETA_CLEARANCE | 17 | 0.335158 | 0.951322 | 0.533582 | 1 |
| GOBP_REGULATION_OF_T_CELL_MEDIATED_IMMUNITY | 76 | 0.247422 | 0.950526 | 0.584 | 1 |
| GOBP_SKELETAL_SYSTEM_DEVELOPMENT | 462 | 0.193268 | 0.950468 | 1 | 1 |
| GOBP_REGULATION_OF_RNA_SPLICING | 141 | 0.221163 | 0.949847 | 0.575343 | 1 |
| GOBP_STEM_CELL_DIFFERENTIATION | 240 | 0.213846 | 0.949216 | 0.586207 | 1 |
| GOBP_GLAND_MORPHOGENESIS | 108 | 0.23959 | 0.949156 | 0.549296 | 1 |
| GOBP_EMBRYONIC_PLACENTA_DEVELOPMENT | 76 | 0.248477 | 0.948921 | 0.574324 | 1 |
| GOBP_REGULATION_OF_VIRAL_TRANSCRIPTION | 42 | 0.280398 | 0.94885 | 0.535 | 1 |
| GOBP_NEGATIVE_REGULATION_OF_VASCULAR_ASSOCIATED_SMOOTH_ | 20 | 0.322789 | 0.947428 | 0.529183 | 1 |
| GOBP_SYNAPTIC_VESICLE_EXOCYTOSIS | 102 | 0.243205 | 0.946989 | 0.575758 | 1 |
| GOBP_POSITIVE_REGULATION_OF_ENDOTHELIAL_CELL_APOPTOTIC_PR | 17 | 0.35026 | 0.946615 | 0.537037 | 1 |
| GOBP_TRABECULA_FORMATION | 21 | 0.323068 | 0.945573 | 0.556391 | 1 |
| GOBP_REGULATION_OF_PROTEIN_BINDING | 192 | 0.215364 | 0.945049 | 0.642857 | 1 |
| GOBP_MICROTUBULE_ANCHORING | 27 | 0.300115 | 0.944855 | 0.541485 | 1 |
| GOBP_SKELETAL_MUSCLE_CELL_PROLIFERATION | 15 | 0.346619 | 0.944726 | 0.542484 | 1 |
| GOBP_REGULATION_OF_PROTEIN_KINASE_A_SIGNALING | 21 | 0.317678 | 0.944431 | 0.545455 | 1 |
| GOBP_ACTIVATION_OF_MAPK_ACTIVITY | 151 | 0.229213 | 0.944219 | 0.648148 | 1 |
| GOBP_RESPONSE_TO_REACTIVE_OXYGEN_SPECIES | 210 | 0.220645 | 0.944099 | 0.472222 | 1 |
| GOBP_CELL_DIFFERENTIATION_INVOLVED_IN_METANEPHROS_DEVELO | 24 | 0.319462 | 0.944044 | 0.493617 | 1 |
| GOBP_RESPIRATORY_SYSTEM_PROCESS | 29 | 0.300544 | 0.943176 | 0.533632 | 1 |
| GOBP_EPITHELIAL_CELL_MORPHOGENESIS | 31 | 0.304737 | 0.941652 | 0.540541 | 1 |
| GOBP_ADENYLATE_CYCLASE_ACTIVATING_ADRENERGIC_RECEPTOR_S | 21 | 0.32647 | 0.941616 | 0.522088 | 1 |
| GOBP_TRABECULA_MORPHOGENESIS | 42 | 0.272059 | 0.940916 | 0.576191 | 1 |

| | | | | | |
|---|---|---|---|---|---|
| GOBP_REGULATION_OF_ENDOTHELIAL_CELL_CHEMOTAXIS | 20 | -0.418603 | -1.024673 | 0.463549 | 1 |
| GOBP_STEROID_BIOSYNTHETIC_PROCESS | 177 | -0.312762 | -1.024609 | 0.42918 | 1 |
| GOBP_REGULATION_OF_NMDA_RECEPTOR_ACTIVITY | 34 | -0.384375 | -1.024197 | 0.426415 | 1 |
| GOBP_NADPH_REGENERATION | 17 | -0.436022 | -1.023097 | 0.46371 | 1 |
| GOBP_ALCOHOL_BIOSYNTHETIC_PROCESS | 144 | -0.316333 | -1.022864 | 0.426087 | 1 |
| GOBP_B_CELL_APOPTOTIC_PROCESS | 25 | -0.408824 | -1.022761 | 0.455026 | 1 |
| GOBP_MAINTENANCE_OF_LOCATION_IN_CELL | 202 | -0.314099 | -1.022561 | 0.426639 | 1 |
| GOBP_REGULATION_OF_ALPHA_BETA_T_CELL_ACTIVATION | 98 | -0.330237 | -1.021615 | 0.457965 | 1 |
| GOBP_I_KAPPAB_PHOSPHORYLATION | 20 | -0.419013 | -1.02158 | 0.468286 | 1 |
| GOBP_TRACHEA_DEVELOPMENT | 17 | -0.432242 | -1.020066 | 0.449931 | 1 |
| GOBP_NEGATIVE_REGULATION_OF_INTERLEUKIN_1_BETA_PRODUCTION | 27 | -0.400242 | -1.019258 | 0.46912 | 1 |
| GOBP_NEGATIVE_REGULATION_OF_INTERLEUKIN_1_PRODUCTION | 32 | -0.386979 | -1.019191 | 0.458278 | 1 |
| GOBP_NEGATIVE_REGULATION_OF_LYMPHOCYTE_ACTIVATION | 146 | -0.315079 | -1.018804 | 0.465443 | 1 |
| GOBP_OVARIAN_FOLLICLE_DEVELOPMENT | 48 | -0.355747 | -1.018062 | 0.464286 | 1 |
| GOBP_POSITIVE_REGULATION_OF_LIPASE_ACTIVITY | 67 | -0.348946 | -1.017897 | 0.458673 | 1 |
| GOBP_NEGATIVE_REGULATION_OF_CELLULAR_AMIDE_METABOLIC_PROCESS | 166 | -0.31195 | -1.017507 | 0.44958 | 1 |
| GOBP_REGULATION_OF_LEUKOCYTE_DIFFERENTIATION | 266 | -0.30529 | -1.017141 | 0.448099 | 1 |
| GOBP_3_UTR_MEDIATED_MRNA_STABILIZATION | 19 | -0.427453 | -1.01712 | 0.459864 | 1 |
| GOBP_POSITIVE_REGULATION_OF_LEUKOCYTE_CELL_CELL_ADHESION | 227 | -0.30634 | -1.016674 | 0.432124 | 1 |
| GOBP_REGULATION_OF_CELL_KILLING | 97 | -0.326577 | -1.016562 | 0.457014 | 1 |
| GOBP_NODAL_SIGNALING_PATHWAY | 15 | -0.451177 | -1.016246 | 0.472934 | 1 |
| GOBP_STEROID_METABOLIC_PROCESS | 300 | -0.301819 | -1.015625 | 0.445585 | 1 |
| GOBP_L_GLUTAMATE_IMPORT_ACROSS_PLASMA_MEMBRANE | 15 | -0.437725 | -1.015 | 0.447945 | 1 |
| GOBP_VITAMIN_D_METABOLIC_PROCESS | 18 | -0.425445 | -1.014958 | 0.461644 | 1 |
| GOBP_NEGATIVE_REGULATION_OF_ATPASE_ACTIVITY | 18 | -0.423345 | -1.014853 | 0.456522 | 1 |
| GOBP_NEGATIVE_REGULATION_OF_LYMPHOCYTE_MEDIATED_IMMUNITY | 44 | -0.361686 | -1.014816 | 0.453202 | 1 |
| GOBP_ALPHA_AMINO_ACID_BIOSYNTHETIC_PROCESS | 66 | -0.341083 | -1.014459 | 0.468187 | 1 |
| GOBP_SULFATE_TRANSPORT | 17 | -0.426739 | -1.014119 | 0.449591 | 1 |
| GOBP_CRANIAL_NERVE_DEVELOPMENT | 41 | -0.365803 | -1.013605 | 0.454315 | 1 |
| GOBP_ENDONUCLEOLYTIC_CLEAVAGE_INVOLVED_IN_RRNA_PROCESSING | 15 | -0.445987 | -1.012271 | 0.463415 | 1 |
| GOBP_GLYCOSYLCERAMIDE_METABOLIC_PROCESS | 17 | -0.430239 | -1.012151 | 0.47067 | 1 |
| GOBP_CELLULAR_COMPONENT_ASSEMBLY_INVOLVED_IN_MORPHOGENESIS | 92 | -0.326633 | -1.011889 | 0.455044 | 1 |
| GOBP_CELL_JUNCTION_DISASSEMBLY | 20 | -0.418541 | -1.010483 | 0.471748 | 1 |
| GOBP_PROTEIN_LOCALIZATION_TO_POSTSYNAPSE | 39 | -0.36827 | -1.010472 | 0.47088 | 1 |
| GOBP_CENTRIOLE_ASSEMBLY | 44 | -0.358754 | -1.010074 | 0.474969 | 1 |
| GOBP_BASE_CONVERSION_OR_SUBSTITUTION_EDITING | 18 | -0.42015 | -1.01003 | 0.47191 | 1 |
| GOBP_PHOSPHATIDYLETHANOLAMINE_ACYL_CHAIN_REMODELING | 25 | -0.396021 | -1.010018 | 0.461436 | 1 |
| GOBP_REGULATION_OF_LIPID_TRANSPORT | 135 | -0.313332 | -1.009733 | 0.456803 | 1 |
| GOBP_NEGATIVE_REGULATION_OF_T_CELL_APOPTOTIC_PROCESS | 20 | -0.417128 | -1.009458 | 0.465211 | 1 |
| GOBP_DEVELOPMENTAL_INDUCTION | 31 | -0.384012 | -1.009066 | 0.450649 | 1 |
| GOBP_2_OXOGLUTARATE_METABOLIC_PROCESS | 16 | -0.430926 | -1.008917 | 0.468056 | 1 |
| GOBP_CILIUM_OR_FLAGELLUM_DEPENDENT_CELL_MOTILITY | 112 | -0.322097 | -1.008185 | 0.476295 | 1 |
| GOBP_POSITIVE_REGULATION_OF_EXTRACELLULAR_MATRIX_ORGANIZATION | 23 | -0.405603 | -1.007907 | 0.471074 | 1 |
| GOBP_RESPONSE_TO_ORGANOPHOSPHORUS | 127 | -0.31495 | -1.007475 | 0.456897 | 1 |
| GOBP_LYMPHOCYTE_APOPTOTIC_PROCESS | 69 | -0.343153 | -1.007428 | 0.456471 | 1 |
| GOBP_RETINOIC_ACID_METABOLIC_PROCESS | 29 | -0.389235 | -1.007139 | 0.462939 | 1 |
| GOBP_DETECTION_OF_LIGHT_STIMULUS | 65 | -0.337984 | -1.006766 | 0.468281 | 1 |
| GOBP_ALCOHOL_METABOLIC_PROCESS | 351 | -0.297146 | -1.0065 | 0.486294 | 1 |
| GOBP_RESPONSE_TO_THYROID_HORMONE | 22 | -0.405823 | -1.006357 | 0.474553 | 1 |
| GOBP_ANIMAL_ORGAN_REGENERATION | 70 | -0.336381 | -1.006042 | 0.474556 | 1 |
| GOBP_REGULATION_OF_INFLAMMATORY_RESPONSE | 316 | -0.297455 | -1.006037 | 0.48934 | 1 |
| GOBP_REGULATION_OF_MICROTUBULE_BASED_MOVEMENT | 47 | -0.362214 | -1.006025 | 0.459854 | 1 |
| GOBP_SARCOMERE_ORGANIZATION | 35 | -0.372772 | -1.004865 | 0.490933 | 1 |
| GOBP_SULFUR_COMPOUND_BIOSYNTHETIC_PROCESS | 188 | -0.306363 | -1.004749 | 0.465409 | 1 |
| GOBP_EMBRYONIC_DIGESTIVE_TRACT_DEVELOPMENT | 31 | -0.373557 | -1.00445 | 0.467581 | 1 |
| GOBP_T_HELPER_17_TYPE_IMMUNE_RESPONSE | 31 | -0.386108 | -1.00429 | 0.476684 | 1 |
| GOBP_POTASSIUM_ION_IMPORT_ACROSS_PLASMA_MEMBRANE | 44 | -0.363398 | -1.004226 | 0.455224 | 1 |
| GOBP_BONE_MORPHOGENESIS | 85 | -0.328399 | -1.004072 | 0.487239 | 1 |
| GOBP_ATTACHMENT_OF_MITOTIC_SPINDLE_MICROTUBULES_TO_KINETOCHO | 15 | -0.438428 | -1.003838 | 0.487805 | 1 |
| GOBP_REGULATION_OF_HORMONE_METABOLIC_PROCESS | 33 | -0.374948 | -1.003616 | 0.465582 | 1 |
| GOBP_MALE_MEIOSIS_I | 20 | -0.408485 | -1.002744 | 0.479396 | 1 |
| GOBP_MIDBRAIN_DEVELOPMENT | 81 | -0.329935 | -1.002526 | 0.496599 | 1 |
| GOBP_PROTEIN_LOCALIZATION_TO_CILIUM | 61 | -0.339747 | -1.001222 | 0.476758 | 1 |
| GOBP_TOLL_LIKE_RECEPTOR_4_SIGNALING_PATHWAY | 38 | -0.370546 | -1.001021 | 0.460576 | 1 |
| GOBP_LYMPHOCYTE_HOMEOSTASIS | 57 | -0.35002 | -0.999665 | 0.489893 | 1 |
| GOBP_ANATOMICAL_STRUCTURE_REGRESSION | 15 | -0.438734 | -0.998898 | 0.475456 | 1 |
| GOBP_NEGATIVE_REGULATION_OF_IMMUNE_EFFECTOR_PROCESS | 120 | -0.314632 | -0.997691 | 0.48855 | 1 |
| GOBP_MITOTIC_RECOMBINATION | 24 | -0.396553 | -0.99768 | 0.487871 | 1 |
| GOBP_INNATE_IMMUNE_RESPONSE_ACTIVATING_SIGNAL_TRANSDUCTION | 114 | -0.317751 | -0.997572 | 0.493407 | 1 |
| GOBP_SOMATIC_DIVERSIFICATION_OF_IMMUNE_RECEPTORS | 74 | -0.331259 | -0.997457 | 0.481914 | 1 |
| GOBP_VASCULAR_ENDOTHELIAL_CELL_PROLIFERATION | 23 | -0.399041 | -0.997373 | 0.47263 | 1 |
| GOBP_REGULATION_OF_CELL_MATURATION | 20 | -0.412112 | -0.997233 | 0.492657 | 1 |
| GOBP_SARCOPLASMIC_RETICULUM_CALCIUM_ION_TRANSPORT | 35 | -0.372726 | -0.997197 | 0.488432 | 1 |
| GOBP_POSITIVE_REGULATION_OF_EPIDERMAL_CELL_DIFFERENTIATION | 25 | -0.394725 | -0.997182 | 0.476067 | 1 |
| GOBP_RESPONSE_TO_IMMOBILIZATION_STRESS | 21 | -0.410762 | -0.996736 | 0.490979 | 1 |
| GOBP_POSITIVE_REGULATION_OF_BEHAVIOR | 21 | -0.403533 | -0.996535 | 0.472042 | 1 |
| GOBP_REGULATION_OF_MICROGLIAL_CELL_ACTIVATION | 18 | -0.423703 | -0.99637 | 0.514325 | 1 |
| GOBP_POSITIVE_REGULATION_OF_CELL_CYCLE_G2_M_PHASE_TRANSITION | 29 | -0.37916 | -0.995959 | 0.49172 | 1 |
| GOBP_REGULATION_OF_MACROPHAGE_CHEMOTAXIS | 27 | -0.390514 | -0.995564 | 0.467192 | 1 |
| GOBP_NEGATIVE_REGULATION_OF_MRNA_CATABOLIC_PROCESS | 53 | -0.345829 | -0.995079 | 0.491337 | 1 |
| GOBP_PROTEIN_LOCALIZATION_TO_CHROMOSOME_CENTROMERIC_REGION | 25 | -0.398545 | -0.993322 | 0.503303 | 1 |
| GOBP_BLASTOCYST_FORMATION | 32 | -0.373372 | -0.993032 | 0.490765 | 1 |
| GOBP_MYELIN_ASSEMBLY | 19 | -0.418975 | -0.992586 | 0.497347 | 1 |
| GOBP_REGULATION_OF_DNA_TEMPLATED_TRANSCRIPTION_ELONGATION | 51 | -0.345288 | -0.992388 | 0.49022 | 1 |
| GOBP_CHOLESTEROL_EFFLUX | 51 | -0.351094 | -0.992325 | 0.469586 | 1 |
| GOBP_REGULATION_OF_CALCIUM_ION_TRANSPORT_INTO_CYTOSOL | 90 | -0.320532 | -0.992242 | 0.482759 | 1 |
| GOBP_REGULATION_OF_SPINDLE_CHECKPOINT | 15 | -0.441295 | -0.992003 | 0.49723 | 1 |
| GOBP_REGULATION_OF_GENE_EXPRESSION_BY_GENETIC_IMPRINTING | 15 | -0.428629 | -0.991868 | 0.502069 | 1 |
| GOBP_RECOMBINATIONAL_REPAIR | 138 | -0.312236 | -0.991165 | 0.501066 | 1 |
| GOBP_SOMATIC_DIVERSIFICATION_OF_IMMUNOGLOBULINS | 64 | -0.340053 | -0.990964 | 0.502359 | 1 |
| GOBP_FEMALE_GAMETE_GENERATION | 119 | -0.314027 | -0.990168 | 0.501654 | 1 |
| GOBP_NUCLEOSIDE_SALVAGE | 16 | -0.42991 | -0.98978 | 0.492958 | 1 |
| GOBP_ADENYLATE_CYCLASE_ACTIVATING_G_PROTEIN_COUPLED_RECEPTO | 124 | -0.30964 | -0.989639 | 0.497859 | 1 |
| GOBP_POSTSYNAPTIC_SIGNAL_TRANSDUCTION | 31 | -0.378733 | -0.989045 | 0.485751 | 1 |
| GOBP_PEPTIDYL_THREONINE_DEPHOSPHORYLATION | 22 | -0.403821 | -0.988886 | 0.495401 | 1 |
| GOBP_REGULATION_OF_OSTEOCLAST_DIFFERENTIATION | 60 | -0.336944 | -0.98887 | 0.497636 | 1 |
| GOBP_POSITIVE_REGULATION_OF_CELL_CYCLE_PHASE_TRANSITION | 92 | -0.321271 | -0.988687 | 0.504515 | 1 |
| GOBP_GLYCEROLIPID_METABOLIC_PROCESS | 408 | -0.290097 | -0.98857 | 0.545181 | 1 |
| GOBP_GLYCINE_METABOLIC_PROCESS | 19 | -0.418472 | -0.988498 | 0.493844 | 1 |
| GOBP_S_ADENOSYLMETHIONINE_METABOLIC_PROCESS | 15 | -0.428594 | -0.988448 | 0.507023 | 1 |
| GOBP_MONOCARBOXYLIC_ACID_BIOSYNTHETIC_PROCESS | 206 | -0.297137 | -0.988279 | 0.522774 | 1 |
| GOBP_ADENYLATE_CYCLASE_MODULATING_G_PROTEIN_COUPLED_RECEPT | 202 | -0.298748 | -0.988112 | 0.523958 | 1 |
| GOBP_LIPID_IMPORT_INTO_CELL | 34 | -0.364905 | -0.988028 | 0.488947 | 1 |
| GOBP_DNA_METHYLATION_DEPENDENT_HETEROCHROMATIN_ASSEMBLY | 19 | -0.411934 | -0.988005 | 0.497215 | 1 |
| GOBP_MULTICELLULAR_ORGANISMAL_HOMEOSTASIS | 499 | -0.286974 | -0.987897 | 0.564694 | 1 |
| GOBP_IMMUNOGLOBULIN_PRODUCTION_INVOLVED_IN_IMMUNOGLOBULIN_M | 54 | -0.342024 | -0.987847 | 0.510949 | 1 |
| GOBP_PEPTIDE_CATABOLIC_PROCESS | 28 | -0.37339 | -0.98651 | 0.470745 | 1 |
| GOBP_SECONDARY_PALATE_DEVELOPMENT | 24 | -0.38838 | -0.986275 | 0.477749 | 1 |
| GOBP_REGULATION_OF_NUCLEOBASE_CONTAINING_COMPOUND_TRANSPOR | 15 | -0.431683 | -0.986132 | 0.515027 | 1 |
| GOBP_POSITIVE_REGULATION_OF_AMYLOID_BETA_FORMATION | 17 | -0.41686 | -0.985845 | 0.464187 | 1 |
| GOBP_ANATOMICAL_STRUCTURE_HOMEOSTASIS | 453 | -0.287634 | -0.985538 | 0.552314 | 1 |
| GOBP_POSITIVE_REGULATION_OF_PROTEIN_KINASE_B_SIGNALING | 159 | -0.303266 | -0.985354 | 0.51383 | 1 |
| GOBP_CELLULAR_MODIFIED_AMINO_ACID_METABOLIC_PROCESS | 190 | -0.298835 | -0.985339 | 0.520042 | 1 |
| GOBP_CELL_CYCLE_DNA_REPLICATION | 64 | -0.333208 | -0.984487 | 0.501163 | 1 |
| GOBP_LYMPHOCYTE_ACTIVATION_INVOLVED_IN_IMMUNE_RESPONSE | 170 | -0.302225 | -0.983871 | 0.516613 | 1 |
| GOBP_GLYCOLIPID_CATABOLIC_PROCESS | 15 | -0.431408 | -0.983497 | 0.513399 | 1 |
| GOBP_VITAMIN_TRANSMEMBRANE_TRANSPORT | 15 | -0.434432 | -0.983293 | 0.487555 | 1 |
| GOBP_NEGATIVE_REGULATION_OF_CHROMATIN_ORGANIZATION | 59 | -0.33784 | -0.983246 | 0.489024 | 1 |
| GOBP_REGULATION_OF_DIGESTIVE_SYSTEM_PROCESS | 33 | -0.373943 | -0.98317 | 0.509554 | 1 |
| GOBP_REGULATION_OF_NEUROBLAST_PROLIFERATION | 28 | -0.381329 | -0.982888 | 0.492147 | 1 |
| GOBP_POSITIVE_REGULATION_OF_LIPID_BIOSYNTHETIC_PROCESS | 76 | -0.326381 | -0.982668 | 0.502268 | 1 |
| GOBP_NEGATIVE_REGULATION_OF_TRANSLATIONAL_INITIATION | 19 | -0.41012 | -0.982295 | 0.493741 | 1 |
| GOBP_CELLULAR_LIPID_CATABOLIC_PROCESS | 220 | -0.297985 | -0.982222 | 0.527893 | 1 |
| GOBP_REGULATION_OF_MITOTIC_SISTER_CHROMATID_SEGREGATION | 44 | -0.349566 | -0.981413 | 0.486683 | 1 |
| GOBP_CALCIUM_ION_TRANSMEMBRANE_TRANSPORT | 287 | -0.291075 | -0.981149 | 0.546099 | 1 |
| GOBP_CHROMOSOME_SEGREGATION | 324 | -0.290008 | -0.980972 | 0.520855 | 1 |
| GOBP_POSITIVE_REGULATION_OF_TRANSLATIONAL_INITIATION | 25 | -0.391291 | -0.980895 | 0.492 | 1 |
| GOBP_SECONDARY_METABOLIC_PROCESS | 52 | -0.339212 | -0.979944 | 0.494611 | 1 |
| GOBP_RESPONSE_TO_FATTY_ACID | 57 | -0.337645 | -0.979257 | 0.51962 | 1 |
| GOBP_NEGATIVE_REGULATION_OF_ACTIN_FILAMENT_POLYMERIZATION | 62 | -0.33383 | -0.979014 | 0.517039 | 1 |
| GOBP_POSITIVE_REGULATION_OF_NEURON_DIFFERENTIATION | 83 | -0.321508 | -0.979 | 0.523375 | 1 |
| GOBP_POSITIVE_REGULATION_OF_IMMUNE_EFFECTOR_PROCESS | 211 | -0.299098 | -0.978879 | 0.532575 | 1 |
| GOBP_REGULATION_OF_LEUKOCYTE_APOPTOTIC_PROCESS | 77 | -0.325532 | -0.978831 | 0.519119 | 1 |
| GOBP_INTERLEUKIN_6_PRODUCTION | 131 | -0.306686 | -0.978633 | 0.556277 | 1 |
| GOBP_POSITIVE_REGULATION_OF_RELEASE_OF_SEQUESTERED_CALCIUM_IO | 39 | -0.357176 | -0.97846 | 0.513141 | 1 |
| GOBP_REGULATION_OF_REPRODUCTIVE_PROCESS | 144 | -0.304817 | -0.978454 | 0.537623 | 1 |
| GOBP_POSITIVE_REGULATION_OF_LEUKOCYTE_MIGRATION | 130 | -0.308602 | -0.977995 | 0.529475 | 1 |
| GOBP_RESPONSE_TO_AMINE | 41 | -0.359677 | -0.976962 | 0.523284 | 1 |

| | | | | | |
|---|---|---|---|---|---|
| GOBP_T_CELL_MEDIATED_IMMUNITY | 99 | 0.239518 | 0.940846 | 0.598361 | 1 |
| GOBP_VIRAL_GENOME_REPLICATION | 126 | 0.231164 | 0.940559 | 0.581081 | 1 |
| GOBP_REGULATION_OF_SYNAPTIC_VESICLE_EXOCYTOSIS | 55 | 0.254424 | 0.940407 | 0.570513 | 1 |
| GOBP_VESICLE_DOCKING | 65 | 0.256788 | 0.939988 | 0.626667 | 1 |
| GOBP_CELLULAR_RESPONSE_TO_NUTRIENT | 37 | 0.279535 | 0.93959 | 0.56 | 1 |
| GOBP_HAIR_CELL_DIFFERENTIATION | 39 | 0.283584 | 0.938434 | 0.572165 | 1 |
| GOBP_NEGATIVE_REGULATION_OF_PROTEIN_MODIFICATION_BY_SMALL | 85 | 0.240847 | 0.938249 | 0.598291 | 1 |
| GOBP_REGULATION_OF_INTRACELLULAR_ESTROGEN_RECEPTOR_SIGN | 39 | 0.272893 | 0.938186 | 0.539604 | 1 |
| GOBP_PHOSPHOLIPID_DEPHOSPHORYLATION | 44 | 0.269167 | 0.938099 | 0.55615 | 1 |
| GOBP_CELLULAR_RESPONSE_TO_EXOGENOUS_DSRNA | 17 | 0.342115 | 0.937511 | 0.535461 | 1 |
| GOBP_REGULATION_OF_MRNA_SPLICING_VIA_SPLICEOSOME | 97 | 0.231092 | 0.937202 | 0.608333 | 1 |
| GOBP_RESPONSE_TO_TEMPERATURE_STIMULUS | 223 | 0.210782 | 0.937164 | 0.625 | 1 |
| GOBP_APOPTOTIC_CELL_CLEARANCE | 50 | 0.258538 | 0.937155 | 0.637931 | 1 |
| GOBP_SYNAPSE_MATURATION | 24 | 0.307702 | 0.936595 | 0.541502 | 1 |
| GOBP_POSITIVE_REGULATION_OF_MAST_CELL_ACTIVATION | 19 | 0.322564 | 0.936554 | 0.525547 | 1 |
| GOBP_SPERM_AXONEME_ASSEMBLY | 22 | 0.31899 | 0.936319 | 0.531381 | 1 |
| GOBP_REGULATION_OF_BONE_MINERALIZATION | 70 | 0.251066 | 0.936286 | 0.572519 | 1 |
| GOBP_NEPHRON_TUBULE_FORMATION | 17 | 0.338625 | 0.936007 | 0.531599 | 1 |
| GOBP_POSITIVE_REGULATION_OF_VASOCONSTRICTION | 27 | 0.304837 | 0.935375 | 0.582031 | 1 |
| GOBP_REGULATION_OF_DENDRITIC_SPINE_DEVELOPMENT | 60 | 0.260203 | 0.935239 | 0.621622 | 1 |
| GOBP_NOTCH_SIGNALING_PATHWAY | 175 | 0.219801 | 0.935133 | 0.666667 | 1 |
| GOBP_REGULATION_OF_AXONOGENESIS | 149 | 0.220807 | 0.934003 | 0.614035 | 1 |
| GOBP_POSITIVE_REGULATION_OF_PROTEIN_MATURATION | 24 | 0.300035 | 0.933824 | 0.57529 | 1 |
| GOBP_POSITIVE_REGULATION_OF_EPITHELIAL_CELL_MIGRATION | 157 | 0.219334 | 0.933572 | 0.696429 | 1 |
| GOBP_RESPONSE_TO_UV | 143 | 0.21748 | 0.932231 | 0.657143 | 1 |
| GOBP_REGULATION_OF_EXTENT_OF_CELL_GROWTH | 101 | 0.230713 | 0.930754 | 0.602151 | 1 |
| GOBP_REGULATION_OF_VIRAL_LIFE_CYCLE | 141 | 0.222729 | 0.930122 | 0.680556 | 1 |
| GOBP_DNA_DAMAGE_RESPONSE_SIGNAL_TRANSDUCTION_RESULTING | 18 | 0.34429 | 0.929793 | 0.531136 | 1 |
| GOBP_REGULATION_OF_AUTOPHAGY | 325 | 0.205072 | 0.92879 | 0.785714 | 1 |
| GOBP_COCHLEA_MORPHOGENESIS | 19 | 0.324358 | 0.928045 | 0.564103 | 1 |
| GOBP_REGULATION_OF_LONG_TERM_NEURONAL_SYNAPTIC_PLASTICIT | 21 | 0.322729 | 0.927732 | 0.524648 | 1 |
| GOBP_REGULATION_OF_ANIMAL_ORGAN_FORMATION | 26 | 0.312551 | 0.927486 | 0.58547 | 1 |
| GOBP_FOCAL_ADHESION_ASSEMBLY | 84 | 0.235525 | 0.926607 | 0.62931 | 1 |
| GOBP_PTERIDINE_CONTAINING_COMPOUND_BIOSYNTHETIC_PROCESS | 15 | 0.340832 | 0.925666 | 0.552189 | 1 |
| GOBP_PLASMA_MEMBRANE_PHOSPHOLIPID_SCRAMBLING | 19 | 0.319937 | 0.925213 | 0.578313 | 1 |
| GOBP_REGULATION_OF_KILLING_OF_CELLS_OF_OTHER_ORGANISM | 16 | 0.335116 | 0.925067 | 0.539286 | 1 |
| GOBP_POSITIVE_REGULATION_OF_CARDIAC_MUSCLE_CELL_PROLIFER | 23 | 0.311842 | 0.923938 | 0.550725 | 1 |
| GOBP_POSITIVE_REGULATION_OF_NITRIC_OXIDE_METABOLIC_PROCES | 36 | 0.278902 | 0.923859 | 0.579487 | 1 |
| GOBP_RESPONSE_TO_PAIN | 25 | 0.300574 | 0.92232 | 0.581897 | 1 |
| GOBP_RESPONSE_TO_VIRUS | 325 | 0.203502 | 0.922219 | 0.85 | 1 |
| GOBP_REGULATION_OF_VASCULAR_ASSOCIATED_SMOOTH_MUSCLE_C | 55 | 0.259047 | 0.921989 | 0.633333 | 1 |
| GOBP_P38MAPK_CASCADE | 44 | 0.263866 | 0.921709 | 0.583333 | 1 |
| GOBP_PROTEIN_K48_LINKED_DEUBIQUITINATION | 34 | 0.277471 | 0.920526 | 0.57767 | 1 |
| GOBP_ENTRY_INTO_HOST | 148 | 0.217209 | 0.920515 | 0.746269 | 1 |
| GOBP_PROTEIN_LOCALIZATION_TO_CELL_PERIPHERY | 320 | 0.199831 | 0.918417 | 0.846154 | 1 |
| GOBP_DEFENSE_RESPONSE_TO_VIRUS | 235 | 0.203932 | 0.918052 | 0.848485 | 1 |
| GOBP_SUPEROXIDE_ANION_GENERATION | 33 | 0.285987 | 0.917614 | 0.606635 | 1 |
| GOBP_INHIBITORY_SYNAPSE_ASSEMBLY | 15 | 0.34687 | 0.917254 | 0.552901 | 1 |
| GOBP_RESPIRATORY_GASEOUS_EXCHANGE_BY_RESPIRATORY_SYSTE | 59 | 0.252368 | 0.91456 | 0.628571 | 1 |
| GOBP_RESPONSE_TO_HEAT | 157 | 0.223129 | 0.914403 | 0.728814 | 1 |
| GOBP_MUSCLE_CELL_MIGRATION | 80 | 0.233106 | 0.913851 | 0.649573 | 1 |
| GOBP_GLUTAMATE_RECEPTOR_SIGNALING_PATHWAY | 52 | 0.255312 | 0.912741 | 0.615385 | 1 |
| GOBP_SMOOTHENED_SIGNALING_PATHWAY | 128 | 0.22063 | 0.912443 | 0.682353 | 1 |
| GOBP_POSITIVE_REGULATION_OF_STRESS_ACTIVATED_PROTEIN_KINAS | 115 | 0.222022 | 0.912281 | 0.707071 | 1 |
| GOBP_REGULATION_OF_MRNA_PROCESSING | 136 | 0.213975 | 0.91215 | 0.6875 | 1 |
| GOBP_BIOLOGICAL_PROCESS_INVOLVED_IN_INTERACTION_WITH_HOST | 211 | 0.208978 | 0.911464 | 0.755556 | 1 |
| GOBP_VIRAL_TRANSLATION | 16 | 0.334684 | 0.911192 | 0.565543 | 1 |
| GOBP_RESPONSE_TO_AUDITORY_STIMULUS | 21 | 0.30696 | 0.910804 | 0.582707 | 1 |
| GOBP_POSITIVE_REGULATION_OF_SMOOTHENED_SIGNALING_PATHWAY | 34 | 0.271616 | 0.908292 | 0.621145 | 1 |
| GOBP_MOVEMENT_IN_HOST_ENVIRONMENT | 170 | 0.208338 | 0.908124 | 0.8 | 1 |
| GOBP_N_GLYCAN_PROCESSING | 18 | 0.322541 | 0.907978 | 0.544803 | 1 |
| GOBP_POSITIVE_REGULATION_OF_HETEROTYPIC_CELL_CELL_ADHESIO | 16 | 0.327374 | 0.907836 | 0.598616 | 1 |
| GOBP_AORTA_MORPHOGENESIS | 29 | 0.289571 | 0.907669 | 0.6 | 1 |
| GOBP_REGULATION_OF_AMPA_RECEPTOR_ACTIVITY | 22 | 0.315474 | 0.907634 | 0.561069 | 1 |
| GOBP_EPHRIN_RECEPTOR_SIGNALING_PATHWAY | 86 | 0.231698 | 0.907534 | 0.743119 | 1 |
| GOBP_RIBONUCLEOTIDE_CATABOLIC_PROCESS | 39 | 0.261442 | 0.906724 | 0.654971 | 1 |
| GOBP_REGULATION_OF_NEUTROPHIL_MIGRATION | 41 | 0.26318 | 0.906659 | 0.637755 | 1 |
| GOBP_BIOMINERALIZATION | 144 | 0.220725 | 0.905698 | 0.725 | 1 |
| GOBP_NEUROTRANSMITTER_RECEPTOR_TRANSPORT_TO_PLASMA_ME | 16 | 0.334904 | 0.905482 | 0.567376 | 1 |
| GOBP_REGULATION_OF_POTASSIUM_ION_TRANSPORT | 91 | 0.230287 | 0.904632 | 0.733333 | 1 |
| GOBP_NEGATIVE_REGULATION_OF_OXIDATIVE_STRESS_INDUCED_INTR | 19 | 0.321996 | 0.904619 | 0.556 | 1 |
| GOBP_HEART_TRABECULA_MORPHOGENESIS | 29 | 0.299518 | 0.904115 | 0.625532 | 1 |
| GOBP_RESPONSE_TO_ACID_CHEMICAL | 111 | 0.218844 | 0.903938 | 0.701149 | 1 |
| GOBP_NEGATIVE_REGULATION_OF_WOUND_HEALING | 65 | 0.244924 | 0.903527 | 0.692308 | 1 |
| GOBP_MUSCLE_CELL_APOPTOTIC_PROCESS | 64 | 0.252103 | 0.903099 | 0.638743 | 1 |
| GOBP_NEGATIVE_REGULATION_OF_ENDOTHELIAL_CELL_APOPTOTIC_PI | 30 | 0.284958 | 0.903061 | 0.665236 | 1 |
| GOBP_NEURON_PROJECTION_EXTENSION | 157 | 0.21485 | 0.90087 | 0.716667 | 1 |
| GOBP_CYTOPLASMIC_SEQUESTERING_OF_PROTEIN | 22 | 0.302959 | 0.900604 | 0.605469 | 1 |
| GOBP_CIRCADIAN_REGULATION_OF_GENE_EXPRESSION | 64 | 0.239722 | 0.900063 | 0.705128 | 1 |
| GOBP_VENTRICULAR_SYSTEM_DEVELOPMENT | 27 | 0.29145 | 0.899775 | 0.584746 | 1 |
| GOBP_LYSOSOME_LOCALIZATION | 74 | 0.233402 | 0.899687 | 0.692308 | 1 |
| GOBP_NEGATIVE_REGULATION_OF_SMALL_MOLECULE_METABOLIC_PR | 89 | 0.231801 | 0.899555 | 0.704762 | 1 |
| GOBP_POSITIVE_REGULATION_OF_NOTCH_SIGNALING_PATHWAY | 53 | 0.249966 | 0.898445 | 0.664804 | 1 |
| GOBP_RESPONSE_TO_STARVATION | 186 | 0.202265 | 0.897853 | 0.72093 | 1 |
| GOBP_DE_NOVO_PROTEIN_FOLDING | 43 | 0.261034 | 0.897623 | 0.610526 | 1 |
| GOBP_DENDRITIC_SPINE_DEVELOPMENT | 98 | 0.230893 | 0.89731 | 0.743119 | 1 |
| GOBP_STARTLE_RESPONSE | 21 | 0.310587 | 0.896842 | 0.595238 | 1 |
| GOBP_LABYRINTHINE_LAYER_MORPHOGENESIS | 20 | 0.304961 | 0.896198 | 0.593284 | 1 |
| GOBP_POSITIVE_REGULATION_OF_PEPTIDYL_SERINE_PHOSPHORYLATI | 90 | 0.231877 | 0.895261 | 0.72381 | 1 |
| GOBP_POSITIVE_REGULATION_OF_TRANSCRIPTION_FROM_RNA_POLYM | 21 | 0.309398 | 0.894897 | 0.598361 | 1 |
| GOBP_TISSUE_REMODELING | 156 | 0.213768 | 0.894569 | 0.706897 | 1 |
| GOBP_SYNAPTIC_TRANSMISSION_GLUTAMATERGIC | 82 | 0.239879 | 0.89419 | 0.691781 | 1 |
| GOBP_REGULATION_OF_DENDRITE_MORPHOGENESIS | 63 | 0.241815 | 0.894041 | 0.659091 | 1 |
| GOBP_POSITIVE_REGULATION_OF_GROWTH | 228 | 0.204054 | 0.8938 | 0.9 | 1 |
| GOBP_CELLULAR_CARBOHYDRATE_BIOSYNTHETIC_PROCESS | 82 | 0.230374 | 0.89284 | 0.692308 | 1 |
| GOBP_PROTEIN_LOCALIZATION_TO_CELL_JUNCTION | 104 | 0.216249 | 0.892172 | 0.731959 | 1 |
| GOBP_G_PROTEIN_COUPLED_GLUTAMATE_RECEPTOR_SIGNALING_PAT | 15 | 0.336791 | 0.891665 | 0.598039 | 1 |
| GOBP_REGULATION_OF_BONE_REMODELING | 49 | 0.249709 | 0.890442 | 0.659091 | 1 |
| GOBP_LIPOPROTEIN_METABOLIC_PROCESS | 129 | 0.210214 | 0.890234 | 0.806452 | 1 |
| GOBP_REGULATION_OF_ANIMAL_ORGAN_MORPHOGENESIS | 175 | 0.20623 | 0.889622 | 0.8 | 1 |
| GOBP_PROTEIN_TRIMERIZATION | 16 | 0.330682 | 0.888702 | 0.596552 | 1 |
| GOBP_RESPONSE_TO_TOPOLOGICALLY_INCORRECT_PROTEIN | 200 | 0.199227 | 0.887509 | 0.893617 | 1 |
| GOBP_VENTRICULAR_CARDIAC_MUSCLE_CELL_MEMBRANE_REPOLARI | 24 | 0.294141 | 0.88686 | 0.616936 | 1 |
| GOBP_NEGATIVE_REGULATION_OF_ORGAN_GROWTH | 33 | 0.281334 | 0.886755 | 0.641148 | 1 |
| GOBP_REGULATION_OF_PROTEIN_LOCALIZATION_TO_CELL_PERIPHERY | 115 | 0.210095 | 0.885859 | 0.783505 | 1 |
| GOBP_PURINE_CONTAINING_COMPOUND_CATABOLIC_PROCESS | 49 | 0.248985 | 0.88525 | 0.718447 | 1 |
| GOBP_NEGATIVE_REGULATION_OF_T_CELL_MEDIATED_IMMUNITY | 21 | 0.296579 | 0.884745 | 0.626761 | 1 |
| GOBP_REGULATION_OF_NEURON_PROJECTION_ARBORIZATION | 15 | 0.342086 | 0.884534 | 0.589928 | 1 |
| GOBP_NEGATIVE_REGULATION_OF_I_KAPPAB_KINASE_NF_KAPPAB_SIG | 43 | 0.256983 | 0.884162 | 0.671875 | 1 |
| GOBP_TRNA_THREONYLCARBAMOYLADENOSINE_METABOLIC_PROCES | 15 | 0.325749 | 0.883527 | 0.619926 | 1 |
| GOBP_XENOBIOTIC_TRANSPORT | 40 | 0.267531 | 0.883085 | 0.659794 | 1 |
| GOBP_AMEBOIDAL_TYPE_CELL_MIGRATION | 411 | 0.194187 | 0.883064 | 1 | 1 |
| GOBP_MUCOPOLYSACCHARIDE_METABOLIC_PROCESS | 114 | 0.21704 | 0.881592 | 0.784091 | 1 |
| GOBP_POSITIVE_REGULATION_OF_CHONDROCYTE_DIFFERENTIATION | 17 | 0.309766 | 0.881132 | 0.632353 | 1 |
| GOBP_PROTEIN_HETEROOLIGOMERIZATION | 24 | 0.293141 | 0.880565 | 0.606838 | 1 |
| GOBP_REGULATION_OF_CELLULAR_RESPONSE_TO_VASCULAR_ENDOT | 22 | 0.306572 | 0.880396 | 0.609375 | 1 |
| GOBP_POSITIVE_REGULATION_OF_MAP_KINASE_ACTIVITY | 226 | 0.204505 | 0.880299 | 0.884615 | 1 |
| GOBP_REGULATION_OF_EPITHELIAL_CELL_MIGRATION | 236 | 0.19752 | 0.880266 | 0.826087 | 1 |
| GOBP_CYTOKINE_PRODUCTION_INVOLVED_IN_IMMUNE_RESPONSE | 91 | 0.226961 | 0.880126 | 0.792079 | 1 |
| GOBP_DEFINITIVE_HEMOPOIESIS | 19 | 0.297991 | 0.87994 | 0.648438 | 1 |
| GOBP_REGULATION_OF_PROTEIN_LOCALIZATION_TO_MEMBRANE | 187 | 0.198549 | 0.879779 | 0.844828 | 1 |
| GOBP_BLOOD_VESSEL_REMODELING | 40 | 0.260921 | 0.878882 | 0.703704 | 1 |
| GOBP_REACTIVE_NITROGEN_SPECIES_METABOLIC_PROCESS | 71 | 0.232515 | 0.878258 | 0.794326 | 1 |
| GOBP_PEPTIDYL_ARGININE_MODIFICATION | 25 | 0.290456 | 0.877449 | 0.654321 | 1 |
| GOBP_PROTEIN_KINASE_A_SIGNALING | 31 | 0.274953 | 0.877089 | 0.689076 | 1 |
| GOBP_PROTEIN_NEDDYLATION | 17 | 0.321969 | 0.87604 | 0.619403 | 1 |
| GOBP_EPIBOLY | 34 | 0.267062 | 0.874294 | 0.645631 | 1 |
| GOBP_T_HELPER_2_CELL_DIFFERENTIATION | 15 | 0.327932 | 0.873459 | 0.626712 | 1 |
| GOBP_REGULATION_OF_OLIGODENDROCYTE_DIFFERENTIATION | 32 | 0.273302 | 0.873352 | 0.682028 | 1 |
| GOBP_MEMBRANE_DEPOLARIZATION_DURING_ACTION_POTENTIAL | 34 | 0.272052 | 0.872887 | 0.686957 | 1 |
| GOBP_NEGATIVE_REGULATION_OF_CELLULAR_CATABOLIC_PROCESS | 239 | 0.195877 | 0.872216 | 0.884615 | 1 |
| GOBP_NEGATIVE_REGULATION_OF_EPITHELIAL_CELL_MIGRATION | 70 | 0.235021 | 0.871529 | 0.785714 | 1 |
| GOBP_REGULATION_OF_FIBROBLAST_MIGRATION | 34 | 0.267274 | 0.871356 | 0.659483 | 1 |
| GOBP_POSITIVE_REGULATION_OF_MICROTUBULE_POLYMERIZATION_OF | 33 | 0.271113 | 0.86977 | 0.714953 | 1 |
| GOBP_FILOPODIUM_ASSEMBLY | 63 | 0.236585 | 0.868847 | 0.735099 | 1 |
| GOBP_REGULATION_OF_MYOTUBE_DIFFERENTIATION | 36 | 0.265661 | 0.867807 | 0.669903 | 1 |
| GOBP_MUSCLE_CELL_PROLIFERATION | 184 | 0.202818 | 0.865562 | 0.910714 | 1 |
| GOBP_NEGATIVE_REGULATION_OF_CALCIUM_ION_TRANSPORT | 51 | 0.238499 | 0.865179 | 0.727273 | 1 |

| | | | | | |
|---|---|---|---|---|---|
| GOBP_RESPONSE_TO_GAMMA_RADIATION | 52 | -0.340746 | -0.976641 | 0.51199 | 1 |
| GOBP_QUINONE_METABOLIC_PROCESS | 34 | -0.364313 | -0.976423 | 0.515152 | 1 |
| GOBP_REGULATION_OF_PHOSPHOLIPASE_ACTIVITY | 68 | -0.326392 | -0.975859 | 0.502359 | 1 |
| GOBP_REGULATION_OF_RESPONSE_TO_BIOTIC_STIMULUS | 396 | -0.28756 | -0.97513 | 0.582828 | 1 |
| GOBP_SULFUR_COMPOUND_METABOLIC_PROCESS | 366 | -0.287477 | -0.974757 | 0.576768 | 1 |
| GOBP_CHLORIDE_TRANSPORT | 102 | -0.312047 | -0.974093 | 0.513631 | 1 |
| GOBP_PLASMA_LIPOPROTEIN_PARTICLE_CLEARANCE | 50 | -0.345852 | -0.973766 | 0.509203 | 1 |
| GOBP_POSITIVE_REGULATION_OF_IMMUNOGLOBULIN_PRODUCTION | 43 | -0.350605 | -0.973727 | 0.51358 | 1 |
| GOBP_INORGANIC_ION_IMPORT_ACROSS_PLASMA_MEMBRANE | 94 | -0.313793 | -0.97317 | 0.538462 | 1 |
| GOBP_CEREBRAL_CORTEX_DEVELOPMENT | 101 | -0.314532 | -0.973154 | 0.523446 | 1 |
| GOBP_PHOSPHATIDYLSERINE_METABOLIC_PROCESS | 35 | -0.359239 | -0.972573 | 0.539642 | 1 |
| GOBP_REGULATION_OF_RELEASE_OF_SEQUESTERED_CALCIUM_ION_INTO_C | 72 | -0.326811 | -0.971379 | 0.539884 | 1 |
| GOBP_REGULATION_OF_HORMONE_LEVELS | 480 | -0.285148 | -0.971334 | 0.60201 | 1 |
| GOBP_RESPONSE_TO_ALCOHOL | 220 | -0.293757 | -0.971334 | 0.551901 | 1 |
| GOBP_LIPID_CATABOLIC_PROCESS | 322 | -0.28981 | -0.970703 | 0.59334 | 1 |
| GOBP_SENSORY_SYSTEM_DEVELOPMENT | 345 | -0.287284 | -0.969703 | 0.577393 | 1 |
| GOBP_NEGATIVE_REGULATION_OF_MUSCLE_HYPERTROPHY | 28 | -0.369661 | -0.96968 | 0.544021 | 1 |
| GOBP_FOLIC_ACID_CONTAINING_COMPOUND_METABOLIC_PROCESS | 27 | -0.375022 | -0.969594 | 0.529644 | 1 |
| GOBP_REGULATION_OF_ALCOHOL_BIOSYNTHETIC_PROCESS | 68 | -0.325235 | -0.969501 | 0.531866 | 1 |
| GOBP_REGULATION_OF_IMMUNOGLOBULIN_PRODUCTION | 65 | -0.332166 | -0.96913 | 0.534747 | 1 |
| GOBP_PHOTORECEPTOR_CELL_MAINTENANCE | 43 | -0.344777 | -0.969049 | 0.514815 | 1 |
| GOBP_POSITIVE_REGULATION_OF_CALCIUM_ION_TRANSPORT | 110 | -0.309691 | -0.969019 | 0.55298 | 1 |
| GOBP_G_PROTEIN_COUPLED_ACETYLCHOLINE_RECEPTOR_SIGNALING_PATH | 17 | -0.412005 | -0.968918 | 0.538462 | 1 |
| GOBP_REGULATION_OF_MYELINATION | 40 | -0.361376 | -0.968708 | 0.514321 | 1 |
| GOBP_REGULATION_OF_RAC_PROTEIN_SIGNAL_TRANSDUCTION | 18 | -0.405207 | -0.96858 | 0.521924 | 1 |
| GOBP_GLUCOCORTICOID_BIOSYNTHETIC_PROCESS | 16 | -0.414178 | -0.968487 | 0.529412 | 1 |
| GOBP_NEGATIVE_REGULATION_OF_DOUBLE_STRAND_BREAK_REPAIR_VIA_H | 19 | -0.40869 | -0.968169 | 0.539919 | 1 |
| GOBP_ORGANIC_ACID_TRANSPORT | 303 | -0.28989 | -0.96759 | 0.588957 | 1 |
| GOBP_CENTRAL_NERVOUS_SYSTEM_NEURON_DEVELOPMENT | 78 | -0.322271 | -0.967384 | 0.54388 | 1 |
| GOBP_NEGATIVE_REGULATION_OF_ERBB_SIGNALING_PATHWAY | 52 | -0.336256 | -0.96699 | 0.538835 | 1 |
| GOBP_REGULATION_OF_FATTY_ACID_BIOSYNTHETIC_PROCESS | 42 | -0.34987 | -0.966666 | 0.533499 | 1 |
| GOBP_MITOTIC_NUCLEAR_DIVISION | 289 | -0.287736 | -0.966633 | 0.583418 | 1 |
| GOBP_REGULATION_OF_HIPPO_SIGNALING | 19 | -0.403075 | -0.966587 | 0.52573 | 1 |
| GOBP_SCHWANN_CELL_DEVELOPMENT | 28 | -0.370278 | -0.965822 | 0.53599 | 1 |
| GOBP_MONOCARBOXYLIC_ACID_TRANSPORT | 162 | -0.296127 | -0.96503 | 0.567597 | 1 |
| GOBP_NEURON_PROJECTION_EXTENSION_INVOLVED_IN_NEURON_PROJECT | 37 | -0.358199 | -0.964846 | 0.526841 | 1 |
| GOBP_PHOSPHATIDYLINOSITOL_ACYL_CHAIN_REMODELING | 17 | -0.410143 | -0.964744 | 0.51511 | 1 |
| GOBP_POSITIVE_REGULATION_OF_DNA_TEMPLATED_TRANSCRIPTION_INITIAT | 35 | -0.360095 | -0.96412 | 0.510076 | 1 |
| GOBP_AMINO_ACID_TRANSPORT | 144 | -0.299104 | -0.963984 | 0.556391 | 1 |
| GOBP_PROTEIN_CATABOLIC_PROCESS_IN_THE_VACUOLE | 18 | -0.405101 | -0.963482 | 0.53022 | 1 |
| GOBP_SOMATIC_RECOMBINATION_OF_IMMUNOGLOBULIN_GENE_SEGMENTS | 54 | -0.331173 | -0.962683 | 0.553086 | 1 |
| GOBP_ADAPTIVE_THERMOGENESIS | 150 | -0.300398 | -0.962571 | 0.546524 | 1 |
| GOBP_PYRIDINE_CONTAINING_COMPOUND_BIOSYNTHETIC_PROCESS | 34 | -0.35851 | -0.962463 | 0.534439 | 1 |
| GOBP_CELLULAR_MODIFIED_AMINO_ACID_BIOSYNTHETIC_PROCESS | 48 | -0.338811 | -0.962153 | 0.522947 | 1 |
| GOBP_GLOMERULAR_EPITHELIUM_DEVELOPMENT | 23 | -0.385071 | -0.961427 | 0.539683 | 1 |
| GOBP_REGULATION_OF_CELL_CELL_ADHESION | 418 | -0.282683 | -0.960774 | 0.603622 | 1 |
| GOBP_OSTEOCLAST_DEVELOPMENT | 16 | -0.417729 | -0.960469 | 0.545202 | 1 |
| GOBP_INNER_EAR_RECEPTOR_CELL_STEREOCILIUM_ORGANIZATION | 28 | -0.37118 | -0.960184 | 0.53121 | 1 |
| GOBP_REGULATION_OF_LEUKOCYTE_MEDIATED_IMMUNITY | 207 | -0.291224 | -0.960184 | 0.569038 | 1 |
| GOBP_OSTEOCLAST_DIFFERENTIATION | 89 | -0.311131 | -0.959914 | 0.552392 | 1 |
| GOBP_AMYLOID_PRECURSOR_PROTEIN_METABOLIC_PROCESS | 71 | -0.320521 | -0.958946 | 0.561118 | 1 |
| GOBP_NEGATIVE_REGULATION_OF_NATURAL_KILLER_CELL_MEDIATED_IMMU | 19 | -0.399539 | -0.958726 | 0.516529 | 1 |
| GOBP_DOPAMINE_METABOLIC_PROCESS | 34 | -0.355742 | -0.958635 | 0.537255 | 1 |
| GOBP_REGULATION_OF_METAL_ION_TRANSPORT | 229 | -0.288496 | -0.958375 | 0.592133 | 1 |
| GOBP_POSITIVE_REGULATION_OF_T_CELL_MIGRATION | 29 | -0.370819 | -0.95819 | 0.542595 | 1 |
| GOBP_CELLULAR_RESPONSE_TO_VIRUS | 62 | -0.327989 | -0.957677 | 0.545562 | 1 |
| GOBP_ANTEROGRADE_AXONAL_TRANSPORT | 47 | -0.344392 | -0.957486 | 0.545894 | 1 |
| GOBP_REGULATION_OF_HOMOTYPIC_CELL_CELL_ADHESION | 27 | -0.37005 | -0.957435 | 0.523745 | 1 |
| GOBP_NEURAL_RETINA_DEVELOPMENT | 66 | -0.320213 | -0.957366 | 0.556471 | 1 |
| GOBP_LONG_CHAIN_FATTY_ACID_IMPORT_INTO_CELL | 17 | -0.408577 | -0.956368 | 0.537517 | 1 |
| GOBP_POSITIVE_REGULATION_OF_CHEMOKINE_PRODUCTION | 56 | -0.325355 | -0.956289 | 0.547124 | 1 |
| GOBP_POSITIVE_REGULATION_OF_MITOTIC_CELL_CYCLE | 106 | -0.30365 | -0.956288 | 0.555921 | 1 |
| GOBP_HISTONE_H3_K9_MODIFICATION | 44 | -0.346314 | -0.955753 | 0.53034 | 1 |
| GOBP_POSITIVE_REGULATION_OF_MACROPHAGE_CHEMOTAXIS | 17 | -0.406725 | -0.955481 | 0.540215 | 1 |
| GOBP_NATURAL_KILLER_CELL_DIFFERENTIATION | 17 | -0.406678 | -0.955481 | 0.533333 | 1 |
| GOBP_CELLULAR_RESPONSE_TO_ESTRADIOL_STIMULUS | 33 | -0.361888 | -0.955205 | 0.554278 | 1 |
| GOBP_RESPONSE_TO_INTERLEUKIN_12 | 49 | -0.338451 | -0.955044 | 0.551766 | 1 |
| GOBP_DETECTION_OF_STIMULUS | 279 | -0.285735 | -0.954792 | 0.620123 | 1 |
| GOBP_C21_STEROID_HORMONE_BIOSYNTHETIC_PROCESS | 23 | -0.375971 | -0.954501 | 0.551491 | 1 |
| GOBP_NEGATIVE_REGULATION_OF_FIBROBLAST_PROLIFERATION | 30 | -0.359662 | -0.954283 | 0.545216 | 1 |
| GOBP_LACTATION | 44 | -0.335285 | -0.95383 | 0.531756 | 1 |
| GOBP_CELL_VOLUME_HOMEOSTASIS | 29 | -0.366493 | -0.953129 | 0.53951 | 1 |
| GOBP_CAMP_MEDIATED_SIGNALING | 58 | -0.32468 | -0.952944 | 0.559952 | 1 |
| GOBP_PROTEIN_MATURATION_BY_IRON_SULFUR_CLUSTER_TRANSFER | 15 | -0.424878 | -0.952849 | 0.537931 | 1 |
| GOBP_MYELOID_DENDRITIC_CELL_ACTIVATION | 29 | -0.369261 | -0.952795 | 0.540184 | 1 |
| GOBP_CD4_POSITIVE_OR_CD8_POSITIVE_ALPHA_BETA_T_CELL_LINEAGE_CC | 22 | -0.380075 | -0.952652 | 0.54251 | 1 |
| GOBP_TUMOR_NECROSIS_FACTOR_SUPERFAMILY_CYTOKINE_PRODUCTION | 143 | -0.29711 | -0.952602 | 0.572805 | 1 |
| GOBP_POSITIVE_REGULATION_OF_HEART_RATE | 25 | -0.377197 | -0.952432 | 0.546559 | 1 |
| GOBP_APICAL_PROTEIN_LOCALIZATION | 15 | -0.419664 | -0.951827 | 0.541444 | 1 |
| GOBP_VACUOLAR_ACIDIFICATION | 23 | -0.376995 | -0.951616 | 0.530559 | 1 |
| GOBP_MUSCLE_ORGAN_MORPHOGENESIS | 67 | -0.319764 | -0.951584 | 0.530121 | 1 |
| GOBP_UNSATURATED_FATTY_ACID_BIOSYNTHETIC_PROCESS | 46 | -0.340503 | -0.951468 | 0.562963 | 1 |
| GOBP_CELLULAR_OXIDANT_DETOXIFICATION | 91 | -0.310367 | -0.951352 | 0.571268 | 1 |
| GOBP_POSITIVE_REGULATION_OF_CELL_MIGRATION_INVOLVED_IN_SPROUTI | 25 | -0.381079 | -0.951331 | 0.547145 | 1 |
| GOBP_ORGANIC_ANION_TRANSPORT | 348 | -0.281529 | -0.951183 | 0.622222 | 1 |
| GOBP_ALPHA_BETA_T_CELL_PROLIFERATION | 37 | -0.348566 | -0.950323 | 0.541087 | 1 |
| GOBP_REGULATION_OF_DNA_RECOMBINATION | 101 | -0.304723 | -0.949917 | 0.576405 | 1 |
| GOBP_REGULATION_OF_INSULIN_LIKE_GROWTH_FACTOR_RECEPTOR_SIGNA | 21 | -0.391191 | -0.949867 | 0.556742 | 1 |
| GOBP_POSITIVE_REGULATION_OF_ANIMAL_ORGAN_MORPHOGENESIS | 30 | -0.360121 | -0.949563 | 0.547619 | 1 |
| GOBP_ACID_SECRETION | 36 | -0.349676 | -0.949309 | 0.565968 | 1 |
| GOBP_ORGANIC_ACID_TRANSMEMBRANE_TRANSPORT | 147 | -0.294382 | -0.948828 | 0.599567 | 1 |
| GOBP_REGULATION_OF_CHOLESTEROL_BIOSYNTHETIC_PROCESS | 43 | -0.341577 | -0.948654 | 0.54625 | 1 |
| GOBP_RESPONSE_TO_HYDROPEROXIDE | 16 | -0.39864 | -0.947871 | 0.559066 | 1 |
| GOBP_NEGATIVE_REGULATION_OF_SMOOTH_MUSCLE_CELL_DIFFERENTIATIO | 17 | -0.406278 | -0.947411 | 0.554945 | 1 |
| GOBP_REGULATION_OF_TOLERANCE_INDUCTION | 18 | -0.395217 | -0.947354 | 0.54212 | 1 |
| GOBP_PYRIMIDINE_NUCLEOSIDE_CATABOLIC_PROCESS | 21 | -0.387524 | -0.947124 | 0.54608 | 1 |
| GOBP_DIVALENT_INORGANIC_CATION_HOMEOSTASIS | 469 | -0.278269 | -0.945964 | 0.669339 | 1 |
| GOBP_OXIDATIVE_DEMETHYLATION | 18 | -0.400773 | -0.945348 | 0.542857 | 1 |
| GOBP_T_HELPER_CELL_LINEAGE_COMMITMENT | 17 | -0.408848 | -0.945141 | 0.57423 | 1 |
| GOBP_CHEMOKINE_PRODUCTION | 87 | -0.309104 | -0.944949 | 0.574246 | 1 |
| GOBP_MIDBRAIN_DOPAMINERGIC_NEURON_DIFFERENTIATION | 16 | -0.412065 | -0.944908 | 0.543018 | 1 |
| GOBP_CALCIUM_ION_TRANSPORT | 381 | -0.278612 | -0.944124 | 0.653885 | 1 |
| GOBP_RESPONSE_TO_SALT_STRESS | 25 | -0.373167 | -0.943865 | 0.562092 | 1 |
| GOBP_EYE_MORPHOGENESIS | 139 | -0.296729 | -0.943477 | 0.588486 | 1 |
| GOBP_NEGATIVE_REGULATION_OF_PROTEIN_POLYMERIZATION | 76 | -0.313341 | -0.943312 | 0.571268 | 1 |
| GOBP_POSITIVE_REGULATION_OF_SODIUM_ION_TRANSPORT | 35 | -0.351286 | -0.942459 | 0.572312 | 1 |
| GOBP_RESPONSE_TO_ACETYLCHOLINE | 26 | -0.368476 | -0.942006 | 0.56 | 1 |
| GOBP_TERTIARY_ALCOHOL_METABOLIC_PROCESS | 18 | -0.399503 | -0.94184 | 0.549072 | 1 |
| GOBP_POSITIVE_REGULATION_OF_BLOOD_PRESSURE | 29 | -0.36487 | -0.941711 | 0.56147 | 1 |
| GOBP_ODONTOGENESIS_OF_DENTIN_CONTAINING_TOOTH | 78 | -0.311513 | -0.941235 | 0.592765 | 1 |
| GOBP_MACROPHAGE_MIGRATION | 52 | -0.331352 | -0.941166 | 0.575829 | 1 |
| GOBP_POSITIVE_REGULATION_OF_SODIUM_ION_TRANSMEMBRANE_TRANSP( | 21 | -0.383149 | -0.940796 | 0.556017 | 1 |
| GOBP_DNA_DEMETHYLATION | 26 | -0.367003 | -0.940629 | 0.571053 | 1 |
| GOBP_NUCLEOBASE_CONTAINING_SMALL_MOLECULE_INTERCONVERSION | 27 | -0.360335 | -0.940394 | 0.571613 | 1 |
| GOBP_PURINE_CONTAINING_COMPOUND_SALVAGE | 15 | -0.405074 | -0.93936 | 0.550136 | 1 |
| GOBP_POSITIVE_REGULATION_OF_SIGNALING_RECEPTOR_ACTIVITY | 41 | -0.339847 | -0.93925 | 0.55377 | 1 |
| GOBP_ERK1_AND_ERK2_CASCADE | 291 | -0.277602 | -0.939011 | 0.646999 | 1 |
| GOBP_POSITIVE_REGULATION_OF_SMALL_MOLECULE_METABOLIC_PROCESS | 123 | -0.29719 | -0.938981 | 0.614799 | 1 |
| GOBP_REGULATION_OF_MACROPHAGE_ACTIVATION | 55 | -0.321445 | -0.938485 | 0.580189 | 1 |
| GOBP_TELOMERE_ORGANIZATION | 166 | -0.291089 | -0.937773 | 0.616999 | 1 |
| GOBP_HISTONE_PHOSPHORYLATION | 39 | -0.346676 | -0.937686 | 0.585276 | 1 |
| GOBP_RHODOPSIN_MEDIATED_SIGNALING_PATHWAY | 26 | -0.364881 | -0.937174 | 0.561697 | 1 |
| GOBP_ENDOCRINE_SYSTEM_DEVELOPMENT | 110 | -0.298313 | -0.937107 | 0.590708 | 1 |
| GOBP_SOMATIC_DIVERSIFICATION_OF_IMMUNE_RECEPTORS_VIA_SOMATIC_M | 16 | -0.408471 | -0.937104 | 0.572414 | 1 |
| GOBP_REGULATION_OF_MEMBRANE_PROTEIN_ECTODOMAIN_PROTEOLYSIS | 23 | -0.37321 | -0.93689 | 0.562834 | 1 |
| GOBP_POSITIVE_REGULATION_OF_INTERLEUKIN_6_PRODUCTION | 79 | -0.308757 | -0.936487 | 0.594318 | 1 |
| GOBP_NATURAL_KILLER_CELL_ACTIVATION | 67 | -0.316179 | -0.936442 | 0.566194 | 1 |
| GOBP_REGULATION_OF_FATTY_ACID_OXIDATION | 32 | -0.35209 | -0.935964 | 0.57197 | 1 |
| GOBP_REGULATION_OF_INTEGRIN_ACTIVATION | 17 | -0.397017 | -0.935647 | 0.549237 | 1 |
| GOBP_T_CELL_ACTIVATION_INVOLVED_IN_IMMUNE_RESPONSE | 95 | -0.303069 | -0.934263 | 0.580317 | 1 |
| GOBP_ERBB2_SIGNALING_PATHWAY | 32 | -0.350577 | -0.934194 | 0.556833 | 1 |
| GOBP_POSITIVE_REGULATION_OF_CALCIUM_MEDIATED_SIGNALING | 28 | -0.357818 | -0.934131 | 0.567183 | 1 |
| GOBP_PROTEIN_K11_LINKED_UBIQUITINATION | 29 | -0.352588 | -0.933143 | 0.580929 | 1 |
| GOBP_CELL_AGGREGATION | 19 | -0.392713 | -0.933012 | 0.570042 | 1 |
| GOBP_DOUBLE_STRAND_BREAK_REPAIR | 258 | -0.28142 | -0.93273 | 0.668699 | 1 |
| GOBP_POSITIVE_REGULATION_OF_BONE_RESORPTION | 18 | -0.395567 | -0.932659 | 0.552668 | 1 |
| GOBP_INTERLEUKIN_12_PRODUCTION | 58 | -0.321914 | -0.932634 | 0.593447 | 1 |
| GOBP_HISTONE_H3_K9_METHYLATION | 33 | -0.346898 | -0.931626 | 0.571975 | 1 |

| | | | | | |
|---|---|---|---|---|---|
| GOBP_ESTABLISHMENT_OF_PROTEIN_LOCALIZATION_TO_TELOMERE | 18 | 0.310768 | 0.864869 | 0.652 | 1 |
| GOBP_BLASTOCYST_DEVELOPMENT | 89 | 0.214624 | 0.864406 | 0.826087 | 1 |
| GOBP_BONE_REMODELING | 85 | 0.225342 | 0.864326 | 0.788321 | 1 |
| GOBP_LIPID_TRANSLOCATION | 48 | 0.246744 | 0.863895 | 0.73913 | 1 |
| GOBP_REGULATION_OF_CHOLESTEROL_EFFLUX | 34 | 0.26926 | 0.863722 | 0.686567 | 1 |
| GOBP_EMBRYONIC_FORELIMB_MORPHOGENESIS | 29 | 0.275397 | 0.863285 | 0.669528 | 1 |
| GOBP_ESTABLISHMENT_OF_PROTEIN_LOCALIZATION_TO_POSTSYNAPTI | 15 | 0.332113 | 0.862393 | 0.619048 | 1 |
| GOBP_REGULATION_OF_MESENCHYMAL_CELL_PROLIFERATION | 26 | 0.280829 | 0.861951 | 0.67623 | 1 |
| GOBP_POSITIVE_REGULATION_OF_MYELOID_CELL_DIFFERENTIATION | 98 | 0.220633 | 0.8612 | 0.790476 | 1 |
| GOBP_COPII_COATED_VESICLE_CARGO_LOADING | 15 | 0.320756 | 0.86022 | 0.683502 | 1 |
| GOBP_REGULATION_OF_STEM_CELL_PROLIFERATION | 38 | 0.256658 | 0.860213 | 0.714976 | 1 |
| GOBP_NEGATIVE_REGULATION_OF_ENDOPLASMIC_RETICULUM_STRES | 16 | 0.311261 | 0.860065 | 0.598592 | 1 |
| GOBP_PROTEIN_LOCALIZATION_TO_PLASMA_MEMBRANE | 271 | 0.189375 | 0.859495 | 0.967742 | 1 |
| GOBP_PROTEIN_ADP_RIBOSYLATION | 34 | 0.271452 | 0.859191 | 0.688312 | 1 |
| GOBP_POSITIVE_REGULATION_OF_CELL_DEVELOPMENT | 278 | 0.188116 | 0.85912 | 0.954545 | 1 |
| GOBP_REGULATION_OF_RECEPTOR_LOCALIZATION_TO_SYNAPSE | 17 | 0.303353 | 0.858657 | 0.632143 | 1 |
| GOBP_POSITIVE_REGULATION_OF_ERBB_SIGNALING_PATHWAY | 39 | 0.254732 | 0.857636 | 0.711443 | 1 |
| GOBP_REGULATION_OF_LONG_TERM_SYNAPTIC_POTENTIATION | 43 | 0.254281 | 0.857576 | 0.692737 | 1 |
| GOBP_ACTIVATION_OF_CYSTEINE_TYPE_ENDOPEPTIDASE_ACTIVITY | 15 | 0.320706 | 0.856996 | 0.649254 | 1 |
| GOBP_REGULATION_OF_CHONDROCYTE_DIFFERENTIATION | 47 | 0.255086 | 0.85692 | 0.718391 | 1 |
| GOBP_REGULATION_OF_ERAD_PATHWAY | 19 | 0.308928 | 0.856347 | 0.695122 | 1 |
| GOBP_AMYLOID_BETA_METABOLIC_PROCESS | 55 | 0.240895 | 0.856133 | 0.778443 | 1 |
| GOBP_NEGATIVE_REGULATION_OF_OXIDOREDUCTASE_ACTIVITY | 23 | 0.283384 | 0.855794 | 0.67623 | 1 |
| GOBP_REGULATION_OF_NEURON_PROJECTION_REGENERATION | 29 | 0.272048 | 0.855632 | 0.733906 | 1 |
| GOBP_REGULATION_OF_ENDOPLASMIC_RETICULUM_STRESS_INDUCED | 28 | 0.275457 | 0.855573 | 0.671296 | 1 |
| GOBP_GLUTAMINE_FAMILY_AMINO_ACID_CATABOLIC_PROCESS | 24 | 0.273988 | 0.854406 | 0.651452 | 1 |
| GOBP_MAST_CELL_ACTIVATION_INVOLVED_IN_IMMUNE_RESPONSE | 47 | 0.24113 | 0.853975 | 0.765714 | 1 |
| GOBP_NEGATIVE_REGULATION_OF_CELLULAR_PROTEIN_CATABOLIC_P | 73 | 0.227455 | 0.853824 | 0.786765 | 1 |
| GOBP_MICROTUBULE_BUNDLE_FORMATION | 102 | 0.208726 | 0.853598 | 0.836735 | 1 |
| GOBP_FIBRINOLYSIS | 21 | 0.294949 | 0.853446 | 0.677165 | 1 |
| GOBP_NEGATIVE_REGULATION_OF_DNA_BIOSYNTHETIC_PROCESS | 38 | 0.258887 | 0.852212 | 0.711864 | 1 |
| GOBP_PROTEIN_HYDROXYLATION | 27 | 0.267319 | 0.848895 | 0.702128 | 1 |
| GOBP_INOSITOL_LIPID_MEDIATED_SIGNALING | 188 | 0.193125 | 0.847812 | 0.931818 | 1 |
| GOBP_NEGATIVE_REGULATION_OF_MYELOID_LEUKOCYTE_DIFFERENTI/ | 46 | 0.243095 | 0.847657 | 0.755682 | 1 |
| GOBP_MYELIN_MAINTENANCE | 16 | 0.309369 | 0.847503 | 0.644599 | 1 |
| GOBP_REGULATION_OF_MYOBLAST_DIFFERENTIATION | 46 | 0.256488 | 0.847075 | 0.755814 | 1 |
| GOBP_G0_TO_G1_TRANSITION | 44 | 0.243979 | 0.846975 | 0.762431 | 1 |
| GOBP_PRI_MIRNA_TRANSCRIPTION_BY_RNA_POLYMERASE_II | 52 | 0.246668 | 0.846635 | 0.802548 | 1 |
| GOBP_POSITIVE_REGULATION_OF_CELL_SUBSTRATE_ADHESION | 120 | 0.205951 | 0.845715 | 0.865169 | 1 |
| GOBP_POSITIVE_REGULATION_OF_T_CELL_CYTOKINE_PRODUCTION | 20 | 0.292958 | 0.845274 | 0.701818 | 1 |
| GOBP_POSITIVE_REGULATION_OF_GLIOGENESIS | 56 | 0.235289 | 0.844474 | 0.719745 | 1 |
| GOBP_AXONEME_ASSEMBLY | 71 | 0.229669 | 0.842803 | 0.818898 | 1 |
| GOBP_RESPONSE_TO_STIMULUS_INVOLVED_IN_REGULATION_OF_MUS | 16 | 0.307065 | 0.842417 | 0.699324 | 1 |
| GOBP_LIPOPROTEIN_CATABOLIC_PROCESS | 15 | 0.304351 | 0.842036 | 0.684783 | 1 |
| GOBP_BONE_CELL_DEVELOPMENT | 32 | 0.260576 | 0.84191 | 0.736111 | 1 |
| GOBP_REGULATION_OF_CELLULAR_RESPONSE_TO_INSULIN_STIMULUS | 60 | 0.231166 | 0.840071 | 0.780488 | 1 |
| GOBP_RESPONSE_TO_LEAD_ION | 21 | 0.29115 | 0.83985 | 0.660448 | 1 |
| GOBP_REGULATION_OF_WNT_SIGNALING_PATHWAY_PLANAR_CELL_PO | 15 | 0.311411 | 0.839502 | 0.666667 | 1 |
| GOBP_POSITIVE_REGULATION_OF_AUTOPHAGY | 119 | 0.207891 | 0.83919 | 0.887324 | 1 |
| GOBP_POSITIVE_REGULATION_OF_BIOMINERALIZATION | 46 | 0.240744 | 0.839069 | 0.758065 | 1 |
| GOBP_PSEUDOPODIUM_ORGANIZATION | 17 | 0.307921 | 0.838937 | 0.664032 | 1 |
| GOBP_POSITIVE_REGULATION_OF_NEUROGENESIS | 207 | 0.191044 | 0.838148 | 0.921053 | 1 |
| GOBP_POTASSIUM_ION_EXPORT_ACROSS_PLASMA_MEMBRANE | 16 | 0.305847 | 0.837766 | 0.682657 | 1 |
| GOBP_POSITIVE_REGULATION_OF_PLASMA_MEMBRANE_BOUNDED_CEI | 105 | 0.210338 | 0.835071 | 0.835165 | 1 |
| GOBP_POSITIVE_REGULATION_OF_CARTILAGE_DEVELOPMENT | 27 | 0.276695 | 0.834272 | 0.708661 | 1 |
| GOBP_MICROVILLUS_ORGANIZATION | 24 | 0.281927 | 0.834077 | 0.707763 | 1 |
| GOBP_CELLULAR_RESPONSE_TO_EXTRACELLULAR_STIMULUS | 226 | 0.191921 | 0.833451 | 0.971429 | 1 |
| GOBP_REGULATION_OF_OXIDOREDUCTASE_ACTIVITY | 97 | 0.208379 | 0.832799 | 0.846847 | 1 |
| GOBP_ZYMOGEN_ACTIVATION | 53 | 0.236123 | 0.832693 | 0.810458 | 1 |
| GOBP_REGULATION_OF_EPITHELIAL_CELL_APOPTOTIC_PROCESS | 79 | 0.221782 | 0.832328 | 0.848 | 1 |
| GOBP_PROTEIN_K63_LINKED_DEUBIQUITINATION | 34 | 0.257566 | 0.832151 | 0.755319 | 1 |
| GOBP_REGULATION_OF_DENDRITE_DEVELOPMENT | 95 | 0.211827 | 0.831715 | 0.880342 | 1 |
| GOBP_CELL_SUBSTRATE_ADHESION | 347 | 0.182493 | 0.830871 | 1 | 1 |
| GOBP_CELL_SUBSTRATE_JUNCTION_ORGANIZATION | 105 | 0.214787 | 0.830365 | 0.858586 | 1 |
| GOBP_POSITIVE_REGULATION_OF_JNK_CASCADE | 84 | 0.213601 | 0.829537 | 0.848 | 1 |
| GOBP_POSITIVE_REGULATION_OF_ERYTHROCYTE_DIFFERENTIATION | 31 | 0.258497 | 0.829012 | 0.748918 | 1 |
| GOBP_PLASMA_MEMBRANE_TUBULATION | 20 | 0.286848 | 0.828666 | 0.723577 | 1 |
| GOBP_REGULATION_OF_ERYTHROCYTE_DIFFERENTIATION | 46 | 0.231779 | 0.827944 | 0.798913 | 1 |
| GOBP_POSITIVE_REGULATION_OF_CELLULAR_RESPONSE_TO_TRANSF( | 27 | 0.269768 | 0.827308 | 0.729614 | 1 |
| GOBP_RESPONSE_TO_MISFOLDED_PROTEIN | 24 | 0.278214 | 0.827165 | 0.712551 | 1 |
| GOBP_AMMONIUM_ION_METABOLIC_PROCESS | 22 | 0.279992 | 0.826891 | 0.72428 | 1 |
| GOBP_NUCLEOTIDE_EXCISION_REPAIR_DNA_INCISION | 39 | 0.247683 | 0.826248 | 0.766169 | 1 |
| GOBP_CELLULAR_RESPONSE_TO_STARVATION | 153 | 0.199538 | 0.825903 | 0.952381 | 1 |
| GOBP_MITOTIC_G1_S_TRANSITION_CHECKPOINT | 62 | 0.230086 | 0.825286 | 0.84106 | 1 |
| GOBP_REGULATION_OF_SODIUM_ION_TRANSMEMBRANE_TRANSPORT | 62 | 0.222919 | 0.825233 | 0.871166 | 1 |
| GOBP_NUCLEOTIDE_SUGAR_METABOLIC_PROCESS | 37 | 0.244838 | 0.825001 | 0.75 | 1 |
| GOBP_NEGATIVE_REGULATION_OF_CELL_PROJECTION_ORGANIZATION | 171 | 0.191739 | 0.824988 | 0.969231 | 1 |
| GOBP_POSITIVE_REGULATION_OF_OXIDOREDUCTASE_ACTIVITY | 54 | 0.221708 | 0.82421 | 0.837209 | 1 |
| GOBP_NEGATIVE_REGULATION_OF_NOTCH_SIGNALING_PATHWAY | 38 | 0.243339 | 0.824142 | 0.785366 | 1 |
| GOBP_REGULATION_OF_SODIUM_ION_TRANSPORT | 85 | 0.214595 | 0.823635 | 0.879699 | 1 |
| GOBP_CATECHOLAMINE_UPTAKE | 17 | 0.299342 | 0.823337 | 0.739777 | 1 |
| GOBP_REGULATION_OF_RECEPTOR_SIGNALING_PATHWAY_VIA_STAT | 82 | 0.211131 | 0.822906 | 0.840909 | 1 |
| GOBP_RESPONSE_TO_X_RAY | 32 | 0.259344 | 0.822241 | 0.777778 | 1 |
| GOBP_NEGATIVE_REGULATION_OF_LOCOMOTION | 305 | 0.18445 | 0.822168 | 1 | 1 |
| GOBP_SELECTIVE_AUTOPHAGY | 62 | 0.225313 | 0.822144 | 0.849057 | 1 |
| GOBP_RESPONSE_TO_FIBROBLAST_GROWTH_FACTOR | 132 | 0.206799 | 0.821885 | 0.965517 | 1 |
| GOBP_MITOTIC_CELL_CYCLE_ARREST | 16 | 0.299461 | 0.82187 | 0.714789 | 1 |
| GOBP_NEGATIVE_REGULATION_OF_ENDOCYTOSIS | 48 | 0.234673 | 0.820966 | 0.846561 | 1 |
| GOBP_FORELIMB_MORPHOGENESIS | 36 | 0.247091 | 0.820673 | 0.748879 | 1 |
| GOBP_NEGATIVE_REGULATION_OF_GLUCOSE_TRANSMEMBRANE_TRAN | 15 | 0.305718 | 0.819488 | 0.705036 | 1 |
| GOBP_CELLULAR_RESPONSE_TO_DSRNA | 21 | 0.286352 | 0.819204 | 0.756303 | 1 |
| GOBP_HISTONE_H3_DEACETYLATION | 19 | 0.281803 | 0.818664 | 0.744526 | 1 |
| GOBP_POSITIVE_REGULATION_OF_HISTONE_DEACETYLATION | 17 | 0.290797 | 0.815298 | 0.732852 | 1 |
| GOBP_NEGATIVE_REGULATION_OF_FIBROBLAST_GROWTH_FACTOR_RE | 19 | 0.277349 | 0.814521 | 0.707865 | 1 |
| GOBP_MODULATION_OF_PROCESS_OF_OTHER_ORGANISM_INVOLVED_ | 94 | 0.201139 | 0.813923 | 0.91453 | 1 |
| GOBP_POSITIVE_REGULATION_OF_LIPID_KINASE_ACTIVITY | 37 | 0.246332 | 0.813561 | 0.830275 | 1 |
| GOBP_MITOCHONDRIAL_CYTOCHROME_C_OXIDASE_ASSEMBLY | 21 | 0.277241 | 0.813129 | 0.757576 | 1 |
| GOBP_MIDDLE_EAR_MORPHOGENESIS | 17 | 0.287735 | 0.812604 | 0.74216 | 1 |
| GOBP_REGULATION_OF_SODIUM_ION_TRANSMEMBRANE_TRANSPORTE | 52 | 0.227514 | 0.812506 | 0.816092 | 1 |
| GOBP_MAMMARY_GLAND_MORPHOGENESIS | 39 | 0.245392 | 0.812102 | 0.816327 | 1 |
| GOBP_AMINE_CATABOLIC_PROCESS | 29 | 0.261125 | 0.812017 | 0.75 | 1 |
| GOBP_PROTEIN_LOCALIZATION_TO_SYNAPSE | 75 | 0.212194 | 0.811222 | 0.857143 | 1 |
| GOBP_DETECTION_OF_OTHER_ORGANISM | 18 | 0.287788 | 0.810618 | 0.75 | 1 |
| GOBP_NEGATIVE_REGULATION_OF_HISTONE_ACETYLATION | 16 | 0.302826 | 0.810178 | 0.716263 | 1 |
| GOBP_REGULATION_OF_MICROTUBULE_DEPOLYMERIZATION | 29 | 0.261064 | 0.809558 | 0.761062 | 1 |
| GOBP_VESICLE_ORGANIZATION | 334 | 0.186723 | 0.809071 | 1 | 1 |
| GOBP_SMOOTH_MUSCLE_CELL_MIGRATION | 67 | 0.218523 | 0.806337 | 0.875969 | 1 |
| GOBP_REGULATION_OF_MYELOID_LEUKOCYTE_MEDIATED_IMMUNITY | 52 | 0.229479 | 0.806306 | 0.845745 | 1 |
| GOBP_REGULATION_OF_RESPONSE_TO_CYTOKINE_STIMULUS | 170 | 0.184181 | 0.806186 | 0.956522 | 1 |
| GOBP_PHOSPHATIDYLINOSITOL_3_KINASE_SIGNALING | 146 | 0.191738 | 0.805443 | 0.964912 | 1 |
| GOBP_POSITIVE_REGULATION_OF_TRANSCRIPTION_FROM_RNA_POLYM | 24 | 0.277894 | 0.805337 | 0.772908 | 1 |
| GOBP_RESPONSE_TO_OXIDATIVE_STRESS | 421 | 0.173081 | 0.804947 | 1 | 1 |
| GOBP_NUCLEOTIDE_EXCISION_REPAIR_DNA_DUPLEX_UNWINDING | 22 | 0.272989 | 0.804849 | 0.761092 | 1 |
| GOBP_MAINTENANCE_OF_PROTEIN_LOCATION_IN_NUCLEUS | 23 | 0.277821 | 0.804723 | 0.761719 | 1 |
| GOBP_NCRNA_TRANSCRIPTION | 107 | 0.198879 | 0.804497 | 0.90099 | 1 |
| GOBP_NEGATIVE_REGULATION_OF_NEURON_PROJECTION_DEVELOPMI | 129 | 0.194889 | 0.803815 | 0.935065 | 1 |
| GOBP_REGULATION_OF_SMOOTHENED_SIGNALING_PATHWAY | 75 | 0.209816 | 0.803442 | 0.891026 | 1 |
| GOBP_APICAL_JUNCTION_ASSEMBLY | 81 | 0.202151 | 0.803088 | 0.871212 | 1 |
| GOBP_REGULATION_OF_MEMBRANE_LIPID_DISTRIBUTION | 54 | 0.225567 | 0.803056 | 0.824242 | 1 |
| GOBP_ACTIVATION_OF_PROTEIN_KINASE_B_ACTIVITY | 32 | 0.252937 | 0.802545 | 0.745536 | 1 |
| GOBP_INTEGRIN_MEDIATED_SIGNALING_PATHWAY | 106 | 0.197806 | 0.801506 | 0.923913 | 1 |
| GOBP_NEGATIVE_REGULATION_OF_TRANSPORTER_ACTIVITY | 71 | 0.208832 | 0.801295 | 0.892086 | 1 |
| GOBP_POSITIVE_REGULATION_OF_APOPTOTIC_SIGNALING_PATHWAY | 120 | 0.197763 | 0.801104 | 0.949495 | 1 |
| GOBP_SPHINGOID_METABOLIC_PROCESS | 22 | 0.274192 | 0.800055 | 0.749004 | 1 |
| GOBP_INTRINSIC_APOPTOTIC_SIGNALING_PATHWAY_IN_RESPONSE_TO | 100 | 0.201154 | 0.798541 | 0.896907 | 1 |
| GOBP_INCLUSION_BODY_ASSEMBLY | 22 | 0.259 | 0.798497 | 0.8 | 1 |
| GOBP_SEROTONIN_TRANSPORT | 18 | 0.283976 | 0.797833 | 0.763838 | 1 |
| GOBP_POSITIVE_REGULATION_OF_VIRAL_TRANSCRIPTION | 26 | 0.257247 | 0.797612 | 0.785156 | 1 |
| GOBP_REGULATION_OF_CELL_SUBSTRATE_ADHESION | 207 | 0.178256 | 0.796437 | 0.959184 | 1 |
| GOBP_NEGATIVE_REGULATION_OF_NECROTIC_CELL_DEATH | 18 | 0.284798 | 0.795419 | 0.766917 | 1 |
| GOBP_REGULATION_OF_RESPONSE_TO_DNA_DAMAGE_STIMULUS | 219 | 0.182668 | 0.795038 | 1 | 1 |
| GOBP_REGULATION_OF_REACTIVE_OXYGEN_SPECIES_METABOLIC_PR( | 179 | 0.1847 | 0.794514 | 0.981482 | 1 |
| GOBP_NEGATIVE_REGULATION_OF_CELL_MIGRATION_INVOLVED_IN_SP | 18 | 0.289873 | 0.794195 | 0.745583 | 1 |
| GOBP_MULTICELLULAR_ORGANISMAL_MOVEMENT | 52 | 0.219607 | 0.792711 | 0.883041 | 1 |
| GOBP_CELLULAR_RESPONSE_TO_GROWTH_HORMONE_STIMULUS | 21 | 0.27067 | 0.79133 | 0.757692 | 1 |
| GOBP_FUCOSE_METABOLIC_PROCESS | 15 | 0.29996 | 0.790834 | 0.730104 | 1 |
| GOBP_REGULATION_OF_MUSCLE_ORGAN_DEVELOPMENT | 38 | 0.238434 | 0.790436 | 0.850962 | 1 |
| GOBP_REGULATION_OF_STEM_CELL_DIFFERENTIATION | 109 | 0.191126 | 0.789789 | 0.963636 | 1 |

| | | | | | |
|---|---|---|---|---|---|
| GOBP_REGULATION_OF_NEURONAL_SYNAPTIC_PLASTICITY | 44 | -0.337704 | -0.93134 | 0.582707 | 1 |
| GOBP_REGULATION_OF_HISTONE_METHYLATION | 59 | -0.321695 | -0.930914 | 0.615385 | 1 |
| GOBP_TRANSITION_METAL_ION_TRANSPORT | 120 | -0.295343 | -0.930902 | 0.635965 | 1 |
| GOBP_POSITIVE_REGULATION_OF_CELL_KILLING | 62 | -0.316309 | -0.930609 | 0.572609 | 1 |
| GOBP_SKELETAL_MUSCLE_ORGAN_DEVELOPMENT | 148 | -0.287766 | -0.930534 | 0.626582 | 1 |
| GOBP_REGULATION_OF_UBIQUITIN_PROTEIN_LIGASE_ACTIVITY | 23 | -0.369083 | -0.930286 | 0.567604 | 1 |
| GOBP_POSITIVE_REGULATION_OF_MULTICELLULAR_ORGANISM_GROWTH | 27 | -0.361186 | -0.93002 | 0.565563 | 1 |
| GOBP_WNT_SIGNALING_PATHWAY_CALCIUM_MODULATING_PATHWAY | 37 | -0.340472 | -0.929311 | 0.558714 | 1 |
| GOBP_EMBRYONIC_CRANIAL_SKELETON_MORPHOGENESIS | 45 | -0.331726 | -0.929133 | 0.590799 | 1 |
| GOBP_MITOCHONDRIAL_CALCIUM_ION_HOMEOSTASIS | 26 | -0.363442 | -0.929116 | 0.573092 | 1 |
| GOBP_TRIGLYCERIDE_BIOSYNTHETIC_PROCESS | 37 | -0.342473 | -0.92826 | 0.591687 | 1 |
| GOBP_REGULATION_OF_CELL_CELL_ADHESION_MEDIATED_BY_CADHERIN | 20 | -0.380816 | -0.928122 | 0.565158 | 1 |
| GOBP_AMINO_ACID_TRANSMEMBRANE_TRANSPORT | 89 | -0.300042 | -0.928092 | 0.6 | 1 |
| GOBP_CIRCADIAN_RHYTHM | 198 | -0.284946 | -0.927819 | 0.651945 | 1 |
| GOBP_REGULATION_OF_NUCLEOTIDE_BIOSYNTHETIC_PROCESS | 38 | -0.337147 | -0.927754 | 0.57572 | 1 |
| GOBP_BRANCHING_INVOLVED_IN_SALIVARY_GLAND_MORPHOGENESIS | 20 | -0.383882 | -0.927729 | 0.557952 | 1 |
| GOBP_PROTEIN_HOMOTETRAMERIZATION | 56 | -0.319279 | -0.92755 | 0.601675 | 1 |
| GOBP_REGULATION_OF_ANDROGEN_RECEPTOR_SIGNALING_PATHWAY | 24 | -0.368881 | -0.927353 | 0.576248 | 1 |
| GOBP_CYCLIC_NUCLEOTIDE_MEDIATED_SIGNALING | 80 | -0.306642 | -0.927139 | 0.60113 | 1 |
| GOBP_INACTIVATION_OF_MAPK_ACTIVITY | 24 | -0.366728 | -0.92666 | 0.570081 | 1 |
| GOBP_GENETIC_IMPRINTING | 23 | -0.369048 | -0.926083 | 0.570856 | 1 |
| GOBP_ACTIVATION_OF_PHOSPHOLIPASE_C_ACTIVITY | 30 | -0.35806 | -0.925609 | 0.583875 | 1 |
| GOBP_GLOMERULUS_DEVELOPMENT | 60 | -0.315455 | -0.925424 | 0.605701 | 1 |
| GOBP_NEGATIVE_REGULATION_OF_IMMUNE_RESPONSE | 139 | -0.287969 | -0.925345 | 0.643703 | 1 |
| GOBP_POSITIVE_REGULATION_OF_CYTOKINE_PRODUCTION | 416 | -0.272234 | -0.924438 | 0.711268 | 1 |
| GOBP_LIPID_OXIDATION | 109 | -0.291701 | -0.924336 | 0.617744 | 1 |
| GOBP_NEGATIVE_REGULATION_OF_ANION_TRANSMEMBRANE_TRANSPORT | 31 | -0.349789 | -0.924136 | 0.600756 | 1 |
| GOBP_PYRIMIDINE_NUCLEOTIDE_METABOLIC_PROCESS | 49 | -0.325778 | -0.923991 | 0.593596 | 1 |
| GOBP_INSULIN_LIKE_GROWTH_FACTOR_RECEPTOR_SIGNALING_PATHWAY | 32 | -0.346157 | -0.9237 | 0.591146 | 1 |
| GOBP_CELLULAR_RESPONSE_TO_TOXIC_SUBSTANCE | 111 | -0.294172 | -0.923682 | 0.631929 | 1 |
| GOBP_ACTIVATION_OF_GTPASE_ACTIVITY | 106 | -0.295174 | -0.923368 | 0.635241 | 1 |
| GOBP_ORGANIC_HYDROXY_COMPOUND_CATABOLIC_PROCESS | 72 | -0.306096 | -0.923316 | 0.586826 | 1 |
| GOBP_ACTIVATION_OF_INNATE_IMMUNE_RESPONSE | 142 | -0.288 | -0.923204 | 0.638229 | 1 |
| GOBP_CELLULAR_ALDEHYDE_METABOLIC_PROCESS | 60 | -0.314951 | -0.923096 | 0.612485 | 1 |
| GOBP_ASPARTATE_FAMILY_AMINO_ACID_METABOLIC_PROCESS | 49 | -0.322213 | -0.923024 | 0.616587 | 1 |
| GOBP_LEUKOCYTE_APOPTOTIC_PROCESS | 99 | -0.295134 | -0.923011 | 0.623077 | 1 |
| GOBP_NEGATIVE_REGULATION_OF_INTERLEUKIN_2_PRODUCTION | 26 | -0.36748 | -0.922882 | 0.596232 | 1 |
| GOBP_NEGATIVE_REGULATION_OF_RECEPTOR_MEDIATED_ENDOCYTOSIS | 29 | -0.346167 | -0.921795 | 0.569604 | 1 |
| GOBP_CELLULAR_RESPONSE_TO_ALKALOID | 29 | -0.360608 | -0.921303 | 0.585302 | 1 |
| GOBP_CELL_REDOX_HOMEOSTASIS | 43 | -0.331314 | -0.920886 | 0.617827 | 1 |
| GOBP_REGULATION_OF_DNA_BINDING | 114 | -0.293081 | -0.920311 | 0.632337 | 1 |
| GOBP_RESPONSE_TO_ANGIOTENSIN | 23 | -0.372045 | -0.920247 | 0.580733 | 1 |
| GOBP_HISTONE_DEUBIQUITINATION | 23 | -0.371914 | -0.920058 | 0.585333 | 1 |
| GOBP_THYROID_GLAND_DEVELOPMENT | 21 | -0.374087 | -0.919996 | 0.584106 | 1 |
| GOBP_REGULATION_OF_BLOOD_PRESSURE | 163 | -0.282233 | -0.919868 | 0.653887 | 1 |
| GOBP_MALE_GAMETE_GENERATION | 472 | -0.271128 | -0.919317 | 0.751509 | 1 |
| GOBP_ONE_CARBON_METABOLIC_PROCESS | 39 | -0.332384 | -0.918734 | 0.607868 | 1 |
| GOBP_OUTFLOW_TRACT_SEPTUM_MORPHOGENESIS | 25 | -0.364343 | -0.918309 | 0.606952 | 1 |
| GOBP_ORGAN_INDUCTION | 21 | -0.375687 | -0.917627 | 0.597841 | 1 |
| GOBP_NEGATIVE_REGULATION_OF_MEIOTIC_CELL_CYCLE | 15 | -0.398958 | -0.917463 | 0.581233 | 1 |
| GOBP_REGULATION_OF_LEUKOCYTE_CHEMOTAXIS | 115 | -0.291153 | -0.917446 | 0.636564 | 1 |
| GOBP_POSITIVE_REGULATION_OF_PROTEIN_DEPHOSPHORYLATION | 45 | -0.331314 | -0.917265 | 0.599278 | 1 |
| GOBP_REGULATION_OF_TRANSCRIPTION_INITIATION_FROM_RNA_POLYMERA | 34 | -0.340263 | -0.917064 | 0.621232 | 1 |
| GOBP_REGULATION_OF_DELAYED_RECTIFIER_POTASSIUM_CHANNEL_ACTIVI | 18 | -0.388794 | -0.917012 | 0.590846 | 1 |
| GOBP_NEGATIVE_REGULATION_OF_DNA_RECOMBINATION | 42 | -0.331101 | -0.916904 | 0.592308 | 1 |
| GOBP_MULTICELLULAR_ORGANISM_GROWTH | 127 | -0.286884 | -0.915753 | 0.658966 | 1 |
| GOBP_NEGATIVE_REGULATION_OF_ADAPTIVE_IMMUNE_RESPONSE | 51 | -0.326194 | -0.915593 | 0.622549 | 1 |
| GOBP_PHOSPHATIDYLINOSITOL_PHOSPHORYLATION | 56 | -0.314834 | -0.915381 | 0.605735 | 1 |
| GOBP_POSITIVE_REGULATION_OF_INTERLEUKIN_10_PRODUCTION | 37 | -0.337324 | -0.915295 | 0.591139 | 1 |
| GOBP_RESPONSE_TO_MANGANESE_ION | 20 | -0.378317 | -0.915205 | 0.600533 | 1 |
| GOBP_POSITIVE_REGULATION_OF_LIPID_METABOLIC_PROCESS | 137 | -0.286147 | -0.914656 | 0.652361 | 1 |
| GOBP_PROTEIN_LOCALIZATION_TO_CHROMATIN | 29 | -0.352445 | -0.914132 | 0.60502 | 1 |
| GOBP_NEGATIVE_REGULATION_OF_CELL_KILLING | 26 | -0.360184 | -0.912992 | 0.612946 | 1 |
| GOBP_POSITIVE_REGULATION_OF_MACROPHAGE_ACTIVATION | 26 | -0.359694 | -0.912927 | 0.604806 | 1 |
| GOBP_CD8_POSITIVE_ALPHA_BETA_T_CELL_ACTIVATION | 26 | -0.358602 | -0.912718 | 0.578187 | 1 |
| GOBP_PYRIMIDINE_RIBONUCLEOTIDE_METABOLIC_PROCESS | 29 | -0.354736 | -0.911848 | 0.612739 | 1 |
| GOBP_REGULATION_OF_PATTERN_RECOGNITION_RECEPTOR_SIGNALING_PA | 98 | -0.294569 | -0.911825 | 0.654525 | 1 |
| GOBP_BONE_GROWTH | 25 | -0.352184 | -0.911749 | 0.599459 | 1 |
| GOBP_SEMAPHORIN_PLEXIN_SIGNALING_PATHWAY | 40 | -0.334372 | -0.911238 | 0.615385 | 1 |
| GOBP_REGULATION_OF_CD4_POSITIVE_ALPHA_BETA_T_CELL_DIFFERENTIAT | 50 | -0.318583 | -0.911143 | 0.621065 | 1 |
| GOBP_REGULATION_OF_PHOSPHOLIPID_BIOSYNTHETIC_PROCESS | 15 | -0.399072 | -0.910997 | 0.601108 | 1 |
| GOBP_ALPHA_AMINO_ACID_METABOLIC_PROCESS | 185 | -0.278071 | -0.910844 | 0.696747 | 1 |
| GOBP_ACUTE_INFLAMMATORY_RESPONSE | 95 | -0.297569 | -0.909843 | 0.637615 | 1 |
| GOBP_CELLULAR_RESPONSE_TO_KETONE | 88 | -0.297716 | -0.908873 | 0.643101 | 1 |
| GOBP_POSITIVE_REGULATION_OF_JUN_KINASE_ACTIVITY | 67 | -0.306017 | -0.908116 | 0.64 | 1 |
| GOBP_DETECTION_OF_TEMPERATURE_STIMULUS | 21 | -0.368766 | -0.907984 | 0.61518 | 1 |
| GOBP_REGULATION_OF_PHOSPHOPROTEIN_PHOSPHATASE_ACTIVITY | 115 | -0.286691 | -0.90757 | 0.666293 | 1 |
| GOBP_REGULATION_OF_JUN_KINASE_ACTIVITY | 81 | -0.295476 | -0.907103 | 0.649044 | 1 |
| GOBP_SPINDLE_ASSEMBLY | 114 | -0.287029 | -0.906535 | 0.661926 | 1 |
| GOBP_RNA_PHOSPHODIESTER_BOND_HYDROLYSIS_ENDONUCLEOLYTIC | 76 | -0.299976 | -0.906496 | 0.637631 | 1 |
| GOBP_REGULATION_OF_SYNAPTIC_PLASTICITY | 170 | -0.279322 | -0.906083 | 0.691332 | 1 |
| GOBP_POSITIVE_REGULATION_OF_MITOCHONDRIAL_OUTER_MEMBRANE_PEI | 35 | -0.340057 | -0.906011 | 0.617076 | 1 |
| GOBP_NEGATIVE_REGULATION_OF_LEUKOCYTE_APOPTOTIC_PROCESS | 45 | -0.321094 | -0.90595 | 0.625 | 1 |
| GOBP_NEURON_PROJECTION_GUIDANCE | 269 | -0.270098 | -0.905643 | 0.727829 | 1 |
| GOBP_REGULATION_OF_STEROL_TRANSPORT | 55 | -0.315965 | -0.904978 | 0.63835 | 1 |
| GOBP_SMAD_PROTEIN_SIGNAL_TRANSDUCTION | 71 | -0.303352 | -0.904676 | 0.647332 | 1 |
| GOBP_CELLULAR_RESPONSE_TO_INTERFERON_BETA | 19 | -0.373046 | -0.90453 | 0.605333 | 1 |
| GOBP_REGULATION_OF_PROTEIN_MATURATION | 60 | -0.307491 | -0.904146 | 0.646429 | 1 |
| GOBP_NEGATIVE_REGULATION_OF_MAP_KINASE_ACTIVITY | 71 | -0.299699 | -0.903863 | 0.633721 | 1 |
| GOBP_ADHESION_OF_SYMBIONT_TO_HOST | 15 | -0.391749 | -0.903394 | 0.60462 | 1 |
| GOBP_POSITIVE_REGULATION_OF_RECEPTOR_INTERNALIZATION | 25 | -0.359337 | -0.903316 | 0.608983 | 1 |
| GOBP_NEGATIVE_CHEMOTAXIS | 45 | -0.325035 | -0.903219 | 0.63171 | 1 |
| GOBP_LABYRINTHINE_LAYER_BLOOD_VESSEL_DEVELOPMENT | 18 | -0.381025 | -0.902652 | 0.609396 | 1 |
| GOBP_REGULATION_OF_FEEDING_BEHAVIOR | 21 | -0.370541 | -0.902573 | 0.602476 | 1 |
| GOBP_MUSCLE_ORGAN_DEVELOPMENT | 297 | -0.268585 | -0.902526 | 0.746421 | 1 |
| GOBP_DOPAMINE_RECEPTOR_SIGNALING_PATHWAY | 37 | -0.331925 | -0.901938 | 0.627353 | 1 |
| GOBP_HEART_FORMATION | 28 | -0.347948 | -0.901694 | 0.631926 | 1 |
| GOBP_INORGANIC_ANION_TRANSPORT | 158 | -0.278465 | -0.901302 | 0.678762 | 1 |
| GOBP_SUBSTANTIA_NIGRA_DEVELOPMENT | 42 | -0.321701 | -0.901301 | 0.624688 | 1 |
| GOBP_POSITIVE_CHEMOTAXIS | 57 | -0.30979 | -0.900888 | 0.639506 | 1 |
| GOBP_HINDLIMB_MORPHOGENESIS | 33 | -0.335347 | -0.900417 | 0.626444 | 1 |
| GOBP_SENSORY_ORGAN_MORPHOGENESIS | 234 | -0.270881 | -0.900124 | 0.71473 | 1 |
| GOBP_POSITIVE_REGULATION_OF_GENE_EXPRESSION_EPIGENETIC | 56 | -0.31083 | -0.899848 | 0.644005 | 1 |
| GOBP_MRNA_MODIFICATION | 24 | -0.364187 | -0.899427 | 0.622517 | 1 |
| GOBP_CELLULAR_IRON_ION_HOMEOSTASIS | 65 | -0.300693 | -0.899228 | 0.64294 | 1 |
| GOBP_CELL_CYCLE_G2_M_PHASE_TRANSITION | 270 | -0.269148 | -0.899103 | 0.734359 | 1 |
| GOBP_REGULATION_OF_PRODUCTION_OF_MOLECULAR_MEDIATOR_OF_IMMU | 140 | -0.283353 | -0.898898 | 0.715665 | 1 |
| GOBP_NEURAL_TUBE_PATTERNING | 34 | -0.334089 | -0.898415 | 0.627451 | 1 |
| GOBP_ALCOHOL_CATABOLIC_PROCESS | 52 | -0.313714 | -0.898253 | 0.621749 | 1 |
| GOBP_SENSORY_PERCEPTION_OF_TEMPERATURE_STIMULUS | 22 | -0.364984 | -0.897929 | 0.610959 | 1 |
| GOBP_POSITIVE_REGULATION_OF_SPROUTING_ANGIOGENESIS | 21 | -0.365696 | -0.897652 | 0.621951 | 1 |
| GOBP_NEGATIVE_REGULATION_OF_ACTIN_FILAMENT_DEPOLYMERIZATION | 44 | -0.323704 | -0.897633 | 0.64799 | 1 |
| GOBP_POSITIVE_REGULATION_OF_ENDOTHELIAL_CELL_PROLIFERATION | 94 | -0.287808 | -0.897229 | 0.674972 | 1 |
| GOBP_CELLULAR_RESPONSE_TO_THYROID_HORMONE_STIMULUS | 15 | -0.394706 | -0.896882 | 0.618105 | 1 |
| GOBP_VASCULAR_PROCESS_IN_CIRCULATORY_SYSTEM | 235 | -0.269205 | -0.896076 | 0.722394 | 1 |
| GOBP_POSITIVE_REGULATION_OF_LEUKOCYTE_MEDIATED_IMMUNITY | 124 | -0.280697 | -0.89562 | 0.67861 | 1 |
| GOBP_ACTIVATED_T_CELL_PROLIFERATION | 43 | -0.318739 | -0.895418 | 0.636475 | 1 |
| GOBP_IRON_ION_HOMEOSTASIS | 81 | -0.296609 | -0.894735 | 0.677928 | 1 |
| GOBP_POSITIVE_REGULATION_OF_TRANSPORTER_ACTIVITY | 109 | -0.285449 | -0.894067 | 0.706992 | 1 |
| GOBP_MITOCHONDRION_LOCALIZATION | 47 | -0.317555 | -0.893073 | 0.658537 | 1 |
| GOBP_CELLULAR_KETONE_METABOLIC_PROCESS | 231 | -0.268837 | -0.892859 | 0.737874 | 1 |
| GOBP_REGULATION_OF_EXTRACELLULAR_MATRIX_ORGANIZATION | 45 | -0.322373 | -0.89254 | 0.643564 | 1 |
| GOBP_REGULATION_OF_T_CELL_MIGRATION | 41 | -0.32518 | -0.891771 | 0.641124 | 1 |
| GOBP_NEGATIVE_REGULATION_OF_T_CELL_PROLIFERATION | 64 | -0.308142 | -0.89118 | 0.671378 | 1 |
| GOBP_REGULATION_OF_SYSTEMIC_ARTERIAL_BLOOD_PRESSURE_MEDIATEI | 43 | -0.314534 | -0.891073 | 0.656863 | 1 |
| GOBP_CYTOSKELETON_DEPENDENT_CYTOKINESIS | 97 | -0.286544 | -0.891071 | 0.672666 | 1 |
| GOBP_DICARBOXYLIC_ACID_CATABOLIC_PROCESS | 16 | -0.378147 | -0.890942 | 0.629681 | 1 |
| GOBP_BONE_MATURATION | 21 | -0.366586 | -0.890839 | 0.629781 | 1 |
| GOBP_PYRIMIDINE_CONTAINING_COMPOUND_METABOLIC_PROCESS | 82 | -0.295937 | -0.890833 | 0.66257 | 1 |
| GOBP_STEROL_METABOLIC_PROCESS | 152 | -0.277046 | -0.89014 | 0.708243 | 1 |
| GOBP_BLOOD_VESSEL_ENDOTHELIAL_CELL_PROLIFERATION_INVOLVED_IN_ | 31 | -0.339193 | -0.89012 | 0.638633 | 1 |
| GOBP_STEROL_TRANSPORT | 105 | -0.285127 | -0.890063 | 0.675854 | 1 |
| GOBP_CARBOHYDRATE_PHOSPHORYLATION | 24 | -0.362284 | -0.889973 | 0.631234 | 1 |
| GOBP_NEGATIVE_REGULATION_OF_RNA_CATABOLIC_PROCESS | 64 | -0.301674 | -0.889933 | 0.67063 | 1 |
| GOBP_SPINDLE_ORGANIZATION | 180 | -0.273831 | -0.889713 | 0.741015 | 1 |
| GOBP_RELEASE_OF_SEQUESTERED_CALCIUM_ION_INTO_CYTOSOL_BY_END( | 31 | -0.337915 | -0.889432 | 0.621728 | 1 |
| GOBP_ODONTOGENESIS | 116 | -0.281082 | -0.889201 | 0.707828 | 1 |
| GOBP_RHYTHMIC_PROCESS | 276 | -0.264773 | -0.888974 | 0.768839 | 1 |

| | | | | | |
|---|---|---|---|---|---|
| GOBP_POLY_A_PLUS_MRNA_EXPORT_FROM_NUCLEUS | 21 | 0.261377 | 0.788614 | 0.780877 | 1 |
| GOBP_BODY_MORPHOGENESIS | 46 | 0.232045 | 0.787972 | 0.860215 | 1 |
| GOBP_REGULATION_OF_BIOMINERALIZATION | 88 | 0.201212 | 0.785243 | 0.943396 | 1 |
| GOBP_HEPATOCYTE_GROWTH_FACTOR_RECEPTOR_SIGNALING_PATHW | 16 | 0.281416 | 0.785227 | 0.750809 | 1 |
| GOBP_POSITIVE_REGULATION_OF_VASCULAR_PERMEABILITY | 15 | 0.291651 | 0.78403 | 0.734082 | 1 |
| GOBP_NEUROTRANSMITTER_RECEPTOR_TRANSPORT | 41 | 0.227892 | 0.783566 | 0.846154 | 1 |
| GOBP_REGULATION_OF_CELL_CYCLE_ARREST | 103 | 0.195458 | 0.783529 | 0.967391 | 1 |
| GOBP_REGULATION_OF_ACTIN_FILAMENT_BASED_PROCESS | 378 | 0.172336 | 0.782692 | 1 | 1 |
| GOBP_REGULATION_OF_ERBB_SIGNALING_PATHWAY | 92 | 0.203123 | 0.782364 | 0.943548 | 1 |
| GOBP_NEGATIVE_REGULATION_OF_MONONUCLEAR_CELL_MIGRATION | 23 | 0.260944 | 0.781798 | 0.788845 | 1 |
| GOBP_RESPIRATORY_CHAIN_COMPLEX_IV_ASSEMBLY | 25 | 0.260542 | 0.781178 | 0.786008 | 1 |
| GOBP_INTERLEUKIN_5_PRODUCTION | 16 | 0.284076 | 0.781132 | 0.756667 | 1 |
| GOBP_REGULATION_OF_VASCULOGENESIS | 16 | 0.27925 | 0.780625 | 0.768116 | 1 |
| GOBP_REGULATION_OF_EPITHELIAL_TO_MESENCHYMAL_TRANSITION | 94 | 0.200663 | 0.780127 | 0.930435 | 1 |
| GOBP_TISSUE_MIGRATION | 307 | 0.178213 | 0.779994 | 1 | 1 |
| GOBP_VIRAL_RELEASE_FROM_HOST_CELL | 33 | 0.247426 | 0.778506 | 0.798077 | 1 |
| GOBP_POSITIVE_REGULATION_OF_PROTEIN_LOCALIZATION_TO_CELL_P | 65 | 0.208715 | 0.778012 | 0.917722 | 1 |
| GOBP_NUCLEOSIDE_PHOSPHATE_CATABOLIC_PROCESS | 80 | 0.202359 | 0.777976 | 0.900901 | 1 |
| GOBP_NEGATIVE_REGULATION_OF_CALCIUM_ION_TRANSMEMBRANE_T | 34 | 0.231113 | 0.77644 | 0.820513 | 1 |
| GOBP_NEGATIVE_REGULATION_OF_CELL_GROWTH | 178 | 0.188536 | 0.776428 | 0.978261 | 1 |
| GOBP_MULTI_ORGANISM_LOCALIZATION | 66 | 0.211174 | 0.776378 | 0.935484 | 1 |
| GOBP_REGULATION_OF_THE_FORCE_OF_HEART_CONTRACTION | 25 | 0.259754 | 0.775841 | 0.794521 | 1 |
| GOBP_RECEPTOR_CLUSTERING | 52 | 0.216289 | 0.775713 | 0.904192 | 1 |
| GOBP_SIGNAL_TRANSDUCTION_IN_RESPONSE_TO_DNA_DAMAGE | 128 | 0.189095 | 0.774771 | 0.973684 | 1 |
| GOBP_CYTOPLASMIC_SEQUESTERING_OF_TRANSCRIPTION_FACTOR | 15 | 0.292743 | 0.774024 | 0.787671 | 1 |
| GOBP_NEGATIVE_REGULATION_OF_COLD_INDUCED_THERMOGENESIS | 47 | 0.223656 | 0.773763 | 0.880435 | 1 |
| GOBP_REGULATION_OF_MEMBRANE_DEPOLARIZATION | 39 | 0.23674 | 0.773191 | 0.889423 | 1 |
| GOBP_POSITIVE_REGULATION_OF_DNA_DAMAGE_RESPONSE_SIGNAL_T | 15 | 0.28845 | 0.772583 | 0.775317 | 1 |
| GOBP_CHAPERONE_MEDIATED_PROTEIN_COMPLEX_ASSEMBLY | 21 | 0.265736 | 0.77249 | 0.783582 | 1 |
| GOBP_POSITIVE_REGULATION_OF_ENDOTHELIAL_CELL_MIGRATION | 115 | 0.186678 | 0.772263 | 0.945205 | 1 |
| GOBP_ENERGY_DERIVATION_BY_OXIDATION_OF_ORGANIC_COMPOUND | 269 | 0.166482 | 0.772176 | 1 | 1 |
| GOBP_IONOTROPIC_GLUTAMATE_RECEPTOR_SIGNALING_PATHWAY | 26 | 0.251819 | 0.772042 | 0.828571 | 1 |
| GOBP_TERMINATION_OF_RNA_POLYMERASE_II_TRANSCRIPTION | 35 | 0.240785 | 0.771867 | 0.876238 | 1 |
| GOBP_CELLULAR_RESPONSE_TO_REACTIVE_OXYGEN_SPECIES | 146 | 0.185042 | 0.771723 | 0.985507 | 1 |
| GOBP_NEURON_PROJECTION_ORGANIZATION | 89 | 0.196664 | 0.771384 | 0.916667 | 1 |
| GOBP_HISTONE_DEACETYLATION | 74 | 0.196665 | 0.771289 | 0.92 | 1 |
| GOBP_REGULATION_OF_RHO_PROTEIN_SIGNAL_TRANSDUCTION | 82 | 0.19946 | 0.770681 | 0.948718 | 1 |
| GOBP_POSITIVE_REGULATION_OF_POTASSIUM_ION_TRANSPORT | 44 | 0.22656 | 0.770169 | 0.810811 | 1 |
| GOBP_PROTEIN_LOCALIZATION_TO_MICROTUBULE | 18 | 0.270618 | 0.770163 | 0.793478 | 1 |
| GOBP_POSITIVE_REGULATION_OF_VASCULAR_ENDOTHELIAL_CELL_PR | 17 | 0.278654 | 0.76992 | 0.802974 | 1 |
| GOBP_ATP_SYNTHESIS_COUPLED_PROTON_TRANSPORT | 27 | 0.248076 | 0.769784 | 0.839827 | 1 |
| GOBP_REGULATION_OF_NUCLEASE_ACTIVITY | 22 | 0.27306 | 0.769426 | 0.812749 | 1 |
| GOBP_NEGATIVE_REGULATION_OF_CYTOKINE_PRODUCTION_INVOLVED | 24 | 0.248994 | 0.768519 | 0.827004 | 1 |
| GOBP_CELLULAR_RESPONSE_TO_ACID_CHEMICAL | 66 | 0.206691 | 0.768161 | 0.966443 | 1 |
| GOBP_CELLULAR_RESPONSE_TO_CAMP | 53 | 0.213453 | 0.768123 | 0.883041 | 1 |
| GOBP_PROTEIN_AUTOUBIQUITINATION | 72 | 0.199348 | 0.767973 | 0.972727 | 1 |
| GOBP_NEGATIVE_REGULATION_OF_AMYLOID_PRECURSOR_PROTEIN_C | 23 | 0.257218 | 0.767795 | 0.838462 | 1 |
| GOBP_REGULATION_OF_MUSCLE_HYPERTROPHY | 55 | 0.216157 | 0.766807 | 0.888889 | 1 |
| GOBP_ENDOTHELIAL_CELL_MIGRATION | 221 | 0.174002 | 0.766667 | 1 | 1 |
| GOBP_REGULATION_OF_PROTEIN_TARGETING_TO_MEMBRANE | 35 | 0.233424 | 0.766201 | 0.840909 | 1 |
| GOBP_PROTEIN_REFOLDING | 23 | 0.25775 | 0.765212 | 0.822835 | 1 |
| GOBP_ERROR_PRONE_TRANSLESION_SYNTHESIS | 21 | 0.256365 | 0.76425 | 0.811538 | 1 |
| GOBP_RETINA_VASCULATURE_DEVELOPMENT_IN_CAMERA_TYPE_EYE | 18 | 0.269986 | 0.763724 | 0.780488 | 1 |
| GOBP_CELLULAR_RESPONSE_TO_EXTERNAL_STIMULUS | 293 | 0.172736 | 0.763715 | 1 | 1 |
| GOBP_CHONDROITIN_SULFATE_BIOSYNTHETIC_PROCESS | 25 | 0.253343 | 0.763444 | 0.812766 | 1 |
| GOBP_VESICLE_LOCALIZATION | 223 | 0.171506 | 0.762004 | 1 | 1 |
| GOBP_SOMITOGENESIS | 60 | 0.209101 | 0.761707 | 0.908537 | 1 |
| GOBP_POSITIVE_REGULATION_OF_COAGULATION | 21 | 0.269015 | 0.760935 | 0.811475 | 1 |
| GOBP_MODIFIED_AMINO_ACID_TRANSPORT | 29 | 0.245491 | 0.760891 | 0.834043 | 1 |
| GOBP_NEGATIVE_REGULATION_OF_AUTOPHAGOSOME_ASSEMBLY | 15 | 0.280242 | 0.760849 | 0.812721 | 1 |
| GOBP_NEGATIVE_REGULATION_OF_LEUKOCYTE_MIGRATION | 43 | 0.225539 | 0.760714 | 0.887755 | 1 |
| GOBP_MAMMARY_GLAND_DUCT_MORPHOGENESIS | 28 | 0.237473 | 0.760646 | 0.863248 | 1 |
| GOBP_POSITIVE_REGULATION_OF_FIBROBLAST_PROLIFERATION | 45 | 0.213963 | 0.759945 | 0.926554 | 1 |
| GOBP_POSITIVE_REGULATION_OF_ENDOTHELIAL_CELL_CHEMOTAXIS | 15 | 0.285184 | 0.759751 | 0.798013 | 1 |
| GOBP_NADH_METABOLIC_PROCESS | 41 | 0.224519 | 0.758643 | 0.888325 | 1 |
| GOBP_CARDIOLIPIN_METABOLIC_PROCESS | 18 | 0.268654 | 0.758166 | 0.792 | 1 |
| GOBP_PROTEIN_FOLDING | 215 | 0.169559 | 0.757736 | 1 | 1 |
| GOBP_DNA_DOUBLE_STRAND_BREAK_PROCESSING | 24 | 0.24501 | 0.754796 | 0.86017 | 1 |
| GOBP_RESPONSE_TO_HYDROGEN_PEROXIDE | 126 | 0.179308 | 0.754535 | 0.986301 | 1 |
| GOBP_KERATINOCYTE_MIGRATION | 18 | 0.264597 | 0.753435 | 0.84585 | 1 |
| GOBP_INTRINSIC_APOPTOTIC_SIGNALING_PATHWAY_BY_P53_CLASS_M | 73 | 0.197341 | 0.752581 | 0.9375 | 1 |
| GOBP_POSITIVE_REGULATION_OF_STRIATED_MUSCLE_CELL_DIFFEREN | 40 | 0.221713 | 0.751542 | 0.897561 | 1 |
| GOBP_NEGATIVE_REGULATION_OF_PROTEIN_CATABOLIC_PROCESS | 114 | 0.185647 | 0.751058 | 1 | 1 |
| GOBP_ACTIVATION_OF_CYSTEINE_TYPE_ENDOPEPTIDASE_ACTIVITY_IN | 86 | 0.195443 | 0.750979 | 0.982609 | 1 |
| GOBP_REGULATION_OF_MITOCHONDRIAL_MEMBRANE_POTENTIAL | 70 | 0.1984 | 0.750655 | 0.97973 | 1 |
| GOBP_BRANCHING_INVOLVED_IN_MAMMARY_GLAND_DUCT_MORPHOG | 20 | 0.256895 | 0.749875 | 0.821293 | 1 |
| GOBP_NEGATIVE_REGULATION_OF_NIK_NF_KAPPAB_SIGNALING | 27 | 0.238628 | 0.749228 | 0.85124 | 1 |
| GOBP_RESPONSE_TO_AMINO_ACID_STARVATION | 49 | 0.216238 | 0.747441 | 0.857143 | 1 |
| GOBP_REGULATION_OF_CARDIAC_MUSCLE_CONTRACTION_BY_CALCIU | 25 | 0.244136 | 0.747427 | 0.850807 | 1 |
| GOBP_PROSTAGLANDIN_TRANSPORT | 17 | 0.271155 | 0.745337 | 0.822742 | 1 |
| GOBP_INTRINSIC_APOPTOTIC_SIGNALING_PATHWAY_IN_RESPONSE_TO | 43 | 0.215169 | 0.745256 | 0.925234 | 1 |
| GOBP_NEGATIVE_REGULATION_OF_RESPONSE_TO_WOUNDING | 80 | 0.199035 | 0.744839 | 0.944 | 1 |
| GOBP_NUCLEOBASE_CONTAINING_SMALL_MOLECULE_CATABOLIC_PRO | 48 | 0.215059 | 0.74434 | 0.881356 | 1 |
| GOBP_POSITIVE_REGULATION_OF_COLLAGEN_METABOLIC_PROCESS | 27 | 0.237396 | 0.743195 | 0.895833 | 1 |
| GOBP_ANGIOGENESIS_INVOLVED_IN_WOUND_HEALING | 25 | 0.251862 | 0.742359 | 0.834646 | 1 |
| GOBP_SNO_S_RNA_METABOLIC_PROCESS | 15 | 0.278792 | 0.742176 | 0.821192 | 1 |
| GOBP_POSITIVE_REGULATION_OF_CANONICAL_WNT_SIGNALING_PATHW | 142 | 0.177154 | 0.742095 | 1 | 1 |
| GOBP_PROTEIN_LOCALIZATION_TO_CHROMOSOME_TELOMERIC_REGIO | 30 | 0.235531 | 0.742057 | 0.851351 | 1 |
| GOBP_NEGATIVE_REGULATION_OF_UBIQUITIN_DEPENDENT_PROTEIN_C | 47 | 0.211648 | 0.739279 | 0.918239 | 1 |
| GOBP_PROTEIN_PEPTIDYL_PROLYL_ISOMERIZATION | 36 | 0.221572 | 0.739086 | 0.919192 | 1 |
| GOBP_MODULATION_OF_PROCESS_OF_OTHER_ORGANISM | 110 | 0.184885 | 0.738899 | 0.980952 | 1 |
| GOBP_SUBSTRATE_DEPENDENT_CELL_MIGRATION | 24 | 0.248107 | 0.738656 | 0.850202 | 1 |
| GOBP_INTRACELLULAR_ESTROGEN_RECEPTOR_SIGNALING_PATHWAY | 53 | 0.203156 | 0.738009 | 0.936047 | 1 |
| GOBP_HEAD_MORPHOGENESIS | 33 | 0.222401 | 0.737648 | 0.894495 | 1 |
| GOBP_POSITIVE_REGULATION_OF_DENDRITE_DEVELOPMENT | 15 | 0.269935 | 0.737454 | 0.824219 | 1 |
| GOBP_EPITHELIAL_TO_MESENCHYMAL_TRANSITION_INVOLVED_IN_END | 16 | 0.267416 | 0.737394 | 0.808664 | 1 |
| GOBP_STRESS_ACTIVATED_PROTEIN_KINASE_SIGNALING_CASCADE | 266 | 0.169564 | 0.736828 | 1 | 1 |
| GOBP_POSITIVE_REGULATION_OF_PROTEIN_AUTOPHOSPHORYLATION | 26 | 0.235244 | 0.73617 | 0.860465 | 1 |
| GOBP_NUCLEOBASE_CONTAINING_SMALL_MOLECULE_BIOSYNTHETIC_ | 112 | 0.182996 | 0.735338 | 1 | 1 |
| GOBP_REGULATION_OF_NITRIC_OXIDE_SYNTHASE_ACTIVITY | 47 | 0.213317 | 0.734166 | 0.892655 | 1 |
| GOBP_MATING_BEHAVIOR | 18 | 0.259672 | 0.733196 | 0.858736 | 1 |
| GOBP_FIBROBLAST_MIGRATION | 43 | 0.217805 | 0.729959 | 0.919355 | 1 |
| GOBP_POSITIVE_REGULATION_OF_NIK_NF_KAPPAB_SIGNALING | 66 | 0.192249 | 0.729954 | 0.948052 | 1 |
| GOBP_NEGATIVE_REGULATION_OF_RHO_PROTEIN_SIGNAL_TRANSDUC | 21 | 0.252306 | 0.729358 | 0.861004 | 1 |
| GOBP_PEPTIDYL_SERINE_MODIFICATION | 311 | 0.159204 | 0.729048 | 1 | 1 |
| GOBP_MEMBRANE_DEPOLARIZATION | 80 | 0.190237 | 0.72837 | 0.984849 | 1 |
| GOBP_NEGATIVE_REGULATION_OF_CALCINEURIN_MEDIATED_SIGNALIN | 15 | 0.27211 | 0.726971 | 0.862903 | 1 |
| GOBP_PIGMENTATION | 89 | 0.184816 | 0.726783 | 0.973214 | 1 |
| GOBP_NEGATIVE_REGULATION_OF_LIPID_STORAGE | 21 | 0.248885 | 0.726033 | 0.824818 | 1 |
| GOBP_TRANSLESION_SYNTHESIS | 42 | 0.215507 | 0.725498 | 0.918033 | 1 |
| GOBP_REGULATION_OF_PHOSPHATIDYLINOSITOL_3_KINASE_SIGNALING | 120 | 0.178225 | 0.724902 | 1 | 1 |
| GOBP_REGULATION_OF_SYSTEMIC_ARTERIAL_BLOOD_PRESSURE_BY_ | 22 | 0.246755 | 0.723567 | 0.846154 | 1 |
| GOBP_REGULATION_OF_NITRIC_OXIDE_METABOLIC_PROCESS | 54 | 0.201663 | 0.723219 | 0.937853 | 1 |
| GOBP_ESTABLISHMENT_OR_MAINTENANCE_OF_CELL_POLARITY | 210 | 0.165387 | 0.72251 | 1 | 1 |
| GOBP_REGULATION_OF_GLUCOSE_IMPORT | 51 | 0.200883 | 0.722185 | 0.921053 | 1 |
| GOBP_REGULATION_OF_JNK_CASCADE | 127 | 0.172154 | 0.722013 | 0.988235 | 1 |
| GOBP_UBIQUITIN_DEPENDENT_ERAD_PATHWAY | 79 | 0.189189 | 0.72187 | 0.992424 | 1 |
| GOBP_PROTEIN_STABILIZATION | 184 | 0.163893 | 0.721442 | 0.977273 | 1 |
| GOBP_JNK_CASCADE | 186 | 0.163187 | 0.721415 | 1 | 1 |
| GOBP_POSITIVE_REGULATION_OF_AXON_EXTENSION | 37 | 0.213033 | 0.721231 | 0.933628 | 1 |
| GOBP_LYMPHOID_PROGENITOR_CELL_DIFFERENTIATION | 20 | 0.243724 | 0.721151 | 0.855513 | 1 |
| GOBP_POSITIVE_REGULATION_OF_PRI_MIRNA_TRANSCRIPTION_BY_RN | 40 | 0.21404 | 0.720382 | 0.947115 | 1 |
| GOBP_INTRACILIARY_TRANSPORT | 53 | 0.20059 | 0.720256 | 0.967914 | 1 |
| GOBP_POSITIVE_REGULATION_OF_GLYCOGEN_METABOLIC_PROCESS | 16 | 0.265702 | 0.719981 | 0.830935 | 1 |
| GOBP_REGULATION_OF_GLUCOSE_TRANSMEMBRANE_TRANSPORT | 67 | 0.191526 | 0.719274 | 0.964539 | 1 |
| GOBP_REGULATION_OF_RESPONSE_TO_WOUNDING | 146 | 0.171295 | 0.719125 | 0.985507 | 1 |
| GOBP_PROTEIN_LOCALIZATION_TO_GOLGI_APPARATUS | 29 | 0.229959 | 0.718169 | 0.87013 | 1 |
| GOBP_VESICLE_CYTOSKELETAL_TRAFFICKING | 71 | 0.188283 | 0.717557 | 0.975 | 1 |
| GOBP_UBIQUINONE_METABOLIC_PROCESS | 18 | 0.255211 | 0.716366 | 0.849462 | 1 |
| GOBP_REGULATION_OF_PROTEASOMAL_UBIQUITIN_DEPENDENT_PROT | 130 | 0.170173 | 0.715383 | 0.988764 | 1 |
| GOBP_REGULATION_OF_GLUCAN_BIOSYNTHETIC_PROCESS | 27 | 0.232107 | 0.714251 | 0.875 | 1 |
| GOBP_RESPONSE_TO_IRON_ION | 27 | 0.227836 | 0.713754 | 0.910638 | 1 |
| GOBP_MITOCHONDRIAL_ATP_SYNTHESIS_COUPLED_PROTON_TRANSP | 22 | 0.248025 | 0.713655 | 0.879828 | 1 |
| GOBP_POSITIVE_REGULATION_OF_CHOLESTEROL_EFFLUX | 21 | 0.244437 | 0.713644 | 0.876812 | 1 |
| GOBP_CELLULAR_RESPONSE_TO_LIGHT_STIMULUS | 123 | 0.177704 | 0.713361 | 0.9875 | 1 |
| GOBP_REGULATION_OF_WOUND_HEALING | 117 | 0.179524 | 0.712565 | 0.989362 | 1 |
| GOBP_SECONDARY_METABOLITE_BIOSYNTHETIC_PROCESS | 20 | 0.24668 | 0.712241 | 0.887097 | 1 |
| GOBP_MITOCHONDRIAL_FISSION | 42 | 0.209442 | 0.711606 | 0.93617 | 1 |

| | | | | | |
|---|---|---|---|---|---|
| GOBP_TOLL_LIKE_RECEPTOR_SIGNALING_PATHWAY | 149 | -0.277418 | -0.888038 | 0.719525 | 1 |
| GOBP_LIPOSACCHARIDE_METABOLIC_PROCESS | 108 | -0.280058 | -0.887979 | 0.695797 | 1 |
| GOBP_MAINTENANCE_OF_CELL_POLARITY | 18 | -0.378148 | -0.887415 | 0.644898 | 1 |
| GOBP_INTERSTRAND_CROSS_LINK_REPAIR | 57 | -0.309424 | -0.886908 | 0.660628 | 1 |
| GOBP_INNERVATION | 23 | -0.358769 | -0.886471 | 0.637216 | 1 |
| GOBP_LIPID_PHOSPHORYLATION | 71 | -0.296438 | -0.886417 | 0.682726 | 1 |
| GOBP_RELAXATION_OF_MUSCLE | 28 | -0.345066 | -0.88626 | 0.637681 | 1 |
| GOBP_REGULATION_OF_ACUTE_INFLAMMATORY_RESPONSE | 45 | -0.315908 | -0.886257 | 0.653137 | 1 |
| GOBP_REGULATION_OF_CD4_POSITIVE_ALPHA_BETA_T_CELL_ACTIVATION | 64 | -0.302497 | -0.886257 | 0.662428 | 1 |
| GOBP_CELLULAR_RESPONSE_TO_RETINOIC_ACID | 64 | -0.301223 | -0.885792 | 0.662353 | 1 |
| GOBP_ADULT_LOCOMOTORY_BEHAVIOR | 60 | -0.302885 | -0.885337 | 0.665476 | 1 |
| GOBP_CELLULAR_METABOLIC_COMPOUND_SALVAGE | 34 | -0.33199 | -0.885229 | 0.680466 | 1 |
| GOBP_ACTIVATION_OF_JUN_KINASE_ACTIVITY | 39 | -0.325055 | -0.884531 | 0.644362 | 1 |
| GOBP_ASPARTATE_FAMILY_AMINO_ACID_BIOSYNTHETIC_PROCESS | 20 | -0.366048 | -0.883776 | 0.64248 | 1 |
| GOBP_CAMERA_TYPE_EYE_MORPHOGENESIS | 112 | -0.281934 | -0.883272 | 0.700779 | 1 |
| GOBP_DETECTION_OF_MECHANICAL_STIMULUS_INVOLVED_IN_SENSORY_PE | 31 | -0.333539 | -0.883017 | 0.624196 | 1 |
| GOBP_POSITIVE_REGULATION_OF_CATION_TRANSMEMBRANE_TRANSPORT | 136 | -0.277013 | -0.882911 | 0.711809 | 1 |
| GOBP_METAL_ION_TRANSPORT | 456 | -0.257582 | -0.882898 | 0.820281 | 1 |
| GOBP_POSITIVE_REGULATION_OF_GTPASE_ACTIVITY | 393 | -0.259766 | -0.882552 | 0.820565 | 1 |
| GOBP_MONOCARBOXYLIC_ACID_CATABOLIC_PROCESS | 129 | -0.27798 | -0.882018 | 0.730263 | 1 |
| GOBP_RESPONSE_TO_LEUKEMIA_INHIBITORY_FACTOR | 78 | -0.290202 | -0.881895 | 0.680556 | 1 |
| GOBP_SENSORY_ORGAN_DEVELOPMENT | 498 | -0.257035 | -0.881106 | 0.844689 | 1 |
| GOBP_ENDOCHONDRAL_BONE_MORPHOGENESIS | 51 | -0.308837 | -0.881024 | 0.665463 | 1 |
| GOBP_POSITIVE_REGULATION_OF_ACTIN_NUCLEATION | 15 | -0.385296 | -0.880576 | 0.625889 | 1 |
| GOBP_CENTRAL_NERVOUS_SYSTEM_PROJECTION_NEURON_AXONOGENESIS | 26 | -0.341877 | -0.879993 | 0.668831 | 1 |
| GOBP_POSITIVE_REGULATION_OF_HEMOPOIESIS | 148 | -0.274986 | -0.879409 | 0.740938 | 1 |
| GOBP_ASPARTATE_FAMILY_AMINO_ACID_CATABOLIC_PROCESS | 23 | -0.353292 | -0.878921 | 0.654206 | 1 |
| GOBP_RESPONSE_TO_ISCHEMIA | 47 | -0.315878 | -0.878629 | 0.665025 | 1 |
| GOBP_REGULATION_OF_OSTEOBLAST_PROLIFERATION | 24 | -0.348654 | -0.878372 | 0.655084 | 1 |
| GOBP_VITAMIN_TRANSPORT | 36 | -0.324301 | -0.878292 | 0.6525 | 1 |
| GOBP_POSITIVE_REGULATION_OF_CARBOHYDRATE_METABOLIC_PROCESS | 71 | -0.290242 | -0.877528 | 0.679245 | 1 |
| GOBP_NEGATIVE_REGULATION_OF_ERK1_AND_ERK2_CASCADE | 70 | -0.294657 | -0.877045 | 0.706358 | 1 |
| GOBP_REGULATION_OF_T_CELL_APOPTOTIC_PROCESS | 32 | -0.327602 | -0.876524 | 0.677707 | 1 |
| GOBP_ENERGY_HOMEOSTASIS | 35 | -0.328683 | -0.876451 | 0.645443 | 1 |
| GOBP_INFLAMMATORY_CELL_APOPTOTIC_PROCESS | 19 | -0.365639 | -0.876348 | 0.636111 | 1 |
| GOBP_PHOSPHOLIPID_METABOLIC_PROCESS | 402 | -0.259774 | -0.875944 | 0.824899 | 1 |
| GOBP_POSITIVE_REGULATION_OF_ERK1_AND_ERK2_CASCADE | 191 | -0.265361 | -0.875602 | 0.775079 | 1 |
| GOBP_CORTICAL_ACTIN_CYTOSKELETON_ORGANIZATION | 38 | -0.31762 | -0.875444 | 0.67362 | 1 |
| GOBP_T_CELL_LINEAGE_COMMITMENT | 26 | -0.347735 | -0.87536 | 0.667107 | 1 |
| GOBP_INTERFERON_ALPHA_PRODUCTION | 28 | -0.339214 | -0.875007 | 0.666223 | 1 |
| GOBP_REGULATION_OF_HEMOPOIESIS | 392 | -0.257924 | -0.874634 | 0.816532 | 1 |
| GOBP_POSITIVE_REGULATION_OF_ACUTE_INFLAMMATORY_RESPONSE | 26 | -0.345169 | -0.873595 | 0.677898 | 1 |
| GOBP_REGULATION_OF_CHEMOTAXIS | 212 | -0.264354 | -0.873229 | 0.764828 | 1 |
| GOBP_POSITIVE_REGULATION_OF_CALCIUM_ION_TRANSMEMBRANE_TRANSF | 68 | -0.293163 | -0.87317 | 0.725 | 1 |
| GOBP_REGULATION_OF_ADAPTIVE_IMMUNE_RESPONSE | 166 | -0.268591 | -0.873104 | 0.753977 | 1 |
| GOBP_POSITIVE_REGULATION_OF_GLIAL_CELL_PROLIFERATION | 16 | -0.378472 | -0.873088 | 0.65 | 1 |
| GOBP_CELLULAR_RESPONSE_TO_CARBOHYDRATE_STIMULUS | 125 | -0.27511 | -0.872988 | 0.725597 | 1 |
| GOBP_MITOTIC_METAPHASE_PLATE_CONGRESSION | 50 | -0.30474 | -0.872878 | 0.657635 | 1 |
| GOBP_OUTER_DYNEIN_ARM_ASSEMBLY | 17 | -0.367524 | -0.871928 | 0.660944 | 1 |
| GOBP_MATURATION_OF_LSU_RRNA_FROM_TRICISTRONIC_RRNA_TRANSCRIF | 15 | -0.378195 | -0.871721 | 0.663408 | 1 |
| GOBP_GLYCOSIDE_METABOLIC_PROCESS | 21 | -0.354222 | -0.871226 | 0.627778 | 1 |
| GOBP_RAP_PROTEIN_SIGNAL_TRANSDUCTION | 15 | -0.375637 | -0.871189 | 0.652299 | 1 |
| GOBP_3_UTR_MEDIATED_MRNA_DESTABILIZATION | 16 | -0.373309 | -0.87093 | 0.650499 | 1 |
| GOBP_AXO_DENDRITIC_TRANSPORT | 69 | -0.290179 | -0.870887 | 0.714452 | 1 |
| GOBP_REGULATION_OF_MITOCHONDRIAL_OUTER_MEMBRANE_PERMEABILIZ | 44 | -0.311322 | -0.870672 | 0.671216 | 1 |
| GOBP_MONOVALENT_INORGANIC_CATION_HOMEOSTASIS | 154 | -0.269409 | -0.870329 | 0.767417 | 1 |
| GOBP_INOSITOL_PHOSPHATE_BIOSYNTHETIC_PROCESS | 25 | -0.34246 | -0.870279 | 0.655172 | 1 |
| GOBP_MAMMARY_GLAND_DEVELOPMENT | 129 | -0.270544 | -0.869895 | 0.730473 | 1 |
| GOBP_PHENOL_CONTAINING_COMPOUND_METABOLIC_PROCESS | 97 | -0.281363 | -0.868137 | 0.729143 | 1 |
| GOBP_PROTEIN_INSERTION_INTO_MITOCHONDRIAL_MEMBRANE_INVOLVED_I | 30 | -0.330759 | -0.868044 | 0.671815 | 1 |
| GOBP_REGULATION_OF_STEM_CELL_POPULATION_MAINTENANCE | 29 | -0.33131 | -0.867399 | 0.679172 | 1 |
| GOBP_REGULATION_OF_MEGAKARYOCYTE_DIFFERENTIATION | 74 | -0.292046 | -0.867345 | 0.700578 | 1 |
| GOBP_POSITIVE_REGULATION_OF_CALCIUM_ION_TRANSMEMBRANE_TRANSF | 37 | -0.322353 | -0.866937 | 0.682957 | 1 |
| GOBP_CENTROSOME_DUPLICATION | 71 | -0.288833 | -0.866561 | 0.699177 | 1 |
| GOBP_REGULATION_OF_ACTIN_NUCLEATION | 30 | -0.329848 | -0.866018 | 0.69697 | 1 |
| GOBP_SPLICEOSOMAL_SNRNP_ASSEMBLY | 46 | -0.309455 | -0.865051 | 0.705314 | 1 |
| GOBP_KIDNEY_EPITHELIUM_DEVELOPMENT | 129 | -0.27204 | -0.864847 | 0.751092 | 1 |
| GOBP_REGULATION_OF_ACTIN_FILAMENT_LENGTH | 183 | -0.265061 | -0.863984 | 0.796842 | 1 |
| GOBP_NEPHRON_EPITHELIUM_DEVELOPMENT | 105 | -0.274441 | -0.863753 | 0.718232 | 1 |
| GOBP_PHOSPHATIDYLCHOLINE_CATABOLIC_PROCESS | 15 | -0.376539 | -0.862597 | 0.665722 | 1 |
| GOBP_RETINA_DEVELOPMENT_IN_CAMERA_TYPE_EYE | 140 | -0.268904 | -0.862247 | 0.742639 | 1 |
| GOBP_LONG_CHAIN_FATTY_ACYL_COA_BIOSYNTHETIC_PROCESS | 19 | -0.363545 | -0.861779 | 0.65722 | 1 |
| GOBP_NEGATIVE_REGULATION_OF_NEURON_DEATH | 186 | -0.262638 | -0.861773 | 0.773013 | 1 |
| GOBP_TETRAPYRROLE_BIOSYNTHETIC_PROCESS | 28 | -0.33429 | -0.861366 | 0.6781 | 1 |
| GOBP_POSITIVE_REGULATION_OF_LYASE_ACTIVITY | 18 | -0.363445 | -0.861218 | 0.668493 | 1 |
| GOBP_SPINAL_CORD_MOTOR_NEURON_DIFFERENTIATION | 24 | -0.340508 | -0.860691 | 0.669774 | 1 |
| GOBP_POSITIVE_REGULATION_OF_ACTIVATED_T_CELL_PROLIFERATION | 24 | -0.337239 | -0.860396 | 0.648438 | 1 |
| GOBP_SALIVARY_GLAND_DEVELOPMENT | 31 | -0.325441 | -0.859877 | 0.673858 | 1 |
| GOBP_LOCOMOTORY_BEHAVIOR | 151 | -0.265829 | -0.859091 | 0.749203 | 1 |
| GOBP_LONG_CHAIN_FATTY_ACYL_COA_METABOLIC_PROCESS | 25 | -0.344257 | -0.858802 | 0.696405 | 1 |
| GOBP_ETHER_METABOLIC_PROCESS | 26 | -0.334071 | -0.858648 | 0.674232 | 1 |
| GOBP_RIBOSOMAL_SMALL_SUBUNIT_ASSEMBLY | 21 | -0.358562 | -0.858532 | 0.661685 | 1 |
| GOBP_ORGANIC_HYDROXY_COMPOUND_BIOSYNTHETIC_PROCESS | 234 | -0.259125 | -0.857235 | 0.81321 | 1 |
| GOBP_POSITIVE_REGULATION_OF_TRANSMEMBRANE_TRANSPORT | 196 | -0.260599 | -0.85675 | 0.790747 | 1 |
| GOBP_COGNITION | 261 | -0.25625 | -0.856728 | 0.803498 | 1 |
| GOBP_RESPONSE_TO_ALKALOID | 92 | -0.279655 | -0.856488 | 0.736959 | 1 |
| GOBP_REGULATION_OF_DEFENSE_RESPONSE_TO_BACTERIUM | 15 | -0.372589 | -0.856488 | 0.662411 | 1 |
| GOBP_NON_RECOMBINATIONAL_REPAIR | 101 | -0.276871 | -0.855951 | 0.75419 | 1 |
| GOBP_POSITIVE_REGULATION_OF_CHEMOTAXIS | 135 | -0.267854 | -0.85585 | 0.761955 | 1 |
| GOBP_PYRIDINE_CONTAINING_COMPOUND_METABOLIC_PROCESS | 40 | -0.313826 | -0.855422 | 0.721945 | 1 |
| GOBP_MEMBRANE_LIPID_METABOLIC_PROCESS | 209 | -0.257378 | -0.854374 | 0.795407 | 1 |
| GOBP_MONOSACCHARIDE_BIOSYNTHETIC_PROCESS | 90 | -0.278673 | -0.854092 | 0.741163 | 1 |
| GOBP_INTERMEMBRANE_LIPID_TRANSFER | 44 | -0.302271 | -0.853626 | 0.706683 | 1 |
| GOBP_VESICLE_TRANSPORT_ALONG_MICROTUBULE | 46 | -0.301628 | -0.853526 | 0.688119 | 1 |
| GOBP_STEROL_HOMEOSTASIS | 77 | -0.286357 | -0.853417 | 0.70907 | 1 |
| GOBP_REGULATION_OF_CELL_DIVISION | 157 | -0.261037 | -0.853338 | 0.761653 | 1 |
| GOBP_MAINTENANCE_OF_CELL_NUMBER | 140 | -0.269138 | -0.853163 | 0.769313 | 1 |
| GOBP_CAMERA_TYPE_EYE_DEVELOPMENT | 290 | -0.254952 | -0.852573 | 0.840771 | 1 |
| GOBP_REGULATION_OF_GLIOGENESIS | 81 | -0.281748 | -0.852289 | 0.756849 | 1 |
| GOBP_ETHANOL_METABOLIC_PROCESS | 20 | -0.353474 | -0.852137 | 0.682065 | 1 |
| GOBP_REGULATION_OF_ASTROCYTE_DIFFERENTIATION | 21 | -0.34523 | -0.851734 | 0.677808 | 1 |
| GOBP_PROTEIN_KINASE_C_SIGNALING | 28 | -0.32624 | -0.851401 | 0.690415 | 1 |
| GOBP_CELL_CELL_ADHESION_MEDIATED_BY_INTEGRIN | 17 | -0.361511 | -0.850647 | 0.662483 | 1 |
| GOBP_SUBPALLIUM_DEVELOPMENT | 22 | -0.343983 | -0.849971 | 0.673497 | 1 |
| GOBP_NEGATIVE_REGULATION_OF_LEUKOCYTE_CELL_CELL_ADHESION | 125 | -0.26756 | -0.849428 | 0.773275 | 1 |
| GOBP_POSITIVE_REGULATION_OF_PROTEIN_POLYMERIZATION | 132 | -0.266831 | -0.848843 | 0.761803 | 1 |
| GOBP_INOSITOL_PHOSPHATE_METABOLIC_PROCESS | 68 | -0.284844 | -0.84829 | 0.726737 | 1 |
| GOBP_GLYCOSYLATION | 256 | -0.25531 | -0.848261 | 0.81893 | 1 |
| GOBP_PATTERN_RECOGNITION_RECEPTOR_SIGNALING_PATHWAY | 199 | -0.256543 | -0.847437 | 0.799583 | 1 |
| GOBP_REGULATION_OF_MAMMARY_GLAND_EPITHELIAL_CELL_PROLIFERATIO | 17 | -0.361761 | -0.847387 | 0.691576 | 1 |
| GOBP_REGULATION_OF_SMOOTH_MUSCLE_CELL_DIFFERENTIATION | 31 | -0.318213 | -0.8461 | 0.707379 | 1 |
| GOBP_REGULATION_OF_NUCLEAR_TRANSCRIBED_MRNA_CATABOLIC_PROC | 16 | -0.365926 | -0.845777 | 0.682192 | 1 |
| GOBP_POSITIVE_REGULATION_OF_ALPHA_BETA_T_CELL_ACTIVATION | 64 | -0.287581 | -0.84568 | 0.757432 | 1 |
| GOBP_CELLULAR_RESPONSE_TO_PROSTAGLANDIN_STIMULUS | 20 | -0.353222 | -0.84537 | 0.692109 | 1 |
| GOBP_TYROSINE_PHOSPHORYLATION_OF_STAT_PROTEIN | 75 | -0.282398 | -0.845203 | 0.741789 | 1 |
| GOBP_REGULATION_OF_MRNA_CATABOLIC_PROCESS | 200 | -0.255811 | -0.845062 | 0.821616 | 1 |
| GOBP_POSITIVE_REGULATION_OF_EPITHELIAL_CELL_DIFFERENTIATION | 50 | -0.302096 | -0.844785 | 0.730159 | 1 |
| GOBP_POSITIVE_REGULATION_OF_PEPTIDYL_THREONINE_PHOSPHORYLATIO | 28 | -0.327541 | -0.844666 | 0.693657 | 1 |
| GOBP_NEGATIVE_REGULATION_OF_CHEMOTAXIS | 58 | -0.293087 | -0.844385 | 0.732057 | 1 |
| GOBP_RECEPTOR_SIGNALING_PATHWAY_VIA_STAT | 143 | -0.262743 | -0.84418 | 0.793869 | 1 |
| GOBP_GLYCEROLIPID_BIOSYNTHETIC_PROCESS | 255 | -0.251422 | -0.843621 | 0.851546 | 1 |
| GOBP_MAINTENANCE_OF_GASTROINTESTINAL_EPITHELIUM | 20 | -0.349518 | -0.843546 | 0.718876 | 1 |
| GOBP_DNA_STRAND_ELONGATION_INVOLVED_IN_DNA_REPLICATION | 19 | -0.35366 | -0.843346 | 0.680614 | 1 |
| GOBP_L_ALPHA_AMINO_ACID_TRANSMEMBRANE_TRANSPORT | 60 | -0.289331 | -0.843249 | 0.737981 | 1 |
| GOBP_EMBRYONIC_SKELETAL_SYSTEM_DEVELOPMENT | 115 | -0.267974 | -0.842534 | 0.774054 | 1 |
| GOBP_REGULATION_OF_CELLULAR_RESPIRATION | 21 | -0.340091 | -0.842404 | 0.688202 | 1 |
| GOBP_REGULATION_OF_MICROTUBULE_POLYMERIZATION | 50 | -0.293787 | -0.842267 | 0.706167 | 1 |
| GOBP_DETECTION_OF_ABIOTIC_STIMULUS | 130 | -0.263163 | -0.842249 | 0.792696 | 1 |
| GOBP_REGULATION_OF_LIPID_BIOSYNTHETIC_PROCESS | 183 | -0.258339 | -0.842246 | 0.810437 | 1 |
| GOBP_NUCLEOTIDE_BINDING_OLIGOMERIZATION_DOMAIN_CONTAINING_2_SI | 16 | -0.359574 | -0.841092 | 0.707015 | 1 |
| GOBP_EOSINOPHIL_CHEMOTAXIS | 17 | -0.359283 | -0.841042 | 0.705489 | 1 |
| GOBP_POSITIVE_REGULATION_OF_EPITHELIAL_CELL_PROLIFERATION | 184 | -0.254333 | -0.840962 | 0.820296 | 1 |
| GOBP_NEGATIVE_REGULATION_OF_CYTOKINE_PRODUCTION | 278 | -0.247716 | -0.840272 | 0.861366 | 1 |
| GOBP_NEPHRON_DEVELOPMENT | 136 | -0.260749 | -0.840229 | 0.793326 | 1 |
| GOBP_INORGANIC_ANION_TRANSMEMBRANE_TRANSPORT | 111 | -0.267201 | -0.839138 | 0.763657 | 1 |
| GOBP_CILIUM_MOVEMENT | 146 | -0.261893 | -0.839107 | 0.811625 | 1 |
| GOBP_NEGATIVE_REGULATION_OF_MUSCLE_CELL_DIFFERENTIATION | 56 | -0.290876 | -0.838893 | 0.730588 | 1 |
| GOBP_CENTRAL_NERVOUS_SYSTEM_NEURON_AXONOGENESIS | 34 | -0.310805 | -0.838482 | 0.714643 | 1 |

| | | | | | |
|---|---|---|---|---|---|
| GOBP_NEGATIVE_REGULATION_OF_PROTEASOMAL_PROTEIN_CATABOL | 47 | 0.20421 | 0.710141 | 0.93617 | 1 |
| GOBP_REGULATION_OF_CD8_POSITIVE_ALPHA_BETA_T_CELL_ACTIVAT | 19 | 0.259379 | 0.709861 | 0.838951 | 1 |
| GOBP_REGULATION_OF_STRESS_ACTIVATED_PROTEIN_KINASE_SIGNAL | 176 | 0.165593 | 0.70804 | 1 | 1 |
| GOBP_INTRACILIARY_TRANSPORT_INVOLVED_IN_CILIUM_ASSEMBLY | 40 | 0.210028 | 0.707943 | 0.945274 | 1 |
| GOBP_POSITIVE_REGULATION_OF_NITRIC_OXIDE_SYNTHASE_ACTIVITY | 23 | 0.23395 | 0.707034 | 0.877049 | 1 |
| GOBP_RNA_CAPPING | 34 | 0.218894 | 0.706703 | 0.940594 | 1 |
| GOBP_REGULATION_OF_WNT_SIGNALING_PATHWAY | 353 | 0.153493 | 0.706663 | 1 | 1 |
| GOBP_POSTREPLICATION_REPAIR | 52 | 0.197221 | 0.705528 | 0.955128 | 1 |
| GOBP_PLATELET_AGGREGATION | 62 | 0.185352 | 0.704457 | 0.947368 | 1 |
| GOBP_NEGATIVE_REGULATION_OF_CATABOLIC_PROCESS | 293 | 0.153632 | 0.7032 | 1 | 1 |
| GOBP_CELL_CELL_SIGNALING_BY_WNT | 499 | 0.151632 | 0.702171 | 1 | 1 |
| GOBP_POLYOL_BIOSYNTHETIC_PROCESS | 61 | 0.19051 | 0.701368 | 0.976744 | 1 |
| GOBP_REGULATION_OF_VOLTAGE_GATED_CALCIUM_CHANNEL_ACTIVIT | 33 | 0.212987 | 0.701239 | 0.91 | 1 |
| GOBP_VESICLE_MEDIATED_TRANSPORT_TO_THE_PLASMA_MEMBRANE | 91 | 0.173869 | 0.700182 | 0.981482 | 1 |
| GOBP_NEGATIVE_REGULATION_OF_SMOOTHENED_SIGNALING_PATHWA | 31 | 0.22576 | 0.699993 | 0.948936 | 1 |
| GOBP_TELOMERASE_RNA_LOCALIZATION | 19 | 0.245539 | 0.69885 | 0.891386 | 1 |
| GOBP_NEGATIVE_REGULATION_OF_NUCLEOCYTOPLASMIC_TRANSPORT | 20 | 0.230423 | 0.698621 | 0.878788 | 1 |
| GOBP_TRICARBOXYLIC_ACID_CYCLE | 33 | 0.213458 | 0.698599 | 0.914692 | 1 |
| GOBP_MAINTENANCE_OF_BLOOD_BRAIN_BARRIER | 33 | 0.218359 | 0.698531 | 0.943478 | 1 |
| GOBP_POSITIVE_REGULATION_OF_GLUCOSE_IMPORT | 33 | 0.214933 | 0.697528 | 0.932692 | 1 |
| GOBP_STRIATUM_DEVELOPMENT | 17 | 0.255699 | 0.697444 | 0.860294 | 1 |
| GOBP_REGULATION_OF_ENDOTHELIAL_CELL_DIFFERENTIATION | 33 | 0.213849 | 0.697013 | 0.934426 | 1 |
| GOBP_INTRA_GOLGI_VESICLE_MEDIATED_TRANSPORT | 34 | 0.211625 | 0.696572 | 0.926829 | 1 |
| GOBP_PEPTIDYL_SERINE_DEPHOSPHORYLATION | 19 | 0.245761 | 0.695741 | 0.869403 | 1 |
| GOBP_POSITIVE_REGULATION_OF_RESPONSE_TO_CYTOKINE_STIMULU | 56 | 0.191366 | 0.695619 | 0.974684 | 1 |
| GOBP_REGULATION_OF_OXIDATIVE_STRESS_INDUCED_INTRINSIC_APO | 27 | 0.221503 | 0.694838 | 0.915556 | 1 |
| GOBP_PROTEIN_DEMETHYLATION | 31 | 0.219001 | 0.693106 | 0.929204 | 1 |
| GOBP_UDP_N_ACETYLGLUCOSAMINE_METABOLIC_PROCESS | 16 | 0.25606 | 0.692987 | 0.847518 | 1 |
| GOBP_POSITIVE_REGULATION_OF_RESPONSE_TO_WOUNDING | 61 | 0.186807 | 0.692621 | 0.93865 | 1 |
| GOBP_REGULATION_OF_GOLGI_ORGANIZATION | 17 | 0.248629 | 0.692251 | 0.869231 | 1 |
| GOBP_RNA_SPLICING_VIA_ENDONUCLEOLYTIC_CLEAVAGE_AND_LIGAT | 16 | 0.262124 | 0.691061 | 0.89313 | 1 |
| GOBP_POSITIVE_REGULATION_OF_PROTEIN_DEACETYLATION | 22 | 0.237951 | 0.690322 | 0.883459 | 1 |
| GOBP_RESPONSE_TO_EPIDERMAL_GROWTH_FACTOR | 46 | 0.200075 | 0.68992 | 0.952096 | 1 |
| GOBP_REGULATION_OF_PROTEIN_LOCALIZATION_TO_PLASMA_MEMBRA | 96 | 0.177656 | 0.689003 | 1 | 1 |
| GOBP_NEGATIVE_REGULATION_OF_CALCIUM_MEDIATED_SIGNALING | 19 | 0.240729 | 0.688386 | 0.876712 | 1 |
| GOBP_POSITIVE_REGULATION_OF_MONOOXYGENASE_ACTIVITY | 31 | 0.215558 | 0.688235 | 0.932203 | 1 |
| GOBP_ACTIN_FILAMENT_SEVERING | 16 | 0.246163 | 0.688049 | 0.897887 | 1 |
| GOBP_REGULATION_OF_CHROMATIN_BINDING | 22 | 0.230561 | 0.687708 | 0.889362 | 1 |
| GOBP_POSITIVE_REGULATION_BY_HOST_OF_VIRAL_TRANSCRIPTION | 16 | 0.25132 | 0.687507 | 0.887641 | 1 |
| GOBP_PROTEIN_DEGLYCOSYLATION | 28 | 0.220612 | 0.687127 | 0.929752 | 1 |
| GOBP_REGULATION_OF_GLUCONEOGENESIS | 43 | 0.201459 | 0.686623 | 0.951872 | 1 |
| GOBP_MODULATION_BY_HOST_OF_SYMBIONT_PROCESS | 60 | 0.18786 | 0.683077 | 0.965517 | 1 |
| GOBP_REGULATION_OF_NIK_NF_KAPPAB_SIGNALING | 95 | 0.170725 | 0.682327 | 0.991071 | 1 |
| GOBP_POSITIVE_REGULATION_OF_SKELETAL_MUSCLE_TISSUE_DEVELC | 17 | 0.243891 | 0.681414 | 0.885376 | 1 |
| GOBP_REGULATION_OF_MEMBRANE_INVAGINATION | 16 | 0.243964 | 0.680143 | 0.885185 | 1 |
| GOBP_POSITIVE_REGULATION_OF_MUSCLE_CELL_DIFFERENTIATION | 69 | 0.181345 | 0.679638 | 0.993421 | 1 |
| GOBP_POSITIVE_REGULATION_OF_LEUKOCYTE_ADHESION_TO_VASCUL | 20 | 0.228368 | 0.679242 | 0.903226 | 1 |
| GOBP_NEGATIVE_REGULATION_OF_CYCLIN_DEPENDENT_PROTEIN_KIN | 30 | 0.210197 | 0.678706 | 0.944 | 1 |
| GOBP_PURINE_NUCLEOTIDE_TRANSPORT | 24 | 0.230664 | 0.675579 | 0.929461 | 1 |
| GOBP_GLYCOLYTIC_PROCESS_THROUGH_FRUCTOSE_6_PHOSPHATE | 27 | 0.220767 | 0.674972 | 0.930894 | 1 |
| GOBP_REGULATION_OF_MRNA_3_END_PROCESSING | 28 | 0.220328 | 0.674848 | 0.922764 | 1 |
| GOBP_EMBRYONIC_HEMOPOIESIS | 22 | 0.226125 | 0.672251 | 0.947368 | 1 |
| GOBP_NEGATIVE_REGULATION_OF_PROTEIN_SERINE_THREONINE_KIN | 132 | 0.159658 | 0.671558 | 1 | 1 |
| GOBP_NEGATIVE_REGULATION_OF_CELL_CYCLE_G1_S_PHASE_TRANS | 103 | 0.167365 | 0.67145 | 0.990476 | 1 |
| GOBP_NEGATIVE_REGULATION_OF_INTRACELLULAR_SIGNAL_TRANSDU | 476 | 0.151781 | 0.671263 | 1 | 1 |
| GOBP_ERYTHROCYTE_HOMEOSTASIS | 119 | 0.163007 | 0.67037 | 1 | 1 |
| GOBP_REGULATION_OF_CYSTEINE_TYPE_ENDOPEPTIDASE_ACTIVITY_IN | 17 | 0.241603 | 0.670216 | 0.854406 | 1 |
| GOBP_PIGMENT_CELL_DIFFERENTIATION | 32 | 0.207956 | 0.67021 | 0.933054 | 1 |
| GOBP_REGULATION_OF_PROTEIN_LOCALIZATION_TO_CELL_SURFACE | 38 | 0.200096 | 0.669801 | 0.959538 | 1 |
| GOBP_REGULATION_OF_NECROTIC_CELL_DEATH | 33 | 0.209746 | 0.669566 | 0.942857 | 1 |
| GOBP_GOLGI_TO_PLASMA_MEMBRANE_TRANSPORT | 57 | 0.184633 | 0.669438 | 0.982143 | 1 |
| GOBP_ESTABLISHMENT_OF_PROTEIN_LOCALIZATION_TO_VACUOLE | 48 | 0.187846 | 0.668186 | 0.930636 | 1 |
| GOBP_RAB_PROTEIN_SIGNAL_TRANSDUCTION | 21 | 0.222426 | 0.668007 | 0.916031 | 1 |
| GOBP_POSITIVE_REGULATION_OF_T_HELPER_CELL_DIFFERENTIATION | 21 | 0.225958 | 0.667956 | 0.908397 | 1 |
| GOBP_MITOCHONDRIAL_ELECTRON_TRANSPORT_CYTOCHROME_C_TO | 21 | 0.227876 | 0.667738 | 0.92 | 1 |
| GOBP_PHOSPHATIDYLGLYCEROL_METABOLIC_PROCESS | 36 | 0.206682 | 0.66643 | 0.985916 | 1 |
| GOBP_DNA_DAMAGE_RESPONSE_DETECTION_OF_DNA_DAMAGE | 38 | 0.199816 | 0.663909 | 0.955 | 1 |
| GOBP_CELL_ADHESION_MOLECULE_PRODUCTION | 15 | 0.245481 | 0.663443 | 0.887324 | 1 |
| GOBP_PRIMARY_NEURAL_TUBE_FORMATION | 88 | 0.174832 | 0.662976 | 1 | 1 |
| GOBP_POSITIVE_REGULATION_OF_GLUCOSE_TRANSMEMBRANE_TRANS | 39 | 0.194138 | 0.662536 | 0.946429 | 1 |
| GOBP_POSITIVE_REGULATION_OF_WNT_SIGNALING_PATHWAY | 174 | 0.152702 | 0.662001 | 1 | 1 |
| GOBP_CHEMOKINE_C_X_C_MOTIF_LIGAND_2_PRODUCTION | 15 | 0.254388 | 0.661213 | 0.878049 | 1 |
| GOBP_STRESS_FIBER_ASSEMBLY | 101 | 0.166879 | 0.660974 | 0.990291 | 1 |
| GOBP_REGULATION_OF_MITOCHONDRIAL_GENE_EXPRESSION | 29 | 0.210425 | 0.660767 | 0.944206 | 1 |
| GOBP_REGULATION_OF_INCLUSION_BODY_ASSEMBLY | 15 | 0.244682 | 0.658434 | 0.9319 | 1 |
| GOBP_MODULATION_BY_SYMBIONT_OF_HOST_CELLULAR_PROCESS | 24 | 0.224506 | 0.658311 | 0.919492 | 1 |
| GOBP_REGULATION_OF_RAS_PROTEIN_SIGNAL_TRANSDUCTION | 187 | 0.155382 | 0.658071 | 1 | 1 |
| GOBP_LYMPHANGIOGENESIS | 17 | 0.234643 | 0.658047 | 0.931035 | 1 |
| GOBP_POSITIVE_REGULATION_OF_MITOCHONDRIAL_FISSION | 17 | 0.241102 | 0.657195 | 0.894531 | 1 |
| GOBP_REGULATION_OF_TRANSLATION_IN_RESPONSE_TO_STRESS | 21 | 0.224847 | 0.656501 | 0.941406 | 1 |
| GOBP_PYRIMIDINE_NUCLEOSIDE_TRIPHOSPHATE_METABOLIC_PROCES | 24 | 0.216053 | 0.653474 | 0.946619 | 1 |
| GOBP_REGULATION_OF_MACROAUTOPHAGY | 160 | 0.154678 | 0.653113 | 1 | 1 |
| GOBP_PYRIMIDINE_RIBONUCLEOSIDE_TRIPHOSPHATE_METABOLIC_PR | 18 | 0.231401 | 0.651934 | 0.91958 | 1 |
| GOBP_PROTEIN_TARGETING_TO_LYSOSOME | 23 | 0.227142 | 0.651564 | 0.925926 | 1 |
| GOBP_MULTIVESICULAR_BODY_ORGANIZATION | 31 | 0.203154 | 0.648481 | 0.977778 | 1 |
| GOBP_REGULATION_OF_CELL_SUBSTRATE_JUNCTION_ORGANIZATION | 68 | 0.177193 | 0.647787 | 0.991597 | 1 |
| GOBP_CELLULAR_RESPONSE_TO_MECHANICAL_STIMULUS | 73 | 0.17038 | 0.647526 | 1 | 1 |
| GOBP_NEURAL_TUBE_FORMATION | 96 | 0.160245 | 0.647164 | 0.991453 | 1 |
| GOBP_REGULATION_OF_DNA_DAMAGE_CHECKPOINT | 15 | 0.239483 | 0.646962 | 0.911765 | 1 |
| GOBP_NEGATIVE_REGULATION_OF_INTERLEUKIN_10_PRODUCTION | 19 | 0.223893 | 0.646576 | 0.909434 | 1 |
| GOBP_REGULATION_OF_SKELETAL_MUSCLE_CELL_DIFFERENTIATION | 18 | 0.225523 | 0.645601 | 0.922535 | 1 |
| GOBP_HISTONE_UBIQUITINATION | 46 | 0.179939 | 0.644215 | 1 | 1 |
| GOBP_MITOTIC_G2_DNA_DAMAGE_CHECKPOINT | 23 | 0.214809 | 0.644087 | 0.93254 | 1 |
| GOBP_HISTONE_MONOUBIQUITINATION | 29 | 0.207487 | 0.641462 | 0.952 | 1 |
| GOBP_MITOCHONDRIAL_FUSION | 28 | 0.203701 | 0.640529 | 0.933333 | 1 |
| GOBP_REGULATION_OF_PROTEIN_SUMOYLATION | 22 | 0.211634 | 0.640179 | 0.957265 | 1 |
| GOBP_NEGATIVE_REGULATION_OF_DNA_METABOLIC_PROCESS | 118 | 0.1583 | 0.63907 | 1 | 1 |
| GOBP_ENDOPLASMIC_RETICULUM_TO_GOLGI_VESICLE_MEDIATED_TRA | 202 | 0.143374 | 0.638811 | 1 | 1 |
| GOBP_LYMPH_VESSEL_DEVELOPMENT | 26 | 0.213076 | 0.638698 | 0.953846 | 1 |
| GOBP_NEGATIVE_REGULATION_OF_PEPTIDYL_SERINE_PHOSPHORYLAT | 28 | 0.202096 | 0.637648 | 0.949791 | 1 |
| GOBP_PEPTIDYL_LYSINE_TRIMETHYLATION | 43 | 0.186455 | 0.637046 | 1 | 1 |
| GOBP_PROTEIN_TARGETING_TO_VACUOLE | 40 | 0.187784 | 0.633595 | 0.968421 | 1 |
| GOBP_TRANSFORMING_GROWTH_FACTOR_BETA_RECEPTOR_SIGNALIN | 185 | 0.146489 | 0.633336 | 1 | 1 |
| GOBP_MACROMOLECULE_DEACYLATION | 105 | 0.154924 | 0.628464 | 1 | 1 |
| GOBP_PROTEIN_DEMANNOSYLATION | 20 | 0.221457 | 0.627281 | 0.9375 | 1 |
| GOBP_GOLGI_TO_ENDOSOME_TRANSPORT | 17 | 0.224769 | 0.627121 | 0.948905 | 1 |
| GOBP_HISTONE_H4_K5_ACETYLATION | 16 | 0.228074 | 0.626306 | 0.937729 | 1 |
| GOBP_ESTABLISHMENT_OF_TISSUE_POLARITY | 122 | 0.150738 | 0.622901 | 1 | 1 |
| GOBP_REGULATION_OF_GRANULOCYTE_DIFFERENTIATION | 15 | 0.232116 | 0.622755 | 0.907407 | 1 |
| GOBP_MAINTENANCE_OF_PROTEIN_LOCALIZATION_IN_ORGANELLE | 42 | 0.182192 | 0.622689 | 0.97619 | 1 |
| GOBP_NUCLEOTIDE_EXCISION_REPAIR_DNA_DAMAGE_RECOGNITION | 23 | 0.203036 | 0.622342 | 0.954751 | 1 |
| GOBP_REGULATION_OF_URINE_VOLUME | 20 | 0.212094 | 0.620558 | 0.958015 | 1 |
| GOBP_NEGATIVE_REGULATION_OF_PEPTIDYL_LYSINE_ACETYLATION | 19 | 0.220142 | 0.619431 | 0.953069 | 1 |
| GOBP_NUCLEOSIDE_MONOPHOSPHATE_BIOSYNTHETIC_PROCESS | 41 | 0.183586 | 0.619052 | 0.979487 | 1 |
| GOBP_NEGATIVE_REGULATION_OF_PROTEIN_ACETYLATION | 21 | 0.207636 | 0.618728 | 0.937269 | 1 |
| GOBP_COLLAGEN_BIOSYNTHETIC_PROCESS | 45 | 0.175626 | 0.618165 | 0.975124 | 1 |
| GOBP_REGULATION_OF_CELL_PROJECTION_ASSEMBLY | 181 | 0.141634 | 0.61729 | 1 | 1 |
| GOBP_NEUROTROPHIN_TRK_RECEPTOR_SIGNALING_PATHWAY | 32 | 0.193664 | 0.615258 | 0.971015 | 1 |
| GOBP_REGULATION_OF_PROTEIN_AUTOPHOSPHORYLATION | 46 | 0.17719 | 0.614861 | 0.98125 | 1 |
| GOBP_THYMOCYTE_APOPTOTIC_PROCESS | 16 | 0.221 | 0.613979 | 0.945017 | 1 |
| GOBP_NEGATIVE_REGULATION_OF_INTRACELLULAR_TRANSPORT | 53 | 0.168374 | 0.611006 | 0.988506 | 1 |
| GOBP_REGULATION_OF_CELL_CYCLE_G1_S_PHASE_TRANSITION | 162 | 0.141598 | 0.610254 | 1 | 1 |
| GOBP_MICROTUBULE_ORGANIZING_CENTER_LOCALIZATION | 30 | 0.189118 | 0.608738 | 0.982833 | 1 |
| GOBP_ENDOTHELIAL_CELL_APOPTOTIC_PROCESS | 54 | 0.171946 | 0.608284 | 0.986395 | 1 |
| GOBP_ENDOPLASMIC_RETICULUM_MANNOSE_TRIMMING | 16 | 0.220503 | 0.608139 | 0.932308 | 1 |
| GOBP_LAMELLIPODIUM_MORPHOGENESIS | 16 | 0.223036 | 0.606949 | 0.954225 | 1 |
| GOBP_RESPONSE_TO_HEPATOCYTE_GROWTH_FACTOR | 17 | 0.219858 | 0.605792 | 0.955224 | 1 |
| GOBP_VESICLE_CARGO_LOADING | 25 | 0.198832 | 0.60379 | 0.95339 | 1 |
| GOBP_NEGATIVE_REGULATION_OF_RECEPTOR_SIGNALING_PATHWAY_ | 22 | 0.208564 | 0.602345 | 0.970149 | 1 |
| GOBP_PROTEIN_ACYLATION | 232 | 0.138828 | 0.602139 | 1 | 1 |
| GOBP_MYELOID_CELL_HOMEOSTASIS | 143 | 0.141365 | 0.601606 | 1 | 1 |
| GOBP_NEGATIVE_REGULATION_OF_INTERFERON_GAMMA_PRODUCTION | 36 | 0.181913 | 0.600111 | 1 | 1 |
| GOBP_LIPID_DROPLET_ORGANIZATION | 27 | 0.188285 | 0.598805 | 0.959821 | 1 |
| GOBP_POSITIVE_REGULATION_OF_EPITHELIAL_CELL_APOPTOTIC_PROC | 32 | 0.187296 | 0.598629 | 0.995283 | 1 |
| GOBP_PROTEIN_DNA_COMPLEX_DISASSEMBLY | 18 | 0.208915 | 0.5967 | 0.968992 | 1 |
| GOBP_CELLULAR_RESPONSE_TO_TOPOLOGICALLY_INCORRECT_PROTI | 164 | 0.13882 | 0.596508 | 1 | 1 |
| GOBP_GLUTAMINE_FAMILY_AMINO_ACID_BIOSYNTHETIC_PROCESS | 17 | 0.2167 | 0.595656 | 0.978947 | 1 |
| GOBP_MICROTUBULE_DEPOLYMERIZATION | 43 | 0.17259 | 0.59416 | 0.994764 | 1 |

| | | | | | |
|---|---|---|---|---|---|
| GOBP_ERBB_SIGNALING_PATHWAY | 141 | -0.262555 | -0.83848 | 0.802139 | 1 |
| GOBP_POSITIVE_REGULATION_OF_VASCULAR_ASSOCIATED_SMOOTH_MUSC | 16 | -0.36399 | -0.838422 | 0.701228 | 1 |
| GOBP_SKELETAL_SYSTEM_MORPHOGENESIS | 204 | -0.254863 | -0.838052 | 0.845041 | 1 |
| GOBP_POSITIVE_REGULATION_OF_MYELOID_LEUKOCYTE_DIFFERENTIATION | 55 | -0.291326 | -0.83799 | 0.731618 | 1 |
| GOBP_GLYCEROL_ETHER_METABOLIC_PROCESS | 21 | -0.339016 | -0.83737 | 0.72973 | 1 |
| GOBP_POSITIVE_REGULATION_OF_NEUROBLAST_PROLIFERATION | 20 | -0.34955 | -0.837225 | 0.718626 | 1 |
| GOBP_ORGANIC_ACID_CATABOLIC_PROCESS | 253 | -0.250939 | -0.836734 | 0.855236 | 1 |
| GOBP_BODY_FLUID_SECRETION | 80 | -0.277338 | -0.83617 | 0.767689 | 1 |
| GOBP_METAPHASE_PLATE_CONGRESSION | 63 | -0.285277 | -0.83582 | 0.736534 | 1 |
| GOBP_POSITIVE_REGULATION_OF_TELOMERE_CAPPING | 17 | -0.355996 | -0.835793 | 0.682561 | 1 |
| GOBP_NEGATIVE_REGULATION_OF_RESPONSE_TO_BIOTIC_STIMULUS | 93 | -0.270921 | -0.835445 | 0.767735 | 1 |
| GOBP_REGULATION_OF_G_PROTEIN_COUPLED_RECEPTOR_SIGNALING_PATH | 133 | -0.262595 | -0.835321 | 0.808556 | 1 |
| GOBP_NEGATIVE_REGULATION_OF_SMOOTH_MUSCLE_CELL_MIGRATION | 21 | -0.338691 | -0.835141 | 0.716033 | 1 |
| GOBP_EMBRYONIC_HINDLIMB_MORPHOGENESIS | 26 | -0.322325 | -0.834776 | 0.718708 | 1 |
| GOBP_REGULATION_OF_CALCIUM_MEDIATED_SIGNALING | 63 | -0.285414 | -0.834607 | 0.748517 | 1 |
| GOBP_INSULIN_SECRETION_INVOLVED_IN_CELLULAR_RESPONSE_TO_GLUC | 61 | -0.286859 | -0.83456 | 0.747929 | 1 |
| GOBP_NLRP3_INFLAMMASOME_COMPLEX_ASSEMBLY | 15 | -0.362264 | -0.834436 | 0.705556 | 1 |
| GOBP_LIPID_LOCALIZATION | 470 | -0.244907 | -0.834377 | 0.903226 | 1 |
| GOBP_REGULATION_OF_EARLY_ENDOSOME_TO_LATE_ENDOSOME_TRANSP | 18 | -0.347842 | -0.834298 | 0.693351 | 1 |
| GOBP_GLUTAMATE_METABOLIC_PROCESS | 30 | -0.319341 | -0.834098 | 0.710938 | 1 |
| GOBP_NUCLEIC_ACID_PHOSPHODIESTER_BOND_HYDROLYSIS | 289 | -0.249303 | -0.833878 | 0.868852 | 1 |
| GOBP_POSITIVE_REGULATION_OF_CELLULAR_CARBOHYDRATE_METABOLIC_ | 48 | -0.293852 | -0.833574 | 0.739237 | 1 |
| GOBP_GLYCOSPHINGOLIPID_BIOSYNTHETIC_PROCESS | 27 | -0.324206 | -0.833364 | 0.714474 | 1 |
| GOBP_LIPID_MODIFICATION | 264 | -0.250644 | -0.832813 | 0.870276 | 1 |
| GOBP_ANIMAL_ORGAN_MATURATION | 25 | -0.325537 | -0.83275 | 0.706112 | 1 |
| GOBP_EPIDERMAL_GROWTH_FACTOR_RECEPTOR_SIGNALING_PATHWAY | 119 | -0.264147 | -0.831835 | 0.784141 | 1 |
| GOBP_DEVELOPMENT_OF_PRIMARY_FEMALE_SEXUAL_CHARACTERISTICS | 91 | -0.270115 | -0.831531 | 0.786353 | 1 |
| GOBP_NEGATIVE_REGULATION_OF_INTERLEUKIN_8_PRODUCTION | 20 | -0.342079 | -0.831319 | 0.714094 | 1 |
| GOBP_ACTIN_FILAMENT_DEPOLYMERIZATION | 58 | -0.283169 | -0.830984 | 0.752998 | 1 |
| GOBP_REGULATION_OF_CALCIUM_ION_IMPORT | 34 | -0.310706 | -0.83083 | 0.73731 | 1 |
| GOBP_CELLULAR_AMINO_ACID_METABOLIC_PROCESS | 323 | -0.24624 | -0.830577 | 0.87753 | 1 |
| GOBP_T_CELL_HOMEOSTASIS | 36 | -0.307342 | -0.830268 | 0.732234 | 1 |
| GOBP_SYNAPSE_ASSEMBLY | 164 | -0.252968 | -0.829027 | 0.846731 | 1 |
| GOBP_OLIGODENDROCYTE_DIFFERENTIATION | 84 | -0.269728 | -0.828911 | 0.788863 | 1 |
| GOBP_REGULATION_OF_CYTOPLASMIC_TRANSPORT | 27 | -0.320386 | -0.82864 | 0.707851 | 1 |
| GOBP_REGULATION_OF_PROTEIN_DEPHOSPHORYLATION | 138 | -0.258544 | -0.828607 | 0.809576 | 1 |
| GOBP_FERTILIZATION | 162 | -0.256515 | -0.82835 | 0.82234 | 1 |
| GOBP_REGULATION_OF_PEPTIDYL_THREONINE_PHOSPHORYLATION | 44 | -0.295276 | -0.827989 | 0.731343 | 1 |
| GOBP_GLOMERULAR_EPITHELIAL_CELL_DIFFERENTIATION | 20 | -0.33936 | -0.827455 | 0.715803 | 1 |
| GOBP_NEGATIVE_REGULATION_OF_ANTIGEN_RECEPTOR_MEDIATED_SIGNAL | 29 | -0.318254 | -0.82717 | 0.721945 | 1 |
| GOBP_DETECTION_OF_EXTERNAL_BIOTIC_STIMULUS | 23 | -0.328348 | -0.826114 | 0.713126 | 1 |
| GOBP_GLIOGENESIS | 265 | -0.246843 | -0.825979 | 0.864286 | 1 |
| GOBP_ALDITOL_METABOLIC_PROCESS | 21 | -0.337388 | -0.825853 | 0.711665 | 1 |
| GOBP_APOPTOTIC_PROCESS_INVOLVED_IN_MORPHOGENESIS | 22 | -0.336769 | -0.825567 | 0.696065 | 1 |
| GOBP_POSITIVE_REGULATION_OF_PEPTIDYL_TYROSINE_PHOSPHORYLATION | 184 | -0.252716 | -0.825527 | 0.843284 | 1 |
| GOBP_GLYCOPROTEIN_METABOLIC_PROCESS | 395 | -0.242776 | -0.825154 | 0.905242 | 1 |
| GOBP_POSITIVE_REGULATION_OF_CELL_CYCLE | 365 | -0.244377 | -0.825128 | 0.899295 | 1 |
| GOBP_POLYOL_CATABOLIC_PROCESS | 25 | -0.324612 | -0.824925 | 0.730245 | 1 |
| GOBP_NEUROTRANSMITTER_TRANSPORT | 198 | -0.252573 | -0.824888 | 0.847193 | 1 |
| GOBP_IRON_ION_TRANSMEMBRANE_TRANSPORT | 19 | -0.344032 | -0.824819 | 0.710067 | 1 |
| GOBP_REGULATION_OF_DENDRITIC_SPINE_MORPHOGENESIS | 45 | -0.295362 | -0.824386 | 0.739445 | 1 |
| GOBP_FOLIC_ACID_METABOLIC_PROCESS | 20 | -0.337977 | -0.824354 | 0.722598 | 1 |
| GOBP_RESPONSE_TO_KETONE | 186 | -0.247982 | -0.823839 | 0.829832 | 1 |
| GOBP_ANIMAL_ORGAN_FORMATION | 60 | -0.282371 | -0.823773 | 0.758007 | 1 |
| GOBP_REGULATION_OF_KERATINOCYTE_PROLIFERATION | 31 | -0.318437 | -0.82358 | 0.72875 | 1 |
| GOBP_BILE_ACID_METABOLIC_PROCESS | 45 | -0.295802 | -0.823368 | 0.729864 | 1 |
| GOBP_CELL_PROLIFERATION_IN_HINDBRAIN | 16 | -0.352061 | -0.822959 | 0.72 | 1 |
| GOBP_EYE_PHOTORECEPTOR_CELL_DEVELOPMENT | 31 | -0.309577 | -0.822878 | 0.726452 | 1 |
| GOBP_MUSCLE_TISSUE_DEVELOPMENT | 342 | -0.243155 | -0.822595 | 0.878147 | 1 |
| GOBP_REGULATION_OF_CALCIUM_ION_TRANSMEMBRANE_TRANSPORT | 136 | -0.257728 | -0.821836 | 0.823974 | 1 |
| GOBP_SPHINGOLIPID_METABOLIC_PROCESS | 163 | -0.252405 | -0.821642 | 0.824524 | 1 |
| GOBP_APOPTOTIC_PROCESS_INVOLVED_IN_DEVELOPMENT | 34 | -0.311709 | -0.821564 | 0.725849 | 1 |
| GOBP_POSITIVE_REGULATION_OF_CELL_CYCLE_PROCESS | 275 | -0.245848 | -0.82151 | 0.890593 | 1 |
| GOBP_OSTEOBLAST_PROLIFERATION | 29 | -0.31521 | -0.821414 | 0.739018 | 1 |
| GOBP_GLYCOSPHINGOLIPID_METABOLIC_PROCESS | 65 | -0.277621 | -0.821094 | 0.762615 | 1 |
| GOBP_REGULATION_OF_ISOTYPE_SWITCHING | 35 | -0.307109 | -0.821002 | 0.74005 | 1 |
| GOBP_REGULATION_OF_PHOSPHATASE_ACTIVITY | 170 | -0.250179 | -0.820381 | 0.849156 | 1 |
| GOBP_POSITIVE_REGULATION_OF_AMYLOID_PRECURSOR_PROTEIN_CATABO | 22 | -0.332347 | -0.820012 | 0.731904 | 1 |
| GOBP_MODULATION_BY_VIRUS_OF_HOST_CELLULAR_PROCESS | 18 | -0.344134 | -0.819894 | 0.72541 | 1 |
| GOBP_PROTEIN_PROCESSING | 210 | -0.248752 | -0.819553 | 0.872518 | 1 |
| GOBP_BILE_ACID_BIOSYNTHETIC_PROCESS | 36 | -0.307271 | -0.819394 | 0.744544 | 1 |
| GOBP_POSITIVE_REGULATION_OF_LYMPHOCYTE_MIGRATION | 36 | -0.299745 | -0.819211 | 0.747423 | 1 |
| GOBP_POSITIVE_REGULATION_OF_HISTONE_METHYLATION | 34 | -0.307393 | -0.818882 | 0.748744 | 1 |
| GOBP_EPITHELIAL_CELL_PROLIFERATION | 378 | -0.24043 | -0.818737 | 0.90433 | 1 |
| GOBP_GLYCEROPHOSPHOLIPID_METABOLIC_PROCESS | 336 | -0.24222 | -0.817916 | 0.899295 | 1 |
| GOBP_SPROUTING_ANGIOGENESIS | 130 | -0.25442 | -0.817655 | 0.823402 | 1 |
| GOBP_NUCLEOSIDE_CATABOLIC_PROCESS | 32 | -0.310926 | -0.817469 | 0.745806 | 1 |
| GOBP_DEMETHYLATION | 67 | -0.276681 | -0.817361 | 0.778409 | 1 |
| GOBP_FATTY_ACID_TRANSPORT | 141 | -0.25283 | -0.817311 | 0.830636 | 1 |
| GOBP_RNA_SURVEILLANCE | 15 | -0.361098 | -0.817302 | 0.727891 | 1 |
| GOBP_WATER_SOLUBLE_VITAMIN_METABOLIC_PROCESS | 81 | -0.271853 | -0.817083 | 0.79883 | 1 |
| GOBP_RESPONSE_TO_DRUG | 340 | -0.241833 | -0.81657 | 0.90202 | 1 |
| GOBP_GLYCOSYL_COMPOUND_METABOLIC_PROCESS | 126 | -0.256297 | -0.8165 | 0.840812 | 1 |
| GOBP_REGULATION_OF_CELLULAR_CARBOHYDRATE_METABOLIC_PROCESS | 136 | -0.252519 | -0.816047 | 0.815016 | 1 |
| GOBP_REGULATION_OF_SYSTEMIC_ARTERIAL_BLOOD_PRESSURE_BY_HORM | 33 | -0.307198 | -0.815805 | 0.745199 | 1 |
| GOBP_MULTI_MULTICELLULAR_ORGANISM_PROCESS | 185 | -0.248313 | -0.815728 | 0.840504 | 1 |
| GOBP_CELL_JUNCTION_MAINTENANCE | 32 | -0.306737 | -0.815686 | 0.75 | 1 |
| GOBP_FIBROBLAST_APOPTOTIC_PROCESS | 20 | -0.338381 | -0.815592 | 0.731844 | 1 |
| GOBP_CELLULAR_RESPONSE_TO_DEXAMETHASONE_STIMULUS | 28 | -0.310119 | -0.815237 | 0.743968 | 1 |
| GOBP_POSITIVE_REGULATION_OF_ANION_TRANSMEMBRANE_TRANSPORT | 55 | -0.281432 | -0.815065 | 0.779805 | 1 |
| GOBP_POSITIVE_REGULATION_OF_CYTOKINESIS | 32 | -0.31074 | -0.814242 | 0.748744 | 1 |
| GOBP_REGULATION_OF_GASTRULATION | 16 | -0.354363 | -0.814212 | 0.726483 | 1 |
| GOBP_GLUCOCORTICOID_METABOLIC_PROCESS | 23 | -0.325891 | -0.81404 | 0.733696 | 1 |
| GOBP_OSTEOBLAST_DEVELOPMENT | 17 | -0.350406 | -0.813593 | 0.743017 | 1 |
| GOBP_POSITIVE_REGULATION_OF_CATION_CHANNEL_ACTIVITY | 68 | -0.272818 | -0.813588 | 0.786618 | 1 |
| GOBP_MITOCHONDRIAL_OUTER_MEMBRANE_PERMEABILIZATION | 55 | -0.281639 | -0.813325 | 0.775904 | 1 |
| GOBP_SYNAPTIC_TRANSMISSION_CHOLINERGIC | 27 | -0.317801 | -0.813049 | 0.740741 | 1 |
| GOBP_FATTY_ACID_TRANSMEMBRANE_TRANSPORT | 49 | -0.281452 | -0.812864 | 0.783582 | 1 |
| GOBP_T_CELL_APOPTOTIC_PROCESS | 47 | -0.290125 | -0.812608 | 0.767689 | 1 |
| GOBP_REGULATION_OF_NERVOUS_SYSTEM_DEVELOPMENT | 400 | -0.238446 | -0.812362 | 0.933735 | 1 |
| GOBP_INTESTINAL_ABSORPTION | 34 | -0.304189 | -0.811429 | 0.748072 | 1 |
| GOBP_RECEPTOR_LOCALIZATION_TO_SYNAPSE | 48 | -0.284318 | -0.810867 | 0.768844 | 1 |
| GOBP_REGULATION_OF_MAST_CELL_ACTIVATION_INVOLVED_IN_IMMUNE_RE | 30 | -0.308097 | -0.810738 | 0.752604 | 1 |
| GOBP_NEGATIVE_REGULATION_OF_FATTY_ACID_METABOLIC_PROCESS | 25 | -0.320791 | -0.81047 | 0.756972 | 1 |
| GOBP_REGULATION_OF_ANATOMICAL_STRUCTURE_SIZE | 478 | -0.238285 | -0.810408 | 0.939819 | 1 |
| GOBP_NUCLEOTIDE_TRANSMEMBRANE_TRANSPORT | 15 | -0.356804 | -0.810197 | 0.721519 | 1 |
| GOBP_RETROGRADE_AXONAL_TRANSPORT | 21 | -0.337889 | -0.810063 | 0.727027 | 1 |
| GOBP_REGULATION_OF_ANION_TRANSMEMBRANE_TRANSPORT | 101 | -0.261316 | -0.809967 | 0.813824 | 1 |
| GOBP_REGULATION_OF_PROTEIN_POLYUBIQUITINATION | 24 | -0.31748 | -0.809523 | 0.753662 | 1 |
| GOBP_REGULATION_OF_GTPASE_ACTIVITY | 467 | -0.236433 | -0.808969 | 0.929789 | 1 |
| GOBP_MAINTENANCE_OF_SYNAPSE_STRUCTURE | 16 | -0.346163 | -0.808959 | 0.731742 | 1 |
| GOBP_NEGATIVE_REGULATION_OF_CALCIUM_ION_TRANSPORT_INTO_CYTOS | 16 | -0.346712 | -0.808851 | 0.741573 | 1 |
| GOBP_REGULATION_OF_ATP_BIOSYNTHETIC_PROCESS | 20 | -0.331446 | -0.808108 | 0.728034 | 1 |
| GOBP_RNA_PHOSPHODIESTER_BOND_HYDROLYSIS | 149 | -0.249972 | -0.807778 | 0.844609 | 1 |
| GOBP_MACROPHAGE_ACTIVATION_INVOLVED_IN_IMMUNE_RESPONSE | 17 | -0.339238 | -0.807322 | 0.727151 | 1 |
| GOBP_OLEFINIC_COMPOUND_BIOSYNTHETIC_PROCESS | 20 | -0.332674 | -0.807151 | 0.721333 | 1 |
| GOBP_REGULATION_OF_CHOLESTEROL_METABOLIC_PROCESS | 55 | -0.280497 | -0.807062 | 0.783432 | 1 |
| GOBP_REGULATION_OF_MEMBRANE_POTENTIAL | 402 | -0.236658 | -0.806443 | 0.925628 | 1 |
| GOBP_SNARE_COMPLEX_ASSEMBLY | 18 | -0.341636 | -0.806399 | 0.734752 | 1 |
| GOBP_SEX_DETERMINATION | 18 | -0.335838 | -0.806057 | 0.734973 | 1 |
| GOBP_REGULATION_OF_TRANSLATIONAL_INITIATION | 75 | -0.26781 | -0.805816 | 0.796948 | 1 |
| GOBP_NEGATIVE_REGULATION_OF_CELL_AGING | 25 | -0.320201 | -0.805724 | 0.720899 | 1 |
| GOBP_NEGATIVE_REGULATION_OF_TOLL_LIKE_RECEPTOR_SIGNALING_PATH | 39 | -0.29264 | -0.805046 | 0.76 | 1 |
| GOBP_REGULATION_OF_POSTSYNAPTIC_NEUROTRANSMITTER_RECEPTOR_A | 15 | -0.352675 | -0.805045 | 0.741667 | 1 |
| GOBP_CYTOKINESIS | 162 | -0.245742 | -0.805028 | 0.864979 | 1 |
| GOBP_PURINE_CONTAINING_COMPOUND_METABOLIC_PROCESS | 441 | -0.236459 | -0.804887 | 0.940763 | 1 |
| GOBP_POSITIVE_REGULATION_OF_LYMPHOCYTE_DIFFERENTIATION | 98 | -0.260238 | -0.804624 | 0.817486 | 1 |
| GOBP_RIBOSOMAL_SMALL_SUBUNIT_BIOGENESIS | 75 | -0.266691 | -0.804582 | 0.813084 | 1 |
| GOBP_ATRIAL_CARDIAC_MUSCLE_CELL_TO_AV_NODE_CELL_SIGNALING | 18 | -0.338848 | -0.804253 | 0.744595 | 1 |
| GOBP_INTRA_S_DNA_DAMAGE_CHECKPOINT | 17 | -0.342377 | -0.804095 | 0.753681 | 1 |
| GOBP_SINGLE_STRANDED_VIRAL_RNA_REPLICATION_VIA_DOUBLE_STRANDI | 20 | -0.328649 | -0.803793 | 0.702667 | 1 |
| GOBP_MAINTENANCE_OF_LOCATION | 307 | -0.238736 | -0.803689 | 0.918699 | 1 |
| GOBP_PHOSPHATE_ION_TRANSPORT | 24 | -0.321516 | -0.803676 | 0.75 | 1 |
| GOBP_SYNAPTIC_VESICLE_CYTOSKELETAL_TRANSPORT | 19 | -0.341066 | -0.803587 | 0.742507 | 1 |
| GOBP_NEGATIVE_REGULATION_OF_RESPONSE_TO_OXIDATIVE_STRESS | 20 | -0.332716 | -0.803568 | 0.735135 | 1 |
| GOBP_REGULATION_OF_CELL_CYCLE_G2_M_PHASE_TRANSITION | 214 | -0.243618 | -0.803447 | 0.882292 | 1 |
| GOBP_REACTIVE_OXYGEN_SPECIES_BIOSYNTHETIC_PROCESS | 110 | -0.256138 | -0.803428 | 0.850279 | 1 |
| GOBP_SISTER_CHROMATID_COHESION | 60 | -0.27411 | -0.803383 | 0.793632 | 1 |

| | | | | | |
|---|---|---|---|---|---|
| GOBP_PHOSPHATIDYLINOSITOL_DEPHOSPHORYLATION | 30 | 0.189796 | 0.593919 | 0.978541 | 1 |
| GOBP_TAIL_ANCHORED_MEMBRANE_PROTEIN_INSERTION_INTO_ER_MI | 16 | 0.223773 | 0.592481 | 0.927757 | 1 |
| GOBP_NEGATIVE_REGULATION_OF_PROTEIN_LOCALIZATION_TO_NUCLE | 29 | 0.186408 | 0.587948 | 0.96729 | 1 |
| GOBP_RESPONSE_TO_DSRNA | 38 | 0.176482 | 0.586412 | 0.984772 | 1 |
| GOBP_VIRION_ASSEMBLY | 40 | 0.169422 | 0.585555 | 0.99505 | 1 |
| GOBP_ESTABLISHMENT_OF_MITOCHONDRION_LOCALIZATION | 25 | 0.187955 | 0.585519 | 0.983122 | 1 |
| GOBP_PROTEIN_MONOUBIQUITINATION | 66 | 0.15327 | 0.584039 | 1 | 1 |
| GOBP_ERAD_PATHWAY | 101 | 0.147675 | 0.583972 | 1 | 1 |
| GOBP_POSITIVE_REGULATION_OF_RESPONSE_TO_ENDOPLASMIC_RETI | 34 | 0.17493 | 0.583477 | 0.988827 | 1 |
| GOBP_CELLULAR_RESPONSE_TO_STEROL_DEPLETION | 16 | 0.217721 | 0.581941 | 0.966443 | 1 |
| GOBP_NEURON_DEATH_IN_RESPONSE_TO_OXIDATIVE_STRESS | 26 | 0.193736 | 0.581467 | 0.987805 | 1 |
| GOBP_REGULATION_OF_PRODUCTION_OF_SMALL_RNA_INVOLVED_IN_( | 23 | 0.195246 | 0.581133 | 0.969957 | 1 |
| GOBP_NEGATIVE_REGULATION_OF_JNK_CASCADE | 35 | 0.175217 | 0.580723 | 0.972477 | 1 |
| GOBP_NEGATIVE_REGULATION_OF_MACROAUTOPHAGY | 35 | 0.177998 | 0.580533 | 0.979695 | 1 |
| GOBP_RESPONSE_TO_HYPEROXIA | 22 | 0.19597 | 0.579428 | 0.962406 | 1 |
| GOBP_NEGATIVE_REGULATION_OF_CHEMOKINE_PRODUCTION | 21 | 0.197774 | 0.578537 | 0.975177 | 1 |
| GOBP_REGULATION_OF_AUTOPHAGOSOME_ASSEMBLY | 39 | 0.177563 | 0.578344 | 0.995098 | 1 |
| GOBP_DENDRITIC_SPINE_MAINTENANCE | 18 | 0.205624 | 0.575953 | 0.967273 | 1 |
| GOBP_POSITIVE_REGULATION_OF_MITOCHONDRIAL_TRANSLATION | 16 | 0.217639 | 0.573583 | 0.967626 | 1 |
| GOBP_COENZYME_A_METABOLIC_PROCESS | 15 | 0.212228 | 0.573264 | 0.96831 | 1 |
| GOBP_REGULATION_OF_PROTEASOMAL_PROTEIN_CATABOLIC_PROCES | 179 | 0.135483 | 0.571599 | 1 | 1 |
| GOBP_REGULATION_OF_DNA_TEMPLATED_TRANSCRIPTION_IN_RESPON | 114 | 0.137588 | 0.570346 | 1 | 1 |
| GOBP_AMINO_ACID_ACTIVATION | 49 | 0.162988 | 0.570299 | 0.995098 | 1 |
| GOBP_POSITIVE_REGULATION_OF_POSTTRANSCRIPTIONAL_GENE_SILE | 29 | 0.183356 | 0.569046 | 0.990431 | 1 |
| GOBP_POSITIVE_REGULATION_OF_SUBSTRATE_ADHESION_DEPENDENT | 41 | 0.169248 | 0.567459 | 0.995074 | 1 |
| GOBP_REGULATION_OF_PROTEIN_STABILITY | 286 | 0.121213 | 0.567167 | 1 | 1 |
| GOBP_VESICLE_TARGETING | 95 | 0.145232 | 0.566956 | 1 | 1 |
| GOBP_ARF_PROTEIN_SIGNAL_TRANSDUCTION | 19 | 0.198305 | 0.564399 | 0.971429 | 1 |
| GOBP_REGULATION_OF_VACUOLE_ORGANIZATION | 47 | 0.16403 | 0.564345 | 1 | 1 |
| GOBP_REGULATION_OF_DNA_REPAIR | 129 | 0.140041 | 0.564074 | 1 | 1 |
| GOBP_CORONARY_VASCULATURE_MORPHOGENESIS | 15 | 0.20699 | 0.562667 | 0.959854 | 1 |
| GOBP_REGULATION_OF_TYPE_I_INTERFERON_MEDIATED_SIGNALING_P | 33 | 0.175653 | 0.562622 | 0.985782 | 1 |
| GOBP_RESPONSE_TO_INCREASED_OXYGEN_LEVELS | 29 | 0.179183 | 0.562255 | 1 | 1 |
| GOBP_PLATELET_DERIVED_GROWTH_FACTOR_RECEPTOR_SIGNALING_ | 55 | 0.159437 | 0.562095 | 1 | 1 |
| GOBP_CHROMATIN_DISASSEMBLY | 18 | 0.197017 | 0.560079 | 0.977012 | 1 |
| GOBP_EXPLORATION_BEHAVIOR | 17 | 0.200768 | 0.556429 | 0.975 | 1 |
| GOBP_NUCLEAR_TRANSCRIBED_MRNA_CATABOLIC_PROCESS_EXONU( | 35 | 0.173861 | 0.555722 | 0.985714 | 1 |
| GOBP_IMMUNOLOGICAL_MEMORY_PROCESS | 17 | 0.197614 | 0.555686 | 0.968421 | 1 |
| GOBP_FATTY_ACID_BETA_OXIDATION_USING_ACYL_COA_OXIDASE | 15 | 0.204208 | 0.554273 | 0.956364 | 1 |
| GOBP_FATTY_ACID_BETA_OXIDATION | 74 | 0.145451 | 0.553966 | 1 | 1 |
| GOBP_REGULATION_OF_MONOCYTE_DIFFERENTIATION | 19 | 0.197117 | 0.553662 | 0.97037 | 1 |
| GOBP_CYTOSOLIC_TRANSPORT | 166 | 0.126188 | 0.551239 | 1 | 1 |
| GOBP_RNA_SPLICING | 450 | 0.121466 | 0.551 | 1 | 1 |
| GOBP_NEGATIVE_REGULATION_OF_INTRACELLULAR_PROTEIN_TRANSP | 40 | 0.160849 | 0.549175 | 1 | 1 |
| GOBP_REGULATION_OF_PROTON_TRANSPORT | 22 | 0.184561 | 0.547474 | 0.972656 | 1 |
| GOBP_REGULATION_OF_HISTONE_DEACETYLATION | 26 | 0.181408 | 0.547352 | 0.991701 | 1 |
| GOBP_NEGATIVE_REGULATION_OF_PROTEOLYSIS_INVOLVED_IN_CELLU | 62 | 0.150054 | 0.547001 | 1 | 1 |
| GOBP_CELL_DEATH_IN_RESPONSE_TO_HYDROGEN_PEROXIDE | 24 | 0.180383 | 0.546667 | 0.977012 | 1 |
| GOBP_CELLULAR_RESPONSE_TO_LIPOPROTEIN_PARTICLE_STIMULUS | 28 | 0.174286 | 0.54629 | 0.991342 | 1 |
| GOBP_ATP_SYNTHESIS_COUPLED_ELECTRON_TRANSPORT | 99 | 0.134815 | 0.546038 | 1 | 1 |
| GOBP_NCRNA_EXPORT_FROM_NUCLEUS | 38 | 0.164421 | 0.545531 | 0.995536 | 1 |
| GOBP_AUTOPHAGOSOME_MATURATION | 41 | 0.160138 | 0.545424 | 0.995496 | 1 |
| GOBP_VESICLE_BUDDING_FROM_MEMBRANE | 109 | 0.137849 | 0.545241 | 1 | 1 |
| GOBP_VIRAL_BUDDING | 26 | 0.186599 | 0.544883 | 0.97479 | 1 |
| GOBP_PREASSEMBLY_OF_GPI_ANCHOR_IN_ER_MEMBRANE | 16 | 0.201269 | 0.544288 | 0.978799 | 1 |
| GOBP_GOLGI_TO_PLASMA_MEMBRANE_PROTEIN_TRANSPORT | 38 | 0.165496 | 0.544142 | 1 | 1 |
| GOBP_MONOSACCHARIDE_CATABOLIC_PROCESS | 58 | 0.149262 | 0.543598 | 1 | 1 |
| GOBP_VASCULAR_ASSOCIATED_SMOOTH_MUSCLE_CELL_DIFFERENTIA | 27 | 0.172583 | 0.543228 | 0.991189 | 1 |
| GOBP_REGULATION_OF_MITOCHONDRIAL_FISSION | 27 | 0.180205 | 0.541505 | 0.986957 | 1 |
| GOBP_POSITIVE_REGULATION_OF_VIRAL_PROCESS | 91 | 0.134124 | 0.539368 | 1 | 1 |
| GOBP_DEOXYRIBONUCLEOSIDE_TRIPHOSPHATE_METABOLIC_PROCESS | 15 | 0.198969 | 0.53876 | 0.979522 | 1 |
| GOBP_NEGATIVE_REGULATION_OF_PHAGOCYTOSIS | 21 | 0.183516 | 0.535849 | 0.995918 | 1 |
| GOBP_TETRAPYRROLE_METABOLIC_PROCESS | 56 | 0.148593 | 0.535626 | 1 | 1 |
| GOBP_DNA_SYNTHESIS_INVOLVED_IN_DNA_REPAIR | 52 | 0.150908 | 0.53537 | 1 | 1 |
| GOBP_REGULATION_OF_DOUBLE_STRAND_BREAK_REPAIR_VIA_NONHO | 28 | 0.167742 | 0.534092 | 0.992157 | 1 |
| GOBP_AUTOPHAGOSOME_ORGANIZATION | 97 | 0.13535 | 0.532577 | 1 | 1 |
| GOBP_COPII_COATED_VESICLE_BUDDING | 73 | 0.142963 | 0.531176 | 0.992958 | 1 |
| GOBP_MITOTIC_DNA_INTEGRITY_CHECKPOINT | 104 | 0.136147 | 0.530646 | 1 | 1 |
| GOBP_OLIGOSACCHARIDE_LIPID_INTERMEDIATE_BIOSYNTHETIC_PROC | 21 | 0.184241 | 0.529953 | 0.984791 | 1 |
| GOBP_FUCOSYLATION | 16 | 0.190596 | 0.528818 | 0.989796 | 1 |
| GOBP_MATURATION_OF_LSU_RRNA | 28 | 0.175613 | 0.527631 | 0.996094 | 1 |
| GOBP_IRON_SULFUR_CLUSTER_ASSEMBLY | 23 | 0.178612 | 0.527057 | 0.991342 | 1 |
| GOBP_PHOSPHORYLATED_CARBOHYDRATE_DEPHOSPHORYLATION | 16 | 0.197379 | 0.526863 | 0.979094 | 1 |
| GOBP_TRANSCRIPTION_ELONGATION_FROM_RNA_POLYMERASE_II_PR( | 84 | 0.135303 | 0.525117 | 1 | 1 |
| GOBP_REGULATION_OF_PROTEIN_DEACETYLATION | 41 | 0.150201 | 0.52176 | 1 | 1 |
| GOBP_REGULATION_OF_PROTEIN_MODIFICATION_BY_SMALL_PROTEIN_ | 229 | 0.120004 | 0.520677 | 1 | 1 |
| GOBP_AMP_METABOLIC_PROCESS | 16 | 0.193281 | 0.520384 | 0.992565 | 1 |
| GOBP_RNA_POLYADENYLATION | 45 | 0.149236 | 0.519567 | 1 | 1 |
| GOBP_HISTONE_H2A_MONOUBIQUITINATION | 17 | 0.18992 | 0.519449 | 0.984436 | 1 |
| GOBP_AEROBIC_RESPIRATION | 85 | 0.134737 | 0.518885 | 0.992701 | 1 |
| GOBP_MISMATCH_REPAIR | 36 | 0.157094 | 0.515552 | 1 | 1 |
| GOBP_POSITIVE_REGULATION_OF_VIRAL_LIFE_CYCLE | 32 | 0.163443 | 0.515288 | 0.995392 | 1 |
| GOBP_SIGNAL_TRANSDUCTION_INVOLVED_IN_CELL_CYCLE_CHECKPOI | 73 | 0.134572 | 0.514863 | 1 | 1 |
| GOBP_PROTEIN_TARGETING_TO_MITOCHONDRION | 100 | 0.128783 | 0.512812 | 1 | 1 |
| GOBP_VACUOLE_ORGANIZATION | 173 | 0.121368 | 0.512534 | 1 | 1 |
| GOBP_MITOCHONDRIAL_ELECTRON_TRANSPORT_NADH_TO_UBIQUINON | 54 | 0.140696 | 0.511328 | 1 | 1 |
| GOBP_RESPONSE_TO_ARSENIC_CONTAINING_SUBSTANCE | 30 | 0.152608 | 0.508427 | 0.995671 | 1 |
| GOBP_REGULATION_OF_PROTEIN_LOCALIZATION_TO_NUCLEUS | 128 | 0.123264 | 0.50513 | 1 | 1 |
| GOBP_VIRAL_BUDDING_VIA_HOST_ESCRT_COMPLEX | 21 | 0.174818 | 0.504968 | 0.987805 | 1 |
| GOBP_LENS_MORPHOGENESIS_IN_CAMERA_TYPE_EYE | 17 | 0.183528 | 0.50492 | 0.992593 | 1 |
| GOBP_REGULATION_OF_MRNA_POLYADENYLATION | 17 | 0.179682 | 0.501364 | 0.989437 | 1 |
| GOBP_ENDOSOMAL_TRANSPORT | 242 | 0.109434 | 0.50129 | 1 | 1 |
| GOBP_ER_NUCLEUS_SIGNALING_PATHWAY | 52 | 0.141153 | 0.501232 | 1 | 1 |
| GOBP_MANNOSYLATION | 36 | 0.148188 | 0.497826 | 1 | 1 |
| GOBP_RNA_SPLICING_VIA_TRANSESTERIFICATION_REACTIONS | 359 | 0.107529 | 0.494966 | 1 | 1 |
| GOBP_VESICLE_TARGETING_TO_FROM_OR_WITHIN_GOLGI | 74 | 0.12742 | 0.492946 | 1 | 1 |
| GOBP_IMMUNE_RESPONSE_TO_TUMOR_CELL | 20 | 0.162719 | 0.492644 | 0.989051 | 1 |
| GOBP_MICROTUBULE_BASED_TRANSPORT | 186 | 0.112285 | 0.490109 | 1 | 1 |
| GOBP_PROTEIN_K48_LINKED_UBIQUITINATION | 61 | 0.132923 | 0.488941 | 1 | 1 |
| GOBP_RESPONSE_TO_STEROL_DEPLETION | 18 | 0.173945 | 0.488213 | 0.982014 | 1 |
| GOBP_NEGATIVE_REGULATION_OF_CELLULAR_PROTEIN_LOCALIZATION | 109 | 0.121432 | 0.486413 | 1 | 1 |
| GOBP_RETROGRADE_TRANSPORT_ENDOSOME_TO_GOLGI | 91 | 0.127362 | 0.4856 | 1 | 1 |
| GOBP_REGULATION_OF_TORC1_SIGNALING | 39 | 0.145666 | 0.484247 | 1 | 1 |
| GOBP_RETROGRADE_VESICLE_MEDIATED_TRANSPORT_GOLGI_TO_END | 86 | 0.12161 | 0.483237 | 1 | 1 |
| GOBP_TORC1_SIGNALING | 47 | 0.139813 | 0.480756 | 1 | 1 |
| GOBP_NEGATIVE_REGULATION_OF_TORC1_SIGNALING | 17 | 0.17578 | 0.480314 | 1 | 1 |
| GOBP_CELLULAR_RESPONSE_TO_ESTROGEN_STIMULUS | 15 | 0.178281 | 0.47568 | 0.996564 | 1 |
| GOBP_TOLL_LIKE_RECEPTOR_3_SIGNALING_PATHWAY | 19 | 0.166044 | 0.475118 | 0.996241 | 1 |
| GOBP_RECEPTOR_CATABOLIC_PROCESS | 35 | 0.143298 | 0.472732 | 1 | 1 |
| GOBP_REGULATION_OF_ENDOPLASMIC_RETICULUM_UNFOLDED_PROT | 28 | 0.153175 | 0.470186 | 1 | 1 |
| GOBP_VACUOLAR_TRANSPORT | 147 | 0.114179 | 0.470064 | 1 | 1 |
| GOBP_REGULATION_OF_UBIQUITIN_DEPENDENT_PROTEIN_CATABOLIC_ | 159 | 0.114368 | 0.469604 | 1 | 1 |
| GOBP_MACROAUTOPHAGY | 308 | 0.102865 | 0.468304 | 1 | 1 |
| GOBP_RESPONSE_TO_ENDOPLASMIC_RETICULUM_STRESS | 289 | 0.100917 | 0.466406 | 1 | 1 |
| GOBP_REGULATION_OF_CYTOPLASMIC_TRANSLATION | 19 | 0.159419 | 0.460099 | 1 | 1 |
| GOBP_LYSOSOMAL_TRANSPORT | 105 | 0.115426 | 0.454223 | 1 | 1 |
| GOBP_AUTOPHAGY_OF_MITOCHONDRION | 74 | 0.11818 | 0.454141 | 1 | 1 |
| GOBP_CYTOCHROME_COMPLEX_ASSEMBLY | 35 | 0.139841 | 0.451362 | 1 | 1 |
| GOBP_ENDOSOME_TO_LYSOSOME_TRANSPORT | 51 | 0.130248 | 0.450627 | 1 | 1 |
| GOBP_NADH_DEHYDROGENASE_COMPLEX_ASSEMBLY | 65 | 0.121832 | 0.450484 | 1 | 1 |
| GOBP_PROTEIN_MANNOSYLATION | 23 | 0.148109 | 0.432064 | 0.995935 | 1 |
| GOBP_GOLGI_VESICLE_TRANSPORT | 365 | 0.09405 | 0.427862 | 1 | 1 |
| GOBP_POSITIVE_REGULATION_OF_PROTEASOMAL_UBIQUITIN_DEPENDE | 87 | 0.110732 | 0.426365 | 1 | 1 |
| GOBP_NUCLEOTIDE_EXCISION_REPAIR | 106 | 0.106495 | 0.426361 | 1 | 1 |
| GOBP_ORGANELLE_DISASSEMBLY | 106 | 0.105636 | 0.421391 | 1 | 1 |
| GOBP_CELLULAR_RESPIRATION | 182 | 0.096467 | 0.417085 | 1 | 1 |
| GOBP_HISTONE_H2A_UBIQUITINATION | 26 | 0.136985 | 0.41564 | 1 | 1 |
| GOBP_IRE1_MEDIATED_UNFOLDED_PROTEIN_RESPONSE | 65 | 0.111876 | 0.41309 | 1 | 1 |
| GOBP_REGULATION_OF_EXOSOMAL_SECRETION | 16 | 0.148322 | 0.404968 | 1 | 1 |
| GOBP_ENDOPLASMIC_RETICULUM_UNFOLDED_PROTEIN_RESPONSE | 123 | 0.097719 | 0.403318 | 1 | 1 |
| GOBP_PRODUCTION_OF_SMALL_RNA_INVOLVED_IN_GENE_SILENCING_ | 53 | 0.110957 | 0.393582 | 1 | 1 |
| GOBP_MEMBRANE_PROTEIN_INTRACELLULAR_DOMAIN_PROTEOLYSIS | 20 | 0.136962 | 0.389109 | 0.996094 | 1 |
| GOBP_POSITIVE_REGULATION_OF_FOCAL_ADHESION_ASSEMBLY | 24 | 0.124352 | 0.384172 | 1 | 1 |
| GOBP_EXTRACELLULAR_VESICLE_BIOGENESIS | 22 | 0.127497 | 0.366326 | 1 | 1 |
| GOBP_POSITIVE_REGULATION_OF_CELL_SUBSTRATE_JUNCTION_ORGA | 29 | 0.111897 | 0.365657 | 1 | 1 |
| GOBP_MITOCHONDRIAL_RESPIRATORY_CHAIN_COMPLEX_ASSEMBLY | 100 | 0.089181 | 0.360601 | 1 | 0.999949 |

| | | | | | |
|---|---|---|---|---|---|
| GOBP_INFLAMMATORY_RESPONSE_TO_ANTIGENIC_STIMULUS | 51 | -0.279277 | -0.80327 | 0.774115 | 1 |
| GOBP_EOSINOPHIL_MIGRATION | 21 | -0.328696 | -0.803168 | 0.753662 | 1 |
| GOBP_EXOCRINE_SYSTEM_DEVELOPMENT | 38 | -0.289348 | -0.802977 | 0.772251 | 1 |
| GOBP_L_AMINO_ACID_TRANSPORT | 63 | -0.270792 | -0.802954 | 0.80376 | 1 |
| GOBP_AMINO_ACID_IMPORT | 47 | -0.284628 | -0.802824 | 0.777509 | 1 |
| GOBP_REGULATION_OF_DNA_TEMPLATED_TRANSCRIPTION_INITIATION | 41 | -0.290937 | -0.80237 | 0.78125 | 1 |
| GOBP_REGULATION_OF_EPIDERMIS_DEVELOPMENT | 64 | -0.272186 | -0.802088 | 0.80355 | 1 |
| GOBP_NEGATIVE_REGULATION_OF_LYMPHOCYTE_DIFFERENTIATION | 51 | -0.278659 | -0.801936 | 0.792271 | 1 |
| GOBP_GMP_METABOLIC_PROCESS | 15 | -0.347255 | -0.801897 | 0.744799 | 1 |
| GOBP_ESTABLISHMENT_OR_MAINTENANCE_OF_BIPOLAR_CELL_POLARITY | 44 | -0.280614 | -0.801698 | 0.783951 | 1 |
| GOBP_NEGATIVE_REGULATION_OF_CELL_DEVELOPMENT | 162 | -0.248462 | -0.800531 | 0.874866 | 1 |
| GOBP_MRNA_CIS_SPLICING_VIA_SPLICEOSOME | 28 | -0.308602 | -0.800368 | 0.767742 | 1 |
| GOBP_RECEPTOR_RECYCLING | 32 | -0.300289 | -0.80031 | 0.766452 | 1 |
| GOBP_GLYCOPROTEIN_BIOSYNTHETIC_PROCESS | 325 | -0.236336 | -0.799555 | 0.926978 | 1 |
| GOBP_RESPONSE_TO_MUSCLE_STRETCH | 21 | -0.328258 | -0.79933 | 0.746988 | 1 |
| GOBP_DENDRITIC_CELL_DIFFERENTIATION | 46 | -0.285645 | -0.799222 | 0.792191 | 1 |
| GOBP_RESPONSE_TO_DEXAMETHASONE | 36 | -0.289159 | -0.799003 | 0.771176 | 1 |
| GOBP_NEGATIVE_REGULATION_OF_MYELOID_CELL_DIFFERENTIATION | 86 | -0.259702 | -0.798918 | 0.811725 | 1 |
| GOBP_AGING | 281 | -0.237639 | -0.798717 | 0.907881 | 1 |
| GOBP_TRNA_WOBBLE_BASE_MODIFICATION | 20 | -0.326274 | -0.798324 | 0.735414 | 1 |
| GOBP_REGULATION_OF_SISTER_CHROMATID_COHESION | 22 | -0.325126 | -0.798065 | 0.769923 | 1 |
| GOBP_REGULATION_OF_TELOMERASE_ACTIVITY | 47 | -0.283285 | -0.797561 | 0.800971 | 1 |
| GOBP_POSITIVE_REGULATION_OF_CYTOSKELETON_ORGANIZATION | 216 | -0.240985 | -0.797395 | 0.888083 | 1 |
| GOBP_ESTABLISHMENT_OR_MAINTENANCE_OF_MONOPOLAR_CELL_POLARIT | 23 | -0.324404 | -0.797091 | 0.755047 | 1 |
| GOBP_SODIUM_ION_TRANSMEMBRANE_TRANSPORT | 160 | -0.246992 | -0.796774 | 0.866667 | 1 |
| GOBP_SKELETAL_MUSCLE_CONTRACTION | 42 | -0.292836 | -0.796481 | 0.803238 | 1 |
| GOBP_NEURON_RECOGNITION | 46 | -0.282139 | -0.796394 | 0.779557 | 1 |
| GOBP_GLYCOLIPID_BIOSYNTHETIC_PROCESS | 64 | -0.271713 | -0.796353 | 0.80485 | 1 |
| GOBP_POSITIVE_REGULATION_OF_TUMOR_NECROSIS_FACTOR_SUPERFAMIL | 86 | -0.259021 | -0.796259 | 0.825167 | 1 |
| GOBP_REGULATION_OF_PLATELET_ACTIVATION | 33 | -0.293706 | -0.796207 | 0.790404 | 1 |
| GOBP_REGULATION_OF_LIPID_LOCALIZATION | 166 | -0.247114 | -0.796079 | 0.879274 | 1 |
| GOBP_ARACHIDONIC_ACID_SECRETION | 23 | -0.31783 | -0.795912 | 0.754054 | 1 |
| GOBP_POSITIVE_REGULATION_OF_TELOMERASE_ACTIVITY | 33 | -0.297199 | -0.795882 | 0.774275 | 1 |
| GOBP_MYD88_DEPENDENT_TOLL_LIKE_RECEPTOR_SIGNALING_PATHWAY | 34 | -0.295325 | -0.795836 | 0.765847 | 1 |
| GOBP_NEGATIVE_REGULATION_OF_T_CELL_RECEPTOR_SIGNALING_PATHWA | 22 | -0.323053 | -0.795807 | 0.746667 | 1 |
| GOBP_NEGATIVE_REGULATION_OF_NERVOUS_SYSTEM_DEVELOPMENT | 128 | -0.25078 | -0.795614 | 0.838919 | 1 |
| GOBP_REGULATION_OF_PROTEIN_POLYMERIZATION | 222 | -0.239775 | -0.795353 | 0.88785 | 1 |
| GOBP_REGULATION_OF_DNA_BINDING_TRANSCRIPTION_FACTOR_ACTIVITY | 415 | -0.235009 | -0.795082 | 0.941414 | 1 |
| GOBP_REGULATION_OF_CELLULAR_KETONE_METABOLIC_PROCESS | 165 | -0.245418 | -0.794482 | 0.880378 | 1 |
| GOBP_CRANIAL_NERVE_MORPHOGENESIS | 23 | -0.318798 | -0.794451 | 0.74867 | 1 |
| GOBP_POSITIVE_REGULATION_OF_MRNA_CATABOLIC_PROCESS | 50 | -0.284218 | -0.794305 | 0.792128 | 1 |
| GOBP_GLIAL_CELL_DIFFERENTIATION | 196 | -0.241632 | -0.794282 | 0.889468 | 1 |
| GOBP_NEGATIVE_REGULATION_OF_INNATE_IMMUNE_RESPONSE | 60 | -0.271678 | -0.794065 | 0.808737 | 1 |
| GOBP_NEGATIVE_REGULATION_OF_PRODUCTION_OF_MOLECULAR_MEDIATO | 37 | -0.293067 | -0.793393 | 0.78875 | 1 |
| GOBP_POSITIVE_REGULATION_OF_CILIUM_ASSEMBLY | 24 | -0.312175 | -0.793337 | 0.758667 | 1 |
| GOBP_POSITIVE_REGULATION_OF_CELL_ADHESION | 417 | -0.233838 | -0.792299 | 0.954683 | 1 |
| GOBP_NEGATIVE_REGULATION_OF_LIPID_LOCALIZATION | 51 | -0.278828 | -0.792077 | 0.795181 | 1 |
| GOBP_REGULATION_OF_OXIDATIVE_PHOSPHORYLATION | 24 | -0.314833 | -0.792028 | 0.750678 | 1 |
| GOBP_ANAPHASE_PROMOTING_COMPLEX_DEPENDENT_CATABOLIC_PROCES | 81 | -0.260545 | -0.79083 | 0.828704 | 1 |
| GOBP_SPLICEOSOMAL_COMPLEX_ASSEMBLY | 66 | -0.26924 | -0.790787 | 0.812717 | 1 |
| GOBP_NEGATIVE_REGULATION_OF_TRANSCRIPTION_REGULATORY_REGION_ | 19 | -0.328823 | -0.790643 | 0.768345 | 1 |
| GOBP_PURINE_CONTAINING_COMPOUND_BIOSYNTHETIC_PROCESS | 199 | -0.239704 | -0.79038 | 0.903564 | 1 |
| GOBP_LONG_CHAIN_FATTY_ACID_TRANSPORT | 65 | -0.267132 | -0.790365 | 0.833931 | 1 |
| GOBP_CHROMOSOME_LOCALIZATION | 77 | -0.263584 | -0.790345 | 0.842045 | 1 |
| GOBP_DNA_LIGATION | 15 | -0.342662 | -0.790291 | 0.744986 | 1 |
| GOBP_REGULATION_OF_CELL_FATE_COMMITMENT | 27 | -0.313805 | -0.790167 | 0.793997 | 1 |
| GOBP_POSITIVE_REGULATION_OF_CELL_AGING | 15 | -0.342423 | -0.789973 | 0.758087 | 1 |
| GOBP_NEGATIVE_REGULATION_OF_PROTEIN_CONTAINING_COMPLEX_ASSEM | 133 | -0.245129 | -0.789846 | 0.855422 | 1 |
| GOBP_ADRENERGIC_RECEPTOR_SIGNALING_PATHWAY | 25 | -0.310217 | -0.789815 | 0.77047 | 1 |
| GOBP_RESPONSE_TO_CAMP | 92 | -0.256537 | -0.789438 | 0.846748 | 1 |
| GOBP_T_HELPER_1_CELL_DIFFERENTIATION | 18 | -0.336268 | -0.789233 | 0.773533 | 1 |
| GOBP_TOLL_LIKE_RECEPTOR_9_SIGNALING_PATHWAY | 25 | -0.31603 | -0.788989 | 0.786189 | 1 |
| GOBP_POSITIVE_REGULATION_OF_RELEASE_OF_CYTOCHROME_C_FROM_MIT | 27 | -0.308358 | -0.788051 | 0.774406 | 1 |
| GOBP_VESICLE_MEDIATED_TRANSPORT_BETWEEN_ENDOSOMAL_COMPARTM | 43 | -0.28267 | -0.787679 | 0.789731 | 1 |
| GOBP_DNA_TEMPLATED_TRANSCRIPTION_INITIATION | 239 | -0.236388 | -0.787665 | 0.918557 | 1 |
| GOBP_PROTON_TRANSMEMBRANE_TRANSPORT | 150 | -0.243951 | -0.787368 | 0.888298 | 1 |
| GOBP_DNA_DEALKYLATION | 32 | -0.296187 | -0.787364 | 0.775408 | 1 |
| GOBP_PROTEIN_KINASE_B_SIGNALING | 241 | -0.23619 | -0.786962 | 0.923947 | 1 |
| GOBP_NEUROINFLAMMATORY_RESPONSE | 36 | -0.293237 | -0.786705 | 0.786164 | 1 |
| GOBP_HISTONE_H4_K16_ACETYLATION | 21 | -0.32073 | -0.78648 | 0.781818 | 1 |
| GOBP_RIBONUCLEOSIDE_MONOPHOSPHATE_METABOLIC_PROCESS | 55 | -0.271647 | -0.786259 | 0.802171 | 1 |
| GOBP_MICROGLIAL_CELL_ACTIVATION | 44 | -0.283892 | -0.786118 | 0.800752 | 1 |
| GOBP_REGULATION_OF_MONONUCLEAR_CELL_MIGRATION | 108 | -0.251734 | -0.786106 | 0.859513 | 1 |
| GOBP_REGULATION_OF_CYSTEINE_TYPE_ENDOPEPTIDASE_ACTIVITY | 223 | -0.237554 | -0.78598 | 0.926471 | 1 |
| GOBP_REGULATION_OF_CARDIAC_MUSCLE_CELL_DIFFERENTIATION | 19 | -0.327618 | -0.785884 | 0.761379 | 1 |
| GOBP_REGULATION_OF_LEUKOCYTE_MIGRATION | 198 | -0.238596 | -0.785597 | 0.902618 | 1 |
| GOBP_NEGATIVE_REGULATION_OF_OXIDATIVE_STRESS_INDUCED_NEURON_ | 16 | -0.341969 | -0.785374 | 0.755274 | 1 |
| GOBP_REGULATION_OF_EPIDERMAL_GROWTH_FACTOR_ACTIVATED_RECEPT | 30 | -0.299459 | -0.785229 | 0.794041 | 1 |
| GOBP_NEGATIVE_REGULATION_OF_PROTEIN_DEPHOSPHORYLATION | 62 | -0.269178 | -0.78514 | 0.82503 | 1 |
| GOBP_HISTONE_H3_K27_METHYLATION | 17 | -0.331383 | -0.785012 | 0.761494 | 1 |
| GOBP_POSITIVE_REGULATION_OF_INSULIN_SECRETION_INVOLVED_IN_CELLU | 31 | -0.302084 | -0.784646 | 0.81306 | 1 |
| GOBP_REPLACEMENT_OSSIFICATION | 28 | -0.301375 | -0.784105 | 0.79359 | 1 |
| GOBP_PURINE_CONTAINING_COMPOUND_TRANSMEMBRANE_TRANSPORT | 16 | -0.335645 | -0.783898 | 0.776056 | 1 |
| GOBP_PHOTORECEPTOR_CELL_DEVELOPMENT | 45 | -0.280404 | -0.78364 | 0.789281 | 1 |
| GOBP_RIBOSE_PHOSPHATE_BIOSYNTHETIC_PROCESS | 190 | -0.239028 | -0.783573 | 0.899577 | 1 |
| GOBP_POSITIVE_REGULATION_OF_CALCIUM_ION_IMPORT | 16 | -0.339045 | -0.783331 | 0.748954 | 1 |
| GOBP_NEGATIVE_REGULATION_OF_AXON_EXTENSION_INVOLVED_IN_AXON_( | 26 | -0.305663 | -0.783242 | 0.793103 | 1 |
| GOBP_NEGATIVE_REGULATION_OF_CELLULAR_SENESCENCE | 19 | -0.324199 | -0.783152 | 0.764626 | 1 |
| GOBP_NUCLEOTIDE_BINDING_DOMAIN_LEUCINE_RICH_REPEAT_CONTAINING | 41 | -0.282792 | -0.782851 | 0.794521 | 1 |
| GOBP_PORPHYRIN_CONTAINING_COMPOUND_METABOLIC_PROCESS | 37 | -0.289058 | -0.782694 | 0.802267 | 1 |
| GOBP_POSITIVE_REGULATION_OF_DNA_METABOLIC_PROCESS | 195 | -0.240484 | -0.782575 | 0.914761 | 1 |
| GOBP_CYTOPLASMIC_MICROTUBULE_ORGANIZATION | 62 | -0.26443 | -0.782184 | 0.826139 | 1 |
| GOBP_REGULATION_OF_NEUROGENESIS | 328 | -0.233103 | -0.782004 | 0.92638 | 1 |
| GOBP_LONG_TERM_SYNAPTIC_DEPRESSION | 29 | -0.305094 | -0.781724 | 0.77892 | 1 |
| GOBP_POSITIVE_REGULATION_OF_ADAPTIVE_IMMUNE_RESPONSE | 101 | -0.250993 | -0.781718 | 0.871766 | 1 |
| GOBP_NEGATIVE_REGULATION_OF_STRIATED_MUSCLE_CELL_DIFFERENTIAT | 34 | -0.290392 | -0.781709 | 0.799496 | 1 |
| GOBP_SODIUM_ION_TRANSPORT | 224 | -0.2354 | -0.781218 | 0.914079 | 1 |
| GOBP_PYRIMIDINE_RIBONUCLEOTIDE_BIOSYNTHETIC_PROCESS | 22 | -0.313346 | -0.780813 | 0.779392 | 1 |
| GOBP_AXONAL_FASCICULATION | 19 | -0.324163 | -0.780152 | 0.779661 | 1 |
| GOBP_LEUKOCYTE_TETHERING_OR_ROLLING | 30 | -0.30005 | -0.779444 | 0.797484 | 1 |
| GOBP_PROTEIN_INSERTION_INTO_MEMBRANE | 74 | -0.258762 | -0.779275 | 0.849425 | 1 |
| GOBP_COTRANSLATIONAL_PROTEIN_TARGETING_TO_MEMBRANE | 105 | -0.25026 | -0.779208 | 0.866446 | 1 |
| GOBP_NEGATIVE_REGULATION_OF_CYSTEINE_TYPE_ENDOPEPTIDASE_ACTIV | 75 | -0.260456 | -0.779206 | 0.841014 | 1 |
| GOBP_POSITIVE_REGULATION_OF_STEM_CELL_PROLIFERATION | 22 | -0.318954 | -0.77903 | 0.783634 | 1 |
| GOBP_REGULATION_OF_PHOSPHOLIPASE_C_ACTIVITY | 44 | -0.27997 | -0.778976 | 0.809102 | 1 |
| GOBP_REGULATION_OF_STEROID_METABOLIC_PROCESS | 110 | -0.250096 | -0.778968 | 0.864745 | 1 |
| GOBP_REGULATION_OF_GLIAL_CELL_DIFFERENTIATION | 58 | -0.266561 | -0.778157 | 0.827798 | 1 |
| GOBP_POSITIVE_REGULATION_OF_DENDRITE_MORPHOGENESIS | 35 | -0.289775 | -0.778142 | 0.809045 | 1 |
| GOBP_NEGATIVE_REGULATION_OF_LIPID_TRANSPORT | 34 | -0.290759 | -0.777217 | 0.821159 | 1 |
| GOBP_RIBOSOMAL_LARGE_SUBUNIT_BIOGENESIS | 72 | -0.260542 | -0.776714 | 0.842775 | 1 |
| GOBP_PYRIMIDINE_CONTAINING_COMPOUND_CATABOLIC_PROCESS | 37 | -0.284995 | -0.776543 | 0.810983 | 1 |
| GOBP_REGULATION_OF_CARBOHYDRATE_CATABOLIC_PROCESS | 87 | -0.252186 | -0.776308 | 0.85017 | 1 |
| GOBP_MESODERMAL_CELL_DIFFERENTIATION | 29 | -0.30332 | -0.776289 | 0.788312 | 1 |
| GOBP_GLIAL_CELL_DEVELOPMENT | 106 | -0.250995 | -0.776137 | 0.874016 | 1 |
| GOBP_REGULATION_OF_MAST_CELL_ACTIVATION | 41 | -0.28297 | -0.775867 | 0.802005 | 1 |
| GOBP_AXON_DEVELOPMENT | 492 | -0.225416 | -0.775225 | 0.971916 | 1 |
| GOBP_NEGATIVE_REGULATION_OF_INFLAMMATORY_RESPONSE | 124 | -0.243088 | -0.774971 | 0.878689 | 1 |
| GOBP_REGULATION_OF_HORMONE_BIOSYNTHETIC_PROCESS | 20 | -0.323409 | -0.774855 | 0.767538 | 1 |
| GOBP_RESPONSE_TO_COLD | 46 | -0.278199 | -0.774147 | 0.823245 | 1 |
| GOBP_RETINA_MORPHOGENESIS_IN_CAMERA_TYPE_EYE | 57 | -0.263581 | -0.774028 | 0.828979 | 1 |
| GOBP_ESTABLISHMENT_OF_ENDOTHELIAL_BARRIER | 48 | -0.278205 | -0.773569 | 0.830732 | 1 |
| GOBP_PORE_COMPLEX_ASSEMBLY | 20 | -0.318219 | -0.773281 | 0.76066 | 1 |
| GOBP_REGULATION_OF_T_HELPER_CELL_DIFFERENTIATION | 38 | -0.283689 | -0.772664 | 0.816121 | 1 |
| GOBP_NEGATIVE_REGULATION_OF_EPITHELIAL_CELL_PROLIFERATION | 124 | -0.244244 | -0.772537 | 0.87365 | 1 |
| GOBP_MITOTIC_SPINDLE_ORGANIZATION | 117 | -0.244873 | -0.772501 | 0.87965 | 1 |
| GOBP_PEPTIDYL_TYROSINE_MODIFICATION | 357 | -0.228485 | -0.772161 | 0.968655 | 1 |
| GOBP_POSITIVE_REGULATION_OF_CELL_ADHESION_MEDIATED_BY_INTEGRIN | 21 | -0.315522 | -0.772011 | 0.81241 | 1 |
| GOBP_PHOSPHOLIPID_TRANSPORT | 88 | -0.252844 | -0.771775 | 0.872461 | 1 |
| GOBP_NEGATIVE_REGULATION_OF_CELL_ADHESION | 270 | -0.230202 | -0.771768 | 0.935845 | 1 |
| GOBP_NEURAL_PRECURSOR_CELL_PROLIFERATION | 134 | -0.241743 | -0.771761 | 0.883978 | 1 |
| GOBP_EMBRYONIC_PATTERN_SPECIFICATION | 60 | -0.258882 | -0.771158 | 0.827014 | 1 |
| GOBP_HORMONE_TRANSPORT | 293 | -0.228756 | -0.771118 | 0.945344 | 1 |
| GOBP_CELLULAR_GLUCOSE_HOMEOSTASIS | 131 | -0.242606 | -0.771047 | 0.901182 | 1 |
| GOBP_REGULATION_OF_RECEPTOR_RECYCLING | 23 | -0.311122 | -0.770743 | 0.776871 | 1 |
| GOBP_LOW_DENSITY_LIPOPROTEIN_PARTICLE_CLEARANCE | 34 | -0.287404 | -0.77074 | 0.784314 | 1 |
| GOBP_CELLULAR_RESPONSE_TO_DRUG | 61 | -0.26193 | -0.770273 | 0.830389 | 1 |
| GOBP_GLUCOSE_METABOLIC_PROCESS | 196 | -0.234183 | -0.770086 | 0.922756 | 1 |
| GOBP_REGULATION_OF_CELL_CYCLE_CHECKPOINT | 31 | -0.292084 | -0.769958 | 0.819693 | 1 |

| | | | | | |
|---|---|---|---|---|---|
| GOBP_DNA_STRAND_ELONGATION | 26 | -0.303595 | -0.769883 | 0.792732 | 1 |
| GOBP_SYNAPSE_ORGANIZATION | 400 | -0.225242 | -0.769582 | 0.968876 | 1 |
| GOBP_DENDRITE_SELF_AVOIDANCE | 17 | -0.329826 | -0.769553 | 0.77973 | 1 |
| GOBP_REGULATION_OF_LIPID_METABOLIC_PROCESS | 372 | -0.225561 | -0.76884 | 0.965657 | 1 |
| GOBP_SMOOTH_MUSCLE_CELL_APOPTOTIC_PROCESS | 23 | -0.305841 | -0.768743 | 0.791395 | 1 |
| GOBP_CALCIUM_ION_REGULATED_EXOCYTOSIS_OF_NEUROTRANSMITTER | 17 | -0.33388 | -0.767814 | 0.79212 | 1 |
| GOBP_POSITIVE_REGULATION_OF_EMBRYONIC_DEVELOPMENT | 19 | -0.316436 | -0.767506 | 0.783083 | 1 |
| GOBP_RAC_PROTEIN_SIGNAL_TRANSDUCTION | 39 | -0.278586 | -0.767474 | 0.827543 | 1 |
| GOBP_REGULATION_OF_CARBOHYDRATE_METABOLIC_PROCESS | 196 | -0.233628 | -0.767336 | 0.928274 | 1 |
| GOBP_PEPTIDYL_LYSINE_DIMETHYLATION | 21 | -0.314001 | -0.767295 | 0.789618 | 1 |
| GOBP_RNA_DESTABILIZATION | 34 | -0.285193 | -0.767138 | 0.819462 | 1 |
| GOBP_AXONEMAL_DYNEIN_COMPLEX_ASSEMBLY | 31 | -0.293059 | -0.76681 | 0.808184 | 1 |
| GOBP_POSITIVE_REGULATION_OF_SMALL_GTPASE_MEDIATED_SIGNAL_TRAN | 76 | -0.255413 | -0.766141 | 0.862587 | 1 |
| GOBP_V_D_J_RECOMBINATION | 15 | -0.332828 | -0.766126 | 0.792958 | 1 |
| GOBP_ACUTE_INFLAMMATORY_RESPONSE_TO_ANTIGENIC_STIMULUS | 21 | -0.312473 | -0.765918 | 0.789614 | 1 |
| GOBP_POTASSIUM_ION_TRANSPORT | 211 | -0.233158 | -0.765894 | 0.91684 | 1 |
| GOBP_MYELOID_DENDRITIC_CELL_DIFFERENTIATION | 20 | -0.313316 | -0.765839 | 0.765252 | 1 |
| GOBP_REGULATION_OF_SYSTEMIC_ARTERIAL_BLOOD_PRESSURE_BY_CIRCU | 18 | -0.325091 | -0.765614 | 0.776685 | 1 |
| GOBP_MYOFIBRIL_ASSEMBLY | 55 | -0.263424 | -0.765537 | 0.848131 | 1 |
| GOBP_PROTEIN_MATURATION | 278 | -0.229294 | -0.765431 | 0.955193 | 1 |
| GOBP_NEGATIVE_REGULATION_OF_PROTEIN_KINASE_B_SIGNALING | 46 | -0.274602 | -0.765246 | 0.836585 | 1 |
| GOBP_MACROPHAGE_CHEMOTAXIS | 38 | -0.280016 | -0.765218 | 0.824561 | 1 |
| GOBP_CELLULAR_CARBOHYDRATE_METABOLIC_PROCESS | 273 | -0.22864 | -0.765214 | 0.951894 | 1 |
| GOBP_MONOSACCHARIDE_METABOLIC_PROCESS | 255 | -0.22787 | -0.764699 | 0.934426 | 1 |
| GOBP_MAST_CELL_ACTIVATION | 60 | -0.257954 | -0.764578 | 0.840047 | 1 |
| GOBP_RESPONSE_TO_PROTOZOAN | 22 | -0.310157 | -0.764552 | 0.802419 | 1 |
| GOBP_POSITIVE_REGULATION_OF_PROTEIN_IMPORT | 40 | -0.282631 | -0.764004 | 0.823009 | 1 |
| GOBP_POSITIVE_REGULATION_OF_PEPTIDE_SECRETION | 142 | -0.239872 | -0.763935 | 0.911669 | 1 |
| GOBP_POSITIVE_REGULATION_OF_ESTABLISHMENT_OF_PROTEIN_LOCALIZAT | 58 | -0.263709 | -0.763904 | 0.860382 | 1 |
| GOBP_MEGAKARYOCYTE_DIFFERENTIATION | 91 | -0.248413 | -0.76372 | 0.865622 | 1 |
| GOBP_NEGATIVE_REGULATION_OF_T_HELPER_CELL_DIFFERENTIATION | 18 | -0.32463 | -0.76355 | 0.811653 | 1 |
| GOBP_POSITIVE_REGULATION_OF_TYROSINE_PHOSPHORYLATION_OF_STAT_ | 61 | -0.26098 | -0.763387 | 0.851508 | 1 |
| GOBP_REGULATION_OF_SYNAPTIC_TRANSMISSION_GLUTAMATERGIC | 62 | -0.265106 | -0.763266 | 0.838184 | 1 |
| GOBP_POSITIVE_REGULATION_OF_INTERLEUKIN_12_PRODUCTION | 39 | -0.281356 | -0.763011 | 0.837388 | 1 |
| GOBP_REGULATION_OF_SPINDLE_ASSEMBLY | 25 | -0.298922 | -0.762976 | 0.802853 | 1 |
| GOBP_POSITIVE_REGULATION_OF_MITOCHONDRION_ORGANIZATION | 83 | -0.250922 | -0.762918 | 0.857974 | 1 |
| GOBP_POSITIVE_REGULATION_OF_MESENCHYMAL_CELL_PROLIFERATION | 21 | -0.310445 | -0.761937 | 0.789905 | 1 |
| GOBP_REGULATION_OF_EPITHELIAL_CELL_DIFFERENTIATION_INVOLVED_IN_I | 18 | -0.324069 | -0.761807 | 0.797886 | 1 |
| GOBP_REGULATION_OF_PEPTIDE_HORMONE_SECRETION | 196 | -0.231123 | -0.761725 | 0.920252 | 1 |
| GOBP_POSITIVE_REGULATION_OF_SMOOTH_MUSCLE_CELL_PROLIFERATION | 85 | -0.249919 | -0.761616 | 0.865826 | 1 |
| GOBP_CARBOHYDRATE_HOMEOSTASIS | 207 | -0.231331 | -0.761416 | 0.922037 | 1 |
| GOBP_INTESTINAL_LIPID_ABSORPTION | 16 | -0.326471 | -0.761222 | 0.767635 | 1 |
| GOBP_NEGATIVE_REGULATION_OF_CD4_POSITIVE_ALPHA_BETA_T_CELL_DIF | 22 | -0.307689 | -0.760994 | 0.793103 | 1 |
| GOBP_REGULATION_OF_ALTERNATIVE_MRNA_SPLICING_VIA_SPLICEOSOME | 55 | -0.262958 | -0.760873 | 0.843897 | 1 |
| GOBP_POSITIVE_REGULATION_OF_TRANSFORMING_GROWTH_FACTOR_BETA_ | 21 | -0.317755 | -0.760653 | 0.814324 | 1 |
| GOBP_RESPONSE_TO_PLATELET_DERIVED_GROWTH_FACTOR | 19 | -0.316605 | -0.760545 | 0.794278 | 1 |
| GOBP_REGULATION_OF_LYMPHOCYTE_MIGRATION | 61 | -0.26179 | -0.760089 | 0.848629 | 1 |
| GOBP_CERAMIDE_METABOLIC_PROCESS | 99 | -0.243274 | -0.76006 | 0.880361 | 1 |
| GOBP_REGULATION_OF_HISTONE_H3_K4_METHYLATION | 26 | -0.300731 | -0.759926 | 0.798693 | 1 |
| GOBP_PHOSPHATIDYLCHOLINE_BIOSYNTHETIC_PROCESS | 37 | -0.27624 | -0.759333 | 0.818296 | 1 |
| GOBP_NEGATIVE_REGULATION_OF_SMOOTH_MUSCLE_CELL_PROLIFERATION | 43 | -0.273987 | -0.758806 | 0.828183 | 1 |
| GOBP_SULFUR_COMPOUND_TRANSPORT | 48 | -0.265596 | -0.758742 | 0.837065 | 1 |
| GOBP_NUCLEOSIDE_METABOLIC_PROCESS | 101 | -0.245165 | -0.758322 | 0.880979 | 1 |
| GOBP_ORGANELLE_INHERITANCE | 15 | -0.328887 | -0.758252 | 0.791724 | 1 |
| GOBP_NEGATIVE_REGULATION_OF_CYTOKINE_PRODUCTION_INVOLVED_IN_II | 22 | -0.306262 | -0.758086 | 0.813333 | 1 |
| GOBP_REGULATION_OF_REACTIVE_OXYGEN_SPECIES_BIOSYNTHETIC_PROC | 87 | -0.247745 | -0.757917 | 0.879587 | 1 |
| GOBP_EPITHELIAL_CELL_DIFFERENTIATION_INVOLVED_IN_KIDNEY_DEVELOP | 47 | -0.266619 | -0.757871 | 0.830935 | 1 |
| GOBP_POSITIVE_REGULATION_OF_PHAGOCYTOSIS | 62 | -0.258644 | -0.757753 | 0.843897 | 1 |
| GOBP_NEUROMUSCULAR_SYNAPTIC_TRANSMISSION | 22 | -0.303769 | -0.757752 | 0.816107 | 1 |
| GOBP_RUFFLE_ASSEMBLY | 42 | -0.269299 | -0.756891 | 0.834356 | 1 |
| GOBP_NEGATIVE_REGULATION_OF_DNA_BINDING_TRANSCRIPTION_FACTOR_ | 156 | -0.235051 | -0.756845 | 0.925373 | 1 |
| GOBP_MESENCHYMAL_CELL_DIFFERENTIATION | 221 | -0.227004 | -0.756663 | 0.928945 | 1 |
| GOBP_REGULATION_OF_CENTROSOME_CYCLE | 48 | -0.267042 | -0.756164 | 0.850877 | 1 |
| GOBP_NEGATIVE_REGULATION_OF_RECEPTOR_SIGNALING_PATHWAY_VIA_S | 24 | -0.29553 | -0.756032 | 0.814077 | 1 |
| GOBP_PEPTIDE_HORMONE_SECRETION | 232 | -0.226203 | -0.754522 | 0.940452 | 1 |
| GOBP_NEGATIVE_REGULATION_OF_CELL_CYCLE_G2_M_PHASE_TRANSITION | 107 | -0.240779 | -0.754494 | 0.899123 | 1 |
| GOBP_RNA_DECAPPING | 16 | -0.32276 | -0.754337 | 0.798571 | 1 |
| GOBP_KERATAN_SULFATE_BIOSYNTHETIC_PROCESS | 28 | -0.29035 | -0.754216 | 0.826701 | 1 |
| GOBP_SPINDLE_LOCALIZATION | 52 | -0.264481 | -0.753705 | 0.845969 | 1 |
| GOBP_CARBOHYDRATE_BIOSYNTHETIC_PROCESS | 198 | -0.230488 | -0.753666 | 0.942105 | 1 |
| GOBP_PHOSPHATIDYLINOSITOL_METABOLIC_PROCESS | 183 | -0.231345 | -0.753661 | 0.923738 | 1 |
| GOBP_REGULATION_OF_GRANULOCYTE_CHEMOTAXIS | 48 | -0.265254 | -0.75359 | 0.847356 | 1 |
| GOBP_POSITIVE_REGULATION_OF_CELL_DIVISION | 74 | -0.247413 | -0.753569 | 0.873256 | 1 |
| GOBP_BONE_RESORPTION | 59 | -0.259146 | -0.753494 | 0.869617 | 1 |
| GOBP_MITOTIC_CYTOKINESIS | 70 | -0.248923 | -0.75333 | 0.885057 | 1 |
| GOBP_ENDOCRINE_PROCESS | 73 | -0.249918 | -0.75329 | 0.872979 | 1 |
| GOBP_POSITIVE_REGULATION_OF_INTERFERON_BETA_PRODUCTION | 33 | -0.281394 | -0.753283 | 0.845758 | 1 |
| GOBP_REGULATION_OF_CLATHRIN_DEPENDENT_ENDOCYTOSIS | 19 | -0.311814 | -0.753167 | 0.811141 | 1 |
| GOBP_RESPONSE_TO_COCAINE | 39 | -0.280312 | -0.753135 | 0.830635 | 1 |
| GOBP_NEGATIVE_REGULATION_OF_MAPK_CASCADE | 164 | -0.232348 | -0.752824 | 0.921444 | 1 |
| GOBP_MITOTIC_SPINDLE_ASSEMBLY | 65 | -0.251631 | -0.752687 | 0.864929 | 1 |
| GOBP_NEGATIVE_REGULATION_OF_ATP_METABOLIC_PROCESS | 28 | -0.288046 | -0.752572 | 0.809769 | 1 |
| GOBP_PROTEIN_TRANSPORT_ALONG_MICROTUBULE | 68 | -0.253217 | -0.752219 | 0.862471 | 1 |
| GOBP_REGULATION_OF_ORGAN_GROWTH | 89 | -0.244865 | -0.751323 | 0.889515 | 1 |
| GOBP_CELLULAR_RESPONSE_TO_ARSENIC_CONTAINING_SUBSTANCE | 18 | -0.316327 | -0.75117 | 0.825676 | 1 |
| GOBP_HEMATOPOIETIC_STEM_CELL_DIFFERENTIATION | 87 | -0.246199 | -0.75093 | 0.881336 | 1 |
| GOBP_RNA_CATABOLIC_PROCESS | 401 | -0.221481 | -0.750841 | 0.963783 | 1 |
| GOBP_MECHANORECEPTOR_DIFFERENTIATION | 58 | -0.25678 | -0.750828 | 0.871886 | 1 |
| GOBP_REGULATION_OF_CELLULAR_AMIDE_METABOLIC_PROCESS | 413 | -0.22123 | -0.750777 | 0.969849 | 1 |
| GOBP_POLARIZED_EPITHELIAL_CELL_DIFFERENTIATION | 22 | -0.300011 | -0.750363 | 0.815115 | 1 |
| GOBP_CARTILAGE_DEVELOPMENT | 172 | -0.230449 | -0.75023 | 0.930962 | 1 |
| GOBP_PTERIDINE_CONTAINING_COMPOUND_METABOLIC_PROCESS | 33 | -0.279729 | -0.750199 | 0.823163 | 1 |
| GOBP_REGULATION_OF_PLATELET_DERIVED_GROWTH_FACTOR_RECEPTOR_ | 22 | -0.304248 | -0.750162 | 0.812 | 1 |
| GOBP_NEGATIVE_REGULATION_OF_MITOTIC_CELL_CYCLE | 297 | -0.221371 | -0.750047 | 0.969543 | 1 |
| GOBP_POSITIVE_REGULATION_OF_ANION_TRANSPORT | 460 | -0.22082 | -0.748705 | 0.97994 | 1 |
| GOBP_REGULATION_OF_CATION_TRANSMEMBRANE_TRANSPORT | 320 | -0.221677 | -0.747576 | 0.969666 | 1 |
| GOBP_SEX_DIFFERENTIATION | 239 | -0.225166 | -0.747507 | 0.948025 | 1 |
| GOBP_POSITIVE_REGULATION_OF_INTRACELLULAR_TRANSPORT | 216 | -0.226086 | -0.747136 | 0.95243 | 1 |
| GOBP_CALCIUM_ION_IMPORT_INTO_CYTOSOL | 21 | -0.303099 | -0.747016 | 0.814714 | 1 |
| GOBP_NEGATIVE_REGULATION_OF_HISTONE_METHYLATION | 21 | -0.303506 | -0.746996 | 0.817105 | 1 |
| GOBP_REGULATION_OF_GLIAL_CELL_PROLIFERATION | 29 | -0.285556 | -0.746966 | 0.805951 | 1 |
| GOBP_ORGANOPHOSPHATE_ESTER_TRANSPORT | 125 | -0.235462 | -0.74695 | 0.90642 | 1 |
| GOBP_CELL_CELL_JUNCTION_ASSEMBLY | 138 | -0.234182 | -0.746533 | 0.913326 | 1 |
| GOBP_MICROTUBULE_BASED_MOVEMENT | 351 | -0.220477 | -0.746279 | 0.977778 | 1 |
| GOBP_NEGATIVE_REGULATION_OF_REACTIVE_OXYGEN_SPECIES_METABOLIC | 54 | -0.259444 | -0.746192 | 0.863473 | 1 |
| GOBP_TRANSLATIONAL_INITIATION | 187 | -0.226866 | -0.745965 | 0.944964 | 1 |
| GOBP_LIPID_HOMEOSTASIS | 139 | -0.233901 | -0.745964 | 0.922414 | 1 |
| GOBP_METHYLATION | 342 | -0.222009 | -0.745945 | 0.974671 | 1 |
| GOBP_NEGATIVE_REGULATION_OF_CELLULAR_RESPONSE_TO_INSULIN_STIM | 35 | -0.278637 | -0.745908 | 0.840731 | 1 |
| GOBP_POSITIVE_REGULATION_OF_PRODUCTION_OF_MOLECULAR_MEDIATOR | 96 | -0.245161 | -0.745217 | 0.904707 | 1 |
| GOBP_GANGLIOSIDE_METABOLIC_PROCESS | 23 | -0.298279 | -0.744805 | 0.827078 | 1 |
| GOBP_L_GLUTAMATE_TRANSMEMBRANE_TRANSPORT | 24 | -0.298457 | -0.744686 | 0.808786 | 1 |
| GOBP_ESTABLISHMENT_OF_PROTEIN_LOCALIZATION_TO_MITOCHONDRIAL_M | 53 | -0.254316 | -0.744513 | 0.874101 | 1 |
| GOBP_REGULATION_OF_TISSUE_REMODELING | 71 | -0.249708 | -0.744304 | 0.872792 | 1 |
| GOBP_POSITIVE_REGULATION_OF_INTRACELLULAR_PROTEIN_TRANSPORT | 178 | -0.227646 | -0.744206 | 0.936306 | 1 |
| GOBP_RUFFLE_ORGANIZATION | 54 | -0.256586 | -0.7442 | 0.869723 | 1 |
| GOBP_PSEUDOURIDINE_SYNTHESIS | 18 | -0.317166 | -0.744091 | 0.829932 | 1 |
| GOBP_MATURATION_OF_5_8S_RRNA_FROM_TRICISTRONIC_RRNA_TRANSCRI | 24 | -0.294209 | -0.74378 | 0.812336 | 1 |
| GOBP_LAMELLIPODIUM_ASSEMBLY | 68 | -0.251763 | -0.743015 | 0.872812 | 1 |
| GOBP_PHAGOSOME_ACIDIFICATION | 27 | -0.290064 | -0.742738 | 0.826772 | 1 |
| GOBP_P_BODY_ASSEMBLY | 21 | -0.304656 | -0.742333 | 0.816733 | 1 |
| GOBP_POSITIVE_REGULATION_OF_PATTERN_RECOGNITION_RECEPTOR_SIGN | 41 | -0.271829 | -0.742301 | 0.841772 | 1 |
| GOBP_REGULATION_OF_CHROMATIN_ORGANIZATION | 178 | -0.227755 | -0.741848 | 0.952128 | 1 |
| GOBP_ENDOTHELIAL_CELL_PROLIFERATION | 151 | -0.229264 | -0.741786 | 0.933761 | 1 |
| GOBP_INTERLEUKIN_13_PRODUCTION | 18 | -0.311201 | -0.741458 | 0.83 | 1 |
| GOBP_RESPONSE_TO_RETINOIC_ACID | 101 | -0.237244 | -0.740992 | 0.898096 | 1 |
| GOBP_PHOSPHOLIPID_BIOSYNTHETIC_PROCESS | 260 | -0.221818 | -0.740748 | 0.955988 | 1 |
| GOBP_MATURATION_OF_SSU_RRNA_FROM_TRICISTRONIC_RRNA_TRANSCRI | 37 | -0.268928 | -0.740343 | 0.857868 | 1 |
| GOBP_ESTABLISHMENT_OF_PROTEIN_LOCALIZATION_TO_CHROMOSOME | 27 | -0.292017 | -0.740273 | 0.832902 | 1 |
| GOBP_RESPONSE_TO_PEPTIDE_HORMONE | 403 | -0.217064 | -0.740002 | 0.987879 | 1 |
| GOBP_RIBONUCLEOPROTEIN_COMPLEX_SUBUNIT_ORGANIZATION | 211 | -0.223335 | -0.739334 | 0.950515 | 1 |
| GOBP_NEGATIVE_REGULATION_OF_ORGANELLE_ORGANIZATION | 334 | -0.218423 | -0.73906 | 0.971631 | 1 |
| GOBP_POSITIVE_REGULATION_OF_GLYCOPROTEIN_METABOLIC_PROCESS | 25 | -0.288644 | -0.738703 | 0.834403 | 1 |
| GOBP_REGULATION_OF_DEPHOSPHORYLATION | 204 | -0.225211 | -0.738643 | 0.948905 | 1 |
| GOBP_RESPONSE_TO_ETHANOL | 119 | -0.234886 | -0.738293 | 0.918771 | 1 |
| GOBP_POSITIVE_REGULATION_OF_DENDRITIC_SPINE_MORPHOGENESIS | 19 | -0.305701 | -0.73829 | 0.831933 | 1 |
| GOBP_DENDRITIC_SPINE_MORPHOGENESIS | 60 | -0.252438 | -0.738224 | 0.877962 | 1 |
| GOBP_POSITIVE_REGULATION_OF_NF_KAPPAB_TRANSCRIPTION_FACTOR_AC | 156 | -0.226891 | -0.737981 | 0.945711 | 1 |

| | | | | | |
|---|---|---|---|---|---|
| GOBP_SEQUESTERING_OF_TRIGLYCERIDE | 16 | -0.325012 | -0.737553 | 0.826399 | 1 |
| GOBP_REGULATION_OF_SMALL_MOLECULE_METABOLIC_PROCESS | 403 | -0.217138 | -0.737143 | 0.986922 | 1 |
| GOBP_REGULATION_OF_CHROMOSOME_ORGANIZATION | 267 | -0.221361 | -0.737116 | 0.969481 | 1 |
| GOBP_MAMMARY_GLAND_ALVEOLUS_DEVELOPMENT | 17 | -0.317982 | -0.73709 | 0.824684 | 1 |
| GOBP_RENAL_SYSTEM_PROCESS_INVOLVED_IN_REGULATION_OF_SYSTEMIC | 22 | -0.299859 | -0.73698 | 0.826613 | 1 |
| GOBP_REGULATION_OF_MYELOID_LEUKOCYTE_DIFFERENTIATION | 111 | -0.237014 | -0.736959 | 0.923418 | 1 |
| GOBP_REGULATION_OF_MICROTUBULE_BASED_PROCESS | 224 | -0.221066 | -0.73681 | 0.964912 | 1 |
| GOBP_ACTIN_POLYMERIZATION_OR_DEPOLYMERIZATION | 211 | -0.223596 | -0.736279 | 0.963312 | 1 |
| GOBP_REGULATION_OF_TRANSPORTER_ACTIVITY | 263 | -0.219267 | -0.735884 | 0.954309 | 1 |
| GOBP_TRANSCRIPTION_INITIATION_FROM_RNA_POLYMERASE_II_PROMOTER | 183 | -0.22375 | -0.735812 | 0.949738 | 1 |
| GOBP_REGULATION_OF_PHAGOCYTOSIS | 88 | -0.242423 | -0.735533 | 0.911665 | 1 |
| GOBP_MITOCHONDRION_DISTRIBUTION | 16 | -0.311209 | -0.734949 | 0.82027 | 1 |
| GOBP_POSITIVE_REGULATION_OF_RECEPTOR_MEDIATED_ENDOCYTOSIS | 50 | -0.261186 | -0.734704 | 0.893463 | 1 |
| GOBP_HISTONE_H3_K4_METHYLATION | 50 | -0.254944 | -0.734331 | 0.877278 | 1 |
| GOBP_REGULATION_OF_GLUCOSE_METABOLIC_PROCESS | 112 | -0.231526 | -0.734294 | 0.92613 | 1 |
| GOBP_REGULATION_OF_HORMONE_SECRETION | 244 | -0.219933 | -0.733787 | 0.965128 | 1 |
| GOBP_MAINTENANCE_OF_PROTEIN_LOCATION_IN_CELL | 63 | -0.249302 | -0.733634 | 0.894919 | 1 |
| GOBP_NEGATIVE_REGULATION_OF_CELL_CELL_ADHESION | 175 | -0.224557 | -0.733456 | 0.938947 | 1 |
| GOBP_RESPONSE_TO_TUMOR_NECROSIS_FACTOR | 303 | -0.218596 | -0.733307 | 0.970468 | 1 |
| GOBP_REGULATION_OF_PEPTIDYL_TYROSINE_PHOSPHORYLATION | 248 | -0.220195 | -0.732986 | 0.967942 | 1 |
| GOBP_SPHINGOLIPID_BIOSYNTHETIC_PROCESS | 108 | -0.23387 | -0.73278 | 0.908886 | 1 |
| GOBP_ESTABLISHMENT_OF_PROTEIN_LOCALIZATION_TO_ENDOPLASMIC_RET | 119 | -0.231266 | -0.73268 | 0.931818 | 1 |
| GOBP_AXON_ENSHEATHMENT_IN_CENTRAL_NERVOUS_SYSTEM | 18 | -0.308343 | -0.732432 | 0.82345 | 1 |
| GOBP_NEGATIVE_REGULATION_OF_PROTEIN_LOCALIZATION_TO_CELL_PERIP | 23 | -0.291751 | -0.732367 | 0.830912 | 1 |
| GOBP_TYPE_B_PANCREATIC_CELL_PROLIFERATION | 19 | -0.309958 | -0.732095 | 0.830688 | 1 |
| GOBP_RESPONSE_TO_ISOQUINOLINE_ALKALOID | 26 | -0.288953 | -0.731644 | 0.832258 | 1 |
| GOBP_MEMBRANE_LIPID_CATABOLIC_PROCESS | 37 | -0.268445 | -0.731614 | 0.865711 | 1 |
| GOBP_ORGANOPHOSPHATE_CATABOLIC_PROCESS | 149 | -0.227744 | -0.731251 | 0.929638 | 1 |
| GOBP_ESTABLISHMENT_OF_EPITHELIAL_CELL_APICAL_BASAL_POLARITY | 15 | -0.320833 | -0.730961 | 0.84233 | 1 |
| GOBP_POSITIVE_REGULATION_OF_CELL_GROWTH | 149 | -0.22584 | -0.730726 | 0.939426 | 1 |
| GOBP_CYTOPLASMIC_TRANSLATION | 99 | -0.233543 | -0.730701 | 0.932203 | 1 |
| GOBP_REGULATION_OF_HAIR_CYCLE | 26 | -0.286605 | -0.730566 | 0.839523 | 1 |
| GOBP_INTERLEUKIN_10_PRODUCTION | 57 | -0.251165 | -0.729649 | 0.897094 | 1 |
| GOBP_CERAMIDE_BIOSYNTHETIC_PROCESS | 67 | -0.247187 | -0.729649 | 0.90625 | 1 |
| GOBP_REGULATION_OF_TRANSLATION_NCRNA_MEDIATED | 28 | -0.281077 | -0.729301 | 0.838254 | 1 |
| GOBP_REGULATION_OF_ESTABLISHMENT_OR_MAINTENANCE_OF_CELL_POLA | 25 | -0.285501 | -0.729252 | 0.828309 | 1 |
| GOBP_CLEAVAGE_INVOLVED_IN_RRNA_PROCESSING | 27 | -0.278944 | -0.728979 | 0.837981 | 1 |
| GOBP_RESPONSE_TO_PROSTAGLANDIN_E | 23 | -0.294464 | -0.728588 | 0.839189 | 1 |
| GOBP_SMOOTH_MUSCLE_CELL_PROLIFERATION | 132 | -0.229596 | -0.728025 | 0.930207 | 1 |
| GOBP_REGULATION_OF_KERATINOCYTE_DIFFERENTIATION | 38 | -0.266809 | -0.72787 | 0.868421 | 1 |
| GOBP_INTERLEUKIN_8_PRODUCTION | 80 | -0.24025 | -0.727562 | 0.909091 | 1 |
| GOBP_PROTEIN_MODIFICATION_BY_SMALL_PROTEIN_REMOVAL | 274 | -0.217029 | -0.727403 | 0.971399 | 1 |
| GOBP_VASCULAR_TRANSPORT | 86 | -0.23888 | -0.72734 | 0.910532 | 1 |
| GOBP_NUCLEOSIDE_MONOPHOSPHATE_METABOLIC_PROCESS | 72 | -0.244445 | -0.726893 | 0.911494 | 1 |
| GOBP_PROTEIN_LOCALIZATION_TO_CHROMOSOME | 90 | -0.236161 | -0.726056 | 0.9098 | 1 |
| GOBP_NEGATIVE_REGULATION_OF_ALPHA_BETA_T_CELL_DIFFERENTIATION | 26 | -0.288293 | -0.725394 | 0.841755 | 1 |
| GOBP_NEGATIVE_REGULATION_OF_TRANSFERASE_ACTIVITY | 281 | -0.217916 | -0.725228 | 0.973442 | 1 |
| GOBP_MEMBRANE_PROTEIN_ECTODOMAIN_PROTEOLYSIS | 42 | -0.256348 | -0.724998 | 0.879093 | 1 |
| GOBP_NEGATIVE_REGULATION_OF_CD4_POSITIVE_ALPHA_BETA_T_CELL_AC | 31 | -0.273848 | -0.724961 | 0.866317 | 1 |
| GOBP_REGULATION_OF_ACTIN_FILAMENT_ORGANIZATION | 267 | -0.21716 | -0.72461 | 0.979339 | 1 |
| GOBP_REGULATION_OF_GLYCOPROTEIN_METABOLIC_PROCESS | 46 | -0.257907 | -0.724346 | 0.87202 | 1 |
| GOBP_POSITIVE_REGULATION_OF_G_PROTEIN_COUPLED_RECEPTOR_SIGNA | 24 | -0.287812 | -0.724155 | 0.859355 | 1 |
| GOBP_REGULATION_OF_STEROID_BIOSYNTHETIC_PROCESS | 83 | -0.236436 | -0.723458 | 0.918203 | 1 |
| GOBP_NEGATIVE_REGULATION_OF_PROTEIN_LOCALIZATION_TO_MEMBRANE | 28 | -0.278314 | -0.723436 | 0.868831 | 1 |
| GOBP_SYNAPTIC_VESICLE_LOCALIZATION | 54 | -0.252754 | -0.72342 | 0.911765 | 1 |
| GOBP_MONOVALENT_INORGANIC_ANION_HOMEOSTASIS | 15 | -0.311721 | -0.723366 | 0.83711 | 1 |
| GOBP_PYRIMIDINE_CONTAINING_COMPOUND_BIOSYNTHETIC_PROCESS | 39 | -0.264051 | -0.722742 | 0.869889 | 1 |
| GOBP_REGULATION_OF_CELL_AGING | 49 | -0.255893 | -0.722725 | 0.890229 | 1 |
| GOBP_POSITIVE_REGULATION_OF_DNA_BIOSYNTHETIC_PROCESS | 65 | -0.244886 | -0.722684 | 0.889535 | 1 |
| GOBP_MULTI_ORGANISM_CELLULAR_PROCESS | 16 | -0.310409 | -0.722431 | 0.829235 | 1 |
| GOBP_NUCLEAR_TRANSCRIBED_MRNA_CATABOLIC_PROCESS_DEADENYLAT | 75 | -0.239409 | -0.722141 | 0.895439 | 1 |
| GOBP_MESENCHYMAL_CELL_PROLIFERATION | 37 | -0.263837 | -0.721806 | 0.883249 | 1 |
| GOBP_REGULATION_OF_T_HELPER_1_TYPE_IMMUNE_RESPONSE | 26 | -0.286798 | -0.721274 | 0.857143 | 1 |
| GOBP_RENAL_FILTRATION | 25 | -0.287689 | -0.721005 | 0.860343 | 1 |
| GOBP_REGULATION_OF_RYANODINE_SENSITIVE_CALCIUM_RELEASE_CHANN | 26 | -0.281722 | -0.720967 | 0.843164 | 1 |
| GOBP_POSITIVE_REGULATION_OF_CELLULAR_EXTRAVASATION | 16 | -0.314943 | -0.720628 | 0.851034 | 1 |
| GOBP_GLAND_DEVELOPMENT | 394 | -0.212767 | -0.720279 | 0.986842 | 1 |
| GOBP_INDOLALKYLAMINE_METABOLIC_PROCESS | 16 | -0.309453 | -0.720188 | 0.838889 | 1 |
| GOBP_PEPTIDYL_LYSINE_METHYLATION | 118 | -0.228283 | -0.720158 | 0.938731 | 1 |
| GOBP_POSITIVE_REGULATION_OF_TELOMERE_MAINTENANCE | 51 | -0.250015 | -0.719946 | 0.895262 | 1 |
| GOBP_RESPONSE_TO_INSULIN | 262 | -0.217551 | -0.719907 | 0.978484 | 1 |
| GOBP_NEGATIVE_REGULATION_OF_LIPID_METABOLIC_PROCESS | 86 | -0.234695 | -0.719742 | 0.933107 | 1 |
| GOBP_NEGATIVE_REGULATION_OF_DEFENSE_RESPONSE | 196 | -0.220455 | -0.719198 | 0.96358 | 1 |
| GOBP_PH_REDUCTION | 55 | -0.249113 | -0.718967 | 0.902381 | 1 |
| GOBP_PROTEIN_COMPLEX_OLIGOMERIZATION | 222 | -0.216575 | -0.718813 | 0.980311 | 1 |
| GOBP_CELLULAR_MONOVALENT_INORGANIC_CATION_HOMEOSTASIS | 112 | -0.228392 | -0.718679 | 0.943459 | 1 |
| GOBP_BLOOD_VESSEL_ENDOTHELIAL_CELL_MIGRATION | 125 | -0.224766 | -0.718526 | 0.947883 | 1 |
| GOBP_POSITIVE_REGULATION_OF_INTERLEUKIN_8_PRODUCTION | 53 | -0.250935 | -0.718526 | 0.878897 | 1 |
| GOBP_CERAMIDE_CATABOLIC_PROCESS | 20 | -0.292108 | -0.718506 | 0.835526 | 1 |
| GOBP_INDOLE_CONTAINING_COMPOUND_METABOLIC_PROCESS | 22 | -0.290759 | -0.717748 | 0.8531 | 1 |
| GOBP_REGULATION_OF_MUSCLE_SYSTEM_PROCESS | 216 | -0.217111 | -0.717615 | 0.974066 | 1 |
| GOBP_PROTEIN_POLYMERIZATION | 291 | -0.21168 | -0.717585 | 0.977528 | 1 |
| GOBP_GLYCOPROTEIN_CATABOLIC_PROCESS | 25 | -0.284118 | -0.717563 | 0.850544 | 1 |
| GOBP_VIRAL_RNA_GENOME_REPLICATION | 35 | -0.267943 | -0.717273 | 0.87284 | 1 |
| GOBP_REGULATION_OF_CELL_DEVELOPMENT | 459 | -0.21021 | -0.716916 | 0.986948 | 1 |
| GOBP_REGULATION_OF_MAP_KINASE_ACTIVITY | 298 | -0.213988 | -0.716574 | 0.982653 | 1 |
| GOBP_RESPONSE_TO_INTERLEUKIN_1 | 194 | -0.218468 | -0.716489 | 0.969886 | 1 |
| GOBP_REGULATION_OF_CENTRIOLE_REPLICATION | 21 | -0.294335 | -0.716324 | 0.866481 | 1 |
| GOBP_MEMBRANE_LIPID_BIOSYNTHETIC_PROCESS | 145 | -0.223008 | -0.716251 | 0.956429 | 1 |
| GOBP_ENDOTHELIAL_CELL_DEVELOPMENT | 66 | -0.24567 | -0.715884 | 0.913652 | 1 |
| GOBP_POSITIVE_REGULATION_OF_MUSCLE_CELL_APOPTOTIC_PROCESS | 23 | -0.285565 | -0.715582 | 0.831341 | 1 |
| GOBP_REGULATION_OF_OXIDATIVE_STRESS_INDUCED_NEURON_DEATH | 22 | -0.286333 | -0.715386 | 0.854111 | 1 |
| GOBP_REGULATION_OF_VASOCONSTRICTION | 56 | -0.245354 | -0.715323 | 0.902469 | 1 |
| GOBP_NUCLEOSIDE_PHOSPHATE_BIOSYNTHETIC_PROCESS | 256 | -0.213639 | -0.71523 | 0.97205 | 1 |
| GOBP_NEGATIVE_REGULATION_OF_INTRACELLULAR_STEROID_HORMONE_RI | 35 | -0.257728 | -0.715217 | 0.86352 | 1 |
| GOBP_MITOCHONDRIAL_RNA_PROCESSING | 17 | -0.303382 | -0.714959 | 0.856946 | 1 |
| GOBP_CALCIUM_ION_IMPORT | 76 | -0.235933 | -0.714781 | 0.924743 | 1 |
| GOBP_REGULATION_OF_MYELOID_CELL_DIFFERENTIATION | 241 | -0.213795 | -0.714454 | 0.968976 | 1 |
| GOBP_PLATELET_ACTIVATION | 152 | -0.220784 | -0.714084 | 0.95279 | 1 |
| GOBP_LENS_FIBER_CELL_DIFFERENTIATION | 29 | -0.27504 | -0.713882 | 0.869565 | 1 |
| GOBP_NEGATIVE_REGULATION_OF_OSSIFICATION | 30 | -0.273685 | -0.713751 | 0.869063 | 1 |
| GOBP_POTASSIUM_ION_HOMEOSTASIS | 27 | -0.280167 | -0.713504 | 0.849807 | 1 |
| GOBP_PHOSPHATIDYLGLYCEROL_ACYL_CHAIN_REMODELING | 17 | -0.304413 | -0.71225 | 0.828929 | 1 |
| GOBP_NECROPTOTIC_PROCESS | 37 | -0.263515 | -0.711911 | 0.879012 | 1 |
| GOBP_CELLULAR_RESPONSE_TO_GAMMA_RADIATION | 27 | -0.280204 | -0.711682 | 0.859873 | 1 |
| GOBP_REGULATION_OF_GENERATION_OF_PRECURSOR_METABOLITES_AND_ | 153 | -0.221323 | -0.71166 | 0.963197 | 1 |
| GOBP_MICROTUBULE_POLYMERIZATION | 77 | -0.234762 | -0.711585 | 0.925495 | 1 |
| GOBP_CILIARY_BASAL_BODY_PLASMA_MEMBRANE_DOCKING | 95 | -0.229831 | -0.711447 | 0.924634 | 1 |
| GOBP_NEGATIVE_REGULATION_OF_OSTEOBLAST_DIFFERENTIATION | 43 | -0.259752 | -0.711314 | 0.879699 | 1 |
| GOBP_PROTEIN_PHOSPHOPANTETHEINYLATION | 301 | -0.211478 | -0.711311 | 0.987805 | 1 |
| GOBP_NUCLEOLAR_LARGE_RRNA_TRANSCRIPTION_BY_RNA_POLYMERASE_ | 17 | -0.301336 | -0.71036 | 0.840541 | 1 |
| GOBP_PROTEIN_DEPHOSPHORYLATION | 315 | -0.21026 | -0.710353 | 0.984772 | 1 |
| GOBP_G_PROTEIN_COUPLED_RECEPTOR_INTERNALIZATION | 15 | -0.306271 | -0.709634 | 0.85266 | 1 |
| GOBP_MRNA_TRANSPORT | 148 | -0.223117 | -0.709543 | 0.965812 | 1 |
| GOBP_REGULATION_OF_MUSCLE_CELL_DIFFERENTIATION | 133 | -0.221124 | -0.709427 | 0.951087 | 1 |
| GOBP_RESPONSE_TO_GROWTH_HORMONE | 34 | -0.267751 | -0.709334 | 0.891277 | 1 |
| GOBP_MYELOID_LEUKOCYTE_DIFFERENTIATION | 195 | -0.216388 | -0.709268 | 0.971757 | 1 |
| GOBP_PHAGOSOME_MATURATION | 45 | -0.251137 | -0.709245 | 0.901113 | 1 |
| GOBP_REGULATION_OF_POLYSACCHARIDE_METABOLIC_PROCESS | 42 | -0.251694 | -0.709148 | 0.901496 | 1 |
| GOBP_REGULATION_OF_MITOCHONDRIAL_MEMBRANE_PERMEABILITY | 70 | -0.238074 | -0.708998 | 0.915332 | 1 |
| GOBP_KERATAN_SULFATE_METABOLIC_PROCESS | 33 | -0.265951 | -0.708853 | 0.868812 | 1 |
| GOBP_CHOLESTEROL_STORAGE | 18 | -0.301466 | -0.708651 | 0.848361 | 1 |
| GOBP_PEPTIDYL_TYROSINE_AUTOPHOSPHORYLATION | 21 | -0.292346 | -0.708485 | 0.864247 | 1 |
| GOBP_SMALL_MOLECULE_CATABOLIC_PROCESS | 417 | -0.208524 | -0.707938 | 0.995964 | 1 |
| GOBP_CELLULAR_RESPONSE_TO_INSULIN_STIMULUS | 212 | -0.215991 | -0.707291 | 0.976141 | 1 |
| GOBP_REPRODUCTIVE_BEHAVIOR | 25 | -0.275804 | -0.706899 | 0.858258 | 1 |
| GOBP_REGULATION_OF_ER_TO_GOLGI_VESICLE_MEDIATED_TRANSPORT | 15 | -0.312291 | -0.706741 | 0.8331 | 1 |
| GOBP_POSITIVE_REGULATION_OF_PROTEIN_DEPOLYMERIZATION | 19 | -0.299881 | -0.706309 | 0.864901 | 1 |
| GOBP_OSTEOBLAST_DIFFERENTIATION | 207 | -0.212559 | -0.706229 | 0.974093 | 1 |
| GOBP_POSITIVE_REGULATION_OF_VASCULATURE_DEVELOPMENT | 150 | -0.218281 | -0.706228 | 0.951974 | 1 |
| GOBP_LUNG_MORPHOGENESIS | 47 | -0.24902 | -0.705799 | 0.900606 | 1 |
| GOBP_ENDOPLASMIC_RETICULUM_TUBULAR_NETWORK_ORGANIZATION | 18 | -0.298289 | -0.705111 | 0.859155 | 1 |
| GOBP_REGULATION_OF_CYCLIN_DEPENDENT_PROTEIN_KINASE_ACTIVITY | 101 | -0.226423 | -0.704855 | 0.946488 | 1 |
| GOBP_NEURONAL_STEM_CELL_POPULATION_MAINTENANCE | 19 | -0.29066 | -0.704587 | 0.843243 | 1 |
| GOBP_POSITIVE_REGULATION_OF_G1_S_TRANSITION_OF_MITOTIC_CELL_CYC | 35 | -0.258256 | -0.704093 | 0.881748 | 1 |
| GOBP_NEGATIVE_REGULATION_OF_HEMOPOIESIS | 98 | -0.229423 | -0.703923 | 0.923248 | 1 |
| GOBP_MACROMOLECULE_METHYLATION | 291 | -0.210316 | -0.70384 | 0.987743 | 1 |
| GOBP_POSITIVE_REGULATION_OF_PROTEIN_CONTAINING_COMPLEX_DISASSI | 35 | -0.260821 | -0.703658 | 0.900498 | 1 |

| | | | | | |
|---|---|---|---|---|---|
| GOBP_REGULATION_OF_TRANSLATIONAL_INITIATION_IN_RESPONSE_TO_STR | 15 | -0.303806 | -0.703423 | 0.85035 | 1 |
| GOBP_POSITIVE_REGULATION_OF_NUCLEOTIDE_METABOLIC_PROCESS | 40 | -0.258571 | -0.702801 | 0.882716 | 1 |
| GOBP_CELLULAR_AMINO_ACID_CATABOLIC_PROCESS | 103 | -0.224397 | -0.702144 | 0.962096 | 1 |
| GOBP_DETECTION_OF_BIOTIC_STIMULUS | 34 | -0.258954 | -0.701527 | 0.86802 | 1 |
| GOBP_POSITIVE_REGULATION_OF_REGULATORY_T_CELL_DIFFERENTIATION | 16 | -0.306228 | -0.701322 | 0.848066 | 1 |
| GOBP_VASCULAR_ASSOCIATED_SMOOTH_MUSCLE_CONTRACTION | 23 | -0.278571 | -0.701247 | 0.879945 | 1 |
| GOBP_NUCLEOTIDE_PHOSPHORYLATION | 129 | -0.219361 | -0.700455 | 0.962285 | 1 |
| GOBP_BONE_DEVELOPMENT | 186 | -0.212084 | -0.700449 | 0.978947 | 1 |
| GOBP_REGULATION_OF_PHOSPHOLIPID_METABOLIC_PROCESS | 34 | -0.264978 | -0.699983 | 0.890196 | 1 |
| GOBP_NUCLEOTIDE_TRANSPORT | 31 | -0.269452 | -0.699811 | 0.909934 | 1 |
| GOBP_PEPTIDYL_CYSTEINE_MODIFICATION | 46 | -0.249187 | -0.6995 | 0.90036 | 1 |
| GOBP_DEVELOPMENT_OF_PRIMARY_SEXUAL_CHARACTERISTICS | 197 | -0.21547 | -0.699233 | 0.975967 | 1 |
| GOBP_REGULATION_OF_CELL_CYCLE_PHASE_TRANSITION | 436 | -0.203934 | -0.699214 | 0.992972 | 1 |
| GOBP_EMBRYONIC_SKELETAL_SYSTEM_MORPHOGENESIS | 89 | -0.227001 | -0.69844 | 0.951057 | 1 |
| GOBP_REGULATION_OF_WATER_LOSS_VIA_SKIN | 25 | -0.275665 | -0.698196 | 0.886719 | 1 |
| GOBP_REGULATION_OF_CYTOKINESIS | 82 | -0.226913 | -0.698109 | 0.942726 | 1 |
| GOBP_MESENCHYME_MORPHOGENESIS | 49 | -0.246413 | -0.698 | 0.921951 | 1 |
| GOBP_POSITIVE_REGULATION_OF_REACTIVE_OXYGEN_SPECIES_BIOSYNTHE | 50 | -0.24617 | -0.697215 | 0.92561 | 1 |
| GOBP_NEGATIVE_REGULATION_OF_PEPTIDYL_THREONINE_PHOSPHORYLATI | 19 | -0.288436 | -0.696143 | 0.875173 | 1 |
| GOBP_TRANSCRIPTION_INITIATION_FROM_RNA_POLYMERASE_I_PROMOTER | 38 | -0.25282 | -0.695306 | 0.900517 | 1 |
| GOBP_POSITIVE_REGULATION_OF_PROTEIN_SERINE_THREONINE_KINASE_AC | 305 | -0.205312 | -0.694735 | 0.990872 | 1 |
| GOBP_VOCALIZATION_BEHAVIOR | 17 | -0.290462 | -0.694727 | 0.844729 | 1 |
| GOBP_MICROTUBULE_POLYMERIZATION_OR_DEPOLYMERIZATION | 113 | -0.218432 | -0.694715 | 0.956763 | 1 |
| GOBP_RIBONUCLEOSIDE_METABOLIC_PROCESS | 70 | -0.232673 | -0.69418 | 0.936563 | 1 |
| GOBP_REGULATION_OF_CELLULAR_COMPONENT_SIZE | 364 | -0.205865 | -0.694164 | 0.995964 | 1 |
| GOBP_PROTEIN_AUTOPHOSPHORYLATION | 223 | -0.209432 | -0.694111 | 0.985477 | 1 |
| GOBP_ALTERNATIVE_MRNA_SPLICING_VIA_SPLICEOSOME | 69 | -0.231249 | -0.693657 | 0.941793 | 1 |
| GOBP_NEGATIVE_REGULATION_OF_KINASE_ACTIVITY | 252 | -0.208817 | -0.693646 | 0.985552 | 1 |
| GOBP_RIBOSOME_BIOGENESIS | 306 | -0.204975 | -0.692655 | 0.994893 | 1 |
| GOBP_NEGATIVE_REGULATION_OF_TELOMERE_MAINTENANCE_VIA_TELOMER | 27 | -0.271941 | -0.692488 | 0.889785 | 1 |
| GOBP_CELLULAR_RESPONSE_TO_LOW_DENSITY_LIPOPROTEIN_PARTICLE_S | 17 | -0.289316 | -0.692262 | 0.868852 | 1 |
| GOBP_GLUTATHIONE_METABOLIC_PROCESS | 59 | -0.235709 | -0.692105 | 0.926683 | 1 |
| GOBP_HOMEOSTASIS_OF_NUMBER_OF_CELLS | 244 | -0.207347 | -0.692038 | 0.983454 | 1 |
| GOBP_MUSCLE_FIBER_DEVELOPMENT | 50 | -0.245174 | -0.691996 | 0.916868 | 1 |
| GOBP_POSITIVE_REGULATION_OF_PROTEIN_CONTAINING_COMPLEX_ASSEMI | 253 | -0.208091 | -0.691961 | 0.989648 | 1 |
| GOBP_REGULATION_OF_NEURON_PROJECTION_DEVELOPMENT | 417 | -0.202287 | -0.691183 | 0.996985 | 1 |
| GOBP_CELLULAR_EXTRAVASATION | 69 | -0.232173 | -0.690748 | 0.929642 | 1 |
| GOBP_REGULATION_OF_DOUBLE_STRAND_BREAK_REPAIR_VIA_HOMOLOGOU | 47 | -0.24449 | -0.690682 | 0.898428 | 1 |
| GOBP_TRANSFERRIN_TRANSPORT | 35 | -0.256339 | -0.690331 | 0.888748 | 1 |
| GOBP_RESPONSE_TO_CORTICOSTEROID | 141 | -0.218228 | -0.690289 | 0.963243 | 1 |
| GOBP_POSITIVE_REGULATION_OF_BINDING | 164 | -0.212843 | -0.690171 | 0.966209 | 1 |
| GOBP_NEGATIVE_REGULATION_OF_PROGRAMMED_NECROTIC_CELL_DEATH | 15 | -0.302085 | -0.690011 | 0.841611 | 1 |
| GOBP_PYRUVATE_METABOLIC_PROCESS | 145 | -0.214369 | -0.690003 | 0.966986 | 1 |
| GOBP_STRIATED_MUSCLE_CELL_DEVELOPMENT | 96 | -0.221203 | -0.689438 | 0.957014 | 1 |
| GOBP_NUCLEAR_TRANSCRIBED_MRNA_CATABOLIC_PROCESS | 206 | -0.208742 | -0.689205 | 0.985371 | 1 |
| GOBP_POSITIVE_REGULATION_OF_TRANSCRIPTION_BY_RNA_POLYMERASE_I | 23 | -0.278446 | -0.689072 | 0.870588 | 1 |
| GOBP_POSITIVE_REGULATION_OF_LAMELLIPODIUM_ASSEMBLY | 28 | -0.267365 | -0.687955 | 0.869452 | 1 |
| GOBP_POSITIVE_REGULATION_OF_GRANULOCYTE_CHEMOTAXIS | 27 | -0.263626 | -0.687932 | 0.898817 | 1 |
| GOBP_REGULATION_OF_STRIATED_MUSCLE_CELL_DIFFERENTIATION | 78 | -0.227134 | -0.686637 | 0.941312 | 1 |
| GOBP_GLYCOSYL_COMPOUND_CATABOLIC_PROCESS | 42 | -0.243057 | -0.686615 | 0.92725 | 1 |
| GOBP_REGULATION_OF_NUCLEOTIDE_METABOLIC_PROCESS | 114 | -0.220547 | -0.686481 | 0.953057 | 1 |
| GOBP_POSITIVE_REGULATION_OF_DNA_TEMPLATED_TRANSCRIPTION_ELONG | 26 | -0.26968 | -0.686448 | 0.880309 | 1 |
| GOBP_BLASTODERM_SEGMENTATION | 16 | -0.298265 | -0.686381 | 0.868493 | 1 |
| GOBP_REGULATION_OF_OSSIFICATION | 104 | -0.220907 | -0.685847 | 0.959025 | 1 |
| GOBP_TOOTH_MINERALIZATION | 22 | -0.279246 | -0.685648 | 0.869507 | 1 |
| GOBP_NEGATIVE_REGULATION_OF_DEPHOSPHORYLATION | 105 | -0.222375 | -0.685496 | 0.955172 | 1 |
| GOBP_RRNA_CONTAINING_RIBONUCLEOPROTEIN_COMPLEX_EXPORT_FROM | 15 | -0.301259 | -0.685313 | 0.859116 | 1 |
| GOBP_GRANULOCYTE_DIFFERENTIATION | 32 | -0.257872 | -0.685181 | 0.882353 | 1 |
| GOBP_GLIAL_CELL_PROLIFERATION | 44 | -0.24429 | -0.685043 | 0.918757 | 1 |
| GOBP_DEOXYRIBONUCLEOTIDE_METABOLIC_PROCESS | 39 | -0.249679 | -0.684788 | 0.88806 | 1 |
| GOBP_POSITIVE_REGULATION_OF_CELL_CYCLE_G1_S_PHASE_TRANSITION | 44 | -0.247434 | -0.684571 | 0.927044 | 1 |
| GOBP_NEGATIVE_REGULATION_OF_LYMPHOCYTE_MIGRATION | 15 | -0.293585 | -0.684571 | 0.870317 | 1 |
| GOBP_SEGMENTATION | 88 | -0.227953 | -0.684506 | 0.952055 | 1 |
| GOBP_RRNA_CATABOLIC_PROCESS | 20 | -0.283363 | -0.684392 | 0.873469 | 1 |
| GOBP_REGULATION_OF_RESPONSE_TO_INTERFERON_GAMMA | 24 | -0.276103 | -0.68434 | 0.890339 | 1 |
| GOBP_REGULATION_OF_AMYLOID_PRECURSOR_PROTEIN_CATABOLIC_PROC | 42 | -0.24767 | -0.684166 | 0.921308 | 1 |
| GOBP_NEUROTROPHIN_SIGNALING_PATHWAY | 38 | -0.252959 | -0.684008 | 0.922403 | 1 |
| GOBP_ENDOCARDIAL_CUSHION_MORPHOGENESIS | 33 | -0.257258 | -0.683897 | 0.913317 | 1 |
| GOBP_HYALURONAN_METABOLIC_PROCESS | 38 | -0.252617 | -0.683832 | 0.891927 | 1 |
| GOBP_N_TERMINAL_PROTEIN_AMINO_ACID_MODIFICATION | 29 | -0.265069 | -0.68345 | 0.896154 | 1 |
| GOBP_MORPHOGENESIS_OF_EMBRYONIC_EPITHELIUM | 138 | -0.212969 | -0.682991 | 0.970842 | 1 |
| GOBP_CHONDROCYTE_DIFFERENTIATION | 100 | -0.221947 | -0.682892 | 0.969663 | 1 |
| GOBP_CELLULAR_SENESCENCE | 67 | -0.231688 | -0.682482 | 0.936842 | 1 |
| GOBP_GLYCOSYL_COMPOUND_BIOSYNTHETIC_PROCESS | 42 | -0.248615 | -0.682123 | 0.92734 | 1 |
| GOBP_POSITIVE_REGULATION_OF_VIRAL_GENOME_REPLICATION | 31 | -0.254221 | -0.682107 | 0.89857 | 1 |
| GOBP_CELL_CELL_JUNCTION_ORGANIZATION | 200 | -0.206323 | -0.681926 | 0.979232 | 1 |
| GOBP_POSITIVE_REGULATION_OF_INTERFERON_ALPHA_PRODUCTION | 22 | -0.2771 | -0.681561 | 0.895861 | 1 |
| GOBP_STRIATED_MUSCLE_ADAPTATION | 41 | -0.246767 | -0.681356 | 0.913097 | 1 |
| GOBP_REGULATION_OF_TELOMERE_CAPPING | 26 | -0.267518 | -0.681043 | 0.89857 | 1 |
| GOBP_MICROTUBULE_ORGANIZING_CENTER_ORGANIZATION | 139 | -0.212942 | -0.681013 | 0.97637 | 1 |
| GOBP_RESPONSE_TO_ESTROGEN | 68 | -0.228268 | -0.680751 | 0.948395 | 1 |
| GOBP_APPENDAGE_MORPHOGENESIS | 130 | -0.212998 | -0.680738 | 0.972163 | 1 |
| GOBP_REGULATION_OF_CARDIAC_MUSCLE_CELL_ACTION_POTENTIAL | 26 | -0.26925 | -0.680505 | 0.898374 | 1 |
| GOBP_SYNAPTONEMAL_COMPLEX_ORGANIZATION | 22 | -0.272792 | -0.680051 | 0.9042 | 1 |
| GOBP_POSITIVE_REGULATION_OF_GLYCOLYTIC_PROCESS | 22 | -0.272203 | -0.680005 | 0.894665 | 1 |
| GOBP_REGULATION_OF_PHOSPHATIDYLINOSITOL_3_KINASE_ACTIVITY | 55 | -0.234548 | -0.679844 | 0.937577 | 1 |
| GOBP_NEGATIVE_REGULATION_OF_PROTEIN_BINDING | 93 | -0.220747 | -0.679715 | 0.959866 | 1 |
| GOBP_SNRNA_PROCESSING | 26 | -0.265203 | -0.679537 | 0.897001 | 1 |
| GOBP_FIBROBLAST_PROLIFERATION | 78 | -0.223387 | -0.679344 | 0.943089 | 1 |
| GOBP_HISTONE_METHYLATION | 125 | -0.213227 | -0.679123 | 0.972797 | 1 |
| GOBP_POST_EMBRYONIC_DEVELOPMENT | 74 | -0.228201 | -0.678611 | 0.940141 | 1 |
| GOBP_HOMOTYPIC_CELL_CELL_ADHESION | 85 | -0.221144 | -0.678446 | 0.960452 | 1 |
| GOBP_ENDOPLASMIC_RETICULUM_CALCIUM_ION_HOMEOSTASIS | 25 | -0.269029 | -0.678207 | 0.889774 | 1 |
| GOBP_REGULATION_OF_RECEPTOR_MEDIATED_ENDOCYTOSIS | 100 | -0.218739 | -0.677571 | 0.96829 | 1 |
| GOBP_MYELOID_CELL_DIFFERENTIATION | 399 | -0.199051 | -0.677417 | 0.994934 | 1 |
| GOBP_INNER_EAR_RECEPTOR_CELL_DEVELOPMENT | 38 | -0.248532 | -0.677288 | 0.915924 | 1 |
| GOBP_MODULATION_BY_HOST_OF_VIRAL_PROCESS | 29 | -0.261302 | -0.676787 | 0.923476 | 1 |
| GOBP_MORPHOGENESIS_OF_AN_EPITHELIAL_FOLD | 23 | -0.272259 | -0.676745 | 0.888591 | 1 |
| GOBP_REGULATION_OF_NUCLEAR_TRANSCRIBED_MRNA_POLY_A_TAIL_SHO | 15 | -0.297052 | -0.676412 | 0.858333 | 1 |
| GOBP_MICROTUBULE_NUCLEATION | 30 | -0.257248 | -0.675846 | 0.90736 | 1 |
| GOBP_POSITIVE_REGULATION_OF_MYOBLAST_DIFFERENTIATION | 17 | -0.289346 | -0.675719 | 0.882353 | 1 |
| GOBP_POSITIVE_REGULATION_OF_NEURON_PROJECTION_DEVELOPMENT | 147 | -0.209489 | -0.675621 | 0.980769 | 1 |
| GOBP_REGULATION_OF_LYSOSOMAL_LUMEN_PH | 15 | -0.296292 | -0.675337 | 0.897143 | 1 |
| GOBP_NEURON_DEATH | 329 | -0.19949 | -0.675254 | 0.993896 | 1 |
| GOBP_POSITIVE_REGULATION_OF_DOUBLE_STRAND_BREAK_REPAIR | 40 | -0.243371 | -0.675191 | 0.910013 | 1 |
| GOBP_LAMELLIPODIUM_ORGANIZATION | 85 | -0.21921 | -0.675131 | 0.964953 | 1 |
| GOBP_MITOTIC_DNA_REPLICATION | 16 | -0.288204 | -0.674352 | 0.89548 | 1 |
| GOBP_FIBROBLAST_GROWTH_FACTOR_RECEPTOR_SIGNALING_PATHWAY | 101 | -0.216433 | -0.674129 | 0.969037 | 1 |
| GOBP_SPHINGOMYELIN_METABOLIC_PROCESS | 15 | -0.297632 | -0.673731 | 0.87395 | 1 |
| GOBP_NEGATIVE_REGULATION_OF_EPITHELIAL_TO_MESENCHYMAL_TRANSIT | 37 | -0.246254 | -0.673174 | 0.909794 | 1 |
| GOBP_NEGATIVE_REGULATION_OF_CELL_SUBSTRATE_ADHESION | 59 | -0.230578 | -0.67269 | 0.948565 | 1 |
| GOBP_LOCALIZATION_WITHIN_MEMBRANE | 71 | -0.226486 | -0.672569 | 0.962543 | 1 |
| GOBP_DOSAGE_COMPENSATION | 17 | -0.285868 | -0.672507 | 0.870095 | 1 |
| GOBP_POSITIVE_REGULATION_OF_NUCLEOCYTOPLASMIC_TRANSPORT | 61 | -0.226441 | -0.672455 | 0.947561 | 1 |
| GOBP_SYNAPTIC_VESICLE_PRIMING | 19 | -0.280308 | -0.672127 | 0.887248 | 1 |
| GOBP_CARBOHYDRATE_DERIVATIVE_TRANSPORT | 79 | -0.222388 | -0.671447 | 0.958573 | 1 |
| GOBP_REGULATION_OF_INFLAMMATORY_RESPONSE_TO_ANTIGENIC_STIMUL | 28 | -0.265097 | -0.671372 | 0.911111 | 1 |
| GOBP_NEGATIVE_REGULATION_OF_CARBOHYDRATE_METABOLIC_PROCESS | 49 | -0.236898 | -0.671207 | 0.942928 | 1 |
| GOBP_NEGATIVE_REGULATION_OF_MICROTUBULE_POLYMERIZATION_OR_DE | 37 | -0.24956 | -0.67072 | 0.908755 | 1 |
| GOBP_REGULATION_OF_ATP_METABOLIC_PROCESS | 114 | -0.213174 | -0.670267 | 0.977124 | 1 |
| GOBP_MACROPHAGE_ACTIVATION | 96 | -0.216269 | -0.670053 | 0.956619 | 1 |
| GOBP_NEURAL_CREST_CELL_DIFFERENTIATION | 87 | -0.217811 | -0.669777 | 0.960586 | 1 |
| GOBP_MAMMARY_GLAND_EPITHELIUM_DEVELOPMENT | 63 | -0.22896 | -0.669592 | 0.953811 | 1 |
| GOBP_NEGATIVE_REGULATION_OF_ENDOTHELIAL_CELL_PROLIFERATION | 42 | -0.242603 | -0.66948 | 0.920954 | 1 |
| GOBP_REGULATION_OF_RIG_I_SIGNALING_PATHWAY | 18 | -0.277889 | -0.669459 | 0.903179 | 1 |
| GOBP_INTRACELLULAR_LIPID_TRANSPORT | 45 | -0.237676 | -0.668761 | 0.928571 | 1 |
| GOBP_HISTONE_H3_K4_TRIMETHYLATION | 16 | -0.288076 | -0.668702 | 0.885315 | 1 |
| GOBP_SYNAPTIC_VESICLE_TRANSPORT | 43 | -0.241596 | -0.66864 | 0.92629 | 1 |
| GOBP_OSSIFICATION | 374 | -0.198241 | -0.668595 | 0.995943 | 1 |
| GOBP_EPITHELIAL_TUBE_FORMATION | 124 | -0.21231 | -0.668455 | 0.982721 | 1 |
| GOBP_PROTEIN_LOCALIZATION_TO_ENDOSOME | 25 | -0.262149 | -0.668428 | 0.894668 | 1 |
| GOBP_REGULATION_OF_MICROTUBULE_POLYMERIZATION_OR_DEPOLYMERIZ | 81 | -0.218916 | -0.668396 | 0.953143 | 1 |
| GOBP_PURINE_NUCLEOSIDE_MONOPHOSPHATE_METABOLIC_PROCESS | 42 | -0.240048 | -0.668084 | 0.924649 | 1 |
| GOBP_REGULATION_OF_CYTOSKELETON_ORGANIZATION | 498 | -0.19448 | -0.667887 | 0.998994 | 1 |
| GOBP_REGULATION_OF_T_HELPER_17_TYPE_IMMUNE_RESPONSE | 20 | -0.275637 | -0.667832 | 0.890836 | 1 |
| GOBP_POSITIVE_REGULATION_OF_GLUCOSE_METABOLIC_PROCESS | 36 | -0.249092 | -0.667832 | 0.92711 | 1 |
| GOBP_VASCULAR_ASSOCIATED_SMOOTH_MUSCLE_CELL_MIGRATION | 22 | -0.272481 | -0.667788 | 0.895095 | 1 |
| GOBP_GLUTATHIONE_DERIVATIVE_METABOLIC_PROCESS | 21 | -0.266401 | -0.667692 | 0.90566 | 1 |

| | | | | | |
|---|---|---|---|---|---|
| GOBP_EPITHELIAL_TO_MESENCHYMAL_TRANSITION | 145 | -0.206887 | -0.667068 | 0.972133 | 1 |
| GOBP_NEGATIVE_REGULATION_OF_PROTEIN_TYROSINE_KINASE_ACTIVITY | 30 | -0.252926 | -0.666521 | 0.917408 | 1 |
| GOBP_POSITIVE_REGULATION_OF_PEPTIDASE_ACTIVITY | 191 | -0.202413 | -0.666515 | 0.983122 | 1 |
| GOBP_RIBONUCLEOSIDE_CATABOLIC_PROCESS | 23 | -0.266423 | -0.666314 | 0.91069 | 1 |
| GOBP_POSITIVE_REGULATION_OF_CYSTEINE_TYPE_ENDOPEPTIDASE_ACTIVI | 149 | -0.209092 | -0.665981 | 0.985946 | 1 |
| GOBP_CENTROSOME_SEPARATION | 15 | -0.289209 | -0.665941 | 0.886713 | 1 |
| GOBP_ASTROCYTE_DIFFERENTIATION | 69 | -0.226634 | -0.665903 | 0.951049 | 1 |
| GOBP_REGULATION_OF_LAMELLIPODIUM_ASSEMBLY | 37 | -0.244953 | -0.665585 | 0.925466 | 1 |
| GOBP_DEOXYRIBOSE_PHOSPHATE_CATABOLIC_PROCESS | 26 | -0.258513 | -0.664944 | 0.907068 | 1 |
| GOBP_NEGATIVE_REGULATION_OF_POSTTRANSCRIPTIONAL_GENE_SILENCIN | 19 | -0.274502 | -0.664743 | 0.899301 | 1 |
| GOBP_ALPHA_AMINO_ACID_CATABOLIC_PROCESS | 85 | -0.217901 | -0.664623 | 0.971461 | 1 |
| GOBP_CELLULAR_RESPONSE_TO_PEPTIDE_HORMONE_STIMULUS | 299 | -0.198207 | -0.664549 | 0.996951 | 1 |
| GOBP_TUBE_FORMATION | 139 | -0.207103 | -0.664225 | 0.981522 | 1 |
| GOBP_HISTONE_H3_ACETYLATION | 59 | -0.225964 | -0.663393 | 0.939803 | 1 |
| GOBP_HEME_METABOLIC_PROCESS | 31 | -0.251954 | -0.663191 | 0.911425 | 1 |
| GOBP_NON_MOTILE_CILIUM_ASSEMBLY | 54 | -0.231491 | -0.663055 | 0.947625 | 1 |
| GOBP_ESTABLISHMENT_OF_EPITHELIAL_CELL_POLARITY | 29 | -0.253069 | -0.663002 | 0.929487 | 1 |
| GOBP_NEGATIVE_REGULATION_OF_STEM_CELL_DIFFERENTIATION | 20 | -0.270241 | -0.662953 | 0.894231 | 1 |
| GOBP_PYRIMIDINE_NUCLEOTIDE_BIOSYNTHETIC_PROCESS | 29 | -0.253866 | -0.662368 | 0.908602 | 1 |
| GOBP_RESPONSE_TO_PEPTIDE | 483 | -0.193855 | -0.662084 | 0.998996 | 1 |
| GOBP_INSULIN_RECEPTOR_SIGNALING_PATHWAY | 135 | -0.208407 | -0.661945 | 0.976344 | 1 |
| GOBP_REGULATION_OF_CELL_ADHESION_MEDIATED_BY_INTEGRIN | 47 | -0.232854 | -0.661499 | 0.940171 | 1 |
| GOBP_NEGATIVE_REGULATION_OF_VASCULAR_PERMEABILITY | 17 | -0.282301 | -0.661213 | 0.913408 | 1 |
| GOBP_PYRIMIDINE_NUCLEOBASE_METABOLIC_PROCESS | 16 | -0.285047 | -0.660926 | 0.912256 | 1 |
| GOBP_NEGATIVE_REGULATION_OF_RESPONSE_TO_EXTERNAL_STIMULUS | 342 | -0.194659 | -0.660589 | 0.998987 | 1 |
| GOBP_ASTROCYTE_DEVELOPMENT | 39 | -0.243681 | -0.660347 | 0.934426 | 1 |
| GOBP_NEGATIVE_REGULATION_OF_INTERLEUKIN_17_PRODUCTION | 15 | -0.289684 | -0.660123 | 0.911967 | 1 |
| GOBP_APPENDAGE_DEVELOPMENT | 160 | -0.202642 | -0.660101 | 0.989177 | 1 |
| GOBP_GASTRULATION_WITH_MOUTH_FORMING_SECOND | 21 | -0.271346 | -0.659719 | 0.907008 | 1 |
| GOBP_DNA_TEMPLATED_TRANSCRIPTION_ELONGATION | 113 | -0.209595 | -0.659325 | 0.97467 | 1 |
| GOBP_NCRNA_3_END_PROCESSING | 40 | -0.238542 | -0.659197 | 0.934649 | 1 |
| GOBP_REGULATION_OF_MYELOID_CELL_APOPTOTIC_PROCESS | 21 | -0.271156 | -0.659015 | 0.891213 | 1 |
| GOBP_TRNA_MODIFICATION | 88 | -0.214646 | -0.658732 | 0.962457 | 1 |
| GOBP_POSITIVE_REGULATION_OF_PROTEIN_CATABOLIC_PROCESS | 223 | -0.197368 | -0.658549 | 0.987552 | 1 |
| GOBP_NEGATIVE_REGULATION_OF_CYTOSKELETON_ORGANIZATION | 152 | -0.203726 | -0.658351 | 0.987069 | 1 |
| GOBP_NEGATIVE_REGULATION_OF_BINDING | 153 | -0.20332 | -0.65827 | 0.986141 | 1 |
| GOBP_NEGATIVE_REGULATION_OF_NF_KAPPAB_TRANSCRIPTION_FACTOR_A | 83 | -0.214263 | -0.657543 | 0.971429 | 1 |
| GOBP_TUMOR_NECROSIS_FACTOR_MEDIATED_SIGNALING_PATHWAY | 167 | -0.200754 | -0.657298 | 0.995745 | 1 |
| GOBP_POSITIVE_REGULATION_OF_ALPHA_BETA_T_CELL_DIFFERENTIATION | 48 | -0.232741 | -0.656161 | 0.944976 | 1 |
| GOBP_REGULATION_OF_VIRAL_GENOME_REPLICATION | 81 | -0.216884 | -0.656079 | 0.962627 | 1 |
| GOBP_REGULATION_OF_POSTSYNAPSE_ORGANIZATION | 83 | -0.215539 | -0.655857 | 0.971198 | 1 |
| GOBP_REGULATION_OF_INTRACELLULAR_STEROID_HORMONE_RECEPTOR_S | 72 | -0.219375 | -0.655713 | 0.958904 | 1 |
| GOBP_CELLULAR_BIOGENIC_AMINE_METABOLIC_PROCESS | 96 | -0.214015 | -0.655164 | 0.962712 | 1 |
| GOBP_POSITIVE_REGULATION_OF_SMAD_PROTEIN_SIGNAL_TRANSDUCTION | 15 | -0.286289 | -0.654649 | 0.887446 | 1 |
| GOBP_PIGMENT_METABOLIC_PROCESS | 65 | -0.221424 | -0.654368 | 0.956725 | 1 |
| GOBP_REGULATION_OF_DNA_METABOLIC_PROCESS | 340 | -0.193284 | -0.654285 | 0.996964 | 1 |
| GOBP_STEM_CELL_PROLIFERATION | 61 | -0.223779 | -0.654265 | 0.967742 | 1 |
| GOBP_POSITIVE_REGULATION_OF_TRANSCRIPTION_OF_NOTCH_RECEPTOR_T | 18 | -0.27654 | -0.653889 | 0.899729 | 1 |
| GOBP_GLUCOSE_CATABOLIC_PROCESS | 34 | -0.245874 | -0.653873 | 0.926509 | 1 |
| GOBP_POSITIVE_REGULATION_OF_NUCLEOTIDE_BIOSYNTHETIC_PROCESS | 18 | -0.278637 | -0.653569 | 0.896 | 1 |
| GOBP_POSITIVE_REGULATION_OF_PROTEIN_TARGETING_TO_MEMBRANE | 30 | -0.250497 | -0.653436 | 0.933504 | 1 |
| GOBP_EMBRYONIC_APPENDAGE_MORPHOGENESIS | 109 | -0.208703 | -0.653325 | 0.975419 | 1 |
| GOBP_NEURON_APOPTOTIC_PROCESS | 220 | -0.197199 | -0.652862 | 0.994775 | 1 |
| GOBP_REGULATION_OF_PROTEIN_CONTAINING_COMPLEX_ASSEMBLY | 436 | -0.192124 | -0.652778 | 1 | 1 |
| GOBP_REGULATION_OF_VASCULATURE_DEVELOPMENT | 272 | -0.194384 | -0.652716 | 0.997945 | 1 |
| GOBP_MITOTIC_G2_M_TRANSITION_CHECKPOINT | 36 | -0.240542 | -0.652278 | 0.933076 | 1 |
| GOBP_PROTEIN_KINASE_C_ACTIVATING_G_PROTEIN_COUPLED_RECEPTOR_S | 27 | -0.25628 | -0.652265 | 0.913044 | 1 |
| GOBP_MICROTUBULE_CYTOSKELETON_ORGANIZATION_INVOLVED_IN_MITOSI | 142 | -0.202169 | -0.651168 | 0.987097 | 1 |
| GOBP_ACTIN_NUCLEATION | 52 | -0.227149 | -0.651003 | 0.941736 | 1 |
| GOBP_GOLGI_TO_VACUOLE_TRANSPORT | 19 | -0.26936 | -0.650895 | 0.920765 | 1 |
| GOBP_NEGATIVE_REGULATION_OF_ALPHA_BETA_T_CELL_ACTIVATION | 40 | -0.236464 | -0.650805 | 0.932656 | 1 |
| GOBP_REGULATION_OF_TELOMERE_MAINTENANCE_VIA_TELOMERE_LENGTH | 60 | -0.224174 | -0.650514 | 0.954225 | 1 |
| GOBP_MUSCLE_HYPERTROPHY_IN_RESPONSE_TO_STRESS | 20 | -0.269187 | -0.650452 | 0.926431 | 1 |
| GOBP_HYALURONAN_CATABOLIC_PROCESS | 17 | -0.273806 | -0.649794 | 0.902473 | 1 |
| GOBP_MRNA_METHYLATION | 16 | -0.280959 | -0.649629 | 0.916094 | 1 |
| GOBP_CELL_CELL_ADHESION_MEDIATED_BY_CADHERIN | 27 | -0.252405 | -0.649582 | 0.898327 | 1 |
| GOBP_POSITIVE_REGULATION_OF_I_KAPPAB_KINASE_NF_KAPPAB_SIGNALINC | 181 | -0.199087 | -0.648672 | 0.988506 | 1 |
| GOBP_REGULATION_OF_GLYCOGEN_METABOLIC_PROCESS | 34 | -0.246146 | -0.648597 | 0.939589 | 1 |
| GOBP_UROGENITAL_SYSTEM_DEVELOPMENT | 313 | -0.193857 | -0.648073 | 0.997974 | 1 |
| GOBP_REGULATION_OF_DEFENSE_RESPONSE_TO_VIRUS_BY_VIRUS | 29 | -0.24671 | -0.647478 | 0.936061 | 1 |
| GOBP_NUCLEAR_TRANSCRIBED_MRNA_CATABOLIC_PROCESS_NONSENSE_M | 120 | -0.202383 | -0.646589 | 0.980562 | 1 |
| GOBP_ADAPTATION_OF_SIGNALING_PATHWAY | 20 | -0.266202 | -0.646252 | 0.905277 | 1 |
| GOBP_NEGATIVE_REGULATION_OF_SUPRAMOLECULAR_FIBER_ORGANIZATIO | 153 | -0.200288 | -0.646234 | 0.988323 | 1 |
| GOBP_PURINE_NUCLEOSIDE_METABOLIC_PROCESS | 60 | -0.219295 | -0.646189 | 0.964158 | 1 |
| GOBP_VENTRICULAR_SEPTUM_MORPHOGENESIS | 36 | -0.241383 | -0.645572 | 0.931846 | 1 |
| GOBP_REGULATION_OF_LIPID_KINASE_ACTIVITY | 67 | -0.217503 | -0.645513 | 0.970692 | 1 |
| GOBP_SOMITE_DEVELOPMENT | 79 | -0.214746 | -0.645028 | 0.975807 | 1 |
| GOBP_REGULATION_OF_INTRACELLULAR_PROTEIN_TRANSPORT | 246 | -0.192498 | -0.644736 | 0.997963 | 1 |
| GOBP_CELL_CYCLE_G1_S_PHASE_TRANSITION | 264 | -0.191564 | -0.644701 | 1 | 1 |
| GOBP_PEPTIDYL_THREONINE_MODIFICATION | 122 | -0.204773 | -0.644637 | 0.989119 | 1 |
| GOBP_NEGATIVE_REGULATION_OF_AXON_EXTENSION | 43 | -0.230402 | -0.644497 | 0.94802 | 1 |
| GOBP_SUBSTRATE_ADHESION_DEPENDENT_CELL_SPREADING | 108 | -0.206208 | -0.644374 | 0.978912 | 1 |
| GOBP_POSITIVE_REGULATION_OF_DNA_BINDING_TRANSCRIPTION_FACTOR_A | 259 | -0.194257 | -0.644286 | 0.997957 | 1 |
| GOBP_ASYMMETRIC_CELL_DIVISION | 17 | -0.277963 | -0.643313 | 0.910364 | 1 |
| GOBP_REGULATION_OF_ESTABLISHMENT_OF_PROTEIN_LOCALIZATION_TO_M | 72 | -0.215511 | -0.643272 | 0.959724 | 1 |
| GOBP_RIBONUCLEOPROTEIN_COMPLEX_BIOGENESIS | 452 | -0.187736 | -0.643221 | 0.998993 | 1 |
| GOBP_PROTEIN_ACETYLATION | 190 | -0.195779 | -0.642766 | 0.996849 | 1 |
| GOBP_CHONDROITIN_SULFATE_PROTEOGLYCAN_METABOLIC_PROCESS | 43 | -0.230714 | -0.642723 | 0.950662 | 1 |
| GOBP_POSITIVE_REGULATION_OF_CATABOLIC_PROCESS | 438 | -0.18801 | -0.642692 | 0.997988 | 1 |
| GOBP_RESPONSE_TO_EPINEPHRINE | 15 | -0.283162 | -0.642305 | 0.902266 | 1 |
| GOBP_NEGATIVE_REGULATION_OF_G_PROTEIN_COUPLED_RECEPTOR_SIGNA | 45 | -0.232389 | -0.642245 | 0.948529 | 1 |
| GOBP_PHOSPHOLIPID_HOMEOSTASIS | 15 | -0.285365 | -0.642028 | 0.901099 | 1 |
| GOBP_MRNA_EXPORT_FROM_NUCLEUS | 111 | -0.202468 | -0.641935 | 0.982437 | 1 |
| GOBP_MIDBODY_ABSCISSION | 17 | -0.269248 | -0.641855 | 0.917007 | 1 |
| GOBP_POSITIVE_REGULATION_OF_STEROL_TRANSPORT | 30 | -0.243192 | -0.641534 | 0.939314 | 1 |
| GOBP_REGULATION_OF_BINDING | 346 | -0.189911 | -0.641352 | 1 | 1 |
| GOBP_REGULATION_OF_TRANSLATIONAL_ELONGATION | 19 | -0.272944 | -0.641037 | 0.924242 | 1 |
| GOBP_MITOCHONDRION_MORPHOGENESIS | 19 | -0.263636 | -0.640591 | 0.913397 | 1 |
| GOBP_REGULATION_OF_PROTEIN_SERINE_THREONINE_KINASE_ACTIVITY | 474 | -0.187131 | -0.639516 | 0.998994 | 1 |
| GOBP_PROTEIN_LOCALIZATION_TO_ENDOPLASMIC_RETICULUM | 147 | -0.199909 | -0.639452 | 0.993631 | 1 |
| GOBP_SODIUM_ION_IMPORT_ACROSS_PLASMA_MEMBRANE | 19 | -0.262021 | -0.639247 | 0.913462 | 1 |
| GOBP_HETEROTYPIC_CELL_CELL_ADHESION | 60 | -0.219047 | -0.638864 | 0.976359 | 1 |
| GOBP_REGULATION_OF_OSTEOBLAST_DIFFERENTIATION | 115 | -0.202573 | -0.638745 | 0.986871 | 1 |
| GOBP_BRANCHED_CHAIN_AMINO_ACID_METABOLIC_PROCESS | 23 | -0.257086 | -0.638667 | 0.920107 | 1 |
| GOBP_SMOOTH_MUSCLE_TISSUE_DEVELOPMENT | 19 | -0.265077 | -0.637945 | 0.909825 | 1 |
| GOBP_PLASMA_MEMBRANE_ORGANIZATION | 111 | -0.203249 | -0.637899 | 0.981441 | 1 |
| GOBP_MATURATION_OF_SSU_RRNA | 50 | -0.224145 | -0.63573 | 0.961298 | 1 |
| GOBP_NEGATIVE_REGULATION_OF_DNA_BINDING | 51 | -0.225097 | -0.635689 | 0.958184 | 1 |
| GOBP_REGULATION_OF_VIRAL_INDUCED_CYTOPLASMIC_PATTERN_RECOGN | 21 | -0.263088 | -0.635632 | 0.909814 | 1 |
| GOBP_RESPONSE_TO_PROSTAGLANDIN | 30 | -0.243423 | -0.635414 | 0.928205 | 1 |
| GOBP_REGULATION_OF_RUFFLE_ASSEMBLY | 28 | -0.241948 | -0.635385 | 0.928757 | 1 |
| GOBP_NEGATIVE_REGULATION_OF_RESPONSE_TO_CYTOKINE_STIMULUS | 63 | -0.215198 | -0.635176 | 0.972813 | 1 |
| GOBP_INTERFERON_BETA_PRODUCTION | 51 | -0.22003 | -0.635022 | 0.959707 | 1 |
| GOBP_PROTEIN_N_LINKED_GLYCOSYLATION | 73 | -0.210666 | -0.634906 | 0.980256 | 1 |
| GOBP_VASCULAR_ENDOTHELIAL_GROWTH_FACTOR_SIGNALING_PATHWAY | 41 | -0.232981 | -0.634864 | 0.955 | 1 |
| GOBP_CELL_JUNCTION_ASSEMBLY | 407 | -0.18568 | -0.634581 | 1 | 1 |
| GOBP_CELL_GROWTH | 447 | -0.185281 | -0.634497 | 1 | 1 |
| GOBP_REGULATION_OF_PROTEIN_IMPORT | 59 | -0.220286 | -0.63438 | 0.975089 | 1 |
| GOBP_REGULATION_OF_CALCIUM_ION_TRANSMEMBRANE_TRANSPORTER_A | 79 | -0.208901 | -0.634153 | 0.973744 | 1 |
| GOBP_CELLULAR_RESPONSE_TO_CORTICOSTEROID_STIMULUS | 56 | -0.220353 | -0.633974 | 0.968331 | 1 |
| GOBP_MRNA_CLEAVAGE | 22 | -0.257849 | -0.633839 | 0.932524 | 1 |
| GOBP_CONNECTIVE_TISSUE_DEVELOPMENT | 225 | -0.193449 | -0.633338 | 0.995855 | 1 |
| GOBP_NUCLEOTIDE_EXCISION_REPAIR_DNA_GAP_FILLING | 23 | -0.254109 | -0.633187 | 0.926121 | 1 |
| GOBP_INOSITOL_PHOSPHATE_CATABOLIC_PROCESS | 18 | -0.268061 | -0.632549 | 0.921271 | 1 |
| GOBP_MITOTIC_CYTOKINETIC_PROCESS | 23 | -0.25677 | -0.632383 | 0.935696 | 1 |
| GOBP_PHOTOPERIODISM | 31 | -0.236922 | -0.632043 | 0.949153 | 1 |
| GOBP_TRNA_METHYLATION | 40 | -0.228193 | -0.631514 | 0.963338 | 1 |
| GOBP_RIBONUCLEOSIDE_MONOPHOSPHATE_BIOSYNTHETIC_PROCESS | 33 | -0.235543 | -0.631358 | 0.942602 | 1 |
| GOBP_NEGATIVE_REGULATION_OF_TRANSMEMBRANE_TRANSPORT | 107 | -0.202877 | -0.630497 | 0.987778 | 1 |
| GOBP_LENS_DEVELOPMENT_IN_CAMERA_TYPE_EYE | 69 | -0.2116 | -0.629805 | 0.971231 | 1 |
| GOBP_HEPATICOBILIARY_SYSTEM_DEVELOPMENT | 134 | -0.197435 | -0.629443 | 0.988108 | 1 |
| GOBP_GUANOSINE_CONTAINING_COMPOUND_METABOLIC_PROCESS | 40 | -0.2307 | -0.628511 | 0.963795 | 1 |
| GOBP_AMINE_METABOLIC_PROCESS | 152 | -0.194251 | -0.628235 | 0.996746 | 1 |
| GOBP_T_HELPER_1_TYPE_IMMUNE_RESPONSE | 40 | -0.227174 | -0.627629 | 0.955696 | 1 |
| GOBP_CYTOPLASMIC_PATTERN_RECOGNITION_RECEPTOR_SIGNALING_PATH | 70 | -0.211308 | -0.627269 | 0.974148 | 1 |
| GOBP_REGULATION_OF_SUPRAMOLECULAR_FIBER_ORGANIZATION | 359 | -0.185717 | -0.626119 | 0.998989 | 1 |
| GOBP_NEGATIVE_REGULATION_OF_PEPTIDYL_TYROSINE_PHOSPHORYLATION | 54 | -0.220345 | -0.625994 | 0.977968 | 1 |
| GOBP_FATTY_ACID_CATABOLIC_PROCESS | 105 | -0.201001 | -0.625947 | 0.988814 | 1 |

| | | | | | |
|---|---|---|---|---|---|
| GOBP_VISUAL_BEHAVIOR | 45 | -0.219897 | -0.625664 | 0.972521 | 1 |
| GOBP_CELL_AGING | 102 | -0.201655 | -0.624634 | 0.991121 | 1 |
| GOBP_NUCLEOBASE_CONTAINING_COMPOUND_TRANSPORT | 244 | -0.188329 | -0.624555 | 0.99691 | 1 |
| GOBP_MELANOCYTE_DIFFERENTIATION | 22 | -0.252156 | -0.622712 | 0.924966 | 1 |
| GOBP_PROTEASOMAL_UBIQUITIN_INDEPENDENT_PROTEIN_CATABOLIC_PROC | 21 | -0.250007 | -0.622445 | 0.928475 | 1 |
| GOBP_NCRNA_METABOLIC_PROCESS | 476 | -0.183129 | -0.622312 | 1 | 1 |
| GOBP_CYTOKINETIC_PROCESS | 38 | -0.228289 | -0.622265 | 0.961202 | 1 |
| GOBP_NIK_NF_KAPPAB_SIGNALING | 170 | -0.191095 | -0.621593 | 0.998938 | 1 |
| GOBP_RIG_I_SIGNALING_PATHWAY | 24 | -0.248102 | -0.621538 | 0.933244 | 1 |
| GOBP_ACTIVATION_OF_NF_KAPPAB_INDUCING_KINASE_ACTIVITY | 18 | -0.259641 | -0.620977 | 0.924451 | 1 |
| GOBP_POSITIVE_REGULATION_OF_LAMELLIPODIUM_ORGANIZATION | 36 | -0.230213 | -0.620974 | 0.957973 | 1 |
| GOBP_REGULATION_OF_KETONE_BIOSYNTHETIC_PROCESS | 17 | -0.26132 | -0.620836 | 0.940217 | 1 |
| GOBP_POSITIVE_REGULATION_OF_ATP_METABOLIC_PROCESS | 39 | -0.228543 | -0.620363 | 0.95125 | 1 |
| GOBP_REGULATION_OF_SIGNAL_TRANSDUCTION_BY_P53_CLASS_MEDIATOR | 175 | -0.188366 | -0.620253 | 0.99791 | 1 |
| GOBP_NONRIBOSOMAL_PEPTIDE_BIOSYNTHETIC_PROCESS | 19 | -0.257353 | -0.619715 | 0.918508 | 1 |
| GOBP_MUSCLE_ADAPTATION | 100 | -0.19918 | -0.618996 | 0.986364 | 1 |
| GOBP_POSITIVE_REGULATION_OF_INTRINSIC_APOPTOTIC_SIGNALING_PATHW | 58 | -0.214885 | -0.618332 | 0.981971 | 1 |
| GOBP_POSITIVE_REGULATION_OF_AXONOGENESIS | 76 | -0.204898 | -0.618328 | 0.978434 | 1 |
| GOBP_REGULATION_OF_TELOMERE_MAINTENANCE | 79 | -0.203688 | -0.618131 | 0.980942 | 1 |
| GOBP_POSITIVE_REGULATION_OF_DNA_REPAIR | 73 | -0.207679 | -0.618002 | 0.982896 | 1 |
| GOBP_MITOTIC_CELL_CYCLE_CHECKPOINT | 156 | -0.192738 | -0.617972 | 0.992553 | 1 |
| GOBP_POSITIVE_REGULATION_OF_TYPE_I_INTERFERON_PRODUCTION | 77 | -0.203459 | -0.617459 | 0.982877 | 1 |
| GOBP_HISTONE_H2A_ACETYLATION | 17 | -0.264194 | -0.617336 | 0.945873 | 1 |
| GOBP_PRODUCTION_OF_MOLECULAR_MEDIATOR_INVOLVED_IN_INFLAMMATO | 67 | -0.207855 | -0.616345 | 0.980163 | 1 |
| GOBP_PROTEIN_IMPORT_INTO_MITOCHONDRIAL_MATRIX | 20 | -0.252435 | -0.615829 | 0.925676 | 1 |
| GOBP_NEGATIVE_REGULATION_OF_SINGLE_STRANDED_VIRAL_RNA_REPLICA | 16 | -0.26852 | -0.614646 | 0.929461 | 1 |
| GOBP_PEPTIDYL_ASPARAGINE_MODIFICATION | 31 | -0.232841 | -0.614615 | 0.951469 | 1 |
| GOBP_REGULATION_OF_ATPASE_ACTIVITY | 79 | -0.204571 | -0.614548 | 0.982778 | 1 |
| GOBP_HEMOGLOBIN_METABOLIC_PROCESS | 15 | -0.266894 | -0.614122 | 0.918733 | 1 |
| GOBP_POSITIVE_REGULATION_OF_PROTEIN_LOCALIZATION_TO_MEMBRANE | 125 | -0.192583 | -0.61405 | 0.995699 | 1 |
| GOBP_POSITIVE_REGULATION_OF_TELOMERASE_RNA_LOCALIZATION_TO_CA | 15 | -0.265351 | -0.613729 | 0.934174 | 1 |
| GOBP_ACTIN_FILAMENT_BUNDLE_ORGANIZATION | 153 | -0.18798 | -0.613426 | 0.994664 | 1 |
| GOBP_NEGATIVE_REGULATION_OF_UBIQUITIN_PROTEIN_TRANSFERASE_ACT | 16 | -0.267062 | -0.612996 | 0.932203 | 1 |
| GOBP_NEGATIVE_REGULATION_OF_TRANSFORMING_GROWTH_FACTOR_BETA | 78 | -0.201568 | -0.612257 | 0.987165 | 1 |
| GOBP_DICARBOXYLIC_ACID_METABOLIC_PROCESS | 92 | -0.19845 | -0.612153 | 0.993341 | 1 |
| GOBP_PEPTIDYL_LYSINE_MODIFICATION | 376 | -0.179657 | -0.611529 | 1 | 1 |
| GOBP_ACTIN_FILAMENT_ORGANIZATION | 424 | -0.180174 | -0.611362 | 1 | 1 |
| GOBP_REGULATION_OF_HAIR_FOLLICLE_DEVELOPMENT | 17 | -0.261958 | -0.611321 | 0.92 | 1 |
| GOBP_GLUTAMINE_FAMILY_AMINO_ACID_METABOLIC_PROCESS | 71 | -0.204818 | -0.611267 | 0.978749 | 1 |
| GOBP_SKELETAL_MUSCLE_ADAPTATION | 23 | -0.250465 | -0.6107 | 0.950993 | 1 |
| GOBP_ORGANELLE_TRANSPORT_ALONG_MICROTUBULE | 81 | -0.200022 | -0.609979 | 0.986364 | 1 |
| GOBP_ACTIN_FILAMENT_BASED_TRANSPORT | 21 | -0.243663 | -0.609692 | 0.944664 | 1 |
| GOBP_REGULATION_OF_MUSCLE_ADAPTATION | 80 | -0.200757 | -0.608183 | 0.985311 | 1 |
| GOBP_LEUKOCYTE_ADHESION_TO_VASCULAR_ENDOTHELIAL_CELL | 45 | -0.217302 | -0.607778 | 0.966061 | 1 |
| GOBP_NUCLEOBASE_BIOSYNTHETIC_PROCESS | 18 | -0.25784 | -0.60713 | 0.927273 | 1 |
| GOBP_POSITIVE_REGULATION_OF_PROTEOLYSIS | 355 | -0.178564 | -0.607049 | 1 | 1 |
| GOBP_PEROXISOMAL_TRANSPORT | 72 | -0.201476 | -0.606372 | 0.989498 | 1 |
| GOBP_GTP_METABOLIC_PROCESS | 24 | -0.244012 | -0.605757 | 0.950392 | 1 |
| GOBP_REPLICATIVE_SENESCENCE | 16 | -0.264381 | -0.605506 | 0.93892 | 1 |
| GOBP_MYELOID_CELL_APOPTOTIC_PROCESS | 26 | -0.240837 | -0.604566 | 0.948886 | 1 |
| GOBP_INTRACELLULAR_RECEPTOR_SIGNALING_PATHWAY | 261 | -0.17889 | -0.60298 | 1 | 1 |
| GOBP_REGULATION_OF_MICROTUBULE_CYTOSKELETON_ORGANIZATION | 137 | -0.187409 | -0.601991 | 0.995685 | 1 |
| GOBP_INNER_MITOCHONDRIAL_MEMBRANE_ORGANIZATION | 56 | -0.210223 | -0.601852 | 0.973621 | 1 |
| GOBP_REGULATION_OF_TRANSLATIONAL_FIDELITY | 20 | -0.246895 | -0.600854 | 0.948824 | 1 |
| GOBP_NEGATIVE_REGULATION_OF_SMALL_GTPASE_MEDIATED_SIGNAL_TRAI | 56 | -0.20891 | -0.600281 | 0.979064 | 1 |
| GOBP_REGULATION_OF_RESPONSE_TO_TUMOR_CELL | 16 | -0.257285 | -0.599859 | 0.935897 | 1 |
| GOBP_NEGATIVE_REGULATION_OF_NUCLEOTIDE_METABOLIC_PROCESS | 22 | -0.239584 | -0.59858 | 0.94709 | 1 |
| GOBP_ENDOCYTIC_RECYCLING | 50 | -0.209997 | -0.598283 | 0.977053 | 1 |
| GOBP_REGULATION_OF_COLLAGEN_METABOLIC_PROCESS | 41 | -0.216469 | -0.59775 | 0.973418 | 1 |
| GOBP_GPI_ANCHOR_METABOLIC_PROCESS | 32 | -0.223983 | -0.59757 | 0.973111 | 1 |
| GOBP_POSITIVE_REGULATION_OF_PROTEIN_LOCALIZATION_TO_NUCLEUS | 84 | -0.197142 | -0.596898 | 0.988453 | 1 |
| GOBP_REGULATION_OF_ACTIN_FILAMENT_BUNDLE_ASSEMBLY | 98 | -0.191381 | -0.595861 | 0.988764 | 1 |
| GOBP_POSITIVE_REGULATION_OF_TELOMERE_MAINTENANCE_VIA_TELOMERI | 36 | -0.220707 | -0.595833 | 0.976601 | 1 |
| GOBP_CARBOHYDRATE_CATABOLIC_PROCESS | 186 | -0.18388 | -0.595765 | 1 | 1 |
| GOBP_SCF_DEPENDENT_PROTEASOMAL_UBIQUITIN_DEPENDENT_PROTEIN_( | 91 | -0.195714 | -0.595747 | 0.991041 | 1 |
| GOBP_SPHINGOLIPID_MEDIATED_SIGNALING_PATHWAY | 15 | -0.260151 | -0.595536 | 0.938462 | 1 |
| GOBP_CELL_CYCLE_ARREST | 227 | -0.178396 | -0.595477 | 1 | 1 |
| GOBP_RESPONSE_TO_RADIATION | 428 | -0.17591 | -0.595418 | 1 | 1 |
| GOBP_MITOCHONDRIAL_MEMBRANE_ORGANIZATION | 139 | -0.185462 | -0.59456 | 0.992497 | 1 |
| GOBP_REGULATION_OF_INTRACELLULAR_TRANSPORT | 343 | -0.175551 | -0.594227 | 1 | 1 |
| GOBP_REGULATION_OF_CILIUM_ASSEMBLY | 57 | -0.204287 | -0.593926 | 0.981287 | 1 |
| GOBP_REGULATION_OF_HEMATOPOIETIC_STEM_CELL_DIFFERENTIATION | 73 | -0.200256 | -0.5933 | 0.994145 | 1 |
| GOBP_ASSOCIATIVE_LEARNING | 64 | -0.20218 | -0.592936 | 0.986858 | 1 |
| GOBP_REGULATION_OF_PLATELET_AGGREGATION | 19 | -0.253013 | -0.592883 | 0.947226 | 1 |
| GOBP_LEARNING | 125 | -0.186324 | -0.591893 | 0.992424 | 1 |
| GOBP_RHO_PROTEIN_SIGNAL_TRANSDUCTION | 131 | -0.186172 | -0.591567 | 0.99676 | 1 |
| GOBP_POSITIVE_REGULATION_OF_NON_CANONICAL_WNT_SIGNALING_PATHW | 15 | -0.259723 | -0.591437 | 0.947218 | 1 |
| GOBP_EPITHELIAL_CELL_CELL_ADHESION | 17 | -0.254368 | -0.589974 | 0.945983 | 1 |
| GOBP_POSITIVE_REGULATION_OF_BLOOD_VESSEL_ENDOTHELIAL_CELL_MIG | 62 | -0.200585 | -0.589756 | 0.985932 | 1 |
| GOBP_CELLULAR_RESPONSE_TO_PEPTIDE | 360 | -0.175326 | -0.589595 | 1 | 1 |
| GOBP_PYRIMIDINE_RIBONUCLEOSIDE_TRIPHOSPHATE_BIOSYNTHETIC_PROC | 15 | -0.25809 | -0.589211 | 0.949106 | 1 |
| GOBP_GROOMING_BEHAVIOR | 15 | -0.25989 | -0.588649 | 0.947218 | 1 |
| GOBP_PLACENTA_BLOOD_VESSEL_DEVELOPMENT | 29 | -0.226839 | -0.587989 | 0.965333 | 1 |
| GOBP_RESPONSE_TO_TRANSFORMING_GROWTH_FACTOR_BETA | 240 | -0.175665 | -0.587568 | 1 | 1 |
| GOBP_NEGATIVE_REGULATION_OF_CELL_CYCLE_PHASE_TRANSITION | 243 | -0.177421 | -0.587519 | 1 | 1 |
| GOBP_RNA_POLYMERASE_II_PREINITIATION_COMPLEX_ASSEMBLY | 23 | -0.238078 | -0.587086 | 0.951936 | 1 |
| GOBP_BLASTOCYST_GROWTH | 19 | -0.239154 | -0.586952 | 0.938028 | 1 |
| GOBP_RESPONSE_TO_OXYGEN_LEVELS | 368 | -0.172798 | -0.586695 | 1 | 1 |
| GOBP_REGULATION_OF_ENDOTHELIAL_CELL_MIGRATION | 175 | -0.18024 | -0.586686 | 0.997872 | 1 |
| GOBP_CELLULAR_RESPONSE_TO_VASCULAR_ENDOTHELIAL_GROWTH_FACT | 63 | -0.200287 | -0.586446 | 0.984634 | 1 |
| GOBP_REGULATION_OF_ACTIN_CYTOSKELETON_REORGANIZATION | 37 | -0.216908 | -0.58636 | 0.967541 | 1 |
| GOBP_POSITIVE_REGULATION_OF_CYTOKINE_PRODUCTION_INVOLVED_IN_IN | 19 | -0.24141 | -0.586051 | 0.943775 | 1 |
| GOBP_POSITIVE_REGULATION_OF_NEUTROPHIL_MIGRATION | 29 | -0.226617 | -0.585983 | 0.958495 | 1 |
| GOBP_NEGATIVE_REGULATION_OF_CATION_TRANSMEMBRANE_TRANSPORT | 78 | -0.195574 | -0.585836 | 0.990836 | 1 |
| GOBP_RESPONSE_TO_NICOTINE | 39 | -0.214208 | -0.585701 | 0.973316 | 1 |
| GOBP_REGULATION_OF_ARP2_3_COMPLEX_MEDIATED_ACTIN_NUCLEATION | 21 | -0.238678 | -0.585453 | 0.945578 | 1 |
| GOBP_NUCLEUS_ORGANIZATION | 118 | -0.18637 | -0.584809 | 0.998889 | 1 |
| GOBP_PROGRAMMED_NECROTIC_CELL_DEATH | 45 | -0.204972 | -0.584553 | 0.982346 | 1 |
| GOBP_RNA_PHOSPHODIESTER_BOND_HYDROLYSIS_EXONUCLEOLYTIC | 43 | -0.209033 | -0.584264 | 0.97764 | 1 |
| GOBP_DEPHOSPHORYLATION | 474 | -0.169456 | -0.583955 | 1 | 1 |
| GOBP_REGULATION_OF_MRNA_METABOLIC_PROCESS | 320 | -0.173882 | -0.583855 | 1 | 1 |
| GOBP_REGULATION_OF_RESPIRATORY_GASEOUS_EXCHANGE | 15 | -0.258666 | -0.583717 | 0.94528 | 1 |
| GOBP_POSITIVE_REGULATION_OF_TORC1_SIGNALING | 17 | -0.2486 | -0.583414 | 0.942701 | 1 |
| GOBP_PROTEIN_LOCALIZATION_TO_MICROTUBULE_ORGANIZING_CENTER | 35 | -0.217961 | -0.583193 | 0.972152 | 1 |
| GOBP_ROOF_OF_MOUTH_DEVELOPMENT | 84 | -0.19112 | -0.582908 | 0.993182 | 1 |
| GOBP_MEMBRANE_FISSION | 15 | -0.254797 | -0.58271 | 0.939107 | 1 |
| GOBP_SPLEEN_DEVELOPMENT | 31 | -0.222954 | -0.582524 | 0.963522 | 1 |
| GOBP_INTERLEUKIN_6_MEDIATED_SIGNALING_PATHWAY | 22 | -0.23554 | -0.581879 | 0.963109 | 1 |
| GOBP_ASTROCYTE_ACTIVATION | 22 | -0.234723 | -0.581171 | 0.955671 | 1 |
| GOBP_REGULATION_OF_CELL_SIZE | 170 | -0.180266 | -0.580955 | 1 | 1 |
| GOBP_TOXIN_TRANSPORT | 39 | -0.209294 | -0.58095 | 0.986146 | 1 |
| GOBP_REGULATION_OF_RECEPTOR_INTERNALIZATION | 51 | -0.20531 | -0.58061 | 0.978049 | 1 |
| GOBP_REGULATION_OF_CELLULAR_AMINE_METABOLIC_PROCESS | 72 | -0.194415 | -0.580544 | 0.993159 | 1 |
| GOBP_PROSTANOID_BIOSYNTHETIC_PROCESS | 27 | -0.225963 | -0.580292 | 0.969415 | 1 |
| GOBP_EXIT_FROM_MITOSIS | 26 | -0.225927 | -0.580105 | 0.958549 | 1 |
| GOBP_NEGATIVE_REGULATION_OF_PHOSPHORYLATION | 384 | -0.171778 | -0.579944 | 1 | 1 |
| GOBP_REGULATION_OF_POSTSYNAPTIC_MEMBRANE_NEUROTRANSMITTER_F | 54 | -0.203341 | -0.579192 | 0.986683 | 1 |
| GOBP_NAD_BIOSYNTHETIC_PROCESS | 28 | -0.228708 | -0.579159 | 0.974194 | 1 |
| GOBP_RESPONSE_TO_ANTIBIOTIC | 43 | -0.207759 | -0.578723 | 0.983648 | 1 |
| GOBP_POSITIVE_REGULATION_OF_HISTONE_H3_K4_METHYLATION | 16 | -0.249271 | -0.578459 | 0.951724 | 1 |
| GOBP_POSITIVE_REGULATION_OF_PEPTIDYL_LYSINE_ACETYLATION | 34 | -0.216443 | -0.578409 | 0.967949 | 1 |
| GOBP_POST_TRANSLATIONAL_PROTEIN_MODIFICATION | 347 | -0.170459 | -0.578278 | 1 | 1 |
| GOBP_REGULATION_OF_UBIQUITIN_PROTEIN_TRANSFERASE_ACTIVITY | 51 | -0.20345 | -0.578081 | 0.982989 | 1 |
| GOBP_FORMATION_OF_CYTOPLASMIC_TRANSLATION_INITIATION_COMPLEX | 16 | -0.249529 | -0.577223 | 0.952974 | 1 |
| GOBP_RNA_EXPORT_FROM_NUCLEUS | 137 | -0.179698 | -0.576491 | 1 | 1 |
| GOBP_IMPORT_INTO_NUCLEUS | 158 | -0.179395 | -0.576456 | 1 | 1 |
| GOBP_REGULATION_OF_PROTEIN_DEPOLYMERIZATION | 84 | -0.188139 | -0.576161 | 0.997704 | 1 |
| GOBP_NEGATIVE_REGULATION_OF_PLASMA_MEMBRANE_BOUNDED_CELL_P | 28 | -0.224567 | -0.575921 | 0.966887 | 1 |
| GOBP_POSITIVE_REGULATION_OF_CELL_CYCLE_ARREST | 79 | -0.189344 | -0.57585 | 0.996475 | 1 |
| GOBP_REGULATION_OF_CAMP_MEDIATED_SIGNALING | 34 | -0.216778 | -0.575662 | 0.973618 | 1 |
| GOBP_PEPTIDYL_LYSINE_ACETYLATION | 162 | -0.179977 | -0.574787 | 1 | 1 |
| GOBP_LIVER_MORPHOGENESIS | 21 | -0.2367 | -0.574716 | 0.964798 | 1 |
| GOBP_NEGATIVE_REGULATION_OF_TELOMERE_MAINTENANCE | 35 | -0.209699 | -0.573876 | 0.97416 | 1 |
| GOBP_REGULATION_OF_CELLULAR_AMINO_ACID_METABOLIC_PROCESS | 58 | -0.200072 | -0.573313 | 0.984394 | 1 |
| GOBP_REGULATION_OF_MONOOXYGENASE_ACTIVITY | 57 | -0.19548 | -0.572546 | 0.990408 | 1 |
| GOBP_MITOCHONDRIAL_TRANSPORT | 266 | -0.170176 | -0.572493 | 1 | 1 |
| GOBP_NEGATIVE_REGULATION_OF_PROTEIN_CONTAINING_COMPLEX_DISASS | 80 | -0.189348 | -0.572078 | 0.993111 | 1 |
| GOBP_AXON_EXTENSION | 111 | -0.180342 | -0.570823 | 0.998901 | 1 |

| | | | | | |
|---|---|---|---|---|---|
| GOBP_INTRACELLULAR_STEROL_TRANSPORT | 25 | -0.227056 | -0.570635 | 0.973475 | 1 |
| GOBP_NON_CANONICAL_WNT_SIGNALING_PATHWAY | 151 | -0.177591 | -0.569536 | 0.998934 | 1 |
| GOBP_VENTRICULAR_CARDIAC_MUSCLE_CELL_ACTION_POTENTIAL | 33 | -0.213381 | -0.569469 | 0.978723 | 1 |
| GOBP_REGULATION_OF_CELL_MORPHOGENESIS_INVOLVED_IN_DIFFERENTIA | 94 | -0.185615 | -0.569183 | 0.997763 | 1 |
| GOBP_CARBOHYDRATE_TRANSPORT | 138 | -0.17882 | -0.568924 | 1 | 1 |
| GOBP_POSITIVE_REGULATION_OF_CELL_MORPHOGENESIS_INVOLVED_IN_DIF | 78 | -0.188342 | -0.568894 | 0.989643 | 1 |
| GOBP_PROTEIN_DEPOLYMERIZATION | 110 | -0.180795 | -0.568833 | 0.994547 | 1 |
| GOBP_REGULATION_OF_BICELLULAR_TIGHT_JUNCTION_ASSEMBLY | 23 | -0.226997 | -0.568814 | 0.953867 | 1 |
| GOBP_I_KAPPAB_KINASE_NF_KAPPAB_SIGNALING | 275 | -0.169552 | -0.567726 | 1 | 1 |
| GOBP_CARBOHYDRATE_TRANSMEMBRANE_TRANSPORT | 105 | -0.181727 | -0.566687 | 0.997743 | 1 |
| GOBP_POSITIVE_REGULATION_OF_ESTABLISHMENT_OF_PROTEIN_LOCALIZAT | 319 | -0.168434 | -0.566511 | 1 | 1 |
| GOBP_PHAGOLYSOSOME_ASSEMBLY | 15 | -0.246848 | -0.566355 | 0.96832 | 1 |
| GOBP_NAD_METABOLIC_PROCESS | 48 | -0.201129 | -0.566099 | 0.978443 | 1 |
| GOBP_IN_UTERO_EMBRYONIC_DEVELOPMENT | 329 | -0.167339 | -0.566019 | 1 | 1 |
| GOBP_PEROXISOME_ORGANIZATION | 84 | -0.185341 | -0.565858 | 0.995439 | 1 |
| GOBP_RETROGRADE_TRANSPORT_ENDOSOME_TO_PLASMA_MEMBRANE | 31 | -0.215965 | -0.565473 | 0.975452 | 1 |
| GOBP_HEMATOPOIETIC_PROGENITOR_CELL_DIFFERENTIATION | 158 | -0.174835 | -0.565281 | 0.998937 | 1 |
| GOBP_POSITIVE_REGULATION_OF_CELLULAR_PROTEIN_LOCALIZATION | 290 | -0.167458 | -0.565247 | 1 | 1 |
| GOBP_NEGATIVE_REGULATION_OF_AXONOGENESIS | 66 | -0.189093 | -0.564612 | 0.99409 | 1 |
| GOBP_POSITIVE_REGULATION_OF_ORGANELLE_ASSEMBLY | 66 | -0.190054 | -0.563334 | 0.996471 | 1 |
| GOBP_REGULATION_OF_PROTEIN_EXIT_FROM_ENDOPLASMIC_RETICULUM | 27 | -0.219573 | -0.562796 | 0.981936 | 1 |
| GOBP_CELL_CYCLE_CHECKPOINT | 204 | -0.169931 | -0.562164 | 1 | 1 |
| GOBP_PYRIMIDINE_NUCLEOTIDE_CATABOLIC_PROCESS | 19 | -0.233444 | -0.561626 | 0.955882 | 1 |
| GOBP_REGULATION_OF_DNA_BIOSYNTHETIC_PROCESS | 106 | -0.180017 | -0.561545 | 0.996729 | 1 |
| GOBP_COVALENT_CHROMATIN_MODIFICATION | 443 | -0.164804 | -0.560506 | 1 | 1 |
| GOBP_POSITIVE_REGULATION_OF_WOUND_HEALING | 51 | -0.197271 | -0.560266 | 0.987908 | 1 |
| GOBP_TRNA_PROCESSING | 128 | -0.17619 | -0.558551 | 0.996739 | 1 |
| GOBP_POSITIVE_REGULATION_OF_PROTEIN_ACETYLATION | 43 | -0.201903 | -0.558305 | 0.986352 | 1 |
| GOBP_ESTABLISHMENT_OF_RNA_LOCALIZATION | 194 | -0.1685 | -0.557852 | 1 | 1 |
| GOBP_POSITIVE_REGULATION_OF_CELLULAR_PROTEIN_CATABOLIC_PROCES | 149 | -0.174101 | -0.557328 | 1 | 1 |
| GOBP_PIGMENT_BIOSYNTHETIC_PROCESS | 55 | -0.190081 | -0.556976 | 0.990454 | 1 |
| GOBP_REGULATION_OF_EMBRYONIC_DEVELOPMENT | 59 | -0.19259 | -0.556577 | 0.992857 | 1 |
| GOBP_REGULATION_OF_HEMATOPOIETIC_PROGENITOR_CELL_DIFFERENTIAT | 88 | -0.182603 | -0.556452 | 0.997768 | 1 |
| GOBP_NEGATIVE_REGULATION_OF_PHOSPHORUS_METABOLIC_PROCESS | 496 | -0.162821 | -0.556184 | 1 | 1 |
| GOBP_RNA_3_END_PROCESSING | 140 | -0.173466 | -0.556132 | 0.997872 | 1 |
| GOBP_REGULATION_OF_FATTY_ACID_BETA_OXIDATION | 19 | -0.232008 | -0.556112 | 0.959889 | 1 |
| GOBP_POSITIVE_REGULATION_OF_NEURON_MIGRATION | 15 | -0.246729 | -0.556111 | 0.966292 | 1 |
| GOBP_NEGATIVE_REGULATION_OF_LEUKOCYTE_CHEMOTAXIS | 18 | -0.236129 | -0.555965 | 0.971311 | 1 |
| GOBP_REGULATION_OF_CELL_SHAPE | 150 | -0.171963 | -0.555912 | 0.995763 | 1 |
| GOBP_LYMPH_VESSEL_MORPHOGENESIS | 21 | -0.226684 | -0.555468 | 0.975643 | 1 |
| GOBP_RESPONSE_TO_LIGHT_STIMULUS | 302 | -0.165774 | -0.554885 | 1 | 1 |
| GOBP_NUCLEAR_TRANSPORT | 334 | -0.163843 | -0.554728 | 1 | 1 |
| GOBP_REGULATION_OF_ACTOMYOSIN_STRUCTURE_ORGANIZATION | 93 | -0.181183 | -0.5547 | 0.997765 | 1 |
| GOBP_PURINE_NUCLEOSIDE_MONOPHOSPHATE_BIOSYNTHETIC_PROCESS | 23 | -0.222555 | -0.554639 | 0.977992 | 1 |
| GOBP_ERYTHROCYTE_DEVELOPMENT | 32 | -0.205412 | -0.553636 | 0.975703 | 1 |
| GOBP_POSITIVE_REGULATION_OF_CELL_PROJECTION_ORGANIZATION | 333 | -0.163901 | -0.552736 | 1 | 1 |
| GOBP_POSITIVE_REGULATION_OF_CYCLASE_ACTIVITY | 17 | -0.234002 | -0.552669 | 0.958791 | 1 |
| GOBP_NUCLEOSIDE_DIPHOSPHATE_METABOLIC_PROCESS | 152 | -0.171876 | -0.552178 | 1 | 1 |
| GOBP_REGULATION_OF_SUBSTRATE_ADHESION_DEPENDENT_CELL_SPREAD | 57 | -0.192143 | -0.551837 | 0.986585 | 1 |
| GOBP_MEMBRANE_DOCKING | 181 | -0.167155 | -0.550587 | 1 | 1 |
| GOBP_POSITIVE_REGULATION_OF_PROTEIN_MODIFICATION_BY_SMALL_PROT | 132 | -0.173851 | -0.549873 | 0.99891 | 1 |
| GOBP_POSITIVE_REGULATION_OF_MEMBRANE_PERMEABILITY | 64 | -0.18941 | -0.54956 | 0.990555 | 1 |
| GOBP_REGULATION_OF_LAMELLIPODIUM_ORGANIZATION | 49 | -0.193158 | -0.548887 | 0.98917 | 1 |
| GOBP_CYTOKINE_PRODUCTION_INVOLVED_IN_INFLAMMATORY_RESPONSE | 50 | -0.189997 | -0.54827 | 0.992823 | 1 |
| GOBP_POSITIVE_REGULATION_OF_CD4_POSITIVE_ALPHA_BETA_T_CELL_ACT | 37 | -0.204583 | -0.547904 | 0.98708 | 1 |
| GOBP_REGULATION_OF_HISTONE_MODIFICATION | 137 | -0.172198 | -0.547875 | 0.998928 | 1 |
| GOBP_ENDOPLASMIC_RETICULUM_TO_CYTOSOL_TRANSPORT | 28 | -0.212607 | -0.547463 | 0.988266 | 1 |
| GOBP_POLYOL_METABOLIC_PROCESS | 125 | -0.172945 | -0.547012 | 0.997852 | 1 |
| GOBP_ADIPOSE_TISSUE_DEVELOPMENT | 34 | -0.20366 | -0.546838 | 0.983139 | 1 |
| GOBP_SNRNA_METABOLIC_PROCESS | 37 | -0.20089 | -0.54661 | 0.991117 | 1 |
| GOBP_REGULATION_OF_SPINDLE_ORGANIZATION | 39 | -0.198854 | -0.546475 | 0.988736 | 1 |
| GOBP_ENDOSOME_TRANSPORT_VIA_MULTIVESICULAR_BODY_SORTING_PATI | 31 | -0.205553 | -0.54607 | 0.9825 | 1 |
| GOBP_HISTONE_MRNA_METABOLIC_PROCESS | 24 | -0.215484 | -0.545876 | 0.981699 | 1 |
| GOBP_CHONDROITIN_SULFATE_PROTEOGLYCAN_BIOSYNTHETIC_PROCESS | 30 | -0.206247 | -0.545867 | 0.988095 | 1 |
| GOBP_REGULATION_OF_CELL_GROWTH_INVOLVED_IN_CARDIAC_MUSCLE_C | 18 | -0.229586 | -0.545152 | 0.978571 | 1 |
| GOBP_TELOMERE_MAINTENANCE_VIA_TELOMERE_LENGTHENING | 79 | -0.180268 | -0.544783 | 0.995413 | 1 |
| GOBP_REGULATION_OF_SMALL_GTPASE_MEDIATED_SIGNAL_TRANSDUCTION | 315 | -0.160374 | -0.54476 | 1 | 1 |
| GOBP_REGULATION_OF_CARBOHYDRATE_BIOSYNTHETIC_PROCESS | 89 | -0.176884 | -0.544483 | 0.998865 | 1 |
| GOBP_REGULATION_OF_MITOTIC_SPINDLE_ASSEMBLY | 18 | -0.236193 | -0.544277 | 0.966942 | 1 |
| GOBP_GLUCOSE_IMPORT | 66 | -0.186161 | -0.543978 | 0.993961 | 1 |
| GOBP_CARDIAC_EPITHELIAL_TO_MESENCHYMAL_TRANSITION | 30 | -0.20568 | -0.543384 | 0.984375 | 1 |
| GOBP_NEGATIVE_REGULATION_OF_TOR_SIGNALING | 43 | -0.198371 | -0.543075 | 0.99375 | 1 |
| GOBP_PEPTIDYL_TYROSINE_DEPHOSPHORYLATION | 100 | -0.174889 | -0.542996 | 0.997735 | 1 |
| GOBP_POSITIVE_REGULATION_OF_PROTEIN_LOCALIZATION_TO_CELL_SURFA | 16 | -0.234003 | -0.542981 | 0.962963 | 1 |
| GOBP_REGULATION_OF_MITOCHONDRION_ORGANIZATION | 148 | -0.169487 | -0.542349 | 0.99893 | 1 |
| GOBP_NEURON_MATURATION | 43 | -0.194146 | -0.541284 | 0.987516 | 1 |
| GOBP_ACTOMYOSIN_STRUCTURE_ORGANIZATION | 183 | -0.164465 | -0.541007 | 1 | 1 |
| GOBP_PROTEIN_SUMOYLATION | 77 | -0.178675 | -0.540109 | 0.996536 | 1 |
| GOBP_POSITIVE_REGULATION_OF_PROTEIN_EXPORT_FROM_NUCLEUS | 20 | -0.222215 | -0.539363 | 0.972727 | 1 |
| GOBP_REGULATION_OF_VASCULAR_PERMEABILITY | 41 | -0.195787 | -0.539067 | 0.989529 | 1 |
| GOBP_NEURAL_TUBE_DEVELOPMENT | 144 | -0.167552 | -0.5382 | 0.997866 | 1 |
| GOBP_REGULATION_OF_CELL_MORPHOGENESIS | 301 | -0.16072 | -0.537353 | 1 | 1 |
| GOBP_NUCLEAR_EXPORT | 196 | -0.163213 | -0.535759 | 1 | 1 |
| GOBP_CELLULAR_RESPONSE_TO_CHEMICAL_STRESS | 327 | -0.158488 | -0.535533 | 1 | 1 |
| GOBP_POSITIVE_REGULATION_OF_DOUBLE_STRAND_BREAK_REPAIR_VIA_NO | 16 | -0.23024 | -0.535051 | 0.968706 | 1 |
| GOBP_MITOCHONDRIAL_GENE_EXPRESSION | 165 | -0.164008 | -0.535008 | 1 | 1 |
| GOBP_NEGATIVE_REGULATION_OF_CELL_CYCLE_PROCESS | 329 | -0.15712 | -0.534082 | 1 | 1 |
| GOBP_SHORT_CHAIN_FATTY_ACID_METABOLIC_PROCESS | 17 | -0.22795 | -0.533903 | 0.969655 | 1 |
| GOBP_POSITIVE_REGULATION_OF_RESPONSE_TO_DNA_DAMAGE_STIMULUS | 106 | -0.171326 | -0.533599 | 1 | 1 |
| GOBP_NECROTIC_CELL_DEATH | 57 | -0.18404 | -0.532893 | 0.994012 | 1 |
| GOBP_SMALL_GTPASE_MEDIATED_SIGNAL_TRANSDUCTION | 496 | -0.156725 | -0.532493 | 1 | 1 |
| GOBP_NEGATIVE_REGULATION_OF_CELL_JUNCTION_ASSEMBLY | 30 | -0.20484 | -0.532488 | 0.989376 | 1 |
| GOBP_POSITIVE_REGULATION_OF_TRANSLATION | 122 | -0.166635 | -0.531258 | 1 | 1 |
| GOBP_LYTIC_VACUOLE_ORGANIZATION | 70 | -0.17936 | -0.530342 | 0.997783 | 1 |
| GOBP_MIRNA_METABOLIC_PROCESS | 27 | -0.209075 | -0.52978 | 0.984694 | 1 |
| GOBP_ENTRAINMENT_OF_CIRCADIAN_CLOCK | 31 | -0.198377 | -0.52684 | 0.98852 | 1 |
| GOBP_INTERLEUKIN_1_MEDIATED_SIGNALING_PATHWAY | 100 | -0.170789 | -0.526677 | 1 | 1 |
| GOBP_PROTEIN_LOCALIZATION_TO_MITOCHONDRION | 146 | -0.162805 | -0.526431 | 1 | 1 |
| GOBP_KETONE_BIOSYNTHETIC_PROCESS | 39 | -0.193033 | -0.526143 | 0.993758 | 1 |
| GOBP_POSITIVE_REGULATION_OF_MRNA_METABOLIC_PROCESS | 80 | -0.173132 | -0.526073 | 0.998828 | 1 |
| GOBP_POSITIVE_REGULATION_OF_CHROMOSOME_ORGANIZATION | 162 | -0.160721 | -0.525649 | 1 | 1 |
| GOBP_TRANSFORMING_GROWTH_FACTOR_BETA_PRODUCTION | 42 | -0.187809 | -0.525116 | 0.991283 | 1 |
| GOBP_MORPHOGENESIS_OF_A_POLARIZED_EPITHELIUM | 143 | -0.161192 | -0.52428 | 1 | 1 |
| GOBP_AMYLOID_PRECURSOR_PROTEIN_CATABOLIC_PROCESS | 54 | -0.181929 | -0.523862 | 0.997625 | 1 |
| GOBP_CELLULAR_RESPONSE_TO_ABIOTIC_STIMULUS | 316 | -0.156652 | -0.523683 | 1 | 1 |
| GOBP_NEGATIVE_REGULATION_OF_ACTIN_FILAMENT_BUNDLE_ASSEMBLY | 31 | -0.198131 | -0.521893 | 0.989744 | 1 |
| GOBP_HIPPO_SIGNALING | 45 | -0.185581 | -0.520726 | 0.995031 | 1 |
| GOBP_RAS_PROTEIN_SIGNAL_TRANSDUCTION | 334 | -0.1546 | -0.520275 | 1 | 1 |
| GOBP_POSITIVE_REGULATION_OF_UBIQUITIN_PROTEIN_TRANSFERASE_ACTIV | 32 | -0.195955 | -0.520207 | 0.989583 | 1 |
| GOBP_REGULATION_OF_TRANSCRIPTION_BY_RNA_POLYMERASE_III | 23 | -0.209119 | -0.520126 | 0.982872 | 1 |
| GOBP_HISTONE_H4_ACETYLATION | 66 | -0.175078 | -0.519754 | 0.998804 | 1 |
| GOBP_RECEPTOR_METABOLIC_PROCESS | 161 | -0.160181 | -0.519483 | 1 | 1 |
| GOBP_TRANSPORT_ALONG_MICROTUBULE | 156 | -0.159549 | -0.519159 | 1 | 1 |
| GOBP_FACE_DEVELOPMENT | 44 | -0.188351 | -0.518873 | 0.998759 | 1 |
| GOBP_CRISTAE_FORMATION | 35 | -0.192597 | -0.518478 | 0.997439 | 1 |
| GOBP_NEGATIVE_REGULATION_OF_CANONICAL_WNT_SIGNALING_PATHWAY | 172 | -0.158105 | -0.515612 | 1 | 1 |
| GOBP_REGULATION_OF_PROGRAMMED_NECROTIC_CELL_DEATH | 22 | -0.204121 | -0.515188 | 0.983007 | 1 |
| GOBP_NUCLEOSIDE_TRIPHOSPHATE_METABOLIC_PROCESS | 107 | -0.163354 | -0.514752 | 1 | 1 |
| GOBP_RNA_5_END_PROCESSING | 23 | -0.203649 | -0.513616 | 0.99085 | 1 |
| GOBP_RECEPTOR_INTERNALIZATION | 105 | -0.165332 | -0.513576 | 0.99889 | 1 |
| GOBP_NUCLEAR_TRANSCRIBED_MRNA_POLY_A_TAIL_SHORTENING | 34 | -0.192102 | -0.513429 | 0.992395 | 1 |
| GOBP_MITOCHONDRIAL_TRANSLATIONAL_TERMINATION | 89 | -0.166744 | -0.513342 | 1 | 1 |
| GOBP_MATURATION_OF_5_8S_RRNA | 35 | -0.192198 | -0.513317 | 0.995019 | 1 |
| GOBP_PROTEIN_K63_LINKED_UBIQUITINATION | 56 | -0.175965 | -0.512188 | 0.996466 | 1 |
| GOBP_MULTIVESICULAR_BODY_SORTING_PATHWAY | 37 | -0.188275 | -0.511968 | 0.996319 | 1 |
| GOBP_CELLULAR_COMPONENT_MAINTENANCE | 60 | -0.174562 | -0.511441 | 1 | 1 |
| GOBP_TELOMERE_MAINTENANCE_VIA_SEMI_CONSERVATIVE_REPLICATION | 27 | -0.200688 | -0.511163 | 0.992032 | 1 |
| GOBP_POSITIVE_REGULATION_OF_ACTIN_FILAMENT_BUNDLE_ASSEMBLY | 61 | -0.175288 | -0.510667 | 0.998829 | 1 |
| GOBP_CELLULAR_RESPONSE_TO_FLUID_SHEAR_STRESS | 17 | -0.216579 | -0.509968 | 0.971154 | 1 |
| GOBP_ENDOMEMBRANE_SYSTEM_ORGANIZATION | 464 | -0.149732 | -0.50889 | 1 | 1 |
| GOBP_CELLULAR_RESPONSE_TO_OXYGEN_LEVELS | 216 | -0.153006 | -0.507767 | 1 | 1 |
| GOBP_BASE_EXCISION_REPAIR | 39 | -0.185615 | -0.507178 | 0.996207 | 1 |
| GOBP_TYPE_I_INTERFERON_PRODUCTION | 126 | -0.159424 | -0.506697 | 1 | 1 |
| GOBP_SEGMENT_SPECIFICATION | 16 | -0.215332 | -0.505882 | 0.986207 | 1 |
| GOBP_DNA_BIOSYNTHETIC_PROCESS | 185 | -0.155781 | -0.505476 | 1 | 1 |
| GOBP_NEGATIVE_REGULATION_OF_RESPONSE_TO_REACTIVE_OXYGEN_SPEC | 16 | -0.213973 | -0.50465 | 0.9809 | 1 |
| GOBP_PHOSPHATIDYLINOSITOL_BIOSYNTHETIC_PROCESS | 115 | -0.158264 | -0.503611 | 0.998899 | 1 |
| GOBP_EPITHELIAL_CELL_APOPTOTIC_PROCESS | 101 | -0.15996 | -0.502536 | 1 | 1 |

| | | | | | |
|---|---|---|---|---|---|
| GOBP_ERROR_FREE_TRANSLESION_SYNTHESIS | 22 | -0.206554 | -0.50149 | 0.989262 | 1 |
| GOBP_TOR_SIGNALING | 118 | -0.157567 | -0.501173 | 0.998902 | 1 |
| GOBP_POSITIVE_REGULATION_OF_MUSCLE_TISSUE_DEVELOPMENT | 21 | -0.204143 | -0.500741 | 0.98269 | 1 |
| GOBP_REGULATION_OF_TRANSCRIPTION_ELONGATION_FROM_RNA_POLYME | 30 | -0.189315 | -0.500726 | 0.989664 | 1 |
| GOBP_CANONICAL_WNT_SIGNALING_PATHWAY | 320 | -0.149435 | -0.500446 | 1 | 1 |
| GOBP_RNA_LOCALIZATION | 227 | -0.15047 | -0.499063 | 1 | 1 |
| GOBP_NEGATIVE_REGULATION_OF_WNT_SIGNALING_PATHWAY | 205 | -0.151235 | -0.497537 | 1 | 1 |
| GOBP_REGULATION_OF_MEMBRANE_PERMEABILITY | 81 | -0.161479 | -0.496769 | 1 | 1 |
| GOBP_PYRIMIDINE_NUCLEOSIDE_TRIPHOSPHATE_BIOSYNTHETIC_PROCESS | 19 | -0.203069 | -0.495851 | 0.990237 | 1 |
| GOBP_PROTEIN_CONTAINING_COMPLEX_DISASSEMBLY | 323 | -0.147013 | -0.494744 | 1 | 1 |
| GOBP_REGULATION_OF_NUCLEOCYTOPLASMIC_TRANSPORT | 103 | -0.158322 | -0.493889 | 1 | 1 |
| GOBP_PROTEIN_INSERTION_INTO_ER_MEMBRANE | 19 | -0.205962 | -0.49375 | 0.991837 | 1 |
| GOBP_NCRNA_CATABOLIC_PROCESS | 35 | -0.184273 | -0.492878 | 0.996212 | 1 |
| GOBP_POSITIVE_REGULATION_OF_TOR_SIGNALING | 40 | -0.179444 | -0.492557 | 0.995068 | 1 |
| GOBP_MAINTENANCE_OF_PROTEIN_LOCATION | 90 | -0.160097 | -0.492103 | 1 | 1 |
| GOBP_CYTOPLASMIC_PATTERN_RECOGNITION_RECEPTOR_SIGNALING_PATH | 30 | -0.188364 | -0.492082 | 0.990838 | 1 |
| GOBP_MITOCHONDRIAL_CALCIUM_ION_TRANSMEMBRANE_TRANSPORT | 25 | -0.194664 | -0.492045 | 0.986877 | 1 |
| GOBP_VIRAL_LIFE_CYCLE | 338 | -0.146239 | -0.4915 | 1 | 1 |
| GOBP_RESPONSE_TO_IONIZING_RADIATION | 141 | -0.15288 | -0.491243 | 1 | 1 |
| GOBP_NCRNA_PROCESSING | 384 | -0.143352 | -0.489874 | 1 | 1 |
| GOBP_POSITIVE_REGULATION_OF_PROTEIN_EXIT_FROM_ENDOPLASMIC_RET | 15 | -0.217871 | -0.488929 | 0.984441 | 1 |
| GOBP_APOPTOTIC_MITOCHONDRIAL_CHANGES | 112 | -0.156365 | -0.488445 | 1 | 1 |
| GOBP_LABYRINTHINE_LAYER_DEVELOPMENT | 42 | -0.175989 | -0.487084 | 1 | 1 |
| GOBP_REGULATION_OF_SPHINGOLIPID_BIOSYNTHETIC_PROCESS | 18 | -0.20436 | -0.486402 | 0.985856 | 1 |
| GOBP_TRNA_TRANSPORT | 37 | -0.181208 | -0.48636 | 0.994943 | 1 |
| GOBP_NEGATIVE_REGULATION_OF_HISTONE_MODIFICATION | 43 | -0.176397 | -0.48627 | 0.995025 | 1 |
| GOBP_REGULATION_OF_POLYSACCHARIDE_BIOSYNTHETIC_PROCESS | 35 | -0.180313 | -0.485113 | 0.997319 | 1 |
| GOBP_POSITIVE_REGULATION_OF_STRESS_FIBER_ASSEMBLY | 52 | -0.169869 | -0.483903 | 0.995215 | 1 |
| GOBP_NEGATIVE_REGULATION_OF_FAT_CELL_DIFFERENTIATION | 49 | -0.171462 | -0.481901 | 0.998794 | 1 |
| GOBP_TRNA_METABOLIC_PROCESS | 177 | -0.147964 | -0.481892 | 1 | 1 |
| GOBP_DNA_INTEGRITY_CHECKPOINT | 152 | -0.147741 | -0.481708 | 1 | 1 |
| GOBP_REGULATION_OF_NON_CANONICAL_WNT_SIGNALING_PATHWAY | 25 | -0.188169 | -0.481239 | 0.996005 | 1 |
| GOBP_PROTEIN_TRANSMEMBRANE_IMPORT_INTO_INTRACELLULAR_ORGANE | 36 | -0.175329 | -0.481016 | 0.995019 | 1 |
| GOBP_POSITIVE_REGULATION_OF_PROTEOLYSIS_INVOLVED_IN_CELLULAR_F | 126 | -0.1506 | -0.480729 | 0.998893 | 1 |
| GOBP_INTRINSIC_APOPTOTIC_SIGNALING_PATHWAY | 274 | -0.143603 | -0.480519 | 1 | 1 |
| GOBP_PROTEIN_QUALITY_CONTROL_FOR_MISFOLDED_OR_INCOMPLETELY_S | 28 | -0.184868 | -0.480146 | 0.990814 | 1 |
| GOBP_TERMINATION_OF_RNA_POLYMERASE_I_TRANSCRIPTION | 31 | -0.184807 | -0.479315 | 0.994798 | 1 |
| GOBP_POSITIVE_REGULATION_OF_CD4_POSITIVE_ALPHA_BETA_T_CELL_DIFF | 31 | -0.181434 | -0.478846 | 0.993614 | 1 |
| GOBP_CORTICAL_CYTOSKELETON_ORGANIZATION | 58 | -0.162788 | -0.476553 | 0.998808 | 1 |
| GOBP_NUCLEAR_ENVELOPE_ORGANIZATION | 51 | -0.167302 | -0.476347 | 0.997619 | 1 |
| GOBP_PYRIMIDINE_DEOXYRIBONUCLEOTIDE_METABOLIC_PROCESS | 22 | -0.193705 | -0.475913 | 0.993324 | 1 |
| GOBP_REGULATION_OF_PROTEIN_CONTAINING_COMPLEX_DISASSEMBLY | 119 | -0.150441 | -0.475168 | 1 | 1 |
| GOBP_MYELOID_CELL_DEVELOPMENT | 66 | -0.159462 | -0.471364 | 0.996458 | 1 |
| GOBP_PROTEIN_DESTABILIZATION | 46 | -0.168354 | -0.471073 | 1 | 1 |
| GOBP_TRANSLATIONAL_ELONGATION | 133 | -0.148569 | -0.469613 | 1 | 1 |
| GOBP_REGULATION_OF_PROTEIN_ACETYLATION | 72 | -0.155417 | -0.468748 | 0.998855 | 1 |
| GOBP_ELECTRON_TRANSPORT_CHAIN | 174 | -0.142674 | -0.46852 | 1 | 1 |
| GOBP_RNA_MODIFICATION | 166 | -0.144304 | -0.467808 | 1 | 1 |
| GOBP_G2_DNA_DAMAGE_CHECKPOINT | 34 | -0.174817 | -0.467082 | 0.99871 | 1 |
| GOBP_REGULATION_OF_VASCULAR_ASSOCIATED_SMOOTH_MUSCLE_CELL_[ | 16 | -0.19989 | -0.466019 | 0.989362 | 1 |
| GOBP_CELLULAR_RESPONSE_TO_RADIATION | 179 | -0.143226 | -0.465284 | 1 | 1 |
| GOBP_PROTEIN_IMPORT | 189 | -0.141628 | -0.464611 | 1 | 1 |
| GOBP_REGULATION_OF_CARDIAC_MUSCLE_CONTRACTION_BY_REGULATION | 20 | -0.191257 | -0.46342 | 0.994616 | 1 |
| GOBP_POSITIVE_REGULATION_OF_CHROMATIN_ORGANIZATION | 98 | -0.15141 | -0.463412 | 1 | 1 |
| GOBP_NEGATIVE_REGULATION_OF_STRESS_ACTIVATED_PROTEIN_KINASE_S | 45 | -0.166432 | -0.462945 | 1 | 1 |
| GOBP_RRNA_METABOLIC_PROCESS | 237 | -0.138402 | -0.462799 | 1 | 1 |
| GOBP_PROTEIN_CONTAINING_COMPLEX_LOCALIZATION | 274 | -0.137816 | -0.461656 | 1 | 1 |
| GOBP_POSITIVE_REGULATION_OF_CELLULAR_AMIDE_METABOLIC_PROCESS | 150 | -0.142103 | -0.461476 | 1 | 1 |
| GOBP_TRANSCRIPTION_PREINITIATION_COMPLEX_ASSEMBLY | 35 | -0.173657 | -0.460886 | 0.998792 | 1 |
| GOBP_MITOCHONDRIAL_TRANSMEMBRANE_TRANSPORT | 107 | -0.147998 | -0.460787 | 1 | 1 |
| GOBP_REGULATION_OF_AUTOPHAGY_OF_MITOCHONDRION | 31 | -0.176563 | -0.460719 | 0.997368 | 1 |
| GOBP_MRNA_3_END_PROCESSING | 98 | -0.147882 | -0.460582 | 1 | 1 |
| GOBP_REGULATION_OF_PROTEIN_CATABOLIC_PROCESS | 375 | -0.135425 | -0.459008 | 1 | 1 |
| GOBP_PROTEIN_EXIT_FROM_ENDOPLASMIC_RETICULUM | 47 | -0.162287 | -0.45745 | 1 | 1 |
| GOBP_CLATHRIN_DEPENDENT_ENDOCYTOSIS | 46 | -0.164033 | -0.457125 | 0.997512 | 1 |
| GOBP_CELLULAR_PROTEIN_COMPLEX_DISASSEMBLY | 219 | -0.136992 | -0.456746 | 1 | 1 |
| GOBP_MYD88_INDEPENDENT_TOLL_LIKE_RECEPTOR_SIGNALING_PATHWAY | 34 | -0.171472 | -0.456607 | 1 | 1 |
| GOBP_NEGATIVE_REGULATION_OF_MITOCHONDRION_ORGANIZATION | 44 | -0.163676 | -0.455678 | 1 | 1 |
| GOBP_ESTABLISHMENT_OF_ORGANELLE_LOCALIZATION | 424 | -0.133301 | -0.45563 | 1 | 1 |
| GOBP_ATP_METABOLIC_PROCESS | 306 | -0.135871 | -0.455259 | 1 | 1 |
| GOBP_INTEGRATED_STRESS_RESPONSE_SIGNALING | 27 | -0.176099 | -0.453171 | 0.996068 | 1 |
| GOBP_NEGATIVE_REGULATION_OF_ORGANELLE_ASSEMBLY | 41 | -0.167377 | -0.453134 | 0.998745 | 1 |
| GOBP_REGULATION_OF_PEPTIDYL_LYSINE_ACETYLATION | 62 | -0.155907 | -0.452882 | 1 | 1 |
| GOBP_VIRAL_GENE_EXPRESSION | 198 | -0.137342 | -0.452483 | 1 | 1 |
| GOBP_INTRINSIC_APOPTOTIC_SIGNALING_PATHWAY_IN_RESPONSE_TO_END( | 59 | -0.155265 | -0.452285 | 1 | 1 |
| GOBP_PURINE_NUCLEOSIDE_BIOSYNTHETIC_PROCESS | 22 | -0.183266 | -0.450794 | 0.99169 | 1 |
| GOBP_PROTEIN_METHYLATION | 163 | -0.13719 | -0.449956 | 1 | 1 |
| GOBP_CILIUM_ORGANIZATION | 397 | -0.132391 | -0.449879 | 1 | 1 |
| GOBP_REGULATION_OF_CELLULAR_RESPONSE_TO_TRANSFORMING_GROWT | 119 | -0.143005 | -0.449837 | 1 | 1 |
| GOBP_CELLULAR_RESPONSE_TO_HYDROGEN_PEROXIDE | 81 | -0.148358 | -0.449498 | 0.99884 | 1 |
| GOBP_PEROXISOMAL_MEMBRANE_TRANSPORT | 20 | -0.183538 | -0.446107 | 0.99581 | 1 |
| GOBP_CELLULAR_RESPONSE_TO_IONIZING_RADIATION | 64 | -0.152619 | -0.44488 | 1 | 1 |
| GOBP_PROTEIN_TARGETING_TO_MEMBRANE | 201 | -0.135455 | -0.443617 | 1 | 1 |
| GOBP_NEGATIVE_REGULATION_OF_GTPASE_ACTIVITY | 44 | -0.157713 | -0.440104 | 1 | 1 |
| GOBP_NEGATIVE_REGULATION_OF_ESTABLISHMENT_OF_PROTEIN_LOCALIZA | 111 | -0.141203 | -0.437238 | 1 | 1 |
| GOBP_REGULATION_OF_DOUBLE_STRAND_BREAK_REPAIR | 85 | -0.143786 | -0.437198 | 1 | 1 |
| GOBP_POSITIVE_REGULATION_OF_PROTEIN_TARGETING_TO_MITOCHONDRIO | 32 | -0.164856 | -0.437171 | 0.997475 | 1 |
| GOBP_PROTEIN_LOCALIZATION_TO_CYTOSKELETON | 58 | -0.14851 | -0.435881 | 0.998812 | 1 |
| GOBP_POSITIVE_REGULATION_OF_UBIQUITIN_DEPENDENT_PROTEIN_CATABO | 103 | -0.140545 | -0.435518 | 1 | 1 |
| GOBP_REGULATION_OF_TRANSCRIPTION_FROM_RNA_POLYMERASE_II_PROM | 76 | -0.142206 | -0.435086 | 1 | 1 |
| GOBP_RIBONUCLEOSIDE_TRIPHOSPHATE_BIOSYNTHETIC_PROCESS | 72 | -0.143977 | -0.432636 | 1 | 1 |
| GOBP_REGULATION_OF_PROTEIN_TARGETING_TO_MITOCHONDRION | 44 | -0.153316 | -0.431464 | 1 | 1 |
| GOBP_PROTEIN_TRANSMEMBRANE_TRANSPORT | 60 | -0.145446 | -0.429907 | 1 | 1 |
| GOBP_CONVERGENT_EXTENSION | 16 | -0.184027 | -0.428825 | 0.993141 | 1 |
| GOBP_NEGATIVE_REGULATION_OF_TYPE_I_INTERFERON_PRODUCTION | 45 | -0.151838 | -0.428084 | 1 | 1 |
| GOBP_ESTABLISHMENT_OF_PROTEIN_LOCALIZATION_TO_MEMBRANE | 343 | -0.12494 | -0.427091 | 1 | 1 |
| GOBP_PROTEIN_LOCALIZATION_TO_LYSOSOME | 42 | -0.155 | -0.426102 | 1 | 1 |
| GOBP_CYTOSKELETON_DEPENDENT_INTRACELLULAR_TRANSPORT | 194 | -0.128874 | -0.425122 | 1 | 1 |
| GOBP_RESPONSE_TO_TUMOR_CELL | 27 | -0.163649 | -0.423272 | 0.998638 | 1 |
| GOBP_CYTOPLASMIC_TRANSLATIONAL_INITIATION | 34 | -0.157361 | -0.421125 | 1 | 1 |
| GOBP_STRESS_GRANULE_ASSEMBLY | 23 | -0.169516 | -0.420531 | 0.99866 | 1 |
| GOBP_ACTIN_CYTOSKELETON_REORGANIZATION | 101 | -0.13668 | -0.419721 | 1 | 1 |
| GOBP_NEGATIVE_REGULATION_OF_DNA_REPAIR | 36 | -0.155278 | -0.419492 | 1 | 1 |
| GOBP_LATE_ENDOSOME_TO_VACUOLE_TRANSPORT | 24 | -0.163948 | -0.415827 | 0.998636 | 1 |
| GOBP_NEGATIVE_REGULATION_OF_TELOMERE_MAINTENANCE_VIA_TELOMEF | 20 | -0.170409 | -0.414746 | 0.997241 | 1 |
| GOBP_REGULATION_OF_PROTEIN_TARGETING | 80 | -0.135643 | -0.414664 | 1 | 1 |
| GOBP_TELOMERE_MAINTENANCE_VIA_RECOMBINATION | 15 | -0.182436 | -0.414361 | 0.99435 | 1 |
| GOBP_POSITIVE_REGULATION_OF_SIGNAL_TRANSDUCTION_BY_P53_CLASS_[ | 22 | -0.168511 | -0.414159 | 0.994498 | 1 |
| GOBP_SIGNAL_TRANSDUCTION_BY_P53_CLASS_MEDIATOR | 258 | -0.123679 | -0.414092 | 1 | 1 |
| GOBP_REGULATION_OF_TOR_SIGNALING | 96 | -0.133574 | -0.413831 | 1 | 1 |
| GOBP_RIBONUCLEOSIDE_TRIPHOSPHATE_METABOLIC_PROCESS | 86 | -0.135657 | -0.411866 | 1 | 1 |
| GOBP_REGULATION_OF_PROTEIN_EXPORT_FROM_NUCLEUS | 32 | -0.158559 | -0.411178 | 1 | 1 |
| GOBP_POSITIVE_REGULATION_OF_ATPASE_ACTIVITY | 56 | -0.142059 | -0.410937 | 1 | 1 |
| GOBP_MONOCYTE_CHEMOTACTIC_PROTEIN_1_PRODUCTION | 16 | -0.172985 | -0.408707 | 1 | 1 |
| GOBP_LACTATE_METABOLIC_PROCESS | 15 | -0.171961 | -0.402503 | 1 | 1 |
| GOBP_CELLULAR_RESPONSE_TO_GLUCOSE_STARVATION | 47 | -0.140633 | -0.399105 | 1 | 1 |
| GOBP_ANTIGEN_PROCESSING_AND_PRESENTATION_OF_PEPTIDE_OR_POLYS | 101 | -0.127723 | -0.398277 | 1 | 1 |
| GOBP_NUCLEOSIDE_TRIPHOSPHATE_BIOSYNTHETIC_PROCESS | 83 | -0.13036 | -0.39653 | 1 | 1 |
| GOBP_NEGATIVE_REGULATION_OF_PLATELET_ACTIVATION | 17 | -0.1693 | -0.395731 | 0.998645 | 1 |
| GOBP_MITOCHONDRIAL_TRANSLATION | 134 | -0.124792 | -0.395084 | 1 | 1 |
| GOBP_REGULATION_OF_CELLULAR_PROTEIN_CATABOLIC_PROCESS | 247 | -0.118707 | -0.394391 | 1 | 1 |
| GOBP_PROTEIN_TARGETING | 430 | -0.11523 | -0.393623 | 1 | 1 |
| GOBP_ENDOSOME_ORGANIZATION | 80 | -0.128059 | -0.393409 | 1 | 1 |
| GOBP_GOLGI_ORGANIZATION | 141 | -0.122624 | -0.39236 | 1 | 1 |
| GOBP_RRNA_TRANSCRIPTION | 33 | -0.148918 | -0.391914 | 1 | 1 |
| GOBP_TRANSLATIONAL_TERMINATION | 105 | -0.124939 | -0.391625 | 1 | 1 |
| GOBP_LATE_ENDOSOME_TO_VACUOLE_TRANSPORT_VIA_MULTIVESICULAR_[ | 17 | -0.164603 | -0.390977 | 1 | 1 |
| GOBP_PROTEIN_POLYUBIQUITINATION | 330 | -0.116775 | -0.390785 | 1 | 1 |
| GOBP_RNA_INTERFERENCE | 16 | -0.168774 | -0.390774 | 1 | 1 |
| GOBP_PROTEIN_O_LINKED_MANNOSYLATION | 18 | -0.160871 | -0.388386 | 1 | 1 |
| GOBP_NEGATIVE_REGULATION_OF_DEFENSE_RESPONSE_TO_VIRUS | 23 | -0.155099 | -0.387432 | 0.997312 | 1 |
| GOBP_OXIDATIVE_PHOSPHORYLATION | 146 | -0.119239 | -0.386395 | 1 | 1 |
| GOBP_RNA_DEPENDENT_DNA_BIOSYNTHETIC_PROCESS | 69 | -0.130155 | -0.386338 | 1 | 1 |
| GOBP_POST_GOLGI_VESICLE_MEDIATED_TRANSPORT | 103 | -0.123708 | -0.384199 | 1 | 1 |
| GOBP_ESTABLISHMENT_OF_PROTEIN_LOCALIZATION_TO_PLASMA_MEMBRAN | 58 | -0.131286 | -0.379989 | 1 | 1 |
| GOBP_ATP_BIOSYNTHETIC_PROCESS | 55 | -0.129032 | -0.374582 | 1 | 1 |
| GOBP_BETA_CATENIN_TCF_COMPLEX_ASSEMBLY | 31 | -0.142588 | -0.373083 | 1 | 1 |
| GOBP_REGULATION_OF_ORGANELLE_ASSEMBLY | 184 | -0.114885 | -0.372765 | 1 | 1 |

| | | | | | |
|---|---|---|---|---|---|
| GOBP_REGULATION_OF_ENDOTHELIAL_CELL_DEVELOPMENT | 17 | -0.154425 | -0.369329 | 1 | 1 |
| GOBP_VASCULAR_WOUND_HEALING | 17 | -0.153795 | -0.361385 | 1 | 1 |
| GOBP_PROTEIN_LOCALIZATION_TO_NUCLEUS | 268 | -0.104717 | -0.350529 | 1 | 1 |
| GOBP_TRANSCRIPTION_COUPLED_NUCLEOTIDE_EXCISION_REPAIR | 73 | -0.11395 | -0.341853 | 1 | 1 |
| GOBP_TRNA_5_END_PROCESSING | 16 | -0.143796 | -0.337649 | 1 | 1 |
| GOBP_PROTEIN_LOCALIZATION_TO_VACUOLE | 67 | -0.114165 | -0.33749 | 1 | 1 |
| GOBP_RNA_METHYLATION | 84 | -0.107771 | -0.329165 | 1 | 1 |
| GOBP_PROTEASOMAL_PROTEIN_CATABOLIC_PROCESS | 474 | -0.096265 | -0.328538 | 1 | 1 |
| GOBP_TRANSCRIPTION_BY_RNA_POLYMERASE_III | 47 | -0.10705 | -0.300838 | 1 | 1 |
| GOBP_ARP2_3_COMPLEX_MEDIATED_ACTIN_NUCLEATION | 39 | -0.107782 | -0.294428 | 1 | 1 |
| GOBP_DNA_TEMPLATED_TRANSCRIPTION_TERMINATION | 75 | -0.094642 | -0.284263 | 1 | 1 |
| GOBP_RESPIRATORY_ELECTRON_TRANSPORT_CHAIN | 115 | -0.089826 | -0.280932 | 1 | 0.999999 |

**Table S12. GSEA analysis of RNA-seq data in $SF3B1^{MT}$ versus $SF3B1^{WT}$ UVM patient cells.** See table S16 for patient information.

**GSEA Positive** Hematopoietic/Immune pathways are highlighted in red.

| Gene sets | SIZE | ES | NES | NOM p-val | FDR q-val |
|---|---|---|---|---|---|
| GOBP_RNA_SPLICING_VIA_TRANSESTERIFICATION_REACTIONS | 355 | 0.142072 | --- | --- | 1 |
| GOBP_RNA_CATABOLIC_PROCESS | 398 | 0.200266 | --- | --- | 1 |
| GOBP_RNA_LOCALIZATION | 226 | 0.100093 | --- | --- | 1 |
| GOBP_CHROMOSOME_SEGREGATION | 321 | 0.23192 | --- | --- | 1 |
| GOBP_RNA_SPLICING | 443 | 0.148349 | --- | --- | 1 |
| GOBP_NEGATIVE_REGULATION_OF_CELL_CYCLE_PROCESS | 323 | 0.133336 | --- | --- | 1 |
| GOBP_RRNA_METABOLIC_PROCESS | 236 | 0.311791 | --- | --- | 1 |
| GOBP_RIBONUCLEOPROTEIN_COMPLEX_BIOGENESIS | 450 | 0.275226 | --- | --- | 1 |
| GOBP_REGULATION_OF_CHROMOSOME_ORGANIZATION | 264 | 0.14286 | --- | --- | 1 |
| GOBP_NCRNA_PROCESSING | 383 | 0.261688 | --- | --- | 1 |
| GOBP_NCRNA_METABOLIC_PROCESS | 472 | 0.226867 | --- | --- | 1 |
| GOBP_RIBOSOME_BIOGENESIS | 306 | 0.327322 | --- | --- | 1 |
| GOBP_PROTEIN_DNA_COMPLEX_SUBUNIT_ORGANIZATION | 241 | 0.169436 | --- | --- | 1 |
| GOBP_NUCLEIC_ACID_PHOSPHODIESTER_BOND_HYDROLYSIS | 287 | 0.179942 | --- | --- | 1 |
| GOBP_NUCLEAR_CHROMOSOME_SEGREGATION | 259 | 0.223306 | --- | --- | 1 |
| GOBP_NUCLEAR_TRANSCRIBED_MRNA_CATABOLIC_PROCESS_NONSENSE_MEDIATE | 120 | 0.566505 | 2.487 | 0 | 0 |
| GOBP_COTRANSLATIONAL_PROTEIN_TARGETING_TO_MEMBRANE | 104 | 0.523855 | 2.25 | 0 | 0.004613 |
| GOBP_RIBOSOMAL_LARGE_SUBUNIT_BIOGENESIS | 72 | 0.513471 | 2.118 | 0 | 0.026384 |
| GOBP_VIRAL_RELEASE_FROM_HOST_CELL | 32 | 0.589655 | 2.046 | 0 | 0.051569 |
| GOBP_ESTABLISHMENT_OF_PROTEIN_LOCALIZATION_TO_ENDOPLASMIC_RETICULUM | 118 | 0.45481 | 1.973 | 0 | 0.083908 |
| GOBP_NUCLEAR_TRANSCRIBED_MRNA_CATABOLIC_PROCESS | 206 | 0.426964 | 1.935 | 0 | 0.10736 |
| GOBP_RIBOSOMAL_LARGE_SUBUNIT_ASSEMBLY | 28 | 0.57402 | 1.899 | 0 | 0.132888 |
| GOBP_VIRAL_GENE_EXPRESSION | 198 | 0.398914 | 1.866 | 0 | 0.159495 |
| GOBP_TRANSLATIONAL_INITIATION | 187 | 0.410112 | 1.815 | 0 | 0.219601 |
| GOBP_BETA_CATENIN_DESTRUCTION_COMPLEX_DISASSEMBLY | 19 | 0.568278 | 1.77 | 0.014563 | 0.285341 |
| GOBP_HOMOLOGOUS_CHROMOSOME_PAIRING_AT_MEIOSIS | 44 | 0.466641 | 1.74 | 0 | 0.331726 |
| GOBP_TELOMERASE_RNA_LOCALIZATION | 19 | 0.559446 | 1.728 | 0.018868 | 0.333455 |
| GOBP_ATP_SYNTHESIS_COUPLED_ELECTRON_TRANSPORT | 99 | 0.414271 | 1.715 | 0 | 0.339672 |
| GOBP_CYTOPLASMIC_TRANSLATION | 99 | 0.401428 | 1.704 | 0 | 0.339798 |
| GOBP_OXIDATIVE_PHOSPHORYLATION | 144 | 0.357901 | 1.695 | 0 | 0.335468 |
| GOBP_PROTEIN_LOCALIZATION_TO_ENDOPLASMIC_RETICULUM | 146 | 0.388675 | 1.645 | 0 | 0.454113 |
| GOBP_SISTER_CHROMATID_COHESION | 58 | 0.419138 | 1.639 | 0 | 0.443816 |
| GOBP_SYNAPTONEMAL_COMPLEX_ORGANIZATION | 21 | 0.534043 | 1.624 | 0.022857 | 0.463308 |
| GOBP_CELLULAR_RESPONSE_TO_STEROL_DEPLETION | 16 | 0.561673 | 1.621 | 0.010204 | 0.448315 |
| GOBP_PROTEIN_DEMETHYLATION | 33 | 0.468828 | 1.617 | 0 | 0.438955 |
| GOBP_PROTEIN_DNA_COMPLEX_DISASSEMBLY | 18 | 0.527042 | 1.595 | 0.030928 | 0.484622 |
| GOBP_NEGATIVE_REGULATION_OF_ANDROGEN_RECEPTOR_SIGNALING_PATHWAY | 15 | 0.54218 | 1.589 | 0.037657 | 0.481479 |
| GOBP_RNA_5_END_PROCESSING | 23 | 0.49918 | 1.587 | 0.013072 | 0.466192 |
| GOBP_REGULATION_OF_MEIOTIC_NUCLEAR_DIVISION | 26 | 0.475032 | 1.582 | 0.00625 | 0.460678 |
| GOBP_RESPIRATORY_ELECTRON_TRANSPORT_CHAIN | 115 | 0.352532 | 1.581 | 0 | 0.445326 |
| GOBP_RIBOSOME_ASSEMBLY | 64 | 0.403973 | 1.579 | 0 | 0.434111 |
| GOBP_LIPID_DIGESTION | 16 | 0.548534 | 1.57 | 0.040816 | 0.441894 |
| GOBP_MATURATION_OF_LSU_RRNA | 28 | 0.471689 | 1.566 | 0.021127 | 0.438266 |
| GOBP_MITOCHONDRIAL_ELECTRON_TRANSPORT_CYTOCHROME_C_TO_OXYGEN | 21 | 0.505897 | 1.566 | 0.051136 | 0.423859 |
| GOBP_CHROMATIN_DISASSEMBLY | 18 | 0.521483 | 1.564 | 0.038043 | 0.413437 |
| GOBP_PHOTOPERIODISM | 32 | 0.461913 | 1.557 | 0.025478 | 0.418268 |
| GOBP_MITOCHONDRIAL_ELECTRON_TRANSPORT_NADH_TO_UBIQUINONE | 54 | 0.399543 | 1.55 | 0.012821 | 0.424862 |
| GOBP_CHROMOSOME_ORGANIZATION_INVOLVED_IN_MEIOTIC_CELL_CYCLE | 59 | 0.398815 | 1.541 | 0.016667 | 0.435039 |
| GOBP_HOMOLOGOUS_CHROMOSOME_SEGREGATION | 52 | 0.420222 | 1.537 | 0.020619 | 0.434807 |
| GOBP_POSITIVE_REGULATION_OF_TELOMERASE_RNA_LOCALIZATION_TO_CAJAL_BO | 15 | 0.547139 | 1.527 | 0.070796 | 0.447055 |
| GOBP_SPERM_CAPACITATION | 20 | 0.504747 | 1.514 | 0.035 | 0.471108 |
| GOBP_MITOCHONDRIAL_TRANSLATION | 134 | 0.341046 | 1.512 | 0 | 0.465229 |
| GOBP_PEPTIDYL_LYSINE_DIMETHYLATION | 21 | 0.483757 | 1.505 | 0.034483 | 0.472391 |
| GOBP_CORNIFICATION | 56 | 0.394116 | 1.493 | 0.039474 | 0.49503 |
| GOBP_PSEUDOURIDINE_SYNTHESIS | 18 | 0.484253 | 1.48 | 0.04878 | 0.521907 |
| GOBP_CRISTAE_FORMATION | 35 | 0.422395 | 1.476 | 0.024793 | 0.523075 |
| GOBP_RRNA_TRANSCRIPTION | 33 | 0.42676 | 1.475 | 0.036496 | 0.510899 |
| GOBP_NEGATIVE_REGULATION_OF_EXECUTION_PHASE_OF_APOPTOSIS | 18 | 0.489264 | 1.473 | 0.076531 | 0.507049 |
| GOBP_REGULATION_OF_TRANSLATIONAL_FIDELITY | 20 | 0.485757 | 1.472 | 0.031414 | 0.497472 |
| GOBP_NUCLEOTIDE_EXCISION_REPAIR_DNA_INCISION | 39 | 0.416623 | 1.463 | 0.022727 | 0.510931 |
| GOBP_KERATINIZATION | 71 | 0.352206 | 1.462 | 0.022727 | 0.501764 |
| GOBP_CYTOPLASMIC_TRANSLATIONAL_INITIATION | 34 | 0.419365 | 1.455 | 0.033058 | 0.51294 |
| GOBP_RIBOSOMAL_SMALL_SUBUNIT_BIOGENESIS | 75 | 0.342797 | 1.435 | 0 | 0.561405 |
| GOBP_MEIOTIC_CHROMOSOME_SEGREGATION | 79 | 0.352121 | 1.432 | 0.029412 | 0.559077 |
| GOBP_NUCLEOTIDE_EXCISION_REPAIR_PREINCISION_COMPLEX_ASSEMBLY | 29 | 0.431428 | 1.409 | 0.06338 | 0.621257 |
| GOBP_MEIOSIS_I_CELL_CYCLE_PROCESS | 114 | 0.322392 | 1.409 | 0 | 0.611083 |
| GOBP_ERROR_FREE_TRANSLESION_SYNTHESIS | 22 | 0.458884 | 1.403 | 0.073171 | 0.619934 |
| GOBP_VIRAL_BUDDING_VIA_HOST_ESCRT_COMPLEX | 21 | 0.445777 | 1.402 | 0.06701 | 0.610978 |
| GOBP_MITOCHONDRIAL_ATP_SYNTHESIS_COUPLED_PROTON_TRANSPORT | 22 | 0.452661 | 1.39 | 0.064706 | 0.643631 |
| GOBP_MITOCHONDRIAL_RESPIRATORY_CHAIN_COMPLEX_ASSEMBLY | 100 | 0.329494 | 1.389 | 0.045455 | 0.634367 |
| GOBP_TRANSCRIPTION_INITIATION_FROM_RNA_POLYMERASE_I_PROMOTER | 38 | 0.398331 | 1.374 | 0.07563 | 0.674322 |
| GOBP_VIRION_ASSEMBLY | 40 | 0.381612 | 1.366 | 0.048544 | 0.690741 |
| GOBP_CLEAVAGE_INVOLVED_IN_RRNA_PROCESSING | 27 | 0.424912 | 1.365 | 0.102041 | 0.685319 |
| GOBP_MITOCHONDRIAL_GENE_EXPRESSION | 165 | 0.308435 | 1.364 | 0 | 0.677419 |
| GOBP_NUCLEAR_TRANSCRIBED_MRNA_CATABOLIC_PROCESS_EXONUCLEOLYTIC | 35 | 0.390137 | 1.359 | 0.076336 | 0.680755 |
| GOBP_TRANSLATIONAL_TERMINATION | 105 | 0.323288 | 1.353 | 0.045455 | 0.694186 |
| GOBP_NUCLEOTIDE_EXCISION_REPAIR_DNA_DUPLEX_UNWINDING | 22 | 0.418162 | 1.346 | 0.077348 | 0.706858 |
| GOBP_PURINE_NUCLEOSIDE_MONOPHOSPHATE_BIOSYNTHETIC_PROCESS | 23 | 0.421987 | 1.344 | 0.075269 | 0.70502 |
| GOBP_NEGATIVE_REGULATION_OF_UBIQUITIN_DEPENDENT_PROTEIN_CATABOLIC_P | 47 | 0.354597 | 1.341 | 0.049383 | 0.704875 |
| GOBP_NADH_DEHYDROGENASE_COMPLEX_ASSEMBLY | 65 | 0.335029 | 1.334 | 0.058824 | 0.720019 |
| GOBP_VIRAL_TRANSLATION | 16 | 0.466788 | 1.332 | 0.110577 | 0.715776 |
| GOBP_MITOCHONDRIAL_TRANSLATIONAL_TERMINATION | 89 | 0.3221 | 1.326 | 0.029412 | 0.730145 |
| GOBP_IRON_SULFUR_CLUSTER_ASSEMBLY | 23 | 0.423913 | 1.323 | 0.122699 | 0.730516 |
| GOBP_STRESS_GRANULE_ASSEMBLY | 23 | 0.421136 | 1.322 | 0.11976 | 0.723907 |
| GOBP_NUCLEAR_MEMBRANE_ORGANIZATION | 17 | 0.434996 | 1.313 | 0.120603 | 0.746481 |
| GOBP_GLYCEROPHOSPHOLIPID_CATABOLIC_PROCESS | 30 | 0.397984 | 1.297 | 0.102941 | 0.799544 |
| GOBP_TERMINATION_OF_RNA_POLYMERASE_I_TRANSCRIPTION | 31 | 0.395844 | 1.294 | 0.090226 | 0.803037 |
| GOBP_UBIQUITIN_DEPENDENT_PROTEIN_CATABOLIC_PROCESS_VIA_THE_MULTIVES | 23 | 0.401974 | 1.29 | 0.120219 | 0.806308 |
| GOBP_GLUTATHIONE_DERIVATIVE_METABOLIC_PROCESS | 20 | 0.408136 | 1.286 | 0.164835 | 0.815905 |
| GOBP_CHROMATIN_SILENCING | 61 | 0.324276 | 1.284 | 0.044776 | 0.812431 |
| GOBP_NUCLEOLAR_LARGE_RRNA_TRANSCRIPTION_BY_RNA_POLYMERASE_I | 17 | 0.420146 | 1.277 | 0.19598 | 0.829264 |
| GOBP_WATER_TRANSPORT | 18 | 0.431651 | 1.266 | 0.152632 | 0.863917 |
| GOBP_MEIOTIC_CELL_CYCLE_PROCESS | 170 | 0.250902 | 1.264 | 0 | 0.865319 |
| GOBP_TRANSCRIPTION_COUPLED_NUCLEOTIDE_EXCISION_REPAIR | 73 | 0.305419 | 1.261 | 0.019231 | 0.864866 |
| GOBP_GLYCOLIPID_CATABOLIC_PROCESS | 15 | 0.467568 | 1.258 | 0.170404 | 0.868834 |
| GOBP_FATTY_ACID_BETA_OXIDATION_USING_ACYL_COA_OXIDASE | 15 | 0.462103 | 1.257 | 0.173516 | 0.861936 |
| GOBP_MRNA_MODIFICATION | 24 | 0.404416 | 1.254 | 0.111801 | 0.862416 |
| GOBP_INNER_MITOCHONDRIAL_MEMBRANE_ORGANIZATION | 56 | 0.324226 | 1.253 | 0.108108 | 0.856699 |
| GOBP_POSITIVE_REGULATION_OF_GENE_EXPRESSION_EPIGENETIC | 55 | 0.337578 | 1.253 | 0.077922 | 0.847516 |
| GOBP_TRNA_5_END_PROCESSING | 16 | 0.433883 | 1.252 | 0.161137 | 0.839411 |
| GOBP_REGULATION_OF_MEIOTIC_CELL_CYCLE | 41 | 0.338655 | 1.251 | 0.096154 | 0.834124 |
| GOBP_PROTEIN_IMPORT_INTO_MITOCHONDRIAL_MATRIX | 20 | 0.406676 | 1.246 | 0.133721 | 0.84571 |
| GOBP_VIRAL_BUDDING | 26 | 0.390865 | 1.242 | 0.144737 | 0.855126 |
| GOBP_REGULATION_OF_CHROMOSOME_SEGREGATION | 82 | 0.297494 | 1.242 | 0.108696 | 0.845942 |
| GOBP_MITOTIC_SPINDLE_ASSEMBLY | 64 | 0.320056 | 1.24 | 0.081633 | 0.844375 |
| GOBP_TRANSCRIPTION_ELONGATION_FROM_RNA_POLYMERASE_I_PROMOTER | 30 | 0.37101 | 1.238 | 0.176471 | 0.84442 |
| GOBP_ATP_SYNTHESIS_COUPLED_PROTON_TRANSPORT | 27 | 0.379191 | 1.238 | 0.128834 | 0.835639 |
| GOBP_AEROBIC_RESPIRATION | 85 | 0.305197 | 1.236 | 0.090909 | 0.832182 |
| GOBP_FORMATION_OF_CYTOPLASMIC_TRANSLATION_INITIATION_COMPLEX | 16 | 0.416935 | 1.233 | 0.197309 | 0.833801 |
| GOBP_HISTONE_H3_DEACETYLATION | 19 | 0.409103 | 1.232 | 0.201117 | 0.832208 |
| GOBP_RNA_INTERFERENCE | 16 | 0.425001 | 1.219 | 0.179487 | 0.879052 |
| GOBP_DOSAGE_COMPENSATION | 17 | 0.422855 | 1.219 | 0.205556 | 0.871234 |
| GOBP_CILIUM_OR_FLAGELLUM_DEPENDENT_CELL_MOTILITY | 103 | 0.283463 | 1.217 | 0.125 | 0.867774 |
| GOBP_PHOSPHOLIPID_CATABOLIC_PROCESS | 45 | 0.335875 | 1.215 | 0.095745 | 0.869291 |
| GOBP_RIBONUCLEOPROTEIN_COMPLEX_SUBUNIT_ORGANIZATION | 209 | 0.260099 | 1.208 | 0 | 0.888427 |
| GOBP_GLOBAL_GENOME_NUCLEOTIDE_EXCISION_REPAIR | 26 | 0.36958 | 1.208 | 0.19708 | 0.880328 |
| GOBP_DNA_TEMPLATED_TRANSCRIPTION_TERMINATION | 75 | 0.287094 | 1.206 | 0.055556 | 0.878865 |
| GOBP_HISTONE_DEACETYLATION | 75 | 0.29838 | 1.201 | 0.112903 | 0.895624 |
| GOBP_ADIPOSE_TISSUE_DEVELOPMENT | 36 | 0.353439 | 1.199 | 0.135417 | 0.894222 |
| GOBP_PHOSPHATIDYLCHOLINE_CATABOLIC_PROCESS | 15 | 0.425689 | 1.198 | 0.24 | 0.887441 |
| GOBP_HISTONE_DEUBIQUITINATION | 23 | 0.384456 | 1.197 | 0.20915 | 0.883228 |
| GOBP_REGULATION_OF_ANDROGEN_RECEPTOR_SIGNALING_PATHWAY | 25 | 0.370836 | 1.197 | 0.182432 | 0.876297 |
| GOBP_REGULATION_OF_EXECUTION_PHASE_OF_APOPTOSIS | 26 | 0.379286 | 1.197 | 0.205479 | 0.870447 |
| GOBP_MATURATION_OF_5_8S_RRNA_FROM_TRICISTRONIC_RRNA_TRANSCRIPT_SSU | 24 | 0.383643 | 1.196 | 0.172043 | 0.864663 |
| GOBP_RNA_SPLICING_VIA_ENDONUCLEOLYTIC_CLEAVAGE_AND_LIGATION | 16 | 0.41842 | 1.195 | 0.233607 | 0.860011 |
| GOBP_SARCOMERE_ORGANIZATION | 31 | 0.343093 | 1.192 | 0.173554 | 0.868233 |
| GOBP_INTERSTRAND_CROSS_LINK_REPAIR | 56 | 0.308019 | 1.191 | 0.137931 | 0.864587 |
| GOBP_NEGATIVE_REGULATION_OF_GENE_EXPRESSION_EPIGENETIC | 98 | 0.277043 | 1.183 | 0.076923 | 0.887754 |
| GOBP_NUCLEOTIDE_EXCISION_REPAIR | 105 | 0.274294 | 1.182 | 0.130435 | 0.887112 |
| GOBP_HISTONE_H2A_ACETYLATION | 17 | 0.407663 | 1.173 | 0.259887 | 0.916969 |
| GOBP_LOW_DENSITY_LIPOPROTEIN_PARTICLE_CLEARANCE | 31 | 0.344591 | 1.171 | 0.177778 | 0.917738 |
| GOBP_SPERM_MOTILITY | 78 | 0.291757 | 1.171 | 0.166667 | 0.910675 |
| GOBP_CELLULAR_RESPONSE_TO_LOW_DENSITY_LIPOPROTEIN_PARTICLE_STIMULU | 18 | 0.381329 | 1.169 | 0.215054 | 0.91007 |
| GOBP_MEMBRANE_LIPID_CATABOLIC_PROCESS | 35 | 0.334036 | 1.169 | 0.185185 | 0.904504 |
| GOBP_SPINDLE_LOCALIZATION | 50 | 0.311099 | 1.169 | 0.151163 | 0.897982 |
| GOBP_OOGENESIS | 74 | 0.2822 | 1.166 | 0.108108 | 0.899595 |
| GOBP_INTESTINAL_ABSORPTION | 31 | 0.34093 | 1.164 | 0.228814 | 0.902365 |
| GOBP_MATURATION_OF_LSU_RRNA_FROM_TRICISTRONIC_RRNA_TRANSCRIPT_SSU | 15 | 0.418959 | 1.16 | 0.22439 | 0.913184 |
| GOBP_FLUID_TRANSPORT | 27 | 0.348896 | 1.151 | 0.220126 | 0.943931 |
| GOBP_GLYCEROLIPID_CATABOLIC_PROCESS | 61 | 0.305438 | 1.15 | 0.183099 | 0.940524 |
| GOBP_RIBOSOMAL_SMALL_SUBUNIT_ASSEMBLY | 21 | 0.373996 | 1.147 | 0.242938 | 0.946079 |
| GOBP_ENDONUCLEOLYTIC_CLEAVAGE_INVOLVED_IN_RRNA_PROCESSING | 15 | 0.414318 | 1.145 | 0.278947 | 0.948189 |
| GOBP_BASE_CONVERSION_OR_SUBSTITUTION_EDITING | 17 | 0.400364 | 1.136 | 0.276923 | 0.981292 |
| GOBP_PROTEIN_TRANSMEMBRANE_IMPORT_INTO_INTRACELLULAR_ORGANELLE | 36 | 0.327321 | 1.135 | 0.207407 | 0.979571 |
| GOBP_MEIOTIC_CELL_CYCLE | 224 | 0.212929 | 1.13 | 0 | 0.991994 |
| GOBP_PHOSPHATIDYLGLYCEROL_ACYL_CHAIN_REMODELING | 15 | 0.410387 | 1.129 | 0.299559 | 0.989201 |
| GOBP_RESPONSE_TO_PLATELET_DERIVED_GROWTH_FACTOR | 19 | 0.384593 | 1.119 | 0.255208 | 1 |
| GOBP_CERAMIDE_CATABOLIC_PROCESS | 20 | 0.362721 | 1.117 | 0.264865 | 1 |

**GSEA Negative.** Hematopoietic/Immune pathway are highlighted green.

| Gene sets | SIZE | ES | NES | NOM p-val | FDR q-val |
|---|---|---|---|---|---|
| GOBP_HUMORAL_IMMUNE_RESPONSE_MEDIATED_BY_CIRCULATING_IMMUNO | 122 | -0.882296 | -2.490951 | 0 | 0 |
| GOBP_COMPLEMENT_ACTIVATION | 136 | -0.845277 | -2.450629 | 0 | 0 |
| GOBP_B_CELL_MEDIATED_IMMUNITY | 188 | -0.831756 | -2.42813 | 0 | 0 |
| GOBP_PHAGOCYTOSIS_RECOGNITION | 75 | -0.872486 | -2.41998 | 0 | 0 |
| GOBP_B_CELL_RECEPTOR_SIGNALING_PATHWAY | 112 | -0.838893 | -2.401427 | 0 | 0 |
| GOBP_REGULATION_OF_HUMORAL_IMMUNE_RESPONSE | 112 | -0.836795 | -2.398314 | 0 | 0 |
| GOBP_POSITIVE_REGULATION_OF_B_CELL_ACTIVATION | 126 | -0.831579 | -2.395378 | 0 | 0 |
| GOBP_REGULATION_OF_COMPLEMENT_ACTIVATION | 93 | -0.846296 | -2.382944 | 0 | 0 |
| GOBP_FC_RECEPTOR_MEDIATED_STIMULATORY_SIGNALING_PATHWAY | 132 | -0.808262 | -2.312371 | 0 | 0 |
| GOBP_MEMBRANE_INVAGINATION | 123 | -0.806589 | -2.310302 | 0 | 0 |
| GOBP_REGULATION_OF_B_CELL_ACTIVATION | 169 | -0.792107 | -2.301625 | 0 | 0 |
| GOBP_IMMUNOGLOBULIN_PRODUCTION | 146 | -0.794577 | -2.295804 | 0 | 0 |
| GOBP_FC_EPSILON_RECEPTOR_SIGNALING_PATHWAY | 155 | -0.787072 | -2.278247 | 0 | 0 |
| GOBP_LYMPHOCYTE_MEDIATED_IMMUNITY | 307 | -0.775904 | -2.267775 | 0 | 0 |
| GOBP_HUMORAL_IMMUNE_RESPONSE | 268 | -0.765267 | -2.24661 | 0 | 0 |
| GOBP_ADAPTIVE_IMMUNE_RESPONSE_BASED_ON_SOMATIC_RECOMBINATIO | 317 | -0.755123 | -2.235376 | 0 | 0 |
| GOBP_ANTIGEN_RECEPTOR_MEDIATED_SIGNALING_PATHWAY | 295 | -0.738025 | -2.185732 | 0 | 0 |
| GOBP_FC_RECEPTOR_SIGNALING_PATHWAY | 224 | -0.738151 | -2.164808 | 0 | 0 |
| GOBP_B_CELL_ACTIVATION | 281 | -0.728171 | -2.138151 | 0 | 0 |
| GOBP_PHAGOCYTOSIS | 340 | -0.728022 | -2.135186 | 0 | 0 |
| GOBP_PHOTOTRANSDUCTION | 51 | -0.802032 | -2.134853 | 0 | 0 |
| GOBP_CELL_RECOGNITION | 193 | -0.734504 | -2.124121 | 0 | 0 |
| GOBP_DEFENSE_RESPONSE_TO_BACTERIUM | 234 | -0.720476 | -2.123265 | 0 | 0 |
| GOBP_IMMUNE_RESPONSE_REGULATING_SIGNALING_PATHWAY | 467 | -0.712192 | -2.113614 | 0 | 0 |
| GOBP_PRODUCTION_OF_MOLECULAR_MEDIATOR_OF_IMMUNE_RESPONSE | 227 | -0.719226 | -2.102047 | 0 | 0 |
| GOBP_REGULATION_OF_IMMUNE_EFFECTOR_PROCESS | 413 | -0.709093 | -2.094148 | 0 | 0 |
| GOBP_POSITIVE_REGULATION_OF_CELL_ACTIVATION | 371 | -0.704515 | -2.086086 | 0 | 0 |
| GOBP_LYMPHOCYTE_CHEMOTAXIS | 54 | -0.770718 | -2.075704 | 0 | 0 |
| GOBP_RETINA_HOMEOSTASIS | 76 | -0.749843 | -2.074338 | 0 | 0 |
| GOBP_DETECTION_OF_LIGHT_STIMULUS | 68 | -0.757831 | -2.072068 | 0 | 0 |
| GOBP_RECEPTOR_MEDIATED_ENDOCYTOSIS | 302 | -0.690327 | -2.065075 | 0 | 0 |
| GOBP_LYMPHOCYTE_MIGRATION | 107 | -0.722484 | -2.064157 | 0 | 0 |
| GOBP_REGULATION_OF_LYMPHOCYTE_ACTIVATION | 460 | -0.695879 | -2.061545 | 0 | 0 |
| GOBP_SENSORY_PERCEPTION_OF_LIGHT_STIMULUS | 201 | -0.692764 | -2.014986 | 0 | 0 |
| GOBP_POSITIVE_REGULATION_OF_CELL_KILLING | 57 | -0.735326 | -1.99362 | 0 | 0 |
| GOBP_LEUKOCYTE_MIGRATION | 461 | -0.670377 | -1.984774 | 0 | 0 |
| GOBP_RESPONSE_TO_CHEMOKINE | 84 | -0.705621 | -1.984205 | 0 | 0 |
| GOBP_NEUTROPHIL_CHEMOTAXIS | 92 | -0.704124 | -1.97754 | 0 | 0 |
| GOBP_POSITIVE_REGULATION_OF_NATURAL_KILLER_CELL_MEDIATED_CYTO | 23 | -0.820567 | -1.971033 | 0 | 0 |
| GOBP_MONONUCLEAR_CELL_MIGRATION | 173 | -0.680093 | -1.967871 | 0 | 0 |
| GOBP_PHOTOTRANSDUCTION_VISIBLE_LIGHT | 33 | -0.775754 | -1.965725 | 0 | 0 |
| GOBP_T_CELL_MIGRATION | 61 | -0.719437 | -1.95523 | 0 | 3.36E-05 |
| GOBP_NEUTROPHIL_MIGRATION | 111 | -0.673122 | -1.946896 | 0 | 6.62E-05 |
| GOBP_MONOCYTE_CHEMOTAXIS | 57 | -0.723552 | -1.946513 | 0 | 8.15E-05 |
| GOBP_NATURAL_KILLER_CELL_MEDIATED_IMMUNITY | 61 | -0.713612 | -1.945122 | 0 | 8.02E-05 |
| GOBP_POSITIVE_REGULATION_OF_NATURAL_KILLER_CELL_MEDIATED_IMMUI | 28 | -0.778846 | -1.942867 | 0 | 7.89E-05 |
| GOBP_REGULATORY_T_CELL_DIFFERENTIATION | 33 | -0.759841 | -1.931521 | 0 | 7.76E-05 |
| GOBP_REGULATION_OF_LYMPHOCYTE_MEDIATED_IMMUNITY | 144 | -0.6627 | -1.925801 | 0 | 1.07E-04 |
| GOBP_REGULATION_OF_CELL_KILLING | 90 | -0.682471 | -1.924987 | 0 | 1.05E-04 |
| GOBP_POSITIVE_REGULATION_OF_T_CELL_PROLIFERATION | 92 | -0.683341 | -1.911445 | 0 | 1.18E-04 |
| GOBP_REGULATION_OF_SYNAPTIC_PLASTICITY | 172 | -0.650948 | -1.906912 | 0 | 1.46E-04 |
| GOBP_PHOTORECEPTOR_CELL_DIFFERENTIATION | 59 | -0.697049 | -1.906308 | 0 | 1.44E-04 |
| GOBP_POSITIVE_T_CELL_SELECTION | 34 | -0.755243 | -1.90453 | 0 | 1.56E-04 |
| GOBP_REGULATION_OF_T_CELL_DIFFERENTIATION | 136 | -0.65748 | -1.898382 | 0 | 1.95E-04 |
| GOBP_PHOTORECEPTOR_CELL_DEVELOPMENT | 45 | -0.716469 | -1.89794 | 0 | 1.92E-04 |
| GOBP_REGULATION_OF_PHOSPHOLIPASE_C_ACTIVITY | 44 | -0.724827 | -1.897194 | 0 | 1.90E-04 |
| GOBP_NEURAL_RETINA_DEVELOPMENT | 67 | -0.695499 | -1.897028 | 0 | 1.87E-04 |
| GOBP_GRANULOCYTE_CHEMOTAXIS | 113 | -0.664263 | -1.896953 | 0 | 1.85E-04 |
| GOBP_T_CELL_DIFFERENTIATION | 230 | -0.648028 | -1.891189 | 0 | 1.95E-04 |
| GOBP_REGULATION_OF_T_CELL_ACTIVATION | 304 | -0.638807 | -1.890324 | 0 | 1.92E-04 |
| GOBP_DETECTION_OF_ABIOTIC_STIMULUS | 130 | -0.651476 | -1.889358 | 0 | 1.90E-04 |
| GOBP_LEUKOCYTE_CHEMOTAXIS | 204 | -0.649075 | -1.88834 | 0 | 2.00E-04 |
| GOBP_NEGATIVE_REGULATION_OF_LEUKOCYTE_MEDIATED_IMMUNITY | 50 | -0.708902 | -1.886779 | 0 | 1.97E-04 |
| GOBP_POSITIVE_REGULATION_OF_LEUKOCYTE_CELL_CELL_ADHESION | 222 | -0.64374 | -1.88374 | 0 | 2.31E-04 |
| GOBP_POSITIVE_REGULATION_OF_LYMPHOCYTE_DIFFERENTIATION | 95 | -0.667496 | -1.881132 | 0 | 2.52E-04 |
| GOBP_POSITIVE_REGULATION_OF_LEUKOCYTE_MEDIATED_IMMUNITY | 116 | -0.664691 | -1.880488 | 0 | 2.73E-04 |
| GOBP_REGULATION_OF_NMDA_RECEPTOR_ACTIVITY | 34 | -0.747393 | -1.879616 | 0 | 2.93E-04 |
| GOBP_INTERLEUKIN_12_PRODUCTION | 56 | -0.697409 | -1.879147 | 0 | 2.90E-04 |
| GOBP_LEUKOCYTE_MEDIATED_CYTOTOXICITY | 100 | -0.666911 | -1.878622 | 0 | 2.86E-04 |
| GOBP_REGULATION_OF_LEUKOCYTE_MEDIATED_CYTOTOXICITY | 72 | -0.678089 | -1.878215 | 0 | 2.94E-04 |
| GOBP_POSITIVE_REGULATION_OF_IMMUNE_EFFECTOR_PROCESS | 199 | -0.63914 | -1.870148 | 0 | 3.69E-04 |
| GOBP_T_CELL_PROLIFERATION | 182 | -0.640175 | -1.86938 | 0 | 3.98E-04 |
| GOBP_REGULATION_OF_LYMPHOCYTE_DIFFERENTIATION | 163 | -0.645947 | -1.86902 | 0 | 3.94E-04 |
| GOBP_POSITIVE_REGULATION_OF_HUMORAL_IMMUNE_RESPONSE | 16 | -0.83562 | -1.868866 | 0 | 3.89E-04 |
| GOBP_EYE_PHOTORECEPTOR_CELL_DEVELOPMENT | 31 | -0.750936 | -1.868584 | 0 | 4.07E-04 |
| GOBP_INTERFERON_GAMMA_PRODUCTION | 98 | -0.658713 | -1.868355 | 0 | 4.02E-04 |
| GOBP_REGULATION_OF_LEUKOCYTE_MEDIATED_IMMUNITY | 194 | -0.634821 | -1.867891 | 0 | 3.98E-04 |
| GOBP_NEUROINFLAMMATORY_RESPONSE | 35 | -0.722997 | -1.866378 | 0 | 4.45E-04 |
| GOBP_RHODOPSIN_MEDIATED_SIGNALING_PATHWAY | 28 | -0.757748 | -1.861973 | 0 | 5.22E-04 |
| GOBP_B_CELL_PROLIFERATION | 75 | -0.672534 | -1.860152 | 0 | 5.37E-04 |
| GOBP_MEMBRANE_DEPOLARIZATION_DURING_ACTION_POTENTIAL | 34 | -0.730617 | -1.859193 | 0 | 5.52E-04 |
| GOBP_RESPIRATORY_BURST | 34 | -0.746317 | -1.858618 | 0 | 5.56E-04 |
| GOBP_NERVE_DEVELOPMENT | 64 | -0.67935 | -1.855756 | 0 | 5.70E-04 |
| GOBP_RESPONSE_TO_INTERFERON_GAMMA | 184 | -0.633242 | -1.852787 | 0 | 5.74E-04 |
| GOBP_T_CELL_SELECTION | 46 | -0.699698 | -1.852775 | 0 | 5.68E-04 |
| GOBP_CELL_KILLING | 139 | -0.643801 | -1.851762 | 0 | 5.72E-04 |
| GOBP_PHOTORECEPTOR_CELL_MAINTENANCE | 44 | -0.710275 | -1.851669 | 0 | 5.66E-04 |
| GOBP_REGULATION_OF_LYMPHOCYTE_MIGRATION | 60 | -0.6875 | -1.851072 | 0 | 5.89E-04 |
| GOBP_T_CELL_ACTIVATION | 430 | -0.626983 | -1.849978 | 0 | 5.92E-04 |
| GOBP_ALPHA_BETA_T_CELL_PROLIFERATION | 34 | -0.726344 | -1.849046 | 0 | 6.23E-04 |
| GOBP_T_CELL_CHEMOTAXIS | 24 | -0.780231 | -1.848992 | 0 | 6.26E-04 |
| GOBP_LEUKOCYTE_CELL_CELL_ADHESION | 333 | -0.622437 | -1.848323 | 0 | 6.48E-04 |
| GOBP_SYNAPTIC_TRANSMISSION_GLUTAMATERGIC | 85 | -0.65901 | -1.847532 | 0 | 6.59E-04 |
| GOBP_ALPHA_BETA_T_CELL_ACTIVATION | 137 | -0.638485 | -1.845431 | 0 | 7.24E-04 |
| GOBP_REGULATION_OF_MONONUCLEAR_CELL_MIGRATION | 106 | -0.652123 | -1.845068 | 0 | 7.26E-04 |
| GOBP_POSITIVE_REGULATION_OF_LEUKOCYTE_PROLIFERATION | 135 | -0.641115 | -1.844357 | 0 | 7.28E-04 |
| GOBP_DENDRITIC_CELL_CHEMOTAXIS | 23 | -0.778803 | -1.841936 | 0 | 7.47E-04 |
| GOBP_REGULATION_OF_T_CELL_MEDIATED_IMMUNITY | 70 | -0.665747 | -1.841694 | 0 | 7.41E-04 |
| GOBP_NATURAL_KILLER_CELL_ACTIVATION | 59 | -0.683505 | -1.840297 | 0 | 7.59E-04 |
| GOBP_GRANULOCYTE_MIGRATION | 135 | -0.639341 | -1.838521 | 0 | 7.70E-04 |
| GOBP_T_CELL_MEDIATED_IMMUNITY | 93 | -0.650599 | -1.838408 | 0 | 7.71E-04 |
| GOBP_RETINA_DEVELOPMENT_IN_CAMERA_TYPE_EYE | 141 | -0.640624 | -1.836606 | 0 | 7.73E-04 |
| GOBP_LEUKOCYTE_MIGRATION_INVOLVED_IN_INFLAMMATORY_RESPONSE | 17 | -0.806293 | -1.83592 | 0 | 7.74E-04 |
| GOBP_REGULATION_OF_T_CELL_MIGRATION | 40 | -0.694346 | -1.832001 | 0 | 8.57E-04 |
| GOBP_REGULATION_OF_LYMPHOCYTE_CHEMOTAXIS | 25 | -0.763878 | -1.831543 | 0 | 8.66E-04 |
| GOBP_ACTIVATION_OF_PHOSPHOLIPASE_C_ACTIVITY | 30 | -0.73913 | -1.831461 | 0 | 8.59E-04 |
| GOBP_CELL_CHEMOTAXIS | 268 | -0.622963 | -1.830712 | 0 | 8.67E-04 |
| GOBP_REGULATION_OF_LEUKOCYTE_PROLIFERATION | 221 | -0.628463 | -1.825532 | 0 | 9.70E-04 |
| GOBP_POSITIVE_REGULATION_OF_INTERFERON_GAMMA_PRODUCTION | 59 | -0.677099 | -1.823421 | 0 | 0.001001 |
| GOBP_POSITIVE_REGULATION_OF_LYMPHOCYTE_MIGRATION | 36 | -0.711893 | -1.822626 | 0 | 0.001001 |
| GOBP_NEGATIVE_REGULATION_OF_IMMUNE_EFFECTOR_PROCESS | 115 | -0.643456 | -1.821415 | 0 | 0.001031 |
| GOBP_T_CELL_RECEPTOR_SIGNALING_PATHWAY | 198 | -0.617021 | -1.820329 | 0 | 0.001061 |
| GOBP_NEUROTRANSMITTER_RECEPTOR_LOCALIZATION_TO_POSTSYNAPTIC_ | 20 | -0.764886 | -1.820314 | 0.001222 | 0.001053 |
| GOBP_POSITIVE_REGULATION_OF_PHAGOCYTOSIS | 63 | -0.66815 | -1.81974 | 0 | 0.001059 |
| GOBP_REGULATION_OF_PRODUCTION_OF_MOLECULAR_MEDIATOR_OF_IMMU | 132 | -0.623034 | -1.816576 | 0 | 0.001147 |
| GOBP_RETINA_MORPHOGENESIS_IN_CAMERA_TYPE_EYE | 57 | -0.673816 | -1.816429 | 0 | 0.001138 |
| GOBP_EYE_PHOTORECEPTOR_CELL_DIFFERENTIATION | 42 | -0.690502 | -1.816307 | 0 | 0.001137 |
| GOBP_POSITIVE_REGULATION_OF_ALPHA_BETA_T_CELL_ACTIVATION | 62 | -0.663821 | -1.815399 | 0 | 0.001158 |
| GOBP_NEGATIVE_REGULATION_OF_LYMPHOCYTE_MEDIATED_IMMUNITY | 42 | -0.696224 | -1.813706 | 0 | 0.001163 |
| GOBP_MYELOID_LEUKOCYTE_MIGRATION | 198 | -0.620625 | -1.813535 | 0 | 0.001162 |
| GOBP_REGULATION_OF_T_CELL_RECEPTOR_SIGNALING_PATHWAY | 40 | -0.705442 | -1.813212 | 0 | 0.00116 |
| GOBP_POSITIVE_REGULATION_OF_ALPHA_BETA_T_CELL_PROLIFERATION | 20 | -0.777516 | -1.812023 | 0.00123 | 0.001201 |
| GOBP_DENDRITIC_CELL_MIGRATION | 28 | -0.747291 | -1.811501 | 0 | 0.001213 |
| GOBP_REGULATION_OF_NATURAL_KILLER_CELL_MEDIATED_IMMUNITY | 44 | -0.695122 | -1.807556 | 0 | 0.001378 |
| GOBP_CELLULAR_DEFENSE_RESPONSE | 46 | -0.692676 | -1.807548 | 0 | 0.001368 |
| GOBP_SEQUESTERING_OF_CALCIUM_ION | 111 | -0.635337 | -1.807262 | 0 | 0.001359 |
| GOBP_RENAL_FILTRATION | 24 | -0.747318 | -1.807253 | 0 | 0.001349 |
| GOBP_REGULATION_OF_ALPHA_BETA_T_CELL_ACTIVATION | 93 | -0.648249 | -1.806171 | 0 | 0.0014 |
| GOBP_NEGATIVE_REGULATION_OF_IMMUNE_SYSTEM_PROCESS | 362 | -0.60625 | -1.805963 | 0 | 0.001397 |
| GOBP_REGULATION_OF_B_CELL_PROLIFERATION | 57 | -0.665975 | -1.805866 | 0 | 0.001387 |
| GOBP_LEUKOCYTE_PROLIFERATION | 274 | -0.615847 | -1.803718 | 0 | 0.001457 |
| GOBP_POSITIVE_REGULATION_OF_CELL_CELL_ADHESION | 261 | -0.615457 | -1.803196 | 0 | 0.00146 |
| GOBP_RESPONSE_TO_PROTOZOAN | 20 | -0.781188 | -1.802942 | 0 | 0.00147 |
| GOBP_REGULATION_OF_LEUKOCYTE_DIFFERENTIATION | 253 | -0.613435 | -1.800419 | 0 | 0.001544 |
| GOBP_MONONUCLEAR_CELL_DIFFERENTIATION | 370 | -0.610384 | -1.799262 | 0 | 0.001572 |
| GOBP_BONE_RESORPTION | 57 | -0.672086 | -1.799138 | 0 | 0.001562 |
| GOBP_REGULATION_OF_CATION_CHANNEL_ACTIVITY | 167 | -0.615607 | -1.798068 | 0 | 0.001602 |
| GOBP_LYMPHOCYTE_COSTIMULATION | 58 | -0.667727 | -1.797306 | 0 | 0.001654 |
| GOBP_CYTOKINE_PRODUCTION_INVOLVED_IN_IMMUNE_RESPONSE | 89 | -0.640915 | -1.796261 | 0 | 0.0017 |
| GOBP_GAMMA_DELTA_T_CELL_ACTIVATION | 18 | -0.782779 | -1.794008 | 0 | 0.001789 |
| GOBP_L_GLUTAMATE_TRANSMEMBRANE_TRANSPORT | 24 | -0.753106 | -1.791003 | 0 | 0.001857 |
| GOBP_POSITIVE_REGULATION_OF_HEMOPOIESIS | 140 | -0.62221 | -1.789357 | 0 | 0.001907 |
| GOBP_RESPONSE_TO_AUDITORY_STIMULUS | 21 | -0.764983 | -1.788973 | 0 | 0.001907 |
| GOBP_NEGATIVE_REGULATION_OF_INTERLEUKIN_12_PRODUCTION | 17 | -0.787162 | -1.788026 | 0 | 0.001913 |
| GOBP_NEGATIVE_REGULATION_OF_LEUKOCYTE_MIGRATION | 42 | -0.683243 | -1.78742 | 0 | 0.001949 |
| GOBP_CELL_CELL_ADHESION_MEDIATED_BY_INTEGRIN | 17 | -0.785959 | -1.784679 | 0 | 0.002015 |
| GOBP_POSITIVE_REGULATION_OF_CELL_ADHESION_MEDIATED_BY_INTEGRIN | 21 | -0.759309 | -1.784374 | 0 | 0.00202 |
| GOBP_LONG_TERM_SYNAPTIC_POTENTIATION | 79 | -0.6445 | -1.783608 | 0 | 0.002079 |

| | | | | | |
|---|---|---|---|---|---|
| GOBP_PHOSPHATIDYLCHOLINE_ACYL_CHAIN_REMODELING | 26 | 0.337511 | 1.113 | 0.248175 | 1 |
| GOBP_KERATINOCYTE_MIGRATION | 16 | 0.396331 | 1.113 | 0.293839 | 1 |
| GOBP_MICROTUBULE_ANCHORING | 26 | 0.343511 | 1.112 | 0.285714 | 1 |
| GOBP_NEGATIVE_REGULATION_OF_PROTEOLYSIS_INVOLVED_IN_CELLULAR_PROTEI | 62 | 0.280961 | 1.108 | 0.210526 | 1 |
| GOBP_NUCLEOBASE_BIOSYNTHETIC_PROCESS | 18 | 0.370155 | 1.106 | 0.305556 | 1 |
| GOBP_REGULATION_OF_MEGAKARYOCYTE_DIFFERENTIATION | 71 | 0.268393 | 1.106 | 0.240741 | 1 |
| GOBP_RESPONSE_TO_HEPATOCYTE_GROWTH_FACTOR | 17 | 0.375723 | 1.106 | 0.288462 | 1 |
| GOBP_RESPONSE_TO_STEROL_DEPLETION | 18 | 0.381186 | 1.104 | 0.299517 | 1 |
| GOBP_ERBB2_SIGNALING_PATHWAY | 31 | 0.333339 | 1.103 | 0.28777 | 1 |
| GOBP_NEGATIVE_REGULATION_OF_NUCLEAR_DIVISION | 52 | 0.282038 | 1.102 | 0.231707 | 1 |
| GOBP_REGULATION_OF_CENTROSOME_CYCLE | 48 | 0.298432 | 1.101 | 0.225806 | 1 |
| GOBP_NCRNA_TRANSCRIPTION | 108 | 0.25483 | 1.096 | 0.238095 | 1 |
| GOBP_PEROXISOME_PROLIFERATOR_ACTIVATED_RECEPTOR_SIGNALING_PATHWAY | 20 | 0.357444 | 1.094 | 0.310976 | 1 |
| GOBP_CELLULAR_RESPONSE_TO_ARSENIC_CONTAINING_SUBSTANCE | 18 | 0.373291 | 1.091 | 0.321101 | 1 |
| GOBP_REGULATION_OF_CALCINEURIN_MEDIATED_SIGNALING | 30 | 0.316063 | 1.09 | 0.285714 | 1 |
| GOBP_RRNA_MODIFICATION | 40 | 0.301556 | 1.088 | 0.298246 | 1 |
| GOBP_NUCLEOTIDE_EXCISION_REPAIR_DNA_DAMAGE_RECOGNITION | 23 | 0.352343 | 1.086 | 0.281768 | 1 |
| GOBP_REGULATION_OF_PROTEIN_SUMOYLATION | 22 | 0.349439 | 1.086 | 0.324176 | 1 |
| GOBP_CELL_DIFFERENTIATION_INVOLVED_IN_EMBRYONIC_PLACENTA_DEVELOPMEI | 22 | 0.352741 | 1.086 | 0.305085 | 1 |
| GOBP_LACTATION | 42 | 0.290138 | 1.084 | 0.26 | 1 |
| GOBP_REGULATION_OF_CELL_CELL_ADHESION_MEDIATED_BY_CADHERIN | 20 | 0.354358 | 1.083 | 0.333333 | 1 |
| GOBP_G0_TO_G1_TRANSITION | 44 | 0.293653 | 1.08 | 0.298969 | 1 |
| GOBP_RNA_CAPPING | 34 | 0.309303 | 1.078 | 0.316547 | 1 |
| GOBP_SHORT_CHAIN_FATTY_ACID_METABOLIC_PROCESS | 17 | 0.37378 | 1.077 | 0.333333 | 1 |
| GOBP_GLUTAMINE_METABOLIC_PROCESS | 22 | 0.328566 | 1.075 | 0.362069 | 1 |
| GOBP_TERMINATION_OF_RNA_POLYMERASE_II_TRANSCRIPTION | 35 | 0.304552 | 1.074 | 0.310606 | 1 |
| GOBP_POSITIVE_REGULATION_OF_VIRAL_TRANSCRIPTION | 26 | 0.328466 | 1.072 | 0.315068 | 1 |
| GOBP_LINOLEIC_ACID_METABOLIC_PROCESS | 20 | 0.335235 | 1.071 | 0.354651 | 1 |
| GOBP_MULTIVESICULAR_BODY_ORGANIZATION | 31 | 0.308105 | 1.068 | 0.335664 | 1 |
| GOBP_TRNA_METABOLIC_PROCESS | 177 | 0.237894 | 1.067 | 0.166667 | 1 |
| GOBP_REGULATION_OF_VIRAL_TRANSCRIPTION | 42 | 0.295845 | 1.067 | 0.343137 | 1 |
| GOBP_RRNA_CATABOLIC_PROCESS | 19 | 0.35648 | 1.063 | 0.329949 | 1 |
| GOBP_REGULATION_OF_URINE_VOLUME | 19 | 0.349576 | 1.061 | 0.377451 | 1 |
| GOBP_CHROMATIN_ORGANIZATION_INVOLVED_IN_REGULATION_OF_TRANSCRIPTION | 124 | 0.243751 | 1.06 | 0.428571 | 1 |
| GOBP_RIBONUCLEOSIDE_MONOPHOSPHATE_BIOSYNTHETIC_PROCESS | 33 | 0.296904 | 1.057 | 0.377049 | 1 |
| GOBP_INNER_DYNEIN_ARM_ASSEMBLY | 15 | 0.378935 | 1.051 | 0.377682 | 1 |
| GOBP_FEMALE_GAMETE_GENERATION | 115 | 0.233237 | 1.05 | 0.363636 | 1 |
| GOBP_PHOSPHATIDIC_ACID_METABOLIC_PROCESS | 47 | 0.284322 | 1.045 | 0.333333 | 1 |
| GOBP_MATURATION_OF_SSU_RRNA | 50 | 0.279512 | 1.043 | 0.388889 | 1 |
| GOBP_TRANSCRIPTION_BY_RNA_POLYMERASE_I | 68 | 0.258775 | 1.041 | 0.301887 | 1 |
| GOBP_MATURATION_OF_SSU_RRNA_FROM_TRICISTRONIC_RRNA_TRANSCRIPT_SSU | 37 | 0.294198 | 1.041 | 0.390625 | 1 |
| GOBP_ATTACHMENT_OF_SPINDLE_MICROTUBULES_TO_KINETOCHORE | 35 | 0.293103 | 1.04 | 0.366412 | 1 |
| GOBP_CILIUM_MOVEMENT | 137 | 0.240409 | 1.039 | 0.411765 | 1 |
| GOBP_LACTATE_METABOLIC_PROCESS | 15 | 0.370821 | 1.038 | 0.401869 | 1 |
| GOBP_PHOSPHATIDYLGLYCEROL_METABOLIC_PROCESS | 34 | 0.297661 | 1.036 | 0.390728 | 1 |
| GOBP_NEUTRAL_LIPID_CATABOLIC_PROCESS | 36 | 0.292274 | 1.036 | 0.338583 | 1 |
| GOBP_REGULATION_OF_CENTRIOLE_REPLICATION | 21 | 0.33403 | 1.035 | 0.4125 | 1 |
| GOBP_NUCLEOTIDE_EXCISION_REPAIR_PREINCISION_COMPLEX_STABILIZATION | 21 | 0.344671 | 1.032 | 0.383784 | 1 |
| GOBP_AMINO_ACID_ACTIVATION | 49 | 0.271358 | 1.03 | 0.357895 | 1 |
| GOBP_REGULATION_OF_TELOMERASE_ACTIVITY | 47 | 0.26916 | 1.03 | 0.404762 | 1 |
| GOBP_ATP_BIOSYNTHETIC_PROCESS | 55 | 0.264775 | 1.029 | 0.364706 | 1 |
| GOBP_PHOSPHATIDYLINOSITOL_ACYL_CHAIN_REMODELING | 15 | 0.373609 | 1.029 | 0.400862 | 1 |
| GOBP_REGULATION_OF_CHROMATIN_SILENCING | 15 | 0.357635 | 1.021 | 0.405622 | 1 |
| GOBP_TRANSLATIONAL_ELONGATION | 133 | 0.237748 | 1.021 | 0.333333 | 1 |
| GOBP_PROTEIN_CONTAINING_COMPLEX_REMODELING | 29 | 0.304444 | 1.018 | 0.410072 | 1 |
| GOBP_CENTROSOME_SEPARATION | 15 | 0.359667 | 1.018 | 0.385714 | 1 |
| GOBP_POSITIVE_REGULATION_OF_CHROMOSOME_SEPARATION | 17 | 0.355807 | 1.017 | 0.419048 | 1 |
| GOBP_PROTEIN_TARGETING_TO_MEMBRANE | 201 | 0.21964 | 1.016 | 0.666667 | 1 |
| GOBP_CYTOKINETIC_PROCESS | 38 | 0.285899 | 1.014 | 0.433628 | 1 |
| GOBP_FRUCTOSE_METABOLIC_PROCESS | 15 | 0.356291 | 1.014 | 0.419192 | 1 |
| GOBP_DNA_UNWINDING_INVOLVED_IN_DNA_REPLICATION | 16 | 0.360069 | 1.014 | 0.4 | 1 |
| GOBP_MRNA_METHYLATION | 16 | 0.353183 | 1.01 | 0.437811 | 1 |
| GOBP_PINOCYTOSIS | 22 | 0.32144 | 1.004 | 0.442529 | 1 |
| GOBP_REGULATION_OF_EXTRACELLULAR_MATRIX_ASSEMBLY | 15 | 0.351047 | 1.001 | 0.422535 | 1 |
| GOBP_REGULATION_OF_REPRODUCTIVE_PROCESS | 128 | 0.231011 | 1.001 | 0.4 | 1 |
| GOBP_MIDBODY_ABSCISSION | 17 | 0.354005 | 1 | 0.458937 | 1 |
| GOBP_REGULATION_OF_SPINDLE_ASSEMBLY | 25 | 0.301253 | 0.999 | 0.447368 | 1 |
| GOBP_OUTER_DYNEIN_ARM_ASSEMBLY | 18 | 0.341387 | 0.999 | 0.461538 | 1 |
| GOBP_RENAL_ABSORPTION | 15 | 0.357336 | 0.997 | 0.432203 | 1 |
| GOBP_MRNA_CLEAVAGE | 22 | 0.304377 | 0.997 | 0.451977 | 1 |
| GOBP_HYDROGEN_PEROXIDE_CATABOLIC_PROCESS | 29 | 0.302173 | 0.996 | 0.503759 | 1 |
| GOBP_TRNA_PROCESSING | 128 | 0.22933 | 0.993 | 0.538462 | 1 |
| GOBP_S_ADENOSYLMETHIONINE_METABOLIC_PROCESS | 15 | 0.34408 | 0.991 | 0.453782 | 1 |
| GOBP_EXIT_FROM_MITOSIS | 26 | 0.303442 | 0.987 | 0.409639 | 1 |
| GOBP_MITOPHAGY | 24 | 0.313739 | 0.985 | 0.509554 | 1 |
| GOBP_REGULATION_OF_KERATINOCYTE_PROLIFERATION | 29 | 0.297934 | 0.985 | 0.417323 | 1 |
| GOBP_PHOSPHATIDYLSERINE_ACYL_CHAIN_REMODELING | 19 | 0.338386 | 0.985 | 0.421053 | 1 |
| GOBP_SPLICEOSOMAL_SNRNP_ASSEMBLY | 45 | 0.268891 | 0.983 | 0.472527 | 1 |
| GOBP_SPINDLE_ASSEMBLY | 113 | 0.213931 | 0.978 | 0.473684 | 1 |
| GOBP_REGULATION_OF_GENE_EXPRESSION_EPIGENETIC | 174 | 0.215802 | 0.976 | 0.5 | 1 |
| GOBP_O_GLYCAN_PROCESSING | 54 | 0.251325 | 0.974 | 0.5 | 1 |
| GOBP_RNA_MODIFICATION | 165 | 0.214422 | 0.972 | 0.636364 | 1 |
| GOBP_DNA_DAMAGE_RESPONSE_DETECTION_OF_DNA_DAMAGE | 38 | 0.277399 | 0.967 | 0.520325 | 1 |
| GOBP_MITOTIC_METAPHASE_PLATE_CONGRESSION | 50 | 0.260472 | 0.961 | 0.574713 | 1 |
| GOBP_NEGATIVE_REGULATION_OF_CHROMOSOME_ORGANIZATION | 87 | 0.224189 | 0.958 | 0.703704 | 1 |
| GOBP_POSITIVE_REGULATION_OF_HISTONE_DEACETYLATION | 17 | 0.339497 | 0.957 | 0.513761 | 1 |
| GOBP_HISTONE_H4_K5_ACETYLATION | 16 | 0.333097 | 0.954 | 0.481132 | 1 |
| GOBP_CELLULAR_RESPIRATION | 182 | 0.215544 | 0.95 | 0.75 | 1 |
| GOBP_CELLULAR_RESPONSE_TO_FLUID_SHEAR_STRESS | 17 | 0.312181 | 0.95 | 0.530387 | 1 |
| GOBP_AXONEMAL_DYNEIN_COMPLEX_ASSEMBLY | 32 | 0.26836 | 0.95 | 0.592593 | 1 |
| GOBP_REGULATION_OF_AUTOPHAGY_OF_MITOCHONDRION | 31 | 0.28013 | 0.95 | 0.528986 | 1 |
| GOBP_BASE_EXCISION_REPAIR | 39 | 0.263103 | 0.948 | 0.54955 | 1 |
| GOBP_ACETYL_COA_METABOLIC_PROCESS | 37 | 0.278798 | 0.946 | 0.495413 | 1 |
| GOBP_THIOESTER_METABOLIC_PROCESS | 99 | 0.222493 | 0.942 | 0.611111 | 1 |
| GOBP_CELL_AGGREGATION | 17 | 0.32422 | 0.942 | 0.549296 | 1 |
| GOBP_NEGATIVE_REGULATION_OF_DNA_RECOMBINATION | 41 | 0.254254 | 0.937 | 0.565657 | 1 |
| GOBP_OLIGOSACCHARIDE_BIOSYNTHETIC_PROCESS | 16 | 0.322575 | 0.936 | 0.517094 | 1 |
| GOBP_SNRNA_METABOLIC_PROCESS | 36 | 0.26407 | 0.933 | 0.612245 | 1 |
| GOBP_POSITIVE_REGULATION_OF_DNA_TEMPLATED_TRANSCRIPTION_ELONGATION | 26 | 0.28469 | 0.933 | 0.578947 | 1 |
| GOBP_NUCLEOSIDE_MONOPHOSPHATE_BIOSYNTHETIC_PROCESS | 41 | 0.262104 | 0.932 | 0.590476 | 1 |
| GOBP_ENTRAINMENT_OF_CIRCADIAN_CLOCK | 32 | 0.272345 | 0.931 | 0.587302 | 1 |
| GOBP_ESTABLISHMENT_OF_SPINDLE_ORIENTATION | 32 | 0.27388 | 0.93 | 0.553719 | 1 |
| GOBP_CHROMOSOME_LOCALIZATION | 77 | 0.229133 | 0.93 | 0.622222 | 1 |
| GOBP_PROTEIN_TRANSMEMBRANE_TRANSPORT | 60 | 0.233612 | 0.93 | 0.589744 | 1 |
| GOBP_NEGATIVE_REGULATION_OF_PROTEASOMAL_UBIQUITIN_DEPENDENT_PROTEI | 35 | 0.257493 | 0.93 | 0.683333 | 1 |
| GOBP_MITOCHONDRIAL_TRANSMEMBRANE_TRANSPORT | 105 | 0.211095 | 0.922 | 0.681818 | 1 |
| GOBP_RIBONUCLEOSIDE_TRIPHOSPHATE_BIOSYNTHETIC_PROCESS | 72 | 0.238156 | 0.921 | 0.660714 | 1 |
| GOBP_REGULATION_OF_DNA_METHYLATION | 18 | 0.31373 | 0.917 | 0.559406 | 1 |
| GOBP_MITOCHONDRIAL_RNA_METABOLIC_PROCESS | 47 | 0.24774 | 0.917 | 0.649485 | 1 |
| GOBP_CELLULAR_RESPONSE_TO_LIPOPROTEIN_PARTICLE_STIMULUS | 29 | 0.271281 | 0.917 | 0.647482 | 1 |
| GOBP_MITOTIC_CYTOKINETIC_PROCESS | 23 | 0.292758 | 0.915 | 0.6 | 1 |
| GOBP_HISTONE_H2A_MONOUBIQUITINATION | 17 | 0.318235 | 0.915 | 0.546667 | 1 |
| GOBP_SISTER_CHROMATID_SEGREGATION | 190 | 0.225794 | 0.913 | 0.8 | 1 |
| GOBP_NEGATIVE_REGULATION_OF_MRNA_PROCESSING | 29 | 0.276076 | 0.911 | 0.540741 | 1 |
| GOBP_DNA_METHYLATION_DEPENDENT_HETEROCHROMATIN_ASSEMBLY | 18 | 0.302972 | 0.91 | 0.584906 | 1 |
| GOBP_MITOTIC_SPINDLE_ORGANIZATION | 116 | 0.213018 | 0.908 | 0.684211 | 1 |
| GOBP_ERROR_PRONE_TRANSLESION_SYNTHESIS | 21 | 0.303188 | 0.907 | 0.65 | 1 |
| GOBP_REGULATION_OF_NUCLEAR_TRANSCRIBED_MRNA_CATABOLIC_PROCESS_DE | 16 | 0.307835 | 0.907 | 0.587379 | 1 |
| GOBP_BENZENE_CONTAINING_COMPOUND_METABOLIC_PROCESS | 23 | 0.285832 | 0.907 | 0.643678 | 1 |
| GOBP_CELL_CELL_ADHESION_MEDIATED_BY_CADHERIN | 27 | 0.276385 | 0.905 | 0.620438 | 1 |
| GOBP_REGULATION_OF_SISTER_CHROMATID_COHESION | 22 | 0.285218 | 0.904 | 0.566138 | 1 |
| GOBP_REGULATION_OF_MITOTIC_SISTER_CHROMATID_SEGREGATION | 43 | 0.242006 | 0.903 | 0.685714 | 1 |
| GOBP_CHROMOSOME_CONDENSATION | 40 | 0.248243 | 0.9 | 0.708333 | 1 |
| GOBP_NEGATIVE_REGULATION_OF_DNA_TEMPLATED_TRANSCRIPTION_ELONGATION | 17 | 0.305579 | 0.9 | 0.622449 | 1 |
| GOBP_REGULATION_OF_WATER_LOSS_VIA_SKIN | 26 | 0.27384 | 0.9 | 0.647059 | 1 |
| GOBP_SNRNA_PROCESSING | 25 | 0.276504 | 0.899 | 0.639053 | 1 |
| GOBP_NCRNA_3_END_PROCESSING | 39 | 0.247219 | 0.895 | 0.625 | 1 |
| GOBP_REGULATION_OF_ERAD_PATHWAY | 19 | 0.307282 | 0.895 | 0.622642 | 1 |
| GOBP_NUCLEOTIDE_EXCISION_REPAIR_DNA_GAP_FILLING | 23 | 0.277611 | 0.894 | 0.637838 | 1 |
| GOBP_POSITIVE_REGULATION_OF_ATPASE_ACTIVITY | 57 | 0.233224 | 0.891 | 0.78 | 1 |
| GOBP_PROTEIN_AUTOUBIQUITINATION | 70 | 0.218433 | 0.889 | 0.769231 | 1 |
| GOBP_RESPIRATORY_CHAIN_COMPLEX_IV_ASSEMBLY | 25 | 0.281024 | 0.888 | 0.633721 | 1 |
| GOBP_REGULATION_OF_MRNA_POLYADENYLATION | 17 | 0.316512 | 0.883 | 0.608696 | 1 |
| GOBP_LATE_ENDOSOME_TO_VACUOLE_TRANSPORT | 24 | 0.274784 | 0.879 | 0.65625 | 1 |
| GOBP_CYTOCHROME_COMPLEX_ASSEMBLY | 35 | 0.24845 | 0.878 | 0.736434 | 1 |
| GOBP_LIPOPROTEIN_CATABOLIC_PROCESS | 15 | 0.311797 | 0.878 | 0.657658 | 1 |
| GOBP_ENDOPLASMIC_RETICULUM_MANNOSE_TRIMMING | 16 | 0.29996 | 0.877 | 0.637615 | 1 |
| GOBP_NUCLEOSIDE_BISPHOSPHATE_METABOLIC_PROCESS | 129 | 0.19032 | 0.876 | 0.875 | 1 |
| GOBP_SEQUESTERING_OF_TRIGLYCERIDE | 16 | 0.30048 | 0.872 | 0.626866 | 1 |
| GOBP_CELLULAR_RESPONSE_TO_PROSTAGLANDIN_E_STIMULUS | 15 | 0.311895 | 0.871 | 0.577114 | 1 |
| GOBP_CHAPERONE_MEDIATED_PROTEIN_COMPLEX_ASSEMBLY | 20 | 0.294134 | 0.87 | 0.641304 | 1 |
| GOBP_MITOCHONDRIAL_TRANSCRIPTION | 17 | 0.298177 | 0.869 | 0.655022 | 1 |
| GOBP_PEPTIDYL_LYSINE_TRIMETHYLATION | 44 | 0.232558 | 0.869 | 0.771429 | 1 |
| GOBP_MRNA_CIS_SPLICING_VIA_SPLICEOSOME | 26 | 0.267197 | 0.868 | 0.6625 | 1 |
| GOBP_POSITIVE_REGULATION_OF_ACTIN_NUCLEATION | 15 | 0.315824 | 0.868 | 0.632653 | 1 |
| GOBP_DNA_REPLICATION_DEPENDENT_NUCLEOSOME_ORGANIZATION | 23 | 0.277284 | 0.866 | 0.682051 | 1 |
| GOBP_NUCLEAR_TRANSCRIBED_MRNA_POLY_A_TAIL_SHORTENING | 34 | 0.249413 | 0.865 | 0.746154 | 1 |
| GOBP_PURINE_CONTAINING_COMPOUND_TRANSMEMBRANE_TRANSPORT | 16 | 0.303744 | 0.864 | 0.630332 | 1 |
| GOBP_HISTONE_UBIQUITINATION | 46 | 0.237806 | 0.862 | 0.802326 | 1 |
| GOBP_NEGATIVE_REGULATION_OF_MACROAUTOPHAGY | 35 | 0.242261 | 0.861 | 0.75 | 1 |
| GOBP_ACTIN_FILAMENT_SEVERING | 16 | 0.300061 | 0.86 | 0.646154 | 1 |

| | | | | | |
|---|---|---|---|---|---|
| GOBP_REGULATION_OF_LEUKOCYTE_MIGRATION | 195 | -0.614924 | -1.78304 | 0 | 0.002095 |
| GOBP_LOCALIZATION_WITHIN_MEMBRANE | 74 | -0.640416 | -1.78239 | 0 | 0.002112 |
| GOBP_NEGATIVE_REGULATION_OF_REGULATED_SECRETORY_PATHWAY | 22 | -0.751314 | -1.782269 | 0 | 0.002105 |
| GOBP_POSITIVE_REGULATION_OF_PRODUCTION_OF_MOLECULAR_MEDIATOF | 91 | -0.628585 | -1.782177 | 0 | 0.00211 |
| GOBP_ALPHA_BETA_T_CELL_DIFFERENTIATION | 100 | -0.623683 | -1.780955 | 0 | 0.00216 |
| GOBP_REGULATION_OF_LEUKOCYTE_CHEMOTAXIS | 114 | -0.619215 | -1.78086 | 0 | 0.002147 |
| GOBP_NEGATIVE_REGULATION_OF_PRODUCTION_OF_MOLECULAR_MEDIATO | 35 | -0.692516 | -1.780628 | 0 | 0.00214 |
| GOBP_REGULATION_OF_SYNAPTIC_TRANSMISSION_GLUTAMATERGIC | 65 | -0.64864 | -1.780531 | 0 | 0.002145 |
| GOBP_NEGATIVE_REGULATION_OF_LEUKOCYTE_PROLIFERATION | 81 | -0.639737 | -1.779784 | 0 | 0.002155 |
| GOBP_EYE_MORPHOGENESIS | 140 | -0.61851 | -1.777697 | 0 | 0.00222 |
| GOBP_NEGATIVE_REGULATION_OF_MONONUCLEAR_CELL_MIGRATION | 23 | -0.736459 | -1.77522 | 0 | 0.002368 |
| GOBP_REGULATION_OF_PHOSPHOLIPASE_ACTIVITY | 67 | -0.637141 | -1.775072 | 0 | 0.00236 |
| GOBP_NEGATIVE_REGULATION_OF_T_CELL_RECEPTOR_SIGNALING_PATHWA | 22 | -0.735818 | -1.774022 | 0 | 0.002401 |
| GOBP_NEGATIVE_REGULATION_OF_LEUKOCYTE_CELL_CELL_ADHESION | 118 | -0.617067 | -1.773664 | 0 | 0.002404 |
| GOBP_REGULATION_OF_ANTIGEN_RECEPTOR_MEDIATED_SIGNALING_PATHW | 63 | -0.649853 | -1.7731 | 0 | 0.002439 |
| GOBP_POSITIVE_REGULATION_OF_SYNAPTIC_TRANSMISSION | 139 | -0.619065 | -1.771882 | 0 | 0.002501 |
| GOBP_REGULATION_OF_CYTOSOLIC_CALCIUM_ION_CONCENTRATION | 307 | -0.602821 | -1.77058 | 0 | 0.00253 |
| GOBP_NEURONAL_ACTION_POTENTIAL | 31 | -0.705942 | -1.770561 | 0 | 0.002516 |
| GOBP_POSITIVE_REGULATION_OF_T_CELL_MIGRATION | 29 | -0.719086 | -1.768047 | 0 | 0.002629 |
| GOBP_POSITIVE_REGULATION_OF_LYMPHOCYTE_CHEMOTAXIS | 19 | -0.7692 | -1.767948 | 0 | 0.00262 |
| GOBP_REGULATION_OF_ADAPTIVE_IMMUNE_RESPONSE | 158 | -0.614695 | -1.765901 | 0 | 0.002742 |
| GOBP_T_CELL_MEDIATED_CYTOTOXICITY | 42 | -0.666881 | -1.765595 | 0 | 0.002732 |
| GOBP_MEMBRANE_DEPOLARIZATION_DURING_CARDIAC_MUSCLE_CELL_ACT | 20 | -0.757242 | -1.765529 | 0 | 0.002723 |
| GOBP_POSITIVE_REGULATION_OF_INTERLEUKIN_12_PRODUCTION | 37 | -0.6876 | -1.764734 | 0 | 0.002765 |
| GOBP_NEGATIVE_REGULATION_OF_LYMPHOCYTE_ACTIVATION | 138 | -0.614207 | -1.764297 | 0 | 0.002801 |
| GOBP_REGULATION_OF_NEUROTRANSMITTER_RECEPTOR_ACTIVITY | 75 | -0.637139 | -1.764248 | 0 | 0.002791 |
| GOBP_TRANSMISSION_OF_NERVE_IMPULSE | 61 | -0.64499 | -1.764095 | 0.001064 | 0.002782 |
| GOBP_REGULATION_OF_PHAGOCYTOSIS | 88 | -0.625176 | -1.76345 | 0 | 0.002803 |
| GOBP_CELL_ADHESION_MEDIATED_BY_INTEGRIN | 71 | -0.636926 | -1.763023 | 0 | 0.002813 |
| GOBP_T_CELL_ACTIVATION_INVOLVED_IN_IMMUNE_RESPONSE | 88 | -0.627386 | -1.762323 | 0 | 0.002853 |
| GOBP_POSITIVE_REGULATION_OF_T_CELL_MEDIATED_IMMUNITY | 47 | -0.66585 | -1.759511 | 0 | 0.003032 |
| GOBP_CALCIUM_ION_TRANSPORT_INTO_CYTOSOL | 142 | -0.613352 | -1.759389 | 0 | 0.003021 |
| GOBP_MICROGLIAL_CELL_ACTIVATION | 42 | -0.670445 | -1.758381 | 0 | 0.003114 |
| GOBP_MATURE_B_CELL_DIFFERENTIATION_INVOLVED_IN_IMMUNE_RESPONS | 22 | -0.728496 | -1.756481 | 0.001214 | 0.003254 |
| GOBP_RESPONSE_TO_AMYLOID_BETA | 44 | -0.661072 | -1.754039 | 0 | 0.003384 |
| GOBP_ANTIBACTERIAL_HUMORAL_RESPONSE | 39 | -0.683251 | -1.753698 | 0 | 0.003401 |
| GOBP_TOLERANCE_INDUCTION | 26 | -0.726902 | -1.753465 | 0 | 0.003393 |
| GOBP_CD8_POSITIVE_ALPHA_BETA_T_CELL_ACTIVATION | 25 | -0.734156 | -1.75145 | 0 | 0.003491 |
| GOBP_NEGATIVE_REGULATION_OF_CELL_ACTIVATION | 180 | -0.602339 | -1.751429 | 0 | 0.003474 |
| GOBP_NEGATIVE_REGULATION_OF_IMMUNE_RESPONSE | 131 | -0.613003 | -1.751059 | 0 | 0.003476 |
| GOBP_CRANIAL_NERVE_DEVELOPMENT | 40 | -0.674681 | -1.748855 | 0 | 0.003647 |
| GOBP_POSITIVE_REGULATION_OF_MONONUCLEAR_CELL_MIGRATION | 59 | -0.656314 | -1.748236 | 0 | 0.003676 |
| GOBP_REGULATION_OF_B_CELL_DIFFERENTIATION | 28 | -0.711055 | -1.747681 | 0 | 0.003696 |
| GOBP_SODIUM_ION_EXPORT_ACROSS_PLASMA_MEMBRANE | 15 | -0.788489 | -1.747423 | 0.001266 | 0.003687 |
| GOBP_POSITIVE_REGULATION_OF_CYTOKINE_PRODUCTION_INVOLVED_IN_IM | 53 | -0.647546 | -1.745722 | 0 | 0.003859 |
| GOBP_REGULATION_OF_RELEASE_OF_SEQUESTERED_CALCIUM_ION_INTO_C | 69 | -0.633153 | -1.744703 | 0 | 0.00391 |
| GOBP_POSITIVE_REGULATION_OF_INTERLEUKIN_1_PRODUCTION | 57 | -0.647635 | -1.742032 | 0 | 0.004162 |
| GOBP_REGULATION_OF_T_CELL_MEDIATED_CYTOTOXICITY | 34 | -0.695519 | -1.741934 | 0 | 0.004146 |
| GOBP_REGULATION_OF_BONE_REMODELING | 47 | -0.656868 | -1.740238 | 0 | 0.004222 |
| GOBP_POSITIVE_REGULATION_OF_LEUKOCYTE_MIGRATION | 129 | -0.606544 | -1.740124 | 0 | 0.004211 |
| GOBP_MATURE_B_CELL_DIFFERENTIATION | 27 | -0.713075 | -1.739753 | 0 | 0.004214 |
| GOBP_NEGATIVE_REGULATION_OF_T_CELL_PROLIFERATION | 61 | -0.641814 | -1.739549 | 0 | 0.004222 |
| GOBP_POSITIVE_REGULATION_OF_LEUKOCYTE_CHEMOTAXIS | 87 | -0.628049 | -1.739435 | 0 | 0.004211 |
| GOBP_POSITIVE_REGULATION_OF_SYNAPSE_ASSEMBLY | 57 | -0.651698 | -1.738082 | 0 | 0.004302 |
| GOBP_REGULATION_OF_TRANS_SYNAPTIC_SIGNALING | 393 | -0.586225 | -1.736941 | 0 | 0.004397 |
| GOBP_DIENCEPHALON_DEVELOPMENT | 59 | -0.636204 | -1.736723 | 0 | 0.004408 |
| GOBP_POSITIVE_REGULATION_OF_TUMOR_NECROSIS_FACTOR_SUPERFAMIL | 81 | -0.614355 | -1.735845 | 0 | 0.004493 |
| GOBP_REGULATION_OF_LEUKOCYTE_APOPTOTIC_PROCESS | 74 | -0.631316 | -1.734992 | 0 | 0.004542 |
| GOBP_POSITIVE_REGULATION_OF_B_CELL_PROLIFERATION | 36 | -0.676327 | -1.734775 | 0 | 0.004539 |
| GOBP_NEUROTRANSMITTER_RECEPTOR_TRANSPORT | 45 | -0.645021 | -1.734553 | 0.001101 | 0.004536 |
| GOBP_REGULATION_OF_TYPE_2_IMMUNE_RESPONSE | 27 | -0.699284 | -1.733823 | 0 | 0.004572 |
| GOBP_COCHLEA_MORPHOGENESIS | 19 | -0.744048 | -1.733787 | 0.001261 | 0.004552 |
| GOBP_NEGATIVE_REGULATION_OF_ADAPTIVE_IMMUNE_RESPONSE | 49 | -0.652484 | -1.733676 | 0.001096 | 0.004536 |
| GOBP_CYTOSOLIC_CALCIUM_ION_TRANSPORT | 163 | -0.591188 | -1.733461 | 0 | 0.004537 |
| GOBP_EOSINOPHIL_MIGRATION | 20 | -0.735456 | -1.732334 | 0.001224 | 0.004606 |
| GOBP_LEUKOCYTE_APOPTOTIC_PROCESS | 95 | -0.614105 | -1.732279 | 0 | 0.004598 |
| GOBP_NEGATIVE_REGULATION_OF_ANTIGEN_RECEPTOR_MEDIATED_SIGNAL | 29 | -0.705826 | -1.732098 | 0.001181 | 0.004595 |
| GOBP_REGULATION_OF_POSTSYNAPTIC_MEMBRANE_NEUROTRANSMITTER_I | 58 | -0.632941 | -1.730998 | 0 | 0.004683 |
| GOBP_POSITIVE_REGULATION_OF_INFLAMMATORY_RESPONSE | 114 | -0.608269 | -1.730109 | 0 | 0.00475 |
| GOBP_GLUTAMATE_RECEPTOR_SIGNALING_PATHWAY | 52 | -0.64506 | -1.729903 | 0 | 0.004738 |
| GOBP_LYMPHOCYTE_ACTIVATION_INVOLVED_IN_IMMUNE_RESPONSE | 159 | -0.596023 | -1.728843 | 0 | 0.004812 |
| GOBP_T_HELPER_1_TYPE_IMMUNE_RESPONSE | 38 | -0.661544 | -1.726663 | 0.003386 | 0.005045 |
| GOBP_POSITIVE_REGULATION_OF_CYCLASE_ACTIVITY | 19 | -0.751147 | -1.726539 | 0.002445 | 0.005032 |
| GOBP_NEUROTRANSMITTER_TRANSPORT | 199 | -0.59497 | -1.725177 | 0 | 0.005165 |
| GOBP_NEGATIVE_REGULATION_OF_SYNAPTIC_TRANSMISSION | 61 | -0.632015 | -1.725132 | 0 | 0.005143 |
| GOBP_POSITIVE_REGULATION_OF_LEUKOCYTE_ADHESION_TO_VASCULAR_E | 20 | -0.750767 | -1.724879 | 0 | 0.005158 |
| GOBP_MULTICELLULAR_ORGANISMAL_SIGNALING | 191 | -0.587789 | -1.724271 | 0 | 0.005229 |
| GOBP_TYPE_2_IMMUNE_RESPONSE | 33 | -0.680473 | -1.722899 | 0 | 0.005355 |
| GOBP_NATURAL_KILLER_CELL_DIFFERENTIATION | 16 | -0.76683 | -1.722845 | 0.001276 | 0.005333 |
| GOBP_POSITIVE_REGULATION_OF_LIPASE_ACTIVITY | 65 | -0.633059 | -1.722525 | 0 | 0.005338 |
| GOBP_REGULATION_OF_FIBROBLAST_GROWTH_FACTOR_RECEPTOR_SIGNAL | 27 | -0.713021 | -1.722415 | 0 | 0.005321 |
| GOBP_POSITIVE_REGULATION_OF_CHEMOTAXIS | 133 | -0.601188 | -1.722254 | 0 | 0.005311 |
| GOBP_MESENCHYMAL_CELL_PROLIFERATION | 37 | -0.659107 | -1.721695 | 0 | 0.005359 |
| GOBP_INNERVATION | 23 | -0.72945 | -1.7216 | 0 | 0.005353 |
| GOBP_POSITIVE_REGULATION_OF_FATTY_ACID_BIOSYNTHETIC_PROCESS | 17 | -0.753097 | -1.719378 | 0 | 0.005572 |
| GOBP_NEUROTRANSMITTER_RECEPTOR_INTERNALIZATION | 24 | -0.721195 | -1.719252 | 0 | 0.005566 |
| GOBP_CYTOLYSIS | 22 | -0.729088 | -1.719147 | 0.001232 | 0.005555 |
| GOBP_REGULATION_OF_SYNAPSE_ASSEMBLY | 90 | -0.614274 | -1.718327 | 0 | 0.00559 |
| GOBP_POSITIVE_REGULATION_OF_CHEMOKINE_PRODUCTION | 54 | -0.637576 | -1.717984 | 0.00108 | 0.005603 |
| GOBP_CHRONIC_INFLAMMATORY_RESPONSE | 18 | -0.765941 | -1.71742 | 0 | 0.005626 |
| GOBP_REGULATION_OF_NEUTROPHIL_MIGRATION | 41 | -0.661053 | -1.715889 | 0 | 0.005767 |
| GOBP_NEGATIVE_REGULATION_OF_CYTOKINE_PRODUCTION_INVOLVED_IN_I | 23 | -0.731666 | -1.715247 | 0 | 0.005865 |
| GOBP_HETEROPHILIC_CELL_CELL_ADHESION_VIA_PLASMA_MEMBRANE_CEL | 43 | -0.648606 | -1.714498 | 0 | 0.005921 |
| GOBP_POSITIVE_REGULATION_OF_LYASE_ACTIVITY | 19 | -0.741824 | -1.713849 | 0 | 0.005946 |
| GOBP_PROTEIN_LOCALIZATION_TO_POSTSYNAPSE | 43 | -0.655562 | -1.713391 | 0 | 0.005979 |
| GOBP_LEUKOCYTE_DIFFERENTIATION | 472 | -0.581798 | -1.713388 | 0 | 0.005956 |
| GOBP_SENSORY_SYSTEM_DEVELOPMENT | 348 | -0.579724 | -1.709093 | 0 | 0.00644 |
| GOBP_ODONTOGENESIS_OF_DENTIN_CONTAINING_TOOTH | 75 | -0.612611 | -1.70866 | 0 | 0.006456 |
| GOBP_REGULATION_OF_ALPHA_BETA_T_CELL_DIFFERENTIATION | 63 | -0.62158 | -1.708289 | 0.001071 | 0.006465 |
| GOBP_REGULATION_OF_CALCIUM_ION_TRANSMEMBRANE_TRANSPORT | 131 | -0.592581 | -1.707997 | 0 | 0.006473 |
| GOBP_LYMPHOCYTE_HOMEOSTASIS | 56 | -0.640278 | -1.707962 | 0 | 0.006452 |
| GOBP_NEURON_MIGRATION | 135 | -0.596074 | -1.707121 | 0 | 0.006522 |
| GOBP_EMBRYONIC_APPENDAGE_MORPHOGENESIS | 105 | -0.597867 | -1.706956 | 0 | 0.006519 |
| GOBP_NEGATIVE_REGULATION_OF_LEUKOCYTE_APOPTOTIC_PROCESS | 43 | -0.658024 | -1.706745 | 0.001093 | 0.006531 |
| GOBP_CAMERA_TYPE_EYE_MORPHOGENESIS | 113 | -0.601323 | -1.706659 | 0 | 0.006528 |
| GOBP_T_CELL_APOPTOTIC_PROCESS | 44 | -0.652061 | -1.706256 | 0 | 0.006547 |
| GOBP_CHEMOKINE_PRODUCTION | 87 | -0.607497 | -1.705749 | 0 | 0.006558 |
| GOBP_REGULATION_OF_T_CELL_CHEMOTAXIS | 15 | -0.773335 | -1.705356 | 0 | 0.006566 |
| GOBP_POSITIVE_REGULATION_OF_REGULATORY_T_CELL_DIFFERENTIATION | 15 | -0.757936 | -1.704841 | 0.002581 | 0.006634 |
| GOBP_CELL_FATE_DETERMINATION | 36 | -0.667102 | -1.702817 | 0 | 0.006923 |
| GOBP_REGULATION_OF_MONOCYTE_CHEMOTAXIS | 25 | -0.703002 | -1.701796 | 0.001183 | 0.007052 |
| GOBP_DETECTION_OF_LIGHT_STIMULUS_INVOLVED_IN_SENSORY_PERCEPTI | 18 | -0.757315 | -1.701541 | 0.002541 | 0.007058 |
| GOBP_MACROPHAGE_ACTIVATION | 91 | -0.610396 | -1.701457 | 0 | 0.007054 |
| GOBP_LYMPH_NODE_DEVELOPMENT | 17 | -0.744877 | -1.701366 | 0.002494 | 0.007042 |
| GOBP_REGULATION_OF_ANTIGEN_PROCESSING_AND_PRESENTATION | 20 | -0.722459 | -1.700617 | 0 | 0.0071 |
| GOBP_GLUTAMATE_SECRETION | 38 | -0.668008 | -1.699733 | 0.001131 | 0.007205 |
| GOBP_POSITIVE_REGULATION_OF_PHOSPHATIDYLINOSITOL_3_KINASE_SIGNA | 85 | -0.61409 | -1.698298 | 0 | 0.007379 |
| GOBP_RETINA_LAYER_FORMATION | 24 | -0.687572 | -1.696338 | 0.001182 | 0.007606 |
| GOBP_POSITIVE_REGULATION_OF_T_CELL_MEDIATED_CYTOTOXICITY | 25 | -0.696231 | -1.696112 | 0.001176 | 0.007606 |
| GOBP_CAMERA_TYPE_EYE_DEVELOPMENT | 292 | -0.575653 | -1.694713 | 0 | 0.007746 |
| GOBP_INTERFERON_GAMMA_MEDIATED_SIGNALING_PATHWAY | 87 | -0.607606 | -1.694549 | 0 | 0.007746 |
| GOBP_CD4_POSITIVE_ALPHA_BETA_T_CELL_ACTIVATION | 90 | -0.608944 | -1.694392 | 0 | 0.007746 |
| GOBP_MEMBRANE_DEPOLARIZATION | 82 | -0.603262 | -1.693927 | 0 | 0.007793 |
| GOBP_POSITIVE_REGULATION_OF_SYNAPTIC_TRANSMISSION_GLUTAMATERG | 29 | -0.685234 | -1.693668 | 0.002336 | 0.007812 |
| GOBP_MYELOID_DENDRITIC_CELL_ACTIVATION | 28 | -0.690407 | -1.693187 | 0 | 0.007859 |
| GOBP_TISSUE_HOMEOSTASIS | 245 | -0.574849 | -1.692504 | 0 | 0.007948 |
| GOBP_SENSORY_PERCEPTION | 492 | -0.568549 | -1.691075 | 0 | 0.008169 |
| GOBP_POSITIVE_REGULATION_OF_ERK1_AND_ERK2_CASCADE | 180 | -0.581581 | -1.689328 | 0 | 0.008389 |
| GOBP_REGULATION_OF_CD8_POSITIVE_ALPHA_BETA_T_CELL_ACTIVATION | 19 | -0.732659 | -1.688937 | 0.001244 | 0.008423 |
| GOBP_SENSORY_ORGAN_MORPHOGENESIS | 228 | -0.574376 | -1.688801 | 0 | 0.008411 |
| GOBP_FATTY_ACID_TRANSMEMBRANE_TRANSPORT | 49 | -0.634703 | -1.688344 | 0 | 0.008464 |
| GOBP_EOSINOPHIL_CHEMOTAXIS | 16 | -0.761055 | -1.688246 | 0.001259 | 0.008458 |
| GOBP_POSITIVE_REGULATION_OF_ADAPTIVE_IMMUNE_RESPONSE | 95 | -0.600523 | -1.688054 | 0 | 0.008452 |
| GOBP_B_CELL_DIFFERENTIATION | 115 | -0.592227 | -1.687883 | 0 | 0.008443 |
| GOBP_POSITIVE_REGULATION_OF_CELL_ADHESION | 405 | -0.572207 | -1.687707 | 0 | 0.008441 |
| GOBP_CD4_POSITIVE_ALPHA_BETA_T_CELL_DIFFERENTIATION | 75 | -0.610684 | -1.687665 | 0 | 0.008422 |
| GOBP_NEUROTRANSMITTER_SECRETION | 151 | -0.583837 | -1.686746 | 0 | 0.008567 |
| GOBP_MAST_CELL_ACTIVATION_INVOLVED_IN_IMMUNE_RESPONSE | 44 | -0.647461 | -1.685397 | 0.002193 | 0.008772 |
| GOBP_INTERLEUKIN_1_PRODUCTION | 99 | -0.594303 | -1.684696 | 0 | 0.008852 |
| GOBP_POSITIVE_REGULATION_OF_CYTOKINE_PRODUCTION | 394 | -0.569037 | -1.684538 | 0 | 0.008838 |
| GOBP_REGULATION_OF_CATION_TRANSMEMBRANE_TRANSPORT | 317 | -0.568599 | -1.684382 | 0 | 0.008847 |
| GOBP_REGULATION_OF_T_CELL_APOPTOTIC_PROCESS | 30 | -0.683862 | -1.683404 | 0.002334 | 0.008992 |
| GOBP_PROTEIN_LOCALIZATION_TO_SYNAPSE | 79 | -0.613475 | -1.682442 | 0 | 0.009117 |
| GOBP_REGULATION_OF_SYNAPSE_STRUCTURE_OR_ACTIVITY | 205 | -0.574094 | -1.682147 | 0 | 0.009156 |
| GOBP_CALCIUM_ION_TRANSMEMBRANE_TRANSPORT | 282 | -0.574167 | -1.681074 | 0 | 0.009307 |
| GOBP_LEARNING | 124 | -0.584205 | -1.680906 | 0 | 0.009315 |
| GOBP_ADENYLATE_CYCLASE_ACTIVATING_G_PROTEIN_COUPLED_RECEPTOI | 118 | -0.586002 | -1.679951 | 0 | 0.009465 |
| GOBP_REGULATION_OF_CELL_ADHESION_MEDIATED_BY_INTEGRIN | 47 | -0.632377 | -1.679782 | 0.002183 | 0.009465 |
| GOBP_NEGATIVE_REGULATION_OF_HEMOPOIESIS | 92 | -0.597087 | -1.679574 | 0 | 0.009475 |
| GOBP_T_CELL_DIFFERENTIATION_IN_THYMUS | 70 | -0.608191 | -1.679112 | 0 | 0.009531 |
| GOBP_CD4_POSITIVE_OR_CD8_POSITIVE_ALPHA_BETA_T_CELL_LINEAGE_CO | 20 | -0.708903 | -1.678989 | 0.003736 | 0.009509 |

| | | | | | |
|---|---|---|---|---|---|
| GOBP_CHROMATIN_REMODELING | 190 | 0.192349 | 0.86 | 1 | 1 |
| GOBP_HISTONE_H3_ACETYLATION | 60 | 0.218448 | 0.859 | 0.830509 | 1 |
| GOBP_OXIDATIVE_DEMETHYLATION | 17 | 0.295785 | 0.853 | 0.7 | 1 |
| GOBP_CELLULAR_RESPONSE_TO_DSRNA | 20 | 0.277859 | 0.853 | 0.673913 | 1 |
| GOBP_REGULATION_OF_LIPOPROTEIN_PARTICLE_CLEARANCE | 16 | 0.300645 | 0.851 | 0.648649 | 1 |
| GOBP_REGULATION_OF_MITOCHONDRIAL_GENE_EXPRESSION | 29 | 0.257718 | 0.85 | 0.729323 | 1 |
| GOBP_CELL_CYCLE_CHECKPOINT | 203 | 0.189997 | 0.85 | 1 | 1 |
| GOBP_METAPHASE_PLATE_CONGRESSION | 63 | 0.21456 | 0.848 | 0.884058 | 1 |
| GOBP_LATE_ENDOSOME_TO_VACUOLE_TRANSPORT_VIA_MULTIVESICULAR_BODY_S | 17 | 0.296657 | 0.846 | 0.647887 | 1 |
| GOBP_MEIOTIC_CHROMOSOME_SEPARATION | 24 | 0.261147 | 0.846 | 0.741936 | 1 |
| GOBP_TELOMERE_MAINTENANCE_VIA_TELOMERE_LENGTHENING | 79 | 0.209944 | 0.844 | 0.923077 | 1 |
| GOBP_HISTONE_MONOUBIQUITINATION | 29 | 0.250066 | 0.843 | 0.776119 | 1 |
| GOBP_DEMETHYLATION | 66 | 0.205815 | 0.843 | 0.895833 | 1 |
| GOBP_REGULATION_OF_HISTONE_DEACETYLATION | 26 | 0.255236 | 0.843 | 0.756098 | 1 |
| GOBP_REGULATION_OF_CELL_MATURATION | 20 | 0.278303 | 0.841 | 0.724324 | 1 |
| GOBP_RNA_SURVEILLANCE | 15 | 0.30278 | 0.834 | 0.679348 | 1 |
| GOBP_REGULATION_OF_NUCLEAR_TRANSCRIBED_MRNA_POLY_A_TAIL_SHORTENIN | 15 | 0.307822 | 0.832 | 0.733945 | 1 |
| GOBP_SNO_S_RNA_METABOLIC_PROCESS | 15 | 0.291366 | 0.83 | 0.725225 | 1 |
| GOBP_POSITIVE_REGULATION_OF_MITOCHONDRIAL_TRANSLATION | 16 | 0.295088 | 0.826 | 0.704762 | 1 |
| GOBP_POSITIVE_REGULATION_OF_CELL_CYCLE_G2_M_PHASE_TRANSITION | 29 | 0.240427 | 0.826 | 0.791045 | 1 |
| GOBP_CHROMATIN_ASSEMBLY_OR_DISASSEMBLY | 182 | 0.173529 | 0.825 | 1 | 1 |
| GOBP_N_TERMINAL_PROTEIN_AMINO_ACID_MODIFICATION | 27 | 0.251997 | 0.825 | 0.738255 | 1 |
| GOBP_MICROTUBULE_CYTOSKELETON_ORGANIZATION_INVOLVED_IN_MITOSIS | 140 | 0.187547 | 0.823 | 1 | 1 |
| GOBP_HOMOLOGOUS_RECOMBINATION | 57 | 0.205472 | 0.822 | 0.855263 | 1 |
| GOBP_MIRNA_METABOLIC_PROCESS | 27 | 0.249336 | 0.819 | 0.790419 | 1 |
| GOBP_NUCLEOSIDE_TRIPHOSPHATE_BIOSYNTHETIC_PROCESS | 83 | 0.19549 | 0.819 | 0.96875 | 1 |
| GOBP_OLIGOSACCHARIDE_LIPID_INTERMEDIATE_BIOSYNTHETIC_PROCESS | 21 | 0.263298 | 0.819 | 0.791444 | 1 |
| GOBP_METHIONINE_METABOLIC_PROCESS | 16 | 0.284216 | 0.819 | 0.702326 | 1 |
| GOBP_DIACYLGLYCEROL_METABOLIC_PROCESS | 22 | 0.260627 | 0.819 | 0.75 | 1 |
| GOBP_ANDROGEN_RECEPTOR_SIGNALING_PATHWAY | 37 | 0.232264 | 0.818 | 0.807018 | 1 |
| GOBP_NEGATIVE_REGULATION_OF_ENDOPLASMIC_RETICULUM_STRESS_INDUCED_ | 15 | 0.289683 | 0.817 | 0.726829 | 1 |
| GOBP_RDNA_HETEROCHROMATIN_ASSEMBLY | 28 | 0.248103 | 0.808 | 0.736487 | 1 |
| GOBP_POSITIVE_REGULATION_OF_SUBSTRATE_ADHESION_DEPENDENT_CELL_SPRE | 38 | 0.231131 | 0.806 | 0.779661 | 1 |
| GOBP_NEGATIVE_REGULATION_OF_OXIDATIVE_STRESS_INDUCED_INTRINSIC_APOPT | 19 | 0.267836 | 0.802 | 0.717172 | 1 |
| GOBP_RECEPTOR_CATABOLIC_PROCESS | 35 | 0.230042 | 0.799 | 0.838462 | 1 |
| GOBP_INTRA_S_DNA_DAMAGE_CHECKPOINT | 17 | 0.276326 | 0.798 | 0.763033 | 1 |
| GOBP_LIPID_DROPLET_ORGANIZATION | 26 | 0.244913 | 0.798 | 0.827586 | 1 |
| GOBP_RNA_POLYADENYLATION | 46 | 0.219214 | 0.794 | 0.865385 | 1 |
| GOBP_SOMATIC_DIVERSIFICATION_OF_IMMUNE_RECEPTORS_VIA_SOMATIC_MUTATIC | 15 | 0.275792 | 0.792 | 0.766816 | 1 |
| GOBP_NEGATIVE_REGULATION_OF_CELLULAR_PROTEIN_CATABOLIC_PROCESS | 73 | 0.210098 | 0.791 | 0.878049 | 1 |
| GOBP_POSITIVE_REGULATION_OF_MAST_CELL_ACTIVATION | 16 | 0.272793 | 0.79 | 0.781659 | 1 |
| GOBP_LUNG_CELL_DIFFERENTIATION | 17 | 0.267896 | 0.788 | 0.782609 | 1 |
| GOBP_NUCLEOSOME_ORGANIZATION | 145 | 0.174662 | 0.787 | 1 | 1 |
| GOBP_RIBONUCLEOSIDE_TRIPHOSPHATE_METABOLIC_PROCESS | 86 | 0.183418 | 0.785 | 0.956522 | 1 |
| GOBP_TELOMERE_ORGANIZATION | 163 | 0.17365 | 0.782 | 1 | 1 |
| GOBP_DNA_TEMPLATED_TRANSCRIPTION_ELONGATION | 113 | 0.174932 | 0.78 | 0.9375 | 1 |
| GOBP_REGULATION_OF_OXIDATIVE_STRESS_INDUCED_INTRINSIC_APOPTOTIC_SIGN | 27 | 0.246638 | 0.78 | 0.821656 | 1 |
| GOBP_GLUCOSE_CATABOLIC_PROCESS | 35 | 0.222006 | 0.78 | 0.810345 | 1 |
| GOBP_KINETOCHORE_ORGANIZATION | 22 | 0.248661 | 0.778 | 0.860606 | 1 |
| GOBP_REGULATION_OF_SPINDLE_ORGANIZATION | 39 | 0.214152 | 0.776 | 0.864865 | 1 |
| GOBP_MATURATION_OF_5_8S_RRNA | 35 | 0.220518 | 0.775 | 0.882813 | 1 |
| GOBP_REGULATION_OF_MITOTIC_SPINDLE_ASSEMBLY | 18 | 0.254663 | 0.775 | 0.79 | 1 |
| GOBP_CHROMOSOME_SEPARATION | 91 | 0.173374 | 0.774 | 1 | 1 |
| GOBP_MITOCHONDRIAL_RNA_PROCESSING | 17 | 0.260951 | 0.773 | 0.79798 | 1 |
| GOBP_HISTONE_H2A_UBIQUITINATION | 26 | 0.237596 | 0.773 | 0.869565 | 1 |
| GOBP_MISMATCH_REPAIR | 36 | 0.21696 | 0.772 | 0.903509 | 1 |
| GOBP_MITOTIC_SISTER_CHROMATID_COHESION | 26 | 0.235159 | 0.77 | 0.833333 | 1 |
| GOBP_ENDOSOME_TRANSPORT_VIA_MULTIVESICULAR_BODY_SORTING_PATHWAY | 31 | 0.225297 | 0.77 | 0.814516 | 1 |
| GOBP_POSITIVE_REGULATION_OF_HISTONE_METHYLATION | 34 | 0.220834 | 0.769 | 0.844961 | 1 |
| GOBP_MITOCHONDRIAL_CYTOCHROME_C_OXIDASE_ASSEMBLY | 21 | 0.248296 | 0.769 | 0.858537 | 1 |
| GOBP_REGULATION_OF_CHOLESTEROL_BIOSYNTHETIC_PROCESS | 43 | 0.213344 | 0.765 | 0.892157 | 1 |
| GOBP_MICROTUBULE_NUCLEATION | 30 | 0.219266 | 0.763 | 0.876812 | 1 |
| GOBP_POLYAMINE_METABOLIC_PROCESS | 17 | 0.261198 | 0.763 | 0.774359 | 1 |
| GOBP_NEGATIVE_REGULATION_OF_METAPHASE_ANAPHASE_TRANSITION_OF_CELL_ | 40 | 0.211389 | 0.76 | 0.912621 | 1 |
| GOBP_PROTEIN_MATURATION_BY_IRON_SULFUR_CLUSTER_TRANSFER | 15 | 0.278272 | 0.76 | 0.800905 | 1 |
| GOBP_CELLULAR_RESPONSE_TO_EXOGENOUS_DSRNA | 16 | 0.271546 | 0.759 | 0.799107 | 1 |
| GOBP_BARBED_END_ACTIN_FILAMENT_CAPPING | 25 | 0.238613 | 0.754 | 0.877551 | 1 |
| GOBP_BLASTOCYST_FORMATION | 32 | 0.229814 | 0.752 | 0.838983 | 1 |
| GOBP_REGULATION_OF_CHROMOSOME_SEPARATION | 67 | 0.185601 | 0.75 | 0.951613 | 1 |
| GOBP_DNA_PACKAGING | 196 | 0.164406 | 0.749 | 1 | 1 |
| GOBP_REGULATION_OF_TELOMERE_MAINTENANCE | 79 | 0.173142 | 0.747 | 0.980769 | 1 |
| GOBP_CENTROSOME_DUPLICATION | 71 | 0.183991 | 0.747 | 1 | 1 |
| GOBP_PROTEIN_NEDDYLATION | 17 | 0.245067 | 0.745 | 0.826733 | 1 |
| GOBP_NEGATIVE_REGULATION_OF_CHROMATIN_ORGANIZATION | 57 | 0.188554 | 0.741 | 0.960526 | 1 |
| GOBP_ENDOPLASMIC_RETICULUM_TO_CYTOSOL_TRANSPORT | 28 | 0.227214 | 0.737 | 0.869281 | 1 |
| GOBP_NEGATIVE_REGULATION_OF_VIRAL_TRANSCRIPTION | 15 | 0.264977 | 0.734 | 0.827907 | 1 |
| GOBP_NEGATIVE_REGULATION_OF_INTRACELLULAR_STEROID_HORMONE_RECEPTC | 35 | 0.211201 | 0.733 | 0.951613 | 1 |
| GOBP_NCRNA_CATABOLIC_PROCESS | 34 | 0.210515 | 0.733 | 0.903226 | 1 |
| GOBP_CHROMATIN_REMODELING_AT_CENTROMERE | 43 | 0.199463 | 0.732 | 0.938776 | 1 |
| GOBP_MITOTIC_SISTER_CHROMATID_SEGREGATION | 159 | 0.157287 | 0.727 | 1 | 1 |
| GOBP_ORGANELLE_DISASSEMBLY | 106 | 0.171339 | 0.727 | 1 | 1 |
| GOBP_LEFT_RIGHT_PATTERN_FORMATION | 19 | 0.243616 | 0.726 | 0.867021 | 1 |
| GOBP_PROTEIN_LOCALIZATION_TO_KINETOCHORE | 19 | 0.248102 | 0.726 | 0.855072 | 1 |
| GOBP_ESTABLISHMENT_OF_MITOTIC_SPINDLE_LOCALIZATION | 32 | 0.211409 | 0.722 | 0.910448 | 1 |
| GOBP_PROTEIN_TARGETING_TO_MITOCHONDRION | 100 | 0.175475 | 0.722 | 1 | 1 |
| GOBP_INTRINSIC_APOPTOTIC_SIGNALING_PATHWAY_IN_RESPONSE_TO_DNA_DAMA( | 44 | 0.190932 | 0.72 | 0.980583 | 1 |
| GOBP_STEROL_BIOSYNTHETIC_PROCESS | 71 | 0.180983 | 0.72 | 0.98 | 1 |
| GOBP_NEGATIVE_REGULATION_OF_DNA_METABOLIC_PROCESS | 115 | 0.158221 | 0.719 | 1 | 1 |
| GOBP_REGULATION_OF_CHROMATIN_BINDING | 23 | 0.229079 | 0.714 | 0.921569 | 1 |
| GOBP_N_TERMINAL_PROTEIN_AMINO_ACID_ACETYLATION | 16 | 0.241804 | 0.714 | 0.886878 | 1 |
| GOBP_HISTONE_H4_K16_ACETYLATION | 21 | 0.233229 | 0.714 | 0.9 | 1 |
| GOBP_SNRNA_TRANSCRIPTION | 75 | 0.176752 | 0.71 | 1 | 1 |
| GOBP_NUCLEOSOME_ASSEMBLY | 111 | 0.157074 | 0.71 | 1 | 1 |
| GOBP_DRUG_CATABOLIC_PROCESS | 16 | 0.246029 | 0.707 | 0.853535 | 1 |
| GOBP_HISTONE_H4_ACETYLATION | 68 | 0.168976 | 0.707 | 0.981132 | 1 |
| GOBP_DOUBLE_STRAND_BREAK_REPAIR | 256 | 0.149991 | 0.706 | 1 | 1 |
| GOBP_RNA_PHOSPHODIESTER_BOND_HYDROLYSIS_ENDONUCLEOLYTIC | 75 | 0.175964 | 0.705 | 1 | 1 |
| GOBP_TAIL_ANCHORED_MEMBRANE_PROTEIN_INSERTION_INTO_ER_MEMBRANE | 16 | 0.251781 | 0.704 | 0.886364 | 1 |
| GOBP_MACROMOLECULE_DEACYLATION | 106 | 0.172212 | 0.703 | 1 | 1 |
| GOBP_RNA_METHYLATION | 84 | 0.165876 | 0.701 | 0.971429 | 1 |
| GOBP_POSITIVE_REGULATION_OF_UBIQUITIN_PROTEIN_TRANSFERASE_ACTIVITY | 31 | 0.20478 | 0.699 | 0.942446 | 1 |
| GOBP_MONOSACCHARIDE_BIOSYNTHETIC_PROCESS | 89 | 0.167192 | 0.698 | 1 | 1 |
| GOBP_ESTABLISHMENT_OF_MITOTIC_SPINDLE_ORIENTATION | 27 | 0.207467 | 0.696 | 0.942529 | 1 |
| GOBP_ALDITOL_METABOLIC_PROCESS | 21 | 0.228189 | 0.69 | 0.897297 | 1 |
| GOBP_VESICLE_CARGO_LOADING | 24 | 0.207546 | 0.687 | 0.939394 | 1 |
| GOBP_REGULATION_OF_PROTON_TRANSPORT | 21 | 0.216813 | 0.683 | 0.917526 | 1 |
| GOBP_TRANSCRIPTION_ELONGATION_FROM_RNA_POLYMERASE_II_PROMOTER | 84 | 0.156323 | 0.683 | 0.97561 | 1 |
| GOBP_RNA_PHOSPHODIESTER_BOND_HYDROLYSIS | 146 | 0.153642 | 0.68 | 1 | 1 |
| GOBP_COENZYME_A_METABOLIC_PROCESS | 15 | 0.243231 | 0.68 | 0.882883 | 1 |
| GOBP_PEPTIDYL_GLUTAMIC_ACID_MODIFICATION | 28 | 0.20173 | 0.672 | 0.956522 | 1 |
| GOBP_REGULATION_OF_TRANSCRIPTION_FROM_RNA_POLYMERASE_II_PROMOTER_ | 76 | 0.166339 | 0.668 | 0.983051 | 1 |
| GOBP_REGULATION_OF_TELOMERE_MAINTENANCE_VIA_TELOMERE_LENGTHENING | 60 | 0.16499 | 0.667 | 0.981482 | 1 |
| GOBP_ATTACHMENT_OF_MITOTIC_SPINDLE_MICROTUBULES_TO_KINETOCHORE | 15 | 0.236772 | 0.667 | 0.918919 | 1 |
| GOBP_REGULATION_OF_TRANSCRIPTION_ELONGATION_FROM_RNA_POLYMERASE_I | 30 | 0.201522 | 0.665 | 0.916667 | 1 |
| GOBP_DNA_REPLICATION_INITIATION | 38 | 0.18435 | 0.665 | 0.951456 | 1 |
| GOBP_RNA_3_END_PROCESSING | 140 | 0.151386 | 0.664 | 1 | 1 |
| GOBP_POSITIVE_REGULATION_OF_METAPHASE_ANAPHASE_TRANSITION_OF_CELL_( | 15 | 0.241625 | 0.664 | 0.918269 | 1 |
| GOBP_MULTIVESICULAR_BODY_SORTING_PATHWAY | 37 | 0.186613 | 0.663 | 0.984375 | 1 |
| GOBP_POSTREPLICATION_REPAIR | 52 | 0.171449 | 0.66 | 0.987179 | 1 |
| GOBP_ENERGY_DERIVATION_BY_OXIDATION_OF_ORGANIC_COMPOUNDS | 265 | 0.152405 | 0.658 | 1 | 1 |
| GOBP_HISTONE_H3_K27_METHYLATION | 17 | 0.22169 | 0.658 | 0.927835 | 1 |
| GOBP_MANNOSYLATION | 36 | 0.191516 | 0.658 | 0.980392 | 1 |
| GOBP_REGULATION_OF_CILIUM_DEPENDENT_CELL_MOTILITY | 15 | 0.236311 | 0.657 | 0.908676 | 1 |
| GOBP_TRNA_MODIFICATION | 88 | 0.151915 | 0.655 | 1 | 1 |
| GOBP_REGULATION_OF_INTRACELLULAR_STEROID_HORMONE_RECEPTOR_SIGNALI | 72 | 0.162846 | 0.655 | 1 | 1 |
| GOBP_AUTOPHAGY_OF_MITOCHONDRION | 74 | 0.169936 | 0.655 | 0.975 | 1 |
| GOBP_RNA_DEPENDENT_DNA_BIOSYNTHETIC_PROCESS | 69 | 0.161242 | 0.653 | 1 | 1 |
| GOBP_REGULATION_OF_ENDOPLASMIC_RETICULUM_UNFOLDED_PROTEIN_RESPON: | 27 | 0.197802 | 0.651 | 0.953947 | 1 |
| GOBP_SPINDLE_ORGANIZATION | 179 | 0.139059 | 0.648 | 1 | 1 |
| GOBP_NUCLEAR_TRANSCRIBED_MRNA_CATABOLIC_PROCESS_DEADENYLATION_DE | 75 | 0.152587 | 0.646 | 1 | 1 |
| GOBP_DNA_CONFORMATION_CHANGE | 306 | 0.129197 | 0.646 | 1 | 1 |
| GOBP_REGULATION_OF_DOUBLE_STRAND_BREAK_REPAIR_VIA_HOMOLOGOUS_REC | 47 | 0.169486 | 0.646 | 0.978261 | 1 |
| GOBP_MITOTIC_CYTOKINESIS | 69 | 0.156161 | 0.641 | 0.98 | 1 |
| GOBP_PROTEIN_LOCALIZATION_TO_CELL_CELL_JUNCTION | 21 | 0.208668 | 0.64 | 0.946524 | 1 |
| GOBP_PROTEIN_LOCALIZATION_TO_CHROMATIN | 29 | 0.193216 | 0.64 | 0.961039 | 1 |
| GOBP_REGULATION_OF_DNA_DAMAGE_CHECKPOINT | 15 | 0.229251 | 0.64 | 0.937239 | 1 |
| GOBP_SIGNAL_TRANSDUCTION_INVOLVED_IN_CELL_CYCLE_CHECKPOINT | 73 | 0.157959 | 0.637 | 1 | 1 |
| GOBP_REGULATION_OF_UBIQUITIN_DEPENDENT_PROTEIN_CATABOLIC_PROCESS | 157 | 0.142402 | 0.637 | 1 | 1 |
| GOBP_LIPOPROTEIN_BIOSYNTHETIC_PROCESS | 92 | 0.154418 | 0.634 | 1 | 1 |
| GOBP_NEGATIVE_REGULATION_OF_ERBB_SIGNALING_PATHWAY | 50 | 0.169386 | 0.634 | 0.987654 | 1 |
| GOBP_INTRINSIC_APOPTOTIC_SIGNALING_PATHWAY_IN_RESPONSE_TO_OXIDATIVE_ | 43 | 0.171959 | 0.634 | 0.977012 | 1 |
| GOBP_PIGMENT_GRANULE_ORGANIZATION | 28 | 0.192167 | 0.632 | 0.956204 | 1 |
| GOBP_SYNAPTIC_VESICLE_CYTOSKELETAL_TRANSPORT | 19 | 0.204439 | 0.631 | 0.937824 | 1 |
| GOBP_REGULATION_OF_DNA_TEMPLATED_TRANSCRIPTION_ELONGATION | 51 | 0.160106 | 0.628 | 0.988506 | 1 |
| GOBP_HISTONE_MRNA_METABOLIC_PROCESS | 24 | 0.194246 | 0.628 | 0.981132 | 1 |
| GOBP_MRNA_3_END_PROCESSING | 99 | 0.15111 | 0.627 | 1 | 1 |
| GOBP_MAINTENANCE_OF_PROTEIN_LOCATION_IN_NUCLEUS | 22 | 0.196503 | 0.625 | 0.953488 | 1 |
| GOBP_RESPONSE_TO_MITOCHONDRIAL_DEPOLARISATION | 20 | 0.206176 | 0.624 | 0.966981 | 1 |
| GOBP_SPLICEOSOMAL_TRI_SNRNP_COMPLEX_ASSEMBLY | 18 | 0.211177 | 0.623 | 0.949438 | 1 |
| GOBP_NUCLEOTIDE_SUGAR_BIOSYNTHETIC_PROCESS | 23 | 0.203222 | 0.622 | 0.96319 | 1 |
| GOBP_RESPONSE_TO_LEAD_ION | 21 | 0.200637 | 0.621 | 0.970414 | 1 |

| | | | | | |
|---|---|---|---|---|---|
| GOBP_PERICARDIUM_DEVELOPMENT | 18 | -0.730688 | -1.677386 | 0.001252 | 0.009733 |
| GOBP_LIPOPOLYSACCHARIDE_MEDIATED_SIGNALING_PATHWAY | 55 | -0.626883 | -1.676562 | 0.001072 | 0.00983 |
| GOBP_REGULATION_OF_CELL_CELL_ADHESION | 398 | -0.568049 | -1.676451 | 0 | 0.009817 |
| GOBP_ENTEROENDOCRINE_CELL_DIFFERENTIATION | 25 | -0.693039 | -1.676242 | 0 | 0.00984 |
| GOBP_SYNAPTIC_MEMBRANE_ADHESION | 24 | -0.700473 | -1.675892 | 0 | 0.00984 |
| GOBP_NEGATIVE_REGULATION_OF_INTERFERON_GAMMA_PRODUCTION | 33 | -0.665877 | -1.674831 | 0.001133 | 0.01005 |
| GOBP_CRANIAL_NERVE_MORPHOGENESIS | 22 | -0.711763 | -1.67441 | 0.001202 | 0.010096 |
| GOBP_ACIDIC_AMINO_ACID_TRANSPORT | 64 | -0.617002 | -1.674202 | 0 | 0.010095 |
| GOBP_POSITIVE_REGULATION_OF_CATION_CHANNEL_ACTIVITY | 69 | -0.600433 | -1.674095 | 0 | 0.010087 |
| GOBP_REGULATION_OF_NEUROTRANSMITTER_TRANSPORT | 96 | -0.595984 | -1.674081 | 0.00103 | 0.010062 |
| GOBP_T_CELL_DIFFERENTIATION_INVOLVED_IN_IMMUNE_RESPONSE | 67 | -0.614235 | -1.673998 | 0 | 0.010058 |
| GOBP_INTERLEUKIN_2_PRODUCTION | 59 | -0.617638 | -1.672708 | 0.001067 | 0.010287 |
| GOBP_COGNITION | 254 | -0.565497 | -1.671976 | 0 | 0.010394 |
| GOBP_CELL_FATE_SPECIFICATION | 78 | -0.600457 | -1.671755 | 0 | 0.010415 |
| GOBP_CALCIUM_MEDIATED_SIGNALING | 183 | -0.574117 | -1.671684 | 0 | 0.010397 |
| GOBP_LEUKOCYTE_HOMEOSTASIS | 78 | -0.607237 | -1.671264 | 0 | 0.010459 |
| GOBP_POSITIVE_REGULATION_OF_RELEASE_OF_SEQUESTERED_CALCIUM_IC | 36 | -0.658614 | -1.671009 | 0.00225 | 0.010489 |
| GOBP_DIVALENT_INORGANIC_CATION_HOMEOSTASIS | 450 | -0.561496 | -1.670976 | 0 | 0.010466 |
| GOBP_POSITIVE_REGULATION_OF_CREB_TRANSCRIPTION_FACTOR_ACTIVITY | 18 | -0.735975 | -1.670505 | 0.006305 | 0.01055 |
| GOBP_TUMOR_NECROSIS_FACTOR_SUPERFAMILY_CYTOKINE_PRODUCTION | 136 | -0.576066 | -1.670375 | 0 | 0.010527 |
| GOBP_ADENYLATE_CYCLASE_MODULATING_G_PROTEIN_COUPLED_RECEPT( | 195 | -0.572626 | -1.670294 | 0 | 0.010521 |
| GOBP_AMINE_TRANSPORT | 85 | -0.601748 | -1.669609 | 0 | 0.01063 |
| GOBP_POSITIVE_REGULATION_OF_NEURON_DIFFERENTIATION | 81 | -0.599847 | -1.669452 | 0 | 0.010633 |
| GOBP_L_GLUTAMATE_IMPORT_ACROSS_PLASMA_MEMBRANE | 15 | -0.753077 | -1.668868 | 0.001274 | 0.01069 |
| GOBP_INORGANIC_ION_IMPORT_ACROSS_PLASMA_MEMBRANE | 92 | -0.587964 | -1.668629 | 0 | 0.010675 |
| GOBP_LONG_TERM_SYNAPTIC_DEPRESSION | 30 | -0.666683 | -1.668051 | 0.001136 | 0.010763 |
| GOBP_GANGLIOSIDE_BIOSYNTHETIC_PROCESS | 15 | -0.753172 | -1.666573 | 0.003704 | 0.011016 |
| GOBP_REGULATION_OF_CD4_POSITIVE_ALPHA_BETA_T_CELL_ACTIVATION | 59 | -0.615531 | -1.665808 | 0.002158 | 0.011144 |
| GOBP_POTASSIUM_ION_HOMEOSTASIS | 27 | -0.68022 | -1.66573 | 0.003601 | 0.011129 |
| GOBP_REGULATION_OF_CALCIUM_ION_DEPENDENT_EXOCYTOSIS | 40 | -0.641731 | -1.664179 | 0.00224 | 0.011418 |
| GOBP_IMPORT_INTO_CELL | 219 | -0.569909 | -1.664113 | 0 | 0.011393 |
| GOBP_NEGATIVE_REGULATION_OF_B_CELL_ACTIVATION | 33 | -0.66384 | -1.663864 | 0.00114 | 0.011416 |
| GOBP_POSITIVE_REGULATION_OF_CATION_TRANSMEMBRANE_TRANSPORT | 134 | -0.579711 | -1.663146 | 0 | 0.011488 |
| GOBP_REGULATION_OF_LIPASE_ACTIVITY | 87 | -0.591601 | -1.662625 | 0 | 0.011555 |
| GOBP_CELLULAR_EXTRAVASATION | 68 | -0.606466 | -1.662088 | 0.001056 | 0.011651 |
| GOBP_HINDBRAIN_DEVELOPMENT | 127 | -0.582586 | -1.662079 | 0 | 0.011618 |
| GOBP_T_HELPER_CELL_LINEAGE_COMMITMENT | 15 | -0.746792 | -1.661777 | 0.003817 | 0.01164 |
| GOBP_MAST_CELL_ACTIVATION | 56 | -0.625608 | -1.661317 | 0 | 0.011675 |
| GOBP_DOPAMINE_TRANSPORT | 43 | -0.640674 | -1.660592 | 0 | 0.011754 |
| GOBP_CELLULAR_RESPONSE_TO_CALCIUM_ION | 79 | -0.598301 | -1.658851 | 0 | 0.012123 |
| GOBP_DICARBOXYLIC_ACID_CATABOLIC_PROCESS | 17 | -0.736929 | -1.658814 | 0.005096 | 0.012097 |
| GOBP_VITAMIN_D_METABOLIC_PROCESS | 18 | -0.721692 | -1.658487 | 0.004963 | 0.012147 |
| GOBP_POSITIVE_REGULATION_OF_GLYCOPROTEIN_METABOLIC_PROCESS | 25 | -0.681874 | -1.658156 | 0.002323 | 0.012183 |
| GOBP_ANTIMICROBIAL_HUMORAL_RESPONSE | 89 | -0.594459 | -1.658044 | 0 | 0.012167 |
| GOBP_REGULATION_OF_IMMUNOGLOBULIN_PRODUCTION | 60 | -0.612872 | -1.657436 | 0 | 0.012262 |
| GOBP_T_CELL_CYTOKINE_PRODUCTION | 33 | -0.657976 | -1.65645 | 0.002288 | 0.012434 |
| GOBP_SENSORY_ORGAN_DEVELOPMENT | 497 | -0.556065 | -1.656279 | 0 | 0.012431 |
| GOBP_RECEPTOR_LOCALIZATION_TO_SYNAPSE | 52 | -0.621643 | -1.655993 | 0.001095 | 0.012456 |
| GOBP_EMBRYONIC_SKELETAL_SYSTEM_DEVELOPMENT | 106 | -0.577522 | -1.655052 | 0 | 0.012626 |
| GOBP_PYROPTOSIS | 18 | -0.721753 | -1.653837 | 0.002506 | 0.012815 |
| GOBP_APOPTOTIC_PROCESS_INVOLVED_IN_MORPHOGENESIS | 22 | -0.701506 | -1.653686 | 0.003636 | 0.012815 |
| GOBP_NEGATIVE_REGULATION_OF_T_CELL_MEDIATED_IMMUNITY | 20 | -0.714481 | -1.653324 | 0.001239 | 0.012853 |
| GOBP_BONE_REMODELING | 83 | -0.591855 | -1.653222 | 0.001043 | 0.012826 |
| GOBP_REGULATION_OF_TISSUE_REMODELING | 65 | -0.600672 | -1.652895 | 0.001062 | 0.012838 |
| GOBP_INTERLEUKIN_10_PRODUCTION | 53 | -0.613654 | -1.652413 | 0 | 0.012918 |
| GOBP_CARDIAC_CONDUCTION | 143 | -0.579863 | -1.651445 | 0 | 0.013084 |
| GOBP_MEMORY | 102 | -0.582618 | -1.651256 | 0 | 0.01308 |
| GOBP_POSITIVE_REGULATION_OF_OLIGODENDROCYTE_DIFFERENTIATION | 22 | -0.701178 | -1.651253 | 0.005051 | 0.013045 |
| GOBP_REGULATION_OF_CHEMOTAXIS | 208 | -0.565144 | -1.651015 | 0 | 0.013046 |
| GOBP_REGULATION_OF_NEUTROPHIL_CHEMOTAXIS | 31 | -0.655119 | -1.650933 | 0 | 0.013026 |
| GOBP_SUBPALLIUM_DEVELOPMENT | 23 | -0.689418 | -1.650129 | 0.001202 | 0.01314 |
| GOBP_REGULATION_OF_CD4_POSITIVE_ALPHA_BETA_T_CELL_DIFFERENTIAT | 47 | -0.633511 | -1.649782 | 0.002162 | 0.013174 |
| GOBP_POSITIVE_REGULATION_OF_ALPHA_BETA_T_CELL_DIFFERENTIATION | 46 | -0.622441 | -1.649704 | 0.00111 | 0.01315 |
| GOBP_SYNAPSE_ORGANIZATION | 398 | -0.557076 | -1.649331 | 0 | 0.013191 |
| GOBP_REGULATION_OF_AMPA_RECEPTOR_ACTIVITY | 26 | -0.680821 | -1.649034 | 0 | 0.013233 |
| GOBP_LIMBIC_SYSTEM_DEVELOPMENT | 84 | -0.593044 | -1.64828 | 0 | 0.013388 |
| GOBP_POSITIVE_REGULATION_OF_NERVOUS_SYSTEM_PROCESS | 31 | -0.665159 | -1.647766 | 0.001166 | 0.013441 |
| GOBP_REGULATION_OF_NERVOUS_SYSTEM_PROCESS | 121 | -0.582799 | -1.647672 | 0 | 0.013423 |
| GOBP_CAMERA_TYPE_EYE_PHOTORECEPTOR_CELL_DIFFERENTIATION | 25 | -0.679202 | -1.647226 | 0.002398 | 0.013486 |
| GOBP_VOCALIZATION_BEHAVIOR | 18 | -0.718835 | -1.646197 | 0.003778 | 0.013679 |
| GOBP_CALCIUM_ION_REGULATED_EXOCYTOSIS | 62 | -0.612878 | -1.646148 | 0 | 0.013654 |
| GOBP_LYMPHOCYTE_APOPTOTIC_PROCESS | 65 | -0.603853 | -1.64613 | 0.001075 | 0.013618 |
| GOBP_SYNAPSE_ASSEMBLY | 165 | -0.56407 | -1.645537 | 0 | 0.013742 |
| GOBP_POSITIVE_REGULATION_OF_ACUTE_INFLAMMATORY_RESPONSE | 23 | -0.689789 | -1.645293 | 0.00361 | 0.013756 |
| GOBP_IONOTROPIC_GLUTAMATE_RECEPTOR_SIGNALING_PATHWAY | 26 | -0.676563 | -1.644409 | 0.002384 | 0.013929 |
| GOBP_DENDRITIC_CELL_DIFFERENTIATION | 45 | -0.625187 | -1.644288 | 0.002186 | 0.013908 |
| GOBP_NEGATIVE_REGULATION_OF_EXOCYTOSIS | 30 | -0.660742 | -1.643951 | 0.002309 | 0.013968 |
| GOBP_DETECTION_OF_BIOTIC_STIMULUS | 32 | -0.65536 | -1.643629 | 0 | 0.014006 |
| GOBP_POSITIVE_REGULATION_OF_CD4_POSITIVE_ALPHA_BETA_T_CELL_ACT | 35 | -0.648612 | -1.643303 | 0 | 0.014047 |
| GOBP_THYMIC_T_CELL_SELECTION | 22 | -0.697663 | -1.641818 | 0.006046 | 0.014382 |
| GOBP_MACROPHAGE_MIGRATION | 52 | -0.617578 | -1.641732 | 0.001089 | 0.01438 |
| GOBP_ANIMAL_ORGAN_FORMATION | 56 | -0.605103 | -1.640849 | 0.001078 | 0.014518 |
| GOBP_REGULATION_OF_BONE_MINERALIZATION | 71 | -0.596118 | -1.640536 | 0.001065 | 0.014554 |
| GOBP_POSITIVE_REGULATION_OF_SIGNALING_RECEPTOR_ACTIVITY | 41 | -0.634304 | -1.640262 | 0.00339 | 0.014581 |
| GOBP_CALCIUM_ION_IMPORT_INTO_CYTOSOL | 20 | -0.701516 | -1.639902 | 0.0025 | 0.014621 |
| GOBP_SIALYLATION | 20 | -0.704908 | -1.639837 | 0 | 0.014594 |
| GOBP_POSITIVE_REGULATION_OF_NEUTROPHIL_MIGRATION | 29 | -0.663733 | -1.63956 | 0.007946 | 0.014639 |
| GOBP_POSITIVE_REGULATION_OF_TYPE_2_IMMUNE_RESPONSE | 16 | -0.743568 | -1.63901 | 0.002506 | 0.014725 |
| GOBP_INFLAMMATORY_RESPONSE_TO_ANTIGENIC_STIMULUS | 45 | -0.620575 | -1.637907 | 0 | 0.014948 |
| GOBP_ACTION_POTENTIAL | 126 | -0.575399 | -1.637904 | 0 | 0.014911 |
| GOBP_RECEPTOR_INTERNALIZATION | 107 | -0.577195 | -1.637211 | 0 | 0.015062 |
| GOBP_REGULATION_OF_B_CELL_MEDIATED_IMMUNITY | 52 | -0.609112 | -1.63712 | 0 | 0.015049 |
| GOBP_RETROGRADE_AXONAL_TRANSPORT | 21 | -0.695668 | -1.63705 | 0.007134 | 0.015038 |
| GOBP_MIDDLE_EAR_MORPHOGENESIS | 16 | -0.73259 | -1.636229 | 0.002594 | 0.015199 |
| GOBP_REGULATION_OF_INFLAMMATORY_RESPONSE | 302 | -0.553645 | -1.636094 | 0 | 0.015188 |
| GOBP_REGULATION_OF_TOLERANCE_INDUCTION | 17 | -0.722638 | -1.636058 | 0 | 0.015165 |
| GOBP_EMBRYONIC_SKELETAL_SYSTEM_MORPHOGENESIS | 80 | -0.58531 | -1.635911 | 0 | 0.015173 |
| GOBP_REGULATION_OF_POSTSYNAPTIC_MEMBRANE_POTENTIAL | 125 | -0.571575 | -1.635474 | 0 | 0.015255 |
| GOBP_DICARBOXYLIC_ACID_TRANSPORT | 87 | -0.58888 | -1.635272 | 0 | 0.01526 |
| GOBP_GLOMERULUS_DEVELOPMENT | 57 | -0.610149 | -1.634815 | 0 | 0.015381 |
| GOBP_NEGATIVE_REGULATION_OF_LEUKOCYTE_CHEMOTAXIS | 18 | -0.713254 | -1.634646 | 0.002538 | 0.015365 |
| GOBP_REGULATION_OF_LYMPHOCYTE_APOPTOTIC_PROCESS | 49 | -0.621877 | -1.634395 | 0.001112 | 0.015374 |
| GOBP_CENTRAL_NERVOUS_SYSTEM_PROJECTION_NEURON_AXONOGENESIS | 26 | -0.674301 | -1.633433 | 0.006002 | 0.015591 |
| GOBP_RESPONSE_TO_TYPE_I_INTERFERON | 84 | -0.579529 | -1.633358 | 0.00105 | 0.015562 |
| GOBP_DEFENSE_RESPONSE_TO_GRAM_POSITIVE_BACTERIUM | 65 | -0.601669 | -1.633051 | 0.001067 | 0.015601 |
| GOBP_NEUROMUSCULAR_PROCESS | 81 | -0.589873 | -1.632361 | 0.001059 | 0.015791 |
| GOBP_POSITIVE_REGULATION_OF_MYELOID_LEUKOCYTE_DIFFERENTIATION | 48 | -0.616959 | -1.63182 | 0.002188 | 0.015854 |
| GOBP_REGULATION_OF_T_HELPER_1_TYPE_IMMUNE_RESPONSE | 24 | -0.671231 | -1.631745 | 0.003555 | 0.015842 |
| GOBP_REGULATION_OF_TOLL_LIKE_RECEPTOR_SIGNALING_PATHWAY | 68 | -0.587363 | -1.631044 | 0 | 0.01597 |
| GOBP_REGULATION_OF_CELLULAR_EXTRAVASATION | 32 | -0.64459 | -1.63099 | 0.002302 | 0.015955 |
| GOBP_REGULATION_OF_PRESYNAPSE_ORGANIZATION | 29 | -0.650539 | -1.630606 | 0.002387 | 0.016035 |
| GOBP_ENDODERM_FORMATION | 51 | -0.608447 | -1.629827 | 0.005495 | 0.016178 |
| GOBP_LEUKOCYTE_TETHERING_OR_ROLLING | 29 | -0.66403 | -1.629062 | 0.002389 | 0.016325 |
| GOBP_EXPORT_ACROSS_PLASMA_MEMBRANE | 56 | -0.600865 | -1.628848 | 0.002176 | 0.016332 |
| GOBP_REGULATION_OF_NEUROTRANSMITTER_LEVELS | 202 | -0.558884 | -1.628795 | 0 | 0.016307 |
| GOBP_POSITIVE_REGULATION_OF_IMMUNOGLOBULIN_PRODUCTION | 39 | -0.632922 | -1.628014 | 0.002242 | 0.016491 |
| GOBP_CELL_CELL_SIGNALING_INVOLVED_IN_CARDIAC_CONDUCTION | 31 | -0.637846 | -1.627681 | 0.009195 | 0.016545 |
| GOBP_MONOAMINE_TRANSPORT | 71 | -0.59172 | -1.62761 | 0.001054 | 0.016533 |
| GOBP_POSITIVE_REGULATION_OF_NUCLEOTIDE_METABOLIC_PROCESS | 42 | -0.619291 | -1.627555 | 0.003356 | 0.016513 |
| GOBP_SECOND_MESSENGER_MEDIATED_SIGNALING | 278 | -0.553979 | -1.62699 | 0 | 0.016637 |
| GOBP_POSITIVE_REGULATION_OF_T_CELL_CYTOKINE_PRODUCTION | 20 | -0.697596 | -1.626472 | 0.008495 | 0.016753 |
| GOBP_POSITIVE_REGULATION_OF_CALCIUM_ION_TRANSMEMBRANE_TRANSF | 65 | -0.59382 | -1.62616 | 0.002128 | 0.016794 |
| GOBP_PURINERGIC_NUCLEOTIDE_RECEPTOR_SIGNALING_PATHWAY | 29 | -0.663768 | -1.626152 | 0.002323 | 0.016758 |
| GOBP_POSITIVE_REGULATION_OF_SMALL_GTPASE_MEDIATED_SIGNAL_TRAN | 74 | -0.591868 | -1.625974 | 0 | 0.01677 |
| GOBP_COLLAGEN_CATABOLIC_PROCESS | 41 | -0.629724 | -1.624757 | 0.001106 | 0.017078 |
| GOBP_DETECTION_OF_STIMULUS | 263 | -0.553952 | -1.624693 | 0 | 0.01705 |
| GOBP_REGULATION_OF_CALCIUM_ION_TRANSMEMBRANE_TRANSPORTER_A | 78 | -0.583222 | -1.623738 | 0 | 0.017262 |
| GOBP_POSITIVE_REGULATION_OF_CD4_POSITIVE_ALPHA_BETA_T_CELL_DIFF | 29 | -0.662161 | -1.62292 | 0.003534 | 0.017436 |
| GOBP_STRIATUM_DEVELOPMENT | 17 | -0.70972 | -1.621692 | 0.010243 | 0.017706 |
| GOBP_NEPHRON_MORPHOGENESIS | 67 | -0.591283 | -1.621671 | 0 | 0.01767 |
| GOBP_DETECTION_OF_TEMPERATURE_STIMULUS | 20 | -0.711659 | -1.620467 | 0.004802 | 0.017977 |
| GOBP_POSITIVE_REGULATION_OF_CELL_JUNCTION_ASSEMBLY | 93 | -0.573682 | -1.61976 | 0 | 0.018158 |
| GOBP_POSITIVE_REGULATION_OF_PROTEIN_TYROSINE_KINASE_ACTIVITY | 58 | -0.600566 | -1.619442 | 0.005376 | 0.018215 |
| GOBP_REGULATION_OF_GLYCOPROTEIN_METABOLIC_PROCESS | 46 | -0.617285 | -1.618991 | 0.002217 | 0.018294 |
| GOBP_ADAPTATION_OF_SIGNALING_PATHWAY | 19 | -0.699706 | -1.618976 | 0.003778 | 0.018258 |
| GOBP_ODONTOGENESIS | 112 | -0.56957 | -1.61889 | 0 | 0.018249 |
| GOBP_REGULATION_OF_REGULATED_SECRETORY_PATHWAY | 132 | -0.564129 | -1.618774 | 0 | 0.018243 |
| GOBP_REGULATION_OF_CALCIUM_ION_TRANSPORT_INTO_CYTOSOL | 87 | -0.576827 | -1.61801 | 0 | 0.018441 |
| GOBP_NEGATIVE_REGULATION_OF_CELL_CELL_ADHESION | 164 | -0.555433 | -1.617894 | 0 | 0.018436 |
| GOBP_RESPONSE_TO_NICOTINE | 38 | -0.624179 | -1.617198 | 0.002278 | 0.018567 |
| GOBP_NEGATIVE_REGULATION_OF_LYMPHOCYTE_MIGRATION | 15 | -0.728569 | -1.616755 | 0.00641 | 0.018673 |
| GOBP_REGULATION_OF_TRANSPORTER_ACTIVITY | 263 | -0.548642 | -1.616497 | 0 | 0.018689 |
| GOBP_INTERLEUKIN_1_BETA_PRODUCTION | 85 | -0.580418 | -1.616194 | 0 | 0.018753 |
| GOBP_UTERUS_DEVELOPMENT | 18 | -0.716486 | -1.616141 | 0.006203 | 0.018725 |
| GOBP_REGULATION_OF_SKELETAL_MUSCLE_CELL_DIFFERENTIATION | 18 | -0.7087 | -1.616097 | 0.004825 | 0.018696 |
| GOBP_POSITIVE_REGULATION_OF_DEFENSE_RESPONSE | 322 | -0.544016 | -1.615408 | 0 | 0.018871 |
| GOBP_RELEASE_OF_SEQUESTERED_CALCIUM_ION_INTO_CYTOSOL_BY_END | 31 | -0.644031 | -1.614588 | 0.001136 | 0.019073 |
| GOBP_IMPORT_ACROSS_PLASMA_MEMBRANE | 153 | -0.55762 | -1.614387 | 0 | 0.019098 |
| GOBP_MACROPHAGE_CHEMOTAXIS | 38 | -0.615448 | -1.61343 | 0.002299 | 0.019298 |
| GOBP_PROXIMAL_DISTAL_PATTERN_FORMATION | 28 | -0.655957 | -1.613275 | 0.005931 | 0.019306 |
| GOBP_REGULATION_OF_MORPHOGENESIS_OF_AN_EPITHELIUM | 56 | -0.597448 | -1.612998 | 0.002144 | 0.019337 |

| | | | | | |
|---|---|---|---|---|---|
| GOBP_MITOTIC_CELL_CYCLE_CHECKPOINT | 156 | 0.141154 | 0.62 | 1 | 1 |
| GOBP_HETEROCHROMATIN_ORGANIZATION | 66 | 0.151835 | 0.619 | 0.985916 | 1 |
| GOBP_PYRIMIDINE_RIBONUCLEOSIDE_TRIPHOSPHATE_BIOSYNTHETIC_PROCESS | 15 | 0.223047 | 0.614 | 0.940171 | 1 |
| GOBP_MITOTIC_RECOMBINATION | 24 | 0.193204 | 0.614 | 0.99422 | 1 |
| GOBP_REGULATION_OF_DOUBLE_STRAND_BREAK_REPAIR | 85 | 0.153461 | 0.614 | 1 | 1 |
| GOBP_POSITIVE_REGULATION_OF_TELOMERE_MAINTENANCE | 51 | 0.164924 | 0.607 | 0.989474 | 1 |
| GOBP_NEGATIVE_REGULATION_OF_TELOMERE_MAINTENANCE | 35 | 0.170696 | 0.607 | 0.991379 | 1 |
| GOBP_POSITIVE_REGULATION_OF_PROTEIN_TARGETING_TO_MITOCHONDRION | 32 | 0.181617 | 0.606 | 0.977941 | 1 |
| GOBP_CELLULAR_PIGMENTATION | 52 | 0.161846 | 0.602 | 1 | 1 |
| GOBP_POSITIVE_REGULATION_OF_TELOMERE_CAPPING | 17 | 0.203959 | 0.601 | 0.955665 | 1 |
| GOBP_TRANSLESION_SYNTHESIS | 42 | 0.16363 | 0.6 | 0.981651 | 1 |
| GOBP_REGULATION_OF_MRNA_3_END_PROCESSING | 28 | 0.184201 | 0.599 | 0.993976 | 1 |
| GOBP_PROTEIN_SUMOYLATION | 77 | 0.141051 | 0.597 | 1 | 1 |
| GOBP_AUTOPHAGOSOME_MATURATION | 40 | 0.161857 | 0.593 | 0.99 | 1 |
| GOBP_NEGATIVE_REGULATION_OF_DNA_REPAIR | 36 | 0.16756 | 0.592 | 0.991597 | 1 |
| GOBP_PROTEIN_LOCALIZATION_TO_CHROMOSOME_CENTROMERIC_REGION | 25 | 0.191732 | 0.591 | 0.985401 | 1 |
| GOBP_HISTONE_EXCHANGE | 51 | 0.159696 | 0.591 | 1 | 1 |
| GOBP_RESOLUTION_OF_MEIOTIC_RECOMBINATION_INTERMEDIATES | 18 | 0.199279 | 0.59 | 0.971698 | 1 |
| GOBP_CARDIOLIPIN_METABOLIC_PROCESS | 18 | 0.199372 | 0.589 | 0.97619 | 1 |
| GOBP_PROTEIN_INSERTION_INTO_ER_MEMBRANE | 19 | 0.198009 | 0.587 | 0.981928 | 1 |
| GOBP_DNA_REPLICATION_CHECKPOINT | 17 | 0.193036 | 0.584 | 0.957143 | 1 |
| GOBP_NUCLEAR_ENVELOPE_ORGANIZATION | 51 | 0.154373 | 0.584 | 0.973333 | 1 |
| GOBP_PYRIMIDINE_RIBONUCLEOSIDE_TRIPHOSPHATE_METABOLIC_PROCESS | 18 | 0.197704 | 0.583 | 0.967213 | 1 |
| GOBP_RECOMBINATIONAL_REPAIR | 137 | 0.13019 | 0.583 | 1 | 1 |
| GOBP_POSITIVE_REGULATION_OF_TELOMERE_MAINTENANCE_VIA_TELOMERE_LENG | 36 | 0.164814 | 0.581 | 1 | 1 |
| GOBP_ORGANELLE_INHERITANCE | 15 | 0.206291 | 0.58 | 0.980861 | 1 |
| GOBP_POSITIVE_REGULATION_OF_CHROMOSOME_ORGANIZATION | 161 | 0.127175 | 0.577 | 1 | 1 |
| GOBP_PROTEIN_QUALITY_CONTROL_FOR_MISFOLDED_OR_INCOMPLETELY_SYNTHE | 28 | 0.170422 | 0.576 | 0.985401 | 1 |
| GOBP_FORMATION_OF_EXTRACHROMOSOMAL_CIRCULAR_DNA | 15 | 0.208547 | 0.575 | 0.974138 | 1 |
| GOBP_CENTRIOLE_ASSEMBLY | 44 | 0.158445 | 0.574 | 1 | 1 |
| GOBP_CENTROMERE_COMPLEX_ASSEMBLY | 52 | 0.146803 | 0.573 | 1 | 1 |
| GOBP_NUCLEAR_ENVELOPE_REASSEMBLY | 18 | 0.196828 | 0.571 | 0.955801 | 1 |
| GOBP_PROTEIN_LOCALIZATION_TO_CHROMOSOME_TELOMERIC_REGION | 30 | 0.170658 | 0.57 | 1 | 1 |
| GOBP_MODULATION_BY_SYMBIONT_OF_HOST_CELLULAR_PROCESS | 22 | 0.183661 | 0.567 | 0.977528 | 1 |
| GOBP_NEGATIVE_REGULATION_OF_MRNA_SPLICING_VIA_SPLICEOSOME | 20 | 0.185206 | 0.564 | 0.978836 | 1 |
| GOBP_REGULATION_OF_UBIQUITIN_PROTEIN_TRANSFERASE_ACTIVITY | 50 | 0.148359 | 0.562 | 1 | 1 |
| GOBP_NEGATIVE_REGULATION_OF_RNA_SPLICING | 25 | 0.174291 | 0.56 | 0.993631 | 1 |
| GOBP_DNA_INTEGRITY_CHECKPOINT | 151 | 0.12085 | 0.557 | 1 | 1 |
| GOBP_MITOCHONDRIAL_MEMBRANE_ORGANIZATION | 139 | 0.127838 | 0.556 | 1 | 1 |
| GOBP_RRNA_METHYLATION | 28 | 0.169448 | 0.556 | 0.985816 | 1 |
| GOBP_MITOCHONDRIAL_TRANSPORT | 264 | 0.12284 | 0.554 | 1 | 1 |
| GOBP_CHAPERONE_COFACTOR_DEPENDENT_PROTEIN_REFOLDING | 34 | 0.161159 | 0.553 | 1 | 1 |
| GOBP_NEGATIVE_REGULATION_OF_TELOMERE_MAINTENANCE_VIA_TELOMERE_LEN | 27 | 0.170636 | 0.547 | 0.99359 | 1 |
| GOBP_NEGATIVE_REGULATION_OF_TELOMERE_MAINTENANCE_VIA_TELOMERASE | 20 | 0.182139 | 0.544 | 0.993939 | 1 |
| GOBP_POSITIVE_REGULATION_OF_TELOMERASE_ACTIVITY | 33 | 0.157221 | 0.544 | 1 | 1 |
| GOBP_MITOCHONDRIAL_FUSION | 28 | 0.164225 | 0.543 | 1 | 1 |
| GOBP_ATP_DEPENDENT_CHROMATIN_REMODELING | 82 | 0.126848 | 0.542 | 1 | 1 |
| GOBP_MICROTUBULE_ORGANIZING_CENTER_ORGANIZATION | 139 | 0.120707 | 0.541 | 1 | 1 |
| GOBP_TETRAPYRROLE_BIOSYNTHETIC_PROCESS | 28 | 0.160785 | 0.539 | 1 | 1 |
| GOBP_NON_RECOMBINATIONAL_REPAIR | 100 | 0.125683 | 0.536 | 1 | 1 |
| GOBP_INTRA_GOLGI_VESICLE_MEDIATED_TRANSPORT | 33 | 0.156424 | 0.535 | 0.986577 | 1 |
| GOBP_REGULATION_OF_TELOMERE_CAPPING | 26 | 0.164741 | 0.525 | 0.994152 | 1 |
| GOBP_DNA_REPLICATION_INDEPENDENT_NUCLEOSOME_ORGANIZATION | 50 | 0.136981 | 0.524 | 1 | 1 |
| GOBP_RRNA_CONTAINING_RIBONUCLEOPROTEIN_COMPLEX_EXPORT_FROM_NUCLE | 15 | 0.183153 | 0.523 | 0.971545 | 1 |
| GOBP_METAPHASE_ANAPHASE_TRANSITION_OF_CELL_CYCLE | 62 | 0.133002 | 0.518 | 1 | 1 |
| GOBP_PROTEIN_LOCALIZATION_TO_CHROMOSOME | 90 | 0.121696 | 0.515 | 1 | 1 |
| GOBP_DNA_GEOMETRIC_CHANGE | 112 | 0.118836 | 0.508 | 1 | 1 |
| GOBP_TELOMERE_MAINTENANCE_VIA_RECOMBINATION | 15 | 0.177884 | 0.506 | 0.990431 | 1 |
| GOBP_ER_NUCLEUS_SIGNALING_PATHWAY | 50 | 0.128229 | 0.503 | 1 | 1 |
| GOBP_TELOMERE_MAINTENANCE_VIA_SEMI_CONSERVATIVE_REPLICATION | 27 | 0.153534 | 0.498 | 1 | 1 |
| GOBP_NEGATIVE_REGULATION_OF_SINGLE_STRANDED_VIRAL_RNA_REPLICATION_V | 15 | 0.182755 | 0.496 | 0.995496 | 1 |
| GOBP_REGULATION_OF_CHROMATIN_ASSEMBLY_OR_DISASSEMBLY | 23 | 0.159844 | 0.494 | 1 | 1 |
| GOBP_MITOTIC_CHROMOSOME_CONDENSATION | 15 | 0.173421 | 0.484 | 0.990698 | 1 |
| GOBP_CILIARY_BASAL_BODY_PLASMA_MEMBRANE_DOCKING | 95 | 0.115505 | 0.482 | 1 | 1 |
| GOBP_TRNA_TRANSPORT | 37 | 0.132833 | 0.479 | 1 | 1 |
| GOBP_PROTEIN_MONOUBIQUITINATION | 66 | 0.119378 | 0.472 | 1 | 1 |
| GOBP_P_BODY_ASSEMBLY | 21 | 0.149283 | 0.465 | 1 | 1 |
| GOBP_NEGATIVE_REGULATION_OF_MRNA_METABOLIC_PROCESS | 79 | 0.113726 | 0.461 | 1 | 1 |
| GOBP_DE_NOVO_PROTEIN_FOLDING | 43 | 0.126658 | 0.459 | 1 | 1 |
| GOBP_UBIQUITIN_DEPENDENT_ERAD_PATHWAY | 78 | 0.109955 | 0.457 | 1 | 1 |
| GOBP_CELL_CYCLE_DNA_REPLICATION | 63 | 0.111878 | 0.451 | 1 | 1 |
| GOBP_RNA_EXPORT_FROM_NUCLEUS | 136 | 0.107056 | 0.45 | 1 | 1 |
| GOBP_NUCLEAR_EXPORT | 195 | 0.096934 | 0.446 | 1 | 1 |
| GOBP_POSITIVE_REGULATION_OF_RESPONSE_TO_ENDOPLASMIC_RETICULUM_STRE | 33 | 0.130792 | 0.443 | 1 | 1 |
| GOBP_REGULATION_OF_AUTOPHAGOSOME_ASSEMBLY | 39 | 0.123929 | 0.438 | 1 | 1 |
| GOBP_RNA_DECAPPING | 16 | 0.144203 | 0.425 | 1 | 1 |
| GOBP_REGULATION_OF_VACUOLE_ORGANIZATION | 47 | 0.105216 | 0.405 | 1 | 1 |
| GOBP_NCRNA_EXPORT_FROM_NUCLEUS | 38 | 0.102663 | 0.361 | 1 | 0.99999 |

| | | | | | |
|---|---|---|---|---|---|
| GOBP_NEUROMUSCULAR_PROCESS_CONTROLLING_BALANCE | 38 | -0.631008 | -1.612935 | 0.004484 | 0.019322 |
| GOBP_REGULATION_OF_MYELOID_LEUKOCYTE_DIFFERENTIATION | 102 | -0.568058 | -1.61276 | 0 | 0.019345 |
| GOBP_KIDNEY_MESENCHYME_DEVELOPMENT | 18 | -0.700523 | -1.612476 | 0.01627 | 0.019404 |
| GOBP_N_GLYCAN_PROCESSING | 19 | -0.700638 | -1.612429 | 0.004896 | 0.019367 |
| GOBP_INTEGRIN_ACTIVATION | 25 | -0.666987 | -1.610617 | 0.003584 | 0.019921 |
| GOBP_SKELETAL_SYSTEM_MORPHOGENESIS | 192 | -0.548183 | -1.609888 | 0 | 0.020106 |
| GOBP_INTERLEUKIN_4_PRODUCTION | 31 | -0.639505 | -1.609415 | 0.006936 | 0.02021 |
| GOBP_REGULATION_OF_DENDRITE_EXTENSION | 25 | -0.661275 | -1.60764 | 0.003667 | 0.020733 |
| GOBP_CARDIAC_MUSCLE_CELL_ACTION_POTENTIAL | 71 | -0.584889 | -1.607247 | 0 | 0.020803 |
| GOBP_REGULATION_OF_HEMOPOIESIS | 373 | -0.543601 | -1.607182 | 0 | 0.020769 |
| GOBP_VASCULAR_WOUND_HEALING | 16 | -0.72011 | -1.60718 | 0.005013 | 0.020725 |
| GOBP_VESICLE_MEDIATED_TRANSPORT_IN_SYNAPSE | 190 | -0.547594 | -1.607055 | 0 | 0.02072 |
| GOBP_DOPAMINE_SECRETION | 30 | -0.644624 | -1.606592 | 0.006897 | 0.020806 |
| GOBP_NEGATIVE_REGULATION_OF_GLIAL_CELL_DIFFERENTIATION | 17 | -0.706118 | -1.60612 | 0.007398 | 0.020933 |
| GOBP_CARDIAC_ATRIUM_MORPHOGENESIS | 26 | -0.660513 | -1.606009 | 0.006083 | 0.020924 |
| GOBP_T_CELL_HOMEOSTASIS | 35 | -0.635319 | -1.605753 | 0.003444 | 0.020966 |
| GOBP_APPENDAGE_MORPHOGENESIS | 125 | -0.558947 | -1.605415 | 0 | 0.021011 |
| GOBP_INTERLEUKIN_6_PRODUCTION | 127 | -0.559074 | -1.605089 | 0 | 0.021065 |
| GOBP_REGULATION_OF_OSSIFICATION | 103 | -0.562053 | -1.603906 | 0 | 0.021359 |
| GOBP_POSITIVE_REGULATION_OF_RESPONSE_TO_EXTERNAL_STIMULUS | 453 | -0.543156 | -1.603598 | 0 | 0.021418 |
| GOBP_REGULATION_OF_KIDNEY_DEVELOPMENT | 27 | -0.65413 | -1.602766 | 0.005889 | 0.021711 |
| GOBP_REGULATION_OF_MONOCYTE_DIFFERENTIATION | 18 | -0.688192 | -1.602087 | 0.008578 | 0.021903 |
| GOBP_REGULATION_OF_BIOMINERALIZATION | 86 | -0.57838 | -1.601866 | 0 | 0.021927 |
| GOBP_REGULATION_OF_LONG_TERM_SYNAPTIC_POTENTIATION | 43 | -0.61862 | -1.601858 | 0 | 0.021887 |
| GOBP_REGULATION_OF_MEMBRANE_POTENTIAL | 404 | -0.541682 | -1.60161 | 0 | 0.021926 |
| GOBP_PHARYNGEAL_SYSTEM_DEVELOPMENT | 22 | -0.689141 | -1.600982 | 0.002433 | 0.022093 |
| GOBP_HETEROTYPIC_CELL_CELL_ADHESION | 56 | -0.592054 | -1.600747 | 0 | 0.022136 |
| GOBP_REGULATION_OF_CYCLASE_ACTIVITY | 42 | -0.616186 | -1.600434 | 0.005501 | 0.022209 |
| GOBP_FORMATION_OF_PRIMARY_GERM_LAYER | 105 | -0.566145 | -1.600345 | 0 | 0.022193 |
| GOBP_ENDODERMAL_CELL_DIFFERENTIATION | 42 | -0.610327 | -1.599914 | 0.002208 | 0.02229 |
| GOBP_POSITIVE_REGULATION_OF_RHO_PROTEIN_SIGNAL_TRANSDUCTION | 32 | -0.636281 | -1.599904 | 0.004556 | 0.022247 |
| GOBP_REGULATION_OF_NATURAL_KILLER_CELL_ACTIVATION | 28 | -0.651417 | -1.599803 | 0.001149 | 0.022235 |
| GOBP_DEFENSE_RESPONSE_TO_GRAM_NEGATIVE_BACTERIUM | 50 | -0.597149 | -1.599443 | 0.002181 | 0.022297 |
| GOBP_RESPONSE_TO_DIETARY_EXCESS | 20 | -0.699462 | -1.599092 | 0.00618 | 0.022358 |
| GOBP_FOREBRAIN_DEVELOPMENT | 328 | -0.543852 | -1.598929 | 0 | 0.022366 |
| GOBP_POTASSIUM_ION_TRANSPORT | 211 | -0.550342 | -1.598757 | 0 | 0.022367 |
| GOBP_ADULT_BEHAVIOR | 107 | -0.558791 | -1.598449 | 0 | 0.02242 |
| GOBP_POSITIVE_REGULATION_OF_BONE_MINERALIZATION | 38 | -0.617174 | -1.598367 | 0.005688 | 0.022406 |
| GOBP_POSTSYNAPTIC_NEUROTRANSMITTER_RECEPTOR_INTERNALIZATION | 17 | -0.708531 | -1.596609 | 0.007426 | 0.022969 |
| GOBP_TONGUE_DEVELOPMENT | 19 | -0.700917 | -1.596258 | 0.007435 | 0.023038 |
| GOBP_ADENYLATE_CYCLASE_INHIBITING_G_PROTEIN_COUPLED_RECEPTOR | 73 | -0.572315 | -1.595903 | 0.002073 | 0.023133 |
| GOBP_POSITIVE_REGULATION_OF_OSSIFICATION | 47 | -0.611149 | -1.595541 | 0.007692 | 0.023196 |
| GOBP_NEGATIVE_REGULATION_OF_INTERLEUKIN_2_PRODUCTION | 27 | -0.649242 | -1.595376 | 0.009592 | 0.023201 |
| GOBP_PEPTIDE_CROSS_LINKING | 28 | -0.650727 | -1.595193 | 0.004802 | 0.02322 |
| GOBP_CELL_FATE_COMMITMENT | 219 | -0.543709 | -1.595154 | 0 | 0.02319 |
| GOBP_TYPE_B_PANCREATIC_CELL_DEVELOPMENT | 20 | -0.685413 | -1.594817 | 0.01087 | 0.023253 |
| GOBP_FOREBRAIN_GENERATION_OF_NEURONS | 50 | -0.599744 | -1.594719 | 0.0022 | 0.023251 |
| GOBP_REGULATION_OF_OLIGODENDROCYTE_DIFFERENTIATION | 35 | -0.626509 | -1.594223 | 0.002294 | 0.02336 |
| GOBP_L_ALPHA_AMINO_ACID_TRANSMEMBRANE_TRANSPORT | 58 | -0.592082 | -1.593826 | 0.001074 | 0.023473 |
| GOBP_DETECTION_OF_EXTERNAL_BIOTIC_STIMULUS | 21 | -0.669546 | -1.593695 | 0.006098 | 0.023473 |
| GOBP_POSITIVE_REGULATION_OF_PEPTIDYL_TYROSINE_PHOSPHORYLATION | 169 | -0.5453 | -1.593616 | 0 | 0.023455 |
| GOBP_VENTRICULAR_CARDIAC_MUSCLE_CELL_MEMBRANE_REPOLARIZATIO | 23 | -0.667967 | -1.593108 | 0.007194 | 0.023585 |
| GOBP_REGULATION_OF_VESICLE_FUSION | 23 | -0.681073 | -1.59302 | 0.003659 | 0.023556 |
| GOBP_SMOOTH_MUSCLE_CONTRACTION | 87 | -0.566073 | -1.591897 | 0.001022 | 0.02397 |
| GOBP_POSITIVE_REGULATION_OF_BONE_RESORPTION | 17 | -0.70477 | -1.591761 | 0.008794 | 0.023987 |
| GOBP_APOPTOTIC_PROCESS_INVOLVED_IN_DEVELOPMENT | 34 | -0.621541 | -1.58954 | 0.010345 | 0.024755 |
| GOBP_REGULATION_OF_GLIAL_CELL_DIFFERENTIATION | 61 | -0.584619 | -1.589137 | 0.004287 | 0.024854 |
| GOBP_MACROPHAGE_ACTIVATION_INVOLVED_IN_IMMUNE_RESPONSE | 17 | -0.709082 | -1.589124 | 0.008805 | 0.02481 |
| GOBP_NEURON_FATE_COMMITMENT | 49 | -0.602222 | -1.588601 | 0.001106 | 0.02492 |
| GOBP_CEREBRAL_CORTEX_NEURON_DIFFERENTIATION | 19 | -0.690857 | -1.5881 | 0.01148 | 0.025053 |
| GOBP_CELLULAR_RESPONSE_TO_MOLECULE_OF_BACTERIAL_ORIGIN | 171 | -0.548191 | -1.587994 | 0 | 0.025036 |
| GOBP_NEUROTROPHIN_SIGNALING_PATHWAY | 37 | -0.616171 | -1.587851 | 0.013621 | 0.025063 |
| GOBP_POSITIVE_REGULATION_OF_ANTIGEN_RECEPTOR_MEDIATED_SIGNALIN | 21 | -0.685025 | -1.587378 | 0.009889 | 0.025196 |
| GOBP_NEPHRON_DEVELOPMENT | 129 | -0.559983 | -1.587162 | 0 | 0.025223 |
| GOBP_ANTIGEN_PROCESSING_AND_PRESENTATION_OF_ENDOGENOUS_ANTI | 25 | -0.664924 | -1.587118 | 0.004624 | 0.025185 |
| GOBP_CALCIUM_ION_TRANSPORT | 375 | -0.538754 | -1.58656 | 0 | 0.025351 |
| GOBP_POSITIVE_REGULATION_OF_LIPID_KINASE_ACTIVITY | 36 | -0.621284 | -1.586371 | 0.004561 | 0.025361 |
| GOBP_ANTERIOR_POSTERIOR_PATTERN_SPECIFICATION | 171 | -0.547073 | -1.585772 | 0 | 0.025512 |
| GOBP_CELLULAR_RESPONSE_TO_THYROID_HORMONE_STIMULUS | 15 | -0.708392 | -1.585738 | 0.011658 | 0.025483 |
| GOBP_CENTRAL_NERVOUS_SYSTEM_NEURON_DIFFERENTIATION | 165 | -0.543407 | -1.585615 | 0 | 0.025489 |
| GOBP_VITAMIN_BIOSYNTHETIC_PROCESS | 19 | -0.686792 | -1.585533 | 0.007426 | 0.025474 |
| GOBP_SODIUM_ION_TRANSMEMBRANE_TRANSPORT | 154 | -0.547806 | -1.584048 | 0 | 0.025971 |
| GOBP_BEHAVIOR | 486 | -0.530507 | -1.583836 | 0 | 0.026032 |
| GOBP_POSITIVE_REGULATION_OF_EXCITATORY_POSTSYNAPTIC_POTENTIAL | 27 | -0.639998 | -1.583436 | 0.010664 | 0.026126 |
| GOBP_CELL_COMMUNICATION_INVOLVED_IN_CARDIAC_CONDUCTION | 55 | -0.583587 | -1.582695 | 0.002158 | 0.026318 |
| GOBP_REGULATION_OF_MACROPHAGE_ACTIVATION | 52 | -0.590027 | -1.582188 | 0.0022 | 0.026482 |
| GOBP_REGULATION_OF_G_PROTEIN_COUPLED_RECEPTOR_SIGNALING_PATH | 135 | -0.552843 | -1.581835 | 0 | 0.026559 |
| GOBP_REGULATION_OF_ACUTE_INFLAMMATORY_RESPONSE | 42 | -0.618023 | -1.581802 | 0.004367 | 0.026517 |
| GOBP_POSITIVE_REGULATION_OF_LEUKOCYTE_APOPTOTIC_PROCESS | 23 | -0.670224 | -1.581604 | 0.006068 | 0.026542 |
| GOBP_DENDRITE_EXTENSION | 34 | -0.635367 | -1.581585 | 0.002275 | 0.026506 |
| GOBP_NEGATIVE_REGULATION_OF_LEUKOCYTE_MEDIATED_CYTOTOXICITY | 21 | -0.672089 | -1.580879 | 0.014833 | 0.026724 |
| GOBP_ERK1_AND_ERK2_CASCADE | 276 | -0.53796 | -1.580402 | 0 | 0.026867 |
| GOBP_REGULATION_OF_INFLAMMATORY_RESPONSE_TO_ANTIGENIC_STIMUL | 26 | -0.641142 | -1.580255 | 0.006928 | 0.026874 |
| GOBP_PHOSPHATIDYLINOSITOL_3_KINASE_SIGNALING | 141 | -0.54633 | -1.580111 | 0 | 0.026867 |
| GOBP_REGULATION_OF_LEUKOCYTE_ADHESION_TO_VASCULAR_ENDOTHELI | 27 | -0.643736 | -1.579371 | 0.004624 | 0.027072 |
| GOBP_NEGATIVE_REGULATION_OF_INTERLEUKIN_6_PRODUCTION | 40 | -0.609208 | -1.579268 | 0.004484 | 0.027056 |
| GOBP_EXTERNAL_ENCAPSULATING_STRUCTURE_ORGANIZATION | 369 | -0.533895 | -1.579058 | 0 | 0.027083 |
| GOBP_POSITIVE_REGULATION_OF_BIOMINERALIZATION | 45 | -0.592075 | -1.579045 | 0.008696 | 0.02704 |
| GOBP_RENAL_SYSTEM_PROCESS | 107 | -0.556965 | -1.578805 | 0 | 0.027071 |
| GOBP_CHONDROITIN_SULFATE_PROTEOGLYCAN_METABOLIC_PROCESS | 42 | -0.607994 | -1.578584 | 0.003272 | 0.027116 |
| GOBP_NEUROTROPHIN_TRK_RECEPTOR_SIGNALING_PATHWAY | 31 | -0.643409 | -1.578537 | 0.004608 | 0.027083 |
| GOBP_RESPONSE_TO_MOLECULE_OF_BACTERIAL_ORIGIN | 296 | -0.533877 | -1.577998 | 0 | 0.027253 |
| GOBP_NEGATIVE_REGULATION_OF_CELL_ADHESION | 259 | -0.534085 | -1.57745 | 0 | 0.027446 |
| GOBP_NEGATIVE_REGULATION_OF_OSTEOCLAST_DIFFERENTIATION | 27 | -0.650375 | -1.577156 | 0.005903 | 0.02751 |
| GOBP_GLYCOSPHINGOLIPID_BIOSYNTHETIC_PROCESS | 28 | -0.64641 | -1.575968 | 0.003567 | 0.027918 |
| GOBP_POSITIVE_REGULATION_OF_PROTEIN_KINASE_B_SIGNALING | 152 | -0.545871 | -1.575851 | 0 | 0.027913 |
| GOBP_NEGATIVE_REGULATION_OF_CELL_KILLING | 24 | -0.660306 | -1.57585 | 0.011891 | 0.027863 |
| GOBP_COCHLEA_DEVELOPMENT | 41 | -0.608225 | -1.575513 | 0.003333 | 0.027956 |
| GOBP_SYNAPTIC_VESICLE_EXOCYTOSIS | 103 | -0.551772 | -1.575365 | 0.00102 | 0.027978 |
| GOBP_CELLULAR_RESPONSE_TO_LIGHT_STIMULUS | 126 | -0.55121 | -1.57365 | 0 | 0.028627 |
| GOBP_VASCULAR_ASSOCIATED_SMOOTH_MUSCLE_CONTRACTION | 23 | -0.649526 | -1.573594 | 0.01348 | 0.028605 |
| GOBP_B_CELL_ACTIVATION_INVOLVED_IN_IMMUNE_RESPONSE | 73 | -0.577865 | -1.573323 | 0.002105 | 0.028682 |
| GOBP_NEGATIVE_REGULATION_OF_TUMOR_NECROSIS_FACTOR_SUPERFAMI | 53 | -0.595556 | -1.572361 | 0.001094 | 0.02899 |
| GOBP_REGULATION_OF_GRANULOCYTE_CHEMOTAXIS | 48 | -0.598071 | -1.572281 | 0.003268 | 0.028978 |
| GOBP_STARTLE_RESPONSE | 20 | -0.670712 | -1.572192 | 0.012469 | 0.028958 |
| GOBP_POSITIVE_REGULATION_OF_CYTOSOLIC_CALCIUM_ION_CONCENTRATI | 30 | -0.63138 | -1.5719 | 0.004619 | 0.029019 |
| GOBP_SPINAL_CORD_DEVELOPMENT | 88 | -0.556403 | -1.57157 | 0.001031 | 0.029112 |
| GOBP_METENCEPHALON_DEVELOPMENT | 93 | -0.559631 | -1.571382 | 0 | 0.029161 |
| GOBP_METANEPHRIC_NEPHRON_DEVELOPMENT | 36 | -0.614031 | -1.571347 | 0.009174 | 0.029122 |
| GOBP_BRAIN_MORPHOGENESIS | 35 | -0.626582 | -1.571315 | 0.00565 | 0.029083 |
| GOBP_POSITIVE_REGULATION_OF_INTERLEUKIN_6_PRODUCTION | 78 | -0.562376 | -1.571237 | 0.001053 | 0.029066 |
| GOBP_FORELIMB_MORPHOGENESIS | 34 | -0.625707 | -1.570997 | 0.006857 | 0.029109 |
| GOBP_REGULATION_OF_MYELOID_LEUKOCYTE_MEDIATED_IMMUNITY | 50 | -0.594796 | -1.570918 | 0.005459 | 0.029093 |
| GOBP_POSITIVE_REGULATION_OF_MACROPHAGE_MIGRATION | 24 | -0.646653 | -1.569392 | 0.013142 | 0.029663 |
| GOBP_NEURON_FATE_SPECIFICATION | 22 | -0.658777 | -1.569374 | 0.011335 | 0.02962 |
| GOBP_METAL_ION_TRANSPORT | 450 | -0.529897 | -1.569371 | 0 | 0.029571 |
| GOBP_REGULATION_OF_T_HELPER_CELL_DIFFERENTIATION | 35 | -0.622977 | -1.568935 | 0.003352 | 0.029702 |
| GOBP_POTASSIUM_ION_IMPORT_ACROSS_PLASMA_MEMBRANE | 45 | -0.598581 | -1.568707 | 0.002193 | 0.029725 |
| GOBP_NEGATIVE_REGULATION_OF_NEURON_DIFFERENTIATION | 58 | -0.579046 | -1.568604 | 0.001062 | 0.029732 |
| GOBP_METANEPHROS_DEVELOPMENT | 76 | -0.565332 | -1.568539 | 0.003135 | 0.029706 |
| GOBP_NEPHRON_TUBULE_FORMATION | 17 | -0.690166 | -1.568323 | 0.018939 | 0.029729 |
| GOBP_NEURON_RECOGNITION | 46 | -0.603608 | -1.566896 | 0.006586 | 0.030298 |
| GOBP_CARDIAC_ATRIUM_DEVELOPMENT | 32 | -0.621493 | -1.566778 | 0.008055 | 0.030296 |
| GOBP_REGULATION_OF_SIGNALING_RECEPTOR_ACTIVITY | 168 | -0.538067 | -1.566585 | 0 | 0.030334 |
| GOBP_CELLULAR_RESPONSE_TO_AMYLOID_BETA | 36 | -0.616718 | -1.565737 | 0.005656 | 0.030633 |
| GOBP_DENDRITE_SELF_AVOIDANCE | 17 | -0.694617 | -1.565421 | 0.011321 | 0.0307 |
| GOBP_REGULATION_OF_EXOCYTOSIS | 189 | -0.536662 | -1.565322 | 0 | 0.030692 |
| GOBP_NEGATIVE_REGULATION_OF_CALCIUM_ION_TRANSMEMBRANE_TRANS | 30 | -0.632985 | -1.564999 | 0.005875 | 0.030759 |
| GOBP_B_CELL_HOMEOSTASIS | 28 | -0.649212 | -1.564843 | 0.005794 | 0.030778 |
| GOBP_NEGATIVE_REGULATION_OF_B_CELL_PROLIFERATION | 17 | -0.689396 | -1.564775 | 0.010178 | 0.030756 |
| GOBP_OSTEOCLAST_DIFFERENTIATION | 85 | -0.55238 | -1.564618 | 0 | 0.030765 |
| GOBP_INTERFERON_ALPHA_PRODUCTION | 28 | -0.626323 | -1.563427 | 0.009292 | 0.031236 |
| GOBP_NEGATIVE_REGULATION_OF_T_CELL_APOPTOTIC_PROCESS | 19 | -0.678176 | -1.563147 | 0.012121 | 0.031297 |
| GOBP_EAR_DEVELOPMENT | 184 | -0.533056 | -1.562844 | 0 | 0.031371 |
| GOBP_PROSTANOID_BIOSYNTHETIC_PROCESS | 26 | -0.647376 | -1.562586 | 0.010502 | 0.031455 |
| GOBP_NEGATIVE_REGULATION_OF_CATION_CHANNEL_ACTIVITY | 39 | -0.606407 | -1.561763 | 0.00453 | 0.031771 |
| GOBP_POSITIVE_REGULATION_OF_TRANSPORTER_ACTIVITY | 108 | -0.54715 | -1.561422 | 0 | 0.031851 |
| GOBP_POSITIVE_REGULATION_OF_CALCIUM_ION_TRANSPORT_INTO_CYTOSC | 48 | -0.598894 | -1.561054 | 0.001115 | 0.031947 |
| GOBP_ASTROCYTE_DEVELOPMENT | 39 | -0.601612 | -1.560893 | 0.005682 | 0.031965 |
| GOBP_BIOLOGICAL_PROCESS_INVOLVED_IN_INTRASPECIES_INTERACTION_B | 48 | -0.591248 | -1.560881 | 0.003268 | 0.031913 |
| GOBP_REGULATION_OF_METAL_ION_TRANSPORT | 224 | -0.535101 | -1.560824 | 0 | 0.03189 |
| GOBP_SARCOPLASMIC_RETICULUM_CALCIUM_ION_TRANSPORT | 34 | -0.622341 | -1.560578 | 0.015801 | 0.03195 |
| GOBP_NEGATIVE_REGULATION_OF_STRIATED_MUSCLE_CELL_DIFFERENTIAT | 31 | -0.621414 | -1.559281 | 0.009357 | 0.032508 |
| GOBP_REGULATION_OF_NEURON_DIFFERENTIATION | 173 | -0.538431 | -1.559254 | 0 | 0.032469 |
| GOBP_REGIONALIZATION | 287 | -0.530666 | -1.558927 | 0 | 0.032584 |
| GOBP_SKELETAL_SYSTEM_DEVELOPMENT | 447 | -0.528007 | -1.55742 | 0 | 0.033211 |
| GOBP_POSITIVE_REGULATION_OF_RESPONSE_TO_BIOTIC_STIMULUS | 228 | -0.529856 | -1.55725 | 0 | 0.03323 |
| GOBP_INTERLEUKIN_8_PRODUCTION | 78 | -0.561891 | -1.556639 | 0 | 0.033463 |
| GOBP_ANION_HOMEOSTASIS | 29 | -0.626668 | -1.556548 | 0.009434 | 0.033459 |
| GOBP_CALCIUM_ION_REGULATED_EXOCYTOSIS_OF_NEUROTRANSMITTER | 17 | -0.688546 | -1.555956 | 0.021465 | 0.033685 |
| GOBP_FAT_SOLUBLE_VITAMIN_METABOLIC_PROCESS | 39 | -0.604033 | -1.55579 | 0.00565 | 0.033702 |

| | | | | | |
|---|---|---|---|---|---|
| GOBP_LEUKOCYTE_ADHESION_TO_VASCULAR_ENDOTHELIAL_CELL | 44 | -0.588588 | -1.555746 | 0.006682 | 0.033666 |
| GOBP_POSITIVE_REGULATION_OF_MESENCHYMAL_CELL_PROLIFERATION | 21 | -0.66352 | -1.555725 | 0.009604 | 0.033626 |
| GOBP_POSITIVE_REGULATION_OF_MONOCYTE_CHEMOTAXIS | 18 | -0.682893 | -1.555398 | 0.010152 | 0.033728 |
| GOBP_TELENCEPHALON_DEVELOPMENT | 215 | -0.534496 | -1.555207 | 0 | 0.033764 |
| GOBP_REGULATION_OF_GLIOGENESIS | 83 | -0.557836 | -1.555017 | 0 | 0.033791 |
| GOBP_ENDODERM_DEVELOPMENT | 72 | -0.56239 | -1.554956 | 0.001056 | 0.033759 |
| GOBP_NEURON_PROJECTION_GUIDANCE | 268 | -0.529813 | -1.554664 | 0 | 0.033833 |
| GOBP_MESONEPHRIC_TUBULE_MORPHOGENESIS | 55 | -0.577173 | -1.554047 | 0.003279 | 0.034039 |
| GOBP_DEOXYRIBONUCLEOSIDE_MONOPHOSPHATE_METABOLIC_PROCESS | 16 | -0.693609 | -1.55386 | 0.012563 | 0.034074 |
| GOBP_APOPTOTIC_CELL_CLEARANCE | 49 | -0.587164 | -1.55377 | 0.003293 | 0.034064 |
| GOBP_EPITHELIAL_CELL_DIFFERENTIATION_INVOLVED_IN_KIDNEY_DEVELOP | 45 | -0.605133 | -1.553582 | 0.007804 | 0.034129 |
| GOBP_POSITIVE_REGULATION_OF_DENDRITE_DEVELOPMENT | 15 | -0.696417 | -1.55285 | 0.021656 | 0.034403 |
| GOBP_NEGATIVE_REGULATION_OF_DEFENSE_RESPONSE | 190 | -0.529606 | -1.552711 | 0 | 0.034402 |
| GOBP_MESENCHYMAL_TO_EPITHELIAL_TRANSITION | 18 | -0.680904 | -1.552512 | 0.009768 | 0.034428 |
| GOBP_POSITIVE_REGULATION_OF_GLYCOLYTIC_PROCESS | 23 | -0.646331 | -1.552352 | 0.017073 | 0.034436 |
| GOBP_REGULATION_OF_BLOOD_CIRCULATION | 261 | -0.524472 | -1.551154 | 0 | 0.034934 |
| GOBP_GLANDULAR_EPITHELIAL_CELL_DEVELOPMENT | 25 | -0.642648 | -1.550915 | 0.011947 | 0.035002 |
| GOBP_NEGATIVE_REGULATION_OF_INNATE_IMMUNE_RESPONSE | 58 | -0.576912 | -1.550672 | 0.005359 | 0.035041 |
| GOBP_AMINO_ACID_TRANSMEMBRANE_TRANSPORT | 87 | -0.562839 | -1.549891 | 0.001038 | 0.03532 |
| GOBP_REGULATION_OF_CARDIAC_CONDUCTION | 55 | -0.581 | -1.549792 | 0.00432 | 0.03532 |
| GOBP_POSITIVE_REGULATION_OF_TRANSMEMBRANE_TRANSPORT | 191 | -0.531334 | -1.549739 | 0 | 0.035286 |
| GOBP_REGULATION_OF_LYASE_ACTIVITY | 44 | -0.593856 | -1.549603 | 0.006536 | 0.035298 |
| GOBP_CGMP_METABOLIC_PROCESS | 16 | -0.696179 | -1.549236 | 0.014925 | 0.035384 |
| GOBP_L_AMINO_ACID_TRANSPORT | 61 | -0.564162 | -1.548961 | 0.004264 | 0.035447 |
| GOBP_POSITIVE_REGULATION_OF_INTERLEUKIN_2_PRODUCTION | 30 | -0.621758 | -1.548186 | 0.011494 | 0.035766 |
| GOBP_NEURON_PROJECTION_ORGANIZATION | 89 | -0.552057 | -1.547651 | 0.001037 | 0.035982 |
| GOBP_REGULATION_OF_PHOSPHATIDYLINOSITOL_3_KINASE_SIGNALING | 117 | -0.543202 | -1.547566 | 0 | 0.035987 |
| GOBP_MACROPHAGE_DIFFERENTIATION | 41 | -0.601384 | -1.547528 | 0.00905 | 0.035949 |
| GOBP_BONE_MINERALIZATION | 107 | -0.546055 | -1.547084 | 0 | 0.036107 |
| GOBP_NEPHRON_EPITHELIUM_DEVELOPMENT | 99 | -0.550862 | -1.547046 | 0 | 0.036075 |
| GOBP_NEGATIVE_REGULATION_OF_CYTOKINE_PRODUCTION | 261 | -0.529224 | -1.546556 | 0 | 0.03622 |
| GOBP_PHOSPHOLIPASE_C_ACTIVATING_G_PROTEIN_COUPLED_RECEPTOR_ | 85 | -0.550196 | -1.546478 | 0.003106 | 0.036203 |
| GOBP_POSITIVE_REGULATION_OF_INTERLEUKIN_10_PRODUCTION | 35 | -0.606317 | -1.546324 | 0.014823 | 0.036209 |
| GOBP_NEURON_CELL_CELL_ADHESION | 17 | -0.682308 | -1.546173 | 0.020936 | 0.036236 |
| GOBP_REGULATION_OF_HEART_CONTRACTION | 222 | -0.530102 | -1.546089 | 0 | 0.036228 |
| GOBP_MOTOR_NEURON_AXON_GUIDANCE | 29 | -0.624809 | -1.545961 | 0.013857 | 0.036248 |
| GOBP_RENAL_TUBULE_DEVELOPMENT | 85 | -0.550965 | -1.545939 | 0 | 0.036205 |
| GOBP_POSITIVE_REGULATION_OF_EPITHELIAL_CELL_PROLIFERATION | 173 | -0.530557 | -1.545931 | 0 | 0.03615 |
| GOBP_REGULATION_OF_MACROPHAGE_MIGRATION | 38 | -0.597701 | -1.545911 | 0.006787 | 0.036101 |
| GOBP_INFLAMMATORY_CELL_APOPTOTIC_PROCESS | 19 | -0.673504 | -1.545526 | 0.016332 | 0.03623 |
| GOBP_REGULATION_OF_INNATE_IMMUNE_RESPONSE | 288 | -0.525205 | -1.545456 | 0 | 0.036221 |
| GOBP_REGULATION_OF_LIPID_KINASE_ACTIVITY | 67 | -0.556669 | -1.544872 | 0.006349 | 0.03646 |
| GOBP_ANGIOGENESIS_INVOLVED_IN_WOUND_HEALING | 24 | -0.633148 | -1.544566 | 0.018846 | 0.036556 |
| GOBP_NEGATIVE_REGULATION_OF_CALCIUM_ION_TRANSMEMBRANE_TRANS | 34 | -0.611719 | -1.544515 | 0.011481 | 0.036523 |
| GOBP_POSITIVE_REGULATION_OF_CALCIUM_ION_DEPENDENT_EXOCYTOSIS | 18 | -0.665007 | -1.544331 | 0.020487 | 0.036552 |
| GOBP_CELL_FATE_COMMITMENT_INVOLVED_IN_FORMATION_OF_PRIMARY_G | 26 | -0.638962 | -1.544306 | 0.009185 | 0.036505 |
| GOBP_NEGATIVE_REGULATION_OF_RESPONSE_TO_BIOTIC_STIMULUS | 91 | -0.550182 | -1.544169 | 0 | 0.036514 |
| GOBP_EMBRYONIC_FORELIMB_MORPHOGENESIS | 28 | -0.631769 | -1.543847 | 0.008314 | 0.036629 |
| GOBP_INHIBITORY_SYNAPSE_ASSEMBLY | 16 | -0.708306 | -1.543376 | 0.017857 | 0.036824 |
| GOBP_G_PROTEIN_COUPLED_RECEPTOR_SIGNALING_PATHWAY_COUPLED_ | 43 | -0.597209 | -1.543272 | 0.00452 | 0.036826 |
| GOBP_HIPPOCAMPUS_DEVELOPMENT | 65 | -0.565697 | -1.541883 | 0.005359 | 0.037494 |
| GOBP_NEGATIVE_REGULATION_OF_MYELOID_LEUKOCYTE_DIFFERENTIATION | 44 | -0.587127 | -1.541846 | 0.006615 | 0.037458 |
| GOBP_REGULATION_OF_MYOBLAST_FUSION | 17 | -0.670389 | -1.541642 | 0.009889 | 0.037482 |
| GOBP_NEGATIVE_REGULATION_OF_ALPHA_BETA_T_CELL_ACTIVATION | 37 | -0.604379 | -1.541246 | 0.006726 | 0.037643 |
| GOBP_POSITIVE_REGULATION_OF_B_CELL_MEDIATED_IMMUNITY | 36 | -0.605966 | -1.541235 | 0.005695 | 0.037593 |
| GOBP_EMBRYONIC_DIGIT_MORPHOGENESIS | 49 | -0.587821 | -1.540948 | 0.007675 | 0.037699 |
| GOBP_INORGANIC_ANION_TRANSMEMBRANE_TRANSPORT | 108 | -0.540197 | -1.540707 | 0.001014 | 0.037763 |
| GOBP_PROTEIN_KINASE_C_SIGNALING | 28 | -0.621743 | -1.540566 | 0.015439 | 0.037774 |
| GOBP_EAR_MORPHOGENESIS | 97 | -0.548572 | -1.539728 | 0.001031 | 0.038192 |
| GOBP_NEGATIVE_REGULATION_OF_LYMPHOCYTE_APOPTOTIC_PROCESS | 29 | -0.627549 | -1.539094 | 0.010251 | 0.038437 |
| GOBP_FATTY_ACID_TRANSPORT | 134 | -0.532135 | -1.538318 | 0 | 0.038751 |
| GOBP_REGULATION_OF_B_CELL_RECEPTOR_SIGNALING_PATHWAY | 25 | -0.646181 | -1.53728 | 0.012077 | 0.039232 |
| GOBP_CELLULAR_RESPONSE_TO_BIOTIC_STIMULUS | 195 | -0.526289 | -1.536712 | 0 | 0.039497 |
| GOBP_BODY_MORPHOGENESIS | 46 | -0.584801 | -1.536554 | 0.005388 | 0.039524 |
| GOBP_GLIOGENESIS | 265 | -0.52118 | -1.535972 | 0 | 0.039773 |
| GOBP_MULTICELLULAR_ORGANISMAL_RESPONSE_TO_STRESS | 59 | -0.572502 | -1.535935 | 0.00107 | 0.039734 |
| GOBP_RELAXATION_OF_MUSCLE | 28 | -0.633234 | -1.535587 | 0.015029 | 0.039872 |
| GOBP_REGULATION_OF_OSTEOCLAST_DIFFERENTIATION | 56 | -0.567268 | -1.535483 | 0.003202 | 0.039867 |
| GOBP_KIDNEY_MORPHOGENESIS | 83 | -0.558437 | -1.53537 | 0 | 0.039859 |
| GOBP_REGULATION_OF_HEART_RATE | 88 | -0.55372 | -1.535296 | 0.002068 | 0.03984 |
| GOBP_ENDOTHELIAL_CELL_CHEMOTAXIS | 26 | -0.626343 | -1.534852 | 0.014151 | 0.040002 |
| GOBP_NEGATIVE_REGULATION_OF_NATURAL_KILLER_CELL_MEDIATED_IMMU | 18 | -0.669218 | -1.534835 | 0.010283 | 0.03995 |
| GOBP_POSITIVE_REGULATION_OF_INTERLEUKIN_8_PRODUCTION | 52 | -0.575191 | -1.534262 | 0.004362 | 0.040203 |
| GOBP_PITUITARY_GLAND_DEVELOPMENT | 34 | -0.608523 | -1.533902 | 0.00905 | 0.040335 |
| GOBP_GANGLIOSIDE_METABOLIC_PROCESS | 24 | -0.651517 | -1.533786 | 0.011862 | 0.040353 |
| GOBP_MYELOID_DENDRITIC_CELL_DIFFERENTIATION | 19 | -0.67731 | -1.533642 | 0.012285 | 0.040361 |
| GOBP_ZINC_ION_HOMEOSTASIS | 37 | -0.599058 | -1.533457 | 0.011507 | 0.040398 |
| GOBP_POSITIVE_REGULATION_OF_PHOSPHATASE_ACTIVITY | 32 | -0.613777 | -1.533425 | 0.006912 | 0.040348 |
| GOBP_REGULATION_OF_RECEPTOR_BINDING | 22 | -0.645054 | -1.533369 | 0.015588 | 0.040328 |
| GOBP_REGULATION_OF_BMP_SIGNALING_PATHWAY | 82 | -0.554319 | -1.53331 | 0.002066 | 0.040299 |
| GOBP_PRESYNAPSE_ORGANIZATION | 47 | -0.585038 | -1.532578 | 0.005495 | 0.040653 |
| GOBP_HEAD_MORPHOGENESIS | 33 | -0.605536 | -1.531901 | 0.017401 | 0.040941 |
| GOBP_CHEMICAL_SYNAPTIC_TRANSMISSION_POSTSYNAPTIC | 96 | -0.541891 | -1.531881 | 0 | 0.040895 |
| GOBP_MORPHOGENESIS_OF_AN_EPITHELIAL_SHEET | 57 | -0.567951 | -1.531094 | 0.008639 | 0.041243 |
| GOBP_ACUTE_INFLAMMATORY_RESPONSE | 87 | -0.5491 | -1.53101 | 0.002079 | 0.04123 |
| GOBP_PATTERN_SPECIFICATION_PROCESS | 382 | -0.51826 | -1.530365 | 0 | 0.041569 |
| GOBP_REGULATION_OF_CARDIAC_MUSCLE_CONTRACTION_BY_REGULATION | 20 | -0.652183 | -1.530249 | 0.014724 | 0.041563 |
| GOBP_CATECHOLAMINE_SECRETION | 47 | -0.582649 | -1.53018 | 0.005501 | 0.041531 |
| GOBP_REGULATION_OF_PHOSPHATIDYLINOSITOL_3_KINASE_ACTIVITY | 56 | -0.575694 | -1.528885 | 0.003219 | 0.042201 |
| GOBP_NEURON_MATURATION | 45 | -0.585332 | -1.528793 | 0.002205 | 0.042193 |
| GOBP_POSITIVE_REGULATION_OF_INTERLEUKIN_4_PRODUCTION | 24 | -0.636259 | -1.528742 | 0.021454 | 0.042149 |
| GOBP_CYCLIC_NUCLEOTIDE_BIOSYNTHETIC_PROCESS | 21 | -0.648261 | -1.528601 | 0.020732 | 0.042156 |
| GOBP_PHOSPHATIDYLETHANOLAMINE_METABOLIC_PROCESS | 30 | -0.614981 | -1.528326 | 0.012791 | 0.042249 |
| GOBP_CELLULAR_TRANSITION_METAL_ION_HOMEOSTASIS | 108 | -0.532637 | -1.528303 | 0.001024 | 0.042197 |
| GOBP_NEGATIVE_REGULATION_OF_BLOOD_PRESSURE | 34 | -0.597381 | -1.528141 | 0.016037 | 0.042227 |
| GOBP_BICARBONATE_TRANSPORT | 40 | -0.589886 | -1.528049 | 0.011211 | 0.042209 |
| GOBP_ICOSANOID_BIOSYNTHETIC_PROCESS | 41 | -0.590378 | -1.527551 | 0.011364 | 0.042451 |
| GOBP_ANTIMICROBIAL_HUMORAL_IMMUNE_RESPONSE_MEDIATED_BY_ANTIM | 47 | -0.572413 | -1.527549 | 0.00765 | 0.042392 |
| GOBP_POSITIVE_REGULATION_OF_MYELOID_CELL_DIFFERENTIATION | 89 | -0.544157 | -1.527538 | 0.001048 | 0.042343 |
| GOBP_PROTEIN_KINASE_B_SIGNALING | 231 | -0.525293 | -1.527479 | 0 | 0.042314 |
| GOBP_NEGATIVE_REGULATION_OF_LYMPHOCYTE_DIFFERENTIATION | 47 | -0.582318 | -1.527272 | 0.010056 | 0.042365 |
| GOBP_NEGATIVE_REGULATION_OF_EXTRINSIC_APOPTOTIC_SIGNALING_PATH | 37 | -0.596987 | -1.52584 | 0.011377 | 0.043059 |
| GOBP_INTERLEUKIN_17_PRODUCTION | 30 | -0.623984 | -1.52568 | 0.012896 | 0.043076 |
| GOBP_REGULATION_OF_RECEPTOR_MEDIATED_ENDOCYTOSIS | 95 | -0.542285 | -1.525662 | 0.002077 | 0.043025 |
| GOBP_NEGATIVE_REGULATION_OF_EPITHELIAL_CELL_DIFFERENTIATION | 37 | -0.598768 | -1.525588 | 0.018994 | 0.043008 |
| GOBP_AXON_DEVELOPMENT | 488 | -0.514304 | -1.525513 | 0 | 0.042979 |
| GOBP_SENSORY_PERCEPTION_OF_TEMPERATURE_STIMULUS | 22 | -0.634014 | -1.525455 | 0.024184 | 0.04295 |
| GOBP_REGULATION_OF_SYNAPTIC_VESICLE_EXOCYTOSIS | 55 | -0.560617 | -1.524928 | 0.009719 | 0.043162 |
| GOBP_LIPID_IMPORT_INTO_CELL | 34 | -0.61036 | -1.524674 | 0.005669 | 0.043207 |
| GOBP_GLIAL_CELL_DIFFERENTIATION | 196 | -0.520491 | -1.52423 | 0 | 0.043391 |
| GOBP_REGULATION_OF_MACROPHAGE_DIFFERENTIATION | 18 | -0.670204 | -1.524209 | 0.018325 | 0.043341 |
| GOBP_POSITIVE_REGULATION_OF_MONOOXYGENASE_ACTIVITY | 30 | -0.602431 | -1.523787 | 0.018605 | 0.043517 |
| GOBP_NLRP3_INFLAMMASOME_COMPLEX_ASSEMBLY | 15 | -0.681896 | -1.523369 | 0.019405 | 0.043676 |
| GOBP_INOSITOL_LIPID_MEDIATED_SIGNALING | 182 | -0.525185 | -1.523317 | 0 | 0.043639 |
| GOBP_REGULATION_OF_TRANSCRIPTION_REGULATORY_REGION_DNA_BINDI | 50 | -0.567891 | -1.523238 | 0.007479 | 0.043626 |
| GOBP_POSITIVE_REGULATION_OF_MORPHOGENESIS_OF_AN_EPITHELIUM | 32 | -0.596634 | -1.522967 | 0.021226 | 0.043697 |
| GOBP_REGULATION_OF_MICROGLIAL_CELL_ACTIVATION | 17 | -0.667789 | -1.522712 | 0.024675 | 0.04378 |
| GOBP_REGULATION_OF_CELL_JUNCTION_ASSEMBLY | 186 | -0.525584 | -1.52223 | 0 | 0.044 |
| GOBP_MAINTENANCE_OF_LOCATION_IN_CELL | 196 | -0.519959 | -1.522135 | 0 | 0.044002 |
| GOBP_TISSUE_REMODELING | 145 | -0.531156 | -1.521072 | 0.001012 | 0.044531 |
| GOBP_ANTIGEN_PROCESSING_AND_PRESENTATION | 225 | -0.519197 | -1.520893 | 0 | 0.044586 |
| GOBP_BONE_CELL_DEVELOPMENT | 31 | -0.612261 | -1.520273 | 0.019767 | 0.044869 |
| GOBP_LUNG_ALVEOLUS_DEVELOPMENT | 35 | -0.584529 | -1.519927 | 0.017143 | 0.044975 |
| GOBP_REGULATION_OF_RECEPTOR_SIGNALING_PATHWAY_VIA_STAT | 74 | -0.55181 | -1.519514 | 0.004215 | 0.045137 |
| GOBP_REGULATION_OF_SYNCYTIUM_FORMATION_BY_PLASMA_MEMBRANE_ | 23 | -0.640987 | -1.519469 | 0.021302 | 0.045098 |
| GOBP_RESPONSE_TO_TUMOR_NECROSIS_FACTOR | 290 | -0.513967 | -1.519223 | 0 | 0.045172 |
| GOBP_RECEPTOR_METABOLIC_PROCESS | 163 | -0.523871 | -1.518809 | 0.001006 | 0.045366 |
| GOBP_MESONEPHROS_DEVELOPMENT | 85 | -0.540402 | -1.518674 | 0.005155 | 0.045375 |
| GOBP_CYCLIC_NUCLEOTIDE_METABOLIC_PROCESS | 38 | -0.587637 | -1.517792 | 0.011737 | 0.045834 |
| GOBP_MONOCARBOXYLIC_ACID_TRANSPORT | 155 | -0.514797 | -1.517575 | 0 | 0.045898 |
| GOBP_REGULATION_OF_HEART_RATE_BY_CARDIAC_CONDUCTION | 39 | -0.589273 | -1.517416 | 0.012346 | 0.04594 |
| GOBP_STEM_CELL_DIVISION | 24 | -0.635787 | -1.517389 | 0.021687 | 0.045892 |
| GOBP_ASTROCYTE_ACTIVATION | 22 | -0.64264 | -1.517104 | 0.019277 | 0.045998 |
| GOBP_HEART_PROCESS | 264 | -0.515786 | -1.516846 | 0 | 0.046076 |
| GOBP_DENDRITE_MORPHOGENESIS | 137 | -0.530384 | -1.516444 | 0 | 0.04624 |
| GOBP_VASOCONSTRICTION | 72 | -0.54554 | -1.516276 | 0.002079 | 0.046263 |
| GOBP_DENDRITE_DEVELOPMENT | 229 | -0.51429 | -1.516232 | 0 | 0.046229 |
| GOBP_SPHINGOLIPID_MEDIATED_SIGNALING_PATHWAY | 15 | -0.684435 | -1.516003 | 0.029487 | 0.046288 |
| GOBP_MORPHOGENESIS_OF_AN_EPITHELIAL_FOLD | 20 | -0.647644 | -1.515901 | 0.019324 | 0.046272 |
| GOBP_TERPENOID_METABOLIC_PROCESS | 101 | -0.535913 | -1.51568 | 0 | 0.046343 |
| GOBP_REGULATION_OF_DENDRITE_DEVELOPMENT | 94 | -0.539326 | -1.515329 | 0.001032 | 0.046496 |
| GOBP_REGULATION_OF_NERVOUS_SYSTEM_DEVELOPMENT | 401 | -0.510165 | -1.515187 | 0 | 0.046538 |
| GOBP_PANCREAS_DEVELOPMENT | 64 | -0.557535 | -1.51428 | 0.003226 | 0.046979 |
| GOBP_NEGATIVE_REGULATION_OF_CD4_POSITIVE_ALPHA_BETA_T_CELL_AC | 28 | -0.61596 | -1.514104 | 0.015222 | 0.047016 |
| GOBP_EMBRYONIC_ORGAN_MORPHOGENESIS | 252 | -0.516146 | -1.514079 | 0.001004 | 0.046964 |
| GOBP_ASSOCIATIVE_LEARNING | 63 | -0.556155 | -1.513884 | 0.006417 | 0.047041 |
| GOBP_CD4_POSITIVE_ALPHA_BETA_T_CELL_CYTOKINE_PRODUCTION | 16 | -0.681507 | -1.513834 | 0.02551 | 0.047005 |
| GOBP_TOLL_LIKE_RECEPTOR_9_SIGNALING_PATHWAY | 26 | -0.616881 | -1.513766 | 0.024038 | 0.046977 |
| GOBP_RESPONSE_TO_INTERLEUKIN_4 | 33 | -0.603418 | -1.513639 | 0.015099 | 0.046986 |
| GOBP_REGULATION_OF_EPITHELIAL_CELL_DIFFERENTIATION_INVOLVED_IN_ | 18 | -0.669214 | -1.513483 | 0.008706 | 0.047012 |
| GOBP_POSITIVE_REGULATION_OF_GRANULOCYTE_CHEMOTAXIS | 27 | -0.619179 | -1.512686 | 0.016222 | 0.04745 |

| | | | | | |
|---|---|---|---|---|---|
| GOBP_CELL_COMMUNICATION_BY_ELECTRICAL_COUPLING_INVOLVED_IN_CA | 24 | -0.626164 | -1.512658 | 0.01816 | 0.047407 |
| GOBP_POSITIVE_REGULATION_OF_MACROPHAGE_CHEMOTAXIS | 17 | -0.685286 | -1.512581 | 0.012739 | 0.04738 |
| GOBP_METAL_ION_EXPORT | 45 | -0.581582 | -1.512413 | 0.010989 | 0.047416 |
| GOBP_CHLORIDE_TRANSPORT | 99 | -0.538235 | -1.512314 | 0.001025 | 0.047419 |
| GOBP_POSITIVE_REGULATION_OF_RESPONSE_TO_CYTOKINE_STIMULUS | 55 | -0.565729 | -1.512165 | 0.0044 | 0.047462 |
| GOBP_POSITIVE_REGULATION_OF_MACROPHAGE_ACTIVATION | 24 | -0.621143 | -1.511339 | 0.02497 | 0.047836 |
| GOBP_REGULATION_OF_VENTRICULAR_CARDIAC_MUSCLE_CELL_MEMBRANI | 19 | -0.654299 | -1.511232 | 0.025 | 0.047834 |
| GOBP_REGULATION_OF_RESPONSE_TO_BIOTIC_STIMULUS | 379 | -0.509321 | -1.510968 | 0 | 0.047954 |
| GOBP_CALCIUM_ION_IMPORT | 74 | -0.54964 | -1.510896 | 0.001049 | 0.047948 |
| GOBP_POSITIVE_REGULATION_OF_CALCIUM_ION_TRANSPORT | 108 | -0.533969 | -1.509371 | 0.001026 | 0.048737 |
| GOBP_POSITIVE_REGULATION_OF_NEURON_PROJECTION_DEVELOPMENT | 146 | -0.524147 | -1.509047 | 0 | 0.048876 |
| GOBP_NEGATIVE_REGULATION_OF_TRANSPORTER_ACTIVITY | 70 | -0.549129 | -1.50824 | 0.007407 | 0.049297 |
| GOBP_RESPONSE_TO_NITRIC_OXIDE | 19 | -0.667562 | -1.50806 | 0.022613 | 0.049354 |
| GOBP_CARDIAC_MUSCLE_CELL_CONTRACTION | 68 | -0.546668 | -1.508047 | 0.005252 | 0.049299 |
| GOBP_AMINO_ACID_TRANSPORT | 141 | -0.522494 | -1.50781 | 0 | 0.049381 |
| GOBP_BRANCHING_MORPHOGENESIS_OF_AN_EPITHELIAL_TUBE | 131 | -0.525296 | -1.507038 | 0 | 0.04978 |
| GOBP_REGULATION_OF_NEURONAL_SYNAPTIC_PLASTICITY | 44 | -0.573261 | -1.504702 | 0.013408 | 0.051122 |
| GOBP_NEGATIVE_REGULATION_OF_TISSUE_REMODELING | 18 | -0.656831 | -1.504621 | 0.025126 | 0.051102 |
| GOBP_NEGATIVE_REGULATION_OF_RESPONSE_TO_EXTERNAL_STIMULUS | 329 | -0.513479 | -1.504384 | 0 | 0.05121 |
| GOBP_ACUTE_INFLAMMATORY_RESPONSE_TO_ANTIGENIC_STIMULUS | 19 | -0.653684 | -1.504378 | 0.027638 | 0.05115 |
| GOBP_RESPONSE_TO_NERVE_GROWTH_FACTOR | 49 | -0.573132 | -1.50344 | 0.010787 | 0.051654 |
| GOBP_GLIAL_CELL_MIGRATION | 49 | -0.573504 | -1.502435 | 0.013001 | 0.052233 |
| GOBP_REGULATION_OF_INTEGRIN_ACTIVATION | 17 | -0.662671 | -1.502181 | 0.029449 | 0.052321 |
| GOBP_REGULATION_OF_MACROPHAGE_CHEMOTAXIS | 27 | -0.619842 | -1.502129 | 0.024155 | 0.052284 |
| GOBP_LOCOMOTORY_BEHAVIOR | 143 | -0.521277 | -1.502013 | 0 | 0.052284 |
| GOBP_ACTIVATION_OF_ADENYLATE_CYCLASE_ACTIVITY | 32 | -0.591301 | -1.501566 | 0.018433 | 0.052484 |
| GOBP_METANEPHRIC_RENAL_VESICLE_MORPHOGENESIS | 15 | -0.674624 | -1.501016 | 0.027487 | 0.052769 |
| GOBP_VASCULAR_TRANSPORT | 87 | -0.536437 | -1.500939 | 0.005219 | 0.052746 |
| GOBP_ORGANIC_ACID_TRANSMEMBRANE_TRANSPORT | 144 | -0.521644 | -1.500805 | 0 | 0.052764 |
| GOBP_T_CELL_LINEAGE_COMMITMENT | 25 | -0.614595 | -1.500066 | 0.03436 | 0.053177 |
| GOBP_POSITIVE_REGULATION_OF_MAPK_CASCADE | 480 | -0.505616 | -1.499874 | 0 | 0.053235 |
| GOBP_INNER_EAR_MORPHOGENESIS | 78 | -0.535415 | -1.499591 | 0.004175 | 0.053343 |
| GOBP_NEGATIVE_REGULATION_OF_CATION_TRANSMEMBRANE_TRANSPORT | 77 | -0.539243 | -1.499205 | 0.005258 | 0.053547 |
| GOBP_FOREBRAIN_CELL_MIGRATION | 51 | -0.56497 | -1.498958 | 0.015317 | 0.053645 |
| GOBP_VASCULAR_PROCESS_IN_CIRCULATORY_SYSTEM | 232 | -0.510304 | -1.498699 | 0 | 0.053754 |
| GOBP_PROTEIN_LOCALIZATION_TO_CELL_JUNCTION | 108 | -0.529356 | -1.498653 | 0.002041 | 0.05371 |
| GOBP_MODULATION_OF_EXCITATORY_POSTSYNAPTIC_POTENTIAL | 42 | -0.580868 | -1.498526 | 0.013058 | 0.053739 |
| GOBP_ENSHEATHMENT_OF_NEURONS | 122 | -0.524309 | -1.498462 | 0.001026 | 0.053706 |
| GOBP_GLIAL_CELL_DEVELOPMENT | 106 | -0.53005 | -1.497511 | 0.002073 | 0.054283 |
| GOBP_DETECTION_OF_OTHER_ORGANISM | 16 | -0.672785 | -1.496941 | 0.026185 | 0.05459 |
| GOBP_CARDIAC_MUSCLE_CELL_MEMBRANE_REPOLARIZATION | 33 | -0.595553 | -1.496436 | 0.020713 | 0.054813 |
| GOBP_AMINOGLYCAN_CATABOLIC_PROCESS | 61 | -0.557257 | -1.496329 | 0.007503 | 0.05481 |
| GOBP_AMINO_ACID_IMPORT | 45 | -0.577486 | -1.496124 | 0.012155 | 0.054849 |
| GOBP_NEGATIVE_REGULATION_OF_MACROPHAGE_ACTIVATION | 17 | -0.66096 | -1.49512 | 0.033766 | 0.05544 |
| GOBP_REGULATION_OF_DENDRITE_MORPHOGENESIS | 63 | -0.555591 | -1.494959 | 0.008574 | 0.055475 |
| GOBP_NEGATIVE_REGULATION_OF_NERVOUS_SYSTEM_DEVELOPMENT | 128 | -0.522868 | -1.494416 | 0 | 0.0558 |
| GOBP_POSITIVE_REGULATION_OF_GLIOGENESIS | 59 | -0.553277 | -1.494388 | 0.013786 | 0.055751 |
| GOBP_CARDIAC_MUSCLE_CELL_ACTION_POTENTIAL_INVOLVED_IN_CONTRA | 50 | -0.560943 | -1.494303 | 0.012987 | 0.055742 |
| GOBP_CHONDROITIN_SULFATE_PROTEOGLYCAN_BIOSYNTHETIC_PROCESS | 30 | -0.593346 | -1.494053 | 0.02331 | 0.055866 |
| GOBP_CARTILAGE_DEVELOPMENT | 170 | -0.513499 | -1.493818 | 0 | 0.055964 |
| GOBP_POSITIVE_REGULATION_OF_NITRIC_OXIDE_METABOLIC_PROCESS | 33 | -0.591357 | -1.493681 | 0.02069 | 0.055973 |
| GOBP_BONE_MORPHOGENESIS | 83 | -0.53623 | -1.493239 | 0.004158 | 0.056192 |
| GOBP_MYELOID_LEUKOCYTE_DIFFERENTIATION | 184 | -0.50767 | -1.4929 | 0 | 0.056314 |
| GOBP_COLLAGEN_METABOLIC_PROCESS | 93 | -0.529239 | -1.492317 | 0.001029 | 0.056631 |
| GOBP_RESPONSE_TO_VIRUS | 314 | -0.505089 | -1.492302 | 0 | 0.056569 |
| GOBP_CELL_DIFFERENTIATION_IN_HINDBRAIN | 18 | -0.652229 | -1.491838 | 0.031056 | 0.056791 |
| GOBP_RETINAL_GANGLION_CELL_AXON_GUIDANCE | 18 | -0.637735 | -1.491756 | 0.027363 | 0.05678 |
| GOBP_MULTICELLULAR_ORGANISMAL_HOMEOSTASIS | 486 | -0.502521 | -1.491687 | 0 | 0.056758 |
| GOBP_ENDOCARDIAL_CUSHION_DEVELOPMENT | 42 | -0.572266 | -1.490869 | 0.012332 | 0.057265 |
| GOBP_CELL_DIFFERENTIATION_INVOLVED_IN_KIDNEY_DEVELOPMENT | 55 | -0.552911 | -1.489706 | 0.009626 | 0.057972 |
| GOBP_POSTSYNAPSE_ASSEMBLY | 30 | -0.598111 | -1.489702 | 0.025641 | 0.057905 |
| GOBP_GLOMERULAR_EPITHELIUM_DEVELOPMENT | 21 | -0.627643 | -1.488416 | 0.031175 | 0.058659 |
| GOBP_MYELOID_LEUKOCYTE_CYTOKINE_PRODUCTION | 26 | -0.607511 | -1.488338 | 0.022327 | 0.058654 |
| GOBP_RETINOIC_ACID_METABOLIC_PROCESS | 25 | -0.617467 | -1.488154 | 0.032787 | 0.058729 |
| GOBP_P38MAPK_CASCADE | 44 | -0.563185 | -1.487707 | 0.019868 | 0.058957 |
| GOBP_ENDOCRINE_SYSTEM_DEVELOPMENT | 103 | -0.524687 | -1.487626 | 0.001024 | 0.058955 |
| GOBP_INORGANIC_ANION_TRANSPORT | 155 | -0.510933 | -1.485799 | 0 | 0.060183 |
| GOBP_REGULATION_OF_DEFENSE_RESPONSE_TO_VIRUS_BY_HOST | 38 | -0.573535 | -1.48568 | 0.017877 | 0.060196 |
| GOBP_NEGATIVE_REGULATION_OF_FIBROBLAST_GROWTH_FACTOR_RECEPT | 18 | -0.653935 | -1.485654 | 0.032581 | 0.060144 |
| GOBP_ASTROCYTE_DIFFERENTIATION | 67 | -0.544041 | -1.485591 | 0.006403 | 0.060103 |
| GOBP_KIDNEY_EPITHELIUM_DEVELOPMENT | 123 | -0.517016 | -1.485141 | 0 | 0.060373 |
| GOBP_ISOPRENOID_METABOLIC_PROCESS | 120 | -0.514666 | -1.484848 | 0.001019 | 0.060497 |
| GOBP_OLIGODENDROCYTE_DIFFERENTIATION | 87 | -0.529693 | -1.484631 | 0.003074 | 0.060585 |
| GOBP_REGULATION_OF_SENSORY_PERCEPTION | 30 | -0.593746 | -1.484431 | 0.023095 | 0.060665 |
| GOBP_AMINOGLYCAN_METABOLIC_PROCESS | 162 | -0.509635 | -1.483959 | 0.001008 | 0.06091 |
| GOBP_REGULATION_OF_NEURON_MIGRATION | 38 | -0.579079 | -1.4831 | 0.01891 | 0.061443 |
| GOBP_REGULATION_OF_TUBE_SIZE | 116 | -0.513991 | -1.481913 | 0.002024 | 0.062185 |
| GOBP_NEGATIVE_REGULATION_OF_CYSTEINE_TYPE_ENDOPEPTIDASE_ACTIV | 74 | -0.539188 | -1.481025 | 0.008448 | 0.062754 |
| GOBP_OSTEOCLAST_DEVELOPMENT | 15 | -0.666758 | -1.480611 | 0.027708 | 0.062973 |
| GOBP_CENTRAL_NERVOUS_SYSTEM_NEURON_AXONOGENESIS | 34 | -0.587021 | -1.480378 | 0.02381 | 0.063071 |
| GOBP_BRANCHING_INVOLVED_IN_BLOOD_VESSEL_MORPHOGENESIS | 31 | -0.594023 | -1.479976 | 0.03555 | 0.063306 |
| GOBP_POSITIVE_REGULATION_OF_AMINE_TRANSPORT | 31 | -0.591348 | -1.479914 | 0.017689 | 0.063282 |
| GOBP_REGULATION_OF_RECEPTOR_INTERNALIZATION | 50 | -0.549023 | -1.479452 | 0.020833 | 0.063553 |
| GOBP_DEFENSE_RESPONSE_TO_VIRUS | 225 | -0.506692 | -1.479273 | 0 | 0.063607 |
| GOBP_FACE_DEVELOPMENT | 44 | -0.556817 | -1.479052 | 0.01652 | 0.063666 |
| GOBP_POSITIVE_REGULATION_OF_POTASSIUM_ION_TRANSPORT | 43 | -0.563549 | -1.478895 | 0.014428 | 0.06372 |
| GOBP_POSITIVE_REGULATION_OF_OSTEOBLAST_DIFFERENTIATION | 60 | -0.544491 | -1.478895 | 0.007511 | 0.063645 |
| GOBP_RESPONSE_TO_ELECTRICAL_STIMULUS | 40 | -0.572003 | -1.478566 | 0.022222 | 0.063807 |
| GOBP_SENSORY_PERCEPTION_OF_PAIN | 85 | -0.534135 | -1.478325 | 0.006205 | 0.063915 |
| GOBP_ZINC_ION_TRANSPORT | 27 | -0.595407 | -1.478308 | 0.033058 | 0.063851 |
| GOBP_REGULATION_OF_DEFENSE_RESPONSE_TO_VIRUS_BY_VIRUS | 29 | -0.594139 | -1.478288 | 0.021557 | 0.063788 |
| GOBP_POST_ANAL_TAIL_MORPHOGENESIS | 15 | -0.675312 | -1.477832 | 0.040208 | 0.064042 |
| GOBP_REGULATION_OF_LEUKOCYTE_DEGRANULATION | 40 | -0.571392 | -1.477642 | 0.025843 | 0.064078 |
| GOBP_POSITIVE_REGULATION_OF_NERVOUS_SYSTEM_DEVELOPMENT | 254 | -0.50221 | -1.477539 | 0 | 0.064081 |
| GOBP_FEEDING_BEHAVIOR | 68 | -0.539427 | -1.477234 | 0.005285 | 0.064224 |
| GOBP_PROTEIN_LOCALIZATION_TO_LYSOSOME | 46 | -0.552093 | -1.476404 | 0.020856 | 0.064775 |
| GOBP_RETINA_VASCULATURE_DEVELOPMENT_IN_CAMERA_TYPE_EYE | 18 | -0.646975 | -1.476103 | 0.027919 | 0.064958 |
| GOBP_METANEPHROS_MORPHOGENESIS | 29 | -0.596614 | -1.475966 | 0.023148 | 0.064982 |
| GOBP_NEPHRON_TUBULE_EPITHELIAL_CELL_DIFFERENTIATION | 15 | -0.670884 | -1.475799 | 0.048656 | 0.065035 |
| GOBP_HYPOTHALAMUS_DEVELOPMENT | 15 | -0.672109 | -1.475745 | 0.037284 | 0.064997 |
| GOBP_REGULATION_OF_MEMBRANE_DEPOLARIZATION | 39 | -0.563244 | -1.475426 | 0.025901 | 0.065177 |
| GOBP_MESENCHYME_MORPHOGENESIS | 49 | -0.552605 | -1.474929 | 0.02 | 0.065463 |
| GOBP_POSITIVE_REGULATION_OF_ENDOTHELIAL_CELL_PROLIFERATION | 85 | -0.530659 | -1.4749 | 0.003083 | 0.065413 |
| GOBP_NEGATIVE_REGULATION_OF_BMP_SIGNALING_PATHWAY | 44 | -0.563526 | -1.474452 | 0.018973 | 0.065628 |
| GOBP_REGULATION_OF_CARDIAC_MUSCLE_CELL_MEMBRANE_REPOLARIZAT | 23 | -0.620222 | -1.474298 | 0.026036 | 0.065665 |
| GOBP_POSITIVE_REGULATION_OF_RECEPTOR_MEDIATED_ENDOCYTOSIS | 48 | -0.556457 | -1.473363 | 0.027263 | 0.066295 |
| GOBP_POSITIVE_REGULATION_OF_BROWN_FAT_CELL_DIFFERENTIATION | 15 | -0.664231 | -1.473159 | 0.035714 | 0.066362 |
| GOBP_REGULATION_OF_SYSTEMIC_ARTERIAL_BLOOD_PRESSURE_MEDIATED | 39 | -0.578636 | -1.47283 | 0.017837 | 0.066559 |
| GOBP_POSITIVE_REGULATION_OF_RESPONSE_TO_WOUNDING | 61 | -0.537555 | -1.471823 | 0.009424 | 0.067217 |
| GOBP_DETECTION_OF_STIMULUS_INVOLVED_IN_SENSORY_PERCEPTION | 104 | -0.516417 | -1.471779 | 0.001026 | 0.067172 |
| GOBP_REGULATION_OF_PATTERN_RECOGNITION_RECEPTOR_SIGNALING_PA | 94 | -0.523057 | -1.471421 | 0.004128 | 0.067367 |
| GOBP_SEGMENTATION | 82 | -0.523605 | -1.470635 | 0.001038 | 0.067905 |
| GOBP_ANTIGEN_PROCESSING_AND_PRESENTATION_OF_ENDOGENOUS_PEPT | 19 | -0.624588 | -1.470525 | 0.03287 | 0.067914 |
| GOBP_RESPONSE_TO_CORTICOSTERONE | 17 | -0.635932 | -1.47051 | 0.037783 | 0.067846 |
| GOBP_POSITIVE_REGULATION_OF_FIBROBLAST_PROLIFERATION | 41 | -0.567015 | -1.470373 | 0.016949 | 0.067884 |
| GOBP_REGULATION_OF_CYSTEINE_TYPE_ENDOPEPTIDASE_ACTIVITY | 220 | -0.498153 | -1.469762 | 0 | 0.068295 |
| GOBP_NEGATIVE_REGULATION_OF_GLIOGENESIS | 29 | -0.594202 | -1.469426 | 0.036643 | 0.068493 |
| GOBP_REGULATION_OF_KILLING_OF_CELLS_OF_OTHER_ORGANISM | 16 | -0.64794 | -1.469293 | 0.039241 | 0.068517 |
| GOBP_DIGESTIVE_SYSTEM_DEVELOPMENT | 118 | -0.513021 | -1.469197 | 0.001022 | 0.068507 |
| GOBP_SODIUM_ION_TRANSPORT | 213 | -0.502191 | -1.469144 | 0 | 0.068465 |
| GOBP_POSTSYNAPSE_ORGANIZATION | 156 | -0.507768 | -1.468983 | 0 | 0.068499 |
| GOBP_NEGATIVE_REGULATION_OF_CELL_MATRIX_ADHESION | 35 | -0.573583 | -1.468781 | 0.034169 | 0.068574 |
| GOBP_RECEPTOR_SIGNALING_PATHWAY_VIA_STAT | 128 | -0.513758 | -1.468702 | 0 | 0.068556 |
| GOBP_MORPHOGENESIS_OF_A_BRANCHING_STRUCTURE | 170 | -0.503916 | -1.468112 | 0 | 0.068937 |
| GOBP_POSITIVE_REGULATION_OF_SMOOTH_MUSCLE_CELL_PROLIFERATION | 79 | -0.526213 | -1.468021 | 0.006192 | 0.068932 |
| GOBP_SUBSTRATE_DEPENDENT_CELL_MIGRATION | 24 | -0.609763 | -1.467891 | 0.037485 | 0.068948 |
| GOBP_RESPONSE_TO_LIGHT_STIMULUS | 304 | -0.493742 | -1.467858 | 0 | 0.068891 |
| GOBP_POSITIVE_REGULATION_OF_STEM_CELL_DIFFERENTIATION | 17 | -0.645535 | -1.467777 | 0.038847 | 0.068884 |
| GOBP_EMBRYONIC_CAMERA_TYPE_EYE_MORPHOGENESIS | 24 | -0.608991 | -1.467659 | 0.035067 | 0.068901 |
| GOBP_NEGATIVE_REGULATION_OF_BIOMINERALIZATION | 26 | -0.604255 | -1.467546 | 0.026036 | 0.06891 |
| GOBP_REGULATION_OF_DEFENSE_RESPONSE_TO_VIRUS | 65 | -0.532754 | -1.467528 | 0.009544 | 0.068848 |
| GOBP_POSITIVE_REGULATION_OF_INTERFERON_ALPHA_PRODUCTION | 22 | -0.624327 | -1.467516 | 0.027947 | 0.06878 |
| GOBP_POSITIVE_REGULATION_OF_GLIAL_CELL_DIFFERENTIATION | 41 | -0.565905 | -1.467216 | 0.024444 | 0.068938 |
| GOBP_CELL_PROLIFERATION_IN_HINDBRAIN | 16 | -0.655215 | -1.466366 | 0.034256 | 0.069505 |
| GOBP_DETECTION_OF_TEMPERATURE_STIMULUS_INVOLVED_IN_SENSORY_F | 16 | -0.657036 | -1.466127 | 0.036943 | 0.069614 |
| GOBP_RESPONSE_TO_BMP | 149 | -0.506218 | -1.46582 | 0.001007 | 0.069759 |
| GOBP_NEGATIVE_REGULATION_OF_CELL_DEVELOPMENT | 161 | -0.510155 | -1.465705 | 0 | 0.06977 |
| GOBP_NEGATIVE_REGULATION_OF_PEPTIDASE_ACTIVITY | 195 | -0.50517 | -1.465506 | 0 | 0.069839 |
| GOBP_ENDOCARDIAL_CUSHION_MORPHOGENESIS | 33 | -0.574806 | -1.464774 | 0.025581 | 0.070327 |
| GOBP_REGULATION_OF_BRANCHING_INVOLVED_IN_URETERIC_BUD_MORPH | 19 | -0.628893 | -1.464148 | 0.0399 | 0.070732 |
| GOBP_REGULATION_OF_MAST_CELL_ACTIVATION_INVOLVED_IN_IMMUNE_RE | 28 | -0.58002 | -1.463813 | 0.034158 | 0.07092 |
| GOBP_RESPONSE_TO_ACETYLCHOLINE | 26 | -0.60201 | -1.463753 | 0.03436 | 0.070899 |
| GOBP_MHC_CLASS_II_BIOSYNTHETIC_PROCESS | 15 | -0.672011 | -1.46339 | 0.041667 | 0.071135 |
| GOBP_REACTIVE_OXYGEN_SPECIES_BIOSYNTHETIC_PROCESS | 107 | -0.510738 | -1.46268 | 0.004082 | 0.071605 |
| GOBP_EXPLORATION_BEHAVIOR | 18 | -0.643227 | -1.462627 | 0.034568 | 0.07156 |
| GOBP_REGULATION_OF_NEURON_PROJECTION_DEVELOPMENT | 413 | -0.4933 | -1.462541 | 0 | 0.071552 |
| GOBP_MONOVALENT_INORGANIC_CATION_HOMEOSTASIS | 149 | -0.506303 | -1.462431 | 0.002018 | 0.071558 |
| GOBP_CELLULAR_RESPONSE_TO_REACTIVE_NITROGEN_SPECIES | 17 | -0.650955 | -1.462206 | 0.036524 | 0.071679 |
| GOBP_NEGATIVE_REGULATION_OF_INFLAMMATORY_RESPONSE | 118 | -0.509799 | -1.460528 | 0.004094 | 0.072932 |
| GOBP_REGULATION_OF_ASTROCYTE_DIFFERENTIATION | 20 | -0.616145 | -1.460305 | 0.037944 | 0.073041 |
| GOBP_REGULATION_OF_MESENCHYMAL_CELL_PROLIFERATION | 26 | -0.60113 | -1.460037 | 0.028235 | 0.07319 |

| | | | | | |
|---|---|---|---|---|---|
| GOBP_POSITIVE_REGULATION_OF_REGULATED_SECRETORY_PATHWAY | 47 | -0.546564 | -1.459832 | 0.018681 | 0.07328 |
| GOBP_POSITIVE_REGULATION_OF_CARTILAGE_DEVELOPMENT | 27 | -0.602516 | -1.459284 | 0.030916 | 0.07369 |
| GOBP_NEGATIVE_REGULATION_OF_NOTCH_SIGNALING_PATHWAY | 37 | -0.575291 | -1.459103 | 0.020045 | 0.073766 |
| GOBP_REGULATION_OF_GLIAL_CELL_MIGRATION | 18 | -0.636457 | -1.459078 | 0.033248 | 0.073701 |
| GOBP_VENTRAL_SPINAL_CORD_DEVELOPMENT | 40 | -0.567042 | -1.458814 | 0.023702 | 0.073826 |
| GOBP_CHONDROCYTE_DEVELOPMENT | 28 | -0.587535 | -1.458766 | 0.033373 | 0.073783 |
| GOBP_MEMBRANE_REPOLARIZATION | 41 | -0.559776 | -1.4587 | 0.023333 | 0.073764 |
| GOBP_REGULATION_OF_VESICLE_MEDIATED_TRANSPORT | 486 | -0.492961 | -1.457645 | 0 | 0.07455 |
| GOBP_SYNAPTIC_TRANSMISSION_GABAERGIC | 43 | -0.562469 | -1.457516 | 0.020902 | 0.074568 |
| GOBP_ENDOTHELIAL_CELL_APOPTOTIC_PROCESS | 50 | -0.544289 | -1.457467 | 0.020541 | 0.074535 |
| GOBP_TELENCEPHALON_GLIAL_CELL_MIGRATION | 22 | -0.612687 | -1.457397 | 0.041463 | 0.074508 |
| GOBP_SIGNAL_RELEASE | 441 | -0.486997 | -1.45632 | 0 | 0.075296 |
| GOBP_CELLULAR_MONOVALENT_INORGANIC_CATION_HOMEOSTASIS | 109 | -0.512265 | -1.455598 | 0.006141 | 0.075825 |
| GOBP_REGULATION_OF_POSTSYNAPTIC_NEUROTRANSMITTER_RECEPTOR_A | 16 | -0.650395 | -1.455113 | 0.040404 | 0.076155 |
| GOBP_FEMALE_GENITALIA_DEVELOPMENT | 15 | -0.667992 | -1.455086 | 0.041772 | 0.076094 |
| GOBP_EPITHELIAL_CELL_PROLIFERATION | 353 | -0.49384 | -1.454878 | 0 | 0.076193 |
| GOBP_ARTERY_MORPHOGENESIS | 61 | -0.534839 | -1.454484 | 0.01921 | 0.076449 |
| GOBP_BIOLOGICAL_PROCESS_INVOLVED_IN_INTERACTION_WITH_SYMBIONT | 75 | -0.522187 | -1.454335 | 0.008359 | 0.076504 |
| GOBP_GASTRULATION | 165 | -0.502941 | -1.454029 | 0.001007 | 0.076662 |
| GOBP_CLATHRIN_COAT_ASSEMBLY | 18 | -0.635431 | -1.453771 | 0.039312 | 0.076777 |
| GOBP_NEGATIVE_REGULATION_OF_G_PROTEIN_COUPLED_RECEPTOR_SIGNA | 45 | -0.557245 | -1.453091 | 0.027293 | 0.077272 |
| GOBP_ORGANIC_ACID_TRANSPORT | 287 | -0.492778 | -1.452249 | 0 | 0.077939 |
| GOBP_REGULATION_OF_MORPHOGENESIS_OF_A_BRANCHING_STRUCTURE | 44 | -0.554376 | -1.451651 | 0.020971 | 0.078348 |
| GOBP_REGULATION_OF_HORMONE_METABOLIC_PROCESS | 33 | -0.571139 | -1.451612 | 0.028769 | 0.078301 |
| GOBP_POSITIVE_REGULATION_OF_EPITHELIAL_TO_MESENCHYMAL_TRANSITI | 48 | -0.551113 | -1.451493 | 0.023939 | 0.07832 |
| GOBP_GLOMERULAR_EPITHELIAL_CELL_DIFFERENTIATION | 18 | -0.632811 | -1.451272 | 0.034308 | 0.078419 |
| GOBP_REGULATION_OF_PEPTIDYL_TYROSINE_PHOSPHORYLATION | 232 | -0.49465 | -1.451236 | 0 | 0.078362 |
| GOBP_CELLULAR_RESPONSE_TO_ESTRADIOL_STIMULUS | 33 | -0.577938 | -1.450637 | 0.035469 | 0.078799 |
| GOBP_CELLULAR_RESPONSE_TO_INORGANIC_SUBSTANCE | 199 | -0.496011 | -1.450482 | 0 | 0.078864 |
| GOBP_DENDRITIC_SPINE_MORPHOGENESIS | 60 | -0.535667 | -1.450268 | 0.011616 | 0.078952 |
| GOBP_NEGATIVE_REGULATION_OF_CELLULAR_RESPONSE_TO_GROWTH_FAC | 87 | -0.519518 | -1.449814 | 0.009326 | 0.079292 |
| GOBP_SECONDARY_PALATE_DEVELOPMENT | 24 | -0.59587 | -1.449664 | 0.043103 | 0.079337 |
| GOBP_POSITIVE_REGULATION_OF_FATTY_ACID_METABOLIC_PROCESS | 29 | -0.582664 | -1.449556 | 0.036428 | 0.079354 |
| GOBP_EMBRYONIC_EYE_MORPHOGENESIS | 32 | -0.571367 | -1.449534 | 0.026651 | 0.079287 |
| GOBP_CATECHOL_CONTAINING_COMPOUND_BIOSYNTHETIC_PROCESS | 18 | -0.643457 | -1.449377 | 0.034869 | 0.079344 |
| GOBP_POSTSYNAPTIC_SPECIALIZATION_ORGANIZATION | 31 | -0.576523 | -1.44916 | 0.038642 | 0.079449 |
| GOBP_PURINE_NUCLEOBASE_METABOLIC_PROCESS | 18 | -0.63056 | -1.448836 | 0.049751 | 0.079668 |
| GOBP_MUSCLE_CELL_PROLIFERATION | 174 | -0.498037 | -1.448321 | 0 | 0.079981 |
| GOBP_ADULT_LOCOMOTORY_BEHAVIOR | 56 | -0.529377 | -1.448221 | 0.016129 | 0.079987 |
| GOBP_POSITIVE_REGULATION_OF_TRANSCRIPTION_REGULATORY_REGION_D | 24 | -0.598242 | -1.447943 | 0.035885 | 0.080125 |
| GOBP_BROWN_FAT_CELL_DIFFERENTIATION | 43 | -0.554469 | -1.447405 | 0.018579 | 0.080487 |
| GOBP_REGULATION_OF_HORMONE_BIOSYNTHETIC_PROCESS | 20 | -0.62515 | -1.447103 | 0.038462 | 0.080671 |
| GOBP_REGULATION_OF_RESPONSE_TO_CYTOKINE_STIMULUS | 166 | -0.498418 | -1.44676 | 0.002016 | 0.080895 |
| GOBP_SPINAL_CORD_PATTERNING | 18 | -0.643683 | -1.446755 | 0.045792 | 0.080816 |
| GOBP_NEGATIVE_REGULATION_OF_NEURON_APOPTOTIC_PROCESS | 126 | -0.507506 | -1.446639 | 0.007064 | 0.08084 |
| GOBP_TUMOR_NECROSIS_FACTOR_MEDIATED_SIGNALING_PATHWAY | 162 | -0.504665 | -1.446574 | 0 | 0.080819 |
| GOBP_ENDOCRINE_PANCREAS_DEVELOPMENT | 40 | -0.562029 | -1.446284 | 0.020157 | 0.081007 |
| GOBP_ENDOTHELIAL_CELL_PROLIFERATION | 137 | -0.499446 | -1.445364 | 0.001011 | 0.081723 |
| GOBP_GENITALIA_DEVELOPMENT | 35 | -0.56707 | -1.445291 | 0.036117 | 0.081702 |
| GOBP_EMBRYONIC_DIGESTIVE_TRACT_DEVELOPMENT | 29 | -0.588815 | -1.445176 | 0.031746 | 0.081717 |
| GOBP_ANTIGEN_PROCESSING_AND_PRESENTATION_OF_PEPTIDE_OR_POLYS | 99 | -0.511702 | -1.444941 | 0.010256 | 0.081863 |
| GOBP_CELL_DIFFERENTIATION_IN_SPINAL_CORD | 43 | -0.54965 | -1.444833 | 0.030405 | 0.081872 |
| GOBP_SPINAL_CORD_MOTOR_NEURON_DIFFERENTIATION | 27 | -0.584919 | -1.444463 | 0.036385 | 0.08209 |
| GOBP_LIPID_EXPORT_FROM_CELL | 75 | -0.523379 | -1.444399 | 0.011518 | 0.08208 |
| GOBP_NEUROPEPTIDE_SIGNALING_PATHWAY | 76 | -0.524576 | -1.443806 | 0.016615 | 0.082482 |
| GOBP_BIOMINERALIZATION | 144 | -0.505556 | -1.442939 | 0 | 0.083092 |
| GOBP_CELL_MIGRATION_INVOLVED_IN_HEART_DEVELOPMENT | 17 | -0.637546 | -1.442643 | 0.053284 | 0.083267 |
| GOBP_KILLING_OF_CELLS_OF_OTHER_ORGANISM | 40 | -0.552897 | -1.441757 | 0.030405 | 0.083978 |
| GOBP_RETINOL_METABOLIC_PROCESS | 41 | -0.557751 | -1.44087 | 0.025901 | 0.084674 |
| GOBP_NEGATIVE_REGULATION_OF_SECRETION | 135 | -0.501507 | -1.440759 | 0.004049 | 0.08469 |
| GOBP_ACTIN_CYTOSKELETON_REORGANIZATION | 99 | -0.506194 | -1.43947 | 0.008222 | 0.085755 |
| GOBP_PROTEIN_HETEROOLIGOMERIZATION | 23 | -0.601762 | -1.439284 | 0.043587 | 0.085847 |
| GOBP_RESPONSE_TO_DOPAMINE | 68 | -0.525317 | -1.439241 | 0.016931 | 0.085794 |
| GOBP_REGULATION_OF_PHOSPHOLIPID_BIOSYNTHETIC_PROCESS | 15 | -0.649032 | -1.439194 | 0.045455 | 0.085756 |
| GOBP_POSITIVE_REGULATION_OF_HEART_RATE | 24 | -0.599402 | -1.438198 | 0.054054 | 0.086576 |
| GOBP_POSITIVE_REGULATION_OF_REACTIVE_OXYGEN_SPECIES_BIOSYNTHE | 47 | -0.541387 | -1.43816 | 0.036504 | 0.086513 |
| GOBP_CARBOXYLIC_ACID_TRANSPORT | 245 | -0.48826 | -1.437573 | 0 | 0.086963 |
| GOBP_POSITIVE_REGULATION_OF_VASCULATURE_DEVELOPMENT | 145 | -0.50022 | -1.43747 | 0.002024 | 0.086984 |
| GOBP_NECROPTOTIC_PROCESS | 37 | -0.555079 | -1.437092 | 0.042824 | 0.087244 |
| GOBP_REGULATION_OF_CELLULAR_RESPONSE_TO_GROWTH_FACTOR_STIMU | 253 | -0.493051 | -1.436887 | 0 | 0.087341 |
| GOBP_NEGATIVE_REGULATION_OF_CHEMOTAXIS | 57 | -0.535587 | -1.436751 | 0.021459 | 0.087373 |
| GOBP_PEPTIDYL_TYROSINE_MODIFICATION | 340 | -0.490778 | -1.436647 | 0 | 0.087375 |
| GOBP_POSITIVE_REGULATION_OF_CHONDROCYTE_DIFFERENTIATION | 17 | -0.630938 | -1.436459 | 0.04625 | 0.087481 |
| GOBP_REGULATION_OF_EPITHELIAL_CELL_APOPTOTIC_PROCESS | 75 | -0.520199 | -1.436206 | 0.009414 | 0.087578 |
| GOBP_PLATELET_DEGRANULATION | 115 | -0.501261 | -1.435979 | 0.009174 | 0.0877 |
| GOBP_REGULATION_OF_ENDOTHELIAL_CELL_CHEMOTAXIS | 18 | -0.637889 | -1.435752 | 0.036524 | 0.087839 |
| GOBP_APPENDAGE_DEVELOPMENT | 156 | -0.500712 | -1.43564 | 0.001009 | 0.087851 |
| GOBP_CHONDROCYTE_DIFFERENTIATION | 100 | -0.511378 | -1.435473 | 0.010277 | 0.087907 |
| GOBP_REGULATION_OF_ACTIN_CYTOSKELETON_REORGANIZATION | 36 | -0.567954 | -1.43539 | 0.038331 | 0.087905 |
| GOBP_REGULATION_OF_MYELINATION | 40 | -0.555369 | -1.435143 | 0.032294 | 0.088064 |
| GOBP_UNSATURATED_FATTY_ACID_BIOSYNTHETIC_PROCESS | 45 | -0.540112 | -1.434254 | 0.027655 | 0.0888 |
| GOBP_IMMUNOGLOBULIN_PRODUCTION_INVOLVED_IN_IMMUNOGLOBULIN_MI | 51 | -0.540346 | -1.434211 | 0.020697 | 0.088759 |
| GOBP_NEGATIVE_CHEMOTAXIS | 46 | -0.542175 | -1.433653 | 0.026432 | 0.089179 |
| GOBP_REGULATION_OF_VASCULATURE_DEVELOPMENT | 264 | -0.490126 | -1.433412 | 0 | 0.089321 |
| GOBP_ANTIGEN_PROCESSING_AND_PRESENTATION_OF_PEPTIDE_ANTIGEN | 187 | -0.492444 | -1.433179 | 0.001007 | 0.089447 |
| GOBP_MACROPHAGE_CYTOKINE_PRODUCTION | 15 | -0.647271 | -1.432959 | 0.062901 | 0.089563 |
| GOBP_POSITIVE_CHEMOTAXIS | 57 | -0.528413 | -1.432782 | 0.028723 | 0.08963 |
| GOBP_REGULATION_OF_OSTEOBLAST_DIFFERENTIATION | 113 | -0.504398 | -1.431952 | 0 | 0.090277 |
| GOBP_SODIUM_ION_HOMEOSTASIS | 46 | -0.535213 | -1.431227 | 0.041758 | 0.090858 |
| GOBP_REGULATION_OF_NEUROGENESIS | 329 | -0.482702 | -1.431175 | 0 | 0.090818 |
| GOBP_LYSOSOME_LOCALIZATION | 71 | -0.514792 | -1.431102 | 0.017186 | 0.090799 |
| GOBP_NEUROTRANSMITTER_REUPTAKE | 28 | -0.575693 | -1.431002 | 0.041909 | 0.0908 |
| GOBP_RENAL_SYSTEM_PROCESS_INVOLVED_IN_REGULATION_OF_SYSTEMIC | 20 | -0.620545 | -1.430857 | 0.048662 | 0.090841 |
| GOBP_SEROTONIN_RECEPTOR_SIGNALING_PATHWAY | 19 | -0.623307 | -1.430036 | 0.046135 | 0.091539 |
| GOBP_RESPIRATORY_SYSTEM_PROCESS | 29 | -0.576214 | -1.429894 | 0.038462 | 0.09156 |
| GOBP_ACTIVATION_OF_PROTEIN_KINASE_B_ACTIVITY | 31 | -0.580514 | -1.428865 | 0.045828 | 0.092414 |
| GOBP_REGULATION_OF_TOLL_LIKE_RECEPTOR_4_SIGNALING_PATHWAY | 21 | -0.606419 | -1.428258 | 0.070574 | 0.092904 |
| GOBP_REGULATION_OF_EPITHELIAL_CELL_DIFFERENTIATION | 131 | -0.500717 | -1.427909 | 0.002024 | 0.09311 |
| GOBP_RETINAL_METABOLIC_PROCESS | 15 | -0.647907 | -1.427383 | 0.058385 | 0.093522 |
| GOBP_POSITIVE_REGULATION_OF_DEVELOPMENTAL_GROWTH | 152 | -0.498719 | -1.426929 | 0.003033 | 0.093874 |
| GOBP_SKELETAL_MUSCLE_CELL_DIFFERENTIATION | 58 | -0.52182 | -1.426876 | 0.020106 | 0.09382 |
| GOBP_REGULATION_OF_CARDIOCYTE_DIFFERENTIATION | 27 | -0.581322 | -1.426824 | 0.048578 | 0.093773 |
| GOBP_METANEPHRIC_NEPHRON_MORPHOGENESIS | 24 | -0.585067 | -1.42665 | 0.051916 | 0.093839 |
| GOBP_NEGATIVE_REGULATION_OF_CALCIUM_ION_TRANSPORT | 49 | -0.54535 | -1.426536 | 0.0299 | 0.093858 |
| GOBP_NEGATIVE_REGULATION_OF_RESPONSE_TO_CYTOKINE_STIMULUS | 60 | -0.531451 | -1.42591 | 0.027897 | 0.094381 |
| GOBP_POSITIVE_REGULATION_OF_CELL_DIVISION | 69 | -0.511833 | -1.425864 | 0.01797 | 0.094336 |
| GOBP_UROGENITAL_SYSTEM_DEVELOPMENT | 301 | -0.482873 | -1.425104 | 0 | 0.094978 |
| GOBP_OUTFLOW_TRACT_MORPHOGENESIS | 68 | -0.517081 | -1.424705 | 0.016985 | 0.095286 |
| GOBP_MEMBRANE_RAFT_ORGANIZATION | 24 | -0.586395 | -1.424107 | 0.055425 | 0.095802 |
| GOBP_XENOBIOTIC_TRANSPORT | 37 | -0.550839 | -1.423263 | 0.042529 | 0.096482 |
| GOBP_REGULATION_OF_BLOOD_PRESSURE | 154 | -0.49183 | -1.423261 | 0.001009 | 0.09639 |
| GOBP_REGULATION_OF_TYPE_I_INTERFERON_MEDIATED_SIGNALING_PATHW | 33 | -0.564347 | -1.423193 | 0.038106 | 0.096364 |
| GOBP_ANTIGEN_PROCESSING_AND_PRESENTATION_OF_EXOGENOUS_PEPTI | 79 | -0.512291 | -1.422707 | 0.010331 | 0.096773 |
| GOBP_REGULATION_OF_CARDIAC_MUSCLE_CONTRACTION_BY_CALCIUM_ION | 25 | -0.586189 | -1.422593 | 0.059032 | 0.096791 |
| GOBP_NEGATIVE_REGULATION_OF_HYDROLASE_ACTIVITY | 380 | -0.481133 | -1.422576 | 0 | 0.096716 |
| GOBP_POSITIVE_REGULATION_OF_OXIDOREDUCTASE_ACTIVITY | 52 | -0.533467 | -1.421822 | 0.024973 | 0.097315 |
| GOBP_POSITIVE_REGULATION_OF_PROTEIN_EXIT_FROM_ENDOPLASMIC_RET | 15 | -0.639084 | -1.421765 | 0.051702 | 0.097282 |
| GOBP_POSITIVE_REGULATION_OF_P38MAPK_CASCADE | 26 | -0.574922 | -1.421579 | 0.061033 | 0.097352 |
| GOBP_PALLIUM_DEVELOPMENT | 143 | -0.493773 | -1.42075 | 0.003018 | 0.098096 |
| GOBP_POSITIVE_REGULATION_OF_CALCIUM_MEDIATED_SIGNALING | 27 | -0.584345 | -1.4207 | 0.049939 | 0.098064 |
| GOBP_POSITIVE_REGULATION_OF_LIPID_METABOLIC_PROCESS | 130 | -0.486455 | -1.420686 | 0.007085 | 0.097982 |
| GOBP_RENAL_VESICLE_DEVELOPMENT | 19 | -0.611383 | -1.420525 | 0.064242 | 0.098048 |
| GOBP_G_PROTEIN_COUPLED_GLUTAMATE_RECEPTOR_SIGNALING_PATHWAY | 15 | -0.64076 | -1.420116 | 0.05443 | 0.098355 |
| GOBP_MESODERM_MORPHOGENESIS | 62 | -0.521339 | -1.419363 | 0.018104 | 0.09899 |
| GOBP_INTERLEUKIN_6_MEDIATED_SIGNALING_PATHWAY | 22 | -0.599396 | -1.419026 | 0.057178 | 0.099243 |
| GOBP_FEAR_RESPONSE | 33 | -0.561573 | -1.418939 | 0.052632 | 0.099227 |
| GOBP_NEGATIVE_REGULATION_OF_TRANSPORT | 379 | -0.475843 | -1.418785 | 0 | 0.099311 |
| GOBP_REGULATION_OF_PROTEIN_TYROSINE_KINASE_ACTIVITY | 92 | -0.506048 | -1.418781 | 0.009288 | 0.09922 |
| GOBP_NEGATIVE_REGULATION_OF_BONE_MINERALIZATION | 17 | -0.625713 | -1.417848 | 0.060837 | 0.100067 |
| GOBP_POSITIVE_REGULATION_OF_RECEPTOR_SIGNALING_PATHWAY_VIA_ST | 38 | -0.55372 | -1.416728 | 0.041387 | 0.101141 |
| GOBP_SMOOTH_MUSCLE_CELL_PROLIFERATION | 124 | -0.494679 | -1.416727 | 0.004077 | 0.101046 |
| GOBP_REGULATION_OF_TRANSMEMBRANE_RECEPTOR_PROTEIN_SERINE_TH | 225 | -0.484094 | -1.416695 | 0 | 0.100978 |
| GOBP_PEPTIDE_CATABOLIC_PROCESS | 26 | -0.586995 | -1.416514 | 0.055819 | 0.101075 |
| GOBP_POSITIVE_REGULATION_OF_GLYCOGEN_METABOLIC_PROCESS | 16 | -0.637701 | -1.416163 | 0.06125 | 0.101308 |
| GOBP_NEGATIVE_REGULATION_OF_LIPID_CATABOLIC_PROCESS | 22 | -0.608004 | -1.41602 | 0.0489 | 0.101356 |
| GOBP_REGULATION_OF_LIPOPOLYSACCHARIDE_MEDIATED_SIGNALING_PATH | 22 | -0.600732 | -1.415921 | 0.066106 | 0.101388 |
| GOBP_CORONARY_VASCULATURE_MORPHOGENESIS | 15 | -0.633101 | -1.415753 | 0.065789 | 0.101479 |
| GOBP_REGULATION_OF_VASOCONSTRICTION | 55 | -0.527144 | -1.415583 | 0.031522 | 0.101549 |
| GOBP_RESPONSE_TO_THYROID_HORMONE | 22 | -0.600648 | -1.41522 | 0.059952 | 0.101806 |
| GOBP_HINDLIMB_MORPHOGENESIS | 32 | -0.561566 | -1.415069 | 0.053592 | 0.101853 |
| GOBP_POSITIVE_REGULATION_OF_PROTEIN_LOCALIZATION_TO_CELL_SURFA | 16 | -0.629461 | -1.414867 | 0.062025 | 0.101952 |
| GOBP_REGULATION_OF_RYANODINE_SENSITIVE_CALCIUM_RELEASE_CHANN | 26 | -0.590266 | -1.414726 | 0.04717 | 0.101994 |
| GOBP_POSITIVE_REGULATION_OF_CALCIUM_ION_TRANSMEMBRANE_TRANSF | 37 | -0.548648 | -1.414349 | 0.033143 | 0.102264 |
| GOBP_SYNAPSE_MATURATION | 24 | -0.578612 | -1.412651 | 0.06383 | 0.103869 |
| GOBP_BONE_DEVELOPMENT | 182 | -0.487966 | -1.412514 | 0.001004 | 0.103923 |
| GOBP_POSITIVE_REGULATION_OF_NUCLEOTIDE_BIOSYNTHETIC_PROCESS | 19 | -0.619539 | -1.412292 | 0.055767 | 0.104052 |
| GOBP_ACUTE_PHASE_RESPONSE | 33 | -0.555863 | -1.411475 | 0.048499 | 0.104825 |
| GOBP_REGULATION_OF_NIK_NF_KAPPAB_SIGNALING | 90 | -0.500806 | -1.411467 | 0.021516 | 0.104736 |
| GOBP_REGULATION_OF_BROWN_FAT_CELL_DIFFERENTIATION | 21 | -0.59652 | -1.411369 | 0.06747 | 0.104725 |
| GOBP_ORGANIC_ANION_TRANSPORT | 327 | -0.478738 | -1.411341 | 0 | 0.104654 |
| GOBP_REGULATION_OF_OXIDOREDUCTASE_ACTIVITY | 95 | -0.495748 | -1.411084 | 0.019588 | 0.104818 |

| | | | | | |
|---|---|---|---|---|---|
| GOBP_MESODERMAL_CELL_DIFFERENTIATION | 28 | -0.565548 | -1.410538 | 0.053676 | 0.105278 |
| GOBP_PROTEIN_TRIMERIZATION | 17 | -0.621669 | -1.410433 | 0.068408 | 0.105294 |
| GOBP_NEGATIVE_REGULATION_OF_TRANSMEMBRANE_TRANSPORT | 106 | -0.50085 | -1.41013 | 0.006135 | 0.105492 |
| GOBP_REGULATION_OF_REACTIVE_OXYGEN_SPECIES_BIOSYNTHETIC_PROC | 84 | -0.504543 | -1.40984 | 0.015496 | 0.105673 |
| GOBP_CD40_SIGNALING_PATHWAY | 15 | -0.639409 | -1.409659 | 0.047558 | 0.105754 |
| GOBP_POSITIVE_REGULATION_OF_DNA_DAMAGE_RESPONSE_SIGNAL_TRANS | 15 | -0.641238 | -1.409481 | 0.071429 | 0.105848 |
| GOBP_VESICLE_FUSION_TO_PLASMA_MEMBRANE | 20 | -0.608328 | -1.408534 | 0.063119 | 0.106733 |
| GOBP_POSITIVE_REGULATION_OF_MEMBRANE_PROTEIN_ECTODOMAIN_PROT | 15 | -0.62761 | -1.408281 | 0.071141 | 0.106918 |
| GOBP_CARDIOCYTE_DIFFERENTIATION | 116 | -0.495778 | -1.407966 | 0.005097 | 0.107117 |
| GOBP_TOLL_LIKE_RECEPTOR_SIGNALING_PATHWAY | 144 | -0.48509 | -1.407699 | 0.006054 | 0.107281 |
| GOBP_NEGATIVE_REGULATION_OF_TRANSMEMBRANE_RECEPTOR_PROTEIN_ | 119 | -0.494925 | -1.40713 | 0.006085 | 0.107767 |
| GOBP_NEGATIVE_REGULATION_OF_AMYLOID_PRECURSOR_PROTEIN_CATAB | 23 | -0.584953 | -1.407074 | 0.049757 | 0.107718 |
| GOBP_REGULATION_OF_MAST_CELL_ACTIVATION | 38 | -0.551461 | -1.406384 | 0.048753 | 0.108371 |
| GOBP_CALCIUM_MEDIATED_SIGNALING_USING_INTRACELLULAR_CALCIUM_S | 22 | -0.588446 | -1.406095 | 0.058111 | 0.108561 |
| GOBP_NEURAL_NUCLEUS_DEVELOPMENT | 56 | -0.524897 | -1.405996 | 0.032909 | 0.108582 |
| GOBP_ATRIAL_SEPTUM_DEVELOPMENT | 21 | -0.597254 | -1.405866 | 0.059322 | 0.108609 |
| GOBP_FOREBRAIN_REGIONALIZATION | 17 | -0.625037 | -1.405672 | 0.060102 | 0.1087 |
| GOBP_REGULATION_OF_CALCIUM_MEDIATED_SIGNALING | 61 | -0.515812 | -1.405598 | 0.020234 | 0.108674 |
| GOBP_BLOOD_VESSEL_ENDOTHELIAL_CELL_PROLIFERATION_INVOLVED_IN_ | 21 | -0.582197 | -1.405461 | 0.060197 | 0.108723 |
| GOBP_NEGATIVE_REGULATION_OF_CELL_SUBSTRATE_ADHESION | 57 | -0.520243 | -1.404602 | 0.028785 | 0.109551 |
| GOBP_TRANSITION_METAL_ION_HOMEOSTASIS | 129 | -0.488013 | -1.404426 | 0.004057 | 0.109646 |
| GOBP_PATHWAY_RESTRICTED_SMAD_PROTEIN_PHOSPHORYLATION | 60 | -0.5136 | -1.404411 | 0.031915 | 0.109561 |
| GOBP_GLANDULAR_EPITHELIAL_CELL_DIFFERENTIATION | 41 | -0.540094 | -1.404156 | 0.029573 | 0.109738 |
| GOBP_NEGATIVE_REGULATION_OF_NEURON_DEATH | 180 | -0.482849 | -1.403489 | 0.002008 | 0.110349 |
| GOBP_PROTEOGLYCAN_METABOLIC_PROCESS | 85 | -0.504077 | -1.403303 | 0.01343 | 0.110456 |
| GOBP_TRANSITION_METAL_ION_TRANSPORT | 119 | -0.48996 | -1.402888 | 0.00611 | 0.110779 |
| GOBP_AMYLOID_PRECURSOR_PROTEIN_METABOLIC_PROCESS | 71 | -0.506553 | -1.402331 | 0.02328 | 0.111297 |
| GOBP_NEGATIVE_REGULATION_OF_TOLL_LIKE_RECEPTOR_SIGNALING_PATH | 37 | -0.550742 | -1.402247 | 0.034169 | 0.111289 |
| GOBP_CELLULAR_RESPONSE_TO_RADIATION | 182 | -0.484323 | -1.402182 | 0.003006 | 0.11126 |
| GOBP_POSITIVE_REGULATION_OF_MYOBLAST_FUSION | 15 | -0.634791 | -1.402024 | 0.071336 | 0.111339 |
| GOBP_INTEGRIN_MEDIATED_SIGNALING_PATHWAY | 106 | -0.495946 | -1.401737 | 0.012245 | 0.111541 |
| GOBP_CELLULAR_RESPONSE_TO_ABIOTIC_STIMULUS | 316 | -0.478145 | -1.40144 | 0.001 | 0.111724 |
| GOBP_POSITIVE_REGULATION_OF_CYSTEINE_TYPE_ENDOPEPTIDASE_ACTIVI | 147 | -0.482358 | -1.400965 | 0.009119 | 0.112094 |
| GOBP_NEGATIVE_REGULATION_OF_MUSCLE_CELL_DIFFERENTIATION | 51 | -0.526568 | -1.400906 | 0.039742 | 0.112064 |
| GOBP_COPPER_ION_TRANSPORT | 16 | -0.613295 | -1.400733 | 0.069136 | 0.112159 |
| GOBP_PROTEOGLYCAN_BIOSYNTHETIC_PROCESS | 61 | -0.514564 | -1.400318 | 0.036364 | 0.112522 |
| GOBP_RESPONSE_TO_INTERFERON_BETA | 25 | -0.576169 | -1.400055 | 0.053333 | 0.112688 |
| GOBP_REGULATION_OF_SYSTEMIC_ARTERIAL_BLOOD_PRESSURE_BY_HORM | 30 | -0.56634 | -1.399815 | 0.043478 | 0.112853 |
| GOBP_MAINTENANCE_OF_SYNAPSE_STRUCTURE | 16 | -0.630151 | -1.399203 | 0.063131 | 0.113439 |
| GOBP_CYCLIC_NUCLEOTIDE_MEDIATED_SIGNALING | 78 | -0.494491 | -1.399049 | 0.023933 | 0.113499 |
| GOBP_RESPONSE_TO_PAIN | 22 | -0.585864 | -1.398647 | 0.076831 | 0.113841 |
| GOBP_NEGATIVE_REGULATION_OF_SMOOTH_MUSCLE_CELL_MIGRATION | 20 | -0.600893 | -1.398397 | 0.068096 | 0.113991 |
| GOBP_POSITIVE_REGULATION_OF_CARDIAC_MUSCLE_CELL_PROLIFERATION | 23 | -0.585496 | -1.398374 | 0.089791 | 0.113907 |
| GOBP_URETER_DEVELOPMENT | 18 | -0.608247 | -1.398057 | 0.081321 | 0.114179 |
| GOBP_CELLULAR_RESPONSE_TO_INTERFERON_BETA | 16 | -0.626258 | -1.397141 | 0.066411 | 0.115052 |
| GOBP_CENTRAL_NERVOUS_SYSTEM_NEURON_DEVELOPMENT | 79 | -0.500066 | -1.3971 | 0.022013 | 0.114999 |
| GOBP_ANATOMICAL_STRUCTURE_HOMEOSTASIS | 443 | -0.471352 | -1.396868 | 0 | 0.115161 |
| GOBP_POSITIVE_REGULATION_OF_CELL_MIGRATION_INVOLVED_IN_SPROUTI | 20 | -0.606781 | -1.396614 | 0.063119 | 0.115331 |
| GOBP_GLYCOLIPID_BIOSYNTHETIC_PROCESS | 66 | -0.504811 | -1.39656 | 0.031088 | 0.115284 |
| GOBP_CARDIAC_MUSCLE_CONTRACTION | 125 | -0.489711 | -1.396403 | 0.008188 | 0.115381 |
| GOBP_PROTEIN_KINASE_C_ACTIVATING_G_PROTEIN_COUPLED_RECEPTOR_ | 25 | -0.572612 | -1.395955 | 0.062426 | 0.115775 |
| GOBP_LYMPHOID_PROGENITOR_CELL_DIFFERENTIATION | 20 | -0.602037 | -1.395632 | 0.066015 | 0.116062 |
| GOBP_MYD88_DEPENDENT_TOLL_LIKE_RECEPTOR_SIGNALING_PATHWAY | 35 | -0.549449 | -1.395597 | 0.039503 | 0.115997 |
| GOBP_REGULATION_OF_POTASSIUM_ION_TRANSMEMBRANE_TRANSPORTER | 54 | -0.512306 | -1.395545 | 0.047619 | 0.115957 |
| GOBP_REGULATION_OF_PEPTIDASE_ACTIVITY | 384 | -0.471014 | -1.395284 | 0 | 0.116191 |
| GOBP_ACTIN_MEDIATED_CELL_CONTRACTION | 111 | -0.486432 | -1.395052 | 0.012397 | 0.116361 |
| GOBP_REGULATION_OF_CARDIAC_MUSCLE_CONTRACTION | 69 | -0.507309 | -1.394394 | 0.021322 | 0.116956 |
| GOBP_VASCULAR_ENDOTHELIAL_GROWTH_FACTOR_PRODUCTION | 37 | -0.541419 | -1.394251 | 0.046243 | 0.117006 |
| GOBP_ENDOCARDIAL_CUSHION_FORMATION | 24 | -0.581365 | -1.393554 | 0.07177 | 0.117667 |
| GOBP_POSTSYNAPTIC_SPECIALIZATION_ASSEMBLY | 21 | -0.598156 | -1.39329 | 0.066185 | 0.117869 |
| GOBP_REGULATION_OF_NITRIC_OXIDE_METABOLIC_PROCESS | 51 | -0.526157 | -1.392794 | 0.040087 | 0.11834 |
| GOBP_NEURAL_CREST_CELL_DIFFERENTIATION | 86 | -0.49508 | -1.392755 | 0.02079 | 0.118279 |
| GOBP_CELL_COMMUNICATION_BY_ELECTRICAL_COUPLING | 30 | -0.558362 | -1.39253 | 0.065116 | 0.118405 |
| GOBP_POSITIVE_REGULATION_OF_NOTCH_SIGNALING_PATHWAY | 52 | -0.518084 | -1.392526 | 0.037199 | 0.118304 |
| GOBP_NEUROTRANSMITTER_UPTAKE | 34 | -0.552752 | -1.392401 | 0.053899 | 0.118356 |
| GOBP_AMINOGLYCAN_BIOSYNTHETIC_PROCESS | 110 | -0.485629 | -1.392391 | 0.012283 | 0.118262 |
| GOBP_NEGATIVE_REGULATION_OF_DEFENSE_RESPONSE_TO_VIRUS | 24 | -0.578492 | -1.392117 | 0.057528 | 0.118476 |
| GOBP_PROGRAMMED_NECROTIC_CELL_DEATH | 45 | -0.530946 | -1.391913 | 0.044053 | 0.118593 |
| GOBP_NEGATIVE_REGULATION_OF_OXIDOREDUCTASE_ACTIVITY | 23 | -0.58132 | -1.391611 | 0.064118 | 0.118819 |
| GOBP_REGULATION_OF_SODIUM_ION_TRANSMEMBRANE_TRANSPORT | 60 | -0.512343 | -1.391561 | 0.03617 | 0.118763 |
| GOBP_POSITIVE_REGULATION_OF_NITRIC_OXIDE_SYNTHASE_ACTIVITY | 22 | -0.591214 | -1.390511 | 0.061151 | 0.119812 |
| GOBP_REGULATION_OF_FATTY_ACID_BIOSYNTHETIC_PROCESS | 39 | -0.544297 | -1.390234 | 0.040495 | 0.120013 |
| GOBP_POSITIVE_REGULATION_OF_ACTIVATED_T_CELL_PROLIFERATION | 20 | -0.602398 | -1.390012 | 0.067081 | 0.120122 |
| GOBP_SIGNAL_TRANSDUCTION_INVOLVED_IN_REGULATION_OF_GENE_EXPR | 16 | -0.61425 | -1.389154 | 0.083969 | 0.120935 |
| GOBP_HOMEOSTASIS_OF_NUMBER_OF_CELLS | 242 | -0.47454 | -1.389003 | 0.001 | 0.120998 |
| GOBP_ACTIVATED_T_CELL_PROLIFERATION | 38 | -0.531848 | -1.388603 | 0.057208 | 0.121337 |
| GOBP_NEUROMUSCULAR_JUNCTION_DEVELOPMENT | 45 | -0.522388 | -1.38832 | 0.052632 | 0.121578 |
| GOBP_DERMATAN_SULFATE_PROTEOGLYCAN_METABOLIC_PROCESS | 16 | -0.624675 | -1.388241 | 0.0825 | 0.121581 |
| GOBP_CELL_SURFACE_RECEPTOR_SIGNALING_PATHWAY_INVOLVED_IN_HEA | 27 | -0.563859 | -1.38789 | 0.066829 | 0.121878 |
| GOBP_RECEPTOR_CLUSTERING | 51 | -0.514553 | -1.38709 | 0.040481 | 0.122696 |
| GOBP_REGULATION_OF_MYOBLAST_DIFFERENTIATION | 45 | -0.526407 | -1.386968 | 0.052805 | 0.122738 |
| GOBP_REGULATION_OF_HETEROTYPIC_CELL_CELL_ADHESION | 22 | -0.584665 | -1.386544 | 0.06152 | 0.123151 |
| GOBP_STRIATED_MUSCLE_CELL_PROLIFERATION | 60 | -0.51215 | -1.38631 | 0.034745 | 0.123301 |
| GOBP_NEGATIVE_REGULATION_OF_ENDOTHELIAL_CELL_APOPTOTIC_PROCE | 26 | -0.571556 | -1.385674 | 0.054588 | 0.123931 |
| GOBP_OLFACTORY_LOBE_DEVELOPMENT | 26 | -0.555114 | -1.385335 | 0.06228 | 0.124224 |
| GOBP_CEREBRAL_CORTEX_CELL_MIGRATION | 38 | -0.536034 | -1.38503 | 0.047458 | 0.12448 |
| GOBP_NEURON_APOPTOTIC_PROCESS | 216 | -0.471076 | -1.384582 | 0 | 0.124886 |
| GOBP_CELLULAR_IRON_ION_HOMEOSTASIS | 64 | -0.51123 | -1.384418 | 0.030884 | 0.124967 |
| GOBP_SPROUTING_ANGIOGENESIS | 120 | -0.484866 | -1.384338 | 0.015322 | 0.124952 |
| GOBP_NEUROTRANSMITTER_BIOSYNTHETIC_PROCESS | 15 | -0.633019 | -1.384136 | 0.069231 | 0.125067 |
| GOBP_REGULATION_OF_ADENYLATE_CYCLASE_ACTIVITY | 29 | -0.560213 | -1.383967 | 0.05694 | 0.125154 |
| GOBP_ORGANIC_HYDROXY_COMPOUND_CATABOLIC_PROCESS | 65 | -0.504208 | -1.383804 | 0.03095 | 0.125223 |
| GOBP_VITAMIN_TRANSPORT | 35 | -0.550573 | -1.383601 | 0.044218 | 0.125356 |
| GOBP_POSITIVE_REGULATION_OF_DEPHOSPHORYLATION | 57 | -0.510901 | -1.383156 | 0.037675 | 0.125778 |
| GOBP_POSITIVE_REGULATION_OF_NEUROGENESIS | 209 | -0.472602 | -1.383111 | 0.00501 | 0.125717 |
| GOBP_OSTEOBLAST_DIFFERENTIATION | 205 | -0.47005 | -1.38309 | 0 | 0.125632 |
| GOBP_OLIGODENDROCYTE_DEVELOPMENT | 40 | -0.536672 | -1.382242 | 0.047566 | 0.126546 |
| GOBP_CATECHOL_CONTAINING_COMPOUND_METABOLIC_PROCESS | 46 | -0.525558 | -1.38207 | 0.049834 | 0.126643 |
| GOBP_NEGATIVE_REGULATION_OF_LOCOMOTION | 297 | -0.469176 | -1.381726 | 0 | 0.126962 |
| GOBP_MEMBRANE_FUSION | 159 | -0.473118 | -1.381616 | 0.004016 | 0.126966 |
| GOBP_POSITIVE_REGULATION_OF_POTASSIUM_ION_TRANSMEMBRANE_TRAN | 23 | -0.576519 | -1.380866 | 0.063625 | 0.127753 |
| GOBP_CELL_JUNCTION_DISASSEMBLY | 20 | -0.592004 | -1.380727 | 0.066667 | 0.127803 |
| GOBP_POSITIVE_REGULATION_OF_OSTEOCLAST_DIFFERENTIATION | 21 | -0.591935 | -1.380438 | 0.072202 | 0.128048 |
| GOBP_POSITIVE_REGULATION_OF_CELL_GROWTH | 147 | -0.477264 | -1.3802 | 0.010081 | 0.128244 |
| GOBP_EMBRYONIC_ORGAN_DEVELOPMENT | 379 | -0.466999 | -1.379822 | 0 | 0.128581 |
| GOBP_CELLULAR_SODIUM_ION_HOMEOSTASIS | 17 | -0.61348 | -1.379305 | 0.08 | 0.129095 |
| GOBP_RESPIRATORY_SYSTEM_DEVELOPMENT | 181 | -0.47819 | -1.379274 | 0.00501 | 0.129016 |
| GOBP_REGULATION_OF_NEURAL_PRECURSOR_CELL_PROLIFERATION | 78 | -0.493175 | -1.379201 | 0.021119 | 0.128991 |
| GOBP_LYMPH_VESSEL_DEVELOPMENT | 26 | -0.563198 | -1.379095 | 0.0732 | 0.129027 |
| GOBP_REGULATION_OF_NOTCH_SIGNALING_PATHWAY | 101 | -0.48535 | -1.37874 | 0.018519 | 0.129327 |
| GOBP_NEGATIVE_REGULATION_OF_BLOOD_CIRCULATION | 19 | -0.590407 | -1.378157 | 0.085678 | 0.129882 |
| GOBP_NEGATIVE_REGULATION_OF_PROTEIN_MATURATION | 24 | -0.574412 | -1.377657 | 0.059952 | 0.130349 |
| GOBP_MYELOID_CELL_DEVELOPMENT | 65 | -0.506574 | -1.377217 | 0.037935 | 0.130801 |
| GOBP_DORSAL_VENTRAL_PATTERN_FORMATION | 69 | -0.494527 | -1.37715 | 0.039112 | 0.130778 |
| GOBP_POSITIVE_REGULATION_OF_WOUND_HEALING | 51 | -0.520681 | -1.377112 | 0.042484 | 0.130714 |
| GOBP_PHASIC_SMOOTH_MUSCLE_CONTRACTION | 18 | -0.593924 | -1.376812 | 0.08761 | 0.130968 |
| GOBP_CHONDROITIN_SULFATE_BIOSYNTHETIC_PROCESS | 25 | -0.576096 | -1.376724 | 0.085337 | 0.130964 |
| GOBP_POSITIVE_REGULATION_OF_CYTOKINE_PRODUCTION_INVOLVED_IN_IN | 18 | -0.601595 | -1.376646 | 0.080503 | 0.13095 |
| GOBP_CARDIAC_MUSCLE_CELL_PROLIFERATION | 47 | -0.51717 | -1.37644 | 0.051819 | 0.131092 |
| GOBP_TERPENOID_BIOSYNTHETIC_PROCESS | 15 | -0.629895 | -1.376242 | 0.067775 | 0.131199 |
| GOBP_VISUAL_BEHAVIOR | 44 | -0.51905 | -1.376069 | 0.057332 | 0.131298 |
| GOBP_PHENOL_CONTAINING_COMPOUND_BIOSYNTHETIC_PROCESS | 40 | -0.532445 | -1.375996 | 0.064994 | 0.131268 |
| GOBP_CELLULAR_PROTEIN_CONTAINING_COMPLEX_LOCALIZATION | 20 | -0.583584 | -1.375991 | 0.084767 | 0.131165 |
| GOBP_REGULATION_OF_VOLTAGE_GATED_CALCIUM_CHANNEL_ACTIVITY | 33 | -0.545241 | -1.375249 | 0.071672 | 0.13195 |
| GOBP_REGULATION_OF_ACTIN_FILAMENT_BASED_MOVEMENT | 36 | -0.531155 | -1.37495 | 0.058219 | 0.132199 |
| GOBP_TOLL_LIKE_RECEPTOR_4_SIGNALING_PATHWAY | 38 | -0.539882 | -1.374692 | 0.049539 | 0.132403 |
| GOBP_CELL_JUNCTION_ASSEMBLY | 403 | -0.464252 | -1.374637 | 0 | 0.132353 |
| GOBP_RESPONSE_TO_INTERLEUKIN_1 | 190 | -0.472779 | -1.374575 | 0.003021 | 0.132304 |
| GOBP_ROOF_OF_MOUTH_DEVELOPMENT | 85 | -0.493157 | -1.374561 | 0.022587 | 0.132205 |
| GOBP_POSITIVE_REGULATION_OF_MYELINATION | 16 | -0.61201 | -1.374548 | 0.094268 | 0.132109 |
| GOBP_REGULATION_OF_CARTILAGE_DEVELOPMENT | 61 | -0.510494 | -1.373582 | 0.043526 | 0.1332 |
| GOBP_T_HELPER_2_CELL_DIFFERENTIATION | 15 | -0.621718 | -1.373368 | 0.074349 | 0.133389 |
| GOBP_SMAD_PROTEIN_SIGNAL_TRANSDUCTION | 70 | -0.499536 | -1.371348 | 0.036765 | 0.135764 |
| GOBP_NEGATIVE_REGULATION_OF_VASCULATURE_DEVELOPMENT | 95 | -0.486077 | -1.371019 | 0.015448 | 0.136041 |
| GOBP_POSITIVE_REGULATION_OF_COLD_INDUCED_THERMOGENESIS | 91 | -0.490398 | -1.371019 | 0.023663 | 0.135928 |
| GOBP_POSITIVE_REGULATION_OF_LIPID_BIOSYNTHETIC_PROCESS | 72 | -0.499425 | -1.370618 | 0.029258 | 0.136301 |
| GOBP_RESPONSE_TO_ALCOHOL | 209 | -0.467298 | -1.369719 | 0.002004 | 0.13733 |
| GOBP_REGULATION_OF_EXTRINSIC_APOPTOTIC_SIGNALING_PATHWAY_VIA_D | 54 | -0.505467 | -1.369338 | 0.054585 | 0.137664 |
| GOBP_MESODERM_DEVELOPMENT | 108 | -0.484428 | -1.368938 | 0.01127 | 0.138022 |
| GOBP_REGULATION_OF_PH | 99 | -0.484305 | -1.368822 | 0.024666 | 0.138058 |
| GOBP_OSTEOBLAST_PROLIFERATION | 30 | -0.547568 | -1.367822 | 0.063879 | 0.139221 |
| GOBP_POSITIVE_REGULATION_OF_MULTI_ORGANISM_PROCESS | 17 | -0.606076 | -1.367707 | 0.099747 | 0.139274 |
| GOBP_POSITIVE_REGULATION_OF_STRESS_ACTIVATED_PROTEIN_KINASE_SI | 110 | -0.48228 | -1.367294 | 0.017294 | 0.139659 |
| GOBP_NEGATIVE_REGULATION_OF_ION_TRANSPORT | 268 | -0.465981 | -1.367212 | 0.001 | 0.139633 |
| GOBP_NEURON_DEATH | 322 | -0.46533 | -1.367009 | 0 | 0.139791 |
| GOBP_REGULATION_OF_ERYTHROCYTE_DIFFERENTIATION | 45 | -0.518308 | -1.366961 | 0.072527 | 0.139732 |
| GOBP_REGULATION_OF_ODONTOGENESIS | 15 | -0.623122 | -1.366851 | 0.096899 | 0.139759 |
| GOBP_AMYLOID_BETA_METABOLIC_PROCESS | 55 | -0.502454 | -1.366495 | 0.052859 | 0.140077 |
| GOBP_CELLULAR_RESPONSE_TO_ZINC_ION | 20 | -0.585001 | -1.366156 | 0.087282 | 0.140386 |
| GOBP_NEGATIVE_REGULATION_OF_AMINE_TRANSPORT | 22 | -0.575269 | -1.366005 | 0.093525 | 0.140461 |
| GOBP_REGULATION_OF_MEMBRANE_PROTEIN_ECTODOMAIN_PROTEOLYSIS | 23 | -0.584396 | -1.365113 | 0.081535 | 0.141422 |

| | | | | | |
|---|---|---|---|---|---|
| GOBP_CEREBRAL_CORTEX_DEVELOPMENT | 101 | -0.483785 | -1.365083 | 0.020492 | 0.141339 |
| GOBP_OSSIFICATION | 369 | -0.4602 | -1.364447 | 0 | 0.141999 |
| GOBP_TOLL_LIKE_RECEPTOR_3_SIGNALING_PATHWAY | 19 | -0.582403 | -1.363963 | 0.092683 | 0.142524 |
| GOBP_HINDBRAIN_MORPHOGENESIS | 40 | -0.530707 | -1.363863 | 0.061179 | 0.142522 |
| GOBP_DEVELOPMENTAL_GROWTH_INVOLVED_IN_MORPHOGENESIS | 216 | -0.467429 | -1.363334 | 0.006006 | 0.143052 |
| GOBP_MUCOPOLYSACCHARIDE_METABOLIC_PROCESS | 111 | -0.47668 | -1.363291 | 0.023517 | 0.142988 |
| GOBP_MUSCLE_CONTRACTION | 315 | -0.461107 | -1.363277 | 0 | 0.142891 |
| GOBP_PERIPHERAL_NERVOUS_SYSTEM_DEVELOPMENT | 71 | -0.494168 | -1.363261 | 0.033368 | 0.142796 |
| GOBP_POSITIVE_REGULATION_OF_TYROSINE_PHOSPHORYLATION_OF_STAT_ | 53 | -0.508967 | -1.362949 | 0.045113 | 0.14309 |
| GOBP_HEMATOPOIETIC_STEM_CELL_PROLIFERATION | 23 | -0.562236 | -1.362674 | 0.09816 | 0.143305 |
| GOBP_RESPONSE_TO_MINERALOCORTICOID | 27 | -0.56418 | -1.362595 | 0.07783 | 0.143277 |
| GOBP_REGULATION_OF_ANIMAL_ORGAN_FORMATION | 24 | -0.573806 | -1.362535 | 0.07497 | 0.143242 |
| GOBP_MUSCLE_CELL_MIGRATION | 78 | -0.491046 | -1.362356 | 0.028243 | 0.143359 |
| GOBP_SOMITE_DEVELOPMENT | 75 | -0.491977 | -1.361944 | 0.045121 | 0.14376 |
| GOBP_CELLULAR_RESPONSE_TO_CADMIUM_ION | 34 | -0.537311 | -1.361846 | 0.073563 | 0.143755 |
| GOBP_RESPONSE_TO_MECHANICAL_STIMULUS | 192 | -0.4627 | -1.361615 | 0.004028 | 0.14394 |
| GOBP_REGULATION_OF_CHONDROCYTE_DIFFERENTIATION | 47 | -0.509631 | -1.361387 | 0.051254 | 0.144102 |
| GOBP_POSITIVE_REGULATION_OF_ORGAN_GROWTH | 44 | -0.520122 | -1.361014 | 0.072464 | 0.144514 |
| GOBP_REGULATION_OF_MEMBRANE_REPOLARIZATION | 30 | -0.553041 | -1.360791 | 0.075669 | 0.144666 |
| GOBP_NEURON_PROJECTION_EXTENSION | 159 | -0.469134 | -1.360361 | 0.010081 | 0.145096 |
| GOBP_DIGESTIVE_TRACT_MORPHOGENESIS | 38 | -0.524398 | -1.360328 | 0.084459 | 0.145031 |
| GOBP_T_HELPER_17_TYPE_IMMUNE_RESPONSE | 28 | -0.560232 | -1.360237 | 0.068287 | 0.145037 |
| GOBP_MAINTENANCE_OF_LOCATION | 299 | -0.461076 | -1.360143 | 0 | 0.145041 |
| GOBP_PRODUCTION_OF_MOLECULAR_MEDIATOR_INVOLVED_IN_INFLAMMATO | 66 | -0.500528 | -1.360038 | 0.040426 | 0.145065 |
| GOBP_POSITIVE_REGULATION_OF_MYELOID_LEUKOCYTE_CYTOKINE_PRODU | 16 | -0.604299 | -1.359558 | 0.086634 | 0.145571 |
| GOBP_REGULATION_OF_RHO_PROTEIN_SIGNAL_TRANSDUCTION | 81 | -0.482496 | -1.35949 | 0.043979 | 0.145539 |
| GOBP_POSITIVE_REGULATION_OF_SMALL_MOLECULE_METABOLIC_PROCESS | 122 | -0.475715 | -1.359292 | 0.023469 | 0.145678 |
| GOBP_PYRIMIDINE_NUCLEOTIDE_CATABOLIC_PROCESS | 19 | -0.5742 | -1.359007 | 0.106796 | 0.145938 |
| GOBP_AMYLOID_BETA_CLEARANCE | 35 | -0.530922 | -1.358587 | 0.07564 | 0.146366 |
| GOBP_NEGATIVE_REGULATION_OF_ACTIN_FILAMENT_BUNDLE_ASSEMBLY | 30 | -0.548062 | -1.358424 | 0.077907 | 0.146478 |
| GOBP_NEGATIVE_REGULATION_OF_SMOOTH_MUSCLE_CELL_PROLIFERATION | 41 | -0.517249 | -1.357547 | 0.078022 | 0.14752 |
| GOBP_REGULATION_OF_RAS_PROTEIN_SIGNAL_TRANSDUCTION | 186 | -0.461983 | -1.357268 | 0.005025 | 0.147773 |
| GOBP_NEGATIVE_REGULATION_OF_EPITHELIAL_CELL_APOPTOTIC_PROCESS | 39 | -0.528272 | -1.356886 | 0.066667 | 0.148155 |
| GOBP_ACTIVATION_OF_PROTEIN_KINASE_A_ACTIVITY | 17 | -0.609111 | -1.356699 | 0.084263 | 0.148254 |
| GOBP_KERATAN_SULFATE_BIOSYNTHETIC_PROCESS | 27 | -0.56097 | -1.356602 | 0.091232 | 0.148283 |
| GOBP_RESPONSE_TO_CALCIUM_ION | 135 | -0.476964 | -1.356437 | 0.015167 | 0.148391 |
| GOBP_REGULATION_OF_EMBRYONIC_DEVELOPMENT | 59 | -0.49738 | -1.356362 | 0.057797 | 0.14837 |
| GOBP_POSITIVE_REGULATION_OF_GROWTH | 225 | -0.462978 | -1.356189 | 0.005015 | 0.14848 |
| GOBP_POSITIVE_REGULATION_OF_EPITHELIAL_CELL_APOPTOTIC_PROCESS | 32 | -0.537662 | -1.355798 | 0.077636 | 0.148867 |
| GOBP_ENDOCHONDRAL_BONE_MORPHOGENESIS | 51 | -0.508808 | -1.355282 | 0.05646 | 0.149462 |
| GOBP_REGULATION_OF_T_HELPER_17_TYPE_IMMUNE_RESPONSE | 17 | -0.599937 | -1.354809 | 0.107053 | 0.149978 |
| GOBP_POSITIVE_REGULATION_OF_PEPTIDYL_SERINE_PHOSPHORYLATION | 85 | -0.481429 | -1.354592 | 0.038502 | 0.150155 |
| GOBP_REGULATION_OF_AMYLOID_PRECURSOR_PROTEIN_CATABOLIC_PROC | 42 | -0.521325 | -1.354569 | 0.069274 | 0.150059 |
| GOBP_ADENYLATE_CYCLASE_ACTIVATING_ADRENERGIC_RECEPTOR_SIGNAL | 21 | -0.580832 | -1.354191 | 0.086852 | 0.150416 |
| GOBP_NEGATIVE_REGULATION_OF_RECEPTOR_SIGNALING_PATHWAY_VIA_S | 24 | -0.560864 | -1.353711 | 0.098592 | 0.150921 |
| GOBP_MUSCLE_ORGAN_MORPHOGENESIS | 65 | -0.49645 | -1.35229 | 0.039488 | 0.152684 |
| GOBP_BRANCH_ELONGATION_OF_AN_EPITHELIUM | 15 | -0.615961 | -1.352217 | 0.080357 | 0.152659 |
| GOBP_POSITIVE_REGULATION_OF_VASCULAR_ENDOTHELIAL_CELL_PROLIFE | 15 | -0.614295 | -1.351612 | 0.098972 | 0.15334 |
| GOBP_MONOCYTE_DIFFERENTIATION | 31 | -0.541825 | -1.350831 | 0.06582 | 0.154273 |
| GOBP_NUCLEOBASE_METABOLIC_PROCESS | 33 | -0.537715 | -1.350685 | 0.075231 | 0.154333 |
| GOBP_COLUMNAR_CUBOIDAL_EPITHELIAL_CELL_DEVELOPMENT | 36 | -0.524332 | -1.350668 | 0.083429 | 0.154238 |
| GOBP_DEFINITIVE_HEMOPOIESIS | 19 | -0.578575 | -1.349962 | 0.108747 | 0.155066 |
| GOBP_CELLULAR_RESPONSE_TO_LIPID | 497 | -0.453959 | -1.349934 | 0 | 0.15498 |
| GOBP_SIGNALING_RECEPTOR_LIGAND_PRECURSOR_PROCESSING | 27 | -0.552999 | -1.349619 | 0.085984 | 0.155276 |
| GOBP_NEGATIVE_REGULATION_OF_ALPHA_BETA_T_CELL_DIFFERENTIATION | 25 | -0.559828 | -1.34955 | 0.086085 | 0.155251 |
| GOBP_EXTRACELLULAR_MATRIX_DISASSEMBLY | 69 | -0.493405 | -1.349494 | 0.050538 | 0.155202 |
| GOBP_CEREBRAL_CORTEX_RADIALLY_ORIENTED_CELL_MIGRATION | 29 | -0.545861 | -1.349432 | 0.096552 | 0.155161 |
| GOBP_CARDIAC_MUSCLE_CELL_DIFFERENTIATION | 91 | -0.475565 | -1.349108 | 0.046392 | 0.15551 |
| GOBP_ACTIVATION_OF_CYSTEINE_TYPE_ENDOPEPTIDASE_ACTIVITY_INVOLVE | 86 | -0.48219 | -1.348204 | 0.039501 | 0.156588 |
| GOBP_MEGAKARYOCYTE_DEVELOPMENT | 16 | -0.599764 | -1.34803 | 0.110971 | 0.156677 |
| GOBP_ARTERY_DEVELOPMENT | 86 | -0.478867 | -1.347975 | 0.036496 | 0.156629 |
| GOBP_DOPAMINE_METABOLIC_PROCESS | 33 | -0.529645 | -1.347462 | 0.075028 | 0.157161 |
| GOBP_PROTEIN_LOCALIZATION_TO_VACUOLE | 70 | -0.489924 | -1.347199 | 0.058201 | 0.157402 |
| GOBP_CELL_SUBSTRATE_ADHESION | 339 | -0.457635 | -1.347117 | 0.001 | 0.157386 |
| GOBP_REGULATION_OF_POTASSIUM_ION_TRANSPORT | 89 | -0.484106 | -1.346994 | 0.026123 | 0.157423 |
| GOBP_ESTABLISHMENT_OF_MITOCHONDRION_LOCALIZATION | 25 | -0.555153 | -1.34621 | 0.09466 | 0.158417 |
| GOBP_NEURON_CELLULAR_HOMEOSTASIS | 32 | -0.535062 | -1.346176 | 0.074408 | 0.158336 |
| GOBP_NEURON_PROJECTION_ARBORIZATION | 28 | -0.551989 | -1.345982 | 0.087861 | 0.15848 |
| GOBP_POSITIVE_REGULATION_OF_VASCULAR_ENDOTHELIAL_GROWTH_FACT | 27 | -0.553415 | -1.345712 | 0.075029 | 0.158713 |
| GOBP_RESPONSE_TO_CATECHOLAMINE | 85 | -0.480971 | -1.345564 | 0.037229 | 0.158767 |
| GOBP_ADRENERGIC_RECEPTOR_SIGNALING_PATHWAY | 25 | -0.561801 | -1.344917 | 0.099648 | 0.159536 |
| GOBP_INSULIN_SECRETION | 192 | -0.463167 | -1.344867 | 0.007007 | 0.159471 |
| GOBP_REGULATION_OF_NUCLEOTIDE_BIOSYNTHETIC_PROCESS | 39 | -0.52132 | -1.344594 | 0.085682 | 0.159738 |
| GOBP_POSITIVE_REGULATION_OF_I_KAPPAB_KINASE_NF_KAPPAB_SIGNALING | 178 | -0.464037 | -1.344459 | 0.006018 | 0.15982 |
| GOBP_HORMONE_METABOLIC_PROCESS | 182 | -0.462587 | -1.344263 | 0.008048 | 0.159961 |
| GOBP_FIBROBLAST_GROWTH_FACTOR_RECEPTOR_SIGNALING_PATHWAY | 97 | -0.476847 | -1.343333 | 0.027778 | 0.161189 |
| GOBP_REGULATION_OF_CELL_DEVELOPMENT | 452 | -0.453803 | -1.343185 | 0 | 0.161266 |
| GOBP_ACTIN_FILAMENT_BASED_MOVEMENT | 141 | -0.468182 | -1.342804 | 0.017189 | 0.161653 |
| GOBP_RESPONSE_TO_DRUG | 327 | -0.456273 | -1.342453 | 0.001001 | 0.161996 |
| GOBP_ISOPRENOID_BIOSYNTHETIC_PROCESS | 28 | -0.546778 | -1.34189 | 0.088339 | 0.162644 |
| GOBP_POSITIVE_REGULATION_OF_NATURAL_KILLER_CELL_ACTIVATION | 17 | -0.595669 | -1.341825 | 0.118321 | 0.162611 |
| GOBP_DETECTION_OF_MECHANICAL_STIMULUS_INVOLVED_IN_SENSORY_PE | 29 | -0.54074 | -1.341706 | 0.101636 | 0.162646 |
| GOBP_NEGATIVE_REGULATION_OF_DEPHOSPHORYLATION | 103 | -0.473027 | -1.341567 | 0.034765 | 0.162729 |
| GOBP_MESODERMAL_CELL_FATE_COMMITMENT | 15 | -0.612473 | -1.340814 | 0.108696 | 0.163647 |
| GOBP_NEGATIVE_REGULATION_OF_EXTRINSIC_APOPTOTIC_SIGNALING_PATH | 94 | -0.477324 | -1.339827 | 0.038855 | 0.164894 |
| GOBP_LENS_DEVELOPMENT_IN_CAMERA_TYPE_EYE | 70 | -0.487998 | -1.339655 | 0.059811 | 0.16501 |
| GOBP_NEGATIVE_REGULATION_OF_INTERLEUKIN_10_PRODUCTION | 16 | -0.608305 | -1.339431 | 0.096203 | 0.165236 |
| GOBP_PROSTATE_GLAND_MORPHOGENESIS | 25 | -0.552396 | -1.339149 | 0.098341 | 0.16549 |
| GOBP_MUSCLE_SYSTEM_PROCESS | 395 | -0.451065 | -1.33914 | 0.003 | 0.165372 |
| GOBP_REGULATION_OF_PRESYNAPTIC_CYTOSOLIC_CALCIUM_ION_CONCENT | 16 | -0.601726 | -1.338938 | 0.105732 | 0.165514 |
| GOBP_REGULATION_OF_BEHAVIOR | 51 | -0.501108 | -1.337865 | 0.065591 | 0.166885 |
| GOBP_POSITIVE_REGULATION_OF_ENDOCYTOSIS | 94 | -0.476751 | -1.337687 | 0.046249 | 0.167035 |
| GOBP_CHONDROCYTE_PROLIFERATION | 19 | -0.589754 | -1.337514 | 0.10533 | 0.167145 |
| GOBP_REGULATION_OF_ENDOTHELIAL_CELL_DEVELOPMENT | 17 | -0.585963 | -1.337069 | 0.104459 | 0.167659 |
| GOBP_POSITIVE_REGULATION_OF_SPROUTING_ANGIOGENESIS | 21 | -0.57417 | -1.336883 | 0.102781 | 0.167801 |
| GOBP_NEURAL_CREST_CELL_MIGRATION | 52 | -0.503235 | -1.336793 | 0.065957 | 0.167787 |
| GOBP_POSITIVE_REGULATION_OF_FILOPODIUM_ASSEMBLY | 30 | -0.539598 | -1.336121 | 0.099885 | 0.168648 |
| GOBP_NEGATIVE_REGULATION_OF_INTERLEUKIN_8_PRODUCTION | 20 | -0.580471 | -1.335777 | 0.10567 | 0.169009 |
| GOBP_STRIATED_MUSCLE_CONTRACTION | 161 | -0.459835 | -1.335569 | 0.013131 | 0.169143 |
| GOBP_NECROTIC_CELL_DEATH | 57 | -0.493501 | -1.335183 | 0.066381 | 0.16959 |
| GOBP_REPRODUCTIVE_BEHAVIOR | 27 | -0.546132 | -1.334968 | 0.105758 | 0.169776 |
| GOBP_POSITIVE_REGULATION_OF_PHOSPHOPROTEIN_PHOSPHATASE_ACTIVI | 20 | -0.584802 | -1.334917 | 0.100245 | 0.169732 |
| GOBP_EPITHELIAL_CELL_APOPTOTIC_PROCESS | 97 | -0.475731 | -1.334715 | 0.034161 | 0.169907 |
| GOBP_CELLULAR_RESPONSE_TO_ALKALOID | 29 | -0.542632 | -1.334696 | 0.081657 | 0.169806 |
| GOBP_REGULATION_OF_RESPONSE_TO_WOUNDING | 141 | -0.465955 | -1.334649 | 0.012183 | 0.169743 |
| GOBP_INNER_EAR_RECEPTOR_CELL_DEVELOPMENT | 38 | -0.519815 | -1.334459 | 0.0823 | 0.169903 |
| GOBP_REGULATION_OF_ENDOTHELIAL_CELL_DIFFERENTIATION | 33 | -0.525218 | -1.334081 | 0.090058 | 0.170311 |
| GOBP_PHENOL_CONTAINING_COMPOUND_METABOLIC_PROCESS | 96 | -0.471978 | -1.334074 | 0.041237 | 0.17019 |
| GOBP_REGULATION_OF_PEPTIDYL_SERINE_PHOSPHORYLATION | 118 | -0.466416 | -1.333476 | 0.034979 | 0.170927 |
| GOBP_EPITHELIAL_TO_MESENCHYMAL_TRANSITION_INVOLVED_IN_ENDOCAR | 16 | -0.59134 | -1.333305 | 0.120668 | 0.171029 |
| GOBP_REGULATION_OF_SUPEROXIDE_METABOLIC_PROCESS | 28 | -0.536958 | -1.332791 | 0.088235 | 0.171629 |
| GOBP_POSITIVE_REGULATION_OF_POTASSIUM_ION_TRANSMEMBRANE_TRAN | 36 | -0.525832 | -1.33274 | 0.073363 | 0.171565 |
| GOBP_ANTEROGRADE_AXONAL_TRANSPORT | 47 | -0.504173 | -1.332572 | 0.075991 | 0.171664 |
| GOBP_NEUROTRANSMITTER_METABOLIC_PROCESS | 26 | -0.550328 | -1.332331 | 0.090047 | 0.171887 |
| GOBP_REGULATION_OF_CELL_FATE_COMMITMENT | 26 | -0.544353 | -1.331966 | 0.10084 | 0.172279 |
| GOBP_REGULATION_OF_PHOSPHATASE_ACTIVITY | 167 | -0.45902 | -1.331416 | 0.009045 | 0.173002 |
| GOBP_POSITIVE_REGULATION_OF_BLOOD_PRESSURE | 29 | -0.540814 | -1.329582 | 0.092527 | 0.17544 |
| GOBP_REGULATION_OF_POSTSYNAPSE_ORGANIZATION | 84 | -0.480059 | -1.329539 | 0.048958 | 0.175369 |
| GOBP_POSITIVE_REGULATION_OF_DNA_BINDING | 53 | -0.492074 | -1.329392 | 0.078891 | 0.175454 |
| GOBP_GLYCOPROTEIN_CATABOLIC_PROCESS | 24 | -0.559438 | -1.329354 | 0.103912 | 0.175379 |
| GOBP_RESPONSE_TO_CORTICOSTEROID | 137 | -0.463504 | -1.329291 | 0.020202 | 0.175346 |
| GOBP_POSITIVE_REGULATION_OF_NIK_NF_KAPPAB_SIGNALING | 63 | -0.491013 | -1.329285 | 0.055135 | 0.175221 |
| GOBP_RESPONSE_TO_RADIATION | 429 | -0.449592 | -1.329103 | 0.001 | 0.175373 |
| GOBP_POSITIVE_REGULATION_OF_PEPTIDASE_ACTIVITY | 188 | -0.459269 | -1.328689 | 0.013092 | 0.175832 |
| GOBP_POSITIVE_REGULATION_OF_ERBB_SIGNALING_PATHWAY | 37 | -0.521563 | -1.328634 | 0.083908 | 0.175778 |
| GOBP_CELLULAR_RESPONSE_TO_VASCULAR_ENDOTHELIAL_GROWTH_FACT | 58 | -0.488786 | -1.328264 | 0.07105 | 0.176187 |
| GOBP_ANTIGEN_PROCESSING_AND_PRESENTATION_OF_PEPTIDE_ANTIGEN_ | 98 | -0.467823 | -1.32825 | 0.037949 | 0.176075 |
| GOBP_POSITIVE_REGULATION_OF_PEPTIDYL_THREONINE_PHOSPHORYLATIO | 27 | -0.536905 | -1.328134 | 0.105018 | 0.176092 |
| GOBP_SMOOTH_MUSCLE_CELL_MIGRATION | 65 | -0.491048 | -1.327355 | 0.068349 | 0.1771 |
| GOBP_SYNCYTIUM_FORMATION | 48 | -0.502128 | -1.327182 | 0.058178 | 0.177231 |
| GOBP_ENDOCRINE_PROCESS | 71 | -0.48514 | -1.327128 | 0.056842 | 0.177191 |
| GOBP_NEURAL_PRECURSOR_CELL_PROLIFERATION | 132 | -0.460477 | -1.326585 | 0.020284 | 0.177867 |
| GOBP_MAINTENANCE_OF_GASTROINTESTINAL_EPITHELIUM | 19 | -0.575537 | -1.326141 | 0.106383 | 0.178362 |
| GOBP_REGULATION_OF_SODIUM_ION_TRANSMEMBRANE_TRANSPORTER_AC | 51 | -0.497562 | -1.325469 | 0.076419 | 0.179181 |
| GOBP_RHO_PROTEIN_SIGNAL_TRANSDUCTION | 129 | -0.462476 | -1.325031 | 0.012109 | 0.179651 |
| GOBP_POSITIVE_REGULATION_OF_VASCULAR_ASSOCIATED_SMOOTH_MUSCI | 33 | -0.527721 | -1.324818 | 0.098083 | 0.17981 |
| GOBP_INTERLEUKIN_13_PRODUCTION | 17 | -0.583087 | -1.323275 | 0.120838 | 0.182008 |
| GOBP_HEART_TRABECULA_MORPHOGENESIS | 27 | -0.540505 | -1.322872 | 0.098131 | 0.182469 |
| GOBP_POSITIVE_REGULATION_OF_ORGANIC_ACID_TRANSPORT | 31 | -0.53564 | -1.32226 | 0.102825 | 0.183249 |
| GOBP_NONRIBOSOMAL_PEPTIDE_BIOSYNTHETIC_PROCESS | 17 | -0.58666 | -1.322099 | 0.121483 | 0.183375 |
| GOBP_NEGATIVE_REGULATION_OF_DNA_BINDING | 52 | -0.499414 | -1.321977 | 0.072131 | 0.18343 |
| GOBP_REGULATION_OF_FATTY_ACID_TRANSPORT | 41 | -0.511011 | -1.321731 | 0.092511 | 0.183653 |
| GOBP_POSITIVE_REGULATION_OF_BLOOD_CIRCULATION | 36 | -0.522212 | -1.321496 | 0.088864 | 0.183876 |
| GOBP_NEGATIVE_REGULATION_OF_CD4_POSITIVE_ALPHA_BETA_T_CELL_DIF | 21 | -0.561861 | -1.321316 | 0.126643 | 0.184013 |
| GOBP_IMMUNE_RESPONSE_TO_TUMOR_CELL | 18 | -0.572164 | -1.321011 | 0.124842 | 0.184351 |
| GOBP_ADAPTIVE_THERMOGENESIS | 147 | -0.458548 | -1.320674 | 0.026342 | 0.184738 |
| GOBP_POSITIVE_REGULATION_OF_HEART_GROWTH | 34 | -0.517305 | -1.320529 | 0.103255 | 0.184826 |
| GOBP_REGULATION_OF_DNA_BINDING | 115 | -0.463388 | -1.320385 | 0.034588 | 0.184921 |
| GOBP_DENDRITIC_SPINE_DEVELOPMENT | 97 | -0.467832 | -1.319842 | 0.044057 | 0.185617 |
| GOBP_ATRIOVENTRICULAR_VALVE_DEVELOPMENT | 25 | -0.543722 | -1.319494 | 0.120425 | 0.18601 |
| GOBP_REGULATION_OF_NEUROBLAST_PROLIFERATION | 28 | -0.52984 | -1.319245 | 0.122664 | 0.186265 |

| | | | | | |
|---|---|---|---|---|---|
| GOBP_NITRIC_OXIDE_MEDIATED_SIGNAL_TRANSDUCTION | 23 | -0.542428 | -1.31924 | 0.110571 | 0.186135 |
| GOBP_CELL_CELL_ADHESION_VIA_PLASMA_MEMBRANE_ADHESION_MOLECU | 258 | -0.449902 | -1.319167 | 0.004004 | 0.186117 |
| GOBP_POSITIVE_REGULATION_OF_PATHWAY_RESTRICTED_SMAD_PROTEIN_ | 44 | -0.502728 | -1.319092 | 0.082599 | 0.186081 |
| GOBP_REGULATION_OF_TRIGLYCERIDE_METABOLIC_PROCESS | 31 | -0.524949 | -1.318971 | 0.103248 | 0.186137 |
| GOBP_REGULATION_OF_MAP_KINASE_ACTIVITY | 293 | -0.444804 | -1.318692 | 0.002 | 0.186402 |
| GOBP_NEGATIVE_REGULATION_OF_GTPASE_ACTIVITY | 44 | -0.50199 | -1.318676 | 0.092018 | 0.186294 |
| GOBP_CGMP_MEDIATED_SIGNALING | 25 | -0.548187 | -1.318446 | 0.11007 | 0.186519 |
| GOBP_IRON_ION_TRANSPORT | 74 | -0.4777 | -1.318178 | 0.057411 | 0.186767 |
| GOBP_POSITIVE_REGULATION_OF_ACTIN_CYTOSKELETON_REORGANIZATION | 17 | -0.585376 | -1.318072 | 0.133683 | 0.186791 |
| GOBP_GLYCOSPHINGOLIPID_METABOLIC_PROCESS | 65 | -0.487514 | -1.318059 | 0.07173 | 0.186671 |
| GOBP_MECHANORECEPTOR_DIFFERENTIATION | 58 | -0.486587 | -1.318026 | 0.083153 | 0.186584 |
| GOBP_NEURONAL_STEM_CELL_POPULATION_MAINTENANCE | 19 | -0.571128 | -1.317689 | 0.127295 | 0.186974 |
| GOBP_REGULATION_OF_AXONOGENESIS | 149 | -0.452999 | -1.317148 | 0.023116 | 0.18768 |
| GOBP_GLIAL_CELL_PROLIFERATION | 44 | -0.499742 | -1.317133 | 0.078977 | 0.187577 |
| GOBP_HOMEOSTASIS_OF_NUMBER_OF_CELLS_WITHIN_A_TISSUE | 24 | -0.548901 | -1.317082 | 0.109049 | 0.187528 |
| GOBP_MITOCHONDRIAL_CALCIUM_ION_TRANSMEMBRANE_TRANSPORT | 25 | -0.547627 | -1.31681 | 0.118149 | 0.187819 |
| GOBP_REGULATION_OF_RESPONSE_TO_EXTRACELLULAR_STIMULUS | 23 | -0.551242 | -1.316794 | 0.115202 | 0.187701 |
| GOBP_PROTEIN_COMPLEX_OLIGOMERIZATION | 218 | -0.452668 | -1.316536 | 0.009018 | 0.187944 |
| GOBP_POSITIVE_REGULATION_OF_MAP_KINASE_ACTIVITY | 223 | -0.453153 | -1.316498 | 0.002004 | 0.187854 |
| GOBP_AORTA_MORPHOGENESIS | 29 | -0.527539 | -1.316102 | 0.112791 | 0.188316 |
| GOBP_REGULATION_OF_MEMBRANE_INVAGINATION | 16 | -0.589215 | -1.316077 | 0.125945 | 0.188213 |
| GOBP_SULFUR_COMPOUND_CATABOLIC_PROCESS | 49 | -0.497398 | -1.315991 | 0.081588 | 0.188205 |
| GOBP_AMYLOID_PRECURSOR_PROTEIN_CATABOLIC_PROCESS | 54 | -0.492243 | -1.315823 | 0.079656 | 0.188306 |
| GOBP_ASPARTATE_FAMILY_AMINO_ACID_CATABOLIC_PROCESS | 23 | -0.562957 | -1.315674 | 0.100719 | 0.188402 |
| GOBP_RESPONSE_TO_METAL_ION | 325 | -0.444655 | -1.315621 | 0.004 | 0.188355 |
| GOBP_PROTEIN_HOMOOLIGOMERIZATION | 167 | -0.452709 | -1.315603 | 0.017172 | 0.188246 |
| GOBP_EXOCYTIC_PROCESS | 76 | -0.46948 | -1.314635 | 0.057351 | 0.189607 |
| GOBP_CEREBELLAR_CORTEX_DEVELOPMENT | 41 | -0.508505 | -1.314436 | 0.098655 | 0.189782 |
| GOBP_CONNECTIVE_TISSUE_DEVELOPMENT | 222 | -0.450069 | -1.313986 | 0.011011 | 0.190322 |
| GOBP_REGULATION_OF_DEVELOPMENTAL_GROWTH | 292 | -0.446621 | -1.313516 | 0.004 | 0.190925 |
| GOBP_RESPIRATORY_GASEOUS_EXCHANGE_BY_RESPIRATORY_SYSTEM | 59 | -0.486254 | -1.313454 | 0.072121 | 0.190886 |
| GOBP_POSITIVE_REGULATION_OF_MUSCLE_HYPERTROPHY | 26 | -0.542327 | -1.313354 | 0.131802 | 0.190906 |
| GOBP_POSITIVE_REGULATION_OF_AMINO_ACID_TRANSPORT | 15 | -0.596658 | -1.313232 | 0.137709 | 0.19096 |
| GOBP_DENDRITIC_SPINE_MAINTENANCE | 18 | -0.581048 | -1.313217 | 0.128269 | 0.19086 |
| GOBP_NEGATIVE_REGULATION_OF_NEURAL_PRECURSOR_CELL_PROLIFERA | 19 | -0.572019 | -1.313049 | 0.131127 | 0.190967 |
| GOBP_POSITIVE_REGULATION_OF_PROTEIN_AUTOPHOSPHORYLATION | 26 | -0.537186 | -1.312977 | 0.10967 | 0.190954 |
| GOBP_POTASSIUM_ION_EXPORT_ACROSS_PLASMA_MEMBRANE | 16 | -0.59074 | -1.312725 | 0.132911 | 0.191204 |
| GOBP_ACTIVATION_OF_MAPKK_ACTIVITY | 50 | -0.489755 | -1.312353 | 0.087757 | 0.191628 |
| GOBP_LIPID_LOCALIZATION | 446 | -0.439826 | -1.312153 | 0.001 | 0.191805 |
| GOBP_AUTONOMIC_NERVOUS_SYSTEM_DEVELOPMENT | 41 | -0.502058 | -1.312131 | 0.094714 | 0.191697 |
| GOBP_CAMP_METABOLIC_PROCESS | 21 | -0.554494 | -1.312055 | 0.121951 | 0.191683 |
| GOBP_CELL_MIGRATION_INVOLVED_IN_SPROUTING_ANGIOGENESIS | 53 | -0.491934 | -1.312035 | 0.083793 | 0.191577 |
| GOBP_REGULATION_OF_HORMONE_LEVELS | 447 | -0.443227 | -1.31202 | 0 | 0.191462 |
| GOBP_CAMP_MEDIATED_SIGNALING | 56 | -0.485932 | -1.311852 | 0.085129 | 0.191604 |
| GOBP_EMBRYONIC_HINDLIMB_MORPHOGENESIS | 26 | -0.534777 | -1.311681 | 0.11032 | 0.191729 |
| GOBP_SEROTONIN_TRANSPORT | 18 | -0.576722 | -1.311233 | 0.130967 | 0.192289 |
| GOBP_POSITIVE_REGULATION_OF_T_HELPER_CELL_DIFFERENTIATION | 19 | -0.576935 | -1.311164 | 0.121437 | 0.19226 |
| GOBP_REGULATION_OF_ANIMAL_ORGAN_MORPHOGENESIS | 170 | -0.451418 | -1.311082 | 0.021169 | 0.192268 |
| GOBP_MICROVILLUS_ORGANIZATION | 24 | -0.541038 | -1.310951 | 0.127098 | 0.192321 |
| GOBP_REGULATION_OF_GASTRULATION | 17 | -0.584779 | -1.310921 | 0.116688 | 0.192233 |
| GOBP_PYRIMIDINE_NUCLEOSIDE_MONOPHOSPHATE_METABOLIC_PROCESS | 18 | -0.57516 | -1.310294 | 0.139057 | 0.193047 |
| GOBP_REGULATION_OF_PHOSPHOLIPID_METABOLIC_PROCESS | 33 | -0.524757 | -1.309933 | 0.105023 | 0.193457 |
| GOBP_POSITIVE_REGULATION_OF_NITRIC_OXIDE_SYNTHASE_BIOSYNTHETIC_ | 15 | -0.603234 | -1.309826 | 0.129199 | 0.193485 |
| GOBP_MODULATION_OF_PROCESS_OF_OTHER_ORGANISM | 106 | -0.461977 | -1.309079 | 0.04111 | 0.194503 |
| GOBP_NIK_NF_KAPPAB_SIGNALING | 164 | -0.449784 | -1.308226 | 0.032193 | 0.19574 |
| GOBP_REACTIVE_NITROGEN_SPECIES_METABOLIC_PROCESS | 68 | -0.473693 | -1.308043 | 0.070231 | 0.195874 |
| GOBP_WOUND_HEALING | 468 | -0.440706 | -1.308003 | 0 | 0.195798 |
| GOBP_LIPID_TRANSLOCATION | 45 | -0.500889 | -1.307966 | 0.104235 | 0.19571 |
| GOBP_I_KAPPAB_KINASE_NF_KAPPAB_SIGNALING | 271 | -0.443656 | -1.307668 | 0.008008 | 0.196042 |
| GOBP_POSITIVE_REGULATION_OF_MITOTIC_NUCLEAR_DIVISION | 38 | -0.511296 | -1.30728 | 0.105499 | 0.196499 |
| GOBP_PEPTIDE_HORMONE_SECRETION | 225 | -0.44512 | -1.307158 | 0.014042 | 0.196542 |
| GOBP_POSITIVE_REGULATION_OF_REACTIVE_OXYGEN_SPECIES_METABOLIC | 90 | -0.468747 | -1.306669 | 0.061983 | 0.197119 |
| GOBP_FIBRINOLYSIS | 17 | -0.57127 | -1.305972 | 0.144291 | 0.198097 |
| GOBP_CELLULAR_RESPONSE_TO_COPPER_ION | 23 | -0.537352 | -1.305553 | 0.132775 | 0.198629 |
| GOBP_MESENCHYMAL_CELL_DIFFERENTIATION | 217 | -0.444334 | -1.305465 | 0.012012 | 0.198626 |
| GOBP_PH_REDUCTION | 56 | -0.485079 | -1.305346 | 0.087889 | 0.198679 |
| GOBP_ESTABLISHMENT_OF_PROTEIN_LOCALIZATION_TO_VACUOLE | 52 | -0.489278 | -1.304862 | 0.078919 | 0.199312 |
| GOBP_POSITIVE_REGULATION_OF_CELL_PROJECTION_ORGANIZATION | 331 | -0.444541 | -1.304763 | 0 | 0.199322 |
| GOBP_RELAXATION_OF_CARDIAC_MUSCLE | 17 | -0.577805 | -1.304663 | 0.145704 | 0.199342 |
| GOBP_LONG_TERM_MEMORY | 32 | -0.522269 | -1.304581 | 0.111239 | 0.199323 |
| GOBP_FEMALE_SEX_DIFFERENTIATION | 98 | -0.462826 | -1.304418 | 0.060575 | 0.199461 |
| GOBP_NEGATIVE_REGULATION_OF_TRANSCRIPTION_REGULATORY_REGION_ | 20 | -0.55912 | -1.304087 | 0.133498 | 0.199834 |
| GOBP_POSITIVE_REGULATION_OF_CELLULAR_EXTRAVASATION | 16 | -0.576396 | -1.303703 | 0.141604 | 0.200293 |
| GOBP_TEMPERATURE_HOMEOSTASIS | 163 | -0.446126 | -1.30366 | 0.022155 | 0.200229 |
| GOBP_PATTERN_RECOGNITION_RECEPTOR_SIGNALING_PATHWAY | 192 | -0.448647 | -1.303591 | 0.018145 | 0.200188 |
| GOBP_RESPONSE_TO_INTERLEUKIN_6 | 39 | -0.508118 | -1.303241 | 0.110731 | 0.20058 |
| GOBP_CERAMIDE_BIOSYNTHETIC_PROCESS | 66 | -0.472955 | -1.302959 | 0.06424 | 0.200882 |
| GOBP_MODULATION_BY_SYMBIONT_OF_ENTRY_INTO_HOST | 44 | -0.496005 | -1.302618 | 0.097374 | 0.201269 |
| GOBP_NEGATIVE_REGULATION_OF_HORMONE_SECRETION | 54 | -0.486297 | -1.302576 | 0.104925 | 0.201206 |
| GOBP_CELLULAR_RESPONSE_TO_ALCOHOL | 80 | -0.470724 | -1.302082 | 0.064315 | 0.201864 |
| GOBP_LYMPHANGIOGENESIS | 17 | -0.568012 | -1.301704 | 0.144458 | 0.202352 |
| GOBP_PLATELET_ACTIVATION | 144 | -0.449166 | -1.30124 | 0.031377 | 0.20296 |
| GOBP_CELLULAR_HORMONE_METABOLIC_PROCESS | 110 | -0.458837 | -1.300815 | 0.054583 | 0.203501 |
| GOBP_POSITIVE_REGULATION_OF_JNK_CASCADE | 80 | -0.468005 | -1.300512 | 0.075393 | 0.203854 |
| GOBP_MYOBLAST_DIFFERENTIATION | 74 | -0.468735 | -1.300478 | 0.074658 | 0.203774 |
| GOBP_IRON_ION_TRANSMEMBRANE_TRANSPORT | 19 | -0.561315 | -1.299947 | 0.150193 | 0.204484 |
| GOBP_REGULATION_OF_RESPIRATORY_GASEOUS_EXCHANGE | 17 | -0.574161 | -1.299945 | 0.142506 | 0.204346 |
| GOBP_REGULATION_OF_SYSTEMIC_ARTERIAL_BLOOD_PRESSURE | 77 | -0.464793 | -1.299785 | 0.070466 | 0.204473 |
| GOBP_DEOXYRIBOSE_PHOSPHATE_CATABOLIC_PROCESS | 26 | -0.533804 | -1.299577 | 0.136087 | 0.204672 |
| GOBP_PROSTATE_GLAND_DEVELOPMENT | 38 | -0.505267 | -1.299448 | 0.129213 | 0.204758 |
| GOBP_TYPE_I_INTERFERON_PRODUCTION | 124 | -0.452554 | -1.299375 | 0.033367 | 0.204739 |
| GOBP_HORMONE_TRANSPORT | 284 | -0.441462 | -1.299151 | 0.008016 | 0.204954 |
| GOBP_NEGATIVE_REGULATION_OF_MYELOID_CELL_DIFFERENTIATION | 81 | -0.465753 | -1.298373 | 0.070825 | 0.206078 |
| GOBP_POSITIVE_REGULATION_OF_GTPASE_ACTIVITY | 388 | -0.440091 | -1.298355 | 0.004 | 0.205974 |
| GOBP_SPECIFICATION_OF_ANIMAL_ORGAN_IDENTITY | 32 | -0.513864 | -1.297788 | 0.104265 | 0.206754 |
| GOBP_REGULATION_OF_THE_FORCE_OF_HEART_CONTRACTION | 24 | -0.541968 | -1.297475 | 0.136364 | 0.207103 |
| GOBP_OLIGOSACCHARIDE_METABOLIC_PROCESS | 50 | -0.486526 | -1.297427 | 0.095861 | 0.20703 |
| GOBP_REGULATION_OF_RESPONSE_TO_FOOD | 16 | -0.575494 | -1.297046 | 0.17298 | 0.207476 |
| GOBP_RESPONSE_TO_COLD | 47 | -0.491702 | -1.296931 | 0.103183 | 0.207505 |
| GOBP_REGULATION_OF_MONOOXYGENASE_ACTIVITY | 56 | -0.481105 | -1.296879 | 0.101842 | 0.207452 |
| GOBP_CELLULAR_RESPONSE_TO_REACTIVE_OXYGEN_SPECIES | 147 | -0.448499 | -1.296714 | 0.043522 | 0.207562 |
| GOBP_HEART_FORMATION | 28 | -0.529029 | -1.296691 | 0.110329 | 0.207467 |
| GOBP_PEPTIDYL_TYROSINE_DEPHOSPHORYLATION | 98 | -0.462419 | -1.296287 | 0.052632 | 0.207976 |
| GOBP_FOREBRAIN_NEURON_DEVELOPMENT | 24 | -0.538055 | -1.296169 | 0.144737 | 0.208018 |
| GOBP_ALPHA_AMINO_ACID_CATABOLIC_PROCESS | 83 | -0.465196 | -1.295826 | 0.06911 | 0.208461 |
| GOBP_REGULATION_OF_AMYLOID_BETA_CLEARANCE | 16 | -0.578642 | -1.295777 | 0.148825 | 0.208405 |
| GOBP_REGULATION_OF_WOUND_HEALING | 113 | -0.454518 | -1.295668 | 0.052469 | 0.208427 |
| GOBP_DEVELOPMENTAL_CELL_GROWTH | 207 | -0.444013 | -1.295574 | 0.02004 | 0.208446 |
| GOBP_SENSORY_PERCEPTION_OF_MECHANICAL_STIMULUS | 154 | -0.449888 | -1.295212 | 0.02834 | 0.208907 |
| GOBP_TYROSINE_PHOSPHORYLATION_OF_STAT_PROTEIN | 67 | -0.473598 | -1.295027 | 0.095789 | 0.209077 |
| GOBP_NEGATIVE_REGULATION_OF_ANION_TRANSPORT | 178 | -0.446998 | -1.294914 | 0.011066 | 0.209116 |
| GOBP_RESPONSE_TO_MAGNESIUM_ION | 15 | -0.59234 | -1.294907 | 0.140306 | 0.20899 |
| GOBP_HEART_VALVE_DEVELOPMENT | 60 | -0.476739 | -1.294835 | 0.095494 | 0.208963 |
| GOBP_ORGAN_GROWTH | 142 | -0.449922 | -1.294507 | 0.035354 | 0.209374 |
| GOBP_POSITIVE_REGULATION_OF_EXOCYTOSIS | 78 | -0.466921 | -1.294417 | 0.075472 | 0.209377 |
| GOBP_WOUND_HEALING_SPREADING_OF_EPIDERMAL_CELLS | 18 | -0.563551 | -1.294239 | 0.13017 | 0.20954 |
| GOBP_CELLULAR_RESPONSE_TO_DRUG | 60 | -0.483167 | -1.293659 | 0.079702 | 0.210348 |
| GOBP_REGULATION_OF_GTPASE_ACTIVITY | 462 | -0.433068 | -1.293212 | 0.002 | 0.210929 |
| GOBP_GASTRULATION_WITH_MOUTH_FORMING_SECOND | 22 | -0.55068 | -1.293096 | 0.123786 | 0.210995 |
| GOBP_CELLULAR_RESPONSE_TO_VIRUS | 59 | -0.476202 | -1.293016 | 0.098818 | 0.21098 |
| GOBP_REGULATION_OF_HAIR_FOLLICLE_DEVELOPMENT | 16 | -0.586091 | -1.292831 | 0.144638 | 0.211157 |
| GOBP_REGULATION_OF_SMALL_GTPASE_MEDIATED_SIGNAL_TRANSDUCTION | 313 | -0.436919 | -1.292643 | 0.006006 | 0.211301 |
| GOBP_CELL_MATRIX_ADHESION | 217 | -0.440151 | -1.292501 | 0.022066 | 0.211377 |
| GOBP_TRANSFORMING_GROWTH_FACTOR_BETA_PRODUCTION | 39 | -0.507367 | -1.292498 | 0.114827 | 0.211241 |
| GOBP_LIVER_REGENERATION | 30 | -0.522104 | -1.292295 | 0.11954 | 0.211414 |
| GOBP_MITOCHONDRIAL_GENOME_MAINTENANCE | 21 | -0.55735 | -1.291908 | 0.147059 | 0.211905 |
| GOBP_REGULATION_OF_PROTEIN_LOCALIZATION_TO_CELL_SURFACE | 37 | -0.506406 | -1.291247 | 0.118363 | 0.212781 |
| GOBP_TRANSMEMBRANE_RECEPTOR_PROTEIN_SERINE_THREONINE_KINASE | 319 | -0.436042 | -1.291141 | 0.006 | 0.212815 |
| GOBP_STRIATED_MUSCLE_CELL_APOPTOTIC_PROCESS | 34 | -0.512144 | -1.291064 | 0.127148 | 0.212805 |
| GOBP_NEGATIVE_REGULATION_OF_LYASE_ACTIVITY | 23 | -0.546563 | -1.290996 | 0.14873 | 0.212785 |
| GOBP_DEOXYRIBONUCLEOTIDE_METABOLIC_PROCESS | 39 | -0.494602 | -1.290756 | 0.126096 | 0.213057 |
| GOBP_REGULATION_OF_SMOOTH_MUSCLE_CONTRACTION | 49 | -0.486126 | -1.290634 | 0.109902 | 0.213118 |
| GOBP_CARDIAC_SEPTUM_MORPHOGENESIS | 62 | -0.476085 | -1.290407 | 0.092827 | 0.213335 |
| GOBP_POSITIVE_REGULATION_OF_LONG_TERM_SYNAPTIC_POTENTIATION | 22 | -0.540845 | -1.290206 | 0.150551 | 0.213509 |
| GOBP_VESICLE_DOCKING | 65 | -0.467108 | -1.289469 | 0.088983 | 0.214577 |
| GOBP_RAC_PROTEIN_SIGNAL_TRANSDUCTION | 39 | -0.501425 | -1.28946 | 0.114736 | 0.214454 |
| GOBP_CARDIAC_SEPTUM_DEVELOPMENT | 94 | -0.455845 | -1.289104 | 0.070842 | 0.21491 |
| GOBP_NEGATIVE_REGULATION_OF_POTASSIUM_ION_TRANSPORT | 28 | -0.522911 | -1.289062 | 0.120092 | 0.21484 |
| GOBP_REGULATION_OF_MEMBRANE_LIPID_DISTRIBUTION | 51 | -0.480095 | -1.28905 | 0.103104 | 0.214716 |
| GOBP_NEGATIVE_REGULATION_OF_RECEPTOR_SIGNALING_PATHWAY_VIA_JA | 23 | -0.540713 | -1.288716 | 0.123623 | 0.215108 |
| GOBP_MEMBRANE_BIOGENESIS | 40 | -0.490532 | -1.288435 | 0.119639 | 0.215403 |
| GOBP_CELLULAR_MODIFIED_AMINO_ACID_CATABOLIC_PROCESS | 26 | -0.529351 | -1.288412 | 0.135266 | 0.215302 |
| GOBP_REGULATION_OF_STRESS_ACTIVATED_PROTEIN_KINASE_SIGNALING_( | 171 | -0.444805 | -1.288373 | 0.029116 | 0.215213 |
| GOBP_CYTOKINE_PRODUCTION_INVOLVED_IN_INFLAMMATORY_RESPONSE | 49 | -0.488107 | -1.288038 | 0.103939 | 0.215629 |
| GOBP_VENTRICULAR_CARDIAC_MUSCLE_CELL_ACTION_POTENTIAL | 32 | -0.514958 | -1.287916 | 0.121354 | 0.215657 |
| GOBP_CELLULAR_RESPONSE_TO_OXYGEN_RADICAL | 22 | -0.543887 | -1.287252 | 0.128329 | 0.216595 |
| GOBP_CELLULAR_RESPONSE_TO_GLUCAGON_STIMULUS | 23 | -0.545068 | -1.28691 | 0.154964 | 0.21699 |
| GOBP_NEGATIVE_REGULATION_OF_CALCIUM_ION_TRANSPORT_INTO_CYTOS | 16 | -0.585501 | -1.286857 | 0.146789 | 0.216931 |
| GOBP_EXCITATORY_SYNAPSE_ASSEMBLY | 26 | -0.533276 | -1.286641 | 0.155213 | 0.217148 |
| GOBP_VASCULAR_ENDOTHELIAL_GROWTH_FACTOR_SIGNALING_PATHWAY | 39 | -0.500939 | -1.286586 | 0.118757 | 0.2171 |
| GOBP_POSITIVE_REGULATION_OF_NEURAL_PRECURSOR_CELL_PROLIFERAT | 47 | -0.490226 | -1.28657 | 0.111475 | 0.21698 |

| | | | | | |
|---|---|---|---|---|---|
| GOBP_REGULATION_OF_STEM_CELL_POPULATION_MAINTENANCE | 29 | -0.515611 | -1.285793 | 0.141869 | 0.21815 |
| GOBP_GLUCOSAMINE_CONTAINING_COMPOUND_METABOLIC_PROCESS | 25 | -0.533335 | -1.285565 | 0.13396 | 0.218362 |
| GOBP_PROTEIN_LOCALIZATION_TO_ENDOSOME | 25 | -0.532455 | -1.285508 | 0.130332 | 0.218313 |
| GOBP_NEGATIVE_REGULATION_OF_LIPID_METABOLIC_PROCESS | 85 | -0.468556 | -1.285392 | 0.078594 | 0.218373 |
| GOBP_RESPONSE_TO_PH | 37 | -0.504314 | -1.285287 | 0.130137 | 0.218399 |
| GOBP_REGULATION_OF_MYELOID_CELL_DIFFERENTIATION | 226 | -0.438156 | -1.285203 | 0.021063 | 0.218417 |
| GOBP_CEREBELLAR_CORTEX_MORPHOGENESIS | 31 | -0.513912 | -1.285125 | 0.123414 | 0.218421 |
| GOBP_ENDOTHELIUM_DEVELOPMENT | 124 | -0.445423 | -1.28507 | 0.046653 | 0.218378 |
| GOBP_SOMATIC_STEM_CELL_POPULATION_MAINTENANCE | 62 | -0.471261 | -1.285048 | 0.10432 | 0.218278 |
| GOBP_REGULATION_OF_OSTEOBLAST_PROLIFERATION | 25 | -0.539063 | -1.284668 | 0.129717 | 0.218756 |
| GOBP_ARF_PROTEIN_SIGNAL_TRANSDUCTION | 19 | -0.554885 | -1.284571 | 0.16791 | 0.218786 |
| GOBP_NOSE_DEVELOPMENT | 15 | -0.577699 | -1.284309 | 0.153061 | 0.219067 |
| GOBP_NEGATIVE_REGULATION_OF_T_HELPER_CELL_DIFFERENTIATION | 17 | -0.562956 | -1.284119 | 0.161006 | 0.219218 |
| GOBP_POSITIVE_REGULATION_OF_CELL_DEVELOPMENT | 273 | -0.435923 | -1.283897 | 0.006 | 0.219451 |
| GOBP_REGULATION_OF_CELL_SUBSTRATE_ADHESION | 200 | -0.439197 | -1.283615 | 0.031093 | 0.21979 |
| GOBP_REGULATION_OF_PROTEIN_LOCALIZATION_TO_SYNAPSE | 26 | -0.532914 | -1.2836 | 0.124138 | 0.219673 |
| GOBP_POSITIVE_REGULATION_OF_FOCAL_ADHESION_ASSEMBLY | 24 | -0.531322 | -1.283466 | 0.135167 | 0.219765 |
| GOBP_NEGATIVE_REGULATION_OF_RESPONSE_TO_WOUNDING | 74 | -0.470324 | -1.283405 | 0.082105 | 0.219725 |
| GOBP_EPITHELIAL_TUBE_BRANCHING_INVOLVED_IN_LUNG_MORPHOGENESI | 24 | -0.533148 | -1.282826 | 0.14841 | 0.220528 |
| GOBP_POSITIVE_REGULATION_OF_ANIMAL_ORGAN_MORPHOGENESIS | 29 | -0.519941 | -1.282801 | 0.137324 | 0.220432 |
| GOBP_NEURON_PROJECTION_REGENERATION | 54 | -0.47678 | -1.282739 | 0.116812 | 0.220389 |
| GOBP_RESPONSE_TO_ETHANOL | 112 | -0.458024 | -1.282377 | 0.054934 | 0.220864 |
| GOBP_POSITIVE_REGULATION_OF_CARDIOCYTE_DIFFERENTIATION | 15 | -0.578635 | -1.282282 | 0.145455 | 0.22088 |
| GOBP_PRIMARY_ALCOHOL_METABOLIC_PROCESS | 81 | -0.46114 | -1.282238 | 0.074844 | 0.220821 |
| GOBP_POSITIVE_REGULATION_OF_EPITHELIAL_CELL_DIFFERENTIATION | 48 | -0.484052 | -1.281848 | 0.120482 | 0.22133 |
| GOBP_RESPONSE_TO_ORGANOPHOSPHORUS | 126 | -0.4484 | -1.281595 | 0.040527 | 0.221631 |
| GOBP_SEX_DIFFERENTIATION | 227 | -0.436554 | -1.281569 | 0.02004 | 0.221529 |
| GOBP_POSITIVE_REGULATION_OF_MYOBLAST_DIFFERENTIATION | 15 | -0.579211 | -1.281493 | 0.17707 | 0.221503 |
| GOBP_MYOTUBE_DIFFERENTIATION | 94 | -0.450871 | -1.280337 | 0.071281 | 0.223308 |
| GOBP_POSITIVE_REGULATION_OF_BMP_SIGNALING_PATHWAY | 31 | -0.514327 | -1.28013 | 0.143513 | 0.223524 |
| GOBP_PHOSPHATE_ION_TRANSPORT | 22 | -0.542748 | -1.280005 | 0.143382 | 0.223606 |
| GOBP_CARDIOBLAST_DIFFERENTIATION | 16 | -0.567003 | -1.279964 | 0.165554 | 0.223527 |
| GOBP_POSITIVE_REGULATION_OF_FAT_CELL_DIFFERENTIATION | 58 | -0.478744 | -1.279463 | 0.099147 | 0.224203 |
| GOBP_GAMMA_AMINOBUTYRIC_ACID_SIGNALING_PATHWAY | 26 | -0.518583 | -1.279458 | 0.148936 | 0.224072 |
| GOBP_POSITIVE_REGULATION_OF_COAGULATION | 20 | -0.550254 | -1.279177 | 0.145985 | 0.224392 |
| GOBP_POSTSYNAPTIC_SIGNAL_TRANSDUCTION | 31 | -0.513059 | -1.278998 | 0.143692 | 0.224545 |
| GOBP_RAS_PROTEIN_SIGNAL_TRANSDUCTION | 332 | -0.432471 | -1.27874 | 0.007 | 0.224842 |
| GOBP_CARDIAC_MUSCLE_TISSUE_DEVELOPMENT | 181 | -0.433577 | -1.278719 | 0.033099 | 0.224733 |
| GOBP_EPIBOLY | 34 | -0.505177 | -1.278554 | 0.153226 | 0.224876 |
| GOBP_REPLACEMENT_OSSIFICATION | 28 | -0.515502 | -1.278554 | 0.150943 | 0.224732 |
| GOBP_CELLULAR_RESPONSE_TO_MECHANICAL_STIMULUS | 73 | -0.461131 | -1.27838 | 0.095339 | 0.22488 |
| GOBP_POSITIVE_REGULATION_OF_TRANSMEMBRANE_RECEPTOR_PROTEIN_ | 99 | -0.451642 | -1.277997 | 0.059794 | 0.225391 |
| GOBP_MYD88_INDEPENDENT_TOLL_LIKE_RECEPTOR_SIGNALING_PATHWAY | 34 | -0.510735 | -1.277907 | 0.12761 | 0.225388 |
| GOBP_AMINE_METABOLIC_PROCESS | 151 | -0.43573 | -1.27745 | 0.032291 | 0.226039 |
| GOBP_NEGATIVE_REGULATION_OF_OSTEOBLAST_DIFFERENTIATION | 43 | -0.484143 | -1.277313 | 0.137555 | 0.226121 |
| GOBP_RESPONSE_TO_ZINC_ION | 51 | -0.475725 | -1.277238 | 0.122271 | 0.226114 |
| GOBP_STEM_CELL_DIFFERENTIATION | 236 | -0.438087 | -1.277178 | 0.017034 | 0.226085 |
| GOBP_PROTEIN_LOCALIZATION_TO_GOLGI_APPARATUS | 27 | -0.514047 | -1.276905 | 0.133178 | 0.226395 |
| GOBP_POSITIVE_REGULATION_OF_DNA_BINDING_TRANSCRIPTION_FACTOR_ | 253 | -0.434901 | -1.276567 | 0.011022 | 0.226833 |
| GOBP_NEGATIVE_REGULATION_OF_PROTEOLYSIS | 278 | -0.430775 | -1.276316 | 0.011 | 0.227116 |
| GOBP_EPITHELIAL_STRUCTURE_MAINTENANCE | 25 | -0.515773 | -1.275995 | 0.157452 | 0.227522 |
| GOBP_COLLAGEN_FIBRIL_ORGANIZATION | 53 | -0.47598 | -1.275972 | 0.117647 | 0.227417 |
| GOBP_TRABECULA_MORPHOGENESIS | 40 | -0.48745 | -1.275892 | 0.110497 | 0.22742 |
| GOBP_REGULATION_OF_EPITHELIAL_TO_MESENCHYMAL_TRANSITION | 92 | -0.456269 | -1.275193 | 0.068252 | 0.228472 |
| GOBP_REGULATION_OF_STRIATED_MUSCLE_CELL_DIFFERENTIATION | 72 | -0.46292 | -1.27505 | 0.097689 | 0.228563 |
| GOBP_NEGATIVE_REGULATION_OF_PROTEIN_DEPHOSPHORYLATION | 60 | -0.470221 | -1.274904 | 0.103373 | 0.228678 |
| GOBP_SMALL_GTPASE_MEDIATED_SIGNAL_TRANSDUCTION | 493 | -0.43055 | -1.2747 | 0.001 | 0.228867 |
| GOBP_HEART_GROWTH | 80 | -0.458221 | -1.274127 | 0.093684 | 0.22968 |
| GOBP_SPLEEN_DEVELOPMENT | 31 | -0.515345 | -1.273972 | 0.14386 | 0.229764 |
| GOBP_RENAL_SYSTEM_VASCULATURE_DEVELOPMENT | 23 | -0.537669 | -1.273598 | 0.154567 | 0.230285 |
| GOBP_REGULATION_OF_MUSCLE_CONTRACTION | 141 | -0.44108 | -1.273325 | 0.039394 | 0.230626 |
| GOBP_POSITIVE_REGULATION_OF_NUCLEAR_DIVISION | 50 | -0.480578 | -1.272905 | 0.108253 | 0.231164 |
| GOBP_CEREBELLAR_CORTEX_FORMATION | 22 | -0.537232 | -1.27273 | 0.15684 | 0.231315 |
| GOBP_REGULATION_OF_DNA_DAMAGE_RESPONSE_SIGNAL_TRANSDUCTION_ | 34 | -0.497496 | -1.272515 | 0.149714 | 0.231552 |
| GOBP_NEGATIVE_REGULATION_OF_CELL_JUNCTION_ASSEMBLY | 30 | -0.506891 | -1.272024 | 0.164144 | 0.23224 |
| GOBP_DOPAMINE_RECEPTOR_SIGNALING_PATHWAY | 38 | -0.500596 | -1.271783 | 0.129545 | 0.232509 |
| GOBP_REGULATION_OF_STRIATED_MUSCLE_CONTRACTION | 83 | -0.455189 | -1.271022 | 0.094105 | 0.23367 |
| GOBP_RESPONSE_TO_ACIDIC_PH | 22 | -0.534987 | -1.270519 | 0.167494 | 0.23441 |
| GOBP_VITAMIN_METABOLIC_PROCESS | 121 | -0.447644 | -1.270339 | 0.051777 | 0.234554 |
| GOBP_RESPONSE_TO_MURAMYL_DIPEPTIDE | 20 | -0.541505 | -1.270155 | 0.171149 | 0.234716 |
| GOBP_REGULATION_OF_MYOTUBE_DIFFERENTIATION | 31 | -0.508149 | -1.270008 | 0.166098 | 0.234837 |
| GOBP_RESPONSE_TO_INORGANIC_SUBSTANCE | 498 | -0.427517 | -1.269975 | 0.002 | 0.234751 |
| GOBP_REGULATION_OF_ENDOCYTOSIS | 193 | -0.438429 | -1.269462 | 0.033166 | 0.235511 |
| GOBP_IRON_ION_HOMEOSTASIS | 80 | -0.454629 | -1.269083 | 0.088174 | 0.236031 |
| GOBP_NEGATIVE_REGULATION_OF_PEPTIDE_SECRETION | 61 | -0.465216 | -1.268788 | 0.098291 | 0.236405 |
| GOBP_POSITIVE_REGULATION_OF_GLIAL_CELL_PROLIFERATION | 16 | -0.574794 | -1.2681 | 0.158236 | 0.237467 |
| GOBP_REGULATION_OF_EXTENT_OF_CELL_GROWTH | 102 | -0.448422 | -1.267815 | 0.072449 | 0.237853 |
| GOBP_REGULATION_OF_ORGANIC_ACID_TRANSPORT | 53 | -0.47164 | -1.267814 | 0.117904 | 0.237705 |
| GOBP_SPHINGOLIPID_BIOSYNTHETIC_PROCESS | 107 | -0.450377 | -1.26747 | 0.072375 | 0.238167 |
| GOBP_HEMATOPOIETIC_PROGENITOR_CELL_DIFFERENTIATION | 156 | -0.440487 | -1.267125 | 0.051463 | 0.238638 |
| GOBP_REGULATION_OF_PEPTIDYL_THREONINE_PHOSPHORYLATION | 43 | -0.48722 | -1.266986 | 0.132203 | 0.238739 |
| GOBP_CELLULAR_RESPONSE_TO_ACID_CHEMICAL | 65 | -0.460757 | -1.266259 | 0.117271 | 0.239868 |
| GOBP_REGULATION_OF_LONG_TERM_NEURONAL_SYNAPTIC_PLASTICITY | 21 | -0.53342 | -1.266158 | 0.155194 | 0.239906 |
| GOBP_BASEMENT_MEMBRANE_ASSEMBLY | 15 | -0.573212 | -1.266025 | 0.162549 | 0.239982 |
| GOBP_NEGATIVE_REGULATION_OF_SMALL_GTPASE_MEDIATED_SIGNAL_TRA | 56 | -0.466567 | -1.265825 | 0.125677 | 0.240177 |
| GOBP_NEGATIVE_REGULATION_OF_NF_KAPPAB_TRANSCRIPTION_FACTOR_A | 82 | -0.458749 | -1.265474 | 0.094595 | 0.240658 |
| GOBP_REGULATION_OF_PROTEIN_EXIT_FROM_ENDOPLASMIC_RETICULUM | 27 | -0.519697 | -1.263776 | 0.157773 | 0.243544 |
| GOBP_ENERGY_HOMEOSTASIS | 35 | -0.493251 | -1.263697 | 0.15411 | 0.243538 |
| GOBP_SUPEROXIDE_METABOLIC_PROCESS | 63 | -0.465778 | -1.263628 | 0.118407 | 0.243508 |
| GOBP_CELL_CELL_RECOGNITION | 64 | -0.46168 | -1.263549 | 0.108626 | 0.24349 |
| GOBP_POSITIVE_REGULATION_OF_AXONOGENESIS | 76 | -0.459322 | -1.263247 | 0.095388 | 0.243873 |
| GOBP_REGULATION_OF_GLYCOGEN_METABOLIC_PROCESS | 34 | -0.492547 | -1.2632 | 0.147465 | 0.243812 |
| GOBP_EPITHELIAL_TUBE_MORPHOGENESIS | 293 | -0.426619 | -1.262629 | 0.013013 | 0.244625 |
| GOBP_NEGATIVE_REGULATION_OF_ORGANIC_ACID_TRANSPORT | 17 | -0.557086 | -1.262518 | 0.167482 | 0.244677 |
| GOBP_EMBRYONIC_PLACENTA_MORPHOGENESIS | 23 | -0.530632 | -1.261726 | 0.158273 | 0.245947 |
| GOBP_CRANIAL_SKELETAL_SYSTEM_DEVELOPMENT | 61 | -0.465733 | -1.261698 | 0.106092 | 0.245851 |
| GOBP_AXONAL_FASCICULATION | 19 | -0.545895 | -1.261451 | 0.17037 | 0.246143 |
| GOBP_NEGATIVE_REGULATION_OF_GLYCOPROTEIN_METABOLIC_PROCESS | 15 | -0.572843 | -1.261115 | 0.183673 | 0.246589 |
| GOBP_REGULATION_OF_DELAYED_RECTIFIER_POTASSIUM_CHANNEL_ACTIVI | 18 | -0.55371 | -1.260561 | 0.180348 | 0.247435 |
| GOBP_INNER_EAR_RECEPTOR_CELL_STEREOCILIUM_ORGANIZATION | 28 | -0.513063 | -1.260548 | 0.160188 | 0.2473 |
| GOBP_MATING | 35 | -0.491106 | -1.260009 | 0.155153 | 0.248133 |
| GOBP_RESPONSE_TO_RETINOIC_ACID | 97 | -0.441126 | -1.259685 | 0.090062 | 0.248592 |
| GOBP_LYMPH_VESSEL_MORPHOGENESIS | 21 | -0.530865 | -1.259248 | 0.17596 | 0.249214 |
| GOBP_REGULATION_OF_DENDRITIC_SPINE_MORPHOGENESIS | 45 | -0.479819 | -1.258691 | 0.151584 | 0.250054 |
| GOBP_NEGATIVE_REGULATION_OF_WOUND_HEALING | 60 | -0.462409 | -1.258594 | 0.131579 | 0.250083 |
| GOBP_SLEEP | 23 | -0.530159 | -1.258204 | 0.167874 | 0.250658 |
| GOBP_ENTRY_INTO_HOST | 145 | -0.435994 | -1.258046 | 0.05136 | 0.250786 |
| GOBP_NEGATIVE_REGULATION_OF_MYOBLAST_DIFFERENTIATION | 24 | -0.534148 | -1.257486 | 0.155152 | 0.251634 |
| GOBP_REGULATION_OF_NUCLEOTIDE_METABOLIC_PROCESS | 114 | -0.439134 | -1.2574 | 0.064024 | 0.251633 |
| GOBP_ORGANELLE_MEMBRANE_FUSION | 109 | -0.439953 | -1.257192 | 0.075433 | 0.251826 |
| GOBP_SENSORY_PERCEPTION_OF_TASTE | 39 | -0.490924 | -1.256318 | 0.161863 | 0.253235 |
| GOBP_MESENCHYME_DEVELOPMENT | 263 | -0.426219 | -1.256263 | 0.019019 | 0.253182 |
| GOBP_POSITIVE_REGULATION_OF_CELL_SUBSTRATE_ADHESION | 114 | -0.438226 | -1.25621 | 0.082569 | 0.253114 |
| GOBP_ATRIAL_CARDIAC_MUSCLE_CELL_TO_AV_NODE_CELL_SIGNALING | 18 | -0.559606 | -1.25612 | 0.20875 | 0.253103 |
| GOBP_FAT_CELL_DIFFERENTIATION | 201 | -0.432033 | -1.255947 | 0.041082 | 0.253253 |
| GOBP_POSITIVE_REGULATION_OF_MYELOID_LEUKOCYTE_MEDIATED_IMMUNI | 17 | -0.553955 | -1.255818 | 0.186312 | 0.253331 |
| GOBP_NEGATIVE_REGULATION_OF_STEROID_METABOLIC_PROCESS | 22 | -0.534056 | -1.255171 | 0.178218 | 0.25436 |
| GOBP_CARDIAC_MYOFIBRIL_ASSEMBLY | 16 | -0.563618 | -1.254741 | 0.178893 | 0.25499 |
| GOBP_COAGULATION | 300 | -0.423027 | -1.252867 | 0.018 | 0.258275 |
| GOBP_PHOSPHATIDYLETHANOLAMINE_BIOSYNTHETIC_PROCESS | 15 | -0.570831 | -1.252866 | 0.174782 | 0.258119 |
| GOBP_PLASMA_MEMBRANE_ORGANIZATION | 107 | -0.44354 | -1.252261 | 0.086735 | 0.259056 |
| GOBP_MONOCYTE_CHEMOTACTIC_PROTEIN_1_PRODUCTION | 16 | -0.563066 | -1.252002 | 0.185515 | 0.259345 |
| GOBP_RESPONSE_TO_PURINE_CONTAINING_COMPOUND | 140 | -0.434653 | -1.251156 | 0.082745 | 0.260758 |
| GOBP_REGULATION_OF_AMINO_ACID_TRANSPORT | 29 | -0.510356 | -1.251138 | 0.156431 | 0.260633 |
| GOBP_REGULATION_OF_ANATOMICAL_STRUCTURE_SIZE | 471 | -0.419545 | -1.25054 | 0.002 | 0.261605 |
| GOBP_REGULATION_OF_ACTION_POTENTIAL | 48 | -0.473793 | -1.250458 | 0.147987 | 0.261616 |
| GOBP_NEGATIVE_REGULATION_OF_INTERLEUKIN_1_PRODUCTION | 30 | -0.501577 | -1.250355 | 0.154639 | 0.261663 |
| GOBP_NEGATIVE_REGULATION_OF_NEURON_PROJECTION_DEVELOPMENT | 129 | -0.436708 | -1.250011 | 0.072654 | 0.262106 |
| GOBP_RESPONSE_TO_CAMP | 91 | -0.447467 | -1.249915 | 0.10219 | 0.262134 |
| GOBP_NEGATIVE_REGULATION_OF_AXONOGENESIS | 66 | -0.457876 | -1.249858 | 0.135106 | 0.262087 |
| GOBP_PEPTIDE_SECRETION | 362 | -0.420575 | -1.248782 | 0.01 | 0.263984 |
| GOBP_POSITIVE_REGULATION_OF_PLASMA_MEMBRANE_BOUNDED_CELL_PR | 104 | -0.445979 | -1.248515 | 0.088776 | 0.264314 |
| GOBP_DETECTION_OF_STIMULUS_INVOLVED_IN_SENSORY_PERCEPTION_OF_ | 22 | -0.522299 | -1.248139 | 0.184874 | 0.26489 |
| GOBP_POSITIVE_REGULATION_OF_ERYTHROCYTE_DIFFERENTIATION | 31 | -0.495855 | -1.247985 | 0.180667 | 0.265033 |
| GOBP_STRIATED_MUSCLE_CELL_DIFFERENTIATION | 230 | -0.42542 | -1.247852 | 0.02002 | 0.265118 |
| GOBP_DEVELOPMENTAL_MATURATION | 248 | -0.424363 | -1.247138 | 0.027 | 0.266301 |
| GOBP_RESPONSE_TO_ATP | 29 | -0.501493 | -1.247038 | 0.171192 | 0.266314 |
| GOBP_STRESS_RESPONSE_TO_METAL_ION | 15 | -0.570026 | -1.246843 | 0.182519 | 0.266534 |
| GOBP_EXCRETION | 51 | -0.464006 | -1.24619 | 0.146104 | 0.267641 |
| GOBP_REGULATION_OF_JUN_KINASE_ACTIVITY | 80 | -0.446553 | -1.246188 | 0.114703 | 0.267482 |
| GOBP_SECONDARY_METABOLIC_PROCESS | 50 | -0.460111 | -1.246132 | 0.133551 | 0.267426 |
| GOBP_POSITIVE_REGULATION_OF_PROTEIN_DEPOLYMERIZATION | 19 | -0.534591 | -1.245757 | 0.193827 | 0.267962 |
| GOBP_ACTIVATION_OF_PROTEIN_KINASE_ACTIVITY | 314 | -0.422375 | -1.245612 | 0.015015 | 0.268076 |
| GOBP_CARDIAC_CHAMBER_MORPHOGENESIS | 107 | -0.438305 | -1.244552 | 0.097938 | 0.269996 |
| GOBP_CELL_DIFFERENTIATION_INVOLVED_IN_METANEPHROS_DEVELOPMEN | 23 | -0.513872 | -1.243923 | 0.181384 | 0.271031 |
| GOBP_REGULATION_OF_RENAL_SYSTEM_PROCESS | 28 | -0.511791 | -1.243238 | 0.175148 | 0.272141 |
| GOBP_RESPONSE_TO_FIBROBLAST_GROWTH_FACTOR | 128 | -0.435596 | -1.243208 | 0.080203 | 0.272037 |
| GOBP_RENAL_WATER_HOMEOSTASIS | 32 | -0.498733 | -1.243185 | 0.165882 | 0.271918 |
| GOBP_REGULATION_OF_MUSCLE_SYSTEM_PROCESS | 207 | -0.426562 | -1.243125 | 0.042042 | 0.271874 |
| GOBP_RESPONSE_TO_TEMPERATURE_STIMULUS | 224 | -0.424824 | -1.242725 | 0.03006 | 0.272442 |
| GOBP_T_HELPER_1_CELL_DIFFERENTIATION | 18 | -0.539022 | -1.242653 | 0.19975 | 0.272408 |

| | | | | | |
|---|---|---|---|---|---|
| GOBP_RESPONSE_TO_AXON_INJURY | 71 | -0.453829 | -1.242468 | 0.132556 | 0.272622 |
| GOBP_POSITIVE_REGULATION_OF_CILIUM_ASSEMBLY | 24 | -0.51895 | -1.24246 | 0.173965 | 0.272473 |
| GOBP_DEVELOPMENT_OF_PRIMARY_SEXUAL_CHARACTERISTICS | 188 | -0.422971 | -1.240334 | 0.050251 | 0.276338 |
| GOBP_KERATAN_SULFATE_METABOLIC_PROCESS | 32 | -0.491973 | -1.239971 | 0.172924 | 0.276875 |
| GOBP_PROSTANOID_METABOLIC_PROCESS | 41 | -0.478986 | -1.239192 | 0.153504 | 0.27817 |
| GOBP_MYOBLAST_FUSION | 35 | -0.491233 | -1.238704 | 0.198404 | 0.278949 |
| GOBP_CATECHOLAMINE_UPTAKE | 16 | -0.553981 | -1.238653 | 0.204005 | 0.278878 |
| GOBP_VESICLE_DOCKING_INVOLVED_IN_EXOCYTOSIS | 44 | -0.471391 | -1.238645 | 0.183114 | 0.278732 |
| GOBP_AXON_ENSHEATHMENT_IN_CENTRAL_NERVOUS_SYSTEM | 18 | -0.530725 | -1.238488 | 0.196701 | 0.278862 |
| GOBP_EMBRYONIC_PATTERN_SPECIFICATION | 57 | -0.460546 | -1.238164 | 0.142857 | 0.279334 |
| GOBP_NEGATIVE_REGULATION_OF_MUSCLE_CONTRACTION | 19 | -0.541376 | -1.237996 | 0.194548 | 0.279483 |
| GOBP_SOMITOGENESIS | 57 | -0.458157 | -1.237875 | 0.155005 | 0.279522 |
| GOBP_REGULATION_OF_REACTIVE_OXYGEN_SPECIES_METABOLIC_PROCESS | 172 | -0.425488 | -1.237797 | 0.054381 | 0.279515 |
| GOBP_NEUROBLAST_PROLIFERATION | 51 | -0.459041 | -1.237038 | 0.150549 | 0.280864 |
| GOBP_INNER_EAR_AUDITORY_RECEPTOR_CELL_DIFFERENTIATION | 31 | -0.49484 | -1.236746 | 0.183371 | 0.281276 |
| GOBP_REGULATION_OF_PLATELET_ACTIVATION | 31 | -0.490063 | -1.235979 | 0.192714 | 0.282644 |
| GOBP_RESPONSE_TO_KETONE | 178 | -0.425726 | -1.235159 | 0.069208 | 0.284114 |
| GOBP_LABYRINTHINE_LAYER_MORPHOGENESIS | 19 | -0.540873 | -1.23507 | 0.207809 | 0.284119 |
| GOBP_REGULATION_OF_FILOPODIUM_ASSEMBLY | 46 | -0.470001 | -1.234927 | 0.16383 | 0.284242 |
| GOBP_NEGATIVE_REGULATION_OF_CELL_PROJECTION_ORGANIZATION | 171 | -0.426609 | -1.234907 | 0.063317 | 0.284111 |
| GOBP_RESPONSE_TO_OXYGEN_RADICAL | 26 | -0.512408 | -1.234323 | 0.176259 | 0.285095 |
| GOBP_ENDOTHELIAL_CELL_MIGRATION | 207 | -0.422509 | -1.234271 | 0.036181 | 0.285027 |
| GOBP_EMBRYONIC_AXIS_SPECIFICATION | 29 | -0.50222 | -1.234212 | 0.189066 | 0.284992 |
| GOBP_OLEFINIC_COMPOUND_METABOLIC_PROCESS | 100 | -0.43482 | -1.234054 | 0.101957 | 0.285107 |
| GOBP_ANIMAL_ORGAN_MATURATION | 25 | -0.510028 | -1.232868 | 0.182464 | 0.287242 |
| GOBP_CELLULAR_AMINO_ACID_CATABOLIC_PROCESS | 101 | -0.434689 | -1.232832 | 0.120494 | 0.287135 |
| GOBP_SMOOTHENED_SIGNALING_PATHWAY | 128 | -0.430364 | -1.232817 | 0.089249 | 0.286998 |
| GOBP_REGULATION_OF_CELL_SHAPE | 148 | -0.422437 | -1.231524 | 0.083754 | 0.289392 |
| GOBP_POSITIVE_REGULATION_OF_DEFENSE_RESPONSE_TO_VIRUS_BY_HOS | 29 | -0.50409 | -1.231327 | 0.188235 | 0.289646 |
| GOBP_GLUTAMINE_FAMILY_AMINO_ACID_CATABOLIC_PROCESS | 25 | -0.505004 | -1.230783 | 0.183432 | 0.29057 |
| GOBP_PEPTIDYL_TYROSINE_AUTOPHOSPHORYLATION | 20 | -0.525384 | -1.230689 | 0.206135 | 0.290579 |
| GOBP_REGULATION_OF_JNK_CASCADE | 123 | -0.432924 | -1.230472 | 0.084435 | 0.290849 |
| GOBP_EMBRYONIC_CRANIAL_SKELETON_MORPHOGENESIS | 44 | -0.468103 | -1.23004 | 0.168701 | 0.291521 |
| GOBP_REGULATION_OF_FAT_CELL_DIFFERENTIATION | 118 | -0.435218 | -1.229807 | 0.102459 | 0.291806 |
| GOBP_BASEMENT_MEMBRANE_ORGANIZATION | 29 | -0.494665 | -1.229595 | 0.189535 | 0.292081 |
| GOBP_SENSORY_PERCEPTION_OF_CHEMICAL_STIMULUS | 83 | -0.441675 | -1.229411 | 0.134096 | 0.292295 |
| GOBP_THYROID_HORMONE_METABOLIC_PROCESS | 23 | -0.513742 | -1.229254 | 0.20386 | 0.292456 |
| GOBP_POSITIVE_REGULATION_OF_PEPTIDYL_LYSINE_ACETYLATION | 34 | -0.484083 | -1.228436 | 0.171558 | 0.29391 |
| GOBP_ORGANIC_HYDROXY_COMPOUND_TRANSPORT | 231 | -0.422621 | -1.228188 | 0.042169 | 0.294243 |
| GOBP_REGULATION_OF_GLIAL_CELL_PROLIFERATION | 29 | -0.500349 | -1.22813 | 0.190476 | 0.294194 |
| GOBP_NOTCH_SIGNALING_PATHWAY | 176 | -0.422079 | -1.227975 | 0.073441 | 0.294342 |
| GOBP_AXO_DENDRITIC_TRANSPORT | 69 | -0.444404 | -1.227686 | 0.137173 | 0.294764 |
| GOBP_REGULATION_OF_LIPID_METABOLIC_PROCESS | 360 | -0.41469 | -1.227527 | 0.021021 | 0.294927 |
| GOBP_CARDIAC_CELL_DEVELOPMENT | 66 | -0.451035 | -1.227233 | 0.156051 | 0.295315 |
| GOBP_REGULATION_OF_PHOSPHOPROTEIN_PHOSPHATASE_ACTIVITY | 112 | -0.432795 | -1.227229 | 0.118126 | 0.295147 |
| GOBP_PHAGOSOME_MATURATION | 45 | -0.470478 | -1.227222 | 0.182819 | 0.294986 |
| GOBP_ORGAN_INDUCTION | 20 | -0.524607 | -1.227209 | 0.211111 | 0.294844 |
| GOBP_ICOSANOID_METABOLIC_PROCESS | 94 | -0.436098 | -1.227203 | 0.122257 | 0.294685 |
| GOBP_MITOCHONDRION_LOCALIZATION | 47 | -0.468339 | -1.226815 | 0.18587 | 0.295312 |
| GOBP_LENS_FIBER_CELL_DIFFERENTIATION | 30 | -0.498623 | -1.226667 | 0.193666 | 0.295444 |
| GOBP_ACTIN_FILAMENT_BASED_TRANSPORT | 21 | -0.52607 | -1.226105 | 0.201722 | 0.29639 |
| GOBP_OVULATION_CYCLE_PROCESS | 36 | -0.486272 | -1.226072 | 0.177778 | 0.296282 |
| GOBP_REGULATION_OF_PROTEIN_TARGETING_TO_MEMBRANE | 35 | -0.478575 | -1.226051 | 0.176201 | 0.29615 |
| GOBP_RESPONSE_TO_INTERFERON_ALPHA | 21 | -0.529195 | -1.225846 | 0.21781 | 0.296372 |
| GOBP_REGULATION_OF_PROTEIN_MATURATION | 57 | -0.452725 | -1.225842 | 0.164163 | 0.296209 |
| GOBP_POSITIVE_REGULATION_OF_TRANSFORMING_GROWTH_FACTOR_BETA | 19 | -0.538283 | -1.225186 | 0.208122 | 0.29737 |
| GOBP_EXTRINSIC_APOPTOTIC_SIGNALING_PATHWAY_VIA_DEATH_DOMAIN_R | 79 | -0.442231 | -1.224969 | 0.138686 | 0.297623 |
| GOBP_RESPONSE_TO_REACTIVE_OXYGEN_SPECIES | 211 | -0.417302 | -1.224947 | 0.057229 | 0.297488 |
| GOBP_MUSCLE_TISSUE_DEVELOPMENT | 333 | -0.414164 | -1.22438 | 0.022 | 0.298437 |
| GOBP_REGULATION_OF_DNA_BINDING_TRANSCRIPTION_FACTOR_ACTIVITY | 407 | -0.41304 | -1.224165 | 0.015 | 0.298703 |
| GOBP_STRESS_ACTIVATED_PROTEIN_KINASE_SIGNALING_CASCADE | 262 | -0.415428 | -1.223928 | 0.023023 | 0.299013 |
| GOBP_NEGATIVE_REGULATION_OF_INTRACELLULAR_TRANSPORT | 52 | -0.458116 | -1.223685 | 0.170442 | 0.299356 |
| GOBP_POSITIVE_REGULATION_OF_NF_KAPPAB_TRANSCRIPTION_FACTOR_AC | 154 | -0.424447 | -1.223565 | 0.080122 | 0.29944 |
| GOBP_STABILIZATION_OF_MEMBRANE_POTENTIAL | 15 | -0.545134 | -1.223211 | 0.222938 | 0.299969 |
| GOBP_PYRIMIDINE_DEOXYRIBONUCLEOTIDE_METABOLIC_PROCESS | 22 | -0.511628 | -1.223065 | 0.229216 | 0.300109 |
| GOBP_SYNAPTIC_TRANSMISSION_DOPAMINERGIC | 21 | -0.52023 | -1.222853 | 0.208283 | 0.300377 |
| GOBP_CELLULAR_RESPONSE_TO_LEPTIN_STIMULUS | 17 | -0.54002 | -1.222509 | 0.206211 | 0.300902 |
| GOBP_REGULATION_OF_INSULIN_LIKE_GROWTH_FACTOR_RECEPTOR_SIGNA | 18 | -0.538921 | -1.221741 | 0.202179 | 0.302276 |
| GOBP_PYRIMIDINE_NUCLEOSIDE_CATABOLIC_PROCESS | 20 | -0.52921 | -1.221607 | 0.203562 | 0.302383 |
| GOBP_REGULATION_OF_PROTEIN_LOCALIZATION_TO_MEMBRANE | 187 | -0.418704 | -1.22052 | 0.058058 | 0.304369 |
| GOBP_REGULATION_OF_LIPID_BIOSYNTHETIC_PROCESS | 176 | -0.423159 | -1.220491 | 0.073515 | 0.304261 |
| GOBP_POSITIVE_REGULATION_OF_NEURON_DEATH | 89 | -0.433942 | -1.22027 | 0.120868 | 0.304536 |
| GOBP_POSITIVE_REGULATION_OF_ATP_METABOLIC_PROCESS | 39 | -0.475997 | -1.220254 | 0.190909 | 0.304389 |
| GOBP_POSITIVE_REGULATION_OF_CELLULAR_CARBOHYDRATE_METABOLIC_ | 47 | -0.466467 | -1.220182 | 0.179431 | 0.304355 |
| GOBP_PROTEIN_PROCESSING | 201 | -0.415935 | -1.220089 | 0.066265 | 0.304375 |
| GOBP_PULMONARY_VALVE_DEVELOPMENT | 21 | -0.522013 | -1.219725 | 0.209756 | 0.304937 |
| GOBP_POSITIVE_REGULATION_OF_VASOCONSTRICTION | 25 | -0.500527 | -1.21934 | 0.214541 | 0.305579 |
| GOBP_POSITIVE_REGULATION_OF_JUN_KINASE_ACTIVITY | 66 | -0.44311 | -1.219216 | 0.155391 | 0.305677 |
| GOBP_MULTICELLULAR_ORGANISM_GROWTH | 121 | -0.423618 | -1.218937 | 0.117409 | 0.306079 |
| GOBP_LONG_CHAIN_FATTY_ACID_TRANSPORT | 62 | -0.446935 | -1.218456 | 0.177402 | 0.306904 |
| GOBP_RESPONSE_TO_VITAMIN_D | 31 | -0.491593 | -1.218401 | 0.196778 | 0.306854 |
| GOBP_GLAND_MORPHOGENESIS | 104 | -0.432124 | -1.218167 | 0.12115 | 0.307152 |
| GOBP_REGULATION_OF_ISOTYPE_SWITCHING | 34 | -0.482972 | -1.217292 | 0.208046 | 0.308794 |
| GOBP_NEGATIVE_REGULATION_OF_OSSIFICATION | 30 | -0.493088 | -1.217287 | 0.211494 | 0.30863 |
| GOBP_PLASMA_MEMBRANE_PHOSPHOLIPID_SCRAMBLING | 16 | -0.546316 | -1.217031 | 0.217503 | 0.309 |
| GOBP_NUCLEOSIDE_CATABOLIC_PROCESS | 31 | -0.480895 | -1.216642 | 0.219767 | 0.309627 |
| GOBP_REGULATION_OF_LIPID_CATABOLIC_PROCESS | 50 | -0.459958 | -1.215938 | 0.186349 | 0.310912 |
| GOBP_REGULATION_OF_ORGAN_GROWTH | 86 | -0.432416 | -1.215841 | 0.153846 | 0.310947 |
| GOBP_CARDIAC_CHAMBER_DEVELOPMENT | 143 | -0.420735 | -1.215657 | 0.100908 | 0.311155 |
| GOBP_MUSCLE_ORGAN_DEVELOPMENT | 291 | -0.408972 | -1.214957 | 0.038 | 0.312475 |
| GOBP_ARACHIDONIC_ACID_METABOLIC_PROCESS | 45 | -0.466877 | -1.214796 | 0.190949 | 0.312643 |
| GOBP_MALE_SEX_DIFFERENTIATION | 134 | -0.422576 | -1.214285 | 0.102123 | 0.313514 |
| GOBP_COPULATION | 17 | -0.541547 | -1.213807 | 0.215837 | 0.314323 |
| GOBP_CARBOHYDRATE_DERIVATIVE_CATABOLIC_PROCESS | 182 | -0.417333 | -1.213343 | 0.076229 | 0.315088 |
| GOBP_MITOCHONDRION_DISTRIBUTION | 16 | -0.55046 | -1.213305 | 0.229713 | 0.314998 |
| GOBP_NEGATIVE_REGULATION_OF_PHOSPHORUS_METABOLIC_PROCESS | 488 | -0.406009 | -1.213284 | 0.015 | 0.314866 |
| GOBP_SYNAPTIC_TRANSMISSION_CHOLINERGIC | 26 | -0.498932 | -1.213226 | 0.231618 | 0.314805 |
| GOBP_REGULATION_OF_SPROUTING_ANGIOGENESIS | 39 | -0.472835 | -1.213109 | 0.203315 | 0.314879 |
| GOBP_POSITIVE_REGULATION_OF_AMYLOID_PRECURSOR_PROTEIN_CATABO | 22 | -0.521009 | -1.212813 | 0.218487 | 0.315307 |
| GOBP_REGULATION_OF_STEM_CELL_DIFFERENTIATION | 108 | -0.426061 | -1.212799 | 0.123722 | 0.315155 |
| GOBP_HEART_MORPHOGENESIS | 224 | -0.415897 | -1.212796 | 0.058175 | 0.31498 |
| GOBP_POSITIVE_REGULATION_OF_KERATINOCYTE_DIFFERENTIATION | 17 | -0.538368 | -1.212772 | 0.22093 | 0.314846 |
| GOBP_RESPONSE_TO_AMINE | 42 | -0.465478 | -1.212092 | 0.213732 | 0.316065 |
| GOBP_REGULATION_OF_NITRIC_OXIDE_SYNTHASE_ACTIVITY | 45 | -0.464669 | -1.21203 | 0.186608 | 0.316025 |
| GOBP_ICOSANOID_SECRETION | 32 | -0.481866 | -1.210769 | 0.202067 | 0.318539 |
| GOBP_POSTSYNAPTIC_MEMBRANE_ORGANIZATION | 36 | -0.470326 | -1.210534 | 0.207441 | 0.318853 |
| GOBP_REGULATION_OF_CELLULAR_RESPONSE_TO_VASCULAR_ENDOTHELIA | 20 | -0.521987 | -1.210284 | 0.255346 | 0.319193 |
| GOBP_LUNG_MORPHOGENESIS | 45 | -0.459252 | -1.210053 | 0.206402 | 0.31951 |
| GOBP_NEGATIVE_REGULATION_OF_DNA_BINDING_TRANSCRIPTION_FACTOR_ | 154 | -0.415672 | -1.209736 | 0.106533 | 0.320007 |
| GOBP_RESPONSE_TO_LITHIUM_ION | 18 | -0.528777 | -1.209598 | 0.227684 | 0.320128 |
| GOBP_RIBONUCLEOTIDE_CATABOLIC_PROCESS | 39 | -0.464184 | -1.208613 | 0.186592 | 0.322001 |
| GOBP_REGULATION_OF_HORMONE_SECRETION | 236 | -0.409355 | -1.208532 | 0.069069 | 0.321994 |
| GOBP_REGULATION_OF_DENDRITIC_SPINE_DEVELOPMENT | 59 | -0.445244 | -1.20846 | 0.163617 | 0.32196 |
| GOBP_CELLULAR_RESPONSE_TO_CAMP | 51 | -0.452975 | -1.207733 | 0.189845 | 0.323298 |
| GOBP_ESTABLISHMENT_OF_EPITHELIAL_CELL_POLARITY | 29 | -0.491285 | -1.207578 | 0.211085 | 0.323437 |
| GOBP_POSITIVE_REGULATION_OF_ENDOTHELIAL_CELL_APOPTOTIC_PROCES | 17 | -0.529183 | -1.207526 | 0.242804 | 0.323368 |
| GOBP_REGULATION_OF_DEPHOSPHORYLATION | 201 | -0.413506 | -1.207298 | 0.082578 | 0.323665 |
| GOBP_NEGATIVE_REGULATION_OF_CHEMOKINE_PRODUCTION | 23 | -0.502698 | -1.207188 | 0.215854 | 0.32371 |
| GOBP_COLUMNAR_CUBOIDAL_EPITHELIAL_CELL_DIFFERENTIATION | 71 | -0.434667 | -1.207179 | 0.170301 | 0.323545 |
| GOBP_G_PROTEIN_COUPLED_ACETYLCHOLINE_RECEPTOR_SIGNALING_PATH | 17 | -0.535633 | -1.207125 | 0.220253 | 0.323466 |
| GOBP_POSITIVE_REGULATION_OF_ENDOTHELIAL_CELL_MIGRATION | 105 | -0.424332 | -1.207118 | 0.124871 | 0.3233 |
| GOBP_POSITIVE_REGULATION_OF_PROTEIN_DEPHOSPHORYLATION | 44 | -0.458951 | -1.206289 | 0.190164 | 0.324858 |
| GOBP_CELL_GROWTH | 441 | -0.405978 | -1.206241 | 0.017 | 0.324776 |
| GOBP_INOSITOL_PHOSPHATE_BIOSYNTHETIC_PROCESS | 24 | -0.504611 | -1.205798 | 0.23031 | 0.325563 |
| GOBP_REGULATION_OF_RESPONSE_TO_REACTIVE_OXYGEN_SPECIES | 30 | -0.487568 | -1.205328 | 0.202797 | 0.326427 |
| GOBP_EPIDERMIS_MORPHOGENESIS | 28 | -0.49284 | -1.20479 | 0.2142 | 0.327366 |
| GOBP_UNSATURATED_FATTY_ACID_METABOLIC_PROCESS | 93 | -0.427888 | -1.204454 | 0.139896 | 0.327908 |
| GOBP_JNK_CASCADE | 183 | -0.415897 | -1.204226 | 0.079317 | 0.328206 |
| GOBP_REGULATION_OF_WNT_SIGNALING_PATHWAY | 348 | -0.407406 | -1.203821 | 0.036 | 0.328882 |
| GOBP_NEGATIVE_REGULATION_OF_STRIATED_MUSCLE_CELL_APOPTOTIC_PF | 22 | -0.515862 | -1.20379 | 0.221951 | 0.328778 |
| GOBP_AUDITORY_RECEPTOR_CELL_DEVELOPMENT | 17 | -0.528134 | -1.203434 | 0.235661 | 0.329388 |
| GOBP_REGULATION_OF_CELL_MATRIX_ADHESION | 117 | -0.418445 | -1.203211 | 0.123984 | 0.329676 |
| GOBP_CELLULAR_RESPONSE_TO_FATTY_ACID | 36 | -0.469535 | -1.203084 | 0.20595 | 0.329768 |
| GOBP_NEGATIVE_REGULATION_OF_WNT_SIGNALING_PATHWAY | 201 | -0.413226 | -1.203029 | 0.081162 | 0.3297 |
| GOBP_REGULATION_OF_KERATINOCYTE_DIFFERENTIATION | 36 | -0.467275 | -1.202883 | 0.212838 | 0.32985 |
| GOBP_ANTERIOR_POSTERIOR_AXIS_SPECIFICATION | 39 | -0.472239 | -1.202508 | 0.205682 | 0.330433 |
| GOBP_PROTEIN_ACTIVATION_CASCADE | 20 | -0.517176 | -1.202307 | 0.240048 | 0.330669 |
| GOBP_POSITIVE_REGULATION_OF_TOLL_LIKE_RECEPTOR_SIGNALING_PATHW | 26 | -0.495411 | -1.202125 | 0.232311 | 0.330866 |
| GOBP_NEGATIVE_REGULATION_OF_BINDING | 150 | -0.414546 | -1.202074 | 0.115152 | 0.3308 |
| GOBP_POSITIVE_REGULATION_OF_PROTEIN_SERINE_THREONINE_KINASE_AC | 300 | -0.407518 | -1.201977 | 0.029 | 0.330822 |
| GOBP_PEPTIDYL_PROLINE_HYDROXYLATION | 15 | -0.555323 | -1.201934 | 0.234456 | 0.330731 |
| GOBP_REPRODUCTIVE_SYSTEM_DEVELOPMENT | 364 | -0.408395 | -1.201623 | 0.035 | 0.331227 |
| GOBP_POSITIVE_REGULATION_OF_SIGNAL_TRANSDUCTION_BY_P53_CLASS_ | 22 | -0.51368 | -1.201413 | 0.249066 | 0.331487 |
| GOBP_EXTRACELLULAR_MATRIX_ASSEMBLY | 44 | -0.461057 | -1.201367 | 0.197987 | 0.331407 |
| GOBP_POSITIVE_REGULATION_OF_CELL_SUBSTRATE_JUNCTION_ORGANIZAT | 29 | -0.490493 | -1.201257 | 0.236047 | 0.331457 |
| GOBP_FIBROBLAST_PROLIFERATION | 73 | -0.432053 | -1.200057 | 0.166316 | 0.333927 |
| GOBP_POSITIVE_REGULATION_OF_BLOOD_VESSEL_ENDOTHELIAL_CELL_MIG | 54 | -0.452742 | -1.199952 | 0.192771 | 0.333967 |
| GOBP_REGULATION_OF_CALCIUM_ION_IMPORT | 33 | -0.482953 | -1.199726 | 0.219653 | 0.334281 |
| GOBP_AROMATIC_AMINO_ACID_FAMILY_METABOLIC_PROCESS | 29 | -0.483871 | -1.199433 | 0.218713 | 0.334742 |
| GOBP_LYTIC_VACUOLE_ORGANIZATION | 70 | -0.433317 | -1.199197 | 0.176594 | 0.335089 |
| GOBP_RESPONSE_TO_STEROL | 30 | -0.477297 | -1.199192 | 0.231043 | 0.334915 |

| | | | | | |
|---|---|---|---|---|---|
| GOBP_POSITIVE_REGULATION_OF_PROTEIN_LOCALIZATION_TO_MEMBRANE | 125 | -0.422612 | -1.199159 | 0.123858 | 0.334806 |
| GOBP_NITRIC_OXIDE_SYNTHASE_BIOSYNTHETIC_PROCESS | 19 | -0.519095 | -1.198943 | 0.236745 | 0.335097 |
| GOBP_RESPONSE_TO_NUTRIENT | 155 | -0.413973 | -1.198916 | 0.120724 | 0.334982 |
| GOBP_DEVELOPMENT_OF_PRIMARY_FEMALE_SEXUAL_CHARACTERISTICS | 85 | -0.43073 | -1.198708 | 0.160752 | 0.335238 |
| GOBP_REGULATION_OF_MULTICELLULAR_ORGANISM_GROWTH | 56 | -0.449264 | -1.198495 | 0.193757 | 0.335513 |
| GOBP_PRESYNAPTIC_ENDOCYTOSIS | 59 | -0.446554 | -1.197967 | 0.184829 | 0.336497 |
| GOBP_REGULATION_OF_ACTIN_FILAMENT_BASED_PROCESS | 372 | -0.403261 | -1.197617 | 0.039 | 0.337102 |
| GOBP_MODULATION_BY_HOST_OF_SYMBIONT_PROCESS | 59 | -0.437005 | -1.196837 | 0.198062 | 0.338607 |
| GOBP_CERAMIDE_METABOLIC_PROCESS | 97 | -0.422734 | -1.196577 | 0.155992 | 0.338981 |
| GOBP_REGULATION_OF_HOMOTYPIC_CELL_CELL_ADHESION | 25 | -0.486358 | -1.196363 | 0.243437 | 0.339275 |
| GOBP_MEMBRANE_LIPID_BIOSYNTHETIC_PROCESS | 145 | -0.414845 | -1.196342 | 0.121581 | 0.339133 |
| GOBP_REGENERATION | 174 | -0.411305 | -1.195774 | 0.096677 | 0.340196 |
| GOBP_EPHRIN_RECEPTOR_SIGNALING_PATHWAY | 86 | -0.426295 | -1.195538 | 0.156863 | 0.340534 |
| GOBP_SOMATIC_RECOMBINATION_OF_IMMUNOGLOBULIN_GENE_SEGMENTS | 52 | -0.447686 | -1.194712 | 0.19779 | 0.342213 |
| GOBP_REGULATION_OF_NEUROTRANSMITTER_UPTAKE | 15 | -0.537508 | -1.193977 | 0.247776 | 0.343608 |
| GOBP_MALE_GENITALIA_DEVELOPMENT | 15 | -0.546156 | -1.193959 | 0.251938 | 0.343461 |
| GOBP_MYELOID_CELL_DIFFERENTIATION | 383 | -0.402966 | -1.193733 | 0.031 | 0.343775 |
| GOBP_REGULATION_OF_CELL_MIGRATION_INVOLVED_IN_SPROUTING_ANGIO | 38 | -0.46454 | -1.193439 | 0.202703 | 0.344223 |
| GOBP_REGULATION_OF_CARDIAC_MUSCLE_CELL_DIFFERENTIATION | 17 | -0.53695 | -1.192876 | 0.24875 | 0.345286 |
| GOBP_FILOPODIUM_ASSEMBLY | 63 | -0.434587 | -1.1924 | 0.195122 | 0.346148 |
| GOBP_ANATOMICAL_STRUCTURE_MATURATION | 198 | -0.412335 | -1.192379 | 0.090271 | 0.345998 |
| GOBP_REGULATION_OF_CARDIAC_MUSCLE_CELL_PROLIFERATION | 38 | -0.461445 | -1.192208 | 0.216426 | 0.346171 |
| GOBP_RESPONSE_TO_OSMOTIC_STRESS | 78 | -0.425228 | -1.192178 | 0.20021 | 0.346052 |
| GOBP_ENDOCRINE_HORMONE_SECRETION | 43 | -0.457844 | -1.191616 | 0.226519 | 0.347079 |
| GOBP_SUBSTRATE_ADHESION_DEPENDENT_CELL_SPREADING | 105 | -0.417361 | -1.191226 | 0.161523 | 0.34776 |
| GOBP_MEMBRANE_REPOLARIZATION_DURING_ACTION_POTENTIAL | 25 | -0.494418 | -1.191076 | 0.219688 | 0.347893 |
| GOBP_POSITIVE_REGULATION_OF_CARBOHYDRATE_METABOLIC_PROCESS | 71 | -0.435172 | -1.190897 | 0.179622 | 0.348098 |
| GOBP_REGULATION_OF_BODY_FLUID_LEVELS | 443 | -0.400511 | -1.190384 | 0.029 | 0.349065 |
| GOBP_NEGATIVE_REGULATION_OF_VIRAL_LIFE_CYCLE | 19 | -0.52049 | -1.190121 | 0.25 | 0.34946 |
| GOBP_RESPONSE_TO_TUMOR_CELL | 25 | -0.495675 | -1.190045 | 0.272397 | 0.349438 |
| GOBP_REGULATION_OF_KETONE_BIOSYNTHETIC_PROCESS | 17 | -0.5273 | -1.189772 | 0.247104 | 0.349875 |
| GOBP_CARDIAC_MUSCLE_TISSUE_MORPHOGENESIS | 53 | -0.444808 | -1.189418 | 0.214363 | 0.350503 |
| GOBP_ORGANISM_EMERGENCE_FROM_PROTECTIVE_STRUCTURE | 21 | -0.506201 | -1.18872 | 0.231138 | 0.351826 |
| GOBP_NEGATIVE_REGULATION_OF_EPITHELIAL_CELL_PROLIFERATION | 115 | -0.417496 | -1.188277 | 0.159836 | 0.352645 |
| GOBP_NOREPINEPHRINE_TRANSPORT | 20 | -0.505162 | -1.18826 | 0.247884 | 0.352494 |
| GOBP_GRANULOCYTE_DIFFERENTIATION | 28 | -0.472473 | -1.187826 | 0.25 | 0.353254 |
| GOBP_MIDBRAIN_DEVELOPMENT | 77 | -0.430708 | -1.187647 | 0.182303 | 0.353482 |
| GOBP_POSITIVE_REGULATION_OF_CELL_MATRIX_ADHESION | 53 | -0.443985 | -1.187074 | 0.207383 | 0.354545 |
| GOBP_RESPONSE_TO_FOOD | 31 | -0.471477 | -1.18688 | 0.227273 | 0.354811 |
| GOBP_CORONARY_VASCULATURE_DEVELOPMENT | 41 | -0.46025 | -1.186836 | 0.211221 | 0.35472 |
| GOBP_REGULATION_OF_CARDIAC_MUSCLE_CELL_ACTION_POTENTIAL | 25 | -0.488685 | -1.186751 | 0.243874 | 0.354724 |
| GOBP_VASCULAR_ENDOTHELIAL_CELL_PROLIFERATION | 20 | -0.510068 | -1.18616 | 0.249694 | 0.355815 |
| GOBP_REGULATION_OF_POSITIVE_CHEMOTAXIS | 25 | -0.500695 | -1.186054 | 0.256348 | 0.355879 |
| GOBP_AMYLOID_FIBRIL_FORMATION | 23 | -0.499594 | -1.185659 | 0.245169 | 0.35656 |
| GOBP_CELLULAR_RESPONSE_TO_CHOLESTEROL | 16 | -0.537883 | -1.185516 | 0.268041 | 0.356675 |
| GOBP_VACUOLAR_ACIDIFICATION | 23 | -0.500831 | -1.185071 | 0.239808 | 0.357472 |
| GOBP_PULMONARY_VALVE_MORPHOGENESIS | 17 | -0.525451 | -1.184953 | 0.243446 | 0.357549 |
| GOBP_POSITIVE_REGULATION_OF_SMOOTH_MUSCLE_CELL_MIGRATION | 37 | -0.458037 | -1.183867 | 0.245132 | 0.359758 |
| GOBP_VENTRICULAR_SEPTUM_DEVELOPMENT | 62 | -0.436464 | -1.183534 | 0.199575 | 0.360348 |
| GOBP_REGULATION_OF_PEPTIDE_HORMONE_SECRETION | 190 | -0.404347 | -1.182879 | 0.106533 | 0.361659 |
| GOBP_AMINO_SUGAR_METABOLIC_PROCESS | 40 | -0.45436 | -1.18283 | 0.241808 | 0.361584 |
| GOBP_V_D_J_RECOMBINATION | 16 | -0.525818 | -1.182764 | 0.245547 | 0.361544 |
| GOBP_REGULATION_OF_HEART_GROWTH | 61 | -0.43307 | -1.182524 | 0.199573 | 0.361893 |
| GOBP_MUSCLE_FIBER_DEVELOPMENT | 48 | -0.446686 | -1.182437 | 0.238612 | 0.36189 |
| GOBP_DETECTION_OF_MECHANICAL_STIMULUS | 44 | -0.44951 | -1.182379 | 0.218855 | 0.361836 |
| GOBP_CELLULAR_BIOGENIC_AMINE_METABOLIC_PROCESS | 94 | -0.421927 | -1.182335 | 0.186143 | 0.361747 |
| GOBP_ESTABLISHMENT_OR_MAINTENANCE_OF_CELL_POLARITY | 209 | -0.404863 | -1.180799 | 0.083 | 0.36503 |
| GOBP_RESPONSE_TO_IRON_ION | 26 | -0.489168 | -1.180412 | 0.245215 | 0.365721 |
| GOBP_POSITIVE_REGULATION_OF_GLUCOSE_METABOLIC_PROCESS | 36 | -0.459944 | -1.17971 | 0.259551 | 0.367118 |
| GOBP_DETECTION_OF_CHEMICAL_STIMULUS | 79 | -0.425494 | -1.179642 | 0.187954 | 0.367069 |
| GOBP_REGULATION_OF_PROTEIN_SERINE_THREONINE_KINASE_ACTIVITY | 468 | -0.396761 | -1.179562 | 0.03 | 0.367053 |
| GOBP_NEGATIVE_REGULATION_OF_LIPID_BIOSYNTHETIC_PROCESS | 49 | -0.440588 | -1.179169 | 0.223416 | 0.367725 |
| GOBP_ADRENAL_GLAND_DEVELOPMENT | 19 | -0.514447 | -1.179024 | 0.262701 | 0.367844 |
| GOBP_NEGATIVE_REGULATION_OF_PROTEIN_LOCALIZATION_TO_CELL_PERIF | 23 | -0.496124 | -1.178674 | 0.256257 | 0.368443 |
| GOBP_DOPAMINERGIC_NEURON_DIFFERENTIATION | 33 | -0.465713 | -1.178249 | 0.242697 | 0.369208 |
| GOBP_NUCLEOTIDE_BINDING_DOMAIN_LEUCINE_RICH_REPEAT_CONTAINING | 40 | -0.452301 | -1.177923 | 0.234375 | 0.369783 |
| GOBP_TYPE_B_PANCREATIC_CELL_PROLIFERATION | 19 | -0.5049 | -1.177639 | 0.265823 | 0.370221 |
| GOBP_REGULATION_OF_CELL_SIZE | 170 | -0.401484 | -1.176701 | 0.134673 | 0.372174 |
| GOBP_EXOCRINE_SYSTEM_DEVELOPMENT | 35 | -0.459268 | -1.176001 | 0.243678 | 0.373608 |
| GOBP_REGULATION_OF_CELL_MORPHOGENESIS | 297 | -0.402076 | -1.175757 | 0.065065 | 0.373988 |
| GOBP_POSITIVE_REGULATION_OF_TYPE_I_INTERFERON_PRODUCTION | 76 | -0.420739 | -1.175688 | 0.214361 | 0.373949 |
| GOBP_PROTEIN_HYDROXYLATION | 27 | -0.485465 | -1.175608 | 0.237028 | 0.37394 |
| GOBP_NEGATIVE_REGULATION_OF_TYPE_I_INTERFERON_PRODUCTION | 45 | -0.442438 | -1.175429 | 0.23847 | 0.374122 |
| GOBP_REGULATION_OF_RECEPTOR_LOCALIZATION_TO_SYNAPSE | 17 | -0.520074 | -1.175209 | 0.283505 | 0.374441 |
| GOBP_REGULATION_OF_NEURON_PROJECTION_ARBORIZATION | 15 | -0.527838 | -1.175191 | 0.273779 | 0.374288 |
| GOBP_RESPONSE_TO_ACID_CHEMICAL | 111 | -0.405582 | -1.174649 | 0.174975 | 0.375361 |
| GOBP_REGULATION_OF_SODIUM_ION_TRANSPORT | 82 | -0.423859 | -1.174142 | 0.184183 | 0.376333 |
| GOBP_REGULATION_OF_SYSTEMIC_ARTERIAL_BLOOD_PRESSURE_BY_RENIN | 18 | -0.514887 | -1.174057 | 0.271833 | 0.376312 |
| GOBP_PROTEIN_DEPHOSPHORYLATION | 308 | -0.395532 | -1.173237 | 0.08 | 0.37794 |
| GOBP_TRANSCYTOSIS | 20 | -0.500524 | -1.173082 | 0.278186 | 0.378101 |
| GOBP_MAINTENANCE_OF_BLOOD_BRAIN_BARRIER | 33 | -0.46224 | -1.172963 | 0.260069 | 0.378182 |
| GOBP_POSITIVE_REGULATION_OF_PROTEOLYSIS | 352 | -0.396749 | -1.172744 | 0.055055 | 0.378512 |
| GOBP_MAMMARY_GLAND_DUCT_MORPHOGENESIS | 25 | -0.492117 | -1.172169 | 0.256532 | 0.37969 |
| GOBP_EPITHELIAL_TO_MESENCHYMAL_TRANSITION | 143 | -0.404845 | -1.171667 | 0.163984 | 0.380677 |
| GOBP_MATING_BEHAVIOR | 20 | -0.504216 | -1.171653 | 0.248775 | 0.380507 |
| GOBP_RESPONSE_TO_ESTRADIOL | 114 | -0.407963 | -1.170656 | 0.173824 | 0.382621 |
| GOBP_REGULATION_OF_NEURON_PROJECTION_REGENERATION | 28 | -0.481533 | -1.170558 | 0.249714 | 0.382654 |
| GOBP_POSITIVE_REGULATION_OF_RECEPTOR_INTERNALIZATION | 24 | -0.491547 | -1.170547 | 0.2723 | 0.382483 |
| GOBP_MYELOID_CELL_APOPTOTIC_PROCESS | 25 | -0.491023 | -1.170493 | 0.25895 | 0.382413 |
| GOBP_NEGATIVE_REGULATION_OF_POTASSIUM_ION_TRANSMEMBRANE_TRA | 22 | -0.496619 | -1.169781 | 0.268005 | 0.383887 |
| GOBP_ICOSANOID_TRANSPORT | 40 | -0.461456 | -1.169393 | 0.214765 | 0.384631 |
| GOBP_NEURON_PROJECTION_EXTENSION_INVOLVED_IN_NEURON_PROJECT | 37 | -0.45907 | -1.169343 | 0.25558 | 0.38455 |
| GOBP_GLAND_DEVELOPMENT | 378 | -0.395079 | -1.169288 | 0.057 | 0.384487 |
| GOBP_CELL_VOLUME_HOMEOSTASIS | 29 | -0.479155 | -1.169239 | 0.261959 | 0.384409 |
| GOBP_MITOCHONDRIAL_DEPOLARIZATION | 22 | -0.497728 | -1.168644 | 0.263415 | 0.385619 |
| GOBP_CALCIUM_INDEPENDENT_CELL_CELL_ADHESION_VIA_PLASMA_MEMBI | 19 | -0.505104 | -1.168193 | 0.265152 | 0.386495 |
| GOBP_MAINTENANCE_OF_CELL_NUMBER | 137 | -0.406497 | -1.168077 | 0.166835 | 0.386545 |
| GOBP_BLOOD_VESSEL_REMODELING | 36 | -0.456264 | -1.167485 | 0.230337 | 0.38771 |
| GOBP_NEGATIVE_REGULATION_OF_FAT_CELL_DIFFERENTIATION | 45 | -0.443297 | -1.167355 | 0.259711 | 0.387814 |
| GOBP_AMELOGENESIS | 19 | -0.500848 | -1.167347 | 0.272616 | 0.387626 |
| GOBP_TERTIARY_ALCOHOL_METABOLIC_PROCESS | 17 | -0.511793 | -1.167239 | 0.290566 | 0.3877 |
| GOBP_POSITIVE_REGULATION_OF_PROTEIN_CONTAINING_COMPLEX_ASSEMI | 246 | -0.399557 | -1.167211 | 0.1001 | 0.387571 |
| GOBP_DIGESTION | 106 | -0.41063 | -1.166235 | 0.185904 | 0.389691 |
| GOBP_NEGATIVE_REGULATION_OF_ADENYLATE_CYCLASE_ACTIVITY | 17 | -0.520347 | -1.164328 | 0.274969 | 0.394018 |
| GOBP_OUTFLOW_TRACT_SEPTUM_MORPHOGENESIS | 24 | -0.48476 | -1.164264 | 0.264242 | 0.393983 |
| GOBP_REGULATION_OF_NON_CANONICAL_WNT_SIGNALING_PATHWAY | 25 | -0.481558 | -1.163661 | 0.264463 | 0.395224 |
| GOBP_RESPONSE_TO_STEROID_HORMONE | 304 | -0.395165 | -1.163345 | 0.081 | 0.395761 |
| GOBP_LABYRINTHINE_LAYER_BLOOD_VESSEL_DEVELOPMENT | 17 | -0.515369 | -1.162218 | 0.304293 | 0.398307 |
| GOBP_METANEPHRIC_TUBULE_DEVELOPMENT | 20 | -0.495857 | -1.162032 | 0.261557 | 0.398571 |
| GOBP_REGULATION_OF_FATTY_ACID_METABOLIC_PROCESS | 76 | -0.418652 | -1.161952 | 0.218978 | 0.398538 |
| GOBP_NEGATIVE_REGULATION_OF_PROTEIN_LOCALIZATION_TO_MEMBRANE | 28 | -0.470957 | -1.161695 | 0.262791 | 0.398964 |
| GOBP_POSITIVE_REGULATION_OF_ACTIN_FILAMENT_POLYMERIZATION | 94 | -0.412105 | -1.161662 | 0.194444 | 0.398831 |
| GOBP_CELL_PROLIFERATION_INVOLVED_IN_KIDNEY_DEVELOPMENT | 17 | -0.50693 | -1.161319 | 0.270677 | 0.399462 |
| GOBP_NEGATIVE_REGULATION_OF_MUSCLE_CELL_APOPTOTIC_PROCESS | 35 | -0.459022 | -1.161001 | 0.263098 | 0.400015 |
| GOBP_RESPONSE_TO_ALKALOID | 90 | -0.408763 | -1.159103 | 0.213915 | 0.404287 |
| GOBP_REGULATION_OF_MULTI_ORGANISM_PROCESS | 41 | -0.449273 | -1.159095 | 0.245475 | 0.404095 |
| GOBP_CELLULAR_AMINO_ACID_BIOSYNTHETIC_PROCESS | 74 | -0.417611 | -1.159082 | 0.22584 | 0.403924 |
| GOBP_FIBROBLAST_APOPTOTIC_PROCESS | 20 | -0.494651 | -1.15896 | 0.289186 | 0.404009 |
| GOBP_SKELETAL_MUSCLE_ADAPTATION | 21 | -0.498359 | -1.158777 | 0.266258 | 0.404241 |
| GOBP_EMBRYONIC_HEART_TUBE_MORPHOGENESIS | 66 | -0.423433 | -1.158137 | 0.224101 | 0.405584 |
| GOBP_MEMBRANE_PROTEIN_ECTODOMAIN_PROTEOLYSIS | 42 | -0.44819 | -1.158124 | 0.241263 | 0.405403 |
| GOBP_REGULATION_OF_PRODUCTION_OF_SMALL_RNA_INVOLVED_IN_GENE_ | 23 | -0.491009 | -1.158122 | 0.285369 | 0.405199 |
| GOBP_REGULATION_OF_SYNAPTIC_VESICLE_ENDOCYTOSIS | 15 | -0.516758 | -1.157689 | 0.29359 | 0.406055 |
| GOBP_GLYCOPROTEIN_BIOSYNTHETIC_PROCESS | 319 | -0.391043 | -1.157349 | 0.098098 | 0.406693 |
| GOBP_EMBRYONIC_CAMERA_TYPE_EYE_DEVELOPMENT | 32 | -0.462286 | -1.157307 | 0.290323 | 0.406574 |
| GOBP_LABYRINTHINE_LAYER_DEVELOPMENT | 40 | -0.441556 | -1.156888 | 0.271205 | 0.407398 |
| GOBP_REGULATION_OF_CELLULAR_RESPIRATION | 21 | -0.49037 | -1.156435 | 0.286935 | 0.408295 |
| GOBP_ACTIVATION_OF_JUN_KINASE_ACTIVITY | 38 | -0.456213 | -1.156283 | 0.265101 | 0.408464 |
| GOBP_INOSITOL_PHOSPHATE_METABOLIC_PROCESS | 66 | -0.424249 | -1.156064 | 0.234672 | 0.40878 |
| GOBP_ORGANIC_CATION_TRANSPORT | 32 | -0.458343 | -1.155432 | 0.263036 | 0.410119 |
| GOBP_REGULATION_OF_COLLAGEN_METABOLIC_PROCESS | 38 | -0.445994 | -1.155141 | 0.267261 | 0.410609 |
| GOBP_GLYCOPROTEIN_METABOLIC_PROCESS | 387 | -0.391351 | -1.155122 | 0.072 | 0.410452 |
| GOBP_SPECIFICATION_OF_SYMMETRY | 116 | -0.406954 | -1.154731 | 0.204268 | 0.411223 |
| GOBP_BILE_ACID_AND_BILE_SALT_TRANSPORT | 26 | -0.465383 | -1.153527 | 0.273672 | 0.413995 |
| GOBP_REGULATION_OF_PROTEIN_KINASE_A_SIGNALING | 20 | -0.49755 | -1.153448 | 0.298377 | 0.41397 |
| GOBP_NEGATIVE_REGULATION_OF_RESPONSE_TO_OXIDATIVE_STRESS | 20 | -0.499474 | -1.153301 | 0.265031 | 0.4141 |
| GOBP_DEPHOSPHORYLATION | 467 | -0.386751 | -1.153089 | 0.068 | 0.414389 |
| GOBP_NEGATIVE_REGULATION_OF_CANONICAL_WNT_SIGNALING_PATHWAY | 168 | -0.39014 | -1.152685 | 0.168177 | 0.415184 |
| GOBP_POSITIVE_REGULATION_OF_TRANSLATIONAL_INITIATION | 24 | -0.481812 | -1.152551 | 0.287247 | 0.415289 |
| GOBP_CELL_REDOX_HOMEOSTASIS | 43 | -0.444703 | -1.152355 | 0.263216 | 0.415546 |
| GOBP_REGULATION_OF_MUSCLE_CELL_DIFFERENTIATION | 125 | -0.400147 | -1.151847 | 0.190283 | 0.416601 |
| GOBP_AMMONIUM_ION_METABOLIC_PROCESS | 23 | -0.477722 | -1.15164 | 0.304401 | 0.41689 |
| GOBP_NEGATIVE_REGULATION_OF_MAP_KINASE_ACTIVITY | 69 | -0.41999 | -1.15078 | 0.23673 | 0.418685 |
| GOBP_ENGULFMENT_OF_APOPTOTIC_CELL | 19 | -0.491706 | -1.150753 | 0.27599 | 0.418537 |
| GOBP_REGULATION_OF_PEPTIDE_SECRETION | 275 | -0.392913 | -1.150411 | 0.116116 | 0.419154 |
| GOBP_ACTIVATION_OF_MAPK_ACTIVITY | 148 | -0.398896 | -1.150295 | 0.176946 | 0.419211 |
| GOBP_RESPONSE_TO_TOXIC_SUBSTANCE | 221 | -0.393787 | -1.15027 | 0.144289 | 0.419072 |
| GOBP_NEGATIVE_REGULATION_OF_VIRAL_GENOME_REPLICATION | 50 | -0.437914 | -1.150183 | 0.256522 | 0.419074 |
| GOBP_AXON_EXTENSION | 112 | -0.404557 | -1.150149 | 0.204268 | 0.418943 |
| GOBP_ORGAN_OR_TISSUE_SPECIFIC_IMMUNE_RESPONSE | 28 | -0.471249 | -1.14997 | 0.291139 | 0.419165 |
| GOBP_SUBSTANTIA_NIGRA_DEVELOPMENT | 42 | -0.436293 | -1.149955 | 0.274123 | 0.418986 |

| | | | | | |
|---|---|---|---|---|---|
| GOBP_POSITIVE_REGULATION_OF_EMBRYONIC_DEVELOPMENT | 19 | -0.500698 | -1.149753 | 0.290798 | 0.419269 |
| GOBP_SOMATIC_DIVERSIFICATION_OF_IMMUNE_RECEPTORS | 72 | -0.415971 | -1.149337 | 0.24866 | 0.420082 |
| GOBP_CORPUS_CALLOSUM_DEVELOPMENT | 15 | -0.522257 | -1.148914 | 0.320053 | 0.420863 |
| GOBP_RESPONSE_TO_COCAINE | 37 | -0.451593 | -1.148349 | 0.274576 | 0.42207 |
| GOBP_VASCULAR_ENDOTHELIAL_GROWTH_FACTOR_RECEPTOR_SIGNALING_ | 89 | -0.41246 | -1.148344 | 0.218201 | 0.421873 |
| GOBP_REGULATION_OF_MYELOID_CELL_APOPTOTIC_PROCESS | 20 | -0.492729 | -1.14777 | 0.302096 | 0.423054 |
| GOBP_CELL_CELL_SIGNALING_BY_WNT | 492 | -0.385028 | -1.147493 | 0.063 | 0.423522 |
| GOBP_HOMOTYPIC_CELL_CELL_ADHESION | 80 | -0.411536 | -1.146841 | 0.241667 | 0.424903 |
| GOBP_PROTEIN_TETRAMERIZATION | 79 | -0.413613 | -1.146275 | 0.248183 | 0.426083 |
| GOBP_MUSCLE_CELL_DIFFERENTIATION | 308 | -0.38904 | -1.146046 | 0.109 | 0.426451 |
| GOBP_CARDIAC_VENTRICLE_DEVELOPMENT | 108 | -0.398211 | -1.145667 | 0.219136 | 0.427204 |
| GOBP_RESPONSE_TO_FATTY_ACID | 57 | -0.42743 | -1.145378 | 0.270358 | 0.427716 |
| GOBP_ESTABLISHMENT_OF_CELL_POLARITY | 135 | -0.398346 | -1.14469 | 0.210898 | 0.429193 |
| GOBP_CEREBELLAR_PURKINJE_CELL_LAYER_DEVELOPMENT | 21 | -0.493105 | -1.144287 | 0.307971 | 0.429953 |
| GOBP_POSITIVE_REGULATION_OF_STEROID_BIOSYNTHETIC_PROCESS | 15 | -0.523708 | -1.143298 | 0.319042 | 0.432213 |
| GOBP_REGULATION_OF_CELL_PROJECTION_ASSEMBLY | 181 | -0.398351 | -1.143034 | 0.167839 | 0.432635 |
| GOBP_AMEBOIDAL_TYPE_CELL_MIGRATION | 392 | -0.386676 | -1.142924 | 0.094 | 0.432698 |
| GOBP_GLUCOCORTICOID_METABOLIC_PROCESS | 20 | -0.496754 | -1.142103 | 0.292593 | 0.434598 |
| GOBP_ACTIVATION_OF_GTPASE_ACTIVITY | 104 | -0.404477 | -1.141549 | 0.23306 | 0.435774 |
| GOBP_MAMMARY_GLAND_MORPHOGENESIS | 36 | -0.449361 | -1.141547 | 0.28341 | 0.435561 |
| GOBP_REGULATION_OF_VASCULAR_PERMEABILITY | 41 | -0.428191 | -1.141354 | 0.283465 | 0.435822 |
| GOBP_ARGININE_METABOLIC_PROCESS | 18 | -0.489199 | -1.141352 | 0.309877 | 0.435612 |
| GOBP_RESPONSE_TO_ISOQUINOLINE_ALKALOID | 25 | -0.474338 | -1.141061 | 0.302462 | 0.43612 |
| GOBP_RESPONSE_TO_INTERLEUKIN_12 | 45 | -0.438074 | -1.140803 | 0.275824 | 0.436496 |
| GOBP_REGULATION_OF_T_CELL_DIFFERENTIATION_IN_THYMUS | 22 | -0.481563 | -1.140788 | 0.302381 | 0.436319 |
| GOBP_VASCULAR_ASSOCIATED_SMOOTH_MUSCLE_CELL_MIGRATION | 20 | -0.494968 | -1.140731 | 0.304819 | 0.436243 |
| GOBP_TISSUE_REGENERATION | 64 | -0.415516 | -1.140564 | 0.264392 | 0.436434 |
| GOBP_PYRIMIDINE_NUCLEOBASE_METABOLIC_PROCESS | 16 | -0.511006 | -1.140457 | 0.316186 | 0.436484 |
| GOBP_CARDIAC_VENTRICLE_MORPHOGENESIS | 64 | -0.419223 | -1.13956 | 0.270386 | 0.438473 |
| GOBP_NEGATIVE_REGULATION_OF_REACTIVE_OXYGEN_SPECIES_BIOSYNTHE | 27 | -0.469275 | -1.138675 | 0.307514 | 0.440422 |
| GOBP_REGULATION_OF_ENDOTHELIAL_CELL_MIGRATION | 161 | -0.397628 | -1.138065 | 0.186869 | 0.441765 |
| GOBP_REGULATION_OF_PROTEIN_AUTOPHOSPHORYLATION | 45 | -0.432843 | -1.137335 | 0.285872 | 0.44337 |
| GOBP_REGULATION_OF_RAC_PROTEIN_SIGNAL_TRANSDUCTION | 18 | -0.499065 | -1.135753 | 0.304668 | 0.44717 |
| GOBP_MODULATION_BY_HOST_OF_VIRAL_PROCESS | 28 | -0.45473 | -1.135482 | 0.314079 | 0.447618 |
| GOBP_REGULATION_OF_COLLATERAL_SPROUTING | 17 | -0.500347 | -1.135428 | 0.31864 | 0.447538 |
| GOBP_POSITIVE_REGULATION_OF_AXON_EXTENSION | 37 | -0.44555 | -1.134358 | 0.303167 | 0.449994 |
| GOBP_NEGATIVE_REGULATION_OF_PEPTIDE_HORMONE_SECRETION | 37 | -0.44543 | -1.134288 | 0.290102 | 0.449947 |
| GOBP_CEREBELLAR_PURKINJE_CELL_LAYER_MORPHOGENESIS | 15 | -0.510256 | -1.134222 | 0.308271 | 0.449916 |
| GOBP_LIPOSACCHARIDE_METABOLIC_PROCESS | 109 | -0.402144 | -1.133966 | 0.219612 | 0.450346 |
| GOBP_MOVEMENT_IN_HOST_ENVIRONMENT | 166 | -0.39098 | -1.133877 | 0.195565 | 0.45034 |
| GOBP_GLYCOSYL_COMPOUND_CATABOLIC_PROCESS | 41 | -0.431142 | -1.133589 | 0.285391 | 0.450833 |
| GOBP_ORGANELLE_FUSION | 144 | -0.389954 | -1.133356 | 0.218623 | 0.451198 |
| GOBP_REACTIVE_OXYGEN_SPECIES_METABOLIC_PROCESS | 256 | -0.387302 | -1.133105 | 0.15 | 0.451578 |
| GOBP_POSITIVE_REGULATION_OF_SODIUM_ION_TRANSPORT | 34 | -0.442883 | -1.133074 | 0.305239 | 0.451422 |
| GOBP_INCLUSION_BODY_ASSEMBLY | 22 | -0.469975 | -1.131772 | 0.323208 | 0.454527 |
| GOBP_NEGATIVE_REGULATION_OF_SMALL_MOLECULE_METABOLIC_PROCES | 85 | -0.406815 | -1.131751 | 0.25387 | 0.45435 |
| GOBP_REGULATION_OF_EXTRINSIC_APOPTOTIC_SIGNALING_PATHWAY | 142 | -0.393785 | -1.130966 | 0.212916 | 0.456117 |
| GOBP_REGULATION_OF_PROTEIN_DEPHOSPHORYLATION | 135 | -0.397685 | -1.128816 | 0.226721 | 0.461384 |
| GOBP_SKELETAL_MUSCLE_ORGAN_DEVELOPMENT | 144 | -0.390806 | -1.128805 | 0.228891 | 0.461182 |
| GOBP_NEGATIVE_REGULATION_OF_INTRINSIC_APOPTOTIC_SIGNALING_PATH\ | 30 | -0.454457 | -1.128768 | 0.317497 | 0.461046 |
| GOBP_NEGATIVE_REGULATION_OF_PHOSPHORYLATION | 378 | -0.382008 | -1.128754 | 0.109 | 0.460853 |
| GOBP_NEGATIVE_REGULATION_OF_RECEPTOR_MEDIATED_ENDOCYTOSIS | 26 | -0.463718 | -1.128721 | 0.321812 | 0.460718 |
| GOBP_PEPTIDYL_SERINE_MODIFICATION | 304 | -0.383361 | -1.128652 | 0.145 | 0.460661 |
| GOBP_GLUTAMATE_METABOLIC_PROCESS | 32 | -0.453129 | -1.128366 | 0.315068 | 0.461164 |
| GOBP_SALIVARY_GLAND_DEVELOPMENT | 29 | -0.454726 | -1.128164 | 0.325882 | 0.46144 |
| GOBP_CELLULAR_RESPONSE_TO_CORTICOSTEROID_STIMULUS | 55 | -0.418035 | -1.128125 | 0.288546 | 0.461316 |
| GOBP_POSITIVE_REGULATION_OF_DENDRITIC_SPINE_MORPHOGENESIS | 19 | -0.487719 | -1.127523 | 0.322196 | 0.462637 |
| GOBP_REGULATION_OF_SMOOTHENED_SIGNALING_PATHWAY | 75 | -0.406621 | -1.127086 | 0.281217 | 0.46352 |
| GOBP_POSITIVE_REGULATION_OF_LAMELLIPODIUM_ASSEMBLY | 27 | -0.461237 | -1.126217 | 0.309662 | 0.465546 |
| GOBP_PHOSPHATIDYLINOSITOL_BIOSYNTHETIC_PROCESS | 113 | -0.394518 | -1.125961 | 0.251277 | 0.465984 |
| GOBP_INSULIN_LIKE_GROWTH_FACTOR_RECEPTOR_SIGNALING_PATHWAY | 29 | -0.455676 | -1.125954 | 0.308585 | 0.465772 |
| GOBP_AROMATIC_AMINO_ACID_FAMILY_CATABOLIC_PROCESS | 19 | -0.496583 | -1.125835 | 0.315271 | 0.465829 |
| GOBP_AORTA_DEVELOPMENT | 51 | -0.422893 | -1.1258 | 0.293668 | 0.465677 |
| GOBP_CELLULAR_RESPONSE_TO_EXTERNAL_STIMULUS | 293 | -0.380501 | -1.125767 | 0.159 | 0.465542 |
| GOBP_REGULATION_OF_FATTY_ACID_BETA_OXIDATION | 18 | -0.486964 | -1.125046 | 0.33583 | 0.467221 |
| GOBP_LIPID_PHOSPHORYLATION | 69 | -0.408668 | -1.124258 | 0.29767 | 0.468995 |
| GOBP_HEAT_GENERATION | 15 | -0.511092 | -1.12416 | 0.322745 | 0.468999 |
| GOBP_GOLGI_TO_ENDOSOME_TRANSPORT | 17 | -0.502501 | -1.122475 | 0.356868 | 0.473107 |
| GOBP_REGULATION_OF_CYSTEINE_TYPE_ENDOPEPTIDASE_ACTIVITY_INVOLV | 17 | -0.501047 | -1.122356 | 0.340796 | 0.473186 |
| GOBP_SEMI_LUNAR_VALVE_DEVELOPMENT | 40 | -0.44497 | -1.122203 | 0.307522 | 0.473339 |
| GOBP_RUFFLE_ORGANIZATION | 54 | -0.421733 | -1.120882 | 0.301846 | 0.476515 |
| GOBP_EXTRINSIC_APOPTOTIC_SIGNALING_PATHWAY | 204 | -0.384797 | -1.120866 | 0.194389 | 0.476317 |
| GOBP_TISSUE_MIGRATION | 289 | -0.379782 | -1.120756 | 0.175 | 0.476374 |
| GOBP_METANEPHRIC_EPITHELIUM_DEVELOPMENT | 21 | -0.4706 | -1.120681 | 0.311656 | 0.476326 |
| GOBP_REGULATION_OF_ALTERNATIVE_MRNA_SPLICING_VIA_SPLICEOSOME | 56 | -0.416928 | -1.120304 | 0.304629 | 0.477067 |
| GOBP_REGULATION_OF_BICELLULAR_TIGHT_JUNCTION_ASSEMBLY | 23 | -0.474179 | -1.119861 | 0.324614 | 0.477978 |
| GOBP_NEGATIVE_REGULATION_OF_INTERLEUKIN_1_BETA_PRODUCTION | 24 | -0.467208 | -1.119696 | 0.31829 | 0.478174 |
| GOBP_RESPONSE_TO_EXTRACELLULAR_STIMULUS | 438 | -0.378259 | -1.119633 | 0.13 | 0.478111 |
| GOBP_REGULATION_OF_CELLULAR_KETONE_METABOLIC_PROCESS | 160 | -0.390017 | -1.11951 | 0.227503 | 0.478193 |
| GOBP_REGULATION_OF_CILIUM_ASSEMBLY | 58 | -0.416275 | -1.119323 | 0.301604 | 0.478462 |
| GOBP_REGULATION_OF_VASCULAR_ENDOTHELIAL_GROWTH_FACTOR_RECE | 26 | -0.457594 | -1.119108 | 0.343284 | 0.478801 |
| GOBP_POSITIVE_REGULATION_OF_PROTEIN_TARGETING_TO_MEMBRANE | 30 | -0.45052 | -1.119038 | 0.32288 | 0.478762 |
| GOBP_REGULATION_OF_COAGULATION | 62 | -0.415106 | -1.118927 | 0.288889 | 0.478812 |
| GOBP_PROSTAGLANDIN_TRANSPORT | 15 | -0.505069 | -1.118653 | 0.35 | 0.479285 |
| GOBP_NEGATIVE_REGULATION_OF_RESPONSE_TO_REACTIVE_OXYGEN_SPE( | 16 | -0.50081 | -1.118282 | 0.335443 | 0.479993 |
| GOBP_MULTI_MULTICELLULAR_ORGANISM_PROCESS | 178 | -0.383824 | -1.117949 | 0.204819 | 0.480614 |
| GOBP_INTERLEUKIN_7_MEDIATED_SIGNALING_PATHWAY | 23 | -0.472205 | -1.117941 | 0.345196 | 0.480404 |
| GOBP_STRIATED_MUSCLE_ADAPTATION | 38 | -0.440062 | -1.117888 | 0.329055 | 0.480304 |
| GOBP_REGULATION_OF_PROTEIN_KINASE_C_SIGNALING | 16 | -0.490344 | -1.117832 | 0.346683 | 0.480215 |
| GOBP_ZYMOGEN_ACTIVATION | 51 | -0.423247 | -1.117744 | 0.3061 | 0.480221 |
| GOBP_POSITIVE_REGULATION_OF_G_PROTEIN_COUPLED_RECEPTOR_SIGNA | 24 | -0.459225 | -1.117735 | 0.332938 | 0.480013 |
| GOBP_REGULATION_OF_ACTIN_NUCLEATION | 30 | -0.447117 | -1.117374 | 0.328 | 0.480757 |
| GOBP_NEGATIVE_REGULATION_OF_DEVELOPMENTAL_GROWTH | 101 | -0.391 | -1.116966 | 0.288205 | 0.481571 |
| GOBP_SECRETORY_GRANULE_ORGANIZATION | 31 | -0.444658 | -1.116623 | 0.330654 | 0.482267 |
| GOBP_REGULATION_OF_ENDOCRINE_PROCESS | 32 | -0.450482 | -1.115892 | 0.331389 | 0.483955 |
| GOBP_NEGATIVE_REGULATION_OF_COAGULATION | 43 | -0.427712 | -1.115823 | 0.311732 | 0.48391 |
| GOBP_DEVELOPMENTAL_INDUCTION | 29 | -0.457408 | -1.115391 | 0.343423 | 0.484823 |
| GOBP_PYRIMIDINE_RIBONUCLEOTIDE_METABOLIC_PROCESS | 29 | -0.449323 | -1.115217 | 0.332168 | 0.485025 |
| GOBP_POST_EMBRYONIC_DEVELOPMENT | 73 | -0.401689 | -1.114212 | 0.291579 | 0.487418 |
| GOBP_MODIFIED_AMINO_ACID_TRANSPORT | 28 | -0.452437 | -1.112865 | 0.327103 | 0.490733 |
| GOBP_SKIN_EPIDERMIS_DEVELOPMENT | 75 | -0.39959 | -1.112364 | 0.296646 | 0.491826 |
| GOBP_WATER_HOMEOSTASIS | 70 | -0.407187 | -1.1121 | 0.314854 | 0.492295 |
| GOBP_POSITIVE_REGULATION_OF_PROTEIN_CONTAINING_COMPLEX_DISASSI | 34 | -0.436371 | -1.111654 | 0.335982 | 0.493222 |
| GOBP_NEGATIVE_REGULATION_OF_VIRAL_ENTRY_INTO_HOST_CELL | 17 | -0.489388 | -1.11067 | 0.337107 | 0.495617 |
| GOBP_C4_DICARBOXYLATE_TRANSPORT | 23 | -0.460995 | -1.109668 | 0.328979 | 0.498 |
| GOBP_REGULATION_OF_SUPEROXIDE_ANION_GENERATION | 19 | -0.473453 | -1.109568 | 0.334573 | 0.498019 |
| GOBP_VIRAL_GENOME_REPLICATION | 122 | -0.387803 | -1.109067 | 0.283838 | 0.499152 |
| GOBP_POSITIVE_REGULATION_OF_DENDRITIC_SPINE_DEVELOPMENT | 41 | -0.425962 | -1.10901 | 0.321267 | 0.499068 |
| GOBP_SINGLE_FERTILIZATION | 119 | -0.387657 | -1.108882 | 0.263265 | 0.499169 |
| GOBP_POSITIVE_REGULATION_OF_STEROID_METABOLIC_PROCESS | 22 | -0.46956 | -1.10852 | 0.368921 | 0.499837 |
| GOBP_CELL_MATURATION | 149 | -0.385429 | -1.108108 | 0.246465 | 0.500684 |
| GOBP_EMBRYONIC_HEART_TUBE_DEVELOPMENT | 73 | -0.403263 | -1.108027 | 0.303704 | 0.50064 |
| GOBP_SECONDARY_METABOLITE_BIOSYNTHETIC_PROCESS | 22 | -0.468383 | -1.108017 | 0.344282 | 0.500432 |
| GOBP_VERY_LONG_CHAIN_FATTY_ACID_METABOLIC_PROCESS | 32 | -0.440015 | -1.107488 | 0.336 | 0.501578 |
| GOBP_POSITIVE_REGULATION_OF_NON_CANONICAL_WNT_SIGNALING_PATHV | 15 | -0.50342 | -1.106947 | 0.361675 | 0.502774 |
| GOBP_POSITIVE_REGULATION_OF_MACROAUTOPHAGY | 58 | -0.410081 | -1.10673 | 0.316249 | 0.50311 |
| GOBP_REGULATION_OF_LAMELLIPODIUM_ASSEMBLY | 36 | -0.429631 | -1.106455 | 0.343607 | 0.503599 |
| GOBP_POSITIVE_REGULATION_OF_STRIATED_MUSCLE_CELL_DIFFERENTIATIO | 36 | -0.427814 | -1.105943 | 0.342593 | 0.504738 |
| GOBP_RESPONSE_TO_OXYGEN_LEVELS | 370 | -0.373909 | -1.105755 | 0.173173 | 0.504987 |
| GOBP_POSITIVE_REGULATION_OF_PROTEIN_LOCALIZATION_TO_CELL_PERIPH | 65 | -0.398043 | -1.104878 | 0.309322 | 0.507113 |
| GOBP_AMINE_BIOSYNTHETIC_PROCESS | 35 | -0.432545 | -1.104804 | 0.33677 | 0.507064 |
| GOBP_ADHERENS_JUNCTION_ORGANIZATION | 65 | -0.404756 | -1.104143 | 0.317359 | 0.508539 |
| GOBP_PHAGOSOME_ACIDIFICATION | 27 | -0.456971 | -1.103983 | 0.343195 | 0.508715 |
| GOBP_PROTEIN_AUTOPHOSPHORYLATION | 220 | -0.3755 | -1.103442 | 0.235 | 0.509986 |
| GOBP_NEGATIVE_REGULATION_OF_PROTEIN_BINDING | 89 | -0.390227 | -1.103228 | 0.315897 | 0.510305 |
| GOBP_ONE_CARBON_METABOLIC_PROCESS | 39 | -0.430051 | -1.102766 | 0.344062 | 0.511307 |
| GOBP_POSITIVE_REGULATION_OF_ISOTYPE_SWITCHING | 25 | -0.457916 | -1.102677 | 0.32426 | 0.511306 |
| GOBP_REGULATION_OF_RETINOIC_ACID_RECEPTOR_SIGNALING_PATHWAY | 16 | -0.494121 | -1.102652 | 0.374517 | 0.511129 |
| GOBP_BONE_MATURATION | 21 | -0.467839 | -1.102461 | 0.36409 | 0.511398 |
| GOBP_REGULATION_OF_TUMOR_NECROSIS_FACTOR_MEDIATED_SIGNALING_ | 57 | -0.411691 | -1.102105 | 0.333695 | 0.512068 |
| GOBP_ENDOTHELIAL_CELL_DEVELOPMENT | 65 | -0.402443 | -1.102023 | 0.342797 | 0.512043 |
| GOBP_RESPONSE_TO_AMPHETAMINE | 30 | -0.448714 | -1.101944 | 0.347079 | 0.512016 |
| GOBP_NEGATIVE_REGULATION_OF_KINASE_ACTIVITY | 250 | -0.376693 | -1.101261 | 0.248 | 0.51364 |
| GOBP_POSITIVE_REGULATION_OF_COLLAGEN_METABOLIC_PROCESS | 24 | -0.46238 | -1.10121 | 0.341578 | 0.513531 |
| GOBP_NEGATIVE_REGULATION_OF_LIPID_TRANSPORT | 29 | -0.446464 | -1.100912 | 0.342012 | 0.514092 |
| GOBP_SPHINGOLIPID_METABOLIC_PROCESS | 160 | -0.382202 | -1.100669 | 0.290161 | 0.514471 |
| GOBP_POSITIVE_REGULATION_OF_SECRETION | 269 | -0.371958 | -1.100458 | 0.228228 | 0.51478 |
| GOBP_NEGATIVE_REGULATION_OF_CARTILAGE_DEVELOPMENT | 25 | -0.455198 | -1.100381 | 0.353155 | 0.514738 |
| GOBP_REGULATION_OF_GLUCAN_BIOSYNTHETIC_PROCESS | 27 | -0.446416 | -1.099896 | 0.348754 | 0.51577 |
| GOBP_ESTABLISHMENT_OF_EPITHELIAL_CELL_APICAL_BASAL_POLARITY | 15 | -0.498916 | -1.099661 | 0.366324 | 0.51614 |
| GOBP_AXIS_ELONGATION | 25 | -0.462427 | -1.099267 | 0.343337 | 0.516943 |
| GOBP_THYMUS_DEVELOPMENT | 42 | -0.423824 | -1.099257 | 0.342481 | 0.516727 |
| GOBP_ARACHIDONIC_ACID_SECRETION | 21 | -0.46849 | -1.098461 | 0.371605 | 0.518617 |
| GOBP_NEGATIVE_REGULATION_OF_ENDOCYTOSIS | 45 | -0.420384 | -1.097963 | 0.340045 | 0.519674 |
| GOBP_CANONICAL_WNT_SIGNALING_PATHWAY | 314 | -0.368414 | -1.097956 | 0.216 | 0.519447 |
| GOBP_SYNAPTIC_VESICLE_RECYCLING | 72 | -0.399338 | -1.097931 | 0.331562 | 0.519271 |
| GOBP_MEMBRANE_PROTEIN_PROTEOLYSIS | 60 | -0.404944 | -1.097477 | 0.335456 | 0.520213 |
| GOBP_LEUKOTRIENE_METABOLIC_PROCESS | 26 | -0.452238 | -1.097269 | 0.355217 | 0.520539 |
| GOBP_NEUROTRANSMITTER_RECEPTOR_TRANSPORT_TO_PLASMA_MEMBRA | 16 | -0.487844 | -1.097233 | 0.36829 | 0.520394 |
| GOBP_SYMPATHETIC_NERVOUS_SYSTEM_DEVELOPMENT | 20 | -0.477528 | -1.0966 | 0.394314 | 0.521817 |
| GOBP_ESTABLISHMENT_OR_MAINTENANCE_OF_MONOPOLAR_CELL_POLARIT | 23 | -0.458845 | -1.096514 | 0.357569 | 0.521791 |

| | | | | | |
|---|---|---|---|---|---|
| GOBP_NEGATIVE_REGULATION_OF_LIPID_LOCALIZATION | 45 | -0.423036 | -1.096428 | 0.330065 | 0.521767 |
| GOBP_MAMMARY_GLAND_EPITHELIUM_DEVELOPMENT | 57 | -0.406771 | -1.09633 | 0.328294 | 0.521817 |
| GOBP_REGULATION_OF_LIPID_LOCALIZATION | 155 | -0.374186 | -1.095963 | 0.277497 | 0.52257 |
| GOBP_DICARBOXYLIC_ACID_METABOLIC_PROCESS | 93 | -0.391535 | -1.094861 | 0.324435 | 0.525259 |
| GOBP_PHOSPHOLIPID_TRANSPORT | 84 | -0.394814 | -1.094284 | 0.307933 | 0.526613 |
| GOBP_APOPTOTIC_DNA_FRAGMENTATION | 23 | -0.456481 | -1.093916 | 0.387931 | 0.527351 |
| GOBP_MUSCLE_HYPERTROPHY | 77 | -0.390048 | -1.093796 | 0.323711 | 0.527397 |
| GOBP_CORTICAL_ACTIN_CYTOSKELETON_ORGANIZATION | 38 | -0.429912 | -1.093724 | 0.353604 | 0.527348 |
| GOBP_PROTEIN_LOCALIZATION_TO_CELL_SURFACE | 63 | -0.39651 | -1.09353 | 0.328421 | 0.527642 |
| GOBP_SULFATE_TRANSPORT | 16 | -0.497982 | -1.093474 | 0.366841 | 0.527539 |
| GOBP_MORPHOGENESIS_OF_AN_ENDOTHELIUM | 18 | -0.481094 | -1.093207 | 0.353523 | 0.528014 |
| GOBP_POSITIVE_REGULATION_OF_NEUROBLAST_PROLIFERATION | 19 | -0.46758 | -1.093121 | 0.375921 | 0.528013 |
| GOBP_RESPONSE_TO_PEPTIDE | 473 | -0.370274 | -1.093072 | 0.184 | 0.527892 |
| GOBP_NEGATIVE_REGULATION_OF_SIGNALING_RECEPTOR_ACTIVITY | 56 | -0.405475 | -1.092914 | 0.341649 | 0.528095 |
| GOBP_REGULATION_OF_ESTABLISHMENT_OR_MAINTENANCE_OF_CELL_POLA | 25 | -0.448792 | -1.09216 | 0.347107 | 0.529875 |
| GOBP_EPITHELIAL_CELL_DEVELOPMENT | 193 | -0.371734 | -1.091053 | 0.281563 | 0.53258 |
| GOBP_GLUCOSE_6_PHOSPHATE_METABOLIC_PROCESS | 23 | -0.458048 | -1.090969 | 0.369249 | 0.532558 |
| GOBP_POSITIVE_REGULATION_OF_PATTERN_RECOGNITION_RECEPTOR_SIGN | 39 | -0.424853 | -1.090686 | 0.362416 | 0.53308 |
| GOBP_POSITIVE_REGULATION_OF_LEUKOCYTE_DEGRANULATION | 20 | -0.459606 | -1.090483 | 0.352357 | 0.533344 |
| GOBP_REGULATION_OF_PROTEIN_LOCALIZATION_TO_PLASMA_MEMBRANE | 95 | -0.388496 | -1.090313 | 0.318841 | 0.533563 |
| GOBP_REGULATION_OF_PROTEIN_LOCALIZATION_TO_CELL_PERIPHERY | 115 | -0.377864 | -1.089657 | 0.314227 | 0.535101 |
| GOBP_OSTEOBLAST_DEVELOPMENT | 16 | -0.498979 | -1.089554 | 0.363057 | 0.535109 |
| GOBP_BLOOD_VESSEL_ENDOTHELIAL_CELL_MIGRATION | 112 | -0.378044 | -1.089045 | 0.315951 | 0.536252 |
| GOBP_N_ACETYLGLUCOSAMINE_METABOLIC_PROCESS | 19 | -0.478891 | -1.088863 | 0.375 | 0.536526 |
| GOBP_REGULATION_OF_CELLULAR_AMINE_METABOLIC_PROCESS | 73 | -0.394991 | -1.088017 | 0.329449 | 0.538527 |
| GOBP_NEGATIVE_REGULATION_OF_VIRAL_PROCESS | 79 | -0.392044 | -1.087932 | 0.343487 | 0.538502 |
| GOBP_REGULATION_OF_ACTIN_FILAMENT_BUNDLE_ASSEMBLY | 97 | -0.388414 | -1.087781 | 0.331612 | 0.538653 |
| GOBP_STEROID_CATABOLIC_PROCESS | 22 | -0.455203 | -1.087477 | 0.375592 | 0.539265 |
| GOBP_CELLULAR_RESPONSE_TO_TOXIC_SUBSTANCE | 111 | -0.380606 | -1.087452 | 0.327818 | 0.539078 |
| GOBP_NEGATIVE_REGULATION_OF_CYTOKINE_PRODUCTION_INVOLVED_IN_I | 22 | -0.465454 | -1.087295 | 0.372738 | 0.539251 |
| GOBP_MAGNESIUM_ION_TRANSPORT | 16 | -0.489398 | -1.087148 | 0.390704 | 0.539393 |
| GOBP_NEGATIVE_REGULATION_OF_ENDOTHELIAL_CELL_PROLIFERATION | 37 | -0.420385 | -1.087147 | 0.368658 | 0.539147 |
| GOBP_RESPONSE_TO_CADMIUM_ION | 56 | -0.399816 | -1.087114 | 0.330108 | 0.538986 |
| GOBP_CELLULAR_RESPONSE_TO_CHEMICAL_STRESS | 327 | -0.368997 | -1.087068 | 0.224 | 0.538861 |
| GOBP_CONVERGENT_EXTENSION | 16 | -0.486838 | -1.087052 | 0.380832 | 0.53865 |
| GOBP_MUSCLE_CELL_DEVELOPMENT | 144 | -0.378011 | -1.086216 | 0.295363 | 0.540637 |
| GOBP_BRANCHED_CHAIN_AMINO_ACID_METABOLIC_PROCESS | 23 | -0.448619 | -1.085829 | 0.399281 | 0.541429 |
| GOBP_RESPONSE_TO_VITAMIN | 83 | -0.38439 | -1.08581 | 0.35544 | 0.541238 |
| GOBP_REGULATION_OF_VIRAL_LIFE_CYCLE | 137 | -0.378087 | -1.08484 | 0.315416 | 0.543601 |
| GOBP_DEVELOPMENTAL_PIGMENTATION | 42 | -0.415202 | -1.084764 | 0.344676 | 0.543566 |
| GOBP_REGULATION_OF_RESPONSE_TO_INTERFERON_GAMMA | 23 | -0.45971 | -1.084623 | 0.380779 | 0.543691 |
| GOBP_ANIMAL_ORGAN_REGENERATION | 68 | -0.39633 | -1.084369 | 0.356092 | 0.544156 |
| GOBP_REGULATION_OF_VIRAL_GENOME_REPLICATION | 79 | -0.38547 | -1.08402 | 0.356621 | 0.544846 |
| GOBP_REGULATION_OF_CELLULAR_COMPONENT_SIZE | 360 | -0.365912 | -1.083949 | 0.234 | 0.544807 |
| GOBP_RESPONSE_TO_TRANSFORMING_GROWTH_FACTOR_BETA | 236 | -0.369641 | -1.083862 | 0.264264 | 0.544787 |
| GOBP_NEURON_DEATH_IN_RESPONSE_TO_OXIDATIVE_STRESS | 25 | -0.447142 | -1.08358 | 0.38981 | 0.545269 |
| GOBP_REGULATION_OF_INCLUSION_BODY_ASSEMBLY | 15 | -0.485711 | -1.083114 | 0.38665 | 0.546231 |
| GOBP_REGULATION_OF_EPIDERMIS_DEVELOPMENT | 61 | -0.402192 | -1.082682 | 0.358591 | 0.547158 |
| GOBP_RESPONSE_TO_COPPER_ION | 37 | -0.420281 | -1.0825 | 0.379545 | 0.547388 |
| GOBP_PROTEIN_CONTAINING_COMPLEX_LOCALIZATION | 276 | -0.366604 | -1.082465 | 0.269539 | 0.547223 |
| GOBP_WNT_SIGNALING_PATHWAY_CALCIUM_MODULATING_PATHWAY | 37 | -0.421216 | -1.082091 | 0.372014 | 0.547987 |
| GOBP_REGULATION_OF_SYNAPTIC_TRANSMISSION_GABAERGIC | 29 | -0.44831 | -1.081873 | 0.367491 | 0.548346 |
| GOBP_NEGATIVE_REGULATION_OF_CARDIAC_MUSCLE_TISSUE_GROWTH | 24 | -0.453403 | -1.081639 | 0.382953 | 0.5487 |
| GOBP_REGULATION_OF_BINDING | 340 | -0.366051 | -1.081284 | 0.231231 | 0.549398 |
| GOBP_REGULATION_OF_RIG_I_SIGNALING_PATHWAY | 17 | -0.477219 | -1.081282 | 0.400249 | 0.549151 |
| GOBP_MULTI_ORGANISM_CELLULAR_PROCESS | 16 | -0.489543 | -1.081271 | 0.37468 | 0.548935 |
| GOBP_POSITIVE_REGULATION_OF_AMYLOID_BETA_FORMATION | 17 | -0.480502 | -1.081241 | 0.395494 | 0.548752 |
| GOBP_NUCLEOTIDE_BINDING_OLIGOMERIZATION_DOMAIN_CONTAINING_2_SI | 15 | -0.483201 | -1.081199 | 0.375318 | 0.548619 |
| GOBP_PROTEIN_CATABOLIC_PROCESS_IN_THE_VACUOLE | 18 | -0.466776 | -1.081047 | 0.3829 | 0.548782 |
| GOBP_NEGATIVE_REGULATION_OF_ESTABLISHMENT_OF_PROTEIN_LOCALIZA | 109 | -0.379573 | -1.081017 | 0.347561 | 0.548629 |
| GOBP_PLATELET_MORPHOGENESIS | 21 | -0.460655 | -1.080601 | 0.381356 | 0.549484 |
| GOBP_POSITIVE_REGULATION_OF_VIRAL_LIFE_CYCLE | 32 | -0.431706 | -1.080515 | 0.373288 | 0.549473 |
| GOBP_NEGATIVE_REGULATION_OF_CELLULAR_PROTEIN_LOCALIZATION | 107 | -0.380484 | -1.080021 | 0.331976 | 0.550533 |
| GOBP_NEGATIVE_REGULATION_OF_RHO_PROTEIN_SIGNAL_TRANSDUCTION | 21 | -0.460358 | -1.07989 | 0.384522 | 0.550623 |
| GOBP_RESPONSE_TO_FUNGUS | 34 | -0.425059 | -1.079749 | 0.362791 | 0.550739 |
| GOBP_REGULATION_OF_VASCULAR_ASSOCIATED_SMOOTH_MUSCLE_CELL_I | 51 | -0.406975 | -1.079726 | 0.376523 | 0.550547 |
| GOBP_REGULATION_OF_STEROID_BIOSYNTHETIC_PROCESS | 81 | -0.383217 | -1.07698 | 0.352941 | 0.55785 |
| GOBP_REGULATION_OF_LIPID_TRANSPORT | 126 | -0.377949 | -1.07667 | 0.310797 | 0.558417 |
| GOBP_OVULATION_CYCLE | 54 | -0.397197 | -1.076517 | 0.372826 | 0.558556 |
| GOBP_NEGATIVE_REGULATION_OF_EPITHELIAL_TO_MESENCHYMAL_TRANSIT | 35 | -0.420545 | -1.076439 | 0.379661 | 0.558512 |
| GOBP_PROTEIN_MATURATION | 268 | -0.36616 | -1.076396 | 0.277 | 0.558365 |
| GOBP_HEPATOCYTE_DIFFERENTIATION | 15 | -0.49143 | -1.075915 | 0.400498 | 0.559387 |
| GOBP_ESTABLISHMENT_OR_MAINTENANCE_OF_BIPOLAR_CELL_POLARITY | 44 | -0.407747 | -1.075482 | 0.37763 | 0.560261 |
| GOBP_REGULATION_OF_INSULIN_RECEPTOR_SIGNALING_PATHWAY | 65 | -0.396502 | -1.075249 | 0.355603 | 0.560641 |
| GOBP_2_OXOGLUTARATE_METABOLIC_PROCESS | 16 | -0.48255 | -1.075205 | 0.398524 | 0.560494 |
| GOBP_REGULATION_OF_PROTEIN_CONTAINING_COMPLEX_ASSEMBLY | 426 | -0.364186 | -1.075121 | 0.253 | 0.560464 |
| GOBP_MELANOCYTE_DIFFERENTIATION | 22 | -0.450872 | -1.075023 | 0.381773 | 0.560482 |
| GOBP_BRANCHING_INVOLVED_IN_SALIVARY_GLAND_MORPHOGENESIS | 19 | -0.467631 | -1.074063 | 0.389447 | 0.562849 |
| GOBP_FERTILIZATION | 149 | -0.369488 | -1.074062 | 0.341734 | 0.562599 |
| GOBP_REGULATION_OF_PEPTIDYL_LYSINE_ACETYLATION | 62 | -0.392709 | -1.073832 | 0.3742 | 0.562973 |
| GOBP_MUSCLE_ADAPTATION | 96 | -0.382447 | -1.073733 | 0.352214 | 0.562997 |
| GOBP_HAIR_CELL_DIFFERENTIATION | 39 | -0.414768 | -1.072937 | 0.384966 | 0.564929 |
| GOBP_POSITIVE_REGULATION_OF_PROTEIN_ACETYLATION | 43 | -0.415492 | -1.07292 | 0.366292 | 0.564709 |
| GOBP_REGULATION_OF_PROTEIN_POLYUBIQUITINATION | 25 | -0.443254 | -1.072821 | 0.384795 | 0.564698 |
| GOBP_FOLIC_ACID_CONTAINING_COMPOUND_METABOLIC_PROCESS | 27 | -0.440437 | -1.072651 | 0.399072 | 0.564897 |
| GOBP_REGULATION_OF_HAIR_CYCLE | 25 | -0.442371 | -1.072593 | 0.382249 | 0.564805 |
| GOBP_RESPONSE_TO_OXIDATIVE_STRESS | 418 | -0.361525 | -1.07215 | 0.256 | 0.56576 |
| GOBP_SODIUM_ION_IMPORT_ACROSS_PLASMA_MEMBRANE | 17 | -0.471364 | -1.071544 | 0.40613 | 0.567158 |
| GOBP_PEPTIDYL_SERINE_DEPHOSPHORYLATION | 19 | -0.464562 | -1.07138 | 0.409836 | 0.567368 |
| GOBP_AXIS_SPECIFICATION | 70 | -0.388613 | -1.071153 | 0.37422 | 0.567722 |
| GOBP_PEPTIDYL_PROLINE_MODIFICATION | 53 | -0.400203 | -1.070651 | 0.371739 | 0.568867 |
| GOBP_HORMONE_BIOSYNTHETIC_PROCESS | 62 | -0.399359 | -1.068901 | 0.39089 | 0.573469 |
| GOBP_RESPONSE_TO_INTERLEUKIN_7 | 33 | -0.420445 | -1.068887 | 0.390465 | 0.573242 |
| GOBP_PHOSPHATIDYLINOSITOL_PHOSPHORYLATION | 55 | -0.39503 | -1.06873 | 0.392199 | 0.57341 |
| GOBP_POSITIVE_REGULATION_OF_LIPID_CATABOLIC_PROCESS | 20 | -0.461485 | -1.068707 | 0.392327 | 0.573223 |
| GOBP_MUSCLE_CELL_APOPTOTIC_PROCESS | 63 | -0.396894 | -1.06854 | 0.369819 | 0.573423 |
| GOBP_POSITIVE_REGULATION_OF_EPIDERMAL_CELL_DIFFERENTIATION | 24 | -0.448152 | -1.068166 | 0.402161 | 0.574172 |
| GOBP_NEGATIVE_REGULATION_OF_GROWTH | 232 | -0.364274 | -1.06796 | 0.312249 | 0.574465 |
| GOBP_SUPEROXIDE_ANION_GENERATION | 33 | -0.426831 | -1.067819 | 0.398851 | 0.574586 |
| GOBP_POSITIVE_REGULATION_OF_SMOOTHENED_SIGNALING_PATHWAY | 34 | -0.416757 | -1.067126 | 0.390909 | 0.576223 |
| GOBP_I_KAPPAB_PHOSPHORYLATION | 19 | -0.4605 | -1.066694 | 0.404255 | 0.577117 |
| GOBP_REGULATION_OF_NECROTIC_CELL_DEATH | 33 | -0.424541 | -1.066653 | 0.38806 | 0.576967 |
| GOBP_CELLULAR_ALDEHYDE_METABOLIC_PROCESS | 58 | -0.390683 | -1.066545 | 0.381915 | 0.577 |
| GOBP_POSITIVE_REGULATION_OF_NEURON_APOPTOTIC_PROCESS | 51 | -0.397779 | -1.0664 | 0.388708 | 0.577139 |
| GOBP_NEGATIVE_REGULATION_OF_PHAGOCYTOSIS | 20 | -0.466108 | -1.065628 | 0.40709 | 0.579056 |
| GOBP_NEGATIVE_REGULATION_OF_NUCLEOTIDE_METABOLIC_PROCESS | 21 | -0.451849 | -1.065288 | 0.410224 | 0.579715 |
| GOBP_CELLULAR_RESPONSE_TO_HYDROGEN_PEROXIDE | 81 | -0.384014 | -1.064715 | 0.393782 | 0.581006 |
| GOBP_RESPONSE_TO_ARSENIC_CONTAINING_SUBSTANCE | 30 | -0.434237 | -1.06444 | 0.414894 | 0.581478 |
| GOBP_ALPHA_AMINO_ACID_METABOLIC_PROCESS | 182 | -0.363551 | -1.064048 | 0.349398 | 0.582267 |
| GOBP_ANATOMICAL_STRUCTURE_REGRESSION | 15 | -0.481767 | -1.063983 | 0.411538 | 0.582183 |
| GOBP_MITOCHONDRIAL_FISSION | 42 | -0.40293 | -1.063951 | 0.383682 | 0.582016 |
| GOBP_SOMATIC_DIVERSIFICATION_OF_IMMUNOGLOBULINS | 61 | -0.394606 | -1.063918 | 0.386096 | 0.581839 |
| GOBP_REGULATION_OF_SMALL_MOLECULE_METABOLIC_PROCESS | 397 | -0.356363 | -1.06391 | 0.289 | 0.581599 |
| GOBP_ACTIVATION_OF_INNATE_IMMUNE_RESPONSE | 136 | -0.366737 | -1.063836 | 0.364279 | 0.581538 |
| GOBP_PHOSPHATIDYLINOSITOL_METABOLIC_PROCESS | 178 | -0.364933 | -1.063536 | 0.343404 | 0.582106 |
| GOBP_NEUTRAL_AMINO_ACID_TRANSPORT | 41 | -0.406436 | -1.063416 | 0.40404 | 0.582169 |
| GOBP_POSITIVE_REGULATION_OF_ALCOHOL_BIOSYNTHETIC_PROCESS | 18 | -0.466171 | -1.062332 | 0.407814 | 0.584869 |
| GOBP_REGULATION_OF_TRIGLYCERIDE_BIOSYNTHETIC_PROCESS | 18 | -0.472286 | -1.061914 | 0.408416 | 0.585723 |
| GOBP_HISTONE_H3_K4_TRIMETHYLATION | 16 | -0.476984 | -1.060259 | 0.41519 | 0.590009 |
| GOBP_NON_CANONICAL_WNT_SIGNALING_PATHWAY | 150 | -0.370817 | -1.059714 | 0.336694 | 0.591262 |
| GOBP_POSITIVE_REGULATION_OF_VASCULAR_ASSOCIATED_SMOOTH_MUSCI | 15 | -0.480092 | -1.059668 | 0.415212 | 0.591119 |
| GOBP_NEGATIVE_REGULATION_OF_PROTEIN_SERINE_THREONINE_KINASE_A | 130 | -0.368234 | -1.05854 | 0.363452 | 0.5939 |
| GOBP_CELLULAR_RESPONSE_TO_EXTRACELLULAR_STIMULUS | 226 | -0.358947 | -1.057903 | 0.356426 | 0.595395 |
| GOBP_CYTOPLASMIC_MICROTUBULE_ORGANIZATION | 61 | -0.390036 | -1.057448 | 0.399575 | 0.59638 |
| GOBP_REGULATION_OF_B_CELL_APOPTOTIC_PROCESS | 18 | -0.470898 | -1.056794 | 0.420253 | 0.597916 |
| GOBP_TUBE_FORMATION | 137 | -0.368067 | -1.056771 | 0.386018 | 0.59771 |
| GOBP_MYOBLAST_PROLIFERATION | 17 | -0.467097 | -1.056365 | 0.426209 | 0.598557 |
| GOBP_VESICLE_ORGANIZATION | 334 | -0.35826 | -1.056116 | 0.327327 | 0.598973 |
| GOBP_PLACENTA_BLOOD_VESSEL_DEVELOPMENT | 27 | -0.423042 | -1.055778 | 0.40355 | 0.599643 |
| GOBP_POSITIVE_REGULATION_OF_INTERFERON_BETA_PRODUCTION | 33 | -0.422949 | -1.055538 | 0.417341 | 0.600038 |
| GOBP_ACTIN_FILAMENT_BUNDLE_ORGANIZATION | 152 | -0.362732 | -1.055179 | 0.403827 | 0.600727 |
| GOBP_SMOOTH_MUSCLE_TISSUE_DEVELOPMENT | 19 | -0.463847 | -1.055048 | 0.413203 | 0.600806 |
| GOBP_POSITIVE_REGULATION_OF_WNT_SIGNALING_PATHWAY | 172 | -0.362749 | -1.055042 | 0.356137 | 0.600557 |
| GOBP_PIRNA_METABOLIC_PROCESS | 16 | -0.467009 | -1.054296 | 0.40375 | 0.602276 |
| GOBP_PROTEIN_TARGETING_TO_LYSOSOME | 23 | -0.437649 | -1.053735 | 0.413915 | 0.603584 |
| GOBP_REGULATION_OF_CELL_DIVISION | 150 | -0.36301 | -1.053212 | 0.387714 | 0.604789 |
| GOBP_CALCIUM_DEPENDENT_CELL_CELL_ADHESION_VIA_PLASMA_MEMBRA | 40 | -0.409867 | -1.053085 | 0.408638 | 0.604854 |
| GOBP_AXONEME_ASSEMBLY | 70 | -0.381673 | -1.05306 | 0.403967 | 0.604658 |
| GOBP_PYRIMIDINE_NUCLEOSIDE_METABOLIC_PROCESS | 33 | -0.416192 | -1.052829 | 0.425287 | 0.605036 |
| GOBP_NEGATIVE_REGULATION_OF_TORC1_SIGNALING | 17 | -0.465505 | -1.051899 | 0.430203 | 0.607269 |
| GOBP_TRACHEA_DEVELOPMENT | 16 | -0.478955 | -1.051594 | 0.437975 | 0.607855 |
| GOBP_NEGATIVE_REGULATION_OF_APOPTOTIC_SIGNALING_PATHWAY | 207 | -0.362153 | -1.05112 | 0.378378 | 0.608845 |
| GOBP_RECEPTOR_RECYCLING | 32 | -0.423394 | -1.050326 | 0.410047 | 0.61078 |
| GOBP_VITAMIN_TRANSMEMBRANE_TRANSPORT | 15 | -0.470382 | -1.049429 | 0.433162 | 0.612948 |
| GOBP_TRANSFERRIN_TRANSPORT | 35 | -0.414171 | -1.048641 | 0.423651 | 0.614804 |
| GOBP_DETOXIFICATION | 123 | -0.366744 | -1.048241 | 0.397959 | 0.615636 |
| GOBP_NEGATIVE_REGULATION_OF_RELEASE_OF_CYTOCHROME_C_FROM_M | 16 | -0.469912 | -1.047423 | 0.401797 | 0.617672 |
| GOBP_REGULATION_OF_CARBOHYDRATE_CATABOLIC_PROCESS | 86 | -0.372971 | -1.047254 | 0.406868 | 0.617863 |
| GOBP_CARBOHYDRATE_TRANSPORT | 131 | -0.36963 | -1.047142 | 0.404858 | 0.617898 |
| GOBP_NEGATIVE_REGULATION_OF_NIK_NF_KAPPAB_SIGNALING | 25 | -0.430065 | -1.04679 | 0.423709 | 0.618592 |
| GOBP_CORTICAL_CYTOSKELETON_ORGANIZATION | 58 | -0.387594 | -1.046106 | 0.418182 | 0.620213 |

| | | | | | |
|---|---|---|---|---|---|
| GOBP_MAMMARY_GLAND_DEVELOPMENT | 121 | -0.366479 | -1.045402 | 0.422405 | 0.621894 |
| GOBP_POSITIVE_REGULATION_OF_CELLULAR_PROTEIN_LOCALIZATION | 289 | -0.356298 | -1.045333 | 0.358358 | 0.621813 |
| GOBP_GLUTATHIONE_METABOLIC_PROCESS | 55 | -0.389225 | -1.044424 | 0.419806 | 0.623965 |
| GOBP_COBALAMIN_METABOLIC_PROCESS | 20 | -0.451985 | -1.043796 | 0.425584 | 0.625418 |
| GOBP_ESTROGEN_METABOLIC_PROCESS | 26 | -0.428137 | -1.043294 | 0.428049 | 0.626516 |
| GOBP_MYOTUBE_CELL_DEVELOPMENT | 30 | -0.419982 | -1.043153 | 0.420256 | 0.626626 |
| GOBP_CELLULAR_RESPONSE_TO_VITAMIN | 28 | -0.423615 | -1.042555 | 0.442604 | 0.627957 |
| GOBP_VENTRICULAR_CARDIAC_MUSCLE_TISSUE_DEVELOPMENT | 47 | -0.396801 | -1.042011 | 0.407002 | 0.629154 |
| GOBP_REGULATION_OF_MUSCLE_HYPERTROPHY | 54 | -0.389508 | -1.041913 | 0.423119 | 0.629152 |
| GOBP_ACTIN_FILAMENT_ORGANIZATION | 418 | -0.349978 | -1.041037 | 0.36 | 0.631287 |
| GOBP_VENTRICULAR_SYSTEM_DEVELOPMENT | 27 | -0.425098 | -1.041017 | 0.43617 | 0.631064 |
| GOBP_CELL_CELL_JUNCTION_ASSEMBLY | 133 | -0.362083 | -1.040979 | 0.416582 | 0.630891 |
| GOBP_ALCOHOL_CATABOLIC_PROCESS | 47 | -0.39482 | -1.040592 | 0.429047 | 0.631665 |
| GOBP_NEGATIVE_REGULATION_OF_TRANSFORMING_GROWTH_FACTOR_BETA | 77 | -0.379866 | -1.040192 | 0.416842 | 0.632481 |
| GOBP_CHAPERONE_MEDIATED_AUTOPHAGY | 16 | -0.465542 | -1.039591 | 0.440806 | 0.63388 |
| GOBP_CYTOPLASMIC_PATTERN_RECOGNITION_RECEPTOR_SIGNALING_PATH | 68 | -0.379456 | -1.039269 | 0.419872 | 0.634502 |
| GOBP_CELLULAR_RESPONSE_TO_KETONE | 85 | -0.371273 | -1.037573 | 0.429897 | 0.63879 |
| GOBP_REGULATION_OF_APPETITE | 17 | -0.457843 | -1.037322 | 0.431498 | 0.639177 |
| GOBP_PYRIMIDINE_CONTAINING_COMPOUND_CATABOLIC_PROCESS | 36 | -0.400096 | -1.036476 | 0.417226 | 0.641214 |
| GOBP_CELL_CELL_JUNCTION_ORGANIZATION | 195 | -0.355059 | -1.036439 | 0.395582 | 0.641048 |
| GOBP_NEGATIVE_REGULATION_OF_REACTIVE_OXYGEN_SPECIES_METABOLIC | 54 | -0.389496 | -1.036113 | 0.425785 | 0.641637 |
| GOBP_NEGATIVE_REGULATION_OF_TRANSFERASE_ACTIVITY | 279 | -0.351454 | -1.035777 | 0.397397 | 0.642333 |
| GOBP_POSITIVE_REGULATION_OF_ORGANELLE_ASSEMBLY | 64 | -0.386315 | -1.035615 | 0.437037 | 0.642483 |
| GOBP_LONG_CHAIN_FATTY_ACID_METABOLIC_PROCESS | 102 | -0.362823 | -1.03542 | 0.437755 | 0.642733 |
| GOBP_DETECTION_OF_CHEMICAL_STIMULUS_INVOLVED_IN_SENSORY_PERC | 22 | -0.44111 | -1.035079 | 0.453771 | 0.643383 |
| GOBP_NEGATIVE_REGULATION_OF_CHONDROCYTE_DIFFERENTIATION | 21 | -0.442538 | -1.035017 | 0.439904 | 0.643257 |
| GOBP_REGULATION_OF_INTRACELLULAR_TRANSPORT | 339 | -0.350512 | -1.034831 | 0.384 | 0.643449 |
| GOBP_ASPARTATE_FAMILY_AMINO_ACID_METABOLIC_PROCESS | 49 | -0.391882 | -1.034768 | 0.436182 | 0.643343 |
| GOBP_PROTEIN_LOCALIZATION_TO_CELL_PERIPHERY | 320 | -0.350479 | -1.034746 | 0.395 | 0.643126 |
| GOBP_CELLULAR_RESPONSE_TO_STEROID_HORMONE_STIMULUS | 186 | -0.353448 | -1.033921 | 0.425852 | 0.645145 |
| GOBP_RESPONSE_TO_CAFFEINE | 17 | -0.457675 | -1.033771 | 0.456604 | 0.645293 |
| GOBP_POLYOL_BIOSYNTHETIC_PROCESS | 60 | -0.381783 | -1.033698 | 0.423118 | 0.645222 |
| GOBP_NEGATIVE_REGULATION_OF_MAPK_CASCADE | 161 | -0.356072 | -1.033384 | 0.425832 | 0.645801 |
| GOBP_POSITIVE_REGULATION_OF_MULTICELLULAR_ORGANISM_GROWTH | 26 | -0.425207 | -1.032957 | 0.452381 | 0.6467 |
| GOBP_REGULATION_OF_PROTEIN_BINDING | 185 | -0.351918 | -1.032909 | 0.430862 | 0.646542 |
| GOBP_CARBOHYDRATE_TRANSMEMBRANE_TRANSPORT | 100 | -0.365474 | -1.032524 | 0.436793 | 0.647336 |
| GOBP_MORPHOGENESIS_OF_EMBRYONIC_EPITHELIUM | 137 | -0.356593 | -1.032203 | 0.454545 | 0.647938 |
| GOBP_EPIDERMIS_DEVELOPMENT | 289 | -0.34858 | -1.03133 | 0.408408 | 0.650013 |
| GOBP_POSITIVE_REGULATION_OF_AUTOPHAGY | 117 | -0.363387 | -1.031075 | 0.45 | 0.65043 |
| GOBP_SENSORY_PERCEPTION_OF_SMELL | 40 | -0.398878 | -1.030851 | 0.44921 | 0.650802 |
| GOBP_RIBONUCLEOSIDE_MONOPHOSPHATE_METABOLIC_PROCESS | 55 | -0.384865 | -1.030321 | 0.432933 | 0.651984 |
| GOBP_FOLIC_ACID_METABOLIC_PROCESS | 20 | -0.444203 | -1.030261 | 0.46402 | 0.651881 |
| GOBP_CELLULAR_OXIDANT_DETOXIFICATION | 92 | -0.368364 | -1.029964 | 0.455108 | 0.652408 |
| GOBP_HYDROGEN_PEROXIDE_BIOSYNTHETIC_PROCESS | 16 | -0.465887 | -1.029681 | 0.463659 | 0.652913 |
| GOBP_SMOOTH_MUSCLE_CELL_DIFFERENTIATION | 55 | -0.380039 | -1.029529 | 0.458874 | 0.653056 |
| GOBP_CARTILAGE_DEVELOPMENT_INVOLVED_IN_ENDOCHONDRAL_BONE_M | 25 | -0.423555 | -1.029343 | 0.452071 | 0.653283 |
| GOBP_AGING | 272 | -0.347365 | -1.029178 | 0.425276 | 0.653461 |
| GOBP_BRANCHING_INVOLVED_IN_MAMMARY_GLAND_DUCT_MORPHOGENES | 18 | -0.443976 | -1.028198 | 0.467581 | 0.655856 |
| GOBP_SENSORY_PERCEPTION_OF_BITTER_TASTE | 21 | -0.437402 | -1.02772 | 0.471322 | 0.656877 |
| GOBP_MICROTUBULE_BUNDLE_FORMATION | 101 | -0.359134 | -1.027582 | 0.440904 | 0.656978 |
| GOBP_MYELOID_CELL_HOMEOSTASIS | 141 | -0.355127 | -1.027579 | 0.444332 | 0.656704 |
| GOBP_CELLULAR_RESPONSE_TO_OXYGEN_LEVELS | 219 | -0.351539 | -1.027489 | 0.438315 | 0.656673 |
| GOBP_ALCOHOL_METABOLIC_PROCESS | 338 | -0.347713 | -1.027379 | 0.403 | 0.656687 |
| GOBP_COLLATERAL_SPROUTING | 23 | -0.435145 | -1.026925 | 0.459135 | 0.657637 |
| GOBP_NUCLEOSIDE_MONOPHOSPHATE_METABOLIC_PROCESS | 72 | -0.369537 | -1.026505 | 0.448637 | 0.658504 |
| GOBP_PLATELET_AGGREGATION | 58 | -0.374522 | -1.026316 | 0.458511 | 0.658737 |
| GOBP_POSITIVE_REGULATION_OF_MUSCLE_CELL_DIFFERENTIATION | 65 | -0.374688 | -1.02609 | 0.44086 | 0.659067 |
| GOBP_THYROID_HORMONE_GENERATION | 19 | -0.449428 | -1.026023 | 0.457921 | 0.658958 |
| GOBP_ERYTHROCYTE_HOMEOSTASIS | 117 | -0.35778 | -1.026004 | 0.463739 | 0.658724 |
| GOBP_TRANSFORMING_GROWTH_FACTOR_BETA_RECEPTOR_SIGNALING_PAT | 183 | -0.354727 | -1.025416 | 0.432866 | 0.660003 |
| GOBP_HOMOPHILIC_CELL_ADHESION_VIA_PLASMA_MEMBRANE_ADHESION_M | 161 | -0.355745 | -1.025273 | 0.436052 | 0.660093 |
| GOBP_ACTIVATION_OF_NF_KAPPAB_INDUCING_KINASE_ACTIVITY | 18 | -0.450513 | -1.024383 | 0.468434 | 0.662216 |
| GOBP_NUCLEOBASE_CONTAINING_SMALL_MOLECULE_INTERCONVERSION | 27 | -0.422537 | -1.024333 | 0.456805 | 0.662072 |
| GOBP_REGULATION_OF_CYTOSKELETON_ORGANIZATION | 492 | -0.345335 | -1.023282 | 0.419 | 0.664682 |
| GOBP_MEMBRANE_LIPID_METABOLIC_PROCESS | 207 | -0.352566 | -1.023148 | 0.440321 | 0.664753 |
| GOBP_CELLULAR_KETONE_METABOLIC_PROCESS | 219 | -0.35028 | -1.022308 | 0.440882 | 0.666696 |
| GOBP_LIVER_MORPHOGENESIS | 21 | -0.436112 | -1.021972 | 0.476601 | 0.667323 |
| GOBP_SERINE_FAMILY_AMINO_ACID_BIOSYNTHETIC_PROCESS | 20 | -0.441687 | -1.021417 | 0.465201 | 0.668526 |
| GOBP_POLYOL_METABOLIC_PROCESS | 123 | -0.356027 | -1.020765 | 0.460526 | 0.670046 |
| GOBP_POSITIVE_REGULATION_OF_EPITHELIAL_CELL_MIGRATION | 145 | -0.353892 | -1.020345 | 0.464903 | 0.67088 |
| GOBP_CELLULAR_RESPONSE_TO_RETINOIC_ACID | 60 | -0.378289 | -1.020221 | 0.449786 | 0.670932 |
| GOBP_TOOTH_MINERALIZATION | 19 | -0.439176 | -1.01986 | 0.453518 | 0.67162 |
| GOBP_NUCLEAR_MIGRATION | 19 | -0.446785 | -1.019467 | 0.469849 | 0.672407 |
| GOBP_MICROVILLUS_ASSEMBLY | 16 | -0.459846 | -1.019434 | 0.479349 | 0.672215 |
| GOBP_INTERLEUKIN_1_MEDIATED_SIGNALING_PATHWAY | 99 | -0.364047 | -1.018892 | 0.460905 | 0.673393 |
| GOBP_NEGATIVE_REGULATION_OF_GLUCOSE_TRANSMEMBRANE_TRANSPOR | 15 | -0.454137 | -1.018793 | 0.479034 | 0.673365 |
| GOBP_DNA_CATABOLIC_PROCESS | 38 | -0.396477 | -1.017504 | 0.451613 | 0.676597 |
| GOBP_TOXIN_TRANSPORT | 37 | -0.399635 | -1.017254 | 0.475356 | 0.676984 |
| GOBP_STRIATED_MUSCLE_CELL_DEVELOPMENT | 91 | -0.359587 | -1.017124 | 0.470649 | 0.677036 |
| GOBP_ACTIN_POLYMERIZATION_OR_DEPOLYMERIZATION | 207 | -0.350658 | -1.016811 | 0.453908 | 0.677609 |
| GOBP_POSITIVE_REGULATION_OF_EPIDERMIS_DEVELOPMENT | 28 | -0.417078 | -1.016526 | 0.474676 | 0.678087 |
| GOBP_REGULATION_OF_CELLULAR_RESPONSE_TO_TRANSFORMING_GROWT | 117 | -0.355587 | -1.016372 | 0.471193 | 0.678216 |
| GOBP_NEGATIVE_REGULATION_OF_COLD_INDUCED_THERMOGENESIS | 46 | -0.384359 | -1.015442 | 0.468062 | 0.680453 |
| GOBP_RESPONSE_TO_GLUCAGON | 34 | -0.403142 | -1.015386 | 0.476625 | 0.680317 |
| GOBP_SPHINGOID_METABOLIC_PROCESS | 22 | -0.421404 | -1.015123 | 0.466019 | 0.680722 |
| GOBP_NEGATIVE_REGULATION_OF_CELL_GROWTH | 176 | -0.349145 | -1.014423 | 0.453172 | 0.682327 |
| GOBP_DIGESTIVE_SYSTEM_PROCESS | 81 | -0.369629 | -1.01426 | 0.496875 | 0.682465 |
| GOBP_NEGATIVE_REGULATION_OF_STEM_CELL_DIFFERENTIATION | 20 | -0.435857 | -1.014134 | 0.484812 | 0.682507 |
| GOBP_POSITIVE_REGULATION_OF_SMOOTH_MUSCLE_CONTRACTION | 23 | -0.426453 | -1.014048 | 0.471143 | 0.682465 |
| GOBP_COLLAGEN_BIOSYNTHETIC_PROCESS | 42 | -0.39065 | -1.013196 | 0.490628 | 0.684482 |
| GOBP_PROTEIN_INSERTION_INTO_MITOCHONDRIAL_MEMBRANE_INVOLVED_I | 30 | -0.409275 | -1.013131 | 0.453287 | 0.684366 |
| GOBP_NEGATIVE_REGULATION_OF_DNA_DEPENDENT_DNA_REPLICATION | 17 | -0.452316 | -1.012497 | 0.474265 | 0.685831 |
| GOBP_GMP_METABOLIC_PROCESS | 15 | -0.457666 | -1.012039 | 0.506345 | 0.686785 |
| GOBP_POSITIVE_REGULATION_OF_DNA_RECOMBINATION | 43 | -0.38673 | -1.011366 | 0.474725 | 0.688299 |
| GOBP_SCHWANN_CELL_DIFFERENTIATION | 36 | -0.399944 | -1.011023 | 0.472345 | 0.688922 |
| GOBP_REGULATION_OF_ACTIN_FILAMENT_ORGANIZATION | 262 | -0.345023 | -1.010292 | 0.46046 | 0.690592 |
| GOBP_CELLULAR_COMPONENT_ASSEMBLY_INVOLVED_IN_MORPHOGENESIS | 87 | -0.35941 | -1.009838 | 0.501023 | 0.691557 |
| GOBP_REGULATION_OF_INTRACELLULAR_PROTEIN_TRANSPORT | 244 | -0.347356 | -1.009214 | 0.496 | 0.692957 |
| GOBP_TOLL_LIKE_RECEPTOR_2_SIGNALING_PATHWAY | 16 | -0.456657 | -1.009168 | 0.492248 | 0.692787 |
| GOBP_NEGATIVE_REGULATION_OF_SIGNAL_TRANSDUCTION_IN_ABSENCE_O | 28 | -0.41224 | -1.009166 | 0.466903 | 0.6925 |
| GOBP_OVARIAN_FOLLICLE_DEVELOPMENT | 46 | -0.380501 | -1.009131 | 0.456667 | 0.692317 |
| GOBP_B_CELL_APOPTOTIC_PROCESS | 24 | -0.423789 | -1.008815 | 0.485346 | 0.692831 |
| GOBP_POSITIVE_REGULATION_OF_LIPID_TRANSPORT | 76 | -0.36183 | -1.007722 | 0.491803 | 0.695451 |
| GOBP_PEPTIDYL_ASPARAGINE_MODIFICATION | 31 | -0.403646 | -1.007658 | 0.486487 | 0.695313 |
| GOBP_MEMBRANE_PROTEIN_INTRACELLULAR_DOMAIN_PROTEOLYSIS | 20 | -0.43345 | -1.007481 | 0.499373 | 0.695502 |
| GOBP_REGULATION_OF_WNT_SIGNALING_PATHWAY_PLANAR_CELL_POLARIT | 15 | -0.460595 | -1.007439 | 0.498079 | 0.695315 |
| GOBP_NADP_METABOLIC_PROCESS | 34 | -0.398107 | -1.006059 | 0.501718 | 0.698744 |
| GOBP_POSITIVE_REGULATION_OF_CYTOSKELETON_ORGANIZATION | 212 | -0.345526 | -1.005898 | 0.490491 | 0.698869 |
| GOBP_PHOSPHOLIPID_DEPHOSPHORYLATION | 44 | -0.383479 | -1.005759 | 0.498874 | 0.698938 |
| GOBP_ALTERNATIVE_MRNA_SPLICING_VIA_SPLICEOSOME | 70 | -0.361865 | -1.005469 | 0.48666 | 0.699423 |
| GOBP_NEGATIVE_REGULATION_OF_JNK_CASCADE | 35 | -0.394451 | -1.004995 | 0.494904 | 0.700348 |
| GOBP_TRANSEPITHELIAL_TRANSPORT | 29 | -0.408402 | -1.00446 | 0.5053 | 0.70146 |
| GOBP_CARBOHYDRATE_HOMEOSTASIS | 201 | -0.343372 | -1.003785 | 0.511066 | 0.702932 |
| GOBP_GLUTAMINE_FAMILY_AMINO_ACID_METABOLIC_PROCESS | 71 | -0.364019 | -1.00347 | 0.490526 | 0.703472 |
| GOBP_OLEFINIC_COMPOUND_BIOSYNTHETIC_PROCESS | 19 | -0.430957 | -1.003093 | 0.490148 | 0.704159 |
| GOBP_VIRAL_LIFE_CYCLE | 331 | -0.338939 | -1.00299 | 0.496497 | 0.704134 |
| GOBP_HEPARAN_SULFATE_PROTEOGLYCAN_METABOLIC_PROCESS | 30 | -0.399841 | -1.002326 | 0.489461 | 0.705593 |
| GOBP_NEGATIVE_REGULATION_OF_ATPASE_ACTIVITY | 18 | -0.439794 | -1.002152 | 0.503798 | 0.705774 |
| GOBP_ORGANIC_ACID_CATABOLIC_PROCESS | 244 | -0.342381 | -1.001788 | 0.526527 | 0.706456 |
| GOBP_SULFUR_COMPOUND_BIOSYNTHETIC_PROCESS | 181 | -0.343565 | -1.001179 | 0.504514 | 0.707758 |
| GOBP_REGULATION_OF_ACTIN_FILAMENT_LENGTH | 179 | -0.348062 | -1.000974 | 0.514141 | 0.70801 |
| GOBP_EMBRYO_IMPLANTATION | 41 | -0.38654 | -1.000962 | 0.497792 | 0.707749 |
| GOBP_PLASMINOGEN_ACTIVATION | 18 | -0.435877 | -1.000412 | 0.481061 | 0.708975 |
| GOBP_CELLULAR_RESPONSE_TO_PEPTIDE | 356 | -0.338581 | -1.000071 | 0.519 | 0.70958 |
| GOBP_GLYCOSYLATION | 250 | -0.339729 | -1.000022 | 0.512 | 0.709412 |
| GOBP_SIGNAL_TRANSDUCTION_IN_ABSENCE_OF_LIGAND | 61 | -0.368574 | -0.999923 | 0.520474 | 0.70939 |
| GOBP_REGULATION_OF_EXOSOMAL_SECRETION | 16 | -0.451347 | -0.999893 | 0.485498 | 0.709173 |
| GOBP_MYELIN_MAINTENANCE | 16 | -0.442746 | -0.999886 | 0.484395 | 0.708896 |
| GOBP_RESPONSE_TO_LEPTIN | 22 | -0.425795 | -0.99972 | 0.488568 | 0.709047 |
| GOBP_POSITIVE_REGULATION_OF_ANION_TRANSPORT | 449 | -0.335846 | -0.999581 | 0.5 | 0.709132 |
| GOBP_LAMELLIPODIUM_ASSEMBLY | 67 | -0.36169 | -0.99947 | 0.488889 | 0.709139 |
| GOBP_VENTRICULAR_SEPTUM_MORPHOGENESIS | 35 | -0.3875 | -0.999014 | 0.472414 | 0.710038 |
| GOBP_BIOLOGICAL_PROCESS_INVOLVED_IN_INTERACTION_WITH_HOST | 204 | -0.340406 | -0.998837 | 0.502508 | 0.710204 |
| GOBP_PURINE_CONTAINING_COMPOUND_BIOSYNTHETIC_PROCESS | 201 | -0.342283 | -0.998116 | 0.515516 | 0.711813 |
| GOBP_ORGANOPHOSPHATE_ESTER_TRANSPORT | 118 | -0.351461 | -0.998004 | 0.521783 | 0.711824 |
| GOBP_REGULATION_OF_HEMATOPOIETIC_PROGENITOR_CELL_DIFFERENTIAT | 88 | -0.355577 | -0.997765 | 0.520747 | 0.712146 |
| GOBP_STEROID_HORMONE_SECRETION | 19 | -0.430014 | -0.997459 | 0.492519 | 0.712676 |
| GOBP_MICROTUBULE_DEPOLYMERIZATION | 43 | -0.385667 | -0.997214 | 0.490524 | 0.713012 |
| GOBP_REGULATION_OF_ALCOHOL_BIOSYNTHETIC_PROCESS | 67 | -0.365667 | -0.996683 | 0.507447 | 0.714112 |
| GOBP_RESPONSE_TO_SALT_STRESS | 23 | -0.417209 | -0.996438 | 0.497573 | 0.714457 |
| GOBP_POSITIVE_REGULATION_OF_CYTOKINESIS | 29 | -0.399789 | -0.996401 | 0.511521 | 0.714263 |
| GOBP_REGULATION_OF_EPITHELIAL_CELL_MIGRATION | 220 | -0.339952 | -0.996124 | 0.520603 | 0.714705 |
| GOBP_OOCYTE_DIFFERENTIATION | 42 | -0.382696 | -0.996077 | 0.503326 | 0.714532 |
| GOBP_RESPONSE_TO_PROSTAGLANDIN | 30 | -0.406866 | -0.995928 | 0.516129 | 0.714628 |
| GOBP_DIOL_METABOLIC_PROCESS | 29 | -0.404163 | -0.995819 | 0.513642 | 0.714619 |
| GOBP_VESICLE_LOCALIZATION | 223 | -0.340996 | -0.99561 | 0.512024 | 0.714883 |
| GOBP_RHYTHMIC_PROCESS | 268 | -0.338449 | -0.995459 | 0.537 | 0.714962 |
| GOBP_PRI_MIRNA_TRANSCRIPTION_BY_RNA_POLYMERASE_II | 51 | -0.375458 | -0.994413 | 0.489011 | 0.717389 |
| GOBP_SKIN_DEVELOPMENT | 249 | -0.339719 | -0.994249 | 0.523524 | 0.717529 |
| GOBP_RESPONSE_TO_HYDROGEN_PEROXIDE | 126 | -0.345371 | -0.993938 | 0.527495 | 0.718051 |
| GOBP_PRODUCTION_OF_SMALL_RNA_INVOLVED_IN_GENE_SILENCING_BY_R | 53 | -0.367624 | -0.992701 | 0.512035 | 0.721081 |

| | | | | | |
|---|---|---|---|---|---|
| GOBP_SYNAPTIC_VESICLE_LOCALIZATION | 54 | -0.370834 | -0.992426 | 0.504329 | 0.721528 |
| GOBP_RESPONSE_TO_PEPTIDE_HORMONE | 394 | -0.335749 | -0.990994 | 0.553 | 0.725007 |
| GOBP_CLATHRIN_DEPENDENT_ENDOCYTOSIS | 46 | -0.378594 | -0.990801 | 0.518072 | 0.72521 |
| GOBP_ACTIN_NUCLEATION | 51 | -0.368426 | -0.990731 | 0.503212 | 0.725098 |
| GOBP_POSITIVE_REGULATION_OF_INSULIN_SECRETION | 65 | -0.359973 | -0.989918 | 0.523355 | 0.726871 |
| GOBP_POSITIVE_REGULATION_OF_DENDRITE_MORPHOGENESIS | 35 | -0.385371 | -0.989812 | 0.497156 | 0.726845 |
| GOBP_INTESTINAL_EPITHELIAL_CELL_DIFFERENTIATION | 19 | -0.432983 | -0.989776 | 0.504414 | 0.726641 |
| GOBP_CARDIAC_RIGHT_VENTRICLE_MORPHOGENESIS | 19 | -0.426902 | -0.989479 | 0.502481 | 0.727134 |
| GOBP_ECTODERM_DEVELOPMENT | 18 | -0.429835 | -0.989414 | 0.516469 | 0.727001 |
| GOBP_REGULATION_OF_CELLULAR_AMINO_ACID_METABOLIC_PROCESS | 59 | -0.361524 | -0.9894 | 0.498929 | 0.726742 |
| GOBP_SPERM_EGG_RECOGNITION | 46 | -0.373898 | -0.989212 | 0.499455 | 0.726913 |
| GOBP_DRUG_METABOLIC_PROCESS | 31 | -0.39149 | -0.988367 | 0.522779 | 0.728831 |
| GOBP_WALKING_BEHAVIOR | 23 | -0.412122 | -0.988282 | 0.509501 | 0.728758 |
| GOBP_MODULATION_OF_PROCESS_OF_OTHER_ORGANISM_INVOLVED_IN_SY | 90 | -0.352005 | -0.987969 | 0.525773 | 0.729291 |
| GOBP_PURINE_NUCLEOSIDE_METABOLIC_PROCESS | 60 | -0.362246 | -0.987649 | 0.519108 | 0.729836 |
| GOBP_NEGATIVE_REGULATION_OF_INTRINSIC_APOPTOTIC_SIGNALING_PATH | 20 | -0.423011 | -0.987549 | 0.510204 | 0.729786 |
| GOBP_CELLULAR_RESPONSE_TO_STEROL | 20 | -0.425981 | -0.987513 | 0.505632 | 0.729586 |
| GOBP_C21_STEROID_HORMONE_METABOLIC_PROCESS | 33 | -0.387232 | -0.987336 | 0.504525 | 0.729755 |
| GOBP_HEPATICOBILIARY_SYSTEM_DEVELOPMENT | 133 | -0.343516 | -0.987251 | 0.554095 | 0.729665 |
| GOBP_REGULATION_OF_NUCLEOBASE_CONTAINING_COMPOUND_TRANSPOR | 15 | -0.437867 | -0.987165 | 0.519104 | 0.729582 |
| GOBP_CELLULAR_AMINO_ACID_METABOLIC_PROCESS | 320 | -0.335019 | -0.986847 | 0.559 | 0.730115 |
| GOBP_STEROID_HORMONE_BIOSYNTHETIC_PROCESS | 42 | -0.376735 | -0.986766 | 0.513483 | 0.730021 |
| GOBP_REGULATION_OF_BIOLOGICAL_PROCESS_INVOLVED_IN_SYMBIOTIC_IN | 184 | -0.339676 | -0.984915 | 0.534604 | 0.73456 |
| GOBP_NEGATIVE_REGULATION_OF_ANION_TRANSMEMBRANE_TRANSPORT | 31 | -0.402794 | -0.984612 | 0.507532 | 0.735043 |
| GOBP_SMALL_MOLECULE_CATABOLIC_PROCESS | 401 | -0.332736 | -0.984449 | 0.584 | 0.735158 |
| GOBP_EPITHELIAL_TUBE_FORMATION | 123 | -0.348393 | -0.983904 | 0.535642 | 0.736231 |
| GOBP_PROTEIN_LOCALIZATION_TO_PLASMA_MEMBRANE | 267 | -0.333192 | -0.98357 | 0.55956 | 0.736779 |
| GOBP_NUCLEUS_LOCALIZATION | 23 | -0.412694 | -0.983343 | 0.508537 | 0.737048 |
| GOBP_HEMATOPOIETIC_STEM_CELL_DIFFERENTIATION | 87 | -0.353244 | -0.982791 | 0.550052 | 0.738148 |
| GOBP_CELL_PROLIFERATION_IN_FOREBRAIN | 24 | -0.409 | -0.982692 | 0.507937 | 0.738105 |
| GOBP_MITOCHONDRIAL_CALCIUM_ION_HOMEOSTASIS | 26 | -0.403504 | -0.982428 | 0.513221 | 0.738477 |
| GOBP_EATING_BEHAVIOR | 21 | -0.415805 | -0.981872 | 0.501801 | 0.739573 |
| GOBP_REGULATION_OF_CLATHRIN_DEPENDENT_ENDOCYTOSIS | 19 | -0.425493 | -0.981584 | 0.52956 | 0.740013 |
| GOBP_LENS_MORPHOGENESIS_IN_CAMERA_TYPE_EYE | 18 | -0.424915 | -0.98147 | 0.51375 | 0.740005 |
| GOBP_POSITIVE_REGULATION_OF_LIPID_LOCALIZATION | 98 | -0.349054 | -0.981282 | 0.541882 | 0.74017 |
| GOBP_REGULATION_OF_ARP2_3_COMPLEX_MEDIATED_ACTIN_NUCLEATION | 21 | -0.415355 | -0.981212 | 0.504926 | 0.740047 |
| GOBP_NEGATIVE_REGULATION_OF_CALCIUM_MEDIATED_SIGNALING | 18 | -0.433537 | -0.981117 | 0.519695 | 0.74 |
| GOBP_CELLULAR_MODIFIED_AMINO_ACID_METABOLIC_PROCESS | 180 | -0.335336 | -0.980783 | 0.565962 | 0.740557 |
| GOBP_RESPONSE_TO_EPIDERMAL_GROWTH_FACTOR | 45 | -0.368181 | -0.980166 | 0.534807 | 0.741832 |
| GOBP_NEUROMUSCULAR_SYNAPTIC_TRANSMISSION | 21 | -0.421713 | -0.980076 | 0.503001 | 0.741764 |
| GOBP_VESICLE_CYTOSKELETAL_TRAFFICKING | 71 | -0.349705 | -0.979618 | 0.524416 | 0.742615 |
| GOBP_EPITHELIAL_CELL_MORPHOGENESIS | 30 | -0.391286 | -0.979145 | 0.516052 | 0.743465 |
| GOBP_PIGMENT_METABOLIC_PROCESS | 66 | -0.355377 | -0.978873 | 0.535525 | 0.743896 |
| GOBP_REGULATION_OF_PROTEIN_ACETYLATION | 72 | -0.355312 | -0.978648 | 0.572176 | 0.744179 |
| GOBP_CELLULAR_COMPONENT_MAINTENANCE | 60 | -0.365943 | -0.976339 | 0.559742 | 0.749734 |
| GOBP_INOSITOL_PHOSPHATE_MEDIATED_SIGNALING | 49 | -0.370135 | -0.975934 | 0.540393 | 0.750458 |
| GOBP_REGULATION_OF_APOPTOTIC_SIGNALING_PATHWAY | 323 | -0.329128 | -0.975648 | 0.594 | 0.750906 |
| GOBP_MOLTING_CYCLE | 89 | -0.345338 | -0.975174 | 0.558115 | 0.7518 |
| GOBP_COPPER_ION_HOMEOSTASIS | 16 | -0.439135 | -0.975155 | 0.525 | 0.751546 |
| GOBP_LIPID_MODIFICATION | 252 | -0.334235 | -0.97514 | 0.587174 | 0.751278 |
| GOBP_RESPONSE_TO_CARBOHYDRATE | 200 | -0.33464 | -0.974582 | 0.578313 | 0.752434 |
| GOBP_POSITIVE_REGULATION_OF_CANONICAL_WNT_SIGNALING_PATHWAY | 140 | -0.337546 | -0.974505 | 0.55499 | 0.752319 |
| GOBP_REGULATION_OF_ACTOMYOSIN_STRUCTURE_ORGANIZATION | 91 | -0.349968 | -0.974082 | 0.560496 | 0.753056 |
| GOBP_NEGATIVE_REGULATION_OF_ERK1_AND_ERK2_CASCADE | 67 | -0.351967 | -0.973608 | 0.543039 | 0.753961 |
| GOBP_PROTEIN_ADP_RIBOSYLATION | 34 | -0.385821 | -0.973605 | 0.531866 | 0.753667 |
| GOBP_CELLULAR_RESPONSE_TO_PH | 20 | -0.423459 | -0.973574 | 0.529412 | 0.753442 |
| GOBP_POSITIVE_REGULATION_OF_PROTEIN_CATABOLIC_PROCESS | 221 | -0.332114 | -0.973336 | 0.595787 | 0.75373 |
| GOBP_POSITIVE_REGULATION_OF_PRI_MIRNA_TRANSCRIPTION_BY_RNA_POL | 39 | -0.373338 | -0.972585 | 0.529345 | 0.755318 |
| GOBP_REGULATION_OF_ATP_METABOLIC_PROCESS | 113 | -0.339254 | -0.97226 | 0.551935 | 0.755843 |
| GOBP_NEGATIVE_REGULATION_OF_SMOOTHENED_SIGNALING_PATHWAY | 31 | -0.388489 | -0.972182 | 0.544601 | 0.755758 |
| GOBP_REGULATION_OF_INTRINSIC_APOPTOTIC_SIGNALING_PATHWAY_BY_P5 | 27 | -0.397853 | -0.972087 | 0.537545 | 0.755684 |
| GOBP_POSITIVE_REGULATION_OF_PROTEIN_POLYMERIZATION | 127 | -0.338529 | -0.97192 | 0.57332 | 0.755787 |
| GOBP_ESTABLISHMENT_OF_ORGANELLE_LOCALIZATION | 419 | -0.326923 | -0.971476 | 0.617 | 0.756572 |
| GOBP_INTRINSIC_APOPTOTIC_SIGNALING_PATHWAY_IN_RESPONSE_TO_DNA | 100 | -0.345118 | -0.971456 | 0.561421 | 0.756313 |
| GOBP_REGULATION_OF_CELL_SUBSTRATE_JUNCTION_ORGANIZATION | 68 | -0.354623 | -0.971386 | 0.5625 | 0.756187 |
| GOBP_POSITIVE_REGULATION_OF_INTRACELLULAR_PROTEIN_TRANSPORT | 178 | -0.33399 | -0.971095 | 0.583249 | 0.756613 |
| GOBP_REGULATION_OF_CELL_MORPHOGENESIS_INVOLVED_IN_DIFFERENTIA | 91 | -0.342154 | -0.971057 | 0.54068 | 0.756409 |
| GOBP_PTERIDINE_CONTAINING_COMPOUND_METABOLIC_PROCESS | 33 | -0.385812 | -0.970964 | 0.536697 | 0.756333 |
| GOBP_REGULATION_OF_CELL_GROWTH_INVOLVED_IN_CARDIAC_MUSCLE_C | 17 | -0.427813 | -0.970636 | 0.513924 | 0.756869 |
| GOBP_NEGATIVE_REGULATION_OF_LIPID_STORAGE | 20 | -0.414589 | -0.970532 | 0.524814 | 0.756818 |
| GOBP_PYRIMIDINE_NUCLEOTIDE_METABOLIC_PROCESS | 49 | -0.365787 | -0.969245 | 0.571584 | 0.759747 |
| GOBP_RESPONSE_TO_HYDROPEROXIDE | 16 | -0.430875 | -0.968942 | 0.545006 | 0.760198 |
| GOBP_NUCLEOTIDE_PHOSPHORYLATION | 127 | -0.335115 | -0.968928 | 0.568667 | 0.759933 |
| GOBP_REGULATION_OF_EXTRINSIC_APOPTOTIC_SIGNALING_PATHWAY_IN_AE | 37 | -0.374241 | -0.968718 | 0.533784 | 0.760152 |
| GOBP_FIBROBLAST_MIGRATION | 43 | -0.369938 | -0.968204 | 0.538462 | 0.761112 |
| GOBP_BASIC_AMINO_ACID_TRANSPORT | 18 | -0.424742 | -0.96761 | 0.5399 | 0.762296 |
| GOBP_POSITIVE_REGULATION_OF_INTRACELLULAR_TRANSPORT | 215 | -0.330745 | -0.967156 | 0.597598 | 0.763145 |
| GOBP_INSULIN_SECRETION_INVOLVED_IN_CELLULAR_RESPONSE_TO_GLUC | 61 | -0.358483 | -0.967076 | 0.556854 | 0.763037 |
| GOBP_HORMONE_MEDIATED_SIGNALING_PATHWAY | 171 | -0.333999 | -0.966539 | 0.574874 | 0.764035 |
| GOBP_SCHWANN_CELL_DEVELOPMENT | 28 | -0.394567 | -0.966376 | 0.540888 | 0.764132 |
| GOBP_BONE_GROWTH | 25 | -0.399507 | -0.966318 | 0.542654 | 0.763966 |
| GOBP_REGULATION_OF_CYTOPLASMIC_TRANSPORT | 27 | -0.39683 | -0.966282 | 0.533333 | 0.763754 |
| GOBP_NEGATIVE_REGULATION_OF_AXON_EXTENSION | 43 | -0.368964 | -0.965325 | 0.536913 | 0.765736 |
| GOBP_POSITIVE_REGULATION_OF_CATABOLIC_PROCESS | 432 | -0.324192 | -0.965018 | 0.651 | 0.766174 |
| GOBP_OOCYTE_MATURATION | 22 | -0.412817 | -0.964799 | 0.544803 | 0.766416 |
| GOBP_CELLULAR_RESPONSE_TO_OSMOTIC_STRESS | 39 | -0.374325 | -0.964493 | 0.537683 | 0.766863 |
| GOBP_FATTY_ACID_DERIVATIVE_METABOLIC_PROCESS | 70 | -0.350412 | -0.962489 | 0.563365 | 0.771557 |
| GOBP_TRABECULA_FORMATION | 19 | -0.418467 | -0.962135 | 0.55985 | 0.772121 |
| GOBP_RESPONSE_TO_VITAMIN_A | 17 | -0.427977 | -0.962008 | 0.525721 | 0.77213 |
| GOBP_NEGATIVE_REGULATION_OF_NUCLEOCYTOPLASMIC_TRANSPORT | 20 | -0.414475 | -0.961911 | 0.541353 | 0.772051 |
| GOBP_STEM_CELL_PROLIFERATION | 58 | -0.356488 | -0.961246 | 0.560299 | 0.773384 |
| GOBP_PROTEIN_POLYMERIZATION | 282 | -0.327809 | -0.961189 | 0.62963 | 0.77322 |
| GOBP_PROTEASOMAL_UBIQUITIN_INDEPENDENT_PROTEIN_CATABOLIC_PRO | 21 | -0.410996 | -0.960492 | 0.560644 | 0.774604 |
| GOBP_LIPID_CATABOLIC_PROCESS | 299 | -0.326631 | -0.960446 | 0.63964 | 0.774403 |
| GOBP_REGULATION_OF_PROGRAMMED_NECROTIC_CELL_DEATH | 22 | -0.406619 | -0.960308 | 0.553903 | 0.774434 |
| GOBP_FOAM_CELL_DIFFERENTIATION | 32 | -0.395121 | -0.960106 | 0.548951 | 0.774623 |
| GOBP_POSITIVE_REGULATION_OF_MUSCLE_CELL_APOPTOTIC_PROCESS | 22 | -0.404901 | -0.959906 | 0.53644 | 0.774833 |
| GOBP_REGULATION_OF_PROTEIN_POLYMERIZATION | 216 | -0.329312 | -0.959315 | 0.61005 | 0.775947 |
| GOBP_REGULATION_OF_PLATELET_AGGREGATION | 18 | -0.424072 | -0.958802 | 0.531888 | 0.776904 |
| GOBP_REGULATION_OF_STEROID_METABOLIC_PROCESS | 104 | -0.336351 | -0.958533 | 0.590299 | 0.77726 |
| GOBP_INNATE_IMMUNE_RESPONSE_ACTIVATING_SIGNAL_TRANSDUCTION | 108 | -0.339197 | -0.958524 | 0.609907 | 0.776974 |
| GOBP_ENERGY_RESERVE_METABOLIC_PROCESS | 79 | -0.345503 | -0.958204 | 0.589583 | 0.777423 |
| GOBP_SYNAPTIC_VESICLE_PRIMING | 19 | -0.41568 | -0.956484 | 0.55886 | 0.78133 |
| GOBP_NEGATIVE_REGULATION_OF_OXIDATIVE_STRESS_INDUCED_NEURON_ | 15 | -0.430921 | -0.956153 | 0.564168 | 0.781841 |
| GOBP_SECRETION_BY_TISSUE | 28 | -0.390781 | -0.956114 | 0.531585 | 0.781629 |
| GOBP_PEROXISOMAL_MEMBRANE_TRANSPORT | 20 | -0.407333 | -0.956066 | 0.540441 | 0.78144 |
| GOBP_EPIDERMAL_CELL_DIFFERENTIATION | 194 | -0.328213 | -0.955665 | 0.620342 | 0.782062 |
| GOBP_POSITIVE_REGULATION_OF_CALCIUM_ION_IMPORT | 17 | -0.425195 | -0.954896 | 0.552296 | 0.783604 |
| GOBP_DIOL_BIOSYNTHETIC_PROCESS | 22 | -0.406778 | -0.95474 | 0.558442 | 0.783669 |
| GOBP_ORGANIC_HYDROXY_COMPOUND_BIOSYNTHETIC_PROCESS | 230 | -0.325636 | -0.954221 | 0.641566 | 0.784599 |
| GOBP_INTERLEUKIN_5_PRODUCTION | 15 | -0.436422 | -0.953979 | 0.555698 | 0.784877 |
| GOBP_NEGATIVE_REGULATION_OF_PROTEIN_LOCALIZATION_TO_NUCLEUS | 29 | -0.390588 | -0.952341 | 0.555948 | 0.788512 |
| GOBP_PROTEIN_MANNOSYLATION | 23 | -0.400668 | -0.9513 | 0.556638 | 0.79072 |
| GOBP_NEGATIVE_REGULATION_OF_ORGAN_GROWTH | 31 | -0.381142 | -0.950908 | 0.574661 | 0.791348 |
| GOBP_STEROL_HOMEOSTASIS | 74 | -0.345691 | -0.950588 | 0.585185 | 0.7918 |
| GOBP_REGULATION_OF_HEMATOPOIETIC_STEM_CELL_DIFFERENTIATION | 73 | -0.348337 | -0.950135 | 0.582983 | 0.792566 |
| GOBP_ESTABLISHMENT_OF_ENDOTHELIAL_BARRIER | 48 | -0.359457 | -0.950118 | 0.591009 | 0.792292 |
| GOBP_SERINE_FAMILY_AMINO_ACID_METABOLIC_PROCESS | 41 | -0.367314 | -0.950074 | 0.569061 | 0.792093 |
| GOBP_POSITIVE_REGULATION_OF_SODIUM_ION_TRANSMEMBRANE_TRANSP | 21 | -0.408271 | -0.950028 | 0.556501 | 0.791887 |
| GOBP_ACTIN_FILAMENT_DEPOLYMERIZATION | 56 | -0.351552 | -0.949764 | 0.577049 | 0.792234 |
| GOBP_AMINO_ACID_BETAINE_METABOLIC_PROCESS | 17 | -0.415993 | -0.94964 | 0.572716 | 0.792233 |
| GOBP_NEGATIVE_REGULATION_OF_PEPTIDYL_TYROSINE_PHOSPHORYLATION | 54 | -0.351653 | -0.949526 | 0.577297 | 0.792173 |
| GOBP_REGULATION_OF_MITOCHONDRIAL_FISSION | 27 | -0.391174 | -0.949188 | 0.571765 | 0.792675 |
| GOBP_POSITIVE_REGULATION_OF_ESTABLISHMENT_OF_PROTEIN_LOCALIZAT | 316 | -0.321311 | -0.948764 | 0.685371 | 0.793386 |
| GOBP_CELLULAR_COMPONENT_DISASSEMBLY_INVOLVED_IN_EXECUTION_PH | 34 | -0.371377 | -0.948669 | 0.558559 | 0.79332 |
| GOBP_REGULATION_OF_FIBROBLAST_MIGRATION | 34 | -0.370183 | -0.948066 | 0.564965 | 0.794426 |
| GOBP_INSULIN_RECEPTOR_SIGNALING_PATHWAY | 136 | -0.325167 | -0.947814 | 0.629442 | 0.794707 |
| GOBP_POSITIVE_REGULATION_OF_PROTEIN_DEACETYLATION | 22 | -0.400213 | -0.946822 | 0.579832 | 0.796762 |
| GOBP_REGULATION_OF_EXTRACELLULAR_MATRIX_DISASSEMBLY | 16 | -0.419428 | -0.946403 | 0.593242 | 0.797461 |
| GOBP_RESPONSE_TO_MONOSACCHARIDE | 177 | -0.324319 | -0.946295 | 0.632899 | 0.79739 |
| GOBP_PEPTIDYL_THREONINE_MODIFICATION | 120 | -0.330047 | -0.946291 | 0.617828 | 0.797087 |
| GOBP_MYOFIBRIL_ASSEMBLY | 51 | -0.350715 | -0.946044 | 0.574353 | 0.797366 |
| GOBP_RESPONSE_TO_PROSTAGLANDIN_E | 23 | -0.392276 | -0.945739 | 0.557178 | 0.797777 |
| GOBP_REGULATION_OF_MICROTUBULE_DEPOLYMERIZATION | 29 | -0.379146 | -0.945575 | 0.576429 | 0.797868 |
| GOBP_VACUOLE_ORGANIZATION | 171 | -0.326224 | -0.94545 | 0.654894 | 0.797836 |
| GOBP_PHOSPHATIDYLETHANOLAMINE_ACYL_CHAIN_REMODELING | 22 | -0.397941 | -0.945277 | 0.57037 | 0.797938 |
| GOBP_REGULATION_OF_INTRINSIC_APOPTOTIC_SIGNALING_PATHWAY_IN_RE | 37 | -0.363791 | -0.944836 | 0.553167 | 0.79868 |
| GOBP_INTERMEDIATE_FILAMENT_ORGANIZATION | 20 | -0.403939 | -0.94443 | 0.544118 | 0.799327 |
| GOBP_INTERFERON_BETA_PRODUCTION | 52 | -0.351476 | -0.943232 | 0.577706 | 0.801813 |
| GOBP_REPLICATIVE_SENESCENCE | 16 | -0.418955 | -0.942764 | 0.579214 | 0.802604 |
| GOBP_CELLULAR_GLUCOSE_HOMEOSTASIS | 131 | -0.329906 | -0.942512 | 0.637462 | 0.802899 |
| GOBP_RESPONSE_TO_IMMOBILIZATION_STRESS | 21 | -0.404909 | -0.942381 | 0.56747 | 0.802895 |
| GOBP_HEMATOPOIETIC_STEM_CELL_HOMEOSTASIS | 15 | -0.430082 | -0.942268 | 0.565161 | 0.802837 |
| GOBP_SYNAPTIC_VESICLE_TRANSPORT | 43 | -0.359702 | -0.941675 | 0.577481 | 0.803893 |
| GOBP_PHOSPHORYLATED_CARBOHYDRATE_DEPHOSPHORYLATION | 16 | -0.424997 | -0.941436 | 0.581515 | 0.80413 |
| GOBP_NEGATIVE_REGULATION_OF_INTRACELLULAR_SIGNAL_TRANSDUCTION | 468 | -0.316632 | -0.94137 | 0.7 | 0.803972 |
| GOBP_POLY_A_PLUS_MRNA_EXPORT_FROM_NUCLEUS | 19 | -0.409891 | -0.940765 | 0.554601 | 0.805094 |
| GOBP_RESPONSE_TO_ACTIVITY | 63 | -0.347042 | -0.940675 | 0.609375 | 0.804984 |
| GOBP_REGULATION_OF_SMAD_PROTEIN_SIGNAL_TRANSDUCTION | 24 | -0.393122 | -0.939618 | 0.586826 | 0.807159 |
| GOBP_NEGATIVE_REGULATION_OF_I_KAPPAB_KINASE_NF_KAPPAB_SIGNALIN | 43 | -0.361268 | -0.939582 | 0.583333 | 0.806921 |
| GOBP_PROTEIN_PEPTIDYL_PROLYL_ISOMERIZATION | 37 | -0.370638 | -0.938699 | 0.610419 | 0.808673 |

| | | | | | |
|---|---|---|---|---|---|
| GOBP_INNATE_IMMUNE_RESPONSE_IN_MUCOSA | 18 | -0.410046 | -0.938428 | 0.578616 | 0.808969 |
| GOBP_REGULATION_OF_PROTEIN_IMPORT | 58 | -0.345717 | -0.938394 | 0.585653 | 0.808733 |
| GOBP_REGULATION_OF_PROTEIN_TARGETING | 80 | -0.335595 | -0.93823 | 0.628931 | 0.808792 |
| GOBP_CELLULAR_RESPONSE_TO_PEPTIDE_HORMONE_STIMULUS | 295 | -0.317768 | -0.937375 | 0.705706 | 0.810461 |
| GOBP_REGULATION_OF_ATP_BIOSYNTHETIC_PROCESS | 20 | -0.404423 | -0.93726 | 0.579657 | 0.810395 |
| GOBP_C21_STEROID_HORMONE_BIOSYNTHETIC_PROCESS | 23 | -0.393944 | -0.936533 | 0.582933 | 0.81172 |
| GOBP_RIBONUCLEOSIDE_METABOLIC_PROCESS | 69 | -0.34628 | -0.93642 | 0.611345 | 0.81168 |
| GOBP_ENDOPLASMIC_RETICULUM_TUBULAR_NETWORK_ORGANIZATION | 18 | -0.400208 | -0.935515 | 0.586381 | 0.813438 |
| GOBP_EMBRYONIC_PLACENTA_DEVELOPMENT | 74 | -0.340227 | -0.935041 | 0.622642 | 0.814224 |
| GOBP_PROTEIN_DEMANNOSYLATION | 20 | -0.398406 | -0.934657 | 0.569007 | 0.814799 |
| GOBP_REGULATION_OF_AUTOPHAGY | 323 | -0.315686 | -0.934458 | 0.707 | 0.81494 |
| GOBP_FOCAL_ADHESION_ASSEMBLY | 84 | -0.33358 | -0.934292 | 0.630753 | 0.815006 |
| GOBP_INTRINSIC_APOPTOTIC_SIGNALING_PATHWAY | 270 | -0.317935 | -0.934055 | 0.707 | 0.815242 |
| GOBP_NEGATIVE_REGULATION_OF_CYTOSKELETON_ORGANIZATION | 148 | -0.32339 | -0.933966 | 0.640324 | 0.815129 |
| GOBP_MULTICELLULAR_ORGANISM_AGING | 28 | -0.378718 | -0.933828 | 0.576471 | 0.815142 |
| GOBP_PRIMARY_NEURAL_TUBE_FORMATION | 87 | -0.330757 | -0.933285 | 0.633265 | 0.816086 |
| GOBP_RESPONSE_TO_GONADOTROPIN | 20 | -0.400605 | -0.932534 | 0.590571 | 0.817449 |
| GOBP_NEGATIVE_REGULATION_OF_MUSCLE_HYPERTROPHY | 27 | -0.377916 | -0.932463 | 0.586931 | 0.817297 |
| GOBP_REGULATION_OF_PROTEIN_EXPORT_FROM_NUCLEUS | 32 | -0.371554 | -0.932375 | 0.603624 | 0.817185 |
| GOBP_MAMMARY_GLAND_EPITHELIAL_CELL_PROLIFERATION | 24 | -0.387713 | -0.931634 | 0.600239 | 0.818572 |
| GOBP_NUCLEOSIDE_METABOLIC_PROCESS | 100 | -0.331941 | -0.930884 | 0.649177 | 0.81996 |
| GOBP_CELL_SUBSTRATE_JUNCTION_ORGANIZATION | 105 | -0.325558 | -0.93077 | 0.639796 | 0.819892 |
| GOBP_NEGATIVE_REGULATION_OF_LIPASE_ACTIVITY | 15 | -0.422885 | -0.93046 | 0.578608 | 0.820274 |
| GOBP_APICAL_PROTEIN_LOCALIZATION | 15 | -0.419773 | -0.92978 | 0.590331 | 0.821478 |
| GOBP_HYALURONAN_CATABOLIC_PROCESS | 16 | -0.42005 | -0.929223 | 0.59542 | 0.822429 |
| GOBP_REGULATION_OF_FATTY_ACID_OXIDATION | 31 | -0.373755 | -0.928958 | 0.606729 | 0.82271 |
| GOBP_REGULATION_OF_ERBB_SIGNALING_PATHWAY | 90 | -0.329153 | -0.928797 | 0.628218 | 0.822758 |
| GOBP_CALCINEURIN_MEDIATED_SIGNALING | 41 | -0.355024 | -0.928542 | 0.590656 | 0.82301 |
| GOBP_RIG_I_SIGNALING_PATHWAY | 23 | -0.394306 | -0.927898 | 0.60628 | 0.824129 |
| GOBP_REGULATION_OF_POLYSACCHARIDE_METABOLIC_PROCESS | 42 | -0.351529 | -0.927628 | 0.606061 | 0.824427 |
| GOBP_ACYLGLYCEROL_HOMEOSTASIS | 27 | -0.382995 | -0.926348 | 0.587958 | 0.82694 |
| GOBP_REGULATION_OF_SUPRAMOLECULAR_FIBER_ORGANIZATION | 353 | -0.312819 | -0.926166 | 0.747 | 0.827004 |
| GOBP_CIRCADIAN_RHYTHM | 194 | -0.320241 | -0.925889 | 0.716012 | 0.827309 |
| GOBP_POSITIVE_REGULATION_OF_TRANSCRIPTION_FROM_RNA_POLYMERAS | 21 | -0.397553 | -0.925652 | 0.609816 | 0.827507 |
| GOBP_STEROID_METABOLIC_PROCESS | 277 | -0.315366 | -0.925365 | 0.724 | 0.827834 |
| GOBP_VESICLE_MEDIATED_TRANSPORT_BETWEEN_ENDOSOMAL_COMPARTM | 43 | -0.350778 | -0.925122 | 0.603583 | 0.828058 |
| GOBP_POLYSACCHARIDE_METABOLIC_PROCESS | 103 | -0.326883 | -0.92512 | 0.651235 | 0.827747 |
| GOBP_PURINE_CONTAINING_COMPOUND_METABOLIC_PROCESS | 433 | -0.313045 | -0.925072 | 0.779 | 0.827525 |
| GOBP_AORTIC_VALVE_MORPHOGENESIS | 30 | -0.37189 | -0.924983 | 0.607143 | 0.827399 |
| GOBP_STORE_OPERATED_CALCIUM_ENTRY | 18 | -0.407268 | -0.924806 | 0.576208 | 0.827472 |
| GOBP_ORGANIC_ACID_BIOSYNTHETIC_PROCESS | 294 | -0.31486 | -0.924383 | 0.741 | 0.828081 |
| GOBP_ETHANOL_METABOLIC_PROCESS | 16 | -0.413319 | -0.924371 | 0.589552 | 0.827793 |
| GOBP_LONG_CHAIN_FATTY_ACYL_COA_METABOLIC_PROCESS | 25 | -0.38166 | -0.924081 | 0.584994 | 0.828123 |
| GOBP_REGULATION_OF_CELLULAR_CARBOHYDRATE_METABOLIC_PROCESS | 132 | -0.319518 | -0.923323 | 0.676441 | 0.829509 |
| GOBP_HYALURONAN_METABOLIC_PROCESS | 37 | -0.359783 | -0.922216 | 0.597082 | 0.831623 |
| GOBP_CARDIAC_EPITHELIAL_TO_MESENCHYMAL_TRANSITION | 30 | -0.372899 | -0.921835 | 0.608545 | 0.832136 |
| GOBP_NEGATIVE_REGULATION_OF_SUPRAMOLECULAR_FIBER_ORGANIZATIO | 149 | -0.316343 | -0.921049 | 0.69254 | 0.833572 |
| GOBP_TRIGLYCERIDE_METABOLIC_PROCESS | 88 | -0.325002 | -0.920706 | 0.66114 | 0.834039 |
| GOBP_RESPONSE_TO_ANTIBIOTIC | 43 | -0.349363 | -0.920273 | 0.61978 | 0.834669 |
| GOBP_RESPONSE_TO_XENOBIOTIC_STIMULUS | 101 | -0.325332 | -0.920241 | 0.681633 | 0.834418 |
| GOBP_CELLULAR_RESPONSE_TO_CARBOHYDRATE_STIMULUS | 125 | -0.318531 | -0.920162 | 0.678173 | 0.834263 |
| GOBP_PLACENTA_DEVELOPMENT | 125 | -0.322197 | -0.919912 | 0.677126 | 0.834505 |
| GOBP_BLASTOCYST_GROWTH | 19 | -0.399851 | -0.919613 | 0.598788 | 0.834863 |
| GOBP_KERATINOCYTE_DIFFERENTIATION | 141 | -0.31876 | -0.919556 | 0.679798 | 0.834659 |
| GOBP_PROTEIN_DEPOLYMERIZATION | 108 | -0.323055 | -0.919334 | 0.669036 | 0.834829 |
| GOBP_PURINE_CONTAINING_COMPOUND_CATABOLIC_PROCESS | 49 | -0.348177 | -0.919305 | 0.629956 | 0.834576 |
| GOBP_MATERNAL_PROCESS_INVOLVED_IN_FEMALE_PREGNANCY | 51 | -0.345238 | -0.918027 | 0.630077 | 0.837083 |
| GOBP_ACETYL_COA_BIOSYNTHETIC_PROCESS | 21 | -0.388887 | -0.917782 | 0.609223 | 0.837314 |
| GOBP_REGULATION_OF_CYTOPLASMIC_TRANSLATION | 19 | -0.39945 | -0.916989 | 0.605425 | 0.838711 |
| GOBP_PYRUVATE_METABOLIC_PROCESS | 143 | -0.318843 | -0.91683 | 0.68113 | 0.838736 |
| GOBP_NEGATIVE_REGULATION_OF_CELL_SUBSTRATE_JUNCTION_ORGANIZA | 18 | -0.40946 | -0.916645 | 0.616834 | 0.838833 |
| GOBP_REGULATION_OF_LAMELLIPODIUM_ORGANIZATION | 48 | -0.348533 | -0.916532 | 0.633588 | 0.838763 |
| GOBP_POLARIZED_EPITHELIAL_CELL_DIFFERENTIATION | 22 | -0.386236 | -0.916394 | 0.603774 | 0.838743 |
| GOBP_POSITIVE_REGULATION_OF_EXTRACELLULAR_MATRIX_ORGANIZATION | 23 | -0.385612 | -0.91611 | 0.612559 | 0.839043 |
| GOBP_LAMELLIPODIUM_ORGANIZATION | 84 | -0.325872 | -0.915977 | 0.645898 | 0.83902 |
| GOBP_POSITIVE_REGULATION_OF_G1_S_TRANSITION_OF_MITOTIC_CELL_CYC | 34 | -0.361906 | -0.915964 | 0.631046 | 0.838734 |
| GOBP_FATTY_ACID_BIOSYNTHETIC_PROCESS | 143 | -0.317743 | -0.915528 | 0.690816 | 0.839384 |
| GOBP_EXTRACELLULAR_VESICLE_BIOGENESIS | 22 | -0.383721 | -0.915362 | 0.604043 | 0.839427 |
| GOBP_EXECUTION_PHASE_OF_APOPTOSIS | 88 | -0.327655 | -0.915012 | 0.666667 | 0.839857 |
| GOBP_VASCULOGENESIS | 73 | -0.331853 | -0.914971 | 0.638831 | 0.839628 |
| GOBP_NUCLEOSIDE_PHOSPHATE_BIOSYNTHETIC_PROCESS | 257 | -0.313302 | -0.914883 | 0.750251 | 0.839521 |
| GOBP_REGULATION_OF_RECEPTOR_RECYCLING | 23 | -0.387041 | -0.914344 | 0.608059 | 0.840396 |
| GOBP_GLYCOSYL_COMPOUND_METABOLIC_PROCESS | 124 | -0.317031 | -0.914296 | 0.673449 | 0.840188 |
| GOBP_POSITIVE_REGULATION_OF_RESPONSE_TO_DNA_DAMAGE_STIMULUS | 105 | -0.321071 | -0.913462 | 0.681263 | 0.841693 |
| GOBP_WATER_SOLUBLE_VITAMIN_METABOLIC_PROCESS | 81 | -0.324456 | -0.913446 | 0.656904 | 0.841411 |
| GOBP_REGULATION_OF_LYSOSOMAL_LUMEN_PH | 15 | -0.418545 | -0.912928 | 0.626599 | 0.842196 |
| GOBP_MONOCARBOXYLIC_ACID_BIOSYNTHETIC_PROCESS | 194 | -0.316458 | -0.912828 | 0.718593 | 0.842117 |
| GOBP_REGULATION_OF_CARBOHYDRATE_METABOLIC_PROCESS | 193 | -0.313579 | -0.912627 | 0.715431 | 0.842221 |
| GOBP_POSITIVE_REGULATION_OF_MACROPHAGE_DERIVED_FOAM_CELL_DIF | 15 | -0.408748 | -0.91194 | 0.603234 | 0.843307 |
| GOBP_NEGATIVE_REGULATION_OF_PLASMA_MEMBRANE_BOUNDED_CELL_P | 28 | -0.374254 | -0.911042 | 0.630787 | 0.844874 |
| GOBP_FATTY_ACID_DERIVATIVE_BIOSYNTHETIC_PROCESS | 53 | -0.342475 | -0.911032 | 0.641593 | 0.844575 |
| GOBP_INTRACELLULAR_LIPID_TRANSPORT | 43 | -0.34805 | -0.910736 | 0.632222 | 0.844883 |
| GOBP_POSITIVE_REGULATION_OF_BINDING | 163 | -0.316016 | -0.909558 | 0.70101 | 0.8471 |
| GOBP_REGULATION_OF_SYNAPTIC_VESICLE_RECYCLING | 23 | -0.379498 | -0.909386 | 0.619835 | 0.847135 |
| GOBP_ACID_SECRETION | 34 | -0.358494 | -0.908878 | 0.625142 | 0.84791 |
| GOBP_PHOSPHATIDYLINOSITOL_DEPHOSPHORYLATION | 30 | -0.368382 | -0.908402 | 0.640371 | 0.848563 |
| GOBP_NOTOCHORD_DEVELOPMENT | 17 | -0.398779 | -0.908084 | 0.622307 | 0.848908 |
| GOBP_REGULATION_OF_VASCULAR_ASSOCIATED_SMOOTH_MUSCLE_CELL_[ | 16 | -0.40356 | -0.907784 | 0.616834 | 0.84924 |
| GOBP_NEGATIVE_REGULATION_OF_AUTOPHAGY | 81 | -0.324833 | -0.907737 | 0.669095 | 0.849023 |
| GOBP_REGULATION_OF_NUCLEOCYTOPLASMIC_TRANSPORT | 102 | -0.321615 | -0.907542 | 0.682702 | 0.849122 |
| GOBP_REGULATION_OF_GENERATION_OF_PRECURSOR_METABOLITES_AND_ | 152 | -0.31509 | -0.907482 | 0.719598 | 0.848934 |
| GOBP_REGULATION_OF_FEEDING_BEHAVIOR | 18 | -0.400313 | -0.907004 | 0.619221 | 0.849607 |
| GOBP_CELL_JUNCTION_MAINTENANCE | 32 | -0.363105 | -0.906482 | 0.622066 | 0.850419 |
| GOBP_SULFUR_COMPOUND_METABOLIC_PROCESS | 353 | -0.30524 | -0.906327 | 0.778 | 0.850423 |
| GOBP_REGULATION_OF_VIRAL_INDUCED_CYTOPLASMIC_PATTERN_RECOGN | 20 | -0.389604 | -0.905833 | 0.631835 | 0.851137 |
| GOBP_CELLULAR_RESPONSE_TO_NUTRIENT | 39 | -0.35063 | -0.905793 | 0.635864 | 0.850904 |
| GOBP_PLATELET_DERIVED_GROWTH_FACTOR_RECEPTOR_SIGNALING_PATH | 54 | -0.338125 | -0.905791 | 0.644397 | 0.85059 |
| GOBP_NEGATIVE_REGULATION_OF_INTRACELLULAR_PROTEIN_TRANSPORT | 39 | -0.354052 | -0.905734 | 0.638158 | 0.850392 |
| GOBP_CARBOHYDRATE_PHOSPHORYLATION | 24 | -0.375979 | -0.90543 | 0.628329 | 0.850702 |
| GOBP_REGULATION_OF_MRNA_SPLICING_VIA_SPLICEOSOME | 96 | -0.322454 | -0.90472 | 0.698969 | 0.851856 |
| GOBP_REGULATION_OF_ORGANELLE_ASSEMBLY | 183 | -0.310508 | -0.904683 | 0.729648 | 0.851613 |
| GOBP_POLYOL_CATABOLIC_PROCESS | 25 | -0.374259 | -0.90467 | 0.63981 | 0.851328 |
| GOBP_NUCLEOSIDE_SALVAGE | 16 | -0.402824 | -0.904186 | 0.617949 | 0.851998 |
| GOBP_NEGATIVE_REGULATION_OF_VASCULAR_PERMEABILITY | 16 | -0.401728 | -0.903787 | 0.620222 | 0.852491 |
| GOBP_THYMOCYTE_APOPTOTIC_PROCESS | 16 | -0.409905 | -0.903399 | 0.610969 | 0.852954 |
| GOBP_REGULATION_OF_SKELETAL_MUSCLE_TISSUE_DEVELOPMENT | 22 | -0.376945 | -0.903139 | 0.611779 | 0.853188 |
| GOBP_NEGATIVE_REGULATION_OF_AXON_EXTENSION_INVOLVED_IN_AXON_( | 26 | -0.367651 | -0.902497 | 0.641667 | 0.854198 |
| GOBP_ACTIVATION_OF_CYSTEINE_TYPE_ENDOPEPTIDASE_ACTIVITY | 15 | -0.402609 | -0.902029 | 0.611902 | 0.854844 |
| GOBP_VESICLE_TARGETING_TO_FROM_OR_WITHIN_GOLGI | 74 | -0.323699 | -0.901633 | 0.679245 | 0.855327 |
| GOBP_NAD_BIOSYNTHETIC_PROCESS | 27 | -0.369423 | -0.901159 | 0.637972 | 0.855992 |
| GOBP_REGULATION_OF_PROTEIN_DEPOLYMERIZATION | 82 | -0.324028 | -0.900919 | 0.686583 | 0.856159 |
| GOBP_RESPONSE_TO_DEXAMETHASONE | 35 | -0.350759 | -0.900462 | 0.641808 | 0.856771 |
| GOBP_NEGATIVE_REGULATION_OF_STRESS_ACTIVATED_PROTEIN_KINASE_S | 46 | -0.341629 | -0.900389 | 0.644592 | 0.856609 |
| GOBP_APICAL_JUNCTION_ASSEMBLY | 76 | -0.322505 | -0.900275 | 0.669439 | 0.85654 |
| GOBP_REGULATION_OF_ANION_TRANSMEMBRANE_TRANSPORT | 99 | -0.318502 | -0.899876 | 0.706064 | 0.857061 |
| GOBP_STRESS_FIBER_ASSEMBLY | 100 | -0.317119 | -0.899696 | 0.710175 | 0.857092 |
| GOBP_PHOSPHOLIPID_METABOLIC_PROCESS | 397 | -0.301914 | -0.899523 | 0.824825 | 0.857136 |
| GOBP_RESPONSE_TO_ISCHEMIA | 48 | -0.338823 | -0.89824 | 0.658889 | 0.859425 |
| GOBP_POSITIVE_REGULATION_OF_SODIUM_ION_TRANSMEMBRANE_TRANSP( | 16 | -0.397417 | -0.898172 | 0.619048 | 0.859232 |
| GOBP_POSITIVE_REGULATION_OF_TRIGLYCERIDE_METABOLIC_PROCESS | 18 | -0.38937 | -0.897278 | 0.619524 | 0.860742 |
| GOBP_POSITIVE_REGULATION_OF_PEPTIDE_SECRETION | 136 | -0.313634 | -0.896973 | 0.723404 | 0.861025 |
| GOBP_NEGATIVE_REGULATION_OF_SIGNAL_TRANSDUCTION_BY_P53_CLASS_ | 30 | -0.360619 | -0.89656 | 0.623853 | 0.86153 |
| GOBP_POSITIVE_REGULATION_OF_CELL_AGING | 15 | -0.404919 | -0.895463 | 0.63464 | 0.86344 |
| GOBP_CYTOSOLIC_TRANSPORT | 165 | -0.309312 | -0.894871 | 0.739657 | 0.86432 |
| GOBP_GLYCEROPHOSPHOLIPID_METABOLIC_PROCESS | 331 | -0.301543 | -0.894837 | 0.82 | 0.86407 |
| GOBP_GAS_TRANSPORT | 17 | -0.39352 | -0.894777 | 0.643902 | 0.863867 |
| GOBP_COPII_COATED_VESICLE_BUDDING | 73 | -0.325159 | -0.894648 | 0.697796 | 0.863802 |
| GOBP_SULFATION | 19 | -0.392403 | -0.893938 | 0.628395 | 0.864877 |
| GOBP_NEGATIVE_REGULATION_OF_SODIUM_ION_TRANSPORT | 17 | -0.393658 | -0.893181 | 0.622618 | 0.86607 |
| GOBP_REGULATION_OF_RESPONSE_TO_OXIDATIVE_STRESS | 79 | -0.321978 | -0.892367 | 0.685146 | 0.867358 |
| GOBP_POSITIVE_REGULATION_OF_NEURON_MIGRATION | 15 | -0.407131 | -0.891361 | 0.644133 | 0.868996 |
| GOBP_CELLULAR_RESPONSE_TO_VITAMIN_D | 20 | -0.383911 | -0.891337 | 0.640491 | 0.868721 |
| GOBP_PROTEIN_HOMOTETRAMERIZATION | 52 | -0.332858 | -0.890076 | 0.682981 | 0.87084 |
| GOBP_NEGATIVE_REGULATION_OF_EPITHELIAL_CELL_MIGRATION | 66 | -0.325948 | -0.889909 | 0.676564 | 0.870828 |
| GOBP_SKELETAL_MUSCLE_TISSUE_REGENERATION | 33 | -0.347598 | -0.889888 | 0.641686 | 0.870549 |
| GOBP_REGULATION_OF_EPIDERMAL_GROWTH_FACTOR_ACTIVATED_RECEPT | 28 | -0.358835 | -0.88973 | 0.641705 | 0.870551 |
| GOBP_ESTABLISHMENT_OF_TISSUE_POLARITY | 121 | -0.310069 | -0.888924 | 0.730496 | 0.871801 |
| GOBP_TETRAPYRROLE_METABOLIC_PROCESS | 57 | -0.328793 | -0.888765 | 0.681378 | 0.871799 |
| GOBP_ERYTHROCYTE_DEVELOPMENT | 32 | -0.355 | -0.888552 | 0.665127 | 0.871897 |
| GOBP_MODULATION_BY_HOST_OF_VIRAL_GENOME_REPLICATION | 18 | -0.388467 | -0.887187 | 0.641439 | 0.874224 |
| GOBP_IN_UTERO_EMBRYONIC_DEVELOPMENT | 322 | -0.300559 | -0.886644 | 0.844 | 0.874978 |
| GOBP_AMINE_CATABOLIC_PROCESS | 28 | -0.367548 | -0.886324 | 0.658906 | 0.875275 |
| GOBP_RESPONSE_TO_ESTROGEN | 66 | -0.322741 | -0.886112 | 0.705059 | 0.875368 |
| GOBP_REGULATION_OF_MICROTUBULE_BASED_MOVEMENT | 40 | -0.339516 | -0.885601 | 0.668514 | 0.876032 |
| GOBP_PEPTIDYL_CYSTEINE_MODIFICATION | 46 | -0.331505 | -0.885459 | 0.671823 | 0.875981 |
| GOBP_PLASMA_MEMBRANE_TUBULATION | 20 | -0.375604 | -0.884877 | 0.646914 | 0.876783 |
| GOBP_POSITIVE_REGULATION_OF_HORMONE_SECRETION | 111 | -0.313148 | -0.884662 | 0.726904 | 0.876909 |
| GOBP_PHOSPHOLIPID_BIOSYNTHETIC_PROCESS | 259 | -0.301316 | -0.884009 | 0.820821 | 0.877847 |
| GOBP_CYTOSKELETON_DEPENDENT_INTRACELLULAR_TRANSPORT | 194 | -0.303155 | -0.883961 | 0.774874 | 0.877614 |
| GOBP_EPIDERMAL_GROWTH_FACTOR_RECEPTOR_SIGNALING_PATHWAY | 116 | -0.309061 | -0.882891 | 0.737108 | 0.879309 |
| GOBP_CARBOHYDRATE_CATABOLIC_PROCESS | 182 | -0.303505 | -0.882856 | 0.784925 | 0.879051 |

| | | | | | |
|---|---|---|---|---|---|
| GOBP_CELLULAR_CARBOHYDRATE_METABOLIC_PROCESS | 265 | -0.298104 | -0.882476 | 0.815 | 0.879488 |
| GOBP_ENDOMEMBRANE_SYSTEM_ORGANIZATION | 457 | -0.296072 | -0.881948 | 0.882883 | 0.88019 |
| GOBP_NEURAL_TUBE_DEVELOPMENT | 140 | -0.305385 | -0.881399 | 0.764467 | 0.880943 |
| GOBP_LIPID_HOMEOSTASIS | 132 | -0.303325 | -0.881396 | 0.74645 | 0.880628 |
| GOBP_GLYCINE_METABOLIC_PROCESS | 17 | -0.397723 | -0.881259 | 0.660448 | 0.880577 |
| GOBP_GLYCEROLIPID_METABOLIC_PROCESS | 398 | -0.296554 | -0.880997 | 0.87 | 0.880778 |
| GOBP_POSITIVE_REGULATION_OF_PROTEIN_MATURATION | 24 | -0.362888 | -0.880596 | 0.649473 | 0.881215 |
| GOBP_POSITIVE_REGULATION_OF_ACTIN_FILAMENT_BUNDLE_ASSEMBLY | 61 | -0.324165 | -0.880124 | 0.705945 | 0.881776 |
| GOBP_GASTRIC_ACID_SECRETION | 16 | -0.393032 | -0.879692 | 0.651341 | 0.882293 |
| GOBP_REGULATION_OF_SPHINGOLIPID_BIOSYNTHETIC_PROCESS | 17 | -0.396185 | -0.879357 | 0.649418 | 0.882587 |
| GOBP_POSITIVE_REGULATION_OF_APOPTOTIC_SIGNALING_PATHWAY | 117 | -0.308303 | -0.879028 | 0.744898 | 0.882892 |
| GOBP_NEGATIVE_REGULATION_OF_TOR_SIGNALING | 42 | -0.341634 | -0.878562 | 0.666297 | 0.883467 |
| GOBP_HEART_FIELD_SPECIFICATION | 18 | -0.38805 | -0.878429 | 0.666261 | 0.883381 |
| GOBP_QUINONE_METABOLIC_PROCESS | 30 | -0.352864 | -0.877605 | 0.652989 | 0.884655 |
| GOBP_BODY_FLUID_SECRETION | 77 | -0.314606 | -0.877534 | 0.70433 | 0.884475 |
| GOBP_REGULATION_OF_PROTEIN_LOCALIZATION_TO_NUCLEUS | 126 | -0.301748 | -0.877326 | 0.736789 | 0.884567 |
| GOBP_PHOSPHATIDYLCHOLINE_BIOSYNTHETIC_PROCESS | 37 | -0.341458 | -0.877313 | 0.68018 | 0.884271 |
| GOBP_PURINE_NUCLEOSIDE_BIOSYNTHETIC_PROCESS | 22 | -0.370882 | -0.876857 | 0.658896 | 0.884798 |
| GOBP_NEGATIVE_REGULATION_OF_PROTEIN_CONTAINING_COMPLEX_ASSEM | 130 | -0.30412 | -0.876849 | 0.763211 | 0.884493 |
| GOBP_LONG_CHAIN_FATTY_ACYL_COA_BIOSYNTHETIC_PROCESS | 19 | -0.381821 | -0.875705 | 0.659176 | 0.886336 |
| GOBP_LYSOSOMAL_TRANSPORT | 109 | -0.306726 | -0.875523 | 0.752058 | 0.886351 |
| GOBP_POSITIVE_REGULATION_OF_PROTEIN_IMPORT | 40 | -0.339549 | -0.875491 | 0.696903 | 0.886089 |
| GOBP_MORPHOGENESIS_OF_A_POLARIZED_EPITHELIUM | 142 | -0.304201 | -0.874939 | 0.769697 | 0.886789 |
| GOBP_DEFENSE_RESPONSE_TO_FUNGUS | 21 | -0.373144 | -0.874933 | 0.66787 | 0.886479 |
| GOBP_CELLULAR_SENESCENCE | 67 | -0.321721 | -0.874011 | 0.706383 | 0.887843 |
| GOBP_DNA_CATABOLIC_PROCESS_ENDONUCLEOLYTIC | 30 | -0.352044 | -0.873979 | 0.651462 | 0.887585 |
| GOBP_GENETIC_IMPRINTING | 22 | -0.36827 | -0.873543 | 0.653659 | 0.888085 |
| GOBP_RESPONSE_TO_MANGANESE_ION | 19 | -0.374721 | -0.871928 | 0.659341 | 0.890723 |
| GOBP_PROTEIN_EXIT_FROM_ENDOPLASMIC_RETICULUM | 47 | -0.335361 | -0.871927 | 0.706208 | 0.890402 |
| GOBP_PYRIDINE_CONTAINING_COMPOUND_METABOLIC_PROCESS | 39 | -0.34024 | -0.870046 | 0.706412 | 0.893522 |
| GOBP_PHYSIOLOGICAL_CARDIAC_MUSCLE_HYPERTROPHY | 23 | -0.360428 | -0.86987 | 0.665105 | 0.893514 |
| GOBP_POSITIVE_REGULATION_OF_NUCLEOCYTOPLASMIC_TRANSPORT | 61 | -0.319274 | -0.869767 | 0.708155 | 0.893374 |
| GOBP_RESPONSE_TO_HEAT | 158 | -0.302264 | -0.869656 | 0.781344 | 0.893261 |
| GOBP_RESPONSE_TO_STIMULUS_INVOLVED_IN_REGULATION_OF_MUSCLE_A | 15 | -0.395235 | -0.869348 | 0.645161 | 0.893492 |
| GOBP_INTERMEDIATE_FILAMENT_BASED_PROCESS | 43 | -0.33107 | -0.868778 | 0.696693 | 0.8942 |
| GOBP_ORGANELLE_TRANSPORT_ALONG_MICROTUBULE | 81 | -0.311749 | -0.867802 | 0.723671 | 0.895639 |
| GOBP_INDOLALKYLAMINE_METABOLIC_PROCESS | 15 | -0.389346 | -0.867773 | 0.657143 | 0.895368 |
| GOBP_VASODILATION | 32 | -0.343365 | -0.867452 | 0.681306 | 0.895605 |
| GOBP_MODIFICATION_OF_SYNAPTIC_STRUCTURE | 19 | -0.373298 | -0.867288 | 0.6625 | 0.895574 |
| GOBP_CYTOPLASMIC_SEQUESTERING_OF_TRANSCRIPTION_FACTOR | 15 | -0.390342 | -0.866711 | 0.669221 | 0.896291 |
| GOBP_REGULATION_OF_EXTRACELLULAR_MATRIX_ORGANIZATION | 45 | -0.329811 | -0.866633 | 0.704595 | 0.896099 |
| GOBP_RESPONSE_TO_GAMMA_RADIATION | 52 | -0.325508 | -0.866458 | 0.699566 | 0.896109 |
| GOBP_INTRACELLULAR_RECEPTOR_SIGNALING_PATHWAY | 256 | -0.294642 | -0.86627 | 0.850701 | 0.896124 |
| GOBP_PROTEIN_KINASE_A_SIGNALING | 30 | -0.355889 | -0.866198 | 0.695402 | 0.895946 |
| GOBP_REGULATION_OF_GLUCOSE_TRANSMEMBRANE_TRANSPORT | 65 | -0.314457 | -0.866156 | 0.732409 | 0.895697 |
| GOBP_INOSITOL_PHOSPHATE_CATABOLIC_PROCESS | 18 | -0.379349 | -0.866147 | 0.689951 | 0.895399 |
| GOBP_RNA_DESTABILIZATION | 34 | -0.336495 | -0.866123 | 0.69397 | 0.895122 |
| GOBP_ALCOHOL_BIOSYNTHETIC_PROCESS | 143 | -0.299046 | -0.865267 | 0.792929 | 0.896354 |
| GOBP_CHOLESTEROL_EFFLUX | 47 | -0.323112 | -0.864614 | 0.707506 | 0.897206 |
| GOBP_POSITIVE_REGULATION_OF_MUSCLE_CONTRACTION | 34 | -0.342871 | -0.863663 | 0.695304 | 0.898563 |
| GOBP_MYELIN_ASSEMBLY | 18 | -0.374612 | -0.863151 | 0.663814 | 0.899179 |
| GOBP_RESPONSE_TO_TESTOSTERONE | 39 | -0.33489 | -0.85994 | 0.711387 | 0.904532 |
| GOBP_RESPONSE_TO_DSRNA | 36 | -0.331518 | -0.85942 | 0.698324 | 0.905137 |
| GOBP_INDOLE_CONTAINING_COMPOUND_METABOLIC_PROCESS | 22 | -0.363557 | -0.859319 | 0.666667 | 0.904991 |
| GOBP_GLYCEROLIPID_BIOSYNTHETIC_PROCESS | 252 | -0.293261 | -0.858807 | 0.850701 | 0.905572 |
| GOBP_SEMAPHORIN_PLEXIN_SIGNALING_PATHWAY | 40 | -0.331077 | -0.857992 | 0.690078 | 0.906688 |
| GOBP_RESPONSE_TO_INSULIN | 258 | -0.291082 | -0.857784 | 0.868 | 0.906719 |
| GOBP_REGULATION_OF_CAMP_MEDIATED_SIGNALING | 33 | -0.338449 | -0.857278 | 0.687429 | 0.907228 |
| GOBP_RETINOIC_ACID_RECEPTOR_SIGNALING_PATHWAY | 31 | -0.344189 | -0.857278 | 0.701043 | 0.906903 |
| GOBP_REGULATION_OF_EXCRETION | 19 | -0.37263 | -0.856723 | 0.690887 | 0.907545 |
| GOBP_NUCLEOSIDE_DIPHOSPHATE_METABOLIC_PROCESS | 150 | -0.294937 | -0.856149 | 0.813577 | 0.908196 |
| GOBP_PROTEIN_INSERTION_INTO_MEMBRANE | 74 | -0.312029 | -0.855404 | 0.74814 | 0.909131 |
| GOBP_CELLULAR_RESPONSE_TO_GLUCOSE_STARVATION | 47 | -0.328341 | -0.854065 | 0.725789 | 0.911148 |
| GOBP_POSITIVE_REGULATION_OF_PROTEIN_LOCALIZATION_TO_NUCLEUS | 83 | -0.304123 | -0.853872 | 0.746625 | 0.911167 |
| GOBP_ESTABLISHMENT_OF_PROTEIN_LOCALIZATION_TO_PLASMA_MEMBRAN | 58 | -0.318737 | -0.852612 | 0.74086 | 0.913011 |
| GOBP_PROTEIN_AUTOPROCESSING | 22 | -0.361474 | -0.852426 | 0.690734 | 0.912988 |
| GOBP_REGULATION_OF_TRANSCRIPTION_INVOLVED_IN_G1_S_TRANSITION_C | 35 | -0.333911 | -0.852364 | 0.704651 | 0.912773 |
| GOBP_CELLULAR_GLUCAN_METABOLIC_PROCESS | 69 | -0.311827 | -0.852285 | 0.75 | 0.912585 |
| GOBP_NEURAL_TUBE_FORMATION | 95 | -0.302272 | -0.852202 | 0.778345 | 0.912406 |
| GOBP_PIGMENT_BIOSYNTHETIC_PROCESS | 57 | -0.316788 | -0.851343 | 0.731522 | 0.913516 |
| GOBP_CELL_DEATH_IN_RESPONSE_TO_HYDROGEN_PEROXIDE | 24 | -0.359307 | -0.850227 | 0.718377 | 0.915079 |
| GOBP_PORE_COMPLEX_ASSEMBLY | 20 | -0.358504 | -0.84957 | 0.684401 | 0.915868 |
| GOBP_NEGATIVE_REGULATION_OF_PROTEIN_CONTAINING_COMPLEX_DISASS | 78 | -0.303044 | -0.84931 | 0.75341 | 0.915996 |
| GOBP_POSITIVE_REGULATION_OF_PROTEIN_EXPORT_FROM_NUCLEUS | 20 | -0.365326 | -0.849114 | 0.702602 | 0.916006 |
| GOBP_HISTONE_PHOSPHORYLATION | 39 | -0.33769 | -0.849096 | 0.720801 | 0.915707 |
| GOBP_PIGMENTATION | 88 | -0.30023 | -0.848394 | 0.775489 | 0.916573 |
| GOBP_POSITIVE_REGULATION_OF_STEM_CELL_PROLIFERATION | 21 | -0.356528 | -0.848037 | 0.679042 | 0.916866 |
| GOBP_PROTEIN_DEGLYCOSYLATION | 28 | -0.342434 | -0.847397 | 0.718824 | 0.917605 |
| GOBP_LIPID_STORAGE | 79 | -0.301637 | -0.846718 | 0.767635 | 0.918416 |
| GOBP_INACTIVATION_OF_MAPK_ACTIVITY | 24 | -0.353974 | -0.846518 | 0.699126 | 0.918425 |
| GOBP_POSITIVE_REGULATION_OF_CELL_MORPHOGENESIS_INVOLVED_IN_DIF | 75 | -0.303555 | -0.846476 | 0.751572 | 0.918163 |
| GOBP_POSITIVE_REGULATION_OF_PEPTIDE_HORMONE_SECRETION | 83 | -0.303009 | -0.845882 | 0.75567 | 0.918826 |
| GOBP_VASCULAR_ASSOCIATED_SMOOTH_MUSCLE_CELL_DIFFERENTIATION | 27 | -0.350592 | -0.845856 | 0.699168 | 0.91854 |
| GOBP_TORC1_SIGNALING | 47 | -0.31555 | -0.845848 | 0.739756 | 0.918228 |
| GOBP_RESPONSE_TO_LEUKEMIA_INHIBITORY_FACTOR | 78 | -0.301219 | -0.845586 | 0.760756 | 0.918331 |
| GOBP_LONG_CHAIN_FATTY_ACID_BIOSYNTHETIC_PROCESS | 30 | -0.34363 | -0.844267 | 0.701031 | 0.920138 |
| GOBP_MONOSACCHARIDE_METABOLIC_PROCESS | 253 | -0.287372 | -0.844243 | 0.880642 | 0.919848 |
| GOBP_NEGATIVE_REGULATION_OF_POSTTRANSCRIPTIONAL_GENE_SILENCIN | 19 | -0.369544 | -0.844224 | 0.697704 | 0.919549 |
| GOBP_NEGATIVE_REGULATION_OF_DNA_REPLICATION | 30 | -0.340838 | -0.844082 | 0.717593 | 0.919461 |
| GOBP_TRIGLYCERIDE_BIOSYNTHETIC_PROCESS | 38 | -0.325325 | -0.84353 | 0.738532 | 0.920073 |
| GOBP_POSITIVE_REGULATION_OF_INTRINSIC_APOPTOTIC_SIGNALING_PATHW | 56 | -0.314429 | -0.84346 | 0.746524 | 0.919862 |
| GOBP_ALPHA_AMINO_ACID_BIOSYNTHETIC_PROCESS | 64 | -0.307921 | -0.843415 | 0.766002 | 0.91961 |
| GOBP_POSITIVE_REGULATION_OF_CELL_CYCLE_G1_S_PHASE_TRANSITION | 43 | -0.322185 | -0.842419 | 0.739955 | 0.920922 |
| GOBP_POSITIVE_REGULATION_OF_NEUROTRANSMITTER_TRANSPORT | 22 | -0.362127 | -0.842382 | 0.720482 | 0.920654 |
| GOBP_NEGATIVE_REGULATION_OF_PROTEIN_KINASE_B_SIGNALING | 44 | -0.320796 | -0.842349 | 0.741971 | 0.920386 |
| GOBP_PHAGOLYSOSOME_ASSEMBLY | 15 | -0.384178 | -0.842165 | 0.710692 | 0.920372 |
| GOBP_NEGATIVE_REGULATION_OF_DNA_DAMAGE_RESPONSE_SIGNAL_TRAN | 15 | -0.377547 | -0.841622 | 0.679898 | 0.920922 |
| GOBP_GUANOSINE_CONTAINING_COMPOUND_METABOLIC_PROCESS | 40 | -0.322919 | -0.841478 | 0.719239 | 0.920837 |
| GOBP_REGULATION_OF_MUSCLE_ADAPTATION | 77 | -0.299961 | -0.841034 | 0.755814 | 0.921251 |
| GOBP_LONG_CHAIN_FATTY_ACID_IMPORT_INTO_CELL | 17 | -0.369256 | -0.840452 | 0.698885 | 0.921863 |
| GOBP_CELLULAR_MODIFIED_AMINO_ACID_BIOSYNTHETIC_PROCESS | 46 | -0.318235 | -0.839302 | 0.726571 | 0.923339 |
| GOBP_HYDROGEN_PEROXIDE_METABOLIC_PROCESS | 52 | -0.317484 | -0.839289 | 0.744797 | 0.923031 |
| GOBP_POSITIVE_REGULATION_OF_MUSCLE_TISSUE_DEVELOPMENT | 20 | -0.355433 | -0.838825 | 0.685323 | 0.923458 |
| GOBP_ERBB_SIGNALING_PATHWAY | 138 | -0.291607 | -0.837932 | 0.828921 | 0.924555 |
| GOBP_NUCLEOBASE_CONTAINING_SMALL_MOLECULE_CATABOLIC_PROCESS | 47 | -0.314653 | -0.836034 | 0.727273 | 0.927293 |
| GOBP_STEROID_BIOSYNTHETIC_PROCESS | 168 | -0.28741 | -0.83524 | 0.854985 | 0.92824 |
| GOBP_ANDROGEN_METABOLIC_PROCESS | 25 | -0.347587 | -0.834577 | 0.710094 | 0.928959 |
| GOBP_MULTICELLULAR_ORGANISMAL_MOVEMENT | 50 | -0.314936 | -0.834276 | 0.74295 | 0.929121 |
| GOBP_PSEUDOPODIUM_ORGANIZATION | 17 | -0.372526 | -0.833491 | 0.720643 | 0.930025 |
| GOBP_PYRIDINE_CONTAINING_COMPOUND_BIOSYNTHETIC_PROCESS | 33 | -0.332282 | -0.832847 | 0.735936 | 0.930731 |
| GOBP_REGULATION_OF_PROTEIN_CONTAINING_COMPLEX_DISASSEMBLY | 116 | -0.293986 | -0.832739 | 0.794688 | 0.930564 |
| GOBP_POSITIVE_REGULATION_OF_CELLULAR_RESPONSE_TO_INSULIN_STIMU | 24 | -0.347025 | -0.832497 | 0.721698 | 0.930633 |
| GOBP_MAINTENANCE_OF_CELL_POLARITY | 18 | -0.369685 | -0.832074 | 0.713936 | 0.930974 |
| GOBP_STEROID_HORMONE_MEDIATED_SIGNALING_PATHWAY | 126 | -0.290321 | -0.831566 | 0.82706 | 0.931449 |
| GOBP_MICROTUBULE_BASED_TRANSPORT | 187 | -0.284676 | -0.831553 | 0.864594 | 0.931141 |
| GOBP_REGULATION_OF_CARBOHYDRATE_BIOSYNTHETIC_PROCESS | 87 | -0.296953 | -0.830883 | 0.790843 | 0.931879 |
| GOBP_NEGATIVE_REGULATION_OF_CATABOLIC_PROCESS | 289 | -0.281392 | -0.83008 | 0.90991 | 0.932781 |
| GOBP_REGULATION_OF_PLATELET_DERIVED_GROWTH_FACTOR_RECEPTOR_ | 21 | -0.357615 | -0.829595 | 0.726176 | 0.933206 |
| GOBP_PIGMENT_CELL_DIFFERENTIATION | 32 | -0.331368 | -0.829588 | 0.726962 | 0.932893 |
| GOBP_RELEASE_OF_CYTOCHROME_C_FROM_MITOCHONDRIA | 54 | -0.308705 | -0.828493 | 0.759358 | 0.934282 |
| GOBP_POSITIVE_REGULATION_OF_ANION_TRANSMEMBRANE_TRANSPORT | 53 | -0.308254 | -0.828435 | 0.758324 | 0.934045 |
| GOBP_THYROID_GLAND_DEVELOPMENT | 17 | -0.367212 | -0.827316 | 0.715385 | 0.935471 |
| GOBP_REGULATION_OF_PROTEIN_CATABOLIC_PROCESS | 372 | -0.280165 | -0.826915 | 0.937 | 0.93579 |
| GOBP_EPITHELIAL_CELL_CELL_ADHESION | 17 | -0.36585 | -0.826358 | 0.716285 | 0.936308 |
| GOBP_REGULATION_OF_GLUCOSE_METABOLIC_PROCESS | 110 | -0.291706 | -0.825191 | 0.822746 | 0.937801 |
| GOBP_CHEMOKINE_C_X_C_MOTIF_LIGAND_2_PRODUCTION | 15 | -0.37621 | -0.824889 | 0.731801 | 0.937956 |
| GOBP_MICROTUBULE_ORGANIZING_CENTER_LOCALIZATION | 30 | -0.331209 | -0.824844 | 0.740872 | 0.937685 |
| GOBP_SMOOTH_MUSCLE_CELL_APOPTOTIC_PROCESS | 23 | -0.337129 | -0.824403 | 0.740926 | 0.938026 |
| GOBP_RESPONSE_TO_AMINO_ACID_STARVATION | 49 | -0.312514 | -0.824028 | 0.765339 | 0.938256 |
| GOBP_RESPONSE_TO_UV_B | 15 | -0.374132 | -0.820819 | 0.717982 | 0.942818 |
| GOBP_REGULATION_OF_CAMP_DEPENDENT_PROTEIN_KINASE_ACTIVITY | 19 | -0.355354 | -0.820278 | 0.739506 | 0.943289 |
| GOBP_SKELETAL_MUSCLE_CONTRACTION | 40 | -0.317563 | -0.819493 | 0.755079 | 0.944108 |
| GOBP_NEUTRAL_LIPID_METABOLIC_PROCESS | 112 | -0.288971 | -0.818735 | 0.82551 | 0.944854 |
| GOBP_PROTEIN_N_LINKED_GLYCOSYLATION | 72 | -0.296455 | -0.81855 | 0.789863 | 0.944809 |
| GOBP_REGULATION_OF_MICROTUBULE_POLYMERIZATION | 49 | -0.305072 | -0.816846 | 0.780911 | 0.947029 |
| GOBP_REGULATION_OF_MITOTIC_NUCLEAR_DIVISION | 104 | -0.292059 | -0.816741 | 0.841727 | 0.946862 |
| GOBP_RESPONSE_TO_PROGESTERONE | 40 | -0.31754 | -0.816391 | 0.767232 | 0.947052 |
| GOBP_REGULATION_OF_MUSCLE_ORGAN_DEVELOPMENT | 37 | -0.321237 | -0.814993 | 0.76 | 0.948789 |
| GOBP_ATP_TRANSPORT | 15 | -0.368723 | -0.814943 | 0.724138 | 0.948526 |
| GOBP_PARAXIAL_MESODERM_DEVELOPMENT | 17 | -0.359092 | -0.81485 | 0.723378 | 0.948335 |
| GOBP_REGULATION_OF_DIGESTIVE_SYSTEM_PROCESS | 31 | -0.327488 | -0.814801 | 0.757955 | 0.948086 |
| GOBP_NEGATIVE_REGULATION_OF_CELLULAR_AMIDE_METABOLIC_PROCESS | 163 | -0.28291 | -0.814486 | 0.882947 | 0.94821 |
| GOBP_3_PHOSPHOADENOSINE_5_PHOSPHOSULFATE_METABOLIC_PROCESS | 20 | -0.346193 | -0.814418 | 0.735043 | 0.947978 |
| GOBP_MEMBRANE_DOCKING | 181 | -0.279392 | -0.814269 | 0.881526 | 0.947853 |
| GOBP_LIPID_OXIDATION | 103 | -0.28996 | -0.814213 | 0.835366 | 0.947599 |
| GOBP_REGULATION_OF_CELLULAR_RESPONSE_TO_INSULIN_STIMULUS | 62 | -0.299816 | -0.813912 | 0.792332 | 0.947713 |
| GOBP_RUFFLE_ASSEMBLY | 42 | -0.308181 | -0.813299 | 0.765432 | 0.948265 |
| GOBP_POSITIVE_REGULATION_OF_LAMELLIPODIUM_ORGANIZATION | 35 | -0.316364 | -0.813174 | 0.767816 | 0.948119 |
| GOBP_ACTOMYOSIN_STRUCTURE_ORGANIZATION | 178 | -0.280522 | -0.813081 | 0.887776 | 0.947922 |
| GOBP_NEGATIVE_REGULATION_OF_SMOOTH_MUSCLE_CELL_DIFFERENTIATIO | 16 | -0.36094 | -0.812908 | 0.732304 | 0.947847 |
| GOBP_NEUROEPITHELIAL_CELL_DIFFERENTIATION | 19 | -0.353417 | -0.812293 | 0.726496 | 0.948422 |

| | | | | | |
|---|---|---|---|---|---|
| GOBP_REGULATION_OF_SMOOTH_MUSCLE_CELL_DIFFERENTIATION | 30 | -0.323881 | -0.812159 | 0.759091 | 0.948296 |
| GOBP_REGULATION_OF_OXIDATIVE_STRESS_INDUCED_NEURON_DEATH | 21 | -0.34326 | -0.811743 | 0.746667 | 0.948568 |
| GOBP_MUSCLE_FILAMENT_SLIDING | 34 | -0.320913 | -0.811706 | 0.753757 | 0.948294 |
| GOBP_REGULATION_OF_RELEASE_OF_CYTOCHROME_C_FROM_MITOCHONDI | 43 | -0.31039 | -0.811188 | 0.801552 | 0.948714 |
| GOBP_NEURAL_TUBE_PATTERNING | 32 | -0.32288 | -0.810965 | 0.768349 | 0.948713 |
| GOBP_REGULATION_OF_INTRINSIC_APOPTOTIC_SIGNALING_PATHWAY | 149 | -0.281045 | -0.810896 | 0.864647 | 0.948489 |
| GOBP_POLYSACCHARIDE_CATABOLIC_PROCESS | 24 | -0.342305 | -0.81079 | 0.750886 | 0.94832 |
| GOBP_NEGATIVE_REGULATION_OF_NECROTIC_CELL_DEATH | 18 | -0.352005 | -0.8106 | 0.755945 | 0.948266 |
| GOBP_FATTY_ACID_METABOLIC_PROCESS | 341 | -0.274358 | -0.809853 | 0.954 | 0.949048 |
| GOBP_PROTON_TRANSMEMBRANE_TRANSPORT | 144 | -0.281019 | -0.8098 | 0.862348 | 0.948797 |
| GOBP_REGULATION_OF_CHOLESTEROL_EFFLUX | 31 | -0.325939 | -0.809715 | 0.757932 | 0.948592 |
| GOBP_RESPONSE_TO_X_RAY | 32 | -0.32044 | -0.809153 | 0.764045 | 0.949083 |
| GOBP_POSITIVE_REGULATION_OF_INSULIN_SECRETION_INVOLVED_IN_CELLU | 31 | -0.324247 | -0.807993 | 0.769231 | 0.950394 |
| GOBP_BILE_ACID_METABOLIC_PROCESS | 42 | -0.30903 | -0.807863 | 0.779978 | 0.950246 |
| GOBP_RAB_PROTEIN_SIGNAL_TRANSDUCTION | 21 | -0.34606 | -0.807654 | 0.749698 | 0.9502 |
| GOBP_PROTEIN_LOCALIZATION_TO_NUCLEUS | 261 | -0.274346 | -0.807627 | 0.924 | 0.949911 |
| GOBP_CELLULAR_RESPONSE_TO_STARVATION | 151 | -0.280395 | -0.806539 | 0.872599 | 0.951156 |
| GOBP_PEPTIDYL_ARGININE_MODIFICATION | 25 | -0.330386 | -0.805741 | 0.773428 | 0.951944 |
| GOBP_POSITIVE_REGULATION_OF_DOUBLE_STRAND_BREAK_REPAIR_VIA_NC | 16 | -0.354272 | -0.804683 | 0.726221 | 0.95311 |
| GOBP_SULFUR_COMPOUND_TRANSPORT | 47 | -0.30452 | -0.804677 | 0.781421 | 0.952789 |
| GOBP_GLYCOSIDE_METABOLIC_PROCESS | 20 | -0.343823 | -0.804436 | 0.747848 | 0.952785 |
| GOBP_NEGATIVE_REGULATION_OF_MICROTUBULE_POLYMERIZATION_OR_DE | 37 | -0.308291 | -0.804127 | 0.749164 | 0.9529 |
| GOBP_REGULATION_OF_CIRCADIAN_RHYTHM | 106 | -0.280925 | -0.804 | 0.851434 | 0.952755 |
| GOBP_ACROSOME_REACTION | 32 | -0.320317 | -0.803778 | 0.776391 | 0.952733 |
| GOBP_CELLULAR_RESPONSE_TO_PROSTAGLANDIN_STIMULUS | 20 | -0.346317 | -0.803686 | 0.738007 | 0.952537 |
| GOBP_NEGATIVE_REGULATION_OF_PEPTIDYL_THREONINE_PHOSPHORYLATI | 19 | -0.353029 | -0.803388 | 0.756098 | 0.952621 |
| GOBP_REGULATION_OF_SUBSTRATE_ADHESION_DEPENDENT_CELL_SPREAD | 54 | -0.302573 | -0.802756 | 0.800217 | 0.953203 |
| GOBP_REGULATION_OF_EARLY_ENDOSOME_TO_LATE_ENDOSOME_TRANSPO | 18 | -0.352281 | -0.802491 | 0.723785 | 0.95324 |
| GOBP_TRIGLYCERIDE_CATABOLIC_PROCESS | 28 | -0.325435 | -0.802302 | 0.762632 | 0.953169 |
| GOBP_CARBOHYDRATE_DERIVATIVE_TRANSPORT | 76 | -0.28691 | -0.801874 | 0.820942 | 0.953434 |
| GOBP_3_UTR_MEDIATED_MRNA_DESTABILIZATION | 16 | -0.351516 | -0.80174 | 0.754453 | 0.953302 |
| GOBP_POSITIVE_REGULATION_OF_PROTEIN_BINDING | 80 | -0.288761 | -0.801533 | 0.821946 | 0.953253 |
| GOBP_POSITIVE_REGULATION_OF_SKELETAL_MUSCLE_TISSUE_DEVELOPMEI | 16 | -0.35724 | -0.80146 | 0.751226 | 0.953016 |
| GOBP_POSITIVE_REGULATION_OF_MITOCHONDRION_ORGANIZATION | 83 | -0.290492 | -0.801127 | 0.823406 | 0.95316 |
| GOBP_POSITIVE_REGULATION_OF_STRESS_FIBER_ASSEMBLY | 52 | -0.296101 | -0.800511 | 0.809168 | 0.953682 |
| GOBP_MUSCLE_HYPERTROPHY_IN_RESPONSE_TO_STRESS | 19 | -0.346816 | -0.800339 | 0.76625 | 0.953607 |
| GOBP_ENDOPLASMIC_RETICULUM_TO_GOLGI_VESICLE_MEDIATED_TRANSPO | 198 | -0.273542 | -0.800217 | 0.907908 | 0.953451 |
| GOBP_ENTRY_OF_BACTERIUM_INTO_HOST_CELL | 15 | -0.362976 | -0.799255 | 0.749354 | 0.954495 |
| GOBP_TRANSPORT_ALONG_MICROTUBULE | 156 | -0.274758 | -0.799198 | 0.889676 | 0.954246 |
| GOBP_NEGATIVE_REGULATION_OF_PEPTIDYL_SERINE_PHOSPHORYLATION | 27 | -0.320228 | -0.796948 | 0.773697 | 0.956975 |
| GOBP_STEROL_TRANSPORT | 99 | -0.283428 | -0.796333 | 0.850976 | 0.957483 |
| GOBP_RESPONSE_TO_ANGIOTENSIN | 22 | -0.336843 | -0.795881 | 0.759904 | 0.957793 |
| GOBP_REGULATION_OF_HISTONE_H3_K4_METHYLATION | 25 | -0.330427 | -0.795631 | 0.785467 | 0.957792 |
| GOBP_REGULATION_OF_MICROTUBULE_CYTOSKELETON_ORGANIZATION | 136 | -0.276098 | -0.795012 | 0.867677 | 0.958295 |
| GOBP_NEGATIVE_REGULATION_OF_ACTIN_FILAMENT_DEPOLYMERIZATION | 42 | -0.309628 | -0.795004 | 0.804469 | 0.957976 |
| GOBP_GOLGI_TO_PLASMA_MEMBRANE_PROTEIN_TRANSPORT | 38 | -0.307116 | -0.793749 | 0.770193 | 0.959308 |
| GOBP_REGULATION_OF_ER_TO_GOLGI_VESICLE_MEDIATED_TRANSPORT | 15 | -0.365218 | -0.793061 | 0.759115 | 0.959899 |
| GOBP_VESICLE_TARGETING | 95 | -0.283478 | -0.792444 | 0.864301 | 0.960374 |
| GOBP_CYTOPLASMIC_PATTERN_RECOGNITION_RECEPTOR_SIGNALING_PATH | 29 | -0.320173 | -0.792411 | 0.7625 | 0.960089 |
| GOBP_REGULATION_OF_MITOCHONDRIAL_MEMBRANE_POTENTIAL | 70 | -0.286799 | -0.792394 | 0.822362 | 0.95978 |
| GOBP_POSITIVE_REGULATION_OF_GLUCOSE_TRANSMEMBRANE_TRANSPORT | 37 | -0.30831 | -0.792255 | 0.790487 | 0.959635 |
| GOBP_VESICLE_MEDIATED_TRANSPORT_TO_THE_PLASMA_MEMBRANE | 91 | -0.281688 | -0.791264 | 0.86653 | 0.960613 |
| GOBP_GOLGI_ORGANIZATION | 140 | -0.272976 | -0.790158 | 0.879032 | 0.961734 |
| GOBP_RESPONSE_TO_MUSCLE_ACTIVITY | 23 | -0.330276 | -0.789726 | 0.77431 | 0.961986 |
| GOBP_POSITIVE_REGULATION_OF_NEUROTRANSMITTER_SECRETION | 17 | -0.348552 | -0.789429 | 0.755025 | 0.962038 |
| GOBP_REGULATION_OF_CELL_CYCLE_G1_S_PHASE_TRANSITION | 160 | -0.272395 | -0.789014 | 0.906977 | 0.96228 |
| GOBP_POSITIVE_REGULATION_OF_MITOCHONDRIAL_FISSION | 17 | -0.344142 | -0.788827 | 0.753165 | 0.962206 |
| GOBP_SEX_DETERMINATION | 15 | -0.355674 | -0.788713 | 0.773608 | 0.962039 |
| GOBP_MONOCARBOXYLIC_ACID_CATABOLIC_PROCESS | 121 | -0.27305 | -0.787285 | 0.876518 | 0.963538 |
| GOBP_MITOTIC_G2_DNA_DAMAGE_CHECKPOINT | 23 | -0.331855 | -0.787077 | 0.788462 | 0.96347 |
| GOBP_RESPONSE_TO_STARVATION | 185 | -0.269596 | -0.786424 | 0.918675 | 0.963983 |
| GOBP_CELLULAR_RESPONSE_TO_DEXAMETHASONE_STIMULUS | 27 | -0.316418 | -0.786288 | 0.776744 | 0.963827 |
| GOBP_MEMBRANE_FISSION | 15 | -0.35371 | -0.78526 | 0.765903 | 0.96482 |
| GOBP_PROTEIN_TRANSPORT_ALONG_MICROTUBULE | 68 | -0.282844 | -0.784934 | 0.8579 | 0.964915 |
| GOBP_REGULATION_OF_MACROAUTOPHAGY | 158 | -0.272156 | -0.784766 | 0.89596 | 0.964818 |
| GOBP_POST_GOLGI_VESICLE_MEDIATED_TRANSPORT | 103 | -0.27606 | -0.784281 | 0.874239 | 0.965132 |
| GOBP_CYTOPLASMIC_SEQUESTERING_OF_PROTEIN | 22 | -0.332069 | -0.782359 | 0.773381 | 0.967274 |
| GOBP_TOR_SIGNALING | 116 | -0.271621 | -0.781787 | 0.871327 | 0.96766 |
| GOBP_SULFUR_AMINO_ACID_METABOLIC_PROCESS | 37 | -0.305693 | -0.781275 | 0.80778 | 0.967965 |
| GOBP_ACTIVIN_RECEPTOR_SIGNALING_PATHWAY | 34 | -0.312449 | -0.780576 | 0.816949 | 0.968519 |
| GOBP_CELL_AGING | 101 | -0.277519 | -0.78034 | 0.857436 | 0.968482 |
| GOBP_NAD_METABOLIC_PROCESS | 50 | -0.293092 | -0.779812 | 0.839737 | 0.968811 |
| GOBP_PURINE_NUCLEOSIDE_MONOPHOSPHATE_METABOLIC_PROCESS | 42 | -0.299462 | -0.779755 | 0.813901 | 0.968558 |
| GOBP_POSITIVE_REGULATION_OF_MICROTUBULE_POLYMERIZATION_OR_DEF | 32 | -0.308691 | -0.778921 | 0.79436 | 0.969284 |
| GOBP_REGULATION_OF_INTRINSIC_APOPTOTIC_SIGNALING_PATHWAY_IN_RE | 17 | -0.342156 | -0.778545 | 0.767003 | 0.969443 |
| GOBP_POSITIVE_REGULATION_OF_MEMBRANE_POTENTIAL | 16 | -0.348573 | -0.778527 | 0.762861 | 0.969138 |
| GOBP_ARP2_3_COMPLEX_MEDIATED_ACTIN_NUCLEATION | 39 | -0.300031 | -0.778468 | 0.808607 | 0.968887 |
| GOBP_CELLULAR_RESPONSE_TO_INSULIN_STIMULUS | 211 | -0.2644 | -0.775941 | 0.943888 | 0.971719 |
| GOBP_REGULATION_OF_LIPID_STORAGE | 50 | -0.290914 | -0.775408 | 0.830601 | 0.972062 |
| GOBP_POSITIVE_REGULATION_OF_STEROL_TRANSPORT | 28 | -0.316055 | -0.77534 | 0.811321 | 0.971813 |
| GOBP_KETONE_BIOSYNTHETIC_PROCESS | 38 | -0.2985 | -0.775302 | 0.797539 | 0.971538 |
| GOBP_POSITIVE_REGULATION_OF_MEMBRANE_PERMEABILITY | 64 | -0.283025 | -0.774976 | 0.845011 | 0.97163 |
| GOBP_EXTRACELLULAR_TRANSPORT | 34 | -0.304834 | -0.773945 | 0.80137 | 0.97258 |
| GOBP_LAMELLIPODIUM_MORPHOGENESIS | 16 | -0.345181 | -0.77358 | 0.790668 | 0.972712 |
| GOBP_PROTEIN_TARGETING_TO_VACUOLE | 40 | -0.300469 | -0.772993 | 0.811725 | 0.973085 |
| GOBP_NEGATIVE_REGULATION_OF_REPRODUCTIVE_PROCESS | 44 | -0.293383 | -0.772651 | 0.817686 | 0.97318 |
| GOBP_CELLULAR_RESPONSE_TO_HEAT | 119 | -0.271382 | -0.772628 | 0.888325 | 0.972882 |
| GOBP_POSITIVE_REGULATION_OF_MEIOTIC_CELL_CYCLE | 19 | -0.337067 | -0.772605 | 0.787955 | 0.972579 |
| GOBP_VACUOLAR_TRANSPORT | 151 | -0.26708 | -0.772525 | 0.929293 | 0.972339 |
| GOBP_ASYMMETRIC_CELL_DIVISION | 17 | -0.3416 | -0.770817 | 0.781211 | 0.974091 |
| GOBP_BINDING_OF_SPERM_TO_ZONA_PELLUCIDA | 32 | -0.308572 | -0.769611 | 0.812933 | 0.97522 |
| GOBP_PHOSPHATIDYLCHOLINE_METABOLIC_PROCESS | 82 | -0.277765 | -0.769448 | 0.868476 | 0.975077 |
| GOBP_RESPONSE_TO_MUSCLE_STRETCH | 20 | -0.334455 | -0.766749 | 0.787578 | 0.977994 |
| GOBP_POST_TRANSLATIONAL_PROTEIN_MODIFICATION | 339 | -0.258452 | -0.765892 | 0.978 | 0.978665 |
| GOBP_MEMBRANE_DISASSEMBLY | 15 | -0.351619 | -0.765557 | 0.799499 | 0.97875 |
| GOBP_RIBONUCLEOSIDE_CATABOLIC_PROCESS | 22 | -0.325859 | -0.764328 | 0.81592 | 0.979902 |
| GOBP_PROTEIN_PHOSPHOPANTETHEINYLATION | 297 | -0.259664 | -0.763966 | 0.963964 | 0.97998 |
| GOBP_NEGATIVE_REGULATION_OF_HISTONE_METHYLATION | 21 | -0.327374 | -0.763679 | 0.804136 | 0.979997 |
| GOBP_REGULATION_OF_MICROTUBULE_POLYMERIZATION_OR_DEPOLYMERIZ | 80 | -0.274602 | -0.762906 | 0.868201 | 0.980591 |
| GOBP_REGULATION_OF_CELL_AGING | 48 | -0.286924 | -0.762871 | 0.827051 | 0.980314 |
| GOBP_RENAL_SODIUM_EXCRETION | 18 | -0.334128 | -0.76249 | 0.809102 | 0.980448 |
| GOBP_PYRIMIDINE_RIBONUCLEOTIDE_BIOSYNTHETIC_PROCESS | 22 | -0.324377 | -0.762404 | 0.813028 | 0.98022 |
| GOBP_SULFUR_AMINO_ACID_BIOSYNTHETIC_PROCESS | 17 | -0.339534 | -0.762176 | 0.801508 | 0.980158 |
| GOBP_CELL_CYCLE_ARREST | 222 | -0.259808 | -0.762152 | 0.943832 | 0.979852 |
| GOBP_REGULATION_OF_PROTEIN_STABILITY | 285 | -0.257363 | -0.76196 | 0.965966 | 0.979751 |
| GOBP_REGULATION_OF_STEROL_TRANSPORT | 50 | -0.288757 | -0.761009 | 0.834962 | 0.98052 |
| GOBP_PYRIMIDINE_CONTAINING_COMPOUND_METABOLIC_PROCESS | 81 | -0.273582 | -0.760813 | 0.86225 | 0.980433 |
| GOBP_RESPONSE_TO_HYPEROXIA | 22 | -0.322156 | -0.760578 | 0.793939 | 0.980394 |
| GOBP_CIRCADIAN_SLEEP_WAKE_CYCLE | 18 | -0.334827 | -0.760479 | 0.803727 | 0.980186 |
| GOBP_DNA_METHYLATION | 61 | -0.280882 | -0.759887 | 0.86873 | 0.980536 |
| GOBP_REGULATION_OF_LIPOPROTEIN_METABOLIC_PROCESS | 15 | -0.343447 | -0.758509 | 0.78125 | 0.981786 |
| GOBP_POSITIVE_REGULATION_OF_DNA_METABOLIC_PROCESS | 191 | -0.260665 | -0.758316 | 0.938816 | 0.981682 |
| GOBP_REGULATION_OF_DNA_RECOMBINATION | 99 | -0.269294 | -0.758179 | 0.884735 | 0.9815 |
| GOBP_DNA_DAMAGE_RESPONSE_SIGNAL_TRANSDUCTION_BY_P53_CLASS_M | 105 | -0.263752 | -0.756211 | 0.906571 | 0.983411 |
| GOBP_NEGATIVE_REGULATION_OF_ANOIKIS | 17 | -0.334683 | -0.75524 | 0.783851 | 0.984167 |
| GOBP_RIBOSE_PHOSPHATE_BIOSYNTHETIC_PROCESS | 191 | -0.258736 | -0.754501 | 0.944724 | 0.984707 |
| GOBP_REGULATION_OF_HISTONE_MODIFICATION | 136 | -0.260127 | -0.753893 | 0.927846 | 0.985077 |
| GOBP_MOTILE_CILIUM_ASSEMBLY | 21 | -0.324826 | -0.753736 | 0.82593 | 0.984921 |
| GOBP_NEGATIVE_REGULATION_OF_FIBROBLAST_PROLIFERATION | 29 | -0.307463 | -0.751436 | 0.829268 | 0.987171 |
| GOBP_GOLGI_VESICLE_TRANSPORT | 359 | -0.253618 | -0.751079 | 0.97998 | 0.987249 |
| GOBP_THIOESTER_BIOSYNTHETIC_PROCESS | 52 | -0.280436 | -0.749743 | 0.854821 | 0.98842 |
| GOBP_REGULATION_OF_TOR_SIGNALING | 94 | -0.264689 | -0.747479 | 0.905895 | 0.990565 |
| GOBP_PROTEIN_K63_LINKED_UBIQUITINATION | 56 | -0.274587 | -0.746668 | 0.856223 | 0.991112 |
| GOBP_UDP_N_ACETYLGLUCOSAMINE_METABOLIC_PROCESS | 16 | -0.329173 | -0.745989 | 0.808729 | 0.991505 |
| GOBP_GLYCEROL_ETHER_METABOLIC_PROCESS | 21 | -0.315531 | -0.745608 | 0.834348 | 0.99158 |
| GOBP_REGULATION_OF_OXIDATIVE_PHOSPHORYLATION | 23 | -0.308851 | -0.743624 | 0.806024 | 0.993388 |
| GOBP_RHYTHMIC_BEHAVIOR | 36 | -0.291074 | -0.743235 | 0.83908 | 0.993474 |
| GOBP_NUCLEOSIDE_PHOSPHATE_CATABOLIC_PROCESS | 80 | -0.26772 | -0.743175 | 0.90501 | 0.993201 |
| GOBP_CELLULAR_CARBOHYDRATE_CATABOLIC_PROCESS | 43 | -0.283661 | -0.743116 | 0.847085 | 0.992935 |
| GOBP_POSITIVE_REGULATION_OF_REPRODUCTIVE_PROCESS | 65 | -0.272384 | -0.742831 | 0.876068 | 0.99291 |
| GOBP_NEGATIVE_REGULATION_OF_CELLULAR_CATABOLIC_PROCESS | 236 | -0.252591 | -0.74243 | 0.962851 | 0.993 |
| GOBP_REGULATION_OF_PROTEIN_DEACETYLATION | 41 | -0.288772 | -0.740695 | 0.877462 | 0.9945 |
| GOBP_BILE_ACID_BIOSYNTHETIC_PROCESS | 33 | -0.294305 | -0.740441 | 0.836197 | 0.99445 |
| GOBP_GLUCOSE_METABOLIC_PROCESS | 195 | -0.253056 | -0.73985 | 0.944835 | 0.994749 |
| GOBP_EMBRYONIC_HEMOPOIESIS | 22 | -0.310369 | -0.739554 | 0.818182 | 0.994732 |
| GOBP_REGULATION_OF_MEMBRANE_PERMEABILITY | 81 | -0.266149 | -0.739464 | 0.899687 | 0.994498 |
| GOBP_REGULATION_OF_TORC1_SIGNALING | 39 | -0.2908 | -0.739273 | 0.849776 | 0.994362 |
| GOBP_PLASMA_LIPOPROTEIN_PARTICLE_CLEARANCE | 47 | -0.278503 | -0.737944 | 0.867925 | 0.995417 |
| GOBP_REGULATION_OF_NUCLEASE_ACTIVITY | 22 | -0.316263 | -0.737435 | 0.822384 | 0.995626 |
| GOBP_VESICLE_BUDDING_FROM_MEMBRANE | 109 | -0.259253 | -0.737222 | 0.919715 | 0.99553 |
| GOBP_FATTY_ACID_BETA_OXIDATION | 72 | -0.266069 | -0.736865 | 0.893572 | 0.995583 |
| GOBP_PYRIMIDINE_NUCLEOSIDE_BIOSYNTHETIC_PROCESS | 16 | -0.328721 | -0.736665 | 0.820876 | 0.995458 |
| GOBP_MICROTUBULE_POLYMERIZATION | 75 | -0.266785 | -0.735806 | 0.898002 | 0.996036 |
| GOBP_REGULATION_OF_ESTABLISHMENT_OF_PROTEIN_LOCALIZATION_TO_M | 72 | -0.264911 | -0.73398 | 0.900104 | 0.997606 |
| GOBP_NUCLEOSIDE_BISPHOSPHATE_BIOSYNTHETIC_PROCESS | 66 | -0.267999 | -0.733017 | 0.883966 | 0.998266 |
| GOBP_NLS_BEARING_PROTEIN_IMPORT_INTO_NUCLEUS | 15 | -0.330015 | -0.732787 | 0.828645 | 0.998167 |
| GOBP_PHOSPHATIDYLSERINE_METABOLIC_PROCESS | 32 | -0.29372 | -0.732009 | 0.858295 | 0.998634 |
| GOBP_PYRIMIDINE_NUCLEOTIDE_BIOSYNTHETIC_PROCESS | 29 | -0.29885 | -0.730829 | 0.854214 | 0.999499 |
| GOBP_NEUTRAL_LIPID_BIOSYNTHETIC_PROCESS | 44 | -0.276915 | -0.73042 | 0.858101 | 0.999572 |
| GOBP_ESTABLISHMENT_OF_PIGMENT_GRANULE_LOCALIZATION | 24 | -0.304385 | -0.730126 | 0.843489 | 0.999535 |
| GOBP_DECIDUALIZATION | 18 | -0.3191 | -0.729818 | 0.810845 | 0.999512 |

| | | | | | |
|---|---|---|---|---|---|
| GOBP_NEGATIVE_REGULATION_OF_INTRINSIC_APOPTOTIC_SIGNALING_PATHW | 88 | -0.258414 | -0.729613 | 0.91875 | 0.999392 |
| GOBP_POSITIVE_REGULATION_OF_PROTEOLYSIS_INVOLVED_IN_CELLULAR_P | 125 | -0.254603 | -0.729491 | 0.936928 | 0.999174 |
| GOBP_FEMALE_MEIOTIC_NUCLEAR_DIVISION | 25 | -0.297996 | -0.729394 | 0.833931 | 0.998934 |
| GOBP_NEGATIVE_REGULATION_OF_CELLULAR_RESPONSE_TO_INSULIN_STIM | 36 | -0.282395 | -0.729068 | 0.850403 | 0.998932 |
| GOBP_NEGATIVE_REGULATION_OF_DIGESTIVE_SYSTEM_PROCESS | 15 | -0.327492 | -0.725584 | 0.81306 | 1 |
| GOBP_CELLULAR_RESPONSE_TO_UV | 86 | -0.260512 | -0.725528 | 0.906054 | 1 |
| GOBP_CELLULAR_LIPID_CATABOLIC_PROCESS | 210 | -0.249027 | -0.724405 | 0.962963 | 1 |
| GOBP_POSITIVE_REGULATION_OF_CELL_CYCLE_PROCESS | 265 | -0.245508 | -0.72412 | 0.981964 | 1 |
| GOBP_GENERATION_OF_PRECURSOR_METABOLITES_AND_ENERGY | 496 | -0.245628 | -0.724005 | 0.996 | 1 |
| GOBP_REGULATION_OF_MACROPHAGE_DERIVED_FOAM_CELL_DIFFERENTIAT | 26 | -0.295549 | -0.722397 | 0.844311 | 1 |
| GOBP_POSITIVE_REGULATION_OF_CELL_CYCLE | 355 | -0.245129 | -0.72028 | 0.990991 | 1 |
| GOBP_COPII_COATED_VESICLE_CARGO_LOADING | 15 | -0.324674 | -0.719413 | 0.830808 | 1 |
| GOBP_POSITIVE_REGULATION_OF_TRANSCRIPTION_OF_NOTCH_RECEPTOR_T | 18 | -0.320184 | -0.718301 | 0.844189 | 1 |
| GOBP_ASPARTATE_FAMILY_AMINO_ACID_BIOSYNTHETIC_PROCESS | 20 | -0.306511 | -0.717966 | 0.840633 | 1 |
| GOBP_REGULATION_OF_ACTIVIN_RECEPTOR_SIGNALING_PATHWAY | 21 | -0.304011 | -0.71688 | 0.858364 | 1 |
| GOBP_POSITIVE_REGULATION_OF_VIRAL_PROCESS | 91 | -0.253991 | -0.71658 | 0.917098 | 1 |
| GOBP_CILIUM_ORGANIZATION | 395 | -0.241848 | -0.715551 | 0.997 | 1 |
| GOBP_PROTEIN_LIPID_COMPLEX_ASSEMBLY | 28 | -0.292513 | -0.715127 | 0.870629 | 1 |
| GOBP_RESPONSE_TO_INCREASED_OXYGEN_LEVELS | 29 | -0.281553 | -0.714808 | 0.866434 | 1 |
| GOBP_PROTEIN_LIPID_COMPLEX_SUBUNIT_ORGANIZATION | 47 | -0.268571 | -0.713453 | 0.899123 | 1 |
| GOBP_CIRCADIAN_REGULATION_OF_GENE_EXPRESSION | 62 | -0.262857 | -0.712702 | 0.891374 | 1 |
| GOBP_REGULATION_OF_POLYSACCHARIDE_BIOSYNTHETIC_PROCESS | 35 | -0.276487 | -0.712595 | 0.864155 | 1 |
| GOBP_CELLULAR_PROCESS_INVOLVED_IN_REPRODUCTION_IN_MULTICELLU | 286 | -0.239758 | -0.712329 | 0.981982 | 1 |
| GOBP_NEGATIVE_REGULATION_OF_GLUCONEOGENESIS | 16 | -0.32045 | -0.712153 | 0.85101 | 1 |
| GOBP_TRANSPOSITION | 27 | -0.285444 | -0.711568 | 0.857647 | 1 |
| GOBP_POSITIVE_REGULATION_OF_CELLULAR_PROTEIN_CATABOLIC_PROCES | 148 | -0.245209 | -0.711449 | 0.9615 | 1 |
| GOBP_MALE_MEIOTIC_NUCLEAR_DIVISION | 39 | -0.275829 | -0.710774 | 0.886493 | 1 |
| GOBP_DETERMINATION_OF_ADULT_LIFESPAN | 16 | -0.317722 | -0.709793 | 0.840412 | 1 |
| GOBP_REGULATION_OF_DNA_DEPENDENT_DNA_REPLICATION | 46 | -0.275242 | -0.708944 | 0.88913 | 1 |
| GOBP_REGULATION_OF_NUCLEAR_DIVISION | 126 | -0.24699 | -0.708838 | 0.952381 | 1 |
| GOBP_NEGATIVE_REGULATION_OF_ORGANELLE_ORGANIZATION | 327 | -0.239942 | -0.708517 | 0.99 | 1 |
| GOBP_POSITIVE_REGULATION_OF_MITOTIC_CELL_CYCLE | 105 | -0.247648 | -0.707746 | 0.920164 | 1 |
| GOBP_CELLULAR_METABOLIC_COMPOUND_SALVAGE | 34 | -0.281519 | -0.706291 | 0.884527 | 1 |
| GOBP_IMPORT_INTO_NUCLEUS | 153 | -0.243417 | -0.70581 | 0.955645 | 1 |
| GOBP_NON_MOTILE_CILIUM_ASSEMBLY | 54 | -0.260575 | -0.70579 | 0.903888 | 1 |
| GOBP_NEGATIVE_REGULATION_OF_PROTEIN_POLYMERIZATION | 74 | -0.256463 | -0.705413 | 0.913044 | 1 |
| GOBP_PROTEIN_LOCALIZATION_TO_CILIUM | 61 | -0.259618 | -0.70539 | 0.904198 | 1 |
| GOBP_INTRINSIC_APOPTOTIC_SIGNALING_PATHWAY_BY_P53_CLASS_MEDIAT | 73 | -0.255301 | -0.704186 | 0.920084 | 1 |
| GOBP_SNARE_COMPLEX_ASSEMBLY | 18 | -0.314208 | -0.703902 | 0.85396 | 1 |
| GOBP_REGULATION_OF_MRNA_PROCESSING | 135 | -0.248203 | -0.703808 | 0.961461 | 1 |
| GOBP_ELECTRON_TRANSPORT_CHAIN | 172 | -0.24266 | -0.703798 | 0.962889 | 1 |
| GOBP_PROTEIN_STABILIZATION | 183 | -0.239123 | -0.703611 | 0.976791 | 1 |
| GOBP_FATTY_ACYL_COA_BIOSYNTHETIC_PROCESS | 31 | -0.280729 | -0.703469 | 0.890058 | 1 |
| GOBP_KERATINOCYTE_PROLIFERATION | 37 | -0.275934 | -0.702268 | 0.888889 | 1 |
| GOBP_ETHER_METABOLIC_PROCESS | 26 | -0.283816 | -0.702056 | 0.874266 | 1 |
| GOBP_NEGATIVE_REGULATION_OF_ATP_METABOLIC_PROCESS | 27 | -0.284465 | -0.701508 | 0.886311 | 1 |
| GOBP_AMP_METABOLIC_PROCESS | 16 | -0.310761 | -0.701297 | 0.849176 | 1 |
| GOBP_GLYCOSYL_COMPOUND_BIOSYNTHETIC_PROCESS | 42 | -0.269532 | -0.701149 | 0.890889 | 1 |
| GOBP_GLUCAN_BIOSYNTHETIC_PROCESS | 43 | -0.266457 | -0.700743 | 0.897492 | 1 |
| GOBP_POSITIVE_REGULATION_OF_DNA_REPAIR | 72 | -0.254127 | -0.700732 | 0.931106 | 1 |
| GOBP_REGULATION_OF_HIPPO_SIGNALING | 19 | -0.303186 | -0.700234 | 0.865672 | 1 |
| GOBP_NEGATIVE_REGULATION_OF_TUMOR_NECROSIS_FACTOR_MEDIATED_S | 18 | -0.302985 | -0.699874 | 0.866499 | 1 |
| GOBP_NEGATIVE_REGULATION_OF_OXIDATIVE_STRESS_INDUCED_CELL_DEA | 39 | -0.271503 | -0.699174 | 0.894737 | 1 |
| GOBP_REGULATION_OF_CHOLESTEROL_METABOLIC_PROCESS | 56 | -0.259436 | -0.698449 | 0.905844 | 1 |
| GOBP_POSITIVE_REGULATION_OF_POSTTRANSCRIPTIONAL_GENE_SILENCING | 29 | -0.2809 | -0.694775 | 0.864583 | 1 |
| GOBP_RAP_PROTEIN_SIGNAL_TRANSDUCTION | 15 | -0.314447 | -0.694135 | 0.879333 | 1 |
| GOBP_PIGMENT_GRANULE_LOCALIZATION | 26 | -0.287116 | -0.692977 | 0.880866 | 1 |
| GOBP_MICROTUBULE_POLYMERIZATION_OR_DEPOLYMERIZATION | 111 | -0.244213 | -0.692975 | 0.935451 | 1 |
| GOBP_MUSCLE_CELL_CELLULAR_HOMEOSTASIS | 19 | -0.29909 | -0.692784 | 0.862864 | 1 |
| GOBP_ENDOSOME_ORGANIZATION | 80 | -0.249114 | -0.691749 | 0.926931 | 1 |
| GOBP_STEROL_METABOLIC_PROCESS | 149 | -0.240087 | -0.690852 | 0.962739 | 1 |
| GOBP_MODULATION_BY_VIRUS_OF_HOST_CELLULAR_PROCESS | 17 | -0.306878 | -0.69064 | 0.871891 | 1 |
| GOBP_REGULATION_OF_HYDROGEN_PEROXIDE_METABOLIC_PROCESS | 19 | -0.300242 | -0.689654 | 0.868159 | 1 |
| GOBP_REGULATION_OF_PLASMA_LIPOPROTEIN_PARTICLE_LEVELS | 77 | -0.248701 | -0.689274 | 0.942348 | 1 |
| GOBP_REGULATION_OF_MICROTUBULE_BASED_PROCESS | 216 | -0.236581 | -0.688587 | 0.981982 | 1 |
| GOBP_POSITIVE_REGULATION_OF_RELEASE_OF_CYTOCHROME_C_FROM_MIT | 27 | -0.280864 | -0.688021 | 0.873536 | 1 |
| GOBP_TETRAHYDROFOLATE_METABOLIC_PROCESS | 19 | -0.301495 | -0.688019 | 0.850835 | 1 |
| GOBP_GOLGI_TO_VACUOLE_TRANSPORT | 19 | -0.29911 | -0.687851 | 0.859278 | 1 |
| GOBP_NEGATIVE_REGULATION_OF_VASCULAR_ASSOCIATED_SMOOTH_MUSC | 19 | -0.294392 | -0.687835 | 0.877929 | 1 |
| GOBP_POSITIVE_REGULATION_OF_CELLULAR_AMIDE_METABOLIC_PROCESS | 149 | -0.236304 | -0.687757 | 0.974748 | 1 |
| GOBP_REGULATION_OF_TRANSCRIPTION_BY_RNA_POLYMERASE_III | 23 | -0.287647 | -0.687512 | 0.898409 | 1 |
| GOBP_RETROGRADE_TRANSPORT_ENDOSOME_TO_PLASMA_MEMBRANE | 31 | -0.273747 | -0.68487 | 0.893816 | 1 |
| GOBP_NADPH_REGENERATION | 17 | -0.307492 | -0.683864 | 0.88539 | 1 |
| GOBP_NEGATIVE_REGULATION_OF_CARBOHYDRATE_METABOLIC_PROCESS | 46 | -0.262124 | -0.683433 | 0.888521 | 1 |
| GOBP_CELL_CYCLE_G1_S_PHASE_TRANSITION | 260 | -0.231531 | -0.683186 | 0.99 | 1 |
| GOBP_MONOSACCHARIDE_CATABOLIC_PROCESS | 57 | -0.255817 | -0.682174 | 0.926327 | 1 |
| GOBP_REGULATION_OF_RESPONSE_TO_DNA_DAMAGE_STIMULUS | 218 | -0.23087 | -0.678306 | 0.986987 | 1 |
| GOBP_REGULATION_OF_CELLULAR_AMIDE_METABOLIC_PROCESS | 410 | -0.22911 | -0.677261 | 0.998 | 1 |
| GOBP_REGULATION_OF_HISTONE_METHYLATION | 58 | -0.248879 | -0.677085 | 0.924731 | 1 |
| GOBP_NEGATIVE_REGULATION_OF_MITOCHONDRION_ORGANIZATION | 44 | -0.257408 | -0.676409 | 0.915755 | 1 |
| GOBP_CELL_DEATH_IN_RESPONSE_TO_OXIDATIVE_STRESS | 77 | -0.242636 | -0.676331 | 0.947479 | 1 |
| GOBP_POSITIVE_REGULATION_OF_MRNA_METABOLIC_PROCESS | 78 | -0.241807 | -0.676236 | 0.93886 | 1 |
| GOBP_PORPHYRIN_CONTAINING_COMPOUND_METABOLIC_PROCESS | 37 | -0.263929 | -0.674905 | 0.915368 | 1 |
| GOBP_NEGATIVE_REGULATION_OF_HISTONE_MODIFICATION | 43 | -0.256351 | -0.674846 | 0.919683 | 1 |
| GOBP_GLYCOSYLCERAMIDE_METABOLIC_PROCESS | 17 | -0.299458 | -0.674564 | 0.876106 | 1 |
| GOBP_NUCLEOTIDE_TRANSPORT | 29 | -0.273235 | -0.674559 | 0.900685 | 1 |
| GOBP_SIGNAL_TRANSDUCTION_IN_RESPONSE_TO_DNA_DAMAGE | 127 | -0.235251 | -0.673646 | 0.971602 | 1 |
| GOBP_POSITIVE_REGULATION_OF_MRNA_CATABOLIC_PROCESS | 50 | -0.253454 | -0.673105 | 0.931744 | 1 |
| GOBP_POSITIVE_REGULATION_OF_MITOCHONDRIAL_OUTER_MEMBRANE_PEF | 35 | -0.26433 | -0.672473 | 0.913694 | 1 |
| GOBP_POSITIVE_REGULATION_OF_VIRAL_GENOME_REPLICATION | 31 | -0.268492 | -0.671338 | 0.915691 | 1 |
| GOBP_POSITIVE_REGULATION_OF_EXTRINSIC_APOPTOTIC_SIGNALING_PATHV | 45 | -0.256588 | -0.670919 | 0.922149 | 1 |
| GOBP_PEPTIDYL_L_CYSTEINE_S_PALMITOYLATION | 24 | -0.280884 | -0.670785 | 0.90942 | 1 |
| GOBP_ESTABLISHMENT_OF_PROTEIN_LOCALIZATION_TO_MEMBRANE | 343 | -0.225736 | -0.669863 | 0.997 | 1 |
| GOBP_NEGATIVE_REGULATION_OF_PROGRAMMED_NECROTIC_CELL_DEATH | 15 | -0.298092 | -0.669417 | 0.883963 | 1 |
| GOBP_REGULATION_OF_MITOCHONDRION_ORGANIZATION | 148 | -0.229744 | -0.666857 | 0.971602 | 1 |
| GOBP_HISTONE_H3_K4_METHYLATION | 49 | -0.250036 | -0.666338 | 0.934549 | 1 |
| GOBP_CARBOHYDRATE_BIOSYNTHETIC_PROCESS | 193 | -0.227638 | -0.666096 | 0.986935 | 1 |
| GOBP_REGULATION_OF_RNA_SPLICING | 139 | -0.231933 | -0.664729 | 0.969666 | 1 |
| GOBP_NEGATIVE_REGULATION_OF_PROTEIN_TYROSINE_KINASE_ACTIVITY | 30 | -0.268364 | -0.663347 | 0.897436 | 1 |
| GOBP_POSITIVE_REGULATION_OF_LIPID_STORAGE | 24 | -0.274897 | -0.662848 | 0.905882 | 1 |
| GOBP_CHOLESTEROL_STORAGE | 18 | -0.289713 | -0.662599 | 0.883019 | 1 |
| GOBP_ENDOPLASMIC_RETICULUM_CALCIUM_ION_HOMEOSTASIS | 25 | -0.276801 | -0.662255 | 0.917661 | 1 |
| GOBP_REGULATION_OF_TRANSLATIONAL_INITIATION_IN_RESPONSE_TO_STR | 15 | -0.299891 | -0.661132 | 0.890625 | 1 |
| GOBP_REGULATION_OF_UBIQUITIN_PROTEIN_LIGASE_ACTIVITY | 23 | -0.277768 | -0.661029 | 0.90122 | 1 |
| GOBP_HISTONE_H3_K9_METHYLATION | 33 | -0.260288 | -0.660991 | 0.909194 | 1 |
| GOBP_MAINTENANCE_OF_PROTEIN_LOCATION | 89 | -0.236385 | -0.658139 | 0.962461 | 1 |
| GOBP_DNA_METHYLATION_OR_DEMETHYLATION | 83 | -0.234318 | -0.657125 | 0.958115 | 1 |
| GOBP_LIPOPROTEIN_METABOLIC_PROCESS | 126 | -0.229992 | -0.656851 | 0.975659 | 1 |
| GOBP_ENDOPLASMIC_RETICULUM_ORGANIZATION | 83 | -0.235559 | -0.655312 | 0.954498 | 1 |
| GOBP_PROTEIN_LOCALIZATION_TO_MICROTUBULE | 18 | -0.290325 | -0.654181 | 0.90617 | 1 |
| GOBP_TRANSCRIPTION_INITIATION_FROM_RNA_POLYMERASE_II_PROMOTER | 180 | -0.225524 | -0.652732 | 0.98994 | 1 |
| GOBP_REGULATION_OF_VASCULOGENESIS | 16 | -0.293059 | -0.652296 | 0.892365 | 1 |
| GOBP_PURINE_NUCLEOTIDE_TRANSPORT | 23 | -0.273072 | -0.652127 | 0.907363 | 1 |
| GOBP_ENDOCYTIC_RECYCLING | 48 | -0.244983 | -0.65119 | 0.938998 | 1 |
| GOBP_NEGATIVE_REGULATION_OF_UBIQUITIN_PROTEIN_TRANSFERASE_ACT | 16 | -0.292803 | -0.650805 | 0.906716 | 1 |
| GOBP_FATTY_ACYL_COA_METABOLIC_PROCESS | 41 | -0.253525 | -0.650501 | 0.935094 | 1 |
| GOBP_REGULATION_OF_RUFFLE_ASSEMBLY | 28 | -0.26627 | -0.650169 | 0.923523 | 1 |
| GOBP_POSITIVE_REGULATION_OF_PROTEIN_MODIFICATION_BY_SMALL_PROT | 132 | -0.228092 | -0.649965 | 0.984833 | 1 |
| GOBP_SPERMATID_DIFFERENTIATION | 128 | -0.226086 | -0.64914 | 0.973737 | 1 |
| GOBP_PROTEIN_O_LINKED_GLYCOSYLATION | 98 | -0.230583 | -0.649128 | 0.970954 | 1 |
| GOBP_VESICLE_TRANSPORT_ALONG_MICROTUBULE | 46 | -0.246381 | -0.647715 | 0.942242 | 1 |
| GOBP_MALE_GAMETE_GENERATION | 452 | -0.217007 | -0.646811 | 0.999 | 1 |
| GOBP_REGULATION_OF_STEM_CELL_PROLIFERATION | 36 | -0.251158 | -0.644977 | 0.924571 | 1 |
| GOBP_RESPONSE_TO_UV | 142 | -0.225534 | -0.644477 | 0.982846 | 1 |
| GOBP_REGULATION_OF_CELL_CYCLE_ARREST | 100 | -0.227736 | -0.644438 | 0.975155 | 1 |
| GOBP_NEGATIVE_REGULATION_OF_HISTONE_ACETYLATION | 16 | -0.289762 | -0.644414 | 0.926199 | 1 |
| GOBP_ENDOSOMAL_TRANSPORT | 239 | -0.218415 | -0.644409 | 0.995992 | 1 |
| GOBP_ORGANOPHOSPHATE_CATABOLIC_PROCESS | 147 | -0.222463 | -0.642495 | 0.983822 | 1 |
| GOBP_REGULATION_OF_ANOIKIS | 23 | -0.268414 | -0.64244 | 0.91352 | 1 |
| GOBP_POSITIVE_REGULATION_OF_BEHAVIOR | 18 | -0.280686 | -0.642284 | 0.901619 | 1 |
| GOBP_REGULATION_OF_PROTEIN_TARGETING_TO_MITOCHONDRION | 44 | -0.246454 | -0.641798 | 0.943771 | 1 |
| GOBP_MICROTUBULE_BASED_MOVEMENT | 341 | -0.218197 | -0.641279 | 1 | 1 |
| GOBP_APOPTOTIC_MITOCHONDRIAL_CHANGES | 112 | -0.226645 | -0.640856 | 0.980473 | 1 |
| GOBP_RESPONSE_TO_FLUID_SHEAR_STRESS | 29 | -0.257475 | -0.640424 | 0.938515 | 1 |
| GOBP_MITOTIC_CELL_CYCLE_ARREST | 16 | -0.284227 | -0.6379 | 0.912669 | 1 |
| GOBP_POSITIVE_REGULATION_OF_ESTABLISHMENT_OF_PROTEIN_LOCALIZAT | 58 | -0.234799 | -0.637493 | 0.952026 | 1 |
| GOBP_HEMOGLOBIN_METABOLIC_PROCESS | 16 | -0.27844 | -0.637224 | 0.912258 | 1 |
| GOBP_POLYSACCHARIDE_BIOSYNTHETIC_PROCESS | 72 | -0.230133 | -0.636121 | 0.966562 | 1 |
| GOBP_ESTABLISHMENT_OF_PROTEIN_LOCALIZATION_TO_MITOCHONDRIAL_M | 53 | -0.23706 | -0.634834 | 0.961538 | 1 |
| GOBP_PROTEIN_POLYUBIQUITINATION | 330 | -0.214447 | -0.63377 | 0.997 | 1 |
| GOBP_REGULATION_OF_SPINDLE_CHECKPOINT | 15 | -0.286967 | -0.632372 | 0.919023 | 1 |
| GOBP_REGULATION_OF_ATPASE_ACTIVITY | 79 | -0.228146 | -0.630551 | 0.972107 | 1 |
| GOBP_NEGATIVE_REGULATION_OF_CELLULAR_SENESCENCE | 19 | -0.27197 | -0.629269 | 0.924482 | 1 |
| GOBP_GERM_CELL_DEVELOPMENT | 210 | -0.214808 | -0.626419 | 0.994985 | 1 |
| GOBP_POSITIVE_REGULATION_OF_UBIQUITIN_DEPENDENT_PROTEIN_CATABC | 102 | -0.224166 | -0.625306 | 0.984488 | 1 |
| GOBP_PYRIMIDINE_CONTAINING_COMPOUND_BIOSYNTHETIC_PROCESS | 39 | -0.24328 | -0.625285 | 0.94363 | 1 |
| GOBP_REGULATION_OF_DNA_METABOLIC_PROCESS | 333 | -0.212018 | -0.625037 | 0.999 | 1 |
| GOBP_LUNG_EPITHELIUM_DEVELOPMENT | 30 | -0.254767 | -0.625012 | 0.946324 | 1 |
| GOBP_MODULATION_BY_VIRUS_OF_HOST_PROCESS | 27 | -0.25588 | -0.62482 | 0.938156 | 1 |
| GOBP_DEOXYRIBONUCLEOSIDE_TRIPHOSPHATE_METABOLIC_PROCESS | 15 | -0.281291 | -0.62452 | 0.918367 | 1 |
| GOBP_POSITIVE_REGULATION_OF_TORC1_SIGNALING | 17 | -0.277988 | -0.623433 | 0.922693 | 1 |

| | | | | | |
|---|---|---|---|---|---|
| GOBP_INTRINSIC_APOPTOTIC_SIGNALING_PATHWAY_IN_RESPONSE_TO_END | 57 | -0.229328 | -0.623095 | 0.964247 | 1 |
| GOBP_NUCLEOBASE_CONTAINING_SMALL_MOLECULE_BIOSYNTHETIC_PROC | 112 | -0.22187 | -0.621857 | 0.979487 | 1 |
| GOBP_PTERIDINE_CONTAINING_COMPOUND_BIOSYNTHETIC_PROCESS | 15 | -0.28301 | -0.621605 | 0.906209 | 1 |
| GOBP_REGULATION_OF_TRANSLATIONAL_INITIATION | 74 | -0.222353 | -0.621508 | 0.967505 | 1 |
| GOBP_REGULATION_OF_TRANSLATION_NCRNA_MEDIATED | 25 | -0.257815 | -0.621504 | 0.937956 | 1 |
| GOBP_NEGATIVE_REGULATION_OF_ACTIN_FILAMENT_POLYMERIZATION | 60 | -0.230946 | -0.620555 | 0.969762 | 1 |
| GOBP_REGULATION_OF_LIPOPROTEIN_LIPASE_ACTIVITY | 19 | -0.267331 | -0.620312 | 0.927591 | 1 |
| GOBP_GTP_METABOLIC_PROCESS | 24 | -0.254563 | -0.618718 | 0.945755 | 1 |
| GOBP_AUTOPHAGOSOME_ORGANIZATION | 95 | -0.219297 | -0.618518 | 0.978528 | 1 |
| GOBP_NEGATIVE_REGULATION_OF_CYCLIN_DEPENDENT_PROTEIN_KINASE_A | 30 | -0.248406 | -0.616862 | 0.952928 | 1 |
| GOBP_POSITIVE_REGULATION_OF_DNA_BIOSYNTHETIC_PROCESS | 65 | -0.223734 | -0.615843 | 0.964744 | 1 |
| GOBP_PROTEIN_FOLDING | 217 | -0.209682 | -0.615099 | 0.996994 | 1 |
| GOBP_REGULATION_OF_INTRACELLULAR_ESTROGEN_RECEPTOR_SIGNALING | 39 | -0.239949 | -0.614318 | 0.945495 | 1 |
| GOBP_PURINE_CONTAINING_COMPOUND_SALVAGE | 15 | -0.281562 | -0.61301 | 0.93947 | 1 |
| GOBP_TRNA_THREONYLCARBAMOYLADENOSINE_METABOLIC_PROCESS | 15 | -0.276651 | -0.612234 | 0.935233 | 1 |
| GOBP_NEGATIVE_REGULATION_OF_BLOOD_VESSEL_ENDOTHELIAL_CELL_MIG | 33 | -0.242488 | -0.611751 | 0.961231 | 1 |
| GOBP_REGULATION_OF_CYCLIN_DEPENDENT_PROTEIN_KINASE_ACTIVITY | 101 | -0.217662 | -0.611716 | 0.984488 | 1 |
| GOBP_FATTY_ACID_CATABOLIC_PROCESS | 101 | -0.214856 | -0.607286 | 0.988648 | 1 |
| GOBP_MITOCHONDRIAL_OUTER_MEMBRANE_PERMEABILIZATION | 55 | -0.225918 | -0.605621 | 0.969828 | 1 |
| GOBP_PROTEIN_LOCALIZATION_TO_CYTOSKELETON | 58 | -0.22346 | -0.604048 | 0.974138 | 1 |
| GOBP_NEGATIVE_REGULATION_OF_CELL_CYCLE_ARREST | 20 | -0.259749 | -0.60365 | 0.943489 | 1 |
| GOBP_NEGATIVE_REGULATION_OF_CELL_CYCLE_G1_S_PHASE_TRANSITION | 102 | -0.214062 | -0.603284 | 0.982546 | 1 |
| GOBP_PROTEIN_REFOLDING | 23 | -0.24919 | -0.599311 | 0.934307 | 1 |
| GOBP_MAINTENANCE_OF_PROTEIN_LOCATION_IN_CELL | 62 | -0.219954 | -0.598831 | 0.976571 | 1 |
| GOBP_REGULATION_OF_MRNA_METABOLIC_PROCESS | 317 | -0.201248 | -0.597069 | 0.999 | 1 |
| GOBP_REGULATION_OF_DNA_REPLICATION | 101 | -0.21167 | -0.596377 | 0.987692 | 1 |
| GOBP_POSITIVE_REGULATION_OF_TRANSCRIPTION_INITIATION_FROM_RNA_P | 26 | -0.245089 | -0.595441 | 0.961894 | 1 |
| GOBP_PROTEIN_K63_LINKED_DEUBIQUITINATION | 34 | -0.233591 | -0.595302 | 0.954442 | 1 |
| GOBP_CELLULAR_RESPONSE_TO_ATP | 17 | -0.26669 | -0.595172 | 0.930263 | 1 |
| GOBP_NEGATIVE_REGULATION_OF_MITOTIC_CELL_CYCLE | 294 | -0.203504 | -0.594757 | 0.998 | 1 |
| GOBP_CELLULAR_RESPONSE_TO_GAMMA_RADIATION | 27 | -0.240649 | -0.594595 | 0.960432 | 1 |
| GOBP_CELLULAR_RESPONSE_TO_GROWTH_HORMONE_STIMULUS | 21 | -0.254934 | -0.594528 | 0.948298 | 1 |
| GOBP_REGULATION_OF_CILIUM_MOVEMENT | 25 | -0.244709 | -0.594269 | 0.948565 | 1 |
| GOBP_DNA_DAMAGE_RESPONSE_SIGNAL_TRANSDUCTION_RESULTING_IN_T | 18 | -0.255828 | -0.592118 | 0.954315 | 1 |
| GOBP_POSITIVE_REGULATION_OF_CELLULAR_RESPONSE_TO_TRANSFORMIN | 27 | -0.246091 | -0.59093 | 0.961677 | 1 |
| GOBP_REGULATION_OF_OXIDATIVE_STRESS_INDUCED_CELL_DEATH | 56 | -0.221195 | -0.590401 | 0.979548 | 1 |
| GOBP_ANOIKIS | 33 | -0.233934 | -0.589984 | 0.95672 | 1 |
| GOBP_REGULATION_OF_GLUCONEOGENESIS | 42 | -0.22678 | -0.589863 | 0.973214 | 1 |
| GOBP_DNA_MODIFICATION | 103 | -0.208187 | -0.589371 | 0.987642 | 1 |
| GOBP_POSITIVE_REGULATION_OF_TRANSLATION | 122 | -0.205749 | -0.589118 | 0.99496 | 1 |
| GOBP_PROTEIN_IMPORT | 184 | -0.202448 | -0.588202 | 0.995988 | 1 |
| GOBP_POSITIVE_REGULATION_OF_MRNA_SPLICING_VIA_SPLICEOSOME | 20 | -0.252388 | -0.587128 | 0.94625 | 1 |
| GOBP_REGULATION_OF_CHROMATIN_ORGANIZATION | 175 | -0.201972 | -0.584876 | 0.99799 | 1 |
| GOBP_NEGATIVE_REGULATION_OF_PLATELET_ACTIVATION | 16 | -0.265024 | -0.58316 | 0.958438 | 1 |
| GOBP_PROTEIN_K48_LINKED_DEUBIQUITINATION | 34 | -0.230689 | -0.58302 | 0.971494 | 1 |
| GOBP_GOLGI_TO_PLASMA_MEMBRANE_TRANSPORT | 57 | -0.217065 | -0.58256 | 0.988197 | 1 |
| GOBP_REGULATION_OF_ENDOPLASMIC_RETICULUM_STRESS_INDUCED_INTR | 27 | -0.239925 | -0.582528 | 0.961905 | 1 |
| GOBP_POSITIVE_REGULATION_OF_CHOLESTEROL_EFFLUX | 20 | -0.248804 | -0.582509 | 0.948148 | 1 |
| GOBP_DORSAL_VENTRAL_NEURAL_TUBE_PATTERNING | 20 | -0.249118 | -0.580425 | 0.959509 | 1 |
| GOBP_RNA_POLYMERASE_II_PREINITIATION_COMPLEX_ASSEMBLY | 23 | -0.243661 | -0.579437 | 0.951673 | 1 |
| GOBP_REGULATION_OF_CELLULAR_PROTEIN_CATABOLIC_PROCESS | 245 | -0.196716 | -0.579253 | 0.999 | 1 |
| GOBP_RESPONSE_TO_MISFOLDED_PROTEIN | 24 | -0.24365 | -0.579233 | 0.963986 | 1 |
| GOBP_PEROXISOME_ORGANIZATION | 82 | -0.20449 | -0.575589 | 0.990645 | 1 |
| GOBP_3_UTR_MEDIATED_MRNA_STABILIZATION | 18 | -0.255121 | -0.57558 | 0.95561 | 1 |
| GOBP_NEGATIVE_REGULATION_OF_PROTEIN_CATABOLIC_PROCESS | 114 | -0.202336 | -0.575227 | 0.993933 | 1 |
| GOBP_CELLULAR_CARBOHYDRATE_BIOSYNTHETIC_PROCESS | 79 | -0.205657 | -0.574127 | 0.991753 | 1 |
| GOBP_TRNA_WOBBLE_BASE_MODIFICATION | 20 | -0.24849 | -0.573839 | 0.956151 | 1 |
| GOBP_HISTONE_METHYLATION | 124 | -0.200045 | -0.57382 | 0.998986 | 1 |
| GOBP_MRNA_SPLICE_SITE_SELECTION | 38 | -0.222085 | -0.573562 | 0.970721 | 1 |
| GOBP_MEGAKARYOCYTE_DIFFERENTIATION | 88 | -0.202577 | -0.573084 | 0.99375 | 1 |
| GOBP_POSITIVE_REGULATION_OF_RUFFLE_ASSEMBLY | 15 | -0.256701 | -0.572559 | 0.951407 | 1 |
| GOBP_KINETOCHORE_ASSEMBLY | 17 | -0.251969 | -0.572221 | 0.951899 | 1 |
| GOBP_POSITIVE_REGULATION_OF_MRNA_PROCESSING | 30 | -0.231025 | -0.570607 | 0.966122 | 1 |
| GOBP_INTERMEMBRANE_LIPID_TRANSFER | 44 | -0.2211 | -0.568585 | 0.982262 | 1 |
| GOBP_REGULATION_OF_CELL_CYCLE_PHASE_TRANSITION | 433 | -0.190961 | -0.568011 | 1 | 1 |
| GOBP_RESPONSE_TO_GROWTH_HORMONE | 34 | -0.223593 | -0.567173 | 0.968362 | 1 |
| GOBP_SPERMATID_NUCLEUS_DIFFERENTIATION | 15 | -0.258115 | -0.567167 | 0.961735 | 1 |
| GOBP_PEPTIDYL_THREONINE_DEPHOSPHORYLATION | 22 | -0.240801 | -0.567112 | 0.958688 | 1 |
| GOBP_INTEGRATED_STRESS_RESPONSE_SIGNALING | 25 | -0.230192 | -0.564258 | 0.973269 | 1 |
| GOBP_HEPATOCYTE_GROWTH_FACTOR_RECEPTOR_SIGNALING_PATHWAY | 16 | -0.252829 | -0.56327 | 0.946716 | 1 |
| GOBP_REGULATION_OF_GENE_SILENCING_BY_RNA | 109 | -0.196935 | -0.560537 | 0.997955 | 1 |
| GOBP_REGULATION_OF_PROTEIN_MODIFICATION_BY_SMALL_PROTEIN_CONJ | 230 | -0.19124 | -0.559925 | 0.998999 | 1 |
| GOBP_NEGATIVE_REGULATION_OF_PEPTIDYL_LYSINE_ACETYLATION | 19 | -0.243129 | -0.559674 | 0.965686 | 1 |
| GOBP_NEGATIVE_REGULATION_OF_DNA_BIOSYNTHETIC_PROCESS | 37 | -0.216312 | -0.559355 | 0.988739 | 1 |
| GOBP_REGULATION_OF_HISTONE_H3_K9_METHYLATION | 20 | -0.242341 | -0.558718 | 0.970803 | 1 |
| GOBP_POSITIVE_REGULATION_OF_GLUCOSE_IMPORT | 31 | -0.219412 | -0.557571 | 0.973068 | 1 |
| GOBP_NEGATIVE_REGULATION_OF_RESPONSE_TO_DNA_DAMAGE_STIMULUS | 82 | -0.198461 | -0.557311 | 0.995872 | 1 |
| GOBP_REGULATION_OF_TRANSPOSITION | 23 | -0.233342 | -0.556655 | 0.96337 | 1 |
| GOBP_POSITIVE_REGULATION_OF_DNA_TEMPLATED_TRANSCRIPTION_INITIAT | 33 | -0.221461 | -0.556297 | 0.984018 | 1 |
| GOBP_NEGATIVE_REGULATION_OF_GENE_SILENCING | 30 | -0.219774 | -0.55469 | 0.986127 | 1 |
| GOBP_PEROXISOMAL_TRANSPORT | 70 | -0.201118 | -0.554034 | 0.989328 | 1 |
| GOBP_DNA_DEALKYLATION | 30 | -0.221005 | -0.552689 | 0.974239 | 1 |
| GOBP_REGULATION_OF_DOUBLE_STRAND_BREAK_REPAIR_VIA_NONHOMOLO | 28 | -0.225715 | -0.551111 | 0.976998 | 1 |
| GOBP_RESPONSE_TO_IONIZING_RADIATION | 141 | -0.189764 | -0.551055 | 0.998989 | 1 |
| GOBP_DNA_SYNTHESIS_INVOLVED_IN_DNA_REPAIR | 51 | -0.205698 | -0.55091 | 0.991304 | 1 |
| GOBP_GPI_ANCHOR_METABOLIC_PROCESS | 32 | -0.212875 | -0.549625 | 0.975207 | 1 |
| GOBP_PEPTIDYL_LYSINE_ACETYLATION | 164 | -0.187209 | -0.548559 | 1 | 1 |
| GOBP_INTRACELLULAR_ESTROGEN_RECEPTOR_SIGNALING_PATHWAY | 53 | -0.204982 | -0.548482 | 0.985931 | 1 |
| GOBP_TRICARBOXYLIC_ACID_CYCLE | 33 | -0.21698 | -0.548376 | 0.987327 | 1 |
| GOBP_DNA_LIGATION | 15 | -0.248301 | -0.547989 | 0.973008 | 1 |
| GOBP_GLUTAMINE_FAMILY_AMINO_ACID_BIOSYNTHETIC_PROCESS | 16 | -0.245065 | -0.544755 | 0.969309 | 1 |
| GOBP_REGULATION_OF_DNA_BIOSYNTHETIC_PROCESS | 105 | -0.191988 | -0.542274 | 0.997942 | 1 |
| GOBP_VIRAL_RNA_GENOME_REPLICATION | 33 | -0.214826 | -0.542193 | 0.980769 | 1 |
| GOBP_PROTEIN_K48_LINKED_UBIQUITINATION | 61 | -0.199226 | -0.541386 | 0.990312 | 1 |
| GOBP_NEGATIVE_REGULATION_OF_RESPONSE_TO_ENDOPLASMIC_RETICULU | 41 | -0.208651 | -0.541015 | 0.987832 | 1 |
| GOBP_UBIQUINONE_METABOLIC_PROCESS | 18 | -0.233842 | -0.540933 | 0.964417 | 1 |
| GOBP_MAINTENANCE_OF_PROTEIN_LOCALIZATION_IN_ORGANELLE | 41 | -0.209812 | -0.539642 | 0.988827 | 1 |
| GOBP_POSITIVE_REGULATION_OF_VASCULAR_PERMEABILITY | 15 | -0.244219 | -0.538547 | 0.967062 | 1 |
| GOBP_DNA_DOUBLE_STRAND_BREAK_PROCESSING | 24 | -0.224813 | -0.53854 | 0.97561 | 1 |
| GOBP_MATERNAL_PLACENTA_DEVELOPMENT | 26 | -0.22483 | -0.538359 | 0.974668 | 1 |
| GOBP_ESTABLISHMENT_OF_PROTEIN_LOCALIZATION_TO_TELOMERE | 18 | -0.23556 | -0.538199 | 0.972603 | 1 |
| GOBP_DNA_STRAND_ELONGATION_INVOLVED_IN_DNA_REPLICATION | 19 | -0.235704 | -0.53621 | 0.971985 | 1 |
| GOBP_DNA_BIOSYNTHETIC_PROCESS | 183 | -0.184641 | -0.535071 | 0.998995 | 1 |
| GOBP_NEGATIVE_REGULATION_OF_PROTEIN_ACETYLATION | 21 | -0.227574 | -0.534016 | 0.985594 | 1 |
| GOBP_CELLULAR_PROTEIN_COMPLEX_DISASSEMBLY | 217 | -0.183455 | -0.533357 | 1 | 1 |
| GOBP_NUCLEAR_TRANSPORT | 329 | -0.179389 | -0.53225 | 1 | 1 |
| GOBP_DNA_DEPENDENT_DNA_REPLICATION_MAINTENANCE_OF_FIDELITY | 48 | -0.20192 | -0.531182 | 0.997802 | 1 |
| GOBP_CELLULAR_RESPONSE_TO_IONIZING_RADIATION | 64 | -0.195144 | -0.531126 | 0.99467 | 1 |
| GOBP_NEGATIVE_REGULATION_OF_CELL_AGING | 24 | -0.219742 | -0.529272 | 0.97343 | 1 |
| GOBP_REGULATION_OF_GOLGI_ORGANIZATION | 17 | -0.23407 | -0.527652 | 0.967702 | 1 |
| GOBP_GLUCOSE_IMPORT | 63 | -0.194892 | -0.526872 | 0.995704 | 1 |
| GOBP_SINGLE_STRANDED_VIRAL_RNA_REPLICATION_VIA_DOUBLE_STRANDE | 19 | -0.228081 | -0.525742 | 0.981527 | 1 |
| GOBP_PROTEASOMAL_PROTEIN_CATABOLIC_PROCESS | 472 | -0.176561 | -0.525295 | 1 | 1 |
| GOBP_DNA_STRAND_ELONGATION | 26 | -0.216319 | -0.5245 | 0.983353 | 1 |
| GOBP_PROTEIN_ACETYLATION | 190 | -0.177994 | -0.519197 | 1 | 1 |
| GOBP_HIPPO_SIGNALING | 45 | -0.198267 | -0.518725 | 0.996681 | 1 |
| GOBP_CELL_CYCLE_G2_M_PHASE_TRANSITION | 270 | -0.175994 | -0.518422 | 1 | 1 |
| GOBP_NEGATIVE_REGULATION_OF_PROTEIN_MODIFICATION_BY_SMALL_PRO | 87 | -0.185688 | -0.518393 | 1 | 1 |
| GOBP_NUCLEOTIDE_SUGAR_METABOLIC_PROCESS | 37 | -0.199882 | -0.517549 | 0.988439 | 1 |
| GOBP_DNA_TEMPLATED_TRANSCRIPTION_INITIATION | 236 | -0.176834 | -0.515971 | 1 | 1 |
| GOBP_DNA_DEMETHYLATION | 24 | -0.213576 | -0.514488 | 0.977273 | 1 |
| GOBP_HISTONE_H3_K9_MODIFICATION | 44 | -0.197533 | -0.514487 | 0.99557 | 1 |
| GOBP_POSITIVE_REGULATION_OF_CHROMATIN_ORGANIZATION | 98 | -0.181382 | -0.511498 | 1 | 1 |
| GOBP_NADH_METABOLIC_PROCESS | 42 | -0.195943 | -0.511275 | 0.99442 | 1 |
| GOBP_REGULATION_OF_TRANSLATION_IN_RESPONSE_TO_STRESS | 21 | -0.218803 | -0.510533 | 0.987261 | 1 |
| GOBP_PROTEIN_TARGETING | 429 | -0.171227 | -0.509262 | 1 | 1 |
| GOBP_ANAPHASE_PROMOTING_COMPLEX_DEPENDENT_CATABOLIC_PROCES | 81 | -0.184389 | -0.50902 | 0.995833 | 1 |
| GOBP_MULTI_ORGANISM_LOCALIZATION | 64 | -0.185225 | -0.507448 | 0.994647 | 1 |
| GOBP_DNA_RECOMBINATION | 281 | -0.171636 | -0.506034 | 1 | 1 |
| GOBP_NEGATIVE_REGULATION_OF_FATTY_ACID_METABOLIC_PROCESS | 24 | -0.211312 | -0.5059 | 0.987937 | 1 |
| GOBP_SPERM_AXONEME_ASSEMBLY | 20 | -0.215633 | -0.50536 | 0.986437 | 1 |
| GOBP_GLYCOLYTIC_PROCESS_THROUGH_FRUCTOSE_6_PHOSPHATE | 28 | -0.202602 | -0.503344 | 0.992908 | 1 |
| GOBP_REGULATION_OF_MITOCHONDRIAL_MEMBRANE_PERMEABILITY | 70 | -0.179606 | -0.498706 | 0.997895 | 1 |
| GOBP_PREASSEMBLY_OF_GPI_ANCHOR_IN_ER_MEMBRANE | 16 | -0.220163 | -0.496358 | 0.980989 | 1 |
| GOBP_REGULATION_OF_CYTOKINESIS | 79 | -0.180321 | -0.495593 | 0.996852 | 1 |
| GOBP_PROTEIN_PALMITOYLATION | 33 | -0.195737 | -0.495563 | 0.993088 | 1 |
| GOBP_REPLICATION_FORK_PROCESSING | 39 | -0.192271 | -0.495233 | 0.994524 | 1 |
| GOBP_ESTABLISHMENT_OF_PROTEIN_LOCALIZATION_TO_POSTSYNAPTIC_ME | 15 | -0.222934 | -0.495134 | 0.979408 | 1 |
| GOBP_POSITIVE_REGULATION_OF_CYCLIN_DEPENDENT_PROTEIN_KINASE_A | 33 | -0.196183 | -0.494996 | 0.98603 | 1 |
| GOBP_POSITIVE_REGULATION_OF_CELL_CYCLE_PHASE_TRANSITION | 91 | -0.176528 | -0.494869 | 1 | 1 |
| GOBP_POSITIVE_REGULATION_OF_CELL_CYCLE_ARREST | 77 | -0.179122 | -0.494451 | 1 | 1 |
| GOBP_REGULATION_OF_GLUCOSE_IMPORT | 49 | -0.188886 | -0.493979 | 0.997836 | 1 |
| GOBP_POSITIVE_REGULATION_OF_HISTONE_H3_K4_METHYLATION | 16 | -0.222846 | -0.493487 | 0.979194 | 1 |
| GOBP_MACROAUTOPHAGY | 305 | -0.166766 | -0.492739 | 1 | 1 |
| GOBP_INTRACELLULAR_STEROL_TRANSPORT | 23 | -0.207906 | -0.492105 | 0.986585 | 1 |
| GOBP_REGULATION_OF_MITOCHONDRIAL_OUTER_MEMBRANE_PERMEABILIZA | 44 | -0.187349 | -0.491204 | 0.996622 | 1 |
| GOBP_METHYLATION | 337 | -0.165241 | -0.488964 | 1 | 1 |
| GOBP_BETA_CATENIN_TCF_COMPLEX_ASSEMBLY | 31 | -0.19711 | -0.486282 | 0.988345 | 1 |
| GOBP_INTRACELLULAR_STEROID_HORMONE_RECEPTOR_SIGNALING_PATHW | 105 | -0.17001 | -0.48592 | 0.998966 | 1 |
| GOBP_RESPONSE_TO_TOPOLOGICALLY_INCORRECT_PROTEIN | 198 | -0.166336 | -0.485292 | 1 | 1 |
| GOBP_REGULATION_OF_DNA_REPAIR | 128 | -0.169503 | -0.483851 | 1 | 1 |

| | | | | | |
|---|---|---|---|---|---|
| GOBP_REGULATION_OF_PROTEASOMAL_PROTEIN_CATABOLIC_PROCESS | 179 | -0.165439 | -0.482185 | 1 | 1 |
| GOBP_INTRACILIARY_TRANSPORT_INVOLVED_IN_CILIUM_ASSEMBLY | 40 | -0.185522 | -0.477657 | 0.99557 | 1 |
| GOBP_PROTEIN_ACYLATION | 232 | -0.161871 | -0.477045 | 1 | 1 |
| GOBP_ENDOPLASMIC_RETICULUM_UNFOLDED_PROTEIN_RESPONSE | 121 | -0.166297 | -0.476048 | 1 | 1 |
| GOBP_PROTEIN_MODIFICATION_BY_SMALL_PROTEIN_REMOVAL | 274 | -0.160903 | -0.476004 | 1 | 1 |
| GOBP_PEPTIDYL_LYSINE_METHYLATION | 117 | -0.167889 | -0.475306 | 1 | 1 |
| GOBP_CYTOKINESIS | 156 | -0.164109 | -0.47444 | 1 | 1 |
| GOBP_MITOTIC_G2_M_TRANSITION_CHECKPOINT | 36 | -0.186659 | -0.473963 | 0.998896 | 1 |
| GOBP_PEPTIDYL_LYSINE_MODIFICATION | 377 | -0.158961 | -0.473502 | 1 | 1 |
| GOBP_PROTEIN_O_LINKED_MANNOSYLATION | 18 | -0.210515 | -0.473431 | 0.988778 | 1 |
| GOBP_REGULATION_OF_SIGNAL_TRANSDUCTION_BY_P53_CLASS_MEDIATOR | 175 | -0.163654 | -0.4725 | 1 | 1 |
| GOBP_NEGATIVE_REGULATION_OF_PROTEASOMAL_PROTEIN_CATABOLIC_PR | 48 | -0.177703 | -0.47193 | 0.996739 | 1 |
| GOBP_SCF_DEPENDENT_PROTEASOMAL_UBIQUITIN_DEPENDENT_PROTEIN_( | 91 | -0.168985 | -0.470471 | 0.998968 | 1 |
| GOBP_REGULATION_OF_TRANSCRIPTION_INITIATION_FROM_RNA_POLYMERA | 32 | -0.185104 | -0.469786 | 0.994318 | 1 |
| GOBP_REGULATION_OF_MRNA_CATABOLIC_PROCESS | 198 | -0.15945 | -0.469428 | 1 | 1 |
| GOBP_NUCLEOBASE_CONTAINING_COMPOUND_TRANSPORT | 241 | -0.160063 | -0.467241 | 1 | 1 |
| GOBP_RETROGRADE_VESICLE_MEDIATED_TRANSPORT_GOLGI_TO_ENDOPLA | 84 | -0.168664 | -0.466831 | 0.998966 | 1 |
| GOBP_HYPEROSMOTIC_RESPONSE | 24 | -0.194943 | -0.466803 | 0.996394 | 1 |
| GOBP_REGULATION_OF_INTEGRIN_MEDIATED_SIGNALING_PATHWAY | 15 | -0.207565 | -0.466557 | 0.986962 | 1 |
| GOBP_BLASTOCYST_DEVELOPMENT | 88 | -0.169383 | -0.465301 | 1 | 1 |
| GOBP_POSITIVE_REGULATION_OF_PROTEASOMAL_UBIQUITIN_DEPENDENT_P | 87 | -0.165634 | -0.465242 | 1 | 1 |
| GOBP_POSITIVE_REGULATION_OF_RNA_SPLICING | 34 | -0.182791 | -0.463012 | 0.998856 | 1 |
| GOBP_NEGATIVE_REGULATION_OF_CELL_CYCLE_PHASE_TRANSITION | 241 | -0.157093 | -0.461863 | 1 | 1 |
| GOBP_MACROMOLECULE_METHYLATION | 285 | -0.156544 | -0.461583 | 1 | 1 |
| GOBP_POSITIVE_REGULATION_OF_TRANSCRIPTION_FROM_RNA_POLYMERAS | 23 | -0.19636 | -0.461209 | 0.991576 | 1 |
| GOBP_RESPONSE_TO_ENDOPLASMIC_RETICULUM_STRESS | 283 | -0.155888 | -0.461173 | 1 | 1 |
| GOBP_NEGATIVE_REGULATION_OF_CELL_CYCLE_G2_M_PHASE_TRANSITION | 106 | -0.162832 | -0.461054 | 1 | 1 |
| GOBP_POSITIVE_REGULATION_BY_HOST_OF_VIRAL_TRANSCRIPTION | 16 | -0.208025 | -0.46005 | 0.991117 | 1 |
| GOBP_POSITIVE_REGULATION_OF_TRANSCRIPTION_BY_RNA_POLYMERASE_I | 23 | -0.194758 | -0.460011 | 0.99511 | 1 |
| GOBP_REGULATION_OF_CELL_CYCLE_G2_M_PHASE_TRANSITION | 213 | -0.156712 | -0.458992 | 1 | 1 |
| GOBP_MALE_MEIOSIS_I | 19 | -0.197443 | -0.458461 | 0.991083 | 1 |
| GOBP_PROTEIN_LOCALIZATION_TO_MITOCHONDRION | 145 | -0.158166 | -0.458341 | 1 | 1 |
| GOBP_ATP_METABOLIC_PROCESS | 300 | -0.155374 | -0.458135 | 1 | 1 |
| GOBP_ORGANELLE_FISSION | 460 | -0.152752 | -0.451852 | 1 | 1 |
| GOBP_POSITIVE_REGULATION_OF_DNA_REPLICATION | 36 | -0.175541 | -0.448788 | 0.997725 | 1 |
| GOBP_POSITIVE_REGULATION_OF_SMAD_PROTEIN_SIGNAL_TRANSDUCTION | 15 | -0.203273 | -0.44798 | 0.991003 | 1 |
| GOBP_REGULATION_OF_CHROMATIN_ASSEMBLY | 18 | -0.197275 | -0.445757 | 0.997519 | 1 |
| GOBP_MITOCHONDRION_MORPHOGENESIS | 19 | -0.194219 | -0.445384 | 0.997497 | 1 |
| GOBP_ESTABLISHMENT_OF_PROTEIN_LOCALIZATION_TO_CHROMOSOME | 27 | -0.180621 | -0.443535 | 0.995423 | 1 |
| GOBP_REGULATION_OF_GENE_SILENCING | 125 | -0.155356 | -0.443303 | 1 | 1 |
| GOBP_PROTEIN_LOCALIZATION_TO_MICROTUBULE_ORGANIZING_CENTER | 35 | -0.175127 | -0.443212 | 0.998847 | 1 |
| GOBP_PROTEIN_DESTABILIZATION | 45 | -0.168103 | -0.442534 | 0.997807 | 1 |
| GOBP_POSITIVE_REGULATION_OF_DOUBLE_STRAND_BREAK_REPAIR | 40 | -0.170328 | -0.441715 | 0.996603 | 1 |
| GOBP_NEGATIVE_REGULATION_OF_TRANSLATIONAL_INITIATION | 19 | -0.190989 | -0.44155 | 0.997459 | 1 |
| GOBP_PROTEIN_METHYLATION | 162 | -0.153317 | -0.44082 | 1 | 1 |
| GOBP_TRNA_METHYLATION | 40 | -0.16976 | -0.437707 | 0.998874 | 1 |
| GOBP_REGULATION_OF_RESPONSE_TO_ENDOPLASMIC_RETICULUM_STRESS | 77 | -0.15793 | -0.437219 | 1 | 1 |
| GOBP_POSITIVE_REGULATION_OF_TOR_SIGNALING | 40 | -0.169247 | -0.435281 | 1 | 1 |
| GOBP_COVALENT_CHROMATIN_MODIFICATION | 445 | -0.146577 | -0.434862 | 1 | 1 |
| GOBP_NEGATIVE_REGULATION_OF_DOUBLE_STRAND_BREAK_REPAIR_VIA_H | 19 | -0.184454 | -0.431475 | 0.996193 | 1 |
| GOBP_TELOMERE_CAPPING | 50 | -0.160149 | -0.425824 | 0.998899 | 1 |
| GOBP_SPLICEOSOMAL_COMPLEX_ASSEMBLY | 64 | -0.156508 | -0.425812 | 0.998951 | 1 |
| GOBP_HEME_METABOLIC_PROCESS | 31 | -0.169573 | -0.425313 | 0.997647 | 1 |
| GOBP_MODULATION_BY_SYMBIONT_OF_HOST_PROCESS | 32 | -0.167903 | -0.422455 | 0.997685 | 1 |
| GOBP_NUCLEOSIDE_TRIPHOSPHATE_METABOLIC_PROCESS | 107 | -0.148016 | -0.422036 | 1 | 1 |
| GOBP_REGULATION_OF_CELLULAR_RESPONSE_TO_HEAT | 79 | -0.15126 | -0.420696 | 1 | 1 |
| GOBP_NEGATIVE_REGULATION_OF_MRNA_CATABOLIC_PROCESS | 50 | -0.158389 | -0.419679 | 0.998886 | 1 |
| GOBP_SIGNAL_TRANSDUCTION_BY_P53_CLASS_MEDIATOR | 258 | -0.142398 | -0.41953 | 1 | 1 |
| GOBP_NEGATIVE_REGULATION_OF_CELL_MIGRATION_INVOLVED_IN_SPROUT | 16 | -0.186688 | -0.416817 | 0.997465 | 1 |
| GOBP_PROTEIN_K11_LINKED_UBIQUITINATION | 29 | -0.168969 | -0.415615 | 0.998824 | 1 |
| GOBP_CELLULAR_RESPONSE_TO_TOPOLOGICALLY_INCORRECT_PROTEIN | 161 | -0.142732 | -0.412507 | 1 | 1 |
| GOBP_REGULATION_OF_PROTEASOMAL_UBIQUITIN_DEPENDENT_PROTEIN_C | 130 | -0.144257 | -0.411278 | 1 | 1 |
| GOBP_NEGATIVE_REGULATION_OF_ORGANELLE_ASSEMBLY | 40 | -0.158298 | -0.408929 | 1 | 1 |
| GOBP_CHAPERONE_MEDIATED_PROTEIN_FOLDING | 61 | -0.149826 | -0.406994 | 1 | 1 |
| GOBP_HEPARAN_SULFATE_PROTEOGLYCAN_BIOSYNTHETIC_PROCESS | 24 | -0.167873 | -0.405757 | 1 | 1 |
| GOBP_GENE_SILENCING | 290 | -0.137179 | -0.405174 | 1 | 1 |
| GOBP_SELECTIVE_AUTOPHAGY | 62 | -0.147284 | -0.400977 | 1 | 1 |
| GOBP_REGULATION_OF_DNA_TEMPLATED_TRANSCRIPTION_INITIATION | 39 | -0.153718 | -0.39893 | 1 | 1 |
| GOBP_RETROGRADE_TRANSPORT_ENDOSOME_TO_GOLGI | 90 | -0.140923 | -0.398325 | 1 | 1 |
| GOBP_NUCLEUS_ORGANIZATION | 117 | -0.139441 | -0.398006 | 1 | 1 |
| GOBP_G2_DNA_DAMAGE_CHECKPOINT | 34 | -0.151907 | -0.397632 | 1 | 1 |
| GOBP_TRANSCRIPTION_PREINITIATION_COMPLEX_ASSEMBLY | 35 | -0.156157 | -0.394759 | 1 | 1 |
| GOBP_PYRIMIDINE_NUCLEOSIDE_TRIPHOSPHATE_BIOSYNTHETIC_PROCESS | 19 | -0.170601 | -0.393611 | 1 | 1 |
| GOBP_MITOTIC_G1_S_TRANSITION_CHECKPOINT | 62 | -0.140503 | -0.385528 | 1 | 1 |
| GOBP_MRNA_TRANSPORT | 146 | -0.131388 | -0.379857 | 1 | 1 |
| GOBP_REGULATION_OF_TRANSCRIPTION_BY_RNA_POLYMERASE_I | 33 | -0.150944 | -0.376679 | 1 | 1 |
| GOBP_MITOTIC_DNA_INTEGRITY_CHECKPOINT | 104 | -0.133045 | -0.374082 | 1 | 1 |
| GOBP_IRE1_MEDIATED_UNFOLDED_PROTEIN_RESPONSE | 64 | -0.135753 | -0.372138 | 1 | 1 |
| GOBP_RNA_PHOSPHODIESTER_BOND_HYDROLYSIS_EXONUCLEOLYTIC | 43 | -0.141757 | -0.37056 | 1 | 1 |
| GOBP_MITOTIC_NUCLEAR_DIVISION | 284 | -0.12377 | -0.365085 | 1 | 1 |
| GOBP_NEGATIVE_REGULATION_OF_AUTOPHAGOSOME_ASSEMBLY | 15 | -0.167409 | -0.363943 | 1 | 1 |
| GOBP_INTRACILIARY_TRANSPORT | 53 | -0.134505 | -0.362766 | 1 | 1 |
| GOBP_REGULATION_OF_CELL_CYCLE_CHECKPOINT | 31 | -0.146638 | -0.361885 | 1 | 0.99982 |
| GOBP_REGULATION_OF_DNA_TEMPLATED_TRANSCRIPTION_IN_RESPONSE_T | 113 | -0.125091 | -0.358651 | 1 | 0.99955 |
| GOBP_REGULATION_OF_MAMMARY_GLAND_EPITHELIAL_CELL_PROLIFERATIC | 16 | -0.15444 | -0.353033 | 1 | 0.99929 |
| GOBP_ERAD_PATHWAY | 100 | -0.124948 | -0.35213 | 1 | 0.999002 |
| GOBP_PYRIMIDINE_NUCLEOSIDE_TRIPHOSPHATE_METABOLIC_PROCESS | 24 | -0.145489 | -0.348835 | 1 | 0.998725 |
| GOBP_REGULATION_OF_TRANSLATIONAL_ELONGATION | 19 | -0.145354 | -0.344807 | 0.998753 | 0.998452 |
| GOBP_PROTEIN_CONTAINING_COMPLEX_DISASSEMBLY | 319 | -0.117122 | -0.344254 | 1 | 0.998163 |
| GOBP_MITOTIC_DNA_REPLICATION | 16 | -0.152694 | -0.342392 | 1 | 0.997877 |
| GOBP_ENDOSOME_TO_LYSOSOME_TRANSPORT | 51 | -0.128253 | -0.341229 | 1 | 0.99759 |
| GOBP_NEGATIVE_REGULATION_OF_RNA_CATABOLIC_PROCESS | 61 | -0.123363 | -0.336754 | 1 | 0.997311 |
| GOBP_DNA_DEPENDENT_DNA_REPLICATION | 153 | -0.117205 | -0.333634 | 1 | 0.99703 |
| GOBP_DNA_REPLICATION | 270 | -0.109406 | -0.322054 | 1 | 0.996762 |
| GOBP_CYTOSKELETON_DEPENDENT_CYTOKINESIS | 94 | -0.111817 | -0.315843 | 1 | 0.996478 |
| GOBP_ESTABLISHMENT_OF_RNA_LOCALIZATION | 193 | -0.10705 | -0.311308 | 1 | 0.996193 |
| GOBP_MRNA_EXPORT_FROM_NUCLEUS | 109 | -0.107861 | -0.306178 | 1 | 0.995907 |
| GOBP_TRANSCRIPTION_BY_RNA_POLYMERASE_III | 47 | -0.111983 | -0.299609 | 1 | 0.995622 |
