## Supplementary material for "Chemokine Receptor 1 and its associated immune pathway are downregulated in SF3B1^MT^ blood and non-blood cancers": Table_S13

**Table S13. Shared GSEA negative hematopoietic/immune genes sets in SF3B1 blood cancers MDS, CLL and AML.** Gene sets with p-values < 0.05 and NES (normalized enrichment score) are shown. Gene sets with related functions are shown in different font colors and chemokine-related pathways highlighted in green.

| Gene sets | NES | | |
|---|---|---|---|
| | MDS | AML | CLL |
| POSITIVE REGULATION OF LEUKOCYTE CELL CELL ADHESION | -1.8699853 | -1.401652 | -1.84993 |
| REGULATION OF T CELL ACTIVATION | -1.834335 | -1.2472168 | -1.862288 |
| T CELL ACTIVATION | -1.8081324 | -1.2118152 | -2.011879 |
| LEUKOCYTE CELL CELL ADHESION | -1.7687527 | -1.3579484 | -1.828649 |
| REGULATION OF LYMPHOCYTE ACTIVATION | -1.7484266 | -1.4356874 | -1.625954 |
| LEUKOCYTE PROLIFERATION | -1.7328736 | -1.3966384 | -1.700117 |
| REGULATION OF LEUKOCYTE PROLIFERATION | -1.7164836 | -1.3870379 | -1.697121 |
| POSITIVE REGULATION OF LEUKOCYTE PROLIFERATION | -1.6618149 | -1.4322945 | -1.818309 |
| REGULATION OF IMMUNE EFFECTOR PROCESS | -1.6389902 | -1.3613187 | -1.444493 |
| T CELL PROLIFERATION | -1.6267977 | -1.3067741 | -1.779037 |
| RESPONSE TO CHEMOKINE | -1.6243176 | -2.0462375 | -1.887222 |
| LEUKOCYTE DIFFERENTIATION | -1.6140529 | -1.2849168 | -1.928443 |
| CHEMOKINE PRODUCTION | -1.6083384 | -1.3743702 | -1.476193 |
| LYMPHOCYTE MEDIATED IMMUNITY | -1.6004554 | -1.387201 | -1.457993 |
| REGULATION OF INFLAMMATORY RESPONSE | -1.5625333 | -1.4693024 | -1.727632 |
| POSITIVE REGULATION OF T CELL PROLIFERATION | -1.555971 | -1.3299707 | -1.843569 |
| ADAPTIVE IMMUNE RESPONSE BASED ON SOMATIC RECOMBINATION OF IMMUNE RE( | -1.5094808 | -1.5656606 | -1.38705 |
| INTERLEUKIN 8 PRODUCTION | -1.506621 | -1.3768785 | -1.499182 |
| POSITIVE REGULATION OF INFLAMMATORY RESPONSE | -1.4799402 | -1.7010176 | -1.51438 |
| LEUKOCYTE CHEMOTAXIS | -1.4716723 | -1.8252776 | -1.784172 |
| DEFENSE RESPONSE TO BACTERIUM | -1.4598963 | -2.054695 | -1.485058 |
| MYELOID LEUKOCYTE MIGRATION | -1.4573506 | -2.0261593 | -1.882169 |
| REGULATION OF LEUKOCYTE CHEMOTAXIS | -1.4377856 | -1.3454143 | -1.58316 |
| REGULATION OF LEUKOCYTE MIGRATION | -1.4255416 | -1.3667679 | -1.75948 |
| MONONUCLEAR CELL MIGRATION | -1.4067936 | -1.5750545 | -1.855846 |
| HUMORAL IMMUNE RESPONSE | -1.3939049 | -2.0626159 | -1.506165 |
| LEUKOCYTE MIGRATION | -1.371388 | -1.707669 | -1.589392 |
| MYELOID LEUKOCYTE DIFFERENTIATION | -1.3703489 | -1.5849216 | -1.571566 |
| RESPONSE TO INTERLEUKIN 1 | -1.3359929 | -1.6015897 | -1.690619 |
| GRANULOCYTE MIGRATION | -1.3231283 | -1.9909668 | -1.919634 |

**Table S13. Shared GSEA negative hematopoietic/immune genes sets in non-blood SF3B1 cancers, BRCA and UVM.** Gene sets with p-values < 0.05 and NES (normalized enrichment score) are shown. Gene sets with related functions are shown in different font colors and chemokine-related pathways highlighted in green.

| Gene sets | NES | |
|---|---|---|
| | BRCA | UVM |
| IMMUNOGLOBULIN PRODUCTION | -1.998158 | -2.2958038 |
| HUMORAL IMMUNE RESPONSE MEDIATED BY CIRCULATING IMMUNOGLOBULIN | -1.952248 | -2.4909513 |
| COMPLEMENT ACTIVATION | -1.889209 | -2.450629 |
| REGULATION OF HUMORAL IMMUNE RESPONSE | -1.865636 | -2.3983142 |
| HUMORAL IMMUNE RESPONSE | -1.799556 | -2.2466104 |
| PRODUCTION OF MOLECULAR MEDIATOR OF IMMUNE RESPONSE | -1.770304 | -2.2353764 |
| B CELL RECEPTOR SIGNALING PATHWAY | -1.764235 | -2.4014268 |
| REGULATION OF COMPLEMENT ACTIVATION | -1.743289 | -2.3829439 |
| B CELL MEDIATED IMMUNITY | -1.670891 | -2.4281297 |
| ANTIBACTERIAL HUMORAL RESPONSE | -1.662323 | -1.7536978 |
| POSITIVE REGULATION OF B CELL ACTIVATION | -1.657498 | -2.3953779 |
| REGULATION OF B CELL ACTIVATION | -1.600826 | -2.3016253 |
| ANTIMICROBIAL HUMORAL RESPONSE | -1.589319 | -1.6580435 |
| REGULATION OF B CELL PROLIFERATION | -1.533498 | -1.805866 |
| LYMPHOCYTE MEDIATED IMMUNITY | -1.53163 | -2.2677746 |
| B CELL ACTIVATION | -1.511996 | -2.138151 |
| B CELL PROLIFERATION | -1.495192 | -1.860152 |
| CHRONIC INFLAMMATORY RESPONSE | -1.478995 | -1.7174199 |
| ADAPTIVE IMMUNE RESPONSE BASED ON SOMATIC RECOMBINATION OF IMMUNE RECEPT | -1.477493 | -2.2353764 |
| POSITIVE REGULATION OF MONOCYTE CHEMOTAXIS | -1.454185 | -1.5553976 |
| REGULATION OF LYMPHOCYTE ACTIVATION | -1.266874 | -2.0615451 |
| REGULATION OF IMMUNE EFFECTOR PROCESS | -1.235125 | -2.0941484 |
| LEUKOCYTE MIGRATION | -1.232723 | -1.9847738 |
| PHAGOCYTOSIS RECOGNITION | -1.807779 | -2.4199803 |
| DEFENSE RESPONSE TO BACTERIUM | -1.637951 | -2.123265 |
| FC EPSILON RECEPTOR SIGNALING PATHWAY | -1.60163 | -2.2782469 |
| POSITIVE REGULATION OF HUMORAL IMMUNE RESPONSE | -1.575089 | -1.868866 |
| IMMUNE RESPONSE REGULATING SIGNALING PATHWAY | -1.548723 | -2.1136138 |
| POSITIVE REGULATION OF ANTIGEN RECEPTOR MEDIATED SIGNALING PATHWAY | -1.545938 | -1.5873777 |
| ANTIGEN RECEPTOR MEDIATED SIGNALING PATHWAY | -1.53574 | -2.1857321 |
| FC RECEPTOR SIGNALING PATHWAY | -1.460147 | -2.1648083 |
| REGULATION OF ANTIGEN RECEPTOR MEDIATED SIGNALING PATHWAY | -1.435945 | -1.7730998 |

| **Table S13. Shared GSEA negative hematopoietic/immune genes sets in SF3B1 cancers, MDS, AML, CLL and UVM.** Gene sets with p-valus < 0.05 are shown and Chemokine-related pathways are highlighted in green. |
|---|
| **Gene sets** |
| POSITIVE REGULATION OF LEUKOCYTE CELL CELL ADHESION |
| REGULATION OF T CELL ACTIVATION |
| T CELL ACTIVATION |
| LEUKOCYTE CELL CELL ADHESION |
| REGULATION OF LYMPHOCYTE ACTIVATION |
| LEUKOCYTE PROLIFERATION |
| REGULATION OF LEUKOCYTE PROLIFERATION |
| POSITIVE REGULATION OF LEUKOCYTE PROLIFERATION |
| REGULATION OF IMMUNE EFFECTOR PROCESS |
| T CELL PROLIFERATION |
| RESPONSE TO CHEMOKINE |
| LEUKOCYTE DIFFERENTIATION |
| CHEMOKINE PRODUCTION |
| LYMPHOCYTE MEDIATED IMMUNITY |
| REGULATION OF INFLAMMATORY RESPONSE |
| POSITIVE REGULATION OF T CELL PROLIFERATION |
| ADAPTIVE IMMUNE RESPONSE BASED ON SOMATIC RECOMBINATION OF IMMUNE RECEPTORS BUILT FROM IMMUNOGLOBULIN SUPERFAMILY DOMAINS |
| INTERLEUKIN 8 PRODUCTION |
| POSITIVE REGULATION OF INFLAMMATORY RESPONSE |
| LEUKOCYTE CHEMOTAXIS |
| DEFENSE RESPONSE TO BACTERIUM |
| MYELOID LEUKOCYTE MIGRATION |
| REGULATION OF LEUKOCYTE CHEMOTAXIS |
| REGULATION OF LEUKOCYTE MIGRATION |
| HUMORAL IMMUNE RESPONSE |
| LEUKOCYTE MIGRATION |
| MYELOID LEUKOCYTE DIFFERENTIATION |
| RESPONSE TO INTERLEUKIN 1 |
| GRANULOCYTE MIGRATION |
