## Supplementary material for "Chemokine Receptor 1 and its associated immune pathway are downregulated in SF3B1^MT^ blood and non-blood cancers": Table_S14

**Table S14. Significantly downregulated genes shared between SF3B1$^{MT}$ MDS and genes in the LEUKOCYTE MIGRATION gene set.** FC is fold change.

| Gene | FC |
|---|---|
| CCL7 | -11.7 |
| IL1B | -6 |
| CXCL3 | -5.8 |
| CH25H | -5.1 |
| CXCL5 | -5 |
| CCL2 | -3.5 |
| CXCL8 | -3.5 |
| EDN2 | -3.5 |
| CCL3L1 | -3.4 |
| CD34 | -3.1 |
| TREM2 | -3.1 |
| CD200 | -3 |
| CCL3 | -2.7 |
| WNT5A | -2.7 |
| CHGA | -2.5 |
| SCG2 | -2.5 |
| THBS1 | -2.5 |
| ECM1 | -2.4 |
| GPR183 | -2.4 |
| IL10 | -2.4 |
| PTGER4 | -2.4 |
| CXCL10 | -2.3 |
| GCSAML | -2.3 |
| NINJ1 | -2.3 |
| CCL20 | -2.2 |
| CCL23 | -2.2 |
| ELANE | -2.2 |
| DEFB124 | -2.1 |
| PTPRO | -2.1 |
| ARTN | -2 |
| CYP7B1 | -2 |
| F2RL1 | -2 |
| PLCB1 | -2 |
| S1PR1 | -2 |
| STAP1 | -2 |
| TNF | -2 |
| CCR1 | -1.9 |
| CD74 | -1.9 |
| CNR2 | -1.9 |
| IL27RA | -1.9 |
| MCOLN2 | -1.9 |

| | |
|---|---|
| TREM1 | -1.9 |
| C3AR1 | -1.8 |
| CCL4 | -1.8 |
| CSF1R | -1.8 |
| DUSP1 | -1.8 |
| ITGA3 | -1.8 |
| ITGA7 | -1.8 |
| ITGB7 | -1.8 |
| KIT | -1.8 |
| PDE4B | -1.8 |
| PDGFD | -1.8 |
| PLEC | -1.8 |
| SPNS2 | -1.8 |
| SWAP70 | -1.8 |
| TNFRSF11A | -1.8 |
| ADGRE2 | -1.7 |
| C5 | -1.7 |
| CXCR3 | -1.7 |
| FYN | -1.7 |
| P2RX4 | -1.7 |
| ST3GAL4 | -1.7 |
| ASB2 | -1.6 |
| CCL5 | -1.6 |
| IL12A | -1.6 |
| LRCH1 | -1.6 |
| NBL1 | -1.6 |
| RHOH | -1.6 |
| SLAMF1 | -1.6 |
| SRC | -1.6 |
| ADA | -1.5 |
| AIF1 | -1.5 |
| ANO6 | -1.5 |
| ANXA1 | -1.5 |
| DOCK8 | -1.5 |
| GPR18 | -1.5 |
| NOD2 | -1.5 |
| PECAM1 | -1.5 |
| SPI1 | -1.5 |
| CCL28 | -1.4 |
| IL23A | -1.4 |
| LYN | -1.4 |

**Table S14. Significantly downregulated genes shared between $SF3B1^{MT}$ AML and genes in the LEUKOCYTE MIGRATION gene set.** FC is fold change.

| Gene | FC |
|---|---|
| SCG2 | -83.1 |
| CCL20 | -20.7 |
| PGF | -18.4 |
| TNFAIP6 | -18.1 |
| CCL3L1 | -12.1 |
| SERPINE1 | -11.5 |
| CYP19A1 | -8.3 |
| EDN1 | -7.8 |
| ITGA2 | -7.5 |
| CCL23 | -7.3 |
| TNFSF14 | -7.2 |
| CXCL11 | -6.9 |
| CXCR1 | -6.9 |
| CCL3 | -6.8 |
| IL6 | -6.7 |
| CXCL10 | -6.4 |
| CXCL1 | -5.7 |
| FPR2 | -5.7 |
| PRTN3 | -5.6 |
| CXCR2 | -5.5 |
| IL1B | -5.5 |
| ARTN | -5.4 |
| ITGA7 | -5.4 |
| CXADR | -5.3 |
| ELANE | -4.9 |
| CCL22 | -4.8 |
| CXCL8 | -4.3 |
| C3AR1 | -4.2 |
| PADI2 | -3.7 |
| PTAFR | -3.4 |
| CCR1 | -3.3 |
| RET | -3.3 |
| TREM1 | -3.3 |
| TNFSF11 | -3.2 |
| CSF1 | -3 |
| CCL4 | -2.9 |
| CXCR4 | -2.9 |
| IL1R1 | -2.7 |
| NINJ1 | -2.7 |
| TRPM2 | -2.7 |
| CHST2 | -2.5 |

| | |
|---|---|
| DUSP1 | -2.5 |
| ADAM8 | -2.2 |
| ANXA1 | -2.1 |
| FLT1 | -2.1 |
| SIRPA | -2.1 |
| SRP54 | -2 |
| BSG | -1.8 |
| CALR | -1.8 |
| FADD | -1.6 |
| AKIRIN1 | -1.5 |
| MYH9 | -1.5 |

**Table S14. Significantly downregulated genes shared between SF3B1$^{MT}$ CLL and genes in the LEUKOCYTE MIGRATION gene set.** FC is fold change.

| Gene | FC |
|---|---|
| CCL24 | -6884402.5 |
| MMP9 | -538.5 |
| CCR1 | -256.2 |
| CCL2 | -245.7 |
| S100A9 | -179.6 |
| PLA2G7 | -150.7 |
| SLAMF8 | -117.4 |
| TREM2 | -105.7 |
| C3AR1 | -58.6 |
| S100A8 | -57.7 |
| IL1A | -51.8 |
| CCR5 | -50 |
| CMKLR1 | -45.2 |
| TREM1 | -34.3 |
| CSF1R | -20.3 |
| GCSAM | -19.3 |
| GPR15 | -19.3 |
| CD300A | -18.2 |
| HCK | -17.2 |
| THBS1 | -14.1 |
| IL6R | -12.9 |
| CCL5 | -12.8 |
| ANXA1 | -12.2 |
| IL1R1 | -8.7 |
| PTK2 | -8.2 |
| CRTAM | -7.8 |
| IL1B | -7.5 |
| CX3CR1 | -6.2 |
| LGMN | -4.9 |
| ITGA1 | -4.5 |
| PODXL2 | -4.2 |
| TNFRSF18 | -3.8 |
| ITGB2 | -3.4 |
| GPR183 | -3.2 |
| PGF | -3.2 |
| EPS8 | -3.1 |
| PTPRO | -3.1 |
| HSD3B7 | -3 |
| SRC | -2.1 |
| IL23A | -2 |
| CCR7 | -1.7 |

| | |
|---|---|
| PTGER4 | -1.7 |
| PIKFYVE | -1.5 |

**Table S14. Significantly downregulated genes shared between SF3B1$^{MT}$ BRCA and genes in the LEUKOCYTE MIGRATION gene set.** FC is fold change.

| Gene | FC |
|---|---|
| CHST4 | -6.9 |
| CXCL5 | -4 |
| S100A9 | -3.8 |
| CXCL13 | -3.7 |
| TNFRSF11A | -3.4 |
| CCL18 | -2.9 |
| GPR15 | -2.7 |
| CXCL17 | -2.6 |
| S100A8 | -2.5 |
| SLAMF1 | -2.4 |
| LBP | -2.2 |
| CXCR5 | -2.1 |
| EXT1 | -2 |
| VEGFD | -2 |
| CCR6 | -1.9 |
| CNR2 | -1.9 |
| CX3CL1 | -1.9 |
| SELL | -1.9 |
| CRTAM | -1.8 |
| CYP7B1 | -1.8 |
| GPR18 | -1.8 |
| GPR183 | -1.7 |
| JAML | -1.7 |
| C5 | -1.6 |
| CCR1 | -1.6 |

**Table S14. Significantly downregulated genes shared between SF3B1$^{MT}$ UVM and genes in the LEUKOCYTE MIGRATION gene set.** FC is fold change.

| Gene | FC |
|---|---|
| CXCL9 | -15.7 |
| CXCR3 | -10.8 |
| CXCL13 | -10.7 |
| CRTAM | -10.2 |
| CCL24 | -10.1 |
| CCL22 | -9.9 |
| LCK | -9.3 |
| CXCL10 | -8.8 |
| XCL1 | -8.4 |
| GPR15 | -8.3 |
| RHOH | -8.3 |
| KLRK1 | -7.9 |
| ZAP70 | -7.3 |
| ITGA4 | -6.6 |
| CCL5 | -6.5 |
| CCR5 | -6.2 |
| CCR7 | -5.9 |
| SPN | -5.9 |
| ITGAL | -5.8 |
| SERPINE1 | -5.8 |
| SLAMF1 | -5.8 |
| XCL2 | -5.8 |
| CXCL6 | -5.7 |
| GPR18 | -5.7 |
| FUT7 | -5.6 |
| ECM1 | -5.5 |
| CCL4 | -5.4 |
| CD200R1 | -5.1 |
| TNFSF14 | -4.9 |
| CXCL11 | -4.8 |
| SLAMF8 | -4.8 |
| CCL13 | -4.6 |
| GCSAM | -4.5 |
| CD74 | -4.4 |
| MMP9 | -4.4 |
| CCL26 | -4.3 |
| PTPRO | -4.2 |
| TBX21 | -4.2 |
| VCAM1 | -4.2 |
| ADTRP | -4 |
| MYO1G | -3.9 |

| | |
|---|---|
| CCR2 | -3.7 |
| FPR2 | -3.6 |
| IL10 | -3.6 |
| NOD2 | -3.6 |
| CCL25 | -3.5 |
| CORO1A | -3.5 |
| NCKAP1L | -3.5 |
| PIK3CG | -3.5 |
| ROR2 | -3.5 |
| SELL | -3.5 |
| TNF | -3.5 |
| CNR2 | -3.4 |
| ELANE | -3.4 |
| CD300A | -3.3 |
| PLA2G7 | -3.3 |
| TGFB2 | -3.3 |
| EDN3 | -3.2 |
| IL1B | -3.2 |
| JAML | -3.2 |
| PTN | -3.2 |
| SCG2 | -3.2 |
| TRIM55 | -3.2 |
| VAV1 | -3.2 |
| AIF1 | -3.1 |
| CCL20 | -3.1 |
| CCL23 | -3.1 |
| CMKLR1 | -3.1 |
| IL1R1 | -3.1 |
| RIPK3 | -3.1 |
| WNT5A | -3.1 |
| CCN3 | -3 |
| NBL1 | -3 |
| BMP5 | -2.8 |
| CCR1 | -2.8 |
| GPR183 | -2.8 |
| SELE | -2.8 |
| ALOX5 | -2.7 |
| CHST2 | -2.7 |
| CXCL2 | -2.7 |
| PTAFR | -2.7 |
| SELPLG | -2.7 |
| ADORA1 | -2.6 |
| ANXA1 | -2.6 |
| CCL3 | -2.6 |
| CXCL12 | -2.6 |

| | |
|---|---|
| RAC2 | -2.6 |
| CAMK1D | -2.5 |
| CSF1R | -2.5 |
| PTGER4 | -2.5 |
| C3AR1 | -2.4 |
| CCL18 | -2.4 |
| CSF3R | -2.4 |
| FCER1G | -2.4 |
| LYN | -2.4 |
| MDK | -2.4 |
| BDKRB1 | -2.3 |
| CCL14 | -2.3 |
| FOLR2 | -2.3 |
| GAS6 | -2.3 |
| IL6 | -2.3 |
| KIT | -2.3 |
| ITGB2 | -2.2 |
| LGALS9 | -2.2 |
| MSTN | -2.2 |
| SPNS2 | -2.2 |
| THY1 | -2.2 |
| TNFRSF18 | -2.2 |
| FOXJ1 | -2.1 |
| PLCB1 | -2.1 |
| S100A8 | -2.1 |
| S1PR1 | -2.1 |
| TRPM2 | -2.1 |
| AZU1 | -2 |
| CXCL8 | -2 |
| MCOLN2 | -2 |
| PECAM1 | -2 |
| SLIT2 | -2 |
| SPI1 | -2 |
| TACR1 | -2 |
| TNFSF18 | -2 |
| C5AR2 | -1.9 |
| CCL3L1 | -1.9 |
| CXCL16 | -1.9 |
| CXCR4 | -1.9 |
| HMOX1 | -1.9 |
| HRH1 | -1.9 |
| ITGA1 | -1.9 |
| S100A9 | -1.9 |
| STK39 | -1.9 |
| ADAM8 | -1.8 |

| | |
|---|---|
| ASB2 | -1.8 |
| CCL2 | -1.8 |
| CH25H | -1.8 |
| FFAR2 | -1.8 |
| IL16 | -1.8 |
| ITGA2 | -1.8 |
| SYK | -1.8 |
| DOCK8 | -1.7 |
| HCK | -1.7 |
| SMPD3 | -1.7 |
| TNFRSF11A | -1.7 |
| AKIRIN1 | -1.6 |
| C5AR1 | -1.6 |
| CSF1 | -1.6 |
| CX3CR1 | -1.6 |
| DAPK2 | -1.6 |
| FLT1 | -1.6 |
| ITGA6 | -1.6 |
| JAM2 | -1.6 |
| PREX1 | -1.6 |
| SLC12A2 | -1.6 |
| ADAM10 | -1.5 |
| B4GALT1 | -1.5 |
| IL27RA | -1.5 |
| IL34 | -1.5 |
| ITGA7 | -1.5 |
| LGMN | -1.5 |
| LYST | -1.5 |
| CD34 | -1.4 |
| SOS1 | -1.4 |
| TNFRSF14 | -1.4 |
| TRPM4 | -1.4 |

**Table S14. Significantly downregulated genes shared between SRSF2$^{MT}$ MDS and genes in the LEUKOCYTE MIGRATION gene set.** FC is fold change.

| Gene | FC |
|---|---|
| LEP | -10.5 |
| OLFM4 | -8 |
| CCR2 | -6.3 |
| TACR1 | -4.6 |
| CSF1R | -3.1 |
| SLAMF8 | -2.9 |
| MOSPD2 | -2.6 |
| ELANE | -2.5 |
| P2RY12 | -2.5 |
| PLA2G7 | -2.4 |
| SPN | -2.4 |
| PRTN3 | -2.3 |
| NOD2 | -2 |
| CALR | -1.9 |
| CD74 | -1.8 |
| PECAM1 | -1.8 |

**Table S14. Significantly downregulated genes shared between $U2AF1^{MT}$ MDS and genes in the LEUKOCYTE MIGRATION gene set.** FC is fold change.

| Gene | FC |
|---|---|
| CCL7 | -17.9 |
| LEP | -11.4 |
| OLFM4 | -6.5 |
| CCR2 | -3.9 |
| CXCL2 | -3.2 |
| DEFB124 | -3.1 |
| MMP9 | -2.6 |
| ASB2 | -2.5 |
| IL1B | -2.5 |
| CXCL3 | -2.4 |
| ELANE | -2.3 |
| GPR183 | -2.3 |
| CCR1 | -2.2 |
| CTSG | -2.1 |
| CCN3 | -2 |
| IL16 | -2 |
| ITGA7 | -2 |
| PRTN3 | -2 |
| SIRPA | -2 |
| ANXA1 | -1.9 |
| AZU1 | -1.9 |
| C3AR1 | -1.9 |
| GCNT1 | -1.9 |
| PYCARD | -1.9 |
| IL17RA | -1.8 |
| ITGB2 | -1.8 |
| NOD2 | -1.8 |
| BST1 | -1.7 |
| ITGA9 | -1.7 |
| PECAM1 | -1.7 |
| PTAFR | -1.7 |
| RHOG | -1.6 |
| SWAP70 | -1.6 |
| ADA | -1.5 |
| MSN | -1.5 |
| WDR1 | -1.5 |

**Table S14. Significantly downregulated genes shared between ZRSR2$^{MT}$ MDS and genes in the LEUKOCYTE MIGRATION gene set.** FC is fold change.

| Gene | FC |
|---|---|
| LEP | -59 |
| IL6 | -6.3 |
| CCR2 | -5.7 |
| CCL22 | -5.5 |
| CXCL3 | -4.9 |
| CCL24 | -4.8 |
| SELE | -4.5 |
| EDNRB | -4.3 |
| CXCL2 | -4 |
| TRPV4 | -3.4 |
| CXCL12 | -3.3 |
| CD200R1 | -3 |
| FOLR2 | -3 |
| HMOX1 | -3 |
| IL10 | -3 |
| MMP14 | -2.9 |
| LGMN | -2.6 |
| CCR5 | -2.2 |
| GCNT1 | -2.2 |
| TNFRSF11A | -2.2 |
| ITGA7 | -2.1 |
| CXADR | -2 |
| CCL23 | -1.9 |
| CHST2 | -1.9 |
| NBL1 | -1.9 |
| GOLPH3 | -1.7 |
| SIRPA | -1.7 |
| LGALS3 | -1.6 |
| LRCH1 | -1.5 |
