## Supplementary material for "Chemokine Receptor 1 and its associated immune pathway are downregulated in SF3B1^MT^ blood and non-blood cancers": Table_S15

**Table S15. Downregulated Chemokine Receptors (CCRs) and their ligands (CCLs) in splicing factor cancers.** CCR1 and its CCLs are highlighted in yellow. Fold changes (FCs) are shown.

| CCRs | SF3B1 | SF3B1 | SF3B1 | SF3B1 | SF3B1 | SRSF2 | U2AF1 | ZRSR2 |
|---|---|---|---|---|---|---|---|---|
| | MDS | AML | CLL | BRCA | UVM | MDS | MDS | MDS |
| CCR1 | -1.9 | -3.3 | -256.2 | -1.6 | -2.8 | | -2.2 | |
| CCR2 | | | | | -3.7 | -6.3 | -3.9 | -5.7 |
| CCR4 | | | -15.8 | -1.8 | -6.2 | | | |
| CCR5 | | | -50 | | -6.2 | | | -2.2 |
| CCR6 | | | | -1.9 | | | | |
| CCR7 | | | -1.7 | | -5.9 | | | |
| CCR8 | | | | | | | | |
| CCR9 | -1.9 | | | | | | | |
| CCRL2 | -4.6 | | | | -2.5 | | | |

| Gene | SF3B1 | SF3B1 | SF3B1 | SF3B1 | SF3B1 | SRSF2 | U2AF1 | ZRSR2 |
|---|---|---|---|---|---|---|---|---|
| | MDS | AML | CLL | BRCA | UVM | MDS | MDS | MDS |
| CCL2 | -3.5 | | -245.7 | | -1.8 | | | |
| CCL3 | -2.7 | -6.8 | | | -2.6 | | | |
| CCL3L1 | -3.4 | -12.1 | | | -1.9 | | | |
| CCL4 | -1.8 | -2.9 | | | -5.4 | | | |
| CCL4L2 | -4.4 | -10 | | | -4.1 | | | |
| CCL5 | -1.6 | | -12.8 | | -6.5 | | | |
| CCL7 | -11.7 | | | | | | -17.9 | |
| CCL8 | | | | | | | | |
| CCL13 | | | | | -4.6 | | | |
| CCL14 | | | | | -2.3 | | | |
| CCL18 | | | | -2.9 | -2.4 | | | |
| CCL20 | -2.2 | -20.7 | | | -3.1 | | | |
| CCL22 | | -4.8 | | | -9.9 | | | -5.5 |
| CCL23 | -2.2 | -7.3 | | | -3.1 | | | -1.9 |
| CCL24 | | | -7E+06 | | -10.1 | | | -4.8 |
| CCL25 | | | | | -3.5 | | | |
| CCL26 | | | | | -4.3 | | | |
| CCL28 | -1.4 | | | | | | | |
