## Supplementary material for "Chemokine Receptor 1 and its associated immune pathway are downregulated in SF3B1^MT^ blood and non-blood cancers": Table_S16

**Table S16. SF3B1$^{MT}$ and SF3B1$^{WT}$ patient information in GEO, TCGA and BeatAML cohorts.**

| **MDS-O (GSE85712)** | | |
|---|---|---|
| **Sample ID** | **Tumor Name** | **SF3B1 Mutation** |
| GSM2282084 | Myelodysplastic Syndrome | K666R |
| GSM2282087 | Myelodysplastic Syndrome | K666T |
| GSM2282082 | Myelodysplastic Syndrome | K700E |
| GSM2282083 | Myelodysplastic Syndrome | K700E |
| GSM2282088 | Myelodysplastic Syndrome | K700E |
| GSM2282089 | Myelodysplastic Syndrome | K700E |
| GSM2282081 | Myelodysplastic Syndrome | WT |
| GSM2282085 | Myelodysplastic Syndrome | WT |
| GSM2282086 | Myelodysplastic Syndrome | WT |
| **MDS-Z (GSE128805)** | | |
| **Sample ID** | **Tumor Name** | **SF3B1 Mutation** |
| GSM3685748 | Myelodysplastic Syndrome | K700E |
| GSM3685749 | Myelodysplastic Syndrome | K700E |
| GSM3685750 | Myelodysplastic Syndrome | D781G |
| GSM3685751 | Myelodysplastic Syndrome | H662D |
| GSM3685752 | Myelodysplastic Syndrome | K700E |
| GSM3685762 | Myelodysplastic Syndrome | D781G |
| GSM3685753 | Myelodysplastic Syndrome | WT |
| GSM3685754 | Myelodysplastic Syndrome | WT |
| GSM3685755 | Myelodysplastic Syndrome | WT |
| GSM3685756 | Myelodysplastic Syndrome | WT |
| GSM3685757 | Myelodysplastic Syndrome | WT |
| GSM3685758 | Myelodysplastic Syndrome | WT |
| GSM3685759 | Myelodysplastic Syndrome | WT |
| GSM3685760 | Myelodysplastic Syndrome | WT |
| GSM3685761 | Myelodysplastic Syndrome | WT |
| **MDS-M (GSE128429)** | | |
| **Sample ID** | **Tumor Name** | **SF3B1 Mutation** |
| GSM3674357 | Myelodysplastic Syndrome | H662Q |
| GSM3674358 | Myelodysplastic Syndrome | K666T |
| GSM3674359 | Myelodysplastic Syndrome | K700E |
| GSM3674360 | Myelodysplastic Syndrome | K700E |
| GSM3674361 | Myelodysplastic Syndrome | K666N |
| GSM3674380 | Myelodysplastic Syndrome | WT |
| GSM3674381 | Myelodysplastic Syndrome | WT |
| GSM3674382 | Myelodysplastic Syndrome | WT |
| GSM3674383 | Myelodysplastic Syndrome | WT |
| GSM3674384 | Myelodysplastic Syndrome | WT |
| GSM3674385 | Myelodysplastic Syndrome | WT |

| | | |
|---|---|---|
| GSM3674386 | Myelodysplastic Syndrome | WT |
| GSM3674387 | Myelodysplastic Syndrome | WT |

**AML**

| Sample ID | Tumor Name | SF3B1 Mutation |
|---|---|---|
| 2202 | BEATAML1.0-COHORT | G742N |
| 2254 | BEATAML1.0-COHORT | K700E |
| 2261 | BEATAML1.0-COHORT | K666N |
| 2366 | BEATAML1.0-COHORT | K666Q |
| 2453 | BEATAML1.0-COHORT | H662Q |
| 2482 | BEATAML1.0-COHORT | K666N |
| 2485 | BEATAML1.0-COHORT | K666N |
| 2041 | BEATAML1.0-COHORT | WT |
| 2122 | BEATAML1.0-COHORT | WT |
| 2197 | BEATAML1.0-COHORT | WT |
| 2259 | BEATAML1.0-COHORT | WT |
| 2321 | BEATAML1.0-COHORT | WT |
| 2376 | BEATAML1.0-COHORT | WT |

**CLL (GSE72790)**

| Sample ID | Tumor Name | SF3B1 Mutation |
|---|---|---|
| GSM1872051 | Chronic Lymphocytic Leukemia | K700EIG742D |
| GSM1872052 | Chronic Lymphocytic Leukemia | Y623C |
| GSM1872053 | Chronic Lymphocytic Leukemia | G742D |
| GSM1872054 | Chronic Lymphocytic Leukemia | V701F |
| GSM1872055 | Chronic Lymphocytic Leukemia | G740E |
| GSM1872056 | Chronic Lymphocytic Leukemia | G742DII704F |
| GSM1872057 | Chronic Lymphocytic Leukemia | G742D |
| GSM1872058 | Chronic Lymphocytic Leukemia | WT |
| GSM1872059 | Chronic Lymphocytic Leukemia | WT |
| GSM1872060 | Chronic Lymphocytic Leukemia | WT |
| GSM1872061 | Chronic Lymphocytic Leukemia | WT |
| GSM1872062 | Chronic Lymphocytic Leukemia | WT |
| GSM1872063 | Chronic Lymphocytic Leukemia | WT |

**BRCA**

| Sample ID | Tumor Name | SF3B1 Mutation |
|---|---|---|
| TCGA-A2-A0EV | Breast Invasive Carcinoma | K700E |
| TCGA-BH-A0DQ | Breast Invasive Carcinoma | K700E |
| TCGA-BH-A18M | Breast Invasive Carcinoma | K700E |
| TCGA-BH-A201 | Breast Invasive Carcinoma | K700E |
| TCGA-C8-A26V | Breast Invasive Carcinoma | K700E |
| TCGA-E2-A56Z | Breast Invasive Carcinoma | K700E |
| TCGA-BH-A5J0 | Breast Invasive Carcinoma | K700E |
| TCGA-AN-A0XO | Breast Invasive Carcinoma | K700E |

| | | |
|---|---|---|
| TCGA-EW-A1P5 | Breast Invasive Carcinoma | K700E |
| TCGA-B6-A0WT | Breast Invasive Carcinoma | K700E |
| TCGA-BH-A0HP | Breast Invasive Carcinoma | K666E |
| TCGA-LD-A74U | Breast Invasive Carcinoma | K666N |
| TCGA-AC-A2B8 | Breast Invasive Carcinoma | E622Q |
| TCGA-AO-A0JC | Breast Invasive Carcinoma | E802Q |
| TCGA-AR-A0TY | Breast Invasive Carcinoma | C1059R |
| TCGA-BH-A0HY | Breast Invasive Carcinoma | Y765C |
| TCGA-E9-A1R3 | Breast Invasive Carcinoma | N626D |
| TCGA-B6-A0I2 | Breast Invasive Carcinoma | A633V |
| TCGA-A8-A07P | Breast Invasive Carcinoma | WT |
| TCGA-B6-A0RG | Breast Invasive Carcinoma | WT |
| TCGA-AR-A0TX | Breast Invasive Carcinoma | WT |
| TCGA-D8-A1X8 | Breast Invasive Carcinoma | WT |
| TCGA-E2-A109 | Breast Invasive Carcinoma | WT |
| TCGA-A8-A08T | Breast Invasive Carcinoma | WT |
| TCGA-E9-A1N4 | Breast Invasive Carcinoma | WT |
| TCGA-BH-A203 | Breast Invasive Carcinoma | WT |
| TCGA-XX-A89A | Breast Invasive Carcinoma | WT |
| TCGA-A7-A2KD | Breast Invasive Carcinoma | WT |
| TCGA-BH-A0BW | Breast Invasive Carcinoma | WT |
| TCGA-AC-A4ZE | Breast Invasive Carcinoma | WT |
| TCGA-OL-A66P | Breast Invasive Carcinoma | WT |
| TCGA-AO-A1KR | Breast Invasive Carcinoma | WT |
| TCGA-C8-A134 | Breast Invasive Carcinoma | WT |
| TCGA-A2-A04V | Breast Invasive Carcinoma | WT |
| TCGA-EW-A1IX | Breast Invasive Carcinoma | WT |
| TCGA-A8-A097 | Breast Invasive Carcinoma | WT |

**UVM**

| Sample ID | Tumor Name | SF3B1 Mutation |
|---|---|---|
| TCGA-WC-AA9A | Uveal Melanoma | K666T |
| TCGA-V4-A9E5 | Uveal Melanoma | K666T |
| TCGA-V4-A9EC | Uveal Melanoma | T663P |
| TCGA-WC-A881 | Uveal Melanoma | H662R |
| TCGA-V4-A9ES | Uveal Melanoma | R625H |
| TCGA-WC-AA9E | Uveal Melanoma | R625H |
| TCGA-WC-A885 | Uveal Melanoma | R625H |
| TCGA-V4-A9EZ | Uveal Melanoma | R625H |
| TCGA-V4-A9E9 | Uveal Melanoma | R625H |
| TCGA-V4-A9F4 | Uveal Melanoma | R625H |
| TCGA-VD-A8KH | Uveal Melanoma | R625H |
| TCGA-V4-A9EA | Uveal Melanoma | R625C |
| TCGA-VD-A8K9 | Uveal Melanoma | R625C |
| TCGA-VD-A8KA | Uveal Melanoma | R625C |

| | | |
|---|---|---|
| TCGA-VD-A8KB | Uveal Melanoma | R625C |
| TCGA-V4-A9EW | Uveal Melanoma | R625C |
| TCGA-YZ-A985 | Uveal Melanoma | R625C |
| TCGA-V4-A9EJ | Uveal Melanoma | R625H |
| TCGA-V4-A9EH | Uveal Melanoma | WT |
| TCGA-V4-A9F0 | Uveal Melanoma | WT |
| TCGA-V4-A9EF | Uveal Melanoma | WT |
| TCGA-V3-A9ZY | Uveal Melanoma | WT |
| TCGA-V4-A9EU | Uveal Melanoma | WT |
| TCGA-V4-A9EL | Uveal Melanoma | WT |
| TCGA-V4-A9EE | Uveal Melanoma | WT |
| TCGA-VD-A8KE | Uveal Melanoma | WT |
| TCGA-V4-A9F1 | Uveal Melanoma | WT |
| TCGA-WC-A884 | Uveal Melanoma | WT |
| TCGA-V4-A9EK | Uveal Melanoma | WT |
| TCGA-VD-AA8R | Uveal Melanoma | WT |
| TCGA-V4-A9EX | Uveal Melanoma | WT |
| TCGA-VD-A8KM | Uveal Melanoma | WT |
| TCGA-WC-A87T | Uveal Melanoma | WT |
| TCGA-VD-AA8O | Uveal Melanoma | WT |
| TCGA-YZ-A980 | Uveal Melanoma | WT |
| TCGA-VD-A8KK | Uveal Melanoma | WT |
| TCGA-V3-A9ZX | Uveal Melanoma | WT |
| TCGA-V4-A9F2 | Uveal Melanoma | WT |
| TCGA-RZ-AB0B | Uveal Melanoma | WT |
| TCGA-VD-AA8Q | Uveal Melanoma | WT |
| TCGA-V4-A9EI | Uveal Melanoma | WT |
| TCGA-V4-A9E8 | Uveal Melanoma | WT |
| TCGA-VD-A8KO | Uveal Melanoma | WT |
| TCGA-VD-A8KL | Uveal Melanoma | WT |
| TCGA-VD-A8KN | Uveal Melanoma | WT |
| TCGA-V4-A9F3 | Uveal Melanoma | WT |
| TCGA-WC-A882 | Uveal Melanoma | WT |
| TCGA-V4-A9ET | Uveal Melanoma | WT |
| TCGA-V4-A9E7 | Uveal Melanoma | WT |
| TCGA-V4-A9F5 | Uveal Melanoma | WT |
| TCGA-YZ-A982 | Uveal Melanoma | WT |
| TCGA-VD-AA8M | Uveal Melanoma | WT |
| TCGA-VD-AA8P | Uveal Melanoma | WT |
| TCGA-VD-AA8S | Uveal Melanoma | WT |
| TCGA-WC-A88A | Uveal Melanoma | WT |
| TCGA-VD-AA8T | Uveal Melanoma | WT |
| TCGA-VD-A8KJ | Uveal Melanoma | WT |
| TCGA-YZ-A984 | Uveal Melanoma | WT |
| TCGA-WC-A880 | Uveal Melanoma | WT |

| | | |
|---|---|---|
| TCGA-VD-A8KF | Uveal Melanoma | WT |
| TCGA-WC-A87W | Uveal Melanoma | WT |
| TCGA-VD-AA8N | Uveal Melanoma | WT |
| TCGA-VD-A8K7 | Uveal Melanoma | WT |
| TCGA-V4-A9EQ | Uveal Melanoma | WT |
| TCGA-V4-A9EM | Uveal Melanoma | WT |
| TCGA-V4-A9EY | Uveal Melanoma | WT |
| TCGA-VD-A8KI | Uveal Melanoma | WT |
| TCGA-VD-A8KD | Uveal Melanoma | WT |
| TCGA-V4-A9ED | Uveal Melanoma | WT |
| TCGA-V4-A9F7 | Uveal Melanoma | WT |
| TCGA-WC-A87Y | Uveal Melanoma | WT |
| TCGA-WC-A883 | Uveal Melanoma | WT |
| TCGA-WC-A888 | Uveal Melanoma | WT |
| TCGA-WC-A87U | Uveal Melanoma | WT |
| TCGA-V4-A9EO | Uveal Melanoma | WT |
| TCGA-VD-A8KG | Uveal Melanoma | WT |
| TCGA-V4-A9F8 | Uveal Melanoma | WT |
| TCGA-V4-A9EV | Uveal Melanoma | WT |
| TCGA-YZ-A983 | Uveal Melanoma | WT |
| TCGA-VD-A8K8 | Uveal Melanoma | WT |

**K562 (GSE95011)**

| Sample ID | Cell Line | SF3B1 Mutation |
|---|---|---|
| GSM2477628 | K562 | K700E |
| GSM2477629 | K562 | K700E |
| GSM2477630 | K562 | K700E |
| GSM2477622 | K562 | WT |
| GSM2477623 | K562 | WT |
| GSM2477624 | K562 | WT |

**NALM-6 (GSE72790)**

| Sample ID | Cell Line | SF3B1 Mutation |
|---|---|---|
| GSM1872081 | Nalm-6 | K700E |
| GSM1872082 | Nalm-6 | K700E |
| GSM1872083 | Nalm-6 | K700E |
| GSM1872078 | Nalm-6 | WT |
| GSM1872079 | Nalm-6 | WT |
| GSM1872080 | Nalm-6 | WT |

| **Table S16. U2AF1$^{MT}$, SRSF2$^{MT}$ and ZRSR2$^{MT}$ patient information in GEO.** | | |
|---|---|---|
| **MDS-M (GSE128429)** | | |
| **Sample ID** | **Tumor Name** | **SF3B1 Mutation** |
| GSM3674362 | Myelodysplastic Syndrome | SRSF2P95R |
| GSM3674363 | Myelodysplastic Syndrome | SRSF2P95H |
| GSM3674364 | Myelodysplastic Syndrome | SRSF2P95R |
| GSM3674365 | Myelodysplastic Syndrome | SRSF2P95_R102del |
| GSM3674366 | Myelodysplastic Syndrome | SRSF2P95_R102del |
| GSM3674367 | Myelodysplastic Syndrome | SRSF2R150fs |
| GSM3674368 | Myelodysplastic Syndrome | U2AF1S34F |
| GSM3674369 | Myelodysplastic Syndrome | U2AF1S34Y |
| GSM3674370 | Myelodysplastic Syndrome | U2AF1R156H |
| GSM3674371 | Myelodysplastic Syndrome | U2AF1Q157R |
| GSM3674372 | Myelodysplastic Syndrome | U2AF1Q157P |
| GSM3674373 | Myelodysplastic Syndrome | U2AF1S34F |
| GSM3674374 | Myelodysplastic Syndrome | U2AF1S34F |
| GSM3674375 | Myelodysplastic Syndrome | U2AF1Q157P |
| GSM3674376 | Myelodysplastic Syndrome | ZRSR2Y373X |
| GSM3674377 | Myelodysplastic Syndrome | ZRSR2H330Y |
| GSM3674378 | Myelodysplastic Syndrome | ZRSR2T411fs |
| GSM3674379 | Myelodysplastic Syndrome | ZRSR2E104fs |
