## Supplementary material for "Chemokine Receptor 1 and its associated immune pathway are downregulated in SF3B1^MT^ blood and non-blood cancers": Table_S17

| Table S17. Primers used for qPCR, RT-PCR and Sanger sequencing. | | |
|---|---|---|
| **Gene** | **Forward Primer** | **Reverse Primer** |
| FL1 | ACCTCAGTTACCTCAGGGAAAG | TGTTATTGCCCCAAGCTCCT |
| GAPDH | TGCACCACCACCTGCTTAGC | GGCATGGACTGTGGTCATGAG |
| MEF2C | GCAAGCAAAATCTCCTCCCC | TTTATCCTTTGATTCACTGATGGC |
| TFF3 | TCTGCTGAGGAGTACGTGG | GGATCCTGGAGTCAAAGCAG |
| VAV1 | AGGAGAGTGTAGGTGATGAAGA | CATTCTCCACGCAGTCATACA |
| APPL2-S | GCACTTTAAAGGATCTATTTGG | CTCATTCTCCTTTTTCTTAGGC |
| BRD9-S | AAGTCCTATCCCGACGTTTC | AAAGCAGACTCCGTGGTAAT |
| DVL2-S | AAGTCCTATCCCGACGTTTC | AAAGCAGACTCCGTGGTAAT |
| MAP3K7-S | GATGGAATATGCTGAAGGGG | CACTCCTTGGGAACACTGTA |
| SF3B1 | AATTTGGGCTACTGATTTGGGGA | GCCTTCAAGAAAGCAGCCAAAC |
